## Supplemental Table 1 for "Tectonic Complex Impedes Diffusion through the Ciliary Transition Zone to Ensure Proper Sorting of Membrane Proteins"

| Gene Symbol | Protein FDR Confidence: Combined | Accession | Description | Exp. q-value: Combined | Sum PEP Score | Coverage [%] | # Peptides | # PSMs | # Unique Peptides | # AAs | MW [kDa] | calc. pI | Abundance Ratio: (Tctn1FHet ROS at 3M) / (Tctn1FK | Abundance Ratio: (WT B6 ROS at 3M) / (Tctn1FKO | Abundance Ratio: (Tctn1FHet ROS at 3M) / (WT B6 ROS at 3M) | Abundance Ratio (log2): (Tctn1FHet ROS at 3M) / | Abundance Ratio (log2): KO / HET | Abundance Ratio (log2): (WT B6 ROS at 3M) / (Tctn1FKO ROS at 3M) | AVG Abundance Ratio WTandHet/KO | AVG Abundance Ratio KO/Hetand WT | Abundance Ratio (log2): (Tctn1FHet ROS at 3M) / (WT B6 ROS at 3M) | Abundance Ratio P-Value: (Tctn1FHet ROS at 3M) / (Tctn1FKO ROS at 3M) | Abundance Ratio P-Value: (WT B6 ROS at 3M) / (Tctn1FKO ROS at 3M) | Abundance Ratio P-Value: (Tctn1FHet ROS at 3M) / (WT B6 ROS at 3M) | Abundance Ratio Adj. P-Value: (Tctn1FHet ROS at 3M) / (Tctn1FKO ROS | Abundance Ratio Adj. P-Value: (WT B6 ROS at 3M) / (Tctn1FKO ROS at 3M) | AVG P Value Ratio Adj. WTandHet/KO | Abundance Ratio Adj. P-Value: (Tctn1FHet ROS at 3M) / (WT B6 ROS at 3M) |
|---|---|---|---|---|---|---|---|---|---|---|---|---|---|---|---|---|---|---|---|---|---|---|---|---|---|---|---|---|
| Btc | High | Q05928 | Probetacellulin OS=Mus musculus OX=10090 GN=Btc PE=1 SV=1 | 0 | 14.868 | 20 | 2 | 6 | 2 | 177 | 19.7 | 8.19 | 0.05 | 0.034 | 1.441 | -4.33 | 4.33 | -4.86 | -1.62 | 4.595 | 0.53 | 1E-17 | 1E-17 | 0.012478101 | 9.345E-15 | 2.8035E-14 | 1.869E-14 | 0.183957829 |
| Stx3 | High | Q64704 | Syntaxin-3 OS=Mus musculus OX=10090 GN=Stx3 PE=1 SV=1 | 0 | 266.488 | 54 | 23 | 157 | 23 | 289 | 33.2 | 5.45 | 0.074 | 0.081 | 0.906 | -3.77 | 3.77 | -3.62 | -1.2066667 | 3.695 | -0.14 | 1E-17 | 6.66134E-15 | 0.592857068 | 9.345E-15 | 9.33753E-12 | 4.67344E-12 | 0.976360704 |
| Slco4a1 | High | Q8K078 | Solute carrier organic anion transporter family member 4A1 OS=Mus musculus OX=10090 GN=Slc | 0.001 | 7.82 | 4 | 3 | 3 | 3 | 723 | 77.6 | 7.85 | 0.117 | 0.129 | 0.911 | -3.09 | 3.09 | -2.96 | -0.9866667 | 3.025 | -0.13 | 2.88658E-15 | 1.86638E-10 | 0.601859362 | 2.31215E-12 | 1.15804E-07 | 5.79031E-08 | 0.97826833 |
| Slc38a3 | High | Q9DCP2 | Sodium-coupled neutral amino acid transporter 3 OS=Mus musculus OX=10090 GN=Slc38a3 PE=1 | 0 | 58.459 | 19 | 10 | 43 | 10 | 505 | 55.6 | 7.09 | 0.101 | 0.078 | 1.299 | -3.31 | 3.31 | -3.69 | -1.23 | 3.5 | 0.38 | 1.24345E-14 | 2.44249E-15 | 0.089072917 | 7.74669E-12 | 4.56502E-12 | 6.15585E-12 | 0.5746627 |
| Slc6a6 | High | O35316 | Sodium- and chloride-dependent taurine transporter OS=Mus musculus OX=10090 GN=Slc6a6 PE= | 0 | 51.885 | 20 | 11 | 28 | 9 | 621 | 69.8 | 7.23 | 0.12 | 0.12 | 0.997 | -3.06 | 3.06 | -3.06 | -1.02 | 3.06 | 0 | 9.3392E-13 | 4.49283E-11 | 0.943619599 | 4.76044E-10 | 4.19855E-08 | 2.12308E-08 | 0.992535481 |
| Cetn3 | High | A0A286YD | Centrin-3 (Fragment) OS=Mus musculus OX=10090 GN=Cetn3 PE=1 SV=1 | 0.001 | 4.685 | 16 | 2 | 2 | 2 | 97 | 11.4 | 4.75 | 6.242 | 6.849 | 0.911 | 2.64 | -2.64 | 2.78 | 0.92666667 | -2.71 | -0.13 | 1.53477E-12 | 6.04487E-09 | 0.611864335 | 5.73698E-10 | 1.61398E-06 | 8.07277E-07 | 0.97826833 |
| Antxr2 | High | Q6DFX2 | Anthrax toxin receptor 2 OS=Mus musculus OX=10090 GN=Antxr2 PE=1 SV=1 | 0 | 51.185 | 27 | 11 | 15 | 11 | 487 | 53.2 | 7.56 | 0.123 | 0.086 | 1.423 | -3.03 | 3.03 | -3.54 | -1.18 | 3.285 | 0.51 | 1.66622E-12 | 2.84217E-14 | 0.037584245 | 5.83907E-10 | 3.18721E-11 | 3.0789E-10 | 0.373579725 |
| Podxl | High | Q9R0M4 | Podocalyxin OS=Mus musculus OX=10090 GN=Podxl PE=1 SV=2 | 0 | 21.186 | 20 | 6 | 7 | 6 | 503 | 53.4 | 4.97 | 0.178 | 0.178 | 1.001 | -2.49 | 2.49 | -2.49 | -0.83 | 2.49 | 0 | 2.9321E-12 | 7.15313E-08 | 0.924250094 | 9.67075E-10 | 1.74381E-05 | 8.71953E-06 | 0.992535481 |
| Hcn4 | High | B2RY58 | Hyperpolarization-activated, cyclic nucleotide-gated K+ 4 OS=Mus musculus OX=10090 GN=Hcn4 I | 0 | 9.823 | 2 | 3 | 4 | 1 | 1201 | 129 | 9.04 | 0.184 | 0.2 | 0.923 | -2.44 | 2.44 | -2.32 | -0.7733333 | 2.38 | -0.12 | 3.95439E-12 | 3.69695E-07 | 0.760538421 | 1.23179E-09 | 6.6867E-05 | 3.34341E-05 | 0.992535481 |
| Scml2 | Medium | B1AVB3 | Scm polycomb group protein-like 2 OS=Mus musculus OX=10090 GN=Scml2 PE=1 SV=1 | 0.029 | 1.917 | 1 | 1 | 1 | 1 | 969 | 106.7 | 9.54 | 5.833 | 7.68 | 0.759 | 2.54 | -2.54 | 2.94 | 0.98 | -2.74 | -0.4 | 2.68157E-11 | 2.68495E-10 | 0.111363606 | 6.01422E-09 | 1.15804E-07 | 6.0909E-08 | 0.632640059 |
| Scamp5 | High | Q9JKD3 | Secretory carrier-associated membrane protein 5 OS=Mus musculus OX=10090 GN=Scamp5 PE=1 | 0 | 45.043 | 39 | 5 | 32 | 5 | 235 | 26.1 | 8.54 | 0.14 | 0.143 | 0.974 | -2.84 | 2.84 | -2.8 | -0.9333333 | 2.82 | -0.04 | 3.54226E-11 | 1.51011E-09 | 0.942607627 | 7.63901E-09 | 6.04799E-07 | 3.06219E-07 | 0.992535481 |
| Ndrg3 | High | Q9QYF9 | Protein NDRG3 OS=Mus musculus OX=10090 GN=Ndrg3 PE=1 SV=1 | 0 | 70.684 | 42 | 10 | 24 | 10 | 375 | 41.5 | 5.25 | 0.147 | 0.145 | 1.008 | -2.77 | 2.77 | -2.78 | -0.9266667 | 2.775 | 0.01 | 1.05997E-10 | 1.98697E-09 | 0.997422662 | 2.20121E-08 | 7.4273E-07 | 3.82371E-07 | 0.999917596 |
| Ndrg1 | High | Q62433 | Protein NDRG1 OS=Mus musculus OX=10090 GN=Ndrg1 PE=1 SV=1 | 0 | 119.965 | 33 | 8 | 29 | 8 | 394 | 43 | 6.1 | 0.154 | 0.18 | 0.86 | -2.7 | 2.7 | -2.48 | -0.8266667 | 2.59 | -0.22 | 3.34225E-10 | 8.75154E-08 | 0.41018113 | 6.69285E-08 | 2.04458E-05 | 1.02564E-05 | 0.936397673 |
| Stxbp1 | High | O08599 | Syntaxin-binding protein 1 OS=Mus musculus OX=10090 GN=Stxbp1 PE=1 SV=2 | 0 | 545.288 | 84 | 58 | 319 | 57 | 594 | 67.5 | 6.96 | 0.17 | 0.146 | 1.161 | -2.56 | 2.56 | -2.77 | -0.9233333 | 2.665 | 0.22 | 2.51364E-09 | 2.24068E-09 | 0.288862827 | 4.85999E-07 | 7.85217E-07 | 6.35608E-07 | 0.888636466 |
| Slc16a3 | High | P57787 | Monocarboxylate transporter 4 OS=Mus musculus OX=10090 GN=Slc16a3 PE=1 SV=1 | 0 | 10.325 | 6 | 3 | 6 | 3 | 470 | 50.3 | 7.96 | 0.185 | 0.188 | 0.982 | -2.43 | 2.43 | -2.41 | -0.8033333 | 2.42 | -0.03 | 2.98234E-09 | 1.93762E-07 | 0.914752042 | 5.574E-07 | 3.88008E-05 | 1.96791E-05 | 0.992535481 |
| Tnip1 | High | D3Z3B6 | TNFAIP3-interacting protein 1 (Fragment) OS=Mus musculus OX=10090 GN=Tnip1 PE=1 SV=1 | 0 | 13.272 | 8 | 2 | 3 | 2 | 272 | 29.4 | 5.44 | 0.23 | 0.262 | 0.877 | -2.12 | 2.12 | -1.93 | -0.6433333 | 2.025 | -0.19 | 4.32723E-09 | 2.71412E-05 | 0.459940784 | 7.13611E-07 | 0.00190226 | 0.000951487 | 0.959744396 |
| Spata7 | High | Q80VP2 | Spermatogenesis-associated protein 7 homolog OS=Mus musculus OX=10090 GN=Spata7 PE=1 S' | 0 | 11.905 | 5 | 4 | 7 | 4 | 582 | 65.6 | 6.7 | 4.443 | 5.397 | 0.823 | 2.15 | -2.15 | 2.43 | 0.81 | -2.29 | -0.28 | 8.48896E-09 | 3.68363E-07 | 0.220099222 | 1.32215E-06 | 6.6867E-05 | 3.40946E-05 | 0.805859554 |
| Kcnj14 | High | Q8JZN3 | ATP-sensitive inward rectifier potassium channel 14 OS=Mus musculus OX=10090 GN=Kcnj14 PE= | 0 | 56.165 | 32 | 11 | 19 | 11 | 434 | 47.6 | 5.71 | 0.184 | 0.174 | 1.052 | -2.45 | 2.45 | -2.52 | -0.84 | 2.485 | 0.07 | 1.33339E-08 | 5.2794E-08 | 0.79470411 | 1.96745E-06 | 1.34553E-05 | 7.71135E-06 | 0.992535481 |
| Slc11a2 | Medium | P49282 | Natural resistance-associated macrophage protein 2 OS=Mus musculus OX=10090 GN=Slc11a2 Pl | 0.025 | 1.986 | 2 | 1 | 1 | 1 | 568 | 62.3 | 5.99 | 0.252 | 0.305 | 0.826 | -1.99 | 1.99 | -1.71 | -0.57 | 1.85 | -0.28 | 1.32012E-08 | 0.00015099 | 0.36127728 | 1.96745E-06 | 0.00814037 | 0.004071169 | 0.917317898 |
| Cep68 | High | Q3TRP8 | Centrosomal protein of 68 kDa OS=Mus musculus OX=10090 GN=Cep68 PE=1 SV=1 | 0 | 8.95 | 3 | 1 | 1 | 1 | 633 | 67.7 | 5.94 | 4.297 | 3.709 | 1.159 | 2.1 | -2.1 | 1.89 | 0.63 | -1.995 | 0.21 | 2.26733E-08 | 8.17352E-05 | 0.402802101 | 3.17823E-06 | 0.00492784 | 0.002465509 | 0.93262326 |
| Ptp4a3 | High | Q9D658 | Protein tyrosine phosphatase type IVA 3 OS=Mus musculus OX=10090 GN=Ptp4a3 PE=1 SV=1 | 0.001 | 5.322 | 10 | 2 | 2 | 2 | 173 | 19.6 | 9.32 | 0.244 | 0.22 | 1.111 | -2.04 | 2.04 | -2.19 | -0.73 | 2.115 | 0.15 | 3.94496E-08 | 2.17726E-06 | 0.526508104 | 5.14404E-06 | 0.000277453 | 0.000141298 | 0.964378027 |
| Jam2 | High | Q9JI59 | Junctional adhesion molecule B OS=Mus musculus OX=10090 GN=Jam2 PE=1 SV=1 | 0 | 15.157 | 14 | 4 | 6 | 4 | 298 | 33 | 8.4 | 0.224 | 0.203 | 1.108 | -2.16 | 2.16 | -2.3 | -0.7666667 | 2.23 | 0.15 | 4.18006E-08 | 6.23686E-07 | 0.500383541 | 5.20835E-06 | 9.71391E-05 | 5.11737E-05 | 0.964378027 |
| Tax1bp1 | High | Q3UKC1 | Tax1-binding protein 1 homolog OS=Mus musculus OX=10090 GN=Tax1bp1 PE=1 SV=2 | 0 | 36.166 | 15 | 12 | 14 | 12 | 814 | 93.6 | 5.33 | 0.21 | 0.198 | 1.056 | -2.25 | 2.25 | -2.33 | -0.7766667 | 2.29 | 0.08 | 4.77783E-08 | 4.52345E-07 | 0.736221504 | 5.82376E-06 | 7.92593E-05 | 4.25415E-05 | 0.992535481 |
| Uba5 | High | D6RH76 | Ubiquitin-like modifier-activating enzyme 5 OS=Mus musculus OX=10090 GN=Uba5 PE=1 SV=1 | 0.009 | 2.828 | 12 | 1 | 1 | 1 | 130 | 14.6 | 8.95 | 0.235 | 0.325 | 0.725 | -2.09 | 2.09 | -1.62 | -0.54 | 1.855 | -0.46 | 7.36762E-08 | 0.000282273 | 0.134734014 | 8.26205E-06 | 0.013101929 | 0.006555096 | 0.697570122 |
| Cacng7 | High | P62956 | Voltage-dependent calcium channel gamma-7 subunit OS=Mus musculus OX=10090 GN=Cacng7 I | 0.001 | 5.599 | 9 | 1 | 1 | 1 | 275 | 31 | 7.12 | 0.261 | 0.228 | 1.146 | -1.94 | 1.94 | -2.13 | -0.71 | 2.035 | 0.2 | 7.57268E-08 | 2.03801E-06 | 0.49820919 | 8.3255E-06 | 0.000272074 | 0.0001402 | 0.964378027 |
| Marveld3 | Medium | Q9D956 | MARVEL domain-containing protein 3 OS=Mus musculus OX=10090 GN=Marveld3 PE=2 SV=1 | 0.035 | 1.78 | 2 | 1 | 1 | 1 | 376 | 42.1 | 8.82 | 0.272 | 0.381 | 0.713 | -1.88 | 1.88 | -1.39 | -0.4633333 | 1.635 | -0.49 | 8.11839E-08 | 0.001866749 | 0.076013573 | 8.58864E-06 | 0.064464881 | 0.032236735 | 0.523043272 |
| Slc4a7 | High | F8VQC9 | Anion exchange protein OS=Mus musculus OX=10090 GN=Slc4a7 PE=1 SV=1 | 0 | 45.21 | 11 | 11 | 13 | 9 | 1131 | 127.1 | 6.44 | 0.207 | 0.256 | 0.81 | -2.27 | 2.27 | -1.97 | -0.6566667 | 2.12 | -0.3 | 1.18151E-07 | 1.94716E-05 | 0.19568865 | 1.03512E-05 | 0.001530147 | 0.000770249 | 0.784517504 |
| A2m | High | Q6GQT1 | Alpha-2-macroglobulin-P OS=Mus musculus OX=10090 GN=A2m PE=2 SV=2 | 0 | 80.387 | 17 | 18 | 21 | 18 | 1474 | 164.2 | 6.61 | 0.209 | 0.25 | 0.835 | -2.26 | 2.26 | -2 | -0.6666667 | 2.13 | -0.26 | 1.43126E-07 | 1.47362E-05 | 0.289207035 | 1.21592E-05 | 0.001400438 | 0.000706298 | 0.888636466 |
| Prcd | High | Q00LT2 | Photoreceptor disk component PRCD OS=Mus musculus OX=10090 GN=Prcd PE=1 SV=1 | 0 | 34.363 | 57 | 3 | 17 | 3 | 53 | 5.9 | 9.14 | 3.78 | 3.467 | 1.09 | 1.92 | -1.92 | 1.79 | 0.59666667 | -1.855 | 0.12 | 1.92179E-07 | 0.000193513 | 0.611095742 | 1.56167E-05 | 0.009775019 | 0.004895318 | 0.97826833 |
| Shisa2 | High | Q8QZV2 | Protein shisa-2 homolog OS=Mus musculus OX=10090 GN=Shisa2 PE=2 SV=1 | 0 | 25.886 | 17 | 4 | 5 | 4 | 295 | 31.4 | 7.83 | 0.276 | 0.261 | 1.06 | -1.86 | 1.86 | -1.94 | -0.6466667 | 1.9 | 0.08 | 2.16556E-07 | 2.57105E-05 | 0.733080447 | 1.73461E-05 | 0.001896827 | 0.000957087 | 0.992535481 |
| Ift20 | High | Z4YJL9 | Intraflagellar transport protein 20 homolog (Fragment) OS=Mus musculus OX=10090 GN=Ift20 PE | 0 | 11.204 | 26 | 2 | 2 | 2 | 111 | 12.7 | 5.63 | 3.595 | 3.291 | 1.092 | 1.85 | -1.85 | 1.72 | 0.57333333 | -1.785 | 0.13 | 2.49334E-07 | 0.00025065 | 0.55795079 | 1.94169E-05 | 0.012115463 | 0.00606744 | 0.971568588 |
| Sppl2b | High | Q3TD49 | Signal peptide peptidase-like 2B OS=Mus musculus OX=10090 GN=Sppl2b PE=1 SV=2 | 0 | 16.781 | 10 | 4 | 4 | 4 | 578 | 63.8 | 7.62 | 0.274 | 0.214 | 1.28 | -1.87 | 1.87 | -2.22 | -0.74 | 2.045 | 0.36 | 3.34828E-07 | 1.51153E-06 | 0.109160494 | 2.57175E-05 | 0.00020671 | 0.000116214 | 0.631643849 |
| Cetn2 | High | Q9R1K9 | Centrin-2 OS=Mus musculus OX=10090 GN=Cetn2 PE=1 SV=1 | 0 | 51.335 | 59 | 7 | 14 | 7 | 172 | 19.8 | 5 | 4.377 | 4.143 | 1.056 | 2.13 | -2.13 | 2.05 | 0.68333333 | -2.09 | 0.08 | 4.11188E-07 | 1.92165E-05 | 0.66873122 | 3.11558E-05 | 0.001530147 | 0.000780651 | 0.987489386 |
| Cnnm2 | High | Q3TWN3 | Metal transporter CNNM2 OS=Mus musculus OX=10090 GN=Cnnm2 PE=1 SV=3 | 0 | 41.024 | 11 | 9 | 15 | 5 | 875 | 96.6 | 6.21 | 0.276 | 0.3 | 0.92 | -1.86 | 1.86 | -1.74 | -0.58 | 1.8 | -0.12 | 4.39545E-07 | 0.000157186 | 0.665464398 | 3.28604E-05 | 0.008393729 | 0.004213295 | 0.987366731 |
| Dnajc5 | High | P60904 | DnaJ homolog subfamily C member 5 OS=Mus musculus OX=10090 GN=Dnajc5 PE=1 SV=1 | 0 | 144.981 | 62 | 13 | 70 | 1 | 198 | 22.1 | 5.07 | 0.279 | 0.239 | 1.168 | -1.84 | 1.84 | -2.06 | -0.6866667 | 1.95 | 0.22 | 4.70223E-07 | 7.73225E-06 | 0.32470606 | 3.46913E-05 | 0.000850093 | 0.000442392 | 0.905686588 |
| Rgs20 | High | E9Q720 | Regulator of G-protein-signaling 20 OS=Mus musculus OX=10090 GN=Rgs20 PE=1 SV=1 | 0 | 11.789 | 18 | 5 | 7 | 5 | 372 | 41.6 | 5.26 | 0.251 | 0.257 | 0.977 | -2 | 2 | -1.96 | -0.6533333 | 1.98 | -0.03 | 9.75645E-07 | 2.07899E-05 | 0.956129403 | 7.10447E-05 | 0.001596838 | 0.000833941 | 0.992902921 |
| Nphp1 | High | Q3TWM5 | Nephrocystin-1 OS=Mus musculus OX=10090 GN=Nphp1 PE=1 SV=1 | 0 | 18.804 | 13 | 8 | 9 | 8 | 690 | 77.3 | 5.17 | 4.05 | 4.659 | 0.869 | 2.02 | -2.02 | 2.22 | 0.74 | -2.12 | -0.2 | 1.06065E-06 | 3.58607E-06 | 0.428359826 | 7.43381E-05 | 0.00042781 | 0.000251074 | 0.943389567 |
| Tctn2 | High | Q2MV57 | Tectonic-2 OS=Mus musculus OX=10090 GN=Tctn2 PE=1 SV=2 | 0 | 15.89 | 8 | 4 | 5 | 4 | 700 | 77.1 | 5.66 | 3.49 | 3.668 | 0.952 | 1.8 | -1.8 | 1.88 | 0.62666667 | -1.84 | -0.07 | 1.33673E-06 | 6.87714E-05 | 0.791202565 | 9.25312E-05 | 0.004303888 | 0.002198209 | 0.992535481 |
| Hcn1 | High | O88704 | Potassium/sodium hyperpolarization-activated cyclic nucleotide-gated channel 1 OS=Mus muscul | 0 | 108.848 | 21 | 19 | 33 | 17 | 910 | 102.4 | 8.37 | 0.243 | 0.246 | 0.986 | -2.04 | 2.04 | -2.02 | -0.6733333 | 2.03 | -0.02 | 1.90193E-06 | 1.14707E-05 | 0.994958662 | 0.00013005 | 0.001148507 | 0.000639279 | 0.999917596 |
| Slc12a6 | High | Q924N4 | Solute carrier family 12 member 6 OS=Mus musculus OX=10090 GN=Slc12a6 PE=1 SV=2 | 0 | 51.817 | 9 | 12 | 20 | 7 | 1150 | 127.4 | 7.03 | 0.266 | 0.301 | 0.883 | -1.91 | 1.91 | -1.73 | -0.5766667 | 1.82 | -0.18 | 4.30615E-06 | 0.000164557 | 0.488082366 | 0.000284054 | 0.00854324 | 0.004413647 | 0.964378027 |
| Mdm1 | High | B2RPU6 | Nuclear protein MDM1 OS=Mus musculus OX=10090 GN=Mdm1 PE=1 SV=1 | 0 | 286.823 | 59 | 44 | 104 | 44 | 718 | 80.8 | 9.44 | 3.864 | 3.65 | 1.059 | 1.95 | -1.95 | 1.87 | 0.62333333 | -1.91 | 0.08 | 5.33817E-06 | 0.000102332 | 0.640053383 | 0.000348036 | 0.005976805 | 0.00316242 | 0.983766261 |
| Ofd1 | High | A2AEG2 | Oral-facial-digital syndrome 1 protein homolog OS=Mus musculus OX=10090 GN=Ofd1 PE=1 SV= | 0 | 9.098 | 4 | 3 | 3 | 3 | 1019 | 117.6 | 5.88 | 3.683 | 3.685 | 0.999 | 1.88 | -1.88 | 1.88 | 0.62666667 | -1.88 | 0 | 6.59403E-06 | 6.98509E-05 | 0.960733134 | 0.000420145 | 0.004303888 | 0.002362016 | 0.99314725 |
| Nbr1 | High | K3W4P1 | Next to BRCA1 gene 1 protein OS=Mus musculus OX=10090 GN=Nbr1 PE=1 SV=1 | 0 | 33.787 | 9 | 8 | 9 | 8 | 988 | 110 | 5.1 | 0.266 | 0.211 | 1.258 | -1.91 | 1.91 | -2.24 | -0.7466667 | 2.075 | 0.33 | 6.83305E-06 | 1.21054E-06 | 0.186685496 | 0.000421021 | 0.000169688 | 0.000295354 | 0.77374646 |
| Pla2r1 | Low | Q62028 | Secretory phospholipase A2 receptor OS=Mus musculus OX=10090 GN=Pla2r1 PE=1 SV=1 | 0.052 | 1.441 | 1 | 1 | 1 | 1 | 1487 | 170.4 | 6.62 | 0.337 | 0.253 | 1.333 | -1.57 | 1.57 | -1.98 | -0.66 | 1.775 | 0.41 | 8.39939E-06 | 1.2077E-05 | 0.141839128 | 0.000511906 | 0.001167516 | 0.000839711 | 0.701851981 |
| Spice1 | High | Q8C804 | Spindle and centriole-associated protein 1 OS=Mus musculus OX=10090 GN=Spice1 PE=1 SV=2 | 0 | 21.238 | 9 | 6 | 6 | 6 | 860 | 95.6 | 7.05 | 3.243 | 3.544 | 0.915 | 1.7 | -1.7 | 1.83 | 0.61 | -1.765 | -0.13 | 8.99037E-06 | 0.000147683 | 0.580407667 | 0.000542032 | 0.008039402 | 0.004290717 | 0.972910028 |
| Fgfr1op | High | Q66JX5 | Centrosomal protein 43 OS=Mus musculus OX=10090 GN=Cep43 PE=1 SV=1 | 0.001 | 6.118 | 5 | 2 | 3 | 2 | 399 | 42.7 | 4.81 | 3.291 | 3.698 | 0.89 | 1.72 | -1.72 | 1.89 | 0.63 | -1.805 | -0.17 | 1.01219E-05 | 5.94969E-05 | 0.525089488 | 0.000603761 | 0.003790898 | 0.002197329 | 0.964378027 |
| Ssna1 | Medium | Q9JJ94 | Sjoegren syndrome nuclear autoantigen 1 homolog OS=Mus musculus OX=10090 GN=Ssna1 PE=1 | 0.035 | 1.81 | 6 | 1 | 1 | 1 | 119 | 13.5 | 5.68 | 3.108 | 4.204 | 0.739 | 1.64 | -1.64 | 2.07 | 0.69 | -1.855 | -0.44 | 1.2178E-05 | 8.71593E-06 | 0.10324602 | 0.000689758 | 0.000922079 | 0.000805919 | 0.611945493 |
| Cep120 | High | Q7TSG1 | Centrosomal protein of 120 kDa OS=Mus musculus OX=10090 GN=Cep120 PE=1 SV=1 | 0 | 29.944 | 9 | 9 | 11 | 9 | 988 | 112.5 | 5.94 | 3.581 | 4.072 | 0.879 | 1.84 | -1.84 | 2.03 | 0.67666667 | -1.935 | -0.19 | 1.21632E-05 | 2.43731E-05 | 0.462665916 | 0.000689758 | 0.001846755 | 0.001268256 | 0.959744396 |
| Stx12 | High | Q9ER00 | Syntaxin-12 OS=Mus musculus OX=10090 GN=Stx12 PE=1 SV=1 | 0 | 52.251 | 38 | 9 | 17 | 9 | 274 | 31.2 | 5.44 | 0.281 | 0.243 | 1.156 | -1.83 | 1.83 | -2.04 | -0.68 | 1.935 | 0.21 | 1.35587E-05 | 9.69097E-06 | 0.419527068 | 0.000760237 | 0.001006246 | 0.000883241 | 0.937638605 |
| Poc5 | High | Q9DBS8 | Centrosomal protein POC5 OS=Mus musculus OX=10090 GN=Poc5 PE=2 SV=1 | 0 | 99.572 | 38 | 18 | 25 | 18 | 558 | 60.9 | 8.09 | 3.631 | 3.904 | 0.93 | 1.86 | -1.86 | 1.96 | 0.65333333 | -1.91 | -0.1 | 1.41236E-05 | 4.29216E-05 | 0.714302011 | 0.000784072 | 0.002865019 | 0.001824546 | 0.992535481 |
| Acacb | High | E9Q4Z2 | Acetyl-CoA carboxylase 2 OS=Mus musculus OX=10090 GN=Acacb PE=1 SV=1 | 0 | 19.999 | 3 | 6 | 7 | 1 | 2448 | 275.6 | 6.32 | 0.326 | 0.306 | 1.063 | -1.62 | 1.62 | -1.71 | -0.57 | 1.665 | 0.09 | 1.93853E-05 | 0.000140245 | 0.747234019 | 0.001065622 | 0.007709352 | 0.004387487 | 0.992535481 |
| Wbp2 | High | P97765 | WW domain-binding protein 2 OS=Mus musculus OX=10090 GN=Wbp2 PE=1 SV=1 | 0 | 66.087 | 36 | 8 | 30 | 8 | 261 | 28 | 6.33 | 0.282 | 0.234 | 1.207 | -1.83 | 1.83 | -2.1 | -0.7 | 1.965 | 0.27 | 2.10664E-05 | 5.56613E-06 | 0.189809705 | 0.001146791 | 0.000645353 | 0.000896072 | 0.77990201 |
| Aqp1 | High | Q02013 | Aquaporin-1 OS=Mus musculus OX=10090 GN=Aqp1 PE=1 SV=3 | 0 | 74.063 | 32 | 6 | 33 | 6 | 269 | 28.8 | 7.43 | 0.283 | 0.261 | 1.084 | -1.82 | 1.82 | -1.94 | -0.6466667 | 1.88 | 0.12 | 2.15335E-05 | 2.56752E-05 | 0.604699106 | 0.001160947 | 0.001896827 | 0.001528887 | 0.97826833 |
| Lca5 | High | G5E886 | Lebercilin OS=Mus musculus OX=10090 GN=Lca5 PE=1 SV=1 | 0 | 51.78 | 23 | 14 | 17 | 14 | 704 | 80.1 | 8.16 | 3.437 | 2.67 | 1.287 | 1.78 | -1.78 | 1.42 | 0.47333333 | -1.6 | 0.36 | 2.37631E-05 | 0.003435167 | 0.083952101 | 0.001245229 | 0.101373593 | 0.051309411 | 0.558386036 |
| Rtbdn | Medium | Q8QZY4 | Retbindin OS=Mus musculus OX=10090 GN=Rtbdn PE=1 SV=1 | 0.016 | 2.162 | 4 | 1 | 1 | 1 | 247 | 26.5 | 7.59 | 0.355 | 0.523 | 0.679 | -1.5 | 1.5 | -0.94 | -0.3133333 | 1.22 | -0.56 | 2.36992E-05 | 0.03786587 | 0.021653001 | 0.001245229 | 0.485844242 | 0.243544736 | 0.266831602 |
| Sirpa | High | P97797 | Tyrosine-protein phosphatase non-receptor type substrate 1 OS=Mus musculus OX=10090 GN=Sir | 0 | 43.905 | 23 | 6 | 9 | 3 | 513 | 56.4 | 8.28 | 0.318 | 0.395 | 0.803 | -1.66 | 1.66 | -1.34 | -0.4466667 | 1.5 | -0.32 | 2.45612E-05 | 0.003374575 | 0.22562389 | 0.001275134 | 0.100778647 | 0.05102689 | 0.813917005 |
| Plpp2 | High | Q9DAX2 | Phospholipid phosphatase 2 OS=Mus musculus OX=10090 GN=Plpp2 PE=1 SV=1 | 0 | 8.696 | 11 | 3 | 3 | 3 | 276 | 31.2 | 8.5 | 0.333 | 0.305 | 1.092 | -1.59 | 1.59 | -1.71 | -0.57 | 1.65 | 0.13 | 2.56382E-05 | 0.000117235 | 0.603664361 | 0.00131884 | 0.006639779 | 0.003979309 | 0.97826833 |
| Rp1 | High | P56716 | Oxygen-regulated protein 1 OS=Mus musculus OX=10090 GN=Rp1 PE=1 SV=2 | 0 | 842.295 | 58 | 113 | 311 | 113 | 2095 | 234.2 | 7.53 | 3.487 | 3.471 | 1.004 | 1.8 | -1.8 | 1.8 | 0.6 | -1.8 | 0.01 | 2.60691E-05 | 0.000190551 | 0.897688621 | 0.001328811 | 0.009712895 | 0.005520853 | 0.992535481 |
| Slc30a1 | High | Q60738 | Zinc transporter 1 OS=Mus musculus OX=10090 GN=Slc30a1 PE=1 SV=1 | 0 | 31.083 | 18 | 7 | 11 | 7 | 503 | 54.7 | 6.62 | 0.305 | 0.311 | 0.981 | -1.71 | 1.71 | -1.68 | -0.56 | 1.695 | -0.03 | 3.54271E-05 | 0.000248545 | 0.900960063 | 0.001757873 | 0.012115463 | 0.006936668 | 0.992535481 |
| Crocc | High | Q8CJ40 | Rootletin OS=Mus musculus OX=10090 GN=Crocc PE=1 SV=2 | 0 | 595.741 | 51 | 105 | 226 | 104 | 2009 | 226.8 | 5.55 | 3.407 | 4.336 | 0.786 | 1.77 | -1.77 | 2.12 | 0.70666667 | -1.945 | -0.35 | 3.67155E-05 | 1.01636E-05 | 0.145686911 | 0.001805823 | 0.00103613 | 0.001420976 | 0.706909771 |
| Tspan7 | High | Q62283 | Tetraspanin-7 OS=Mus musculus OX=10090 GN=Tspan7 PE=1 SV=2 | 0 | 16.598 | 10 | 2 | 4 | 2 | 249 | 27.5 | 7.2 | 0.371 | 0.389 | 0.954 | -1.43 | 1.43 | -1.36 | -0.4533333 | 1.395 | -0.07 | 3.86251E-05 | 0.001929952 | 0.79375304 | 0.001883228 | 0.06592534 | 0.033904284 | 0.992535481 |
| Clu | High | Q06890 | Clusterin OS=Mus musculus OX=10090 GN=Clu PE=1 SV=1 | 0 | 81.74 | 35 | 15 | 30 | 15 | 448 | 51.6 | 5.67 | 0.297 | 0.343 | 0.865 | -1.75 | 1.75 | -1.54 | -0.5133333 | 1.645 | -0.21 | 4.43624E-05 | 0.000762892 | 0.420317453 | 0.002144308 | 0.029500247 | 0.015822278 | 0.938558327 |
| Ano2 | Medium | Q8CFW1 | Anoctamin-2 OS=Mus musculus OX=10090 GN=Ano2 PE=1 SV=2 | 0.015 | 2.176 | 1 | 1 | 1 | 1 | 1002 | 114.1 | 7.36 | 0.319 | 0.503 | 0.635 | -1.65 | 1.65 | -0.99 | -0.33 | 1.32 | -0.65 | 4.57663E-05 | 0.025564288 | 0.038152539 | 0.00219326 | 0.399272881 | 0.200733071 | 0.373579725 |
| Drd4 | Medium | P51436 | D(4) dopamine receptor OS=Mus musculus OX=10090 GN=Drd4 PE=1 SV=1 | 0.01 | 2.663 | 10 | 2 | 2 | 2 | 387 | 41.5 | 8.63 | 0.355 | 0.469 | 0.758 | -1.49 | 1.49 | -1.09 | -0.3633333 | 1.29 | -0.4 | 4.90467E-05 | 0.013254647 | 0.18694996 | 0.00233055 | 0.267445207 | 0.134887878 | 0.77374646 |
| Wdr13 | High | S4R225 | WD repeat-containing protein 13 OS=Mus musculus OX=10090 GN=Wdr13 PE=1 SV=1 | 0.002 | 4.183 | 6 | 2 | 2 | 2 | 393 | 43 | 7.46 | 0.344 | 0.303 | 1.134 | -1.54 | 1.54 | -1.72 | -0.5733333 | 1.63 | 0.18 | 5.10492E-05 | 0.000106747 | 0.460618879 | 0.002385276 | 0.006107476 | 0.004246376 | 0.959744396 |
| Advgr1 | High | Q8VHN7 | Adhesion G-protein coupled receptor V1 OS=Mus musculus OX=10090 GN=Adgrv1 PE=1 SV=1 | 0.006 | 3.327 | 0 | 1 | 1 | 1 | 6298 | 687 | 4.75 | 2.734 | 3.132 | 0.873 | 1.45 | -1.45 | 1.65 | 0.55 | -1.55 | -0.2 | 5.91835E-05 | 0.000505845 | 0.417846795 | 0.002697903 | 0.021650944 | 0.012174423 | 0.936397673 |
| Ubap1l-fr | High | A0A6I8MV | Ubiquitin-associated protein 1-like (Fragment) OS=Mus musculus OX=10090 GN=Ubap1l PE=4 SV | 0 | 26.094 | 37 | 6 | 8 | 2 | 243 | 27 | 7.77 | 0.338 | 0.275 | 1.226 | -1.57 | 1.57 | -1.86 | -0.62 | 1.715 | 0.29 | 6.74815E-05 | 5.31967E-05 | 0.243319803 | 0.003051361 | 0.003428434 | 0.003239898 | 0.837310213 |
| Itm2b | High | O89051 | Integral membrane protein 2B OS=Mus musculus OX=10090 GN=Itm2b PE=1 SV=1 | 0.001 | 6.233 | 3 | 1 | 2 | 1 | 266 | 30.2 | 5.3 | 0.377 | 0.486 | 0.775 | -1.41 | 1.41 | -1.04 | -0.3466667 | 1.225 | -0.37 | 6.91918E-05 | 0.016938245 | 0.145996347 | 0.003079033 | 0.305844112 | 0.154461573 | 0.706909771 |
| Cep350 | High | E9Q309 | Centrosome-associated protein 350 OS=Mus musculus OX=10090 GN=Cep350 PE=1 SV=1 | 0 | 15.764 | 2 | 5 | 5 | 5 | 3095 | 346.2 | 6.35 | 2.622 | 2.743 | 0.956 | 1.39 | -1.39 | 1.46 | 0.48666667 | -1.425 | -0.07 | 7.2573E-05 | 0.001751676 | 0.846098826 | 0.003204071 | 0.061771379 | 0.032487725 | 0.992535481 |
| Zfp365 | High | Q8BG89 | Protein ZNF365 OS=Mus musculus OX=10090 GN=Znf365 PE=1 SV=1 | 0.006 | 3.152 | 2 | 1 | 2 | 1 | 408 | 46.8 | 9.14 | 2.615 | 2.531 | 1.033 | 1.39 | -1.39 | 1.34 | 0.44666667 | -1.365 | 0.05 | 8.32135E-05 | 0.004721866 | 0.798185863 | 0.003645142 | 0.128816384 | 0.066230763 | 0.992535481 |
| Rp1l1 | High | Q8CGM2 | Retinitis pigmentosa 1-like 1 protein OS=Mus musculus OX=10090 GN=Rp1l1 PE=1 SV=1 | 0 | 715.996 | 63 | 95 | 254 | 95 | 1859 | 199.6 | 6.48 | 3.169 | 3.175 | 0.998 | 1.66 | -1.66 | 1.67 | 0.55666667 | -1.665 | 0 | 0.000102853 | 0.000544731 | 0.930447589 | 0.004470533 | 0.022933836 | 0.013702184 | 0.992535481 |
| Nr2e3 | High | Q9QXZ7 | Photoreceptor-specific nuclear receptor OS=Mus musculus OX=10090 GN=Nr2e3 PE=1 SV=1 | 0 | 4.728 | 7 | 2 | 2 | 2 | 395 | 43.1 | 7.74 | 2.698 | 2.062 | 1.309 | 1.43 | -1.43 | 1.04 | 0.34666667 | -1.235 | 0.39 | 0.000117224 | 0.029448095 | 0.095112418 | 0.005055976 | 0.436813413 | 0.220934694 | 0.586683527 |
| Atp1b2 | High | P14231 | Sodium/potassium-transporting ATPase subunit beta-2 OS=Mus musculus OX=10090 GN=Atp1b2 | 0 | 159.368 | 41 | 14 | 79 | 14 | 290 | 33.3 | 8.31 | 0.321 | 0.397 | 0.81 | -1.64 | 1.64 | -1.33 | -0.4433333 | 1.485 | -0.3 | 0.000136283 | 0.003503504 | 0.208310657 | 0.005833119 | 0.102019919 | 0.053926519 | 0.801370196 |
| Vps37a | High | Q8CHS8 | Vacuolar protein sorting-associated protein 37A OS=Mus musculus OX=10090 GN=Vps37a PE=1 S | 0 | 43.061 | 24 | 8 | 10 | 8 | 397 | 44.4 | 5.67 | 0.32 | 0.256 | 1.25 | -1.65 | 1.65 | -1.97 | -0.6566667 | 1.81 | 0.32 | 0.000140982 | 1.96488E-05 | 0.182616763 | 0.005988534 | 0.001530147 | 0.003759341 | 0.765268049 |
| Snap25 | High | P60879 | Synaptosomal-associated protein 25 OS=Mus musculus OX=10090 GN=Snap25 PE=1 SV=1 | 0 | 212.817 | 76 | 19 | 98 | 18 | 206 | 23.3 | 4.77 | 0.325 | 0.373 | 0.87 | -1.62 | 1.62 | -1.42 | -0.4733333 | 1.52 | -0.2 | 0.000155347 | 0.001874046 | 0.425280234 | 0.006549084 | 0.064464881 | 0.035506982 | 0.941180024 |
| Atp4a | High | E9QNX7 | Sodium/potassium-transporting ATPase subunit alpha OS=Mus musculus OX=10090 GN=Atp4a PE | 0 | 50.587 | 6 | 8 | 44 | 1 | 1034 | 114 | 5.81 | 0.335 | 0.455 | 0.735 | -1.58 | 1.58 | -1.14 | -0.38 | 1.36 | -0.44 | 0.000167517 | 0.012419668 | 0.071384669 | 0.006890831 | 0.256963386 | 0.131927109 | 0.508467979 |
| Ick | High | Q9JKV2 | Serine/threonine-protein kinase ICK OS=Mus musculus OX=10090 GN=Cilk1 PE=1 SV=2 | 0 | 35.18 | 18 | 10 | 14 | 8 | 629 | 70.5 | 9.82 | 2.901 | 3.068 | 0.945 | 1.54 | -1.54 | 1.62 | 0.54 | -1.58 | -0.08 | 0.000194987 | 0.00079966 | 0.793022412 | 0.007865396 | 0.030295245 | 0.01908032 | 0.992535481 |
| Rpgrip1 | High | E9QQ97 | X-linked retinitis pigmentosa GTPase regulator-interacting protein 1 OS=Mus musculus OX=10090 | 0 | 42.768 | 12 | 13 | 14 | 13 | 1127 | 128.7 | 5.2 | 2.986 | 3.986 | 0.749 | 1.58 | -1.58 | 1.99 | 0.66333333 | -1.785 | -0.42 | 0.000226617 | 3.25238E-05 | 0.091027158 | 0.008948199 | 0.002197119 | 0.005572659 | 0.576060131 |
| Ccdc8 | Low | D3YZV8 | Coiled-coil domain-containing protein 8 homolog OS=Mus musculus OX=10090 GN=Ccdc8 PE=1 S\ | 0.056 | 1.36 | 1 | 1 | 1 | 1 | 685 | 73.9 | 9.09 | 2.469 | 1.97 | 1.253 | 1.3 | -1.3 | 0.98 | 0.32666667 | -1.14 | 0.33 | 0.000243453 | 0.042339698 | 0.141963225 | 0.009545747 | 0.510534809 | 0.260040278 | 0.701851981 |
| Cd47 | High | A0A2R8W | Integrin-associated protein OS=Mus musculus OX=10090 GN=Cd47 PE=1 SV=1 | 0 | 125.539 | 23 | 9 | 64 | 9 | 342 | 37.3 | 7.74 | 0.336 | 0.311 | 1.079 | -1.57 | 1.57 | -1.68 | -0.56 | 1.625 | 0.11 | 0.000247637 | 0.000249102 | 0.554208504 | 0.009642379 | 0.012115463 | 0.010878921 | 0.96775057 |
| Plekhb1 | High | D3Z6S5 | Pleckstrin homology domain-containing family B member 1 (Fragment) OS=Mus musculus OX=10 | 0 | 64.998 | 56 | 9 | 31 | 1 | 149 | 17.2 | 6.19 | 0.397 | 0.564 | 0.704 | -1.33 | 1.33 | -0.83 | -0.2766667 | 1.08 | -0.51 | 0.000252307 | 0.065661776 | 0.044032701 | 0.009756443 | 0.592044692 | 0.300900568 | 0.405968973 |
| Hmgn2 | High | A3KGL9 | Non-histone chromosomal protein HMG-17 OS=Mus musculus OX=10090 GN=Hmgn2 PE=1 SV=1 | 0 | 12.098 | 42 | 3 | 6 | 1 | 96 | 9.7 | 9.73 | 2.554 | 1.656 | 1.542 | 1.35 | -1.35 | 0.73 | 0.24333333 | -1.04 | 0.62 | 0.000281288 | 0.143758028 | 0.012232417 | 0.010585122 | 0.837891129 | 0.424238125 | 0.183058049 |
| Spry2 | High | Q9QXV8 | Protein sprouty homolog 2 OS=Mus musculus OX=10090 GN=Spry2 PE=1 SV=1 | 0 | 12.977 | 16 | 5 | 5 | 5 | 315 | 34.6 | 8.29 | 0.393 | 0.373 | 1.052 | -1.35 | 1.35 | -1.42 | -0.4733333 | 1.385 | 0.07 | 0.000288771 | 0.001870072 | 0.738952453 | 0.01079427 | 0.064464881 | 0.037629575 | 0.992535481 |
| Bub1b | High | Q9Z1S0 | Mitotic checkpoint serine/threonine-protein kinase BUB1 beta OS=Mus musculus OX=10090 GN=I | 0.001 | 4.543 | 1 | 1 | 1 | 1 | 1052 | 118.3 | 5.43 | 0.401 | 0.361 | 1.109 | -1.32 | 1.32 | -1.47 | -0.49 | 1.395 | 0.15 | 0.000291334 | 0.000971751 | 0.570969349 | 0.010817948 | 0.03656784 | 0.023692894 | 0.972910028 |
| Hsbp1 | High | Q9CQZ1 | Heat shock factor-binding protein 1 OS=Mus musculus OX=10090 GN=Hsbp1 PE=1 SV=1 | 0 | 29.011 | 82 | 4 | 9 | 4 | 76 | 8.6 | 4.31 | 2.501 | 3.097 | 0.807 | 1.32 | -1.32 | 1.63 | 0.54333333 | -1.475 | -0.31 | 0.000294659 | 0.00072046 | 0.181111862 | 0.010869427 | 0.028249095 | 0.019559261 | 0.759531947 |
| Cfap20 | High | Q8BTU1 | Cilia- and flagella-associated protein 20 OS=Mus musculus OX=10090 GN=Cfap20 PE=1 SV=1 | 0 | 11.628 | 22 | 4 | 6 | 4 | 193 | 22.7 | 9.76 | 2.54 | 1.869 | 1.359 | 1.34 | -1.34 | 0.9 | 0.3 | -1.12 | 0.44 | 0.00029804 | 0.061825839 | 0.056767104 | 0.010922279 | 0.586560875 | 0.298741577 | 0.457975756 |
| Ccdc181 | High | Q80ZU5 | Coiled-coil domain-containing protein 181 OS=Mus musculus OX=10090 GN=Ccdc181 PE=1 SV=1 | 0 | 38.751 | 19 | 10 | 14 | 10 | 509 | 59.2 | 5.33 | 2.762 | 2.668 | 1.035 | 1.47 | -1.47 | 1.42 | 0.47333333 | -1.445 | 0.05 | 0.000322352 | 0.003453949 | 0.747495712 | 0.011654724 | 0.101394204 | 0.056524464 | 0.992535481 |
| Traf2 | High | P39429 | TNF receptor-associated factor 2 OS=Mus musculus OX=10090 GN=Traf2 PE=1 SV=1 | 0 | 21.056 | 12 | 6 | 10 | 6 | 501 | 56 | 6.96 | 0.381 | 0.457 | 0.835 | -1.39 | 1.39 | -1.13 | -0.3766667 | 1.26 | -0.26 | 0.000325241 | 0.012804971 | 0.303347853 | 0.011654724 | 0.262579646 | 0.137117185 | 0.900241969 |
| B9d1 | High | Q9R1S0 | B9 domain-containing protein 1 OS=Mus musculus OX=10090 GN=B9d1 PE=1 SV=1 | 0 | 9.582 | 15 | 3 | 3 | 3 | 204 | 22.6 | 6.57 | 2.661 | 3.346 | 0.795 | 1.41 | -1.41 | 1.74 | 0.58 | -1.575 | -0.33 | 0.000355689 | 0.000296354 | 0.190002879 | 0.012622448 | 0.013620158 | 0.013121303 | 0.77990201 |
| Dnajc5 | High | G5E8T0 | DnaJ homolog subfamily C member 5 OS=Mus musculus OX=10090 GN=Dnajc5 PE=1 SV=1 | 0 | 123.327 | 54 | 13 | 52 | 1 | 167 | 18.7 | 5.59 | 0.349 | 0.324 | 1.078 | -1.52 | 1.52 | -1.63 | -0.5433333 | 1.575 | 0.11 | 0.000404679 | 0.000396318 | 0.560234369 | 0.01387814 | 0.017636156 | 0.015757148 | 0.972910028 |
| Fam161a- | High | A0A140LH | Protein FAM161A (Fragment) OS=Mus musculus OX=10090 GN=Fam161a PE=1 SV=1 | 0 | 15.378 | 9 | 4 | 4 | 4 | 562 | 65.3 | 9.33 | 2.747 | 2.77 | 0.992 | 1.46 | -1.46 | 1.47 | 0.49 | -1.465 | -0.01 | 0.000410874 | 0.002372723 | 0.97164228 | 0.01387814 | 0.078257978 | 0.046068059 | 0.996524285 |
| Sdcbp-Fr | High | H3BLG5 | Syntenin-1 (Fragment) OS=Mus musculus OX=10090 GN=Sdcbp PE=1 SV=1 | 0.003 | 3.859 | 7 | 2 | 2 | 2 | 249 | 26.8 | 7.42 | 0.397 | 0.402 | 0.986 | -1.33 | 1.33 | -1.31 | -0.4366667 | 1.32 | -0.02 | 0.000408184 | 0.00331302 | 0.932270175 | 0.01387814 | 0.100411377 | 0.057144759 | 0.992535481 |
| Abcg4 | High | Q91WA9 | ATP-binding cassette subfamily G member 4 OS=Mus musculus OX=10090 GN=Abcg4 PE=1 SV=2 | 0 | 8.145 | 7 | 3 | 3 | 3 | 646 | 72.1 | 8.19 | 0.406 | 0.405 | 1.002 | -1.3 | 1.3 | -1.3 | -0.4333333 | 1.3 | 0 | 0.000409589 | 0.00424085 | 0.919503787 | 0.01387814 | 0.120702776 | 0.067290458 | 0.992535481 |
| Ift43 | High | Q9DA69 | Intraflagellar transport protein 43 homolog OS=Mus musculus OX=10090 GN=Ift43 PE=1 SV=2 | 0 | 15.69 | 12 | 2 | 3 | 2 | 206 | 23.5 | 4.69 | 2.53 | 2.115 | 1.196 | 1.34 | -1.34 | 1.08 | 0.36 | -1.21 | 0.26 | 0.000448926 | 0.022683144 | 0.257219066 | 0.015072639 | 0.378524969 | 0.196798804 | 0.85616924 |
| Eif2s1 | High | Q6ZWX6 | Eukaryotic translation initiation factor 2 subunit 1 OS=Mus musculus OX=10090 GN=Eif2s1 PE=1 S | 0 | 4.932 | 6 | 2 | 2 | 2 | 315 | 36.1 | 5.08 | 0.416 | 0.333 | 1.247 | -1.27 | 1.27 | -1.59 | -0.53 | 1.43 | 0.32 | 0.00046775 | 0.000548089 | 0.161831177 | 0.015548009 | 0.022933836 | 0.019240922 | 0.733684585 |
| Wdr19 | High | Q3UGF1 | WD repeat-containing protein 19 OS=Mus musculus OX=10090 GN=Wdr19 PE=1 SV=1 | 0 | 17.622 | 4 | 6 | 6 | 6 | 1341 | 151.4 | 6.57 | 2.772 | 2.297 | 1.207 | 1.47 | -1.47 | 1.2 | 0.4 | -1.335 | 0.27 | 0.000468631 | 0.013725489 | 0.238786091 | 0.015548009 | 0.270981748 | 0.143264878 | 0.830051837 |
| Sema4b | High | Q62179 | Semaphorin-4B OS=Mus musculus OX=10090 GN=Sema4b PE=1 SV=2 | 0 | 11.356 | 4 | 4 | 4 | 4 | 823 | 91.3 | 8.15 | 0.379 | 0.389 | 0.976 | -1.4 | 1.4 | -1.36 | -0.4533333 | 1.38 | -0.04 | 0.000475004 | 0.002851805 | 0.923587943 | 0.015666741 | 0.090339391 | 0.053003066 | 0.992535481 |
| Gnb3 | High | Q61011 | Guanine nucleotide-binding protein G(I)/G(S)/G(T) subunit beta-3 OS=Mus musculus OX=10090 G | 0 | 202.869 | 82 | 18 | 188 | 15 | 340 | 37.2 | 5.69 | 0.359 | 0.399 | 0.9 | -1.48 | 1.48 | -1.33 | -0.4433333 | 1.405 | -0.15 | 0.000572703 | 0.003650615 | 0.565031412 | 0.018778643 | 0.104969226 | 0.061873934 | 0.972910028 |
| Mks1 | High | Q5SW45 | Meckel syndrome type 1 protein homolog OS=Mus musculus OX=10090 GN=Mks1 PE=1 SV=1 | 0 | 10.959 | 6 | 2 | 3 | 2 | 561 | 64.4 | 6.25 | 2.347 | 2.667 | 0.88 | 1.23 | -1.23 | 1.42 | 0.47333333 | -1.325 | -0.18 | 0.000663114 | 0.002492296 | 0.460571031 | 0.021368278 | 0.080983898 | 0.051176088 | 0.959744396 |
| Ift88 | High | Q61371 | Intraflagellar transport protein 88 homolog OS=Mus musculus OX=10090 GN=Ift88 PE=1 SV=2 | 0 | 19.876 | 7 | 5 | 5 | 5 | 824 | 92.9 | 5.9 | 2.433 | 2.05 | 1.187 | 1.28 | -1.28 | 1.04 | 0.34666667 | -1.16 | 0.25 | 0.000736481 | 0.034362685 | 0.31689406 | 0.023462783 | 0.476392832 | 0.249927808 | 0.904943958 |
| Tchp | High | Q3TVW5 | Trichoplein keratin filament-binding protein OS=Mus musculus OX=10090 GN=Tchp PE=1 SV=2 | 0 | 53.168 | 28 | 13 | 17 | 13 | 497 | 60.6 | 6.8 | 2.733 | 2.675 | 1.022 | 1.45 | -1.45 | 1.42 | 0.47333333 | -1.435 | 0.03 | 0.00076744 | 0.003371157 | 0.821641294 | 0.02431093 | 0.100778647 | 0.062544789 | 0.992535481 |
| Cep41 | High | Q99NF3 | Centrosomal protein of 41 kDa OS=Mus musculus OX=10090 GN=Cep41 PE=1 SV=1 | 0 | 19.839 | 23 | 6 | 9 | 6 | 373 | 41.4 | 8.12 | 2.351 | 2.497 | 0.942 | 1.23 | -1.23 | 1.32 | 0.44 | -1.275 | -0.09 | 0.000776925 | 0.006517837 | 0.68036163 | 0.024473142 | 0.167639951 | 0.096056547 | 0.991285312 |
| Ptprf | High | A2A8L5 | Receptor-type tyrosine-protein phosphatase F OS=Mus musculus OX=10090 GN=Ptprf PE=1 SV=1 | 0 | 13.683 | 2 | 4 | 4 | 4 | 1898 | 211.4 | 6.65 | 0.468 | 0.439 | 1.066 | -1.09 | 1.09 | -1.19 | -0.3966667 | 1.14 | 0.09 | 0.00090065 | 0.009101395 | 0.637947043 | 0.028055253 | 0.203312831 | 0.115684042 | 0.983597794 |
| Abhd8 | High | Q8R0P8 | Protein ABHD8 OS=Mus musculus OX=10090 GN=Abhd8 PE=2 SV=1 | 0 | 25.995 | 21 | 6 | 11 | 6 | 439 | 48.2 | 6.64 | 2.348 | 2.682 | 0.875 | 1.23 | -1.23 | 1.42 | 0.47333333 | -1.325 | -0.19 | 0.000915472 | 0.003282434 | 0.3928631 | 0.028359386 | 0.100411377 | 0.064385381 | 0.93064581 |
| Spry3 | High | Q3UUD2 | Protein sprouty homolog 3 OS=Mus musculus OX=10090 GN=Spry3 PE=1 SV=1 | 0 | 13.447 | 13 | 4 | 4 | 4 | 288 | 31.3 | 7.39 | 0.393 | 0.365 | 1.077 | -1.35 | 1.35 | -1.45 | -0.4833333 | 1.4 | 0.11 | 0.000939428 | 0.001489186 | 0.602217734 | 0.028941604 | 0.052847269 | 0.040894436 | 0.97826833 |
| Ift46 | High | A0A1L1SR | Intraflagellar transport protein 46 homolog (Fragment) OS=Mus musculus OX=10090 GN=Ift46 PE | 0 | 9.816 | 28 | 3 | 3 | 3 | 188 | 21.3 | 4.42 | 2.248 | 1.727 | 1.301 | 1.17 | -1.17 | 0.79 | 0.26333333 | -0.98 | 0.38 | 0.000987988 | 0.101257742 | 0.129290009 | 0.029944058 | 0.730697757 | 0.380320907 | 0.683250784 |
| Dync2li1 | High | Q8K0T2 | Cytoplasmic dynein 2 light intermediate chain 1 OS=Mus musculus OX=10090 GN=Dync2li1 PE=1 S | 0.001 | 7.32 | 7 | 3 | 3 | 3 | 351 | 39.4 | 8.09 | 2.247 | 1.864 | 1.205 | 1.17 | -1.17 | 0.9 | 0.3 | -1.035 | 0.27 | 0.001029142 | 0.062757894 | 0.222732602 | 0.031023664 | 0.587451609 | 0.309237637 | 0.808324725 |
| Tctn1 | High | Q8BZ64 | Tectonic-1 OS=Mus musculus OX=10090 GN=Tctn1 PE=1 SV=1 | 0 | 9.728 | 6 | 2 | 3 | 2 | 593 | 63.4 | 7.28 | 2.375 | 2.923 | 0.813 | 1.25 | -1.25 | 1.55 | 0.51666667 | -1.4 | -0.3 | 0.001078685 | 0.001126361 | 0.219608283 | 0.032171212 | 0.041549364 | 0.036860288 | 0.805859554 |
| Ncstn | High | P57716 | Nicastrin OS=Mus musculus OX=10090 GN=Ncstn PE=1 SV=3 | 0 | 34.656 | 11 | 5 | 7 | 5 | 708 | 78.4 | 6.09 | 0.394 | 0.488 | 0.807 | -1.34 | 1.34 | -1.03 | -0.3433333 | 1.185 | -0.31 | 0.001076624 | 0.02237903 | 0.208584514 | 0.032171212 | 0.374564835 | 0.203368023 | 0.801552719 |
| Cep290 | High | Q6A078 | Centrosomal protein of 290 kDa OS=Mus musculus OX=10090 GN=Cep290 PE=1 SV=2 | 0 | 136.316 | 18 | 43 | 52 | 43 | 2472 | 288.9 | 6.1 | 2.613 | 3.071 | 0.851 | 1.39 | -1.39 | 1.62 | 0.54 | -1.505 | -0.23 | 0.001216613 | 0.000791866 | 0.346780736 | 0.034795413 | 0.030204039 | 0.032499726 | 0.909022715 |

| | | | | | | | | | | | | | | | | | | | | | | | | | | | | |
|---|---|---|---|---|---|---|---|---|---|---|---|---|---|---|---|---|---|---|---|---|---|---|---|---|---|---|---|---|
| Slc38a2 | High | Q8CFE6 | Sodium-coupled neutral amino acid transporter 2 OS=Mus musculus OX=10090 GN=Slc38a2 PE=1 | 0.003 | 3.888 | 6 | 2 | 2 | 2 | 504 | 55.5 | 7.94 | 0.444 | 0.424 | 1.049 | -1.17 | 1.17 | -1.24 | -0.4133333 | 1.205 | 0.07 | 0.00121627 | 0.004617487 | 0.763984664 | 0.034795413 | 0.127538172 | 0.081166792 | 0.992535481 |
| Tmem87a | High | A2AQJ7 | Transmembrane protein 87A OS=Mus musculus OX=10090 GN=Tmem87a PE=1 SV=1 | 0 | 4.963 | 2 | 1 | 1 | 1 | 500 | 57 | 8.1 | 2.276 | 2.416 | 0.942 | 1.19 | -1.19 | 1.27 | 0.42333333 | -1.23 | -0.09 | 0.001234936 | 0.007457011 | 0.745912729 | 0.034795413 | 0.174943339 | 0.104869376 | 0.992535481 |
| Mga | Medium | E9QLG3 | MAX gene-associated protein OS=Mus musculus OX=10090 GN=Mga PE=1 SV=1 | 0.047 | 1.537 | 0 | 1 | 1 | 1 | 3042 | 332.9 | 7.58 | 0.46 | 0.402 | 1.142 | -1.12 | 1.12 | -1.31 | -0.4366667 | 1.215 | 0.19 | 0.001275335 | 0.003175556 | 0.446708166 | 0.035400011 | 0.098918583 | 0.067159297 | 0.958608248 |
| Cfap97 | High | Q6ZPR1 | Cilia- and flagella-associated protein 97 OS=Mus musculus OX=10090 GN=Cfap97 PE=1 SV=2 | 0 | 52.853 | 31 | 19 | 24 | 19 | 541 | 60.8 | 9.51 | 2.598 | 2.072 | 1.254 | 1.38 | -1.38 | 1.05 | 0.35 | -1.215 | 0.33 | 0.001300108 | 0.031613429 | 0.11934946 | 0.03573384 | 0.457149627 | 0.246441733 | 0.663223412 |
| Apoe | High | P08226 | Apolipoprotein E OS=Mus musculus OX=10090 GN=Apoe PE=1 SV=2 | 0 | 30.796 | 24 | 8 | 8 | 8 | 311 | 35.8 | 5.68 | 0.422 | 0.514 | 0.821 | -1.25 | 1.25 | -0.96 | -0.32 | 1.105 | -0.28 | 0.001296985 | 0.033172685 | 0.216615012 | 0.03573384 | 0.470474383 | 0.253104111 | 0.804947026 |
| Aplp2 | High | Q06335 | Amyloid-like protein 2 OS=Mus musculus OX=10090 GN=Aplp2 PE=1 SV=4 | 0 | 23.977 | 11 | 6 | 6 | 2 | 707 | 80.4 | 4.7 | 0.419 | 0.462 | 0.908 | -1.25 | 1.25 | -1.12 | -0.3733333 | 1.185 | -0.14 | 0.001329244 | 0.013996216 | 0.594197603 | 0.036356454 | 0.273438276 | 0.154897365 | 0.977451587 |
| Cdip1 | Medium | Q9DB75 | Cell death-inducing p53-target protein 1 OS=Mus musculus OX=10090 GN=Cdip1 PE=2 SV=1 | 0.029 | 1.924 | 3 | 1 | 1 | 1 | 208 | 21.8 | 5.66 | 0.434 | 0.409 | 1.062 | -1.2 | 1.2 | -1.29 | -0.43 | 1.245 | 0.09 | 0.001389184 | 0.003379059 | 0.711279657 | 0.037628764 | 0.100778647 | 0.069203706 | 0.992535481 |
| Plekhb2 | High | Q9QZC7 | Pleckstrin homology domain-containing family B member 2 OS=Mus musculus OX=10090 GN=Ple | 0 | 42.663 | 35 | 7 | 19 | 7 | 221 | 24.6 | 5.6 | 0.389 | 0.397 | 0.98 | -1.36 | 1.36 | -1.33 | -0.4433333 | 1.345 | -0.03 | 0.001501633 | 0.003511654 | 0.968328928 | 0.039859881 | 0.102019919 | 0.0709399 | 0.995075385 |
| Mia3 | Low | Q8BI84 | Transport and Golgi organization protein 1 homolog OS=Mus musculus OX=10090 GN=Mia3 PE=1 | 0.072 | 1.274 | 0 | 1 | 1 | 1 | 1930 | 213.5 | 4.75 | 0.471 | 0.57 | 0.827 | -1.09 | 1.09 | -0.81 | -0.27 | 0.95 | -0.27 | 0.001501225 | 0.058444719 | 0.31895099 | 0.039859881 | 0.571608376 | 0.305734129 | 0.904943958 |
| Ipo5 | High | Q8BKC5 | Importin-5 OS=Mus musculus OX=10090 GN=Ipo5 PE=1 SV=3 | 0 | 48.329 | 14 | 12 | 16 | 12 | 1097 | 123.5 | 4.93 | 0.404 | 0.447 | 0.905 | -1.31 | 1.31 | -1.16 | -0.3866667 | 1.235 | -0.14 | 0.001528038 | 0.010524427 | 0.542940818 | 0.040036012 | 0.230509622 | 0.135272817 | 0.96694628 |
| Csnk1a1 | High | E9Q2U6 | Casein kinase I isoform alpha OS=Mus musculus OX=10090 GN=Csnk1a1 PE=1 SV=1 | 0 | 49.044 | 25 | 9 | 17 | 1 | 353 | 40.6 | 9.51 | 2.291 | 1.904 | 1.203 | 1.2 | -1.2 | 0.93 | 0.31 | -1.065 | 0.27 | 0.001600481 | 0.052818092 | 0.242478658 | 0.041354376 | 0.552520601 | 0.296937488 | 0.836663283 |
| Prepl | Medium | Q8C167 | Prolyl endopeptidase-like OS=Mus musculus OX=10090 GN=Prepl PE=1 SV=1 | 0.011 | 2.384 | 1 | 1 | 1 | 1 | 725 | 83.1 | 6.65 | 2.298 | 2.234 | 1.029 | 1.2 | -1.2 | 1.16 | 0.38666667 | -1.18 | 0.04 | 0.00166808 | 0.014225771 | 0.800409475 | 0.042513286 | 0.275999641 | 0.159256464 | 0.992535481 |
| Tubg1 | Medium | P83887 | Tubulin gamma-1 chain OS=Mus musculus OX=10090 GN=Tubg1 PE=1 SV=1 | 0.01 | 2.597 | 2 | 1 | 1 | 1 | 451 | 51.1 | 6.02 | 2.408 | 2.143 | 1.123 | 1.27 | -1.27 | 1.1 | 0.36666667 | -1.185 | 0.17 | 0.001662601 | 0.024310786 | 0.500759036 | 0.042513286 | 0.384371506 | 0.213442396 | 0.964378027 |
| Slc2a3 | High | P32037 | Solute carrier family 2, facilitated glucose transporter member 3 OS=Mus musculus OX=10090 GI | 0 | 48.364 | 17 | 8 | 21 | 8 | 493 | 53.4 | 4.98 | 0.394 | 0.322 | 1.221 | -1.35 | 1.35 | -1.63 | -0.5433333 | 1.49 | 0.29 | 0.001719325 | 0.000371816 | 0.151273239 | 0.043424564 | 0.016812661 | 0.030118613 | 0.721249193 |
| Vps4a | High | Q8VEJ9 | Vacuolar protein sorting-associated protein 4A OS=Mus musculus OX=10090 GN=Vps4a PE=1 SV= | 0 | 99.187 | 44 | 21 | 27 | 13 | 437 | 48.9 | 7.8 | 0.393 | 0.34 | 1.156 | -1.35 | 1.35 | -1.55 | -0.5166667 | 1.45 | 0.21 | 0.001712255 | 0.00069515 | 0.301566282 | 0.043424564 | 0.027457382 | 0.035440973 | 0.900241969 |
| Efnb1 | High | P52795 | Ephrin-B1 OS=Mus musculus OX=10090 GN=Efnb1 PE=1 SV=1 | 0.005 | 3.526 | 3 | 1 | 1 | 1 | 345 | 37.8 | 9.03 | 0.463 | 0.551 | 0.841 | -1.11 | 1.11 | -0.86 | -0.2866667 | 0.985 | -0.25 | 0.001826084 | 0.047455991 | 0.435546711 | 0.045506004 | 0.520047141 | 0.282776572 | 0.949084077 |
| Rps27a | High | P62983 | Ubiquitin-40S ribosomal protein S27a OS=Mus musculus OX=10090 GN=Rps27a PE=1 SV=2 | 0 | 89.175 | 56 | 10 | 107 | 2 | 156 | 17.9 | 9.64 | 0.401 | 0.333 | 1.203 | -1.32 | 1.32 | -1.59 | -0.53 | 1.455 | 0.27 | 0.002108547 | 0.000542673 | 0.196230053 | 0.052312487 | 0.022933836 | 0.037623161 | 0.784953482 |
| Chmp4b | High | Q9D8B3 | Charged multivesicular body protein 4b OS=Mus musculus OX=10090 GN=Chmp4b PE=1 SV=2 | 0 | 77.119 | 50 | 14 | 41 | 14 | 224 | 24.9 | 4.82 | 0.402 | 0.415 | 0.967 | -1.32 | 1.32 | -1.27 | -0.4233333 | 1.295 | -0.05 | 0.00216264 | 0.005405881 | 0.908009069 | 0.053418166 | 0.141639126 | 0.097528646 | 0.992535481 |
| Pitpnc1 | High | Q8K4R4 | Cytoplasmic phosphatidylinositol transfer protein 1 OS=Mus musculus OX=10090 GN=Pitpnc1 PE= | 0 | 43.975 | 31 | 9 | 10 | 9 | 332 | 38.4 | 6.32 | 2.479 | 3.602 | 0.688 | 1.31 | -1.31 | 1.85 | 0.61666667 | -1.58 | -0.54 | 0.002263039 | 0.000120713 | 0.02542436 | 0.055168963 | 0.006768358 | 0.030968661 | 0.298856161 |
| Cc2d2a | High | Q8CFW7 | Coiled-coil and C2 domain-containing protein 2A OS=Mus musculus OX=10090 GN=Cc2d2a PE=1 S | 0 | 43.335 | 9 | 12 | 12 | 12 | 1633 | 187.4 | 6.83 | 2.39 | 2.823 | 0.847 | 1.26 | -1.26 | 1.5 | 0.5 | -1.38 | -0.24 | 0.002258139 | 0.001952688 | 0.330450876 | 0.055168963 | 0.06592534 | 0.060547152 | 0.906329766 |
| Trf | High | Q921I1 | Serotransferrin OS=Mus musculus OX=10090 GN=Tf PE=1 SV=1 | 0 | 67.814 | 30 | 19 | 26 | 19 | 697 | 76.7 | 7.18 | 0.403 | 0.502 | 0.803 | -1.31 | 1.31 | -0.99 | -0.33 | 1.15 | -0.32 | 0.002251158 | 0.027896956 | 0.187617973 | 0.055168963 | 0.417321937 | 0.23624545 | 0.774149321 |
| Fsip2 | Medium | A2ARZ3 | Fibrous sheath-interacting protein 2 OS=Mus musculus OX=10090 GN=Fsip2 PE=1 SV=3 | 0.03 | 1.857 | 0 | 1 | 1 | 1 | 6995 | 784.4 | 6.51 | 0.469 | 0.487 | 0.963 | -1.09 | 1.09 | -1.04 | -0.3466667 | 1.065 | -0.05 | 0.00229535 | 0.016541648 | 0.845176982 | 0.055714394 | 0.305844112 | 0.180779253 | 0.992535481 |
| Plcxd2 | High | B2RXA1 | PI-PLC X domain-containing protein 2 OS=Mus musculus OX=10090 GN=Plcxd2 PE=2 SV=1 | 0 | 76.101 | 33 | 13 | 24 | 13 | 340 | 38.6 | 8.88 | 2.469 | 2.586 | 0.955 | 1.3 | -1.3 | 1.37 | 0.45666667 | -1.335 | -0.07 | 0.002329141 | 0.004685587 | 0.84428559 | 0.056290922 | 0.128784729 | 0.092537826 | 0.992535481 |
| Cd63-fr | High | A0A1W2P | Tetraspanin (Fragment) OS=Mus musculus OX=10090 GN=Cd63 PE=1 SV=1 | 0.001 | 6.149 | 11 | 2 | 2 | 2 | 164 | 17.9 | 7.5 | 0.498 | 0.475 | 1.049 | -1 | 1 | -1.07 | -0.3566667 | 1.035 | 0.07 | 0.002444004 | 0.01398262 | 0.733384184 | 0.0585621 | 0.273438276 | 0.166000188 | 0.992535481 |
| Vps37c | High | Q8R105 | Vacuolar protein sorting-associated protein 37C OS=Mus musculus OX=10090 GN=Vps37c PE=1 S' | 0 | 22.889 | 21 | 4 | 4 | 4 | 352 | 38.4 | 5.31 | 0.457 | 0.424 | 1.077 | -1.13 | 1.13 | -1.24 | -0.4133333 | 1.185 | 0.11 | 0.002517527 | 0.004889884 | 0.646755907 | 0.060067118 | 0.129022591 | 0.094544855 | 0.985010103 |
| Atp1a3 | High | A0A0G2JG | Sodium/potassium-transporting ATPase subunit alpha OS=Mus musculus OX=10090 GN=Atp1a3 I | 0 | 934.121 | 56 | 67 | 517 | 44 | 1026 | 112.9 | 5.41 | 0.409 | 0.498 | 0.821 | -1.29 | 1.29 | -1.01 | -0.3366667 | 1.15 | -0.29 | 0.002650302 | 0.02624545 | 0.240726104 | 0.062412856 | 0.405394594 | 0.233903725 | 0.832665803 |
| Cldn23 | High | Q9D7D7 | Claudin-23 OS=Mus musculus OX=10090 GN=Cldn23 PE=1 SV=1 | 0.001 | 6.334 | 7 | 3 | 4 | 3 | 296 | 32.4 | 7.55 | 0.478 | 0.582 | 0.822 | -1.06 | 1.06 | -0.78 | -0.26 | 0.92 | -0.28 | 0.00267537 | 0.0748859 | 0.284476473 | 0.06250333 | 0.637155143 | 0.349829236 | 0.886074068 |
| Ndfip1 | High | A0A494BA | NEDD4 family-interacting protein 1 OS=Mus musculus OX=10090 GN=Ndfip1 PE=1 SV=1 | 0.001 | 5.8 | 17 | 1 | 2 | 1 | 167 | 18 | 4.02 | 0.442 | 0.664 | 0.665 | -1.18 | 1.18 | -0.59 | -0.1966667 | 0.885 | -0.59 | 0.002935045 | 0.16359267 | 0.051066979 | 0.067445883 | 0.869444647 | 0.468445265 | 0.427566646 |
| Spata6 | High | Q3U6K5 | Spermatogenesis-associated protein 6 OS=Mus musculus OX=10090 GN=Spata6 PE=1 SV=2 | 0.001 | 5.278 | 4 | 2 | 2 | 2 | 488 | 56.1 | 8.47 | 2.174 | 2.904 | 0.749 | 1.12 | -1.12 | 1.54 | 0.51333333 | -1.33 | -0.42 | 0.003098517 | 0.00121182 | 0.101637671 | 0.07062352 | 0.044121247 | 0.057372383 | 0.610114517 |
| Krt76 | High | Q3UV17 | Keratin, type II cytoskeletal 2 oral OS=Mus musculus OX=10090 GN=Krt76 PE=1 SV=1 | 0 | 14.451 | 8 | 5 | 11 | 1 | 594 | 62.8 | 8.43 | 2.088 | 1.389 | 1.503 | 1.06 | -1.06 | 0.47 | 0.15666667 | -0.765 | 0.59 | 0.003191448 | 0.356216458 | 0.018118841 | 0.072447153 | 0.977313184 | 0.524880169 | 0.237723313 |
| Alms1 | High | Q8K4E0 | Alstrom syndrome protein 1 homolog OS=Mus musculus OX=10090 GN=Alms1 PE=1 SV=2 | 0 | 27.648 | 4 | 11 | 11 | 11 | 3251 | 360 | 6.39 | 2.26 | 2.466 | 0.917 | 1.18 | -1.18 | 1.3 | 0.43333333 | -1.24 | -0.13 | 0.003289521 | 0.007318136 | 0.584443903 | 0.072505492 | 0.174607618 | 0.123556555 | 0.973095093 |
| Ccsap | High | Q8QZT2 | Centriole, cilia and spindle-associated protein OS=Mus musculus OX=10090 GN=Ccsap PE=1 SV=1 | 0 | 10.387 | 11 | 2 | 2 | 2 | 252 | 28.4 | 9.22 | 2.187 | 2.167 | 1.009 | 1.13 | -1.13 | 1.12 | 0.37333333 | -1.125 | 0.01 | 0.003318246 | 0.019925134 | 0.930758475 | 0.072505492 | 0.345883049 | 0.20919427 | 0.992535481 |
| Nme7 | High | Q3UMG6 | Nucleoside diphosphate kinase 7 OS=Mus musculus OX=10090 GN=Nme7 PE=1 SV=1 | 0 | 19.384 | 14 | 5 | 5 | 5 | 395 | 44.4 | 7.12 | 2.326 | 2.123 | 1.096 | 1.22 | -1.22 | 1.09 | 0.36333333 | -1.155 | 0.13 | 0.003299974 | 0.026182493 | 0.597458387 | 0.072505492 | 0.405394594 | 0.238950043 | 0.977451587 |
| Efnb2 | High | F6RSU6 | Ephrin-B2 (Fragment) OS=Mus musculus OX=10090 GN=Efnb2 PE=1 SV=1 | 0.001 | 4.367 | 8 | 1 | 1 | 1 | 218 | 23.9 | 8.87 | 0.507 | 0.513 | 0.989 | -0.98 | 0.98 | -0.96 | -0.32 | 0.97 | -0.02 | 0.003336261 | 0.026694037 | 0.9823416 | 0.072505492 | 0.407239716 | 0.239872604 | 0.997765878 |
| Rad23a | High | Q3TN85 | UV excision repair protein RAD23 homolog A (Fragment) OS=Mus musculus OX=10090 GN=Rad23 | 0.001 | 5.955 | 7 | 2 | 3 | 1 | 322 | 34.8 | 4.69 | 0.481 | 0.533 | 0.903 | -1.06 | 1.06 | -0.91 | -0.3033333 | 0.985 | -0.15 | 0.003290139 | 0.035843434 | 0.588600661 | 0.072505492 | 0.476392832 | 0.274449162 | 0.973095093 |
| Traf3ip1 | High | A0A087W | TRAF3-interacting protein 1 OS=Mus musculus OX=10090 GN=Traf3ip1 PE=1 SV=1 | 0 | 8.949 | 6 | 4 | 4 | 4 | 654 | 74.2 | 7.12 | 2.105 | 2.021 | 1.041 | 1.07 | -1.07 | 1.01 | 0.33666667 | -1.04 | 0.06 | 0.00344949 | 0.034588501 | 0.752664112 | 0.074307178 | 0.476392832 | 0.275350005 | 0.992535481 |
| Mecr | High | Q9DCS3 | Enoyl-[acyl-carrier-protein] reductase, mitochondrial OS=Mus musculus OX=10090 GN=Mecr PE= | 0.005 | 3.788 | 2 | 1 | 1 | 1 | 373 | 40.3 | 9.07 | 2.164 | 1.779 | 1.216 | 1.11 | -1.11 | 0.83 | 0.27666667 | -0.97 | 0.28 | 0.003458922 | 0.084751659 | 0.222946583 | 0.074307178 | 0.682905914 | 0.378606546 | 0.808577935 |
| D8Ertd738 | Medium | Q8R1F0 | Leydig cell tumor 10 kDa protein homolog OS=Mus musculus OX=10090 GN=D8Ertd738e PE=3 SV | 0.01 | 2.695 | 14 | 2 | 3 | 2 | 94 | 10.2 | 11.63 | 2.235 | 1.573 | 1.421 | 1.16 | -1.16 | 0.65 | 0.21666667 | -0.905 | 0.51 | 0.003501238 | 0.183908781 | 0.032399117 | 0.07492916 | 0.895896206 | 0.485412684 | 0.349349706 |
| Lrrc8b | High | Q5DU41 | Volume-regulated anion channel subunit LRRC8B OS=Mus musculus OX=10090 GN=Lrrc8b PE=1 S | 0 | 10.597 | 4 | 3 | 3 | 3 | 803 | 92.1 | 6.7 | 0.484 | 0.482 | 1.004 | -1.05 | 1.05 | -1.05 | -0.35 | 1.05 | 0.01 | 0.003559081 | 0.020282244 | 0.956000404 | 0.075590034 | 0.345911522 | 0.210750778 | 0.992902921 |
| Xiap | High | Q60989 | E3 ubiquitin-protein ligase XIAP OS=Mus musculus OX=10090 GN=Xiap PE=1 SV=2 | 0 | 24.689 | 17 | 6 | 7 | 6 | 496 | 56 | 6.23 | 0.483 | 0.609 | 0.794 | -1.05 | 1.05 | -0.72 | -0.24 | 0.885 | -0.33 | 0.003552983 | 0.107734353 | 0.153293983 | 0.075590034 | 0.759466825 | 0.417528429 | 0.727583768 |
| Nucb2 | High | P81117 | Nucleobindin-2 OS=Mus musculus OX=10090 GN=Nucb2 PE=1 SV=2 | 0.001 | 5.554 | 4 | 1 | 1 | 1 | 420 | 50.3 | 5.15 | 2.11 | 3.032 | 0.696 | 1.08 | -1.08 | 1.6 | 0.53333333 | -1.34 | -0.52 | 0.003657915 | 0.000665687 | 0.039829621 | 0.077104993 | 0.027244573 | 0.052174783 | 0.38570757 |
| Nphp4 | High | P59240 | Nephrocystin-4 OS=Mus musculus OX=10090 GN=Nphp4 PE=1 SV=2 | 0 | 35.518 | 8 | 10 | 11 | 10 | 1425 | 157.2 | 7.77 | 2.375 | 2.599 | 0.914 | 1.25 | -1.25 | 1.38 | 0.46 | -1.315 | -0.13 | 0.003819802 | 0.004452873 | 0.660736835 | 0.080215835 | 0.124836305 | 0.10252607 | 0.98592496 |
| Serinc3 | High | Q9QZI9 | Serine incorporator 3 OS=Mus musculus OX=10090 GN=Serinc3 PE=1 SV=2 | 0 | 48.867 | 13 | 5 | 28 | 5 | 472 | 52.6 | 7.23 | 0.423 | 0.479 | 0.884 | -1.24 | 1.24 | -1.06 | -0.3533333 | 1.15 | -0.18 | 0.003848536 | 0.019003345 | 0.487643357 | 0.080517681 | 0.331936928 | 0.206227305 | 0.964378027 |
| Ank | High | Q9JHZ2 | Progressive ankylosis protein OS=Mus musculus OX=10090 GN=Ankh PE=2 SV=1 | 0.001 | 6.458 | 5 | 2 | 2 | 2 | 492 | 54.3 | 7.88 | 0.48 | 0.547 | 0.876 | -1.06 | 1.06 | -0.87 | -0.29 | 0.965 | -0.19 | 0.003883986 | 0.04485356 | 0.458329235 | 0.080957276 | 0.520047141 | 0.300502208 | 0.959744396 |
| Plekhb1 | High | Q9QYE9 | Pleckstrin homology domain-containing family B member 1 OS=Mus musculus OX=10090 GN=Ple | 0 | 84.871 | 51 | 12 | 37 | 4 | 243 | 27.3 | 6.98 | 0.425 | 0.524 | 0.812 | -1.23 | 1.23 | -0.93 | -0.31 | 1.08 | -0.3 | 0.00405709 | 0.038667904 | 0.212824999 | 0.08363274 | 0.486480307 | 0.285056524 | 0.804011919 |
| Podxl2 | Medium | Q8CAE9 | Podocalyxin-like protein 2 OS=Mus musculus OX=10090 GN=Podxl2 PE=1 SV=2 | 0.035 | 1.79 | 2 | 1 | 1 | 1 | 603 | 64.9 | 4.36 | 0.494 | 0.384 | 1.286 | -1.02 | 1.02 | -1.38 | -0.46 | 1.2 | 0.36 | 0.004521891 | 0.001975291 | 0.194860898 | 0.086417902 | 0.06592534 | 0.076171621 | 0.784517504 |
| Chmp1b | High | Q99LU0 | Charged multivesicular body protein 1b-1 OS=Mus musculus OX=10090 GN=Chmp1b1 PE=1 SV=1 | 0 | 8.746 | 9 | 2 | 3 | 2 | 199 | 22.1 | 8.1 | 0.45 | 0.439 | 1.027 | -1.15 | 1.15 | -1.19 | -0.3966667 | 1.17 | 0.04 | 0.00477637 | 0.008950811 | 0.818748252 | 0.086417902 | 0.200748781 | 0.143583342 | 0.992535481 |
| Csnk1a1 | High | E9Q3W1 | Casein kinase I isoform alpha OS=Mus musculus OX=10090 GN=Csnk1a1 PE=1 SV=1 | 0 | 78.19 | 24 | 9 | 20 | 1 | 365 | 41.9 | 9.6 | 2.326 | 2.337 | 0.995 | 1.22 | -1.22 | 1.22 | 0.40666667 | -1.22 | -0.01 | 0.004471276 | 0.011827348 | 0.944463183 | 0.086417902 | 0.249308049 | 0.167862976 | 0.992535481 |
| Susd3-fr | Medium | D3YY27 | Sushi domain-containing protein 3 (Fragment) OS=Mus musculus OX=10090 GN=Susd3 PE=1 SV= | 0.011 | 2.389 | 6 | 1 | 1 | 1 | 139 | 14.9 | 8.63 | 0.475 | 0.474 | 1.002 | -1.08 | 1.08 | -1.08 | -0.36 | 1.08 | 0 | 0.00464929 | 0.01521177 | 0.927375591 | 0.086417902 | 0.290110179 | 0.188264041 | 0.992535481 |
| Chmp5 | High | Q9D7S9 | Charged multivesicular body protein 5 OS=Mus musculus OX=10090 GN=Chmp5 PE=1 SV=1 | 0 | 42.015 | 46 | 7 | 20 | 7 | 219 | 24.6 | 4.79 | 0.421 | 0.472 | 0.893 | -1.25 | 1.25 | -1.08 | -0.36 | 1.165 | -0.16 | 0.004235142 | 0.016819735 | 0.546975992 | 0.086417902 | 0.305844112 | 0.196131007 | 0.96694628 |
| H2-K1 | High | P01901 | H-2 class I histocompatibility antigen, K-B alpha chain OS=Mus musculus OX=10090 GN=H2-K1 PE | 0.003 | 4.064 | 3 | 1 | 2 | 1 | 369 | 41.3 | 6.39 | 0.487 | 0.543 | 0.896 | -1.04 | 1.04 | -0.88 | -0.2933333 | 0.96 | -0.16 | 0.004839526 | 0.047951556 | 0.551749221 | 0.086417902 | 0.520047141 | 0.303232522 | 0.96694628 |
| Phlpp1 | Medium | Q8CHE4 | PH domain leucine-rich repeat-containing protein phosphatase 1 OS=Mus musculus OX=10090 GN | 0.01 | 2.687 | 0 | 1 | 1 | 1 | 1687 | 182.3 | 6.19 | 0.492 | 0.563 | 0.874 | -1.02 | 1.02 | -0.83 | -0.2766667 | 0.925 | -0.19 | 0.004518136 | 0.055028451 | 0.453483424 | 0.086417902 | 0.567177431 | 0.326797667 | 0.959744396 |
| Gnaq | High | P21279 | Guanine nucleotide-binding protein G(q) subunit alpha OS=Mus musculus OX=10090 GN=Gnaq PE | 0 | 87.202 | 44 | 13 | 32 | 9 | 359 | 42.1 | 5.68 | 0.434 | 0.455 | 0.952 | -1.21 | 1.21 | -1.13 | -0.3766667 | 1.17 | -0.07 | 0.004984454 | 0.012490796 | 0.828960223 | 0.088723289 | 0.257484909 | 0.173104099 | 0.992535481 |
| Polg2 | Low | Q9QZM2 | DNA polymerase subunit gamma-2, mitochondrial OS=Mus musculus OX=10090 GN=Polg2 PE=1 S | 0.069 | 1.305 | 2 | 1 | 1 | 1 | 459 | 51.4 | 8.44 | 2.117 | 1.721 | 1.23 | 1.08 | -1.08 | 0.78 | 0.26 | -0.93 | 0.3 | 0.005055845 | 0.103050614 | 0.184352453 | 0.089709257 | 0.736995911 | 0.413352584 | 0.768475941 |
| Vps28 | High | Q9D1C8 | Vacuolar protein sorting-associated protein 28 homolog OS=Mus musculus OX=10090 GN=Vps28 | 0 | 39.625 | 48 | 9 | 14 | 9 | 221 | 25.4 | 5.54 | 0.457 | 0.384 | 1.19 | -1.13 | 1.13 | -1.38 | -0.46 | 1.255 | 0.25 | 0.005155655 | 0.002513144 | 0.328437966 | 0.091191661 | 0.080983898 | 0.086087779 | 0.905686588 |
| Elmod2 | High | Q8BGF6 | ELMO domain-containing protein 2 OS=Mus musculus OX=10090 GN=Elmod2 PE=1 SV=1 | 0.001 | 6.101 | 6 | 2 | 2 | 2 | 293 | 34.7 | 8.57 | 2.054 | 1.739 | 1.181 | 1.04 | -1.04 | 0.8 | 0.26666667 | -0.92 | 0.24 | 0.00529792 | 0.096480218 | 0.340699257 | 0.093120485 | 0.712733312 | 0.402926899 | 0.907770836 |
| Glb1l2 | High | A0A0R4J0 | Beta-galactosidase OS=Mus musculus OX=10090 GN=Glb1l2 PE=1 SV=1 | 0 | 172.103 | 37 | 20 | 62 | 20 | 652 | 73.9 | 7.87 | 2.281 | 2.921 | 0.781 | 1.19 | -1.19 | 1.55 | 0.51666667 | -1.37 | -0.36 | 0.005485794 | 0.001364858 | 0.134882889 | 0.096121403 | 0.048743678 | 0.07243254 | 0.69768299 |
| Mak | High | Q04859 | Serine/threonine-protein kinase MAK OS=Mus musculus OX=10090 GN=Mak PE=1 SV=2 | 0 | 85.428 | 41 | 22 | 28 | 20 | 622 | 70 | 9.66 | 2.255 | 2.48 | 0.909 | 1.17 | -1.17 | 1.31 | 0.43666667 | -1.24 | -0.14 | 0.006148991 | 0.00693696 | 0.608987259 | 0.106411697 | 0.169849505 | 0.138130601 | 0.97826833 |
| Lztfl1 | High | Q9JHQ5 | Leucine zipper transcription factor-like protein 1 OS=Mus musculus OX=10090 GN=Lztfl1 PE=1 SV | 0 | 111.702 | 51 | 17 | 41 | 17 | 299 | 34.8 | 5.17 | 0.444 | 0.431 | 1.029 | -1.17 | 1.17 | -1.21 | -0.4033333 | 1.19 | 0.04 | 0.006347675 | 0.007693477 | 0.774642198 | 0.109512045 | 0.176353762 | 0.142932903 | 0.992535481 |
| Arhgef18 | High | A0A5F8MI | Rho guanine nucleotide exchange factor 18 OS=Mus musculus OX=10090 GN=Arhgef18 PE=1 SV= | 0 | 4.859 | 2 | 2 | 2 | 2 | 1126 | 125.7 | 6.86 | 0.52 | 0.456 | 1.141 | -0.94 | 0.94 | -1.13 | -0.3766667 | 1.035 | 0.19 | 0.006559262 | 0.009558302 | 0.435864365 | 0.111831547 | 0.210997641 | 0.161414594 | 0.949084077 |
| Elfn1 | High | Q8C8T7 | Protein ELFN1 OS=Mus musculus OX=10090 GN=Elfn1 PE=2 SV=1 | 0.009 | 2.856 | 1 | 1 | 1 | 1 | 828 | 90.8 | 8.18 | 0.465 | 0.586 | 0.795 | -1.1 | 1.1 | -0.77 | -0.2566667 | 0.935 | -0.33 | 0.006561901 | 0.073729669 | 0.303943887 | 0.111831547 | 0.636899913 | 0.37436573 | 0.900241969 |
| Stx7 | High | Q8BH40 | Syntaxin-7 OS=Mus musculus OX=10090 GN=Stx7 PE=1 SV=1 | 0 | 50.116 | 44 | 9 | 20 | 9 | 261 | 29.7 | 5.58 | 0.444 | 0.384 | 1.157 | -1.17 | 1.17 | -1.38 | -0.46 | 1.275 | 0.21 | 0.006998636 | 0.002506751 | 0.344914649 | 0.118196842 | 0.080983898 | 0.09959037 | 0.907770836 |
| Slc39a6 | High | Q8C145 | Zinc transporter ZIP6 OS=Mus musculus OX=10090 GN=Slc39a6 PE=1 SV=1 | 0 | 4.85 | 2 | 2 | 4 | 2 | 765 | 86.3 | 6.84 | 0.49 | 0.337 | 1.454 | -1.03 | 1.03 | -1.57 | -0.5233333 | 1.3 | 0.54 | 0.007093719 | 0.00062982 | 0.013813268 | 0.119442883 | 0.026158531 | 0.072800707 | 0.192664157 |
| Mt3 | High | P28184 | Metallothionein-3 OS=Mus musculus OX=10090 GN=Mt3 PE=1 SV=1 | 0 | 20.645 | 35 | 3 | 8 | 3 | 68 | 7 | 7.47 | 2.181 | 2.037 | 1.071 | 1.12 | -1.12 | 1.03 | 0.34333333 | -1.075 | 0.1 | 0.007499176 | 0.036115127 | 0.60785852 | 0.125558595 | 0.476392832 | 0.300975713 | 0.97826833 |
| Golga5 | High | Q9QYE6 | Golgin subfamily A member 5 OS=Mus musculus OX=10090 GN=Golga5 PE=1 SV=2 | 0.008 | 2.942 | 3 | 2 | 2 | 2 | 729 | 82.3 | 6.23 | 0.53 | 0.731 | 0.726 | -0.92 | 0.92 | -0.45 | -0.15 | 0.685 | -0.46 | 0.007593151 | 0.269933115 | 0.067425495 | 0.125960936 | 0.95334888 | 0.539654908 | 0.499896415 |
| Gtf3c4 | Medium | E9Q2B0 | General transcription factor 3C polypeptide 4 OS=Mus musculus OX=10090 GN=Gtf3c4 PE=1 SV=1 | 0.03 | 1.876 | 2 | 1 | 1 | 1 | 676 | 76.7 | 7.14 | 1.957 | 1.52 | 1.288 | 0.97 | -0.97 | 0.6 | 0.2 | -0.785 | 0.36 | 0.007902535 | 0.226787984 | 0.10281317 | 0.130322092 | 0.935008675 | 0.532665384 | 0.610114517 |
| Scamp2 | High | Q9ERN0 | Secretory carrier-associated membrane protein 2 OS=Mus musculus OX=10090 GN=Scamp2 PE=1 | 0.001 | 6.969 | 6 | 2 | 2 | 2 | 329 | 36.4 | 6.32 | 0.537 | 0.552 | 0.973 | -0.9 | 0.9 | -0.86 | -0.2866667 | 0.88 | -0.04 | 0.008190784 | 0.052134955 | 0.884346356 | 0.134679544 | 0.546393818 | 0.340536681 | 0.992535481 |
| Mrpl22 | Low | Q8BU88 | 39S ribosomal protein L22, mitochondrial OS=Mus musculus OX=10090 GN=Mrpl22 PE=1 SV=1 | 0.054 | 1.413 | 3 | 1 | 1 | 1 | 206 | 23.8 | 9.86 | 0.547 | 0.522 | 1.048 | -0.87 | 0.87 | -0.94 | -0.3133333 | 0.905 | 0.07 | 0.00851973 | 0.037388223 | 0.717246164 | 0.139678737 | 0.480815973 | 0.310247355 | 0.992535481 |
| Tsg101 | High | Q61187 | Tumor susceptibility gene 101 protein OS=Mus musculus OX=10090 GN=Tsg101 PE=1 SV=2 | 0 | 65.134 | 28 | 12 | 21 | 12 | 391 | 44.1 | 6.71 | 0.458 | 0.409 | 1.119 | -1.13 | 1.13 | -1.29 | -0.43 | 1.21 | 0.16 | 0.008682332 | 0.004732687 | 0.410877804 | 0.141929547 | 0.128816384 | 0.135372965 | 0.936397673 |
| Snf8 | High | Q9CZ28 | Vacuolar-sorting protein SNF8 OS=Mus musculus OX=10090 GN=Snf8 PE=1 SV=1 | 0 | 26.555 | 24 | 6 | 7 | 6 | 258 | 28.9 | 6.65 | 0.513 | 0.463 | 1.109 | -0.96 | 0.96 | -1.11 | -0.37 | 1.035 | 0.15 | 0.008775509 | 0.014359271 | 0.541699939 | 0.143035689 | 0.277629069 | 0.210332379 | 0.96694628 |
| Scg2 | High | Q03517 | Secretogranin-2 OS=Mus musculus OX=10090 GN=Scg2 PE=1 SV=1 | 0.002 | 4.221 | 6 | 2 | 2 | 2 | 617 | 70.6 | 4.75 | 0.512 | 0.487 | 1.053 | -0.97 | 0.97 | -1.04 | -0.3466667 | 1.005 | 0.07 | 0.008862172 | 0.017262352 | 0.748016743 | 0.143613292 | 0.305844112 | 0.224728702 | 0.992535481 |
| Krt16 | High | Q9Z2K1 | Keratin, type I cytoskeletal 16 OS=Mus musculus OX=10090 GN=Krt16 PE=1 SV=3 | 0 | 17.465 | 10 | 6 | 10 | 2 | 469 | 51.6 | 5.2 | 1.97 | 2.41 | 0.818 | 0.98 | -0.98 | 1.27 | 0.42333333 | -1.125 | -0.29 | 0.009189535 | 0.007705845 | 0.260466487 | 0.148062426 | 0.176353762 | 0.162208094 | 0.856560464 |
| Cd44 | High | A2APM3 | CD44 antigen OS=Mus musculus OX=10090 GN=Cd44 PE=1 SV=1 | 0 | 12.726 | 9 | 4 | 5 | 4 | 464 | 50.7 | 5.29 | 0.533 | 0.552 | 0.966 | -0.91 | 0.91 | -0.86 | -0.2866667 | 0.885 | -0.05 | 0.009876318 | 0.056366174 | 0.866573922 | 0.152973803 | 0.569450702 | 0.361212253 | 0.992535481 |
| Zc3h18 | High | G3X8T2 | Zinc finger CCCH domain-containing protein 18 OS=Mus musculus OX=10090 GN=Zc3h18 PE=1 SV | 0.001 | 5.61 | 2 | 2 | 2 | 2 | 972 | 108.2 | 8 | 2.085 | 1.428 | 1.46 | 1.06 | -1.06 | 0.51 | 0.17 | -0.785 | 0.55 | 0.009820808 | 0.323202553 | 0.038177489 | 0.152973803 | 0.955820031 | 0.554396917 | 0.373579725 |
| Cfh | High | P06909 | Complement factor H OS=Mus musculus OX=10090 GN=Cfh PE=1 SV=2 | 0 | 9.931 | 2 | 3 | 3 | 3 | 1234 | 139 | 6.99 | 0.508 | 0.411 | 1.236 | -0.98 | 0.98 | -1.28 | -0.4266667 | 1.13 | 0.31 | 0.010204981 | 0.00490134 | 0.16579148 | 0.154669302 | 0.129022591 | 0.141845947 | 0.736602876 |
| Dync2h1 | High | Q45VK7 | Cytoplasmic dynein 2 heavy chain 1 OS=Mus musculus OX=10090 GN=Dync2h1 PE=1 SV=1 | 0 | 35.211 | 3 | 13 | 14 | 13 | 4306 | 492 | 6.6 | 2.126 | 2.088 | 1.018 | 1.09 | -1.09 | 1.06 | 0.35333333 | -1.075 | 0.03 | 0.010206464 | 0.029800245 | 0.810015898 | 0.154669302 | 0.438556356 | 0.296612829 | 0.992535481 |
| Armc9 | High | Q9D2I5 | LisH domain-containing protein ARMC9 OS=Mus musculus OX=10090 GN=Armc9 PE=1 SV=1 | 0 | 10.456 | 5 | 4 | 4 | 4 | 817 | 91.9 | 6.92 | 1.98 | 1.642 | 1.206 | 0.99 | -0.99 | 0.72 | 0.24 | -0.855 | 0.27 | 0.01001996 | 0.150716697 | 0.235718297 | 0.154669302 | 0.847684847 | 0.501177075 | 0.82759705 |
| Ift57 | High | Q8BXG3 | Intraflagellar transport protein 57 homolog OS=Mus musculus OX=10090 GN=Ift57 PE=1 SV=1 | 0.001 | 7.099 | 8 | 4 | 4 | 4 | 429 | 48.7 | 4.98 | 1.983 | 1.824 | 1.087 | 0.99 | -0.99 | 0.87 | 0.29 | -0.93 | 0.12 | 0.010489088 | 0.078737808 | 0.588851605 | 0.158523765 | 0.65511466 | 0.406819213 | 0.973095093 |
| Cep135 | High | Q6P5D4 | Centrosomal protein of 135 kDa OS=Mus musculus OX=10090 GN=Cep135 PE=1 SV=1 | 0 | 64.575 | 18 | 19 | 22 | 19 | 1140 | 133.3 | 6.48 | 2.048 | 2.142 | 0.956 | 1.03 | -1.03 | 1.1 | 0.36666667 | -1.065 | -0.06 | 0.01056834 | 0.02440454 | 0.849534593 | 0.15929216 | 0.384371506 | 0.271831833 | 0.992535481 |
| Ilf3 | High | A0A1L1ST | Interleukin enhancer-binding factor 3 OS=Mus musculus OX=10090 GN=Ilf3 PE=1 SV=1 | 0.006 | 3.425 | 1 | 1 | 1 | 1 | 908 | 97.4 | 8.92 | 0.541 | 0.581 | 0.931 | -0.89 | 0.89 | -0.78 | -0.26 | 0.835 | -0.1 | 0.010664586 | 0.079450324 | 0.701342756 | 0.159456892 | 0.65511466 | 0.407285776 | 0.991285312 |
| Tollip | High | Q9QZ06 | Toll-interacting protein OS=Mus musculus OX=10090 GN=Tollip PE=1 SV=1 | 0 | 48.756 | 45 | 10 | 17 | 10 | 274 | 30.3 | 5.17 | 0.468 | 0.491 | 0.953 | -1.1 | 1.1 | -1.03 | -0.3433333 | 1.065 | -0.07 | 0.010737973 | 0.023430853 | 0.831317455 | 0.160127167 | 0.380802291 | 0.270464729 | 0.992535481 |
| Kras | High | P32883 | GTPase KRas OS=Mus musculus OX=10090 GN=Kras PE=1 SV=1 | 0 | 135.386 | 56 | 9 | 28 | 2 | 189 | 21.6 | 6.77 | 0.447 | 0.398 | 1.122 | -1.16 | 1.16 | -1.33 | -0.4433333 | 1.245 | 0.17 | 0.011054154 | 0.003629014 | 0.496737857 | 0.160156695 | 0.104886 | 0.132521347 | 0.964378027 |
| Brox | High | Q8K2Q7 | BRO1 domain-containing protein BROX OS=Mus musculus OX=10090 GN=Brox PE=1 SV=1 | 0 | 16.621 | 16 | 6 | 7 | 6 | 411 | 46.2 | 7.69 | 0.508 | 0.427 | 1.191 | -0.98 | 0.98 | -1.23 | -0.41 | 1.105 | 0.25 | 0.011030353 | 0.006998389 | 0.312311944 | 0.160156695 | 0.169869988 | 0.165013341 | 0.904943958 |
| Chmp3 | High | Q9CQ10 | Charged multivesicular body protein 3 OS=Mus musculus OX=10090 GN=Chmp3 PE=1 SV=3 | 0 | 29.837 | 56 | 10 | 14 | 10 | 224 | 25.2 | 5.06 | 0.478 | 0.466 | 1.025 | -1.06 | 1.06 | -1.1 | -0.3666667 | 1.08 | 0.04 | 0.010925695 | 0.01530922 | 0.901377046 | 0.160156695 | 0.290978976 | 0.225567835 | 0.992535481 |
| Gm20390 | High | E9PZF0 | Nucleoside diphosphate kinase OS=Mus musculus OX=10090 GN=Gm20390 PE=3 SV=1 | 0 | 47.156 | 54 | 11 | 20 | 3 | 267 | 30.2 | 8.72 | 0.47 | 0.471 | 1 | -1.09 | 1.09 | -1.09 | -0.3633333 | 1.09 | 0 | 0.010997092 | 0.016483191 | 0.929907796 | 0.160156695 | 0.305844112 | 0.233000403 | 0.992535481 |
| Zfpl1 | High | E0CZD9 | Zinc finger protein-like 1 (Fragment) OS=Mus musculus OX=10090 GN=Zfpl1 PE=1 SV=1 | 0.001 | 5.162 | 4 | 1 | 1 | 1 | 242 | 26.8 | 7.11 | 0.534 | 0.778 | 0.687 | -0.9 | 0.9 | -0.36 | -0.12 | 0.63 | -0.54 | 0.011008073 | 0.385208157 | 0.036271136 | 0.160156695 | 0.977313184 | 0.568734939 | 0.370300617 |
| Sirpa | High | E0CX65 | Tyrosine-protein phosphatase non-receptor type substrate 1 OS=Mus musculus OX=10090 GN=Sir | 0 | 37.599 | 47 | 4 | 8 | 1 | 172 | 18.9 | 7.37 | 0.511 | 0.476 | 1.073 | -0.97 | 0.97 | -1.07 | -0.3566667 | 1.02 | 0.1 | 0.011134725 | 0.015647642 | 0.652796415 | 0.160908252 | 0.294417206 | 0.227662729 | 0.985010103 |
| Pcdh15 | High | Q99PJ1 | Protocadherin-15 OS=Mus musculus OX=10090 GN=Pcdh15 PE=1 SV=3 | 0 | 13.212 | 2 | 4 | 4 | 4 | 1943 | 214.6 | 5.15 | 1.942 | 1.555 | 1.249 | 0.96 | -0.96 | 0.64 | 0.21333333 | -0.8 | 0.32 | 0.011287247 | 0.19201673 | 0.166617927 | 0.161860859 | 0.907729584 | 0.534795221 | 0.737935797 |
| Impg2 | High | Q80XH2 | Interphotoreceptor matrix proteoglycan 2 OS=Mus musculus OX=10090 GN=Impg2 PE=1 SV=1 | 0 | 65.605 | 12 | 12 | 22 | 12 | 1243 | 138 | 4.78 | 0.455 | 0.376 | 1.208 | -1.14 | 1.14 | -1.41 | -0.47 | 1.275 | 0.27 | 0.011544033 | 0.002043359 | 0.230862035 | 0.164700741 | 0.067793568 | 0.116247154 | 0.819676582 |
| Stk10 | Medium | O55098 | Serine/threonine-protein kinase 10 OS=Mus musculus OX=10090 GN=Stk10 PE=1 SV=2 | 0.049 | 1.508 | 1 | 1 | 1 | 1 | 966 | 111.8 | 7.15 | 0.542 | 0.598 | 0.906 | -0.88 | 0.88 | -0.74 | -0.2466667 | 0.81 | -0.14 | 0.011536036 | 0.08934288 | 0.664307456 | 0.164700741 | 0.696517952 | 0.430609346 | 0.986647506 |
| Pdcd6ip | High | Q9WU78 | Programmed cell death 6-interacting protein OS=Mus musculus OX=10090 GN=Pdcd6ip PE=1 SV= | 0 | 184.342 | 54 | 42 | 63 | 42 | 869 | 96 | 6.52 | 0.472 | 0.431 | 1.097 | -1.08 | 1.08 | -1.22 | -0.4066667 | 1.15 | 0.13 | 0.011743947 | 0.007606615 | 0.486666978 | 0.167127698 | 0.176353762 | 0.17174073 | 0.964378027 |
| Lpcat1 | High | D6RFA8 | Lysophosphatidylcholine acyltransferase 1 OS=Mus musculus OX=10090 GN=Lpcat1 PE=1 SV=1 | 0.005 | 3.558 | 17 | 1 | 1 | 1 | 46 | 4.9 | 12.22 | 0.527 | 0.578 | 0.911 | -0.92 | 0.92 | -0.79 | -0.2633333 | 0.855 | -0.13 | 0.011859892 | 0.074795979 | 0.605180073 | 0.167502302 | 0.637155143 | 0.402328722 | 0.97826833 |
| Eno3 | High | P21550 | Beta-enolase OS=Mus musculus OX=10090 GN=Eno3 PE=1 SV=3 | 0 | 134.82 | 24 | 8 | 58 | 1 | 434 | 47 | 7.18 | 0.535 | 0.719 | 0.743 | -0.9 | 0.9 | -0.48 | -0.16 | 0.69 | -0.43 | 0.012113929 | 0.272467152 | 0.069585747 | 0.170660296 | 0.955820031 | 0.563240163 | 0.504092101 |
| Luc7l2 | High | E9Q715 | Putative RNA-binding protein Luc7-like 2 OS=Mus musculus OX=10090 GN=Luc7l2 PE=1 SV=1 | 0 | 11.705 | 9 | 2 | 2 | 2 | 272 | 32.6 | 10.01 | 1.94 | 1.324 | 1.465 | 0.96 | -0.96 | 0.41 | 0.13666667 | -0.685 | 0.55 | 0.01257928 | 0.43252749 | 0.014386007 | 0.175890332 | 0.99399649 | 0.584943411 | 0.197218435 |
| Wdr54 | High | Q9R0D8 | WD repeat-containing protein 54 OS=Mus musculus OX=10090 GN=Wdr54 PE=1 SV=1 | 0 | 35.468 | 34 | 7 | 9 | 7 | 334 | 35.6 | 6.21 | 0.491 | 0.343 | 1.432 | -1.03 | 1.03 | -1.54 | -0.5133333 | 1.285 | 0.52 | 0.012836373 | 0.000756452 | 0.037396201 | 0.17903866 | 0.029454338 | 0.104246499 | 0.373579725 |
| Tmsb4x | High | P20065 | Thymosin beta-4 OS=Mus musculus OX=10090 GN=Tmsb4x PE=1 SV=1 | 0 | 17.819 | 40 | 6 | 9 | 6 | 50 | 5.7 | 5.06 | 0.47 | 0.599 | 0.786 | -1.09 | 1.09 | -0.74 | -0.2466667 | 0.915 | -0.35 | 0.012902473 | 0.096877068 | 0.158652806 | 0.17951406 | 0.714723313 | 0.447118686 | 0.733684585 |
| Chmp2a | High | Q9DB34 | Charged multivesicular body protein 2a OS=Mus musculus OX=10090 GN=Chmp2a PE=1 SV=1 | 0 | 81.999 | 42 | 14 | 38 | 14 | 222 | 25.1 | 5.97 | 0.479 | 0.424 | 1.129 | -1.06 | 1.06 | -1.24 | -0.4133333 | 1.15 | 0.18 | 0.013480536 | 0.006637089 | 0.37621567 | 0.186630535 | 0.169155269 | 0.177892902 | 0.92034959 |
| Slc9a1 | High | Q61165 | Sodium/hydrogen exchanger 1 OS=Mus musculus OX=10090 GN=Slc9a1 PE=1 SV=1 | 0.001 | 5.388 | 3 | 2 | 2 | 2 | 820 | 91.4 | 7.12 | 0.521 | 0.431 | 1.208 | -0.94 | 0.94 | -1.21 | -0.4033333 | 1.075 | 0.27 | 0.013449594 | 0.007630159 | 0.222030434 | 0.186630535 | 0.176353762 | 0.181492149 | 0.807447486 |
| Arrb2 | High | Q5F2D9 | Arrestin, beta 2 OS=Mus musculus OX=10090 GN=Arrb2 PE=1 SV=1 | 0 | 10.973 | 9 | 2 | 2 | 2 | 406 | 45.7 | 7.83 | 2.057 | 1.876 | 1.096 | 1.04 | -1.04 | 0.91 | 0.30333333 | -0.975 | 0.13 | 0.013761566 | 0.06464676 | 0.601584554 | 0.188197801 | 0.58835905 | 0.388278426 | 0.97826833 |
| Fcgbp | High | E9Q0B5 | Fc fragment of IgG-binding protein OS=Mus musculus OX=10090 GN=Fcgbp PE=1 SV=1 | 0.003 | 4.077 | 0 | 1 | 4 | 1 | 2583 | 275 | 5.03 | 2.145 | 1.84 | 1.166 | 1.1 | -1.1 | 0.88 | 0.29333333 | -0.99 | 0.22 | 0.013747048 | 0.074361992 | 0.328460036 | 0.188197801 | 0.636899913 | 0.412548857 | 0.905686588 |
| Arpin | High | Q9D0A3 | Arpin OS=Mus musculus OX=10090 GN=Arpin PE=1 SV=1 | 0.001 | 6.778 | 8 | 2 | 2 | 2 | 226 | 25.2 | 5.19 | 0.559 | 0.522 | 1.07 | -0.84 | 0.84 | -0.94 | -0.3133333 | 0.89 | 0.1 | 0.014303109 | 0.033782747 | 0.706700673 | 0.195127813 | 0.473899991 | 0.334513902 | 0.992485323 |
| Ift140 | High | E9PY46 | Intraflagellar transport protein 140 homolog OS=Mus musculus OX=10090 GN=Ift140 PE=1 SV=1 | 0 | 37.533 | 9 | 15 | 15 | 15 | 1464 | 165.8 | 5.91 | 1.98 | 1.807 | 1.096 | 0.99 | -0.99 | 0.85 | 0.28333333 | -0.92 | 0.13 | 0.014688585 | 0.083775881 | 0.491021128 | 0.199416211 | 0.680770092 | 0.440093152 | 0.964378027 |
| B9d2 | High | Q3UK10 | B9 domain-containing protein 2 OS=Mus musculus OX=10090 GN=B9d2 PE=1 SV=2 | 0 | 15.313 | 25 | 3 | 4 | 3 | 175 | 19.2 | 7.58 | 1.994 | 2.467 | 0.808 | 1 | -1 | 1.3 | 0.43333333 | -1.15 | -0.31 | 0.015126443 | 0.007284547 | 0.208858306 | 0.20486465 | 0.174607618 | 0.189736134 | 0.801552719 |
| Zc2hc1a | High | Q8BJH1 | Zinc finger C2HC domain-containing protein 1A OS=Mus musculus OX=10090 GN=Zc2hc1a PE=1 S | 0 | 12.867 | 13 | 3 | 3 | 3 | 324 | 35.1 | 9.88 | 0.523 | 0.508 | 1.03 | -0.93 | 0.93 | -0.98 | -0.3266667 | 0.955 | 0.04 | 0.015516221 | 0.030662629 | 0.804420276 | 0.209637229 | 0.446559375 | 0.328098302 | 0.992535481 |
| Ift81 | High | O35594 | Intraflagellar transport protein 81 homolog OS=Mus musculus OX=10090 GN=Ift81 PE=1 SV=4 | 0 | 56.752 | 26 | 17 | 17 | 17 | 676 | 79.2 | 8.75 | 2.023 | 2.16 | 0.936 | 1.02 | -1.02 | 1.11 | 0.37 | -1.065 | -0.09 | 0.015745389 | 0.022796323 | 0.750458779 | 0.211206686 | 0.379284818 | 0.295245752 | 0.992535481 |
| Cep250 | High | A3KGJ7 | Centrosome-associated protein CEP250 OS=Mus musculus OX=10090 GN=Cep250 PE=1 SV=2 | 0 | 68.477 | 10 | 23 | 24 | 23 | 2435 | 278.8 | 5.05 | 2.061 | 2.035 | 1.013 | 1.04 | -1.04 | 1.02 | 0.34 | -1.03 | 0.02 | 0.016326105 | 0.036360001 | 0.854507653 | 0.217435799 | 0.476392832 | 0.346914316 | 0.992535481 |
| Chmp2b | High | Q8BJF9 | Charged multivesicular body protein 2b OS=Mus musculus OX=10090 GN=Chmp2b PE=1 SV=1 | 0 | 29.027 | 31 | 8 | 13 | 8 | 213 | 23.9 | 8.78 | 0.488 | 0.443 | 1.102 | -1.04 | 1.04 | -1.18 | -0.3933333 | 1.11 | 0.14 | 0.016727143 | 0.009761801 | 0.544317441 | 0.220680215 | 0.214644778 | 0.217662496 | 0.96694628 |
| Nadk2-fr | High | A0A7N9VS | NAD kinase 2, mitochondrial (Fragment) OS=Mus musculus OX=10090 GN=Nadk2 PE=4 SV=1 | 0 | 18.644 | 12 | 3 | 4 | 3 | 343 | 39 | 6.05 | 1.906 | 1.556 | 1.225 | 0.93 | -0.93 | 0.64 | 0.21333333 | -0.785 | 0.29 | 0.016694013 | 0.198204523 | 0.168922698 | 0.220680215 | 0.912882441 | 0.566781328 | 0.738806217 |
| Inpp1 | High | P49442 | Inositol polyphosphate 1-phosphatase OS=Mus musculus OX=10090 GN=Inpp1 PE=1 SV=2 | 0.005 | 3.528 | 2 | 1 | 2 | 1 | 396 | 43.3 | 5.01 | 0.584 | 0.624 | 0.935 | -0.78 | 0.78 | -0.68 | -0.2266667 | 0.73 | -0.1 | 0.016917144 | 0.125525581 | 0.732570142 | 0.222662966 | 0.799286338 | 0.510974652 | 0.992535481 |
| Sectm1a | Low | A2ABP7 | Secreted and transmembrane protein 1A OS=Mus musculus OX=10090 GN=Sectm1a PE=1 SV=1 | 0.077 | 1.229 | 9 | 1 | 1 | 1 | 174 | 19.4 | 8.97 | 0.549 | 0.804 | 0.683 | -0.87 | 0.87 | -0.32 | -0.1066667 | 0.595 | -0.55 | 0.017500794 | 0.427671347 | 0.036455492 | 0.228734155 | 0.99399649 | 0.611365323 | 0.370300617 |
| Cnnm4 | High | Q69ZF7 | Metal transporter CNNM4 OS=Mus musculus OX=10090 GN=Cnnm4 PE=1 SV=2 | 0 | 98.46 | 34 | 25 | 35 | 20 | 771 | 86.6 | 6.07 | 0.493 | 0.394 | 1.252 | -1.02 | 1.02 | -1.34 | -0.4466667 | 1.18 | 0.32 | 0.017586251 | 0.003246141 | 0.12093853 | 0.229316537 | 0.100411377 | 0.164863957 | 0.669174681 |
| Ankrd33 | High | Q8BXP5 | Photoreceptor ankyrin repeat protein OS=Mus musculus OX=10090 GN=Ankrd33 PE=1 SV=2 | 0 | 35.08 | 23 | 7 | 10 | 7 | 385 | 41.6 | 6.71 | 2.06 | 2.223 | 0.927 | 1.04 | -1.04 | 1.15 | 0.38333333 | -1.095 | -0.11 | 0.017655249 | 0.018030584 | 0.695267155 | 0.229682096 | 0.315929647 | 0.272805871 | 0.991285312 |
| Ttc21b | High | Q0HA38 | Tetratricopeptide repeat protein 21B OS=Mus musculus OX=10090 GN=Ttc21b PE=1 SV=1 | 0 | 10.821 | 4 | 6 | 6 | 6 | 1315 | 150.7 | 7.14 | 1.971 | 1.61 | 1.224 | 0.98 | -0.98 | 0.69 | 0.23 | -0.835 | 0.29 | 0.018010462 | 0.169319027 | 0.232815665 | 0.233760789 | 0.874947902 | 0.554354345 | 0.821004674 |
| Tubb2b | High | Q9CWF2 | Tubulin beta-2B chain OS=Mus musculus OX=10090 GN=Tubb2b PE=1 SV=1 | 0 | 334.014 | 73 | 25 | 177 | 1 | 445 | 49.9 | 4.89 | 1.962 | 2.279 | 0.861 | 0.97 | -0.97 | 1.19 | 0.39666667 | -1.08 | -0.22 | 0.018075193 | 0.0146806 | 0.381128595 | 0.23405914 | 0.282866406 | 0.258462773 | 0.923104982 |
| Gars | High | Q9CZD3 | Glycine--tRNA ligase OS=Mus musculus OX=10090 GN=Gars1 PE=1 SV=1 | 0 | 64.98 | 18 | 13 | 25 | 13 | 729 | 81.8 | 6.65 | 0.495 | 0.467 | 1.061 | -1.01 | 1.01 | -1.1 | -0.3666667 | 1.055 | 0.09 | 0.018197625 | 0.015422102 | 0.639033545 | 0.235101568 | 0.292134205 | 0.263617886 | 0.983597794 |

| | | | | | | | | | | | | | | | | | | | | | | | | | | | | |
|---|---|---|---|---|---|---|---|---|---|---|---|---|---|---|---|---|---|---|---|---|---|---|---|---|---|---|---|---|
| Flvcr2 | High | A0A0R4J0 | Feline leukemia virus subgroup C receptor-related protein 2 OS=Mus musculus OX=10090 GN=Flv | 0.009 | 2.767 | 2 | 1 | 2 | 1 | 551 | 60 | 6.42 | 1.825 | 2.143 | 0.852 | 0.87 | -0.87 | 1.1 | 0.36666667 | -0.985 | -0.23 | 0.018393443 | 0.024349656 | 0.304420298 | 0.23654137 | 0.384371506 | 0.310456438 | 0.900241969 |
| Slc22a8 | High | O88909 | Solute carrier family 22 member 8 OS=Mus musculus OX=10090 GN=Slc22a8 PE=1 SV=2 | 0 | 31.754 | 17 | 10 | 13 | 10 | 537 | 59.2 | 8.4 | 1.949 | 1.811 | 1.076 | 0.96 | -0.96 | 0.86 | 0.28666667 | -0.91 | 0.11 | 0.018589956 | 0.082628257 | 0.565898303 | 0.238521475 | 0.673396277 | 0.455958876 | 0.972910028 |
| Tom1l2 | High | Q5SRX1 | TOM1-like protein 2 OS=Mus musculus OX=10090 GN=Tom1l2 PE=1 SV=1 | 0 | 166.679 | 57 | 20 | 46 | 20 | 507 | 55.6 | 4.82 | 0.497 | 0.501 | 0.993 | -1.01 | 1.01 | -1 | -0.3333333 | 1.005 | -0.01 | 0.018880533 | 0.027277696 | 0.956094309 | 0.241696687 | 0.413367685 | 0.327532186 | 0.992902921 |
| Fnta | High | Q61239 | Protein farnesyltransferase/geranylgeranyltransferase type-1 subunit alpha OS=Mus musculus OX | 0 | 9.249 | 10 | 2 | 2 | 2 | 377 | 44 | 4.93 | 0.538 | 0.746 | 0.721 | -0.89 | 0.89 | -0.42 | -0.14 | 0.655 | -0.47 | 0.019022924 | 0.304391752 | 0.073897692 | 0.242964774 | 0.955820031 | 0.599392402 | 0.519879998 |
| Tspan31 | High | Q9CQ88 | Tetraspanin-31 OS=Mus musculus OX=10090 GN=Tspan31 PE=1 SV=1 | 0 | 8.135 | 10 | 2 | 2 | 2 | 210 | 22.7 | 8.44 | 0.57 | 0.513 | 1.111 | -0.81 | 0.81 | -0.96 | -0.32 | 0.885 | 0.15 | 0.019693521 | 0.026728057 | 0.515020104 | 0.250958123 | 0.407239716 | 0.32909892 | 0.964378027 |
| Myo7a | Medium | P97479 | Unconventional myosin-VIIa OS=Mus musculus OX=10090 GN=Myo7a PE=1 SV=2 | 0.011 | 2.468 | 0 | 1 | 1 | 1 | 2215 | 254.8 | 8.54 | 0.555 | 0.532 | 1.043 | -0.85 | 0.85 | -0.91 | -0.3033333 | 0.88 | 0.06 | 0.020076066 | 0.035763949 | 0.794339044 | 0.253528163 | 0.476392832 | 0.364960497 | 0.992535481 |
| Tom1 | High | Q3UDC3 | Target of Myb protein 1 OS=Mus musculus OX=10090 GN=Tom1 PE=1 SV=1 | 0 | 74.073 | 41 | 13 | 22 | 13 | 516 | 57.1 | 4.97 | 0.521 | 0.491 | 1.062 | -0.94 | 0.94 | -1.03 | -0.3433333 | 0.985 | 0.09 | 0.020863624 | 0.023226714 | 0.735885859 | 0.261705463 | 0.379812885 | 0.320759174 | 0.992535481 |
| Vta1 | High | Q9CR26 | Vacuolar protein sorting-associated protein VTA1 homolog OS=Mus musculus OX=10090 GN=Vta | 0 | 34.212 | 27 | 6 | 8 | 6 | 309 | 33.9 | 6.13 | 0.539 | 0.559 | 0.963 | -0.89 | 0.89 | -0.84 | -0.28 | 0.865 | -0.05 | 0.020975849 | 0.061823215 | 0.820525941 | 0.261952272 | 0.586560875 | 0.424256573 | 0.992535481 |
| Ptges2 | High | Q8BWM0 | Prostaglandin E synthase 2 OS=Mus musculus OX=10090 GN=Ptges2 PE=1 SV=3 | 0.006 | 3.384 | 2 | 1 | 1 | 1 | 384 | 43.3 | 9 | 1.831 | 1.611 | 1.136 | 0.87 | -0.87 | 0.69 | 0.23 | -0.78 | 0.18 | 0.020976738 | 0.156414127 | 0.475889416 | 0.261952272 | 0.847684847 | 0.554818559 | 0.962227984 |
| Cdc37 | High | Q61081 | Hsp90 co-chaperone Cdc37 OS=Mus musculus OX=10090 GN=Cdc37 PE=1 SV=1 | 0 | 24.892 | 19 | 6 | 7 | 6 | 379 | 44.6 | 5.34 | 0.559 | 0.731 | 0.764 | -0.84 | 0.84 | -0.45 | -0.15 | 0.645 | -0.39 | 0.021043567 | 0.294580561 | 0.139670828 | 0.262202839 | 0.955820031 | 0.609011435 | 0.701851981 |
| Man2a2 | High | Q8BRK9 | Alpha-mannosidase 2x OS=Mus musculus OX=10090 GN=Man2a2 PE=1 SV=2 | 0 | 171.496 | 37 | 32 | 55 | 32 | 1152 | 130.6 | 6.87 | 1.977 | 2.288 | 0.864 | 0.98 | -0.98 | 1.19 | 0.39666667 | -1.085 | -0.21 | 0.021644027 | 0.01417922 | 0.400336592 | 0.268324868 | 0.275999641 | 0.272162255 | 0.93262326 |
| Tnpo1 | High | Q3TKD0 | Transportin-1 (Fragment) OS=Mus musculus OX=10090 GN=Tnpo1 PE=1 SV=1 | 0 | 10.121 | 4 | 3 | 5 | 1 | 807 | 91.7 | 4.92 | 0.536 | 0.557 | 0.963 | -0.9 | 0.9 | -0.84 | -0.28 | 0.87 | -0.05 | 0.021790324 | 0.05439846 | 0.889544508 | 0.268324868 | 0.561716694 | 0.415020781 | 0.992535481 |
| Vps26a | High | P40336 | Vacuolar protein sorting-associated protein 26A OS=Mus musculus OX=10090 GN=Vps26a PE=1 S | 0.002 | 4.197 | 4 | 1 | 1 | 1 | 327 | 38.1 | 6.57 | 0.596 | 0.635 | 0.938 | -0.75 | 0.75 | -0.65 | -0.2166667 | 0.7 | -0.09 | 0.021822033 | 0.132535396 | 0.765264073 | 0.268324868 | 0.808473523 | 0.538399196 | 0.992535481 |
| Stam2 | High | O88811 | Signal transducing adapter molecule 2 OS=Mus musculus OX=10090 GN=Stam2 PE=1 SV=1 | 0.001 | 5.166 | 3 | 2 | 2 | 1 | 523 | 57.4 | 5.07 | 0.578 | 0.649 | 0.89 | -0.79 | 0.79 | -0.62 | -0.2066667 | 0.705 | -0.17 | 0.021637511 | 0.157806281 | 0.4961 | 0.268324868 | 0.85160714 | 0.559966004 | 0.964378027 |
| Rap1a | High | P62835 | Ras-related protein Rap-1A OS=Mus musculus OX=10090 GN=Rap1a PE=1 SV=1 | 0 | 84.187 | 53 | 11 | 37 | 3 | 184 | 21 | 6.67 | 0.506 | 0.499 | 1.014 | -0.98 | 0.98 | -1 | -0.3333333 | 0.99 | 0.02 | 0.02203429 | 0.026524411 | 0.849854315 | 0.270341929 | 0.407239716 | 0.338790823 | 0.992535481 |
| Eif3l | High | Q8QZY1 | Eukaryotic translation initiation factor 3 subunit L OS=Mus musculus OX=10090 GN=Eif3l PE=1 SV | 0 | 9.662 | 7 | 3 | 3 | 3 | 564 | 66.6 | 6.44 | 1.797 | 1.576 | 1.14 | 0.85 | -0.85 | 0.66 | 0.22 | -0.755 | 0.19 | 0.022094034 | 0.178252803 | 0.399189202 | 0.27048308 | 0.875953959 | 0.57321852 | 0.93262326 |
| Afdn | High | E9Q852 | Afadin OS=Mus musculus OX=10090 GN=Afdn PE=1 SV=2 | 0.001 | 5.578 | 1 | 2 | 2 | 2 | 1805 | 204.4 | 6.24 | 0.586 | 0.603 | 0.973 | -0.77 | 0.77 | -0.73 | -0.2433333 | 0.75 | -0.04 | 0.022541844 | 0.095196067 | 0.873999745 | 0.27298514 | 0.707910274 | 0.490447707 | 0.992535481 |
| Trim28 | High | Q62318 | Transcription intermediary factor 1-beta OS=Mus musculus OX=10090 GN=Trim28 PE=1 SV=3 | 0.001 | 5.848 | 3 | 3 | 3 | 3 | 834 | 88.8 | 5.77 | 1.897 | 1.879 | 1.01 | 0.92 | -0.92 | 0.91 | 0.30333333 | -0.915 | 0.01 | 0.022695676 | 0.058545629 | 0.905853984 | 0.274255724 | 0.571608376 | 0.42293205 | 0.992535481 |
| Pibf1 | High | E9Q6K3 | Progesterone immunomodulatory-binding factor 1 OS=Mus musculus OX=10090 GN=Pibf1 PE=1 S | 0.001 | 6.17 | 2 | 2 | 2 | 2 | 756 | 89.5 | 6.16 | 1.846 | 2.202 | 0.838 | 0.88 | -0.88 | 1.14 | 0.38 | -1.01 | -0.25 | 0.022866778 | 0.018017089 | 0.345533429 | 0.275137397 | 0.315929647 | 0.295533522 | 0.907770836 |
| Slc25a5 | High | P51881 | ADP/ATP translocase 2 OS=Mus musculus OX=10090 GN=Slc25a5 PE=1 SV=3 | 0 | 88.794 | 50 | 17 | 46 | 8 | 298 | 32.9 | 9.73 | 1.9 | 2.045 | 0.929 | 0.93 | -0.93 | 1.03 | 0.34333333 | -0.98 | -0.11 | 0.022819651 | 0.03495922 | 0.708368806 | 0.275137397 | 0.476392832 | 0.375765115 | 0.992507514 |
| Stx8 | High | O88983 | Syntaxin-8 OS=Mus musculus OX=10090 GN=Stx8 PE=1 SV=1 | 0 | 13.132 | 18 | 4 | 4 | 4 | 236 | 26.9 | 5.01 | 0.563 | 0.535 | 1.051 | -0.83 | 0.83 | -0.9 | -0.3 | 0.865 | 0.07 | 0.02319186 | 0.045249075 | 0.756529819 | 0.276086536 | 0.520047141 | 0.398066838 | 0.992535481 |
| Arl4d | High | Q99PE9 | ADP-ribosylation factor-like protein 4D OS=Mus musculus OX=10090 GN=Arl4d PE=2 SV=2 | 0.003 | 3.931 | 7 | 1 | 1 | 1 | 201 | 22.3 | 9.85 | 1.844 | 1.395 | 1.322 | 0.88 | -0.88 | 0.48 | 0.16 | -0.68 | 0.4 | 0.023088365 | 0.339157704 | 0.100486208 | 0.276086536 | 0.976045759 | 0.626066147 | 0.609768578 |
| Serf2 | High | P84102 | Small EDRK-rich factor 2 OS=Mus musculus OX=10090 GN=Serf2 PE=1 SV=1 | 0 | 4.737 | 25 | 2 | 2 | 2 | 59 | 6.9 | 10.45 | 1.835 | 1.482 | 1.239 | 0.88 | -0.88 | 0.57 | 0.19 | -0.725 | 0.31 | 0.024606963 | 0.257739052 | 0.178734641 | 0.291759839 | 0.951072422 | 0.621416131 | 0.755780643 |
| Krt5 | High | Q922U2 | Keratin, type II cytoskeletal 5 OS=Mus musculus OX=10090 GN=Krt5 PE=1 SV=1 | 0 | 42.705 | 18 | 13 | 27 | 5 | 580 | 61.7 | 7.75 | 1.897 | 1.336 | 1.42 | 0.92 | -0.92 | 0.42 | 0.14 | -0.67 | 0.51 | 0.024612521 | 0.422162646 | 0.0182793 | 0.291759839 | 0.99399649 | 0.642878165 | 0.237800551 |
| Ubap1 | High | Q8BH48 | Ubiquitin-associated protein 1 OS=Mus musculus OX=10090 GN=Ubap1 PE=1 SV=1 | 0 | 8.137 | 5 | 3 | 3 | 3 | 502 | 55 | 5.07 | 0.548 | 0.519 | 1.056 | -0.87 | 0.87 | -0.95 | -0.3166667 | 0.91 | 0.08 | 0.024961895 | 0.036170029 | 0.688703372 | 0.292040545 | 0.476392832 | 0.384216688 | 0.991285312 |
| Osmr | High | A0A0R4J2 | Oncostatin-M-specific receptor subunit beta OS=Mus musculus OX=10090 GN=Osmr PE=1 SV=1 | 0 | 11.68 | 6 | 6 | 7 | 6 | 970 | 110 | 6.65 | 0.57 | 0.588 | 0.969 | -0.81 | 0.81 | -0.77 | -0.2566667 | 0.79 | -0.05 | 0.025000796 | 0.086682442 | 0.870447211 | 0.292040545 | 0.68454712 | 0.488293832 | 0.992535481 |
| Rims2 | High | D9HP81 | Regulating synaptic membrane exocytosis 2 OS=Mus musculus OX=10090 GN=Rims2 PE=1 SV=1 | 0.003 | 4.09 | 1 | 2 | 2 | 1 | 1572 | 177.5 | 9.36 | 1.8 | 1.427 | 1.261 | 0.85 | -0.85 | 0.51 | 0.17 | -0.68 | 0.33 | 0.024968706 | 0.304668161 | 0.174992952 | 0.292040545 | 0.955820031 | 0.623930288 | 0.744450291 |
| Rhot1 | High | Q8BG51 | Mitochondrial Rho GTPase 1 OS=Mus musculus OX=10090 GN=Rhot1 PE=1 SV=1 | 0 | 16.663 | 9 | 5 | 6 | 4 | 631 | 72.2 | 6.49 | 1.926 | 2.072 | 0.929 | 0.95 | -0.95 | 1.05 | 0.35 | -1 | -0.11 | 0.025221221 | 0.031640228 | 0.684254255 | 0.294002885 | 0.457149627 | 0.375576256 | 0.991285312 |
| Rock2 | High | P70336 | Rho-associated protein kinase 2 OS=Mus musculus OX=10090 GN=Rock2 PE=1 SV=1 | 0.001 | 7.665 | 2 | 3 | 3 | 3 | 1388 | 160.5 | 5.99 | 0.574 | 0.816 | 0.703 | -0.8 | 0.8 | -0.29 | -0.0966667 | 0.545 | -0.51 | 0.025479787 | 0.454395579 | 0.051905291 | 0.295787095 | 0.995559182 | 0.645673138 | 0.431159947 |
| Vldlr | High | F8WGI9 | Very low-density lipoprotein receptor OS=Mus musculus OX=10090 GN=Vldlr PE=1 SV=1 | 0.002 | 4.153 | 1 | 1 | 1 | 1 | 804 | 88.9 | 4.84 | 0.581 | 0.55 | 1.057 | -0.78 | 0.78 | -0.86 | -0.2866667 | 0.82 | 0.08 | 0.026768816 | 0.049808644 | 0.761595796 | 0.306533837 | 0.52893384 | 0.417733839 | 0.992535481 |
| Ksr1-fr | High | A0A140LH | Kinase suppressor of Ras 1 (Fragment) OS=Mus musculus OX=10090 GN=Ksr1 PE=1 SV=1 | 0.001 | 5.063 | 4 | 2 | 2 | 2 | 432 | 47.7 | 9.67 | 0.57 | 0.556 | 1.026 | -0.81 | 0.81 | -0.85 | -0.2833333 | 0.83 | 0.04 | 0.026842895 | 0.059035104 | 0.862259857 | 0.306533837 | 0.571608376 | 0.439071107 | 0.992535481 |
| Ptges3 | High | D3Z7C6 | Prostaglandin E synthase 3 OS=Mus musculus OX=10090 GN=Ptges3 PE=1 SV=1 | 0 | 19.569 | 31 | 4 | 6 | 4 | 130 | 15 | 4.79 | 0.568 | 0.54 | 1.052 | -0.82 | 0.82 | -0.89 | -0.2966667 | 0.855 | 0.07 | 0.027225551 | 0.048411103 | 0.758360258 | 0.309642318 | 0.523007814 | 0.416325066 | 0.992535481 |
| Fkbp3 | High | Q62446 | Peptidyl-prolyl cis-trans isomerase FKBP3 OS=Mus musculus OX=10090 GN=Fkbp3 PE=1 SV=2 | 0 | 99.56 | 65 | 16 | 37 | 16 | 224 | 25.1 | 9.28 | 1.924 | 1.391 | 1.383 | 0.94 | -0.94 | 0.48 | 0.16 | -0.71 | 0.47 | 0.027484716 | 0.354128714 | 0.028573468 | 0.311957094 | 0.977313184 | 0.644635139 | 0.32300692 |
| Hba-a1; Hl | High | Q91VB8 | Alpha globin 1 OS=Mus musculus OX=10090 GN=Hba-a1 PE=1 SV=1 | 0 | 31.018 | 49 | 4 | 10 | 4 | 142 | 15.1 | 8.22 | 1.921 | 1.791 | 1.073 | 0.94 | -0.94 | 0.84 | 0.28 | -0.89 | 0.1 | 0.028554228 | 0.088880676 | 0.580449931 | 0.323441526 | 0.696517952 | 0.509979739 | 0.972910028 |
| Sfxn5 | High | Q925N0 | Sideroflexin-5 OS=Mus musculus OX=10090 GN=Sfxn5 PE=1 SV=2 | 0 | 33.277 | 26 | 8 | 13 | 8 | 342 | 37.3 | 9.44 | 1.944 | 1.8 | 1.08 | 0.96 | -0.96 | 0.85 | 0.28333333 | -0.905 | 0.11 | 0.028865918 | 0.085919037 | 0.606708922 | 0.325656344 | 0.68454712 | 0.505101732 | 0.97826833 |
| Acadvl | High | P50544 | Very long-chain specific acyl-CoA dehydrogenase, mitochondrial OS=Mus musculus OX=10090 GN | 0 | 14.637 | 7 | 4 | 4 | 4 | 656 | 70.8 | 8.75 | 1.81 | 1.557 | 1.163 | 0.86 | -0.86 | 0.64 | 0.21333333 | -0.75 | 0.22 | 0.029106689 | 0.196287762 | 0.329032206 | 0.327056522 | 0.912882441 | 0.619969481 | 0.905686588 |
| Ubr2 | High | Q6WKZ8 | E3 ubiquitin-protein ligase UBR2 OS=Mus musculus OX=10090 GN=Ubr2 PE=1 SV=2 | 0 | 8.131 | 2 | 3 | 4 | 3 | 1755 | 199 | 6.33 | 0.52 | 0.595 | 0.873 | -0.94 | 0.94 | -0.75 | -0.25 | 0.845 | -0.2 | 0.02948515 | 0.093738742 | 0.448742835 | 0.329393447 | 0.703605253 | 0.51649935 | 0.958608248 |
| Cep104 | High | Q80V31 | Centrosomal protein of 104 kDa OS=Mus musculus OX=10090 GN=Cep104 PE=1 SV=1 | 0 | 14.496 | 6 | 6 | 6 | 5 | 926 | 103.9 | 8.15 | 1.873 | 1.73 | 1.082 | 0.91 | -0.91 | 0.79 | 0.26333333 | -0.85 | 0.11 | 0.029490906 | 0.110648895 | 0.647433609 | 0.329393447 | 0.769444739 | 0.549419093 | 0.985010103 |
| Isoc2a | High | P85094 | Isochorismatase domain-containing protein 2A OS=Mus musculus OX=10090 GN=Isoc2a PE=1 SV= | 0 | 8.376 | 9 | 2 | 4 | 2 | 206 | 22.4 | 8.02 | 1.78 | 1.434 | 1.241 | 0.83 | -0.83 | 0.52 | 0.17333333 | -0.675 | 0.31 | 0.029794852 | 0.308205901 | 0.208257497 | 0.332126707 | 0.955820031 | 0.643973369 | 0.801370196 |
| Lyrm4 | Medium | Q8K215 | LYR motif-containing protein 4 OS=Mus musculus OX=10090 GN=Lyrm4 PE=1 SV=1 | 0.011 | 2.426 | 15 | 1 | 1 | 1 | 91 | 10.8 | 10.13 | 1.815 | 1.656 | 1.096 | 0.86 | -0.86 | 0.73 | 0.24333333 | -0.795 | 0.13 | 0.030071715 | 0.138507171 | 0.581120666 | 0.334547826 | 0.82258226 | 0.578565043 | 0.972910028 |
| Stam | High | P70297 | Signal transducing adapter molecule 1 OS=Mus musculus OX=10090 GN=Stam PE=1 SV=3 | 0.001 | 5.772 | 3 | 2 | 3 | 1 | 548 | 59.7 | 4.84 | 0.578 | 0.729 | 0.793 | -0.79 | 0.79 | -0.46 | -0.1533333 | 0.625 | -0.33 | 0.030231448 | 0.264335083 | 0.201166202 | 0.33499551 | 0.951072422 | 0.643033966 | 0.78697088 |
| Slco1a6 | High | Q99J94 | Solute carrier organic anion transporter family member 1A6 OS=Mus musculus OX=10090 GN=Slc | 0.001 | 5.159 | 3 | 2 | 3 | 1 | 670 | 74.1 | 7.99 | 0.563 | 0.689 | 0.817 | -0.83 | 0.83 | -0.54 | -0.18 | 0.685 | -0.29 | 0.030568628 | 0.207230606 | 0.365456877 | 0.338063697 | 0.912882441 | 0.625473069 | 0.920124252 |
| Micos13 | High | Q8R404 | MICOS complex subunit MIC13 OS=Mus musculus OX=10090 GN=Micos13 PE=1 SV=1 | 0.001 | 5.284 | 23 | 2 | 2 | 2 | 119 | 13.4 | 8.63 | 1.779 | 1.847 | 0.964 | 0.83 | -0.83 | 0.88 | 0.29333333 | -0.855 | -0.05 | 0.030711131 | 0.067864182 | 0.876788967 | 0.338971083 | 0.603034023 | 0.471002553 | 0.992535481 |
| Tulp1 | High | Q9Z273 | Tubby-related protein 1 OS=Mus musculus OX=10090 GN=Tulp1 PE=1 SV=1 | 0 | 152.603 | 55 | 28 | 53 | 25 | 543 | 60.3 | 9.1 | 1.895 | 1.439 | 1.317 | 0.92 | -0.92 | 0.53 | 0.17666667 | -0.725 | 0.4 | 0.031262563 | 0.302715006 | 0.061017565 | 0.344379548 | 0.955820031 | 0.650099789 | 0.476099813 |
| Krt78 | High | E9Q0F0 | Keratin 78 OS=Mus musculus OX=10090 GN=Krt78 PE=1 SV=1 | 0 | 25.259 | 2 | 3 | 12 | 2 | 1068 | 112.2 | 7.97 | 1.782 | 1.183 | 1.506 | 0.83 | -0.83 | 0.24 | 0.08 | -0.535 | 0.59 | 0.031359256 | 0.666593624 | 0.017066251 | 0.344767347 | 0.995559182 | 0.670163264 | 0.228924566 |
| Hibadh | High | Q99L13 | 3-hydroxyisobutyrate dehydrogenase, mitochondrial OS=Mus musculus OX=10090 GN=Hibadh PE= | 0 | 47.028 | 26 | 6 | 8 | 6 | 335 | 35.4 | 8.13 | 1.902 | 1.881 | 1.011 | 0.93 | -0.93 | 0.91 | 0.30333333 | -0.92 | 0.02 | 0.031516185 | 0.063980062 | 0.913870938 | 0.345814577 | 0.58835905 | 0.467086813 | 0.992535481 |
| Hadha | High | Q8BMS1 | Trifunctional enzyme subunit alpha, mitochondrial OS=Mus musculus OX=10090 GN=Hadha PE=1 | 0 | 32.21 | 20 | 9 | 10 | 9 | 763 | 82.6 | 9.14 | 1.848 | 1.781 | 1.038 | 0.89 | -0.89 | 0.83 | 0.27666667 | -0.86 | 0.05 | 0.031867354 | 0.092248488 | 0.84404133 | 0.348984868 | 0.698969289 | 0.523977078 | 0.992535481 |
| Ahi1 | High | A0A1L1SQ | Jouberin OS=Mus musculus OX=10090 GN=Ahi1 PE=1 SV=1 | 0 | 34.089 | 8 | 6 | 11 | 6 | 1005 | 114.9 | 6.9 | 1.828 | 2.017 | 0.906 | 0.87 | -0.87 | 1.01 | 0.33666667 | -0.94 | -0.14 | 0.032250299 | 0.038808252 | 0.546736525 | 0.349640059 | 0.486480307 | 0.418060183 | 0.96694628 |
| Ttpal | High | Q9D3D0 | Alpha-tocopherol transfer protein-like OS=Mus musculus OX=10090 GN=Ttpal PE=1 SV=3 | 0.001 | 4.327 | 6 | 2 | 2 | 2 | 343 | 38.8 | 6.44 | 1.788 | 1.844 | 0.97 | 0.84 | -0.84 | 0.88 | 0.29333333 | -0.86 | -0.04 | 0.032301329 | 0.067746052 | 0.877782342 | 0.349640059 | 0.603034023 | 0.476337041 | 0.992535481 |
| Cadm2-fr | High | G3UZM4 | Cell adhesion molecule 2 (Fragment) OS=Mus musculus OX=10090 GN=Cadm2 PE=1 SV=1 | 0 | 73.164 | 31 | 10 | 18 | 10 | 435 | 47.3 | 5.41 | 0.519 | 0.717 | 0.723 | -0.95 | 0.95 | -0.48 | -0.16 | 0.715 | -0.47 | 0.032192179 | 0.269476902 | 0.046591837 | 0.349640059 | 0.953285167 | 0.651462613 | 0.409467755 |
| Zc3h3 | High | Q8CHP0 | Zinc finger CCCH domain-containing protein 3 OS=Mus musculus OX=10090 GN=Zc3h3 PE=1 SV=1 | 0 | 9.396 | 6 | 4 | 4 | 4 | 950 | 103.2 | 10.58 | 1.756 | 1.735 | 1.012 | 0.81 | -0.81 | 0.79 | 0.26333333 | -0.8 | 0.02 | 0.032551926 | 0.102193073 | 0.923567054 | 0.3516737 | 0.734610978 | 0.543142339 | 0.992535481 |
| Nras | High | P08556 | GTPase NRas OS=Mus musculus OX=10090 GN=Nras PE=1 SV=1 | 0 | 138.526 | 66 | 11 | 33 | 2 | 189 | 21.2 | 5.17 | 0.58 | 0.621 | 0.933 | -0.79 | 0.79 | -0.69 | -0.23 | 0.74 | -0.1 | 0.032877786 | 0.122437257 | 0.688082284 | 0.352477529 | 0.795487485 | 0.573982507 | 0.991285312 |
| Smn1 | Medium | P97801 | Survival motor neuron protein OS=Mus musculus OX=10090 GN=Smn1 PE=1 SV=1 | 0.01 | 2.715 | 3 | 1 | 1 | 1 | 288 | 31.2 | 7.62 | 1.788 | 1.627 | 1.099 | 0.84 | -0.84 | 0.7 | 0.23333333 | -0.77 | 0.14 | 0.032772109 | 0.149174469 | 0.582615844 | 0.352477529 | 0.847684847 | 0.600081188 | 0.973095093 |
| Rnf149 | High | Q3U2C5 | E3 ubiquitin-protein ligase RNF149 OS=Mus musculus OX=10090 GN=Rnf149 PE=1 SV=3 | 0 | 10.931 | 7 | 2 | 2 | 2 | 394 | 42.5 | 7.01 | 0.595 | 0.557 | 1.069 | -0.75 | 0.75 | -0.84 | -0.28 | 0.795 | 0.1 | 0.033046031 | 0.047810268 | 0.682449804 | 0.353605147 | 0.520047141 | 0.436826144 | 0.991285312 |
| Gripap1 | Medium | A2AEW9 | GRIP1-associated protein 1 OS=Mus musculus OX=10090 GN=Gripap1 PE=1 SV=1 | 0.039 | 1.706 | 1 | 1 | 1 | 1 | 792 | 90.7 | 5.17 | 0.554 | 0.83 | 0.667 | -0.85 | 0.85 | -0.27 | -0.09 | 0.56 | -0.58 | 0.033907661 | 0.488377342 | 0.067025511 | 0.360759497 | 0.995559182 | 0.678159339 | 0.499896415 |
| Clauap1-fr | High | D3Z1F2 | Clusterin-associated protein 1 (Fragment) OS=Mus musculus OX=10090 GN=Cluap1 PE=1 SV=9 | 0 | 14.248 | 20 | 5 | 5 | 5 | 187 | 21.2 | 8.97 | 1.716 | 1.299 | 1.321 | 0.78 | -0.78 | 0.38 | 0.12666667 | -0.58 | 0.4 | 0.034998757 | 0.473593303 | 0.101879769 | 0.370260432 | 0.995559182 | 0.682909807 | 0.610114517 |
| Lgalsl | High | Q8VED9 | Galectin-related protein OS=Mus musculus OX=10090 GN=Lgalsl PE=1 SV=1 | 0 | 12.472 | 12 | 2 | 3 | 2 | 172 | 18.9 | 5.35 | 0.562 | 0.494 | 1.138 | -0.83 | 0.83 | -1.02 | -0.34 | 0.925 | 0.19 | 0.035163423 | 0.024334195 | 0.380719607 | 0.370603974 | 0.384371506 | 0.37748774 | 0.923104982 |
| Immt | High | E9Q800 | MICOS complex subunit MIC60 OS=Mus musculus OX=10090 GN=Immt PE=1 SV=1 | 0 | 120.515 | 44 | 29 | 40 | 1 | 679 | 75.6 | 7.8 | 1.758 | 1.939 | 0.907 | 0.81 | -0.81 | 0.96 | 0.32 | -0.885 | -0.14 | 0.035109902 | 0.045658401 | 0.615948812 | 0.370603974 | 0.520047141 | 0.445325558 | 0.97826833 |
| Smg5-fr | High | A0A0A6YV | Protein SMG5 (Fragment) OS=Mus musculus OX=10090 GN=Smg5 PE=1 SV=1 | 0.005 | 3.604 | 4 | 1 | 1 | 1 | 180 | 21 | 7.96 | 1.717 | 1.411 | 1.217 | 0.78 | -0.78 | 0.5 | 0.16666667 | -0.64 | 0.28 | 0.035569851 | 0.316303468 | 0.259509046 | 0.373483435 | 0.955820031 | 0.664651733 | 0.85616924 |
| Krt42 | High | Q6IFX2 | Keratin, type I cytoskeletal 42 OS=Mus musculus OX=10090 GN=Krt42 PE=1 SV=1 | 0 | 47.691 | 20 | 9 | 16 | 6 | 452 | 50.1 | 5.16 | 1.723 | 1.831 | 0.941 | 0.79 | -0.79 | 0.87 | 0.29 | -0.83 | -0.09 | 0.035978205 | 0.076812028 | 0.696072885 | 0.377065036 | 0.650581637 | 0.513823337 | 0.991285312 |
| Chmp1a | High | Q921W0 | Charged multivesicular body protein 1a OS=Mus musculus OX=10090 GN=Chmp1a PE=1 SV=1 | 0 | 19.364 | 27 | 7 | 13 | 7 | 196 | 21.6 | 8.06 | 0.524 | 0.529 | 0.991 | -0.93 | 0.93 | -0.92 | -0.3066667 | 0.925 | -0.01 | 0.036596118 | 0.041471456 | 0.968409584 | 0.382825441 | 0.503577691 | 0.443201566 | 0.995075385 |
| Pon1 | High | P52430 | Serum paraoxonase/arylesterase 1 OS=Mus musculus OX=10090 GN=Pon1 PE=1 SV=2 | 0 | 132.766 | 52 | 13 | 58 | 13 | 355 | 39.5 | 5.22 | 1.858 | 2.272 | 0.818 | 0.89 | -0.89 | 1.18 | 0.39333333 | -1.035 | -0.29 | 0.036862565 | 0.015044999 | 0.231345841 | 0.384894604 | 0.288894893 | 0.336894749 | 0.819676582 |
| Gnas | High | P63094 | Guanine nucleotide-binding protein G(s) subunit alpha isoforms short OS=Mus musculus OX=1009 | 0 | 108.021 | 50 | 19 | 54 | 1 | 394 | 45.6 | 5.96 | 0.612 | 0.755 | 0.811 | -0.71 | 0.71 | -0.4 | -0.1333333 | 0.555 | -0.3 | 0.037112373 | 0.341485588 | 0.237636864 | 0.386065073 | 0.977313184 | 0.681689129 | 0.828110564 |
| Nae1 | High | Q8VBW6 | NEDD8-activating enzyme E1 regulatory subunit OS=Mus musculus OX=10090 GN=Nae1 PE=1 SV= | 0.001 | 6.2 | 5 | 2 | 2 | 2 | 534 | 60.2 | 5.52 | 0.599 | 0.515 | 1.165 | -0.74 | 0.74 | -0.96 | -0.32 | 0.85 | 0.22 | 0.037422877 | 0.029618591 | 0.415997487 | 0.387855955 | 0.437030106 | 0.41244303 | 0.936397673 |
| Rbp3 | High | A0A0R4J0 | Retinol-binding protein 3 OS=Mus musculus OX=10090 GN=Rbp3 PE=1 SV=1 | 0 | 503.702 | 64 | 54 | 177 | 54 | 1234 | 134.4 | 5.11 | 0.539 | 0.555 | 0.972 | -0.89 | 0.89 | -0.85 | -0.2833333 | 0.87 | -0.04 | 0.037896684 | 0.058388573 | 0.931879171 | 0.392041895 | 0.571608376 | 0.481825135 | 0.992535481 |
| Tbk1 | High | Q9WUN2 | Serine/threonine-protein kinase TBK1 OS=Mus musculus OX=10090 GN=Tbk1 PE=1 SV=1 | 0 | 19.992 | 6 | 3 | 4 | 3 | 729 | 83.4 | 6.87 | 0.602 | 0.484 | 1.243 | -0.73 | 0.73 | -1.05 | -0.35 | 0.89 | 0.31 | 0.038209478 | 0.020803075 | 0.219803531 | 0.394549801 | 0.351333856 | 0.372941829 | 0.805859554 |
| Mien1 | Medium | Q9CQ86 | Migration and invasion enhancer 1 OS=Mus musculus OX=10090 GN=Mien1 PE=1 SV=1 | 0.041 | 1.659 | 6 | 1 | 1 | 1 | 115 | 12.3 | 4.51 | 0.603 | 0.573 | 1.054 | -0.73 | 0.73 | -0.8 | -0.2666667 | 0.765 | 0.08 | 0.038371923 | 0.066654277 | 0.770670847 | 0.395498846 | 0.59705936 | 0.496279103 | 0.992535481 |
| Reep6 | High | Q9JM62 | Receptor expression-enhancing protein 6 OS=Mus musculus OX=10090 GN=Reep6 PE=1 SV=1 | 0 | 40.279 | 25 | 4 | 18 | 4 | 201 | 22.2 | 7.33 | 1.78 | 2.021 | 0.881 | 0.83 | -0.83 | 1.02 | 0.34 | -0.925 | -0.18 | 0.039170403 | 0.038242801 | 0.473025891 | 0.402177331 | 0.486480307 | 0.444328819 | 0.961310681 |
| Pitpnb | High | Q8JZZ5 | Phosphatidylinositol transfer protein beta isoform OS=Mus musculus OX=10090 GN=Pitpnb PE=1 | 0 | 47.907 | 55 | 13 | 17 | 11 | 272 | 31.6 | 6.37 | 1.818 | 1.554 | 1.17 | 0.86 | -0.86 | 0.64 | 0.21333333 | -0.75 | 0.23 | 0.039251968 | 0.205711909 | 0.29321528 | 0.402177331 | 0.912882441 | 0.657529886 | 0.893995689 |
| Samd7 | High | Q8C8Y5 | Sterile alpha motif domain-containing protein 7 OS=Mus musculus OX=10090 GN=Samd7 PE=2 S | 0.001 | 5.928 | 3 | 1 | 1 | 1 | 445 | 49.2 | 6.8 | 1.678 | 1.348 | 1.244 | 0.75 | -0.75 | 0.43 | 0.14333333 | -0.59 | 0.32 | 0.039788794 | 0.38828154 | 0.213708885 | 0.405628667 | 0.98165177 | 0.693640218 | 0.804011919 |
| Srsf4 | High | Q542V3 | Serine/arginine-rich-splicing factor 4 OS=Mus musculus OX=10090 GN=Srsf4 PE=1 SV=1 | 0 | 11.365 | 8 | 4 | 4 | 2 | 491 | 56.2 | 11.37 | 1.706 | 1.245 | 1.37 | 0.77 | -0.77 | 0.32 | 0.10666667 | -0.545 | 0.45 | 0.039756231 | 0.557247428 | 0.074157501 | 0.405628667 | 0.995559182 | 0.700593924 | 0.520184986 |
| Dnajb2 | High | Q9QYI5 | DnaJ homolog subfamily B member 2 OS=Mus musculus OX=10090 GN=Dnajb2 PE=1 SV=3 | 0 | 21.052 | 16 | 5 | 9 | 5 | 324 | 35.6 | 5.91 | 0.547 | 0.505 | 1.085 | -0.87 | 0.87 | -0.99 | -0.33 | 0.93 | 0.12 | 0.039921304 | 0.029005694 | 0.636142706 | 0.406240924 | 0.432539704 | 0.419390314 | 0.983597794 |
| Ndfip2 | High | M0QWK1 | NEDD4 family-interacting protein 2 OS=Mus musculus OX=10090 GN=Ndfip2 PE=1 SV=1 | 0 | 16.305 | 8 | 2 | 3 | 2 | 338 | 36.5 | 5.31 | 0.539 | 0.473 | 1.14 | -0.89 | 0.89 | -1.08 | -0.36 | 0.985 | 0.19 | 0.040680978 | 0.017076026 | 0.39861576 | 0.410091165 | 0.305844112 | 0.357967639 | 0.93262326 |
| Nop16 | High | Q9CPT5 | Nucleolar protein 16 OS=Mus musculus OX=10090 GN=Nop16 PE=1 SV=1 | 0.001 | 6.229 | 14 | 2 | 2 | 2 | 178 | 21.1 | 9.91 | 1.716 | 1.754 | 0.978 | 0.78 | -0.78 | 0.81 | 0.27 | -0.795 | -0.03 | 0.040738502 | 0.091350743 | 0.925666641 | 0.410091165 | 0.696875669 | 0.553483417 | 0.992535481 |
| Tex30 | Low | Q3TUU5 | Testis-expressed protein 30 OS=Mus musculus OX=10090 GN=Tex30 PE=1 SV=1 | 0.077 | 1.23 | 3 | 1 | 1 | 1 | 225 | 25.2 | 8.75 | 1.68 | 1.236 | 1.359 | 0.75 | -0.75 | 0.31 | 0.10333333 | -0.53 | 0.44 | 0.040437175 | 0.573920859 | 0.050721457 | 0.410091165 | 0.995559182 | 0.702825173 | 0.427566646 |
| Syne1 | High | Q6ZWR6 | Nesprin-1 OS=Mus musculus OX=10090 GN=Syne1 PE=1 SV=2 | 0 | 51.022 | 2 | 12 | 15 | 10 | 8799 | 1009.3 | 5.59 | 1.839 | 1.792 | 1.026 | 0.88 | -0.88 | 0.84 | 0.28 | -0.86 | 0.04 | 0.041139387 | 0.088617851 | 0.825003033 | 0.413384488 | 0.696517952 | 0.55495122 | 0.992535481 |
| Stx16 | High | Q8BVI5 | Syntaxin-16 OS=Mus musculus OX=10090 GN=Stx16 PE=1 SV=3 | 0 | 12.435 | 6 | 1 | 2 | 1 | 326 | 37.1 | 5.86 | 0.615 | 0.776 | 0.792 | -0.7 | 0.7 | -0.37 | -0.1233333 | 0.535 | -0.34 | 0.042301719 | 0.379142526 | 0.256205494 | 0.422038685 | 0.977313184 | 0.699675934 | 0.855700667 |
| Vti1b | High | Q91XH6 | Vesicle transport through interaction with t-SNAREs 1B homolog OS=Mus musculus OX=10090 GI | 0 | 12.004 | 24 | 4 | 4 | 4 | 232 | 26.7 | 9.04 | 0.636 | 0.734 | 0.867 | -0.65 | 0.65 | -0.45 | -0.15 | 0.55 | -0.21 | 0.042612293 | 0.301838012 | 0.402297693 | 0.42287987 | 0.955820031 | 0.68934995 | 0.93262326 |
| Atp5g1 | Medium | Q9CR84 | ATP synthase F(0) complex subunit C1, mitochondrial OS=Mus musculus OX=10090 GN=Atp5mc1 | 0.02 | 2.031 | 5 | 1 | 4 | 1 | 136 | 14.2 | 10.01 | 1.665 | 1.413 | 1.178 | 0.74 | -0.74 | 0.5 | 0.16666667 | -0.62 | 0.24 | 0.043326209 | 0.329272431 | 0.336407263 | 0.426942093 | 0.965096978 | 0.696019536 | 0.906329766 |
| Ppa2 | High | Q91VM9 | Inorganic pyrophosphatase 2, mitochondrial OS=Mus musculus OX=10090 GN=Ppa2 PE=1 SV=1 | 0 | 15.792 | 22 | 5 | 5 | 5 | 330 | 38.1 | 6.98 | 1.701 | 1.555 | 1.094 | 0.77 | -0.77 | 0.64 | 0.21333333 | -0.705 | 0.13 | 0.044012634 | 0.204671809 | 0.601198201 | 0.431431535 | 0.912882441 | 0.672156988 | 0.978211931 |
| Ubqln2 | High | Q9QZM0 | Ubiquilin-2 OS=Mus musculus OX=10090 GN=Ubqln2 PE=1 SV=2 | 0 | 44.592 | 13 | 6 | 10 | 4 | 638 | 67.3 | 5.22 | 0.588 | 0.518 | 1.137 | -0.77 | 0.77 | -0.95 | -0.3166667 | 0.86 | 0.19 | 0.044687834 | 0.035192997 | 0.460382059 | 0.437285663 | 0.476392832 | 0.456839248 | 0.959744396 |
| Stat3 | High | P42227 | Signal transducer and activator of transcription 3 OS=Mus musculus OX=10090 GN=Stat3 PE=1 SV | 0 | 10.571 | 7 | 4 | 4 | 4 | 770 | 88 | 6.3 | 0.605 | 0.574 | 1.054 | -0.72 | 0.72 | -0.8 | -0.2666667 | 0.76 | 0.08 | 0.044873365 | 0.073931621 | 0.761919804 | 0.437573841 | 0.636899913 | 0.537236877 | 0.992535481 |
| Ift74 | High | Q8BKE9 | Intraflagellar transport protein 74 homolog OS=Mus musculus OX=10090 GN=Ift74 PE=1 SV=2 | 0 | 22.82 | 14 | 8 | 8 | 8 | 600 | 69.3 | 6 | 1.704 | 1.364 | 1.249 | 0.77 | -0.77 | 0.45 | 0.15 | -0.61 | 0.32 | 0.045065297 | 0.385902841 | 0.202672722 | 0.438682498 | 0.978153114 | 0.708417806 | 0.78697088 |
| Hsp90aa1 | High | P07901 | Heat shock protein HSP 90-alpha OS=Mus musculus OX=10090 GN=Hsp90aa1 PE=1 SV=4 | 0 | 359.148 | 62 | 46 | 156 | 30 | 733 | 84.7 | 5.01 | 0.551 | 0.592 | 0.932 | -0.86 | 0.86 | -0.76 | -0.2533333 | 0.81 | -0.1 | 0.045334813 | 0.089915388 | 0.721331025 | 0.440094263 | 0.696517952 | 0.568306107 | 0.992535481 |
| S100a11 | High | P50543 | Protein S100-A11 OS=Mus musculus OX=10090 GN=S100a11 PE=1 SV=1 | 0 | 13.066 | 16 | 1 | 2 | 1 | 98 | 11.1 | 5.45 | 0.596 | 0.813 | 0.733 | -0.75 | 0.75 | -0.3 | -0.1 | 0.525 | -0.45 | 0.045367306 | 0.447743129 | 0.092554369 | 0.440094263 | 0.994630833 | 0.717362548 | 0.581982763 |
| Fus | High | P56959 | RNA-binding protein FUS OS=Mus musculus OX=10090 GN=Fus PE=1 SV=1 | 0 | 49.728 | 17 | 11 | 15 | 11 | 518 | 52.6 | 9.36 | 1.814 | 1.256 | 1.444 | 0.86 | -0.86 | 0.33 | 0.11 | -0.595 | 0.53 | 0.045734309 | 0.539471862 | 0.013621392 | 0.442124603 | 0.995559182 | 0.718841892 | 0.191288138 |
| Vps25 | High | Q9CQ80 | Vacuolar protein-sorting-associated protein 25 OS=Mus musculus OX=10090 GN=Vps25 PE=1 SV= | 0 | 29.156 | 51 | 6 | 7 | 6 | 176 | 20.7 | 6.34 | 0.584 | 0.48 | 1.217 | -0.78 | 0.78 | -1.06 | -0.3533333 | 0.92 | 0.28 | 0.045853858 | 0.019398203 | 0.252002959 | 0.442517353 | 0.337781755 | 0.390149554 | 0.851275262 |
| Mbnl1 | Medium | A0A0A6YV | Muscleblind-like protein 1 OS=Mus musculus OX=10090 GN=Mbnl1 PE=1 SV=1 | 0.025 | 1.993 | 2 | 1 | 1 | 1 | 347 | 37.8 | 8.75 | 1.702 | 1.677 | 1.015 | 0.77 | -0.77 | 0.75 | 0.25 | -0.76 | 0.02 | 0.046984455 | 0.127390361 | 0.86759029 | 0.444252676 | 0.799286338 | 0.621769507 | 0.992535481 |
| Pef1 | High | Q8BFY6 | Peflin OS=Mus musculus OX=10090 GN=Pef1 PE=1 SV=1 | 0 | 26.396 | 25 | 4 | 5 | 4 | 275 | 29.2 | 6.3 | 0.636 | 0.7 | 0.909 | -0.65 | 0.65 | -0.52 | -0.1733333 | 0.585 | -0.14 | 0.046458231 | 0.237721681 | 0.579756727 | 0.444252676 | 0.937374952 | 0.690813814 | 0.972910028 |
| Rab27b | High | Q99P58 | Ras-related protein Rab-27B OS=Mus musculus OX=10090 GN=Rab27b PE=1 SV=3 | 0.001 | 6.959 | 13 | 2 | 2 | 2 | 218 | 24.5 | 5.54 | 0.598 | 0.588 | 1.017 | -0.74 | 0.74 | -0.77 | -0.2566667 | 0.755 | 0.02 | 0.04811842 | 0.080031983 | 0.907426688 | 0.452684535 | 0.656051653 | 0.554368094 | 0.992535481 |
| Glul | High | P15105 | Glutamine synthetase OS=Mus musculus OX=10090 GN=Glul PE=1 SV=6 | 0 | 225.49 | 64 | 25 | 136 | 25 | 373 | 42.1 | 7.08 | 1.798 | 1.498 | 1.201 | 0.85 | -0.85 | 0.58 | 0.19333333 | -0.715 | 0.26 | 0.047964312 | 0.248671091 | 0.201299182 | 0.452684535 | 0.950034749 | 0.701359642 | 0.78697088 |
| Mlycd | Medium | Q99J39 | Malonyl-CoA decarboxylase, mitochondrial OS=Mus musculus OX=10090 GN=Mlycd PE=1 SV=1 | 0.019 | 2.067 | 2 | 1 | 1 | 1 | 492 | 54.7 | 8.94 | 1.702 | 1.487 | 1.144 | 0.77 | -0.77 | 0.57 | 0.19 | -0.67 | 0.19 | 0.048047294 | 0.268439299 | 0.429747482 | 0.452684535 | 0.953285167 | 0.702984851 | 0.944356224 |
| Lrrc8d | High | Q8BGR2 | Volume-regulated anion channel subunit LRRC8D OS=Mus musculus OX=10090 GN=Lrrc8d PE=1 S | 0 | 8.127 | 3 | 3 | 3 | 3 | 859 | 98.1 | 7.44 | 0.554 | 0.528 | 1.049 | -0.85 | 0.85 | -0.92 | -0.3066667 | 0.885 | 0.07 | 0.049509753 | 0.04080534 | 0.729990139 | 0.464216033 | 0.498588435 | 0.481402234 | 0.992535481 |
| Ckmt1 | High | P30275 | Creatine kinase U-type, mitochondrial OS=Mus musculus OX=10090 GN=Ckmt1 PE=1 SV=1 | 0 | 74.199 | 46 | 13 | 24 | 13 | 418 | 47 | 8.16 | 1.822 | 1.67 | 1.091 | 0.87 | -0.87 | 0.74 | 0.24666667 | -0.805 | 0.13 | 0.050225387 | 0.13689774 | 0.509538103 | 0.470139805 | 0.817450082 | 0.643794943 | 0.964378027 |
| Rnf13 | Medium | O54965 | E3 ubiquitin-protein ligase RNF13 OS=Mus musculus OX=10090 GN=Rnf13 PE=1 SV=2 | 0.023 | 2.002 | 2 | 1 | 1 | 1 | 381 | 42.7 | 5.06 | 0.618 | 0.94 | 0.658 | -0.69 | 0.69 | -0.09 | -0.03 | 0.39 | -0.6 | 0.051010378 | 0.768420452 | 0.03451988 | 0.475108292 | 0.995559182 | 0.735333737 | 0.364272011 |
| Gna11 | High | P21278 | Guanine nucleotide-binding protein subunit alpha-11 OS=Mus musculus OX=10090 GN=Gna11 PE | 0 | 57.187 | 40 | 12 | 26 | 8 | 359 | 42 | 5.97 | 0.564 | 0.53 | 1.066 | -0.83 | 0.83 | -0.92 | -0.3066667 | 0.875 | 0.09 | 0.051215774 | 0.041776641 | 0.704049745 | 0.47623026 | 0.505921442 | 0.491075851 | 0.992485323 |
| Sfpq | High | Q8VIJ6 | Splicing factor, proline- and glutamine-rich OS=Mus musculus OX=10090 GN=Sfpq PE=1 SV=1 | 0 | 33.687 | 14 | 10 | 15 | 9 | 699 | 75.4 | 9.44 | 1.798 | 1.285 | 1.399 | 0.85 | -0.85 | 0.36 | 0.12 | -0.605 | 0.48 | 0.051498105 | 0.494603859 | 0.027959334 | 0.478062703 | 0.995559182 | 0.736810942 | 0.317344098 |
| Arsg | High | Q3TYD4 | Arylsulfatase G OS=Mus musculus OX=10090 GN=Arsg PE=2 SV=1 | 0 | 14.201 | 11 | 4 | 4 | 4 | 525 | 57.4 | 6.39 | 1.757 | 1.771 | 0.992 | 0.81 | -0.81 | 0.82 | 0.27333333 | -0.815 | -0.01 | 0.05232797 | 0.095620953 | 0.972570946 | 0.484963515 | 0.708251892 | 0.596607703 | 0.996575243 |
| Lnp; Lnpk | High | Q7TQ95 | Endoplasmic reticulum junction formation protein lunapark OS=Mus musculus OX=10090 GN=Lnp | 0.001 | 4.316 | 4 | 1 | 1 | 1 | 425 | 47.5 | 5.27 | 0.603 | 0.717 | 0.841 | -0.73 | 0.73 | -0.48 | -0.16 | 0.605 | -0.25 | 0.052603662 | 0.265271148 | 0.343290258 | 0.486714081 | 0.951072422 | 0.718893252 | 0.907770836 |
| Cdsn | Low | Q7TPC1 | Corneodesmosin OS=Mus musculus OX=10090 GN=Cdsn PE=2 SV=2 | 0.052 | 1.466 | 2 | 1 | 1 | 1 | 561 | 54.3 | 9.28 | 1.64 | 1.46 | 1.123 | 0.71 | -0.71 | 0.55 | 0.18333333 | -0.63 | 0.17 | 0.053185347 | 0.268588771 | 0.496046325 | 0.489126583 | 0.953285167 | 0.721205875 | 0.964378027 |
| Rbm3 | High | Q8BG13 | RNA-binding protein 3 OS=Mus musculus OX=10090 GN=Rbm3 PE=1 SV=1 | 0.001 | 6.009 | 12 | 1 | 1 | 1 | 154 | 16.8 | 8.38 | 0.629 | 0.943 | 0.668 | -0.67 | 0.67 | -0.09 | -0.03 | 0.38 | -0.58 | 0.053213343 | 0.790320817 | 0.032199761 | 0.489126583 | 0.998402594 | 0.743764589 | 0.347869094 |
| Rbm39 | High | E9Q8F0 | RNA-binding protein 39 OS=Mus musculus OX=10090 GN=Rbm39 PE=1 SV=1 | 0.001 | 7.038 | 5 | 2 | 2 | 2 | 415 | 47 | 10.52 | 1.664 | 1.513 | 1.1 | 0.73 | -0.73 | 0.6 | 0.2 | -0.665 | 0.14 | 0.054091189 | 0.229129135 | 0.566078877 | 0.492352754 | 0.935008675 | 0.713680715 | 0.972910028 |
| Rita1 | High | Q9D1H0 | RBPJ-interacting and tubulin-associated protein 1 OS=Mus musculus OX=10090 GN=Rita1 PE=2 S | 0.002 | 4.127 | 8 | 2 | 2 | 2 | 253 | 27.1 | 10.67 | 1.634 | 1.475 | 1.108 | 0.71 | -0.71 | 0.56 | 0.18666667 | -0.635 | 0.15 | 0.053868896 | 0.256478886 | 0.491549146 | 0.492352754 | 0.951072422 | 0.721712588 | 0.964378027 |
| Plekha2 | High | Q9ERS5 | Pleckstrin homology domain-containing family A member 2 OS=Mus musculus OX=10090 GN=Ple | 0.001 | 7.674 | 5 | 2 | 2 | 2 | 425 | 47.4 | 8.47 | 1.695 | 1.23 | 1.378 | 0.76 | -0.76 | 0.3 | 0.1 | -0.53 | 0.46 | 0.053744112 | 0.573081714 | 0.059391266 | 0.492352754 | 0.995559182 | 0.743955968 | 0.471013901 |
| Ptpmt1 | High | Q66GT5 | Phosphatidylglycerophosphatase and protein-tyrosine phosphatase 1 OS=Mus musculus OX=1009 | 0 | 4.694 | 16 | 2 | 2 | 2 | 193 | 21.9 | 9.72 | 1.627 | 1.599 | 1.018 | 0.7 | -0.7 | 0.68 | 0.22666667 | -0.69 | 0.03 | 0.054767474 | 0.164690002 | 0.888808084 | 0.497700529 | 0.871008827 | 0.684354678 | 0.992535481 |
| Selenbp2 | High | Q63836 | Selenium-binding protein 2 OS=Mus musculus OX=10090 GN=Selenbp2 PE=1 SV=2 | 0 | 36.718 | 29 | 9 | 9 | 9 | 472 | 52.6 | 6.18 | 1.616 | 2.192 | 0.737 | 0.69 | -0.69 | 1.13 | 0.37666667 | -0.91 | -0.44 | 0.055079761 | 0.02024965 | 0.061583936 | 0.498116481 | 0.345911522 | 0.422014001 | 0.476099813 |
| Golph3 | High | Q9CRA5 | Golgi phosphoprotein 3 OS=Mus musculus OX=10090 GN=Golph3 PE=1 SV=1 | 0.001 | 5.742 | 8 | 2 | 2 | 2 | 298 | 33.7 | 6.44 | 0.629 | 0.691 | 0.91 | -0.67 | 0.67 | -0.53 | -0.1766667 | 0.6 | -0.14 | 0.055051387 | 0.205722887 | 0.687080898 | 0.498116481 | 0.912882441 | 0.705499461 | 0.991285312 |
| Rpgr | High | Q8CDM3 | X-linked retinitis pigmentosa GTPase regulator OS=Mus musculus OX=10090 GN=Rpgr PE=1 SV=1 | 0.001 | 5.531 | 2 | 2 | 2 | 2 | 972 | 108.4 | 4.64 | 1.675 | 2.085 | 0.804 | 0.74 | -0.74 | 1.06 | 0.35333333 | -0.9 | -0.32 | 0.055977259 | 0.027910777 | 0.227838126 | 0.503795331 | 0.417321937 | 0.460558634 | 0.816808422 |
| Ncl | High | P09405 | Nucleolin OS=Mus musculus OX=10090 GN=Ncl PE=1 SV=2 | 0 | 116.612 | 40 | 36 | 54 | 36 | 707 | 76.7 | 4.75 | 1.763 | 1.242 | 1.419 | 0.82 | -0.82 | 0.31 | 0.10333333 | -0.565 | 0.51 | 0.056123971 | 0.562765864 | 0.018463012 | 0.504306263 | 0.995559182 | 0.749932722 | 0.238069974 |
| Slc20a1 | High | Q61609 | Sodium-dependent phosphate transporter 1 OS=Mus musculus OX=10090 GN=Slc20a1 PE=1 SV=1 | 0 | 21.62 | 8 | 5 | 7 | 4 | 681 | 74.1 | 6.89 | 0.563 | 0.511 | 1.102 | -0.83 | 0.83 | -0.97 | -0.3233333 | 0.9 | 0.14 | 0.056365064 | 0.031896139 | 0.522415767 | 0.504494509 | 0.457395534 | 0.480945022 | 0.964378027 |
| Ist1 | High | Q9CX00 | IST1 homolog OS=Mus musculus OX=10090 GN=Ist1 PE=1 SV=1 | 0 | 17.932 | 15 | 5 | 6 | 5 | 362 | 39.4 | 5.44 | 0.618 | 0.666 | 0.927 | -0.69 | 0.69 | -0.59 | -0.1966667 | 0.64 | -0.11 | 0.056414849 | 0.183363717 | 0.664918287 | 0.504494509 | 0.895493677 | 0.699994093 | 0.987078855 |
| Map7; Mta | High | D3YWN7 | Ensconsin OS=Mus musculus OX=10090 GN=Map7 PE=1 SV=2 | 0 | 24.466 | 10 | 7 | 9 | 7 | 738 | 82.8 | 9.38 | 1.762 | 1.551 | 1.136 | 0.82 | -0.82 | 0.63 | 0.21 | -0.725 | 0.18 | 0.056396052 | 0.20777275 | 0.412423342 | 0.504494509 | 0.912882441 | 0.708688475 | 0.936397673 |
| | High | A0A452J8 | Enoyl-CoA delta isomerase 1, mitochondrial OS=Mus musculus OX=10090 GN=Eci1 PE=1 SV=1 | 0 | 16.658 | 11 | 2 | 3 | 2 | 286 | 32 | 8.98 | 1.611 | 1.798 | 0.896 | 0.69 | -0.69 | 0.85 | 0.28333333 | -0.77 | -0.16 | 0.056761209 | 0.078039704 | 0.534015511 | 0.505174756 | 0.655042841 | 0.580108798 | 0.96694628 |
| Nek9 | Low | Q8K1R7 | Serine/threonine-protein kinase Nek9 OS=Mus musculus OX=10090 GN=Nek9 PE=1 SV=2 | 0.079 | 1.214 | 1 | 1 | 1 | 1 | 984 | 107.1 | 5.63 | 0.64 | 0.646 | 0.99 | -0.64 | 0.64 | -0.63 | -0.21 | 0.635 | -0.01 | 0.057158633 | 0.135899433 | 0.977063531 | 0.507905638 | 0.817450082 | 0.66267786 | 0.997221123 |
| Lhpp | High | Q9D7I5 | Phospholysine phosphohistidine inorganic pyrophosphate phosphatase OS=Mus musculus OX=100 | 0.003 | 4.056 | 5 | 1 | 1 | 1 | 270 | 29.1 | 5.1 | 0.618 | 0.541 | 1.141 | -0.69 | 0.69 | -0.89 | -0.2966667 | 0.79 | 0.19 | 0.057410924 | 0.046082864 | 0.445040673 | 0.50934027 | 0.520047141 | 0.514693706 | 0.958608248 |

| | | | | | | | | | | | | | | | | | | | | | | | | | | | | |
|---|---|---|---|---|---|---|---|---|---|---|---|---|---|---|---|---|---|---|---|---|---|---|---|---|---|---|---|---|
| Dnajb6 | High | O54946 | DnaJ homolog subfamily B member 6 OS=Mus musculus OX=10090 GN=Dnajb6 PE=1 SV=4 | 0 | 18.231 | 10 | 4 | 7 | 4 | 365 | 39.8 | 9.36 | 1.64 | 1.189 | 1.38 | 0.71 | -0.71 | 0.25 | 0.08333333 | -0.48 | 0.46 | 0.059556989 | 0.656872475 | 0.063901846 | 0.522591604 | 0.995559182 | 0.759075393 | 0.485498168 |
| Cpm | High | Q80V42 | Carboxypeptidase M OS=Mus musculus OX=10090 GN=Cpm PE=1 SV=2 | 0 | 12.783 | 11 | 5 | 5 | 5 | 443 | 50.5 | 7.78 | 1.653 | 1.466 | 1.128 | 0.73 | -0.73 | 0.55 | 0.18333333 | -0.64 | 0.17 | 0.05966191 | 0.277181972 | 0.490086703 | 0.522694269 | 0.955820031 | 0.73925715 | 0.964378027 |
| Stoml2 | High | Q99JB2 | Stomatin-like protein 2, mitochondrial OS=Mus musculus OX=10090 GN=Stoml2 PE=1 SV=1 | 0 | 11.542 | 16 | 3 | 3 | 3 | 353 | 38.4 | 8.87 | 1.618 | 1.284 | 1.261 | 0.69 | -0.69 | 0.36 | 0.12 | -0.525 | 0.33 | 0.059907462 | 0.480971942 | 0.178621502 | 0.524026742 | 0.995559182 | 0.759792962 | 0.755780643 |
| | Medium | G3UX45 | URB2 ribosome biogenesis 2 homolog (S. cerevisiae) OS=Mus musculus OX=10090 GN=Urb2 PE= | 0.038 | 1.708 | 1 | 1 | 1 | 1 | 1296 | 145.6 | 7.27 | 0.63 | 0.688 | 0.916 | -0.67 | 0.67 | -0.54 | -0.18 | 0.605 | -0.13 | 0.060137024 | 0.217771584 | 0.622088664 | 0.524398592 | 0.922239631 | 0.723319112 | 0.97826833 |
| Tomm40l | High | Q9CZR3 | Mitochondrial import receptor subunit TOM40B OS=Mus musculus OX=10090 GN=Tomm40l PE=1 | 0.006 | 3.472 | 3 | 1 | 1 | 1 | 308 | 34 | 7.27 | 1.654 | 1.931 | 0.857 | 0.73 | -0.73 | 0.95 | 0.31666667 | -0.84 | -0.22 | 0.060526377 | 0.046487943 | 0.390510762 | 0.526974218 | 0.520047141 | 0.523510679 | 0.93064581 |
| Lyar | High | Q08288 | Cell growth-regulating nucleolar protein OS=Mus musculus OX=10090 GN=Lyar PE=1 SV=2 | 0.001 | 5.095 | 5 | 2 | 2 | 2 | 388 | 43.7 | 9.5 | 1.648 | 1.944 | 0.848 | 0.72 | -0.72 | 0.96 | 0.32 | -0.84 | -0.24 | 0.06100528 | 0.046532095 | 0.3216412 | 0.527537459 | 0.520047141 | 0.5237923 | 0.905686588 |
| D430042O | High | F6XQI1 | Katanin-interacting protein OS=Mus musculus OX=10090 GN=Katnip PE=1 SV=2 | 0.001 | 6.002 | 1 | 2 | 2 | 2 | 1584 | 176.5 | 5.22 | 1.657 | 1.703 | 0.973 | 0.73 | -0.73 | 0.77 | 0.25666667 | -0.75 | -0.04 | 0.061232586 | 0.121989015 | 0.903800696 | 0.527537459 | 0.795487485 | 0.661512472 | 0.992535481 |
| Flot1 | High | O08917 | Flotillin-1 OS=Mus musculus OX=10090 GN=Flot1 PE=1 SV=1 | 0 | 19.139 | 17 | 6 | 7 | 6 | 428 | 47.5 | 7.15 | 0.615 | 0.637 | 0.966 | -0.7 | 0.7 | -0.65 | -0.2166667 | 0.675 | -0.05 | 0.060856486 | 0.141340664 | 0.788496181 | 0.527537459 | 0.83278102 | 0.68015924 | 0.992535481 |
| | High | A0A0G2JF | Protein FAM133B OS=Mus musculus OX=10090 GN=Fam133b PE=1 SV=1 | 0.001 | 4.397 | 9 | 1 | 1 | 1 | 109 | 13 | 10.42 | 1.655 | 1.333 | 1.242 | 0.73 | -0.73 | 0.41 | 0.13666667 | -0.57 | 0.31 | 0.061249668 | 0.439872857 | 0.217596265 | 0.527537459 | 0.994630833 | 0.761084146 | 0.805520378 |
| Map4; Mt | High | P27546 | Microtubule-associated protein 4 OS=Mus musculus OX=10090 GN=Map4 PE=1 SV=3 | 0 | 270.808 | 35 | 39 | 104 | 12 | 1125 | 117.4 | 4.98 | 1.741 | 1.457 | 1.195 | 0.8 | -0.8 | 0.54 | 0.18 | -0.67 | 0.26 | 0.061545739 | 0.285464861 | 0.211184399 | 0.529274477 | 0.955820031 | 0.742547254 | 0.804011919 |
| Gucy1b3 | High | O54865 | Guanylate cyclase soluble subunit beta-1 OS=Mus musculus OX=10090 GN=Gucy1b1 PE=1 SV=1 | 0 | 13.452 | 9 | 5 | 5 | 5 | 620 | 70.6 | 5.31 | 0.567 | 0.716 | 0.792 | -0.82 | 0.82 | -0.48 | -0.16 | 0.65 | -0.34 | 0.061988122 | 0.266256121 | 0.168808004 | 0.531448628 | 0.951072422 | 0.741260525 | 0.738806217 |
| Xpo5 | High | Q924C1 | Exportin-5 OS=Mus musculus OX=10090 GN=Xpo5 PE=1 SV=1 | 0 | 8.742 | 2 | 2 | 2 | 2 | 1204 | 136.9 | 5.82 | 0.626 | 0.631 | 0.993 | -0.68 | 0.68 | -0.66 | -0.22 | 0.67 | -0.01 | 0.063604712 | 0.127382758 | 0.999248838 | 0.543645765 | 0.799286338 | 0.671466051 | 0.999917596 |
| Erh | Medium | G3UW85 | Enhancer of rudimentary homolog OS=Mus musculus OX=10090 GN=Erh PE=1 SV=1 | 0.035 | 1.788 | 6 | 1 | 2 | 1 | 123 | 14.3 | 5.54 | 1.672 | 2.159 | 0.774 | 0.74 | -0.74 | 1.11 | 0.37 | -0.925 | -0.37 | 0.063814706 | 0.020420316 | 0.107307636 | 0.543782758 | 0.345911522 | 0.44484714 | 0.626743662 |
| Rad23b | High | P54728 | UV excision repair protein RAD23 homolog B OS=Mus musculus OX=10090 GN=Rad23b PE=1 SV= | 0 | 11.123 | 9 | 4 | 5 | 3 | 416 | 43.5 | 4.83 | 0.618 | 0.74 | 0.836 | -0.69 | 0.69 | -0.43 | -0.1433333 | 0.56 | -0.26 | 0.064549917 | 0.312260898 | 0.301414869 | 0.549213024 | 0.955820031 | 0.752516527 | 0.900241969 |
| 1810009A | Medium | Q9D937 | Uncharacterized protein C11orf98 homolog OS=Mus musculus OX=10090 PE=1 SV=1 | 0.034 | 1.813 | 6 | 1 | 1 | 1 | 123 | 14.1 | 11.47 | 1.598 | 2.418 | 0.661 | 0.68 | -0.68 | 1.27 | 0.42333333 | -0.975 | -0.6 | 0.06480741 | 0.006994374 | 0.017784967 | 0.55056841 | 0.169869988 | 0.360219199 | 0.236304052 |
| Hmgn2; L | High | Q5XK38 | Hmgn2 protein OS=Mus musculus OX=10090 GN=Hmgn2 PE=1 SV=1 | 0 | 7.905 | 46 | 3 | 4 | 1 | 92 | 9.6 | 9.99 | 1.714 | 2.492 | 0.688 | 0.78 | -0.78 | 1.32 | 0.44 | -1.05 | -0.54 | 0.065088326 | 0.007030098 | 0.068762051 | 0.552118368 | 0.169904135 | 0.361011251 | 0.499920511 |
| Keap1 | High | Q9Z2X8 | Kelch-like ECH-associated protein 1 OS=Mus musculus OX=10090 GN=Keap1 PE=1 SV=1 | 0.006 | 3.301 | 2 | 1 | 1 | 1 | 624 | 69.5 | 6.44 | 0.627 | 0.561 | 1.119 | -0.67 | 0.67 | -0.84 | -0.28 | 0.755 | 0.16 | 0.06598637 | 0.060283888 | 0.511794941 | 0.557207192 | 0.580776217 | 0.568991705 | 0.964378027 |
| Gnas | High | Z4YKV1 | Guanine nucleotide-binding protein G(s) subunit alpha isoforms short OS=Mus musculus OX=100 | 0 | 102.242 | 50 | 19 | 51 | 1 | 379 | 44.2 | 6.47 | 0.578 | 0.706 | 0.82 | -0.79 | 0.79 | -0.5 | -0.1666667 | 0.645 | -0.29 | 0.065843473 | 0.247813002 | 0.237491543 | 0.557207192 | 0.950034749 | 0.753620971 | 0.828110564 |
| Scamp1 | High | Q8K021 | Secretory carrier-associated membrane protein 1 OS=Mus musculus OX=10090 GN=Scamp1 PE=1 | 0 | 58.357 | 29 | 6 | 15 | 6 | 338 | 38 | 7.71 | 0.601 | 0.698 | 0.861 | -0.74 | 0.74 | -0.52 | -0.1733333 | 0.63 | -0.22 | 0.066387022 | 0.234070434 | 0.358389634 | 0.559747414 | 0.936813316 | 0.748280365 | 0.914600864 |
| Hnrnpul2 | Medium | Q00PI9 | Heterogeneous nuclear ribonucleoprotein U-like protein 2 OS=Mus musculus OX=10090 GN=Hnrn | 0.035 | 1.795 | 1 | 1 | 1 | 1 | 745 | 84.9 | 4.89 | 0.645 | 0.622 | 1.037 | -0.63 | 0.63 | -0.69 | -0.23 | 0.66 | 0.05 | 0.067331359 | 0.109853205 | 0.816716604 | 0.566857249 | 0.768225125 | 0.667541187 | 0.992535481 |
| Slc12a5 | High | Q91V14 | Solute carrier family 12 member 5 OS=Mus musculus OX=10090 GN=Slc12a5 PE=1 SV=2 | 0 | 191.897 | 30 | 32 | 75 | 27 | 1138 | 126.2 | 6.74 | 0.58 | 0.738 | 0.787 | -0.78 | 0.78 | -0.44 | -0.1466667 | 0.61 | -0.35 | 0.067672852 | 0.308762235 | 0.147796071 | 0.568878086 | 0.955820031 | 0.762349059 | 0.712464409 |
| Oxct1 | High | Q9D0K2 | Succinyl-CoA:3-ketoacid coenzyme A transferase 1, mitochondrial OS=Mus musculus OX=10090 G | 0 | 16.049 | 8 | 4 | 6 | 4 | 520 | 56 | 8.53 | 1.608 | 1.918 | 0.838 | 0.69 | -0.69 | 0.94 | 0.31333333 | -0.815 | -0.25 | 0.068381593 | 0.05585268 | 0.271114414 | 0.573117474 | 0.569450702 | 0.571284088 | 0.871850649 |
| Emc8-119 | High | M0QWE2 | Predicted gene, 27021 OS=Mus musculus OX=10090 GN=Gm27021 PE=4 SV=1 | 0 | 17.577 | 31 | 3 | 5 | 2 | 156 | 16.9 | 7.14 | 0.646 | 0.714 | 0.904 | -0.63 | 0.63 | -0.49 | -0.1633333 | 0.56 | -0.14 | 0.069065125 | 0.263215757 | 0.54954686 | 0.576262134 | 0.951072422 | 0.763667278 | 0.96694628 |
| Caskin2 | High | Q8VHK1 | Caskin-2 OS=Mus musculus OX=10090 GN=Caskin2 PE=1 SV=3 | 0 | 8.801 | 3 | 3 | 3 | 2 | 1201 | 126.7 | 7.39 | 1.615 | 1.517 | 1.065 | 0.69 | -0.69 | 0.6 | 0.2 | -0.645 | 0.09 | 0.069776478 | 0.224141195 | 0.699116332 | 0.579700023 | 0.935008675 | 0.757354349 | 0.991285312 |
| Tspan5 | High | P62080 | Tetraspanin-5 OS=Mus musculus OX=10090 GN=Tspan5 PE=1 SV=1 | 0.001 | 7.02 | 5 | 1 | 2 | 1 | 268 | 30.3 | 4.78 | 1.614 | 1.128 | 1.431 | 0.69 | -0.69 | 0.17 | 0.05666667 | -0.43 | 0.52 | 0.069787322 | 0.780481696 | 0.027687044 | 0.579700023 | 0.998402594 | 0.789051309 | 0.315745518 |
| Eml1 | High | D3Z4J9 | Echinoderm microtubule-associated protein-like 1 OS=Mus musculus OX=10090 GN=Eml1 PE=1 S | 0 | 15.089 | 3 | 3 | 3 | 3 | 800 | 88.3 | 7.94 | 1.685 | 1.605 | 1.05 | 0.75 | -0.75 | 0.68 | 0.22666667 | -0.715 | 0.07 | 0.071487134 | 0.177003022 | 0.755212002 | 0.589453469 | 0.875953959 | 0.732703714 | 0.992535481 |
| 2010107E | High | P56379 | ATP synthase subunit ATP5MJ, mitochondrial OS=Mus musculus OX=10090 GN=Atp5mj PE=1 SV= | 0.001 | 6.285 | 34 | 2 | 3 | 2 | 58 | 6.7 | 9.99 | 1.55 | 1.461 | 1.06 | 0.63 | -0.63 | 0.55 | 0.18333333 | -0.59 | 0.08 | 0.07133137 | 0.28109266 | 0.73673171 | 0.589453469 | 0.955820031 | 0.77263675 | 0.992535481 |
| Epb4.1l3; | High | A7YY80 | 130kDa Protein 4.1B MEF cell isoform OS=Mus musculus OX=10090 GN=Epb41l3 PE=1 SV=1 | 0 | 130.674 | 32 | 26 | 39 | 2 | 876 | 97.5 | 5.43 | 0.635 | 0.772 | 0.822 | -0.66 | 0.66 | -0.37 | -0.1233333 | 0.515 | -0.28 | 0.071232606 | 0.355878792 | 0.284905897 | 0.589453469 | 0.977313184 | 0.783383327 | 0.886074068 |
| Lrit1 | High | Q8K099 | Leucine-rich repeat, immunoglobulin-like domain and transmembrane domain-containing protein | 0.001 | 7.189 | 2 | 1 | 1 | 1 | 624 | 68.4 | 7.68 | 0.653 | 0.867 | 0.754 | -0.61 | 0.61 | -0.21 | -0.07 | 0.41 | -0.41 | 0.071730156 | 0.59477088 | 0.135531581 | 0.590588818 | 0.995559182 | 0.793074 | 0.699265243 |
| 1300018I1 | Medium | Q8C190 | VPS9 domain-containing protein 1 OS=Mus musculus OX=10090 GN=Vps9d1 PE=1 SV=1 | 0.016 | 2.164 | 1 | 1 | 1 | 1 | 649 | 71.2 | 6.87 | 0.619 | 0.522 | 1.185 | -0.69 | 0.69 | -0.94 | -0.3133333 | 0.815 | 0.24 | 0.074144383 | 0.033395542 | 0.372677225 | 0.591362101 | 0.470474383 | 0.530918242 | 0.920164724 |
| Fam84b | Medium | D3YXJ5 | LRAT domain-containing 1 OS=Mus musculus OX=10090 GN=Lratd2 PE=1 SV=1 | 0.044 | 1.598 | 4 | 1 | 1 | 1 | 310 | 34.6 | 5.54 | 1.657 | 1.849 | 0.896 | 0.73 | -0.73 | 0.89 | 0.29666667 | -0.81 | -0.16 | 0.074128281 | 0.071926605 | 0.483759267 | 0.591362101 | 0.626230553 | 0.608796327 | 0.964378027 |
| Ablim1 | Medium | E9Q030 | Actin-binding LIM protein 1 OS=Mus musculus OX=10090 GN=Ablim1 PE=1 SV=1 | 0.01 | 2.724 | 2 | 1 | 1 | 1 | 545 | 62 | 9.39 | 0.663 | 0.606 | 1.094 | -0.59 | 0.59 | -0.72 | -0.24 | 0.655 | 0.13 | 0.07312983 | 0.091055405 | 0.58214966 | 0.591362101 | 0.696517952 | 0.643940026 | 0.972910028 |
| Hnrnph3 | Medium | D3Z3N4 | Heterogeneous nuclear ribonucleoprotein H3 OS=Mus musculus OX=10090 GN=Hnrnph3 PE=1 SV= | 0.038 | 1.715 | 3 | 1 | 1 | 1 | 346 | 36.8 | 6.87 | 1.61 | 1.693 | 0.951 | 0.69 | -0.69 | 0.76 | 0.25333333 | -0.725 | -0.07 | 0.073459307 | 0.126303253 | 0.81491546 | 0.591362101 | 0.799286338 | 0.695324219 | 0.992535481 |
| Hist1h1a | High | P43275 | Histone H1.1 OS=Mus musculus OX=10090 GN=H1-1 PE=1 SV=2 | 0 | 25.537 | 21 | 7 | 10 | 5 | 213 | 21.8 | 10.93 | 1.589 | 1.554 | 1.023 | 0.67 | -0.67 | 0.64 | 0.21333333 | -0.655 | 0.03 | 0.073720581 | 0.205533553 | 0.901874276 | 0.591362101 | 0.912882441 | 0.752122271 | 0.992535481 |
| Slco1a4 | High | Q9EP96 | Solute carrier organic anion transporter family member 1A4 OS=Mus musculus OX=10090 GN=Sl | 0 | 16.767 | 9 | 5 | 7 | 4 | 670 | 73.9 | 7.59 | 1.764 | 1.512 | 1.167 | 0.82 | -0.82 | 0.6 | 0.2 | -0.71 | 0.22 | 0.073961337 | 0.237318722 | 0.33496524 | 0.591362101 | 0.937374952 | 0.764368526 | 0.906329766 |
| Ccdc66 | High | Q6NS45 | Coiled-coil domain-containing protein 66 OS=Mus musculus OX=10090 GN=Ccdc66 PE=1 SV=3 | 0 | 9.866 | 5 | 4 | 4 | 4 | 935 | 107.3 | 7.83 | 1.612 | 1.437 | 1.122 | 0.69 | -0.69 | 0.52 | 0.17333333 | -0.605 | 0.17 | 0.07345371 | 0.30488696 | 0.511810776 | 0.591362101 | 0.955820031 | 0.773591066 | 0.964378027 |
| Decr1 | High | Q9CQ62 | 2,4-dienoyl-CoA reductase [(3E)-enoyl-CoA-producing], mitochondrial OS=Mus musculus OX=100 | 0 | 15.968 | 16 | 4 | 4 | 4 | 335 | 36.2 | 8.95 | 1.604 | 1.46 | 1.099 | 0.68 | -0.68 | 0.55 | 0.18333333 | -0.615 | 0.14 | 0.074017938 | 0.280958059 | 0.533798371 | 0.591362101 | 0.955820031 | 0.773591066 | 0.96694628 |
| Dkk3 | High | Q9QUN9 | Dickkopf-related protein 3 OS=Mus musculus OX=10090 GN=Dkk3 PE=2 SV=1 | 0.001 | 5.235 | 7 | 1 | 1 | 1 | 349 | 38.4 | 4.54 | 0.611 | 0.864 | 0.707 | -0.71 | 0.71 | -0.21 | -0.07 | 0.46 | -0.5 | 0.073839481 | 0.570147097 | 0.11619269 | 0.591362101 | 0.995559182 | 0.793460641 | 0.651277329 |
| Khdrbs1 | Medium | Q60749 | KH domain-containing, RNA-binding, signal transduction-associated protein 1 OS=Mus musculus | 0.01 | 2.605 | 2 | 1 | 1 | 1 | 443 | 48.3 | 8.72 | 1.614 | 1.07 | 1.508 | 0.69 | -0.69 | 0.1 | 0.03333333 | -0.395 | 0.59 | 0.074451086 | 0.892740302 | 0.022051249 | 0.592964832 | 0.998402594 | 0.795683713 | 0.268785551 |
| Mavs | Medium | Q8VCF0 | Mitochondrial antiviral-signaling protein OS=Mus musculus OX=10090 GN=Mavs PE=1 SV=1 | 0.041 | 1.682 | 1 | 1 | 1 | 1 | 503 | 53.4 | 6.37 | 1.6 | 1.365 | 1.172 | 0.68 | -0.68 | 0.45 | 0.15 | -0.565 | 0.23 | 0.074809386 | 0.380533001 | 0.381809341 | 0.594973375 | 0.977313184 | 0.786143279 | 0.923955536 |
| Ncam1 | High | E9QB01 | Neural cell adhesion molecule 1 OS=Mus musculus OX=10090 GN=Ncam1 PE=1 SV=2 | 0 | 353.972 | 56 | 41 | 203 | 7 | 849 | 93.6 | 4.93 | 0.589 | 0.636 | 0.927 | -0.76 | 0.76 | -0.65 | -0.2166667 | 0.705 | -0.11 | 0.075800611 | 0.140657024 | 0.695492198 | 0.602002868 | 0.831079719 | 0.716541294 | 0.991285312 |
| Hint3 | High | Q9CPS6 | Histidine triad nucleotide-binding protein 3 OS=Mus musculus OX=10090 GN=Hint3 PE=1 SV=1 | 0 | 15.938 | 22 | 5 | 9 | 5 | 165 | 18.8 | 6.33 | 1.712 | 2.328 | 0.736 | 0.78 | -0.78 | 1.22 | 0.40666667 | -1 | -0.44 | 0.076756703 | 0.012236374 | 0.070863807 | 0.607874061 | 0.256005042 | 0.431939551 | 0.508467979 |
| Gclc | High | P97494 | Glutamate--cysteine ligase catalytic subunit OS=Mus musculus OX=10090 GN=Gclc PE=1 SV=4 | 0.002 | 4.145 | 3 | 1 | 1 | 1 | 637 | 72.5 | 5.9 | 1.667 | 1.766 | 0.944 | 0.74 | -0.74 | 0.82 | 0.27333333 | -0.78 | -0.08 | 0.077505799 | 0.100733563 | 0.833537074 | 0.612087032 | 0.730697757 | 0.671392394 | 0.992535481 |
| Slc12a2 | High | E9QM38 | Solute carrier family 12 member 2 OS=Mus musculus OX=10090 GN=Slc12a2 PE=1 SV=1 | 0 | 100.167 | 23 | 22 | 35 | 22 | 1206 | 130.6 | 6.52 | 0.591 | 0.614 | 0.962 | -0.76 | 0.76 | -0.7 | -0.2333333 | 0.73 | -0.06 | 0.077507008 | 0.114108733 | 0.882577533 | 0.612087032 | 0.77945068 | 0.695768856 | 0.992535481 |
| Atp5k | High | Q06185 | ATP synthase subunit e, mitochondrial OS=Mus musculus OX=10090 GN=Atp5me PE=1 SV=2 | 0 | 13.895 | 34 | 2 | 10 | 2 | 71 | 8.2 | 9.35 | 1.548 | 1.559 | 0.993 | 0.63 | -0.63 | 0.64 | 0.21333333 | -0.635 | -0.01 | 0.078677885 | 0.201783911 | 0.96641701 | 0.620459776 | 0.912882441 | 0.766671108 | 0.994815921 |
| Rab12 | High | A2CG35 | Ras-related protein Rab-12 OS=Mus musculus OX=10090 GN=Rab12 PE=1 SV=1 | 0 | 17.474 | 22 | 6 | 8 | 5 | 291 | 32.2 | 8.84 | 0.628 | 0.698 | 0.9 | -0.67 | 0.67 | -0.52 | -0.1733333 | 0.595 | -0.15 | 0.079736925 | 0.234882982 | 0.511094783 | 0.627047601 | 0.936813316 | 0.781930458 | 0.964378027 |
| Hadh | High | Q61425 | Hydroxyacyl-coenzyme A dehydrogenase, mitochondrial OS=Mus musculus OX=10090 GN=Hadh P | 0.008 | 2.89 | 2 | 1 | 1 | 1 | 314 | 34.4 | 8.65 | 1.62 | 1.445 | 1.121 | 0.7 | -0.7 | 0.53 | 0.17666667 | -0.615 | 0.16 | 0.079862671 | 0.296539301 | 0.437716229 | 0.627156857 | 0.955820031 | 0.791488444 | 0.951638192 |
| BC031181 | High | Q91WE4 | UPF0729 protein C18orf32 homolog OS=Mus musculus OX=10090 PE=3 SV=1 | 0 | 19.275 | 44 | 4 | 9 | 4 | 72 | 8 | 9.35 | 1.684 | 2.052 | 0.821 | 0.75 | -0.75 | 1.04 | 0.34666667 | -0.895 | -0.28 | 0.080350196 | 0.034100303 | 0.263344191 | 0.629222833 | 0.476192035 | 0.552707434 | 0.859470827 |
| Fam82a2; | High | Q3UJU9 | Regulator of microtubule dynamics protein 3 OS=Mus musculus OX=10090 GN=Rmdn3 PE=1 SV= | 0.001 | 7.188 | 7 | 2 | 2 | 2 | 470 | 52 | 5.21 | 1.602 | 1.739 | 0.921 | 0.68 | -0.68 | 0.8 | 0.26666667 | -0.74 | -0.12 | 0.0802906 | 0.104048222 | 0.696838193 | 0.629222833 | 0.741927143 | 0.685574988 | 0.991285312 |
| Copb1 | Low | Q9JIF7 | Coatomer subunit beta OS=Mus musculus OX=10090 GN=Copb1 PE=1 SV=1 | 0.051 | 1.479 | 1 | 1 | 1 | 1 | 953 | 107 | 6 | 0.645 | 0.845 | 0.763 | -0.63 | 0.63 | -0.24 | -0.08 | 0.435 | -0.39 | 0.08052315 | 0.517382464 | 0.123704747 | 0.629697772 | 0.995559182 | 0.812628477 | 0.677192798 |
| Hgs | High | Q99LI8 | Hepatocyte growth factor-regulated tyrosine kinase substrate OS=Mus musculus OX=10090 GN=H | 0 | 15.868 | 6 | 5 | 5 | 5 | 775 | 86 | 6.16 | 0.644 | 0.683 | 0.943 | -0.64 | 0.64 | -0.55 | -0.1833333 | 0.595 | -0.09 | 0.080941551 | 0.209153612 | 0.750668175 | 0.630332331 | 0.912882441 | 0.771607386 | 0.992535481 |
| Nudc | High | O35685 | Nuclear migration protein nudC OS=Mus musculus OX=10090 GN=Nudc PE=1 SV=1 | 0 | 30.245 | 31 | 9 | 10 | 9 | 332 | 38.3 | 5.26 | 0.594 | 0.622 | 0.955 | -0.75 | 0.75 | -0.69 | -0.23 | 0.72 | -0.07 | 0.081078406 | 0.12299919 | 0.80035246 | 0.630522364 | 0.798213491 | 0.714367927 | 0.992535481 |
| Atp5d | High | Q9D3D9 | ATP synthase subunit delta, mitochondrial OS=Mus musculus OX=10090 GN=Atp5f1d PE=1 SV=1 | 0 | 11.674 | 14 | 2 | 4 | 2 | 168 | 17.6 | 5.08 | 1.602 | 1.612 | 0.994 | 0.68 | -0.68 | 0.69 | 0.23 | -0.685 | -0.01 | 0.081419133 | 0.161155706 | 0.96293754 | 0.632295122 | 0.865517285 | 0.748906203 | 0.993960013 |
| Glo1 | High | Q9CPU0 | Lactoylglutathione lyase OS=Mus musculus OX=10090 GN=Glo1 PE=1 SV=3 | 0 | 18.763 | 27 | 5 | 6 | 5 | 184 | 20.8 | 5.47 | 0.606 | 0.565 | 1.072 | -0.72 | 0.72 | -0.82 | -0.2733333 | 0.77 | 0.1 | 0.082069708 | 0.066543642 | 0.696078272 | 0.635986083 | 0.59705936 | 0.616522722 | 0.991285312 |
| ATP8 | High | P03930 | ATP synthase protein 8 OS=Mus musculus OX=10090 GN=Mtatp8 PE=1 SV=1 | 0.001 | 5.411 | 27 | 2 | 3 | 2 | 67 | 7.8 | 9.88 | 1.621 | 1.639 | 0.989 | 0.7 | -0.7 | 0.71 | 0.23666667 | -0.705 | -0.02 | 0.082461545 | 0.152950293 | 0.962715074 | 0.635986083 | 0.847684847 | 0.741835465 | 0.993960013 |
| Nap1l1 | High | Q8BSH9 | Nucleosome assembly protein 1-like 1 OS=Mus musculus OX=10090 GN=Nap1l1 PE=1 SV=1 | 0 | 46.075 | 31 | 8 | 16 | 7 | 383 | 44.6 | 4.56 | 0.633 | 0.757 | 0.837 | -0.66 | 0.66 | -0.4 | -0.1333333 | 0.53 | -0.26 | 0.082349111 | 0.346522062 | 0.286198271 | 0.635986083 | 0.977313184 | 0.806649634 | 0.886074068 |
| Dnajc9 | High | Q91WN1 | DnaJ homolog subfamily C member 9 OS=Mus musculus OX=10090 GN=Dnajc9 PE=1 SV=2 | 0.001 | 6.338 | 15 | 2 | 2 | 2 | 259 | 30 | 5.94 | 1.585 | 1.424 | 1.113 | 0.66 | -0.66 | 0.51 | 0.17 | -0.585 | 0.15 | 0.083147707 | 0.307222808 | 0.526016698 | 0.640397247 | 0.955820031 | 0.798108639 | 0.964378027 |
| Gna13 | High | P27601 | Guanine nucleotide-binding protein subunit alpha-13 OS=Mus musculus OX=10090 GN=Gna13 PE | 0 | 59.179 | 34 | 14 | 40 | 12 | 377 | 44 | 8.21 | 0.597 | 0.579 | 1.032 | -0.74 | 0.74 | -0.79 | -0.2633333 | 0.765 | 0.05 | 0.083652297 | 0.07794647 | 0.760994118 | 0.641189049 | 0.655042841 | 0.648115945 | 0.992535481 |
| Gpnmb | Low | Q99P91 | Transmembrane glycoprotein NMB OS=Mus musculus OX=10090 GN=Gpnmb PE=1 SV=2 | 0.051 | 1.497 | 1 | 1 | 1 | 1 | 574 | 63.6 | 7.61 | 0.678 | 0.728 | 0.931 | -0.56 | 0.56 | -0.46 | -0.1533333 | 0.51 | -0.1 | 0.083707934 | 0.26655646 | 0.73382538 | 0.641189049 | 0.951072422 | 0.796130736 | 0.992535481 |
| Syap1 | High | Q9D5V6 | Synapse-associated protein 1 OS=Mus musculus OX=10090 GN=Syap1 PE=1 SV=1 | 0.001 | 7.467 | 8 | 3 | 4 | 3 | 365 | 41.3 | 4.54 | 0.651 | 0.739 | 0.881 | -0.62 | 0.62 | -0.44 | -0.1466667 | 0.53 | -0.18 | 0.083538719 | 0.288489889 | 0.511041638 | 0.641189049 | 0.955820031 | 0.79850454 | 0.964378027 |
| Rala | High | P63321 | Ras-related protein Ral-A OS=Mus musculus OX=10090 GN=Rala PE=1 SV=1 | 0 | 72.697 | 49 | 9 | 41 | 5 | 206 | 23.5 | 7.11 | 0.598 | 0.616 | 0.971 | -0.74 | 0.74 | -0.7 | -0.2333333 | 0.72 | -0.04 | 0.084608791 | 0.116313528 | 0.928562368 | 0.647205305 | 0.779640783 | 0.713423044 | 0.992535481 |
| Prpf19 | High | Q99KP6 | Pre-mRNA-processing factor 19 OS=Mus musculus OX=10090 GN=Prpf19 PE=1 SV=1 | 0 | 8.738 | 6 | 2 | 3 | 2 | 504 | 55.2 | 6.61 | 1.6 | 1.61 | 0.994 | 0.68 | -0.68 | 0.69 | 0.23 | -0.685 | -0.01 | 0.08568207 | 0.169451874 | 0.980428053 | 0.653631789 | 0.874947902 | 0.764289845 | 0.997765878 |
| Whrn | High | Q80VW5 | Whirlin OS=Mus musculus OX=10090 GN=Whrn PE=1 SV=3 | 0 | 4.966 | 2 | 2 | 2 | 2 | 918 | 98 | 8.22 | 1.608 | 1.478 | 1.088 | 0.68 | -0.68 | 0.56 | 0.18666667 | -0.62 | 0.12 | 0.085673289 | 0.266358592 | 0.556554908 | 0.653631789 | 0.951072422 | 0.802352105 | 0.970638684 |
| Dmrtc2 | Low | Q8CGW9 | Doublesex- and mab-3-related transcription factor C2 OS=Mus musculus OX=10090 GN=Dmrtc2 P | 0.063 | 1.311 | 2 | 1 | 1 | 1 | 370 | 39.1 | 9.42 | 0.689 | 0.596 | 1.156 | -0.54 | 0.54 | -0.75 | -0.25 | 0.645 | 0.21 | 0.086832137 | 0.082072632 | 0.347871014 | 0.657929448 | 0.669841702 | 0.663885575 | 0.909466648 |
| | High | A0A571BD | SRC kinase-signaling inhibitor 1 OS=Mus musculus OX=10090 GN=Srcin1 PE=1 SV=1 | 0 | 21.674 | 6 | 6 | 6 | 6 | 1225 | 131.5 | 9.45 | 0.606 | 0.617 | 0.983 | -0.72 | 0.72 | -0.7 | -0.2333333 | 0.71 | -0.02 | 0.086766127 | 0.116804052 | 0.94975199 | 0.657929448 | 0.779640783 | 0.718785116 | 0.992535481 |
| Uba52; Gr | High | P62984 | Ubiquitin-60S ribosomal protein L40 OS=Mus musculus OX=10090 GN=Uba52 PE=1 SV=2 | 0 | 82.829 | 55 | 9 | 105 | 1 | 128 | 14.7 | 9.83 | 1.597 | 1.292 | 1.235 | 0.68 | -0.68 | 0.37 | 0.12333333 | -0.525 | 0.3 | 0.087978661 | 0.484382832 | 0.145831778 | 0.664819883 | 0.995559182 | 0.830189532 | 0.706909771 |
| Snrnp70 | High | Q62376 | U1 small nuclear ribonucleoprotein 70 kDa OS=Mus musculus OX=10090 GN=Snrnp70 PE=1 SV=2 | 0.001 | 6.126 | 5 | 3 | 3 | 3 | 448 | 52 | 9.94 | 1.644 | 1.471 | 1.118 | 0.72 | -0.72 | 0.56 | 0.18666667 | -0.64 | 0.16 | 0.088365637 | 0.272429884 | 0.519705094 | 0.665055201 | 0.955820031 | 0.810437616 | 0.964378027 |
| LOC10263 | Medium | A0A1W2P | RIKEN cDNA 4932415D10 gene OS=Mus musculus OX=10090 GN=4932415D10Rik PE=4 SV=1 | 0.042 | 1.673 | 0 | 1 | 1 | 1 | 4986 | 551.5 | 8 | 1.572 | 1.091 | 1.441 | 0.65 | -0.65 | 0.13 | 0.04333333 | -0.39 | 0.53 | 0.088248272 | 0.85935427 | 0.029148853 | 0.665055201 | 0.998402594 | 0.831728897 | 0.324925684 |
| Dctn5 | High | Q9QZB9 | Dynactin subunit 5 OS=Mus musculus OX=10090 GN=Dctn5 PE=1 SV=1 | 0 | 4.983 | 6 | 1 | 1 | 1 | 182 | 20.1 | 8.02 | 0.659 | 0.79 | 0.833 | -0.6 | 0.6 | -0.34 | -0.1133333 | 0.47 | -0.26 | 0.088863131 | 0.406968798 | 0.386974091 | 0.667902915 | 0.9855335 | 0.826718208 | 0.93064581 |
| Prkdc | Low | P97313 | DNA-dependent protein kinase catalytic subunit OS=Mus musculus OX=10090 GN=Prkdc PE=1 SV= | 0.051 | 1.484 | 0 | 1 | 1 | 1 | 4128 | 471.2 | 7.12 | 0.649 | 0.493 | 1.316 | -0.62 | 0.62 | -1.02 | -0.34 | 0.82 | 0.4 | 0.09057628 | 0.021287401 | 0.12307408 | 0.678826967 | 0.35839181 | 0.518609389 | 0.677192798 |
| Chchd3 | High | Q9D9P1 | MICOS complex subunit Mic19 OS=Mus musculus OX=10090 GN=Chchd3 PE=1 SV=1 | 0 | 27.748 | 34 | 7 | 11 | 7 | 175 | 20.4 | 8.27 | 1.649 | 1.632 | 1.01 | 0.72 | -0.72 | 0.71 | 0.23666667 | -0.715 | 0.01 | 0.090921893 | 0.156474731 | 0.84771613 | 0.678826967 | 0.847684847 | 0.763255907 | 0.992535481 |
| Rragc | High | Q99K70 | Ras-related GTP-binding protein C OS=Mus musculus OX=10090 GN=Rragc PE=1 SV=1 | 0.003 | 4.092 | 3 | 1 | 1 | 1 | 398 | 44.1 | 5.1 | 0.673 | 0.663 | 1.016 | -0.57 | 0.57 | -0.59 | -0.1966667 | 0.58 | 0.02 | 0.090815692 | 0.158910845 | 0.855251086 | 0.678826967 | 0.854406564 | 0.766616766 | 0.992535481 |
| Zfyve19 | High | Q4KL52 | Abscission/NoCut checkpoint regulator OS=Mus musculus OX=10090 GN=Zfyve19 PE=1 SV=1 | 0 | 9.753 | 10 | 4 | 5 | 4 | 330 | 36.3 | 7.81 | 0.684 | 0.712 | 0.961 | -0.55 | 0.55 | -0.49 | -0.1633333 | 0.52 | -0.06 | 0.091225134 | 0.259574946 | 0.844664215 | 0.679281971 | 0.951072422 | 0.815177197 | 0.992535481 |
| Acaa2 | High | Q8BWT1 | 3-ketoacyl-CoA thiolase, mitochondrial OS=Mus musculus OX=10090 GN=Acaa2 PE=1 SV=3 | 0 | 10.339 | 9 | 3 | 3 | 3 | 397 | 41.8 | 8.09 | 1.584 | 1.22 | 1.298 | 0.66 | -0.66 | 0.29 | 0.09666667 | -0.475 | 0.38 | 0.091680855 | 0.60825833 | 0.096838654 | 0.681769969 | 0.995559182 | 0.838664575 | 0.593414569 |
| Tango6; T | Medium | Q8C3S2 | Transport and Golgi organization protein 6 homolog OS=Mus musculus OX=10090 GN=Tango6 PE | 0.014 | 2.281 | 1 | 1 | 2 | 1 | 1079 | 118.7 | 6.32 | 1.56 | 1.316 | 1.186 | 0.64 | -0.64 | 0.4 | 0.13333333 | -0.52 | 0.25 | 0.092137844 | 0.434444014 | 0.306891138 | 0.684260783 | 0.99399649 | 0.839128636 | 0.903485924 |
| Napb | High | P28663 | Beta-soluble NSF attachment protein OS=Mus musculus OX=10090 GN=Napb PE=1 SV=2 | 0 | 224.255 | 76 | 21 | 68 | 18 | 298 | 33.5 | 5.47 | 0.606 | 0.518 | 1.171 | -0.72 | 0.72 | -0.95 | -0.3166667 | 0.835 | 0.23 | 0.092934431 | 0.035291336 | 0.263288528 | 0.685898128 | 0.476392832 | 0.58114548 | 0.859470827 |
| Pvr | High | Q8K094 | Poliovirus receptor OS=Mus musculus OX=10090 GN=Pvr PE=1 SV=1 | 0 | 8.066 | 9 | 3 | 3 | 3 | 408 | 44.6 | 8.06 | 0.612 | 0.627 | 0.977 | -0.71 | 0.71 | -0.67 | -0.2233333 | 0.69 | -0.03 | 0.092615039 | 0.128692588 | 0.942611247 | 0.685898128 | 0.799286338 | 0.742592233 | 0.992535481 |
| Pcyt1a | High | P49586 | Choline-phosphate cytidylyltransferase A OS=Mus musculus OX=10090 GN=Pcyt1a PE=1 SV=1 | 0.001 | 7.431 | 6 | 2 | 2 | 1 | 367 | 41.6 | 7.03 | 1.654 | 1.371 | 1.206 | 0.73 | -0.73 | 0.45 | 0.15 | -0.59 | 0.27 | 0.092786862 | 0.382049254 | 0.354217487 | 0.685898128 | 0.977313184 | 0.831605656 | 0.912309349 |
| Vti1a | High | Q5FWJ7 | Vesicle transport through interaction with t-SNAREs homolog 1A OS=Mus musculus OX=10090 G | 0 | 9.207 | 19 | 4 | 4 | 4 | 224 | 25.9 | 6.4 | 0.656 | 0.576 | 1.138 | -0.61 | 0.61 | -0.79 | -0.2633333 | 0.7 | 0.19 | 0.093979172 | 0.075790077 | 0.444977305 | 0.687015932 | 0.643871158 | 0.665443545 | 0.958608248 |
| | High | D3YVS7 | Cysteine-rich and transmembrane domain-containing protein 1 (Fragment) OS=Mus musculus OX | 0 | 11.102 | 13 | 2 | 5 | 2 | 84 | 9.2 | 4.44 | 0.663 | 0.619 | 1.071 | -0.59 | 0.59 | -0.69 | -0.23 | 0.64 | 0.1 | 0.093867149 | 0.119582593 | 0.622350035 | 0.687015932 | 0.786053068 | 0.7365345 | 0.97826833 |
| Cntrob | High | Q8CB62 | Centrobin OS=Mus musculus OX=10090 GN=Cntrob PE=2 SV=2 | 0.001 | 7.171 | 3 | 2 | 2 | 2 | 887 | 99.3 | 5.55 | 1.622 | 1.421 | 1.142 | 0.7 | -0.7 | 0.51 | 0.17 | -0.605 | 0.19 | 0.094867471 | 0.330337111 | 0.46503894 | 0.692606653 | 0.967711694 | 0.830159174 | 0.960359242 |
| Enkd1 | High | Q7TSV9 | Enkurin domain-containing protein 1 OS=Mus musculus OX=10090 GN=Enkd1 PE=1 SV=1 | 0.001 | 6.883 | 9 | 3 | 3 | 3 | 346 | 38.9 | 10.32 | 1.553 | 1.457 | 1.066 | 0.64 | -0.64 | 0.54 | 0.18 | -0.59 | 0.09 | 0.095102106 | 0.2813338 | 0.64206439 | 0.693416789 | 0.955820031 | 0.82461841 | 0.985010103 |
| Rs1 | High | Q9Z1L4 | Retinoschisin OS=Mus musculus OX=10090 GN=Rs1 PE=1 SV=1 | 0 | 67.116 | 44 | 10 | 31 | 10 | 224 | 25.6 | 5.66 | 0.609 | 0.754 | 0.808 | -0.72 | 0.72 | -0.41 | -0.1366667 | 0.565 | -0.31 | 0.095638038 | 0.340416772 | 0.202361276 | 0.695515538 | 0.976045759 | 0.835780648 | 0.78697088 |
| Cox4i2 | Medium | Q91W29 | Cytochrome c oxidase subunit 4 isoform 2, mitochondrial OS=Mus musculus OX=10090 GN=Cox4i | 0.047 | 1.531 | 4 | 1 | 1 | 1 | 172 | 20.2 | 9.47 | 1.664 | 1.936 | 0.86 | 0.73 | -0.73 | 0.95 | 0.31666667 | -0.84 | -0.22 | 0.096232607 | 0.050281975 | 0.491884356 | 0.696001929 | 0.53295092 | 0.614476424 | 0.964378027 |
| Rogdi | High | Q3TDK6 | Protein rogdi homolog OS=Mus musculus OX=10090 GN=Rogdi PE=1 SV=2 | 0.009 | 2.802 | 3 | 1 | 1 | 1 | 287 | 32.1 | 8.18 | 0.658 | 0.646 | 1.019 | -0.6 | 0.6 | -0.63 | -0.21 | 0.615 | 0.03 | 0.096098651 | 0.144429292 | 0.897743504 | 0.696001929 | 0.838826712 | 0.767414321 | 0.992535481 |
| | High | G3UXT8 | BCL2-associated athanogene 6 (Fragment) OS=Mus musculus OX=10090 GN=Bag6 PE=1 SV=1 | 0 | 4.973 | 8 | 3 | 3 | 3 | 563 | 59.3 | 6.32 | 0.677 | 0.685 | 0.989 | -0.56 | 0.56 | -0.55 | -0.1833333 | 0.555 | -0.02 | 0.09707036 | 0.212674297 | 0.977301599 | 0.696001929 | 0.913066452 | 0.80453419 | 0.997221123 |
| | Medium | H7BWY7 | RBR-type E3 ubiquitin transferase OS=Mus musculus OX=10090 GN=Arih1 PE=1 SV=1 | 0.044 | 1.606 | 2 | 1 | 1 | 1 | 381 | 42.5 | 4.79 | 0.657 | 0.525 | 1.25 | -0.61 | 0.61 | -0.93 | -0.31 | 0.77 | 0.32 | 0.098363176 | 0.031797459 | 0.208225176 | 0.698129528 | 0.457149627 | 0.577639577 | 0.801370196 |
| Rabgap1 | High | A2AWA9 | Rab GTPase-activating protein 1 OS=Mus musculus OX=10090 GN=Rabgap1 PE=1 SV=1 | 0 | 10.899 | 3 | 4 | 4 | 4 | 1064 | 120.7 | 5.25 | 0.65 | 0.614 | 1.058 | -0.62 | 0.62 | -0.7 | -0.2333333 | 0.66 | 0.08 | 0.09769877 | 0.114068433 | 0.672531653 | 0.698129528 | 0.77945068 | 0.738790104 | 0.990250803 |
| | High | A0A3B2W | Predicted gene, 49804 OS=Mus musculus OX=10090 GN=Gm49804 PE=3 SV=1 | 0 | 27.748 | 28 | 9 | 15 | 1 | 287 | 32.7 | 9.22 | 0.676 | 0.646 | 1.047 | -0.56 | 0.56 | -0.63 | -0.21 | 0.595 | 0.07 | 0.097960643 | 0.140333735 | 0.749855362 | 0.698129528 | 0.830888335 | 0.764508931 | 0.992535481 |
| Pdcd6 | High | P12815 | Programmed cell death protein 6 OS=Mus musculus OX=10090 GN=Pdcd6 PE=1 SV=2 | 0 | 23.675 | 29 | 6 | 7 | 6 | 191 | 21.9 | 5.4 | 0.672 | 0.727 | 0.924 | -0.57 | 0.57 | -0.46 | -0.1533333 | 0.515 | -0.11 | 0.09811372 | 0.287920495 | 0.641904905 | 0.698129528 | 0.955820031 | 0.826974779 | 0.985010103 |
| Ssr3 | Medium | Q9DCF9 | Translocon-associated protein subunit gamma OS=Mus musculus OX=10090 GN=Ssr3 PE=1 SV=1 | 0.038 | 1.715 | 8 | 1 | 1 | 1 | 185 | 21.1 | 9.61 | 0.632 | 1.03 | 0.614 | -0.66 | 0.66 | 0.04 | 0.01333333 | 0.31 | -0.7 | 0.098072881 | 0.981090085 | 0.025553294 | 0.698129528 | 0.998402594 | 0.848266061 | 0.299743343 |
| Lrp2 | High | A2ARV4 | Low-density lipoprotein receptor-related protein 2 OS=Mus musculus OX=10090 GN=Lrp2 PE=1 S | 0.005 | 3.625 | 0 | 2 | 2 | 2 | 4660 | 518.9 | 5.15 | 1.552 | 1.692 | 0.918 | 0.63 | -0.63 | 0.76 | 0.25333333 | -0.695 | -0.12 | 0.099608367 | 0.115006222 | 0.644769272 | 0.702906868 | 0.77945068 | 0.741178774 | 0.985010103 |
| 2410002O | High | Q3TIR1 | Trafficking protein particle complex subunit 13 OS=Mus musculus OX=10090 GN=Trappc13 PE=2 | 0.003 | 3.903 | 4 | 1 | 1 | 1 | 417 | 46.4 | 5.49 | 0.662 | 0.641 | 1.033 | -0.6 | 0.6 | -0.64 | -0.2133333 | 0.62 | 0.05 | 0.099663093 | 0.136592898 | 0.822111003 | 0.702906868 | 0.817450082 | 0.760178475 | 0.992535481 |
| Ndufaf4 | Medium | A2AJX3 | NADH dehydrogenase [ubiquinone] 1 alpha subcomplex assembly factor 4 (Fragment) OS=Mus m | 0.021 | 2.018 | 7 | 1 | 1 | 1 | 127 | 15.2 | 9.99 | 1.57 | 1.468 | 1.069 | 0.65 | -0.65 | 0.55 | 0.18333333 | -0.6 | 0.1 | 0.099553067 | 0.265120718 | 0.640763155 | 0.702906868 | 0.951072422 | 0.826989645 | 0.984047935 |
| Hadhb | High | Q99JY0 | Trifunctional enzyme subunit beta, mitochondrial OS=Mus musculus OX=10090 GN=Hadhb PE=1 S | 0 | 18.391 | 10 | 4 | 5 | 4 | 475 | 51.4 | 9.38 | 1.613 | 1.382 | 1.167 | 0.69 | -0.69 | 0.47 | 0.15666667 | -0.58 | 0.22 | 0.100056706 | 0.364721688 | 0.375946222 | 0.704636625 | 0.977313184 | 0.840974904 | 0.92034959 |
| Pdcl | High | Q9DBX2 | Phosducin-like protein OS=Mus musculus OX=10090 GN=Pdcl PE=1 SV=1 | 0.001 | 4.644 | 6 | 2 | 2 | 1 | 301 | 34.4 | 4.87 | 0.664 | 0.854 | 0.777 | -0.59 | 0.59 | -0.23 | -0.0766667 | 0.41 | -0.36 | 0.100159691 | 0.549950662 | 0.223537221 | 0.704636625 | 0.995559182 | 0.850097903 | 0.80967261 |
| Srp68 | High | Q8BMA6 | Signal recognition particle subunit SRP68 OS=Mus musculus OX=10090 GN=Srp68 PE=1 SV=2 | 0.006 | 3.275 | 3 | 1 | 1 | 1 | 625 | 70.5 | 8.57 | 1.61 | 1.658 | 0.971 | 0.69 | -0.69 | 0.73 | 0.24333333 | -0.71 | -0.04 | 0.100870221 | 0.148156858 | 0.944471013 | 0.707858986 | 0.845081896 | 0.776470441 | 0.992535481 |
| AI314976; | High | Q8R5F3 | ADP-ribose glycohydrolase OARD1 OS=Mus musculus OX=10090 GN=Oard1 PE=1 SV=2 | 0 | 9.954 | 20 | 3 | 4 | 3 | 152 | 17.1 | 8.53 | 1.517 | 1.04 | 1.458 | 0.6 | -0.6 | 0.06 | 0.02 | -0.33 | 0.54 | 0.101768358 | 0.971759058 | 0.031141877 | 0.713255946 | 0.998402594 | 0.85582927 | 0.341707442 |
| Snrpa | High | Q62189 | U1 small nuclear ribonucleoprotein A OS=Mus musculus OX=10090 GN=Snrpa PE=1 SV=3 | 0.001 | 7.897 | 10 | 3 | 4 | 3 | 287 | 31.8 | 9.8 | 1.516 | 1.93 | 0.786 | 0.6 | -0.6 | 0.95 | 0.31666667 | -0.775 | -0.35 | 0.102595337 | 0.053581117 | 0.17016467 | 0.714598824 | 0.556350595 | 0.635474709 | 0.738844686 |
| Cse1l | High | Q9ERK4 | Exportin-2 OS=Mus musculus OX=10090 GN=Cse1l PE=1 SV=1 | 0 | 18.466 | 7 | 6 | 6 | 6 | 971 | 110.4 | 5.77 | 0.657 | 0.715 | 0.919 | -0.61 | 0.61 | -0.48 | -0.16 | 0.545 | -0.12 | 0.102354666 | 0.264115834 | 0.658026315 | 0.714598824 | 0.951072422 | 0.832835623 | 0.985010103 |
| Zc3h15 | High | Q3TIV5 | Zinc finger CCCH domain-containing protein 15 OS=Mus musculus OX=10090 GN=Zc3h15 PE=1 SV | 0 | 9.137 | 6 | 3 | 4 | 3 | 426 | 48.3 | 5.3 | 1.577 | 1.492 | 1.057 | 0.66 | -0.66 | 0.58 | 0.19333333 | -0.62 | 0.08 | 0.10295044 | 0.253608952 | 0.710670551 | 0.716182524 | 0.950034749 | 0.833108637 | 0.992535481 |
| Purg | High | Q8R4E6 | Purine-rich element-binding protein gamma OS=Mus musculus OX=10090 GN=Purg PE=1 SV=1 | 0 | 8.871 | 9 | 4 | 4 | 3 | 350 | 39.9 | 9.51 | 1.573 | 1.359 | 1.158 | 0.65 | -0.65 | 0.44 | 0.14666667 | -0.545 | 0.21 | 0.1034426 | 0.39283137 | 0.406360671 | 0.717825075 | 0.983884835 | 0.850854955 | 0.93629196 |
| Edf1 | High | Q9JMG1 | Endothelial differentiation-related factor 1 OS=Mus musculus OX=10090 GN=Edf1 PE=1 SV=1 | 0 | 11.576 | 18 | 3 | 6 | 3 | 148 | 16.4 | 9.99 | 1.624 | 1.573 | 1.032 | 0.7 | -0.7 | 0.65 | 0.21666667 | -0.675 | 0.05 | 0.103666502 | 0.192166045 | 0.861742634 | 0.718489591 | 0.907729584 | 0.813109588 | 0.992535481 |
| Dsp | High | E9Q557 | Desmoplakin OS=Mus musculus OX=10090 GN=Dsp PE=1 SV=1 | 0 | 15.598 | 2 | 6 | 6 | 6 | 2883 | 332.7 | 6.8 | 1.543 | 2.332 | 0.662 | 0.63 | -0.63 | 1.22 | 0.40666667 | -0.925 | -0.6 | 0.104770184 | 0.011204381 | 0.018469848 | 0.724348241 | 0.23978231 | 0.482065275 | 0.238069974 |
| Ppp6r3 | Medium | G5E8R4 | Serine/threonine-protein phosphatase 6 regulatory subunit 3 OS=Mus musculus OX=10090 GN=P | 0.045 | 1.582 | 1 | 1 | 1 | 1 | 873 | 97.8 | 4.58 | 0.667 | 0.667 | 1.001 | -0.58 | 0.58 | -0.58 | -0.1933333 | 0.58 | 0 | 0.105649464 | 0.166984265 | 0.937523565 | 0.725063089 | 0.871008827 | 0.798035958 | 0.992535481 |
| Slc20a2 | High | Q80UP8 | Sodium-dependent phosphate transporter 2 OS=Mus musculus OX=10090 GN=Slc20a2 PE=1 SV=2 | 0 | 47.963 | 18 | 8 | 10 | 7 | 656 | 70.8 | 6.48 | 0.638 | 0.726 | 0.88 | -0.65 | 0.65 | -0.46 | -0.1533333 | 0.555 | -0.18 | 0.105629826 | 0.284936728 | 0.434835614 | 0.725063089 | 0.955820031 | 0.84044156 | 0.949084077 |
| Btbd1 | Medium | P58544 | BTB/POZ domain-containing protein 1 OS=Mus musculus OX=10090 GN=Btbd1 PE=1 SV=3 | 0.013 | 2.352 | 2 | 1 | 1 | 1 | 488 | 53.2 | 6.1 | 0.641 | 0.743 | 0.863 | -0.64 | 0.64 | -0.43 | -0.1433333 | 0.535 | -0.21 | 0.105832409 | 0.295815988 | 0.523403941 | 0.725115998 | 0.955820031 | 0.840468014 | 0.964378027 |
| Zfp706 | High | Q9D115 | Zinc finger protein 706 OS=Mus musculus OX=10090 GN=Znf706 PE=1 SV=1 | 0.001 | 7.415 | 34 | 4 | 6 | 4 | 76 | 8.5 | 10.01 | 1.611 | 1.27 | 1.268 | 0.69 | -0.69 | 0.34 | 0.11333333 | -0.515 | 0.34 | 0.10591582 | 0.517422604 | 0.15896335 | 0.725115998 | 0.995559182 | 0.86033759 | 0.733684585 |
| Nt5e | High | Q61503 | 5'-nucleotidase OS=Mus musculus OX=10090 GN=Nt5e PE=1 SV=2 | 0 | 223.941 | 54 | 26 | 94 | 26 | 576 | 63.8 | 6.64 | 1.615 | 1.887 | 0.856 | 0.69 | -0.69 | 0.92 | 0.30666667 | -0.805 | -0.22 | 0.106238093 | 0.06265872 | 0.366079279 | 0.726435352 | 0.587451609 | 0.656943481 | 0.920164724 |
| Mlf2 | High | Q99KX1 | Myeloid leukemia factor 2 OS=Mus musculus OX=10090 GN=Mlf2 PE=1 SV=1 | 0 | 17.854 | 24 | 4 | 6 | 4 | 247 | 28 | 6.98 | 1.562 | 1.283 | 1.217 | 0.64 | -0.64 | 0.36 | 0.12 | -0.5 | 0.28 | 0.107051699 | 0.497870824 | 0.215808443 | 0.731107036 | 0.995559182 | 0.863333109 | 0.804011919 |
| Snrpd3 | High | P62320 | Small nuclear ribonucleoprotein Sm D3 OS=Mus musculus OX=10090 GN=Snrpd3 PE=1 SV=1 | 0.008 | 3.035 | 7 | 1 | 1 | 1 | 126 | 13.9 | 10.32 | 1.55 | 1.488 | 1.041 | 0.63 | -0.63 | 0.57 | 0.19 | -0.6 | 0.06 | 0.107238941 | 0.251251958 | 0.764020576 | 0.731494818 | 0.950034749 | 0.840764784 | 0.992535481 |
| Ccdc59 | Low | Q8R2N0 | Thyroid transcription factor 1-associated protein 26 OS=Mus musculus OX=10090 GN=Ccdc59 PE= | 0.079 | 1.215 | 3 | 1 | 1 | 1 | 240 | 28 | 10.13 | 1.532 | 1.497 | 1.023 | 0.62 | -0.62 | 0.58 | 0.19333333 | -0.6 | 0.03 | 0.107577425 | 0.234911726 | 0.860039243 | 0.732912054 | 0.936813316 | 0.834862685 | 0.992535481 |
| H1fx | High | Q80ZM5 | H1 histone family, member X OS=Mus musculus OX=10090 GN=H1f10 PE=1 SV=1 | 0 | 12.558 | 16 | 3 | 3 | 3 | 188 | 20.1 | 11.22 | 1.554 | 1.708 | 0.91 | 0.64 | -0.64 | 0.77 | 0.25666667 | -0.705 | -0.14 | 0.108058128 | 0.119723439 | 0.597713542 | 0.733347301 | 0.786053068 | 0.759700184 | 0.977451587 |
| Pafah1b3 | Medium | Q61205 | Platelet-activating factor acetylhydrolase IB subunit alpha1 OS=Mus musculus OX=10090 GN=Paf | 0.011 | 2.408 | 4 | 1 | 1 | 1 | 232 | 25.8 | 6.93 | 0.65 | 0.739 | 0.88 | -0.62 | 0.62 | -0.44 | -0.1466667 | 0.53 | -0.18 | 0.108164476 | 0.293309362 | 0.564658798 | 0.733347301 | 0.955820031 | 0.844583666 | 0.972910028 |
| Exoc4 | High | O35382 | Exocyst complex component 4 OS=Mus musculus OX=10090 GN=Exoc4 PE=1 SV=2 | 0.001 | 7.865 | 2 | 2 | 2 | 2 | 975 | 110.5 | 6.49 | 1.539 | 1.351 | 1.14 | 0.62 | -0.62 | 0.43 | 0.14333333 | -0.525 | 0.19 | 0.108482596 | 0.388493897 | 0.448980817 | 0.734615843 | 0.98165177 | 0.858133806 | 0.958608248 |

| | | | | | | | | | | | | | | | | | | | | | | | | | | | | |
|---|---|---|---|---|---|---|---|---|---|---|---|---|---|---|---|---|---|---|---|---|---|---|---|---|---|---|---|---|
| D630037F | Medium | Q3URV1 | Protein broad-minded OS=Mus musculus OX=10090 GN=Tbc1d32 PE=1 SV=2 | 0.042 | 1.627 | 1 | 1 | 1 | 1 | 1296 | 148 | 6.15 | 1.53 | 1.363 | 1.122 | 0.61 | -0.61 | 0.45 | 0.15 | -0.53 | 0.17 | 0.108873161 | 0.371723583 | 0.490441627 | 0.736371307 | 0.977313184 | 0.856842246 | 0.964378027 |
| Dbf4 | Medium | Q9QZ41 | Protein DBF4 homolog A OS=Mus musculus OX=10090 GN=Dbf4 PE=1 SV=1 | 0.037 | 1.724 | 3 | 1 | 1 | 1 | 663 | 74.1 | 7.93 | 1.566 | 1.707 | 0.917 | 0.65 | -0.65 | 0.77 | 0.25666667 | -0.71 | -0.12 | 0.109239934 | 0.110843667 | 0.645486227 | 0.737659696 | 0.769444739 | 0.753552217 | 0.985010103 |
| Gspt2 | High | Q149F3 | Eukaryotic peptide chain release factor GTP-binding subunit ERF3B OS=Mus musculus OX=10090 | 0 | 24.664 | 11 | 7 | 9 | 1 | 632 | 69.1 | 5.2 | 0.693 | 0.709 | 0.977 | -0.53 | 0.53 | -0.5 | -0.1666667 | 0.515 | -0.03 | 0.109326771 | 0.233601444 | 0.965220422 | 0.737659696 | 0.936813316 | 0.837236506 | 0.994302941 |
| Ggt1 | High | Q60928 | Glutathione hydrolase 1 proenzyme OS=Mus musculus OX=10090 GN=Ggt1 PE=1 SV=1 | 0 | 81.511 | 36 | 21 | 36 | 21 | 568 | 61.5 | 7.15 | 1.606 | 1.748 | 0.919 | 0.68 | -0.68 | 0.81 | 0.27 | -0.745 | -0.12 | 0.11013066 | 0.103667674 | 0.655927897 | 0.741299666 | 0.740464523 | 0.740882095 | 0.985010103 |
| Cox6a1 | Medium | P43024 | Cytochrome c oxidase subunit 6A1, mitochondrial OS=Mus musculus OX=10090 GN=Cox6a1 PE=1 | 0.01 | 2.702 | 26 | 1 | 1 | 1 | 111 | 12.3 | 9.98 | 1.546 | 1.504 | 1.027 | 0.63 | -0.63 | 0.59 | 0.19666667 | -0.61 | 0.04 | 0.110262872 | 0.23940979 | 0.822524325 | 0.741299666 | 0.937938091 | 0.839618878 | 0.992535481 |
| Ift172 | High | Q6VH22 | Intraflagellar transport protein 172 homolog OS=Mus musculus OX=10090 GN=Ift172 PE=1 SV=1 | 0 | 31.289 | 7 | 10 | 10 | 10 | 1749 | 197.4 | 6.02 | 1.526 | 1.569 | 0.973 | 0.61 | -0.61 | 0.65 | 0.21666667 | -0.63 | -0.04 | 0.110442409 | 0.195198246 | 0.838350386 | 0.741617467 | 0.912882441 | 0.827249954 | 0.992535481 |
| Zfp187; Zs | Medium | Q5RJ54 | Zinc finger and SCAN domain-containing protein 26 OS=Mus musculus OX=10090 GN=Zscan26 PE | 0.011 | 2.518 | 2 | 1 | 1 | 1 | 466 | 53.5 | 8.6 | 1.541 | 1.194 | 1.291 | 0.62 | -0.62 | 0.26 | 0.08666667 | -0.44 | 0.37 | 0.110901182 | 0.651898515 | 0.092906146 | 0.742918669 | 0.995559182 | 0.869238925 | 0.582038839 |
| Cpsf7 | High | Q8BTV2 | Cleavage and polyadenylation specificity factor subunit 7 OS=Mus musculus OX=10090 GN=Cpsf7 | 0.001 | 5.186 | 4 | 2 | 2 | 2 | 471 | 52 | 8 | 1.549 | 1.088 | 1.423 | 0.63 | -0.63 | 0.12 | 0.04 | -0.375 | 0.51 | 0.111206889 | 0.878108807 | 0.019831488 | 0.744077596 | 0.998402594 | 0.871240095 | 0.251572743 |
| Galnt10 | Medium | Q6P9S7 | Polypeptide N-acetylgalactosaminyltransferase 10 OS=Mus musculus OX=10090 GN=Galnt10 PE= | 0.014 | 2.242 | 1 | 1 | 1 | 1 | 603 | 69.1 | 8.43 | 0.651 | 0.615 | 1.058 | -0.62 | 0.62 | -0.7 | -0.2333333 | 0.66 | 0.08 | 0.111560108 | 0.104745506 | 0.743511077 | 0.745551282 | 0.742488054 | 0.744019668 | 0.992535481 |
| Pdc | High | Q9QW08 | Phosducin OS=Mus musculus OX=10090 GN=Pdc PE=2 SV=1 | 0 | 160.501 | 69 | 16 | 66 | 15 | 244 | 28 | 4.65 | 0.623 | 0.53 | 1.175 | -0.68 | 0.68 | -0.92 | -0.3066667 | 0.8 | 0.23 | 0.111777056 | 0.042030353 | 0.254308149 | 0.746111847 | 0.507896962 | 0.627004404 | 0.854859588 |
| Pfkp | High | Q8C605 | ATP-dependent 6-phosphofructokinase OS=Mus musculus OX=10090 GN=Pfkp PE=1 SV=1 | 0 | 332.405 | 50 | 42 | 127 | 34 | 784 | 85.5 | 6.89 | 0.625 | 0.426 | 1.468 | -0.68 | 0.68 | -1.23 | -0.41 | 0.955 | 0.55 | 0.113978991 | 0.006800655 | 0.01007944 | 0.75541395 | 0.169849505 | 0.462631728 | 0.16039109 |
| Rap1b | High | Q99JI6 | Ras-related protein Rap-1b OS=Mus musculus OX=10090 GN=Rap1b PE=1 SV=2 | 0 | 95.125 | 67 | 13 | 42 | 5 | 184 | 20.8 | 5.78 | 0.624 | 0.658 | 0.948 | -0.68 | 0.68 | -0.6 | -0.2 | 0.64 | -0.08 | 0.11365692 | 0.171265626 | 0.809090173 | 0.75541395 | 0.875484829 | 0.815449389 | 0.992535481 |
| Adcy6 | High | F8VQ52 | Adenylate cyclase type 6 OS=Mus musculus OX=10090 GN=Adcy6 PE=1 SV=1 | 0.001 | 7.751 | 2 | 3 | 3 | 3 | 1168 | 130.6 | 7.93 | 0.639 | 0.658 | 0.97 | -0.65 | 0.65 | -0.6 | -0.2 | 0.625 | -0.04 | 0.113830214 | 0.171346856 | 0.884632499 | 0.75541395 | 0.875484829 | 0.815449389 | 0.992535481 |
| Rhoa | High | Q9QUI0 | Transforming protein RhoA OS=Mus musculus OX=10090 GN=Rhoa PE=1 SV=1 | 0 | 113.389 | 57 | 12 | 39 | 4 | 193 | 21.8 | 6.1 | 0.625 | 0.639 | 0.978 | -0.68 | 0.68 | -0.65 | -0.2166667 | 0.665 | -0.03 | 0.114289277 | 0.144447639 | 0.964493785 | 0.756576125 | 0.838826712 | 0.797701418 | 0.994302941 |
| Cpt1a | High | P97742 | Carnitine O-palmitoyltransferase 1, liver isoform OS=Mus musculus OX=10090 GN=Cpt1a PE=1 SV | 0 | 4.942 | 3 | 2 | 3 | 1 | 773 | 88.2 | 8.62 | 1.553 | 1.828 | 0.849 | 0.63 | -0.63 | 0.87 | 0.29 | -0.75 | -0.24 | 0.114736485 | 0.069310047 | 0.361397403 | 0.757747319 | 0.612002256 | 0.684874788 | 0.917317898 |
| Slc25a12 | High | Q8BH59 | Calcium-binding mitochondrial carrier protein Aralar1 OS=Mus musculus OX=10090 GN=Slc25a12 | 0 | 82.404 | 28 | 14 | 22 | 14 | 677 | 74.5 | 8.25 | 1.596 | 1.648 | 0.968 | 0.67 | -0.67 | 0.72 | 0.24 | -0.695 | -0.05 | 0.114979633 | 0.147975833 | 0.914746231 | 0.758459769 | 0.845081896 | 0.801770833 | 0.992535481 |
| Wdr35 | High | Q8BND3 | WD repeat-containing protein 35 OS=Mus musculus OX=10090 GN=Wdr35 PE=1 SV=3 | 0 | 14.035 | 3 | 4 | 4 | 4 | 1181 | 133.9 | 6.34 | 1.532 | 1.224 | 1.251 | 0.62 | -0.62 | 0.29 | 0.09666667 | -0.455 | 0.32 | 0.11530025 | 0.58862922 | 0.161523969 | 0.759680967 | 0.995559182 | 0.877620075 | 0.733684585 |
| Msi2 | High | Q920Q6 | RNA-binding protein Musashi homolog 2 OS=Mus musculus OX=10090 GN=Msi2 PE=1 SV=1 | 0 | 20.571 | 16 | 5 | 7 | 4 | 346 | 36.9 | 8.47 | 1.489 | 1.236 | 1.205 | 0.57 | -0.57 | 0.31 | 0.10333333 | -0.44 | 0.27 | 0.115614543 | 0.572782333 | 0.281527393 | 0.760857682 | 0.995559182 | 0.878208432 | 0.881857034 |
| Got1 | High | P05201 | Aspartate aminotransferase, cytoplasmic OS=Mus musculus OX=10090 GN=Got1 PE=1 SV=3 | 0 | 74.395 | 43 | 17 | 37 | 17 | 413 | 46.2 | 7.14 | 0.627 | 0.647 | 0.97 | -0.67 | 0.67 | -0.63 | -0.21 | 0.65 | -0.04 | 0.117152523 | 0.154689088 | 0.922704284 | 0.769173534 | 0.847684847 | 0.808429191 | 0.992535481 |
| Lamp2 | High | P17047 | Lysosome-associated membrane glycoprotein 2 OS=Mus musculus OX=10090 GN=Lamp2 PE=1 SV | 0.001 | 7.497 | 2 | 1 | 2 | 1 | 415 | 45.7 | 7.39 | 0.682 | 0.958 | 0.712 | -0.55 | 0.55 | -0.06 | -0.02 | 0.305 | -0.49 | 0.117423003 | 0.808017021 | 0.052884251 | 0.770047691 | 0.998402594 | 0.884225143 | 0.438642008 |
| Adk | High | P55264 | Adenosine kinase OS=Mus musculus OX=10090 GN=Adk PE=1 SV=2 | 0.005 | 3.797 | 4 | 2 | 2 | 2 | 361 | 40.1 | 6.21 | 1.542 | 1.637 | 0.942 | 0.62 | -0.62 | 0.71 | 0.23666667 | -0.665 | -0.09 | 0.117613314 | 0.153989829 | 0.751106887 | 0.770394688 | 0.847684847 | 0.809039768 | 0.992535481 |
| Slc6a13 | High | P31649 | Sodium- and chloride-dependent GABA transporter 2 OS=Mus musculus OX=10090 GN=Slc6a13 P | 0 | 52.285 | 13 | 9 | 29 | 8 | 602 | 68.2 | 6.25 | 1.59 | 1.677 | 0.949 | 0.67 | -0.67 | 0.75 | 0.25 | -0.71 | -0.08 | 0.117814378 | 0.133844509 | 0.809954861 | 0.770811224 | 0.813954621 | 0.792382923 | 0.992535481 |
| Mip | High | P51180 | Lens fiber major intrinsic protein OS=Mus musculus OX=10090 GN=Mip PE=2 SV=2 | 0 | 13.474 | 8 | 1 | 4 | 1 | 263 | 28.2 | 9.41 | 0.68 | 0.694 | 0.979 | -0.56 | 0.56 | -0.53 | -0.1766667 | 0.545 | -0.03 | 0.118004185 | 0.207416831 | 0.976095182 | 0.771153225 | 0.912882441 | 0.842017833 | 0.997221123 |
| Rhot2 | High | Q8JZN7 | Mitochondrial Rho GTPase 2 OS=Mus musculus OX=10090 GN=Rhot2 PE=1 SV=1 | 0 | 17.749 | 11 | 5 | 5 | 4 | 620 | 69 | 6 | 1.56 | 1.394 | 1.119 | 0.64 | -0.64 | 0.48 | 0.16 | -0.56 | 0.16 | 0.119268395 | 0.34681954 | 0.439109045 | 0.776699062 | 0.977313184 | 0.877006123 | 0.952819047 |
| | High | A0A668KL | Microtubule-associated protein OS=Mus musculus OX=10090 GN=Map2 PE=1 SV=1 | 0 | 117.976 | 15 | 27 | 49 | 8 | 1989 | 216 | 4.84 | 1.587 | 1.12 | 1.416 | 0.67 | -0.67 | 0.16 | 0.05333333 | -0.415 | 0.5 | 0.119580924 | 0.792786489 | 0.019123833 | 0.776929594 | 0.998402594 | 0.887666094 | 0.245371467 |
| Ndufa4 | High | Q62425 | Cytochrome c oxidase subunit NDUFA4 OS=Mus musculus OX=10090 GN=Ndufa4 PE=1 SV=2 | 0 | 13.277 | 41 | 5 | 9 | 5 | 82 | 9.3 | 9.52 | 1.545 | 1.714 | 0.902 | 0.63 | -0.63 | 0.78 | 0.26 | -0.705 | -0.15 | 0.119776972 | 0.117286892 | 0.570634898 | 0.777214646 | 0.779640783 | 0.778427714 | 0.972910028 |
| Slc1a4 | High | O35874 | Neutral amino acid transporter A OS=Mus musculus OX=10090 GN=Slc1a4 PE=1 SV=1 | 0 | 23.142 | 12 | 5 | 8 | 4 | 532 | 56 | 5.87 | 0.665 | 0.668 | 0.995 | -0.59 | 0.59 | -0.58 | -0.1933333 | 0.585 | -0.01 | 0.119902028 | 0.186152672 | 0.920804237 | 0.777214646 | 0.900299151 | 0.838756898 | 0.992535481 |
| Ipo4 | High | Q8VI75 | Importin-4 OS=Mus musculus OX=10090 GN=Ipo4 PE=1 SV=1 | 0.009 | 2.816 | 1 | 1 | 2 | 1 | 1082 | 119.2 | 5.03 | 0.687 | 0.808 | 0.85 | -0.54 | 0.54 | -0.31 | -0.1033333 | 0.425 | -0.23 | 0.120093634 | 0.441564666 | 0.446901607 | 0.777557746 | 0.994630833 | 0.88609429 | 0.958608248 |
| Hspb11 | High | Q9D6H2 | Intraflagellar transport protein 25 homolog OS=Mus musculus OX=10090 GN=Hspb11 PE=1 SV=2 | 0.001 | 5.919 | 12 | 1 | 1 | 1 | 143 | 16.3 | 5.16 | 1.602 | 1.566 | 1.023 | 0.68 | -0.68 | 0.65 | 0.21666667 | -0.665 | 0.03 | 0.120900277 | 0.196247983 | 0.891458683 | 0.781877572 | 0.912882441 | 0.847380006 | 0.992535481 |
| Sugt1 | High | Q9CX34 | Protein SGT1 homolog OS=Mus musculus OX=10090 GN=Sugt1 PE=1 SV=3 | 0 | 16.763 | 17 | 5 | 5 | 5 | 336 | 38.1 | 5.45 | 0.655 | 0.701 | 0.935 | -0.61 | 0.61 | -0.51 | -0.17 | 0.56 | -0.1 | 0.121197273 | 0.239716643 | 0.729024507 | 0.782895288 | 0.937938091 | 0.860416689 | 0.992535481 |
| | High | A0A0G2JD | DCN1-like protein (Fragment) OS=Mus musculus OX=10090 GN=Dcun1d1 PE=1 SV=1 | 0.001 | 6.367 | 13 | 2 | 2 | 2 | 142 | 16.3 | 5.64 | 0.714 | 0.576 | 1.239 | -0.49 | 0.49 | -0.79 | -0.2633333 | 0.64 | 0.31 | 0.123214068 | 0.06447742 | 0.171772857 | 0.785962775 | 0.58835905 | 0.687160912 | 0.738844686 |
| | High | A0A0A6YY | Neural cell adhesion molecule 1 (Fragment) OS=Mus musculus OX=10090 GN=Ncam1 PE=1 SV=1 | 0 | 432.331 | 55 | 49 | 220 | 15 | 1089 | 116.3 | 4.83 | 0.633 | 0.675 | 0.937 | -0.66 | 0.66 | -0.57 | -0.19 | 0.615 | -0.09 | 0.124660421 | 0.196922318 | 0.751981363 | 0.793670211 | 0.912882441 | 0.853276326 | 0.992535481 |
| Urod | High | P70697 | Uroporphyrinogen decarboxylase OS=Mus musculus OX=10090 GN=Urod PE=1 SV=2 | 0.005 | 3.603 | 3 | 1 | 1 | 1 | 367 | 40.7 | 6.65 | 0.696 | 0.707 | 0.984 | -0.52 | 0.52 | -0.5 | -0.1666667 | 0.51 | -0.02 | 0.12470545 | 0.240564469 | 0.925034731 | 0.793670211 | 0.938986585 | 0.866328398 | 0.992535481 |
| Hist1h1c | High | P15864 | Histone H1.2 OS=Mus musculus OX=10090 GN=H1-2 PE=1 SV=2 | 0 | 82.408 | 47 | 18 | 38 | 5 | 212 | 21.3 | 11 | 1.575 | 1.966 | 0.801 | 0.66 | -0.66 | 0.98 | 0.32666667 | -0.82 | -0.32 | 0.125613446 | 0.046879847 | 0.184030497 | 0.798542619 | 0.520047141 | 0.65929488 | 0.767792634 |
| Hdhd2 | High | Q3UGR5 | Haloacid dehalogenase-like hydrolase domain-containing protein 2 OS=Mus musculus OX=10090 | 0 | 10.877 | 11 | 3 | 3 | 3 | 259 | 28.7 | 6.05 | 0.694 | 0.774 | 0.897 | -0.53 | 0.53 | -0.37 | -0.1233333 | 0.45 | -0.16 | 0.125869694 | 0.383888788 | 0.532162482 | 0.799265427 | 0.977313184 | 0.888289305 | 0.96694628 |
| Dip2b | High | B2RQC7 | DIP2 disco-interacting protein 2 homolog B (Drosophila) OS=Mus musculus OX=10090 GN=Dip2b | 0 | 52.654 | 14 | 15 | 20 | 1 | 1340 | 146.6 | 8.03 | 1.569 | 1.328 | 1.181 | 0.65 | -0.65 | 0.41 | 0.13666667 | -0.53 | 0.24 | 0.126135962 | 0.437881967 | 0.400899485 | 0.800050159 | 0.994630833 | 0.897340496 | 0.93262326 |
| Ralb | High | Q9JIW9 | Ras-related protein Ral-B OS=Mus musculus OX=10090 GN=Ralb PE=1 SV=1 | 0 | 45.47 | 35 | 7 | 22 | 3 | 206 | 23.3 | 6.62 | 0.65 | 0.606 | 1.074 | -0.62 | 0.62 | -0.72 | -0.24 | 0.67 | 0.1 | 0.126440488 | 0.104401733 | 0.651826167 | 0.800861047 | 0.741927143 | 0.771394095 | 0.985010103 |
| Pklr | High | G3X925 | Pyruvate kinase OS=Mus musculus OX=10090 GN=Pklr PE=1 SV=1 | 0 | 9.582 | 3 | 2 | 19 | 1 | 574 | 62.2 | 7.09 | 0.684 | 0.75 | 0.911 | -0.55 | 0.55 | -0.41 | -0.1366667 | 0.48 | -0.13 | 0.126835136 | 0.313977271 | 0.631997244 | 0.800861047 | 0.955820031 | 0.878340539 | 0.982425436 |
| S1pr3 | Medium | Q9Z0U9 | Sphingosine 1-phosphate receptor 3 OS=Mus musculus OX=10090 GN=S1pr3 PE=2 SV=1 | 0.047 | 1.563 | 3 | 1 | 2 | 1 | 378 | 42.2 | 9.55 | 0.663 | 0.467 | 1.421 | -0.59 | 0.59 | -1.1 | -0.3666667 | 0.845 | 0.51 | 0.127844532 | 0.01364437 | 0.080259216 | 0.806326538 | 0.270332091 | 0.538329314 | 0.544917434 |
| Iqcb1 | High | Q8BP00 | IQ calmodulin-binding motif-containing protein 1 OS=Mus musculus OX=10090 GN=Iqcb1 PE=1 SV | 0 | 17.235 | 8 | 4 | 4 | 4 | 598 | 68.7 | 9.31 | 1.479 | 1.763 | 0.839 | 0.56 | -0.56 | 0.82 | 0.27333333 | -0.69 | -0.25 | 0.129740699 | 0.093212419 | 0.334182611 | 0.807044546 | 0.702101282 | 0.754572914 | 0.906329766 |
| Fkbp4 | High | P30416 | Peptidyl-prolyl cis-trans isomerase FKBP4 OS=Mus musculus OX=10090 GN=Fkbp4 PE=1 SV=5 | 0 | 43.806 | 31 | 12 | 14 | 12 | 458 | 51.5 | 5.72 | 0.638 | 0.621 | 1.028 | -0.65 | 0.65 | -0.69 | -0.23 | 0.67 | 0.04 | 0.129807362 | 0.121321238 | 0.847352404 | 0.807044546 | 0.794682457 | 0.800863501 | 0.992535481 |
| Napa | High | Q9DB05 | Alpha-soluble NSF attachment protein OS=Mus musculus OX=10090 GN=Napa PE=1 SV=1 | 0 | 180.108 | 73 | 17 | 45 | 14 | 295 | 33.2 | 5.45 | 0.636 | 0.63 | 1.009 | -0.65 | 0.65 | -0.67 | -0.2233333 | 0.66 | 0.01 | 0.128170511 | 0.132799022 | 0.874326816 | 0.807044546 | 0.808473523 | 0.807759034 | 0.992535481 |
| 5430417L2 | High | A0A1L1ST | InaF motif-containing 2 OS=Mus musculus OX=10090 GN=Inafm2 PE=1 SV=1 | 0.001 | 5.254 | 7 | 1 | 1 | 1 | 152 | 15.6 | 9 | 1.493 | 1.673 | 0.892 | 0.58 | -0.58 | 0.74 | 0.24666667 | -0.66 | -0.16 | 0.129199202 | 0.135467187 | 0.509679903 | 0.807044546 | 0.817450082 | 0.812247314 | 0.964378027 |
| Acat1 | High | Q8QZT1 | Acetyl-CoA acetyltransferase, mitochondrial OS=Mus musculus OX=10090 GN=Acat1 PE=1 SV=1 | 0 | 104.395 | 45 | 16 | 31 | 16 | 424 | 44.8 | 8.51 | 1.567 | 1.663 | 0.943 | 0.65 | -0.65 | 0.73 | 0.24333333 | -0.69 | -0.09 | 0.12973152 | 0.140662503 | 0.779571301 | 0.807044546 | 0.831079719 | 0.819062132 | 0.992535481 |
| Fgd4 | High | Q91ZT5 | FYVE, RhoGEF and PH domain-containing protein 4 OS=Mus musculus OX=10090 GN=Fgd4 PE=1 S | 0.001 | 7.231 | 3 | 2 | 2 | 2 | 766 | 86.5 | 5.95 | 0.684 | 0.654 | 1.046 | -0.55 | 0.55 | -0.61 | -0.2033333 | 0.58 | 0.06 | 0.131196478 | 0.165342806 | 0.730624312 | 0.807044546 | 0.871008827 | 0.839026686 | 0.992535481 |
| Immt | High | Q8CAQ8 | MICOS complex subunit Mic60 OS=Mus musculus OX=10090 GN=Immt PE=1 SV=1 | 0 | 123.617 | 40 | 29 | 41 | 1 | 757 | 83.8 | 6.61 | 1.564 | 1.602 | 0.977 | 0.65 | -0.65 | 0.68 | 0.22666667 | -0.665 | -0.03 | 0.131412818 | 0.174195801 | 0.958540918 | 0.807044546 | 0.875938481 | 0.841491513 | 0.992902921 |
| Rbpms2 | Medium | Q9CWY6 | RNA-binding protein with multiple-splicing 2 OS=Mus musculus OX=10090 GN=Rbpms2 PE=1 SV= | 0.044 | 1.604 | 4 | 1 | 1 | 1 | 168 | 18.3 | 9.29 | 1.459 | 1.541 | 0.947 | 0.55 | -0.55 | 0.62 | 0.20666667 | -0.585 | -0.08 | 0.130999014 | 0.204386797 | 0.80079246 | 0.807044546 | 0.912882441 | 0.859963493 | 0.992535481 |
| Dctn3 | High | Q9Z0Y1 | Dynactin subunit 3 OS=Mus musculus OX=10090 GN=Dctn3 PE=1 SV=2 | 0 | 17.799 | 21 | 3 | 4 | 3 | 186 | 21 | 6.06 | 0.685 | 0.715 | 0.958 | -0.55 | 0.55 | -0.48 | -0.16 | 0.515 | -0.06 | 0.12892503 | 0.264083315 | 0.81887472 | 0.807044546 | 0.951072422 | 0.879058484 | 0.992535481 |
| Lrpprc | High | Q6PB66 | Leucine-rich PPR motif-containing protein, mitochondrial OS=Mus musculus OX=10090 GN=Lrpprc | 0 | 23.546 | 6 | 8 | 8 | 8 | 1392 | 156.5 | 6.83 | 1.442 | 1.328 | 1.086 | 0.53 | -0.53 | 0.41 | 0.13666667 | -0.47 | 0.12 | 0.131179095 | 0.43288861 | 0.631095299 | 0.807044546 | 0.99399649 | 0.900520518 | 0.9821125 |
| Ptms | High | Q9D0J8 | Parathymosin OS=Mus musculus OX=10090 GN=Ptms PE=1 SV=3 | 0.001 | 7.543 | 11 | 1 | 4 | 1 | 101 | 11.4 | 4.22 | 0.685 | 0.846 | 0.81 | -0.55 | 0.55 | -0.24 | -0.08 | 0.395 | -0.3 | 0.130780222 | 0.548140901 | 0.250813404 | 0.807044546 | 0.995559182 | 0.901301864 | 0.850762708 |
| Nsfl1c | High | Q9CZ44 | NSFL1 cofactor p47 OS=Mus musculus OX=10090 GN=Nsfl1c PE=1 SV=1 | 0 | 9.647 | 11 | 3 | 3 | 3 | 370 | 40.7 | 5.15 | 0.715 | 0.876 | 0.817 | -0.48 | 0.48 | -0.19 | -0.0633333 | 0.335 | -0.29 | 0.131042149 | 0.62176234 | 0.212286497 | 0.807044546 | 0.995559182 | 0.901301864 | 0.804011919 |
| Srsf7 | High | Q8BL97 | Serine/arginine-rich splicing factor 7 OS=Mus musculus OX=10090 GN=Srsf7 PE=1 SV=1 | 0.001 | 5.249 | 4 | 1 | 1 | 1 | 267 | 30.8 | 11.9 | 1.48 | 1.248 | 1.186 | 0.57 | -0.57 | 0.32 | 0.10666667 | -0.445 | 0.25 | 0.131183279 | 0.538270956 | 0.316426276 | 0.807044546 | 0.995559182 | 0.901301864 | 0.904943958 |
| Fam179b; | High | Q6A070 | TOG array regulator of axonemal microtubules protein 1 OS=Mus musculus OX=10090 GN=Togar | 0 | 19.629 | 5 | 7 | 8 | 7 | 1776 | 194.8 | 8.35 | 1.53 | 1.237 | 1.236 | 0.61 | -0.61 | 0.31 | 0.10333333 | -0.46 | 0.31 | 0.131279075 | 0.570557616 | 0.217793632 | 0.807044546 | 0.995559182 | 0.901301864 | 0.805520378 |
| | High | A0A0U1R( | Transient receptor potential cation channel subfamily M member 1 OS=Mus musculus OX=10090 | 0.001 | 5.578 | 1 | 2 | 2 | 2 | 1628 | 184.2 | 8.05 | 0.679 | 1.078 | 0.63 | -0.56 | 0.56 | 0.11 | 0.03666667 | 0.225 | -0.67 | 0.130127915 | 0.877290339 | 0.01012635 | 0.807044546 | 0.998402594 | 0.90272357 | 0.16039109 |
| Aip | High | O08915 | AH receptor-interacting protein OS=Mus musculus OX=10090 GN=Aip PE=1 SV=1 | 0 | 12.323 | 13 | 3 | 3 | 3 | 330 | 37.6 | 6.4 | 0.65 | 0.519 | 1.254 | -0.62 | 0.62 | -0.95 | -0.3166667 | 0.785 | 0.33 | 0.139365958 | 0.035892472 | 0.154080456 | 0.812244524 | 0.476392832 | 0.644318678 | 0.728999569 |
| Pex5l | High | D3YYH0 | PEX5-related protein OS=Mus musculus OX=10090 GN=Pex5l PE=1 SV=1 | 0 | 54.132 | 23 | 10 | 15 | 10 | 591 | 65.8 | 5.22 | 0.624 | 0.531 | 1.174 | -0.68 | 0.68 | -0.91 | -0.3033333 | 0.795 | 0.23 | 0.1377533 | 0.042902472 | 0.319240084 | 0.812244524 | 0.514004615 | 0.66312457 | 0.904943958 |
| Nrgn | High | P60761 | Neurogranin OS=Mus musculus OX=10090 GN=Nrgn PE=1 SV=1 | 0.008 | 3.027 | 19 | 1 | 1 | 1 | 78 | 7.5 | 7.05 | 1.511 | 1.922 | 0.786 | 0.6 | -0.6 | 0.94 | 0.31333333 | -0.77 | -0.35 | 0.13408525 | 0.051280671 | 0.206812732 | 0.812244524 | 0.541881751 | 0.677063137 | 0.79972344 |
| Osbp2 | High | Q5QNQ6 | Oxysterol-binding protein 2 OS=Mus musculus OX=10090 GN=Osbp2 PE=1 SV=1 | 0 | 45.834 | 13 | 11 | 19 | 10 | 908 | 101.3 | 7.14 | 1.569 | 1.718 | 0.913 | 0.65 | -0.65 | 0.78 | 0.26 | -0.715 | -0.13 | 0.138828969 | 0.115659527 | 0.625971208 | 0.812244524 | 0.77945068 | 0.795847602 | 0.980123029 |
| Atp5b | High | P56480 | ATP synthase subunit beta, mitochondrial OS=Mus musculus OX=10090 GN=Atp5f1b PE=1 SV=2 | 0 | 323.677 | 76 | 25 | 122 | 25 | 529 | 56.3 | 5.34 | 1.562 | 1.623 | 0.962 | 0.64 | -0.64 | 0.7 | 0.23333333 | -0.67 | -0.06 | 0.13289793 | 0.161771654 | 0.882562042 | 0.812244524 | 0.867993939 | 0.840119231 | 0.992535481 |
| Abca17 | Medium | E9PX95 | ATP-binding cassette sub-family A member 17 OS=Mus musculus OX=10090 GN=Abca17 PE=1 SV | 0.034 | 1.815 | 0 | 1 | 1 | 1 | 1733 | 195.9 | 7.83 | 0.69 | 0.663 | 1.041 | -0.54 | 0.54 | -0.59 | -0.1966667 | 0.565 | 0.06 | 0.136769653 | 0.169465565 | 0.803150523 | 0.812244524 | 0.874947902 | 0.843596213 | 0.992535481 |
| Prep | High | Q9QUR6 | Prolyl endopeptidase OS=Mus musculus OX=10090 GN=Prep PE=1 SV=1 | 0 | 14.102 | 7 | 5 | 5 | 5 | 710 | 80.7 | 5.73 | 0.663 | 0.681 | 0.973 | -0.59 | 0.59 | -0.55 | -0.1833333 | 0.57 | -0.04 | 0.137783939 | 0.206192975 | 0.900956138 | 0.812244524 | 0.912882441 | 0.862563482 | 0.992535481 |
| Hsdl1 | High | Q8BTX9 | Inactive hydroxysteroid dehydrogenase-like protein 1 OS=Mus musculus OX=10090 GN=Hsdl1 PE= | 0 | 17.298 | 17 | 4 | 4 | 4 | 330 | 36.8 | 8.48 | 1.504 | 1.552 | 0.969 | 0.59 | -0.59 | 0.63 | 0.21 | -0.61 | -0.05 | 0.138878637 | 0.206542541 | 0.894018643 | 0.812244524 | 0.912882441 | 0.862563482 | 0.992535481 |
| Mtfp1 | High | Q9CRB8 | Mitochondrial fission process protein 1 OS=Mus musculus OX=10090 GN=Mtfp1 PE=1 SV=1 | 0.006 | 3.386 | 7 | 1 | 1 | 1 | 166 | 18.3 | 8.68 | 1.466 | 1.528 | 0.96 | 0.55 | -0.55 | 0.61 | 0.20333333 | -0.58 | -0.06 | 0.138253237 | 0.21588565 | 0.845650066 | 0.812244524 | 0.920170644 | 0.866207584 | 0.992535481 |
| Atp5c1 | High | Q8C2Q8 | ATP synthase subunit gamma OS=Mus musculus OX=10090 GN=Atp5c1 PE=1 SV=1 | 0 | 71.144 | 42 | 13 | 38 | 13 | 274 | 30.2 | 8.84 | 1.551 | 1.539 | 1.008 | 0.63 | -0.63 | 0.62 | 0.20666667 | -0.625 | 0.01 | 0.138682643 | 0.216215087 | 0.879634278 | 0.812244524 | 0.920170644 | 0.866207584 | 0.992535481 |
| Tsr2 | High | H7BWY8 | Pre-rRNA-processing protein TSR2 homolog OS=Mus musculus OX=10090 GN=Tsr2 PE=1 SV=1 | 0.001 | 7.459 | 13 | 2 | 2 | 2 | 195 | 21.4 | 4.26 | 0.684 | 0.71 | 0.963 | -0.55 | 0.55 | -0.49 | -0.1633333 | 0.52 | -0.05 | 0.139792396 | 0.255891631 | 0.862148858 | 0.812244524 | 0.951072422 | 0.881658473 | 0.992535481 |
| Pafah1b2 | High | Q61206 | Platelet-activating factor acetylhydrolase IB subunit alpha2 OS=Mus musculus OX=10090 GN=Paf | 0 | 8.098 | 15 | 3 | 3 | 3 | 229 | 25.6 | 5.92 | 0.684 | 0.735 | 0.931 | -0.55 | 0.55 | -0.44 | -0.1466667 | 0.495 | -0.1 | 0.132566428 | 0.302778706 | 0.712171206 | 0.812244524 | 0.955820031 | 0.884032277 | 0.992535481 |
| Tbcel | Medium | Q8C5W3 | Tubulin-specific chaperone cofactor E-like protein OS=Mus musculus OX=10090 GN=Tbcel PE=1 S | 0.047 | 1.558 | 2 | 1 | 1 | 1 | 424 | 48 | 5.53 | 0.708 | 0.743 | 0.954 | -0.5 | 0.5 | -0.43 | -0.1433333 | 0.465 | -0.07 | 0.136988846 | 0.314768938 | 0.807581205 | 0.812244524 | 0.955820031 | 0.884032277 | 0.992535481 |
| Dhrs4 | High | Q99LB2 | Dehydrogenase/reductase SDR family member 4 OS=Mus musculus OX=10090 GN=Dhrs4 PE=1 S | 0.001 | 6.361 | 10 | 3 | 3 | 3 | 279 | 29.9 | 9.38 | 1.499 | 1.458 | 1.028 | 0.58 | -0.58 | 0.54 | 0.18 | -0.56 | 0.04 | 0.138666108 | 0.284361285 | 0.91611093 | 0.812244524 | 0.955820031 | 0.884032277 | 0.992535481 |
| Adam17 | High | E9PXU2 | Disintegrin and metalloproteinase domain-containing protein 17 OS=Mus musculus OX=10090 GN | 0.008 | 3.014 | 1 | 1 | 1 | 1 | 846 | 95.2 | 5.88 | 0.726 | 0.776 | 0.936 | -0.46 | 0.46 | -0.37 | -0.1233333 | 0.415 | -0.1 | 0.13926068 | 0.357112816 | 0.745482296 | 0.812244524 | 0.977313184 | 0.894778854 | 0.992535481 |
| Acadm | High | P45952 | Medium-chain specific acyl-CoA dehydrogenase, mitochondrial OS=Mus musculus OX=10090 GN= | 0 | 27.385 | 20 | 6 | 7 | 6 | 421 | 46.5 | 8.37 | 1.478 | 1.381 | 1.07 | 0.56 | -0.56 | 0.47 | 0.15666667 | -0.515 | 0.1 | 0.136811679 | 0.365525758 | 0.720702006 | 0.812244524 | 0.977313184 | 0.894778854 | 0.992535481 |
| Akr1b7 | Medium | P21300 | Aldo-keto reductase family 1 member B7 OS=Mus musculus OX=10090 GN=Akr1b7 PE=1 SV=4 | 0.01 | 2.715 | 3 | 1 | 3 | 1 | 316 | 36 | 7.25 | 0.658 | 0.797 | 0.826 | -0.6 | 0.6 | -0.33 | -0.11 | 0.465 | -0.28 | 0.135765676 | 0.433959418 | 0.259087501 | 0.812244524 | 0.99399649 | 0.903120507 | 0.85616924 |
| Kif3a | High | P28741 | Kinesin-like protein KIF3A OS=Mus musculus OX=10090 GN=Kif3a PE=1 SV=2 | 0 | 38.37 | 17 | 10 | 12 | 9 | 701 | 80.1 | 6.54 | 1.56 | 1.318 | 1.184 | 0.64 | -0.64 | 0.4 | 0.13333333 | -0.52 | 0.24 | 0.135088588 | 0.44669952 | 0.325695986 | 0.812244524 | 0.994630833 | 0.903437679 | 0.905686588 |
| Elavl2 | High | B1AXZ5 | ELAV-like protein OS=Mus musculus OX=10090 GN=Elavl2 PE=1 SV=1 | 0.003 | 3.846 | 4 | 2 | 3 | 2 | 389 | 42.6 | 9.04 | 1.472 | 1.255 | 1.173 | 0.56 | -0.56 | 0.33 | 0.11 | -0.445 | 0.23 | 0.137925918 | 0.541738152 | 0.358147424 | 0.812244524 | 0.995559182 | 0.903901853 | 0.914600864 |
| Rps21 | High | Q9CQR2 | 40S ribosomal protein S21 OS=Mus musculus OX=10090 GN=Rps21 PE=1 SV=1 | 0.006 | 3.258 | 11 | 1 | 1 | 1 | 83 | 9.1 | 8.51 | 1.486 | 1.262 | 1.178 | 0.57 | -0.57 | 0.34 | 0.11333333 | -0.455 | 0.24 | 0.13801665 | 0.516856752 | 0.335489641 | 0.812244524 | 0.995559182 | 0.903901853 | 0.906329766 |
| Aplp2 | High | Q60709 | Amyloid-like protein 2 OS=Mus musculus OX=10090 GN=Aplp2 PE=1 SV=1 | 0 | 18.357 | 10 | 5 | 5 | 1 | 751 | 85.2 | 4.73 | 0.674 | 1.003 | 0.672 | -0.57 | 0.57 | 0 | 0 | 0.285 | -0.57 | 0.139576048 | 0.911421851 | 0.061301043 | 0.812244524 | 0.998402594 | 0.905323559 | 0.476099813 |
| Gm7324 | High | A0A2I3BR | Predicted gene 7324 OS=Mus musculus OX=10090 GN=Gm7324 PE=4 SV=1 | 0.001 | 7.415 | 7 | 3 | 3 | 3 | 388 | 42.2 | 9.99 | 1.491 | 0.966 | 1.543 | 0.58 | -0.58 | -0.05 | -0.0166667 | -0.265 | 0.63 | 0.138573133 | 0.848202177 | 0.010363697 | 0.812244524 | 0.998402594 | 0.905323559 | 0.162316331 |
| Srsf1 | High | H7BX95 | Serine/arginine-rich splicing factor 1 OS=Mus musculus OX=10090 GN=Srsf1 PE=1 SV=1 | 0.001 | 7.6 | 8 | 2 | 2 | 2 | 253 | 28.3 | 10.08 | 1.53 | 1.089 | 1.405 | 0.61 | -0.61 | 0.12 | 0.04 | -0.365 | 0.49 | 0.134998806 | 0.860474615 | 0.02926811 | 0.812244524 | 0.998402594 | 0.905323559 | 0.324962961 |
| Aifm1 | High | B1AU25 | Apoptosis-inducing factor 1, mitochondrial OS=Mus musculus OX=10090 GN=Aifm1 PE=1 SV=1 | 0 | 13.534 | 8 | 4 | 4 | 4 | 608 | 66.1 | 9.11 | 1.523 | 1.558 | 0.978 | 0.61 | -0.61 | 0.64 | 0.21333333 | -0.625 | -0.03 | 0.140688083 | 0.195678492 | 0.932170521 | 0.813802165 | 0.912882441 | 0.863342303 | 0.992535481 |
| Ttc30a2 | High | A2AKQ8 | Tetratricopeptide repeat protein 30A2 OS=Mus musculus OX=10090 GN=Ttc30a2 PE=1 SV=1 | 0 | 13.294 | 8 | 5 | 5 | 5 | 664 | 76.1 | 5.16 | 1.505 | 1.421 | 1.059 | 0.59 | -0.59 | 0.51 | 0.17 | -0.55 | 0.08 | 0.141175414 | 0.320771575 | 0.739098461 | 0.813802165 | 0.955820031 | 0.884811098 | 0.992535481 |
| Aimp1 | High | Q3UZG4 | Aminoacyl tRNA synthase complex-interacting multifunctional protein 1 OS=Mus musculus OX=1( | 0 | 14.832 | 16 | 4 | 5 | 4 | 319 | 35.1 | 8.57 | 1.46 | 1.345 | 1.086 | 0.55 | -0.55 | 0.43 | 0.14333333 | -0.49 | 0.12 | 0.140379667 | 0.410513768 | 0.630794023 | 0.813802165 | 0.986182817 | 0.899992491 | 0.981916182 |
| Cspg5 | High | Q71M36 | Chondroitin sulfate proteoglycan 5 OS=Mus musculus OX=10090 GN=Cspg5 PE=1 SV=2 | 0 | 98.545 | 26 | 11 | 43 | 11 | 566 | 60.4 | 4.54 | 1.547 | 1.259 | 1.229 | 0.63 | -0.63 | 0.33 | 0.11 | -0.48 | 0.3 | 0.141221599 | 0.534844476 | 0.15338042 | 0.813802165 | 0.995559182 | 0.904680674 | 0.727583768 |
| Vamp2 | High | B0QZN5 | Synaptobrevin-2 OS=Mus musculus OX=10090 GN=Vamp2 PE=1 SV=1 | 0 | 65.067 | 25 | 3 | 17 | 3 | 163 | 17.9 | 9.38 | 0.627 | 0.563 | 1.115 | -0.67 | 0.67 | -0.83 | -0.2766667 | 0.75 | 0.16 | 0.143506549 | 0.064502966 | 0.488155379 | 0.819896178 | 0.58835905 | 0.704127614 | 0.964378027 |
| Api5 | High | O35841 | Apoptosis inhibitor 5 OS=Mus musculus OX=10090 GN=Api5 PE=1 SV=2 | 0.001 | 7.302 | 4 | 2 | 2 | 2 | 504 | 56.8 | 5.92 | 1.479 | 1.752 | 0.845 | 0.57 | -0.57 | 0.81 | 0.27 | -0.69 | -0.24 | 0.142943199 | 0.093184489 | 0.333397342 | 0.819896178 | 0.702101282 | 0.76099873 | 0.906329766 |
| Cul1 | High | Q9WTX6 | Cullin-1 OS=Mus musculus OX=10090 GN=Cul1 PE=1 SV=1 | 0 | 34.341 | 15 | 12 | 13 | 12 | 776 | 89.6 | 8 | 0.668 | 0.609 | 1.098 | -0.58 | 0.58 | -0.72 | -0.24 | 0.65 | 0.13 | 0.14308204 | 0.107818012 | 0.575381868 | 0.819896178 | 0.759466825 | 0.789681502 | 0.972910028 |
| Etfa | High | Q99LC5 | Electron transfer flavoprotein subunit alpha, mitochondrial OS=Mus musculus OX=10090 GN=Etfa | 0 | 15.499 | 21 | 5 | 5 | 5 | 333 | 35 | 8.38 | 1.465 | 1.441 | 1.017 | 0.55 | -0.55 | 0.53 | 0.17666667 | -0.54 | 0.02 | 0.142629583 | 0.300964922 | 0.918734399 | 0.819896178 | 0.955820031 | 0.887858104 | 0.992535481 |
| Ube2r2 | Medium | Q6ZWZ2 | Ubiquitin-conjugating enzyme E2 R2 OS=Mus musculus OX=10090 GN=Ube2r2 PE=1 SV=1 | 0.047 | 1.55 | 3 | 1 | 1 | 1 | 238 | 27.1 | 4.42 | 0.689 | 0.982 | 0.702 | -0.54 | 0.54 | -0.03 | -0.01 | 0.285 | -0.51 | 0.142883191 | 0.888701891 | 0.049247966 | 0.819896178 | 0.998402594 | 0.909149386 | 0.427566646 |
| Ensa | High | P60840 | Alpha-endosulfine OS=Mus musculus OX=10090 GN=Ensa PE=1 SV=1 | 0 | 9.821 | 31 | 3 | 4 | 2 | 121 | 13.3 | 7.24 | 0.704 | 0.515 | 1.368 | -0.51 | 0.51 | -0.96 | -0.32 | 0.735 | 0.45 | 0.14587519 | 0.03380774 | 0.073204962 | 0.822493948 | 0.473899991 | 0.64819697 | 0.516302163 |
| Grip1 | Medium | H7BX09 | Glutamate receptor-interacting protein 1 OS=Mus musculus OX=10090 GN=Grip1 PE=1 SV=1 | 0.042 | 1.642 | 1 | 1 | 1 | 1 | 1061 | 114.3 | 5.78 | 0.695 | 0.661 | 1.052 | -0.52 | 0.52 | -0.6 | -0.2 | 0.56 | 0.07 | 0.145957103 | 0.167304531 | 0.742034335 | 0.822493948 | 0.871008827 | 0.846751388 | 0.992535481 |
| Crym | High | O54983 | Ketimine reductase mu-crystallin OS=Mus musculus OX=10090 GN=Crym PE=1 SV=1 | 0 | 30.454 | 34 | 8 | 12 | 8 | 313 | 33.5 | 5.67 | 0.666 | 0.723 | 0.921 | -0.59 | 0.59 | -0.47 | -0.1566667 | 0.53 | -0.12 | 0.145462282 | 0.278988705 | 0.613983932 | 0.822493948 | 0.955820031 | 0.889156989 | 0.97826833 |
| Pcbp4 | High | P57724 | Poly(rC)-binding protein 4 OS=Mus musculus OX=10090 GN=Pcbp4 PE=1 SV=1 | 0 | 18.1 | 15 | 5 | 7 | 4 | 403 | 41.4 | 8.34 | 1.524 | 1.125 | 1.354 | 0.61 | -0.61 | 0.17 | 0.05666667 | -0.39 | 0.44 | 0.144575192 | 0.782407636 | 0.08113121 | 0.822493948 | 0.998402594 | 0.910448271 | 0.548736665 |
| Ppp6c | High | Q9CQR6 | Serine/threonine-protein phosphatase 6 catalytic subunit OS=Mus musculus OX=10090 GN=Ppp6c | 0 | 8.377 | 9 | 3 | 3 | 3 | 305 | 35.1 | 5.69 | 0.694 | 0.682 | 1.017 | -0.53 | 0.53 | -0.55 | -0.1833333 | 0.54 | 0.02 | 0.146493878 | 0.207728354 | 0.918764459 | 0.823862762 | 0.912882441 | 0.868372601 | 0.992535481 |
| Vps37b | High | Q8R0J7 | Vacuolar protein sorting-associated protein 37B OS=Mus musculus OX=10090 GN=Vps37b PE=1 S | 0 | 27.745 | 31 | 6 | 8 | 6 | 285 | 31 | 7.05 | 0.693 | 0.678 | 1.023 | -0.53 | 0.53 | -0.56 | -0.1866667 | 0.545 | 0.03 | 0.146860945 | 0.200688371 | 0.87778659 | 0.824273592 | 0.912882441 | 0.868578016 | 0.992535481 |
| Nlgn2 | High | Q69ZK9 | Neuroligin-2 OS=Mus musculus OX=10090 GN=Nlgn2 PE=1 SV=2 | 0.01 | 2.746 | 1 | 1 | 1 | 1 | 836 | 90.9 | 6.18 | 0.698 | 0.739 | 0.944 | -0.52 | 0.52 | -0.44 | -0.1466667 | 0.48 | -0.08 | 0.147574727 | 0.286671784 | 0.785589659 | 0.827332039 | 0.955820031 | 0.891576035 | 0.992535481 |
| Sucla2 | High | Q9Z2I9 | Succinate--CoA ligase [ADP-forming] subunit beta, mitochondrial OS=Mus musculus OX=10090 G | 0 | 57.588 | 31 | 13 | 15 | 13 | 463 | 50.1 | 7.01 | 1.492 | 1.356 | 1.1 | 0.58 | -0.58 | 0.44 | 0.14666667 | -0.51 | 0.14 | 0.147700976 | 0.395726613 | 0.475807614 | 0.827332039 | 0.984837604 | 0.906084821 | 0.962227984 |
| Mdh2 | High | P08249 | Malate dehydrogenase, mitochondrial OS=Mus musculus OX=10090 GN=Mdh2 PE=1 SV=3 | 0 | 195.645 | 68 | 20 | 72 | 20 | 338 | 35.6 | 8.68 | 1.534 | 1.688 | 0.909 | 0.62 | -0.62 | 0.75 | 0.25 | -0.685 | -0.14 | 0.148760516 | 0.128699762 | 0.608083034 | 0.829950459 | 0.799286338 | 0.814618398 | 0.97826833 |
| Gtf3c1 | High | Q8K284 | General transcription factor 3C polypeptide 1 OS=Mus musculus OX=10090 GN=Gtf3c1 PE=1 SV=2 | 0 | 20.671 | 4 | 8 | 8 | 8 | 2101 | 237.3 | 7.25 | 1.494 | 1.429 | 1.046 | 0.58 | -0.58 | 0.51 | 0.17 | -0.545 | 0.06 | 0.148569578 | 0.313364607 | 0.787979109 | 0.829950459 | 0.955820031 | 0.892885245 | 0.992535481 |
| Fau | High | P62862 | 40S ribosomal protein S30 OS=Mus musculus OX=10090 GN=Fau PE=1 SV=1 | 0.001 | 6.921 | 31 | 2 | 3 | 2 | 59 | 6.6 | 12.15 | 1.468 | 1.178 | 1.246 | 0.55 | -0.55 | 0.24 | 0.08 | -0.395 | 0.32 | 0.148759685 | 0.677215785 | 0.196725258 | 0.829950459 | 0.995559182 | 0.91275482 | 0.784953482 |
| Aars | High | Q8BGQ7 | Alanine--tRNA ligase, cytoplasmic OS=Mus musculus OX=10090 GN=Aars1 PE=1 SV=1 | 0 | 132.779 | 34 | 29 | 43 | 29 | 968 | 106.8 | 5.67 | 0.651 | 0.635 | 1.025 | -0.62 | 0.62 | -0.66 | -0.22 | 0.64 | 0.04 | 0.149297046 | 0.139085445 | 0.795016621 | 0.83211584 | 0.824367961 | 0.8282419 | 0.992535481 |
| Nolc1 | Medium | A0A286YD | Nucleolar and coiled-body phosphoprotein 1 (Fragment) OS=Mus musculus OX=10090 GN=Nolc1 | 0.037 | 1.733 | 1 | 1 | 1 | 1 | 702 | 73.6 | 9.36 | 1.52 | 2.246 | 0.677 | 0.6 | -0.6 | 1.17 | 0.39 | -0.885 | -0.56 | 0.151453632 | 0.017291347 | 0.050579319 | 0.832339849 | 0.305844112 | 0.569091981 | 0.427566646 |
| Dcxr | Medium | A2AC16 | L-xylulose reductase OS=Mus musculus OX=10090 GN=Dcxr PE=1 SV=1 | 0.01 | 2.653 | 3 | 1 | 1 | 1 | 236 | 25 | 6.55 | 0.738 | 0.504 | 1.463 | -0.44 | 0.44 | -0.99 | -0.33 | 0.715 | 0.55 | 0.159964801 | 0.023302235 | 0.020318605 | 0.832339849 | 0.379812885 | 0.606076367 | 0.253733677 |
| Ccdc51 | High | Q3URS9 | Mitochondrial potassium channel OS=Mus musculus OX=10090 GN=Ccdc51 PE=1 SV=1 | 0 | 13.045 | 7 | 2 | 5 | 2 | 406 | 45.1 | 8.09 | 1.453 | 1.936 | 0.751 | 0.54 | -0.54 | 0.95 | 0.31666667 | -0.745 | -0.41 | 0.155177076 | 0.047926404 | 0.127282555 | 0.832339849 | 0.520047141 | 0.676193495 | 0.678506823 |
| Csnk2a1 | High | Q60737 | Casein kinase II subunit alpha OS=Mus musculus OX=10090 GN=Csnk2a1 PE=1 SV=2 | 0 | 41.189 | 44 | 11 | 13 | 11 | 391 | 45.1 | 7.74 | 1.488 | 1.871 | 0.795 | 0.57 | -0.57 | 0.9 | 0.3 | -0.735 | -0.33 | 0.156100758 | 0.066408223 | 0.178207358 | 0.832339849 | 0.59705936 | 0.714699605 | 0.754689318 |
| Fam71d | Low | A0A1W2P | Family with sequence similarity 71, member D (Fragment) OS=Mus musculus OX=10090 GN=Fam | 0.064 | 1.306 | 6 | 1 | 3 | 1 | 109 | 12.4 | 5.2 | 1.451 | 1.748 | 0.83 | 0.54 | -0.54 | 0.81 | 0.27 | -0.675 | -0.27 | 0.160974406 | 0.098536027 | 0.30425471 | 0.832339849 | 0.720327908 | 0.776333879 | 0.900241969 |
| Eno2 | High | P17183 | Gamma-enolase OS=Mus musculus OX=10090 GN=Eno2 PE=1 SV=2 | 0 | 141.995 | 50 | 15 | 55 | 9 | 434 | 47.3 | 5.11 | 0.639 | 0.616 | 1.037 | -0.65 | 0.65 | -0.7 | -0.2333333 | 0.675 | 0.05 | 0.149697701 | 0.115656148 | 0.753728559 | 0.832339849 | 0.77945068 | 0.805895265 | 0.992535481 |
| Prdx2 | High | Q61171 | Peroxiredoxin-2 OS=Mus musculus OX=10090 GN=Prdx2 PE=1 SV=3 | 0 | 74 | 53 | 8 | 27 | 8 | 198 | 21.8 | 5.41 | 0.652 | 0.626 | 1.042 | -0.62 | 0.62 | -0.68 | -0.2266667 | 0.65 | 0.06 | 0.151620867 | 0.127914456 | 0.714216272 | 0.832339849 | 0.799286338 | 0.815813094 | 0.992535481 |
| Qars | High | A0A140LH | Glutaminyl-tRNA synthetase (Fragment) OS=Mus musculus OX=10090 GN=Qars PE=1 SV=1 | 0.001 | 5.32 | 7 | 2 | 2 | 2 | 246 | 27.5 | 8.09 | 0.724 | 0.631 | 1.147 | -0.47 | 0.47 | -0.66 | -0.22 | 0.565 | 0.2 | 0.16117792 | 0.128866568 | 0.42665053 | 0.832339849 | 0.799286338 | 0.815813094 | 0.941822646 |
| Dpp3 | High | Q99KK7 | Dipeptidyl peptidase 3 OS=Mus musculus OX=10090 GN=Dpp3 PE=1 SV=2 | 0 | 15.293 | 9 | 5 | 5 | 5 | 738 | 82.8 | 5.38 | 0.664 | 0.638 | 1.04 | -0.59 | 0.59 | -0.65 | -0.2166667 | 0.62 | 0.06 | 0.153528158 | 0.143203418 | 0.789017655 | 0.832339849 | 0.837891129 | 0.835115489 | 0.992535481 |
| Sars | High | Q8C483 | Seryl-tRNA synthetase OS=Mus musculus OX=10090 GN=Sars PE=1 SV=1 | 0 | 13.849 | 12 | 4 | 5 | 4 | 536 | 61.1 | 7.43 | 0.653 | 0.647 | 1.01 | -0.61 | 0.61 | -0.63 | -0.21 | 0.62 | 0.01 | 0.160342404 | 0.155566668 | 0.913919415 | 0.832339849 | 0.847684847 | 0.840012348 | 0.992535481 |
| Nsf | High | P46460 | Vesicle-fusing ATPase OS=Mus musculus OX=10090 GN=Nsf PE=1 SV=2 | 0 | 362.912 | 70 | 55 | 166 | 55 | 744 | 82.6 | 6.95 | 0.655 | 0.656 | 0.998 | -0.61 | 0.61 | -0.61 | -0.2033333 | 0.61 | 0 | 0.154805802 | 0.167884497 | 0.931297754 | 0.832339849 | 0.873217418 | 0.852778634 | 0.992535481 |
| Akap7 | High | Q7TN79 | A-kinase anchor protein 7 isoform gamma OS=Mus musculus OX=10090 GN=Akap7 PE=1 SV=2 | 0 | 15.511 | 7 | 2 | 4 | 2 | 314 | 35.5 | 7.87 | 0.719 | 0.681 | 1.057 | -0.48 | 0.48 | -0.55 | -0.1833333 | 0.515 | 0.08 | 0.1555796 | 0.205643755 | 0.698650076 | 0.832339849 | 0.912882441 | 0.872611145 | 0.991285312 |
| Timm44 | High | O35857 | Mitochondrial import inner membrane translocase subunit TIM44 OS=Mus musculus OX=10090 G | 0 | 19.488 | 10 | 5 | 5 | 5 | 452 | 51.1 | 8.13 | 1.46 | 1.551 | 0.941 | 0.55 | -0.55 | 0.63 | 0.21 | -0.59 | -0.09 | 0.160717469 | 0.207646777 | 0.690372373 | 0.832339849 | 0.912882441 | 0.872611145 | 0.991285312 |
| Ak4 | High | Q9WUR9 | Adenylate kinase 4, mitochondrial OS=Mus musculus OX=10090 GN=Ak4 PE=1 SV=1 | 0.001 | 6.848 | 13 | 2 | 2 | 2 | 223 | 25 | 7.53 | 1.494 | 1.557 | 0.959 | 0.58 | -0.58 | 0.64 | 0.21333333 | -0.61 | -0.06 | 0.161307126 | 0.203413133 | 0.83019066 | 0.832339849 | 0.912882441 | 0.872611145 | 0.992535481 |
| Inpp5e | High | A2AIX0 | Phosphatidylinositol polyphosphate 5-phosphatase type IV OS=Mus musculus OX=10090 GN=Inpp | 0 | 30.711 | 15 | 8 | 10 | 8 | 605 | 67.2 | 9.07 | 1.526 | 1.54 | 0.991 | 0.61 | -0.61 | 0.62 | 0.20666667 | -0.615 | -0.01 | 0.15145753 | 0.215267623 | 0.970207207 | 0.832339849 | 0.920170644 | 0.876255247 | 0.995598794 |
| Aldh16a1 | High | A0A1B0GS | Aldehyde dehydrogenase family 16 member A1 OS=Mus musculus OX=10090 GN=Aldh16a1 PE=1 | 0.005 | 3.493 | 2 | 1 | 1 | 1 | 802 | 84.7 | 6.32 | 0.713 | 0.692 | 1.03 | -0.49 | 0.49 | -0.53 | -0.1766667 | 0.51 | 0.04 | 0.153727395 | 0.218828573 | 0.862894561 | 0.832339849 | 0.923924555 | 0.878132202 | 0.992535481 |
| Eftud2 | High | G3UZ34 | 116 kDa U5 small nuclear ribonucleoprotein component OS=Mus musculus OX=10090 GN=Eftud2 | 0.001 | 5.124 | 2 | 2 | 5 | 1 | 962 | 108.3 | 5.02 | 1.471 | 1.518 | 0.969 | 0.56 | -0.56 | 0.6 | 0.2 | -0.58 | -0.05 | 0.158104499 | 0.234466149 | 0.929698655 | 0.832339849 | 0.936813316 | 0.884576583 | 0.992535481 |

| | | | | | | | | | | | | | | | | | | | | | | | | | | | | |
|---|---|---|---|---|---|---|---|---|---|---|---|---|---|---|---|---|---|---|---|---|---|---|---|---|---|---|---|---|
| Hnrnpm | High | B8JK33 | Heterogeneous nuclear ribonucleoprotein M OS=Mus musculus OX=10090 GN=Hnrnpm PE=1 SV= | 0 | 14.262 | 9 | 4 | 4 | 4 | 640 | 68.1 | 8.32 | 1.489 | 1.482 | 1.005 | 0.57 | -0.57 | 0.57 | 0.19 | -0.57 | 0.01 | 0.157770469 | 0.258339073 | 0.934753527 | 0.832339849 | 0.951072422 | 0.891706136 | 0.992535481 |
| Acad8 | High | D3YTT4 | Isobutyryl-CoA dehydrogenase, mitochondrial OS=Mus musculus OX=10090 GN=Acad8 PE=1 SV=1 | 0 | 10.6 | 9 | 4 | 4 | 4 | 413 | 45 | 8.02 | 1.47 | 1.44 | 1.021 | 0.56 | -0.56 | 0.53 | 0.17666667 | -0.545 | 0.03 | 0.154085045 | 0.302008563 | 0.85219516 | 0.832339849 | 0.955820031 | 0.89407994 | 0.992535481 |
| Ubqln1 | High | Q8R317 | Ubiquilin-1 OS=Mus musculus OX=10090 GN=Ubqln1 PE=1 SV=1 | 0 | 23.592 | 8 | 4 | 5 | 2 | 582 | 61.9 | 4.94 | 0.682 | 0.766 | 0.891 | -0.55 | 0.55 | -0.38 | -0.1266667 | 0.465 | -0.17 | 0.149604848 | 0.347491105 | 0.623709431 | 0.832339849 | 0.977313184 | 0.904826517 | 0.97840592 |
| Pura | High | P42669 | Transcriptional activator protein Pur-alpha OS=Mus musculus OX=10090 GN=Pura PE=1 SV=1 | 0 | 70.194 | 52 | 12 | 25 | 12 | 321 | 34.9 | 6.44 | 1.529 | 1.389 | 1.101 | 0.61 | -0.61 | 0.47 | 0.15666667 | -0.54 | 0.14 | 0.151976438 | 0.355992928 | 0.472615007 | 0.832339849 | 0.977313184 | 0.904826517 | 0.961211268 |
| Mapre3 | High | Q6PER3 | Microtubule-associated protein RP/EB family member 3 OS=Mus musculus OX=10090 GN=Mapre | 0 | 47.23 | 36 | 12 | 19 | 1 | 281 | 31.9 | 5.54 | 1.545 | 1.397 | 1.106 | 0.63 | -0.63 | 0.48 | 0.16 | -0.555 | 0.14 | 0.154509215 | 0.347341726 | 0.446170391 | 0.832339849 | 0.977313184 | 0.904826517 | 0.958608248 |
| Rpl24 | High | Q8BP67 | 60S ribosomal protein L24 OS=Mus musculus OX=10090 GN=Rpl24 PE=1 SV=2 | 0 | 39.11 | 34 | 9 | 16 | 9 | 157 | 17.8 | 11.25 | 1.533 | 1.361 | 1.126 | 0.62 | -0.62 | 0.45 | 0.15 | -0.535 | 0.17 | 0.150217521 | 0.389476217 | 0.388010106 | 0.832339849 | 0.982598494 | 0.907469172 | 0.93064581 |
| Cobll1 | High | Q3UMF0 | Cordon-bleu protein-like 1 OS=Mus musculus OX=10090 GN=Cobll1 PE=1 SV=2 | 0.006 | 3.39 | 1 | 1 | 1 | 1 | 1273 | 137.3 | 8.16 | 0.718 | 0.783 | 0.917 | -0.48 | 0.48 | -0.35 | -0.1166667 | 0.415 | -0.12 | 0.160816614 | 0.394109785 | 0.62260633 | 0.832339849 | 0.983884835 | 0.908112342 | 0.97826833 |
| Stk16 | Medium | Q05DP1 | Serine/threonine-protein kinase 16 OS=Mus musculus OX=10090 GN=Stk16 PE=1 SV=1 | 0.036 | 1.751 | 3 | 1 | 1 | 1 | 227 | 25.3 | 7.3 | 1.437 | 1.338 | 1.074 | 0.52 | -0.52 | 0.42 | 0.14 | -0.47 | 0.1 | 0.157535106 | 0.404380107 | 0.669069409 | 0.832339849 | 0.9855335 | 0.908936675 | 0.987489386 |
| Cep164 | High | D3YVU3 | Centrosomal protein of 164 kDa OS=Mus musculus OX=10090 GN=Cep164 PE=1 SV=1 | 0 | 9.283 | 2 | 3 | 3 | 3 | 1333 | 149.5 | 5.6 | 0.707 | 0.805 | 0.878 | -0.5 | 0.5 | -0.31 | -0.1033333 | 0.405 | -0.19 | 0.157354131 | 0.431513661 | 0.50455387 | 0.832339849 | 0.99399649 | 0.91316817 | 0.964378027 |
| Nradd | Low | A0A0G2JE | Death domain-containing membrane protein NRADD (Fragment) OS=Mus musculus OX=10090 GI | 0.069 | 1.303 | 12 | 1 | 1 | 1 | 59 | 6.7 | 9.88 | 1.465 | 1.307 | 1.121 | 0.55 | -0.55 | 0.39 | 0.13 | -0.47 | 0.16 | 0.161658301 | 0.447172461 | 0.490530156 | 0.832339849 | 0.994630833 | 0.913485341 | 0.964378027 |
| Tpd52l2 | High | A2AUD5 | Tumor protein D54 OS=Mus musculus OX=10090 GN=Tpd52l2 PE=1 SV=1 | 0 | 26.054 | 20 | 5 | 6 | 1 | 229 | 25.1 | 6.55 | 0.69 | 0.855 | 0.807 | -0.54 | 0.54 | -0.23 | -0.0766667 | 0.385 | -0.31 | 0.158698163 | 0.571249375 | 0.192195036 | 0.832339849 | 0.995559182 | 0.913949516 | 0.784495541 |
| Scgn | High | Q91WD9 | Secretagogin OS=Mus musculus OX=10090 GN=Scgn PE=2 SV=1 | 0 | 55.268 | 38 | 10 | 16 | 10 | 276 | 32.1 | 5.19 | 0.652 | 0.919 | 0.709 | -0.62 | 0.62 | -0.12 | -0.04 | 0.37 | -0.5 | 0.150617736 | 0.729748782 | 0.034562638 | 0.832339849 | 0.995559182 | 0.913949516 | 0.364272011 |
| C2cd5 | High | A0A0N4S\ | C2 domain-containing protein 5 (Fragment) OS=Mus musculus OX=10090 GN=C2cd5 PE=1 SV=3 | 0.005 | 3.653 | 6 | 2 | 2 | 2 | 232 | 26.4 | 5.16 | 0.739 | 0.83 | 0.89 | -0.44 | 0.44 | -0.27 | -0.09 | 0.355 | -0.17 | 0.161284703 | 0.49029665 | 0.50242626 | 0.832339849 | 0.995559182 | 0.913949516 | 0.964378027 |
| Ankrd13a | High | Q80UP5 | Ankyrin repeat domain-containing protein 13A OS=Mus musculus OX=10090 GN=Ankrd13a PE=1 S | 0 | 40.092 | 16 | 7 | 9 | 7 | 588 | 67.1 | 5.08 | 0.699 | 0.911 | 0.768 | -0.52 | 0.52 | -0.13 | -0.0433333 | 0.325 | -0.38 | 0.152066561 | 0.711070366 | 0.133326245 | 0.832339849 | 0.995559182 | 0.913949516 | 0.695404891 |
| Gcat | High | O88986 | 2-amino-3-ketobutyrate coenzyme A ligase, mitochondrial OS=Mus musculus OX=10090 GN=Gcat | 0 | 8.749 | 7 | 2 | 2 | 2 | 416 | 44.9 | 7.33 | 1.463 | 1.19 | 1.23 | 0.55 | -0.55 | 0.25 | 0.08333333 | -0.4 | 0.3 | 0.154837607 | 0.651024763 | 0.189998521 | 0.832339849 | 0.995559182 | 0.913949516 | 0.77990201 |
| Srsf10 | Medium | Q3TFP0 | Serine/arginine-rich-splicing factor 10 OS=Mus musculus OX=10090 GN=Srsf10 PE=1 SV=1 | 0.047 | 1.538 | 4 | 1 | 1 | 1 | 182 | 22.1 | 10.33 | 1.48 | 1.228 | 1.205 | 0.57 | -0.57 | 0.3 | 0.1 | -0.435 | 0.27 | 0.158521829 | 0.586263104 | 0.207215651 | 0.832339849 | 0.995559182 | 0.913949516 | 0.800177794 |
| Hfm1 | Low | D3Z4R1 | Probable ATP-dependent DNA helicase HFM1 OS=Mus musculus OX=10090 GN=Hfm1 PE=3 SV=2 | 0.054 | 1.41 | 0 | 1 | 1 | 1 | 1434 | 161.3 | 7.85 | 1.534 | 1.213 | 1.265 | 0.62 | -0.62 | 0.28 | 0.09333333 | -0.45 | 0.34 | 0.155943876 | 0.618064605 | 0.251473542 | 0.832339849 | 0.995559182 | 0.913949516 | 0.851275262 |
| Camsap2 | Medium | H7BX08 | Calmodulin-regulated spectrin-associated protein 2 OS=Mus musculus OX=10090 GN=Camsap2 P | 0.01 | 2.541 | 1 | 1 | 1 | 1 | 1478 | 166 | 6.84 | 1.517 | 1.276 | 1.189 | 0.6 | -0.6 | 0.35 | 0.11666667 | -0.475 | 0.25 | 0.158032528 | 0.522365363 | 0.360502372 | 0.832339849 | 0.995559182 | 0.913949516 | 0.916290479 |
| Me2 | High | Q99KE1 | NAD-dependent malic enzyme, mitochondrial OS=Mus musculus OX=10090 GN=Me2 PE=1 SV=1 | 0 | 13.214 | 9 | 5 | 5 | 5 | 589 | 65.8 | 7.61 | 1.503 | 1.304 | 1.152 | 0.59 | -0.59 | 0.38 | 0.12666667 | -0.485 | 0.2 | 0.158268059 | 0.466132614 | 0.419940436 | 0.832339849 | 0.995559182 | 0.913949516 | 0.938090049 |
| Msi1 | High | Q61474 | RNA-binding protein Musashi homolog 1 OS=Mus musculus OX=10090 GN=Msi1 PE=1 SV=1 | 0 | 31.318 | 28 | 6 | 7 | 5 | 362 | 39.1 | 7.87 | 1.503 | 1.068 | 1.408 | 0.59 | -0.59 | 0.09 | 0.03 | -0.34 | 0.49 | 0.151515291 | 0.908178603 | 0.034368004 | 0.832339849 | 0.998402594 | 0.915371222 | 0.364272011 |
| Fbl | High | P35550 | rRNA 2'-O-methyltransferase fibrillarin OS=Mus musculus OX=10090 GN=Fbl PE=1 SV=2 | 0.001 | 6.862 | 12 | 4 | 5 | 4 | 327 | 34.3 | 10.24 | 1.56 | 1.064 | 1.466 | 0.64 | -0.64 | 0.09 | 0.03 | -0.365 | 0.55 | 0.160184853 | 0.917307927 | 0.021476059 | 0.832339849 | 0.998402594 | 0.915371222 | 0.265234058 |
| Slc25a3 | High | G5E902 | Phosphate carrier protein, mitochondrial OS=Mus musculus OX=10090 GN=Slc25a3 PE=1 SV=1 | 0 | 84.853 | 31 | 13 | 39 | 13 | 358 | 39.7 | 9.29 | 1.514 | 1.634 | 0.926 | 0.6 | -0.6 | 0.71 | 0.23666667 | -0.655 | -0.11 | 0.162022169 | 0.155279485 | 0.693563657 | 0.832684051 | 0.847684847 | 0.840184449 | 0.991285312 |
| Dync1i2 | High | Q3TPJ8 | Cytoplasmic dynein 1 intermediate chain 2 OS=Mus musculus OX=10090 GN=Dync1i2 PE=1 SV=1 | 0 | 26.535 | 15 | 8 | 11 | 8 | 632 | 70.6 | 5.24 | 0.679 | 0.645 | 1.052 | -0.56 | 0.56 | -0.63 | -0.21 | 0.595 | 0.07 | 0.16237175 | 0.15296916 | 0.781443189 | 0.832953707 | 0.847684847 | 0.840319277 | 0.992535481 |
| Mtx2 | High | O88441 | Metaxin-2 OS=Mus musculus OX=10090 GN=Mtx2 PE=1 SV=1 | 0 | 12.432 | 15 | 3 | 4 | 3 | 263 | 29.7 | 5.63 | 1.472 | 1.746 | 0.843 | 0.56 | -0.56 | 0.8 | 0.26666667 | -0.68 | -0.25 | 0.163022859 | 0.095347162 | 0.334338576 | 0.833078748 | 0.708094752 | 0.77058675 | 0.906329766 |
| Gspt1 | High | Q8R050 | Eukaryotic peptide chain release factor GTP-binding subunit ERF3A OS=Mus musculus OX=10090 | 0 | 29.912 | 15 | 11 | 13 | 5 | 636 | 68.6 | 5.21 | 0.679 | 0.667 | 1.019 | -0.56 | 0.56 | -0.58 | -0.1933333 | 0.57 | 0.03 | 0.163374185 | 0.184262326 | 0.925019084 | 0.833078748 | 0.896839292 | 0.86495902 | 0.992535481 |
| Mindy2 | High | Q6PDI6 | Ubiquitin carboxyl-terminal hydrolase MINDY-2 OS=Mus musculus OX=10090 GN=Mindy2 PE=1 S\ | 0 | 49.513 | 23 | 9 | 12 | 8 | 601 | 65.6 | 4.63 | 0.673 | 0.708 | 0.95 | -0.57 | 0.57 | -0.5 | -0.1666667 | 0.535 | -0.07 | 0.162846664 | 0.252496795 | 0.762120294 | 0.833078748 | 0.950034749 | 0.891556749 | 0.992535481 |
| Lmod1 | High | Q8BVA4 | Leiomodin-1 OS=Mus musculus OX=10090 GN=Lmod1 PE=1 SV=1 | 0.005 | 3.692 | 3 | 1 | 1 | 1 | 595 | 66.3 | 9.19 | 1.439 | 1.483 | 0.971 | 0.53 | -0.53 | 0.57 | 0.19 | -0.55 | -0.04 | 0.16296368 | 0.247989008 | 0.89081617 | 0.833078748 | 0.950034749 | 0.891556749 | 0.992535481 |
| Appl2 | High | Q8K3G9 | DCC-interacting protein 13-beta OS=Mus musculus OX=10090 GN=Appl2 PE=1 SV=1 | 0.008 | 2.977 | 3 | 1 | 1 | 1 | 662 | 73.8 | 5.03 | 0.722 | 0.914 | 0.789 | -0.47 | 0.47 | -0.13 | -0.0433333 | 0.3 | -0.34 | 0.163242236 | 0.713297372 | 0.209794381 | 0.833078748 | 0.995559182 | 0.914318965 | 0.803698897 |
| Fam78b | High | F6ZXE6 | Protein FAM78B (Fragment) OS=Mus musculus OX=10090 GN=Fam78b PE=4 SV=1 | 0.001 | 4.652 | 7 | 2 | 2 | 2 | 254 | 29.1 | 9.47 | 0.741 | 0.931 | 0.796 | -0.43 | 0.43 | -0.1 | -0.0333333 | 0.265 | -0.33 | 0.163881908 | 0.761253294 | 0.15782435 | 0.833078748 | 0.995559182 | 0.914318965 | 0.733365826 |
| Ddx42 | High | Q810A7 | ATP-dependent RNA helicase DDX42 OS=Mus musculus OX=10090 GN=Ddx42 PE=1 SV=3 | 0.001 | 5.98 | 2 | 1 | 1 | 1 | 929 | 101.9 | 6.98 | 1.509 | 1.192 | 1.266 | 0.59 | -0.59 | 0.25 | 0.08333333 | -0.42 | 0.34 | 0.163452077 | 0.659301234 | 0.241154995 | 0.833078748 | 0.995559182 | 0.914318965 | 0.833635055 |
| Zfand2b | High | Q91X58 | AN1-type zinc finger protein 2B OS=Mus musculus OX=10090 GN=Zfand2b PE=1 SV=1 | 0.001 | 6.314 | 9 | 2 | 2 | 2 | 257 | 27.9 | 6.93 | 0.726 | 0.69 | 1.052 | -0.46 | 0.46 | -0.53 | -0.1766667 | 0.495 | 0.07 | 0.165090367 | 0.221299201 | 0.715921872 | 0.837702887 | 0.928185815 | 0.882944351 | 0.992535481 |
| Pfdn4 | High | Q3UWL8 | Prefoldin subunit 4 OS=Mus musculus OX=10090 GN=Pfdn4 PE=1 SV=1 | 0 | 9.413 | 21 | 2 | 2 | 2 | 134 | 15.2 | 4.61 | 0.708 | 0.654 | 1.083 | -0.5 | 0.5 | -0.61 | -0.2033333 | 0.555 | 0.11 | 0.165894524 | 0.147569598 | 0.585469329 | 0.839218264 | 0.845081896 | 0.84215008 | 0.973095093 |
| Atp5l | High | Q9CPQ8 | ATP synthase subunit g, mitochondrial OS=Mus musculus OX=10090 GN=Atp5mg PE=1 SV=1 | 0 | 35.858 | 49 | 4 | 15 | 4 | 103 | 11.4 | 9.74 | 1.508 | 1.598 | 0.944 | 0.59 | -0.59 | 0.68 | 0.22666667 | -0.635 | -0.08 | 0.167070909 | 0.17616506 | 0.791210701 | 0.839218264 | 0.875953959 | 0.857586111 | 0.992535481 |
| | High | A0A0N4S\ | Electron transfer flavoprotein subunit beta OS=Mus musculus OX=10090 GN=Etfbl PE=3 SV=1 | 0 | 20.07 | 34 | 6 | 6 | 6 | 221 | 24.1 | 6.1 | 1.456 | 1.548 | 0.941 | 0.54 | -0.54 | 0.63 | 0.21 | -0.585 | -0.09 | 0.166293644 | 0.20971484 | 0.685665933 | 0.839218264 | 0.912882441 | 0.876050352 | 0.991285312 |
| Csnk1e | Medium | Q3TYE1 | Casein kinase I isoform epsilon OS=Mus musculus OX=10090 GN=Csnk1e PE=1 SV=1 | 0.036 | 1.749 | 2 | 1 | 1 | 1 | 368 | 42.2 | 8.9 | 1.473 | 1.215 | 1.212 | 0.56 | -0.56 | 0.28 | 0.09333333 | -0.42 | 0.28 | 0.167484437 | 0.613100265 | 0.20187522 | 0.839218264 | 0.995559182 | 0.917388723 | 0.78697088 |
| Srsf3 | High | P84104 | Serine/arginine-rich splicing factor 3 OS=Mus musculus OX=10090 GN=Srsf3 PE=1 SV=1 | 0 | 20.862 | 17 | 3 | 8 | 3 | 164 | 19.3 | 11.65 | 1.521 | 1.027 | 1.48 | 0.6 | -0.6 | 0.04 | 0.01333333 | -0.32 | 0.57 | 0.16748411 | 0.997075101 | 0.020081144 | 0.839218264 | 0.998490794 | 0.918854529 | 0.252647811 |
| Smarcad1 | Low | Q04692 | SWI/SNF-related matrix-associated actin-dependent regulator of chromatin subfamily A containi | 0.081 | 1.195 | 1 | 1 | 1 | 1 | 1021 | 116.4 | 5.68 | 0.728 | 0.499 | 1.46 | -0.46 | 0.46 | -1 | -0.3333333 | 0.73 | 0.55 | 0.168011813 | 0.024020674 | 0.037697295 | 0.839623913 | 0.384371506 | 0.61199771 | 0.373579725 |
| Cs | High | Q9CZU6 | Citrate synthase, mitochondrial OS=Mus musculus OX=10090 GN=Cs PE=1 SV=1 | 0 | 76.277 | 41 | 15 | 39 | 15 | 464 | 51.7 | 8.57 | 1.504 | 1.711 | 0.879 | 0.59 | -0.59 | 0.77 | 0.25666667 | -0.68 | -0.19 | 0.168463867 | 0.118419397 | 0.465787981 | 0.839623913 | 0.783916832 | 0.811770372 | 0.960359242 |
| Atp5f1 | High | Q9CQQ7 | ATP synthase F(0) complex subunit B1, mitochondrial OS=Mus musculus OX=10090 GN=Atp5pb P | 0 | 44.321 | 24 | 7 | 19 | 7 | 256 | 28.9 | 9.06 | 1.505 | 1.575 | 0.955 | 0.59 | -0.59 | 0.66 | 0.22 | -0.625 | -0.07 | 0.168219343 | 0.190790553 | 0.844749344 | 0.839623913 | 0.905821916 | 0.872722915 | 0.992535481 |
| Prnp | High | P04925 | Major prion protein OS=Mus musculus OX=10090 GN=Prnp PE=1 SV=2 | 0 | 61.221 | 26 | 8 | 35 | 8 | 254 | 28 | 9.33 | 0.663 | 0.691 | 0.96 | -0.59 | 0.59 | -0.53 | -0.1766667 | 0.56 | -0.06 | 0.168103284 | 0.222485926 | 0.870558898 | 0.839623913 | 0.930954168 | 0.885289041 | 0.992535481 |
| Sncb | High | Q91ZZ3 | Beta-synuclein OS=Mus musculus OX=10090 GN=Sncb PE=1 SV=1 | 0 | 95.274 | 54 | 8 | 30 | 3 | 133 | 14 | 4.37 | 0.663 | 0.956 | 0.694 | -0.59 | 0.59 | -0.06 | -0.02 | 0.325 | -0.53 | 0.167997128 | 0.823471256 | 0.023934614 | 0.839623913 | 0.998402594 | 0.919013254 | 0.284324954 |
| Fam98a | High | Q3TJZ6 | Protein FAM98A OS=Mus musculus OX=10090 GN=Fam98a PE=1 SV=1 | 0 | 10.859 | 8 | 3 | 3 | 3 | 515 | 55 | 8.95 | 1.431 | 1.354 | 1.057 | 0.52 | -0.52 | 0.44 | 0.14666667 | -0.48 | 0.08 | 0.168792103 | 0.399440859 | 0.74366821 | 0.840512718 | 0.985440071 | 0.912976394 | 0.992535481 |
| Rpe | High | Q8VEE0 | Ribulose-phosphate 3-epimerase OS=Mus musculus OX=10090 GN=Rpe PE=1 SV=1 | 0.006 | 3.193 | 4 | 1 | 1 | 1 | 228 | 24.9 | 5.41 | 0.723 | 0.637 | 1.134 | -0.47 | 0.47 | -0.65 | -0.2166667 | 0.56 | 0.18 | 0.169924069 | 0.129569884 | 0.463627251 | 0.844649161 | 0.800990451 | 0.822819806 | 0.959744396 |
| Slc25a27 | High | Q9D6D0 | Solute carrier family 25, member 27 OS=Mus musculus OX=10090 GN=Slc25a27 PE=1 SV=1 | 0.003 | 3.947 | 7 | 1 | 1 | 1 | 322 | 35.8 | 9.39 | 1.451 | 2.009 | 0.722 | 0.54 | -0.54 | 1.01 | 0.33666667 | -0.775 | -0.47 | 0.171964679 | 0.037004201 | 0.065043596 | 0.847523331 | 0.480071477 | 0.663797404 | 0.491508685 |
| Rdh12 | High | Q8BYK4 | Retinol dehydrogenase 12 OS=Mus musculus OX=10090 GN=Rdh12 PE=2 SV=1 | 0 | 44.17 | 30 | 8 | 14 | 8 | 316 | 35.3 | 9.61 | 1.47 | 2.003 | 0.734 | 0.56 | -0.56 | 1 | 0.33333333 | -0.78 | -0.45 | 0.171787146 | 0.040815426 | 0.058149337 | 0.847523331 | 0.498588435 | 0.673055883 | 0.4657565 |
| 2310021P | High | Q3UHH1 | Zinc finger SWIM domain-containing protein 8 OS=Mus musculus OX=10090 GN=Zswim8 PE=1 S\ | 0 | 11.127 | 2 | 3 | 3 | 3 | 1832 | 196.9 | 6.84 | 0.72 | 0.64 | 1.126 | -0.47 | 0.47 | -0.64 | -0.2133333 | 0.555 | 0.17 | 0.171889992 | 0.128576802 | 0.473776788 | 0.847523331 | 0.799286338 | 0.823404834 | 0.962227984 |
| Rap2b | High | P61226 | Ras-related protein Rap-2b OS=Mus musculus OX=10090 GN=Rap2b PE=1 SV=1 | 0 | 32.381 | 42 | 6 | 11 | 3 | 183 | 20.5 | 4.81 | 0.718 | 0.744 | 0.965 | -0.48 | 0.48 | -0.43 | -0.1433333 | 0.455 | -0.05 | 0.171840995 | 0.321139019 | 0.835385434 | 0.847523331 | 0.955820031 | 0.901671681 | 0.992535481 |
| Timmdc1 | High | Q8BUY5 | Complex I assembly factor TIMMDC1, mitochondrial OS=Mus musculus OX=10090 GN=Timmdc1 I | 0.005 | 3.77 | 4 | 1 | 2 | 1 | 285 | 31.8 | 9.29 | 1.453 | 1.325 | 1.097 | 0.54 | -0.54 | 0.41 | 0.13666667 | -0.475 | 0.13 | 0.172013831 | 0.421352737 | 0.568960172 | 0.847523331 | 0.99399649 | 0.92075991 | 0.972910028 |
| Txn2 | High | A2A439 | Thioredoxin, mitochondrial OS=Mus musculus OX=10090 GN=Txn2 PE=1 SV=1 | 0.001 | 5.784 | 14 | 1 | 1 | 1 | 149 | 16.2 | 9.04 | 1.451 | 1.148 | 1.264 | 0.54 | -0.54 | 0.2 | 0.06666667 | -0.37 | 0.34 | 0.171672775 | 0.754176128 | 0.184040182 | 0.847523331 | 0.995559182 | 0.921541256 | 0.767792634 |
| CYTB | Low | P00158 | Cytochrome b OS=Mus musculus OX=10090 GN=Mt-Cyb PE=1 SV=1 | 0.056 | 1.359 | 2 | 1 | 1 | 1 | 381 | 43.2 | 7.97 | 1.443 | 1.332 | 1.083 | 0.53 | -0.53 | 0.41 | 0.13666667 | -0.47 | 0.12 | 0.172610405 | 0.429242473 | 0.571017574 | 0.849716017 | 0.99399649 | 0.921856253 | 0.972910028 |
| | High | O35516 | Neurogenic locus notch homolog protein 2 OS=Mus musculus OX=10090 GN=Notch2 PE=1 SV=2 | 0 | 24.944 | 5 | 9 | 10 | 9 | 2473 | 265.5 | 5.22 | 1.417 | 1.552 | 0.913 | 0.5 | -0.5 | 0.63 | 0.21 | -0.565 | -0.13 | 0.173091368 | 0.207012353 | 0.562308769 | 0.851336228 | 0.912882441 | 0.882109334 | 0.972910028 |
| Hist1h1e | High | P43274 | Histone H1.4 OS=Mus musculus OX=10090 GN=H1-4 PE=1 SV=2 | 0 | 70.662 | 41 | 15 | 34 | 4 | 219 | 22 | 11.11 | 1.445 | 1.858 | 0.777 | 0.53 | -0.53 | 0.89 | 0.29666667 | -0.71 | -0.36 | 0.17405047 | 0.069553269 | 0.139207254 | 0.855303232 | 0.613184245 | 0.734243739 | 0.701851981 |
| Sec24d | Medium | Q6NXL1 | Sec24-related gene family, member D (S. cerevisiae) OS=Mus musculus OX=10090 GN=Sec24d PE | 0.034 | 1.816 | 1 | 1 | 1 | 1 | 1032 | 112.6 | 7.18 | 1.435 | 1.608 | 0.892 | 0.52 | -0.52 | 0.69 | 0.23 | -0.605 | -0.17 | 0.174623494 | 0.158660114 | 0.521793014 | 0.857367715 | 0.854406564 | 0.85588714 | 0.964378027 |
| Cxadr | High | P97792 | Coxsackievirus and adenovirus receptor homolog OS=Mus musculus OX=10090 GN=Cxadr PE=1 SV | 0 | 14.572 | 12 | 4 | 6 | 4 | 365 | 39.9 | 6.96 | 0.69 | 0.732 | 0.942 | -0.54 | 0.54 | -0.45 | -0.15 | 0.495 | -0.09 | 0.175791815 | 0.298055424 | 0.689991728 | 0.861595024 | 0.955820031 | 0.908707527 | 0.991285312 |
| Npm1 | High | Q61937 | Nucleophosmin OS=Mus musculus OX=10090 GN=Npm1 PE=1 SV=1 | 0 | 48.584 | 30 | 13 | 20 | 13 | 292 | 32.5 | 4.77 | 1.477 | 2.019 | 0.731 | 0.56 | -0.56 | 1.01 | 0.33666667 | -0.785 | -0.45 | 0.176695896 | 0.038485082 | 0.055212693 | 0.862558172 | 0.486480307 | 0.67451924 | 0.450649022 |
| Cdkn2c | High | Q60772 | Cyclin-dependent kinase 4 inhibitor C OS=Mus musculus OX=10090 GN=Cdkn2c PE=1 SV=1 | 0.006 | 3.393 | 8 | 2 | 2 | 2 | 168 | 18.1 | 6 | 0.66 | 0.629 | 1.049 | -0.6 | 0.6 | -0.67 | -0.2233333 | 0.635 | 0.07 | 0.176565059 | 0.132228188 | 0.740363916 | 0.862558172 | 0.808473523 | 0.835515848 | 0.992535481 |
| | Medium | A0A1L1SU | Choline transporter-like protein 2 (Fragment) OS=Mus musculus OX=10090 GN=Slc44a2 PE=1 SV= | 0.037 | 1.739 | 9 | 1 | 1 | 1 | 170 | 19.7 | 6.52 | 0.693 | 0.688 | 1.007 | -0.53 | 0.53 | -0.54 | -0.18 | 0.535 | 0.01 | 0.176757507 | 0.200901963 | 0.948370549 | 0.862558172 | 0.912882441 | 0.887720306 | 0.992535481 |
| Luc7l3 | High | Q5SUF2 | Luc7-like protein 3 OS=Mus musculus OX=10090 GN=Luc7l3 PE=1 SV=1 | 0 | 16.509 | 7 | 2 | 3 | 2 | 432 | 51.4 | 9.77 | 1.456 | 1.131 | 1.287 | 0.54 | -0.54 | 0.18 | 0.06 | -0.36 | 0.36 | 0.176397771 | 0.780027069 | 0.102373905 | 0.862558172 | 0.998402594 | 0.930480383 | 0.610114517 |
| Rplp1 | High | P47955 | 60S acidic ribosomal protein P1 OS=Mus musculus OX=10090 GN=Rplp1 PE=1 SV=1 | 0 | 12.082 | 14 | 1 | 4 | 1 | 114 | 11.5 | 4.32 | 1.464 | 1.387 | 1.056 | 0.55 | -0.55 | 0.47 | 0.15666667 | -0.51 | 0.08 | 0.177213581 | 0.359189028 | 0.768452947 | 0.864031781 | 0.977313184 | 0.920672483 | 0.992535481 |
| Arl13b | High | Q640N2 | ADP-ribosylation factor-like protein 13B OS=Mus musculus OX=10090 GN=Arl13b PE=1 SV=2 | 0 | 165.721 | 59 | 24 | 49 | 24 | 427 | 48.1 | 6.32 | 1.491 | 1.469 | 1.015 | 0.58 | -0.58 | 0.55 | 0.18333333 | -0.565 | 0.02 | 0.177923035 | 0.273925966 | 0.84518627 | 0.866737148 | 0.955820031 | 0.91127859 | 0.992535481 |
| Pdcd5 | High | D3Z7Q5 | Programmed cell death protein 5 OS=Mus musculus OX=10090 GN=Pdcd5 PE=1 SV=1 | 0 | 8.368 | 18 | 4 | 6 | 4 | 205 | 22 | 7.87 | 0.721 | 0.657 | 1.097 | -0.47 | 0.47 | -0.61 | -0.2033333 | 0.54 | 0.13 | 0.180213558 | 0.1689387 | 0.589924179 | 0.86728355 | 0.874947902 | 0.871115726 | 0.974007893 |
| | High | E9Q9H3 | DnaJ homolog subfamily C member 2 OS=Mus musculus OX=10090 GN=Dnajc2 PE=1 SV=1 | 0.001 | 4.304 | 2 | 1 | 2 | 1 | 619 | 71.2 | 8.24 | 0.713 | 0.734 | 0.972 | -0.49 | 0.49 | -0.45 | -0.15 | 0.47 | -0.04 | 0.178564752 | 0.291777199 | 0.900263231 | 0.86728355 | 0.955820031 | 0.91155179 | 0.992535481 |
| Plaa | High | P27612 | Phospholipase A-2-activating protein OS=Mus musculus OX=10090 GN=Plaa PE=1 SV=4 | 0.001 | 6.787 | 3 | 3 | 3 | 3 | 794 | 87.2 | 6.14 | 0.734 | 0.74 | 0.992 | -0.45 | 0.45 | -0.43 | -0.1433333 | 0.44 | -0.01 | 0.180144398 | 0.291267645 | 0.98529779 | 0.86728355 | 0.955820031 | 0.91155179 | 0.998604519 |
| | High | F6SJM7 | Fucose mutarotase (Fragment) OS=Mus musculus OX=10090 GN=Fuom PE=1 SV=1 | 0.001 | 4.373 | 10 | 1 | 1 | 1 | 156 | 17.3 | 5.39 | 0.701 | 0.782 | 0.895 | -0.51 | 0.51 | -0.35 | -0.1166667 | 0.43 | -0.16 | 0.180252746 | 0.381644137 | 0.638765363 | 0.86728355 | 0.977313184 | 0.922298367 | 0.983597794 |
| Stx17 | High | Q9D0I4 | Syntaxin-17 OS=Mus musculus OX=10090 GN=Stx17 PE=1 SV=1 | 0 | 12.244 | 12 | 2 | 2 | 2 | 301 | 33.2 | 6.74 | 1.421 | 1.361 | 1.045 | 0.51 | -0.51 | 0.44 | 0.14666667 | -0.475 | 0.06 | 0.179550011 | 0.37859483 | 0.78057271 | 0.86728355 | 0.977313184 | 0.922298367 | 0.992535481 |
| Capn6 | High | O35646 | Calpain-6 OS=Mus musculus OX=10090 GN=Capn6 PE=1 SV=2 | 0 | 29.108 | 15 | 8 | 12 | 8 | 641 | 74.5 | 7.17 | 1.462 | 1.383 | 1.058 | 0.55 | -0.55 | 0.47 | 0.15666667 | -0.51 | 0.08 | 0.180410549 | 0.363739652 | 0.644888355 | 0.86728355 | 0.977313184 | 0.922298367 | 0.985010103 |
| Slc16a6 | High | B1AT66 | Monocarboxylate transporter 7 OS=Mus musculus OX=10090 GN=Slc16a6 PE=1 SV=1 | 0.006 | 3.236 | 3 | 1 | 1 | 1 | 607 | 66.2 | 8.81 | 0.698 | 0.878 | 0.794 | -0.52 | 0.52 | -0.19 | -0.0633333 | 0.355 | -0.33 | 0.179064674 | 0.60724826 | 0.288574618 | 0.86728355 | 0.995559182 | 0.931421366 | 0.888636466 |
| Prkce | High | P16054 | Protein kinase C epsilon type OS=Mus musculus OX=10090 GN=Prkce PE=1 SV=1 | 0 | 26.702 | 14 | 9 | 10 | 7 | 737 | 83.5 | 7.03 | 0.716 | 0.854 | 0.838 | -0.48 | 0.48 | -0.23 | -0.0766667 | 0.355 | -0.25 | 0.179150742 | 0.56844984 | 0.322331553 | 0.86728355 | 0.995559182 | 0.931421366 | 0.905686588 |
| Kif3b | High | Q61771 | Kinesin-like protein KIF3B OS=Mus musculus OX=10090 GN=Kif3b PE=1 SV=1 | 0 | 29.95 | 11 | 9 | 12 | 8 | 747 | 85.2 | 7.69 | 1.498 | 1.136 | 1.319 | 0.58 | -0.58 | 0.18 | 0.06 | -0.38 | 0.4 | 0.180139638 | 0.760338918 | 0.096171341 | 0.86728355 | 0.995559182 | 0.931421366 | 0.590616329 |
| Rpl12 | High | P35979 | 60S ribosomal protein L12 OS=Mus musculus OX=10090 GN=Rpl12 PE=1 SV=2 | 0 | 51.735 | 64 | 9 | 16 | 9 | 165 | 17.8 | 9.42 | 1.463 | 1.207 | 1.212 | 0.55 | -0.55 | 0.27 | 0.09 | -0.41 | 0.28 | 0.180510059 | 0.623966494 | 0.179966293 | 0.86728355 | 0.995559182 | 0.931421366 | 0.756556316 |
| Gorasp2 | High | Q99JX3 | Golgi reassembly-stacking protein 2 OS=Mus musculus OX=10090 GN=Gorasp2 PE=1 SV=3 | 0 | 29.381 | 22 | 8 | 9 | 8 | 451 | 47 | 4.79 | 0.687 | 0.697 | 0.985 | -0.54 | 0.54 | -0.52 | -0.1733333 | 0.53 | -0.02 | 0.181314719 | 0.2336968 | 0.88927289 | 0.86891592 | 0.936813316 | 0.902864618 | 0.992535481 |
| Rpl11 | High | Q9CXW4 | 60S ribosomal protein L11 OS=Mus musculus OX=10090 GN=Rpl11 PE=1 SV=4 | 0 | 16.025 | 17 | 3 | 6 | 3 | 178 | 20.2 | 9.6 | 1.499 | 1.314 | 1.14 | 0.58 | -0.58 | 0.39 | 0.13 | -0.485 | 0.19 | 0.181928635 | 0.452051674 | 0.344396021 | 0.869628182 | 0.995559182 | 0.932593682 | 0.907770836 |
| Pacsin2 | High | Q9WVE8 | Protein kinase C and casein kinase substrate in neurons protein 2 OS=Mus musculus OX=10090 GI | 0 | 64.793 | 30 | 15 | 25 | 6 | 486 | 55.8 | 5.2 | 0.672 | 0.597 | 1.125 | -0.57 | 0.57 | -0.74 | -0.2466667 | 0.655 | 0.17 | 0.182154142 | 0.095537664 | 0.389376731 | 0.869964457 | 0.708251892 | 0.789108174 | 0.93064581 |
| Gm98; My | High | Q3UR85 | Myelin regulatory factor OS=Mus musculus OX=10090 GN=Myrf PE=1 SV=2 | 0.01 | 2.746 | 1 | 1 | 1 | 1 | 1138 | 123.2 | 7.18 | 1.417 | 1.712 | 0.828 | 0.5 | -0.5 | 0.78 | 0.26 | -0.64 | -0.27 | 0.183368287 | 0.113474077 | 0.299482322 | 0.87501786 | 0.77945068 | 0.82723427 | 0.900241969 |
| Gdpd1 | Medium | Q9CRY7 | Lysophospholipase D GDPD1 OS=Mus musculus OX=10090 GN=Gdpd1 PE=1 SV=1 | 0.011 | 2.466 | 3 | 1 | 1 | 1 | 314 | 35.8 | 8.31 | 1.425 | 1.562 | 0.912 | 0.51 | -0.51 | 0.64 | 0.21333333 | -0.575 | -0.13 | 0.183668632 | 0.187539166 | 0.614224629 | 0.875705799 | 0.902623507 | 0.889164653 | 0.97826833 |
| Atp5j | High | P97450 | ATP synthase-coupling factor 6, mitochondrial OS=Mus musculus OX=10090 GN=Atp5pf PE=1 SV= | 0 | 27.398 | 50 | 6 | 11 | 6 | 108 | 12.5 | 9.36 | 1.447 | 1.522 | 0.95 | 0.53 | -0.53 | 0.61 | 0.20333333 | -0.57 | -0.07 | 0.184031842 | 0.229124502 | 0.819950886 | 0.875947827 | 0.935008675 | 0.905478251 | 0.992535481 |
| Slc4a3 | High | P16283 | Anion exchange protein 3 OS=Mus musculus OX=10090 GN=Slc4a3 PE=1 SV=2 | 0.01 | 2.754 | 1 | 1 | 1 | 1 | 1227 | 135.3 | 6.51 | 0.749 | 1.013 | 0.739 | -0.42 | 0.42 | 0.02 | 0.00666667 | 0.2 | -0.44 | 0.183901617 | 0.954189892 | 0.06163737 | 0.875947827 | 0.998402594 | 0.93717521 | 0.476099813 |
| Agpat3 | High | Q9D517 | 1-acyl-sn-glycerol-3-phosphate acyltransferase gamma OS=Mus musculus OX=10090 GN=Agpat3 | 0 | 34.126 | 20 | 9 | 14 | 9 | 376 | 43.3 | 8.51 | 1.467 | 2.093 | 0.701 | 0.55 | -0.55 | 1.07 | 0.35666667 | -0.81 | -0.51 | 0.189176847 | 0.029246232 | 0.028438163 | 0.876837873 | 0.434969823 | 0.655903848 | 0.322126832 |
| Tmem41b | High | Q8K1A5 | Transmembrane protein 41B OS=Mus musculus OX=10090 GN=Tmem41b PE=1 SV=1 | 0.001 | 4.263 | 3 | 1 | 1 | 1 | 291 | 32.4 | 9.31 | 1.422 | 1.957 | 0.727 | 0.51 | -0.51 | 0.97 | 0.32333333 | -0.74 | -0.46 | 0.18967546 | 0.044378148 | 0.06811501 | 0.876837873 | 0.520047141 | 0.698442507 | 0.499896415 |
| Gmppb | High | Q8BTZ7 | Mannose-1-phosphate guanyltransferase beta OS=Mus musculus OX=10090 GN=Gmppb PE=1 SV | 0.001 | 5.789 | 3 | 1 | 1 | 1 | 360 | 39.9 | 6.74 | 0.723 | 0.563 | 1.284 | -0.47 | 0.47 | -0.83 | -0.2766667 | 0.65 | 0.36 | 0.186978368 | 0.054092041 | 0.137075546 | 0.876837873 | 0.560617514 | 0.718727693 | 0.700189449 |
| | High | A0A2I3BQ | Ubiquitin carboxyl-terminal hydrolase (Fragment) OS=Mus musculus OX=10090 GN=Uchl3 PE=1 S | 0 | 8.148 | 3 | 1 | 2 | 1 | 265 | 29.7 | 6.76 | 0.688 | 0.595 | 1.156 | -0.54 | 0.54 | -0.75 | -0.25 | 0.645 | 0.21 | 0.193200234 | 0.093224708 | 0.368037275 | 0.876837873 | 0.702101282 | 0.789469577 | 0.920164724 |
| March5 | High | A0A0R4J1 | E3 ubiquitin-protein ligase MARCHF5 OS=Mus musculus OX=10090 GN=Marchf5 PE=1 SV=1 | 0.009 | 2.835 | 4 | 1 | 1 | 1 | 244 | 27.3 | 8.47 | 1.422 | 1.747 | 0.814 | 0.51 | -0.51 | 0.8 | 0.26666667 | -0.655 | -0.3 | 0.18741941 | 0.102614158 | 0.250322767 | 0.876837873 | 0.734811727 | 0.8058248 | 0.849612441 |
| Dlst | High | Q9D2G2 | Dihydrolipoyllysine-residue succinyltransferase component of 2-oxoglutarate dehydrogenase com | 0 | 36.523 | 17 | 8 | 14 | 8 | 454 | 49 | 8.95 | 1.441 | 1.68 | 0.858 | 0.53 | -0.53 | 0.75 | 0.25 | -0.64 | -0.22 | 0.194257142 | 0.13235661 | 0.373910112 | 0.876837873 | 0.808473523 | 0.842655698 | 0.920164724 |
| Apool | High | Q78IK4 | MICOS complex subunit Mic27 OS=Mus musculus OX=10090 GN=Apool PE=1 SV=1 | 0 | 18.837 | 20 | 4 | 4 | 4 | 265 | 29.2 | 9.31 | 1.464 | 1.643 | 0.891 | 0.55 | -0.55 | 0.72 | 0.24 | -0.635 | -0.17 | 0.187559672 | 0.15063468 | 0.510506279 | 0.876837873 | 0.847684847 | 0.86226136 | 0.964378027 |
| Gls | High | D3Z7P3 | Glutaminase kidney isoform, mitochondrial OS=Mus musculus OX=10090 GN=Gls PE=1 SV=1 | 0 | 70.5 | 25 | 12 | 19 | 12 | 674 | 73.9 | 7.99 | 1.471 | 1.605 | 0.917 | 0.56 | -0.56 | 0.68 | 0.22666667 | -0.62 | -0.13 | 0.192668466 | 0.17217017 | 0.645725684 | 0.876837873 | 0.875484829 | 0.876161351 | 0.985010103 |
| | Medium | E9Q811 | Serine/threonine-protein kinase STK11 OS=Mus musculus OX=10090 GN=Stk11 PE=1 SV=1 | 0.01 | 2.534 | 5 | 1 | 1 | 1 | 310 | 34.6 | 5.44 | 0.724 | 0.671 | 1.079 | -0.47 | 0.47 | -0.58 | -0.1933333 | 0.525 | 0.11 | 0.19451998 | 0.174257348 | 0.651877905 | 0.876837873 | 0.875938481 | 0.876388177 | 0.985010103 |
| Snrpc | High | Q62241 | U1 small nuclear ribonucleoprotein C OS=Mus musculus OX=10090 GN=Snrpc PE=1 SV=1 | 0.006 | 3.425 | 6 | 1 | 2 | 1 | 159 | 17.4 | 9.67 | 1.397 | 1.599 | 0.874 | 0.48 | -0.48 | 0.68 | 0.22666667 | -0.58 | -0.2 | 0.191865022 | 0.173812897 | 0.446144597 | 0.876837873 | 0.875938481 | 0.876388177 | 0.958608248 |
| Atp5o; LO | High | Q9DB20 | ATP synthase subunit O, mitochondrial OS=Mus musculus OX=10090 GN=Atp5po PE=1 SV=1 | 0 | 76.285 | 51 | 12 | 27 | 12 | 213 | 23.3 | 9.99 | 1.469 | 1.587 | 0.926 | 0.56 | -0.56 | 0.67 | 0.22333333 | -0.615 | -0.11 | 0.194428521 | 0.18325038 | 0.691249032 | 0.876837873 | 0.895493677 | 0.886165775 | 0.991285312 |
| | High | A0A0R4J1 | Acyl-peptide hydrolase (Fragment) OS=Mus musculus OX=10090 GN=Apeh PE=1 SV=1 | 0 | 12.008 | 5 | 3 | 3 | 3 | 716 | 79.8 | 5.59 | 0.722 | 0.685 | 1.054 | -0.47 | 0.47 | -0.55 | -0.1833333 | 0.51 | 0.08 | 0.188142272 | 0.191050112 | 0.730234519 | 0.876837873 | 0.905821916 | 0.891329895 | 0.992535481 |
| Adss | High | P46664 | Adenylosuccinate synthetase isozyme 2 OS=Mus musculus OX=10090 GN=Adss2 PE=1 SV=2 | 0.001 | 5.476 | 2 | 1 | 1 | 1 | 456 | 50 | 6.38 | 0.716 | 0.696 | 1.027 | -0.48 | 0.48 | -0.52 | -0.1733333 | 0.5 | 0.04 | 0.192460151 | 0.232013851 | 0.81509993 | 0.876837873 | 0.936813316 | 0.906825594 | 0.992535481 |
| Atp5h | High | Q9DCX2 | ATP synthase subunit d, mitochondrial OS=Mus musculus OX=10090 GN=Atp5pd PE=1 SV=3 | 0 | 64.477 | 81 | 10 | 23 | 10 | 161 | 18.7 | 5.69 | 1.468 | 1.49 | 0.985 | 0.55 | -0.55 | 0.58 | 0.19333333 | -0.565 | -0.02 | 0.195325194 | 0.255403654 | 0.996481737 | 0.876837873 | 0.951072422 | 0.913955148 | 0.999917596 |
| Inpp4a | High | D3YUL8 | Phosphatidylinositol-3,4-bisphosphate 4-phosphatase OS=Mus musculus OX=10090 GN=Inpp4a PE | 0.001 | 7.182 | 2 | 2 | 2 | 2 | 972 | 109 | 7.11 | 0.732 | 0.745 | 0.982 | -0.45 | 0.45 | -0.42 | -0.14 | 0.435 | -0.03 | 0.195634774 | 0.299222637 | 0.941929196 | 0.876837873 | 0.955820031 | 0.916328952 | 0.992535481 |
| Pdhx | High | Q8BKZ9 | Pyruvate dehydrogenase protein X component, mitochondrial OS=Mus musculus OX=10090 GN=P | 0 | 10.076 | 7 | 3 | 3 | 3 | 501 | 54 | 7.75 | 1.456 | 1.422 | 1.024 | 0.54 | -0.54 | 0.51 | 0.17 | -0.525 | 0.03 | 0.192894295 | 0.319800348 | 0.888984286 | 0.876837873 | 0.955820031 | 0.916328952 | 0.992535481 |
| Uqcrfs1 | High | Q9CR68 | Cytochrome b-c1 complex subunit Rieske, mitochondrial OS=Mus musculus OX=10090 GN=Uqcrfs | 0 | 49.194 | 36 | 8 | 13 | 8 | 274 | 29.3 | 8.7 | 1.513 | 1.426 | 1.061 | 0.6 | -0.6 | 0.51 | 0.17 | -0.555 | 0.09 | 0.186716708 | 0.315597658 | 0.69830393 | 0.876837873 | 0.955820031 | 0.916328952 | 0.991285312 |
| Sod2 | High | P09671 | Superoxide dismutase [Mn], mitochondrial OS=Mus musculus OX=10090 GN=Sod2 PE=1 SV=3 | 0 | 63.178 | 45 | 10 | 30 | 10 | 222 | 24.6 | 8.62 | 1.477 | 1.466 | 1.008 | 0.56 | -0.56 | 0.55 | 0.18333333 | -0.555 | 0.01 | 0.188023338 | 0.276606182 | 0.881390696 | 0.876837873 | 0.955820031 | 0.916328952 | 0.992535481 |
| | High | Q5U4F6 | Cytoplasmic dynein 2 intermediate chain 2 OS=Mus musculus OX=10090 GN=Dync2i2 PE=1 SV=3 | 0.001 | 7.732 | 4 | 2 | 2 | 2 | 537 | 58.1 | 6.52 | 1.433 | 1.381 | 1.037 | 0.52 | -0.52 | 0.47 | 0.15666667 | -0.495 | 0.05 | 0.184391114 | 0.36362519 | 0.776386308 | 0.876837873 | 0.977313184 | 0.927075529 | 0.992535481 |
| | High | A0A498W | Asparagine--tRNA ligase OS=Mus musculus OX=10090 GN=Nars PE=1 SV=1 | 0 | 13.799 | 10 | 4 | 5 | 4 | 558 | 64.2 | 5.86 | 0.708 | 0.787 | 0.9 | -0.5 | 0.5 | -0.35 | -0.1166667 | 0.425 | -0.15 | 0.194528146 | 0.412284536 | 0.548734701 | 0.876837873 | 0.988319535 | 0.932578704 | 0.96694628 |
| Cops8 | High | Q8VBV7 | COP9 signalosome complex subunit 8 OS=Mus musculus OX=10090 GN=Cops8 PE=1 SV=1 | 0 | 10.069 | 11 | 2 | 2 | 2 | 209 | 23.2 | 5.2 | 0.721 | 0.802 | 0.898 | -0.47 | 0.47 | -0.32 | -0.1066667 | 0.395 | -0.16 | 0.192430921 | 0.421805151 | 0.568492794 | 0.876837873 | 0.99399649 | 0.935417181 | 0.972910028 |
| Tsfm | High | Q9CX33 | Elongation factor Ts, mitochondrial OS=Mus musculus OX=10090 GN=Tsfm PE=1 SV=1 | 0.005 | 3.57 | 5 | 1 | 1 | 1 | 193 | 21.2 | 9.11 | 1.415 | 1.321 | 1.071 | 0.5 | -0.5 | 0.4 | 0.13333333 | -0.45 | 0.1 | 0.192560979 | 0.432342981 | 0.708933938 | 0.876837873 | 0.99399649 | 0.935417181 | 0.992507514 |
| Gtf2i | High | Q3UHU8 | General transcription factor II-I OS=Mus musculus OX=10090 GN=Gtf2i PE=1 SV=1 | 0 | 11.925 | 4 | 3 | 5 | 3 | 962 | 108.3 | 7.8 | 1.435 | 1.329 | 1.08 | 0.52 | -0.52 | 0.41 | 0.13666667 | -0.465 | 0.11 | 0.192705351 | 0.430914785 | 0.654597169 | 0.876837873 | 0.99399649 | 0.935417181 | 0.985010103 |
| Rpl29; Gm | High | P47915 | 60S ribosomal protein L29 OS=Mus musculus OX=10090 GN=Rpl29 PE=1 SV=2 | 0 | 33.234 | 29 | 7 | 18 | 7 | 160 | 17.6 | 11.84 | 1.474 | 1.319 | 1.118 | 0.56 | -0.56 | 0.4 | 0.13333333 | -0.48 | 0.16 | 0.190631673 | 0.445155799 | 0.414425856 | 0.876837873 | 0.994630833 | 0.935734353 | 0.936397673 |
| Rpl17 | High | Q9CPR4 | 60S ribosomal protein L17 OS=Mus musculus OX=10090 GN=Rpl17 PE=1 SV=3 | 0 | 40.029 | 51 | 9 | 16 | 8 | 184 | 21.4 | 10.18 | 1.48 | 1.319 | 1.121 | 0.57 | -0.57 | 0.4 | 0.13333333 | -0.485 | 0.17 | 0.186281497 | 0.44455182 | 0.401582451 | 0.876837873 | 0.994630833 | 0.935734353 | 0.93262326 |
| Tmem9b | High | Q9JJR8 | Transmembrane protein 9B OS=Mus musculus OX=10090 GN=Tmem9b PE=1 SV=1 | 0.001 | 4.404 | 8 | 2 | 2 | 2 | 199 | 22.6 | 8.18 | 0.707 | 0.861 | 0.821 | -0.5 | 0.5 | -0.22 | -0.0733333 | 0.36 | -0.28 | 0.194822953 | 0.563317559 | 0.37410627 | 0.876837873 | 0.995559182 | 0.936198527 | 0.920164724 |
| Picalm | High | Q7M6Y3 | Phosphatidylinositol-binding clathrin assembly protein OS=Mus musculus OX=10090 GN=Picalm P | 0 | 31.737 | 10 | 7 | 12 | 5 | 660 | 71.5 | 7.91 | 0.731 | 0.841 | 0.869 | -0.45 | 0.45 | -0.25 | -0.0833333 | 0.35 | -0.2 | 0.1926348 | 0.538055573 | 0.365881033 | 0.876837873 | 0.995559182 | 0.936198527 | 0.920164724 |
| Rplp0 | High | P14869 | 60S acidic ribosomal protein P0 OS=Mus musculus OX=10090 GN=Rplp0 PE=1 SV=3 | 0 | 67.427 | 44 | 11 | 21 | 11 | 317 | 34.2 | 6.25 | 1.473 | 1.177 | 1.251 | 0.56 | -0.56 | 0.24 | 0.08 | -0.4 | 0.32 | 0.1917901 | 0.678350796 | 0.12295579 | 0.876837873 | 0.995559182 | 0.936198527 | 0.677192798 |
| | High | A0A0G2JF | Intraflagellar transport protein 22 homolog (Fragment) OS=Mus musculus OX=10090 GN=Ift22 PE | 0 | 11.275 | 26 | 3 | 4 | 3 | 155 | 17.5 | 5.74 | 1.395 | 1.255 | 1.112 | 0.48 | -0.48 | 0.33 | 0.11 | -0.405 | 0.15 | 0.186118901 | 0.541459607 | 0.541798881 | 0.876837873 | 0.995559182 | 0.936198527 | 0.96694628 |
| Rps20 | High | P60867 | 40S ribosomal protein S20 OS=Mus musculus OX=10090 GN=Rps20 PE=1 SV=1 | 0 | 26.995 | 53 | 6 | 11 | 6 | 119 | 13.4 | 9.94 | 1.437 | 1.25 | 1.15 | 0.52 | -0.52 | 0.32 | 0.10666667 | -0.42 | 0.2 | 0.186246143 | 0.549262997 | 0.317574257 | 0.876837873 | 0.995559182 | 0.936198527 | 0.904943958 |
| Rpl15 | High | Q9CZM2 | 60S ribosomal protein L15 OS=Mus musculus OX=10090 GN=Rpl15 PE=1 SV=4 | 0 | 43.133 | 40 | 8 | 13 | 8 | 204 | 24.1 | 11.62 | 1.505 | 1.193 | 1.261 | 0.59 | -0.59 | 0.26 | 0.08666667 | -0.425 | 0.33 | 0.187011312 | 0.647903705 | 0.163957607 | 0.876837873 | 0.995559182 | 0.936198527 | 0.736602876 |
| | High | A0A1B0G | 60S ribosomal protein L13a OS=Mus musculus OX=10090 GN=Gm45713 PE=2 SV=1 | 0 | 24.501 | 12 | 7 | 12 | 7 | 446 | 50.8 | 10.23 | 1.451 | 1.266 | 1.146 | 0.54 | -0.54 | 0.34 | 0.11333333 | -0.44 | 0.2 | 0.187551657 | 0.524451683 | 0.326511046 | 0.876837873 | 0.995559182 | 0.936198527 | 0.905686588 |
| 4833439L1 | Medium | D3Z1F7 | Putative monooxygenase p33MONOX OS=Mus musculus OX=10090 GN=4833439L19Rik PE=1 SV= | 0.01 | 2.66 | 3 | 1 | 1 | 1 | 284 | 30.8 | 9.35 | 1.43 | 1.291 | 1.108 | 0.52 | -0.52 | 0.37 | 0.12333333 | -0.445 | 0.15 | 0.18797076 | 0.484268632 | 0.574401714 | 0.876837873 | 0.995559182 | 0.936198527 | 0.972910028 |
| Dnajb1 | High | Q9QYJ3 | DnaJ homolog subfamily B member 1 OS=Mus musculus OX=10090 GN=Dnajb1 PE=1 SV=3 | 0 | 39.99 | 30 | 10 | 12 | 9 | 340 | 38.1 | 8.63 | 1.491 | 1.276 | 1.168 | 0.58 | -0.58 | 0.35 | 0.11666667 | -0.465 | 0.22 | 0.190296673 | 0.50777378 | 0.291670396 | 0.876837873 | 0.995559182 | 0.936198527 | 0.892844069 |
| | High | A0A2I3BP | Ribosomal protein L36A, pseudogene 1 OS=Mus musculus OX=10090 GN=Rpl36a-ps1 PE=3 SV=1 | 0 | 13.826 | 26 | 5 | 8 | 5 | 106 | 12.4 | 10.51 | 1.48 | 1.28 | 1.156 | 0.57 | -0.57 | 0.36 | 0.12 | -0.465 | 0.21 | 0.18623397 | 0.501419361 | 0.301627943 | 0.876837873 | 0.995559182 | 0.936198527 | 0.900241969 |
| Ppp6r1 | High | Q7TSI3 | Serine/threonine-protein phosphatase 6 regulatory subunit 1 OS=Mus musculus OX=10090 GN=P | 0.005 | 3.724 | 1 | 1 | 1 | 1 | 856 | 94.5 | 4.64 | 1.424 | 1.031 | 1.381 | 0.51 | -0.51 | 0.04 | 0.01333333 | -0.275 | 0.47 | 0.191503538 | 0.991184778 | 0.063080376 | 0.876837873 | 0.998402594 | 0.937620233 | 0.48303291 |
| Abcf1 | High | Q6P542 | ATP-binding cassette sub-family F member 1 OS=Mus musculus OX=10090 GN=Abcf1 PE=1 SV=1 | 0 | 20.208 | 8 | 6 | 8 | 6 | 837 | 94.9 | 6.51 | 1.406 | 1.065 | 1.32 | 0.49 | -0.49 | 0.09 | 0.03 | -0.29 | 0.4 | 0.186680271 | 0.91493812 | 0.100996129 | 0.876837873 | 0.998402594 | 0.937620233 | 0.610114517 |
| Pin4; Gm6 | High | Q9CWW6 | Peptidyl-prolyl cis-trans isomerase NIMA-interacting 4 OS=Mus musculus OX=10090 GN=Pin4 PE= | 0 | 42.71 | 67 | 8 | 14 | 8 | 131 | 13.8 | 9.77 | 1.436 | 1.042 | 1.377 | 0.52 | -0.52 | 0.06 | 0.02 | -0.29 | 0.46 | 0.192594345 | 0.967093991 | 0.030458716 | 0.876837873 | 0.998402594 | 0.937620233 | 0.334866709 |

| | | | | | | | | | | | | | | | | | | | | | | | | | | | | |
|---|---|---|---|---|---|---|---|---|---|---|---|---|---|---|---|---|---|---|---|---|---|---|---|---|---|---|---|---|
| Akr1e1 | High | Q9DCT1 | 1,5-anhydro-D-fructose reductase OS=Mus musculus OX=10090 GN=Akr1e2 PE=1 SV=1 | 0 | 15.256 | 11 | 4 | 7 | 4 | 301 | 34.4 | 7.33 | 1.485 | 1.035 | 1.435 | 0.57 | -0.57 | 0.05 | 0.01666667 | -0.31 | 0.52 | 0.186049117 | 0.985167154 | 0.019980669 | 0.876837873 | 0.998402594 | 0.937620233 | 0.252647811 |
| Agfg2 | High | Q3TBU7 | Arf-GAP domain and FG repeat-containing protein 2 (Fragment) OS=Mus musculus OX=10090 GN | 0 | 11.703 | 8 | 3 | 5 | 2 | 406 | 41.7 | 8.98 | 0.724 | 0.753 | 0.962 | -0.47 | 0.47 | -0.41 | -0.1366667 | 0.44 | -0.06 | 0.196733045 | 0.331611819 | 0.823258346 | 0.878949947 | 0.968621131 | 0.923785539 | 0.992535481 |
| Kifap3 | High | P70188 | Kinesin-associated protein 3 OS=Mus musculus OX=10090 GN=Kifap3 PE=1 SV=1 | 0 | 18.421 | 8 | 5 | 7 | 5 | 793 | 91.2 | 5.11 | 1.414 | 1.001 | 1.412 | 0.5 | -0.5 | 0 | 0 | -0.25 | 0.5 | 0.196307724 | 0.934969318 | 0.040576146 | 0.878949947 | 0.998402594 | 0.93867627 | 0.388243087 |
| Npl | High | Q9DCJ9 | N-acetylneuraminate lyase OS=Mus musculus OX=10090 GN=Npl PE=1 SV=1 | 0 | 9.572 | 12 | 3 | 4 | 3 | 320 | 35.1 | 8.03 | 0.701 | 0.541 | 1.297 | -0.51 | 0.51 | -0.89 | -0.2966667 | 0.7 | 0.38 | 0.197307638 | 0.048517169 | 0.126102092 | 0.879945382 | 0.523145709 | 0.701545546 | 0.678506823 |
| | High | D3YTS3 | RIKEN cDNA D630045J12 gene OS=Mus musculus OX=10090 GN=D630045J12Rik PE=1 SV=2 | 0 | 56.775 | 12 | 16 | 19 | 16 | 1799 | 193.9 | 6.13 | 1.468 | 1.774 | 0.828 | 0.55 | -0.55 | 0.83 | 0.27666667 | -0.69 | -0.27 | 0.199125616 | 0.094381485 | 0.263207964 | 0.879945382 | 0.70644407 | 0.793194726 | 0.859470827 |
| Samm50 | High | Q8BGH2 | Sorting and assembly machinery component 50 homolog OS=Mus musculus OX=10090 GN=Samm | 0 | 23.648 | 17 | 5 | 7 | 5 | 469 | 51.8 | 6.8 | 1.456 | 1.591 | 0.915 | 0.54 | -0.54 | 0.67 | 0.22333333 | -0.605 | -0.13 | 0.200342488 | 0.180715113 | 0.626892805 | 0.879945382 | 0.885725209 | 0.882835296 | 0.980381496 |
| Tsta3 | High | P23591 | GDP-L-fucose synthase OS=Mus musculus OX=10090 GN=Gfus PE=1 SV=3 | 0 | 9.7 | 9 | 2 | 3 | 2 | 321 | 35.9 | 6.74 | 0.758 | 0.733 | 1.034 | -0.4 | 0.4 | -0.45 | -0.15 | 0.425 | 0.05 | 0.199232407 | 0.288385761 | 0.77341461 | 0.879945382 | 0.955820031 | 0.917882706 | 0.992535481 |
| | Medium | D3Z783 | Non-specific serine/threonine protein kinase OS=Mus musculus OX=10090 GN=Akt1 PE=1 SV=2 | 0.019 | 2.064 | 2 | 1 | 1 | 1 | 437 | 50.9 | 6.13 | 1.399 | 1.44 | 0.972 | 0.48 | -0.48 | 0.53 | 0.17666667 | -0.505 | -0.04 | 0.199814869 | 0.301772883 | 0.938253927 | 0.879945382 | 0.955820031 | 0.917882706 | 0.992535481 |
| Hnrnpr | High | Q8VHM5 | Heterogeneous nuclear ribonucleoprotein R OS=Mus musculus OX=10090 GN=Hnrnpr PE=1 SV=1 | 0 | 18.899 | 12 | 7 | 8 | 5 | 632 | 70.8 | 8.13 | 1.409 | 1.404 | 1.004 | 0.5 | -0.5 | 0.49 | 0.16333333 | -0.495 | 0.01 | 0.200213317 | 0.339930025 | 0.97206406 | 0.879945382 | 0.976045759 | 0.92799557 | 0.996575243 |
| Flvcr1; Mf | High | B2RXV4 | Feline leukemia virus subgroup C receptor-related protein 1 OS=Mus musculus OX=10090 GN=Flv | 0.001 | 4.579 | 6 | 2 | 2 | 2 | 560 | 60.4 | 5.68 | 0.714 | 0.761 | 0.939 | -0.49 | 0.49 | -0.39 | -0.13 | 0.44 | -0.09 | 0.199461469 | 0.350192002 | 0.757519562 | 0.879945382 | 0.977313184 | 0.928629283 | 0.992535481 |
| Ubtf | High | A2AWT7 | Nucleolar transcription factor 1 OS=Mus musculus OX=10090 GN=Ubtf PE=1 SV=1 | 0.001 | 6.524 | 2 | 2 | 2 | 2 | 752 | 88.3 | 9.89 | 1.439 | 1.357 | 1.061 | 0.53 | -0.53 | 0.44 | 0.14666667 | -0.485 | 0.09 | 0.200224855 | 0.393329136 | 0.664451753 | 0.879945382 | 0.983884835 | 0.931915109 | 0.986647506 |
| Apoo | High | Q9DCZ4 | MICOS complex subunit Mic26 OS=Mus musculus OX=10090 GN=Apoo PE=1 SV=2 | 0 | 21.43 | 21 | 4 | 7 | 4 | 198 | 22.6 | 9.25 | 1.457 | 1.338 | 1.089 | 0.54 | -0.54 | 0.42 | 0.14 | -0.48 | 0.12 | 0.197631558 | 0.419840571 | 0.623827566 | 0.879945382 | 0.99399649 | 0.936970936 | 0.97840592 |
| Crb1 | High | Q8VHS2 | Protein crumbs homolog 1 OS=Mus musculus OX=10090 GN=Crb1 PE=1 SV=3 | 0 | 48.684 | 7 | 8 | 14 | 8 | 1405 | 153.2 | 5.1 | 0.719 | 1.049 | 0.685 | -0.48 | 0.48 | 0.07 | 0.02333333 | 0.205 | -0.54 | 0.200408463 | 0.951738207 | 0.020096473 | 0.879945382 | 0.998402594 | 0.939173988 | 0.252647811 |
| Agpat1 | High | O35083 | 1-acyl-sn-glycerol-3-phosphate acyltransferase alpha OS=Mus musculus OX=10090 GN=Agpat1 PE | 0.001 | 4.481 | 5 | 1 | 1 | 1 | 285 | 31.7 | 9.14 | 1.458 | 1.047 | 1.392 | 0.54 | -0.54 | 0.07 | 0.02333333 | -0.305 | 0.48 | 0.198181277 | 0.980866711 | 0.072045587 | 0.879945382 | 0.998402594 | 0.939173988 | 0.511341275 |
| | High | E9Q0U7 | Heat shock protein 105 kDa OS=Mus musculus OX=10090 GN=Hsph1 PE=1 SV=1 | 0 | 26.256 | 6 | 5 | 8 | 3 | 817 | 91.6 | 5.59 | 0.696 | 0.727 | 0.957 | -0.52 | 0.52 | -0.46 | -0.1533333 | 0.49 | -0.06 | 0.201032357 | 0.288352816 | 0.825691897 | 0.881304477 | 0.955820031 | 0.918562254 | 0.992535481 |
| Tprgl | High | Q9DBS2 | Tumor protein p63-regulated gene 1-like protein OS=Mus musculus OX=10090 GN=Tprg1l PE=1 S\ | 0 | 43.644 | 42 | 8 | 13 | 8 | 266 | 29.8 | 7.37 | 0.681 | 0.596 | 1.142 | -0.55 | 0.55 | -0.75 | -0.25 | 0.65 | 0.19 | 0.20173122 | 0.094494926 | 0.363883952 | 0.883677307 | 0.70644407 | 0.795060688 | 0.91781256 |
| | High | H3BKG0 | Caveolin-1 OS=Mus musculus OX=10090 GN=Cav1 PE=1 SV=1 | 0.001 | 5.051 | 19 | 2 | 2 | 2 | 93 | 10.4 | 6.05 | 0.693 | 0.708 | 0.978 | -0.53 | 0.53 | -0.5 | -0.1666667 | 0.515 | -0.03 | 0.203029102 | 0.253131706 | 0.943131596 | 0.885213201 | 0.950034749 | 0.917623975 | 0.992535481 |
| Glrx3 | High | Q9CQM9 | Glutaredoxin-3 OS=Mus musculus OX=10090 GN=Glrx3 PE=1 SV=1 | 0.001 | 7.801 | 10 | 3 | 3 | 3 | 337 | 37.8 | 5.59 | 0.719 | 0.742 | 0.97 | -0.48 | 0.48 | -0.43 | -0.1433333 | 0.455 | -0.04 | 0.202457466 | 0.315813405 | 0.883082505 | 0.885213201 | 0.955820031 | 0.920516616 | 0.992535481 |
| | High | A0A3Q4E( | Tetratricopeptide repeat protein 39C OS=Mus musculus OX=10090 GN=Ttc39c PE=1 SV=1 | 0 | 9.383 | 7 | 4 | 4 | 4 | 580 | 65.4 | 6.87 | 1.396 | 1.341 | 1.042 | 0.48 | -0.48 | 0.42 | 0.14 | -0.45 | 0.06 | 0.203382682 | 0.4084243 | 0.748482201 | 0.885377869 | 0.9855335 | 0.935455685 | 0.992535481 |
| Diablo | High | Q9JIQ3 | Diablo homolog, mitochondrial OS=Mus musculus OX=10090 GN=Diablo PE=1 SV=2 | 0.009 | 2.786 | 3 | 1 | 1 | 1 | 237 | 26.8 | 6.37 | 1.408 | 1.809 | 0.779 | 0.49 | -0.49 | 0.85 | 0.28333333 | -0.67 | -0.36 | 0.20603015 | 0.078275781 | 0.12719165 | 0.892550179 | 0.65506314 | 0.77380666 | 0.678506823 |
| Psmd14 | High | O35593 | 26S proteasome non-ATPase regulatory subunit 14 OS=Mus musculus OX=10090 GN=Psmd14 PE= | 0 | 16.265 | 16 | 5 | 8 | 5 | 310 | 34.6 | 6.52 | 0.725 | 0.617 | 1.176 | -0.46 | 0.46 | -0.7 | -0.2333333 | 0.58 | 0.23 | 0.207061529 | 0.117233761 | 0.356509399 | 0.892550179 | 0.779640783 | 0.836095481 | 0.914600864 |
| Prrt1 | High | O35449 | Proline-rich transmembrane protein 1 OS=Mus musculus OX=10090 GN=Prrt1 PE=1 SV=1 | 0 | 13.322 | 5 | 1 | 2 | 1 | 306 | 31.4 | 7.65 | 0.736 | 0.65 | 1.132 | -0.44 | 0.44 | -0.62 | -0.2066667 | 0.53 | 0.18 | 0.207167166 | 0.144516962 | 0.461422977 | 0.892550179 | 0.838826712 | 0.865688446 | 0.959744396 |
| Rab23 | High | Q9D4I9 | Ras-related protein Rab-23 OS=Mus musculus OX=10090 GN=Rab23 PE=1 SV=1 | 0 | 12.647 | 24 | 4 | 4 | 4 | 237 | 26.8 | 7.3 | 0.745 | 0.785 | 0.948 | -0.43 | 0.43 | -0.35 | -0.1166667 | 0.39 | -0.08 | 0.207258843 | 0.408523183 | 0.784247389 | 0.892550179 | 0.9855335 | 0.93904184 | 0.992535481 |
| Mlc1 | Medium | Q8VHK5 | Membrane protein MLC1 OS=Mus musculus OX=10090 GN=Mlc1 PE=1 SV=1 | 0.047 | 1.568 | 3 | 1 | 1 | 1 | 382 | 41.6 | 7.9 | 0.726 | 0.796 | 0.912 | -0.46 | 0.46 | -0.33 | -0.11 | 0.395 | -0.13 | 0.20695978 | 0.432613732 | 0.615739853 | 0.892550179 | 0.99399649 | 0.943273335 | 0.97826833 |
| Dek | High | Q7TNV0 | Protein DEK OS=Mus musculus OX=10090 GN=Dek PE=1 SV=1 | 0.001 | 7.013 | 4 | 2 | 2 | 2 | 380 | 43.1 | 6.86 | 1.399 | 1.22 | 1.146 | 0.48 | -0.48 | 0.29 | 0.09666667 | -0.385 | 0.2 | 0.205889119 | 0.599554996 | 0.433814499 | 0.892550179 | 0.995559182 | 0.944054681 | 0.948052758 |
| | High | G3UY93 | Valyl-tRNA synthetase (Fragment) OS=Mus musculus OX=10090 GN=Vars PE=1 SV=1 | 0 | 12.673 | 3 | 4 | 6 | 4 | 1278 | 141.3 | 7.75 | 1.463 | 1.303 | 1.123 | 0.55 | -0.55 | 0.38 | 0.12666667 | -0.465 | 0.17 | 0.206804939 | 0.467314884 | 0.449128566 | 0.892550179 | 0.995559182 | 0.944054681 | 0.958608248 |
| Ryr2 | High | E9Q401 | Ryanodine receptor 2 OS=Mus musculus OX=10090 GN=Ryr2 PE=1 SV=1 | 0.001 | 5.432 | 1 | 2 | 3 | 2 | 4966 | 564.5 | 6.09 | 0.736 | 1.091 | 0.675 | -0.44 | 0.44 | 0.13 | 0.04333333 | 0.155 | -0.57 | 0.206771984 | 0.861988874 | 0.027738826 | 0.892550179 | 0.998402594 | 0.945476387 | 0.315745518 |
| Hist1h1d | High | P43277 | Histone H1.3 OS=Mus musculus OX=10090 GN=H1-3 PE=1 SV=2 | 0 | 78.792 | 44 | 17 | 39 | 1 | 221 | 22.1 | 11.03 | 1.479 | 1.85 | 0.799 | 0.56 | -0.56 | 0.89 | 0.29666667 | -0.725 | -0.32 | 0.207469745 | 0.071605729 | 0.179515223 | 0.892772725 | 0.625156241 | 0.758964483 | 0.756556316 |
| Pycr2 | Medium | Q922Q4 | Pyrroline-5-carboxylate reductase 2 OS=Mus musculus OX=10090 GN=Pycr2 PE=1 SV=1 | 0.044 | 1.593 | 2 | 1 | 1 | 1 | 320 | 33.6 | 7.77 | 1.41 | 1.409 | 1.001 | 0.5 | -0.5 | 0.49 | 0.16333333 | -0.495 | 0 | 0.208438192 | 0.319439338 | 0.962513449 | 0.896252257 | 0.955820031 | 0.926036144 | 0.993960013 |
| AI314180 | Medium | Q6PDI5 | Proteasome adapter and scaffold protein ECM29 OS=Mus musculus OX=10090 GN=Ecpas PE=1 S\ | 0.042 | 1.649 | 1 | 1 | 1 | 1 | 1840 | 203.6 | 7.06 | 0.715 | 0.564 | 1.268 | -0.48 | 0.48 | -0.83 | -0.2766667 | 0.655 | 0.34 | 0.209224844 | 0.058750278 | 0.246029239 | 0.898861914 | 0.571608376 | 0.735235145 | 0.840125423 |
| Uqcrc1 | High | Q9CZ13 | Cytochrome b-c1 complex subunit 1, mitochondrial OS=Mus musculus OX=10090 GN=Uqcrc1 PE= | 0 | 88.992 | 41 | 15 | 27 | 15 | 480 | 52.8 | 6.21 | 1.415 | 1.551 | 0.913 | 0.5 | -0.5 | 0.63 | 0.21 | -0.565 | -0.13 | 0.209365732 | 0.207717309 | 0.625002368 | 0.898861914 | 0.912882441 | 0.905872177 | 0.979106043 |
| Fabp12 | High | Q9DAK4 | Fatty acid-binding protein 12 OS=Mus musculus OX=10090 GN=Fabp12 PE=2 SV=1 | 0.001 | 7.318 | 19 | 2 | 3 | 2 | 132 | 14.7 | 8.15 | 1.432 | 1.474 | 0.971 | 0.52 | -0.52 | 0.56 | 0.18666667 | -0.54 | -0.04 | 0.209826978 | 0.264938977 | 0.895213015 | 0.899464731 | 0.951072422 | 0.925268577 | 0.992535481 |
| Rpl23a | High | P62751 | 60S ribosomal protein L23a OS=Mus musculus OX=10090 GN=Rpl23a PE=1 SV=1 | 0 | 42.72 | 57 | 12 | 21 | 12 | 156 | 17.7 | 10.45 | 1.45 | 1.276 | 1.136 | 0.54 | -0.54 | 0.35 | 0.11666667 | -0.445 | 0.18 | 0.209757108 | 0.507385535 | 0.355922187 | 0.899464731 | 0.995559182 | 0.947511956 | 0.914600864 |
| Ppp2r2d | High | Q925E7 | Serine/threonine-protein phosphatase 2A 55 kDa regulatory subunit B delta isoform OS=Mus mus | 0 | 14.569 | 11 | 4 | 5 | 2 | 453 | 51.9 | 6.49 | 0.761 | 0.811 | 0.938 | -0.39 | 0.39 | -0.3 | -0.1 | 0.345 | -0.09 | 0.210117175 | 0.433283564 | 0.753822585 | 0.900020626 | 0.99399649 | 0.947008558 | 0.992535481 |
| COX2 | High | P00405 | Cytochrome c oxidase subunit 2 OS=Mus musculus OX=10090 GN=Mtco2 PE=1 SV=1 | 0 | 25.167 | 21 | 4 | 10 | 4 | 227 | 26 | 4.73 | 1.46 | 1.472 | 0.992 | 0.55 | -0.55 | 0.56 | 0.18666667 | -0.555 | -0.01 | 0.210531208 | 0.271257451 | 0.942101017 | 0.900489157 | 0.955364652 | 0.927926905 | 0.992535481 |
| Mcat | Medium | Q8R3F5 | Malonyl-CoA-acyl carrier protein transacylase, mitochondrial OS=Mus musculus OX=10090 GN=M | 0.012 | 2.375 | 2 | 1 | 1 | 1 | 381 | 41.9 | 8.1 | 1.431 | 1.429 | 1.001 | 0.52 | -0.52 | 0.52 | 0.17333333 | -0.52 | 0 | 0.211001078 | 0.323720749 | 0.9314096 | 0.900489157 | 0.955820031 | 0.928154594 | 0.992535481 |
| | High | D3Z5B1 | Zinc finger, BED type-containing 5 (Fragment) OS=Mus musculus OX=10090 GN=Zbed5 PE=4 SV=8 | 0 | 17.853 | 26 | 3 | 3 | 3 | 183 | 19.2 | 8.28 | 1.386 | 1.189 | 1.166 | 0.47 | -0.47 | 0.25 | 0.08333333 | -0.36 | 0.22 | 0.211029562 | 0.661405951 | 0.324420218 | 0.900489157 | 0.995559182 | 0.948024169 | 0.905686588 |
| Smpd3 | High | Q9JJY3 | Sphingomyelin phosphodiesterase 3 OS=Mus musculus OX=10090 GN=Smpd3 PE=1 SV=1 | 0.001 | 6 | 3 | 2 | 2 | 2 | 655 | 71.2 | 5.88 | 0.755 | 0.786 | 0.959 | -0.41 | 0.41 | -0.35 | -0.1166667 | 0.38 | -0.06 | 0.21165071 | 0.384313345 | 0.835026916 | 0.90243685 | 0.977313184 | 0.939875017 | 0.992535481 |
| Pdxp | High | P60487 | Chronophin OS=Mus musculus OX=10090 GN=Pdxp PE=1 SV=1 | 0 | 12.958 | 14 | 3 | 3 | 3 | 292 | 31.5 | 5.74 | 0.761 | 0.775 | 0.982 | -0.39 | 0.39 | -0.37 | -0.1233333 | 0.38 | -0.03 | 0.2118079 | 0.353686609 | 0.985317837 | 0.90243685 | 0.977313184 | 0.939875017 | 0.998604519 |
| Cabp4 | High | Q8VHC5 | Calcium-binding protein 4 OS=Mus musculus OX=10090 GN=Cabp4 PE=1 SV=1 | 0 | 16.533 | 21 | 3 | 4 | 3 | 271 | 30.2 | 5.29 | 1.409 | 0.927 | 1.52 | 0.49 | -0.49 | -0.11 | -0.0366667 | -0.19 | 0.6 | 0.21202108 | 0.741228271 | 0.017348818 | 0.90265922 | 0.995559182 | 0.949109201 | 0.231606717 |
| Rpl39 | Medium | P62892 | 60S ribosomal protein L39 OS=Mus musculus OX=10090 GN=Rpl39 PE=1 SV=2 | 0.026 | 1.968 | 20 | 1 | 1 | 1 | 51 | 6.4 | 12.56 | 1.443 | 1.022 | 1.411 | 0.53 | -0.53 | 0.03 | 0.01 | -0.28 | 0.5 | 0.212624746 | 0.961865264 | 0.060772436 | 0.904542453 | 0.998402594 | 0.951472523 | 0.476099813 |
| Cyc1 | High | Q9D0M3 | Cytochrome c1, heme protein, mitochondrial OS=Mus musculus OX=10090 GN=Cyc1 PE=1 SV=1 | 0 | 28.293 | 25 | 6 | 13 | 6 | 325 | 35.3 | 9.16 | 1.427 | 1.54 | 0.927 | 0.51 | -0.51 | 0.62 | 0.20666667 | -0.565 | -0.11 | 0.212804675 | 0.215208353 | 0.635439767 | 0.904621542 | 0.920170644 | 0.912396093 | 0.983562471 |
| Bag5 | High | Q8CI32 | BAG family molecular chaperone regulator 5 OS=Mus musculus OX=10090 GN=Bag5 PE=1 SV=1 | 0 | 37.76 | 29 | 9 | 12 | 9 | 447 | 50.9 | 6.05 | 1.46 | 1.453 | 1.004 | 0.55 | -0.55 | 0.54 | 0.18 | -0.545 | 0.01 | 0.213050563 | 0.288890872 | 0.993114098 | 0.904980687 | 0.955820031 | 0.930400359 | 0.999917596 |
| Rpl27; LO( | High | P61358 | 60S ribosomal protein L27 OS=Mus musculus OX=10090 GN=Rpl27 PE=1 SV=2 | 0 | 26.731 | 49 | 6 | 14 | 6 | 136 | 15.8 | 10.56 | 1.412 | 1.295 | 1.09 | 0.5 | -0.5 | 0.37 | 0.12333333 | -0.435 | 0.12 | 0.213771 | 0.479962438 | 0.510448153 | 0.90735352 | 0.995559182 | 0.951456351 | 0.964378027 |
| Optn | High | Q8K3K8 | Optineurin OS=Mus musculus OX=10090 GN=Optn PE=1 SV=1 | 0 | 172.448 | 52 | 32 | 74 | 32 | 584 | 67 | 5.26 | 0.693 | 0.526 | 1.318 | -0.53 | 0.53 | -0.93 | -0.31 | 0.73 | 0.4 | 0.217845487 | 0.039535687 | 0.060262571 | 0.908812298 | 0.493711801 | 0.701262049 | 0.474986455 |
| Dennd5a | Medium | Q6PAL8 | DENN domain-containing protein 5A OS=Mus musculus OX=10090 GN=Dennd5a PE=1 SV=1 | 0.038 | 1.721 | 1 | 1 | 1 | 1 | 1287 | 146.6 | 6.7 | 1.396 | 1.799 | 0.776 | 0.48 | -0.48 | 0.85 | 0.28333333 | -0.665 | -0.37 | 0.215568153 | 0.086228135 | 0.140988448 | 0.908812298 | 0.68454712 | 0.796679709 | 0.701851981 |
| Csnk1g3 | High | Q8C4X2 | Casein kinase I isoform gamma-3 OS=Mus musculus OX=10090 GN=Csnk1g3 PE=1 SV=2 | 0 | 50.496 | 27 | 12 | 19 | 6 | 424 | 48.9 | 9.11 | 1.441 | 1.732 | 0.832 | 0.53 | -0.53 | 0.79 | 0.26333333 | -0.66 | -0.27 | 0.21758275 | 0.109887134 | 0.278461051 | 0.908812298 | 0.768225125 | 0.838518711 | 0.876067403 |
| Olfr703 | Medium | Q9EPF5 | Olfactory receptor 703 OS=Mus musculus OX=10090 GN=Olfr703 PE=4 SV=1 | 0.036 | 1.753 | 2 | 1 | 2 | 1 | 319 | 35.5 | 7.9 | 1.38 | 1.714 | 0.805 | 0.46 | -0.46 | 0.78 | 0.26 | -0.62 | -0.31 | 0.218868503 | 0.114710751 | 0.197113341 | 0.908812298 | 0.77945068 | 0.844131489 | 0.784953482 |
| | High | A0A1B0GS | Predicted gene 45808 (Fragment) OS=Mus musculus OX=10090 GN=Gm45808 PE=4 SV=1 | 0 | 24.182 | 19 | 4 | 5 | 2 | 263 | 28.8 | 5.68 | 1.397 | 1.696 | 0.824 | 0.48 | -0.48 | 0.76 | 0.25333333 | -0.62 | -0.28 | 0.219544076 | 0.124805911 | 0.276908765 | 0.908812298 | 0.799286338 | 0.854049318 | 0.876067403 |
| | High | D3Z440 | COP9 signalosome complex subunit 7a (Fragment) OS=Mus musculus OX=10090 GN=Cops7a PE= | 0.001 | 7.896 | 9 | 2 | 2 | 2 | 226 | 25 | 5.68 | 0.741 | 0.65 | 1.14 | -0.43 | 0.43 | -0.62 | -0.2066667 | 0.525 | 0.19 | 0.216187633 | 0.145159379 | 0.447695163 | 0.908812298 | 0.840814708 | 0.874813503 | 0.958608248 |
| Tubb3 | High | Q9ERD7 | Tubulin beta-3 chain OS=Mus musculus OX=10090 GN=Tubb3 PE=1 SV=1 | 0 | 242.152 | 55 | 19 | 118 | 5 | 450 | 50.4 | 4.93 | 1.346 | 1.617 | 0.832 | 0.43 | -0.43 | 0.69 | 0.23 | -0.56 | -0.27 | 0.215236991 | 0.164813512 | 0.271151286 | 0.908812298 | 0.871008827 | 0.889910562 | 0.871850649 |
| Lrrc40 | High | A0A0R4J0 | Leucine-rich repeat-containing protein 40 OS=Mus musculus OX=10090 GN=Lrrc40 PE=1 SV=1 | 0.005 | 3.492 | 3 | 1 | 1 | 1 | 602 | 68 | 6.84 | 0.75 | 0.694 | 1.08 | -0.42 | 0.42 | -0.53 | -0.1766667 | 0.475 | 0.11 | 0.215256231 | 0.208297662 | 0.626693843 | 0.908812298 | 0.912882441 | 0.910847369 | 0.980381496 |
| Pfn1 | High | P62962 | Profilin-1 OS=Mus musculus OX=10090 GN=Pfn1 PE=1 SV=2 | 0 | 35.015 | 54 | 7 | 9 | 7 | 140 | 14.9 | 8.28 | 0.71 | 0.72 | 0.986 | -0.49 | 0.49 | -0.47 | -0.1566667 | 0.48 | -0.02 | 0.215809298 | 0.27359435 | 0.920123254 | 0.908812298 | 0.955820031 | 0.932316164 | 0.992535481 |
| Tubb5 | High | P99024 | Tubulin beta-5 chain OS=Mus musculus OX=10090 GN=Tubb5 PE=1 SV=1 | 0 | 426.941 | 74 | 27 | 217 | 6 | 444 | 49.6 | 4.89 | 1.444 | 1.438 | 1.004 | 0.53 | -0.53 | 0.52 | 0.17333333 | -0.525 | 0.01 | 0.214890541 | 0.303855622 | 0.897504594 | 0.908812298 | 0.955820031 | 0.932316164 | 0.992535481 |
| Ppm1n | High | Q8BGL1 | Probable protein phosphatase 1N OS=Mus musculus OX=10090 GN=Ppm1n PE=2 SV=1 | 0 | 54.341 | 31 | 8 | 15 | 8 | 404 | 43.2 | 6.07 | 1.454 | 1.388 | 1.048 | 0.54 | -0.54 | 0.47 | 0.15666667 | -0.505 | 0.07 | 0.21782692 | 0.358184402 | 0.763119956 | 0.908812298 | 0.977313184 | 0.943062741 | 0.992535481 |
| Got2 | High | P05202 | Aspartate aminotransferase, mitochondrial OS=Mus musculus OX=10090 GN=Got2 PE=1 SV=1 | 0 | 138.01 | 63 | 25 | 49 | 25 | 430 | 47.4 | 9 | 1.44 | 1.355 | 1.062 | 0.53 | -0.53 | 0.44 | 0.14666667 | -0.485 | 0.09 | 0.219090435 | 0.397232654 | 0.624956885 | 0.908812298 | 0.985440071 | 0.947126184 | 0.979106043 |
| Adrm1 | High | Q9JKV1 | Proteasomal ubiquitin receptor ADRM1 OS=Mus musculus OX=10090 GN=Adrm1 PE=1 SV=2 | 0 | 10.392 | 10 | 3 | 6 | 3 | 407 | 42 | 5.07 | 0.758 | 0.799 | 0.948 | -0.4 | 0.4 | -0.32 | -0.1066667 | 0.36 | -0.08 | 0.216386275 | 0.40914843 | 0.785002118 | 0.908812298 | 0.9855335 | 0.947172899 | 0.992535481 |
| Pa2g4 | High | P50580 | Proliferation-associated protein 2G4 OS=Mus musculus OX=10090 GN=Pa2g4 PE=1 SV=3 | 0 | 30.057 | 21 | 7 | 9 | 7 | 394 | 43.7 | 6.86 | 1.487 | 1.145 | 1.298 | 0.57 | -0.57 | 0.2 | 0.06666667 | -0.385 | 0.38 | 0.215884571 | 0.740993312 | 0.1171591 | 0.908812298 | 0.995559182 | 0.95218574 | 0.654114904 |
| Tpgs1 | Medium | Q99MS8 | Tubulin polyglutamylase complex subunit 1 OS=Mus musculus OX=10090 GN=Tpgs1 PE=1 SV=1 | 0.015 | 2.218 | 3 | 1 | 1 | 1 | 303 | 32.6 | 8.94 | 1.377 | 1.286 | 1.071 | 0.46 | -0.46 | 0.36 | 0.12 | -0.41 | 0.1 | 0.219223514 | 0.478583939 | 0.676159321 | 0.908812298 | 0.995559182 | 0.95218574 | 0.990250803 |
| Thoc5 | High | Q5SVF9 | THO complex subunit 5 homolog OS=Mus musculus OX=10090 GN=Thoc5 PE=1 SV=1 | 0.001 | 5.956 | 2 | 1 | 1 | 1 | 635 | 73 | 7.59 | 1.387 | 1.294 | 1.072 | 0.47 | -0.47 | 0.37 | 0.12333333 | -0.42 | 0.1 | 0.219625586 | 0.464680879 | 0.683365574 | 0.908812298 | 0.995559182 | 0.95218574 | 0.991285312 |
| Emc10 | Medium | A0A0X1KG | ER membrane protein complex subunit 10 OS=Mus musculus OX=10090 GN=Emc10 PE=1 SV=1 | 0.01 | 2.614 | 3 | 1 | 1 | 1 | 268 | 28.2 | 6.55 | 0.746 | 1.098 | 0.68 | -0.42 | 0.42 | 0.13 | 0.04333333 | 0.145 | -0.56 | 0.219311865 | 0.843891193 | 0.043222673 | 0.908812298 | 0.998402594 | 0.953607446 | 0.404433694 |
| Rtca; Rtcd | High | Q9D7H3 | RNA 3'-terminal phosphate cyclase OS=Mus musculus OX=10090 GN=RtcA PE=1 SV=2 | 0 | 25.326 | 22 | 5 | 9 | 5 | 366 | 39.2 | 7.9 | 1.453 | 1.041 | 1.396 | 0.54 | -0.54 | 0.06 | 0.02 | -0.3 | 0.48 | 0.215663438 | 0.970873257 | 0.038089597 | 0.908812298 | 0.998402594 | 0.953607446 | 0.373579725 |
| C1qbp | High | Q8R5L1 | Complement component 1 Q subcomponent-binding protein, mitochondrial OS=Mus musculus OX | 0 | 10.325 | 12 | 2 | 3 | 2 | 279 | 31 | 4.87 | 1.393 | 1.719 | 0.811 | 0.48 | -0.48 | 0.78 | 0.26 | -0.63 | -0.3 | 0.219999751 | 0.108692104 | 0.212028782 | 0.908939654 | 0.764663273 | 0.836801464 | 0.804011919 |
| Flna | High | B7FAV1 | Filamin, alpha (Fragment) OS=Mus musculus OX=10090 GN=Flna PE=1 SV=1 | 0 | 27.461 | 5 | 10 | 10 | 9 | 2583 | 274.5 | 5.97 | 0.712 | 0.699 | 1.018 | -0.49 | 0.49 | -0.52 | -0.1733333 | 0.505 | 0.03 | 0.220608023 | 0.236193949 | 0.938987554 | 0.908939654 | 0.937374952 | 0.923157303 | 0.992535481 |
| Tmem62 | High | Q8BXJ9 | Transmembrane protein 62 OS=Mus musculus OX=10090 GN=Tmem62 PE=2 SV=1 | 0 | 22.452 | 14 | 6 | 7 | 6 | 643 | 72.8 | 8.78 | 0.727 | 0.745 | 0.976 | -0.46 | 0.46 | -0.42 | -0.14 | 0.44 | -0.03 | 0.220145935 | 0.322830027 | 0.83691195 | 0.908939654 | 0.955820031 | 0.932379842 | 0.992535481 |
| Rab28 | High | Q99KL7 | Ras-related protein Rab-28 OS=Mus musculus OX=10090 GN=Rab28 PE=1 SV=1 | 0 | 106.705 | 62 | 11 | 38 | 1 | 221 | 24.7 | 5.67 | 1.392 | 1.413 | 0.985 | 0.48 | -0.48 | 0.5 | 0.16666667 | -0.49 | -0.02 | 0.22079112 | 0.32259991 | 0.973202953 | 0.908939654 | 0.955820031 | 0.932379842 | 0.996848549 |
| Heatr6 | Medium | Q6P1G0 | HEAT repeat-containing protein 6 OS=Mus musculus OX=10090 GN=Heatr6 PE=1 SV=1 | 0.019 | 2.063 | 1 | 1 | 1 | 1 | 1184 | 128.8 | 7.24 | 0.752 | 0.805 | 0.933 | -0.41 | 0.41 | -0.31 | -0.1033333 | 0.36 | -0.1 | 0.220786851 | 0.436617745 | 0.764010019 | 0.908939654 | 0.994630833 | 0.951785244 | 0.992535481 |
| Huwe1 | High | Q7TMY8 | E3 ubiquitin-protein ligase HUWE1 OS=Mus musculus OX=10090 GN=Huwe1 PE=1 SV=5 | 0 | 45.101 | 4 | 15 | 16 | 15 | 4377 | 482.3 | 5.22 | 0.688 | 0.643 | 1.07 | -0.54 | 0.54 | -0.64 | -0.2133333 | 0.59 | 0.1 | 0.222238821 | 0.149379177 | 0.591322003 | 0.909433078 | 0.847684847 | 0.878558963 | 0.975147842 |
| Lman2l | High | P59481 | VIP36-like protein OS=Mus musculus OX=10090 GN=Lman2l PE=1 SV=1 | 0.001 | 6.576 | 6 | 2 | 2 | 2 | 347 | 39.9 | 9.41 | 1.415 | 1.515 | 0.935 | 0.5 | -0.5 | 0.6 | 0.2 | -0.55 | -0.1 | 0.22366831 | 0.235080495 | 0.728735442 | 0.909433078 | 0.936813316 | 0.923123197 | 0.992535481 |
| Cdh2 | High | P15116 | Cadherin-2 OS=Mus musculus OX=10090 GN=Cdh2 PE=1 SV=2 | 0 | 29.146 | 14 | 5 | 6 | 5 | 906 | 99.7 | 4.78 | 0.694 | 0.715 | 0.971 | -0.53 | 0.53 | -0.48 | -0.16 | 0.505 | -0.04 | 0.22173681 | 0.264678742 | 0.909767337 | 0.909433078 | 0.951072422 | 0.93025275 | 0.992535481 |
| Pcm1 | High | Q9R0L6 | Pericentriolar material 1 protein OS=Mus musculus OX=10090 GN=Pcm1 PE=1 SV=2 | 0 | 21.657 | 3 | 7 | 7 | 7 | 2025 | 228.7 | 5.01 | 1.397 | 1.439 | 0.971 | 0.48 | -0.48 | 0.53 | 0.17666667 | -0.505 | -0.04 | 0.222307789 | 0.302851802 | 0.830923822 | 0.909433078 | 0.955820031 | 0.932626554 | 0.992535481 |
| Atp5a1 | High | Q03265 | ATP synthase subunit alpha, mitochondrial OS=Mus musculus OX=10090 GN=Atp5f1a PE=1 SV=1 | 0 | 263.89 | 59 | 35 | 137 | 35 | 553 | 59.7 | 9.19 | 1.436 | 1.458 | 0.985 | 0.52 | -0.52 | 0.54 | 0.18 | -0.53 | -0.02 | 0.221830363 | 0.284727943 | 0.995247996 | 0.909433078 | 0.955820031 | 0.932626554 | 0.999917596 |
| Crx | High | A0A0A0M | Cone-rod homeobox protein OS=Mus musculus OX=10090 GN=Crx PE=4 SV=1 | 0 | 12.094 | 6 | 2 | 5 | 2 | 323 | 34.9 | 9.42 | 1.387 | 1.41 | 0.984 | 0.47 | -0.47 | 0.5 | 0.16666667 | -0.485 | -0.02 | 0.223258009 | 0.324965224 | 0.967670061 | 0.909433078 | 0.957981077 | 0.933707078 | 0.994815921 |
| Rpl22; LO( | High | P67984 | 60S ribosomal protein L22 OS=Mus musculus OX=10090 GN=Rpl22 PE=1 SV=2 | 0 | 21.977 | 32 | 4 | 6 | 4 | 128 | 14.8 | 9.19 | 1.449 | 1.318 | 1.099 | 0.53 | -0.53 | 0.4 | 0.13333333 | -0.465 | 0.14 | 0.222173289 | 0.446258575 | 0.546720122 | 0.909433078 | 0.994630833 | 0.952031956 | 0.96694628 |
| Sphkap | Medium | Q6NSW3 | A-kinase anchor protein SPHKAP OS=Mus musculus OX=10090 GN=Sphkap PE=1 SV=2 | 0.022 | 2.007 | 0 | 1 | 1 | 1 | 1687 | 185 | 5.06 | 1.391 | 1.126 | 1.235 | 0.48 | -0.48 | 0.17 | 0.05666667 | -0.325 | 0.3 | 0.223268461 | 0.772121829 | 0.219233425 | 0.909433078 | 0.995559182 | 0.95249613 | 0.805859554 |
| Zdhhc5 | High | Q8VDZ4 | Palmitoyltransferase ZDHHC5 OS=Mus musculus OX=10090 GN=Zdhhc5 PE=1 SV=1 | 0.005 | 3.775 | 1 | 1 | 1 | 1 | 715 | 77.5 | 9.01 | 0.721 | 1.077 | 0.669 | -0.47 | 0.47 | 0.11 | 0.03666667 | 0.18 | -0.58 | 0.222927939 | 0.908075974 | 0.065250466 | 0.909433078 | 0.998402594 | 0.953917836 | 0.491746452 |
| Cox5a | High | P12787 | Cytochrome c oxidase subunit 5A, mitochondrial OS=Mus musculus OX=10090 GN=Cox5a PE=1 SV | 0 | 35.797 | 63 | 8 | 18 | 8 | 146 | 16.1 | 6.54 | 1.434 | 1.54 | 0.931 | 0.52 | -0.52 | 0.62 | 0.20666667 | -0.57 | -0.1 | 0.224314638 | 0.215477709 | 0.717781401 | 0.910003362 | 0.920170644 | 0.915087003 | 0.992535481 |
| Pls1 | High | Q3V0K9 | Plastin-1 OS=Mus musculus OX=10090 GN=Pls1 PE=1 SV=1 | 0 | 8.25 | 7 | 3 | 3 | 2 | 630 | 70.4 | 5.38 | 0.725 | 0.807 | 0.898 | -0.46 | 0.46 | -0.31 | -0.1033333 | 0.385 | -0.16 | 0.224771108 | 0.436619986 | 0.635088497 | 0.910003362 | 0.994630833 | 0.952317097 | 0.983562471 |
| Rps27; Gn | High | Q6ZWU9 | 40S ribosomal protein S27 OS=Mus musculus OX=10090 GN=Rps27 PE=1 SV=3 | 0 | 9.743 | 24 | 3 | 7 | 2 | 84 | 9.5 | 9.45 | 1.415 | 1.214 | 1.165 | 0.5 | -0.5 | 0.28 | 0.09333333 | -0.39 | 0.22 | 0.224192722 | 0.610975332 | 0.393995121 | 0.910003362 | 0.995559182 | 0.952781272 | 0.932122635 |
| | Medium | A0A0J9YU | Protein ABHD11 OS=Mus musculus OX=10090 GN=Abhd11 PE=1 SV=1 | 0.01 | 2.631 | 5 | 1 | 1 | 1 | 158 | 17 | 10.1 | 1.392 | 1.126 | 1.236 | 0.48 | -0.48 | 0.17 | 0.05666667 | -0.325 | 0.31 | 0.224782354 | 0.785338815 | 0.246413377 | 0.910003362 | 0.998402594 | 0.954202978 | 0.840413506 |
| Clic4 | High | Q9QYB1 | Chloride intracellular channel protein 4 OS=Mus musculus OX=10090 GN=Clic4 PE=1 SV=3 | 0.001 | 7.888 | 12 | 2 | 2 | 2 | 253 | 28.7 | 5.59 | 1.422 | 1.539 | 0.924 | 0.51 | -0.51 | 0.62 | 0.20666667 | -0.565 | -0.11 | 0.227854723 | 0.216043071 | 0.661717294 | 0.912820352 | 0.920170644 | 0.916495498 | 0.986260965 |
| Akr1b3 | High | P45376 | Aldo-keto reductase family 1 member B1 OS=Mus musculus OX=10090 GN=Akr1b1 PE=1 SV=3 | 0 | 28.735 | 26 | 7 | 10 | 7 | 316 | 35.7 | 7.18 | 0.751 | 0.725 | 1.037 | -0.41 | 0.41 | -0.46 | -0.1533333 | 0.435 | 0.05 | 0.226868711 | 0.282824554 | 0.839141769 | 0.912820352 | 0.955820031 | 0.934320191 | 0.992535481 |
| Ypel1 | High | Q9ESC7 | Protein yippee-like 1 OS=Mus musculus OX=10090 GN=Ypel1 PE=2 SV=1 | 0.001 | 4.352 | 15 | 2 | 2 | 2 | 118 | 13.5 | 8.27 | 0.752 | 0.734 | 1.024 | -0.41 | 0.41 | -0.45 | -0.15 | 0.43 | 0.03 | 0.22785843 | 0.280054367 | 0.879329763 | 0.912820352 | 0.955820031 | 0.934320191 | 0.992535481 |
| Pitpnm1 | High | O35954 | Membrane-associated phosphatidylinositol transfer protein 1 OS=Mus musculus OX=10090 GN=P | 0 | 49.5 | 14 | 12 | 13 | 12 | 1243 | 134.9 | 6.06 | 0.73 | 0.749 | 0.974 | -0.45 | 0.45 | -0.42 | -0.14 | 0.435 | -0.04 | 0.226668606 | 0.331638372 | 0.822264242 | 0.912820352 | 0.968621131 | 0.940720742 | 0.992535481 |
| Hsd17b10 | High | O08756 | 3-hydroxyacyl-CoA dehydrogenase type-2 OS=Mus musculus OX=10090 GN=Hsd17b10 PE=1 SV=4 | 0 | 7.93 | 14 | 2 | 2 | 2 | 261 | 27.4 | 8.41 | 1.38 | 1.377 | 1.002 | 0.46 | -0.46 | 0.46 | 0.15333333 | -0.46 | 0 | 0.227499494 | 0.37007905 | 0.945155819 | 0.912820352 | 0.977313184 | 0.945066768 | 0.992535481 |
| Camk2d | High | Q6PHZ2 | Calcium/calmodulin-dependent protein kinase type II subunit delta OS=Mus musculus OX=10090 ( | 0 | 124.895 | 46 | 19 | 58 | 1 | 499 | 56.3 | 7.25 | 1.39 | 1.398 | 0.995 | 0.48 | -0.48 | 0.48 | 0.16 | -0.48 | -0.01 | 0.227920188 | 0.345773735 | 0.956155873 | 0.912820352 | 0.977313184 | 0.945066768 | 0.992902921 |
| Pcp4 | High | P63054 | Calmodulin regulator protein PCP4 OS=Mus musculus OX=10090 GN=Pcp4 PE=3 SV=2 | 0 | 11.546 | 50 | 4 | 6 | 4 | 62 | 6.8 | 6.71 | 0.74 | 0.969 | 0.764 | -0.43 | 0.43 | -0.05 | -0.0166667 | 0.24 | -0.39 | 0.227693788 | 0.855561775 | 0.120704881 | 0.912820352 | 0.998402594 | 0.955611473 | 0.669025476 |
| | High | F2Z490 | Diacylglycerol kinase (ATP) OS=Mus musculus OX=10090 GN=Dgke PE=1 SV=1 | 0 | 22.387 | 14 | 5 | 7 | 5 | 406 | 46 | 8.59 | 1.414 | 1.887 | 0.749 | 0.5 | -0.5 | 0.92 | 0.30666667 | -0.71 | -0.42 | 0.233989683 | 0.0627005 | 0.10901729 | 0.915023338 | 0.587451609 | 0.751237473 | 0.631466885 |
| Ndufs7 | High | Q9DC70 | NADH dehydrogenase [ubiquinone] iron-sulfur protein 7, mitochondrial OS=Mus musculus OX=10( | 0.001 | 5.855 | 8 | 2 | 2 | 2 | 224 | 24.7 | 9.92 | 1.403 | 1.867 | 0.752 | 0.49 | -0.49 | 0.9 | 0.3 | -0.695 | -0.41 | 0.241646946 | 0.067476356 | 0.103795665 | 0.915023338 | 0.602452115 | 0.758737726 | 0.612171613 |
| Pla2g7 | High | Q60963 | Platelet-activating factor acetylhydrolase OS=Mus musculus OX=10090 GN=Pla2g7 PE=1 SV=2 | 0 | 34.689 | 19 | 7 | 10 | 7 | 440 | 49.2 | 7.12 | 1.433 | 1.865 | 0.769 | 0.52 | -0.52 | 0.9 | 0.3 | -0.71 | -0.38 | 0.235803911 | 0.067834725 | 0.102604379 | 0.915023338 | 0.603034023 | 0.75902868 | 0.610114517 |
| Pxk | Medium | Q8BX57 | PX domain-containing protein kinase-like protein OS=Mus musculus OX=10090 GN=Pxk PE=1 SV=2 | 0.015 | 2.188 | 2 | 1 | 1 | 1 | 582 | 65.2 | 9.38 | 0.74 | 0.58 | 1.275 | -0.43 | 0.43 | -0.78 | -0.26 | 0.605 | 0.35 | 0.231244834 | 0.079380422 | 0.111029044 | 0.915023338 | 0.65511466 | 0.785068999 | 0.632640059 |
| Uroc1 | Medium | Q3UEL5 | Imidazolonepropionate hydrolase OS=Mus musculus OX=10090 GN=Uroc1 PE=1 SV=1 | 0.014 | 2.246 | 3 | 1 | 1 | 1 | 710 | 78.4 | 7.27 | 1.385 | 1.733 | 0.8 | 0.47 | -0.47 | 0.79 | 0.26333333 | -0.63 | -0.32 | 0.237769979 | 0.104297892 | 0.221556687 | 0.915023338 | 0.741927143 | 0.82847524 | 0.807447486 |
| Csnk2b | High | P67871 | Casein kinase II subunit beta OS=Mus musculus OX=10090 GN=Csnk2b PE=1 SV=1 | 0 | 40.694 | 38 | 6 | 9 | 6 | 215 | 24.9 | 5.55 | 1.374 | 1.707 | 0.805 | 0.46 | -0.46 | 0.77 | 0.25666667 | -0.615 | -0.31 | 0.236628144 | 0.120005985 | 0.182752616 | 0.915023338 | 0.786986619 | 0.851004979 | 0.765268049 |
| Acvr1b | High | Q61271 | Activin receptor type-1B OS=Mus musculus OX=10090 GN=Acvr1b PE=1 SV=1 | 0.003 | 4.096 | 3 | 2 | 2 | 2 | 505 | 56.7 | 7.17 | 0.75 | 0.647 | 1.16 | -0.41 | 0.41 | -0.63 | -0.21 | 0.52 | 0.21 | 0.243409268 | 0.137389951 | 0.370302493 | 0.915023338 | 0.818645544 | 0.866834441 | 0.920164724 |
| Pfdn5 | High | Q9WU28 | Prefoldin subunit 5 OS=Mus musculus OX=10090 GN=Pfdn5 PE=1 SV=1 | 0 | 12.685 | 33 | 5 | 5 | 5 | 154 | 17.3 | 6.33 | 0.711 | 0.653 | 1.09 | -0.49 | 0.49 | -0.62 | -0.2066667 | 0.555 | 0.12 | 0.237835357 | 0.162990823 | 0.571471869 | 0.915023338 | 0.869444647 | 0.892233993 | 0.972910028 |
| Stub1 | High | Q9WUD1 | E3 ubiquitin-protein ligase CHIP OS=Mus musculus OX=10090 GN=Stub1 PE=1 SV=1 | 0 | 32.32 | 33 | 8 | 12 | 8 | 304 | 34.9 | 6.01 | 0.735 | 0.661 | 1.113 | -0.44 | 0.44 | -0.6 | -0.2 | 0.52 | 0.15 | 0.234135868 | 0.174895975 | 0.562201085 | 0.915023338 | 0.875938481 | 0.895480909 | 0.972910028 |
| Htt | High | G3X9H5 | Huntingtin OS=Mus musculus OX=10090 GN=Htt PE=1 SV=1 | 0.001 | 6.587 | 1 | 3 | 3 | 3 | 3120 | 344.6 | 6.28 | 0.754 | 0.686 | 1.1 | -0.41 | 0.41 | -0.54 | -0.18 | 0.475 | 0.14 | 0.241039005 | 0.190497037 | 0.576488808 | 0.915023338 | 0.905821916 | 0.910422627 | 0.972910028 |
| Cox5b | High | P19536 | Cytochrome c oxidase subunit 5B, mitochondrial OS=Mus musculus OX=10090 GN=Cox5b PE=1 SV | 0 | 46.667 | 37 | 7 | 18 | 7 | 128 | 13.8 | 8.38 | 1.419 | 1.519 | 0.934 | 0.51 | -0.51 | 0.6 | 0.2 | -0.555 | -0.1 | 0.237327887 | 0.231504399 | 0.736180599 | 0.915023338 | 0.936813316 | 0.925918327 | 0.992535481 |
| Sdhb | High | Q9CQA3 | Succinate dehydrogenase [ubiquinone] iron-sulfur subunit, mitochondrial OS=Mus musculus OX=1 | 0 | 39.774 | 29 | 11 | 15 | 11 | 282 | 31.8 | 8.68 | 1.419 | 1.503 | 0.944 | 0.5 | -0.5 | 0.59 | 0.19666667 | -0.545 | -0.08 | 0.24229909 | 0.244645714 | 0.793279203 | 0.915023338 | 0.941474619 | 0.928248978 | 0.992535481 |
| Tufm | High | Q8BFR5 | Elongation factor Tu, mitochondrial OS=Mus musculus OX=10090 GN=Tufm PE=1 SV=1 | 0 | 45.545 | 27 | 10 | 12 | 10 | 452 | 49.5 | 7.56 | 1.431 | 1.502 | 0.953 | 0.52 | -0.52 | 0.59 | 0.19666667 | -0.555 | -0.07 | 0.235049199 | 0.24534769 | 0.834048586 | 0.915023338 | 0.943528463 | 0.9292759 | 0.992535481 |
| Gnai3 | High | Q9DC51 | Guanine nucleotide-binding protein G(i) subunit alpha-3 OS=Mus musculus OX=10090 GN=Gnai3 | 0 | 80.846 | 36 | 15 | 91 | 5 | 354 | 40.5 | 5.69 | 0.746 | 0.725 | 1.028 | -0.42 | 0.42 | -0.46 | -0.1533333 | 0.44 | 0.04 | 0.235305559 | 0.284511115 | 0.87365601 | 0.915023338 | 0.955820031 | 0.935421684 | 0.992535481 |
| Cops3 | High | O88543 | COP9 signalosome complex subunit 3 OS=Mus musculus OX=10090 GN=Cops3 PE=1 SV=3 | 0 | 12.205 | 9 | 3 | 4 | 3 | 423 | 47.8 | 6.65 | 0.759 | 0.72 | 1.054 | -0.4 | 0.4 | -0.47 | -0.1566667 | 0.435 | 0.08 | 0.231748307 | 0.275094782 | 0.705985397 | 0.915023338 | 0.955820031 | 0.935421684 | 0.992485323 |
| Ppp1r7 | High | Q3UM45 | Protein phosphatase 1 regulatory subunit 7 OS=Mus musculus OX=10090 GN=Ppp1r7 PE=1 SV=2 | 0 | 33.442 | 25 | 7 | 10 | 7 | 361 | 41.3 | 4.92 | 0.739 | 0.742 | 0.995 | -0.44 | 0.44 | -0.43 | -0.1433333 | 0.435 | -0.01 | 0.233272481 | 0.317346489 | 0.944587861 | 0.915023338 | 0.955820031 | 0.935421684 | 0.992535481 |
| Mab21l1 | High | O70299 | Putative nucleotidyltransferase MAB21L1 OS=Mus musculus OX=10090 GN=Mab21l1 PE=2 SV=1 | 0 | 4.979 | 4 | 2 | 2 | 2 | 359 | 40.9 | 8.66 | 0.762 | 0.73 | 1.043 | -0.39 | 0.39 | -0.45 | -0.15 | 0.42 | 0.06 | 0.231714009 | 0.29333411 | 0.740162166 | 0.915023338 | 0.955820031 | 0.935421684 | 0.992535481 |
| | High | D3Z041 | Long-chain-fatty-acid--CoA ligase 1 OS=Mus musculus OX=10090 GN=Acsl1 PE=1 SV=1 | 0 | 15.53 | 10 | 6 | 7 | 6 | 699 | 78 | 7.47 | 1.377 | 1.429 | 0.964 | 0.46 | -0.46 | 0.51 | 0.17 | -0.485 | -0.05 | 0.243998873 | 0.313218595 | 0.795680049 | 0.915023338 | 0.955820031 | 0.935421684 | 0.992535481 |
| Cc2d1b | High | Q8BRN9 | Coiled-coil and C2 domain-containing protein 1B OS=Mus musculus OX=10090 GN=Cc2d1b PE=1 S | 0.001 | 6.832 | 2 | 2 | 2 | 2 | 848 | 93 | 5.41 | 1.379 | 1.425 | 0.968 | 0.46 | -0.46 | 0.51 | 0.17 | -0.485 | -0.05 | 0.236789666 | 0.304966851 | 0.873851711 | 0.915023338 | 0.955820031 | 0.935421684 | 0.992535481 |
| Acad9 | High | Q8JZN5 | Complex I assembly factor ACAD9, mitochondrial OS=Mus musculus OX=10090 GN=Acad9 PE=1 S' | 0 | 13.562 | 10 | 5 | 5 | 5 | 625 | 68.7 | 7.46 | 1.397 | 1.434 | 0.974 | 0.48 | -0.48 | 0.52 | 0.17333333 | -0.5 | -0.04 | 0.233167202 | 0.307900046 | 0.909624374 | 0.915023338 | 0.955820031 | 0.935421684 | 0.992535481 |
| Mars | High | E9QB02 | Methionine--tRNA ligase, cytoplasmic OS=Mus musculus OX=10090 GN=Mars1 PE=1 SV=1 | 0 | 14.778 | 6 | 5 | 5 | 5 | 910 | 102.3 | 6.92 | 1.398 | 1.443 | 0.969 | 0.48 | -0.48 | 0.53 | 0.17666667 | -0.505 | -0.05 | 0.240741862 | 0.298658483 | 0.879668571 | 0.915023338 | 0.955820031 | 0.935421684 | 0.992535481 |
| | High | G3UX26 | Outer mitochondrial membrane protein porin 2 OS=Mus musculus OX=10090 GN=Vdac2 PE=1 SV= | 0 | 108.443 | 61 | 15 | 33 | 15 | 283 | 30.4 | 7.58 | 1.423 | 1.425 | 0.998 | 0.51 | -0.51 | 0.51 | 0.17 | -0.51 | 0 | 0.234146754 | 0.316586381 | 0.929698081 | 0.915023338 | 0.955820031 | 0.935421684 | 0.992535481 |
| | High | A0A1L1ST | Isocitrate dehydrogenase [NAD] subunit, mitochondrial OS=Mus musculus OX=10090 GN=Idh3a P | 0 | 73.328 | 42 | 16 | 29 | 16 | 384 | 41.5 | 6.93 | 1.42 | 1.44 | 0.986 | 0.51 | -0.51 | 0.53 | 0.17666667 | -0.52 | -0.02 | 0.236714624 | 0.301528922 | 0.992632159 | 0.915023338 | 0.955820031 | 0.935421684 | 0.999783808 |
| | High | F6ZQL0 | Protein LYRIC (Fragment) OS=Mus musculus OX=10090 GN=Mtdh PE=1 SV=1 | 0.005 | 3.82 | 5 | 1 | 1 | 1 | 220 | 24.4 | 9.31 | 1.403 | 1.474 | 0.952 | 0.49 | -0.49 | 0.56 | 0.18666667 | -0.525 | -0.07 | 0.229862216 | 0.278323109 | 0.858893012 | 0.915023338 | 0.955820031 | 0.935421684 | 0.992535481 |
| | High | F8WHP8 | ATP synthase membrane subunit f OS=Mus musculus OX=10090 GN=Atp5j2 PE=1 SV=1 | 0.001 | 6.345 | 16 | 1 | 3 | 1 | 76 | 9 | 9.88 | 1.401 | 1.369 | 1.023 | 0.49 | -0.49 | 0.45 | 0.15 | -0.47 | 0.03 | 0.237716408 | 0.37991137 | 0.868865119 | 0.915023338 | 0.977313184 | 0.946168261 | 0.992535481 |
| Abcb7 | High | Q61102 | Iron-sulfur clusters transporter ABCB7, mitochondrial OS=Mus musculus OX=10090 GN=Abcb7 PE | 0 | 4.875 | 1 | 1 | 1 | 1 | 752 | 82.5 | 9.32 | 1.396 | 1.391 | 1.003 | 0.48 | -0.48 | 0.48 | 0.16 | -0.48 | 0 | 0.243499846 | 0.34994171 | 0.932547799 | 0.915023338 | 0.977313184 | 0.946168261 | 0.992535481 |
| Cyb5r1 | High | Q9DB73 | NADH-cytochrome b5 reductase 1 OS=Mus musculus OX=10090 GN=Cyb5r1 PE=1 SV=1 | 0 | 25.379 | 20 | 6 | 9 | 6 | 305 | 34.1 | 8.87 | 1.372 | 1.363 | 1.007 | 0.46 | -0.46 | 0.45 | 0.15 | -0.455 | 0.01 | 0.243319722 | 0.3875762 | 0.997748431 | 0.915023338 | 0.981101468 | 0.948062403 | 0.999917596 |
| Lta4h | High | P24527 | Leukotriene A-4 hydrolase OS=Mus musculus OX=10090 GN=Lta4h PE=1 SV=4 | 0 | 14.794 | 9 | 5 | 5 | 5 | 611 | 69 | 6.42 | 0.747 | 0.778 | 0.96 | -0.42 | 0.42 | -0.36 | -0.12 | 0.39 | -0.06 | 0.235188414 | 0.393095761 | 0.828810612 | 0.915023338 | 0.983884835 | 0.949454087 | 0.992535481 |

| | | | | | | | | | | | | | | | | | | | | | | | | | | | | |
|---|---|---|---|---|---|---|---|---|---|---|---|---|---|---|---|---|---|---|---|---|---|---|---|---|---|---|---|---|
| Srp9 | High | P49962 | Signal recognition particle 9 kDa protein OS=Mus musculus OX=10090 GN=Srp9 PE=1 SV=2 | 0.007 | 3.068 | 9 | 1 | 1 | 1 | 86 | 10.2 | 7.99 | 0.725 | 0.791 | 0.916 | -0.46 | 0.46 | -0.34 | -0.1133333 | 0.4 | -0.13 | 0.234295139 | 0.398978631 | 0.717307638 | 0.915023338 | 0.985440071 | 0.950231704 | 0.992535481 |
| Creld2 | High | Q9CYA0 | Protein disulfide isomerase Creld2 OS=Mus musculus OX=10090 GN=Creld2 PE=1 SV=1 | 0.001 | 4.67 | 6 | 2 | 2 | 2 | 350 | 38.2 | 4.58 | 1.403 | 1.356 | 1.035 | 0.49 | -0.49 | 0.44 | 0.14666667 | -0.465 | 0.05 | 0.229474122 | 0.408124926 | 0.796553345 | 0.915023338 | 0.9855335 | 0.950278419 | 0.992535481 |
| Gap43 | High | P06837 | Neuromodulin OS=Mus musculus OX=10090 GN=Gap43 PE=1 SV=1 | 0 | 75.328 | 61 | 9 | 23 | 9 | 227 | 23.6 | 4.73 | 0.705 | 0.797 | 0.885 | -0.5 | 0.5 | -0.33 | -0.11 | 0.415 | -0.18 | 0.241077537 | 0.434123036 | 0.491327486 | 0.915023338 | 0.99399649 | 0.954509914 | 0.964378027 |
| | High | E9Q123 | C-terminal-binding protein 2 OS=Mus musculus OX=10090 GN=Ctbp2 PE=4 SV=1 | 0 | 9.131 | 16 | 3 | 3 | 3 | 188 | 20.6 | 8.31 | 0.736 | 0.801 | 0.919 | -0.44 | 0.44 | -0.32 | -0.1066667 | 0.38 | -0.12 | 0.229112676 | 0.427223314 | 0.674921125 | 0.915023338 | 0.99399649 | 0.954509914 | 0.990250803 |
| Cpne5 | High | Q8JZW4 | Copine-5 OS=Mus musculus OX=10090 GN=Cpne5 PE=1 SV=1 | 0 | 11.863 | 7 | 3 | 4 | 1 | 593 | 65.6 | 5.77 | 0.754 | 0.807 | 0.934 | -0.41 | 0.41 | -0.31 | -0.1033333 | 0.36 | -0.1 | 0.243548562 | 0.432073534 | 0.745480325 | 0.915023338 | 0.99399649 | 0.954509914 | 0.992535481 |
| Ckb | High | Q04447 | Creatine kinase B-type OS=Mus musculus OX=10090 GN=Ckb PE=1 SV=1 | 0 | 219.731 | 71 | 21 | 99 | 21 | 381 | 42.7 | 5.67 | 0.699 | 0.803 | 0.871 | -0.52 | 0.52 | -0.32 | -0.1066667 | 0.42 | -0.2 | 0.229932761 | 0.447176526 | 0.430371272 | 0.915023338 | 0.994630833 | 0.954827085 | 0.945198482 |
| | Medium | A0A498W | Thioredoxin-like protein 1 OS=Mus musculus OX=10090 GN=Txnl1 PE=1 SV=1 | 0.017 | 2.131 | 6 | 1 | 1 | 1 | 292 | 32.7 | 5.07 | 0.733 | 0.884 | 0.829 | -0.45 | 0.45 | -0.18 | -0.06 | 0.315 | -0.27 | 0.243408223 | 0.62004341 | 0.412084959 | 0.915023338 | 0.995559182 | 0.95529126 | 0.936397673 |
| Cartpt | High | P56388 | Cocaine- and amphetamine-regulated transcript protein OS=Mus musculus OX=10090 GN=Cartpt | 0.001 | 6.205 | 12 | 1 | 1 | 1 | 129 | 14.3 | 8.87 | 0.728 | 0.928 | 0.784 | -0.46 | 0.46 | -0.11 | -0.0366667 | 0.285 | -0.35 | 0.234203552 | 0.729263506 | 0.260818458 | 0.915023338 | 0.995559182 | 0.95529126 | 0.857215179 |
| Wdr6 | Low | Q99ME2 | WD repeat-containing protein 6 OS=Mus musculus OX=10090 GN=Wdr6 PE=1 SV=1 | 0.077 | 1.23 | 1 | 1 | 1 | 1 | 1125 | 121.8 | 6.98 | 0.754 | 0.931 | 0.809 | -0.41 | 0.41 | -0.1 | -0.0333333 | 0.255 | -0.31 | 0.23480448 | 0.750774851 | 0.277690402 | 0.915023338 | 0.995559182 | 0.95529126 | 0.876067403 |
| | Medium | A0A1L1SS | CCR4-NOT transcription complex subunit 10 OS=Mus musculus OX=10090 GN=Cnot10 PE=1 SV=1 | 0.01 | 2.561 | 2 | 1 | 1 | 1 | 405 | 45 | 7.9 | 1.423 | 0.87 | 1.635 | 0.51 | -0.51 | -0.2 | -0.0666667 | -0.155 | 0.71 | 0.238610833 | 0.585786771 | 0.018230966 | 0.915023338 | 0.995559182 | 0.95529126 | 0.237723313 |
| U2af2 | High | P26369 | Splicing factor U2AF 65 kDa subunit OS=Mus musculus OX=10090 GN=U2af2 PE=1 SV=3 | 0.001 | 6.691 | 4 | 1 | 1 | 1 | 475 | 53.5 | 9.09 | 1.375 | 1.189 | 1.157 | 0.46 | -0.46 | 0.25 | 0.08333333 | -0.355 | 0.21 | 0.237526778 | 0.643634293 | 0.380986069 | 0.915023338 | 0.995559182 | 0.95529126 | 0.923104982 |
| Rps4x | High | P62702 | 40S ribosomal protein S4, X isoform OS=Mus musculus OX=10090 GN=Rps4x PE=1 SV=2 | 0 | 74.43 | 47 | 12 | 23 | 12 | 263 | 29.6 | 10.15 | 1.414 | 1.164 | 1.215 | 0.5 | -0.5 | 0.22 | 0.07333333 | -0.36 | 0.28 | 0.242380847 | 0.703948793 | 0.175584405 | 0.915023338 | 0.995559182 | 0.95529126 | 0.745270069 |
| Dennd4b | High | Q3U1Y4 | DENN domain-containing protein 4B OS=Mus musculus OX=10090 GN=Dennd4b PE=1 SV=2 | 0 | 11.123 | 2 | 2 | 2 | 2 | 1499 | 164.6 | 7.71 | 1.421 | 1.172 | 1.212 | 0.51 | -0.51 | 0.23 | 0.07666667 | -0.37 | 0.28 | 0.233188844 | 0.707288352 | 0.304410001 | 0.915023338 | 0.995559182 | 0.95529126 | 0.900241969 |
| Uck1 | High | P52623 | Uridine-cytidine kinase 1 OS=Mus musculus OX=10090 GN=Uck1 PE=1 SV=2 | 0.001 | 4.503 | 6 | 1 | 1 | 1 | 277 | 31 | 7.88 | 1.355 | 1.228 | 1.104 | 0.44 | -0.44 | 0.3 | 0.1 | -0.37 | 0.14 | 0.230277317 | 0.572338136 | 0.563552151 | 0.915023338 | 0.995559182 | 0.95529126 | 0.972910028 |
| Rpl4 | High | Q9D8E6 | 60S ribosomal protein L4 OS=Mus musculus OX=10090 GN=Rpl4 PE=1 SV=3 | 0 | 115.12 | 57 | 25 | 42 | 25 | 419 | 47.1 | 11 | 1.415 | 1.221 | 1.158 | 0.5 | -0.5 | 0.29 | 0.09666667 | -0.395 | 0.21 | 0.241724574 | 0.597987629 | 0.29459447 | 0.915023338 | 0.995559182 | 0.95529126 | 0.894411787 |
| Arhgap24 | High | D3Z5T4 | Rho GTPase-activating protein 24 OS=Mus musculus OX=10090 GN=Arhgap24 PE=1 SV=1 | 0.001 | 6.305 | 3 | 2 | 2 | 2 | 657 | 73.5 | 8.16 | 1.404 | 1.236 | 1.136 | 0.49 | -0.49 | 0.31 | 0.10333333 | -0.4 | 0.18 | 0.233684264 | 0.572592449 | 0.4114614 | 0.915023338 | 0.995559182 | 0.95529126 | 0.936397673 |
| Rpl10 | High | I7HLV2 | 60S ribosomal protein L10 (Fragment) OS=Mus musculus OX=10090 GN=Rpl10 PE=1 SV=1 | 0 | 34.687 | 27 | 6 | 11 | 6 | 201 | 23.1 | 10.01 | 1.418 | 1.242 | 1.141 | 0.5 | -0.5 | 0.31 | 0.10333333 | -0.405 | 0.19 | 0.240289191 | 0.56243873 | 0.378419538 | 0.915023338 | 0.995559182 | 0.95529126 | 0.922487352 |
| Slc39a12 | High | Q5FWH7 | Zinc transporter ZIP12 OS=Mus musculus OX=10090 GN=Slc39a12 PE=1 SV=1 | 0 | 33.93 | 12 | 7 | 10 | 7 | 689 | 76.2 | 5.77 | 1.43 | 1.228 | 1.164 | 0.52 | -0.52 | 0.3 | 0.1 | -0.41 | 0.22 | 0.242796597 | 0.585851089 | 0.303931255 | 0.915023338 | 0.995559182 | 0.95529126 | 0.900241969 |
| Sirt2 | High | Q8VDQ8 | NAD-dependent protein deacetylase sirtuin-2 OS=Mus musculus OX=10090 GN=Sirt2 PE=1 SV=2 | 0 | 74.497 | 47 | 15 | 22 | 15 | 389 | 43.2 | 5.35 | 1.414 | 1.253 | 1.129 | 0.5 | -0.5 | 0.32 | 0.10666667 | -0.41 | 0.18 | 0.242205762 | 0.545263295 | 0.377341965 | 0.915023338 | 0.995559182 | 0.95529126 | 0.92149669 |
| Rpl36 | High | Q6ZWZ4 | 60S ribosomal protein L36 OS=Mus musculus OX=10090 GN=Rpl36 PE=1 SV=1 | 0 | 14.836 | 20 | 2 | 8 | 2 | 105 | 12.2 | 11.59 | 1.413 | 1.275 | 1.108 | 0.5 | -0.5 | 0.35 | 0.11666667 | -0.425 | 0.15 | 0.242099606 | 0.509529454 | 0.553194091 | 0.915023338 | 0.995559182 | 0.95529126 | 0.96775057 |
| Rps9 | High | Q6ZWN5 | 40S ribosomal protein S9 OS=Mus musculus OX=10090 GN=Rps9 PE=1 SV=3 | 0 | 24.991 | 42 | 11 | 19 | 11 | 194 | 22.6 | 10.65 | 1.425 | 1.279 | 1.114 | 0.51 | -0.51 | 0.35 | 0.11666667 | -0.43 | 0.16 | 0.232382082 | 0.504093866 | 0.425191757 | 0.915023338 | 0.995559182 | 0.95529126 | 0.941180024 |
| Rpl35a-ps | High | O55142 | 60S ribosomal protein L35a OS=Mus musculus OX=10090 GN=Rpl35a PE=1 SV=2 | 0 | 23.084 | 45 | 7 | 12 | 7 | 110 | 12.5 | 10.89 | 1.395 | 1.313 | 1.062 | 0.48 | -0.48 | 0.39 | 0.13 | -0.435 | 0.09 | 0.237103911 | 0.452948909 | 0.643956882 | 0.915023338 | 0.995559182 | 0.95529126 | 0.985010103 |
| Gm13826; | High | Q9D823 | 60S ribosomal protein L37 OS=Mus musculus OX=10090 GN=Rpl37 PE=1 SV=3 | 0.001 | 5.641 | 28 | 4 | 5 | 4 | 97 | 11.1 | 11.74 | 1.403 | 1.304 | 1.076 | 0.49 | -0.49 | 0.38 | 0.12666667 | -0.435 | 0.11 | 0.242502204 | 0.466850522 | 0.650375769 | 0.915023338 | 0.995559182 | 0.95529126 | 0.985010103 |
| Epn1 | High | Q80VP1 | Epsin-1 OS=Mus musculus OX=10090 GN=Epn1 PE=1 SV=3 | 0 | 23.17 | 14 | 6 | 9 | 6 | 575 | 60.2 | 4.81 | 0.744 | 0.988 | 0.753 | -0.43 | 0.43 | -0.02 | -0.0066667 | 0.225 | -0.41 | 0.233375084 | 0.902164145 | 0.094919024 | 0.915023338 | 0.998402594 | 0.956712966 | 0.586204524 |
| Scaper | High | F8VQ70 | S phase cyclin A-associated protein in the ER OS=Mus musculus OX=10090 GN=Scaper PE=1 SV=1 | 0.005 | 3.594 | 1 | 2 | 2 | 2 | 1398 | 157.7 | 7.31 | 1.37 | 0.967 | 1.417 | 0.45 | -0.45 | -0.05 | -0.0166667 | -0.2 | 0.5 | 0.236367887 | 0.846244402 | 0.032588667 | 0.915023338 | 0.998402594 | 0.956712966 | 0.349748955 |
| Brap | High | Q99MP8 | BRCA1-associated protein OS=Mus musculus OX=10090 GN=Brap PE=1 SV=1 | 0.001 | 5.464 | 2 | 1 | 1 | 1 | 591 | 66.9 | 5.64 | 1.409 | 1.014 | 1.389 | 0.49 | -0.49 | 0.02 | 0.00666667 | -0.255 | 0.47 | 0.244136778 | 0.944628262 | 0.083844268 | 0.915023338 | 0.998402594 | 0.956712966 | 0.558386036 |
| | Medium | A0A087W | Death-associated protein 1 OS=Mus musculus OX=10090 GN=Dap PE=1 SV=1 | 0.01 | 2.589 | 10 | 1 | 1 | 1 | 79 | 8.9 | 9.19 | 1.362 | 1.695 | 0.804 | 0.45 | -0.45 | 0.76 | 0.25333333 | -0.605 | -0.32 | 0.245449273 | 0.115309012 | 0.214463955 | 0.917461722 | 0.77945068 | 0.848456201 | 0.804011919 |
| Qdpr; LOC | High | Q8BVI4 | Dihydropteridine reductase OS=Mus musculus OX=10090 GN=Qdpr PE=1 SV=2 | 0 | 55.789 | 43 | 8 | 13 | 8 | 241 | 25.6 | 7.81 | 0.713 | 0.638 | 1.118 | -0.49 | 0.49 | -0.65 | -0.2166667 | 0.57 | 0.16 | 0.245605501 | 0.142625 | 0.44178108 | 0.917461722 | 0.836504579 | 0.876983151 | 0.956765746 |
| Hist1h1b | High | P43276 | Histone H1.5 OS=Mus musculus OX=10090 GN=H1-5 PE=1 SV=2 | 0 | 27.168 | 30 | 9 | 14 | 4 | 223 | 22.6 | 10.92 | 1.388 | 1.841 | 0.754 | 0.47 | -0.47 | 0.88 | 0.29333333 | -0.675 | -0.41 | 0.246217536 | 0.074062989 | 0.075239883 | 0.918524102 | 0.636899913 | 0.777712008 | 0.520184986 |
| Cox7b | Medium | P56393 | Cytochrome c oxidase subunit 7B, mitochondrial OS=Mus musculus OX=10090 GN=Cox7b PE=3 SV | 0.028 | 1.927 | 9 | 1 | 1 | 1 | 80 | 9 | 10.27 | 1.352 | 1.371 | 0.986 | 0.43 | -0.43 | 0.46 | 0.15333333 | -0.445 | -0.02 | 0.246057524 | 0.377768777 | 0.932107139 | 0.918524102 | 0.977313184 | 0.947918643 | 0.992535481 |
| Rpl13 | High | P47963 | 60S ribosomal protein L13 OS=Mus musculus OX=10090 GN=Rpl13 PE=1 SV=3 | 0 | 42.185 | 36 | 8 | 19 | 8 | 211 | 24.3 | 11.55 | 1.409 | 1.282 | 1.099 | 0.5 | -0.5 | 0.36 | 0.12 | -0.43 | 0.14 | 0.247004049 | 0.499068182 | 0.477220938 | 0.920845548 | 0.995559182 | 0.958202365 | 0.963549802 |
| Iars | High | Q8BU30 | Isoleucine--tRNA ligase, cytoplasmic OS=Mus musculus OX=10090 GN=Iars1 PE=1 SV=2 | 0.003 | 3.895 | 1 | 1 | 1 | 1 | 1262 | 144.2 | 6.55 | 1.384 | 1.145 | 1.208 | 0.47 | -0.47 | 0.2 | 0.06666667 | -0.335 | 0.27 | 0.247282641 | 0.741154511 | 0.231528391 | 0.921271608 | 0.995559182 | 0.958415395 | 0.819676582 |
| Ppid | High | Q9CR16 | Peptidyl-prolyl cis-trans isomerase D OS=Mus musculus OX=10090 GN=Ppid PE=1 SV=3 | 0 | 60.786 | 34 | 14 | 20 | 13 | 370 | 40.7 | 7.43 | 0.712 | 0.507 | 1.405 | -0.49 | 0.49 | -0.98 | -0.3266667 | 0.735 | 0.49 | 0.254252444 | 0.029949722 | 0.021952992 | 0.921417129 | 0.439602331 | 0.68050973 | 0.268170868 |
| Syne2 | High | Q6ZWQ0 | Nesprin-2 OS=Mus musculus OX=10090 GN=Syne2 PE=1 SV=2 | 0.001 | 4.594 | 0 | 3 | 4 | 1 | 6874 | 782.2 | 5.33 | 0.78 | 0.538 | 1.45 | -0.36 | 0.36 | -0.89 | -0.2966667 | 0.625 | 0.54 | 0.255269592 | 0.038869837 | 0.022340178 | 0.921417129 | 0.486480307 | 0.703948718 | 0.270201833 |
| Fkbp1a | High | P26883 | Peptidyl-prolyl cis-trans isomerase FKBP1A OS=Mus musculus OX=10090 GN=Fkbp1a PE=1 SV=2 | 0 | 12.98 | 13 | 1 | 4 | 1 | 108 | 11.9 | 8.16 | 0.749 | 0.639 | 1.174 | -0.42 | 0.42 | -0.65 | -0.2166667 | 0.535 | 0.23 | 0.253443626 | 0.136537277 | 0.343971194 | 0.921417129 | 0.817450082 | 0.869433606 | 0.907770836 |
| Ttc39b | High | Q8BYY4 | Tetratricopeptide repeat protein 39B OS=Mus musculus OX=10090 GN=Ttc39b PE=1 SV=1 | 0.001 | 7.498 | 5 | 3 | 3 | 3 | 617 | 70.2 | 6.64 | 1.352 | 1.586 | 0.852 | 0.44 | -0.44 | 0.67 | 0.22333333 | -0.555 | -0.23 | 0.25334061 | 0.18366644 | 0.349013049 | 0.921417129 | 0.895493677 | 0.908455403 | 0.909466648 |
| | High | H3BLI8 | WD repeat domain 17 OS=Mus musculus OX=10090 GN=Wdr17 PE=1 SV=1 | 0 | 182.026 | 35 | 41 | 64 | 41 | 1283 | 142.6 | 6.83 | 0.709 | 0.678 | 1.046 | -0.5 | 0.5 | -0.56 | -0.1866667 | 0.53 | 0.06 | 0.248825624 | 0.20144652 | 0.697244183 | 0.921417129 | 0.912882441 | 0.917149785 | 0.991285312 |
| Ndufc2; LO | High | Q9CQ54 | NADH dehydrogenase [ubiquinone] 1 subunit C2 OS=Mus musculus OX=10090 GN=Ndufc2 PE=1 S | 0 | 14.074 | 25 | 3 | 5 | 3 | 120 | 14.2 | 9.2 | 1.336 | 1.554 | 0.859 | 0.42 | -0.42 | 0.64 | 0.21333333 | -0.53 | -0.22 | 0.251334299 | 0.205374615 | 0.34056596 | 0.921417129 | 0.912882441 | 0.917149785 | 0.907770836 |
| Tfam | High | A0A0R4J1 | Transcription factor A, mitochondrial OS=Mus musculus OX=10090 GN=Tfam PE=1 SV=1 | 0.001 | 4.242 | 9 | 2 | 2 | 2 | 199 | 23.4 | 9.72 | 1.379 | 1.472 | 0.937 | 0.46 | -0.46 | 0.56 | 0.18666667 | -0.51 | -0.09 | 0.253771819 | 0.2709745 | 0.724477189 | 0.921417129 | 0.954967959 | 0.938192544 | 0.992535481 |
| Pgls | High | Q9CQ60 | 6-phosphogluconolactonase OS=Mus musculus OX=10090 GN=Pgls PE=1 SV=1 | 0 | 24.393 | 40 | 6 | 7 | 6 | 257 | 27.2 | 5.85 | 0.758 | 0.732 | 1.035 | -0.4 | 0.4 | -0.45 | -0.15 | 0.425 | 0.05 | 0.254908137 | 0.297598864 | 0.841591876 | 0.921417129 | 0.955820031 | 0.93861858 | 0.992535481 |
| Ptpn23 | High | Q6PB44 | Tyrosine-protein phosphatase non-receptor type 23 OS=Mus musculus OX=10090 GN=Ptpn23 PE= | 0.001 | 5.429 | 2 | 2 | 2 | 2 | 1692 | 185.1 | 6.8 | 0.754 | 0.751 | 1.003 | -0.41 | 0.41 | -0.41 | -0.1366667 | 0.41 | 0 | 0.250374527 | 0.314894881 | 0.911427631 | 0.921417129 | 0.955820031 | 0.93861858 | 0.992535481 |
| Pdpn | High | Q62011 | Podoplanin OS=Mus musculus OX=10090 GN=Pdpn PE=1 SV=2 | 0.001 | 5.253 | 8 | 1 | 1 | 1 | 172 | 18.2 | 5.53 | 0.746 | 0.784 | 0.952 | -0.42 | 0.42 | -0.35 | -0.1166667 | 0.385 | -0.07 | 0.254705258 | 0.396406058 | 0.80738578 | 0.921417129 | 0.985399792 | 0.953408461 | 0.992535481 |
| Atp6v1g2 | High | B0V2H4 | V-type proton ATPase subunit G OS=Mus musculus OX=10090 GN=Atp6v1g2 PE=1 SV=1 | 0 | 46.428 | 80 | 6 | 11 | 6 | 93 | 10.9 | 10.24 | 0.728 | 0.797 | 0.914 | -0.46 | 0.46 | -0.33 | -0.11 | 0.395 | -0.13 | 0.249830614 | 0.434350794 | 0.622211065 | 0.921417129 | 0.99399649 | 0.95770681 | 0.97826833 |
| Eef1d | High | A0A0R4J1 | Elongation factor 1-delta OS=Mus musculus OX=10090 GN=Eef1d PE=1 SV=1 | 0 | 14.413 | 6 | 3 | 4 | 3 | 660 | 72.9 | 6.51 | 1.392 | 1.329 | 1.047 | 0.48 | -0.48 | 0.41 | 0.13666667 | -0.445 | 0.07 | 0.254760112 | 0.432124985 | 0.785305899 | 0.921417129 | 0.99399649 | 0.95770681 | 0.992535481 |
| Actn1 | High | A1BN54 | Alpha actinin 1a OS=Mus musculus OX=10090 GN=Actn1 PE=1 SV=1 | 0 | 45.702 | 15 | 11 | 16 | 3 | 887 | 102.7 | 5.48 | 0.712 | 0.85 | 0.839 | -0.49 | 0.49 | -0.24 | -0.08 | 0.365 | -0.25 | 0.2550325 | 0.558110176 | 0.305877141 | 0.921417129 | 0.995559182 | 0.958488156 | 0.901236536 |
| Ncdn | Medium | Q9Z0E0 | Neurochondrin OS=Mus musculus OX=10090 GN=Ncdn PE=1 SV=1 | 0.034 | 1.83 | 1 | 1 | 1 | 1 | 729 | 78.8 | 5.54 | 0.751 | 0.914 | 0.821 | -0.41 | 0.41 | -0.13 | -0.0433333 | 0.27 | -0.28 | 0.255374036 | 0.713304171 | 0.299391896 | 0.921417129 | 0.995559182 | 0.958488156 | 0.900241969 |
| Gclm | High | O09172 | Glutamate--cysteine ligase regulatory subunit OS=Mus musculus OX=10090 GN=Gclm PE=1 SV=1 | 0 | 10.414 | 13 | 3 | 3 | 3 | 274 | 30.5 | 5.52 | 0.786 | 0.885 | 0.888 | -0.35 | 0.35 | -0.18 | -0.06 | 0.265 | -0.17 | 0.251486118 | 0.641353788 | 0.48397978 | 0.921417129 | 0.995559182 | 0.958488156 | 0.964378027 |
| Tjp1 | High | P39447 | Tight junction protein ZO-1 OS=Mus musculus OX=10090 GN=Tjp1 PE=1 SV=2 | 0.001 | 5.223 | 1 | 1 | 1 | 1 | 1745 | 194.6 | 6.64 | 0.766 | 0.92 | 0.833 | -0.38 | 0.38 | -0.12 | -0.04 | 0.25 | -0.26 | 0.250212074 | 0.7318436 | 0.305469294 | 0.921417129 | 0.995559182 | 0.958488156 | 0.900981762 |
| Samd11 | High | Q1RNF8 | Sterile alpha motif domain-containing protein 11 OS=Mus musculus OX=10090 GN=Samd11 PE=2 | 0.005 | 3.817 | 2 | 1 | 1 | 1 | 542 | 57.7 | 6.28 | 1.411 | 0.893 | 1.579 | 0.5 | -0.5 | -0.16 | -0.0533333 | -0.17 | 0.66 | 0.249495902 | 0.640459236 | 0.024273456 | 0.921417129 | 0.995559182 | 0.958488156 | 0.285927032 |
| Slc29a1 | High | Q9JIM1 | Equilibrative nucleoside transporter 1 OS=Mus musculus OX=10090 GN=Slc29a1 PE=1 SV=3 | 0.001 | 4.482 | 3 | 1 | 1 | 1 | 460 | 50.2 | 8.31 | 1.415 | 0.942 | 1.502 | 0.5 | -0.5 | -0.09 | -0.03 | -0.205 | 0.59 | 0.253333163 | 0.764419003 | 0.044879114 | 0.921417129 | 0.995559182 | 0.958488156 | 0.405968973 |
| Eps8 | High | Q08509 | Epidermal growth factor receptor kinase substrate 8 OS=Mus musculus OX=10090 GN=Eps8 PE=1 | 0 | 10.334 | 4 | 3 | 4 | 3 | 821 | 91.7 | 7.65 | 1.394 | 1.185 | 1.176 | 0.48 | -0.48 | 0.25 | 0.08333333 | -0.365 | 0.23 | 0.249229545 | 0.666080411 | 0.275956787 | 0.921417129 | 0.995559182 | 0.958488156 | 0.876067403 |
| Cnot1 | High | Q6ZQ08 | CCR4-NOT transcription complex subunit 1 OS=Mus musculus OX=10090 GN=Cnot1 PE=1 SV=2 | 0 | 11.046 | 2 | 4 | 4 | 4 | 2375 | 266.6 | 7.11 | 1.361 | 1.241 | 1.096 | 0.44 | -0.44 | 0.31 | 0.10333333 | -0.375 | 0.13 | 0.248942878 | 0.554268146 | 0.568154617 | 0.921417129 | 0.995559182 | 0.958488156 | 0.972910028 |
| Ssx2ip | High | Q8VC66 | Afadin- and alpha-actinin-binding protein OS=Mus musculus OX=10090 GN=Ssx2ip PE=1 SV=1 | 0.002 | 4.167 | 2 | 2 | 2 | 2 | 615 | 70.9 | 6.65 | 1.357 | 1.295 | 1.048 | 0.44 | -0.44 | 0.37 | 0.12333333 | -0.405 | 0.07 | 0.254196821 | 0.484097784 | 0.715632997 | 0.921417129 | 0.995559182 | 0.958488156 | 0.992535481 |
| Spag7 | High | Q7TNE3 | Sperm-associated antigen 7 OS=Mus musculus OX=10090 GN=Spag7 PE=1 SV=1 | 0 | 36.052 | 40 | 8 | 10 | 8 | 227 | 25.9 | 7.43 | 1.43 | 1.264 | 1.132 | 0.52 | -0.52 | 0.34 | 0.11333333 | -0.43 | 0.18 | 0.250454649 | 0.527025492 | 0.456740872 | 0.921417129 | 0.995559182 | 0.958488156 | 0.959744396 |
| Rpl21; Gm | High | Q9CQM8 | 60S ribosomal protein L21 OS=Mus musculus OX=10090 GN=Rpl21 PE=1 SV=1 | 0 | 24.559 | 46 | 8 | 17 | 8 | 160 | 18.6 | 10.49 | 1.405 | 1.309 | 1.073 | 0.49 | -0.49 | 0.39 | 0.13 | -0.44 | 0.1 | 0.251501552 | 0.459286418 | 0.578758459 | 0.921417129 | 0.995559182 | 0.958488156 | 0.972910028 |
| Cplx4 | High | Q80WM3 | Complexin-4 OS=Mus musculus OX=10090 GN=Cplx4 PE=1 SV=2 | 0 | 80.6 | 75 | 10 | 35 | 8 | 160 | 18.3 | 4.72 | 0.71 | 1.035 | 0.685 | -0.5 | 0.5 | 0.05 | 0.01666667 | 0.225 | -0.54 | 0.249559295 | 0.983995944 | 0.019606379 | 0.921417129 | 0.998402594 | 0.959909862 | 0.249847648 |
| Try5 | High | Q9QUK9 | TESP4 OS=Mus musculus OX=10090 GN=Try5 PE=1 SV=1 | 0 | 8.261 | 12 | 2 | 2 | 1 | 246 | 26.3 | 5.3 | 0.747 | 1.078 | 0.693 | -0.42 | 0.42 | 0.11 | 0.03666667 | 0.155 | -0.53 | 0.251591308 | 0.884703361 | 0.03782332 | 0.921417129 | 0.998402594 | 0.959909862 | 0.373579725 |
| | High | A0A0G2JG | Kelch-like protein 18 OS=Mus musculus OX=10090 GN=Klhl18 PE=1 SV=1 | 0.001 | 6.429 | 3 | 2 | 2 | 2 | 509 | 56.4 | 5.77 | 1.36 | 1.021 | 1.332 | 0.44 | -0.44 | 0.03 | 0.01 | -0.235 | 0.41 | 0.252301342 | 0.966812742 | 0.058708515 | 0.921417129 | 0.998402594 | 0.959909862 | 0.4662587 |
| Micall1 | High | Q8BGT6 | MICAL-like protein 1 OS=Mus musculus OX=10090 GN=Micall1 PE=1 SV=3 | 0.003 | 3.845 | 1 | 1 | 1 | 1 | 870 | 94 | 6.68 | 1.391 | 2.04 | 0.682 | 0.48 | -0.48 | 1.03 | 0.34333333 | -0.755 | -0.55 | 0.263469797 | 0.036435277 | 0.055296331 | 0.923229118 | 0.476392832 | 0.699810975 | 0.450649022 |
| Dpp4 | High | P28843 | Dipeptidyl peptidase 4 OS=Mus musculus OX=10090 GN=Dpp4 PE=1 SV=3 | 0.006 | 3.136 | 1 | 1 | 1 | 1 | 760 | 87.4 | 6.42 | 1.366 | 1.966 | 0.695 | 0.45 | -0.45 | 0.97 | 0.32333333 | -0.71 | -0.53 | 0.264676519 | 0.041342461 | 0.044110963 | 0.923229118 | 0.503577691 | 0.713403405 | 0.405968973 |
| Crk | High | Q5ND50 | Adapter molecule crk OS=Mus musculus OX=10090 GN=Crk PE=1 SV=2 | 0.001 | 5.293 | 6 | 1 | 1 | 1 | 261 | 29.3 | 7.14 | 0.742 | 0.599 | 1.238 | -0.43 | 0.43 | -0.74 | -0.2466667 | 0.585 | 0.31 | 0.259816428 | 0.090428213 | 0.284913493 | 0.923229118 | 0.696517952 | 0.809873535 | 0.886074068 |
| Rap1gap | High | A2ALS5 | Rap1 GTPase-activating protein 1 OS=Mus musculus OX=10090 GN=Rap1gap PE=1 SV=2 | 0.001 | 6.325 | 3 | 2 | 2 | 2 | 663 | 73.4 | 5.87 | 1.373 | 1.648 | 0.833 | 0.46 | -0.46 | 0.72 | 0.24 | -0.59 | -0.26 | 0.261111371 | 0.14815228 | 0.286508344 | 0.923229118 | 0.845081896 | 0.884155507 | 0.886074068 |
| Uqcrb | High | Q9CQB4 | Cytochrome b-c1 complex subunit 7 OS=Mus musculus OX=10090 GN=Uqcrb PE=1 SV=1 | 0 | 28.308 | 64 | 8 | 13 | 8 | 111 | 13.6 | 9.11 | 1.394 | 1.628 | 0.856 | 0.48 | -0.48 | 0.7 | 0.23333333 | -0.59 | -0.22 | 0.256261688 | 0.158934554 | 0.368412369 | 0.923229118 | 0.854406564 | 0.888817841 | 0.920164724 |
| Hspd1 | High | P63038 | 60 kDa heat shock protein, mitochondrial OS=Mus musculus OX=10090 GN=Hspd1 PE=1 SV=1 | 0 | 162.008 | 51 | 26 | 51 | 26 | 573 | 60.9 | 6.18 | 1.397 | 1.62 | 0.862 | 0.48 | -0.48 | 0.7 | 0.23333333 | -0.59 | -0.21 | 0.259391245 | 0.163352203 | 0.392643316 | 0.923229118 | 0.869444647 | 0.896336883 | 0.93064581 |
| | High | A0A571BD | AP2-associated protein kinase 1 (Fragment) OS=Mus musculus OX=10090 GN=Aak1 PE=1 SV=1 | 0 | 35.82 | 46 | 8 | 12 | 1 | 237 | 25.2 | 8.66 | 0.766 | 0.692 | 1.106 | -0.39 | 0.39 | -0.53 | -0.1766667 | 0.46 | 0.15 | 0.259157855 | 0.203764895 | 0.532513598 | 0.923229118 | 0.912882441 | 0.918055779 | 0.96694628 |
| Raly | High | Q64012 | RNA-binding protein Raly OS=Mus musculus OX=10090 GN=Raly PE=1 SV=3 | 0 | 4.849 | 9 | 3 | 3 | 3 | 312 | 33.2 | 8.84 | 1.358 | 1.55 | 0.876 | 0.44 | -0.44 | 0.63 | 0.21 | -0.535 | -0.19 | 0.267845864 | 0.195706436 | 0.462363529 | 0.923229118 | 0.912882441 | 0.918055779 | 0.959744396 |
| Ssb | High | P32067 | Lupus La protein homolog OS=Mus musculus OX=10090 GN=Ssb PE=1 SV=1 | 0 | 23.993 | 14 | 6 | 7 | 6 | 415 | 47.7 | 9.77 | 1.38 | 1.549 | 0.891 | 0.47 | -0.47 | 0.63 | 0.21 | -0.55 | -0.17 | 0.260519863 | 0.209042184 | 0.468489654 | 0.923229118 | 0.912882441 | 0.918055779 | 0.961044326 |
| Gpx4 | High | O70325 | Phospholipid hydroperoxide glutathione peroxidase OS=Mus musculus OX=10090 GN=Gpx4 PE=1 S | 0 | 160.862 | 59 | 15 | 77 | 15 | 197 | 22.2 | 8.46 | 1.397 | 1.563 | 0.893 | 0.48 | -0.48 | 0.64 | 0.21333333 | -0.56 | -0.16 | 0.259763217 | 0.199040875 | 0.53115803 | 0.923229118 | 0.912882441 | 0.918055779 | 0.96694628 |
| | High | F8WGD9 | 1-acyl-sn-glycerol-3-phosphate acyltransferase epsilon OS=Mus musculus OX=10090 GN=Agpat5 | 0.001 | 5.089 | 4 | 1 | 1 | 1 | 291 | 33.8 | 9.2 | 1.401 | 1.526 | 0.918 | 0.49 | -0.49 | 0.61 | 0.20333333 | -0.55 | -0.12 | 0.261993412 | 0.229458166 | 0.70273014 | 0.923229118 | 0.935008675 | 0.929118896 | 0.992013705 |
| Cox6b1 | High | P56391 | Cytochrome c oxidase subunit 6B1 OS=Mus musculus OX=10090 GN=Cox6b1 PE=1 SV=2 | 0 | 35.557 | 71 | 7 | 17 | 7 | 86 | 10.1 | 8.72 | 1.397 | 1.504 | 0.929 | 0.48 | -0.48 | 0.59 | 0.19666667 | -0.535 | -0.11 | 0.267896161 | 0.243383236 | 0.707559181 | 0.923229118 | 0.940627712 | 0.931928415 | 0.992507514 |
| Hbs1l | High | Q3TGM7 | HBS1-like protein OS=Mus musculus OX=10090 GN=Hbs1l PE=1 SV=1 | 0.008 | 3.028 | 3 | 2 | 2 | 2 | 679 | 74.8 | 6.35 | 0.797 | 0.729 | 1.093 | -0.33 | 0.33 | -0.46 | -0.1533333 | 0.395 | 0.13 | 0.260366468 | 0.269258796 | 0.517439922 | 0.923229118 | 0.953285167 | 0.938257143 | 0.964378027 |
| Rab8a | High | P55258 | Ras-related protein Rab-8A OS=Mus musculus OX=10090 GN=Rab8a PE=1 SV=2 | 0 | 52.037 | 41 | 11 | 25 | 5 | 207 | 23.7 | 9.07 | 0.717 | 0.725 | 0.989 | -0.48 | 0.48 | -0.46 | -0.1533333 | 0.47 | -0.02 | 0.263863649 | 0.282846788 | 0.97588348 | 0.923229118 | 0.955820031 | 0.939524574 | 0.997221123 |
| Ppp2r2a | High | Q6P1F6 | Serine/threonine-protein phosphatase 2A 55 kDa regulatory subunit B alpha isoform OS=Mus mu | 0 | 20.682 | 13 | 5 | 8 | 3 | 447 | 51.7 | 6.2 | 0.753 | 0.734 | 1.026 | -0.41 | 0.41 | -0.45 | -0.15 | 0.43 | 0.04 | 0.266823547 | 0.300240008 | 0.909454617 | 0.923229118 | 0.955820031 | 0.939524574 | 0.992535481 |
| Gpm6a | High | P35802 | Neuronal membrane glycoprotein M6-a OS=Mus musculus OX=10090 GN=Gpm6a PE=1 SV=1 | 0 | 50.976 | 23 | 9 | 32 | 9 | 278 | 31.1 | 5.27 | 1.392 | 1.437 | 0.969 | 0.48 | -0.48 | 0.52 | 0.17333333 | -0.5 | -0.05 | 0.264288033 | 0.304635882 | 0.916338458 | 0.923229118 | 0.955820031 | 0.939524574 | 0.992535481 |
| Rab24 | High | P35290 | Ras-related protein Rab-24 OS=Mus musculus OX=10090 GN=Rab24 PE=1 SV=2 | 0 | 58.745 | 53 | 9 | 15 | 9 | 203 | 23.1 | 6.23 | 0.732 | 0.747 | 0.979 | -0.45 | 0.45 | -0.42 | -0.14 | 0.435 | -0.03 | 0.257979136 | 0.327656418 | 0.894354136 | 0.923229118 | 0.962372728 | 0.942800923 | 0.992535481 |
| Tmem222 | High | Q8BVA2 | Transmembrane protein 222 OS=Mus musculus OX=10090 GN=Tmem222 PE=1 SV=1 | 0 | 24.756 | 26 | 3 | 6 | 3 | 208 | 23.2 | 6.61 | 0.715 | 0.779 | 0.918 | -0.48 | 0.48 | -0.36 | -0.12 | 0.42 | -0.12 | 0.25734386 | 0.393654556 | 0.637252547 | 0.923229118 | 0.983884835 | 0.953556977 | 0.983597794 |
| Meig1 | High | A2AJG4 | Meiosis-expressed gene 1 protein (Fragment) OS=Mus musculus OX=10090 GN=Meig1 PE=1 SV= | 0.001 | 4.286 | 20 | 1 | 1 | 1 | 65 | 7.8 | 9.52 | 0.762 | 0.806 | 0.945 | -0.39 | 0.39 | -0.31 | -0.1033333 | 0.35 | -0.08 | 0.267264195 | 0.432804812 | 0.811429594 | 0.923229118 | 0.99399649 | 0.958612804 | 0.992535481 |
| Tnfaip3 | High | Q60769 | Tumor necrosis factor alpha-induced protein 3 OS=Mus musculus OX=10090 GN=Tnfaip3 PE=1 SV | 0 | 302.401 | 60 | 46 | 138 | 46 | 775 | 87.6 | 8.16 | 0.716 | 0.819 | 0.874 | -0.48 | 0.48 | -0.29 | -0.0966667 | 0.385 | -0.19 | 0.262687537 | 0.485445516 | 0.443751254 | 0.923229118 | 0.995559182 | 0.95939415 | 0.958608248 |
| Nap1l4 | High | B7ZNL2 | Nap1l4 protein OS=Mus musculus OX=10090 GN=Nap1l4 PE=1 SV=1 | 0 | 30.602 | 23 | 7 | 10 | 6 | 386 | 43.9 | 4.68 | 0.723 | 0.851 | 0.85 | -0.47 | 0.47 | -0.23 | -0.0766667 | 0.35 | -0.23 | 0.2661785 | 0.561196379 | 0.371549877 | 0.923229118 | 0.995559182 | 0.95939415 | 0.920164724 |
| Cadm1 | High | Q8R5M8 | Cell adhesion molecule 1 OS=Mus musculus OX=10090 GN=Cadm1 PE=1 SV=2 | 0 | 50.096 | 25 | 8 | 15 | 2 | 456 | 49.8 | 5.03 | 0.712 | 0.895 | 0.795 | -0.49 | 0.49 | -0.16 | -0.0533333 | 0.325 | -0.33 | 0.265368225 | 0.670640166 | 0.167676357 | 0.923229118 | 0.995559182 | 0.95939415 | 0.738806217 |
| Ppm1a | High | Q9EQE2 | Protein-serine/threonine phosphatase OS=Mus musculus OX=10090 GN=Ppm1a PE=1 SV=1 | 0 | 25.629 | 17 | 5 | 6 | 3 | 323 | 35.9 | 5.67 | 0.75 | 0.858 | 0.875 | -0.41 | 0.41 | -0.22 | -0.0733333 | 0.315 | -0.19 | 0.266835922 | 0.578800069 | 0.409053328 | 0.923229118 | 0.995559182 | 0.95939415 | 0.936397673 |
| Pcdhgb2 | High | Q91XX7 | Protocadherin gamma B2 OS=Mus musculus OX=10090 GN=Pcdhgb2 PE=1 SV=1 | 0 | 10.306 | 4 | 3 | 3 | 1 | 928 | 100.8 | 4.97 | 0.792 | 0.838 | 0.945 | -0.34 | 0.34 | -0.25 | -0.0833333 | 0.295 | -0.08 | 0.266502576 | 0.514804573 | 0.759302414 | 0.923229118 | 0.995559182 | 0.95939415 | 0.992535481 |
| Tpd52 | High | F8WHQ1 | Tumor protein D52 OS=Mus musculus OX=10090 GN=Tpd52 PE=1 SV=1 | 0 | 36.285 | 31 | 6 | 8 | 6 | 247 | 26.9 | 4.96 | 0.753 | 0.892 | 0.845 | -0.41 | 0.41 | -0.17 | -0.0566667 | 0.29 | -0.24 | 0.25991582 | 0.661396851 | 0.350206175 | 0.923229118 | 0.995559182 | 0.95939415 | 0.909466648 |
| Vtn | High | P29788 | Vitronectin OS=Mus musculus OX=10090 GN=Vtn PE=1 SV=2 | 0 | 28.765 | 18 | 7 | 8 | 7 | 478 | 54.8 | 5.88 | 0.746 | 0.928 | 0.804 | -0.42 | 0.42 | -0.11 | -0.0366667 | 0.265 | -0.32 | 0.257964929 | 0.752221717 | 0.168430739 | 0.923229118 | 0.995559182 | 0.95939415 | 0.738806217 |
| | High | J3QNM3 | Protein enabled homolog OS=Mus musculus OX=10090 GN=Enah PE=1 SV=1 | 0 | 26.379 | 14 | 9 | 10 | 9 | 789 | 84.5 | 7.52 | 1.338 | 1.146 | 1.167 | 0.42 | -0.42 | 0.2 | 0.06666667 | -0.31 | 0.22 | 0.262072933 | 0.739333308 | 0.371334077 | 0.923229118 | 0.995559182 | 0.95939415 | 0.920164724 |
| | High | Q5M8N4 | Epimerase family protein SDR39U1 OS=Mus musculus OX=10090 GN=Sdr39u1 PE=1 SV=2 | 0 | 12.526 | 19 | 5 | 5 | 5 | 293 | 31.4 | 9.31 | 1.366 | 1.21 | 1.129 | 0.45 | -0.45 | 0.27 | 0.09 | -0.36 | 0.18 | 0.259028907 | 0.61848724 | 0.475306123 | 0.923229118 | 0.995559182 | 0.95939415 | 0.962227984 |
| Rpl38; Rpl | High | Q9JJI8 | 60S ribosomal protein L38 OS=Mus musculus OX=10090 GN=Rpl38 PE=1 SV=3 | 0 | 16.814 | 51 | 4 | 8 | 4 | 70 | 8.2 | 10.1 | 1.395 | 1.198 | 1.165 | 0.48 | -0.48 | 0.26 | 0.08666667 | -0.37 | 0.22 | 0.262226419 | 0.639654258 | 0.366275095 | 0.923229118 | 0.995559182 | 0.95939415 | 0.920164724 |
| Rpl23; LO | High | P62830 | 60S ribosomal protein L23 OS=Mus musculus OX=10090 GN=Rpl23 PE=1 SV=1 | 0 | 28.508 | 55 | 7 | 11 | 7 | 140 | 14.9 | 10.51 | 1.371 | 1.214 | 1.129 | 0.46 | -0.46 | 0.28 | 0.09333333 | -0.37 | 0.18 | 0.260157378 | 0.610286874 | 0.374104431 | 0.923229118 | 0.995559182 | 0.95939415 | 0.920164724 |
| Rpl34-ps1 | High | Q9D1R9 | 60S ribosomal protein L34 OS=Mus musculus OX=10090 GN=Rpl34 PE=1 SV=2 | 0 | 25.492 | 43 | 7 | 17 | 7 | 117 | 13.3 | 11.47 | 1.393 | 1.206 | 1.155 | 0.48 | -0.48 | 0.27 | 0.09 | -0.375 | 0.21 | 0.263538346 | 0.624876594 | 0.304006123 | 0.923229118 | 0.995559182 | 0.95939415 | 0.900241969 |
| Rpl5 | High | P47962 | 60S ribosomal protein L5 OS=Mus musculus OX=10090 GN=Rpl5 PE=1 SV=3 | 0 | 63.12 | 42 | 13 | 24 | 13 | 297 | 34.4 | 9.77 | 1.4 | 1.222 | 1.146 | 0.49 | -0.49 | 0.29 | 0.09666667 | -0.39 | 0.2 | 0.256436929 | 0.597433401 | 0.328181726 | 0.923229118 | 0.995559182 | 0.95939415 | 0.905686588 |
| | High | A0A1W2P | 40S ribosomal protein S12 OS=Mus musculus OX=10090 GN=Rps12 PE=1 SV=1 | 0 | 50.003 | 49 | 8 | 14 | 8 | 148 | 16 | 8.16 | 1.365 | 1.255 | 1.088 | 0.45 | -0.45 | 0.33 | 0.11 | -0.39 | 0.12 | 0.25728596 | 0.540829495 | 0.520874516 | 0.923229118 | 0.995559182 | 0.95939415 | 0.964378027 |
| Them4 | Medium | Q3UUI3 | Acyl-coenzyme A thioesterase THEM4 OS=Mus musculus OX=10090 GN=Them4 PE=1 SV=1 | 0.014 | 2.245 | 5 | 1 | 1 | 1 | 230 | 26 | 9.64 | 1.36 | 1.264 | 1.076 | 0.44 | -0.44 | 0.34 | 0.11333333 | -0.39 | 0.11 | 0.267376428 | 0.530995308 | 0.59579644 | 0.923229118 | 0.995559182 | 0.95939415 | 0.977451587 |
| Rps25 | High | P62852 | 40S ribosomal protein S25 OS=Mus musculus OX=10090 GN=Rps25 PE=1 SV=1 | 0 | 15.318 | 36 | 6 | 10 | 6 | 125 | 13.7 | 10.11 | 1.397 | 1.249 | 1.119 | 0.48 | -0.48 | 0.32 | 0.10666667 | -0.4 | 0.16 | 0.263920005 | 0.551521409 | 0.421715162 | 0.923229118 | 0.995559182 | 0.95939415 | 0.939192266 |
| Rab3a | High | P63011 | Ras-related protein Rab-3A OS=Mus musculus OX=10090 GN=Rab3a PE=1 SV=1 | 0 | 86.885 | 54 | 10 | 38 | 7 | 220 | 25 | 5.03 | 0.715 | 0.955 | 0.749 | -0.48 | 0.48 | -0.07 | -0.0233333 | 0.275 | -0.42 | 0.260376892 | 0.820894588 | 0.077343715 | 0.923229118 | 0.998402594 | 0.960815856 | 0.529506969 |
| Sh3d21 | Medium | Z4YK11 | SH3 domain-containing protein 21 OS=Mus musculus OX=10090 GN=Sh3d21 PE=1 SV=1 | 0.041 | 1.655 | 1 | 1 | 23 | 1 | 665 | 73.7 | 8.28 | 0.719 | 1.024 | 0.702 | -0.48 | 0.48 | 0.03 | 0.01 | 0.225 | -0.51 | 0.267868707 | 0.989793173 | 0.02914112 | 0.923229118 | 0.998402594 | 0.960815856 | 0.324925684 |
| Grn | High | P28798 | Progranulin OS=Mus musculus OX=10090 GN=Grn PE=1 SV=2 | 0 | 22.733 | 7 | 3 | 5 | 3 | 589 | 63.4 | 6.8 | 0.763 | 1.068 | 0.715 | -0.39 | 0.39 | 0.09 | 0.03 | 0.15 | -0.48 | 0.266329789 | 0.907896454 | 0.039112485 | 0.923229118 | 0.998402594 | 0.960815856 | 0.380996077 |
| Nxnl2 | High | Q9D531 | Nucleoredoxin-like protein 2 OS=Mus musculus OX=10090 GN=Nxnl2 PE=1 SV=1 | 0 | 16.466 | 44 | 5 | 6 | 5 | 156 | 17.6 | 7.34 | 1.364 | 0.994 | 1.372 | 0.45 | -0.45 | -0.01 | -0.0033333 | -0.22 | 0.46 | 0.262344783 | 0.917612292 | 0.061973361 | 0.923229118 | 0.998402594 | 0.960815856 | 0.476659309 |
| Tdrd7 | High | Q8K1H1 | Tudor domain-containing protein 7 OS=Mus musculus OX=10090 GN=Tdrd7 PE=1 SV=1 | 0 | 54.688 | 17 | 17 | 19 | 17 | 1086 | 122.1 | 6.7 | 1.393 | 1.07 | 1.302 | 0.48 | -0.48 | 0.1 | 0.03333333 | -0.29 | 0.38 | 0.263511246 | 0.90426285 | 0.071444268 | 0.923229118 | 0.998402594 | 0.960815856 | 0.508467979 |
| Idh1 | High | O88844 | Isocitrate dehydrogenase [NADP] cytoplasmic OS=Mus musculus OX=10090 GN=Idh1 PE=1 SV=2 | 0 | 210.14 | 72 | 32 | 80 | 30 | 414 | 46.6 | 7.17 | 1.398 | 1.107 | 1.263 | 0.48 | -0.48 | 0.15 | 0.05 | -0.315 | 0.34 | 0.257778765 | 0.821196534 | 0.108186341 | 0.923229118 | 0.998402594 | 0.960815856 | 0.630562179 |
| 2610029G | High | Q9D0B6 | Protein PBDC1 OS=Mus musculus OX=10090 GN=Pbdc1 PE=1 SV=1 | 0 | 9.86 | 16 | 2 | 2 | 2 | 198 | 22.2 | 4.55 | 0.767 | 0.543 | 1.411 | -0.38 | 0.38 | -0.88 | -0.2933333 | 0.63 | 0.5 | 0.268529618 | 0.045847549 | 0.050186424 | 0.923593314 | 0.520047141 | 0.721820228 | 0.427566646 |
| Mrps36 | High | Q9CQX8 | 28S ribosomal protein S36, mitochondrial OS=Mus musculus OX=10090 GN=Mrps36 PE=1 SV=1 | 0 | 17.098 | 49 | 3 | 3 | 3 | 102 | 11.1 | 9.99 | 1.349 | 1.422 | 0.949 | 0.43 | -0.43 | 0.51 | 0.17 | -0.47 | -0.08 | 0.269154891 | 0.320258207 | 0.717577482 | 0.923593314 | 0.955820031 | 0.939706672 | 0.992535481 |
| Cox7a2 | High | P48771 | Cytochrome c oxidase subunit 7A2, mitochondrial OS=Mus musculus OX=10090 GN=Cox7a2 PE=1 | 0 | 15.419 | 28 | 2 | 7 | 2 | 83 | 9.3 | 10.27 | 1.357 | 1.429 | 0.949 | 0.44 | -0.44 | 0.52 | 0.17333333 | -0.48 | -0.08 | 0.269011 | 0.312806768 | 0.724532374 | 0.923593314 | 0.955820031 | 0.939706672 | 0.992535481 |
| Eef1b2 | High | O70251 | Elongation factor 1-beta OS=Mus musculus OX=10090 GN=Eef1b PE=1 SV=5 | 0 | 23.509 | 31 | 8 | 11 | 8 | 225 | 24.7 | 4.69 | 1.365 | 1.32 | 1.034 | 0.45 | -0.45 | 0.4 | 0.13333333 | -0.425 | 0.05 | 0.269022037 | 0.443483608 | 0.751220229 | 0.923593314 | 0.994630833 | 0.959112074 | 0.992535481 |
| Rpl31 | High | P62900 | 60S ribosomal protein L31 OS=Mus musculus OX=10090 GN=Rpl31 PE=1 SV=1 | 0 | 19.652 | 30 | 4 | 8 | 4 | 125 | 14.5 | 10.54 | 1.385 | 1.21 | 1.145 | 0.47 | -0.47 | 0.27 | 0.09 | -0.37 | 0.19 | 0.268646414 | 0.617936357 | 0.387783403 | 0.923593314 | 0.995559182 | 0.959576248 | 0.93064581 |
| Pik3r2 | Medium | O08908 | Phosphatidylinositol 3-kinase regulatory subunit beta OS=Mus musculus OX=10090 GN=Pik3r2 PE | 0.01 | 2.701 | 1 | 1 | 1 | 1 | 722 | 81.2 | 6.07 | 1.356 | 1.126 | 1.205 | 0.44 | -0.44 | 0.17 | 0.05666667 | -0.305 | 0.27 | 0.269070355 | 0.773476935 | 0.275208676 | 0.923593314 | 0.996297996 | 0.959945655 | 0.876067403 |
| Prps1l3 | High | G3UXL2 | Ribose-phosphate diphosphokinase OS=Mus musculus OX=10090 GN=Prps1l3 PE=3 SV=1 | 0 | 48.456 | 38 | 11 | 19 | 5 | 318 | 34.8 | 6.98 | 0.72 | 0.701 | 1.027 | -0.47 | 0.47 | -0.51 | -0.17 | 0.49 | 0.04 | 0.269910373 | 0.239408625 | 0.78521698 | 0.924097568 | 0.937938091 | 0.931017829 | 0.992535481 |
| Tmod2 | High | Q9JKK7 | Tropomodulin-2 OS=Mus musculus OX=10090 GN=Tmod2 PE=1 SV=2 | 0 | 22.752 | 23 | 6 | 6 | 6 | 351 | 39.5 | 5.35 | 0.769 | 0.814 | 0.945 | -0.38 | 0.38 | -0.3 | -0.1 | 0.34 | -0.08 | 0.269568143 | 0.472555319 | 0.692723284 | 0.924097568 | 0.995559182 | 0.959828375 | 0.991285312 |
| Lmnb1 | High | P14733 | Lamin-B1 OS=Mus musculus OX=10090 GN=Lmnb1 PE=1 SV=3 | 0 | 27.535 | 13 | 7 | 7 | 7 | 588 | 66.7 | 5.16 | 1.304 | 1.115 | 1.17 | 0.38 | -0.38 | 0.16 | 0.05333333 | -0.27 | 0.23 | 0.269961087 | 0.804658223 | 0.368499331 | 0.924097568 | 0.998402594 | 0.961250081 | 0.920164724 |
| Pcp4l1 | High | Q6W8Q3 | Purkinje cell protein 4-like protein 1 OS=Mus musculus OX=10090 GN=Pcp4l1 PE=1 SV=1 | 0.002 | 4.15 | 22 | 2 | 2 | 2 | 68 | 7.5 | 5.52 | 0.77 | 1.047 | 0.736 | -0.38 | 0.38 | 0.07 | 0.02333333 | 0.155 | -0.44 | 0.270375913 | 0.956781987 | 0.065182484 | 0.924952862 | 0.998402594 | 0.961677728 | 0.491746452 |
| 6230416C( | Low | A2ARV7 | Predicted gene 14401 OS=Mus musculus OX=10090 GN=Gm14401 PE=1 SV=1 | 0.055 | 1.382 | 3 | 1 | 1 | 1 | 303 | 35 | 8.81 | 0.768 | 0.797 | 0.963 | -0.38 | 0.38 | -0.33 | -0.11 | 0.355 | -0.05 | 0.272179482 | 0.428786004 | 0.860712092 | 0.930555096 | 0.99399649 | 0.962275793 | 0.992535481 |
| Gltp | High | Q9JL62 | Glycolipid transfer protein OS=Mus musculus OX=10090 GN=Gltp PE=1 SV=3 | 0 | 7.986 | 13 | 3 | 3 | 3 | 209 | 23.7 | 7.39 | 0.718 | 0.518 | 1.387 | -0.48 | 0.48 | -0.95 | -0.3166667 | 0.715 | 0.47 | 0.280346995 | 0.035350517 | 0.044519384 | 0.933919361 | 0.476392832 | 0.705156097 | 0.405968973 |
| Prom1 | High | Q8R056 | Prom1 protein OS=Mus musculus OX=10090 GN=Prom1 PE=1 SV=1 | 0 | 283.736 | 38 | 32 | 123 | 1 | 842 | 94.4 | 6.9 | 1.383 | 2.028 | 0.682 | 0.47 | -0.47 | 1.02 | 0.34 | -0.745 | -0.55 | 0.274156745 | 0.037244711 | 0.017855279 | 0.933919361 | 0.480071477 | 0.706995419 | 0.236677417 |
| Ulk3 | High | Q3U3Q1 | Serine/threonine-protein kinase ULK3 OS=Mus musculus OX=10090 GN=Ulk3 PE=2 SV=1 | 0 | 18.604 | 14 | 5 | 5 | 5 | 472 | 53.5 | 7.03 | 0.77 | 0.529 | 1.455 | -0.38 | 0.38 | -0.92 | -0.3066667 | 0.65 | 0.54 | 0.289601267 | 0.041493293 | 0.029391488 | 0.933919361 | 0.503577691 | 0.718748526 | 0.32568789 |
| Blmh | High | Q8R016 | Bleomycin hydrolase OS=Mus musculus OX=10090 GN=Blmh PE=1 SV=1 | 0 | 32.467 | 17 | 7 | 11 | 7 | 455 | 52.5 | 6.48 | 0.728 | 0.548 | 1.329 | -0.46 | 0.46 | -0.87 | -0.29 | 0.665 | 0.41 | 0.285917732 | 0.053468376 | 0.076669768 | 0.933919361 | 0.556350595 | 0.745134978 | 0.526177957 |
| Guk1 | High | Q64520 | Guanylate kinase OS=Mus musculus OX=10090 GN=Guk1 PE=1 SV=2 | 0 | 38.903 | 43 | 10 | 16 | 10 | 198 | 21.9 | 6.55 | 0.721 | 0.587 | 1.228 | -0.47 | 0.47 | -0.77 | -0.2566667 | 0.62 | 0.3 | 0.299220774 | 0.085828061 | 0.194581761 | 0.933919361 | 0.68454712 | 0.80923324 | 0.784517504 |

| | | | | | | | | | | | | | | | | | | | | | | | | | | | | |
|---|---|---|---|---|---|---|---|---|---|---|---|---|---|---|---|---|---|---|---|---|---|---|---|---|---|---|---|---|
| Gng11 | High | P61953 | Guanine nucleotide-binding protein G(I)/G(S)/G(O) subunit gamma-11 OS=Mus musculus OX=10( | 0 | 25.766 | 47 | 5 | 105 | 1 | 73 | 8.5 | 5.54 | 1.362 | 1.771 | 0.769 | 0.45 | -0.45 | 0.82 | 0.27333333 | -0.635 | -0.38 | 0.280135674 | 0.085973475 | 0.113654494 | 0.933919361 | 0.68454712 | 0.80923324 | 0.643956085 |
| Nedd4 | High | P46935 | E3 ubiquitin-protein ligase NEDD4 OS=Mus musculus OX=10090 GN=Nedd4 PE=1 SV=3 | 0 | 68.452 | 25 | 20 | 24 | 18 | 887 | 102.6 | 5.26 | 0.725 | 0.592 | 1.224 | -0.46 | 0.46 | -0.76 | -0.2533333 | 0.61 | 0.29 | 0.280373092 | 0.090513906 | 0.160686879 | 0.933919361 | 0.696517952 | 0.815218656 | 0.733684585 |
| Osbp | High | Q3B7Z2 | Oxysterol-binding protein 1 OS=Mus musculus OX=10090 GN=Osbp PE=1 SV=3 | 0 | 17.648 | 9 | 5 | 5 | 4 | 805 | 88.7 | 7.2 | 0.757 | 0.592 | 1.28 | -0.4 | 0.4 | -0.76 | -0.2533333 | 0.58 | 0.36 | 0.284224049 | 0.089918765 | 0.116794326 | 0.933919361 | 0.696517952 | 0.815218656 | 0.652907063 |
| Ush2a | High | Q2QI47 | Usherin OS=Mus musculus OX=10090 GN=Ush2A PE=1 SV=2 | 0 | 9.324 | 1 | 4 | 4 | 4 | 5193 | 569.3 | 6.81 | 1.373 | 1.741 | 0.789 | 0.46 | -0.46 | 0.8 | 0.26666667 | -0.63 | -0.34 | 0.275088322 | 0.101134597 | 0.145472939 | 0.933919361 | 0.730697757 | 0.832308559 | 0.706909771 |
| Myh9 | High | Q8VDD5 | Myosin-9 OS=Mus musculus OX=10090 GN=Myh9 PE=1 SV=4 | 0 | 69.646 | 12 | 20 | 24 | 13 | 1960 | 226.2 | 5.66 | 0.746 | 0.629 | 1.187 | -0.42 | 0.42 | -0.67 | -0.2233333 | 0.545 | 0.25 | 0.293474586 | 0.131084585 | 0.313621411 | 0.933919361 | 0.805028768 | 0.869474065 | 0.904943958 |
| Vbp1 | High | P61759 | Prefoldin subunit 3 OS=Mus musculus OX=10090 GN=Vbp1 PE=1 SV=2 | 0 | 15.013 | 28 | 6 | 7 | 6 | 196 | 22.4 | 6.28 | 0.769 | 0.63 | 1.221 | -0.38 | 0.38 | -0.67 | -0.2233333 | 0.525 | 0.29 | 0.297301943 | 0.132673916 | 0.261461816 | 0.933919361 | 0.808473523 | 0.871196442 | 0.858323421 |
| Arl8b | High | Q9CQW2 | ADP-ribosylation factor-like protein 8B OS=Mus musculus OX=10090 GN=Arl8b PE=1 SV=1 | 0 | 103.361 | 59 | 10 | 30 | 5 | 186 | 21.5 | 8.43 | 1.411 | 1.65 | 0.855 | 0.5 | -0.5 | 0.72 | 0.24 | -0.61 | -0.23 | 0.275273725 | 0.146874305 | 0.363405722 | 0.933919361 | 0.844640233 | 0.889279797 | 0.91781256 |
| Rdh11 | Medium | Q9R1R8 | Retinol dehydrogenase 11 OS=Mus musculus OX=10090 GN=Rdh11 PE=1 SV=1 | 0.01 | 2.693 | 4 | 1 | 1 | 1 | 300 | 33.2 | 8.79 | 1.335 | 1.623 | 0.823 | 0.42 | -0.42 | 0.7 | 0.23333333 | -0.56 | -0.28 | 0.284545404 | 0.151370462 | 0.275482343 | 0.933919361 | 0.847684847 | 0.890802104 | 0.876067403 |
| Bdh1 | High | Q80XN0 | D-beta-hydroxybutyrate dehydrogenase, mitochondrial OS=Mus musculus OX=10090 GN=Bdh1 PE | 0 | 15.87 | 9 | 3 | 5 | 3 | 343 | 38.3 | 9.01 | 1.35 | 1.633 | 0.827 | 0.43 | -0.43 | 0.71 | 0.23666667 | -0.57 | -0.27 | 0.284707644 | 0.156207635 | 0.262541024 | 0.933919361 | 0.847684847 | 0.890802104 | 0.859470827 |
| Vwa8 | High | Q8CC88 | von Willebrand factor A domain-containing protein 8 OS=Mus musculus OX=10090 GN=Vwa8 PE= | 0 | 4.906 | 1 | 1 | 1 | 1 | 1905 | 213.3 | 6.6 | 1.327 | 1.593 | 0.833 | 0.41 | -0.41 | 0.67 | 0.22333333 | -0.54 | -0.26 | 0.277136042 | 0.169078785 | 0.320947706 | 0.933919361 | 0.874947902 | 0.904433631 | 0.905686588 |
| Hectd1 | High | Q69ZR2 | E3 ubiquitin-protein ligase HECTD1 OS=Mus musculus OX=10090 GN=Hectd1 PE=1 SV=2 | 0 | 15.955 | 2 | 3 | 3 | 3 | 2618 | 289.9 | 5.41 | 0.803 | 0.671 | 1.196 | -0.32 | 0.32 | -0.58 | -0.1933333 | 0.45 | 0.26 | 0.281958728 | 0.171715651 | 0.245746024 | 0.933919361 | 0.875484829 | 0.904702095 | 0.840125423 |
| Tmx2 | High | Q9D710 | Thioredoxin-related transmembrane protein 2 OS=Mus musculus OX=10090 GN=Tmx2 PE=1 SV=1 | 0.001 | 6.77 | 13 | 4 | 4 | 4 | 295 | 33.9 | 8.75 | 1.292 | 1.582 | 0.817 | 0.37 | -0.37 | 0.66 | 0.22 | -0.515 | -0.29 | 0.297501871 | 0.186623567 | 0.232618021 | 0.933919361 | 0.900299151 | 0.917109256 | 0.820823943 |
| | High | A0A2I3BR | Bcl-2-like protein 2 (Fragment) OS=Mus musculus OX=10090 GN=Bcl2l2 PE=3 SV=1 | 0 | 21.159 | 44 | 3 | 5 | 3 | 117 | 12.6 | 5.9 | 1.36 | 1.582 | 0.86 | 0.44 | -0.44 | 0.66 | 0.22 | -0.55 | -0.22 | 0.289458411 | 0.186739417 | 0.418471578 | 0.933919361 | 0.900299151 | 0.917109256 | 0.936674706 |
| Nedd8 | High | P29595 | NEDD8 OS=Mus musculus OX=10090 GN=Nedd8 PE=1 SV=2 | 0 | 10.731 | 17 | 2 | 7 | 2 | 81 | 9 | 7.25 | 0.735 | 0.677 | 1.086 | -0.44 | 0.44 | -0.56 | -0.1866667 | 0.5 | 0.12 | 0.274698295 | 0.199372767 | 0.615371261 | 0.933919361 | 0.912882441 | 0.923400901 | 0.97826833 |
| Abcc4 | Low | E9Q467 | ATP-binding cassette, sub-family C (CFTR/MRP), member 4 OS=Mus musculus OX=10090 GN=Ab | 0.053 | 1.42 | 1 | 1 | 1 | 1 | 1250 | 140.1 | 8.29 | 1.324 | 1.543 | 0.858 | 0.4 | -0.4 | 0.63 | 0.21 | -0.515 | -0.22 | 0.283956115 | 0.202711723 | 0.412920063 | 0.933919361 | 0.912882441 | 0.923400901 | 0.936397673 |
| Smad2 | High | Q62432 | Mothers against decapentaplegic homolog 2 OS=Mus musculus OX=10090 GN=Smad2 PE=1 SV=2 | 0 | 8.499 | 4 | 2 | 2 | 2 | 467 | 52.2 | 6.58 | 1.355 | 1.56 | 0.869 | 0.44 | -0.44 | 0.64 | 0.21333333 | -0.54 | -0.2 | 0.298135343 | 0.195418936 | 0.399226707 | 0.933919361 | 0.912882441 | 0.923400901 | 0.93262326 |
| Akna | High | Q80VW7 | Microtubule organization protein AKNA OS=Mus musculus OX=10090 GN=Akna PE=1 SV=1 | 0 | 9.662 | 3 | 4 | 5 | 4 | 1404 | 153 | 6.19 | 1.363 | 1.529 | 0.891 | 0.45 | -0.45 | 0.61 | 0.20333333 | -0.53 | -0.17 | 0.286553312 | 0.212173018 | 0.50808085 | 0.933919361 | 0.913066452 | 0.923492906 | 0.964378027 |
| Slc25a4 | High | P48962 | ADP/ATP translocase 1 OS=Mus musculus OX=10090 GN=Slc25a4 PE=1 SV=4 | 0 | 139.043 | 53 | 19 | 77 | 10 | 298 | 32.9 | 9.72 | 1.365 | 1.545 | 0.883 | 0.45 | -0.45 | 0.63 | 0.21 | -0.54 | -0.18 | 0.293750444 | 0.211608119 | 0.48396328 | 0.933919361 | 0.913066452 | 0.923492906 | 0.964378027 |
| Ndufs2 | High | Q91WD5 | NADH dehydrogenase [ubiquinone] iron-sulfur protein 2, mitochondrial OS=Mus musculus OX=10( | 0 | 34.381 | 21 | 8 | 8 | 8 | 463 | 52.6 | 6.99 | 1.375 | 1.546 | 0.889 | 0.46 | -0.46 | 0.63 | 0.21 | -0.545 | -0.17 | 0.294357666 | 0.211505018 | 0.515504992 | 0.933919361 | 0.913066452 | 0.923492906 | 0.964378027 |
| Lrba | High | E9Q3Y4 | Lipopolysaccharide-responsive and beige-like anchor protein OS=Mus musculus OX=10090 GN=Lrb | 0 | 20.618 | 2 | 6 | 6 | 4 | 2854 | 316.6 | 5.54 | 0.772 | 0.69 | 1.119 | -0.37 | 0.37 | -0.53 | -0.1766667 | 0.45 | 0.16 | 0.287008605 | 0.221493244 | 0.511364622 | 0.933919361 | 0.928185815 | 0.931052588 | 0.964378027 |
| Tomm5 | Medium | B1AXP6 | Mitochondrial import receptor subunit TOM5 homolog OS=Mus musculus OX=10090 GN=Tomm5 | 0.048 | 1.517 | 14 | 1 | 1 | 1 | 51 | 6 | 9.6 | 1.368 | 1.516 | 0.902 | 0.45 | -0.45 | 0.6 | 0.2 | -0.525 | -0.15 | 0.282157688 | 0.222689896 | 0.557429153 | 0.933919361 | 0.931112786 | 0.932516073 | 0.971568588 |
| Lactb2 | High | Q99KR3 | Endoribonuclease LACTB2 OS=Mus musculus OX=10090 GN=Lactb2 PE=1 SV=1 | 0.001 | 4.632 | 6 | 1 | 1 | 1 | 288 | 32.7 | 6.33 | 0.794 | 0.705 | 1.126 | -0.33 | 0.33 | -0.5 | -0.1666667 | 0.415 | 0.17 | 0.292411902 | 0.226353206 | 0.440180828 | 0.933919361 | 0.935008675 | 0.934464018 | 0.954775204 |
| Uqcrc2 | High | Q9DB77 | Cytochrome b-c1 complex subunit 2, mitochondrial OS=Mus musculus OX=10090 GN=Uqcrc2 PE=1 | 0 | 90.318 | 41 | 17 | 32 | 17 | 453 | 48.2 | 9.25 | 1.377 | 1.527 | 0.902 | 0.46 | -0.46 | 0.61 | 0.20333333 | -0.535 | -0.15 | 0.280619979 | 0.225470613 | 0.571013331 | 0.933919361 | 0.935008675 | 0.934464018 | 0.972910028 |
| Armc1 | Medium | Q9D7A8 | Armadillo repeat-containing protein 1 OS=Mus musculus OX=10090 GN=Armc1 PE=1 SV=1 | 0.011 | 2.513 | 3 | 1 | 1 | 1 | 282 | 31.2 | 5.57 | 1.364 | 1.508 | 0.905 | 0.45 | -0.45 | 0.59 | 0.19666667 | -0.52 | -0.14 | 0.282355807 | 0.240649852 | 0.590272914 | 0.933919361 | 0.938986585 | 0.936452973 | 0.974289145 |
| Uqcrq | High | Q9CQ69 | Cytochrome b-c1 complex subunit 8 OS=Mus musculus OX=10090 GN=Uqcrq PE=1 SV=3 | 0 | 26.108 | 38 | 3 | 7 | 3 | 82 | 9.8 | 10.26 | 1.358 | 1.503 | 0.903 | 0.44 | -0.44 | 0.59 | 0.19666667 | -0.515 | -0.15 | 0.298769837 | 0.244097578 | 0.588161224 | 0.933919361 | 0.94065644 | 0.9372879 | 0.973095093 |
| Dlat | High | Q8BMF4 | Dihydrolipoyllysine-residue acetyltransferase component of pyruvate dehydrogenase complex, mit | 0 | 33.154 | 14 | 7 | 10 | 7 | 642 | 67.9 | 8.57 | 1.399 | 1.493 | 0.937 | 0.48 | -0.48 | 0.58 | 0.19333333 | -0.53 | -0.09 | 0.282979762 | 0.252681334 | 0.699989194 | 0.933919361 | 0.950034749 | 0.941977055 | 0.991285312 |
| Prps2 | High | Q9CS42 | Ribose-phosphate pyrophosphokinase 2 OS=Mus musculus OX=10090 GN=Prps2 PE=1 SV=4 | 0 | 20.972 | 22 | 7 | 9 | 1 | 318 | 34.8 | 6.61 | 0.78 | 0.721 | 1.082 | -0.36 | 0.36 | -0.47 | -0.1566667 | 0.415 | 0.11 | 0.276534885 | 0.261585695 | 0.626788332 | 0.933919361 | 0.951072422 | 0.942495892 | 0.980381496 |
| U2af1 | High | Q9D883 | Splicing factor U2AF 35 kDa subunit OS=Mus musculus OX=10090 GN=U2af1 PE=1 SV=4 | 0.005 | 3.532 | 9 | 2 | 2 | 2 | 239 | 27.8 | 8.81 | 1.331 | 1.468 | 0.907 | 0.41 | -0.41 | 0.55 | 0.18333333 | -0.48 | -0.14 | 0.275868754 | 0.261281538 | 0.594599493 | 0.933919361 | 0.951072422 | 0.942495892 | 0.977451587 |
| Stt3a | Medium | P46978 | Dolichyl-diphosphooligosaccharide--protein glycosyltransferase subunit STT3A OS=Mus musculus | 0.03 | 1.88 | 1 | 1 | 1 | 1 | 705 | 80.5 | 8.1 | 1.339 | 1.47 | 0.911 | 0.42 | -0.42 | 0.56 | 0.18666667 | -0.49 | -0.13 | 0.274728051 | 0.260017822 | 0.60819983 | 0.933919361 | 0.951072422 | 0.942495892 | 0.97826833 |
| Glod4 | High | Q9CPV4 | Glyoxalase domain-containing protein 4 OS=Mus musculus OX=10090 GN=Glod4 PE=1 SV=1 | 0 | 48 | 46 | 12 | 16 | 12 | 298 | 33.3 | 5.47 | 0.754 | 0.717 | 1.052 | -0.41 | 0.41 | -0.48 | -0.16 | 0.445 | 0.07 | 0.280933337 | 0.267950042 | 0.776690993 | 0.933919361 | 0.953285167 | 0.943602264 | 0.992535481 |
| Madd | High | A0A0R4J1 | MAP kinase-activating death domain protein OS=Mus musculus OX=10090 GN=Madd PE=1 SV=1 | 0 | 34.11 | 7 | 11 | 11 | 11 | 1558 | 173.1 | 6.2 | 0.745 | 0.727 | 1.023 | -0.43 | 0.43 | -0.46 | -0.1533333 | 0.445 | 0.03 | 0.290747933 | 0.28841288 | 0.911439414 | 0.933919361 | 0.955820031 | 0.944869696 | 0.992535481 |
| | Medium | D3Z710 | Kinesin light chain OS=Mus musculus OX=10090 GN=Klc3 PE=1 SV=1 | 0.035 | 1.788 | 2 | 1 | 1 | 1 | 505 | 55.6 | 6.77 | 0.776 | 0.739 | 1.05 | -0.37 | 0.37 | -0.44 | -0.1466667 | 0.405 | 0.07 | 0.298247111 | 0.29011601 | 0.704751095 | 0.933919361 | 0.955820031 | 0.944869696 | 0.992485323 |
| | High | A0A2R8VH | Mitochondrial import receptor subunit TOM22 homolog OS=Mus musculus OX=10090 GN=Tomm | 0 | 10.568 | 40 | 2 | 3 | 2 | 89 | 9.9 | 9.98 | 1.341 | 1.413 | 0.949 | 0.42 | -0.42 | 0.5 | 0.16666667 | -0.46 | -0.08 | 0.285410672 | 0.314227894 | 0.772290075 | 0.933919361 | 0.955820031 | 0.944869696 | 0.992535481 |
| Pck2 | High | A0A0R4J0 | Phosphoenolpyruvate carboxykinase (GTP) OS=Mus musculus OX=10090 GN=Pck2 PE=1 SV=1 | 0.008 | 2.957 | 2 | 1 | 1 | 1 | 667 | 73.4 | 7.74 | 1.339 | 1.421 | 0.942 | 0.42 | -0.42 | 0.51 | 0.17 | -0.465 | -0.09 | 0.290079451 | 0.308829996 | 0.733400741 | 0.933919361 | 0.955820031 | 0.944869696 | 0.992535481 |
| Bnip2 | High | G5E8U9 | BCL2/adenovirus E1B 19 kDa protein-interacting protein 2 OS=Mus musculus OX=10090 GN=Bnip | 0.001 | 5.47 | 6 | 2 | 2 | 2 | 348 | 39.8 | 5.22 | 1.336 | 1.423 | 0.938 | 0.42 | -0.42 | 0.51 | 0.17 | -0.465 | -0.09 | 0.292095097 | 0.318885166 | 0.752710002 | 0.933919361 | 0.955820031 | 0.944869696 | 0.992535481 |
| Nqo1 | High | Q64669 | NAD(P)H dehydrogenase [quinone] 1 OS=Mus musculus OX=10090 GN=Nqo1 PE=1 SV=3 | 0 | 45.857 | 36 | 12 | 22 | 12 | 274 | 30.9 | 8.72 | 1.375 | 1.458 | 0.943 | 0.46 | -0.46 | 0.54 | 0.18 | -0.5 | -0.08 | 0.282428946 | 0.28452751 | 0.782433875 | 0.933919361 | 0.955820031 | 0.944869696 | 0.992535481 |
| | High | H7BWX9 | Small ubiquitin-related modifier 2 OS=Mus musculus OX=10090 GN=Sumo2 PE=1 SV=1 | 0 | 9.626 | 55 | 3 | 7 | 3 | 53 | 6 | 8.91 | 0.741 | 0.752 | 0.986 | -0.43 | 0.43 | -0.41 | -0.1366667 | 0.42 | -0.02 | 0.284041433 | 0.336401697 | 0.959078124 | 0.933919361 | 0.975286615 | 0.954602988 | 0.992902921 |
| 1500003O | High | P61022 | Calcineurin B homologous protein 1 OS=Mus musculus OX=10090 GN=Chp1 PE=1 SV=2 | 0 | 10.474 | 17 | 3 | 3 | 3 | 195 | 22.4 | 5.1 | 0.766 | 0.754 | 1.016 | -0.39 | 0.39 | -0.41 | -0.1366667 | 0.4 | 0.02 | 0.296007671 | 0.3406673 | 0.889858489 | 0.933919361 | 0.976045759 | 0.95498256 | 0.992535481 |
| Dlg4 | High | Q62108 | Disks large homolog 4 OS=Mus musculus OX=10090 GN=Dlg4 PE=1 SV=1 | 0 | 21.557 | 10 | 6 | 6 | 4 | 724 | 80.4 | 5.87 | 0.766 | 0.769 | 0.997 | -0.38 | 0.38 | -0.38 | -0.1266667 | 0.38 | 0 | 0.297864912 | 0.372566816 | 0.982598032 | 0.933919361 | 0.977313184 | 0.955616273 | 0.997765878 |
| Lmtk2 | High | Q3TYD6 | Serine/threonine-protein kinase LMTK2 OS=Mus musculus OX=10090 GN=Lmtk2 PE=1 SV=3 | 0 | 4.872 | 2 | 2 | 2 | 2 | 1471 | 160.4 | 4.51 | 0.775 | 0.767 | 1.01 | -0.37 | 0.37 | -0.38 | -0.1266667 | 0.375 | 0.01 | 0.276796533 | 0.36860747 | 0.910995496 | 0.933919361 | 0.977313184 | 0.955616273 | 0.992535481 |
| Slc23a2 | High | Q9EPR4 | Solute carrier family 23 member 2 OS=Mus musculus OX=10090 GN=Slc23a2 PE=1 SV=2 | 0 | 17.538 | 11 | 5 | 5 | 5 | 648 | 70 | 7.5 | 0.79 | 0.764 | 1.034 | -0.34 | 0.34 | -0.39 | -0.13 | 0.365 | 0.05 | 0.297085062 | 0.362055008 | 0.781821301 | 0.933919361 | 0.977313184 | 0.955616273 | 0.992535481 |
| | High | G3UZI2 | Heterogeneous nuclear ribonucleoprotein Q OS=Mus musculus OX=10090 GN=Syncrip PE=1 SV=1 | 0 | 18.404 | 14 | 7 | 8 | 5 | 527 | 58.7 | 7.56 | 1.351 | 1.369 | 0.987 | 0.43 | -0.43 | 0.45 | 0.15 | -0.44 | -0.02 | 0.298970405 | 0.380338739 | 0.967191453 | 0.933919361 | 0.977313184 | 0.955616273 | 0.994815921 |
| Phb2 | High | O35129 | Prohibitin-2 OS=Mus musculus OX=10090 GN=Phb2 PE=1 SV=1 | 0 | 41.215 | 41 | 12 | 17 | 12 | 299 | 33.3 | 9.83 | 1.347 | 1.367 | 0.986 | 0.43 | -0.43 | 0.45 | 0.15 | -0.44 | -0.02 | 0.276615363 | 0.383057211 | 0.993578862 | 0.933919361 | 0.977313184 | 0.955616273 | 0.999917596 |
| Hspa9 | High | P38647 | Stress-70 protein, mitochondrial OS=Mus musculus OX=10090 GN=Hspa9 PE=1 SV=3 | 0 | 117.822 | 38 | 22 | 37 | 21 | 679 | 73.4 | 6.07 | 1.373 | 1.382 | 0.993 | 0.46 | -0.46 | 0.47 | 0.15666667 | -0.465 | -0.01 | 0.284713246 | 0.364472681 | 0.954316821 | 0.933919361 | 0.977313184 | 0.955616273 | 0.992884189 |
| Wwp1 | High | Q8BZZ3 | NEDD4-like E3 ubiquitin-protein ligase WWP1 OS=Mus musculus OX=10090 GN=Wwp1 PE=1 SV= | 0 | 13.805 | 4 | 4 | 7 | 4 | 918 | 104.6 | 6.38 | 0.744 | 0.792 | 0.94 | -0.43 | 0.43 | -0.34 | -0.1133333 | 0.385 | -0.09 | 0.28303369 | 0.423268693 | 0.752487562 | 0.933919361 | 0.99399649 | 0.963957926 | 0.992535481 |
| 2310008H | Low | F6ZLH4 | Lysine-rich nucleolar protein 1 OS=Mus musculus OX=10090 GN=Knop1 PE=1 SV=2 | 0.081 | 1.202 | 4 | 1 | 1 | 1 | 347 | 38.9 | 10.23 | 0.785 | 0.803 | 0.977 | -0.35 | 0.35 | -0.32 | -0.1066667 | 0.335 | -0.03 | 0.297305657 | 0.432458444 | 0.978541043 | 0.933919361 | 0.99399649 | 0.963957926 | 0.997578114 |
| Rbp1 | High | Q00915 | Retinol-binding protein 1 OS=Mus musculus OX=10090 GN=Rbp1 PE=1 SV=2 | 0 | 144.848 | 79 | 19 | 65 | 19 | 135 | 15.8 | 5.25 | 1.373 | 1.335 | 1.028 | 0.46 | -0.46 | 0.42 | 0.14 | -0.44 | 0.04 | 0.284756188 | 0.423006748 | 0.780158722 | 0.933919361 | 0.99399649 | 0.963957926 | 0.992535481 |
| Lonp1 | High | Q8CGK3 | Lon protease homolog, mitochondrial OS=Mus musculus OX=10090 GN=Lonp1 PE=1 SV=2 | 0 | 11.496 | 4 | 4 | 4 | 4 | 949 | 105.8 | 6.57 | 1.341 | 1.316 | 1.018 | 0.42 | -0.42 | 0.4 | 0.13333333 | -0.41 | 0.03 | 0.296597192 | 0.448758491 | 0.915830633 | 0.933919361 | 0.994630833 | 0.964275097 | 0.992535481 |
| Syp | High | Q62277 | Synaptophysin OS=Mus musculus OX=10090 GN=Syp PE=1 SV=2 | 0 | 23.143 | 15 | 5 | 21 | 5 | 314 | 34 | 4.94 | 0.726 | 0.87 | 0.834 | -0.46 | 0.46 | -0.2 | -0.0666667 | 0.33 | -0.26 | 0.283345436 | 0.608899065 | 0.286389076 | 0.933919361 | 0.995559182 | 0.964739271 | 0.886074068 |
| | High | D3YVY0 | E3 ubiquitin-protein ligase MARCHF1 OS=Mus musculus OX=10090 GN=Marchf1 PE=1 SV=1 | 0 | 70.696 | 35 | 15 | 24 | 15 | 540 | 61.2 | 8.21 | 0.729 | 0.902 | 0.808 | -0.46 | 0.46 | -0.15 | -0.05 | 0.305 | -0.31 | 0.288568762 | 0.686574559 | 0.203680718 | 0.933919361 | 0.995559182 | 0.964739271 | 0.789790999 |
| Eif4b | High | Q8BGD9 | Eukaryotic translation initiation factor 4B OS=Mus musculus OX=10090 GN=Eif4b PE=1 SV=1 | 0 | 50.82 | 18 | 8 | 10 | 8 | 611 | 68.8 | 5.67 | 0.741 | 0.885 | 0.837 | -0.43 | 0.43 | -0.18 | -0.06 | 0.305 | -0.26 | 0.277165153 | 0.644817261 | 0.268274172 | 0.933919361 | 0.995559182 | 0.964739271 | 0.869723277 |
| Ufd1l; Ufd | High | P70362 | Ubiquitin recognition factor in ER-associated degradation protein 1 OS=Mus musculus OX=10090 | 0.006 | 3.316 | 7 | 2 | 2 | 2 | 307 | 34.5 | 6.7 | 0.785 | 0.898 | 0.875 | -0.35 | 0.35 | -0.16 | -0.0533333 | 0.255 | -0.19 | 0.285508201 | 0.651806903 | 0.470945653 | 0.933919361 | 0.995559182 | 0.964739271 | 0.961211268 |
| Rgmb | Medium | Q7TQ33 | Repulsive guidance molecule B OS=Mus musculus OX=10090 GN=Rgmb PE=1 SV=1 | 0.041 | 1.638 | 2 | 1 | 1 | 1 | 436 | 47.2 | 6.34 | 0.756 | 1.16 | 0.652 | -0.4 | 0.4 | 0.21 | 0.07 | 0.095 | -0.62 | 0.28334855 | 0.720662623 | 0.038114962 | 0.933919361 | 0.995559182 | 0.964739271 | 0.373579725 |
| Try10 | High | Q792Z1 | Trypsin 10 OS=Mus musculus OX=10090 GN=Try10 PE=1 SV=1 | 0 | 8.204 | 4 | 1 | 3 | 1 | 246 | 26.2 | 5.83 | 0.77 | 1.139 | 0.676 | -0.38 | 0.38 | 0.19 | 0.06333333 | 0.095 | -0.56 | 0.282607834 | 0.753945668 | 0.016763523 | 0.933919361 | 0.995559182 | 0.964739271 | 0.225403054 |
| | High | E0CY16 | Cell adhesion molecule 1 OS=Mus musculus OX=10090 GN=Cadm1 PE=1 SV=1 | 0 | 42.71 | 24 | 7 | 13 | 1 | 390 | 42.9 | 5.72 | 0.756 | 1.231 | 0.614 | -0.4 | 0.4 | 0.3 | 0.1 | 0.05 | -0.7 | 0.292981797 | 0.597377991 | 0.021125952 | 0.933919361 | 0.995559182 | 0.964739271 | 0.261486126 |
| Prrc2c | High | A0A0A0M | Protein PRRC2C OS=Mus musculus OX=10090 GN=Prrc2c PE=1 SV=1 | 0.001 | 4.44 | 1 | 2 | 2 | 2 | 2844 | 310.5 | 9.1 | 1.333 | 1.137 | 1.173 | 0.41 | -0.41 | 0.18 | 0.06 | -0.295 | 0.23 | 0.298695612 | 0.750391481 | 0.337853975 | 0.933919361 | 0.995559182 | 0.964739271 | 0.906329766 |
| | Medium | A0A571BF | K(+) channel subunit beta-1 OS=Mus musculus OX=10090 GN=Kcnab2 PE=1 SV=1 | 0.012 | 2.373 | 2 | 1 | 1 | 1 | 398 | 44 | 9.13 | 1.325 | 1.135 | 1.167 | 0.41 | -0.41 | 0.18 | 0.06 | -0.295 | 0.22 | 0.288252283 | 0.762811033 | 0.373937966 | 0.933919361 | 0.995559182 | 0.964739271 | 0.920164724 |
| | High | G3UX48 | Protein PRRC2A OS=Mus musculus OX=10090 GN=Prrc2a PE=1 SV=1 | 0.001 | 6.166 | 1 | 2 | 2 | 2 | 2069 | 219.3 | 9.25 | 1.343 | 1.148 | 1.17 | 0.43 | -0.43 | 0.2 | 0.06666667 | -0.315 | 0.23 | 0.293170371 | 0.738342177 | 0.314350938 | 0.933919361 | 0.995559182 | 0.964739271 | 0.904943958 |
| Rpl32; Rpl | High | P62911 | 60S ribosomal protein L32 OS=Mus musculus OX=10090 GN=Rpl32 PE=1 SV=2 | 0 | 52.14 | 40 | 8 | 18 | 8 | 135 | 15.9 | 11.33 | 1.365 | 1.139 | 1.198 | 0.45 | -0.45 | 0.19 | 0.06333333 | -0.32 | 0.26 | 0.293440017 | 0.753719344 | 0.205221586 | 0.933919361 | 0.995559182 | 0.964739271 | 0.795215918 |
| Purb | High | O35295 | Transcriptional activator protein Pur-beta OS=Mus musculus OX=10090 GN=Purb PE=1 SV=3 | 0 | 79.814 | 58 | 11 | 18 | 10 | 324 | 33.9 | 5.43 | 1.338 | 1.179 | 1.135 | 0.42 | -0.42 | 0.24 | 0.08 | -0.33 | 0.18 | 0.293597742 | 0.675828082 | 0.358803669 | 0.933919361 | 0.995559182 | 0.964739271 | 0.914600864 |
| Rps2; Rps | High | P25444 | 40S ribosomal protein S2 OS=Mus musculus OX=10090 GN=Rps2 PE=1 SV=3 | 0 | 81.533 | 49 | 13 | 28 | 13 | 293 | 31.2 | 10.24 | 1.379 | 1.148 | 1.202 | 0.46 | -0.46 | 0.2 | 0.06666667 | -0.33 | 0.26 | 0.278188289 | 0.736495322 | 0.19923992 | 0.933919361 | 0.995559182 | 0.964739271 | 0.78697088 |
| Hmgb3 | High | A2AP78 | High mobility group protein B3 (Fragment) OS=Mus musculus OX=10090 GN=Hmgb3 PE=1 SV=1 | 0 | 13.717 | 20 | 3 | 3 | 2 | 159 | 18.3 | 9.77 | 1.362 | 1.168 | 1.166 | 0.45 | -0.45 | 0.22 | 0.07333333 | -0.335 | 0.22 | 0.285671312 | 0.701097398 | 0.300524674 | 0.933919361 | 0.995559182 | 0.964739271 | 0.900241969 |
| Rpl6 | High | P47911 | 60S ribosomal protein L6 OS=Mus musculus OX=10090 GN=Rpl6 PE=1 SV=3 | 0 | 74.039 | 49 | 18 | 36 | 18 | 296 | 33.5 | 10.7 | 1.383 | 1.157 | 1.195 | 0.47 | -0.47 | 0.21 | 0.07 | -0.34 | 0.26 | 0.273925485 | 0.717502722 | 0.211635896 | 0.933919361 | 0.995559182 | 0.964739271 | 0.804011919 |
| Rpl28 | High | P41105 | 60S ribosomal protein L28 OS=Mus musculus OX=10090 GN=Rpl28 PE=1 SV=2 | 0 | 40.562 | 46 | 9 | 19 | 9 | 137 | 15.7 | 12.02 | 1.373 | 1.163 | 1.18 | 0.46 | -0.46 | 0.22 | 0.07333333 | -0.34 | 0.24 | 0.284920528 | 0.70597353 | 0.24232544 | 0.933919361 | 0.995559182 | 0.964739271 | 0.836649471 |
| Rps7 | High | P62082 | 40S ribosomal protein S7 OS=Mus musculus OX=10090 GN=Rps7 PE=2 SV=1 | 0 | 41.548 | 60 | 10 | 17 | 10 | 194 | 22.1 | 10.1 | 1.368 | 1.171 | 1.169 | 0.45 | -0.45 | 0.23 | 0.07666667 | -0.34 | 0.22 | 0.288759649 | 0.69048491 | 0.269282271 | 0.933919361 | 0.995559182 | 0.964739271 | 0.871243909 |
| | Medium | E9PX24 | LYR motif-containing protein 9 (Fragment) OS=Mus musculus OX=10090 GN=Lyrm9 PE=1 SV=1 | 0.029 | 1.906 | 7 | 1 | 1 | 1 | 97 | 11.1 | 6.89 | 1.362 | 1.184 | 1.15 | 0.45 | -0.45 | 0.24 | 0.08 | -0.345 | 0.2 | 0.287446103 | 0.669503826 | 0.344020152 | 0.933919361 | 0.995559182 | 0.964739271 | 0.907770836 |
| Rps8; Gm | High | P62242 | 40S ribosomal protein S8 OS=Mus musculus OX=10090 GN=Rps8 PE=1 SV=2 | 0 | 46.127 | 35 | 8 | 16 | 8 | 208 | 24.2 | 10.32 | 1.381 | 1.198 | 1.153 | 0.47 | -0.47 | 0.26 | 0.08666667 | -0.365 | 0.21 | 0.296350429 | 0.640366286 | 0.317457663 | 0.933919361 | 0.995559182 | 0.964739271 | 0.904943958 |
| Rps3a; Rp | High | P97351 | 40S ribosomal protein S3a OS=Mus musculus OX=10090 GN=Rps3a PE=1 SV=3 | 0 | 79.273 | 63 | 20 | 34 | 20 | 264 | 29.9 | 9.73 | 1.368 | 1.216 | 1.126 | 0.45 | -0.45 | 0.28 | 0.09333333 | -0.365 | 0.17 | 0.289893064 | 0.607965198 | 0.388125068 | 0.933919361 | 0.995559182 | 0.964739271 | 0.93064581 |
| Eif4a3 | High | Q91VC3 | Eukaryotic initiation factor 4A-III OS=Mus musculus OX=10090 GN=Eif4a3 PE=1 SV=3 | 0 | 23.755 | 20 | 7 | 7 | 5 | 411 | 46.8 | 6.73 | 1.366 | 1.222 | 1.118 | 0.45 | -0.45 | 0.29 | 0.09666667 | -0.37 | 0.16 | 0.279466904 | 0.597368977 | 0.481567203 | 0.933919361 | 0.995559182 | 0.964739271 | 0.964378027 |
| Fsd1l | High | Q8BYN5 | FSD1-like protein OS=Mus musculus OX=10090 GN=Fsd1l PE=1 SV=2 | 0 | 32.349 | 17 | 9 | 12 | 9 | 507 | 56.9 | 6.7 | 1.351 | 1.247 | 1.083 | 0.43 | -0.43 | 0.32 | 0.10666667 | -0.375 | 0.12 | 0.279879934 | 0.554389887 | 0.537563078 | 0.933919361 | 0.995559182 | 0.964739271 | 0.96694628 |
| Rpl19; Rpl | High | P84099 | 60S ribosomal protein L19 OS=Mus musculus OX=10090 GN=Rpl19 PE=1 SV=1 | 0 | 38.829 | 35 | 9 | 16 | 9 | 196 | 23.5 | 11.47 | 1.399 | 1.21 | 1.156 | 0.48 | -0.48 | 0.28 | 0.09333333 | -0.38 | 0.21 | 0.274593122 | 0.617746709 | 0.324293627 | 0.933919361 | 0.995559182 | 0.964739271 | 0.905686588 |
| Ubtd1 | High | Q91WB7 | Ubiquitin domain-containing protein 1 OS=Mus musculus OX=10090 GN=Ubtd1 PE=2 SV=1 | 0 | 4.872 | 6 | 1 | 1 | 1 | 227 | 26 | 7.83 | 1.347 | 1.267 | 1.063 | 0.43 | -0.43 | 0.34 | 0.11333333 | -0.385 | 0.09 | 0.288633025 | 0.518350375 | 0.726618046 | 0.933919361 | 0.995559182 | 0.964739271 | 0.992535481 |
| | Low | F7AGB5 | Methyltransferase-like protein 17, mitochondrial (Fragment) OS=Mus musculus OX=10090 GN=M | 0.06 | 1.324 | 3 | 1 | 1 | 1 | 280 | 31.6 | 8.25 | 1.336 | 1.284 | 1.04 | 0.42 | -0.42 | 0.36 | 0.12 | -0.39 | 0.06 | 0.299012406 | 0.495524327 | 0.762118189 | 0.933919361 | 0.995559182 | 0.964739271 | 0.992535481 |
| Cirbp | High | P60824 | Cold-inducible RNA-binding protein OS=Mus musculus OX=10090 GN=Cirbp PE=1 SV=1 | 0 | 16.472 | 24 | 3 | 4 | 3 | 172 | 18.6 | 9.61 | 1.356 | 1.283 | 1.057 | 0.44 | -0.44 | 0.36 | 0.12 | -0.4 | 0.08 | 0.292650962 | 0.497827715 | 0.768583722 | 0.933919361 | 0.995559182 | 0.964739271 | 0.992535481 |
| Lin7a | High | Q8JZS0 | Protein lin-7 homolog A OS=Mus musculus OX=10090 GN=Lin7a PE=1 SV=2 | 0 | 36.02 | 39 | 7 | 11 | 4 | 233 | 26 | 8.72 | 0.768 | 0.956 | 0.803 | -0.38 | 0.38 | -0.07 | -0.0233333 | 0.225 | -0.32 | 0.298719917 | 0.822053484 | 0.222346873 | 0.933919361 | 0.998402594 | 0.966160978 | 0.807447486 |
| Mtap7d3; | Medium | A0A0R4J1 | MAP7 domain-containing protein 3 (Fragment) OS=Mus musculus OX=10090 GN=Map7d3 PE=4 S | 0.026 | 1.959 | 1 | 1 | 1 | 1 | 489 | 54.3 | 9.33 | 0.777 | 0.968 | 0.803 | -0.36 | 0.36 | -0.05 | -0.0166667 | 0.205 | -0.32 | 0.29931391 | 0.851783454 | 0.212297069 | 0.933919361 | 0.998402594 | 0.966160978 | 0.804011919 |
| Eea1 | High | Q8BL66 | Early endosome antigen 1 OS=Mus musculus OX=10090 GN=Eea1 PE=1 SV=2 | 0 | 10.066 | 3 | 3 | 4 | 2 | 1411 | 160.8 | 5.77 | 0.772 | 0.973 | 0.793 | -0.37 | 0.37 | -0.04 | -0.0133333 | 0.205 | -0.33 | 0.298152474 | 0.856196535 | 0.218568527 | 0.933919361 | 0.998402594 | 0.966160978 | 0.805811317 |
| Gfap | High | P03995 | Glial fibrillary acidic protein OS=Mus musculus OX=10090 GN=Gfap PE=1 SV=4 | 0 | 9.269 | 9 | 4 | 6 | 2 | 430 | 49.9 | 5.34 | 0.763 | 0.998 | 0.764 | -0.39 | 0.39 | 0 | 0 | 0.195 | -0.39 | 0.278372226 | 0.907558036 | 0.124971737 | 0.933919361 | 0.998402594 | 0.966160978 | 0.678506823 |
| S100a1 | High | Q91V77 | Protein S100 OS=Mus musculus OX=10090 GN=S100a1 PE=1 SV=1 | 0 | 10.017 | 16 | 1 | 2 | 1 | 94 | 10.5 | 4.59 | 0.785 | 1.004 | 0.781 | -0.35 | 0.35 | 0.01 | 0.00333333 | 0.17 | -0.36 | 0.285613108 | 0.945166677 | 0.17270951 | 0.933919361 | 0.998402594 | 0.966160978 | 0.738844686 |
| Gm4980; | High | Q8C013 | Trophoblast glycoprotein-like OS=Mus musculus OX=10090 GN=Tpbgl PE=1 SV=2 | 0.002 | 4.114 | 3 | 1 | 1 | 1 | 384 | 41.1 | 7.39 | 0.787 | 1.078 | 0.73 | -0.35 | 0.35 | 0.11 | 0.03666667 | 0.12 | -0.45 | 0.296625918 | 0.877048294 | 0.079032666 | 0.933919361 | 0.998402594 | 0.966160978 | 0.539751713 |
| Bzw2 | High | Q91VK1 | Basic leucine zipper and W2 domain-containing protein 2 OS=Mus musculus OX=10090 GN=Bzw2 | 0 | 42.961 | 34 | 14 | 21 | 14 | 419 | 48 | 6.68 | 1.363 | 0.937 | 1.454 | 0.45 | -0.45 | -0.09 | -0.03 | -0.18 | 0.54 | 0.281506637 | 0.776270672 | 0.013885449 | 0.933919361 | 0.998402594 | 0.966160978 | 0.19319035 |
| Hepacam | High | Q640R3 | Hepatocyte cell adhesion molecule OS=Mus musculus OX=10090 GN=Hepacam PE=1 SV=2 | 0.006 | 3.129 | 2 | 1 | 1 | 1 | 418 | 46.3 | 9.42 | 1.353 | 0.991 | 1.365 | 0.44 | -0.44 | -0.01 | -0.0033333 | -0.215 | 0.45 | 0.294376935 | 0.892838163 | 0.051091431 | 0.933919361 | 0.998402594 | 0.966160978 | 0.427566646 |
| Rpl7 | High | P14148 | 60S ribosomal protein L7 OS=Mus musculus OX=10090 GN=Rpl7 PE=1 SV=2 | 0 | 95.227 | 51 | 18 | 36 | 18 | 270 | 31.4 | 10.89 | 1.383 | 1.08 | 1.28 | 0.47 | -0.47 | 0.11 | 0.03666667 | -0.29 | 0.36 | 0.274104215 | 0.880136706 | 0.090712745 | 0.933919361 | 0.998402594 | 0.966160978 | 0.576060131 |
| Arhgef12 | High | Q8R4H2 | Rho guanine nucleotide exchange factor 12 OS=Mus musculus OX=10090 GN=Arhgef12 PE=1 SV= | 0.006 | 3.176 | 1 | 1 | 1 | 1 | 1543 | 172.2 | 5.74 | 1.338 | 1.115 | 1.201 | 0.42 | -0.42 | 0.16 | 0.05333333 | -0.29 | 0.26 | 0.283966254 | 0.800549208 | 0.299729973 | 0.933919361 | 0.998402594 | 0.966160978 | 0.900241969 |
| | High | A0A1D5RL | 60S ribosomal protein L18a OS=Mus musculus OX=10090 GN=Rpl18a PE=1 SV=1 | 0 | 39.11 | 41 | 7 | 17 | 7 | 147 | 17.4 | 10.89 | 1.377 | 1.123 | 1.226 | 0.46 | -0.46 | 0.17 | 0.05666667 | -0.315 | 0.29 | 0.280221555 | 0.786944629 | 0.157335801 | 0.933919361 | 0.998402594 | 0.966160978 | 0.733365826 |
| | High | E9Q2S9 | Coiled-coil domain-containing protein 43 OS=Mus musculus OX=10090 GN=Ccdc43 PE=1 SV=1 | 0.003 | 3.965 | 4 | 1 | 1 | 1 | 222 | 25 | 5.02 | 0.776 | 0.779 | 0.996 | -0.37 | 0.37 | -0.36 | -0.12 | 0.365 | -0.01 | 0.300264725 | 0.370913126 | 0.95293215 | 0.934805283 | 0.977313184 | 0.956059233 | 0.992535481 |
| Tuba1a | High | P68369 | Tubulin alpha-1A chain OS=Mus musculus OX=10090 GN=Tuba1a PE=1 SV=1 | 0 | 399.607 | 62 | 25 | 172 | 6 | 451 | 50.1 | 5.06 | 1.359 | 1.38 | 0.985 | 0.44 | -0.44 | 0.46 | 0.15333333 | -0.45 | -0.02 | 0.300130266 | 0.367148573 | 0.997234633 | 0.934805283 | 0.977313184 | 0.956059233 | 0.999917596 |
| Rps23 | High | P62267 | 40S ribosomal protein S23 OS=Mus musculus OX=10090 GN=Rps23 PE=1 SV=3 | 0 | 33.102 | 30 | 6 | 10 | 6 | 143 | 15.8 | 10.49 | 1.32 | 1.155 | 1.143 | 0.4 | -0.4 | 0.21 | 0.07 | -0.305 | 0.19 | 0.300025356 | 0.721705753 | 0.337685255 | 0.934805283 | 0.995559182 | 0.965182232 | 0.906329766 |
| Daglb | High | Q91WC9 | Diacylglycerol lipase-beta OS=Mus musculus OX=10090 GN=Daglb PE=1 SV=2 | 0.008 | 2.986 | 1 | 1 | 1 | 1 | 669 | 73.9 | 6.73 | 0.783 | 0.757 | 1.034 | -0.35 | 0.35 | -0.4 | -0.1333333 | 0.375 | 0.05 | 0.300501837 | 0.34007095 | 0.820504785 | 0.935024307 | 0.976045759 | 0.955535033 | 0.992535481 |
| Fam162a | High | Q9D6U8 | Protein FAM162A OS=Mus musculus OX=10090 GN=Fam162a PE=1 SV=1 | 0 | 9.805 | 19 | 4 | 4 | 4 | 155 | 17.7 | 9.88 | 1.294 | 1.384 | 0.935 | 0.37 | -0.37 | 0.47 | 0.15666667 | -0.42 | -0.1 | 0.300978545 | 0.361853938 | 0.689221136 | 0.935615217 | 0.977313184 | 0.9564642 | 0.991285312 |
| Alb | High | P07724 | Albumin OS=Mus musculus OX=10090 GN=Alb PE=1 SV=3 | 0 | 49.493 | 22 | 13 | 17 | 13 | 608 | 68.6 | 6.07 | 0.735 | 0.968 | 0.759 | -0.44 | 0.44 | -0.05 | -0.0166667 | 0.245 | -0.4 | 0.301025477 | 0.85169002 | 0.094059827 | 0.935615217 | 0.998402594 | 0.967008905 | 0.583399835 |
| Tab3 | Medium | Q571K4 | TGF-beta-activated kinase 1 and MAP3K7-binding protein 3 OS=Mus musculus OX=10090 GN=Tab | 0.01 | 2.702 | 1 | 1 | 1 | 1 | 716 | 79 | 8.5 | 0.771 | 0.731 | 1.054 | -0.38 | 0.38 | -0.45 | -0.15 | 0.415 | 0.08 | 0.301555902 | 0.286151513 | 0.750174834 | 0.936225882 | 0.955820031 | 0.946022956 | 0.992535481 |
| Slc39a4 | High | Q78IQ7 | Zinc transporter ZIP4 OS=Mus musculus OX=10090 GN=Slc39a4 PE=1 SV=1 | 0 | 57.227 | 24 | 9 | 18 | 9 | 660 | 71 | 5.57 | 1.364 | 1.184 | 1.152 | 0.45 | -0.45 | 0.24 | 0.08 | -0.345 | 0.2 | 0.302237442 | 0.666121297 | 0.370919386 | 0.937303837 | 0.995559182 | 0.966431509 | 0.920164724 |
| Psat1 | High | Q99K85 | Phosphoserine aminotransferase OS=Mus musculus OX=10090 GN=Psat1 PE=1 SV=1 | 0 | 39.084 | 22 | 8 | 12 | 8 | 370 | 40.4 | 8.03 | 1.336 | 1.25 | 1.069 | 0.42 | -0.42 | 0.32 | 0.10666667 | -0.37 | 0.1 | 0.302421694 | 0.549215585 | 0.597347918 | 0.937356794 | 0.995559182 | 0.966457988 | 0.977451587 |
| Prom1 | High | Q8CDK8 | Prominin-1 OS=Mus musculus OX=10090 GN=Prom1 PE=1 SV=1 | 0 | 267.467 | 39 | 32 | 110 | 1 | 827 | 92.7 | 6.55 | 0.798 | 0.778 | 1.027 | -0.33 | 0.33 | -0.36 | -0.12 | 0.345 | 0.04 | 0.302861653 | 0.369450644 | 0.874470586 | 0.938201817 | 0.977313184 | 0.957757501 | 0.992535481 |
| Hspa4 | High | Q3U2G2 | Heat shock 70 kDa protein 4 OS=Mus musculus OX=10090 GN=Hspa4 PE=1 SV=1 | 0 | 145.33 | 48 | 32 | 46 | 29 | 842 | 94.1 | 5.21 | 0.736 | 0.736 | 1.001 | -0.44 | 0.44 | -0.44 | -0.1466667 | 0.44 | 0 | 0.304355139 | 0.304388355 | 0.916824725 | 0.940231001 | 0.955820031 | 0.948025516 | 0.992535481 |
| Arhgap1 | High | Q5FWK3 | Rho GTPase-activating protein 1 OS=Mus musculus OX=10090 GN=Arhgap1 PE=1 SV=1 | 0 | 33.484 | 28 | 9 | 10 | 9 | 439 | 50.4 | 6.44 | 1.316 | 1.455 | 0.905 | 0.4 | -0.4 | 0.54 | 0.18 | -0.47 | -0.14 | 0.304313808 | 0.287613587 | 0.532659467 | 0.940231001 | 0.955820031 | 0.948025516 | 0.96694628 |
| Faim2 | High | Q8K097 | Protein lifeguard 2 OS=Mus musculus OX=10090 GN=Faim2 PE=2 SV=1 | 0.001 | 5.617 | 7 | 1 | 1 | 1 | 317 | 35.2 | 6.92 | 0.766 | 0.93 | 0.824 | -0.38 | 0.38 | -0.1 | -0.0333333 | 0.24 | -0.28 | 0.304248313 | 0.756771663 | 0.308910275 | 0.940231001 | 0.995559182 | 0.967895091 | 0.904943958 |
| Rps19 | High | Q9CZX8 | 40S ribosomal protein S19 OS=Mus musculus OX=10090 GN=Rps19 PE=1 SV=3 | 0 | 45.511 | 55 | 12 | 19 | 12 | 145 | 16.1 | 10.4 | 1.356 | 1.26 | 1.076 | 0.44 | -0.44 | 0.33 | 0.11 | -0.385 | 0.11 | 0.304302338 | 0.533334422 | 0.567027698 | 0.940231001 | 0.995559182 | 0.967895091 | 0.972910028 |
| Rap1gds1 | High | E9Q912 | Rap1 GTPase-GDP dissociation stimulator 1 OS=Mus musculus OX=10090 GN=Rap1gds1 PE=2 SV | 0 | 37.85 | 17 | 8 | 11 | 8 | 607 | 66 | 5.35 | 0.755 | 0.893 | 0.846 | -0.4 | 0.4 | -0.16 | -0.0533333 | 0.28 | -0.24 | 0.304606414 | 0.664332376 | 0.320606625 | 0.940489078 | 0.995559182 | 0.96802413 | 0.905612769 |
| Lamp1 | High | P11438 | Lysosome-associated membrane glycoprotein 1 OS=Mus musculus OX=10090 GN=Lamp1 PE=1 SV | 0.001 | 5.254 | 3 | 1 | 1 | 1 | 406 | 43.8 | 8.4 | 0.786 | 1.038 | 0.757 | -0.35 | 0.35 | 0.05 | 0.01666667 | 0.15 | -0.4 | 0.304873803 | 0.985092399 | 0.170565887 | 0.940796594 | 0.998402594 | 0.969599594 | 0.738844686 |
| Cops2 | High | P61202 | COP9 signalosome complex subunit 2 OS=Mus musculus OX=10090 GN=Cops2 PE=1 SV=1 | 0 | 19.672 | 18 | 7 | 7 | 7 | 443 | 51.6 | 5.53 | 0.753 | 0.729 | 1.032 | -0.41 | 0.41 | -0.46 | -0.1533333 | 0.435 | 0.05 | 0.305246071 | 0.291687754 | 0.870970572 | 0.940920112 | 0.955820031 | 0.948370071 | 0.992535481 |
| Uqcrh | High | P99028 | Cytochrome b-c1 complex subunit 6, mitochondrial OS=Mus musculus OX=10090 GN=Uqcrh PE=1 | 0 | 11.407 | 28 | 2 | 6 | 2 | 89 | 10.4 | 4.87 | 1.329 | 1.454 | 0.914 | 0.41 | -0.41 | 0.54 | 0.18 | -0.475 | -0.13 | 0.305249453 | 0.288476219 | 0.616998991 | 0.940920112 | 0.955820031 | 0.948370071 | 0.97826833 |
| | High | A0A286YE | 40S ribosomal protein S24 OS=Mus musculus OX=10090 GN=Rps24 PE=1 SV=1 | 0 | 24.382 | 23 | 5 | 13 | 5 | 118 | 13.6 | 10.77 | 1.355 | 1.091 | 1.242 | 0.44 | -0.44 | 0.13 | 0.04333333 | -0.285 | 0.31 | 0.305593056 | 0.85716433 | 0.134165079 | 0.941461683 | 0.998402594 | 0.969932139 | 0.697570122 |
| Nploc4 | Medium | P60670 | Nuclear protein localization protein 4 homolog OS=Mus musculus OX=10090 GN=Nploc4 PE=1 SV | 0.018 | 2.074 | 1 | 1 | 1 | 1 | 608 | 68 | 6.46 | 0.786 | 0.654 | 1.202 | -0.35 | 0.35 | -0.61 | -0.2033333 | 0.48 | 0.26 | 0.305908676 | 0.151515849 | 0.273154723 | 0.94146351 | 0.847684847 | 0.894574179 | 0.876067403 |
| | Low | D3Z408 | High affinity cGMP-specific 3',5'-cyclic phosphodiesterase 9A OS=Mus musculus OX=10090 GN=P | 0.08 | 1.21 | 4 | 1 | 1 | 1 | 387 | 45.3 | 5.69 | 1.327 | 1.494 | 0.888 | 0.41 | -0.41 | 0.58 | 0.19333333 | -0.495 | -0.17 | 0.310295089 | 0.237532216 | 0.526649816 | 0.94146351 | 0.937374952 | 0.939419231 | 0.964378027 |
| | High | F8WJ64 | Acyl carrier protein (Fragment) OS=Mus musculus OX=10090 GN=Ndufab1 PE=1 SV=1 | 0 | 15.701 | 26 | 3 | 5 | 3 | 128 | 14.4 | 4.77 | 1.335 | 1.43 | 0.933 | 0.42 | -0.42 | 0.52 | 0.17333333 | -0.47 | -0.1 | 0.307877941 | 0.311424881 | 0.7211034 | 0.94146351 | 0.955820031 | 0.94864177 | 0.992535481 |
| Gdi1 | High | P50396 | Rab GDP dissociation inhibitor alpha OS=Mus musculus OX=10090 GN=Gdi1 PE=1 SV=3 | 0 | 120.874 | 72 | 23 | 36 | 18 | 447 | 50.5 | 5.08 | 0.737 | 0.768 | 0.96 | -0.44 | 0.44 | -0.38 | -0.1266667 | 0.41 | -0.06 | 0.306008555 | 0.369592112 | 0.871584249 | 0.94146351 | 0.977313184 | 0.959388347 | 0.992535481 |
| Tars | High | Q9D0R2 | Threonine--tRNA ligase 1, cytoplasmic OS=Mus musculus OX=10090 GN=Tars1 PE=1 SV=2 | 0 | 19.986 | 10 | 7 | 8 | 6 | 722 | 83.3 | 7.36 | 0.766 | 0.769 | 0.996 | -0.38 | 0.38 | -0.38 | -0.1266667 | 0.38 | -0.01 | 0.306527852 | 0.373550728 | 0.962137707 | 0.94146351 | 0.977313184 | 0.959388347 | 0.993960013 |
| Scai | High | Q8C8N2 | Protein SCAI OS=Mus musculus OX=10090 GN=Scai PE=1 SV=2 | 0.001 | 6.418 | 4 | 3 | 3 | 3 | 606 | 70.2 | 8.6 | 0.774 | 0.872 | 0.887 | -0.37 | 0.37 | -0.2 | -0.0666667 | 0.285 | -0.17 | 0.30930101 | 0.611011606 | 0.536258205 | 0.94146351 | 0.995559182 | 0.968511346 | 0.96694628 |
| | High | G5E8Y0 | Na(+)/Ca(2+)-exchange protein 1 OS=Mus musculus OX=10090 GN=Slc8a1 PE=1 SV=2 | 0.001 | 5.979 | 2 | 2 | 2 | 2 | 970 | 107.9 | 5 | 0.801 | 0.929 | 0.862 | -0.32 | 0.32 | -0.11 | -0.0366667 | 0.215 | -0.21 | 0.308699814 | 0.729150613 | 0.39235271 | 0.94146351 | 0.995559182 | 0.968511346 | 0.93064581 |
| Alyref | High | O08583 | THO complex subunit 4 OS=Mus musculus OX=10090 GN=Alyref PE=1 SV=3 | 0.001 | 4.679 | 7 | 1 | 1 | 1 | 255 | 26.9 | 11.15 | 1.354 | 1.143 | 1.185 | 0.44 | -0.44 | 0.19 | 0.06333333 | -0.315 | 0.25 | 0.306920483 | 0.767191544 | 0.337459185 | 0.94146351 | 0.995559182 | 0.968511346 | 0.906329766 |
| Rpl3 | High | P27659 | 60S ribosomal protein L3 OS=Mus musculus OX=10090 GN=Rpl3 PE=1 SV=3 | 0 | 77.182 | 34 | 24 | 37 | 24 | 403 | 46.1 | 10.21 | 1.352 | 1.187 | 1.139 | 0.44 | -0.44 | 0.25 | 0.08333333 | -0.345 | 0.19 | 0.308400817 | 0.65999981 | 0.347028766 | 0.94146351 | 0.995559182 | 0.968511346 | 0.9092478 |
| Rpl27a | High | P14115 | 60S ribosomal protein L27a OS=Mus musculus OX=10090 GN=Rpl27a PE=1 SV=5 | 0 | 19.193 | 31 | 5 | 10 | 5 | 148 | 16.6 | 11.12 | 1.353 | 1.196 | 1.131 | 0.44 | -0.44 | 0.26 | 0.08666667 | -0.35 | 0.18 | 0.307772364 | 0.64238549 | 0.372647456 | 0.94146351 | 0.995559182 | 0.968511346 | 0.920164724 |
| Rps5 | High | Q91V55 | 40S ribosomal protein S5 OS=Mus musculus OX=10090 GN=Rps5 PE=1 SV=1 | 0 | 44.458 | 33 | 8 | 14 | 8 | 204 | 22.9 | 9.72 | 1.358 | 1.223 | 1.11 | 0.44 | -0.44 | 0.29 | 0.09666667 | -0.365 | 0.15 | 0.307326261 | 0.594996151 | 0.450207202 | 0.94146351 | 0.995559182 | 0.968511346 | 0.959449555 |

| | | | | | | | | | | | | | | | | | | | | | | | | | | | | |
|---|---|---|---|---|---|---|---|---|---|---|---|---|---|---|---|---|---|---|---|---|---|---|---|---|---|---|---|---|
| Rps14 | High | P62264 | 40S ribosomal protein S14 OS=Mus musculus OX=10090 GN=Rps14 PE=1 SV=3 | 0 | 31.203 | 38 | 5 | 13 | 5 | 151 | 16.3 | 10.05 | 1.366 | 1.216 | 1.123 | 0.45 | -0.45 | 0.28 | 0.09333333 | -0.365 | 0.17 | 0.307433656 | 0.60640163 | 0.462468369 | 0.94146351 | 0.995559182 | 0.968511346 | 0.959744396 |
| Scarb1 | High | Q61009 | Scavenger receptor class B member 1 OS=Mus musculus OX=10090 GN=Scarb1 PE=1 SV=1 | 0 | 16.677 | 14 | 6 | 7 | 6 | 509 | 56.7 | 8.09 | 1.354 | 1.264 | 1.071 | 0.44 | -0.44 | 0.34 | 0.11333333 | -0.39 | 0.1 | 0.306524638 | 0.527345541 | 0.696721232 | 0.94146351 | 0.995559182 | 0.968511346 | 0.991285312 |
| Sub1 | High | P11031 | Activated RNA polymerase II transcriptional coactivator p15 OS=Mus musculus OX=10090 GN=Sul | 0 | 16.177 | 26 | 4 | 6 | 4 | 127 | 14.4 | 9.6 | 1.364 | 0.977 | 1.396 | 0.45 | -0.45 | -0.03 | -0.01 | -0.21 | 0.48 | 0.307146657 | 0.875531055 | 0.037527493 | 0.94146351 | 0.998402594 | 0.969933052 | 0.373579725 |
| Hkdc1 | High | Q91W97 | Hexokinase HKDC1 OS=Mus musculus OX=10090 GN=Hkdc1 PE=2 SV=1 | 0 | 8.455 | 3 | 4 | 5 | 2 | 915 | 102.2 | 7.09 | 0.796 | 0.787 | 1.012 | -0.33 | 0.33 | -0.35 | -0.1166667 | 0.34 | 0.02 | 0.311107446 | 0.411057309 | 0.901212745 | 0.942398408 | 0.986643122 | 0.964520765 | 0.992535481 |
| Usp5 | High | P56399 | Ubiquitin carboxyl-terminal hydrolase 5 OS=Mus musculus OX=10090 GN=Usp5 PE=1 SV=1 | 0 | 45.07 | 21 | 15 | 17 | 15 | 858 | 95.8 | 5.01 | 0.757 | 0.851 | 0.89 | -0.4 | 0.4 | -0.23 | -0.0766667 | 0.315 | -0.17 | 0.31104856 | 0.561358363 | 0.449049816 | 0.942398408 | 0.995559182 | 0.968978795 | 0.958608248 |
| Cox6c | High | Q9CPQ1 | Cytochrome c oxidase subunit 6C OS=Mus musculus OX=10090 GN=Cox6c PE=1 SV=3 | 0 | 15.855 | 38 | 5 | 9 | 5 | 76 | 8.5 | 10.14 | 1.362 | 1.447 | 0.941 | 0.45 | -0.45 | 0.53 | 0.17666667 | -0.49 | -0.09 | 0.312084262 | 0.294504828 | 0.789485096 | 0.94457979 | 0.955820031 | 0.95019991 | 0.992535481 |
| Ift52 | High | F6RBL8 | Intraflagellar transport protein 52 homolog (Fragment) OS=Mus musculus OX=10090 GN=Ift52 PE | 0.001 | 6.419 | 9 | 2 | 2 | 2 | 234 | 26.2 | 8.31 | 1.338 | 1.048 | 1.277 | 0.42 | -0.42 | 0.07 | 0.02333333 | -0.245 | 0.35 | 0.312332964 | 0.969993936 | 0.096152716 | 0.94457979 | 0.998402594 | 0.971491192 | 0.590616329 |
| Cox4i1 | High | P19783 | Cytochrome c oxidase subunit 4 isoform 1, mitochondrial OS=Mus musculus OX=10090 GN=Cox4i | 0 | 32.092 | 40 | 6 | 16 | 6 | 169 | 19.5 | 9.23 | 1.357 | 1.408 | 0.964 | 0.44 | -0.44 | 0.49 | 0.16333333 | -0.465 | -0.05 | 0.312589144 | 0.335305039 | 0.890556127 | 0.944598895 | 0.975040602 | 0.959819749 | 0.992535481 |
| Prkaa1 | High | Q3TUQ7 | Acetyl-CoA carboxylase kinase OS=Mus musculus OX=10090 GN=Prkaa1 PE=1 SV=1 | 0 | 10.594 | 7 | 3 | 3 | 3 | 550 | 62.7 | 7.59 | 0.791 | 0.768 | 1.03 | -0.34 | 0.34 | -0.38 | -0.1266667 | 0.36 | 0.04 | 0.313013153 | 0.370048166 | 0.793103508 | 0.944598895 | 0.977313184 | 0.96095604 | 0.992535481 |
| Camkv | High | Q3UHL1 | CaM kinase-like vesicle-associated protein OS=Mus musculus OX=10090 GN=Camkv PE=1 SV=2 | 0 | 54.475 | 38 | 10 | 17 | 10 | 512 | 54.8 | 5.54 | 0.755 | 0.845 | 0.894 | -0.41 | 0.41 | -0.24 | -0.08 | 0.325 | -0.16 | 0.312892329 | 0.545879903 | 0.508162309 | 0.944598895 | 0.995559182 | 0.970079038 | 0.964378027 |
| Rbbp9 | High | O88851 | Putative hydrolase RBBP9 OS=Mus musculus OX=10090 GN=Rbbp9 PE=1 SV=2 | 0 | 9.906 | 19 | 2 | 2 | 2 | 186 | 20.9 | 5.97 | 0.783 | 0.914 | 0.856 | -0.35 | 0.35 | -0.13 | -0.0433333 | 0.24 | -0.22 | 0.313705984 | 0.696502587 | 0.415711052 | 0.946180449 | 0.995559182 | 0.970869815 | 0.936397673 |
| Usp47 | High | A0A1L1SV | Ubiquitin carboxyl-terminal hydrolase 47 OS=Mus musculus OX=10090 GN=Usp47 PE=1 SV=1 | 0 | 10.53 | 2 | 3 | 3 | 3 | 1376 | 157.1 | 5.06 | 0.807 | 0.554 | 1.458 | -0.31 | 0.31 | -0.85 | -0.2833333 | 0.58 | 0.54 | 0.324478811 | 0.047614307 | 0.020940539 | 0.950027282 | 0.520047141 | 0.735037212 | 0.259764608 |
| Ndufv3 | High | Q8BK30 | NADH dehydrogenase [ubiquinone] flavoprotein 3, mitochondrial OS=Mus musculus OX=10090 GN | 0.001 | 6.297 | 16 | 1 | 1 | 1 | 104 | 11.8 | 9.35 | 1.318 | 1.95 | 0.676 | 0.4 | -0.4 | 0.96 | 0.32 | -0.68 | -0.56 | 0.318311513 | 0.045724161 | 0.028978157 | 0.950027282 | 0.520047141 | 0.735037212 | 0.324925684 |
| Nefm | High | P08553 | Neurofilament medium polypeptide OS=Mus musculus OX=10090 GN=Nefm PE=1 SV=4 | 0 | 8.385 | 3 | 2 | 4 | 1 | 848 | 95.9 | 4.77 | 0.795 | 0.569 | 1.399 | -0.33 | 0.33 | -0.81 | -0.27 | 0.57 | 0.48 | 0.32201014 | 0.061684151 | 0.074568703 | 0.950027282 | 0.586560875 | 0.768294078 | 0.520184986 |
| Ndufa5 | High | Q9CPP6 | NADH dehydrogenase [ubiquinone] 1 alpha subcomplex subunit 5 OS=Mus musculus OX=10090 G | 0 | 25.954 | 45 | 5 | 8 | 5 | 116 | 13.4 | 8.1 | 1.317 | 1.699 | 0.775 | 0.4 | -0.4 | 0.76 | 0.25333333 | -0.58 | -0.37 | 0.323093247 | 0.123745841 | 0.123689864 | 0.950027282 | 0.799286338 | 0.87465681 | 0.677192798 |
| Bpgm | High | P15327 | Bisphosphoglycerate mutase OS=Mus musculus OX=10090 GN=Bpgm PE=1 SV=2 | 0 | 13.478 | 7 | 1 | 2 | 1 | 259 | 30 | 7.06 | 0.798 | 0.658 | 1.212 | -0.33 | 0.33 | -0.6 | -0.2 | 0.465 | 0.28 | 0.318241273 | 0.163341817 | 0.274278851 | 0.950027282 | 0.869444647 | 0.909735965 | 0.876067403 |
| Rheb | High | Q921J2 | GTP-binding protein Rheb OS=Mus musculus OX=10090 GN=Rheb PE=1 SV=1 | 0 | 12.57 | 26 | 5 | 6 | 5 | 184 | 20.4 | 5.92 | 0.776 | 0.663 | 1.171 | -0.37 | 0.37 | -0.59 | -0.1966667 | 0.48 | 0.23 | 0.315565585 | 0.178060461 | 0.376057247 | 0.950027282 | 0.875953959 | 0.91299062 | 0.92034959 |
| Map3k7 | High | A2AP92 | Mitogen-activated protein kinase kinase kinase 7 OS=Mus musculus OX=10090 GN=Map3k7 PE=1 | 0.006 | 3.415 | 2 | 1 | 1 | 1 | 518 | 56.7 | 7.24 | 0.786 | 0.679 | 1.157 | -0.35 | 0.35 | -0.56 | -0.1866667 | 0.455 | 0.21 | 0.319124823 | 0.176845407 | 0.378494111 | 0.950027282 | 0.875953959 | 0.91299062 | 0.922487352 |
| Ak3 | High | Q9WTP7 | GTP:AMP phosphotransferase AK3, mitochondrial OS=Mus musculus OX=10090 GN=Ak3 PE=1 SV= | 0.001 | 7.18 | 12 | 3 | 3 | 3 | 227 | 25.4 | 8.84 | 1.301 | 1.476 | 0.881 | 0.38 | -0.38 | 0.56 | 0.18666667 | -0.47 | -0.18 | 0.320789261 | 0.266273136 | 0.488130651 | 0.950027282 | 0.951072422 | 0.950549852 | 0.964378027 |
| Mdh1 | High | P14152 | Malate dehydrogenase, cytoplasmic OS=Mus musculus OX=10090 GN=Mdh1 PE=1 SV=3 | 0 | 162.624 | 48 | 17 | 70 | 17 | 334 | 36.5 | 6.58 | 0.743 | 0.72 | 1.032 | -0.43 | 0.43 | -0.47 | -0.1566667 | 0.45 | 0.05 | 0.319251653 | 0.274644108 | 0.763095603 | 0.950027282 | 0.955820031 | 0.952923656 | 0.992535481 |
| Atp1b3 | High | P97370 | Sodium/potassium-transporting ATPase subunit beta-3 OS=Mus musculus OX=10090 GN=Atp1b3 | 0 | 46.536 | 45 | 10 | 18 | 10 | 278 | 31.8 | 8.51 | 0.742 | 0.724 | 1.024 | -0.43 | 0.43 | -0.47 | -0.1566667 | 0.45 | 0.03 | 0.316376263 | 0.282098727 | 0.798937868 | 0.950027282 | 0.955820031 | 0.952923656 | 0.992535481 |
| Ogfr | High | Q99PG2 | Opioid growth factor receptor OS=Mus musculus OX=10090 GN=Ogfr PE=1 SV=1 | 0 | 8.226 | 4 | 2 | 2 | 2 | 633 | 70.6 | 4.77 | 0.78 | 0.742 | 1.052 | -0.36 | 0.36 | -0.43 | -0.1433333 | 0.395 | 0.07 | 0.319586153 | 0.306846426 | 0.756120942 | 0.950027282 | 0.955820031 | 0.952923656 | 0.992535481 |
| Aimp2 | High | Q8R010 | Aminoacyl tRNA synthase complex-interacting multifunctional protein 2 OS=Mus musculus OX=1( | 0 | 19.655 | 13 | 3 | 4 | 3 | 320 | 35.4 | 7.83 | 1.316 | 1.427 | 0.922 | 0.4 | -0.4 | 0.51 | 0.17 | -0.455 | -0.12 | 0.31912896 | 0.314783743 | 0.672133601 | 0.950027282 | 0.955820031 | 0.952923656 | 0.990250803 |
| Cdhr1 | High | Q8VHP6 | Cadherin-related family member 1 OS=Mus musculus OX=10090 GN=Cdhr1 PE=1 SV=1 | 0 | 207.104 | 42 | 23 | 70 | 23 | 859 | 94 | 5.73 | 1.341 | 1.436 | 0.933 | 0.42 | -0.42 | 0.52 | 0.17333333 | -0.47 | -0.1 | 0.322340816 | 0.305424397 | 0.730754306 | 0.950027282 | 0.955820031 | 0.952923656 | 0.992535481 |
| Dync1i1 | Medium | Q3TYJ3 | Cytoplasmic dynein 1 intermediate chain 1 OS=Mus musculus OX=10090 GN=Dync1i1 PE=1 SV=1 | 0.011 | 2.485 | 3 | 1 | 1 | 1 | 617 | 69.6 | 5.08 | 1.328 | 1.405 | 0.945 | 0.41 | -0.41 | 0.49 | 0.16333333 | -0.45 | -0.08 | 0.32003342 | 0.337573938 | 0.817077595 | 0.950027282 | 0.976045759 | 0.96303652 | 0.992535481 |
| Ppp3r1 | High | Q63810 | Calcineurin subunit B type 1 OS=Mus musculus OX=10090 GN=Ppp3r1 PE=1 SV=3 | 0 | 120.379 | 74 | 12 | 41 | 12 | 170 | 19.3 | 4.81 | 0.744 | 0.761 | 0.977 | -0.43 | 0.43 | -0.39 | -0.13 | 0.41 | -0.03 | 0.320638316 | 0.356420857 | 0.959492324 | 0.950027282 | 0.977313184 | 0.963670233 | 0.992962986 |
| Fam120c | High | Q8C3F2 | Constitutive coactivator of PPAR-gamma-like protein 2 OS=Mus musculus OX=10090 GN=Fam120 | 0 | 13.313 | 4 | 4 | 4 | 4 | 1091 | 119.6 | 8.46 | 1.332 | 1.358 | 0.981 | 0.41 | -0.41 | 0.44 | 0.14666667 | -0.425 | -0.03 | 0.323422183 | 0.384741212 | 0.947467218 | 0.950027282 | 0.977313184 | 0.963670233 | 0.992535481 |
| | High | A0A0A0M | Tubulin alpha chain (Fragment) OS=Mus musculus OX=10090 GN=Tuba4a PE=1 SV=1 | 0 | 364.922 | 58 | 25 | 160 | 6 | 477 | 52.9 | 5.19 | 1.344 | 1.387 | 0.969 | 0.43 | -0.43 | 0.47 | 0.15666667 | -0.45 | -0.05 | 0.318276703 | 0.358381108 | 0.916742175 | 0.950027282 | 0.977313184 | 0.963670233 | 0.992535481 |
| Mvp | High | Q9EQK5 | Major vault protein OS=Mus musculus OX=10090 GN=Mvp PE=1 SV=4 | 0.001 | 5.878 | 4 | 1 | 1 | 1 | 861 | 95.9 | 5.59 | 1.364 | 1.374 | 0.993 | 0.45 | -0.45 | 0.46 | 0.15333333 | -0.455 | -0.01 | 0.320551669 | 0.374922024 | 0.987159859 | 0.950027282 | 0.977313184 | 0.963670233 | 0.998907549 |
| Npc1 | Medium | O35604 | NPC intracellular cholesterol transporter 1 OS=Mus musculus OX=10090 GN=Npc1 PE=1 SV=2 | 0.019 | 2.055 | 1 | 1 | 1 | 1 | 1277 | 142.8 | 5.71 | 0.8 | 0.792 | 1.01 | -0.32 | 0.32 | -0.34 | -0.1133333 | 0.33 | 0.01 | 0.323204114 | 0.417550161 | 0.93907159 | 0.950027282 | 0.99399649 | 0.972011886 | 0.992535481 |
| Pvrl1; Nec | High | Q9JKF6 | Nectin-1 OS=Mus musculus OX=10090 GN=Nectin1 PE=1 SV=3 | 0 | 28.302 | 16 | 7 | 11 | 7 | 515 | 57 | 6.35 | 0.776 | 0.844 | 0.92 | -0.37 | 0.37 | -0.24 | -0.08 | 0.305 | -0.12 | 0.321063594 | 0.544160091 | 0.604781054 | 0.950027282 | 0.995559182 | 0.972793232 | 0.97826833 |
| Prpsap1 | High | Q9D0M1 | Phosphoribosyl pyrophosphate synthase-associated protein 1 OS=Mus musculus OX=10090 GN=Pr | 0 | 17.964 | 16 | 5 | 7 | 3 | 356 | 39.4 | 7.2 | 0.772 | 0.891 | 0.866 | -0.37 | 0.37 | -0.17 | -0.0566667 | 0.27 | -0.21 | 0.318117114 | 0.63793227 | 0.515215421 | 0.950027282 | 0.995559182 | 0.972793232 | 0.964378027 |
| | Medium | A0A1Y7VN | Eukaryotic translation initiation factor 5 (Fragment) OS=Mus musculus OX=10090 GN=Eif5 PE=1 S | 0.018 | 2.098 | 6 | 1 | 1 | 1 | 146 | 16.5 | 8.94 | 0.795 | 0.876 | 0.907 | -0.33 | 0.33 | -0.19 | -0.0633333 | 0.26 | -0.14 | 0.3186434 | 0.622409036 | 0.580455444 | 0.950027282 | 0.995559182 | 0.972793232 | 0.972910028 |
| Arfgap3 | High | A0A0R4J0 | ADP-ribosylation factor GTPase-activating protein 3 OS=Mus musculus OX=10090 GN=Arfgap3 PE | 0 | 8.138 | 5 | 3 | 3 | 2 | 524 | 57.5 | 8.47 | 0.788 | 0.906 | 0.87 | -0.34 | 0.34 | -0.14 | -0.0466667 | 0.24 | -0.2 | 0.324256125 | 0.672758993 | 0.46151717 | 0.950027282 | 0.995559182 | 0.972793232 | 0.959744396 |
| Rdh5 | High | O55240 | Retinol dehydrogenase 5 OS=Mus musculus OX=10090 GN=Rdh5 PE=1 SV=1 | 0 | 22.647 | 26 | 7 | 7 | 7 | 318 | 34.8 | 8.48 | 0.758 | 1.156 | 0.656 | -0.4 | 0.4 | 0.21 | 0.07 | 0.095 | -0.61 | 0.322221715 | 0.720494952 | 0.017118479 | 0.950027282 | 0.995559182 | 0.972793232 | 0.229077117 |
| | Medium | A0A140LH | Mitotic checkpoint protein BUB3 OS=Mus musculus OX=10090 GN=Bub3 PE=1 SV=1 | 0.011 | 2.386 | 3 | 1 | 1 | 1 | 324 | 36.7 | 6.84 | 0.798 | 1.131 | 0.705 | -0.33 | 0.33 | 0.18 | 0.06 | 0.075 | -0.5 | 0.320229259 | 0.760338383 | 0.058263531 | 0.950027282 | 0.995559182 | 0.972793232 | 0.4657565 |
| | Medium | A0A0A6YY | TRAF3-interacting JNK-activating modulator (Fragment) OS=Mus musculus OX=10090 GN=Traf3i | 0.011 | 2.48 | 8 | 1 | 1 | 1 | 147 | 17 | 5 | 1.309 | 1.143 | 1.145 | 0.39 | -0.39 | 0.19 | 0.06333333 | -0.29 | 0.2 | 0.320768056 | 0.736526384 | 0.425151326 | 0.950027282 | 0.995559182 | 0.972793232 | 0.941180024 |
| Rpl26; Gm | High | P61255 | 60S ribosomal protein L26 OS=Mus musculus OX=10090 GN=Rpl26 PE=1 SV=1 | 0 | 20.35 | 39 | 7 | 10 | 7 | 145 | 17.2 | 10.55 | 1.338 | 1.191 | 1.123 | 0.42 | -0.42 | 0.25 | 0.08333333 | -0.335 | 0.17 | 0.318961854 | 0.652563621 | 0.374958823 | 0.950027282 | 0.995559182 | 0.972793232 | 0.92018107 |
| Rps13; Rp | High | P62301 | 40S ribosomal protein S13 OS=Mus musculus OX=10090 GN=Rps13 PE=1 SV=2 | 0 | 33.599 | 50 | 9 | 12 | 9 | 151 | 17.2 | 10.54 | 1.314 | 1.22 | 1.077 | 0.39 | -0.39 | 0.29 | 0.09666667 | -0.34 | 0.11 | 0.32463925 | 0.600701998 | 0.562671963 | 0.950027282 | 0.995559182 | 0.972793232 | 0.972910028 |
| Rnf114 | Medium | Q9ET26 | E3 ubiquitin-protein ligase RNF114 OS=Mus musculus OX=10090 GN=Rnf114 PE=1 SV=2 | 0.042 | 1.61 | 3 | 1 | 1 | 1 | 229 | 25.7 | 7.03 | 1.321 | 1.227 | 1.077 | 0.4 | -0.4 | 0.3 | 0.1 | -0.35 | 0.11 | 0.321442516 | 0.591380056 | 0.598818885 | 0.950027282 | 0.995559182 | 0.972793232 | 0.977451587 |
| Esd | High | Q9R0P3 | S-formylglutathione hydrolase OS=Mus musculus OX=10090 GN=Esd PE=1 SV=1 | 0 | 37.563 | 50 | 7 | 8 | 7 | 282 | 31.3 | 7.12 | 0.757 | 1.025 | 0.738 | -0.4 | 0.4 | 0.04 | 0.01333333 | 0.18 | -0.44 | 0.324252745 | 0.992509854 | 0.089988902 | 0.950027282 | 0.998402594 | 0.974214938 | 0.5746627 |
| Mtch1 | High | Q791T5 | Mitochondrial carrier homolog 1 OS=Mus musculus OX=10090 GN=Mtch1 PE=1 SV=1 | 0.001 | 6.758 | 5 | 2 | 2 | 2 | 389 | 41.5 | 9.32 | 0.796 | 0.988 | 0.805 | -0.33 | 0.33 | -0.02 | -0.0066667 | 0.175 | -0.31 | 0.316868706 | 0.904083431 | 0.225757923 | 0.950027282 | 0.998402594 | 0.974214938 | 0.813917005 |
| Nlgn3 | High | A2AGI2 | Neuroligin-3 OS=Mus musculus OX=10090 GN=Nlgn3 PE=1 SV=1 | 0.005 | 3.627 | 2 | 1 | 1 | 1 | 845 | 93.4 | 6.28 | 0.79 | 1.003 | 0.788 | -0.34 | 0.34 | 0 | 0 | 0.17 | -0.34 | 0.323000091 | 0.93915751 | 0.154588994 | 0.950027282 | 0.998402594 | 0.974214938 | 0.728999569 |
| Nucks1 | High | A0A087W | Nuclear ubiquitous casein and cyclin-dependent kinase substrate 1 OS=Mus musculus OX=10090 ( | 0.005 | 3.798 | 7 | 2 | 2 | 2 | 233 | 26.2 | 5.12 | 1.337 | 0.977 | 1.368 | 0.42 | -0.42 | -0.03 | -0.01 | -0.195 | 0.45 | 0.317650926 | 0.875991331 | 0.050897552 | 0.950027282 | 0.998402594 | 0.974214938 | 0.427566646 |
| Myrip | High | Q8K3I4 | Rab effector MyRIP OS=Mus musculus OX=10090 GN=Myrip PE=1 SV=1 | 0 | 19.682 | 8 | 6 | 8 | 6 | 856 | 94.9 | 6.07 | 1.287 | 1.101 | 1.169 | 0.36 | -0.36 | 0.14 | 0.04666667 | -0.25 | 0.23 | 0.315959218 | 0.833667992 | 0.369093381 | 0.950027282 | 0.998402594 | 0.974214938 | 0.920164724 |
| Pacsin1 | High | Q61644 | Protein kinase C and casein kinase substrate in neurons protein 1 OS=Mus musculus OX=10090 GI | 0 | 105.716 | 44 | 21 | 34 | 19 | 441 | 50.5 | 5.24 | 0.746 | 0.827 | 0.902 | -0.42 | 0.42 | -0.27 | -0.09 | 0.345 | -0.15 | 0.325282825 | 0.504361677 | 0.570375029 | 0.951116084 | 0.995559182 | 0.973337633 | 0.972910028 |
| | High | A0A3B2W | Ribosomal protein OS=Mus musculus OX=10090 GN=Rpl10a PE=1 SV=1 | 0 | 47.625 | 44 | 9 | 21 | 9 | 190 | 21.6 | 9.98 | 1.338 | 1.155 | 1.158 | 0.42 | -0.42 | 0.21 | 0.07 | -0.315 | 0.21 | 0.326198721 | 0.721655648 | 0.295694794 | 0.951116084 | 0.995559182 | 0.973337633 | 0.894411787 |
| Sri | High | Q6P069 | Sorcin OS=Mus musculus OX=10090 GN=Sri PE=1 SV=1 | 0 | 11.638 | 11 | 2 | 5 | 2 | 198 | 21.6 | 5.59 | 0.76 | 0.95 | 0.8 | -0.4 | 0.4 | -0.07 | -0.0233333 | 0.235 | -0.32 | 0.325552765 | 0.807837785 | 0.225990298 | 0.951116084 | 0.998402594 | 0.974759339 | 0.813917005 |
| Psip1 | High | Q99JF8 | PC4 and SFRS1-interacting protein OS=Mus musculus OX=10090 GN=Psip1 PE=1 SV=1 | 0 | 19.213 | 13 | 6 | 6 | 5 | 528 | 59.7 | 9.13 | 1.354 | 1.08 | 1.254 | 0.44 | -0.44 | 0.11 | 0.03666667 | -0.275 | 0.33 | 0.325829228 | 0.879826277 | 0.167998922 | 0.951116084 | 0.998402594 | 0.974759339 | 0.738806217 |
| Paip1 | High | F6Y616 | Polyadenylate-binding protein-interacting protein 1 OS=Mus musculus OX=10090 GN=Paip1 PE=1 | 0.001 | 5.009 | 5 | 2 | 2 | 2 | 484 | 53.9 | 4.78 | 0.778 | 0.835 | 0.932 | -0.36 | 0.36 | -0.26 | -0.0866667 | 0.31 | -0.1 | 0.326771081 | 0.515456578 | 0.701007256 | 0.952289737 | 0.995559182 | 0.973924459 | 0.991285312 |
| Grm2 | High | A0A0J9YV | Metabotropic glutamate receptor 2 OS=Mus musculus OX=10090 GN=Grm2 PE=1 SV=1 | 0.001 | 4.363 | 2 | 1 | 1 | 1 | 594 | 65.7 | 8.28 | 0.795 | 0.885 | 0.899 | -0.33 | 0.33 | -0.18 | -0.06 | 0.255 | -0.15 | 0.32713638 | 0.625763804 | 0.637036982 | 0.952364321 | 0.995559182 | 0.973961751 | 0.983597794 |
| | Medium | D3YWY6 | Mitochondrial pyruvate carrier OS=Mus musculus OX=10090 GN=Mpc1 PE=1 SV=1 | 0.011 | 2.516 | 9 | 1 | 1 | 1 | 85 | 9.7 | 9.39 | 1.313 | 1.468 | 0.894 | 0.39 | -0.39 | 0.55 | 0.18333333 | -0.47 | -0.16 | 0.32807818 | 0.261184191 | 0.528735387 | 0.952633018 | 0.951072422 | 0.95185272 | 0.965674043 |
| Rps3 | High | P62908 | 40S ribosomal protein S3 OS=Mus musculus OX=10090 GN=Rps3 PE=1 SV=1 | 0 | 80.363 | 74 | 17 | 29 | 17 | 243 | 26.7 | 9.66 | 1.336 | 1.156 | 1.157 | 0.42 | -0.42 | 0.21 | 0.07 | -0.315 | 0.21 | 0.327637689 | 0.7206885 | 0.299544576 | 0.952633018 | 0.995559182 | 0.9740961 | 0.900241969 |
| Cdc42 | High | P60766 | Cell division control protein 42 homolog OS=Mus musculus OX=10090 GN=Cdc42 PE=1 SV=2 | 0 | 71.914 | 42 | 8 | 27 | 7 | 191 | 21.2 | 6.55 | 0.747 | 0.737 | 1.014 | -0.42 | 0.42 | -0.44 | -0.1466667 | 0.43 | 0.02 | 0.328746646 | 0.306273328 | 0.847146746 | 0.9530933 | 0.955820031 | 0.954456666 | 0.992535481 |
| Pip5k1c | High | O70161 | Phosphatidylinositol 4-phosphate 5-kinase type-1 gamma OS=Mus musculus OX=10090 GN=Pip5k | 0 | 11.459 | 6 | 4 | 4 | 4 | 661 | 72.4 | 5.59 | 0.792 | 0.902 | 0.878 | -0.34 | 0.34 | -0.15 | -0.05 | 0.245 | -0.19 | 0.32863142 | 0.666845904 | 0.466373836 | 0.9530933 | 0.995559182 | 0.974326241 | 0.960359242 |
| Fbxo6 | High | Q9QZN4 | F-box only protein 6 OS=Mus musculus OX=10090 GN=Fbxo6 PE=1 SV=1 | 0 | 20.294 | 26 | 6 | 8 | 6 | 295 | 34.5 | 8.18 | 0.77 | 0.786 | 0.979 | -0.38 | 0.38 | -0.35 | -0.1166667 | 0.365 | -0.03 | 0.329210223 | 0.409540406 | 0.928240075 | 0.953944041 | 0.9855335 | 0.96973877 | 0.992535481 |
| Prdx3 | High | P20108 | Thioredoxin-dependent peroxide reductase, mitochondrial OS=Mus musculus OX=10090 GN=Prdx | 0 | 15.85 | 31 | 5 | 5 | 5 | 257 | 28.1 | 7.58 | 1.303 | 1.212 | 1.075 | 0.38 | -0.38 | 0.28 | 0.09333333 | -0.33 | 0.1 | 0.329575378 | 0.613507068 | 0.625147072 | 0.954508855 | 0.995559182 | 0.975034019 | 0.979106043 |
| Sdha | High | Q8K2B3 | Succinate dehydrogenase [ubiquinone] flavoprotein subunit, mitochondrial OS=Mus musculus OX= | 0 | 65.162 | 31 | 13 | 18 | 13 | 664 | 72.5 | 7.37 | 1.301 | 1.394 | 0.933 | 0.38 | -0.38 | 0.48 | 0.16 | -0.43 | -0.1 | 0.331149459 | 0.350642427 | 0.728186402 | 0.957128875 | 0.977313184 | 0.96722103 | 0.992535481 |
| Cst3 | High | P21460 | Cystatin-C OS=Mus musculus OX=10090 GN=Cst3 PE=1 SV=2 | 0 | 20.875 | 36 | 4 | 8 | 4 | 140 | 15.5 | 9 | 0.795 | 0.777 | 1.023 | -0.33 | 0.33 | -0.36 | -0.12 | 0.345 | 0.03 | 0.333018373 | 0.390362028 | 0.89980332 | 0.957128875 | 0.983542228 | 0.970335552 | 0.992535481 |
| Rpsa; LOC | High | P14206 | 40S ribosomal protein SA OS=Mus musculus OX=10090 GN=Rpsa PE=1 SV=4 | 0 | 61.599 | 41 | 9 | 18 | 9 | 295 | 32.8 | 4.87 | 1.332 | 1.345 | 0.99 | 0.41 | -0.41 | 0.43 | 0.14333333 | -0.42 | -0.01 | 0.333588916 | 0.410919553 | 0.969697159 | 0.957128875 | 0.986643122 | 0.971885999 | 0.995598794 |
| Mpp6 | High | Q9JLB0 | Protein PALS2 OS=Mus musculus OX=10090 GN=Pals2 PE=1 SV=1 | 0 | 31.454 | 13 | 7 | 8 | 5 | 553 | 62.6 | 6.4 | 0.749 | 0.842 | 0.889 | -0.42 | 0.42 | -0.25 | -0.0833333 | 0.335 | -0.17 | 0.331225529 | 0.540755213 | 0.503532121 | 0.957128875 | 0.995559182 | 0.976344028 | 0.964378027 |
| | High | A0A5F8MI | Mitogen-activated protein kinase OS=Mus musculus OX=10090 GN=Mapk10 PE=1 SV=1 | 0.001 | 6.432 | 5 | 2 | 2 | 2 | 478 | 54.1 | 7.27 | 0.775 | 0.868 | 0.893 | -0.37 | 0.37 | -0.2 | -0.0666667 | 0.285 | -0.16 | 0.334064778 | 0.603288771 | 0.514148785 | 0.957128875 | 0.995559182 | 0.976344028 | 0.964378027 |
| Aco2 | High | Q99KI0 | Aconitate hydratase, mitochondrial OS=Mus musculus OX=10090 GN=Aco2 PE=1 SV=1 | 0 | 215.484 | 52 | 34 | 69 | 34 | 780 | 85.4 | 7.93 | 1.332 | 1.25 | 1.066 | 0.41 | -0.41 | 0.32 | 0.10666667 | -0.365 | 0.09 | 0.333497767 | 0.549892535 | 0.610230472 | 0.957128875 | 0.995559182 | 0.976344028 | 0.97826833 |
| Fundc2 | High | Q9D6K8 | FUN14 domain-containing protein 2 OS=Mus musculus OX=10090 GN=Fundc2 PE=1 SV=1 | 0 | 8.294 | 9 | 1 | 1 | 1 | 151 | 16.6 | 9.7 | 1.3 | 1.287 | 1.01 | 0.38 | -0.38 | 0.36 | 0.12 | -0.37 | 0.01 | 0.332622813 | 0.484140145 | 0.902575088 | 0.957128875 | 0.995559182 | 0.976344028 | 0.992535481 |
| | High | D6RGM3 | Echinoderm microtubule-associated protein-like 2 OS=Mus musculus OX=10090 GN=Eml2 PE=1 S | 0.001 | 5.512 | 4 | 2 | 2 | 2 | 427 | 46.7 | 6.62 | 0.794 | 1.064 | 0.747 | -0.33 | 0.33 | 0.09 | 0.03 | 0.12 | -0.42 | 0.332598851 | 0.929536577 | 0.103616681 | 0.957128875 | 0.998402594 | 0.977765735 | 0.612171613 |
| | High | A0A087W | SH3 domain-binding glutamic acid-rich-like protein OS=Mus musculus OX=10090 GN=Sh3bgrl2 PE | 0 | 16.65 | 41 | 4 | 6 | 4 | 120 | 13.5 | 8.43 | 1.31 | 0.972 | 1.347 | 0.39 | -0.39 | -0.04 | -0.0133333 | -0.175 | 0.43 | 0.332061839 | 0.863100505 | 0.076119727 | 0.957128875 | 0.998402594 | 0.977765735 | 0.523043272 |
| Hdgfrp3; H | High | Q9JMG7 | Hepatoma-derived growth factor-related protein 3 OS=Mus musculus OX=10090 GN=Hdgfl3 PE=1 | 0 | 4.769 | 8 | 2 | 2 | 1 | 202 | 22.4 | 8.4 | 1.282 | 1.118 | 1.147 | 0.36 | -0.36 | 0.16 | 0.05333333 | -0.26 | 0.2 | 0.332645445 | 0.798076563 | 0.436449153 | 0.957128875 | 0.998402594 | 0.977765735 | 0.949727902 |
| Slc25a10; | Low | Q9QZD8 | Mitochondrial dicarboxylate carrier OS=Mus musculus OX=10090 GN=Slc25a10 PE=1 SV=2 | 0.06 | 1.331 | 3 | 1 | 1 | 1 | 287 | 31.7 | 9.32 | 1.307 | 1.683 | 0.777 | 0.39 | -0.39 | 0.75 | 0.25 | -0.57 | -0.36 | 0.334911379 | 0.118979393 | 0.160437301 | 0.959064402 | 0.786053068 | 0.872558735 | 0.733684585 |
| St13 | High | Q99L47 | Hsc70-interacting protein OS=Mus musculus OX=10090 GN=St13 PE=1 SV=1 | 0 | 67.285 | 36 | 14 | 27 | 14 | 371 | 41.6 | 5.26 | 0.75 | 0.83 | 0.904 | -0.41 | 0.41 | -0.27 | -0.09 | 0.34 | -0.15 | 0.335478688 | 0.511182193 | 0.581937185 | 0.959219276 | 0.995559182 | 0.977389229 | 0.972910028 |
| Plscr1 | High | Q9JJ00 | Phospholipid scramblase 1 OS=Mus musculus OX=10090 GN=Plscr1 PE=1 SV=1 | 0 | 9.601 | 11 | 3 | 3 | 3 | 328 | 35.9 | 5.06 | 0.817 | 0.882 | 0.926 | -0.29 | 0.29 | -0.18 | -0.06 | 0.235 | -0.11 | 0.335341891 | 0.621469285 | 0.666101114 | 0.959219276 | 0.995559182 | 0.977389229 | 0.987489386 |
| Usp2 | High | O88623 | Ubiquitin carboxyl-terminal hydrolase 2 OS=Mus musculus OX=10090 GN=Usp2 PE=1 SV=2 | 0.001 | 5.808 | 3 | 1 | 1 | 1 | 613 | 68.8 | 9.23 | 1.333 | 1.04 | 1.282 | 0.41 | -0.41 | 0.06 | 0.02 | -0.235 | 0.36 | 0.335143756 | 0.994722505 | 0.201749412 | 0.959219276 | 0.998402594 | 0.978810935 | 0.78697088 |
| Rcvrn | High | P34057 | Recoverin OS=Mus musculus OX=10090 GN=Rcvrn PE=1 SV=2 | 0 | 159.876 | 83 | 19 | 106 | 19 | 202 | 23.4 | 5.12 | 0.751 | 0.518 | 1.451 | -0.41 | 0.41 | -0.95 | -0.3166667 | 0.68 | 0.54 | 0.336373646 | 0.035193703 | 0.012500077 | 0.960108302 | 0.476392832 | 0.718250567 | 0.183957829 |
| Pde6g | High | P09174 | Retinal rod rhodopsin-sensitive cGMP 3',5'-cyclic phosphodiesterase subunit gamma OS=Mus mu | 0.002 | 4.221 | 17 | 2 | 6 | 2 | 87 | 9.6 | 9.48 | 1.273 | 1.426 | 0.893 | 0.35 | -0.35 | 0.51 | 0.17 | -0.43 | -0.16 | 0.33635171 | 0.315607799 | 0.524724634 | 0.960108302 | 0.955820031 | 0.957964167 | 0.964378027 |
| Uba3 | High | Q8C878 | NEDD8-activating enzyme E1 catalytic subunit OS=Mus musculus OX=10090 GN=Uba3 PE=1 SV=2 | 0 | 10.295 | 10 | 3 | 3 | 3 | 462 | 51.7 | 5.45 | 1.308 | 1.314 | 0.995 | 0.39 | -0.39 | 0.39 | 0.13 | -0.39 | -0.01 | 0.336474552 | 0.448869281 | 0.99220682 | 0.960108302 | 0.994630833 | 0.977369568 | 0.999783808 |
| Arl8a | High | Q8VEH3 | ADP-ribosylation factor-like protein 8A OS=Mus musculus OX=10090 GN=Arl8a PE=1 SV=1 | 0 | 106.278 | 56 | 12 | 29 | 7 | 186 | 21.4 | 7.77 | 1.328 | 1.804 | 0.736 | 0.41 | -0.41 | 0.85 | 0.28333333 | -0.63 | -0.44 | 0.338191206 | 0.084769487 | 0.060997024 | 0.960606936 | 0.682905914 | 0.821756425 | 0.476099813 |
| Pde6a | High | Q8K0A8 | Phosphodiesterase OS=Mus musculus OX=10090 GN=Pde6a PE=1 SV=1 | 0 | 1309.049 | 83 | 82 | 1092 | 61 | 860 | 99.5 | 5.55 | 1.328 | 1.486 | 0.894 | 0.41 | -0.41 | 0.57 | 0.19 | -0.49 | -0.16 | 0.338168382 | 0.258798706 | 0.532332543 | 0.960606936 | 0.951072422 | 0.955839679 | 0.96694628 |
| Atp5e | High | P56382 | ATP synthase subunit epsilon, mitochondrial OS=Mus musculus OX=10090 GN=Atp5f1e PE=1 SV= | 0 | 7.901 | 58 | 4 | 11 | 4 | 52 | 5.8 | 10.01 | 1.336 | 1.462 | 0.914 | 0.42 | -0.42 | 0.55 | 0.18333333 | -0.485 | -0.13 | 0.337002828 | 0.280971677 | 0.6158788 | 0.960606936 | 0.955820031 | 0.958213483 | 0.97826833 |
| Slc25a11 | High | Q9CR62 | Mitochondrial 2-oxoglutarate/malate carrier protein OS=Mus musculus OX=10090 GN=Slc25a11 I | 0 | 41.258 | 27 | 7 | 9 | 7 | 314 | 34.1 | 9.94 | 1.369 | 1.431 | 0.957 | 0.45 | -0.45 | 0.52 | 0.17333333 | -0.485 | -0.06 | 0.338185202 | 0.310631419 | 0.790787 | 0.960606936 | 0.955820031 | 0.958213483 | 0.992535481 |
| Tmem67 | High | E9QNI1 | Meckelin OS=Mus musculus OX=10090 GN=Tmem67 PE=1 SV=1 | 0 | 48.16 | 14 | 13 | 19 | 13 | 995 | 112 | 7.74 | 0.736 | 0.97 | 0.759 | -0.44 | 0.44 | -0.04 | -0.0133333 | 0.24 | -0.4 | 0.337366132 | 0.856982271 | 0.093590718 | 0.960606936 | 0.998402594 | 0.979504765 | 0.582204115 |
| Mogs | High | Q80UM7 | Mannosyl-oligosaccharide glucosidase OS=Mus musculus OX=10090 GN=Mogs PE=1 SV=1 | 0 | 10.776 | 4 | 3 | 3 | 3 | 834 | 91.8 | 9 | 1.285 | 1.558 | 0.825 | 0.36 | -0.36 | 0.64 | 0.21333333 | -0.5 | -0.28 | 0.338850905 | 0.188009229 | 0.276060931 | 0.96199343 | 0.902623507 | 0.932308468 | 0.876067403 |
| Farsb | High | Q9WUA2 | Phenylalanine--tRNA ligase beta subunit OS=Mus musculus OX=10090 GN=Farsb PE=1 SV=2 | 0 | 16.857 | 10 | 6 | 6 | 6 | 589 | 65.7 | 7.12 | 1.368 | 0.979 | 1.397 | 0.45 | -0.45 | -0.03 | -0.01 | -0.21 | 0.48 | 0.339030487 | 0.879983482 | 0.047014293 | 0.962016164 | 0.998402594 | 0.980209379 | 0.411889287 |
| Nrxn3 | High | Q6P9K9 | Neurexin-3 OS=Mus musculus OX=10090 GN=Nrxn3 PE=1 SV=2 | 0 | 46.046 | 8 | 10 | 10 | 3 | 1571 | 173.3 | 6.14 | 0.788 | 0.764 | 1.032 | -0.34 | 0.34 | -0.39 | -0.13 | 0.365 | 0.05 | 0.340276855 | 0.361321026 | 0.885880302 | 0.962325513 | 0.977313184 | 0.969819349 | 0.992535481 |
| | High | A0A571BE | AP2-associated protein kinase 1 (Fragment) OS=Mus musculus OX=10090 GN=Aak1 PE=1 SV=1 | 0 | 30.264 | 15 | 7 | 10 | 3 | 510 | 54 | 5.01 | 0.807 | 0.784 | 1.03 | -0.31 | 0.31 | -0.35 | -0.1166667 | 0.33 | 0.04 | 0.339624397 | 0.379266693 | 0.846845025 | 0.962325513 | 0.977313184 | 0.969819349 | 0.992535481 |
| Cd82 | High | P40237 | CD82 antigen OS=Mus musculus OX=10090 GN=Cd82 PE=1 SV=1 | 0 | 21.359 | 16 | 4 | 8 | 4 | 266 | 29.6 | 5.02 | 1.284 | 1.325 | 0.969 | 0.36 | -0.36 | 0.41 | 0.13666667 | -0.385 | -0.05 | 0.340123489 | 0.436589366 | 0.857141951 | 0.962325513 | 0.994630833 | 0.978478173 | 0.992535481 |
| Egfr | High | Q01279 | Epidermal growth factor receptor OS=Mus musculus OX=10090 GN=Egfr PE=1 SV=1 | 0 | 12.872 | 3 | 4 | 4 | 4 | 1210 | 134.8 | 6.86 | 0.793 | 0.878 | 0.904 | -0.33 | 0.33 | -0.19 | -0.0633333 | 0.26 | -0.15 | 0.339829117 | 0.601895294 | 0.598892641 | 0.962325513 | 0.995559182 | 0.978942348 | 0.977451587 |
| Sepw1; Se | Low | P63300 | Selenoprotein W OS=Mus musculus OX=10090 GN=Selenow PE=1 SV=3 | 0.053 | 1.438 | 8 | 1 | 1 | 1 | 88 | 9.7 | 8.72 | 1.321 | 0.97 | 1.362 | 0.4 | -0.4 | -0.04 | -0.0133333 | -0.18 | 0.45 | 0.340340912 | 0.832677027 | 0.042208963 | 0.962325513 | 0.998402594 | 0.980364054 | 0.399120459 |
| Eif3d | Medium | O70194 | Eukaryotic translation initiation factor 3 subunit D OS=Mus musculus OX=10090 GN=Eif3d PE=1 S\ | 0.015 | 2.174 | 2 | 1 | 1 | 1 | 548 | 63.9 | 6.05 | 1.3 | 1.141 | 1.139 | 0.38 | -0.38 | 0.19 | 0.06333333 | -0.285 | 0.19 | 0.340966873 | 0.745180196 | 0.407815363 | 0.963609503 | 0.995559182 | 0.979584343 | 0.936397673 |
| Armc10 | High | Q9D0L7 | Armadillo repeat-containing protein 10 OS=Mus musculus OX=10090 GN=Armc10 PE=1 SV=1 | 0 | 23.575 | 23 | 6 | 8 | 6 | 306 | 33.3 | 7.99 | 1.329 | 1.599 | 0.831 | 0.41 | -0.41 | 0.68 | 0.22666667 | -0.545 | -0.27 | 0.341967875 | 0.176045263 | 0.268290882 | 0.964090735 | 0.875953959 | 0.920022347 | 0.869723277 |
| LOC10263 | High | P0DN34 | NADH dehydrogenase [ubiquinone] 1 beta subcomplex subunit 1 OS=Mus musculus OX=10090 GN | 0.001 | 5.483 | 30 | 2 | 2 | 2 | 57 | 7 | 8.21 | 1.322 | 1.505 | 0.878 | 0.4 | -0.4 | 0.59 | 0.19666667 | -0.495 | -0.19 | 0.341682324 | 0.237651991 | 0.446559906 | 0.964090735 | 0.937374952 | 0.950732844 | 0.958608248 |
| Idh2 | High | P54071 | Isocitrate dehydrogenase [NADP], mitochondrial OS=Mus musculus OX=10090 GN=Idh2 PE=1 SV= | 0 | 18.474 | 15 | 6 | 7 | 4 | 452 | 50.9 | 8.69 | 1.3 | 1.489 | 0.873 | 0.38 | -0.38 | 0.57 | 0.19 | -0.475 | -0.2 | 0.342242638 | 0.256117484 | 0.402064534 | 0.964090735 | 0.951072422 | 0.957581579 | 0.93262326 |
| Itgb3 | Medium | O54890 | Integrin beta-3 OS=Mus musculus OX=10090 GN=Itgb3 PE=1 SV=2 | 0.035 | 1.771 | 1 | 1 | 1 | 1 | 787 | 86.7 | 5.24 | 1.285 | 1.135 | 1.132 | 0.36 | -0.36 | 0.18 | 0.06 | -0.27 | 0.18 | 0.341879052 | 0.751675206 | 0.462666761 | 0.964090735 | 0.995559182 | 0.979824959 | 0.959744396 |
| Eprs | High | Q8CGC7 | Bifunctional glutamate/proline--tRNA ligase OS=Mus musculus OX=10090 GN=Eprs1 PE=1 SV=4 | 0 | 57.952 | 13 | 17 | 19 | 17 | 1512 | 170 | 7.66 | 1.32 | 1.282 | 1.03 | 0.4 | -0.4 | 0.36 | 0.12 | -0.38 | 0.04 | 0.342512706 | 0.499497629 | 0.771506188 | 0.964090735 | 0.995559182 | 0.979824959 | 0.992535481 |
| Ttc26 | Medium | Q8BS45 | Intraflagellar transport protein 56 OS=Mus musculus OX=10090 GN=Ttc26 PE=1 SV=1 | 0.041 | 1.682 | 1 | 1 | 1 | 1 | 554 | 64.1 | 6.8 | 1.313 | 1.114 | 1.178 | 0.39 | -0.39 | 0.16 | 0.05333333 | -0.275 | 0.24 | 0.342415464 | 0.806583937 | 0.334929258 | 0.964090735 | 0.998402594 | 0.981246665 | 0.906329766 |
| Fbxw11 | High | Q5SRY7 | F-box/WD repeat-containing protein 11 OS=Mus musculus OX=10090 GN=Fbxw11 PE=1 SV=1 | 0.005 | 3.556 | 4 | 2 | 2 | 2 | 542 | 62 | 7.12 | 1.311 | 1.601 | 0.819 | 0.39 | -0.39 | 0.68 | 0.22666667 | -0.535 | -0.29 | 0.343020492 | 0.172223964 | 0.335857769 | 0.965035575 | 0.875484829 | 0.920260202 | 0.906329766 |
| | High | F8WJ41 | 40S ribosomal protein S15a (Fragment) OS=Mus musculus OX=10090 GN=Rps15a PE=1 SV=1 | 0 | 21.367 | 53 | 6 | 14 | 6 | 108 | 12.3 | 10.15 | 1.345 | 1.233 | 1.09 | 0.43 | -0.43 | 0.3 | 0.1 | -0.365 | 0.12 | 0.343881693 | 0.577208061 | 0.617380008 | 0.966488547 | 0.995559182 | 0.981023864 | 0.97826833 |
| Pex14 | Low | Q9R0A0 | Peroxisomal membrane protein PEX14 OS=Mus musculus OX=10090 GN=Pex14 PE=1 SV=1 | 0.052 | 1.451 | 2 | 1 | 1 | 1 | 376 | 41.2 | 5.11 | 1.284 | 1.568 | 0.819 | 0.36 | -0.36 | 0.65 | 0.21666667 | -0.505 | -0.29 | 0.345132265 | 0.182181905 | 0.258330042 | 0.967398981 | 0.891355967 | 0.929377474 | 0.85616924 |
| Khnyn | High | Q80U38 | Protein KHNYN OS=Mus musculus OX=10090 GN=Khnyn PE=2 SV=2 | 0.005 | 3.812 | 2 | 1 | 1 | 1 | 671 | 74.5 | 7.87 | 0.804 | 0.679 | 1.185 | -0.31 | 0.31 | -0.56 | -0.1866667 | 0.435 | 0.24 | 0.346378306 | 0.189740951 | 0.336044715 | 0.967398981 | 0.905821916 | 0.936610449 | 0.906329766 |
| Wnk1 | Medium | P83741 | Serine/threonine-protein kinase WNK1 OS=Mus musculus OX=10090 GN=Wnk1 PE=1 SV=2 | 0.014 | 2.244 | 0 | 1 | 1 | 1 | 2377 | 250.8 | 6.43 | 0.802 | 0.735 | 1.092 | -0.32 | 0.32 | -0.45 | -0.15 | 0.385 | 0.13 | 0.344854324 | 0.285571324 | 0.592267158 | 0.967398981 | 0.955820031 | 0.961609506 | 0.97614402 |
| Prune; Pru | High | Q8BIW1 | Exopolyphosphatase PRUNE1 OS=Mus musculus OX=10090 GN=Prune1 PE=1 SV=1 | 0.006 | 3.081 | 4 | 1 | 1 | 1 | 454 | 50.2 | 5.11 | 0.791 | 0.758 | 1.043 | -0.34 | 0.34 | -0.4 | -0.1333333 | 0.37 | 0.06 | 0.345850371 | 0.343478261 | 0.794456127 | 0.967398981 | 0.977313184 | 0.972356083 | 0.992535481 |
| Hspe1 | High | Q64433 | 10 kDa heat shock protein, mitochondrial OS=Mus musculus OX=10090 GN=Hspe1 PE=1 SV=2 | 0 | 33.737 | 75 | 9 | 17 | 9 | 102 | 11 | 8.35 | 1.323 | 1.393 | 0.949 | 0.4 | -0.4 | 0.48 | 0.16 | -0.44 | -0.07 | 0.344907149 | 0.351540209 | 0.814497326 | 0.967398981 | 0.977313184 | 0.972356083 | 0.992535481 |
| Mblac2 | High | Q8BL86 | Metallo-beta-lactamase domain-containing protein 2 OS=Mus musculus OX=10090 GN=Mblac2 P | 0.001 | 7.828 | 12 | 3 | 3 | 3 | 279 | 31.2 | 6.86 | 0.809 | 0.791 | 1.023 | -0.31 | 0.31 | -0.34 | -0.1133333 | 0.325 | 0.03 | 0.344498747 | 0.416393193 | 0.853735726 | 0.967398981 | 0.99399649 | 0.980697736 | 0.992535481 |
| Cops6 | High | O88545 | COP9 signalosome complex subunit 6 OS=Mus musculus OX=10090 GN=Cops6 PE=1 SV=1 | 0 | 10.635 | 5 | 2 | 3 | 2 | 324 | 35.9 | 5.73 | 0.808 | 0.794 | 1.017 | -0.31 | 0.31 | -0.33 | -0.11 | 0.32 | 0.02 | 0.348173915 | 0.427763244 | 0.865893183 | 0.967398981 | 0.99399649 | 0.980697736 | 0.992535481 |
| Gdap1l1 | High | Q3USC7 | Ganglioside-induced differentiation-associated protein 1-like 1 OS=Mus musculus OX=10090 GN= | 0 | 15.765 | 12 | 3 | 4 | 3 | 367 | 41.9 | 6.48 | 0.8 | 0.807 | 0.991 | -0.32 | 0.32 | -0.31 | -0.1033333 | 0.315 | -0.01 | 0.345822001 | 0.430159106 | 0.998062591 | 0.967398981 | 0.99399649 | 0.980697736 | 0.999917596 |
| Uba1 | High | Q02053 | Ubiquitin-like modifier-activating enzyme 1 OS=Mus musculus OX=10090 GN=Uba1 PE=1 SV=1 | 0 | 161.023 | 33 | 28 | 45 | 28 | 1058 | 117.7 | 5.66 | 0.756 | 0.819 | 0.923 | -0.4 | 0.4 | -0.29 | -0.0966667 | 0.345 | -0.12 | 0.347425758 | 0.484359599 | 0.676997465 | 0.967398981 | 0.995559182 | 0.981479081 | 0.990892882 |
| Stx4a | High | P70452 | Syntaxin-4 OS=Mus musculus OX=10090 GN=Stx4 PE=1 SV=1 | 0 | 37.893 | 39 | 8 | 10 | 8 | 298 | 34.1 | 6.14 | 0.797 | 0.813 | 0.981 | -0.33 | 0.33 | -0.3 | -0.1 | 0.315 | -0.03 | 0.346461883 | 0.470402376 | 0.910527343 | 0.967398981 | 0.995559182 | 0.981479081 | 0.992535481 |
| Hnrnpc | High | Q9Z204 | Heterogeneous nuclear ribonucleoproteins C1/C2 OS=Mus musculus OX=10090 GN=Hnrnpc PE=1 S | 0.003 | 3.873 | 6 | 2 | 2 | 2 | 313 | 34.4 | 5.05 | 1.273 | 1.161 | 1.097 | 0.35 | -0.35 | 0.22 | 0.07333333 | -0.285 | 0.13 | 0.346886674 | 0.698278975 | 0.588125677 | 0.967398981 | 0.995559182 | 0.981479081 | 0.973095093 |
| H3f3a; H3f | High | P84244 | Histone H3.3 OS=Mus musculus OX=10090 GN=H3-3a PE=1 SV=2 | 0 | 9.733 | 35 | 3 | 3 | 3 | 136 | 15.3 | 11.27 | 1.299 | 1.259 | 1.032 | 0.38 | -0.38 | 0.33 | 0.11 | -0.355 | 0.05 | 0.348083049 | 0.526616586 | 0.839987851 | 0.967398981 | 0.995559182 | 0.981479081 | 0.992535481 |
| | High | H3BL62 | Propionyl-CoA carboxylase alpha chain, mitochondrial (Fragment) OS=Mus musculus OX=10090 G | 0.001 | 4.286 | 10 | 2 | 2 | 2 | 183 | 20.3 | 6.57 | 1.306 | 1.304 | 1.001 | 0.38 | -0.38 | 0.38 | 0.12666667 | -0.38 | 0 | 0.347382947 | 0.465834763 | 0.944799421 | 0.967398981 | 0.995559182 | 0.981479081 | 0.992535481 |
| | High | E0CYA3 | Receptor protein-tyrosine kinase OS=Mus musculus OX=10090 GN=Ephb3 PE=1 SV=2 | 0 | 20.106 | 9 | 5 | 6 | 5 | 739 | 81.5 | 6.95 | 1.305 | 1.038 | 1.257 | 0.38 | -0.38 | 0.05 | 0.01666667 | -0.215 | 0.33 | 0.34593425 | 0.971244829 | 0.194220242 | 0.967398981 | 0.998402594 | 0.982900788 | 0.784517504 |
| Raph1 | High | F2Z3U3 | Ras association (RalGDS/AF-6) and pleckstrin homology domains 1 OS=Mus musculus OX=10090 | 0.008 | 2.9 | 1 | 1 | 1 | 1 | 1266 | 136.7 | 8.98 | 1.304 | 1.045 | 1.248 | 0.38 | -0.38 | 0.06 | 0.02 | -0.22 | 0.32 | 0.346480037 | 0.964216683 | 0.221679589 | 0.967398981 | 0.998402594 | 0.982900788 | 0.807447486 |
| Ubl3 | High | Q9Z2M6 | Ubiquitin-like protein 3 OS=Mus musculus OX=10090 GN=Ubl3 PE=1 SV=1 | 0 | 40.752 | 38 | 5 | 13 | 5 | 117 | 13.2 | 6.92 | 1.287 | 1.583 | 0.813 | 0.36 | -0.36 | 0.66 | 0.22 | -0.51 | -0.3 | 0.349152528 | 0.185708842 | 0.216713828 | 0.969157538 | 0.900299151 | 0.934728344 | 0.804947026 |
| Kcnb1 | High | Q03717 | Potassium voltage-gated channel subfamily B member 1 OS=Mus musculus OX=10090 GN=Kcnb1 | 0 | 31.047 | 10 | 8 | 8 | 8 | 857 | 95.5 | 8.16 | 1.324 | 1.267 | 1.045 | 0.4 | -0.4 | 0.34 | 0.11333333 | -0.37 | 0.06 | 0.34910715 | 0.522447783 | 0.811537203 | 0.969157538 | 0.995559182 | 0.98235836 | 0.992535481 |
| Notch1 | High | Q01705 | Neurogenic locus notch homolog protein 1 OS=Mus musculus OX=10090 GN=Notch1 PE=1 SV=3 | 0.001 | 5.224 | 1 | 2 | 2 | 2 | 2531 | 270.7 | 5.1 | 0.809 | 0.851 | 0.951 | -0.31 | 0.31 | -0.23 | -0.0766667 | 0.27 | -0.07 | 0.350079814 | 0.557745344 | 0.786524819 | 0.971250628 | 0.995559182 | 0.983404905 | 0.992535481 |
| Eif4h | High | Q9WUK2 | Eukaryotic translation initiation factor 4H OS=Mus musculus OX=10090 GN=Eif4h PE=1 SV=3 | 0 | 18.836 | 23 | 4 | 6 | 4 | 248 | 27.3 | 7.23 | 0.791 | 0.655 | 1.208 | -0.34 | 0.34 | -0.61 | -0.2033333 | 0.475 | 0.27 | 0.353027431 | 0.166425584 | 0.273786626 | 0.974570736 | 0.871008827 | 0.922789781 | 0.876067403 |
| Pdha1 | High | P35486 | Pyruvate dehydrogenase E1 component subunit alpha, somatic form, mitochondrial OS=Mus musc | 0 | 15.998 | 14 | 6 | 7 | 6 | 390 | 43.2 | 8.19 | 1.273 | 1.444 | 0.882 | 0.35 | -0.35 | 0.53 | 0.17666667 | -0.44 | -0.18 | 0.353667524 | 0.297619584 | 0.447379252 | 0.974570736 | 0.955820031 | 0.965195383 | 0.958608248 |

| | | | | | | | | | | | | | | | | | | | | | | | | | | | | |
|---|---|---|---|---|---|---|---|---|---|---|---|---|---|---|---|---|---|---|---|---|---|---|---|---|---|---|---|---|
| Atf6 | Low | F6VAN0 | Cyclic AMP-dependent transcription factor ATF-6 alpha OS=Mus musculus OX=10090 GN=Atf6 PE= | 0.055 | 1.381 | 2 | 1 | 1 | 1 | 656 | 72.6 | 7.77 | 1.299 | 0.866 | 1.5 | 0.38 | -0.38 | -0.21 | -0.07 | -0.085 | 0.59 | 0.353429626 | 0.578443136 | 0.022451691 | 0.974570736 | 0.995559182 | 0.985064959 | 0.270723942 |
| | High | V9GX87 | Double-stranded RNA-binding protein Staufen homolog 1 OS=Mus musculus OX=10090 GN=Stau1 | 0 | 4.907 | 3 | 1 | 1 | 1 | 436 | 48.7 | 9.86 | 1.293 | 1.202 | 1.076 | 0.37 | -0.37 | 0.27 | 0.09 | -0.32 | 0.11 | 0.353709907 | 0.634529355 | 0.605748203 | 0.974570736 | 0.995559182 | 0.985064959 | 0.97826833 |
| Rps6-ps4; | High | P62754 | 40S ribosomal protein S6 OS=Mus musculus OX=10090 GN=Rps6 PE=1 SV=1 | 0 | 48.543 | 35 | 10 | 18 | 10 | 249 | 28.7 | 10.84 | 1.327 | 1.19 | 1.115 | 0.41 | -0.41 | 0.25 | 0.08333333 | -0.33 | 0.16 | 0.351735744 | 0.654982373 | 0.421773877 | 0.974570736 | 0.995559182 | 0.985064959 | 0.939192266 |
| Iars2 | High | Q8BIJ6 | Isoleucine--tRNA ligase, mitochondrial OS=Mus musculus OX=10090 GN=Iars2 PE=1 SV=1 | 0 | 31.422 | 11 | 8 | 9 | 8 | 1012 | 112.7 | 6.81 | 1.276 | 1.24 | 1.029 | 0.35 | -0.35 | 0.31 | 0.10333333 | -0.33 | 0.04 | 0.352731717 | 0.565967174 | 0.894357626 | 0.974570736 | 0.995559182 | 0.985064959 | 0.992535481 |
| Comtd1 | High | Q8BIG7 | Catechol O-methyltransferase domain-containing protein 1 OS=Mus musculus OX=10090 GN=Con | 0.001 | 4.418 | 7 | 2 | 2 | 2 | 262 | 28.9 | 8.32 | 1.32 | 1.237 | 1.067 | 0.4 | -0.4 | 0.31 | 0.10333333 | -0.355 | 0.09 | 0.353145307 | 0.569123191 | 0.650065033 | 0.974570736 | 0.995559182 | 0.985064959 | 0.985010103 |
| Slc4a8 | High | Q8JZR6 | Electroneutral sodium bicarbonate exchanger 1 OS=Mus musculus OX=10090 GN=Slc4a8 PE=2 SV | 0 | 23.637 | 8 | 7 | 8 | 6 | 1089 | 122.3 | 6.6 | 0.79 | 1.012 | 0.781 | -0.34 | 0.34 | 0.02 | 0.00666667 | 0.16 | -0.36 | 0.353418571 | 0.96029365 | 0.16203664 | 0.974570736 | 0.998402594 | 0.986486665 | 0.733684585 |
| Tmtc1 | Low | Q3UV71 | Protein O-mannosyl-transferase TMTC1 OS=Mus musculus OX=10090 GN=Tmtc1 PE=2 SV=2 | 0.084 | 1.183 | 1 | 1 | 1 | 1 | 942 | 105.7 | 9 | 1.301 | 1.109 | 1.173 | 0.38 | -0.38 | 0.15 | 0.05 | -0.265 | 0.23 | 0.353191636 | 0.819580657 | 0.37286108 | 0.974570736 | 0.998402594 | 0.986486665 | 0.920164724 |
| Loh12cr1; | High | E9Q2W8 | BLOC-1-related complex subunit 5 OS=Mus musculus OX=10090 GN=Borcs5 PE=1 SV=1 | 0.005 | 3.539 | 10 | 2 | 2 | 2 | 196 | 22.6 | 8.84 | 0.757 | 0.652 | 1.16 | -0.4 | 0.4 | -0.62 | -0.2066667 | 0.51 | 0.21 | 0.35449822 | 0.162813855 | 0.339091303 | 0.974787008 | 0.869444647 | 0.922115827 | 0.907534575 |
| Echs1 | High | Q8BH95 | Enoyl-CoA hydratase, mitochondrial OS=Mus musculus OX=10090 GN=Echs1 PE=1 SV=1 | 0 | 26.524 | 27 | 9 | 10 | 9 | 290 | 31.5 | 8.48 | 1.339 | 1.391 | 0.963 | 0.42 | -0.42 | 0.48 | 0.16 | -0.45 | -0.05 | 0.354989696 | 0.354168612 | 0.791769442 | 0.974787008 | 0.977313184 | 0.976050096 | 0.992535481 |
| | Medium | F7D291 | Protein MTSS 2 (Fragment) OS=Mus musculus OX=10090 GN=Mtss2 PE=1 SV=1 | 0.014 | 2.286 | 2 | 1 | 1 | 1 | 364 | 39.1 | 8 | 0.793 | 0.919 | 0.863 | -0.33 | 0.33 | -0.12 | -0.04 | 0.225 | -0.21 | 0.354945034 | 0.70966673 | 0.406444557 | 0.974787008 | 0.995559182 | 0.985173095 | 0.93629196 |
| Eif3g | High | Q9Z1D1 | Eukaryotic translation initiation factor 3 subunit G OS=Mus musculus OX=10090 GN=Eif3g PE=1 S | 0 | 14.805 | 17 | 4 | 4 | 4 | 320 | 35.6 | 5.9 | 1.294 | 1.206 | 1.073 | 0.37 | -0.37 | 0.27 | 0.09 | -0.32 | 0.1 | 0.355005363 | 0.624329269 | 0.676110945 | 0.974787008 | 0.995559182 | 0.985173095 | 0.990250803 |
| Rhoc | High | Q62159 | Rho-related GTP-binding protein RhoC OS=Mus musculus OX=10090 GN=Rhoc PE=1 SV=2 | 0 | 81.181 | 45 | 9 | 26 | 1 | 193 | 22 | 6.58 | 0.787 | 0.687 | 1.145 | -0.34 | 0.34 | -0.54 | -0.18 | 0.44 | 0.2 | 0.358491882 | 0.201994374 | 0.490834151 | 0.974893705 | 0.912882441 | 0.943888073 | 0.964378027 |
| Pde6b | High | P23440 | Rod cGMP-specific 3',5'-cyclic phosphodiesterase subunit beta OS=Mus musculus OX=10090 GN= | 0 | 1308.825 | 80 | 78 | 1071 | 57 | 856 | 98.5 | 5.31 | 1.314 | 1.504 | 0.874 | 0.39 | -0.39 | 0.59 | 0.19666667 | -0.49 | -0.19 | 0.356333384 | 0.243420868 | 0.440979699 | 0.974893705 | 0.940627712 | 0.957760709 | 0.955768524 |
| Ccm2 | High | F7AVU1 | Cerebral cavernous malformations protein 2 homolog OS=Mus musculus OX=10090 GN=Ccm2 PE= | 0 | 10.437 | 8 | 2 | 2 | 2 | 389 | 42.5 | 5 | 1.311 | 1.446 | 0.906 | 0.39 | -0.39 | 0.53 | 0.17666667 | -0.46 | -0.14 | 0.356222549 | 0.290599337 | 0.568599289 | 0.974893705 | 0.955820031 | 0.965356868 | 0.972910028 |
| | High | D3YX62 | Heme oxygenase (biliverdin-producing) (Fragment) OS=Mus musculus OX=10090 GN=Hmox2 PE= | 0 | 14.941 | 26 | 6 | 6 | 6 | 228 | 26.4 | 5.86 | 1.311 | 1.403 | 0.934 | 0.39 | -0.39 | 0.49 | 0.16333333 | -0.44 | -0.1 | 0.358712259 | 0.340543373 | 0.697795746 | 0.974893705 | 0.976045759 | 0.975469732 | 0.991285312 |
| Rab28 | High | A0A0J9YV | Ras-related protein Rab-28 OS=Mus musculus OX=10090 GN=Rab28 PE=1 SV=1 | 0 | 114.367 | 65 | 12 | 45 | 2 | 220 | 24.7 | 5.47 | 1.314 | 1.385 | 0.949 | 0.39 | -0.39 | 0.47 | 0.15666667 | -0.43 | -0.08 | 0.356055564 | 0.360895235 | 0.811643381 | 0.974893705 | 0.977313184 | 0.976103445 | 0.992535481 |
| Ccdc124 | High | Q9D8X2 | Coiled-coil domain-containing protein 124 OS=Mus musculus OX=10090 GN=Ccdc124 PE=1 SV=1 | 0 | 13.632 | 18 | 5 | 6 | 5 | 217 | 25.3 | 9.64 | 1.28 | 1.358 | 0.943 | 0.36 | -0.36 | 0.44 | 0.14666667 | -0.4 | -0.09 | 0.358436114 | 0.393468955 | 0.710407954 | 0.974893705 | 0.983884835 | 0.97938927 | 0.992535481 |
| Vps45 | High | P97390 | Vacuolar protein sorting-associated protein 45 OS=Mus musculus OX=10090 GN=Vps45 PE=1 SV= | 0 | 11.112 | 5 | 3 | 4 | 3 | 570 | 65 | 8.25 | 0.778 | 0.858 | 0.907 | -0.36 | 0.36 | -0.22 | -0.0733333 | 0.29 | -0.14 | 0.359564862 | 0.577978505 | 0.577370382 | 0.974893705 | 0.995559182 | 0.985226443 | 0.972910028 |
| Zdhhc2 | High | P59267 | Palmitoyltransferase ZDHHC2 OS=Mus musculus OX=10090 GN=Zdhhc2 PE=1 SV=1 | 0 | 71.361 | 41 | 13 | 39 | 11 | 366 | 42 | 8.48 | 0.76 | 1.149 | 0.661 | -0.4 | 0.4 | 0.2 | 0.06666667 | 0.1 | -0.6 | 0.356879353 | 0.73347081 | 0.010275763 | 0.974893705 | 0.995559182 | 0.985226443 | 0.16229917 |
| Cplx2 | High | P84086 | Complexin-2 OS=Mus musculus OX=10090 GN=Cplx2 PE=1 SV=1 | 0 | 17.583 | 30 | 3 | 7 | 3 | 134 | 15.4 | 5.08 | 0.794 | 1.178 | 0.673 | -0.33 | 0.33 | 0.24 | 0.08 | 0.045 | -0.57 | 0.357666649 | 0.676075102 | 0.014318522 | 0.974893705 | 0.995559182 | 0.985226443 | 0.197218435 |
| Srsf6 | High | Q3TWW8 | Serine/arginine-rich splicing factor 6 OS=Mus musculus OX=10090 GN=Srsf6 PE=1 SV=1 | 0 | 9.612 | 9 | 3 | 3 | 1 | 339 | 39 | 11.46 | 1.317 | 0.912 | 1.444 | 0.4 | -0.4 | -0.13 | -0.0433333 | -0.135 | 0.53 | 0.357124257 | 0.689129859 | 0.059692586 | 0.974893705 | 0.995559182 | 0.985226443 | 0.472734926 |
| Rps16-ps2 | High | P14131 | 40S ribosomal protein S16 OS=Mus musculus OX=10090 GN=Rps16 PE=1 SV=4 | 0 | 44.61 | 55 | 12 | 19 | 12 | 146 | 16.4 | 10.21 | 1.315 | 1.14 | 1.153 | 0.39 | -0.39 | 0.19 | 0.06333333 | -0.29 | 0.21 | 0.355798801 | 0.752701479 | 0.307560432 | 0.974893705 | 0.995559182 | 0.985226443 | 0.904295409 |
| Nek7 | High | Q3TN15 | Serine/threonine-protein kinase Nek7 OS=Mus musculus OX=10090 GN=Nek7 PE=1 SV=1 | 0 | 12.302 | 15 | 2 | 2 | 2 | 201 | 22.7 | 9.39 | 1.29 | 1.25 | 1.032 | 0.37 | -0.37 | 0.32 | 0.10666667 | -0.345 | 0.05 | 0.35860658 | 0.546659909 | 0.799540069 | 0.974893705 | 0.995559182 | 0.985226443 | 0.992535481 |
| ND2 | Low | P03893 | NADH-ubiquinone oxidoreductase chain 2 OS=Mus musculus OX=10090 GN=Mtnd2 PE=1 SV=2 | 0.053 | 1.435 | 2 | 1 | 1 | 1 | 345 | 38.7 | 9.92 | 1.283 | 1.277 | 1.005 | 0.36 | -0.36 | 0.35 | 0.11666667 | -0.355 | 0.01 | 0.358119327 | 0.495015482 | 0.953345062 | 0.974893705 | 0.995559182 | 0.985226443 | 0.992535481 |
| Prkacb | High | P68181 | cAMP-dependent protein kinase catalytic subunit beta OS=Mus musculus OX=10090 GN=Prkacb P | 0 | 44.613 | 30 | 12 | 15 | 4 | 351 | 40.7 | 8.56 | 1.301 | 1.281 | 1.015 | 0.38 | -0.38 | 0.36 | 0.12 | -0.37 | 0.02 | 0.356675829 | 0.500501356 | 0.900453906 | 0.974893705 | 0.995559182 | 0.985226443 | 0.992535481 |
| Ccdc132; \ | High | Q8CI71 | Syndetin OS=Mus musculus OX=10090 GN=Vps50 PE=1 SV=2 | 0.001 | 7.293 | 1 | 3 | 3 | 3 | 964 | 111.1 | 6.07 | 1.3 | 1.299 | 1.001 | 0.38 | -0.38 | 0.38 | 0.12666667 | -0.38 | 0 | 0.355693317 | 0.459721789 | 0.999234586 | 0.974893705 | 0.995559182 | 0.985226443 | 0.999917596 |
| Omg | High | Q63912 | Oligodendrocyte-myelin glycoprotein OS=Mus musculus OX=10090 GN=Omg PE=1 SV=1 | 0.001 | 6.919 | 5 | 2 | 3 | 2 | 440 | 49.3 | 8.41 | 1.316 | 1.308 | 1.006 | 0.4 | -0.4 | 0.39 | 0.13 | -0.395 | 0.01 | 0.359491096 | 0.46082825 | 0.994459831 | 0.974893705 | 0.995559182 | 0.985226443 | 0.999917596 |
| Tfg | High | Q9Z1A1 | TFG protein OS=Mus musculus OX=10090 GN=Tfg PE=1 SV=1 | 0 | 21.675 | 17 | 4 | 6 | 4 | 397 | 43 | 5.1 | 0.807 | 1.017 | 0.794 | -0.31 | 0.31 | 0.02 | 0.00666667 | 0.145 | -0.33 | 0.359180097 | 0.97265642 | 0.15245969 | 0.974893705 | 0.998402594 | 0.98664815 | 0.725056389 |
| Smpd1 | Medium | Q04519 | Sphingomyelin phosphodiesterase OS=Mus musculus OX=10090 GN=Smpd1 PE=1 SV=2 | 0.039 | 1.697 | 2 | 1 | 1 | 1 | 627 | 69.9 | 7.2 | 0.811 | 1.005 | 0.807 | -0.3 | 0.3 | 0.01 | 0.00333333 | 0.145 | -0.31 | 0.357282637 | 0.946568946 | 0.230776947 | 0.974893705 | 0.998402594 | 0.98664815 | 0.819676582 |
| Ankle2 | Medium | Q6P1H6 | Ankyrin repeat and LEM domain-containing protein 2 OS=Mus musculus OX=10090 GN=Ankle2 PE | 0.045 | 1.592 | 1 | 1 | 1 | 1 | 964 | 106.1 | 7.37 | 1.289 | 1.079 | 1.195 | 0.37 | -0.37 | 0.11 | 0.03666667 | -0.24 | 0.26 | 0.359550958 | 0.89637895 | 0.262562439 | 0.974893705 | 0.998402594 | 0.98664815 | 0.859470827 |
| Rps18 | High | P62270 | 40S ribosomal protein S18 OS=Mus musculus OX=10090 GN=Rps18 PE=1 SV=3 | 0 | 33.975 | 41 | 9 | 18 | 9 | 152 | 17.7 | 10.99 | 1.292 | 1.092 | 1.183 | 0.37 | -0.37 | 0.13 | 0.04333333 | -0.25 | 0.24 | 0.357819581 | 0.853215604 | 0.237003844 | 0.974893705 | 0.998402594 | 0.98664815 | 0.828110564 |
| Nme1 | High | P15532 | Nucleoside diphosphate kinase A OS=Mus musculus OX=10090 GN=Nme1 PE=1 SV=1 | 0 | 56.706 | 74 | 12 | 22 | 4 | 152 | 17.2 | 7.37 | 0.761 | 0.741 | 1.027 | -0.39 | 0.39 | -0.43 | -0.1433333 | 0.41 | 0.04 | 0.359821865 | 0.314542437 | 0.78481522 | 0.975118992 | 0.955820031 | 0.965469511 | 0.992535481 |
| | Medium | A0A286YC | Trafficking protein particle complex 8 OS=Mus musculus OX=10090 GN=Trappc8 PE=1 SV=2 | 0.01 | 2.739 | 1 | 2 | 2 | 2 | 1437 | 160.8 | 7.01 | 0.793 | 0.613 | 1.293 | -0.33 | 0.33 | -0.71 | -0.2366667 | 0.52 | 0.37 | 0.361161441 | 0.112851802 | 0.10935351 | 0.975810853 | 0.77945068 | 0.877630766 | 0.632108382 |
| Paics | High | Q9DCL9 | Multifunctional protein ADE2 OS=Mus musculus OX=10090 GN=Paics PE=1 SV=4 | 0 | 48.095 | 28 | 10 | 14 | 10 | 425 | 47 | 7.23 | 0.788 | 0.678 | 1.163 | -0.34 | 0.34 | -0.56 | -0.1866667 | 0.45 | 0.22 | 0.360757406 | 0.20070679 | 0.376875047 | 0.975810853 | 0.912882441 | 0.944346647 | 0.920757467 |
| Klhdc10 | High | Q6PAR0 | Kelch domain-containing protein 10 OS=Mus musculus OX=10090 GN=Klhdc10 PE=1 SV=1 | 0 | 25.631 | 8 | 3 | 4 | 3 | 439 | 49 | 9.32 | 0.806 | 0.721 | 1.117 | -0.31 | 0.31 | -0.47 | -0.1566667 | 0.39 | 0.16 | 0.36096034 | 0.251545753 | 0.499772383 | 0.975810853 | 0.950034749 | 0.962922801 | 0.964378027 |
| Gprc5b | High | Q923Z0 | G-protein coupled receptor family C group 5 member B OS=Mus musculus OX=10090 GN=Gprc5b | 0.001 | 7.279 | 6 | 2 | 2 | 2 | 410 | 45.9 | 8.38 | 1.28 | 1.405 | 0.911 | 0.36 | -0.36 | 0.49 | 0.16333333 | -0.425 | -0.13 | 0.361469439 | 0.338264789 | 0.646364207 | 0.975810853 | 0.976045759 | 0.975928306 | 0.985010103 |
| Ctbp2 | High | P56546 | C-terminal-binding protein 2 OS=Mus musculus OX=10090 GN=Ctbp2 PE=1 SV=2 | 0 | 19.979 | 11 | 4 | 6 | 4 | 445 | 48.9 | 6.95 | 0.802 | 0.938 | 0.854 | -0.32 | 0.32 | -0.09 | -0.03 | 0.205 | -0.23 | 0.361153271 | 0.778939169 | 0.358642742 | 0.975810853 | 0.998402594 | 0.987106724 | 0.914600864 |
| Rpl14 | High | Q9CR57 | 60S ribosomal protein L14 OS=Mus musculus OX=10090 GN=Rpl14 PE=1 SV=3 | 0 | 34.325 | 30 | 8 | 14 | 8 | 217 | 23.5 | 11.02 | 1.312 | 1.117 | 1.175 | 0.39 | -0.39 | 0.16 | 0.05333333 | -0.275 | 0.23 | 0.3611559 | 0.79941854 | 0.248603564 | 0.975810853 | 0.998402594 | 0.987106724 | 0.845312422 |
| Ppip5k1 | High | A2ARP1 | Inositol hexakisphosphate and diphosphoinositol-pentakisphosphate kinase 1 OS=Mus musculus C | 0.001 | 5.729 | 1 | 2 | 2 | 2 | 1436 | 159.8 | 5.39 | 0.803 | 0.614 | 1.308 | -0.32 | 0.32 | -0.7 | -0.2333333 | 0.51 | 0.39 | 0.362559641 | 0.107523482 | 0.120751522 | 0.977273314 | 0.759299955 | 0.868286634 | 0.669025476 |
| Gprin1 | High | Q3UNH4 | G protein-regulated inducer of neurite outgrowth 1 OS=Mus musculus OX=10090 GN=Gprin1 PE= | 0 | 86.149 | 24 | 23 | 33 | 23 | 932 | 95.4 | 7.93 | 0.763 | 0.829 | 0.919 | -0.39 | 0.39 | -0.27 | -0.09 | 0.33 | -0.12 | 0.363405539 | 0.509774021 | 0.658446533 | 0.977273314 | 0.995559182 | 0.986416248 | 0.985010103 |
| Ckap5 | High | A2AGT5 | Cytoskeleton-associated protein 5 OS=Mus musculus OX=10090 GN=Ckap5 PE=1 SV=1 | 0 | 73.486 | 11 | 20 | 25 | 20 | 2032 | 225.5 | 7.96 | 1.309 | 1.155 | 1.134 | 0.39 | -0.39 | 0.21 | 0.07 | -0.3 | 0.18 | 0.363050927 | 0.721930169 | 0.363809896 | 0.977273314 | 0.995559182 | 0.986416248 | 0.91781256 |
| Rrbp1 | High | A2AVJ7 | Ribosome-binding protein 1 OS=Mus musculus OX=10090 GN=Rrbp1 PE=1 SV=1 | 0 | 84.277 | 28 | 21 | 27 | 21 | 1464 | 158.3 | 9.19 | 0.749 | 1.037 | 0.722 | -0.42 | 0.42 | 0.05 | 0.01666667 | 0.185 | -0.47 | 0.36334034 | 0.978821236 | 0.054208016 | 0.977273314 | 0.998402594 | 0.987837954 | 0.446320628 |
| Cldn1 | High | O88551 | Claudin-1 OS=Mus musculus OX=10090 GN=Cldn1 PE=1 SV=1 | 0.001 | 4.308 | 7 | 1 | 1 | 1 | 211 | 22.9 | 7.91 | 1.324 | 1.418 | 0.934 | 0.4 | -0.4 | 0.5 | 0.16666667 | -0.45 | -0.1 | 0.364035123 | 0.327167672 | 0.76947875 | 0.978052424 | 0.961945011 | 0.969998718 | 0.992535481 |
| Gja1 | High | P23242 | Gap junction alpha-1 protein OS=Mus musculus OX=10090 GN=Gja1 PE=1 SV=2 | 0 | 8.713 | 8 | 2 | 2 | 2 | 382 | 43 | 8.76 | 1.255 | 1.127 | 1.113 | 0.33 | -0.33 | 0.17 | 0.05666667 | -0.25 | 0.15 | 0.364044125 | 0.777816205 | 0.531752266 | 0.978052424 | 0.998402594 | 0.988227509 | 0.96694628 |
| Ccdc104; C | High | Q8C6E0 | Cilia- and flagella-associated protein 36 OS=Mus musculus OX=10090 GN=Cfap36 PE=1 SV=1 | 0 | 98.482 | 58 | 15 | 28 | 15 | 343 | 39.6 | 4.93 | 0.763 | 0.615 | 1.241 | -0.39 | 0.39 | -0.7 | -0.2333333 | 0.545 | 0.31 | 0.364835537 | 0.114671734 | 0.135509618 | 0.979709222 | 0.77945068 | 0.879579951 | 0.699265243 |
| Mcts1 | High | Q9DB27 | Malignant T-cell-amplified sequence 1 OS=Mus musculus OX=10090 GN=Mcts1 PE=1 SV=1 | 0 | 27.361 | 52 | 7 | 8 | 7 | 181 | 20.5 | 8.82 | 1.277 | 0.986 | 1.296 | 0.35 | -0.35 | -0.02 | -0.0066667 | -0.165 | 0.37 | 0.365200217 | 0.896406559 | 0.127303224 | 0.98021906 | 0.998402594 | 0.989310827 | 0.678506823 |
| Folh1 | Medium | O35409 | Glutamate carboxypeptidase 2 OS=Mus musculus OX=10090 GN=Folh1 PE=1 SV=2 | 0.01 | 2.74 | 1 | 1 | 1 | 1 | 752 | 84.5 | 8.1 | 0.806 | 0.941 | 0.856 | -0.31 | 0.31 | -0.09 | -0.03 | 0.2 | -0.22 | 0.365625061 | 0.764996746 | 0.40075918 | 0.980320596 | 0.995559182 | 0.987939889 | 0.93262326 |
| Naglu | High | O88325 | Alpha-N-acetylglucosaminidase OS=Mus musculus OX=10090 GN=Naglu PE=1 SV=1 | 0 | 8.102 | 4 | 2 | 2 | 2 | 739 | 82.5 | 6.61 | 0.804 | 0.992 | 0.81 | -0.32 | 0.32 | -0.01 | -0.0033333 | 0.165 | -0.3 | 0.365762562 | 0.895817269 | 0.263659724 | 0.980320596 | 0.998402594 | 0.989361595 | 0.860000043 |
| Ndufv2 | High | Q9D6J6 | NADH dehydrogenase [ubiquinone] flavoprotein 2, mitochondrial OS=Mus musculus OX=10090 GN | 0 | 20.007 | 22 | 5 | 8 | 5 | 248 | 27.3 | 7.4 | 1.27 | 1.409 | 0.901 | 0.34 | -0.34 | 0.5 | 0.16666667 | -0.42 | -0.15 | 0.366699281 | 0.333683372 | 0.504033136 | 0.980487776 | 0.97322306 | 0.976855418 | 0.964378027 |
| Rpl9-ps6 | High | A0A140T8 | 60S ribosomal protein L9 OS=Mus musculus OX=10090 GN=Rpl9-ps6 PE=3 SV=1 | 0 | 28.285 | 27 | 4 | 9 | 4 | 192 | 21.8 | 9.88 | 1.31 | 1.149 | 1.14 | 0.39 | -0.39 | 0.2 | 0.06666667 | -0.295 | 0.19 | 0.366539452 | 0.732861804 | 0.46633931 | 0.980487776 | 0.995559182 | 0.988023479 | 0.960359242 |
| Cnst | High | Q8CBC4 | Consortin OS=Mus musculus OX=10090 GN=Cnst PE=1 SV=1 | 0.003 | 3.994 | 2 | 1 | 1 | 1 | 711 | 76.8 | 4.59 | 0.788 | 1.02 | 0.773 | -0.34 | 0.34 | 0.03 | 0.01 | 0.155 | -0.37 | 0.366278055 | 0.956435438 | 0.232133198 | 0.980487776 | 0.998402594 | 0.989445185 | 0.820566032 |
| Obp2b | Low | A2BHR0 | Odorant-binding protein 2B OS=Mus musculus OX=10090 GN=Obp2b PE=3 SV=1 | 0.082 | 1.188 | 14 | 1 | 1 | 1 | 176 | 20 | 8.25 | 0.793 | 1.057 | 0.75 | -0.34 | 0.34 | 0.08 | 0.02666667 | 0.13 | -0.41 | 0.36659729 | 0.938009559 | 0.122182494 | 0.980487776 | 0.998402594 | 0.989445185 | 0.674952952 |
| Lhfpl3 | Low | Q9CTN8 | LHFPL tetraspan subfamily member 3 protein OS=Mus musculus OX=10090 GN=Lhfpl3 PE=2 SV=2 | 0.069 | 1.298 | 4 | 1 | 1 | 1 | 222 | 24.7 | 5.95 | 0.817 | 0.689 | 1.186 | -0.29 | 0.29 | -0.54 | -0.18 | 0.415 | 0.25 | 0.367066897 | 0.210388628 | 0.318731162 | 0.980535537 | 0.913066452 | 0.946800994 | 0.904943958 |
| Farsa | High | Q8C0C7 | Phenylalanine--tRNA ligase alpha subunit OS=Mus musculus OX=10090 GN=Farsa PE=1 SV=1 | 0 | 24.29 | 15 | 7 | 8 | 7 | 508 | 57.6 | 8.28 | 1.32 | 0.855 | 1.544 | 0.4 | -0.4 | -0.23 | -0.0766667 | -0.085 | 0.63 | 0.368199385 | 0.571862724 | 0.011106543 | 0.982156971 | 0.995559182 | 0.988858076 | 0.172028691 |
| Rpl35 | High | Q6ZWV7 | 60S ribosomal protein L35 OS=Mus musculus OX=10090 GN=Rpl35 PE=1 SV=1 | 0 | 12.956 | 33 | 5 | 8 | 5 | 123 | 14.5 | 11.05 | 1.31 | 1.041 | 1.259 | 0.39 | -0.39 | 0.06 | 0.02 | -0.225 | 0.33 | 0.368389408 | 0.971071996 | 0.145468684 | 0.982196582 | 0.998402594 | 0.990299588 | 0.706909771 |
| Ogdh | High | Q60597 | 2-oxoglutarate dehydrogenase, mitochondrial OS=Mus musculus OX=10090 GN=Ogdh PE=1 SV=3 | 0 | 63.194 | 20 | 18 | 21 | 12 | 1023 | 116.4 | 6.83 | 1.344 | 1.486 | 0.905 | 0.43 | -0.43 | 0.57 | 0.19 | -0.5 | -0.14 | 0.3699057 | 0.258939648 | 0.547211014 | 0.982967422 | 0.951072422 | 0.967019922 | 0.96694628 |
| Fis1 | High | G3X9U9 | Mitochondrial fission 1 protein OS=Mus musculus OX=10090 GN=Fis1 PE=1 SV=1 | 0 | 50.09 | 34 | 5 | 18 | 5 | 145 | 16.3 | 9.25 | 0.765 | 0.848 | 0.903 | -0.39 | 0.39 | -0.24 | -0.08 | 0.315 | -0.15 | 0.3696633 | 0.553907912 | 0.574912407 | 0.982967422 | 0.995559182 | 0.989263302 | 0.972910028 |
| Chchd6 | High | Q91VN4 | MICOS complex subunit Mic25 OS=Mus musculus OX=10090 GN=Chchd6 PE=1 SV=2 | 0 | 14.711 | 17 | 4 | 7 | 4 | 273 | 29.8 | 8.41 | 1.259 | 1.245 | 1.012 | 0.33 | -0.33 | 0.32 | 0.10666667 | -0.325 | 0.02 | 0.369895407 | 0.558464202 | 0.952744479 | 0.982967422 | 0.995559182 | 0.989263302 | 0.992535481 |
| Bbs1 | High | Q3V3N7 | Bardet-Biedl syndrome 1 protein homolog OS=Mus musculus OX=10090 GN=Bbs1 PE=1 SV=1 | 0 | 524.461 | 82 | 43 | 184 | 43 | 593 | 65.1 | 8.54 | 1.305 | 1.302 | 1.002 | 0.38 | -0.38 | 0.38 | 0.12666667 | -0.38 | 0 | 0.369001284 | 0.468842983 | 0.910510558 | 0.982967422 | 0.995559182 | 0.989263302 | 0.992535481 |
| Copb2 | High | O55029 | Coatomer subunit beta' OS=Mus musculus OX=10090 GN=Copb2 PE=1 SV=2 | 0.001 | 5.062 | 2 | 2 | 2 | 2 | 905 | 102.4 | 5.3 | 0.813 | 0.892 | 0.911 | -0.3 | 0.3 | -0.16 | -0.0533333 | 0.23 | -0.13 | 0.370222568 | 0.663276984 | 0.630346795 | 0.983343411 | 0.995559182 | 0.989451296 | 0.981916182 |
| | High | A0A286YC | Calcium/calmodulin-dependent protein kinase type II subunit gamma (Fragment) OS=Mus muscu | 0 | 44.798 | 34 | 9 | 19 | 1 | 274 | 30 | 8.07 | 0.809 | 0.654 | 1.238 | -0.31 | 0.31 | -0.61 | -0.2033333 | 0.46 | 0.31 | 0.372608614 | 0.146139038 | 0.20650836 | 0.983900096 | 0.843605535 | 0.913752816 | 0.799097568 |
| Pcx | High | G5E8R3 | Pyruvate carboxylase OS=Mus musculus OX=10090 GN=Pcx PE=1 SV=1 | 0 | 46.352 | 7 | 6 | 33 | 6 | 1178 | 129.6 | 6.71 | 1.259 | 1.52 | 0.828 | 0.33 | -0.33 | 0.6 | 0.2 | -0.465 | -0.27 | 0.371678348 | 0.230606198 | 0.24067591 | 0.983900096 | 0.936284541 | 0.960092318 | 0.832665803 |
| Ndufs6 | High | P52503 | NADH dehydrogenase [ubiquinone] iron-sulfur protein 6, mitochondrial OS=Mus musculus OX=10( | 0 | 14.338 | 31 | 3 | 4 | 3 | 116 | 13 | 8.65 | 1.277 | 1.498 | 0.852 | 0.35 | -0.35 | 0.58 | 0.19333333 | -0.465 | -0.23 | 0.372277007 | 0.248762339 | 0.327244602 | 0.983900096 | 0.950034749 | 0.966967423 | 0.905686588 |
| Ndufs4 | High | Q9CXZ1 | NADH dehydrogenase [ubiquinone] iron-sulfur protein 4, mitochondrial OS=Mus musculus OX=10( | 0 | 39.476 | 43 | 7 | 12 | 7 | 175 | 19.8 | 9.99 | 1.271 | 1.43 | 0.889 | 0.35 | -0.35 | 0.52 | 0.17333333 | -0.435 | -0.17 | 0.373590744 | 0.312317265 | 0.471876674 | 0.983900096 | 0.955820031 | 0.969860063 | 0.961211268 |
| Rd3 | High | Q8BRE0 | Protein RD3 OS=Mus musculus OX=10090 GN=Rd3 PE=1 SV=1 | 0 | 46.43 | 71 | 11 | 22 | 11 | 195 | 22.7 | 8.66 | 0.767 | 0.746 | 1.027 | -0.38 | 0.38 | -0.42 | -0.14 | 0.4 | 0.04 | 0.373203507 | 0.325785572 | 0.784996291 | 0.983900096 | 0.959894747 | 0.971897421 | 0.992535481 |
| Nxnl1; LOC | High | Q8VC33 | Nucleoredoxin-like protein 1 OS=Mus musculus OX=10090 GN=Nxnl1 PE=1 SV=1 | 0 | 90.294 | 63 | 12 | 51 | 12 | 217 | 24.9 | 5.05 | 1.302 | 1.331 | 0.979 | 0.38 | -0.38 | 0.41 | 0.13666667 | -0.395 | -0.03 | 0.372326495 | 0.429493581 | 0.970038818 | 0.983900096 | 0.99399649 | 0.988948293 | 0.995598794 |
| Phpt1 | High | Q9DAK9 | 14 kDa phosphohistidine phosphatase OS=Mus musculus OX=10090 GN=Phpt1 PE=1 SV=1 | 0 | 23.17 | 40 | 6 | 11 | 6 | 124 | 14 | 5.53 | 0.783 | 0.863 | 0.907 | -0.35 | 0.35 | -0.21 | -0.07 | 0.28 | -0.14 | 0.371877087 | 0.591379519 | 0.537259996 | 0.983900096 | 0.995559182 | 0.989729639 | 0.96694628 |
| Cntnap2 | High | E9QNF7 | Contactin-associated protein-like 2 OS=Mus musculus OX=10090 GN=Cntnap2 PE=1 SV=1 | 0 | 9.844 | 4 | 4 | 4 | 4 | 1332 | 148.2 | 6.76 | 0.799 | 0.923 | 0.865 | -0.32 | 0.32 | -0.12 | -0.04 | 0.22 | -0.21 | 0.373423347 | 0.740474613 | 0.414546575 | 0.983900096 | 0.995559182 | 0.989729639 | 0.936397673 |
| Fdps | High | Q920E5 | Farnesyl pyrophosphate synthase OS=Mus musculus OX=10090 GN=Fdps PE=1 SV=1 | 0 | 25.902 | 18 | 5 | 6 | 5 | 353 | 40.6 | 5.66 | 0.802 | 0.927 | 0.865 | -0.32 | 0.32 | -0.11 | -0.0366667 | 0.215 | -0.21 | 0.37299777 | 0.749930488 | 0.379193877 | 0.983900096 | 0.995559182 | 0.989729639 | 0.922487352 |
| | Medium | E9PW07 | Voltage-dependent calcium channel beta subunit-associated regulatory protein OS=Mus musculu | 0.01 | 2.661 | 1 | 1 | 1 | 1 | 705 | 75 | 6.21 | 0.82 | 1.054 | 0.778 | -0.29 | 0.29 | 0.08 | 0.02666667 | 0.105 | -0.36 | 0.37218846 | 0.956046041 | 0.160970026 | 0.983900096 | 0.998402594 | 0.991151345 | 0.733684585 |
| Eif3h | High | Q91WK2 | Eukaryotic translation initiation factor 3 subunit H OS=Mus musculus OX=10090 GN=Eif3h PE=1 S | 0 | 17.162 | 13 | 4 | 4 | 4 | 352 | 39.8 | 6.67 | 1.28 | 1.063 | 1.203 | 0.36 | -0.36 | 0.09 | 0.03 | -0.225 | 0.27 | 0.37187459 | 0.918597738 | 0.285290688 | 0.983900096 | 0.998402594 | 0.991151345 | 0.886074068 |
| Macrod1 | High | Q922B1 | ADP-ribose glycohydrolase MACROD1 OS=Mus musculus OX=10090 GN=Macrod1 PE=1 SV=2 | 0.001 | 5.759 | 6 | 2 | 2 | 2 | 323 | 35.3 | 8.85 | 1.302 | 1.117 | 1.166 | 0.38 | -0.38 | 0.16 | 0.05333333 | -0.27 | 0.22 | 0.371528387 | 0.815698372 | 0.288628233 | 0.983900096 | 0.998402594 | 0.991151345 | 0.888636466 |
| Banf1 | High | O54962 | Barrier-to-autointegration factor OS=Mus musculus OX=10090 GN=Banf1 PE=1 SV=1 | 0.003 | 3.887 | 17 | 2 | 2 | 2 | 89 | 10.1 | 6.09 | 0.766 | 0.768 | 0.997 | -0.39 | 0.39 | -0.38 | -0.1266667 | 0.385 | 0 | 0.374106972 | 0.370318442 | 0.970848054 | 0.984334957 | 0.977313184 | 0.980824071 | 0.995891884 |
| Gabra1 | High | P62812 | Gamma-aminobutyric acid receptor subunit alpha-1 OS=Mus musculus OX=10090 GN=Gabra1 PE= | 0 | 13.125 | 12 | 6 | 6 | 6 | 455 | 51.7 | 9.31 | 0.811 | 0.876 | 0.926 | -0.3 | 0.3 | -0.19 | -0.0633333 | 0.245 | -0.11 | 0.374021017 | 0.623450747 | 0.609769432 | 0.984334957 | 0.995559182 | 0.989947069 | 0.97826833 |
| Rhox2e; R | Medium | Q9D4Y3 | Reproductive homeobox 2A OS=Mus musculus OX=10090 GN=Rhox2a PE=2 SV=1 | 0.037 | 1.727 | 9 | 1 | 3 | 1 | 191 | 21.1 | 4.93 | 1.274 | 1.545 | 0.825 | 0.35 | -0.35 | 0.63 | 0.21 | -0.49 | -0.28 | 0.375731971 | 0.207752448 | 0.27639109 | 0.984554489 | 0.912882441 | 0.948718465 | 0.876067403 |
| Ptpra | High | Q91V35 | Receptor-type tyrosine-protein phosphatase alpha OS=Mus musculus OX=10090 GN=Ptpra PE=1 S | 0 | 38.025 | 15 | 10 | 12 | 8 | 793 | 89.8 | 6.64 | 0.771 | 0.741 | 1.041 | -0.37 | 0.37 | -0.43 | -0.1433333 | 0.4 | 0.06 | 0.37524849 | 0.314656154 | 0.783487066 | 0.984554489 | 0.955820031 | 0.97018726 | 0.992535481 |
| Ndufv3 | High | Q3U422 | Complex I-9kD OS=Mus musculus OX=10090 GN=Ndufv3 PE=1 SV=1 | 0 | 49.241 | 33 | 11 | 14 | 11 | 468 | 50.5 | 8.97 | 1.312 | 1.378 | 0.952 | 0.39 | -0.39 | 0.46 | 0.15333333 | -0.425 | -0.07 | 0.375770752 | 0.369833299 | 0.847832275 | 0.984554489 | 0.977313184 | 0.980933836 | 0.992535481 |
| Dctn2 | High | Q99KJ8 | Dynactin subunit 2 OS=Mus musculus OX=10090 GN=Dctn2 PE=1 SV=3 | 0 | 40.396 | 28 | 10 | 12 | 10 | 402 | 44.1 | 5.26 | 0.785 | 0.797 | 0.985 | -0.35 | 0.35 | -0.33 | -0.11 | 0.34 | -0.02 | 0.375456448 | 0.43450431 | 0.913853074 | 0.984554489 | 0.99399649 | 0.989275489 | 0.992535481 |
| Lss | High | Q8BLN5 | Lanosterol synthase OS=Mus musculus OX=10090 GN=Lss PE=1 SV=2 | 0.001 | 6.103 | 2 | 1 | 1 | 1 | 733 | 83.1 | 6.4 | 1.285 | 0.975 | 1.318 | 0.36 | -0.36 | -0.04 | -0.0133333 | -0.16 | 0.4 | 0.375524027 | 0.847044586 | 0.079849992 | 0.984554489 | 0.998402594 | 0.991478541 | 0.544008389 |
| Acp1 | High | Q561M1 | Low molecular weight cytosolic acid phosphatase OS=Mus musculus OX=10090 GN=Acp1 PE=1 S\ | 0.001 | 6.229 | 11 | 2 | 4 | 2 | 158 | 17.9 | 6.54 | 0.797 | 0.68 | 1.174 | -0.33 | 0.33 | -0.56 | -0.1866667 | 0.445 | 0.23 | 0.376721604 | 0.203819595 | 0.351784404 | 0.985204306 | 0.912882441 | 0.949043373 | 0.910644115 |
| | High | A0A1L1ST | CLIP-associating protein 2 OS=Mus musculus OX=10090 GN=Clasp2 PE=1 SV=1 | 0 | 15.315 | 9 | 5 | 5 | 1 | 600 | 66.3 | 6.55 | 1.274 | 1.22 | 1.044 | 0.35 | -0.35 | 0.29 | 0.09666667 | -0.32 | 0.06 | 0.376658328 | 0.584242078 | 0.779888787 | 0.985204306 | 0.995559182 | 0.990381744 | 0.992535481 |
| Sharpin | High | Q91WA6 | Sharpin OS=Mus musculus OX=10090 GN=Sharpin PE=1 SV=1 | 0.001 | 7.398 | 6 | 2 | 2 | 2 | 380 | 39.8 | 5.76 | 0.822 | 0.956 | 0.859 | -0.28 | 0.28 | -0.06 | -0.02 | 0.17 | -0.22 | 0.376454748 | 0.798466079 | 0.40694099 | 0.985204306 | 0.998402594 | 0.99180345 | 0.936397673 |
| Erbb2ip; E | High | B2RUJ2 | Erbb2ip protein OS=Mus musculus OX=10090 GN=Erbin PE=1 SV=1 | 0.001 | 6.139 | 1 | 2 | 2 | 2 | 1294 | 145 | 5.41 | 0.793 | 0.854 | 0.928 | -0.34 | 0.34 | -0.23 | -0.0766667 | 0.285 | -0.11 | 0.377255076 | 0.542941631 | 0.747358786 | 0.985220873 | 0.995559182 | 0.990390027 | 0.992535481 |
| Atg3 | High | Q9CPX6 | Ubiquitin-like-conjugating enzyme ATG3 OS=Mus musculus OX=10090 GN=Atg3 PE=1 SV=1 | 0.006 | 3.249 | 6 | 2 | 2 | 2 | 314 | 35.8 | 4.72 | 0.82 | 0.638 | 1.286 | -0.29 | 0.29 | -0.65 | -0.2166667 | 0.47 | 0.36 | 0.378718541 | 0.137266666 | 0.161889447 | 0.986433382 | 0.818645544 | 0.902539463 | 0.733684585 |
| Mark3 | High | Q03141 | MAP/microtubule affinity-regulating kinase 3 OS=Mus musculus OX=10090 GN=Mark3 PE=1 SV=2 | 0 | 28.648 | 8 | 6 | 9 | 6 | 753 | 84.3 | 9.51 | 0.806 | 0.7 | 1.151 | -0.31 | 0.31 | -0.51 | -0.17 | 0.41 | 0.2 | 0.37864853 | 0.237969873 | 0.419571118 | 0.986433382 | 0.937374952 | 0.961904167 | 0.937638605 |
| Idh3b | High | Q91VA7 | Isocitrate dehydrogenase [NAD] subunit, mitochondrial OS=Mus musculus OX=10090 GN=Idh3b P | 0 | 61.082 | 29 | 12 | 22 | 11 | 384 | 42.2 | 8.6 | 1.31 | 1.386 | 0.945 | 0.39 | -0.39 | 0.47 | 0.15666667 | -0.43 | -0.08 | 0.380358297 | 0.359867139 | 0.819134589 | 0.986433382 | 0.977313184 | 0.981873283 | 0.992535481 |
| Hint1 | High | P70349 | Histidine triad nucleotide-binding protein 1 OS=Mus musculus OX=10090 GN=Hint1 PE=1 SV=3 | 0 | 13.911 | 33 | 3 | 6 | 3 | 126 | 13.8 | 6.87 | 0.788 | 0.779 | 1.012 | -0.34 | 0.34 | -0.36 | -0.12 | 0.35 | 0.02 | 0.379982642 | 0.394466401 | 0.962191356 | 0.986433382 | 0.983884835 | 0.985159109 | 0.993960013 |
| Dennd1a | Medium | Q8K382 | DENN domain-containing protein 1A OS=Mus musculus OX=10090 GN=Dennd1a PE=1 SV=2 | 0.011 | 2.421 | 1 | 1 | 1 | 1 | 1016 | 111.5 | 6.79 | 0.819 | 1.129 | 0.725 | -0.29 | 0.29 | 0.17 | 0.05666667 | 0.06 | -0.46 | 0.378834375 | 0.76571588 | 0.076784009 | 0.986433382 | 0.995559182 | 0.990996282 | 0.526317775 |
| Denr | High | Q9CQJ6 | Density-regulated protein OS=Mus musculus OX=10090 GN=Denr PE=1 SV=1 | 0 | 14.033 | 20 | 4 | 4 | 4 | 198 | 22.2 | 5.3 | 1.286 | 0.886 | 1.453 | 0.36 | -0.36 | -0.18 | -0.06 | -0.09 | 0.54 | 0.380355888 | 0.646440635 | 0.035214948 | 0.986433382 | 0.995559182 | 0.990996282 | 0.367937483 |
| | High | E0CYW1 | Ubiquitin-like protein 5 (Fragment) OS=Mus musculus OX=10090 GN=Ubl5 PE=1 SV=1 | 0.001 | 6.258 | 38 | 2 | 2 | 2 | 56 | 6.5 | 9.66 | 1.28 | 1.177 | 1.088 | 0.36 | -0.36 | 0.24 | 0.08 | -0.3 | 0.12 | 0.38020235 | 0.666281628 | 0.619836869 | 0.986433382 | 0.995559182 | 0.990996282 | 0.97826833 |
| Rpl30; Rpl | High | P62889 | 60S ribosomal protein L30 OS=Mus musculus OX=10090 GN=Rpl30 PE=1 SV=2 | 0 | 17.921 | 26 | 5 | 7 | 5 | 115 | 12.8 | 9.63 | 1.327 | 1.167 | 1.137 | 0.41 | -0.41 | 0.22 | 0.07333333 | -0.315 | 0.19 | 0.378905091 | 0.698138494 | 0.448021928 | 0.986433382 | 0.995559182 | 0.990996282 | 0.958608248 |
| | Low | H7BX74 | Adhesion G protein-coupled receptor B2 OS=Mus musculus OX=10090 GN=Adgrb2 PE=1 SV=1 | 0.075 | 1.244 | 1 | 1 | 1 | 1 | 1544 | 168.3 | 7.47 | 1.268 | 1.697 | 0.747 | 0.34 | -0.34 | 0.76 | 0.25333333 | -0.55 | -0.42 | 0.382342595 | 0.117513046 | 0.102809259 | 0.986560023 | 0.779640783 | 0.883100403 | 0.610114517 |
| Ndufa13 | High | Q9ERS2 | NADH dehydrogenase [ubiquinone] 1 alpha subcomplex subunit 13 OS=Mus musculus OX=10090 ( | 0 | 11.024 | 22 | 3 | 5 | 3 | 144 | 16.8 | 9.48 | 1.287 | 1.423 | 0.904 | 0.36 | -0.36 | 0.51 | 0.17 | -0.435 | -0.15 | 0.381323686 | 0.319171773 | 0.5959255 | 0.986560023 | 0.955820031 | 0.971190027 | 0.977451587 |
| Rab4b | High | Q91ZR1 | Ras-related protein Rab-4B OS=Mus musculus OX=10090 GN=Rab4b PE=1 SV=2 | 0 | 27.207 | 45 | 8 | 11 | 6 | 213 | 23.6 | 6.06 | 0.786 | 0.926 | 0.849 | -0.35 | 0.35 | -0.11 | -0.0366667 | 0.23 | -0.24 | 0.381654704 | 0.747773406 | 0.318516449 | 0.986560023 | 0.995559182 | 0.991059602 | 0.904943958 |
| Opa3 | High | Q505D7 | Optic atrophy 3 protein homolog OS=Mus musculus OX=10090 GN=Opa3 PE=1 SV=1 | 0.001 | 4.538 | 16 | 3 | 3 | 3 | 179 | 20.1 | 8.34 | 1.299 | 1.166 | 1.115 | 0.38 | -0.38 | 0.22 | 0.07333333 | -0.3 | 0.16 | 0.381184806 | 0.707341506 | 0.460434718 | 0.986560023 | 0.995559182 | 0.991059602 | 0.959744396 |
| Rplp2 | High | P99027 | 60S acidic ribosomal protein P2 OS=Mus musculus OX=10090 GN=Rplp2 PE=1 SV=3 | 0 | 78.764 | 71 | 7 | 17 | 7 | 115 | 11.6 | 4.54 | 1.272 | 1.217 | 1.045 | 0.35 | -0.35 | 0.28 | 0.09333333 | -0.315 | 0.06 | 0.382252736 | 0.605873834 | 0.698979054 | 0.986560023 | 0.995559182 | 0.991059602 | 0.991285312 |
| Hdac5 | High | A2AWS5 | Histone deacetylase OS=Mus musculus OX=10090 GN=Hdac5 PE=1 SV=1 | 0.001 | 7.77 | 2 | 2 | 2 | 2 | 1030 | 111.4 | 5.72 | 0.815 | 0.652 | 1.25 | -0.3 | 0.3 | -0.62 | -0.2066667 | 0.46 | 0.32 | 0.385557634 | 0.152493894 | 0.21206905 | 0.992116409 | 0.847684847 | 0.919900628 | 0.804011919 |
| Pcmtd1 | High | P59913 | Protein-L-isoaspartate O-methyltransferase domain-containing protein 1 OS=Mus musculus OX=1 | 0.001 | 6.079 | 5 | 2 | 2 | 2 | 357 | 40.7 | 5.66 | 0.823 | 0.832 | 0.989 | -0.28 | 0.28 | -0.27 | -0.09 | 0.275 | -0.02 | 0.385314692 | 0.505633703 | 0.985093745 | 0.992116409 | 0.995559182 | 0.993837796 | 0.998604519 |
| Tagln3 | High | Q9R1Q8 | Transgelin-3 OS=Mus musculus OX=10090 GN=Tagln3 PE=1 SV=1 | 0 | 26.204 | 40 | 7 | 9 | 7 | 199 | 22.5 | 7.33 | 0.808 | 0.952 | 0.849 | -0.31 | 0.31 | -0.07 | -0.0233333 | 0.19 | -0.24 | 0.386014819 | 0.811757885 | 0.356775021 | 0.992837197 | 0.998402594 | 0.995619896 | 0.914600864 |
| Rasa1 | High | E9PYG6 | RAS p21 protein activator 1 OS=Mus musculus OX=10090 GN=Rasa1 PE=1 SV=1 | 0.001 | 6.398 | 2 | 3 | 3 | 3 | 1038 | 115.4 | 6.6 | 1.291 | 1.811 | 0.713 | 0.37 | -0.37 | 0.86 | 0.28666667 | -0.615 | -0.49 | 0.391140132 | 0.077957047 | 0.089904774 | 0.994143971 | 0.655042841 | 0.824593406 | 0.5746627 |
| Adpgk | High | A0A1L1SS | ADP-dependent glucokinase OS=Mus musculus OX=10090 GN=Adpgk PE=1 SV=1 | 0.003 | 3.95 | 3 | 1 | 1 | 1 | 496 | 53.9 | 5.62 | 1.272 | 1.679 | 0.757 | 0.35 | -0.35 | 0.75 | 0.25 | -0.55 | -0.4 | 0.394569909 | 0.124368211 | 0.132753088 | 0.994143971 | 0.799286338 | 0.896715154 | 0.695121227 |
| Unc119 | High | Q9Z2R6 | Protein unc-119 homolog A OS=Mus musculus OX=10090 GN=Unc119 PE=1 SV=1 | 0 | 42.335 | 40 | 9 | 18 | 9 | 240 | 27 | 6.15 | 0.777 | 0.629 | 1.235 | -0.36 | 0.36 | -0.67 | -0.2233333 | 0.515 | 0.3 | 0.398683717 | 0.131966394 | 0.143631732 | 0.994143971 | 0.808473523 | 0.901308747 | 0.70564779 |
| LOC10004 | High | P62827 | GTP-binding nuclear protein Ran OS=Mus musculus OX=10090 GN=Ran PE=1 SV=3 | 0 | 52.778 | 33 | 8 | 15 | 8 | 216 | 24.4 | 7.49 | 0.778 | 0.633 | 1.229 | -0.36 | 0.36 | -0.66 | -0.22 | 0.51 | 0.3 | 0.40016888 | 0.136886098 | 0.153360509 | 0.994143971 | 0.817450082 | 0.905797027 | 0.727583768 |
| Ccne2 | Low | A2API1 | G1/S-specific cyclin-E2 OS=Mus musculus OX=10090 GN=Ccne2 PE=1 SV=1 | 0.063 | 1.31 | 4 | 1 | 1 | 1 | 403 | 46.6 | 7.84 | 1.272 | 1.65 | 0.771 | 0.35 | -0.35 | 0.72 | 0.24 | -0.535 | -0.38 | 0.390472021 | 0.135689887 | 0.140986361 | 0.994143971 | 0.817450082 | 0.905797027 | 0.701851981 |
| Trove2 | High | O08848 | 60 kDa SS-A/Ro ribonucleoprotein OS=Mus musculus OX=10090 GN=RO60 PE=1 SV=1 | 0 | 30.262 | 15 | 7 | 10 | 7 | 538 | 60.1 | 7.9 | 0.768 | 0.634 | 1.211 | -0.38 | 0.38 | -0.66 | -0.22 | 0.52 | 0.28 | 0.397973921 | 0.137942664 | 0.21826443 | 0.994143971 | 0.821066364 | 0.907605168 | 0.805811317 |
| Fap | High | P97321 | Prolyl endopeptidase FAP OS=Mus musculus OX=10090 GN=Fap PE=1 SV=1 | 0 | 51.834 | 16 | 12 | 18 | 12 | 761 | 87.9 | 7.08 | 1.288 | 1.584 | 0.813 | 0.36 | -0.36 | 0.66 | 0.22 | -0.51 | -0.3 | 0.400493049 | 0.185338773 | 0.217347801 | 0.994143971 | 0.900299151 | 0.947221561 | 0.805520378 |
| Pcyox1 | High | Q9CQF9 | Prenylcysteine oxidase OS=Mus musculus OX=10090 GN=Pcyox1 PE=1 SV=1 | 0 | 10.994 | 6 | 4 | 5 | 4 | 505 | 56.5 | 6.92 | 1.269 | 1.486 | 0.854 | 0.34 | -0.34 | 0.57 | 0.19 | -0.455 | -0.23 | 0.396419057 | 0.259182867 | 0.322241059 | 0.994143971 | 0.951072422 | 0.972608197 | 0.905686588 |
| Marc2; Mc | High | Q922Q1 | Mitochondrial amidoxime reducing component 2 OS=Mus musculus OX=10090 GN=Mtarc2 PE=1 S | 0.001 | 5.831 | 5 | 2 | 2 | 2 | 338 | 38.2 | 8.68 | 1.286 | 1.476 | 0.871 | 0.36 | -0.36 | 0.56 | 0.18666667 | -0.46 | -0.2 | 0.389920031 | 0.267495343 | 0.435720146 | 0.994143971 | 0.952283423 | 0.973213697 | 0.949084077 |
| Zfp330 | Medium | Q922H9 | Zinc finger protein 330 OS=Mus musculus OX=10090 GN=Znf330 PE=1 SV=1 | 0.021 | 2.011 | 3 | 1 | 1 | 1 | 316 | 35.6 | 6.16 | 0.803 | 0.734 | 1.095 | -0.32 | 0.32 | -0.45 | -0.15 | 0.385 | 0.13 | 0.393887299 | 0.291931792 | 0.606962632 | 0.994143971 | 0.955820031 | 0.974982001 | 0.97826833 |
| Stxbp5l | Medium | Q5DQR4 | Syntaxin-binding protein 5-like OS=Mus musculus OX=10090 GN=Stxbp5l PE=1 SV=1 | 0.042 | 1.617 | 1 | 1 | 1 | 1 | 1185 | 131.8 | 6.86 | 0.822 | 0.735 | 1.119 | -0.28 | 0.28 | -0.45 | -0.15 | 0.365 | 0.16 | 0.387297592 | 0.28115183 | 0.504309108 | 0.994143971 | 0.955820031 | 0.974982001 | 0.964378027 |
| Ndufa8 | High | Q9DCJ5 | NADH dehydrogenase [ubiquinone] 1 alpha subcomplex subunit 8 OS=Mus musculus OX=10090 G | 0 | 21.699 | 47 | 7 | 7 | 7 | 172 | 20 | 8.46 | 1.286 | 1.442 | 0.892 | 0.36 | -0.36 | 0.53 | 0.17666667 | -0.445 | -0.17 | 0.400198506 | 0.300118118 | 0.462543513 | 0.994143971 | 0.955820031 | 0.974982001 | 0.959744396 |
| Srgap3 | High | E9QN14 | SLIT-ROBO Rho GTPase-activating protein 3 OS=Mus musculus OX=10090 GN=Srgap3 PE=1 SV=1 | 0.001 | 6.754 | 2 | 3 | 4 | 2 | 1075 | 121.6 | 6.57 | 1.291 | 1.429 | 0.903 | 0.37 | -0.37 | 0.52 | 0.17333333 | -0.445 | -0.15 | 0.399587737 | 0.302769739 | 0.563275052 | 0.994143971 | 0.955820031 | 0.974982001 | 0.972910028 |
| | High | D3Z1U9 | NADH dehydrogenase [ubiquinone] flavoprotein 1, mitochondrial OS=Mus musculus OX=10090 GN | 0 | 38.856 | 28 | 8 | 13 | 8 | 385 | 41.8 | 6.96 | 1.294 | 1.47 | 0.88 | 0.37 | -0.37 | 0.56 | 0.18666667 | -0.465 | -0.18 | 0.394689565 | 0.273456579 | 0.482033247 | 0.994143971 | 0.955820031 | 0.974982001 | 0.964378027 |
| Prdx5 | High | P99029 | Peroxiredoxin-5, mitochondrial OS=Mus musculus OX=10090 GN=Prdx5 PE=1 SV=2 | 0 | 114.621 | 61 | 14 | 34 | 14 | 210 | 21.9 | 8.85 | 1.287 | 1.412 | 0.912 | 0.36 | -0.36 | 0.5 | 0.16666667 | -0.43 | -0.13 | 0.394293311 | 0.331320821 | 0.62001264 | 0.994143971 | 0.968621131 | 0.981382551 | 0.97826833 |
| Pygm | High | Q9WUB3 | Glycogen phosphorylase, muscle form OS=Mus musculus OX=10090 GN=Pygm PE=1 SV=3 | 0 | 127.232 | 42 | 32 | 45 | 23 | 842 | 97.2 | 7.11 | 0.776 | 0.753 | 1.03 | -0.37 | 0.37 | -0.41 | -0.1366667 | 0.39 | 0.04 | 0.395965643 | 0.340082765 | 0.769980418 | 0.994143971 | 0.976045759 | 0.985094865 | 0.992535481 |

| | | | | | | | | | | | | | | | | | | | | | | | | | | | | |
|---|---|---|---|---|---|---|---|---|---|---|---|---|---|---|---|---|---|---|---|---|---|---|---|---|---|---|---|---|
| Actr1a | High | P61164 | Alpha-centractin OS=Mus musculus OX=10090 GN=Actr1a PE=1 SV=1 | 0 | 52.155 | 32 | 8 | 10 | 3 | 376 | 42.6 | 6.64 | 0.79 | 0.774 | 1.021 | -0.34 | 0.34 | -0.37 | -0.1233333 | 0.355 | 0.03 | 0.388905857 | 0.382646684 | 0.931818005 | 0.994143971 | 0.977313184 | 0.985728578 | 0.992535481 |
| Nrd1 | Medium | Q8BHG1 | Nardilysin OS=Mus musculus OX=10090 GN=Nrdc PE=1 SV=1 | 0.015 | 2.18 | 1 | 1 | 1 | 1 | 1161 | 132.8 | 4.87 | 0.827 | 0.771 | 1.073 | -0.27 | 0.27 | -0.38 | -0.1266667 | 0.325 | 0.1 | 0.400736963 | 0.370934641 | 0.69776862 | 0.994143971 | 0.977313184 | 0.985728578 | 0.991285312 |
| D10Wsu52 | High | Q99LF4 | RNA-splicing ligase RtcB homolog OS=Mus musculus OX=10090 GN=Rtcb PE=1 SV=1 | 0 | 35.529 | 25 | 10 | 14 | 10 | 505 | 55.2 | 7.23 | 1.289 | 1.359 | 0.948 | 0.37 | -0.37 | 0.44 | 0.14666667 | -0.405 | -0.08 | 0.401763033 | 0.392237769 | 0.81731042 | 0.994143971 | 0.983884835 | 0.989014403 | 0.992535481 |
| Hp1bp3 | High | Q3TEA8 | Heterochromatin protein 1-binding protein 3 OS=Mus musculus OX=10090 GN=Hp1bp3 PE=1 SV=1 | 0 | 8.693 | 5 | 4 | 4 | 4 | 554 | 60.8 | 9.7 | 0.807 | 0.78 | 1.035 | -0.31 | 0.31 | -0.36 | -0.12 | 0.335 | 0.05 | 0.402851306 | 0.395611464 | 0.816384643 | 0.994143971 | 0.984837604 | 0.989490788 | 0.992535481 |
| Sept5 | High | Q9Z2Q6 | Septin-5 OS=Mus musculus OX=10090 GN=Septin5 PE=1 SV=2 | 0 | 39.087 | 29 | 9 | 14 | 9 | 369 | 42.7 | 6.67 | 0.771 | 0.793 | 0.973 | -0.38 | 0.38 | -0.34 | -0.1133333 | 0.36 | -0.04 | 0.398139008 | 0.424746736 | 0.935551241 | 0.994143971 | 0.99399649 | 0.994070231 | 0.992535481 |
| Sdccag8 | High | Q80UF4 | Serologically defined colon cancer antigen 8 homolog OS=Mus musculus OX=10090 GN=Sdccag8 I | 0.001 | 5.802 | 3 | 2 | 2 | 2 | 717 | 82.9 | 6.52 | 1.299 | 1.339 | 0.97 | 0.38 | -0.38 | 0.42 | 0.14 | -0.4 | -0.04 | 0.395811758 | 0.426916154 | 0.906598828 | 0.994143971 | 0.99399649 | 0.994070231 | 0.992535481 |
| Gucy2f | High | Q5SDA5 | Retinal guanylyl cyclase 2 OS=Mus musculus OX=10090 GN=Gucy2f PE=1 SV=1 | 0 | 675.486 | 66 | 58 | 378 | 50 | 1108 | 124.3 | 7.42 | 1.283 | 1.324 | 0.969 | 0.36 | -0.36 | 0.4 | 0.13333333 | -0.38 | -0.04 | 0.399561609 | 0.438601227 | 0.92016953 | 0.994143971 | 0.994630833 | 0.994387402 | 0.992535481 |
| Ciapin1 | High | Q8WTY4 | Anamorsin OS=Mus musculus OX=10090 GN=Ciapin1 PE=1 SV=1 | 0 | 21.027 | 18 | 6 | 8 | 6 | 309 | 33.4 | 5.2 | 0.808 | 0.813 | 0.994 | -0.31 | 0.31 | -0.3 | -0.1 | 0.305 | -0.01 | 0.403012172 | 0.4705984 | 0.938032488 | 0.994143971 | 0.995559182 | 0.994851576 | 0.992535481 |
| Inpp5a | High | Q3TZT4 | Inositol-polyphosphate 5-phosphatase OS=Mus musculus OX=10090 GN=Inpp5a PE=1 SV=1 | 0 | 10.475 | 10 | 5 | 5 | 5 | 420 | 48.9 | 7.06 | 0.79 | 0.842 | 0.938 | -0.34 | 0.34 | -0.25 | -0.0833333 | 0.295 | -0.09 | 0.400963754 | 0.539489388 | 0.744632671 | 0.994143971 | 0.995559182 | 0.994851576 | 0.992535481 |
| Ptgds | High | O09114 | Prostaglandin-H2 D-isomerase OS=Mus musculus OX=10090 GN=Ptgds PE=1 SV=1 | 0 | 17.052 | 23 | 6 | 10 | 6 | 189 | 21.1 | 8.25 | 0.792 | 0.862 | 0.919 | -0.34 | 0.34 | -0.21 | -0.07 | 0.275 | -0.12 | 0.391606831 | 0.587654939 | 0.612235661 | 0.994143971 | 0.995559182 | 0.994851576 | 0.97826833 |
| Trappc2l | Medium | Q9JME7 | Trafficking protein particle complex subunit 2-like protein OS=Mus musculus OX=10090 GN=Trapp | 0.012 | 2.374 | 6 | 1 | 1 | 1 | 139 | 16 | 6.77 | 0.829 | 0.844 | 0.983 | -0.27 | 0.27 | -0.25 | -0.0833333 | 0.26 | -0.03 | 0.402446083 | 0.536006308 | 0.93165212 | 0.994143971 | 0.995559182 | 0.994851576 | 0.992535481 |
| Necap1 | High | Q9CR95 | Adaptin ear-binding coat-associated protein 1 OS=Mus musculus OX=10090 GN=Necap1 PE=1 SV= | 0 | 12.321 | 22 | 6 | 7 | 6 | 275 | 29.6 | 6.38 | 0.805 | 0.876 | 0.919 | -0.31 | 0.31 | -0.19 | -0.0633333 | 0.25 | -0.12 | 0.386997597 | 0.621933591 | 0.660224761 | 0.994143971 | 0.995559182 | 0.994851576 | 0.98592496 |
| Fabp3 | Medium | P11404 | Fatty acid-binding protein, heart OS=Mus musculus OX=10090 GN=Fabp3 PE=1 SV=5 | 0.036 | 1.768 | 7 | 1 | 1 | 1 | 133 | 14.8 | 6.57 | 0.82 | 0.886 | 0.926 | -0.29 | 0.29 | -0.18 | -0.06 | 0.235 | -0.11 | 0.40031095 | 0.641371595 | 0.663412598 | 0.994143971 | 0.995559182 | 0.994851576 | 0.986647506 |
| Nptx1 | Low | Q62443 | Neuronal pentraxin-1 OS=Mus musculus OX=10090 GN=Nptx1 PE=1 SV=1 | 0.076 | 1.239 | 2 | 1 | 1 | 1 | 432 | 47.1 | 6.44 | 0.821 | 1.21 | 0.679 | -0.28 | 0.28 | 0.28 | 0.09333333 | 0 | -0.56 | 0.3909259 | 0.622406119 | 0.026508407 | 0.994143971 | 0.995559182 | 0.994851576 | 0.308366473 |
| Lphn2; Adg | Low | A0A0G2JD | Adhesion G protein-coupled receptor L2 OS=Mus musculus OX=10090 GN=Adgrl2 PE=1 SV=1 | 0.075 | 1.246 | 1 | 1 | 1 | 1 | 1403 | 157 | 6.54 | 1.303 | 0.86 | 1.515 | 0.38 | -0.38 | -0.22 | -0.0733333 | -0.08 | 0.6 | 0.394472096 | 0.566452953 | 0.045759299 | 0.994143971 | 0.995559182 | 0.994851576 | 0.405968973 |
| Fzd1 | Medium | O70421 | Frizzled-1 OS=Mus musculus OX=10090 GN=Fzd1 PE=1 SV=3 | 0.021 | 2.021 | 1 | 1 | 2 | 1 | 642 | 71.1 | 7.71 | 1.264 | 1.166 | 1.084 | 0.34 | -0.34 | 0.22 | 0.07333333 | -0.28 | 0.12 | 0.399506492 | 0.70900302 | 0.571457227 | 0.994143971 | 0.995559182 | 0.994851576 | 0.972910028 |
| Ube3b | High | Q9ES34 | Ubiquitin-protein ligase E3B OS=Mus musculus OX=10090 GN=Ube3b PE=1 SV=3 | 0.001 | 6.345 | 3 | 3 | 3 | 3 | 1070 | 122.7 | 8.32 | 1.272 | 1.197 | 1.063 | 0.35 | -0.35 | 0.26 | 0.08666667 | -0.305 | 0.09 | 0.400574923 | 0.641312724 | 0.65678344 | 0.994143971 | 0.995559182 | 0.994851576 | 0.985010103 |
| Rpl8 | High | P62918 | 60S ribosomal protein L8 OS=Mus musculus OX=10090 GN=Rpl8 PE=1 SV=2 | 0 | 28.137 | 30 | 9 | 14 | 9 | 257 | 28 | 11.03 | 1.261 | 1.217 | 1.036 | 0.33 | -0.33 | 0.28 | 0.09333333 | -0.305 | 0.05 | 0.387898808 | 0.606214408 | 0.740540622 | 0.994143971 | 0.995559182 | 0.994851576 | 0.992535481 |
| Npr2 | High | Q6VVW5 | Atrial natriuretic peptide receptor 2 OS=Mus musculus OX=10090 GN=Npr2 PE=1 SV=2 | 0 | 26.554 | 3 | 2 | 15 | 1 | 1047 | 117 | 6.87 | 1.27 | 1.257 | 1.01 | 0.34 | -0.34 | 0.33 | 0.11 | -0.335 | 0.01 | 0.393918396 | 0.543652307 | 0.87367561 | 0.994143971 | 0.995559182 | 0.994851576 | 0.992535481 |
| Svip | High | Q3UZP4 | Small VCP/p97-interacting protein OS=Mus musculus OX=10090 GN=Svip PE=3 SV=1 | 0 | 24.022 | 32 | 2 | 12 | 2 | 77 | 8.4 | 8.91 | 1.304 | 1.239 | 1.053 | 0.38 | -0.38 | 0.31 | 0.10333333 | -0.345 | 0.07 | 0.389522541 | 0.568068634 | 0.730194343 | 0.994143971 | 0.995559182 | 0.994851576 | 0.992535481 |
| Ttc8 | High | Q8VD72 | Tetratricopeptide repeat protein 8 OS=Mus musculus OX=10090 GN=Ttc8 PE=1 SV=1 | 0 | 307.721 | 61 | 21 | 96 | 21 | 515 | 58.4 | 6.92 | 1.287 | 1.261 | 1.02 | 0.36 | -0.36 | 0.33 | 0.11 | -0.345 | 0.03 | 0.394329526 | 0.531592123 | 0.817476638 | 0.994143971 | 0.995559182 | 0.994851576 | 0.992535481 |
| Ppfia4 | High | A0A087W | Protein tyrosine phosphatase, receptor type, f polypeptide (PTPRF),-interacting protein (liprin), alp | 0.006 | 3.387 | 1 | 2 | 2 | 1 | 1184 | 133.6 | 6.3 | 1.26 | 1.297 | 0.971 | 0.33 | -0.33 | 0.38 | 0.12666667 | -0.355 | -0.04 | 0.392356831 | 0.461705148 | 0.888746229 | 0.994143971 | 0.995559182 | 0.994851576 | 0.992535481 |
| Eef1g | High | Q9D8N0 | Elongation factor 1-gamma OS=Mus musculus OX=10090 GN=Eef1g PE=1 SV=3 | 0 | 50.503 | 38 | 18 | 21 | 18 | 437 | 50 | 6.74 | 1.288 | 1.279 | 1.007 | 0.37 | -0.37 | 0.35 | 0.11666667 | -0.36 | 0.01 | 0.392115769 | 0.503695024 | 0.882451431 | 0.994143971 | 0.995559182 | 0.994851576 | 0.992535481 |
| Ptrhd1 | High | D3Z4S3 | Putative peptidyl-tRNA hydrolase PTRHD1 OS=Mus musculus OX=10090 GN=Ptrhd1 PE=1 SV=1 | 0.001 | 4.329 | 24 | 2 | 2 | 2 | 140 | 16 | 9.32 | 1.267 | 1.312 | 0.966 | 0.34 | -0.34 | 0.39 | 0.13 | -0.365 | -0.05 | 0.401935714 | 0.45985701 | 0.877401607 | 0.994143971 | 0.995559182 | 0.994851576 | 0.992535481 |
| Bbs5 | High | A2AUC6 | Bardet-Biedl syndrome 5 protein homolog OS=Mus musculus OX=10090 GN=Bbs5 PE=1 SV=1 | 0 | 105.623 | 55 | 16 | 64 | 16 | 320 | 36.5 | 5.35 | 1.285 | 1.311 | 0.98 | 0.36 | -0.36 | 0.39 | 0.13 | -0.375 | -0.03 | 0.397430951 | 0.456505655 | 0.976069264 | 0.994143971 | 0.995559182 | 0.994851576 | 0.997221123 |
| Dmtn; Epb | High | Q9WV69 | Dematin OS=Mus musculus OX=10090 GN=Dmtn PE=1 SV=1 | 0.001 | 7.262 | 6 | 3 | 3 | 3 | 405 | 45.4 | 8.41 | 0.839 | 1.005 | 0.835 | -0.25 | 0.25 | 0.01 | 0.00333333 | 0.12 | -0.26 | 0.399248336 | 0.931601817 | 0.279366598 | 0.994143971 | 0.998402594 | 0.996273283 | 0.876067403 |
| Ppef2 | High | O35385 | Serine/threonine-protein phosphatase with EF-hands 2 OS=Mus musculus OX=10090 GN=Ppef2 P | 0 | 92.896 | 31 | 16 | 20 | 16 | 757 | 86.6 | 7.03 | 1.276 | 0.941 | 1.356 | 0.35 | -0.35 | -0.09 | -0.03 | -0.13 | 0.44 | 0.395363251 | 0.786409446 | 0.04173229 | 0.994143971 | 0.998402594 | 0.996273283 | 0.397271566 |
| Gsk3a | High | Q2NL51 | Glycogen synthase kinase-3 alpha OS=Mus musculus OX=10090 GN=Gsk3a PE=1 SV=2 | 0 | 36.313 | 27 | 7 | 8 | 5 | 490 | 51.6 | 8.81 | 1.275 | 0.989 | 1.29 | 0.35 | -0.35 | -0.02 | -0.0066667 | -0.165 | 0.37 | 0.387104007 | 0.904039432 | 0.135619761 | 0.994143971 | 0.998402594 | 0.996273283 | 0.699265243 |
| Cbr1 | High | P48758 | Carbonyl reductase [NADPH] 1 OS=Mus musculus OX=10090 GN=Cbr1 PE=1 SV=3 | 0 | 61.46 | 69 | 15 | 20 | 15 | 277 | 30.6 | 8.31 | 1.281 | 1.014 | 1.264 | 0.36 | -0.36 | 0.02 | 0.00666667 | -0.19 | 0.34 | 0.402303173 | 0.964411261 | 0.107040877 | 0.994143971 | 0.998402594 | 0.996273283 | 0.626490812 |
| Eif2a | High | Q8BJW6 | Eukaryotic translation initiation factor 2A OS=Mus musculus OX=10090 GN=Eif2a PE=1 SV=2 | 0.001 | 7.512 | 2 | 2 | 2 | 2 | 581 | 64.4 | 8.91 | 1.267 | 1.059 | 1.196 | 0.34 | -0.34 | 0.08 | 0.02666667 | -0.21 | 0.26 | 0.400211014 | 0.94405473 | 0.261149194 | 0.994143971 | 0.998402594 | 0.996273283 | 0.857799373 |
| | High | A0A6I8MX | DnaJ homolog subfamily C member 7 OS=Mus musculus OX=10090 GN=Dnajc7 PE=1 SV=1 | 0 | 48.955 | 31 | 16 | 17 | 16 | 484 | 55.3 | 7.84 | 1.298 | 1.04 | 1.247 | 0.38 | -0.38 | 0.06 | 0.02 | -0.22 | 0.32 | 0.39779566 | 0.971488611 | 0.148620488 | 0.994143971 | 0.998402594 | 0.996273283 | 0.713454689 |
| Pnmal2 | Medium | G3X9N3 | PNMA-like 2 OS=Mus musculus OX=10090 GN=Pnmal2 PE=1 SV=1 | 0.025 | 1.978 | 1 | 1 | 1 | 1 | 661 | 71.7 | 5.17 | 1.258 | 1.089 | 1.155 | 0.33 | -0.33 | 0.12 | 0.04 | -0.225 | 0.21 | 0.400467165 | 0.874298017 | 0.347539354 | 0.994143971 | 0.998402594 | 0.996273283 | 0.909421216 |
| Tub | High | P50586 | Tubby protein OS=Mus musculus OX=10090 GN=Tub PE=1 SV=1 | 0 | 57.529 | 32 | 14 | 22 | 11 | 505 | 55.3 | 8.68 | 1.297 | 1.054 | 1.23 | 0.38 | -0.38 | 0.08 | 0.02666667 | -0.23 | 0.3 | 0.398523715 | 0.939760436 | 0.211302732 | 0.994143971 | 0.998402594 | 0.996273283 | 0.804011919 |
| Fam120a | High | Q6A0A9 | Constitutive coactivator of PPAR-gamma-like protein 1 OS=Mus musculus OX=10090 GN=FAM120 | 0 | 25.977 | 8 | 6 | 6 | 6 | 1112 | 121.6 | 8.92 | 1.275 | 1.084 | 1.177 | 0.35 | -0.35 | 0.12 | 0.04 | -0.235 | 0.23 | 0.402606167 | 0.872108211 | 0.331993291 | 0.994143971 | 0.998402594 | 0.996273283 | 0.906329766 |
| Kcnv2 | High | Q8CFS6 | Potassium voltage-gated channel subfamily V member 2 OS=Mus musculus OX=10090 GN=Kcnv2 | 0.003 | 3.945 | 3 | 1 | 1 | 1 | 562 | 64.4 | 6.2 | 1.265 | 1.126 | 1.123 | 0.34 | -0.34 | 0.17 | 0.05666667 | -0.255 | 0.17 | 0.392853868 | 0.784234216 | 0.451822208 | 0.994143971 | 0.998402594 | 0.996273283 | 0.959744396 |
| Vwa5a | High | Q99KC8 | von Willebrand factor A domain-containing protein 5A OS=Mus musculus OX=10090 GN=Vwa5a F | 0 | 26.036 | 9 | 8 | 11 | 8 | 793 | 87.1 | 6.58 | 0.818 | 0.887 | 0.922 | -0.29 | 0.29 | -0.17 | -0.0566667 | 0.23 | -0.12 | 0.403698641 | 0.650998077 | 0.613148017 | 0.995399419 | 0.995559182 | 0.995479301 | 0.97826833 |
| Gstk1 | Medium | Q9DCM2 | Glutathione S-transferase kappa 1 OS=Mus musculus OX=10090 GN=Gstk1 PE=1 SV=3 | 0.044 | 1.6 | 3 | 1 | 1 | 1 | 226 | 25.7 | 8.88 | 1.249 | 1.462 | 0.854 | 0.32 | -0.32 | 0.55 | 0.18333333 | -0.435 | -0.23 | 0.404262724 | 0.266913272 | 0.396243008 | 0.995476984 | 0.951072422 | 0.973274703 | 0.93262326 |
| Glud1 | High | P26443 | Glutamate dehydrogenase 1, mitochondrial OS=Mus musculus OX=10090 GN=Glud1 PE=1 SV=1 | 0 | 85.898 | 37 | 19 | 27 | 19 | 558 | 61.3 | 8 | 1.28 | 1.43 | 0.895 | 0.36 | -0.36 | 0.52 | 0.17333333 | -0.44 | -0.16 | 0.404257216 | 0.311883765 | 0.539463896 | 0.995476984 | 0.955820031 | 0.975648507 | 0.96694628 |
| G3bp1 | High | P97855 | Ras GTPase-activating protein-binding protein 1 OS=Mus musculus OX=10090 GN=G3bp1 PE=1 S\ | 0 | 9.952 | 5 | 2 | 2 | 2 | 465 | 51.8 | 5.59 | 1.252 | 1.251 | 1.001 | 0.32 | -0.32 | 0.32 | 0.10666667 | -0.32 | 0 | 0.404130445 | 0.548255317 | 0.952172779 | 0.995476984 | 0.995559182 | 0.995518083 | 0.992535481 |
| | Low | A0A087W | Ankyrin-3 (Fragment) OS=Mus musculus OX=10090 GN=Ank3 PE=1 SV=1 | 0.052 | 1.464 | 0 | 1 | 1 | 1 | 2871 | 315.3 | 5.77 | 0.834 | 0.621 | 1.343 | -0.26 | 0.26 | -0.69 | -0.23 | 0.475 | 0.43 | 0.412842397 | 0.116128742 | 0.099137624 | 0.997078122 | 0.779640783 | 0.888359452 | 0.604200714 |
| Cox7c; Gm | High | P17665 | Cytochrome c oxidase subunit 7C, mitochondrial OS=Mus musculus OX=10090 GN=Cox7c PE=1 SV | 0 | 9.835 | 29 | 2 | 3 | 2 | 63 | 7.3 | 11 | 1.272 | 1.538 | 0.827 | 0.35 | -0.35 | 0.62 | 0.20666667 | -0.485 | -0.27 | 0.41261248 | 0.207828442 | 0.242308481 | 0.997078122 | 0.912882441 | 0.954980281 | 0.836649471 |
| Zc4h2 | Low | Q05BE0 | Zc4h2 protein OS=Mus musculus OX=10090 GN=Zc4h2 PE=1 SV=1 | 0.087 | 1.173 | 8 | 1 | 1 | 1 | 196 | 23.1 | 7.18 | 1.252 | 1.528 | 0.819 | 0.32 | -0.32 | 0.61 | 0.20333333 | -0.465 | -0.29 | 0.413216683 | 0.216295625 | 0.268070181 | 0.997078122 | 0.920170644 | 0.958624383 | 0.869723277 |
| Fah | High | P35505 | Fumarylacetoacetase OS=Mus musculus OX=10090 GN=Fah PE=1 SV=2 | 0 | 8.374 | 9 | 2 | 2 | 2 | 419 | 46.1 | 7.18 | 1.26 | 1.529 | 0.824 | 0.33 | -0.33 | 0.61 | 0.20333333 | -0.47 | -0.28 | 0.416958903 | 0.217173332 | 0.281878291 | 0.997078122 | 0.921794757 | 0.959436439 | 0.882463193 |
| Ephx2 | High | P34914 | Bifunctional epoxide hydrolase 2 OS=Mus musculus OX=10090 GN=Ephx2 PE=1 SV=2 | 0 | 10.487 | 7 | 2 | 2 | 2 | 554 | 62.5 | 6.19 | 0.828 | 0.702 | 1.179 | -0.27 | 0.27 | -0.51 | -0.17 | 0.39 | 0.24 | 0.411966468 | 0.220963468 | 0.302126651 | 0.997078122 | 0.928185815 | 0.962631968 | 0.900241969 |
| Ethe1 | High | Q9DCM0 | Persulfide dioxygenase ETHE1, mitochondrial OS=Mus musculus OX=10090 GN=Ethe1 PE=1 SV=2 | 0.001 | 6.313 | 9 | 2 | 2 | 2 | 254 | 27.7 | 7.23 | 1.238 | 1.482 | 0.835 | 0.31 | -0.31 | 0.57 | 0.19 | -0.44 | -0.26 | 0.417205848 | 0.252501279 | 0.333275097 | 0.997078122 | 0.950034749 | 0.973556435 | 0.906329766 |
| Ndufb5 | High | Q9CQH3 | NADH dehydrogenase [ubiquinone] 1 beta subcomplex subunit 5, mitochondrial OS=Mus musculu | 0 | 13.559 | 15 | 3 | 6 | 3 | 189 | 21.7 | 9.41 | 1.251 | 1.494 | 0.838 | 0.32 | -0.32 | 0.58 | 0.19333333 | -0.45 | -0.26 | 0.417716516 | 0.252357058 | 0.275233457 | 0.997078122 | 0.950034749 | 0.973556435 | 0.876067403 |
| Prmt5 | High | A0A0R4J0 | Protein arginine N-methyltransferase 5 OS=Mus musculus OX=10090 GN=Prmt5 PE=1 SV=1 | 0 | 9.466 | 4 | 2 | 3 | 2 | 637 | 72.7 | 6.34 | 0.806 | 0.731 | 1.102 | -0.31 | 0.31 | -0.45 | -0.15 | 0.38 | 0.14 | 0.408542749 | 0.287470258 | 0.56702822 | 0.997078122 | 0.955820031 | 0.976449076 | 0.972910028 |
| Ipo9 | Medium | E9QKZ2 | Importin-9 OS=Mus musculus OX=10090 GN=Ipo9 PE=1 SV=1 | 0.019 | 2.054 | 1 | 1 | 1 | 1 | 1040 | 115.9 | 4.82 | 0.826 | 0.732 | 1.129 | -0.28 | 0.28 | -0.45 | -0.15 | 0.365 | 0.17 | 0.406696921 | 0.286357479 | 0.491696326 | 0.997078122 | 0.955820031 | 0.976449076 | 0.964378027 |
| | Medium | A0A0J9YU | Fibronectin type III domain-containing protein 4 (Fragment) OS=Mus musculus OX=10090 GN=Fn | 0.029 | 1.926 | 8 | 1 | 1 | 1 | 220 | 23.7 | 7.25 | 0.833 | 0.738 | 1.129 | -0.26 | 0.26 | -0.44 | -0.1466667 | 0.35 | 0.17 | 0.41167421 | 0.297997061 | 0.504527393 | 0.997078122 | 0.955820031 | 0.976449076 | 0.964378027 |
| Sdk2 | High | Q6V4S5 | Protein sidekick-2 OS=Mus musculus OX=10090 GN=Sdk2 PE=1 SV=1 | 0.001 | 6.777 | 1 | 2 | 2 | 2 | 2176 | 239.8 | 7.36 | 0.825 | 0.746 | 1.105 | -0.28 | 0.28 | -0.42 | -0.14 | 0.35 | 0.14 | 0.414430445 | 0.302497148 | 0.506935876 | 0.997078122 | 0.955820031 | 0.976449076 | 0.964378027 |
| Dtd1 | High | Q9DD18 | D-aminoacyl-tRNA deacylase 1 OS=Mus musculus OX=10090 GN=Dtd1 PE=1 SV=2 | 0.006 | 3.237 | 4 | 1 | 1 | 1 | 209 | 23.4 | 7.87 | 1.252 | 1.431 | 0.875 | 0.32 | -0.32 | 0.52 | 0.17333333 | -0.42 | -0.19 | 0.412529641 | 0.30934969 | 0.49023529 | 0.997078122 | 0.955820031 | 0.976449076 | 0.964378027 |
| Ddx5 | High | Q61656 | Probable ATP-dependent RNA helicase DDX5 OS=Mus musculus OX=10090 GN=Ddx5 PE=1 SV=2 | 0 | 19.426 | 10 | 6 | 8 | 5 | 614 | 69.2 | 8.92 | 1.288 | 1.467 | 0.878 | 0.36 | -0.36 | 0.55 | 0.18333333 | -0.455 | -0.19 | 0.414128542 | 0.275986775 | 0.43368462 | 0.997078122 | 0.955820031 | 0.976449076 | 0.948052758 |
| Cpt2 | High | P52825 | Carnitine O-palmitoyltransferase 2, mitochondrial OS=Mus musculus OX=10090 GN=Cpt2 PE=1 S\ | 0.001 | 6.985 | 7 | 4 | 4 | 4 | 658 | 73.9 | 8.37 | 1.259 | 1.416 | 0.889 | 0.33 | -0.33 | 0.5 | 0.16666667 | -0.415 | -0.17 | 0.409680931 | 0.326980025 | 0.521554885 | 0.997078122 | 0.961945011 | 0.979511566 | 0.964378027 |
| Bbs7 | High | Q8K2G4 | Bardet-Biedl syndrome 7 protein homolog OS=Mus musculus OX=10090 GN=Bbs7 PE=1 SV=1 | 0 | 438.042 | 66 | 45 | 197 | 45 | 715 | 80.3 | 5.81 | 1.271 | 1.415 | 0.898 | 0.35 | -0.35 | 0.5 | 0.16666667 | -0.425 | -0.15 | 0.417343662 | 0.327506925 | 0.555072926 | 0.997078122 | 0.962372728 | 0.979725425 | 0.968958249 |
| Ide | High | Q9JHR7 | Insulin-degrading enzyme OS=Mus musculus OX=10090 GN=Ide PE=1 SV=1 | 0 | 13.426 | 4 | 4 | 4 | 4 | 1019 | 117.7 | 6.54 | 0.781 | 0.756 | 1.034 | -0.36 | 0.36 | -0.4 | -0.1333333 | 0.38 | 0.05 | 0.415438866 | 0.34553594 | 0.797878474 | 0.997078122 | 0.977313184 | 0.987195653 | 0.992535481 |
| Habp4 | High | Q9JKS5 | Intracellular hyaluronan-binding protein 4 OS=Mus musculus OX=10090 GN=Habp4 PE=1 SV=2 | 0 | 4.862 | 6 | 2 | 2 | 2 | 411 | 45.9 | 6.84 | 0.813 | 0.763 | 1.066 | -0.3 | 0.3 | -0.39 | -0.13 | 0.345 | 0.09 | 0.407928959 | 0.36016008 | 0.696832168 | 0.997078122 | 0.977313184 | 0.987195653 | 0.991285312 |
| Eef1e1 | High | Q9D1M4 | Eukaryotic translation elongation factor 1 epsilon-1 OS=Mus musculus OX=10090 GN=Eef1e1 PE= | 0.001 | 5.81 | 5 | 1 | 2 | 1 | 174 | 19.8 | 8.59 | 1.244 | 1.354 | 0.919 | 0.32 | -0.32 | 0.44 | 0.14666667 | -0.38 | -0.12 | 0.410953725 | 0.398730604 | 0.659815357 | 0.997078122 | 0.985440071 | 0.991259096 | 0.985767308 |
| Hspa4l | High | P48722 | Heat shock 70 kDa protein 4L OS=Mus musculus OX=10090 GN=Hspa4l PE=1 SV=2 | 0 | 163.61 | 43 | 31 | 49 | 29 | 838 | 94.3 | 5.74 | 0.782 | 0.782 | 1 | -0.35 | 0.35 | -0.35 | -0.1166667 | 0.35 | 0 | 0.410603549 | 0.401299775 | 0.91936874 | 0.997078122 | 0.9855335 | 0.991305811 | 0.992535481 |
| Bbs9 | High | Q811G0 | Protein PTHB1 OS=Mus musculus OX=10090 GN=Bbs9 PE=1 SV=2 | 0 | 473.829 | 59 | 47 | 212 | 47 | 885 | 99 | 6.13 | 1.272 | 1.333 | 0.954 | 0.35 | -0.35 | 0.41 | 0.13666667 | -0.38 | -0.07 | 0.416445538 | 0.425795222 | 0.838970297 | 0.997078122 | 0.99399649 | 0.995537306 | 0.992535481 |
| Gbas; Nips | High | Q7TMG8 | Glioblastoma amplified sequence OS=Mus musculus OX=10090 GN=Nipsnap2 PE=1 SV=1 | 0 | 31.324 | 20 | 6 | 11 | 6 | 281 | 32.9 | 9.33 | 1.279 | 1.327 | 0.964 | 0.36 | -0.36 | 0.41 | 0.13666667 | -0.385 | -0.05 | 0.411732565 | 0.434209497 | 0.919382936 | 0.997078122 | 0.99399649 | 0.995537306 | 0.992535481 |
| Clip1 | High | Q922J3 | CAP-Gly domain-containing linker protein 1 OS=Mus musculus OX=10090 GN=Clip1 PE=1 SV=1 | 0.002 | 4.199 | 2 | 3 | 3 | 3 | 1391 | 155.7 | 5.24 | 0.83 | 0.835 | 0.994 | -0.27 | 0.27 | -0.26 | -0.0866667 | 0.265 | -0.01 | 0.414482077 | 0.522356523 | 0.965092506 | 0.997078122 | 0.995559182 | 0.996318652 | 0.994302941 |
| Smc3 | High | Q9CW03 | Structural maintenance of chromosomes protein 3 OS=Mus musculus OX=10090 GN=Smc3 PE=1 S | 0 | 4.879 | 2 | 2 | 2 | 2 | 1217 | 141.5 | 7.18 | 0.824 | 0.872 | 0.946 | -0.28 | 0.28 | -0.2 | -0.0666667 | 0.24 | -0.08 | 0.405488029 | 0.596221274 | 0.8332788 | 0.997078122 | 0.995559182 | 0.996318652 | 0.992535481 |
| Stom | Medium | P54116 | Stomatin OS=Mus musculus OX=10090 GN=Stom PE=1 SV=3 | 0.041 | 1.639 | 6 | 1 | 1 | 1 | 284 | 31.4 | 6.93 | 0.809 | 0.905 | 0.894 | -0.31 | 0.31 | -0.14 | -0.0466667 | 0.225 | -0.16 | 0.415977899 | 0.672201868 | 0.634864998 | 0.997078122 | 0.995559182 | 0.996318652 | 0.983562471 |
| Atp2b3 | High | Q0VF55 | Calcium-transporting ATPase OS=Mus musculus OX=10090 GN=Atp2b3 PE=1 SV=1 | 0 | 97.75 | 11 | 14 | 33 | 1 | 1220 | 134.3 | 5.69 | 0.856 | 0.9 | 0.951 | -0.22 | 0.22 | -0.15 | -0.05 | 0.185 | -0.07 | 0.414782121 | 0.660098186 | 0.806775106 | 0.997078122 | 0.995559182 | 0.996318652 | 0.992535481 |
| | Medium | F6SRV1 | Splicing factor 1 (Fragment) OS=Mus musculus OX=10090 GN=Sf1 PE=1 SV=8 | 0.01 | 2.591 | 8 | 1 | 1 | 1 | 171 | 18 | 9.22 | 0.831 | 1.142 | 0.728 | -0.27 | 0.27 | 0.19 | 0.06333333 | 0.04 | -0.46 | 0.412791596 | 0.748219124 | 0.093146474 | 0.997078122 | 0.995559182 | 0.996318652 | 0.582204115 |
| Nde1 | Medium | Q9CZA6 | Nuclear distribution protein nudE homolog 1 OS=Mus musculus OX=10090 GN=Nde1 PE=1 SV=1 | 0.014 | 2.306 | 3 | 1 | 1 | 1 | 344 | 38.5 | 5.34 | 1.254 | 0.919 | 1.366 | 0.33 | -0.33 | -0.12 | -0.04 | -0.105 | 0.45 | 0.406475712 | 0.71433066 | 0.087090758 | 0.997078122 | 0.995559182 | 0.996318652 | 0.571132024 |
| Rsl1d1 | High | Q8BVY0 | Ribosomal L1 domain-containing protein 1 OS=Mus musculus OX=10090 GN=Rsl1d1 PE=1 SV=1 | 0 | 12.189 | 6 | 3 | 3 | 3 | 452 | 50.4 | 9.98 | 1.238 | 1.149 | 1.078 | 0.31 | -0.31 | 0.2 | 0.06666667 | -0.255 | 0.11 | 0.40811764 | 0.733868799 | 0.650403537 | 0.997078122 | 0.995559182 | 0.996318652 | 0.985010103 |
| Atad1 | High | Q9D5T0 | Outer mitochondrial transmembrane helix translocase OS=Mus musculus OX=10090 GN=Atad1 PI | 0.001 | 6.416 | 8 | 2 | 2 | 2 | 361 | 40.7 | 6.9 | 1.244 | 1.208 | 1.03 | 0.32 | -0.32 | 0.27 | 0.09 | -0.295 | 0.04 | 0.417487496 | 0.61445112 | 0.804018695 | 0.997078122 | 0.995559182 | 0.996318652 | 0.992535481 |
| Nisch | Medium | Q80TM9 | Nischarin OS=Mus musculus OX=10090 GN=Nisch PE=1 SV=2 | 0.045 | 1.582 | 1 | 1 | 1 | 1 | 1593 | 174.9 | 5.16 | 1.234 | 1.229 | 1.004 | 0.3 | -0.3 | 0.3 | 0.1 | -0.3 | 0.01 | 0.414768526 | 0.584970243 | 0.983248878 | 0.997078122 | 0.995559182 | 0.996318652 | 0.997765878 |
| Etfdh | High | Q921G7 | Electron transfer flavoprotein-ubiquinone oxidoreductase, mitochondrial OS=Mus musculus OX=1( | 0 | 11.946 | 7 | 4 | 4 | 4 | 616 | 68 | 7.58 | 1.225 | 1.308 | 0.937 | 0.29 | -0.29 | 0.39 | 0.13 | -0.34 | -0.09 | 0.413151986 | 0.461344308 | 0.68507938 | 0.997078122 | 0.995559182 | 0.996318652 | 0.991285312 |
| 1700019D | High | Q9CPS8 | Small membrane A-kinase anchor protein OS=Mus musculus OX=10090 PE=1 SV=1 | 0.006 | 3.213 | 13 | 1 | 1 | 1 | 106 | 11.9 | 4.25 | 1.253 | 1.272 | 0.985 | 0.33 | -0.33 | 0.35 | 0.11666667 | -0.34 | -0.02 | 0.409582038 | 0.500011595 | 0.950976401 | 0.997078122 | 0.995559182 | 0.996318652 | 0.992535481 |
| Spata5 | High | A0A0A0M | Vesicle-fusing ATPase OS=Mus musculus OX=10090 GN=Spata5 PE=1 SV=1 | 0 | 10.155 | 2 | 2 | 4 | 1 | 892 | 97.1 | 8.24 | 1.277 | 1.297 | 0.985 | 0.35 | -0.35 | 0.37 | 0.12333333 | -0.36 | -0.02 | 0.40649682 | 0.492529885 | 0.999009871 | 0.997078122 | 0.995559182 | 0.996318652 | 0.999917596 |
| | High | A0A1L1SV | Alpha-actinin-4 OS=Mus musculus OX=10090 GN=Actn4 PE=1 SV=1 | 0 | 138.29 | 36 | 27 | 38 | 19 | 932 | 107 | 5.36 | 0.785 | 0.986 | 0.796 | -0.35 | 0.35 | -0.02 | -0.0066667 | 0.185 | -0.33 | 0.416853822 | 0.898221728 | 0.169885626 | 0.997078122 | 0.998402594 | 0.997740358 | 0.738844686 |
| Nebl | High | Q9DC07 | LIM zinc-binding domain-containing Nebulette OS=Mus musculus OX=10090 GN=Nebl PE=1 SV=1 | 0 | 48.418 | 37 | 10 | 18 | 9 | 270 | 31.1 | 8.31 | 0.82 | 0.978 | 0.839 | -0.29 | 0.29 | -0.03 | -0.01 | 0.16 | -0.25 | 0.413296827 | 0.876635562 | 0.301994072 | 0.997078122 | 0.998402594 | 0.997740358 | 0.900241969 |
| Cacna2d4 | High | F8VPL1 | Voltage-dependent calcium channel subunit alpha-2/delta-4 OS=Mus musculus OX=10090 GN=Ca | 0.005 | 3.716 | 1 | 2 | 2 | 2 | 1144 | 128.7 | 5.49 | 0.844 | 0.953 | 0.886 | -0.24 | 0.24 | -0.07 | -0.0233333 | 0.155 | -0.18 | 0.409070538 | 0.815915816 | 0.467248085 | 0.997078122 | 0.998402594 | 0.997740358 | 0.960359242 |
| Dnaja1 | High | P63037 | DnaJ homolog subfamily A member 1 OS=Mus musculus OX=10090 GN=Dnaja1 PE=1 SV=1 | 0 | 20.427 | 17 | 7 | 11 | 7 | 397 | 44.8 | 7.08 | 1.257 | 0.975 | 1.289 | 0.33 | -0.33 | -0.04 | -0.0133333 | -0.145 | 0.37 | 0.408072033 | 0.870645744 | 0.08315066 | 0.997078122 | 0.998402594 | 0.997740358 | 0.557020012 |
| Doc2b | High | P70169 | Double C2-like domain-containing protein beta OS=Mus musculus OX=10090 GN=Doc2b PE=1 SV= | 0.001 | 7.372 | 4 | 2 | 3 | 1 | 412 | 45.8 | 8.07 | 1.26 | 1.016 | 1.241 | 0.33 | -0.33 | 0.02 | 0.00666667 | -0.175 | 0.31 | 0.416827451 | 0.963274892 | 0.222076078 | 0.997078122 | 0.998402594 | 0.997740358 | 0.807447486 |
| Prkaca | High | P05132 | cAMP-dependent protein kinase catalytic subunit alpha OS=Mus musculus OX=10090 GN=Prkaca I | 0 | 78.822 | 42 | 16 | 25 | 8 | 351 | 40.5 | 8.79 | 1.278 | 1.008 | 1.267 | 0.35 | -0.35 | 0.01 | 0.00333333 | -0.18 | 0.34 | 0.407832971 | 0.951632405 | 0.10383007 | 0.997078122 | 0.998402594 | 0.997740358 | 0.612171613 |
| Camsap1 | High | A0A0A0M | Calmodulin-regulated spectrin-associated protein 1 OS=Mus musculus OX=10090 GN=Camsap1 P | 0.001 | 5.092 | 1 | 2 | 2 | 2 | 1582 | 175.9 | 6.9 | 1.257 | 1.053 | 1.193 | 0.33 | -0.33 | 0.07 | 0.02333333 | -0.2 | 0.26 | 0.416078334 | 0.936565597 | 0.30969229 | 0.997078122 | 0.998402594 | 0.997740358 | 0.904943958 |
| Ilf2 | High | Q9CXY6 | Interleukin enhancer-binding factor 2 OS=Mus musculus OX=10090 GN=Ilf2 PE=1 SV=1 | 0 | 13.139 | 13 | 4 | 4 | 4 | 390 | 43 | 5.26 | 1.221 | 1.097 | 1.113 | 0.29 | -0.29 | 0.13 | 0.04333333 | -0.21 | 0.16 | 0.416339708 | 0.843874997 | 0.531849277 | 0.997078122 | 0.998402594 | 0.997740358 | 0.96694628 |
| Gnb2l1; Ra | High | P68040 | Receptor of activated protein C kinase 1 OS=Mus musculus OX=10090 GN=Rack1 PE=1 SV=3 | 0 | 70.04 | 38 | 12 | 20 | 12 | 317 | 35.1 | 7.69 | 1.274 | 1.074 | 1.187 | 0.35 | -0.35 | 0.1 | 0.03333333 | -0.225 | 0.25 | 0.412949761 | 0.894250199 | 0.229142257 | 0.997078122 | 0.998402594 | 0.997740358 | 0.818865925 |
| Pde6d | High | O55057 | Retinal rod rhodopsin-sensitive cGMP 3',5'-cyclic phosphodiesterase subunit delta OS=Mus muscu | 0 | 25.587 | 33 | 4 | 6 | 4 | 150 | 17.3 | 5.67 | 1.282 | 1.088 | 1.178 | 0.36 | -0.36 | 0.12 | 0.04 | -0.24 | 0.24 | 0.410001183 | 0.862875507 | 0.335638459 | 0.997078122 | 0.998402594 | 0.997740358 | 0.906329766 |
| Atp9a | High | Q8C288 | Phospholipid-transporting ATPase OS=Mus musculus OX=10090 GN=Atp9a PE=1 SV=1 | 0.003 | 4.046 | 2 | 2 | 2 | 2 | 1031 | 116.8 | 7.25 | 0.826 | 0.838 | 0.986 | -0.28 | 0.28 | -0.25 | -0.0833333 | 0.265 | -0.02 | 0.418923989 | 0.505362917 | 0.999917596 | 0.997489268 | 0.995559182 | 0.996524225 | 0.999917596 |
| | High | A0A0A6YY | Glutathione peroxidase OS=Mus musculus OX=10090 GN=Gpx1 PE=1 SV=1 | 0 | 13.365 | 27 | 3 | 5 | 3 | 144 | 16.3 | 7.25 | 0.833 | 1.196 | 0.697 | -0.26 | 0.26 | 0.26 | 0.08666667 | 0 | -0.52 | 0.419134067 | 0.631092245 | 0.04649961 | 0.997489268 | 0.995559182 | 0.996524225 | 0.409467755 |
| | High | E9PZ88 | Alpha-mannosidase OS=Mus musculus OX=10090 GN=Man2c1 PE=1 SV=1 | 0 | 13.326 | 4 | 3 | 3 | 3 | 940 | 104.6 | 6.46 | 0.828 | 0.821 | 1.008 | -0.27 | 0.27 | -0.28 | -0.0933333 | 0.275 | 0.01 | 0.419836148 | 0.484727262 | 0.966702716 | 0.997822454 | 0.995559182 | 0.996690818 | 0.994815921 |
| Arvcf | High | P98203 | Armadillo repeat protein deleted in velo-cardio-facial syndrome homolog OS=Mus musculus OX=1 | 0 | 22.045 | 9 | 6 | 6 | 6 | 962 | 105 | 6.64 | 0.836 | 0.862 | 0.969 | -0.26 | 0.26 | -0.21 | -0.07 | 0.235 | -0.05 | 0.419985909 | 0.589525393 | 0.897291791 | 0.997822454 | 0.995559182 | 0.996690818 | 0.992535481 |
| Heph | High | Q9Z0Z4 | Hephaestin OS=Mus musculus OX=10090 GN=Heph PE=1 SV=3 | 0 | 107.242 | 24 | 23 | 34 | 22 | 1157 | 129.6 | 6.13 | 1.269 | 1.376 | 0.923 | 0.34 | -0.34 | 0.46 | 0.15333333 | -0.4 | -0.12 | 0.420226803 | 0.371643558 | 0.674777621 | 0.997971912 | 0.977313184 | 0.987642548 | 0.990250803 |
| Ola1 | High | Q9CZ30 | Obg-like ATPase 1 OS=Mus musculus OX=10090 GN=Ola1 PE=1 SV=1 | 0 | 15.551 | 15 | 6 | 8 | 6 | 396 | 44.7 | 7.81 | 0.803 | 0.741 | 1.084 | -0.32 | 0.32 | -0.43 | -0.1433333 | 0.375 | 0.12 | 0.420894586 | 0.315536584 | 0.644269732 | 0.99856391 | 0.955820031 | 0.97719197 | 0.985010103 |
| Tfrc | High | Q8C872 | Transferrin receptor protein 1 OS=Mus musculus OX=10090 GN=Tfrc PE=1 SV=1 | 0 | 10.524 | 7 | 4 | 4 | 4 | 515 | 57.3 | 6.48 | 1.218 | 1.427 | 0.854 | 0.28 | -0.28 | 0.51 | 0.17 | -0.395 | -0.23 | 0.423496052 | 0.314764426 | 0.353253425 | 0.99856391 | 0.955820031 | 0.97719197 | 0.91150113 |
| Hibch | High | Q8QZS1 | 3-hydroxyisobutyryl-CoA hydrolase, mitochondrial OS=Mus musculus OX=10090 GN=Hibch PE=1 S\ | 0 | 8.433 | 6 | 3 | 3 | 3 | 385 | 43 | 8.06 | 1.26 | 1.462 | 0.862 | 0.33 | -0.33 | 0.55 | 0.18333333 | -0.44 | -0.21 | 0.422383032 | 0.280290826 | 0.404984493 | 0.99856391 | 0.955820031 | 0.97719197 | 0.934464219 |
| Tmx4 | High | Q8C0L0 | Thioredoxin-related transmembrane protein 4 OS=Mus musculus OX=10090 GN=Tmx4 PE=1 SV=2 | 0.001 | 6.886 | 10 | 3 | 3 | 3 | 335 | 37.1 | 4.37 | 1.24 | 1.405 | 0.882 | 0.31 | -0.31 | 0.49 | 0.16333333 | -0.4 | -0.18 | 0.423182691 | 0.33407517 | 0.502141444 | 0.99856391 | 0.97322306 | 0.985893485 | 0.964378027 |
| Ndufs1 | High | Q91VD9 | NADH-ubiquinone oxidoreductase 75 kDa subunit, mitochondrial OS=Mus musculus OX=10090 GN | 0 | 101.147 | 34 | 21 | 35 | 21 | 727 | 79.7 | 5.72 | 1.268 | 1.356 | 0.935 | 0.34 | -0.34 | 0.44 | 0.14666667 | -0.39 | -0.1 | 0.423193215 | 0.396479745 | 0.738465833 | 0.99856391 | 0.985399792 | 0.991981851 | 0.992535481 |
| Atp2b1 | High | G5E829 | Plasma membrane calcium-transporting ATPase 1 OS=Mus musculus OX=10090 GN=Atp2b1 PE=1 | 0 | 210.469 | 32 | 35 | 72 | 18 | 1220 | 134.7 | 5.91 | 0.788 | 0.871 | 0.905 | -0.34 | 0.34 | -0.2 | -0.0666667 | 0.27 | -0.14 | 0.424928391 | 0.610508566 | 0.586230931 | 0.99856391 | 0.995559182 | 0.997061546 | 0.973095093 |
| Flot2 | High | Q60634 | Flotillin-2 OS=Mus musculus OX=10090 GN=Flot2 PE=1 SV=2 | 0 | 36.824 | 20 | 7 | 10 | 7 | 428 | 47 | 5.2 | 0.829 | 0.841 | 0.985 | -0.27 | 0.27 | -0.25 | -0.0833333 | 0.26 | -0.02 | 0.421877275 | 0.537829439 | 0.900824911 | 0.99856391 | 0.995559182 | 0.997061546 | 0.992535481 |
| Drg2 | High | Q9QXB9 | Developmentally-regulated GTP-binding protein 2 OS=Mus musculus OX=10090 GN=Drg2 PE=1 S\ | 0.001 | 6.008 | 6 | 2 | 2 | 2 | 364 | 40.7 | 8.88 | 0.824 | 0.896 | 0.921 | -0.28 | 0.28 | -0.16 | -0.0533333 | 0.22 | -0.12 | 0.421850564 | 0.668331333 | 0.647983401 | 0.99856391 | 0.995559182 | 0.997061546 | 0.985010103 |
| Bcl2l1 | Medium | Q9QWX2 | Apoptosis regulator Bcl-X (Fragment) OS=Mus musculus OX=10090 GN=Bcl2l1 PE=1 SV=1 | 0.015 | 2.178 | 5 | 1 | 1 | 1 | 188 | 21.1 | 4.73 | 0.826 | 0.899 | 0.919 | -0.28 | 0.28 | -0.15 | -0.05 | 0.215 | -0.12 | 0.424245592 | 0.655768498 | 0.646482686 | 0.99856391 | 0.995559182 | 0.997061546 | 0.985010103 |
| | High | D3YWB9 | Contactin-associated protein-like 4 OS=Mus musculus OX=10090 GN=Cntnap4 PE=1 SV=1 | 0.001 | 4.359 | 1 | 1 | 1 | 1 | 1310 | 144.7 | 7.02 | 1.281 | 1.145 | 1.118 | 0.36 | -0.36 | 0.2 | 0.06666667 | -0.28 | 0.16 | 0.424102952 | 0.76001522 | 0.547319707 | 0.99856391 | 0.995559182 | 0.997061546 | 0.96694628 |
| Txn1 | High | P10639 | Thioredoxin OS=Mus musculus OX=10090 GN=Txn PE=1 SV=3 | 0 | 48.546 | 74 | 7 | 21 | 7 | 105 | 11.7 | 4.92 | 1.267 | 1.221 | 1.038 | 0.34 | -0.34 | 0.29 | 0.09666667 | -0.315 | 0.05 | 0.423827015 | 0.598294017 | 0.734882352 | 0.99856391 | 0.995559182 | 0.997061546 | 0.992535481 |
| Fam149a | High | Q8CFV2 | Protein FAM149A OS=Mus musculus OX=10090 GN=Fam149a PE=2 SV=1 | 0 | 10.863 | 5 | 3 | 3 | 3 | 787 | 85 | 7.72 | 1.244 | 1.309 | 0.95 | 0.31 | -0.31 | 0.39 | 0.13 | -0.35 | -0.07 | 0.423756194 | 0.454901155 | 0.773225161 | 0.99856391 | 0.995559182 | 0.997061546 | 0.992535481 |
| Clcn3 | High | Q8K4W8 | Chloride channel protein OS=Mus musculus OX=10090 GN=Clcn3 PE=1 SV=1 | 0.001 | 6.418 | 4 | 3 | 3 | 3 | 791 | 87.9 | 7.03 | 0.813 | 0.973 | 0.836 | -0.3 | 0.3 | -0.04 | -0.0133333 | 0.17 | -0.26 | 0.424665588 | 0.864672521 | 0.315912441 | 0.99856391 | 0.998402594 | 0.998483252 | 0.904943958 |
| Dpy30 | High | Q99LT0 | Protein dpy-30 homolog OS=Mus musculus OX=10090 GN=Dpy30 PE=1 SV=1 | 0.001 | 6.414 | 25 | 2 | 2 | 2 | 99 | 11.2 | 4.88 | 0.857 | 1.077 | 0.795 | -0.22 | 0.22 | 0.11 | 0.03666667 | 0.055 | -0.33 | 0.423433595 | 0.891299093 | 0.160605145 | 0.99856391 | 0.998402594 | 0.998483252 | 0.733684585 |
| Pfdn1 | High | Q9CQF7 | Prefoldin 1 OS=Mus musculus OX=10090 GN=Pfdn1 PE=1 SV=1 | 0 | 11.841 | 35 | 5 | 5 | 5 | 122 | 14.2 | 6.81 | 0.808 | 0.709 | 1.139 | -0.31 | 0.31 | -0.5 | -0.1666667 | 0.405 | 0.19 | 0.426021429 | 0.253661573 | 0.453302247 | 0.999456968 | 0.950034749 | 0.974745859 | 0.959744396 |
| Copa | High | Q8CIE6 | Coatomer subunit alpha OS=Mus musculus OX=10090 GN=Copa PE=1 SV=2 | 0 | 11.261 | 4 | 3 | 4 | 3 | 1224 | 138.3 | 7.65 | 0.815 | 0.949 | 0.86 | -0.29 | 0.29 | -0.08 | -0.0266667 | 0.185 | -0.22 | 0.425999881 | 0.804477772 | 0.37429776 | 0.999456968 | 0.998402594 | 0.998929781 | 0.920164724 |
| Cd177 | High | Q8R2S8 | CD177 antigen OS=Mus musculus OX=10090 GN=Cd177 PE=2 SV=1 | 0.008 | 2.952 | 1 | 1 | 2 | 1 | 817 | 87 | 5.67 | 0.841 | 0.555 | 1.515 | -0.25 | 0.25 | -0.85 | -0.2833333 | 0.55 | 0.6 | 0.43564476 | 0.051615133 | 0.01570741 | 0.999652205 | 0.541958901 | 0.770805553 | 0.213248064 |
| 3110002H | High | Q8VC42 | Regulator of MON1-CCZ1 complex OS=Mus musculus OX=10090 GN=Rmc1 PE=2 SV=1 | 0.006 | 3.217 | 2 | 1 | 1 | 1 | 657 | 74.9 | 7.83 | 0.889 | 0.567 | 1.567 | -0.17 | 0.17 | -0.82 | -0.2733333 | 0.495 | 0.65 | 0.61968299 | 0.062315318 | 0.014222653 | 0.999652205 | 0.587451609 | 0.793551907 | 0.196904733 |
| | Medium | E9PUP9 | Receptor protein-tyrosine kinase OS=Mus musculus OX=10090 GN=Epha5 PE=1 SV=1 | 0.037 | 1.738 | 1 | 1 | 1 | 1 | 1017 | 112.7 | 6.79 | 0.82 | 0.571 | 1.436 | -0.29 | 0.29 | -0.81 | -0.27 | 0.55 | 0.52 | 0.435219684 | 0.065058429 | 0.067426069 | 0.999652205 | 0.58835905 | 0.794005627 | 0.499896415 |
| Strn3 | High | Q9ERG2 | Striatin-3 OS=Mus musculus OX=10090 GN=Strn3 PE=1 SV=1 | 0.005 | 3.653 | 2 | 2 | 2 | 2 | 796 | 87.1 | 5.29 | 0.817 | 0.572 | 1.43 | -0.29 | 0.29 | -0.81 | -0.27 | 0.55 | 0.52 | 0.430683463 | 0.071652552 | 0.022360202 | 0.999652205 | 0.625156241 | 0.812404223 | 0.270201833 |
| Arl1 | High | P61211 | ADP-ribosylation factor-like protein 1 OS=Mus musculus OX=10090 GN=Arl1 PE=1 SV=1 | 0 | 4.718 | 14 | 2 | 2 | 2 | 181 | 20.4 | 5.72 | 1.163 | 1.862 | 0.625 | 0.22 | -0.22 | 0.9 | 0.3 | -0.56 | -0.68 | 0.639246986 | 0.071571687 | 0.023284476 | 0.999652205 | 0.625156241 | 0.812404223 | 0.278965935 |
| Tmx1 | High | Q8VBT0 | Thioredoxin-related transmembrane protein 1 OS=Mus musculus OX=10090 GN=Tmx1 PE=1 SV=1 | 0 | 29.779 | 19 | 7 | 12 | 7 | 278 | 31.4 | 5.29 | 1.23 | 1.833 | 0.671 | 0.3 | -0.3 | 0.87 | 0.29 | -0.585 | -0.58 | 0.453348919 | 0.076356415 | 0.016179123 | 0.999652205 | 0.647701084 | 0.823676644 | 0.218068136 |
| Mast1 | High | Q9R1L5 | Microtubule-associated serine/threonine-protein kinase 1 OS=Mus musculus OX=10090 GN=Mast | 0.008 | 2.919 | 1 | 1 | 1 | 1 | 1570 | 170.9 | 8.44 | 0.873 | 0.594 | 1.469 | -0.2 | 0.2 | -0.75 | -0.25 | 0.475 | 0.56 | 0.531306161 | 0.085157869 | 0.030026722 | 0.999652205 | 0.68454712 | 0.842099662 | 0.330765876 |
| Fam3a | High | Q9D8T0 | Protein FAM3A OS=Mus musculus OX=10090 GN=Fam3a PE=2 SV=1 | 0.006 | 3.129 | 7 | 2 | 3 | 2 | 230 | 25.4 | 8.44 | 1.219 | 1.798 | 0.678 | 0.29 | -0.29 | 0.85 | 0.28333333 | -0.57 | -0.56 | 0.483140702 | 0.08656946 | 0.021867189 | 0.999652205 | 0.68454712 | 0.842099662 | 0.267705959 |
| | Medium | A0A1L1SS | 39S ribosomal protein L4, mitochondrial (Fragment) OS=Mus musculus OX=10090 GN=Mrpl4 PE= | 0.047 | 1.563 | 7 | 1 | 1 | 1 | 264 | 29.4 | 9.23 | 1.214 | 1.792 | 0.677 | 0.28 | -0.28 | 0.84 | 0.28 | -0.56 | -0.56 | 0.556037194 | 0.088050855 | 0.022709675 | 0.999652205 | 0.694375726 | 0.847013965 | 0.273247103 |
| Lpcat2 | High | Q8BYI6 | Lysophosphatidylcholine acyltransferase 2 OS=Mus musculus OX=10090 GN=Lpcat2 PE=1 SV=1 | 0 | 115.375 | 41 | 22 | 42 | 22 | 544 | 60.2 | 5.83 | 1.262 | 1.785 | 0.707 | 0.34 | -0.34 | 0.84 | 0.28 | -0.59 | -0.5 | 0.432285268 | 0.090702979 | 0.032623275 | 0.999652205 | 0.696517952 | 0.848085078 | 0.349748955 |
| Lrrc8a | High | Q80WG5 | Volume-regulated anion channel subunit LRRC8A OS=Mus musculus OX=10090 GN=Lrrc8a PE=1 S | 0 | 14.602 | 10 | 6 | 6 | 6 | 810 | 94.1 | 7.94 | 0.836 | 0.599 | 1.395 | -0.26 | 0.26 | -0.74 | -0.2466667 | 0.5 | 0.48 | 0.482637405 | 0.097686107 | 0.057543559 | 0.999652205 | 0.719745075 | 0.85969864 | 0.463572893 |
| Fbxo27 | Low | Q6DIA9 | F-box only protein 27 OS=Mus musculus OX=10090 GN=Fbxo27 PE=2 SV=1 | 0.059 | 1.338 | 3 | 1 | 1 | 1 | 280 | 31.6 | 8.63 | 0.857 | 0.606 | 1.414 | -0.22 | 0.22 | -0.72 | -0.24 | 0.47 | 0.5 | 0.478045064 | 0.098507042 | 0.051770391 | 0.999652205 | 0.720327908 | 0.859990057 | 0.431159947 |

| | | | | | | | | | | | | | | | | | | | | | | | | | | | | |
|---|---|---|---|---|---|---|---|---|---|---|---|---|---|---|---|---|---|---|---|---|---|---|---|---|---|---|---|---|
| | Medium | S4R2G0 | Diacylglycerol kinase OS=Mus musculus OX=10090 GN=Dgki PE=1 SV=1 | 0.029 | 1.918 | 1 | 1 | 1 | 1 | 841 | 93 | 8.63 | 1.012 | 0.609 | 1.661 | 0.02 | -0.02 | -0.72 | -0.24 | 0.35 | 0.73 | 0.985499843 | 0.100130741 | 0.011348679 | 0.999652205 | 0.728188149 | 0.863920177 | 0.173441884 |
| Psmb7 | High | P70195 | Proteasome subunit beta type-7 OS=Mus musculus OX=10090 GN=Psmb7 PE=1 SV=1 | 0 | 31.806 | 24 | 7 | 15 | 7 | 277 | 29.9 | 7.99 | 0.805 | 0.602 | 1.337 | -0.31 | 0.31 | -0.73 | -0.2433333 | 0.52 | 0.42 | 0.427083445 | 0.100844298 | 0.083461644 | 0.999652205 | 0.730697757 | 0.865174981 | 0.557770483 |
| Blvra | High | Q9CY64 | Biliverdin reductase A OS=Mus musculus OX=10090 GN=Blvra PE=1 SV=1 | 0 | 12.116 | 16 | 4 | 4 | 4 | 295 | 33.5 | 7.02 | 0.795 | 0.603 | 1.319 | -0.33 | 0.33 | -0.73 | -0.2433333 | 0.53 | 0.4 | 0.461228188 | 0.101523215 | 0.085273144 | 0.999652205 | 0.73167181 | 0.865662008 | 0.56316433 |
| Cab39 | High | Q06138 | Calcium-binding protein 39 OS=Mus musculus OX=10090 GN=Cab39 PE=1 SV=2 | 0 | 27.339 | 26 | 9 | 9 | 6 | 341 | 39.8 | 6.89 | 0.844 | 0.604 | 1.399 | -0.24 | 0.24 | -0.73 | -0.2433333 | 0.485 | 0.48 | 0.540332411 | 0.102188672 | 0.04509135 | 0.999652205 | 0.734610978 | 0.867131592 | 0.405968973 |
| Spryd7 | High | Q3TFQ1 | SPRY domain-containing protein 7 OS=Mus musculus OX=10090 GN=Spryd7 PE=1 SV=2 | 0 | 12.516 | 23 | 3 | 3 | 3 | 196 | 21.7 | 6.7 | 1.143 | 1.749 | 0.654 | 0.19 | -0.19 | 0.81 | 0.27 | -0.5 | -0.61 | 0.697681389 | 0.102337132 | 0.010458405 | 0.999652205 | 0.734704609 | 0.867178407 | 0.163343383 |
| Myef2 | Medium | A2ATP6 | Myelin expression factor 2 OS=Mus musculus OX=10090 GN=Myef2 PE=1 SV=2 | 0.015 | 2.188 | 2 | 1 | 1 | 1 | 567 | 61.1 | 8.75 | 1.223 | 1.712 | 0.715 | 0.29 | -0.29 | 0.78 | 0.26 | -0.535 | -0.48 | 0.469999557 | 0.110294494 | 0.071459384 | 0.999652205 | 0.768225125 | 0.883938665 | 0.508467979 |
| Tmem35; | High | Q9D328 | Novel acetylcholine receptor chaperone OS=Mus musculus OX=10090 GN=Tmem35a PE=1 SV=1 | 0.001 | 5.524 | 9 | 1 | 1 | 1 | 167 | 18.5 | 9.99 | 1.216 | 1.702 | 0.715 | 0.28 | -0.28 | 0.77 | 0.25666667 | -0.525 | -0.48 | 0.490702673 | 0.111136458 | 0.06031574 | 0.999652205 | 0.770262199 | 0.884957202 | 0.474986455 |
| Psmb1 | High | O09061 | Proteasome subunit beta type-1 OS=Mus musculus OX=10090 GN=Psmb1 PE=1 SV=1 | 0 | 27.554 | 37 | 6 | 7 | 6 | 240 | 26.4 | 7.81 | 0.862 | 0.612 | 1.409 | -0.21 | 0.21 | -0.71 | -0.2366667 | 0.46 | 0.5 | 0.554920434 | 0.11132368 | 0.049940884 | 0.999652205 | 0.770607254 | 0.88512973 | 0.427566646 |
| Erlin2 | High | Q8BFZ9 | Erlin-2 OS=Mus musculus OX=10090 GN=Erlin2 PE=1 SV=1 | 0 | 46.97 | 35 | 11 | 16 | 11 | 340 | 37.8 | 5.5 | 1.147 | 1.718 | 0.668 | 0.2 | -0.2 | 0.78 | 0.26 | -0.49 | -0.58 | 0.674746453 | 0.115457564 | 0.017614299 | 0.999652205 | 0.77945068 | 0.889551442 | 0.23459234 |
| Sf3b3 | Medium | Q921M3 | Splicing factor 3B subunit 3 OS=Mus musculus OX=10090 GN=Sf3b3 PE=1 SV=1 | 0.016 | 2.149 | 1 | 1 | 1 | 1 | 1217 | 135.5 | 5.26 | 1.24 | 1.707 | 0.726 | 0.31 | -0.31 | 0.77 | 0.25666667 | -0.54 | -0.46 | 0.435221608 | 0.113191237 | 0.056149859 | 0.999652205 | 0.77945068 | 0.889551442 | 0.454949834 |
| Psma3 | High | O70435 | Proteasome subunit alpha type-3 OS=Mus musculus OX=10090 GN=Psma3 PE=1 SV=3 | 0 | 29.978 | 32 | 10 | 14 | 10 | 255 | 28.4 | 5.44 | 0.859 | 0.617 | 1.391 | -0.22 | 0.22 | -0.7 | -0.2333333 | 0.46 | 0.48 | 0.627939387 | 0.117634404 | 0.039139244 | 0.999652205 | 0.779640783 | 0.889646494 | 0.380996077 |
| | Medium | E9Q8X9 | Actin filament-associated protein 1 (Fragment) OS=Mus musculus OX=10090 GN=Afap1 PE=1 SV | 0.025 | 1.999 | 1 | 1 | 1 | 1 | 627 | 69.6 | 8.7 | 0.842 | 0.631 | 1.335 | -0.25 | 0.25 | -0.67 | -0.2233333 | 0.46 | 0.42 | 0.501087621 | 0.119585478 | 0.168399955 | 0.999652205 | 0.786053068 | 0.892852636 | 0.738806217 |
| Bri3bp | High | Q8BXV2 | BRI3-binding protein OS=Mus musculus OX=10090 GN=Bri3bp PE=1 SV=1 | 0.008 | 3.003 | 4 | 1 | 1 | 1 | 253 | 28.2 | 9.52 | 1.089 | 1.671 | 0.652 | 0.12 | -0.12 | 0.74 | 0.24666667 | -0.43 | -0.62 | 0.80030441 | 0.126164488 | 0.016172801 | 0.999652205 | 0.799286338 | 0.899469271 | 0.218068136 |
| Dhcr7 | Low | Q711T1 | 7-dehydrocholesterol reductase (Fragment) OS=Mus musculus OX=10090 GN=Dhcr7 PE=2 SV=1 | 0.054 | 1.404 | 58 | 1 | 1 | 1 | 12 | 1.3 | 10.01 | 1.091 | 1.679 | 0.65 | 0.13 | -0.13 | 0.75 | 0.25 | -0.44 | -0.62 | 0.75105115 | 0.128306984 | 0.013680496 | 0.999652205 | 0.799286338 | 0.899469271 | 0.191288138 |
| Akap9 | Medium | Q70FJ1 | A-kinase anchor protein 9 OS=Mus musculus OX=10090 GN=Akap9 PE=1 SV=2 | 0.01 | 2.642 | 0 | 2 | 2 | 1 | 3797 | 435.9 | 5.03 | 1.176 | 1.701 | 0.692 | 0.23 | -0.23 | 0.77 | 0.25666667 | -0.5 | -0.53 | 0.613684335 | 0.12838517 | 0.084603957 | 0.999652205 | 0.799286338 | 0.899469271 | 0.560726223 |
| Tubb6 | High | Q922F4 | Tubulin beta-6 chain OS=Mus musculus OX=10090 GN=Tubb6 PE=1 SV=1 | 0 | 154.563 | 31 | 13 | 77 | 1 | 447 | 50.1 | 4.89 | 1.252 | 1.684 | 0.744 | 0.32 | -0.32 | 0.75 | 0.25 | -0.535 | -0.43 | 0.450859892 | 0.128495235 | 0.142776392 | 0.999652205 | 0.799286338 | 0.899469271 | 0.703085292 |
| Oat | High | P29758 | Ornithine aminotransferase, mitochondrial OS=Mus musculus OX=10090 GN=Oat PE=1 SV=1 | 0 | 18.483 | 13 | 4 | 5 | 4 | 439 | 48.3 | 6.62 | 1.166 | 1.685 | 0.692 | 0.22 | -0.22 | 0.75 | 0.25 | -0.485 | -0.53 | 0.572313207 | 0.130125828 | 0.042211242 | 0.999652205 | 0.803541319 | 0.901596762 | 0.399120459 |
| Sec23b | High | Q9D662 | Protein transport protein Sec23B OS=Mus musculus OX=10090 GN=Sec23b PE=1 SV=1 | 0 | 12.272 | 6 | 4 | 4 | 3 | 767 | 86.4 | 6.96 | 0.909 | 0.631 | 1.44 | -0.14 | 0.14 | -0.66 | -0.22 | 0.4 | 0.53 | 0.644581909 | 0.134645332 | 0.019572944 | 0.999652205 | 0.817450082 | 0.908551144 | 0.249847648 |
| Thop1 | High | A0A0R4IZ | Thimet oligopeptidase OS=Mus musculus OX=10090 GN=Thop1 PE=1 SV=1 | 0 | 17.096 | 8 | 5 | 5 | 5 | 687 | 78 | 5.99 | 0.911 | 0.633 | 1.438 | -0.14 | 0.14 | -0.66 | -0.22 | 0.4 | 0.52 | 0.766074899 | 0.136811833 | 0.026205308 | 0.999652205 | 0.817450082 | 0.908551144 | 0.306110757 |
| Polr2e | High | Q80UW8 | DNA-directed RNA polymerases I, II, and III subunit RPABC1 OS=Mus musculus OX=10090 GN=Pol | 0 | 10.813 | 14 | 2 | 4 | 2 | 210 | 24.6 | 5.95 | 0.955 | 0.635 | 1.504 | -0.07 | 0.07 | -0.66 | -0.22 | 0.365 | 0.59 | 0.842567927 | 0.13863746 | 0.010880476 | 0.999652205 | 0.82258226 | 0.911117233 | 0.168993987 |
| Dyrk2 | High | Q5U4C9 | Dual specificity tyrosine-phosphorylation-regulated kinase 2 OS=Mus musculus OX=10090 GN=Dy | 0 | 14.923 | 6 | 3 | 4 | 3 | 599 | 66.5 | 9.67 | 1.262 | 1.65 | 0.765 | 0.34 | -0.34 | 0.72 | 0.24 | -0.53 | -0.39 | 0.434478722 | 0.138416041 | 0.093659374 | 0.999652205 | 0.82258226 | 0.911117233 | 0.582204115 |
| Cmpk1 | High | Q9DBP5 | UMP-CMP kinase OS=Mus musculus OX=10090 GN=Cmpk1 PE=1 SV=1 | 0 | 24.112 | 39 | 6 | 9 | 6 | 196 | 22.2 | 5.83 | 0.884 | 0.637 | 1.389 | -0.18 | 0.18 | -0.65 | -0.2166667 | 0.415 | 0.47 | 0.634835725 | 0.141396028 | 0.049440484 | 0.999652205 | 0.83278102 | 0.916216612 | 0.427566646 |
| Hook3 | High | Q8BUK6 | Protein Hook homolog 3 OS=Mus musculus OX=10090 GN=Hook3 PE=1 SV=2 | 0.001 | 6.691 | 3 | 3 | 3 | 3 | 718 | 83.2 | 5.19 | 0.788 | 0.637 | 1.237 | -0.34 | 0.34 | -0.65 | -0.2166667 | 0.495 | 0.31 | 0.429192897 | 0.141839974 | 0.187634908 | 0.999652205 | 0.834519134 | 0.91708567 | 0.774149321 |
| Myl12b | High | Q3THE2 | Myosin regulatory light chain 12B OS=Mus musculus OX=10090 GN=Myl12b PE=1 SV=2 | 0 | 18.947 | 35 | 5 | 6 | 5 | 172 | 19.8 | 4.84 | 0.833 | 0.651 | 1.28 | -0.26 | 0.26 | -0.62 | -0.2066667 | 0.44 | 0.36 | 0.427766396 | 0.14331945 | 0.119050092 | 0.999652205 | 0.837891129 | 0.918771667 | 0.662216135 |
| Xylb | Medium | Q3TNA1 | Xylulose kinase OS=Mus musculus OX=10090 GN=Xylb PE=1 SV=1 | 0.01 | 2.546 | 1 | 1 | 1 | 1 | 551 | 59.5 | 6.9 | 0.902 | 0.648 | 1.392 | -0.15 | 0.15 | -0.63 | -0.21 | 0.39 | 0.48 | 0.59178419 | 0.147975404 | 0.050128803 | 0.999652205 | 0.845081896 | 0.922367051 | 0.427566646 |
| Psmb2 | High | Q9R1P3 | Proteasome subunit beta type-2 OS=Mus musculus OX=10090 GN=Psmb2 PE=1 SV=1 | 0 | 23.183 | 23 | 4 | 7 | 4 | 201 | 22.9 | 7.02 | 0.919 | 0.644 | 1.426 | -0.12 | 0.12 | -0.63 | -0.21 | 0.375 | 0.51 | 0.730994959 | 0.151254905 | 0.03449755 | 0.999652205 | 0.847684847 | 0.923668526 | 0.364272011 |
| Prkaa2 | High | Q8BRK8 | 5'-AMP-activated protein kinase catalytic subunit alpha-2 OS=Mus musculus OX=10090 GN=Prkaa | 0.001 | 6.813 | 3 | 2 | 2 | 2 | 552 | 62 | 7.87 | 0.988 | 0.652 | 1.517 | -0.02 | 0.02 | -0.62 | -0.2066667 | 0.32 | 0.6 | 0.910456152 | 0.150449915 | 0.048048387 | 0.999652205 | 0.847684847 | 0.923668526 | 0.419637552 |
| Gaa | High | P70699 | Lysosomal alpha-glucosidase OS=Mus musculus OX=10090 GN=Gaa PE=1 SV=2 | 0 | 8.891 | 4 | 2 | 2 | 2 | 953 | 106.2 | 5.83 | 1.079 | 1.643 | 0.657 | 0.11 | -0.11 | 0.72 | 0.24 | -0.415 | -0.61 | 0.833640639 | 0.155509018 | 0.035369771 | 0.999652205 | 0.847684847 | 0.923668526 | 0.367937483 |
| Sh3bgrl | High | Q9JJU8 | SH3 domain-binding glutamic acid-rich-like protein OS=Mus musculus OX=10090 GN=Sh3bgrl PE= | 0 | 17.763 | 21 | 3 | 4 | 3 | 114 | 12.8 | 4.92 | 1.134 | 1.645 | 0.689 | 0.18 | -0.18 | 0.72 | 0.24 | -0.45 | -0.54 | 0.660164004 | 0.14984979 | 0.035131695 | 0.999652205 | 0.847684847 | 0.923668526 | 0.367937483 |
| Gde1 | High | Q9JL56 | Glycerophosphodiester phosphodiesterase 1 OS=Mus musculus OX=10090 GN=Gde1 PE=1 SV=1 | 0.001 | 4.612 | 3 | 1 | 1 | 1 | 331 | 37.6 | 6.9 | 1.164 | 1.634 | 0.712 | 0.22 | -0.22 | 0.71 | 0.23666667 | -0.465 | -0.49 | 0.598805139 | 0.155723732 | 0.042029726 | 0.999652205 | 0.847684847 | 0.923668526 | 0.398749023 |
| Grk1 | High | G3X905 | G protein-coupled receptor kinase OS=Mus musculus OX=10090 GN=Grk1 PE=1 SV=1 | 0 | 609.561 | 73 | 42 | 476 | 42 | 564 | 63.8 | 5.54 | 1.22 | 1.646 | 0.741 | 0.29 | -0.29 | 0.72 | 0.24 | -0.505 | -0.43 | 0.501097918 | 0.14912013 | 0.067413311 | 0.999652205 | 0.847684847 | 0.923668526 | 0.499896415 |
| Srp54c | High | E9PXC0 | Signal recognition particle 54 kDa protein OS=Mus musculus OX=10090 GN=Srp54c PE=1 SV=1 | 0 | 48.12 | 27 | 12 | 18 | 12 | 504 | 55.8 | 8.92 | 1.14 | 1.631 | 0.699 | 0.19 | -0.19 | 0.71 | 0.23666667 | -0.45 | -0.52 | 0.618417246 | 0.157203141 | 0.027269081 | 0.999652205 | 0.849407305 | 0.924529755 | 0.315745518 |
| Prkag1 | High | O54950 | 5'-AMP-activated protein kinase subunit gamma-1 OS=Mus musculus OX=10090 GN=Prkag1 PE=1 | 0 | 34.372 | 26 | 8 | 12 | 8 | 330 | 37.5 | 7.14 | 0.915 | 0.653 | 1.403 | -0.13 | 0.13 | -0.62 | -0.2066667 | 0.375 | 0.49 | 0.690329189 | 0.163164601 | 0.045212258 | 0.999652205 | 0.869444647 | 0.934548426 | 0.405968973 |
| M6pr | High | P24668 | Cation-dependent mannose-6-phosphate receptor OS=Mus musculus OX=10090 GN=M6pr PE=1 S | 0 | 9.689 | 13 | 3 | 3 | 3 | 278 | 31.2 | 5.39 | 1.216 | 1.618 | 0.751 | 0.28 | -0.28 | 0.69 | 0.23 | -0.485 | -0.41 | 0.514324526 | 0.164255389 | 0.101870321 | 0.999652205 | 0.870491463 | 0.935071834 | 0.610114517 |
| Setd7 | Medium | Q8VHL1 | Histone-lysine N-methyltransferase SETD7 OS=Mus musculus OX=10090 GN=Setd7 PE=1 SV=2 | 0.01 | 2.568 | 2 | 1 | 1 | 1 | 366 | 40.5 | 4.65 | 1.238 | 1.598 | 0.775 | 0.31 | -0.31 | 0.68 | 0.22666667 | -0.495 | -0.37 | 0.458061567 | 0.164112492 | 0.155850885 | 0.999652205 | 0.870491463 | 0.935071834 | 0.730648756 |
| Ube2n | High | P61089 | Ubiquitin-conjugating enzyme E2 N OS=Mus musculus OX=10090 GN=Ube2n PE=1 SV=1 | 0 | 30.07 | 53 | 8 | 10 | 8 | 152 | 17.1 | 6.57 | 0.927 | 0.655 | 1.414 | -0.11 | 0.11 | -0.61 | -0.2033333 | 0.36 | 0.5 | 0.817035748 | 0.166809564 | 0.019560118 | 0.999652205 | 0.871008827 | 0.935330516 | 0.249847648 |
| Rexo1 | Low | Q7TT28 | RNA exonuclease 1 homolog OS=Mus musculus OX=10090 GN=Rexo1 PE=1 SV=1 | 0.054 | 1.405 | 1 | 1 | 1 | 1 | 1213 | 130.7 | 8.97 | 1.059 | 0.665 | 1.593 | 0.08 | -0.08 | -0.59 | -0.1966667 | 0.255 | 0.67 | 0.894645907 | 0.16696653 | 0.011721151 | 0.999652205 | 0.871008827 | 0.935330516 | 0.178588299 |
| Bphl | Medium | Q8R164 | Valacyclovir hydrolase OS=Mus musculus OX=10090 GN=Bphl PE=1 SV=1 | 0.021 | 2.011 | 3 | 1 | 1 | 1 | 291 | 32.8 | 8.94 | 1.171 | 1.613 | 0.726 | 0.23 | -0.23 | 0.69 | 0.23 | -0.46 | -0.46 | 0.615417322 | 0.165089487 | 0.111606236 | 0.999652205 | 0.871008827 | 0.935330516 | 0.633376688 |
| Rdh13 | High | Q8CEE7 | Retinol dehydrogenase 13 OS=Mus musculus OX=10090 GN=Rdh13 PE=1 SV=1 | 0.001 | 5.608 | 6 | 2 | 2 | 2 | 334 | 36.4 | 8.85 | 1.215 | 1.6 | 0.76 | 0.28 | -0.28 | 0.68 | 0.22666667 | -0.48 | -0.4 | 0.499964719 | 0.169813501 | 0.127675981 | 0.999652205 | 0.875132627 | 0.937392416 | 0.679202302 |
| Ap3m2 | High | Q8R2R9 | AP-3 complex subunit mu-2 OS=Mus musculus OX=10090 GN=Ap3m2 PE=1 SV=1 | 0 | 10.038 | 15 | 4 | 4 | 4 | 418 | 46.9 | 7.56 | 1.164 | 1.607 | 0.724 | 0.22 | -0.22 | 0.68 | 0.22666667 | -0.45 | -0.47 | 0.615014343 | 0.170866778 | 0.075761653 | 0.999652205 | 0.875484829 | 0.937568517 | 0.523043272 |
| Scfd1 | High | Q8BRF7 | Sec1 family domain-containing protein 1 OS=Mus musculus OX=10090 GN=Scfd1 PE=1 SV=1 | 0 | 30.788 | 12 | 5 | 6 | 5 | 639 | 72.3 | 6.38 | 1.074 | 1.604 | 0.669 | 0.1 | -0.1 | 0.68 | 0.22666667 | -0.39 | -0.58 | 0.83022804 | 0.172571335 | 0.027762179 | 0.999652205 | 0.875662873 | 0.937657539 | 0.315745518 |
| Micu1 | High | Q8VCX5 | Calcium uptake protein 1, mitochondrial OS=Mus musculus OX=10090 GN=Micu1 PE=1 SV=1 | 0.001 | 6.064 | 5 | 2 | 2 | 2 | 477 | 54.3 | 8.59 | 1.217 | 1.585 | 0.768 | 0.28 | -0.28 | 0.66 | 0.22 | -0.47 | -0.38 | 0.480879786 | 0.172473164 | 0.138799165 | 0.999652205 | 0.875662873 | 0.937657539 | 0.701851981 |
| Psmb6 | High | Q60692 | Proteasome subunit beta type-6 OS=Mus musculus OX=10090 GN=Psmb6 PE=1 SV=3 | 0 | 24.852 | 33 | 5 | 6 | 5 | 238 | 25.4 | 5.11 | 0.91 | 0.661 | 1.376 | -0.14 | 0.14 | -0.6 | -0.2 | 0.37 | 0.46 | 0.676801134 | 0.175499902 | 0.061872639 | 0.999652205 | 0.875938481 | 0.937795343 | 0.476538303 |
| Dner | Medium | Q8JZM4 | Delta and Notch-like epidermal growth factor-related receptor OS=Mus musculus OX=10090 GN= | 0.047 | 1.567 | 1 | 1 | 1 | 1 | 737 | 78.7 | 5.1 | 1.087 | 1.579 | 0.688 | 0.12 | -0.12 | 0.66 | 0.22 | -0.39 | -0.54 | 0.783503793 | 0.174204378 | 0.043494231 | 0.999652205 | 0.875938481 | 0.937795343 | 0.404826557 |
| | Medium | A0A0G2JE | (Lyso)-N-acylphosphatidylethanolamine lipase (Fragment) OS=Mus musculus OX=10090 GN=Abh | 0.042 | 1.641 | 21 | 1 | 1 | 1 | 56 | 6.5 | 8.21 | 1.104 | 1.605 | 0.688 | 0.14 | -0.14 | 0.68 | 0.22666667 | -0.41 | -0.54 | 0.77435762 | 0.174749947 | 0.061730794 | 0.999652205 | 0.875938481 | 0.937795343 | 0.476099813 |
| Gngt1 | High | Q61012 | Guanine nucleotide-binding protein G(T) subunit gamma-T1 OS=Mus musculus OX=10090 GN=Gr | 0 | 84.736 | 66 | 12 | 284 | 10 | 74 | 8.5 | 4.82 | 1.221 | 1.6 | 0.763 | 0.29 | -0.29 | 0.68 | 0.22666667 | -0.485 | -0.39 | 0.499533216 | 0.17532604 | 0.100872369 | 0.999652205 | 0.875938481 | 0.937795343 | 0.610114517 |
| Praf2 | High | Q9JIG8 | PRA1 family protein 2 OS=Mus musculus OX=10090 GN=Praf2 PE=1 SV=1 | 0 | 4.999 | 10 | 1 | 2 | 1 | 178 | 19.5 | 9.6 | 1.054 | 1.58 | 0.667 | 0.08 | -0.08 | 0.66 | 0.22 | -0.37 | -0.58 | 0.913653301 | 0.176817078 | 0.029760117 | 0.999652205 | 0.875953959 | 0.937803082 | 0.328474358 |
| Ptbp2 | Medium | Q91Z31 | Polypyrimidine tract-binding protein 2 OS=Mus musculus OX=10090 GN=Ptbp2 PE=1 SV=2 | 0.042 | 1.641 | 1 | 1 | 1 | 1 | 531 | 57.5 | 8.66 | 1.181 | 1.585 | 0.745 | 0.24 | -0.24 | 0.66 | 0.22 | -0.45 | -0.42 | 0.567865637 | 0.177881246 | 0.068510456 | 0.999652205 | 0.875953959 | 0.937803082 | 0.499920511 |
| Akr1a1 | High | Q9JII6 | Aldo-keto reductase family 1 member A1 OS=Mus musculus OX=10090 GN=Akr1a1 PE=1 SV=3 | 0 | 50.183 | 38 | 10 | 17 | 10 | 325 | 36.6 | 7.39 | 0.903 | 0.663 | 1.361 | -0.15 | 0.15 | -0.59 | -0.1966667 | 0.37 | 0.44 | 0.733818399 | 0.178810646 | 0.036862962 | 0.999652205 | 0.877925824 | 0.938789015 | 0.373579725 |
| Pgrmc2 | High | Q80UU9 | Membrane-associated progesterone receptor component 2 OS=Mus musculus OX=10090 GN=Pgr | 0 | 13.703 | 19 | 3 | 3 | 2 | 217 | 23.3 | 5.15 | 1.119 | 1.594 | 0.702 | 0.16 | -0.16 | 0.67 | 0.22333333 | -0.415 | -0.51 | 0.739251858 | 0.179017156 | 0.040412999 | 0.999652205 | 0.878170771 | 0.938911488 | 0.388006308 |
| Tmem98 | High | Q91X86 | Transmembrane protein 98 OS=Mus musculus OX=10090 GN=Tmem98 PE=1 SV=1 | 0.001 | 5.04 | 5 | 1 | 1 | 1 | 226 | 24.7 | 4.94 | 1.17 | 1.572 | 0.744 | 0.23 | -0.23 | 0.65 | 0.21666667 | -0.44 | -0.43 | 0.567894144 | 0.1819348 | 0.089821754 | 0.999652205 | 0.890924388 | 0.945288296 | 0.5746627 |
| Hspa5 | High | P20029 | Endoplasmic reticulum chaperone BiP OS=Mus musculus OX=10090 GN=Hspa5 PE=1 SV=3 | 0 | 201.897 | 57 | 36 | 87 | 33 | 655 | 72.4 | 5.16 | 1.075 | 1.587 | 0.677 | 0.1 | -0.1 | 0.67 | 0.22333333 | -0.385 | -0.56 | 0.805589388 | 0.183508901 | 0.015976274 | 0.999652205 | 0.895493677 | 0.947572941 | 0.216374326 |
| Caprin1 | High | Q60865 | Caprin-1 OS=Mus musculus OX=10090 GN=Caprin1 PE=1 SV=2 | 0 | 30.211 | 9 | 5 | 8 | 5 | 707 | 78.1 | 5.25 | 1.14 | 1.579 | 0.722 | 0.19 | -0.19 | 0.66 | 0.22 | -0.425 | -0.47 | 0.661723239 | 0.188187423 | 0.04156759 | 0.999652205 | 0.902623507 | 0.951137856 | 0.396376666 |
| | High | A0A668KL | Serine/threonine-protein phosphatase 2A 55 kDa regulatory subunit B OS=Mus musculus OX=100 | 0.001 | 6.053 | 4 | 2 | 2 | 1 | 501 | 58.2 | 6.7 | 0.998 | 0.682 | 1.464 | 0 | 0 | -0.55 | -0.1833333 | 0.275 | 0.55 | 0.926271432 | 0.189074328 | 0.036432759 | 0.999652205 | 0.905328573 | 0.952490389 | 0.370300617 |
| Eif4a1 | High | P60843 | Eukaryotic initiation factor 4A-I OS=Mus musculus OX=10090 GN=Eif4a1 PE=1 SV=1 | 0 | 52.012 | 28 | 10 | 15 | 3 | 406 | 46.1 | 5.48 | 0.901 | 0.671 | 1.342 | -0.15 | 0.15 | -0.57 | -0.19 | 0.36 | 0.42 | 0.697154451 | 0.190896138 | 0.065913927 | 0.999652205 | 0.905821916 | 0.952737061 | 0.495414731 |
| Rps6ka3 | High | B1AXN9 | Non-specific serine/threonine protein kinase OS=Mus musculus OX=10090 GN=Rps6ka3 PE=1 SV= | 0.001 | 5.13 | 3 | 2 | 2 | 2 | 712 | 80.6 | 7.15 | 0.908 | 0.684 | 1.328 | -0.14 | 0.14 | -0.55 | -0.1833333 | 0.345 | 0.41 | 0.654767479 | 0.191115985 | 0.093508438 | 0.999652205 | 0.905821916 | 0.952737061 | 0.582204115 |
| Psmc3 | High | O88685 | 26S proteasome regulatory subunit 6A OS=Mus musculus OX=10090 GN=Psmc3 PE=1 SV=2 | 0 | 14.404 | 11 | 5 | 5 | 5 | 442 | 49.5 | 5.19 | 0.938 | 0.671 | 1.397 | -0.09 | 0.09 | -0.58 | -0.1933333 | 0.335 | 0.48 | 0.761278975 | 0.190747971 | 0.056229755 | 0.999652205 | 0.905821916 | 0.952737061 | 0.454949834 |
| Dak; Tkfc | High | Q8VC30 | Triokinase/FMN cyclase OS=Mus musculus OX=10090 GN=Tkfc PE=1 SV=1 | 0 | 13.293 | 8 | 3 | 3 | 3 | 578 | 59.7 | 6.92 | 0.87 | 0.686 | 1.269 | -0.2 | 0.2 | -0.54 | -0.18 | 0.37 | 0.34 | 0.542992057 | 0.192340016 | 0.133475937 | 0.999652205 | 0.90778659 | 0.953719397 | 0.695538641 |
| Psma6 | High | Q9QUM9 | Proteasome subunit alpha type-6 OS=Mus musculus OX=10090 GN=Psma6 PE=1 SV=1 | 0 | 31.638 | 33 | 8 | 11 | 8 | 246 | 27.4 | 6.76 | 0.886 | 0.683 | 1.297 | -0.17 | 0.17 | -0.55 | -0.1833333 | 0.36 | 0.37 | 0.735288731 | 0.209825614 | 0.089406396 | 0.999652205 | 0.912882441 | 0.956267323 | 0.5746627 |
| Psmb3 | High | Q9R1P1 | Proteasome subunit beta type-3 OS=Mus musculus OX=10090 GN=Psmb3 PE=1 SV=1 | 0 | 33.72 | 36 | 5 | 7 | 5 | 205 | 22.9 | 6.55 | 0.895 | 0.682 | 1.312 | -0.16 | 0.16 | -0.55 | -0.1833333 | 0.355 | 0.39 | 0.636860764 | 0.207903432 | 0.117243709 | 0.999652205 | 0.912882441 | 0.956267323 | 0.654114904 |
| Cd99l2 | High | A2AP74 | CD99 antigen-like protein 2 OS=Mus musculus OX=10090 GN=Cd99l2 PE=1 SV=1 | 0.001 | 5.485 | 8 | 1 | 1 | 1 | 216 | 23.2 | 4.93 | 0.899 | 0.685 | 1.313 | -0.15 | 0.15 | -0.55 | -0.1833333 | 0.35 | 0.39 | 0.639803026 | 0.198781366 | 0.136940769 | 0.999652205 | 0.912882441 | 0.956267323 | 0.700189449 |
| Nefl | High | P08551 | Neurofilament light polypeptide OS=Mus musculus OX=10090 GN=Nefl PE=1 SV=5 | 0 | 20.353 | 10 | 6 | 8 | 5 | 543 | 61.5 | 4.64 | 0.96 | 0.679 | 1.413 | -0.06 | 0.06 | -0.56 | -0.1866667 | 0.31 | 0.5 | 0.897482299 | 0.202914512 | 0.046836855 | 0.999652205 | 0.912882441 | 0.956267323 | 0.410976909 |
| Psma4 | High | Q9R1P0 | Proteasome subunit alpha type-4 OS=Mus musculus OX=10090 GN=Psma4 PE=1 SV=1 | 0 | 32.918 | 33 | 8 | 10 | 8 | 261 | 29.5 | 7.72 | 0.992 | 0.677 | 1.465 | -0.01 | 0.01 | -0.56 | -0.1866667 | 0.285 | 0.55 | 0.948577276 | 0.19980934 | 0.015627452 | 0.999652205 | 0.912882441 | 0.956267323 | 0.212677478 |
| Eif2d | High | E9PUG7 | Eukaryotic translation initiation factor 2D OS=Mus musculus OX=10090 GN=Eif2d PE=1 SV=1 | 0.001 | 6.203 | 4 | 2 | 2 | 2 | 552 | 60.8 | 7.75 | 1.053 | 0.677 | 1.555 | 0.07 | -0.07 | -0.56 | -0.1866667 | 0.245 | 0.64 | 0.873945144 | 0.199774086 | 0.01364138 | 0.999652205 | 0.912882441 | 0.956267323 | 0.191288138 |
| | Medium | A0A0A6YX | Serine/threonine-protein kinase DCLK2 OS=Mus musculus OX=10090 GN=Dclk2 PE=1 SV=1 | 0.049 | 1.507 | 1 | 1 | 1 | 1 | 711 | 78 | 8.79 | 1.051 | 1.533 | 0.686 | 0.07 | -0.07 | 0.62 | 0.20666667 | -0.345 | -0.54 | 0.903372371 | 0.207909385 | 0.03715943 | 0.999652205 | 0.912882441 | 0.956267323 | 0.373579725 |
| Nipsnap1 | High | O55125 | Protein NipSnap homolog 1 OS=Mus musculus OX=10090 GN=Nipsnap1 PE=1 SV=1 | 0 | 11.711 | 8 | 2 | 3 | 2 | 284 | 33.3 | 9.44 | 1.143 | 1.551 | 0.737 | 0.19 | -0.19 | 0.63 | 0.21 | -0.41 | -0.44 | 0.584520535 | 0.207634418 | 0.08234316 | 0.999652205 | 0.912882441 | 0.956267323 | 0.552270451 |
| Avl9 | Medium | Q80U56 | Late secretory pathway protein AVL9 homolog OS=Mus musculus OX=10090 GN=Avl9 PE=1 SV=2 | 0.019 | 2.066 | 1 | 1 | 1 | 1 | 649 | 72.1 | 6.14 | 1.169 | 1.557 | 0.751 | 0.23 | -0.23 | 0.64 | 0.21333333 | -0.435 | -0.41 | 0.619538879 | 0.203003584 | 0.084475343 | 0.999652205 | 0.912882441 | 0.956267323 | 0.560726223 |
| Acadl | High | A0A0R4J0 | Long-chain specific acyl-CoA dehydrogenase, mitochondrial OS=Mus musculus OX=10090 GN=Aca | 0 | 9.8 | 10 | 4 | 4 | 4 | 430 | 47.9 | 8.15 | 1.188 | 1.557 | 0.763 | 0.25 | -0.25 | 0.64 | 0.21333333 | -0.445 | -0.39 | 0.483327953 | 0.203533022 | 0.097204758 | 0.999652205 | 0.912882441 | 0.956267323 | 0.593711416 |
| Alpl | High | P09242 | Alkaline phosphatase, tissue-nonspecific isozyme OS=Mus musculus OX=10090 GN=Alpl PE=1 SV= | 0 | 13.864 | 6 | 3 | 4 | 3 | 524 | 57.5 | 7.01 | 1.24 | 1.548 | 0.801 | 0.31 | -0.31 | 0.63 | 0.21 | -0.47 | -0.32 | 0.439169795 | 0.208893997 | 0.187248909 | 0.999652205 | 0.912882441 | 0.956267323 | 0.774149321 |
| Ermn | High | Q5EBJ4 | Ermin OS=Mus musculus OX=10090 GN=Ermn PE=1 SV=1 | 0 | 141.755 | 65 | 18 | 67 | 18 | 281 | 32.1 | 4.59 | 1.248 | 1.552 | 0.804 | 0.32 | -0.32 | 0.63 | 0.21 | -0.475 | -0.31 | 0.45464134 | 0.207185451 | 0.192087243 | 0.999652205 | 0.912882441 | 0.956267323 | 0.784495541 |
| Tmem11 | High | E9Q933 | Transmembrane protein 11, mitochondrial OS=Mus musculus OX=10090 GN=Tmem11 PE=1 SV=1 | 0.005 | 3.745 | 9 | 1 | 1 | 1 | 176 | 19.8 | 6.25 | 1.038 | 1.548 | 0.671 | 0.05 | -0.05 | 0.63 | 0.21 | -0.34 | -0.58 | 0.950943086 | 0.210586275 | 0.044679859 | 0.999652205 | 0.913066452 | 0.956359328 | 0.405968973 |
| Bcap31 | High | Q61335 | B-cell receptor-associated protein 31 OS=Mus musculus OX=10090 GN=Bcap31 PE=1 SV=4 | 0 | 20.795 | 18 | 5 | 9 | 5 | 245 | 27.9 | 8.7 | 1.063 | 1.544 | 0.688 | 0.09 | -0.09 | 0.63 | 0.21 | -0.36 | -0.54 | 0.816200419 | 0.212265842 | 0.020886466 | 0.999652205 | 0.913066452 | 0.956359328 | 0.259764608 |
| Ptprj | High | E9Q4S7 | Protein-tyrosine-phosphatase OS=Mus musculus OX=10090 GN=Ptprj PE=1 SV=2 | 0.001 | 7.128 | 2 | 2 | 2 | 2 | 1350 | 148.5 | 7.17 | 1.026 | 0.699 | 1.468 | 0.04 | -0.04 | -0.52 | -0.1733333 | 0.24 | 0.55 | 0.999003407 | 0.216626583 | 0.012432095 | 0.999652205 | 0.920170644 | 0.959911425 | 0.183922842 |
| Lrpap1 | High | P55302 | Alpha-2-macroglobulin receptor-associated protein OS=Mus musculus OX=10090 GN=Lrpap1 PE= | 0 | 14.036 | 20 | 7 | 8 | 7 | 360 | 42.2 | 7.87 | 1.169 | 1.537 | 0.761 | 0.23 | -0.23 | 0.62 | 0.20666667 | -0.425 | -0.39 | 0.614326729 | 0.217704743 | 0.096728342 | 0.999652205 | 0.922239631 | 0.960945918 | 0.593387105 |
| Agk | High | Q9ESW4 | Acylglycerol kinase, mitochondrial OS=Mus musculus OX=10090 GN=Agk PE=1 SV=1 | 0.001 | 6.323 | 5 | 2 | 2 | 2 | 421 | 46.9 | 8.4 | 1.251 | 1.536 | 0.814 | 0.32 | -0.32 | 0.62 | 0.20666667 | -0.47 | -0.3 | 0.444980336 | 0.218589795 | 0.252213173 | 0.999652205 | 0.923924555 | 0.96178838 | 0.851275262 |
| Hsd17b12 | High | O70503 | Very-long-chain 3-oxoacyl-CoA reductase OS=Mus musculus OX=10090 GN=Hsd17b12 PE=1 SV=1 | 0 | 31.369 | 22 | 6 | 8 | 6 | 312 | 34.7 | 9.52 | 1.02 | 1.534 | 0.665 | 0.03 | -0.03 | 0.62 | 0.20666667 | -0.325 | -0.59 | 0.978853239 | 0.219771214 | 0.013001626 | 0.999652205 | 0.92720632 | 0.963429262 | 0.188860411 |
| | High | E9Q5L3 | Short/branched chain-specific acyl-CoA dehydrogenase, mitochondrial OS=Mus musculus OX=100 | 0.001 | 7.774 | 8 | 3 | 3 | 3 | 465 | 51.8 | 8.32 | 0.993 | 1.528 | 0.65 | -0.01 | 0.01 | 0.61 | 0.20333333 | -0.3 | -0.62 | 0.977567796 | 0.220912473 | 0.015473623 | 0.999652205 | 0.928185815 | 0.96391901 | 0.211096356 |
| Ppp5c | High | Q60676 | Serine/threonine-protein phosphatase 5 OS=Mus musculus OX=10090 GN=Ppp5c PE=1 SV=3 | 0 | 23.929 | 13 | 6 | 8 | 6 | 499 | 56.8 | 6.2 | 0.858 | 0.693 | 1.239 | -0.22 | 0.22 | -0.53 | -0.1766667 | 0.375 | 0.31 | 0.530059966 | 0.225313232 | 0.227125577 | 0.999652205 | 0.935008675 | 0.96733044 | 0.816808422 |
| Psma7 | High | Q9Z2U0 | Proteasome subunit alpha type-7 OS=Mus musculus OX=10090 GN=Psma7 PE=1 SV=1 | 0 | 54.181 | 33 | 8 | 17 | 8 | 248 | 27.8 | 8.46 | 0.914 | 0.692 | 1.32 | -0.13 | 0.13 | -0.53 | -0.1766667 | 0.33 | 0.4 | 0.784943728 | 0.224218772 | 0.058570506 | 0.999652205 | 0.935008675 | 0.96733044 | 0.4662587 |
| Hsd17b4 | High | P51660 | Peroxisomal multifunctional enzyme type 2 OS=Mus musculus OX=10090 GN=Hsd17b4 PE=1 SV= | 0 | 32.523 | 17 | 9 | 10 | 9 | 735 | 79.4 | 8.57 | 1.037 | 1.529 | 0.678 | 0.05 | -0.05 | 0.61 | 0.20333333 | -0.33 | -0.56 | 0.982244208 | 0.224066795 | 0.019677304 | 0.999652205 | 0.935008675 | 0.96733044 | 0.250182865 |
| Card14 | Low | Q99KF0 | Caspase recruitment domain-containing protein 14 OS=Mus musculus OX=10090 GN=Card14 PE= | 0.053 | 1.419 | 1 | 1 | 1 | 1 | 999 | 113.4 | 6.1 | 1.14 | 1.519 | 0.751 | 0.19 | -0.19 | 0.6 | 0.2 | -0.395 | -0.41 | 0.66497773 | 0.227057793 | 0.109691028 | 0.999652205 | 0.935008675 | 0.96733044 | 0.632515291 |
| Arfgap1 | High | Q3TGS9 | ADP-ribosylation factor GTPase-activating protein 1 OS=Mus musculus OX=10090 GN=Arfgap1 PE | 0 | 45.86 | 25 | 8 | 13 | 8 | 394 | 43.2 | 5.54 | 1.155 | 1.524 | 0.757 | 0.21 | -0.21 | 0.61 | 0.20333333 | -0.41 | -0.4 | 0.659474964 | 0.227266874 | 0.081975359 | 0.999652205 | 0.935008675 | 0.96733044 | 0.55046208 |
| | Medium | H3BL09 | Protein CASP OS=Mus musculus OX=10090 GN=Cux1 PE=1 SV=2 | 0.021 | 2.011 | 4 | 1 | 1 | 1 | 199 | 23.3 | 5.44 | 1.182 | 1.515 | 0.78 | 0.24 | -0.24 | 0.6 | 0.2 | -0.42 | -0.36 | 0.552821016 | 0.228888381 | 0.16267572 | 0.999652205 | 0.935008675 | 0.96733044 | 0.733684585 |
| Scrn3 | High | Q3TMH2 | Secernin-3 OS=Mus musculus OX=10090 GN=Scrn3 PE=1 SV=1 | 0 | 9.575 | 5 | 2 | 2 | 2 | 418 | 47.6 | 5.66 | 0.937 | 0.696 | 1.347 | -0.09 | 0.09 | -0.52 | -0.1733333 | 0.305 | 0.43 | 0.849438286 | 0.230484295 | 0.088738304 | 0.999652205 | 0.936284541 | 0.967968373 | 0.574544652 |
| Gnb1 | High | P62874 | Guanine nucleotide-binding protein G(I)/G(S)/G(T) subunit beta-1 OS=Mus musculus OX=10090 G | 0 | 597.061 | 76 | 23 | 1464 | 12 | 340 | 37.4 | 6 | 1.252 | 1.521 | 0.823 | 0.32 | -0.32 | 0.6 | 0.2 | -0.46 | -0.28 | 0.446990227 | 0.230093022 | 0.249574984 | 0.999652205 | 0.936284541 | 0.967968373 | 0.848101174 |
| Snx12 | High | Q3TGS7 | Sorting nexin-12 OS=Mus musculus OX=10090 GN=Snx12 PE=1 SV=1 | 0 | 14.376 | 31 | 6 | 7 | 5 | 166 | 19.2 | 7.25 | 0.987 | 0.697 | 1.416 | -0.02 | 0.02 | -0.52 | -0.1733333 | 0.27 | 0.5 | 0.91978493 | 0.232842332 | 0.038158639 | 0.999652205 | 0.936813316 | 0.96823276 | 0.373579725 |
| Slc39a7 | Medium | Q31125 | Zinc transporter SLC39A7 OS=Mus musculus OX=10090 GN=Slc39a7 PE=1 SV=2 | 0.037 | 1.729 | 2 | 1 | 1 | 1 | 476 | 50.6 | 6.87 | 1.055 | 1.498 | 0.704 | 0.08 | -0.08 | 0.58 | 0.19333333 | -0.33 | -0.51 | 0.877161916 | 0.232952072 | 0.058313013 | 0.999652205 | 0.936813316 | 0.96823276 | 0.4657565 |
| Cisd3 | High | Z4YKM2 | CDGSH iron-sulfur domain-containing protein 3, mitochondrial OS=Mus musculus OX=10090 GN= | 0.005 | 3.822 | 12 | 2 | 2 | 2 | 115 | 13.1 | 9.83 | 1.182 | 1.511 | 0.783 | 0.24 | -0.24 | 0.6 | 0.2 | -0.42 | -0.35 | 0.577021236 | 0.232098272 | 0.131601393 | 0.999652205 | 0.936813316 | 0.96823276 | 0.693504708 |
| Mlec | High | Q6ZQI3 | Malectin OS=Mus musculus OX=10090 GN=Mlec PE=1 SV=2 | 0.001 | 5.142 | 6 | 2 | 2 | 2 | 291 | 32.3 | 6.05 | 1.193 | 1.518 | 0.786 | 0.25 | -0.25 | 0.6 | 0.2 | -0.425 | -0.35 | 0.542021667 | 0.232213671 | 0.187526393 | 0.999652205 | 0.936813316 | 0.96823276 | 0.774149321 |
| Hsp90b1 | High | P08113 | Endoplasmin OS=Mus musculus OX=10090 GN=Hsp90b1 PE=1 SV=2 | 0 | 78.627 | 23 | 20 | 32 | 18 | 802 | 92.4 | 4.82 | 1.121 | 1.511 | 0.742 | 0.16 | -0.16 | 0.6 | 0.2 | -0.38 | -0.43 | 0.69824068 | 0.23806348 | 0.068346089 | 0.999652205 | 0.937374952 | 0.968513578 | 0.499920511 |
| Prrt3 | Low | A0A0N4S | Proline-rich transmembrane protein 3 OS=Mus musculus OX=10090 GN=Prrt3 PE=1 SV=1 | 0.072 | 1.27 | 1 | 1 | 1 | 1 | 972 | 101.2 | 7.52 | 1.147 | 1.49 | 0.77 | 0.2 | -0.2 | 0.57 | 0.19 | -0.385 | -0.38 | 0.658157632 | 0.239636672 | 0.147877915 | 0.999652205 | 0.937938091 | 0.968795148 | 0.712464409 |
| Psmb4 | High | P99026 | Proteasome subunit beta type-4 OS=Mus musculus OX=10090 GN=Psmb4 PE=1 SV=1 | 0 | 23.728 | 22 | 4 | 7 | 4 | 264 | 29.1 | 5.64 | 0.959 | 0.701 | 1.367 | -0.06 | 0.06 | -0.51 | -0.17 | 0.285 | 0.45 | 0.843743823 | 0.240463631 | 0.05366612 | 0.999652205 | 0.938986585 | 0.969319395 | 0.443814062 |
| | High | A0A087W | 6-phosphofructo-2-kinase OS=Mus musculus OX=10090 GN=Pfkfb2 PE=1 SV=1 | 0 | 39.492 | 26 | 12 | 14 | 12 | 533 | 61.5 | 7.83 | 0.825 | 0.703 | 1.173 | -0.28 | 0.28 | -0.51 | -0.17 | 0.395 | 0.23 | 0.512737487 | 0.243922364 | 0.278674945 | 0.999652205 | 0.940627712 | 0.970139959 | 0.876067403 |
| Ina | High | P46660 | Alpha-internexin OS=Mus musculus OX=10090 GN=Ina PE=1 SV=3 | 0 | 11.711 | 6 | 3 | 5 | 2 | 501 | 55.3 | 5.4 | 1.08 | 0.716 | 1.508 | 0.11 | -0.11 | -0.48 | -0.16 | 0.185 | 0.59 | 0.816605419 | 0.243786597 | 0.012908359 | 0.999652205 | 0.940627712 | 0.970139959 | 0.18799264 |
| Erp44 | Medium | Q9D1Q6 | Endoplasmic reticulum resident protein 44 OS=Mus musculus OX=10090 GN=Erp44 PE=1 SV=1 | 0.01 | 2.685 | 3 | 1 | 1 | 1 | 406 | 46.8 | 5.27 | 1.115 | 1.491 | 0.747 | 0.16 | -0.16 | 0.58 | 0.19333333 | -0.37 | -0.42 | 0.703787862 | 0.241901223 | 0.097079886 | 0.999652205 | 0.940627712 | 0.970139959 | 0.593711416 |
| Hnrnpd | High | Q60668 | Heterogeneous nuclear ribonucleoprotein D0 OS=Mus musculus OX=10090 GN=Hnrnpd PE=1 SV=2 | 0 | 8.471 | 9 | 3 | 4 | 3 | 355 | 38.3 | 7.81 | 1.13 | 1.505 | 0.75 | 0.18 | -0.18 | 0.59 | 0.19666667 | -0.385 | -0.41 | 0.698680662 | 0.242555672 | 0.088391654 | 0.999652205 | 0.940627712 | 0.970139959 | 0.573625007 |
| Cisd1 | High | Q91WS0 | CDGSH iron-sulfur domain-containing protein 1 OS=Mus musculus OX=10090 GN=Cisd1 PE=1 SV= | 0 | 28.739 | 44 | 5 | 12 | 5 | 108 | 12.1 | 9.06 | 1.224 | 1.504 | 0.814 | 0.29 | -0.29 | 0.59 | 0.19666667 | -0.44 | -0.3 | 0.488457612 | 0.243732112 | 0.219771914 | 0.999652205 | 0.940627712 | 0.970139959 | 0.805859554 |
| Fgf1 | Medium | P61148 | Fibroblast growth factor 1 OS=Mus musculus OX=10090 GN=Fgf1 PE=1 SV=1 | 0.035 | 1.789 | 8 | 1 | 1 | 1 | 155 | 17.4 | 7.02 | 1.254 | 1.49 | 0.841 | 0.33 | -0.33 | 0.58 | 0.19333333 | -0.455 | -0.25 | 0.428575702 | 0.241585357 | 0.327402856 | 0.999652205 | 0.940627712 | 0.970139959 | 0.905686588 |
| Ykt6 | High | Q9CQW1 | Synaptobrevin homolog YKT6 OS=Mus musculus OX=10090 GN=Ykt6 PE=1 SV=1 | 0 | 19.915 | 27 | 6 | 10 | 6 | 198 | 22.3 | 6.35 | 0.881 | 0.704 | 1.251 | -0.18 | 0.18 | -0.51 | -0.17 | 0.345 | 0.32 | 0.595824072 | 0.244633498 | 0.184478084 | 0.999652205 | 0.941474619 | 0.970563412 | 0.768475941 |
| Pter | Medium | A2AUR5 | Parathion hydrolase-related protein (Fragment) OS=Mus musculus OX=10090 GN=Pter PE=1 SV=1 | 0.037 | 1.727 | 3 | 1 | 1 | 1 | 253 | 28.6 | 6.29 | 0.839 | 0.715 | 1.173 | -0.25 | 0.25 | -0.48 | -0.16 | 0.365 | 0.23 | 0.454947738 | 0.253637817 | 0.336475166 | 0.999652205 | 0.950034749 | 0.974843477 | 0.906329766 |
| Snx17 | High | Q8BVL3 | Sorting nexin-17 OS=Mus musculus OX=10090 GN=Snx17 PE=1 SV=2 | 0.001 | 5.032 | 2 | 2 | 2 | 2 | 470 | 52.8 | 7.49 | 0.863 | 0.712 | 1.213 | -0.21 | 0.21 | -0.49 | -0.1633333 | 0.35 | 0.28 | 0.581682405 | 0.251819614 | 0.269519364 | 0.999652205 | 0.950034749 | 0.974843477 | 0.871508117 |
| Cd59a | High | O55186 | CD59A glycoprotein OS=Mus musculus OX=10090 GN=Cd59a PE=2 SV=1 | 0 | 28.585 | 23 | 3 | 14 | 3 | 123 | 13.6 | 7.47 | 0.927 | 0.707 | 1.312 | -0.11 | 0.11 | -0.5 | -0.1666667 | 0.305 | 0.39 | 0.763357535 | 0.250433113 | 0.090892327 | 0.999652205 | 0.950034749 | 0.974843477 | 0.576060131 |
| Ikbkap | High | Q7TT37 | Elongator complex protein 1 OS=Mus musculus OX=10090 GN=Elp1 PE=1 SV=2 | 0 | 11.533 | 3 | 4 | 4 | 4 | 1333 | 149.5 | 6 | 0.926 | 0.715 | 1.294 | -0.11 | 0.11 | -0.48 | -0.16 | 0.295 | 0.37 | 0.685036031 | 0.250771902 | 0.091562104 | 0.999652205 | 0.950034749 | 0.974843477 | 0.578792238 |
| Gga3 | High | S4R2D2 | ADP-ribosylation factor-binding protein GGA3 OS=Mus musculus OX=10090 GN=Gga3 PE=1 SV=1 | 0.001 | 5.02 | 5 | 3 | 3 | 2 | 595 | 64.5 | 5.47 | 0.954 | 0.717 | 1.33 | -0.07 | 0.07 | -0.48 | -0.16 | 0.275 | 0.41 | 0.782990868 | 0.252951597 | 0.109987945 | 0.999652205 | 0.950034749 | 0.974843477 | 0.632515291 |
| Cyp4f13 | Low | B8JK07 | Cytochrome P450, family 4, subfamily f, polypeptide 13 OS=Mus musculus OX=10090 GN=Cyp4f13 | 0.06 | 1.329 | 6 | 1 | 2 | 1 | 414 | 47.3 | 7.23 | 0.99 | 0.716 | 1.382 | -0.01 | 0.01 | -0.48 | -0.16 | 0.245 | 0.47 | 0.923471867 | 0.252317187 | 0.099807058 | 0.999652205 | 0.950034749 | 0.974843477 | 0.606961141 |
| Hist1h2bb | High | Q64475 | Histone H2B type 1-B OS=Mus musculus OX=10090 GN=H2bc3 PE=1 SV=3 | 0 | 25.221 | 52 | 7 | 9 | 7 | 126 | 13.9 | 10.32 | 1.138 | 1.497 | 0.76 | 0.19 | -0.19 | 0.58 | 0.19333333 | -0.385 | -0.4 | 0.667376024 | 0.249792155 | 0.092689604 | 0.999652205 | 0.950034749 | 0.974843477 | 0.581982763 |
| P4hb | High | P09103 | Protein disulfide-isomerase OS=Mus musculus OX=10090 GN=P4hb PE=1 SV=2 | 0 | 89.983 | 54 | 21 | 30 | 21 | 509 | 57 | 4.88 | 1.196 | 1.497 | 0.799 | 0.26 | -0.26 | 0.58 | 0.19333333 | -0.42 | -0.32 | 0.544188587 | 0.249063063 | 0.178151659 | 0.999652205 | 0.950034749 | 0.974843477 | 0.754689318 |
| Ccdc45; Ce | High | A2A5Z4 | Centrosomal protein of 95 kDa (Fragment) OS=Mus musculus OX=10090 GN=Cep95 PE=1 SV=1 | 0.009 | 2.81 | 7 | 1 | 1 | 1 | 121 | 14.8 | 9.55 | 1.233 | 1.48 | 0.833 | 0.3 | -0.3 | 0.57 | 0.19 | -0.435 | -0.26 | 0.441377503 | 0.250773076 | 0.318171188 | 0.999652205 | 0.950034749 | 0.974843477 | 0.904943958 |
| Cd9 | High | P40240 | CD9 antigen OS=Mus musculus OX=10090 GN=Cd9 PE=1 SV=2 | 0 | 19.325 | 21 | 4 | 6 | 4 | 226 | 25.2 | 7.23 | 0.811 | 0.715 | 1.134 | -0.3 | 0.3 | -0.48 | -0.16 | 0.39 | 0.18 | 0.486770156 | 0.264363032 | 0.398144959 | 0.999652205 | 0.951072422 | 0.975362314 | 0.93262326 |
| Vps36 | High | Q91XD6 | Vacuolar protein-sorting-associated protein 36 OS=Mus musculus OX=10090 GN=Vps36 PE=1 SV= | 0 | 26.807 | 24 | 8 | 9 | 8 | 386 | 43.7 | 7.15 | 0.831 | 0.71 | 1.17 | -0.27 | 0.27 | -0.49 | -0.1633333 | 0.38 | 0.23 | 0.429492194 | 0.255821133 | 0.352502053 | 0.999652205 | 0.951072422 | 0.975362314 | 0.910898173 |
| | High | A0A1B0GF | Transforming acidic coiled-coil-containing protein 1 (Fragment) OS=Mus musculus OX=10090 GN | 0.001 | 4.381 | 7 | 1 | 1 | 1 | 146 | 15.5 | 4.81 | 0.825 | 0.727 | 1.135 | -0.28 | 0.28 | -0.46 | -0.1533333 | 0.37 | 0.18 | 0.440577489 | 0.266617736 | 0.4758121 | 0.999652205 | 0.951072422 | 0.975362314 | 0.962227984 |
| Zfyve20; R | Medium | Q80Y56 | Rabenosyn-5 OS=Mus musculus OX=10090 GN=Rbsn PE=1 SV=1 | 0.037 | 1.727 | 1 | 1 | 1 | 1 | 783 | 88.4 | 5.3 | 0.849 | 0.725 | 1.172 | -0.24 | 0.24 | -0.46 | -0.1533333 | 0.35 | 0.23 | 0.514346164 | 0.264704836 | 0.392261039 | 0.999652205 | 0.951072422 | 0.975362314 | 0.93064581 |
| Dnal1; Dna | Medium | Q05A62 | Dynein axonemal light chain 1 OS=Mus musculus OX=10090 GN=Dnal1 PE=1 SV=2 | 0.019 | 2.038 | 5 | 1 | 1 | 1 | 190 | 21.5 | 5.76 | 0.869 | 0.718 | 1.21 | -0.2 | 0.2 | -0.48 | -0.16 | 0.34 | 0.27 | 0.541110581 | 0.26484499 | 0.271180227 | 0.999652205 | 0.951072422 | 0.975362314 | 0.871850649 |
| Hspa2 | High | P17156 | Heat shock-related 70 kDa protein 2 OS=Mus musculus OX=10090 GN=Hspa2 PE=1 SV=2 | 0 | 155.44 | 29 | 23 | 99 | 5 | 633 | 69.6 | 5.67 | 0.881 | 0.71 | 1.239 | -0.18 | 0.18 | -0.49 | -0.1633333 | 0.335 | 0.31 | 0.586436732 | 0.256584039 | 0.216777244 | 0.999652205 | 0.951072422 | 0.975362314 | 0.804947026 |
| Eno1; Gm! | High | P17182 | Alpha-enolase OS=Mus musculus OX=10090 GN=Eno1 PE=1 SV=3 | 0 | 508.152 | 81 | 35 | 302 | 28 | 434 | 47.1 | 6.8 | 0.894 | 0.714 | 1.252 | -0.16 | 0.16 | -0.49 | -0.1633333 | 0.325 | 0.32 | 0.707839237 | 0.262455062 | 0.121017144 | 0.999652205 | 0.951072422 | 0.975362314 | 0.669174681 |

| | | | | | | | | | | | | | | | | | | | | | | | | | | | | |
|---|---|---|---|---|---|---|---|---|---|---|---|---|---|---|---|---|---|---|---|---|---|---|---|---|---|---|---|---|
| Nampt | High | Q99KQ4 | Nicotinamide phosphoribosyltransferase OS=Mus musculus OX=10090 GN=Nampt PE=1 SV=1 | 0 | 15.865 | 10 | 5 | 5 | 5 | 491 | 55.4 | 7.15 | 0.921 | 0.711 | 1.295 | -0.12 | 0.12 | -0.49 | -0.1633333 | 0.305 | 0.37 | 0.710675666 | 0.258112833 | 0.142823996 | 0.999652205 | 0.951072422 | 0.975362314 | 0.703085292 |
| | High | A0A5F8MI | Kinesin-like protein KIF1B OS=Mus musculus OX=10090 GN=Kif1b PE=1 SV=1 | 0 | 24.19 | 4 | 9 | 10 | 4 | 1883 | 211.5 | 5.64 | 0.92 | 0.715 | 1.286 | -0.12 | 0.12 | -0.48 | -0.16 | 0.3 | 0.36 | 0.717401658 | 0.265751996 | 0.133120366 | 0.999652205 | 0.951072422 | 0.975362314 | 0.695121227 |
| Abcf3 | High | Q8K268 | ATP-binding cassette sub-family F member 3 OS=Mus musculus OX=10090 GN=Abcf3 PE=1 SV=1 | 0.001 | 5.29 | 2 | 2 | 2 | 2 | 709 | 79.8 | 6.16 | 0.923 | 0.726 | 1.27 | -0.12 | 0.12 | -0.46 | -0.1533333 | 0.29 | 0.35 | 0.696626601 | 0.263499951 | 0.154317606 | 0.999652205 | 0.951072422 | 0.975362314 | 0.728999569 |
| Prkcz | High | Q02956 | Protein kinase C zeta type OS=Mus musculus OX=10090 GN=Prkcz PE=1 SV=2 | 0 | 8.403 | 4 | 2 | 2 | 2 | 592 | 67.6 | 5.76 | 1.075 | 0.719 | 1.496 | 0.1 | -0.1 | -0.48 | -0.16 | 0.19 | 0.58 | 0.845581465 | 0.257083782 | 0.043278084 | 0.999652205 | 0.951072422 | 0.975362314 | 0.404433694 |
| Arrb1 | High | J3QNU6 | Beta-arrestin-1 OS=Mus musculus OX=10090 GN=Arrb1 PE=1 SV=1 | 0 | 26.743 | 16 | 6 | 74 | 3 | 419 | 47.1 | 6.55 | 0.935 | 1.465 | 0.639 | -0.1 | 0.1 | 0.55 | 0.18333333 | -0.225 | -0.65 | 0.733873645 | 0.263420341 | 0.012243048 | 0.999652205 | 0.951072422 | 0.975362314 | 0.183058049 |
| Cisd2 | High | Q9CQB5 | CDGSH iron-sulfur domain-containing protein 2 OS=Mus musculus OX=10090 GN=Cisd2 PE=1 SV= | 0 | 8.977 | 27 | 4 | 4 | 4 | 135 | 15.2 | 9.51 | 1.066 | 1.488 | 0.717 | 0.09 | -0.09 | 0.57 | 0.19 | -0.33 | -0.48 | 0.788350215 | 0.25712593 | 0.046511322 | 0.999652205 | 0.951072422 | 0.975362314 | 0.409467755 |
| Vapb | High | Q9QY76 | Vesicle-associated membrane protein-associated protein B OS=Mus musculus OX=10090 GN=Vap | 0 | 25.232 | 26 | 5 | 9 | 4 | 243 | 26.9 | 7.78 | 1.069 | 1.477 | 0.724 | 0.1 | -0.1 | 0.56 | 0.18666667 | -0.33 | -0.47 | 0.886094386 | 0.266985552 | 0.042669225 | 0.999652205 | 0.951072422 | 0.975362314 | 0.401420039 |
| Dpm1 | High | O70152 | Dolichol-phosphate mannosyltransferase subunit 1 OS=Mus musculus OX=10090 GN=Dpm1 PE=1 | 0 | 8.027 | 12 | 3 | 4 | 3 | 260 | 29.2 | 9.51 | 1.076 | 1.481 | 0.726 | 0.11 | -0.11 | 0.57 | 0.19 | -0.34 | -0.46 | 0.867519671 | 0.262982988 | 0.061061883 | 0.999652205 | 0.951072422 | 0.975362314 | 0.476099813 |
| Gnb4 | High | P29387 | Guanine nucleotide-binding protein subunit beta-4 OS=Mus musculus OX=10090 GN=Gnb4 PE=1 S | 0 | 127.028 | 27 | 6 | 126 | 2 | 340 | 37.4 | 6.16 | 1.17 | 1.472 | 0.795 | 0.23 | -0.23 | 0.56 | 0.18666667 | -0.395 | -0.33 | 0.593440299 | 0.25822944 | 0.198966506 | 0.999652205 | 0.951072422 | 0.975362314 | 0.78697088 |
| | High | A0A0R3P9 | NADH dehydrogenase [ubiquinone] 1 alpha subcomplex subunit 9, mitochondrial OS=Mus muscul | 0 | 27.594 | 18 | 8 | 9 | 8 | 373 | 42.1 | 9.09 | 1.225 | 1.489 | 0.823 | 0.29 | -0.29 | 0.57 | 0.19 | -0.43 | -0.28 | 0.508102387 | 0.255923566 | 0.221223199 | 0.999652205 | 0.951072422 | 0.975362314 | 0.807447486 |
| Rer1 | High | Q9CQU3 | Protein RER1 OS=Mus musculus OX=10090 GN=Rer1 PE=1 SV=1 | 0.009 | 2.862 | 4 | 1 | 1 | 1 | 196 | 23 | 9.51 | 1.228 | 1.482 | 0.828 | 0.3 | -0.3 | 0.57 | 0.19 | -0.435 | -0.27 | 0.482824851 | 0.262555863 | 0.262792112 | 0.999652205 | 0.951072422 | 0.975362314 | 0.859470827 |
| Alg2 | High | Q9DBE8 | Alpha-1,3/1,6-mannosyltransferase ALG2 OS=Mus musculus OX=10090 GN=Alg2 PE=1 SV=2 | 0 | 15.829 | 12 | 4 | 5 | 4 | 415 | 47.4 | 7.97 | 1.085 | 1.474 | 0.736 | 0.12 | -0.12 | 0.56 | 0.18666667 | -0.34 | -0.44 | 0.841127997 | 0.269120038 | 0.054477698 | 0.999652205 | 0.953285167 | 0.976468686 | 0.44788336 |
| Asah1 | High | Q9WV54 | Acid ceramidase OS=Mus musculus OX=10090 GN=Asah1 PE=1 SV=1 | 0 | 20.308 | 15 | 6 | 6 | 6 | 394 | 44.6 | 8.46 | 1.133 | 1.475 | 0.768 | 0.18 | -0.18 | 0.56 | 0.18666667 | -0.37 | -0.38 | 0.667652781 | 0.268359825 | 0.109756499 | 0.999652205 | 0.953285167 | 0.976468686 | 0.632515291 |
| Pcyox1l | High | Q8C7K6 | Prenylcysteine oxidase-like OS=Mus musculus OX=10090 GN=Pcyox1l PE=1 SV=1 | 0 | 8.456 | 6 | 3 | 3 | 3 | 495 | 54.8 | 7.65 | 1.148 | 1.456 | 0.788 | 0.2 | -0.2 | 0.54 | 0.18 | -0.37 | -0.34 | 0.609441269 | 0.269337501 | 0.189488456 | 0.999652205 | 0.953285167 | 0.976468686 | 0.77990201 |
| | High | A0A2I3BR | Microsomal signal peptidase 12 kDa subunit OS=Mus musculus OX=10090 GN=Spcs1 PE=1 SV=1 | 0.006 | 3.206 | 13 | 1 | 1 | 1 | 102 | 11.8 | 9.31 | 1.049 | 1.463 | 0.717 | 0.07 | -0.07 | 0.55 | 0.18333333 | -0.31 | -0.48 | 0.906455323 | 0.270004998 | 0.068701486 | 0.999652205 | 0.95334888 | 0.976500542 | 0.499920511 |
| Gabarap | High | B1AR50 | Gamma-aminobutyric acid receptor-associated protein OS=Mus musculus OX=10090 GN=Gabara | 0 | 8.284 | 11 | 3 | 4 | 1 | 106 | 12.2 | 9.8 | 0.819 | 0.724 | 1.132 | -0.29 | 0.29 | -0.47 | -0.1566667 | 0.38 | 0.18 | 0.461632926 | 0.281197675 | 0.460181446 | 0.999652205 | 0.955820031 | 0.977736118 | 0.959744396 |
| Napg | High | Q9CWZ7 | Gamma-soluble NSF attachment protein OS=Mus musculus OX=10090 GN=Napg PE=1 SV=1 | 0 | 154.809 | 62 | 19 | 54 | 19 | 312 | 34.7 | 5.41 | 0.801 | 0.742 | 1.079 | -0.32 | 0.32 | -0.43 | -0.1433333 | 0.375 | 0.11 | 0.456966792 | 0.316952067 | 0.55404078 | 0.999652205 | 0.955820031 | 0.977736118 | 0.96775057 |
| H2-Ke2; Pf | High | Q03958 | Prefoldin subunit 6 OS=Mus musculus OX=10090 GN=Pfdn6 PE=1 SV=1 | 0 | 13.389 | 24 | 4 | 4 | 4 | 127 | 14.4 | 8.88 | 0.814 | 0.739 | 1.102 | -0.3 | 0.3 | -0.44 | -0.1466667 | 0.37 | 0.14 | 0.510487362 | 0.310091674 | 0.553516667 | 0.999652205 | 0.955820031 | 0.977736118 | 0.96775057 |
| Cc2d1a | High | Q8K1A6 | Coiled-coil and C2 domain-containing protein 1A OS=Mus musculus OX=10090 GN=Cc2d1a PE=1 S | 0 | 30.87 | 11 | 9 | 10 | 9 | 943 | 103.6 | 7.84 | 0.826 | 0.734 | 1.125 | -0.28 | 0.28 | -0.45 | -0.15 | 0.365 | 0.17 | 0.445296541 | 0.300382682 | 0.494352144 | 0.999652205 | 0.955820031 | 0.977736118 | 0.964378027 |
| Ccbl1; Kya | Medium | Q8BTY1 | Kynurenine--oxoglutarate transaminase 1 OS=Mus musculus OX=10090 GN=Kyat1 PE=1 SV=1 | 0.01 | 2.638 | 2 | 1 | 1 | 1 | 424 | 47.5 | 6.95 | 0.858 | 0.738 | 1.162 | -0.22 | 0.22 | -0.44 | -0.1466667 | 0.33 | 0.22 | 0.500042723 | 0.287369614 | 0.363438749 | 0.999652205 | 0.955820031 | 0.977736118 | 0.91781256 |
| Rbck1 | High | Q9WUB0 | RanBP-type and C3HC4-type zinc finger-containing protein 1 OS=Mus musculus OX=10090 GN=Rb | 0 | 11.823 | 6 | 2 | 2 | 2 | 508 | 57.5 | 6.19 | 0.844 | 0.746 | 1.131 | -0.24 | 0.24 | -0.42 | -0.14 | 0.33 | 0.18 | 0.440447234 | 0.315850756 | 0.462367647 | 0.999652205 | 0.955820031 | 0.977736118 | 0.959744396 |
| Ncs1 | Medium | Q8BNY6 | Neuronal calcium sensor 1 OS=Mus musculus OX=10090 GN=Ncs1 PE=1 SV=3 | 0.015 | 2.211 | 8 | 1 | 1 | 1 | 190 | 21.9 | 4.83 | 0.866 | 0.736 | 1.177 | -0.21 | 0.21 | -0.44 | -0.1466667 | 0.325 | 0.23 | 0.560248543 | 0.281245704 | 0.420860928 | 0.999652205 | 0.955820031 | 0.977736118 | 0.938934618 |
| G6pdx | High | Q00612 | Glucose-6-phosphate 1-dehydrogenase X OS=Mus musculus OX=10090 GN=G6pdx PE=1 SV=3 | 0 | 106.354 | 45 | 20 | 35 | 20 | 515 | 59.2 | 6.49 | 0.898 | 0.72 | 1.246 | -0.16 | 0.16 | -0.47 | -0.1566667 | 0.315 | 0.32 | 0.719518726 | 0.275027254 | 0.128643775 | 0.999652205 | 0.955820031 | 0.977736118 | 0.682408369 |
| Arf3 | High | P61205 | ADP-ribosylation factor 3 OS=Mus musculus OX=10090 GN=Arf3 PE=2 SV=2 | 0 | 91.498 | 48 | 8 | 27 | 5 | 181 | 20.6 | 7.43 | 0.874 | 0.74 | 1.182 | -0.19 | 0.19 | -0.44 | -0.1466667 | 0.315 | 0.24 | 0.652439824 | 0.311824468 | 0.239504212 | 0.999652205 | 0.955820031 | 0.977736118 | 0.831240308 |
| Capn7 | High | Q9R1S8 | Calpain-7 OS=Mus musculus OX=10090 GN=Capn7 PE=2 SV=1 | 0 | 55.897 | 15 | 9 | 11 | 9 | 813 | 92.5 | 7.97 | 0.893 | 0.725 | 1.23 | -0.16 | 0.16 | -0.46 | -0.1533333 | 0.31 | 0.3 | 0.647720422 | 0.284430516 | 0.22071448 | 0.999652205 | 0.955820031 | 0.977736118 | 0.806744518 |
| Ttc1 | High | Q91Z38 | Tetratricopeptide repeat protein 1 OS=Mus musculus OX=10090 GN=Ttc1 PE=1 SV=1 | 0.001 | 7.528 | 6 | 2 | 2 | 2 | 292 | 33.2 | 5.01 | 0.875 | 0.744 | 1.176 | -0.19 | 0.19 | -0.43 | -0.1433333 | 0.31 | 0.23 | 0.522372794 | 0.320809575 | 0.279359431 | 0.999652205 | 0.955820031 | 0.977736118 | 0.876067403 |
| Psma2 | High | P49722 | Proteasome subunit alpha type-2 OS=Mus musculus OX=10090 GN=Psma2 PE=1 SV=3 | 0 | 43.769 | 43 | 6 | 10 | 6 | 234 | 25.9 | 7.43 | 0.899 | 0.726 | 1.238 | -0.15 | 0.15 | -0.46 | -0.1533333 | 0.305 | 0.31 | 0.67411269 | 0.285605877 | 0.192381196 | 0.999652205 | 0.955820031 | 0.977736118 | 0.784495541 |
| Shoc2 | High | O88520 | Leucine-rich repeat protein SHOC-2 OS=Mus musculus OX=10090 GN=Shoc2 PE=1 SV=2 | 0 | 17.298 | 5 | 2 | 3 | 2 | 582 | 64.9 | 8.46 | 0.894 | 0.73 | 1.224 | -0.16 | 0.16 | -0.45 | -0.15 | 0.305 | 0.29 | 0.614445635 | 0.294135241 | 0.224231097 | 0.999652205 | 0.955820031 | 0.977736118 | 0.811661562 |
| Daam1 | High | Q8BPM0 | Disheveled-associated activator of morphogenesis 1 OS=Mus musculus OX=10090 GN=Daam1 PE | 0 | 4.841 | 3 | 3 | 3 | 3 | 1077 | 123.3 | 7.46 | 0.889 | 0.738 | 1.204 | -0.17 | 0.17 | -0.44 | -0.1466667 | 0.305 | 0.27 | 0.57150775 | 0.285523218 | 0.278172239 | 0.999652205 | 0.955820031 | 0.977736118 | 0.876067403 |
| Ppm1b | High | Q99NF7 | Protein-serine/threonine phosphatase OS=Mus musculus OX=10090 GN=Ppm1b PE=1 SV=1 | 0 | 38.285 | 24 | 8 | 10 | 6 | 477 | 52.1 | 5.11 | 0.891 | 0.742 | 1.2 | -0.17 | 0.17 | -0.43 | -0.1433333 | 0.3 | 0.26 | 0.626684811 | 0.317659227 | 0.294368316 | 0.999652205 | 0.955820031 | 0.977736118 | 0.894411787 |
| Pin1 | High | Q9QUR7 | Peptidyl-prolyl cis-trans isomerase NIMA-interacting 1 OS=Mus musculus OX=10090 GN=Pin1 PE= | 0 | 10.573 | 21 | 2 | 3 | 2 | 165 | 18.4 | 8.79 | 0.909 | 0.731 | 1.244 | -0.14 | 0.14 | -0.45 | -0.15 | 0.295 | 0.31 | 0.698028527 | 0.287337248 | 0.220184918 | 0.999652205 | 0.955820031 | 0.977736118 | 0.805859554 |
| Ppa1 | High | Q9D819 | Inorganic pyrophosphatase OS=Mus musculus OX=10090 GN=Ppa1 PE=1 SV=1 | 0 | 145.313 | 66 | 19 | 39 | 19 | 289 | 32.6 | 5.6 | 0.929 | 0.719 | 1.292 | -0.11 | 0.11 | -0.48 | -0.16 | 0.295 | 0.37 | 0.805982352 | 0.271842728 | 0.079752764 | 0.999652205 | 0.955820031 | 0.977736118 | 0.544006989 |
| | High | E9Q9B0 | Kinesin-like protein KIF1C OS=Mus musculus OX=10090 GN=Kif1c PE=1 SV=1 | 0 | 10.279 | 2 | 2 | 2 | 1 | 1112 | 123.7 | 6.93 | 0.902 | 0.737 | 1.223 | -0.15 | 0.15 | -0.44 | -0.1466667 | 0.295 | 0.29 | 0.658916451 | 0.298396272 | 0.2562371 | 0.999652205 | 0.955820031 | 0.977736118 | 0.855700667 |
| Psma5 | High | Q9Z2U1 | Proteasome subunit alpha type-5 OS=Mus musculus OX=10090 GN=Psma5 PE=1 SV=1 | 0 | 35.424 | 32 | 6 | 8 | 6 | 241 | 26.4 | 4.79 | 0.923 | 0.725 | 1.273 | -0.12 | 0.12 | -0.46 | -0.1533333 | 0.29 | 0.35 | 0.70496899 | 0.284485185 | 0.15213031 | 0.999652205 | 0.955820031 | 0.977736118 | 0.724325782 |
| | High | H3BIX0 | Treacle protein (Fragment) OS=Mus musculus OX=10090 GN=Tcof1 PE=1 SV=1 | 0 | 11.578 | 4 | 3 | 3 | 3 | 1056 | 107 | 6.95 | 0.895 | 0.746 | 1.2 | -0.16 | 0.16 | -0.42 | -0.14 | 0.29 | 0.26 | 0.618566523 | 0.293346997 | 0.248301262 | 0.999652205 | 0.955820031 | 0.977736118 | 0.84479683 |
| Pepd | High | Q11136 | Xaa-Pro dipeptidase OS=Mus musculus OX=10090 GN=Pepd PE=1 SV=3 | 0.006 | 3.208 | 2 | 1 | 1 | 1 | 493 | 55 | 5.78 | 0.892 | 0.746 | 1.196 | -0.16 | 0.16 | -0.42 | -0.14 | 0.29 | 0.26 | 0.607129808 | 0.304827567 | 0.29211648 | 0.999652205 | 0.955820031 | 0.977736118 | 0.893073667 |
| Psmc1 | High | P62192 | 26S proteasome regulatory subunit 4 OS=Mus musculus OX=10090 GN=Psmc1 PE=1 SV=1 | 0 | 30.708 | 21 | 9 | 10 | 8 | 440 | 49.2 | 6.21 | 0.929 | 0.727 | 1.278 | -0.11 | 0.11 | -0.46 | -0.1533333 | 0.285 | 0.35 | 0.816023137 | 0.28690615 | 0.108124447 | 0.999652205 | 0.955820031 | 0.977736118 | 0.630562179 |
| Slc7a8 | High | Q9QXW9 | Large neutral amino acids transporter small subunit 2 OS=Mus musculus OX=10090 GN=Slc7a8 Pl | 0 | 25.693 | 9 | 5 | 7 | 5 | 531 | 57.8 | 6.67 | 0.903 | 0.745 | 1.211 | -0.15 | 0.15 | -0.42 | -0.14 | 0.285 | 0.28 | 0.657410534 | 0.323260972 | 0.259137797 | 0.999652205 | 0.955820031 | 0.977736118 | 0.85616924 |
| Aldh7a1 | High | Q9DBF1 | Alpha-aminoadipic semialdehyde dehydrogenase OS=Mus musculus OX=10090 GN=Aldh7a1 PE=1 | 0 | 55.71 | 30 | 14 | 17 | 14 | 539 | 58.8 | 7.47 | 0.929 | 0.734 | 1.266 | -0.11 | 0.11 | -0.45 | -0.15 | 0.28 | 0.34 | 0.807253826 | 0.300875428 | 0.10518423 | 0.999652205 | 0.955820031 | 0.977736118 | 0.618854123 |
| Psmb5 | High | O55234 | Proteasome subunit beta type-5 OS=Mus musculus OX=10090 GN=Psmb5 PE=1 SV=3 | 0 | 34.522 | 30 | 8 | 11 | 8 | 264 | 28.5 | 7.02 | 0.922 | 0.745 | 1.238 | -0.12 | 0.12 | -0.43 | -0.1433333 | 0.275 | 0.31 | 0.713905401 | 0.322102764 | 0.232252237 | 0.999652205 | 0.955820031 | 0.977736118 | 0.820566032 |
| Cand1 | High | Q6ZQ38 | Cullin-associated NEDD8-dissociated protein 1 OS=Mus musculus OX=10090 GN=Cand1 PE=1 SV= | 0 | 153.317 | 34 | 32 | 43 | 32 | 1230 | 136.2 | 5.78 | 0.926 | 0.743 | 1.247 | -0.11 | 0.11 | -0.43 | -0.1433333 | 0.27 | 0.32 | 0.798569905 | 0.318664808 | 0.128452106 | 0.999652205 | 0.955820031 | 0.977736118 | 0.682036893 |
| Psma1 | High | Q9R1P4 | Proteasome subunit alpha type-1 OS=Mus musculus OX=10090 GN=Psma1 PE=1 SV=1 | 0 | 46.92 | 42 | 10 | 17 | 10 | 263 | 29.5 | 6.46 | 0.932 | 0.743 | 1.255 | -0.1 | 0.1 | -0.43 | -0.1433333 | 0.265 | 0.33 | 0.846016316 | 0.318554106 | 0.11381496 | 0.999652205 | 0.955820031 | 0.977736118 | 0.643956085 |
| Arf6 | High | P62331 | ADP-ribosylation factor 6 OS=Mus musculus OX=10090 GN=Arf6 PE=1 SV=2 | 0 | 15.231 | 21 | 3 | 4 | 3 | 175 | 20.1 | 8.95 | 0.942 | 0.735 | 1.282 | -0.09 | 0.09 | -0.44 | -0.1466667 | 0.265 | 0.36 | 0.771279269 | 0.302399179 | 0.143722093 | 0.999652205 | 0.955820031 | 0.977736118 | 0.70564779 |
| Gbe1 | High | Q9D6Y9 | 1,4-alpha-glucan-branching enzyme OS=Mus musculus OX=10090 GN=Gbe1 PE=1 SV=1 | 0 | 62.932 | 23 | 14 | 17 | 14 | 702 | 80.3 | 6.43 | 0.992 | 0.721 | 1.376 | -0.01 | 0.01 | -0.47 | -0.1566667 | 0.24 | 0.46 | 0.944799806 | 0.276039032 | 0.043664265 | 0.999652205 | 0.955820031 | 0.977736118 | 0.404826557 |
| Cyp3a11 | Medium | Q64459 | Cytochrome P450 3A11 OS=Mus musculus OX=10090 GN=Cyp3a11 PE=1 SV=1 | 0.035 | 1.802 | 3 | 1 | 1 | 1 | 504 | 57.8 | 8.82 | 0.976 | 0.736 | 1.325 | -0.04 | 0.04 | -0.44 | -0.1466667 | 0.24 | 0.41 | 0.952800576 | 0.305832731 | 0.104702155 | 0.999652205 | 0.955820031 | 0.977736118 | 0.616664896 |
| Eef2 | High | P58252 | Elongation factor 2 OS=Mus musculus OX=10090 GN=Eef2 PE=1 SV=2 | 0 | 301.013 | 61 | 45 | 119 | 44 | 858 | 95.3 | 6.83 | 0.993 | 0.727 | 1.367 | -0.01 | 0.01 | -0.46 | -0.1533333 | 0.235 | 0.45 | 0.984185744 | 0.286771138 | 0.034515219 | 0.999652205 | 0.955820031 | 0.977736118 | 0.364272011 |
| Yrdc | Medium | Q3U5F4 | YrdC domain-containing protein, mitochondrial OS=Mus musculus OX=10090 GN=Yrdc PE=1 SV=1 | 0.014 | 2.243 | 4 | 1 | 1 | 1 | 280 | 29.4 | 6.07 | 0.977 | 0.754 | 1.295 | -0.03 | 0.03 | -0.41 | -0.1366667 | 0.22 | 0.37 | 0.886141066 | 0.323410074 | 0.168183114 | 0.999652205 | 0.955820031 | 0.977736118 | 0.738806217 |
| | High | A0A1Y7VI | Serine/threonine-protein phosphatase 2A 56 kDa regulatory subunit gamma isoform (Fragment) | 0 | 13.176 | 18 | 4 | 6 | 3 | 283 | 32.8 | 8.29 | 1.025 | 0.725 | 1.414 | 0.04 | -0.04 | -0.46 | -0.1533333 | 0.21 | 0.5 | 0.971724265 | 0.283538871 | 0.047341363 | 0.999652205 | 0.955820031 | 0.977736118 | 0.414107675 |
| Edil3 | High | O35474 | EGF-like repeat and discoidin I-like domain-containing protein 3 OS=Mus musculus OX=10090 GN | 0.001 | 4.362 | 2 | 1 | 1 | 1 | 480 | 53.7 | 7.58 | 1.026 | 0.743 | 1.381 | 0.04 | -0.04 | -0.43 | -0.1433333 | 0.195 | 0.47 | 0.967560286 | 0.308268772 | 0.05988513 | 0.999652205 | 0.955820031 | 0.977736118 | 0.472923839 |
| Cyb5; Cyb5 | High | G5E850 | Cytochrome b5 OS=Mus musculus OX=10090 GN=Cyb5a PE=1 SV=1 | 0.006 | 3.332 | 12 | 1 | 1 | 1 | 98 | 11.1 | 5.3 | 0.889 | 1.44 | 0.618 | -0.17 | 0.17 | 0.53 | 0.17666667 | -0.18 | -0.7 | 0.584231479 | 0.291089998 | 0.012862338 | 0.999652205 | 0.955820031 | 0.977736118 | 0.187810232 |
| | High | A0A494BA | Endoplasmic reticulum-Golgi intermediate compartment protein 1 OS=Mus musculus OX=10090 | 0.001 | 5.087 | 8 | 2 | 2 | 2 | 247 | 27.7 | 7.61 | 0.964 | 1.408 | 0.684 | -0.05 | 0.05 | 0.49 | 0.16333333 | -0.22 | -0.55 | 0.818922708 | 0.322894998 | 0.032184299 | 0.999652205 | 0.955820031 | 0.977736118 | 0.347869094 |
| Ddx39b | High | Q9Z1N5 | Spliceosome RNA helicase Ddx39b OS=Mus musculus OX=10090 GN=Ddx39b PE=1 SV=1 | 0.001 | 6.777 | 6 | 2 | 2 | 2 | 428 | 49 | 5.67 | 0.953 | 1.445 | 0.659 | -0.07 | 0.07 | 0.53 | 0.17666667 | -0.23 | -0.6 | 0.778931552 | 0.292415214 | 0.027703959 | 0.999652205 | 0.955820031 | 0.977736118 | 0.315745518 |
| Cds2 | High | Q99L43 | Phosphatidate cytidylyltransferase 2 OS=Mus musculus OX=10090 GN=Cds2 PE=1 SV=1 | 0 | 20.331 | 12 | 3 | 4 | 3 | 444 | 51.3 | 7.05 | 1.008 | 1.433 | 0.703 | 0.01 | -0.01 | 0.52 | 0.17333333 | -0.265 | -0.51 | 0.949730426 | 0.308366868 | 0.040524271 | 0.999652205 | 0.955820031 | 0.977736118 | 0.388243087 |
| Pgrmc1 | High | O55022 | Membrane-associated progesterone receptor component 1 OS=Mus musculus OX=10090 GN=Pgr | 0 | 18.441 | 19 | 5 | 6 | 4 | 195 | 21.7 | 4.7 | 1.013 | 1.42 | 0.714 | 0.02 | -0.02 | 0.51 | 0.17 | -0.265 | -0.49 | 0.943521344 | 0.322813213 | 0.063862225 | 0.999652205 | 0.955820031 | 0.977736118 | 0.485498168 |
| Uggt1 | High | Q6P5E4 | UDP-glucose:glycoprotein glucosyltransferase 1 OS=Mus musculus OX=10090 GN=Uggt1 PE=1 SV | 0 | 9.186 | 3 | 4 | 4 | 4 | 1551 | 176.3 | 5.62 | 1.051 | 1.422 | 0.739 | 0.07 | -0.07 | 0.51 | 0.17 | -0.29 | -0.44 | 0.926678328 | 0.314476783 | 0.061257979 | 0.999652205 | 0.955820031 | 0.977736118 | 0.476099813 |
| 0610009D | Medium | P59708 | Splicing factor 3B subunit 6 OS=Mus musculus OX=10090 GN=Sf3b6 PE=1 SV=1 | 0.03 | 1.868 | 6 | 1 | 1 | 1 | 125 | 14.6 | 9.38 | 1.089 | 1.428 | 0.763 | 0.12 | -0.12 | 0.51 | 0.17 | -0.315 | -0.39 | 0.743756786 | 0.314194056 | 0.123996012 | 0.999652205 | 0.955820031 | 0.977736118 | 0.677192798 |
| Rcn1 | High | Q05186 | Reticulocalbin-1 OS=Mus musculus OX=10090 GN=Rcn1 PE=1 SV=1 | 0.001 | 7.098 | 8 | 2 | 2 | 2 | 325 | 38.1 | 4.84 | 1.096 | 1.416 | 0.774 | 0.13 | -0.13 | 0.5 | 0.16666667 | -0.315 | -0.37 | 0.787755961 | 0.319758684 | 0.156516806 | 0.999652205 | 0.955820031 | 0.977736118 | 0.73193472 |
| | Medium | A0A0J9YU | Actin-depolymerizing factor (Fragment) OS=Mus musculus OX=10090 GN=Gsn PE=1 SV=1 | 0.047 | 1.546 | 1 | 1 | 1 | 1 | 568 | 62.3 | 6.92 | 1.064 | 1.467 | 0.725 | 0.09 | -0.09 | 0.55 | 0.18333333 | -0.32 | -0.46 | 0.882303215 | 0.280038639 | 0.141846561 | 0.999652205 | 0.955820031 | 0.977736118 | 0.701851981 |
| 8430419L | High | Q8BYI8 | Protein FAM234B OS=Mus musculus OX=10090 GN=Fam234b PE=1 SV=1 | 0.001 | 6.072 | 4 | 2 | 2 | 2 | 624 | 67 | 5.03 | 1.084 | 1.442 | 0.752 | 0.12 | -0.12 | 0.53 | 0.17666667 | -0.325 | -0.41 | 0.799378292 | 0.298273152 | 0.108589926 | 0.999652205 | 0.955820031 | 0.977736118 | 0.631380318 |
| Top2b | High | Q64511 | DNA topoisomerase 2-beta OS=Mus musculus OX=10090 GN=Top2b PE=1 SV=2 | 0 | 21.937 | 4 | 8 | 11 | 8 | 1612 | 181.8 | 8.29 | 1.078 | 1.469 | 0.734 | 0.11 | -0.11 | 0.55 | 0.18333333 | -0.33 | -0.45 | 0.845426637 | 0.27401393 | 0.050645223 | 0.999652205 | 0.955820031 | 0.977736118 | 0.427566646 |
| Atp2a2 | High | O55143 | Sarcoplasmic/endoplasmic reticulum calcium ATPase 2 OS=Mus musculus OX=10090 GN=Atp2a2 | 0 | 111.383 | 21 | 25 | 37 | 25 | 1044 | 114.8 | 5.34 | 1.111 | 1.421 | 0.782 | 0.15 | -0.15 | 0.51 | 0.17 | -0.33 | -0.36 | 0.720967869 | 0.321371341 | 0.137341849 | 0.999652205 | 0.955820031 | 0.977736118 | 0.700189449 |
| Yipf5 | High | Q9EQQ2 | Protein YIPF5 OS=Mus musculus OX=10090 GN=Yipf5 PE=1 SV=1 | 0.001 | 5.409 | 8 | 2 | 2 | 2 | 257 | 27.9 | 4.36 | 1.122 | 1.423 | 0.789 | 0.17 | -0.17 | 0.51 | 0.17 | -0.34 | -0.34 | 0.723678982 | 0.315057413 | 0.215421616 | 0.999652205 | 0.955820031 | 0.977736118 | 0.804011919 |
| Ccdc109a; | High | Q3UMR5 | Calcium uniporter protein, mitochondrial OS=Mus musculus OX=10090 GN=Mcu PE=1 SV=2 | 0 | 8.24 | 15 | 4 | 4 | 4 | 350 | 39.7 | 8.56 | 1.111 | 1.463 | 0.76 | 0.15 | -0.15 | 0.55 | 0.18333333 | -0.35 | -0.4 | 0.738092305 | 0.273615404 | 0.094930214 | 0.999652205 | 0.955820031 | 0.977736118 | 0.586204524 |
| Sorcs2 | High | Q9EPR5 | VPS10 domain-containing receptor SorCS2 OS=Mus musculus OX=10090 GN=Sorcs2 PE=1 SV=2 | 0.001 | 7.823 | 2 | 3 | 3 | 3 | 1159 | 128.8 | 7.34 | 1.162 | 1.406 | 0.827 | 0.22 | -0.22 | 0.49 | 0.16333333 | -0.355 | -0.27 | 0.590784151 | 0.32303692 | 0.283342553 | 0.999652205 | 0.955820031 | 0.977736118 | 0.886074068 |
| 5730469M | High | Q9CYH2 | Peroxiredoxin-like 2A OS=Mus musculus OX=10090 GN=Prxl2a PE=1 SV=2 | 0 | 26.61 | 34 | 7 | 7 | 7 | 218 | 24.4 | 9.17 | 1.138 | 1.444 | 0.788 | 0.19 | -0.19 | 0.53 | 0.17666667 | -0.36 | -0.34 | 0.609396671 | 0.29804694 | 0.172752877 | 0.999652205 | 0.955820031 | 0.977736118 | 0.738844686 |
| Ghitm | High | Q91VC9 | Growth hormone-inducible transmembrane protein OS=Mus musculus OX=10090 GN=Ghitm PE= | 0 | 8.452 | 6 | 2 | 2 | 2 | 346 | 37.3 | 9.8 | 1.171 | 1.418 | 0.825 | 0.23 | -0.23 | 0.5 | 0.16666667 | -0.365 | -0.28 | 0.588586015 | 0.321877826 | 0.257175127 | 0.999652205 | 0.955820031 | 0.977736118 | 0.85616924 |
| Vipr2 | High | P41588 | Vasoactive intestinal polypeptide receptor 2 OS=Mus musculus OX=10090 GN=Vipr2 PE=1 SV=1 | 0 | 8.677 | 5 | 2 | 2 | 2 | 437 | 49.4 | 7.61 | 1.177 | 1.409 | 0.836 | 0.24 | -0.24 | 0.49 | 0.16333333 | -0.365 | -0.26 | 0.575913041 | 0.318696751 | 0.323914829 | 0.999652205 | 0.955820031 | 0.977736118 | 0.905686588 |
| Syngr3 | High | Q8R191 | Synaptogyrin-3 OS=Mus musculus OX=10090 GN=Syngr3 PE=1 SV=1 | 0 | 8.578 | 12 | 2 | 2 | 2 | 229 | 24.5 | 8.18 | 1.149 | 1.453 | 0.791 | 0.2 | -0.2 | 0.54 | 0.18 | -0.37 | -0.34 | 0.655781626 | 0.28936326 | 0.160350978 | 0.999652205 | 0.955820031 | 0.977736118 | 0.733684585 |
| Abhd6 | High | Q8R2Y0 | Monoacylglycerol lipase ABHD6 OS=Mus musculus OX=10090 GN=Abhd6 PE=1 SV=1 | 0 | 8.062 | 7 | 3 | 3 | 3 | 336 | 38.2 | 8.47 | 1.15 | 1.458 | 0.788 | 0.2 | -0.2 | 0.54 | 0.18 | -0.37 | -0.34 | 0.643121266 | 0.283859541 | 0.194192154 | 0.999652205 | 0.955820031 | 0.977736118 | 0.784517504 |
| Fmr1 | High | Q6AXB7 | Synaptic functional regulator FMR1 OS=Mus musculus OX=10090 GN=Fmr1 PE=1 SV=1 | 0.001 | 5.742 | 3 | 2 | 3 | 1 | 614 | 68.8 | 7.42 | 1.184 | 1.419 | 0.835 | 0.24 | -0.24 | 0.5 | 0.16666667 | -0.37 | -0.26 | 0.538700996 | 0.32357922 | 0.316005568 | 0.999652205 | 0.955820031 | 0.977736118 | 0.904943958 |
| Clic6 | High | Q8BHB9 | Chloride intracellular channel protein 6 OS=Mus musculus OX=10090 GN=Clic6 PE=1 SV=1 | 0 | 329.058 | 59 | 28 | 122 | 28 | 596 | 62.8 | 4.32 | 1.171 | 1.439 | 0.814 | 0.23 | -0.23 | 0.52 | 0.17333333 | -0.375 | -0.3 | 0.592401828 | 0.302920777 | 0.220405583 | 0.999652205 | 0.955820031 | 0.977736118 | 0.806140969 |
| Ubap2l | High | Q80X50 | Ubiquitin-associated protein 2-like OS=Mus musculus OX=10090 GN=Ubap2l PE=1 SV=1 | 0 | 21.76 | 5 | 5 | 5 | 5 | 1107 | 116.7 | 7.11 | 1.179 | 1.423 | 0.829 | 0.24 | -0.24 | 0.51 | 0.17 | -0.375 | -0.27 | 0.570536357 | 0.319387176 | 0.291862253 | 0.999652205 | 0.955820031 | 0.977736118 | 0.892844069 |
| | High | A0A286YC | BDNF/NT-3 growth factors receptor OS=Mus musculus OX=10090 GN=Ntrk2 PE=1 SV=1 | 0.001 | 5.163 | 3 | 1 | 1 | 1 | 489 | 54.6 | 6.76 | 1.207 | 1.409 | 0.857 | 0.27 | -0.27 | 0.49 | 0.16333333 | -0.38 | -0.22 | 0.487607129 | 0.32083428 | 0.404491297 | 0.999652205 | 0.955820031 | 0.977736118 | 0.934095017 |
| Ndufa6 | High | Q9CQZ5 | NADH dehydrogenase [ubiquinone] 1 alpha subcomplex subunit 6 OS=Mus musculus OX=10090 G | 0 | 17.153 | 37 | 4 | 4 | 4 | 131 | 15.3 | 10.11 | 1.174 | 1.45 | 0.81 | 0.23 | -0.23 | 0.54 | 0.18 | -0.385 | -0.3 | 0.50938963 | 0.29150857 | 0.231494522 | 0.999652205 | 0.955820031 | 0.977736118 | 0.819676582 |
| Gosr2 | Medium | A2A9I0 | Golgi SNAP receptor complex member 2 OS=Mus musculus OX=10090 GN=Gosr2 PE=1 SV=1 | 0.03 | 1.859 | 4 | 1 | 1 | 1 | 165 | 19.6 | 9.5 | 1.173 | 1.457 | 0.805 | 0.23 | -0.23 | 0.54 | 0.18 | -0.385 | -0.31 | 0.593928563 | 0.277792316 | 0.259976013 | 0.999652205 | 0.955820031 | 0.977736118 | 0.85616924 |
| Acsl6 | High | Q91WC3 | Long-chain-fatty-acid--CoA ligase 6 OS=Mus musculus OX=10090 GN=Acsl6 PE=1 SV=1 | 0 | 55.045 | 28 | 16 | 21 | 16 | 697 | 78 | 7.34 | 1.191 | 1.455 | 0.819 | 0.25 | -0.25 | 0.54 | 0.18 | -0.395 | -0.29 | 0.521223401 | 0.287471175 | 0.235071785 | 0.999652205 | 0.955820031 | 0.977736118 | 0.826767714 |
| Ddost | High | O54734 | Dolichyl-diphosphooligosaccharide--protein glycosyltransferase 48 kDa subunit OS=Mus musculus | 0 | 8.421 | 6 | 3 | 3 | 3 | 441 | 49 | 5.83 | 1.19 | 1.459 | 0.816 | 0.25 | -0.25 | 0.54 | 0.18 | -0.395 | -0.29 | 0.543382751 | 0.283547264 | 0.239572737 | 0.999652205 | 0.955820031 | 0.977736118 | 0.831240308 |
| Ndufs8 | High | Q8K3J1 | NADH dehydrogenase [ubiquinone] iron-sulfur protein 8, mitochondrial OS=Mus musculus OX=100 | 0 | 15.246 | 13 | 3 | 7 | 3 | 212 | 24 | 6.21 | 1.204 | 1.432 | 0.841 | 0.27 | -0.27 | 0.52 | 0.17333333 | -0.395 | -0.25 | 0.506298254 | 0.310103559 | 0.288047173 | 0.999652205 | 0.955820031 | 0.977736118 | 0.887894721 |
| Metap2 | High | O08663 | Methionine aminopeptidase 2 OS=Mus musculus OX=10090 GN=Metap2 PE=1 SV=1 | 0 | 4.809 | 2 | 1 | 1 | 1 | 478 | 52.9 | 5.82 | 1.207 | 1.441 | 0.837 | 0.27 | -0.27 | 0.53 | 0.17666667 | -0.4 | -0.26 | 0.508805904 | 0.29458755 | 0.31048278 | 0.999652205 | 0.955820031 | 0.977736118 | 0.904943958 |
| Ndufb7 | High | Q9CR61 | NADH dehydrogenase [ubiquinone] 1 beta subcomplex subunit 7 OS=Mus musculus OX=10090 GN | 0 | 11.215 | 26 | 3 | 3 | 3 | 137 | 16.3 | 8.18 | 1.204 | 1.462 | 0.823 | 0.27 | -0.27 | 0.55 | 0.18333333 | -0.41 | -0.28 | 0.545331331 | 0.272470413 | 0.257433845 | 0.999652205 | 0.955820031 | 0.977736118 | 0.85616924 |
| Tmem87b | Low | Q8BKU8 | Transmembrane protein 87B OS=Mus musculus OX=10090 GN=Tmem87b PE=2 SV=1 | 0.084 | 1.183 | 2 | 1 | 1 | 1 | 555 | 62.9 | 6 | 1.246 | 1.425 | 0.875 | 0.32 | -0.32 | 0.51 | 0.17 | -0.415 | -0.19 | 0.44274314 | 0.311269501 | 0.487913429 | 0.999652205 | 0.955820031 | 0.977736118 | 0.964378027 |
| Espn | High | B1AWQ4 | Espin OS=Mus musculus OX=10090 GN=Espn PE=1 SV=1 | 0 | 10.745 | 4 | 4 | 6 | 4 | 881 | 94.9 | 7.46 | 1.251 | 1.421 | 0.88 | 0.32 | -0.32 | 0.51 | 0.17 | -0.415 | -0.18 | 0.43277681 | 0.320968338 | 0.488278225 | 0.999652205 | 0.955820031 | 0.977736118 | 0.964378027 |
| Ndufb11 | High | O09111 | NADH dehydrogenase [ubiquinone] 1 beta subcomplex subunit 11, mitochondrial OS=Mus muscul | 0.006 | 3.165 | 7 | 1 | 1 | 1 | 151 | 17.4 | 5.22 | 1.247 | 1.446 | 0.862 | 0.32 | -0.32 | 0.53 | 0.17666667 | -0.425 | -0.21 | 0.438288959 | 0.295590446 | 0.38598484 | 0.999652205 | 0.955820031 | 0.977736118 | 0.929303068 |
| Pdhb | High | Q9D051 | Pyruvate dehydrogenase E1 component subunit beta, mitochondrial OS=Mus musculus OX=10090 | 0 | 34.744 | 31 | 8 | 10 | 8 | 359 | 38.9 | 6.87 | 1.273 | 1.44 | 0.884 | 0.35 | -0.35 | 0.53 | 0.17666667 | -0.44 | -0.18 | 0.43651166 | 0.302020041 | 0.464956918 | 0.999652205 | 0.955820031 | 0.977736118 | 0.960359242 |
| Psmd12 | High | Q9D8W5 | 26S proteasome non-ATPase regulatory subunit 12 OS=Mus musculus OX=10090 GN=Psmd12 PE= | 0 | 15.235 | 6 | 3 | 6 | 3 | 456 | 52.9 | 7.06 | 1.059 | 0.747 | 1.418 | 0.08 | -0.08 | -0.42 | -0.14 | 0.17 | 0.5 | 0.894891652 | 0.325992738 | 0.040953816 | 0.999652205 | 0.960000674 | 0.97982644 | 0.391189174 |
| Slc5a3 | High | Q9JKZ2 | Sodium/myo-inositol cotransporter OS=Mus musculus OX=10090 GN=Slc5a3 PE=1 SV=2 | 0.008 | 3.044 | 2 | 2 | 3 | 2 | 718 | 79.5 | 7.3 | 1.071 | 0.749 | 1.429 | 0.1 | -0.1 | -0.42 | -0.14 | 0.16 | 0.52 | 0.823583505 | 0.327064689 | 0.039617612 | 0.999652205 | 0.961945011 | 0.980798608 | 0.384984321 |
| Map2k2 | High | Q91YS7 | Dual-specificity mitogen-activated protein kinase kinase 2 OS=Mus musculus OX=10090 GN=Map | 0 | 11.189 | 9 | 4 | 5 | 3 | 400 | 44.3 | 7.05 | 1.005 | 0.749 | 1.34 | 0.01 | -0.01 | -0.42 | -0.14 | 0.205 | 0.42 | 0.981037245 | 0.331856821 | 0.086885381 | 0.999652205 | 0.968621131 | 0.984136668 | 0.571121137 |
| Eef1a1; LC | High | P10126 | Elongation factor 1-alpha 1 OS=Mus musculus OX=10090 GN=Eef1a1 PE=1 SV=3 | 0 | 276.163 | 75 | 31 | 140 | 17 | 462 | 50.1 | 9.01 | 0.846 | 0.75 | 1.128 | -0.24 | 0.24 | -0.41 | -0.1366667 | 0.325 | 0.17 | 0.576525926 | 0.333935508 | 0.381979773 | 0.999652205 | 0.97322306 | 0.986437632 | 0.923969192 |
| Mreg | High | Q6NVG5 | Melanoregulin OS=Mus musculus OX=10090 GN=Mreg PE=1 SV=1 | 0.001 | 7.053 | 10 | 2 | 2 | 2 | 214 | 25 | 7.2 | 0.873 | 1.403 | 0.622 | -0.2 | 0.2 | 0.49 | 0.16333333 | -0.145 | -0.69 | 0.540929955 | 0.334127767 | 0.01178711 | 0.999652205 | 0.97322306 | 0.986437632 | 0.179106573 |
| Rffl | High | Q6ZQM0 | E3 ubiquitin-protein ligase rififylin OS=Mus musculus OX=10090 GN=Rffl PE=1 SV=1 | 0 | 21.105 | 21 | 4 | 4 | 4 | 377 | 42.2 | 5.82 | 0.967 | 0.751 | 1.287 | -0.05 | 0.05 | -0.41 | -0.1366667 | 0.23 | 0.36 | 0.932924764 | 0.334930088 | 0.133040216 | 0.999652205 | 0.974547485 | 0.987099845 | 0.695121227 |
| Chmp6 | High | P0C0A3 | Charged multivesicular body protein 6 OS=Mus musculus OX=10090 GN=Chmp6 PE=1 SV=2 | 0 | 57.519 | 35 | 5 | 15 | 5 | 200 | 23.4 | 5.44 | 1.116 | 1.407 | 0.793 | 0.16 | -0.16 | 0.49 | 0.16333333 | -0.325 | -0.34 | 0.752842336 | 0.335795149 | 0.15527384 | 0.999652205 | 0.975040602 | 0.987346404 | 0.730581221 |
| Rufy3 | High | A0A0G2JF | Protein RUFY3 OS=Mus musculus OX=10090 GN=Rufy3 PE=1 SV=1 | 0 | 107.746 | 35 | 21 | 36 | 6 | 669 | 74.7 | 6.19 | 0.975 | 0.753 | 1.294 | -0.04 | 0.04 | -0.41 | -0.1366667 | 0.225 | 0.37 | 0.934120709 | 0.339937135 | 0.0783177 | 0.999652205 | 0.976045759 | 0.987848982 | 0.535521148 |
| Mapre2 | High | E9Q6X0 | Microtubule-associated protein RP/EB family member 2 OS=Mus musculus OX=10090 GN=Mapre | 0 | 43.4 | 47 | 9 | 15 | 8 | 284 | 32.2 | 5.26 | 1.015 | 0.753 | 1.347 | 0.02 | -0.02 | -0.41 | -0.1366667 | 0.195 | 0.43 | 0.969495123 | 0.339319142 | 0.090088909 | 0.999652205 | 0.976045759 | 0.987848982 | 0.5746627 |
| Scarb2 | High | O35114 | Lysosome membrane protein 2 OS=Mus musculus OX=10090 GN=Scarb2 PE=1 SV=3 | 0.001 | 7.411 | 4 | 2 | 3 | 2 | 478 | 54 | 5.1 | 0.991 | 1.406 | 0.705 | -0.01 | 0.01 | 0.49 | 0.16333333 | -0.24 | -0.5 | 0.890656731 | 0.337459531 | 0.042936094 | 0.999652205 | 0.976045759 | 0.987848982 | 0.403254067 |
| Sgk3 | High | Q9ERE3 | Serine/threonine-protein kinase Sgk3 OS=Mus musculus OX=10090 GN=Sgk3 PE=1 SV=1 | 0.008 | 2.936 | 2 | 1 | 1 | 1 | 496 | 57.1 | 7.15 | 1.067 | 1.394 | 0.765 | 0.09 | -0.09 | 0.48 | 0.16 | -0.285 | -0.39 | 0.844680513 | 0.340654902 | 0.155706181 | 0.999652205 | 0.976045759 | 0.987848982 | 0.730581221 |
| Prkcsh | High | O08795 | Glucosidase 2 subunit beta OS=Mus musculus OX=10090 GN=Prkcsh PE=1 SV=1 | 0 | 31.404 | 12 | 8 | 10 | 8 | 521 | 58.8 | 4.46 | 1.115 | 1.406 | 0.793 | 0.16 | -0.16 | 0.49 | 0.16333333 | -0.325 | -0.33 | 0.744480986 | 0.337717648 | 0.147241401 | 0.999652205 | 0.976045759 | 0.987848982 | 0.712323156 |
| Ank3 | High | G5E8K5 | Ankyrin-3 OS=Mus musculus OX=10090 GN=Ank3 PE=1 SV=1 | 0 | 23.248 | 4 | 8 | 9 | 6 | 1961 | 213.9 | 6.7 | 0.789 | 0.769 | 1.026 | -0.34 | 0.34 | -0.38 | -0.1266667 | 0.36 | 0.04 | 0.439827129 | 0.372402757 | 0.855654458 | 0.999652205 | 0.977313184 | 0.988482695 | 0.992535481 |
| Ephb2 | High | P54763 | Ephrin type-B receptor 2 OS=Mus musculus OX=10090 GN=Ephb2 PE=1 SV=3 | 0 | 14.827 | 8 | 6 | 7 | 6 | 986 | 109.8 | 5.71 | 0.806 | 0.773 | 1.042 | -0.31 | 0.31 | -0.37 | -0.1233333 | 0.34 | 0.06 | 0.483277599 | 0.38141592 | 0.750177769 | 0.999652205 | 0.977313184 | 0.988482695 | 0.992535481 |
| Pcmt1 | High | E0CYV0 | Protein-L-isoaspartate O-methyltransferase OS=Mus musculus OX=10090 GN=Pcmt1 PE=1 SV=1 | 0 | 34.877 | 43 | 10 | 14 | 10 | 286 | 30.4 | 7.24 | 0.835 | 0.758 | 1.102 | -0.26 | 0.26 | -0.4 | -0.1333333 | 0.33 | 0.14 | 0.549781809 | 0.348681893 | 0.468926593 | 0.999652205 | 0.977313184 | 0.988482695 | 0.961044326 |
| Stxbp3a; S | High | Q60770 | Syntaxin-binding protein 3 OS=Mus musculus OX=10090 GN=Stxbp3 PE=1 SV=1 | 0 | 82.995 | 33 | 19 | 27 | 19 | 592 | 67.9 | 8.02 | 0.831 | 0.764 | 1.087 | -0.27 | 0.27 | -0.39 | -0.13 | 0.33 | 0.12 | 0.534180315 | 0.361973522 | 0.523133682 | 0.999652205 | 0.977313184 | 0.988482695 | 0.964378027 |
| Park7 | High | Q99LX0 | Parkinson disease protein 7 homolog OS=Mus musculus OX=10090 GN=Park7 PE=1 SV=1 | 0 | 103.681 | 80 | 15 | 49 | 15 | 189 | 20 | 6.77 | 0.826 | 0.767 | 1.077 | -0.28 | 0.28 | -0.38 | -0.1266667 | 0.33 | 0.11 | 0.522881395 | 0.36899862 | 0.563549921 | 0.999652205 | 0.977313184 | 0.988482695 | 0.972910028 |
| Ube2d3 | Medium | P61079 | Ubiquitin-conjugating enzyme E2 D3 OS=Mus musculus OX=10090 GN=Ube2d3 PE=1 SV=1 | 0.041 | 1.675 | 5 | 1 | 1 | 1 | 147 | 16.7 | 7.8 | 0.836 | 0.763 | 1.094 | -0.26 | 0.26 | -0.39 | -0.13 | 0.325 | 0.13 | 0.563676586 | 0.360957946 | 0.577145419 | 0.999652205 | 0.977313184 | 0.988482695 | 0.972910028 |
| Zdhhc20 | High | Q5Y5T1 | Palmitoyltransferase ZDHHC20 OS=Mus musculus OX=10090 GN=Zdhhc20 PE=1 SV=1 | 0 | 19.573 | 14 | 5 | 7 | 3 | 380 | 43.9 | 8.25 | 0.83 | 0.766 | 1.083 | -0.27 | 0.27 | -0.38 | -0.1266667 | 0.325 | 0.12 | 0.430508568 | 0.346501024 | 0.640270607 | 0.999652205 | 0.977313184 | 0.988482695 | 0.983830445 |
| Myo5b | High | G5E8G6 | Unconventional myosin-Vb OS=Mus musculus OX=10090 GN=Myo5b PE=1 SV=1 | 0 | 24.513 | 4 | 10 | 12 | 3 | 1818 | 210.4 | 7.09 | 0.84 | 0.765 | 1.098 | -0.25 | 0.25 | -0.39 | -0.13 | 0.32 | 0.13 | 0.461048559 | 0.355551842 | 0.567735841 | 0.999652205 | 0.977313184 | 0.988482695 | 0.972910028 |
| Nudt9 | High | Q8BVU5 | ADP-ribose pyrophosphatase, mitochondrial OS=Mus musculus OX=10090 GN=Nudt9 PE=1 SV=1 | 0.006 | 3.096 | 3 | 1 | 1 | 1 | 350 | 38.6 | 6.76 | 0.841 | 0.761 | 1.105 | -0.25 | 0.25 | -0.39 | -0.13 | 0.32 | 0.14 | 0.493025617 | 0.347076941 | 0.581603999 | 0.999652205 | 0.977313184 | 0.988482695 | 0.972910028 |
| Anxa2 | High | P07356 | Annexin A2 OS=Mus musculus OX=10090 GN=Anxa2 PE=1 SV=2 | 0 | 47.644 | 42 | 12 | 13 | 12 | 339 | 38.7 | 7.69 | 0.839 | 0.769 | 1.09 | -0.25 | 0.25 | -0.38 | -0.1266667 | 0.315 | 0.12 | 0.511007347 | 0.373070389 | 0.618861752 | 0.999652205 | 0.977313184 | 0.988482695 | 0.97826833 |
| Rasgrf2 | High | P70392 | Ras-specific guanine nucleotide-releasing factor 2 OS=Mus musculus OX=10090 GN=Rasgrf2 PE= | 0.001 | 6.742 | 2 | 2 | 2 | 2 | 1189 | 135.6 | 7.85 | 0.831 | 0.779 | 1.067 | -0.27 | 0.27 | -0.36 | -0.12 | 0.315 | 0.09 | 0.469141571 | 0.385050418 | 0.699044367 | 0.999652205 | 0.977313184 | 0.988482695 | 0.991285312 |
| Pfdn2 | High | O70591 | Prefoldin subunit 2 OS=Mus musculus OX=10090 GN=Pfdn2 PE=1 SV=2 | 0 | 19.876 | 47 | 7 | 9 | 7 | 154 | 16.5 | 6.58 | 0.851 | 0.771 | 1.104 | -0.23 | 0.23 | -0.38 | -0.1266667 | 0.305 | 0.14 | 0.572277077 | 0.376760913 | 0.521611058 | 0.999652205 | 0.977313184 | 0.988482695 | 0.964378027 |
| Dctn1 | High | E9Q3M3 | Dynactin subunit 1 OS=Mus musculus OX=10090 GN=Dctn1 PE=1 SV=1 | 0 | 59.259 | 15 | 17 | 21 | 17 | 1264 | 139.7 | 5.71 | 0.851 | 0.773 | 1.101 | -0.23 | 0.23 | -0.37 | -0.1233333 | 0.3 | 0.14 | 0.588123372 | 0.382219216 | 0.509756319 | 0.999652205 | 0.977313184 | 0.988482695 | 0.964378027 |
| Eif4a2 | High | Q8BTU6 | RNA helicase OS=Mus musculus OX=10090 GN=Eif4a2 PE=1 SV=1 | 0 | 49.935 | 32 | 10 | 13 | 3 | 362 | 41.3 | 5.64 | 0.895 | 0.759 | 1.18 | -0.16 | 0.16 | -0.4 | -0.1333333 | 0.28 | 0.24 | 0.681020584 | 0.351392959 | 0.294928044 | 0.999652205 | 0.977313184 | 0.988482695 | 0.894411787 |
| Hook1 | High | Q8BIL5 | Protein Hook homolog 1 OS=Mus musculus OX=10090 GN=Hook1 PE=1 SV=2 | 0 | 17.137 | 9 | 6 | 6 | 6 | 728 | 84.4 | 5.22 | 0.878 | 0.772 | 1.137 | -0.19 | 0.19 | -0.37 | -0.1233333 | 0.28 | 0.19 | 0.626136523 | 0.380078067 | 0.450983018 | 0.999652205 | 0.977313184 | 0.988482695 | 0.959744396 |
| Ssu72 | Medium | Q9CY97 | RNA polymerase II subunit A C-terminal domain phosphatase SSU72 OS=Mus musculus OX=1009 | 0.015 | 2.219 | 8 | 1 | 1 | 1 | 194 | 22.5 | 5.21 | 0.878 | 0.773 | 1.136 | -0.19 | 0.19 | -0.37 | -0.1233333 | 0.28 | 0.18 | 0.556821752 | 0.365203681 | 0.454411897 | 0.999652205 | 0.977313184 | 0.988482695 | 0.959744396 |
| Rab2b | High | P59279 | Ras-related protein Rab-2B OS=Mus musculus OX=10090 GN=Rab2b PE=1 SV=1 | 0 | 83.487 | 50 | 9 | 31 | 3 | 216 | 24.2 | 6.68 | 0.874 | 0.773 | 1.13 | -0.19 | 0.19 | -0.37 | -0.1233333 | 0.28 | 0.18 | 0.572495424 | 0.381216893 | 0.483657561 | 0.999652205 | 0.977313184 | 0.988482695 | 0.964378027 |
| Cul3 | High | Q9JLV5 | Cullin-3 OS=Mus musculus OX=10090 GN=Cul3 PE=1 SV=1 | 0 | 55.886 | 21 | 14 | 16 | 14 | 768 | 88.9 | 8.46 | 0.885 | 0.771 | 1.147 | -0.18 | 0.18 | -0.37 | -0.1233333 | 0.275 | 0.2 | 0.589399742 | 0.377219135 | 0.448271546 | 0.999652205 | 0.977313184 | 0.988482695 | 0.958608248 |
| Me1 | High | P06801 | NADP-dependent malic enzyme OS=Mus musculus OX=10090 GN=Me1 PE=1 SV=2 | 0 | 266.716 | 70 | 32 | 95 | 32 | 572 | 63.9 | 7.44 | 0.922 | 0.759 | 1.215 | -0.12 | 0.12 | -0.4 | -0.1333333 | 0.26 | 0.28 | 0.78632574 | 0.350741103 | 0.175347265 | 0.999652205 | 0.977313184 | 0.988482695 | 0.74518415 |

| | | | | | | | | | | | | | | | | | | | | | | | | | | | | |
|---|---|---|---|---|---|---|---|---|---|---|---|---|---|---|---|---|---|---|---|---|---|---|---|---|---|---|---|---|
| Cdc34 | Low | A0A1W2P | Ubiquitin-conjugating enzyme E2 R1 OS=Mus musculus OX=10090 GN=Cdc34 PE=1 SV=1 | 0.054 | 1.404 | 6 | 1 | 1 | 1 | 190 | 21.4 | 4.41 | 0.897 | 0.778 | 1.154 | -0.16 | 0.16 | -0.36 | -0.12 | 0.26 | 0.21 | 0.647993757 | 0.372335593 | 0.467204923 | 0.999652205 | 0.977313184 | 0.988482695 | 0.960359242 |
| 4931406C | High | A0A1L1SS | RIKEN cDNA 4931406C07 gene OS=Mus musculus OX=10090 GN=4931406C07Rik PE=1 SV=1 | 0 | 8.449 | 12 | 3 | 4 | 3 | 282 | 31.3 | 6.52 | 0.921 | 0.763 | 1.208 | -0.12 | 0.12 | -0.39 | -0.13 | 0.255 | 0.27 | 0.673602446 | 0.343419827 | 0.209847545 | 0.999652205 | 0.977313184 | 0.988482695 | 0.803698897 |
| | High | A0A5F8M | Aquaporin-4 OS=Mus musculus OX=10090 GN=Aqp4 PE=1 SV=1 | 0 | 29.534 | 18 | 6 | 15 | 6 | 330 | 35.3 | 5.8 | 0.912 | 0.768 | 1.187 | -0.13 | 0.13 | -0.38 | -0.1266667 | 0.255 | 0.25 | 0.759099295 | 0.371562329 | 0.228642239 | 0.999652205 | 0.977313184 | 0.988482695 | 0.81760015 |
| Usp33 | High | A0A0H2U | Ubiquitin carboxyl-terminal hydrolase OS=Mus musculus OX=10090 GN=Usp33 PE=1 SV=1 | 0.001 | 7.766 | 2 | 3 | 3 | 3 | 897 | 101.4 | 5.68 | 0.915 | 0.771 | 1.186 | -0.13 | 0.13 | -0.38 | -0.1266667 | 0.255 | 0.25 | 0.620898197 | 0.354327939 | 0.26735927 | 0.999652205 | 0.977313184 | 0.988482695 | 0.869537952 |
| Cnrip1 | High | Q5M8N0 | CB1 cannabinoid receptor-interacting protein 1 OS=Mus musculus OX=10090 GN=Cnrip1 PE=1 SV | 0 | 17.698 | 18 | 3 | 6 | 3 | 164 | 18.6 | 7.96 | 0.922 | 0.766 | 1.203 | -0.12 | 0.12 | -0.38 | -0.1266667 | 0.25 | 0.27 | 0.719857062 | 0.367137902 | 0.287101889 | 0.999652205 | 0.977313184 | 0.988482695 | 0.886931291 |
| Impdh2 | High | P24547 | Inosine-5'-monophosphate dehydrogenase 2 OS=Mus musculus OX=10090 GN=Impdh2 PE=1 SV= | 0 | 25.435 | 15 | 7 | 9 | 5 | 514 | 55.8 | 7.28 | 0.941 | 0.756 | 1.245 | -0.09 | 0.09 | -0.4 | -0.1333333 | 0.245 | 0.32 | 0.860421899 | 0.345698998 | 0.206441925 | 0.999652205 | 0.977313184 | 0.988482695 | 0.799097568 |
| LOC10230 | High | A0A087W | LBH domain-containing 1 OS=Mus musculus OX=10090 GN=Lbhd1 PE=4 SV=1 | 0 | 36.367 | 46 | 6 | 12 | 6 | 259 | 28.7 | 4.35 | 0.925 | 0.771 | 1.2 | -0.11 | 0.11 | -0.38 | -0.1266667 | 0.245 | 0.26 | 0.752757682 | 0.376558205 | 0.295744147 | 0.999652205 | 0.977313184 | 0.988482695 | 0.894411787 |
| Zcchc11 | High | B2RX14 | Terminal uridylyltransferase 4 OS=Mus musculus OX=10090 GN=Tut4 PE=1 SV=2 | 0.001 | 4.299 | 1 | 2 | 3 | 2 | 1644 | 184.5 | 8.19 | 0.939 | 0.769 | 1.22 | -0.09 | 0.09 | -0.38 | -0.1266667 | 0.235 | 0.29 | 0.846221419 | 0.373596752 | 0.260091982 | 0.999652205 | 0.977313184 | 0.988482695 | 0.85616924 |
| Prkra | High | Q9WTX2 | Interferon-inducible double-stranded RNA-dependent protein kinase activator A OS=Mus musculu | 0 | 20.282 | 15 | 4 | 4 | 4 | 313 | 34.3 | 8.43 | 0.954 | 0.766 | 1.245 | -0.07 | 0.07 | -0.39 | -0.13 | 0.23 | 0.32 | 0.740299357 | 0.343497443 | 0.137090953 | 0.999652205 | 0.977313184 | 0.988482695 | 0.700189449 |
| | High | A0A087W | MAGUK p55 subfamily member 4 OS=Mus musculus OX=10090 GN=Mpp4 PE=1 SV=1 | 0.001 | 7.108 | 6 | 3 | 3 | 3 | 591 | 66.8 | 6.42 | 0.965 | 0.757 | 1.275 | -0.05 | 0.05 | -0.4 | -0.1333333 | 0.225 | 0.35 | 0.818760877 | 0.348151047 | 0.123580697 | 0.999652205 | 0.977313184 | 0.988482695 | 0.677192798 |
| Psmd1 | High | Q3TXS7 | 26S proteasome non-ATPase regulatory subunit 1 OS=Mus musculus OX=10090 GN=Psmd1 PE=1 | 0 | 37.793 | 11 | 10 | 12 | 10 | 953 | 105.7 | 5.39 | 0.949 | 0.766 | 1.239 | -0.07 | 0.07 | -0.38 | -0.1266667 | 0.225 | 0.31 | 0.898898248 | 0.366917254 | 0.138516675 | 0.999652205 | 0.977313184 | 0.988482695 | 0.701851981 |
| Ttll12 | High | Q3UDE2 | Tubulin--tyrosine ligase-like protein 12 OS=Mus musculus OX=10090 GN=Ttll12 PE=1 SV=1 | 0 | 10.544 | 4 | 2 | 2 | 2 | 639 | 74 | 5.63 | 0.956 | 0.764 | 1.252 | -0.06 | 0.06 | -0.39 | -0.13 | 0.225 | 0.32 | 0.850548057 | 0.361382726 | 0.158900709 | 0.999652205 | 0.977313184 | 0.988482695 | 0.733684585 |
| Tlk2 | Medium | O55047 | Serine/threonine-protein kinase tousled-like 2 OS=Mus musculus OX=10090 GN=Tlk2 PE=1 SV=2 | 0.01 | 2.672 | 1 | 1 | 1 | 1 | 718 | 82.2 | 8.27 | 0.958 | 0.774 | 1.237 | -0.06 | 0.06 | -0.37 | -0.1233333 | 0.215 | 0.31 | 0.805380172 | 0.379262277 | 0.2157784 | 0.999652205 | 0.977313184 | 0.988482695 | 0.804011919 |
| | High | D3YWR3 | Adhesion G protein-coupled receptor L3 OS=Mus musculus OX=10090 GN=Adgrl3 PE=1 SV=1 | 0 | 11.008 | 3 | 4 | 7 | 4 | 1268 | 141.9 | 7.01 | 0.986 | 0.758 | 1.3 | -0.02 | 0.02 | -0.4 | -0.1333333 | 0.21 | 0.38 | 0.907862239 | 0.350292582 | 0.127212278 | 0.999652205 | 0.977313184 | 0.988482695 | 0.678506823 |
| Ehd4 | High | Q9EQP2 | EH domain-containing protein 4 OS=Mus musculus OX=10090 GN=Ehd4 PE=1 SV=1 | 0 | 79.155 | 25 | 14 | 27 | 12 | 541 | 61.4 | 6.76 | 0.977 | 0.769 | 1.271 | -0.03 | 0.03 | -0.38 | -0.1266667 | 0.205 | 0.35 | 0.940750314 | 0.372931956 | 0.100278879 | 0.999652205 | 0.977313184 | 0.988482695 | 0.609169746 |
| Rragb | High | Q6NTA4 | Ras-related GTP-binding protein B OS=Mus musculus OX=10090 GN=Rragb PE=1 SV=1 | 0 | 14.151 | 12 | 4 | 5 | 4 | 374 | 43.2 | 6.38 | 1.01 | 0.755 | 1.338 | 0.01 | -0.01 | -0.41 | -0.1366667 | 0.2 | 0.42 | 0.957972752 | 0.342996846 | 0.079969974 | 0.999652205 | 0.977313184 | 0.988482695 | 0.544164616 |
| Ccbl2; Kya | High | Q71RI9 | Kynurenine--oxoglutarate transaminase 3 OS=Mus musculus OX=10090 GN=Kyat3 PE=1 SV=1 | 0.006 | 3.416 | 2 | 1 | 1 | 1 | 455 | 51.1 | 8.37 | 0.968 | 0.784 | 1.236 | -0.05 | 0.05 | -0.35 | -0.1166667 | 0.2 | 0.31 | 0.759866217 | 0.380206261 | 0.162289324 | 0.999652205 | 0.977313184 | 0.988482695 | 0.733684585 |
| Snap47 | High | Q8R570 | Synaptosomal-associated protein 47 OS=Mus musculus OX=10090 GN=Snap47 PE=1 SV=1 | 0 | 32.056 | 21 | 6 | 7 | 6 | 413 | 46.5 | 5.76 | 1.003 | 0.761 | 1.318 | 0 | 0 | -0.39 | -0.13 | 0.195 | 0.4 | 0.997934994 | 0.355965017 | 0.0879149 | 0.999652205 | 0.977313184 | 0.988482695 | 0.573625007 |
| Eif2s2 | Medium | Q99L45 | Eukaryotic translation initiation factor 2 subunit 2 OS=Mus musculus OX=10090 GN=Eif2s2 PE=1 S | 0.011 | 2.49 | 3 | 1 | 1 | 1 | 331 | 38.1 | 5.8 | 0.987 | 0.776 | 1.271 | -0.02 | 0.02 | -0.36 | -0.12 | 0.19 | 0.35 | 0.90449056 | 0.383060538 | 0.16182354 | 0.999652205 | 0.977313184 | 0.988482695 | 0.733684585 |
| Nf1 | High | Q04690 | Neurofibromin OS=Mus musculus OX=10090 GN=Nf1 PE=1 SV=1 | 0 | 13.322 | 2 | 4 | 4 | 4 | 2841 | 319.4 | 7.39 | 1 | 0.78 | 1.282 | 0 | 0 | -0.36 | -0.12 | 0.18 | 0.36 | 0.917004572 | 0.380892732 | 0.1554497 | 0.999652205 | 0.977313184 | 0.988482695 | 0.730581221 |
| Rabgef1 | High | Q9JM13 | Rab5 GDP/GTP exchange factor OS=Mus musculus OX=10090 GN=Rabgef1 PE=1 SV=1 | 0 | 9.544 | 7 | 3 | 3 | 3 | 491 | 56.8 | 6.79 | 1.026 | 0.765 | 1.342 | 0.04 | -0.04 | -0.39 | -0.13 | 0.175 | 0.42 | 0.993671407 | 0.342401537 | 0.053291078 | 0.999652205 | 0.977313184 | 0.988482695 | 0.441363476 |
| Cnbp | Medium | A0A0N4S | Cellular nucleic acid-binding protein OS=Mus musculus OX=10090 GN=Cnbp PE=1 SV=1 | 0.029 | 1.917 | 5 | 1 | 1 | 1 | 171 | 18.8 | 7.71 | 1.018 | 0.771 | 1.321 | 0.03 | -0.03 | -0.38 | -0.1266667 | 0.175 | 0.4 | 0.999171034 | 0.355672548 | 0.165506277 | 0.999652205 | 0.977313184 | 0.988482695 | 0.736602876 |
| BC030336 | High | Q8C784 | Modulator of smoothened protein OS=Mus musculus OX=10090 GN=Mosmo PE=2 SV=2 | 0.005 | 3.612 | 5 | 1 | 2 | 1 | 167 | 18.2 | 8.28 | 1.012 | 0.778 | 1.3 | 0.02 | -0.02 | -0.36 | -0.12 | 0.17 | 0.38 | 0.976029316 | 0.369274167 | 0.199077756 | 0.999652205 | 0.977313184 | 0.988482695 | 0.78697088 |
| Atic | High | Q9CWJ9 | Bifunctional purine biosynthesis protein ATIC OS=Mus musculus OX=10090 GN=Atic PE=1 SV=2 | 0 | 24.989 | 10 | 5 | 6 | 5 | 592 | 64.2 | 6.76 | 1.066 | 0.767 | 1.39 | 0.09 | -0.09 | -0.38 | -0.1266667 | 0.145 | 0.47 | 0.854489513 | 0.348717287 | 0.048437482 | 0.999652205 | 0.977313184 | 0.988482695 | 0.42237786 |
| Arhgap12 | Medium | S4R248 | Rho GTPase-activating protein 12 OS=Mus musculus OX=10090 GN=Arhgap12 PE=1 SV=1 | 0.018 | 2.117 | 1 | 1 | 1 | 1 | 791 | 90 | 8.41 | 1.068 | 0.772 | 1.383 | 0.09 | -0.09 | -0.37 | -0.1233333 | 0.14 | 0.47 | 0.857282007 | 0.356899144 | 0.075176385 | 0.999652205 | 0.977313184 | 0.988482695 | 0.520184986 |
| Map2k7 | High | Q8CE90 | Dual specificity mitogen-activated protein kinase kinase 7 OS=Mus musculus OX=10090 GN=Map | 0.001 | 4.613 | 4 | 2 | 2 | 2 | 535 | 59.3 | 9.11 | 1.106 | 0.767 | 1.441 | 0.15 | -0.15 | -0.38 | -0.1266667 | 0.115 | 0.53 | 0.745298308 | 0.360162036 | 0.033449058 | 0.999652205 | 0.977313184 | 0.988482695 | 0.357917689 |
| Mcl1 | Medium | P97287 | Induced myeloid leukemia cell differentiation protein Mcl-1 homolog OS=Mus musculus OX=1009 | 0.018 | 2.097 | 2 | 1 | 1 | 1 | 331 | 35.2 | 6.2 | 0.898 | 1.369 | 0.656 | -0.15 | 0.15 | 0.45 | 0.15 | -0.15 | -0.61 | 0.659580046 | 0.376517341 | 0.033970359 | 0.999652205 | 0.977313184 | 0.988482695 | 0.362803439 |
| Tubb2a | High | Q7TMM9 | Tubulin beta-2A chain OS=Mus musculus OX=10090 GN=Tubb2a PE=1 SV=1 | 0 | 344.44 | 73 | 25 | 181 | 1 | 445 | 49.9 | 4.89 | 0.996 | 1.37 | 0.727 | -0.01 | 0.01 | 0.45 | 0.15 | -0.22 | -0.46 | 0.991669133 | 0.37846446 | 0.050369331 | 0.999652205 | 0.977313184 | 0.988482695 | 0.427566646 |
| Arsb | High | A0A0R4J1 | Arylsulfatase B OS=Mus musculus OX=10090 GN=Arsb PE=1 SV=1 | 0.008 | 2.987 | 2 | 1 | 1 | 1 | 534 | 59.7 | 7.14 | 0.984 | 1.375 | 0.716 | -0.02 | 0.02 | 0.46 | 0.15333333 | -0.22 | -0.48 | 0.915239375 | 0.359995059 | 0.063243798 | 0.999652205 | 0.977313184 | 0.988482695 | 0.48303291 |
| Fam3c | High | G5E911 | Protein FAM3C OS=Mus musculus OX=10090 GN=Fam3c PE=1 SV=1 | 0 | 23.1 | 16 | 4 | 8 | 4 | 247 | 27 | 8.66 | 1.003 | 1.39 | 0.722 | 0 | 0 | 0.47 | 0.15666667 | -0.235 | -0.47 | 0.948392546 | 0.355665901 | 0.043681125 | 0.999652205 | 0.977313184 | 0.988482695 | 0.404826557 |
| Tapt1 | High | Q4VBD2 | Transmembrane anterior posterior transformation protein 1 OS=Mus musculus OX=10090 GN=Ta | 0.001 | 5.407 | 3 | 1 | 1 | 1 | 564 | 63.9 | 8.22 | 1.027 | 1.384 | 0.742 | 0.04 | -0.04 | 0.47 | 0.15666667 | -0.255 | -0.43 | 0.954140526 | 0.360323933 | 0.089268499 | 0.999652205 | 0.977313184 | 0.988482695 | 0.5746627 |
| Slc27a1 | High | Q60714 | Long-chain fatty acid transport protein 1 OS=Mus musculus OX=10090 GN=Slc27a1 PE=1 SV=1 | 0 | 11.729 | 3 | 2 | 3 | 2 | 646 | 71.2 | 8.38 | 1.052 | 1.366 | 0.77 | 0.07 | -0.07 | 0.45 | 0.15 | -0.26 | -0.38 | 0.864271834 | 0.383568997 | 0.147905934 | 0.999652205 | 0.977313184 | 0.988482695 | 0.712464409 |
| | High | E9PUJ3 | Endosome/lysosome-associated apoptosis and autophagy regulator family member 2 OS=Mus m | 0.001 | 5.602 | 3 | 2 | 2 | 2 | 580 | 64.3 | 5.92 | 1.054 | 1.352 | 0.78 | 0.08 | -0.08 | 0.44 | 0.14666667 | -0.26 | -0.36 | 0.883330621 | 0.38397487 | 0.186985278 | 0.999652205 | 0.977313184 | 0.988482695 | 0.77374646 |
| Cat | High | P24270 | Catalase OS=Mus musculus OX=10090 GN=Cat PE=1 SV=4 | 0 | 16.571 | 10 | 4 | 4 | 4 | 527 | 59.8 | 7.88 | 1.068 | 1.366 | 0.782 | 0.09 | -0.09 | 0.45 | 0.15 | -0.27 | -0.36 | 0.868285758 | 0.381727535 | 0.175819201 | 0.999652205 | 0.977313184 | 0.988482695 | 0.745702164 |
| Calr | High | P14211 | Calreticulin OS=Mus musculus OX=10090 GN=Calr PE=1 SV=1 | 0 | 71.936 | 45 | 17 | 31 | 17 | 416 | 48 | 4.49 | 1.062 | 1.375 | 0.772 | 0.09 | -0.09 | 0.46 | 0.15333333 | -0.275 | -0.37 | 0.836676313 | 0.372985578 | 0.117975221 | 0.999652205 | 0.977313184 | 0.988482695 | 0.656888843 |
| Fkbp8 | High | O35465 | Peptidyl-prolyl cis-trans isomerase FKBP8 OS=Mus musculus OX=10090 GN=Fkbp8 PE=1 SV=2 | 0 | 11.357 | 14 | 4 | 4 | 4 | 402 | 43.5 | 5.16 | 1.054 | 1.392 | 0.757 | 0.08 | -0.08 | 0.48 | 0.16 | -0.28 | -0.4 | 0.838935636 | 0.353278243 | 0.124687573 | 0.999652205 | 0.977313184 | 0.988482695 | 0.678506823 |
| Mylk | High | Q6PDN3 | Myosin light chain kinase, smooth muscle OS=Mus musculus OX=10090 GN=Mylk PE=1 SV=3 | 0 | 11.502 | 1 | 2 | 2 | 2 | 1941 | 212.8 | 6.25 | 1.068 | 1.387 | 0.77 | 0.1 | -0.1 | 0.47 | 0.15666667 | -0.285 | -0.38 | 0.798159721 | 0.358476689 | 0.132528901 | 0.999652205 | 0.977313184 | 0.988482695 | 0.695121227 |
| BC017158 | High | A0A0R4J0 | RUS family member 1 OS=Mus musculus OX=10090 GN=Rusf1 PE=1 SV=1 | 0.001 | 4.387 | 4 | 2 | 2 | 2 | 466 | 50.5 | 7.56 | 1.09 | 1.367 | 0.798 | 0.12 | -0.12 | 0.45 | 0.15 | -0.285 | -0.33 | 0.815302697 | 0.383213143 | 0.199771799 | 0.999652205 | 0.977313184 | 0.988482695 | 0.78697088 |
| Slc19a1 | High | P41438 | Reduced folate transporter OS=Mus musculus OX=10090 GN=Slc19a1 PE=1 SV=1 | 0 | 24.673 | 11 | 4 | 5 | 4 | 512 | 58.1 | 8.62 | 1.089 | 1.373 | 0.793 | 0.12 | -0.12 | 0.46 | 0.15333333 | -0.29 | -0.33 | 0.794332533 | 0.363370915 | 0.192284141 | 0.999652205 | 0.977313184 | 0.988482695 | 0.784495541 |
| Srprb | Low | P47758 | Signal recognition particle receptor subunit beta OS=Mus musculus OX=10090 GN=Srprb PE=1 SV | 0.078 | 1.223 | 3 | 1 | 1 | 1 | 269 | 29.6 | 9.28 | 1.1 | 1.399 | 0.786 | 0.14 | -0.14 | 0.48 | 0.16 | -0.31 | -0.35 | 0.736934422 | 0.343202619 | 0.172417635 | 0.999652205 | 0.977313184 | 0.988482695 | 0.738844686 |
| Ndufa10 | High | Q99LC3 | NADH dehydrogenase [ubiquinone] 1 alpha subcomplex subunit 10, mitochondrial OS=Mus muscu | 0 | 25.902 | 21 | 6 | 7 | 6 | 355 | 40.6 | 7.78 | 1.153 | 1.366 | 0.844 | 0.2 | -0.2 | 0.45 | 0.15 | -0.325 | -0.25 | 0.638957499 | 0.383396242 | 0.297798578 | 0.999652205 | 0.977313184 | 0.988482695 | 0.898684946 |
| | Medium | S4R2U7 | CUGBP Elav-like family member 2 OS=Mus musculus OX=10090 GN=Celf2 PE=1 SV=1 | 0.01 | 2.717 | 2 | 1 | 2 | 1 | 460 | 48.8 | 9.09 | 1.148 | 1.364 | 0.841 | 0.2 | -0.2 | 0.45 | 0.15 | -0.325 | -0.25 | 0.623955056 | 0.376057397 | 0.345979361 | 0.999652205 | 0.977313184 | 0.988482695 | 0.907770836 |
| Vps26b | High | Q8C0E2 | Vacuolar protein sorting-associated protein 26B OS=Mus musculus OX=10090 GN=Vps26b PE=1 S | 0 | 17.652 | 12 | 3 | 5 | 3 | 336 | 39.1 | 7.37 | 1.14 | 1.388 | 0.821 | 0.19 | -0.19 | 0.47 | 0.15666667 | -0.33 | -0.28 | 0.634602634 | 0.357800456 | 0.244883166 | 0.999652205 | 0.977313184 | 0.988482695 | 0.838891386 |
| Ezr | High | P26040 | Ezrin OS=Mus musculus OX=10090 GN=Ezr PE=1 SV=3 | 0 | 416.923 | 66 | 56 | 199 | 42 | 586 | 69.4 | 6.1 | 1.15 | 1.379 | 0.834 | 0.2 | -0.2 | 0.46 | 0.15333333 | -0.33 | -0.26 | 0.636666501 | 0.368083528 | 0.283405211 | 0.999652205 | 0.977313184 | 0.988482695 | 0.886074068 |
| Angel1 | High | Q8VCU0 | Protein angel homolog 1 OS=Mus musculus OX=10090 GN=Angel1 PE=1 SV=2 | 0.001 | 7.896 | 4 | 2 | 2 | 2 | 667 | 75.2 | 4.73 | 1.159 | 1.385 | 0.837 | 0.21 | -0.21 | 0.47 | 0.15666667 | -0.34 | -0.26 | 0.664646544 | 0.365298437 | 0.415928237 | 0.999652205 | 0.977313184 | 0.988482695 | 0.936397673 |
| Slc25a22 | High | Q9D6M3 | Mitochondrial glutamate carrier 1 OS=Mus musculus OX=10090 GN=Slc25a22 PE=1 SV=1 | 0 | 44.87 | 42 | 9 | 14 | 9 | 323 | 34.6 | 9.09 | 1.147 | 1.401 | 0.818 | 0.2 | -0.2 | 0.49 | 0.16333333 | -0.345 | -0.29 | 0.647725069 | 0.342475049 | 0.238339779 | 0.999652205 | 0.977313184 | 0.988482695 | 0.830020494 |
| Tomm40 | High | Q9QYA2 | Mitochondrial import receptor subunit TOM40 homolog OS=Mus musculus OX=10090 GN=Tomm | 0.005 | 3.756 | 2 | 1 | 1 | 1 | 361 | 37.9 | 7.74 | 1.172 | 1.381 | 0.849 | 0.23 | -0.23 | 0.47 | 0.15666667 | -0.35 | -0.24 | 0.594396753 | 0.350562916 | 0.353795793 | 0.999652205 | 0.977313184 | 0.988482695 | 0.911642008 |
| Elmod1 | High | Q3V1U8 | ELMO domain-containing protein 1 OS=Mus musculus OX=10090 GN=Elmod1 PE=1 SV=2 | 0.001 | 5.645 | 6 | 2 | 2 | 2 | 326 | 38 | 8.4 | 1.189 | 1.369 | 0.868 | 0.25 | -0.25 | 0.45 | 0.15 | -0.35 | -0.2 | 0.540694655 | 0.379618275 | 0.433042629 | 0.999652205 | 0.977313184 | 0.988482695 | 0.948052758 |
| Arhgap32 | High | Q811P8 | Rho GTPase-activating protein 32 OS=Mus musculus OX=10090 GN=Arhgap32 PE=1 SV=2 | 0 | 8.848 | 1 | 3 | 3 | 3 | 2089 | 229.6 | 6.86 | 1.202 | 1.383 | 0.869 | 0.26 | -0.26 | 0.47 | 0.15666667 | -0.365 | -0.2 | 0.509425796 | 0.36298494 | 0.450190713 | 0.999652205 | 0.977313184 | 0.988482695 | 0.959449555 |
| Prkcb | High | P68404 | Protein kinase C beta type OS=Mus musculus OX=10090 GN=Prkcb PE=1 SV=4 | 0 | 53.396 | 17 | 11 | 20 | 2 | 671 | 76.7 | 7.01 | 1.213 | 1.378 | 0.881 | 0.28 | -0.28 | 0.46 | 0.15333333 | -0.37 | -0.18 | 0.479975741 | 0.36970143 | 0.445406786 | 0.999652205 | 0.977313184 | 0.988482695 | 0.958608248 |
| 2700060E | High | Q9CQE8 | RNA transcription, translation and transport factor protein OS=Mus musculus OX=10090 GN=RTR | 0 | 16.646 | 26 | 7 | 8 | 7 | 244 | 28.1 | 6.89 | 1.232 | 1.366 | 0.902 | 0.3 | -0.3 | 0.45 | 0.15 | -0.375 | -0.15 | 0.479078589 | 0.383503945 | 0.528324517 | 0.999652205 | 0.977313184 | 0.988482695 | 0.965674043 |
| Phb | High | P67778 | Prohibitin OS=Mus musculus OX=10090 GN=Phb PE=1 SV=1 | 0 | 44.223 | 39 | 10 | 13 | 10 | 272 | 29.8 | 5.76 | 1.238 | 1.367 | 0.905 | 0.31 | -0.31 | 0.45 | 0.15 | -0.38 | -0.14 | 0.485905978 | 0.382178462 | 0.59720971 | 0.999652205 | 0.977313184 | 0.988482695 | 0.977451587 |
| Vdac1 | High | Q60932 | Voltage-dependent anion-selective channel protein 1 OS=Mus musculus OX=10090 GN=Vdac1 PE | 0 | 279.311 | 69 | 18 | 100 | 17 | 296 | 32.3 | 8.43 | 1.244 | 1.387 | 0.897 | 0.31 | -0.31 | 0.47 | 0.15666667 | -0.39 | -0.16 | 0.461071239 | 0.358508286 | 0.545985169 | 0.999652205 | 0.977313184 | 0.988482695 | 0.96694628 |
| Bbip1 | High | J3QMK2 | BBSome-interacting protein 1 OS=Mus musculus OX=10090 GN=Bbip1 PE=1 SV=1 | 0 | 38.179 | 55 | 3 | 17 | 3 | 67 | 7.6 | 8.76 | 1.254 | 1.378 | 0.91 | 0.33 | -0.33 | 0.46 | 0.15333333 | -0.395 | -0.14 | 0.450688576 | 0.369402104 | 0.63531921 | 0.999652205 | 0.977313184 | 0.988482695 | 0.983562471 |
| Car14 | High | Q9WVT6 | Carbonic anhydrase 14 OS=Mus musculus OX=10090 GN=Ca14 PE=1 SV=1 | 0 | 91.343 | 25 | 10 | 40 | 10 | 337 | 37.5 | 6.35 | 1.259 | 1.374 | 0.916 | 0.33 | -0.33 | 0.46 | 0.15333333 | -0.395 | -0.13 | 0.437188268 | 0.374629769 | 0.642932271 | 0.999652205 | 0.977313184 | 0.988482695 | 0.985010103 |
| Sfxn3 | High | Q91V61 | Sideroflexin-3 OS=Mus musculus OX=10090 GN=Sfxn3 PE=1 SV=1 | 0 | 33.524 | 25 | 7 | 11 | 7 | 321 | 35.4 | 9.51 | 1.263 | 1.368 | 0.924 | 0.34 | -0.34 | 0.45 | 0.15 | -0.395 | -0.11 | 0.444602163 | 0.381998986 | 0.652761583 | 0.999652205 | 0.977313184 | 0.988482695 | 0.985010103 |
| Vat1l | High | Q80TB8 | Synaptic vesicle membrane protein VAT-1 homolog-like OS=Mus musculus OX=10090 GN=Vat1l P | 0 | 27.169 | 15 | 6 | 8 | 6 | 417 | 45.8 | 5.06 | 1.006 | 1.364 | 0.737 | 0.01 | -0.01 | 0.45 | 0.15 | -0.23 | -0.44 | 0.98118149 | 0.386740418 | 0.073716814 | 0.999652205 | 0.979427969 | 0.989540087 | 0.519259015 |
| Dbi | High | P31786 | Acyl-CoA-binding protein OS=Mus musculus OX=10090 GN=Dbi PE=1 SV=2 | 0 | 47.657 | 61 | 5 | 12 | 5 | 87 | 10 | 8.82 | 1.177 | 1.362 | 0.864 | 0.24 | -0.24 | 0.45 | 0.15 | -0.345 | -0.21 | 0.609468015 | 0.388468034 | 0.423722614 | 0.999652205 | 0.98165177 | 0.990651988 | 0.940543427 |
| Casp3 | Medium | P70677 | Caspase-3 OS=Mus musculus OX=10090 GN=Casp3 PE=1 SV=1 | 0.01 | 2.605 | 5 | 1 | 1 | 1 | 277 | 31.5 | 6.92 | 1.176 | 0.787 | 1.495 | 0.23 | -0.23 | -0.35 | -0.1166667 | 0.06 | 0.58 | 0.584042637 | 0.389493697 | 0.023416002 | 0.999652205 | 0.982598494 | 0.99112535 | 0.279347924 |
| Tm9sf4 | High | Q8BH24 | Transmembrane 9 superfamily member 4 OS=Mus musculus OX=10090 GN=Tm9sf4 PE=1 SV=1 | 0 | 28.872 | 10 | 7 | 8 | 7 | 643 | 74.6 | 7.23 | 0.962 | 1.361 | 0.707 | -0.06 | 0.06 | 0.44 | 0.14666667 | -0.19 | -0.5 | 0.828799576 | 0.3899113 | 0.04011831 | 0.999652205 | 0.983018282 | 0.991335243 | 0.387364155 |
| Ruvbl2 | High | Q9WTM5 | RuvB-like 2 OS=Mus musculus OX=10090 GN=Ruvbl2 PE=1 SV=3 | 0 | 21.687 | 9 | 5 | 7 | 5 | 463 | 51.1 | 5.64 | 1.043 | 0.777 | 1.342 | 0.06 | -0.06 | -0.36 | -0.12 | 0.15 | 0.42 | 0.912431516 | 0.390469948 | 0.063318921 | 0.999652205 | 0.983542228 | 0.991597217 | 0.48303291 |
| Lypla1 | High | P97823 | Acyl-protein thioesterase 1 OS=Mus musculus OX=10090 GN=Lypla1 PE=1 SV=1 | 0 | 22.907 | 25 | 5 | 9 | 5 | 230 | 24.7 | 6.62 | 1.031 | 0.777 | 1.326 | 0.04 | -0.04 | -0.36 | -0.12 | 0.16 | 0.41 | 0.957478934 | 0.390699193 | 0.085851134 | 0.999652205 | 0.983677761 | 0.991664983 | 0.566314481 |
| | High | A0A171KX | Protein arginine N-methyltransferase 1 OS=Mus musculus OX=10090 GN=Prmt1 PE=1 SV=1 | 0.001 | 5.377 | 6 | 2 | 2 | 2 | 318 | 36.5 | 6.19 | 0.88 | 0.781 | 1.126 | -0.18 | 0.18 | -0.36 | -0.12 | 0.27 | 0.17 | 0.594349702 | 0.39406008 | 0.493697668 | 0.999652205 | 0.983884835 | 0.99176852 | 0.964378027 |
| Cap1 | High | P40124 | Adenylyl cyclase-associated protein 1 OS=Mus musculus OX=10090 GN=Cap1 PE=1 SV=4 | 0 | 73.321 | 35 | 13 | 18 | 12 | 474 | 51.5 | 7.52 | 0.909 | 0.779 | 1.167 | -0.14 | 0.14 | -0.36 | -0.12 | 0.25 | 0.22 | 0.804939209 | 0.393692356 | 0.269261472 | 0.999652205 | 0.983884835 | 0.99176852 | 0.871243909 |
| Akr7a5 | High | Q8CG76 | Aflatoxin B1 aldehyde reductase member 2 OS=Mus musculus OX=10090 GN=Akr7a2 PE=1 SV=3 | 0 | 15.958 | 10 | 2 | 3 | 2 | 367 | 40.6 | 8.12 | 0.94 | 0.79 | 1.189 | -0.09 | 0.09 | -0.34 | -0.1133333 | 0.215 | 0.25 | 0.696354863 | 0.393269987 | 0.240107797 | 0.999652205 | 0.983884835 | 0.99176852 | 0.832581581 |
| Eps15l1 | High | Q60902 | Epidermal growth factor receptor substrate 15-like 1 OS=Mus musculus OX=10090 GN=Eps15l1 P | 0 | 11.862 | 4 | 3 | 3 | 3 | 907 | 99.2 | 5.02 | 1.02 | 1.358 | 0.751 | 0.03 | -0.03 | 0.44 | 0.14666667 | -0.235 | -0.41 | 0.982181064 | 0.393429516 | 0.102200091 | 0.999652205 | 0.983884835 | 0.99176852 | 0.610114517 |
| Kcnj13 | High | P86046 | Inward rectifier potassium channel 13 OS=Mus musculus OX=10090 GN=Kcnj13 PE=3 SV=1 | 0 | 37.297 | 31 | 8 | 20 | 8 | 360 | 40.5 | 6.52 | 1.202 | 1.358 | 0.885 | 0.27 | -0.27 | 0.44 | 0.14666667 | -0.355 | -0.18 | 0.533534246 | 0.394245803 | 0.494109867 | 0.999652205 | 0.983884835 | 0.99176852 | 0.964378027 |
| Gnat1 | High | P20612 | Guanine nucleotide-binding protein G(t) subunit alpha-1 OS=Mus musculus OX=10090 GN=Gnat1 | 0 | 844.738 | 73 | 39 | 2412 | 26 | 350 | 39.9 | 5.62 | 1.194 | 1.356 | 0.881 | 0.26 | -0.26 | 0.44 | 0.14666667 | -0.35 | -0.18 | 0.548098306 | 0.396367294 | 0.472805542 | 0.999652205 | 0.985399792 | 0.992525999 | 0.961211268 |
| Dnajc27 | High | Q8CFP6 | DnaJ homolog subfamily C member 27 OS=Mus musculus OX=10090 GN=Dnajc27 PE=1 SV=1 | 0.005 | 3.525 | 4 | 1 | 1 | 1 | 273 | 30.8 | 8.47 | 0.844 | 0.785 | 1.075 | -0.24 | 0.24 | -0.35 | -0.1166667 | 0.295 | 0.1 | 0.494634033 | 0.399483731 | 0.664086062 | 0.999652205 | 0.985440071 | 0.992546138 | 0.986647506 |
| Pak2 | High | Q8CIN4 | Serine/threonine-protein kinase PAK 2 OS=Mus musculus OX=10090 GN=Pak2 PE=1 SV=1 | 0 | 35.759 | 19 | 10 | 14 | 5 | 524 | 57.9 | 5.77 | 0.882 | 0.781 | 1.129 | -0.18 | 0.18 | -0.36 | -0.12 | 0.27 | 0.18 | 0.721434484 | 0.399092315 | 0.371724596 | 0.999652205 | 0.985440071 | 0.992546138 | 0.920164724 |
| | Medium | A0A2I3BQ | Leucine-rich repeat and calponin homology domain-containing protein 1 OS=Mus musculus OX=1 | 0.01 | 2.571 | 1 | 1 | 1 | 1 | 682 | 76 | 6.09 | 0.882 | 0.785 | 1.123 | -0.18 | 0.18 | -0.35 | -0.1166667 | 0.265 | 0.17 | 0.615275101 | 0.399451665 | 0.498139871 | 0.999652205 | 0.985440071 | 0.992546138 | 0.964378027 |
| Atxn10 | Medium | P28658 | Ataxin-10 OS=Mus musculus OX=10090 GN=Atxn10 PE=1 SV=2 | 0.011 | 2.399 | 2 | 1 | 1 | 1 | 475 | 53.7 | 5.25 | 0.887 | 0.783 | 1.132 | -0.17 | 0.17 | -0.35 | -0.1166667 | 0.26 | 0.18 | 0.573637717 | 0.398036916 | 0.472797217 | 0.999652205 | 0.985440071 | 0.992546138 | 0.961211268 |
| Sec24c | High | G3X972 | Sec24-related gene family, member C (S. cerevisiae) OS=Mus musculus OX=10090 GN=Sec24c PE | 0 | 41.755 | 10 | 9 | 12 | 9 | 1096 | 118.5 | 6.84 | 0.984 | 0.78 | 1.261 | -0.02 | 0.02 | -0.36 | -0.12 | 0.19 | 0.33 | 0.924724023 | 0.397386712 | 0.162364728 | 0.999652205 | 0.985440071 | 0.992546138 | 0.733684585 |
| Basp1 | High | Q91XV3 | Brain acid soluble protein 1 OS=Mus musculus OX=10090 GN=Basp1 PE=1 SV=3 | 0 | 332.356 | 94 | 29 | 167 | 29 | 226 | 22.1 | 4.51 | 1.175 | 1.355 | 0.867 | 0.23 | -0.23 | 0.44 | 0.14666667 | -0.335 | -0.21 | 0.584833351 | 0.397589682 | 0.413775451 | 0.999652205 | 0.985440071 | 0.992546138 | 0.936397673 |
| Cul5 | High | Q9D5V5 | Cullin-5 OS=Mus musculus OX=10090 GN=Cul5 PE=1 SV=3 | 0 | 20.386 | 7 | 5 | 7 | 5 | 780 | 90.9 | 7.81 | 0.826 | 0.783 | 1.055 | -0.28 | 0.28 | -0.35 | -0.1166667 | 0.315 | 0.08 | 0.478745189 | 0.40357673 | 0.761989258 | 0.999652205 | 0.9855335 | 0.992592853 | 0.992535481 |
| Vps33a | High | Q9D2N9 | Vacuolar protein sorting-associated protein 33A OS=Mus musculus OX=10090 GN=Vps33a PE=1 S | 0.001 | 6.973 | 2 | 1 | 2 | 1 | 598 | 67.5 | 7.08 | 0.826 | 0.789 | 1.047 | -0.28 | 0.28 | -0.34 | -0.1133333 | 0.31 | 0.07 | 0.435087224 | 0.408404753 | 0.772237648 | 0.999652205 | 0.9855335 | 0.992592853 | 0.992535481 |
| Tbcb | High | Q9D1E6 | Tubulin-folding cofactor B OS=Mus musculus OX=10090 GN=Tbcb PE=1 SV=2 | 0.001 | 4.256 | 5 | 1 | 1 | 1 | 244 | 27.4 | 5.24 | 0.823 | 0.796 | 1.034 | -0.28 | 0.28 | -0.33 | -0.11 | 0.305 | 0.05 | 0.430037648 | 0.408540384 | 0.802592493 | 0.999652205 | 0.9855335 | 0.992592853 | 0.992535481 |
| Rnh1 | High | A0A1B0G | Ribonuclease inhibitor OS=Mus musculus OX=10090 GN=Rnh1 PE=1 SV=1 | 0 | 50.311 | 33 | 12 | 15 | 12 | 492 | 53.9 | 5.01 | 0.85 | 0.782 | 1.086 | -0.24 | 0.24 | -0.35 | -0.1166667 | 0.295 | 0.12 | 0.550081441 | 0.400942405 | 0.597949018 | 0.999652205 | 0.9855335 | 0.992592853 | 0.977451587 |
| Dync1h1 | High | Q9JHU4 | Cytoplasmic dynein 1 heavy chain 1 OS=Mus musculus OX=10090 GN=Dync1h1 PE=1 SV=2 | 0 | 618.819 | 35 | 143 | 199 | 143 | 4644 | 531.7 | 6.42 | 0.871 | 0.785 | 1.11 | -0.2 | 0.2 | -0.35 | -0.1166667 | 0.275 | 0.15 | 0.644778476 | 0.407522385 | 0.440696907 | 0.999652205 | 0.9855335 | 0.992592853 | 0.955524965 |
| Elp2 | Medium | Q91WG4 | Elongator complex protein 2 OS=Mus musculus OX=10090 GN=Elp2 PE=1 SV=1 | 0.01 | 2.686 | 1 | 1 | 1 | 1 | 831 | 93 | 5.82 | 0.873 | 0.795 | 1.099 | -0.2 | 0.2 | -0.33 | -0.11 | 0.265 | 0.14 | 0.549510473 | 0.406541784 | 0.524100114 | 0.999652205 | 0.9855335 | 0.992592853 | 0.964378027 |
| Cfl1 | High | P18760 | Cofilin-1 OS=Mus musculus OX=10090 GN=Cfl1 PE=1 SV=3 | 0 | 110.102 | 80 | 18 | 43 | 15 | 166 | 18.5 | 8.09 | 0.887 | 0.783 | 1.133 | -0.17 | 0.17 | -0.35 | -0.1166667 | 0.26 | 0.18 | 0.688908338 | 0.402687786 | 0.365323754 | 0.999652205 | 0.9855335 | 0.992592853 | 0.920124252 |
| Cct7 | High | P80313 | T-complex protein 1 subunit eta OS=Mus musculus OX=10090 GN=Cct7 PE=1 SV=1 | 0 | 99.428 | 45 | 22 | 32 | 22 | 544 | 59.6 | 7.84 | 0.902 | 0.786 | 1.148 | -0.15 | 0.15 | -0.35 | -0.1166667 | 0.25 | 0.2 | 0.730500618 | 0.408761497 | 0.322323896 | 0.999652205 | 0.9855335 | 0.992592853 | 0.905686588 |
| | Medium | F6XXE6 | Amylo-1,6-glucosidase, 4-alpha-glucanotransferase (Fragment) OS=Mus musculus OX=10090 GN | 0.011 | 2.481 | 1 | 1 | 1 | 1 | 830 | 93.5 | 7.47 | 0.918 | 0.789 | 1.163 | -0.12 | 0.12 | -0.34 | -0.1133333 | 0.23 | 0.22 | 0.70935632 | 0.409497273 | 0.388325181 | 0.999652205 | 0.9855335 | 0.992592853 | 0.93064581 |
| Mthfd1 | High | Q922D8 | C-1-tetrahydrofolate synthase, cytoplasmic OS=Mus musculus OX=10090 GN=Mthfd1 PE=1 SV=4 | 0 | 48.916 | 18 | 14 | 16 | 14 | 935 | 101.1 | 7.14 | 0.93 | 0.785 | 1.185 | -0.1 | 0.1 | -0.35 | -0.1166667 | 0.225 | 0.24 | 0.866947043 | 0.407861793 | 0.230254807 | 0.999652205 | 0.9855335 | 0.992592853 | 0.819186996 |
| Psmc6 | High | P62334 | 26S proteasome regulatory subunit 10B OS=Mus musculus OX=10090 GN=Psmc6 PE=1 SV=1 | 0 | 23.97 | 20 | 7 | 7 | 7 | 389 | 44.1 | 7.49 | 0.931 | 0.785 | 1.186 | -0.1 | 0.1 | -0.35 | -0.1166667 | 0.225 | 0.25 | 0.738729527 | 0.40682609 | 0.323978277 | 0.999652205 | 0.9855335 | 0.992592853 | 0.905686588 |
| Pcyt2 | High | Q922E4 | Ethanolamine-phosphate cytidylyltransferase OS=Mus musculus OX=10090 GN=Pcyt2 PE=1 SV=1 | 0.008 | 2.871 | 4 | 1 | 1 | 1 | 404 | 45.2 | 6.58 | 0.954 | 0.789 | 1.209 | -0.07 | 0.07 | -0.34 | -0.1133333 | 0.205 | 0.27 | 0.782803549 | 0.408882462 | 0.292740718 | 0.999652205 | 0.9855335 | 0.992592853 | 0.893646499 |
| Psmc2 | High | Q8BVQ9 | 26S proteasome AAA-ATPase subunit RPT1 OS=Mus musculus OX=10090 GN=Psmc2 PE=1 SV=1 | 0 | 27.807 | 17 | 7 | 8 | 7 | 475 | 52.8 | 6.28 | 0.965 | 0.783 | 1.232 | -0.05 | 0.05 | -0.35 | -0.1166667 | 0.2 | 0.3 | 0.855593844 | 0.40376498 | 0.21557346 | 0.999652205 | 0.9855335 | 0.992592853 | 0.804011919 |
| | High | A0A140LI | 26S proteasome AAA-ATPase subunit RPT3 OS=Mus musculus OX=10090 GN=Psmc4 PE=1 SV=1 | 0 | 20.837 | 18 | 6 | 8 | 6 | 387 | 43.5 | 5.26 | 0.99 | 0.784 | 1.263 | -0.01 | 0.01 | -0.35 | -0.1166667 | 0.18 | 0.34 | 0.921507098 | 0.404473368 | 0.162211087 | 0.999652205 | 0.9855335 | 0.992592853 | 0.733684585 |
| 1300018J1 | Low | Q9DBC0 | Protein adenylyltransferase SelO, mitochondrial OS=Mus musculus OX=10090 GN=Selenoo PE=1 S | 0.053 | 1.438 | 1 | 1 | 1 | 1 | 667 | 74.2 | 5.83 | 1.084 | 0.787 | 1.377 | 0.12 | -0.12 | -0.35 | -0.1166667 | 0.115 | 0.46 | 0.810454271 | 0.408370802 | 0.070453185 | 0.999652205 | 0.9855335 | 0.992592853 | 0.508467979 |
| Eftud1; Efl | High | Q8C0D5 | Elongation factor-like GTPase 1 OS=Mus musculus OX=10090 GN=Efl1 PE=1 SV=1 | 0.001 | 4.481 | 2 | 2 | 2 | 2 | 1127 | 125.7 | 6.16 | 1.085 | 1.354 | 0.801 | 0.12 | -0.12 | 0.44 | 0.14666667 | -0.28 | -0.32 | 0.837899518 | 0.401252429 | 0.323412194 | 0.999652205 | 0.9855335 | 0.992592853 | 0.905686588 |
| | High | A0A1D5RL | Microtubule-associated protein 1S OS=Mus musculus OX=10090 GN=Map1s PE=1 SV=1 | 0 | 12.254 | 4 | 3 | 4 | 3 | 864 | 91 | 7.03 | 1.1 | 1.348 | 0.816 | 0.14 | -0.14 | 0.43 | 0.14333333 | -0.285 | -0.29 | 0.781294301 | 0.406910914 | 0.230022657 | 0.999652205 | 0.9855335 | 0.992592853 | 0.819186996 |
| Gstz1 | High | Q9WVL0 | Maleylacetoacetate isomerase OS=Mus musculus OX=10090 GN=Gstz1 PE=1 SV=1 | 0.001 | 5.399 | 9 | 1 | 1 | 1 | 216 | 24.3 | 7.85 | 1.099 | 1.349 | 0.814 | 0.14 | -0.14 | 0.43 | 0.14333333 | -0.285 | -0.3 | 0.780372553 | 0.40351852 | 0.278633135 | 0.999652205 | 0.9855335 | 0.992592853 | 0.876067403 |
| Hist2h2ac | High | Q64523 | Histone H2A type 2-C OS=Mus musculus OX=10090 GN=H2ac20 PE=1 SV=3 | 0 | 8.043 | 22 | 2 | 3 | 1 | 129 | 14 | 10.9 | 1.135 | 1.349 | 0.842 | 0.18 | -0.18 | 0.43 | 0.14333333 | -0.305 | -0.25 | 0.673775638 | 0.408562805 | 0.350199446 | 0.999652205 | 0.9855335 | 0.992592853 | 0.909466648 |
| Gba2 | Medium | Q69ZF3 | Non-lysosomal glucosylceramidase OS=Mus musculus OX=10090 GN=Gba2 PE=1 SV=2 | 0.035 | 1.771 | 1 | 1 | 1 | 1 | 918 | 103.2 | 5.59 | 1.144 | 1.346 | 0.85 | 0.19 | -0.19 | 0.43 | 0.14333333 | -0.31 | -0.23 | 0.650707893 | 0.409275555 | 0.342809251 | 0.999652205 | 0.9855335 | 0.992592853 | 0.907770836 |
| Hdlbp | High | Q8VDJ3 | Vigilin OS=Mus musculus OX=10090 GN=Hdlbp PE=1 SV=1 | 0 | 8.774 | 3 | 4 | 4 | 4 | 1268 | 141.7 | 6.87 | 1.231 | 1.351 | 0.911 | 0.3 | -0.3 | 0.43 | 0.14333333 | -0.365 | -0.13 | 0.477429748 | 0.402794774 | 0.61269677 | 0.999652205 | 0.9855335 | 0.992592853 | 0.97826833 |
| Ndufa7 | High | Q9Z1P6 | NADH dehydrogenase [ubiquinone] 1 alpha subcomplex subunit 7 OS=Mus musculus OX=10090 G | 0 | 41.544 | 67 | 9 | 16 | 9 | 113 | 12.6 | 10.17 | 1.23 | 1.353 | 0.909 | 0.3 | -0.3 | 0.44 | 0.14666667 | -0.37 | -0.14 | 0.505540837 | 0.400582159 | 0.582921682 | 0.999652205 | 0.9855335 | 0.992592853 | 0.973095093 |
| Ndufa2 | High | Q9CQ75 | NADH dehydrogenase [ubiquinone] 1 alpha subcomplex subunit 2 OS=Mus musculus OX=10090 G | 0 | 21.164 | 42 | 5 | 8 | 5 | 99 | 10.9 | 9.99 | 1.252 | 1.345 | 0.931 | 0.32 | -0.32 | 0.43 | 0.14333333 | -0.375 | -0.1 | 0.449517937 | 0.410331888 | 0.653729908 | 0.999652205 | 0.986182817 | 0.992917511 | 0.985010103 |
| Gm3244; | High | Q9CQC7 | NADH dehydrogenase [ubiquinone] 1 beta subcomplex subunit 4 OS=Mus musculus OX=10090 GN | 0 | 11.448 | 46 | 4 | 4 | 4 | 129 | 15.1 | 9.89 | 1.173 | 1.342 | 0.874 | 0.23 | -0.23 | 0.42 | 0.14 | -0.325 | -0.19 | 0.595806434 | 0.414213159 | 0.462256052 | 0.999652205 | 0.992518455 | 0.99608533 | 0.959744396 |
| Slc16a1 | High | P53986 | Monocarboxylate transporter 1 OS=Mus musculus OX=10090 GN=Slc16a1 PE=1 SV=1 | 0 | 126.156 | 22 | 16 | 64 | 16 | 493 | 53.2 | 7.47 | 1.19 | 1.342 | 0.887 | 0.25 | -0.25 | 0.42 | 0.14 | -0.335 | -0.17 | 0.555893819 | 0.414538779 | 0.501195178 | 0.999652205 | 0.992874386 | 0.996263295 | 0.964378027 |
| Vcp | High | Q01853 | Transitional endoplasmic reticulum ATPase OS=Mus musculus OX=10090 GN=Vcp PE=1 SV=4 | 0 | 199.443 | 58 | 37 | 63 | 36 | 806 | 89.3 | 5.26 | 0.831 | 0.794 | 1.048 | -0.27 | 0.27 | -0.33 | -0.11 | 0.3 | 0.07 | 0.536159957 | 0.42653252 | 0.689483623 | 0.999652205 | 0.99399649 | 0.996824347 | 0.991285312 |
| Myh10 | High | Q5SV64 | Myosin-10 OS=Mus musculus OX=10090 GN=Myh10 PE=1 SV=1 | 0 | 100.849 | 15 | 25 | 32 | 18 | 2007 | 232.3 | 5.54 | 0.833 | 0.794 | 1.049 | -0.26 | 0.26 | -0.33 | -0.11 | 0.295 | 0.07 | 0.541364575 | 0.428068302 | 0.68246205 | 0.999652205 | 0.99399649 | 0.996824347 | 0.991285312 |
| | Low | V9GXD9 | Synergin gamma OS=Mus musculus OX=10090 GN=Synrg PE=1 SV=1 | 0.052 | 1.471 | 1 | 1 | 1 | 1 | 1216 | 129.9 | 5.02 | 0.862 | 0.796 | 1.083 | -0.21 | 0.21 | -0.33 | -0.11 | 0.27 | 0.12 | 0.558846573 | 0.42557533 | 0.619088567 | 0.999652205 | 0.99399649 | 0.996824347 | 0.97826833 |
| Ppme1 | High | Q8BVQ5 | Protein phosphatase methylesterase 1 OS=Mus musculus OX=10090 GN=Ppme1 PE=1 SV=5 | 0 | 26.847 | 17 | 5 | 9 | 5 | 386 | 42.2 | 5.97 | 0.877 | 0.791 | 1.109 | -0.19 | 0.19 | -0.34 | -0.1133333 | 0.265 | 0.15 | 0.561145316 | 0.420719326 | 0.537753837 | 0.999652205 | 0.99399649 | 0.996824347 | 0.96694628 |
| Cops4 | High | O88544 | COP9 signalosome complex subunit 4 OS=Mus musculus OX=10090 GN=Cops4 PE=1 SV=1 | 0 | 37.491 | 26 | 8 | 9 | 8 | 406 | 46.3 | 5.83 | 0.879 | 0.791 | 1.112 | -0.19 | 0.19 | -0.34 | -0.1133333 | 0.265 | 0.15 | 0.575958229 | 0.420232889 | 0.56616343 | 0.999652205 | 0.99399649 | 0.996824347 | 0.972910028 |
| Igbp1 | High | Q61249 | Immunoglobulin-binding protein 1 OS=Mus musculus OX=10090 GN=Igbp1 PE=1 SV=1 | 0 | 26.987 | 26 | 6 | 7 | 6 | 340 | 38.9 | 6.18 | 0.872 | 0.797 | 1.095 | -0.2 | 0.2 | -0.33 | -0.11 | 0.265 | 0.13 | 0.616845675 | 0.433412396 | 0.589928804 | 0.999652205 | 0.99399649 | 0.996824347 | 0.974007893 |
| Glra3 | High | G5E811 | Glycine receptor subunit alpha-3 OS=Mus musculus OX=10090 GN=Glra3 PE=1 SV=1 | 0 | 4.718 | 2 | 1 | 1 | 1 | 480 | 55.6 | 8.72 | 0.871 | 0.801 | 1.087 | -0.2 | 0.2 | -0.32 | -0.1066667 | 0.26 | 0.12 | 0.470568004 | 0.419547049 | 0.539763958 | 0.999652205 | 0.99399649 | 0.996824347 | 0.96694628 |
| Slc1a3 | High | P56564 | Excitatory amino acid transporter 1 OS=Mus musculus OX=10090 GN=Slc1a3 PE=1 SV=2 | 0 | 98.608 | 19 | 8 | 34 | 8 | 543 | 59.6 | 8.4 | 0.894 | 0.79 | 1.132 | -0.16 | 0.16 | -0.34 | -0.1133333 | 0.25 | 0.18 | 0.709022872 | 0.418321408 | 0.368367443 | 0.999652205 | 0.99399649 | 0.996824347 | 0.920164724 |
| Sgta | High | Q8BJU0 | Small glutamine-rich tetratricopeptide repeat-containing protein alpha OS=Mus musculus OX=10 | 0 | 39.568 | 26 | 8 | 11 | 8 | 315 | 34.3 | 5.06 | 0.902 | 0.795 | 1.134 | -0.15 | 0.15 | -0.33 | -0.11 | 0.24 | 0.18 | 0.741520532 | 0.430431895 | 0.361957686 | 0.999652205 | 0.99399649 | 0.996824347 | 0.917494008 |
| Zscan29 | Low | A2ARU6 | Zinc finger SCAN domains 29 OS=Mus musculus OX=10090 GN=Zscan29 PE=1 SV=1 | 0.082 | 1.189 | 1 | 1 | 1 | 1 | 548 | 63 | 8.51 | 0.903 | 0.801 | 1.127 | -0.15 | 0.15 | -0.32 | -0.1066667 | 0.235 | 0.17 | 0.556825683 | 0.415649671 | 0.404791094 | 0.999652205 | 0.99399649 | 0.996824347 | 0.934402497 |
| Cct6a | High | Q52KG9 | Chaperonin containing Tcp1, subunit 6a (Zeta) OS=Mus musculus OX=10090 GN=Cct6a PE=2 SV=1 | 0 | 106.024 | 42 | 21 | 35 | 17 | 531 | 58 | 6.9 | 0.922 | 0.791 | 1.166 | -0.12 | 0.12 | -0.34 | -0.1133333 | 0.23 | 0.22 | 0.7869338 | 0.420852968 | 0.276884925 | 0.999652205 | 0.99399649 | 0.996824347 | 0.876067403 |
| Appl1 | High | Q8K3H0 | DCC-interacting protein 13-alpha OS=Mus musculus OX=10090 GN=Appl1 PE=1 SV=1 | 0 | 13.091 | 5 | 3 | 3 | 3 | 707 | 79.3 | 5.41 | 0.932 | 0.792 | 1.177 | -0.1 | 0.1 | -0.34 | -0.1133333 | 0.22 | 0.24 | 0.733634417 | 0.416550554 | 0.334007366 | 0.999652205 | 0.99399649 | 0.996824347 | 0.906329766 |
| Mapk1 | High | P63085 | Mitogen-activated protein kinase 1 OS=Mus musculus OX=10090 GN=Mapk1 PE=1 SV=3 | 0 | 33.091 | 32 | 10 | 13 | 7 | 358 | 41.2 | 6.98 | 0.988 | 0.792 | 1.247 | -0.02 | 0.02 | -0.34 | -0.1133333 | 0.18 | 0.32 | 0.975684974 | 0.424060241 | 0.149533955 | 0.999652205 | 0.99399649 | 0.996824347 | 0.717225739 |
| Psmd6 | High | Q99JI4 | 26S proteasome non-ATPase regulatory subunit 6 OS=Mus musculus OX=10090 GN=Psmd6 PE=1 | 0.001 | 5.422 | 5 | 2 | 2 | 2 | 389 | 45.5 | 5.52 | 0.999 | 0.789 | 1.266 | 0 | 0 | -0.34 | -0.1133333 | 0.17 | 0.34 | 0.948828489 | 0.416643407 | 0.133147886 | 0.999652205 | 0.99399649 | 0.996824347 | 0.695121227 |
| Scoc | Medium | Q78YZ6 | Short coiled-coil protein OS=Mus musculus OX=10090 GN=Scoc PE=1 SV=1 | 0.014 | 2.236 | 8 | 1 | 1 | 1 | 125 | 14.1 | 4.68 | 0.995 | 0.806 | 1.235 | -0.01 | 0.01 | -0.31 | -0.1033333 | 0.16 | 0.3 | 0.907589627 | 0.42173015 | 0.193909549 | 0.999652205 | 0.99399649 | 0.996824347 | 0.784517504 |
| Vcpip1 | Medium | Q8CDG3 | Deubiquitinating protein VCPIP1 OS=Mus musculus OX=10090 GN=Vcpip1 PE=1 SV=1 | 0.034 | 1.814 | 1 | 1 | 1 | 1 | 1220 | 134.4 | 7.17 | 1.011 | 0.796 | 1.27 | 0.02 | -0.02 | -0.33 | -0.11 | 0.155 | 0.35 | 0.959143192 | 0.42157778 | 0.179084321 | 0.999652205 | 0.99399649 | 0.996824347 | 0.756118818 |

| | | | | | | | | | | | | | | | | | | | | | | | | | | | | |
|---|---|---|---|---|---|---|---|---|---|---|---|---|---|---|---|---|---|---|---|---|---|---|---|---|---|---|---|---|
| Lypla2 | High | Q9WTL7 | Acyl-protein thioesterase 2 OS=Mus musculus OX=10090 GN=Lypla2 PE=1 SV=1 | 0 | 39.004 | 37 | 7 | 12 | 7 | 231 | 24.8 | 7.23 | 1.031 | 0.797 | 1.295 | 0.04 | -0.04 | -0.33 | -0.11 | 0.145 | 0.37 | 0.986614304 | 0.433483498 | 0.117544309 | 0.999652205 | 0.99399649 | 0.996824347 | 0.655140097 |
| | High | A0A338P7 | Ubiquitin-conjugating enzyme E2 L3 OS=Mus musculus OX=10090 GN=Ube2l3 PE=1 SV=1 | 0 | 44.333 | 67 | 7 | 9 | 7 | 122 | 14.1 | 8.92 | 1.071 | 0.797 | 1.344 | 0.1 | -0.1 | -0.33 | -0.11 | 0.115 | 0.43 | 0.844379886 | 0.433923754 | 0.064751496 | 0.999652205 | 0.99399649 | 0.996824347 | 0.489961721 |
| Commd3 | High | Q63829 | COMM domain-containing protein 3 OS=Mus musculus OX=10090 GN=Commd3 PE=1 SV=1 | 0.001 | 7.246 | 11 | 2 | 2 | 2 | 195 | 22 | 5.59 | 1.074 | 0.801 | 1.341 | 0.1 | -0.1 | -0.32 | -0.1066667 | 0.11 | 0.42 | 0.831883476 | 0.422079274 | 0.083449944 | 0.999652205 | 0.99399649 | 0.996824347 | 0.557770483 |
| Gpsm2 | High | Q8VDU0 | G-protein-signaling modulator 2 OS=Mus musculus OX=10090 GN=Gpsm2 PE=1 SV=2 | 0 | 18.156 | 11 | 7 | 7 | 7 | 679 | 75.5 | 6.95 | 1.127 | 0.79 | 1.426 | 0.17 | -0.17 | -0.34 | -0.1133333 | 0.085 | 0.51 | 0.641689255 | 0.418150273 | 0.043834763 | 0.999652205 | 0.99399649 | 0.996824347 | 0.404911892 |
| Arhgef11 | High | Q68FM7 | Rho guanine nucleotide exchange factor (GEF) 11 OS=Mus musculus OX=10090 GN=Arhgef11 PE= | 0.001 | 5.448 | 1 | 2 | 2 | 1 | 1552 | 171.7 | 5.58 | 1.238 | 0.799 | 1.55 | 0.31 | -0.31 | -0.32 | -0.1066667 | 0.005 | 0.63 | 0.493547688 | 0.420243729 | 0.034971913 | 0.999652205 | 0.99399649 | 0.996824347 | 0.367894023 |
| Por | High | Q05DV1 | NADPH--hemoprotein reductase OS=Mus musculus OX=10090 GN=Por PE=1 SV=1 | 0 | 32.338 | 20 | 11 | 11 | 11 | 649 | 72.9 | 5.58 | 0.999 | 1.329 | 0.752 | 0 | 0 | 0.41 | 0.13666667 | -0.205 | -0.41 | 0.977062549 | 0.430889386 | 0.094449672 | 0.999652205 | 0.99399649 | 0.996824347 | 0.584524624 |
| Grm8 | Low | Q05BD6 | Glutamate receptor, metabotropic 8 OS=Mus musculus OX=10090 GN=Grm8 PE=1 SV=1 | 0.062 | 1.315 | 1 | 1 | 1 | 1 | 908 | 101.6 | 8.15 | 1.01 | 1.327 | 0.761 | 0.01 | -0.01 | 0.41 | 0.13666667 | -0.21 | -0.39 | 0.959051322 | 0.425779688 | 0.145059102 | 0.999652205 | 0.99399649 | 0.996824347 | 0.706643254 |
| Pcbp3 | High | P57722 | Poly(rC)-binding protein 3 OS=Mus musculus OX=10090 GN=Pcbp3 PE=1 SV=3 | 0 | 71.262 | 39 | 10 | 25 | 4 | 371 | 39.3 | 7.52 | 1.006 | 1.334 | 0.754 | 0.01 | -0.01 | 0.42 | 0.14 | -0.215 | -0.41 | 0.875844389 | 0.417411356 | 0.080274978 | 0.999652205 | 0.99399649 | 0.996824347 | 0.544917434 |
| Vapa | High | Q9WV55 | Vesicle-associated membrane protein-associated protein A OS=Mus musculus OX=10090 GN=Va | 0 | 55.609 | 48 | 12 | 21 | 11 | 249 | 27.8 | 8.4 | 1.014 | 1.328 | 0.764 | 0.02 | -0.02 | 0.41 | 0.13666667 | -0.215 | -0.39 | 0.959782345 | 0.432862427 | 0.101500521 | 0.999652205 | 0.99399649 | 0.996824347 | 0.610114517 |
| Hyou1 | High | Q9JKR6 | Hypoxia up-regulated protein 1 OS=Mus musculus OX=10090 GN=Hyou1 PE=1 SV=1 | 0 | 31.069 | 15 | 12 | 13 | 12 | 999 | 111.1 | 5.19 | 1.016 | 1.331 | 0.763 | 0.02 | -0.02 | 0.41 | 0.13666667 | -0.215 | -0.39 | 0.990872968 | 0.428362939 | 0.113997173 | 0.999652205 | 0.99399649 | 0.996824347 | 0.644336844 |
| Ganab | High | Q8BHN3 | Neutral alpha-glucosidase AB OS=Mus musculus OX=10090 GN=Ganab PE=1 SV=1 | 0 | 48.627 | 16 | 12 | 15 | 12 | 944 | 106.8 | 6.06 | 1.015 | 1.337 | 0.759 | 0.02 | -0.02 | 0.42 | 0.14 | -0.22 | -0.4 | 0.940604086 | 0.42083126 | 0.094005103 | 0.999652205 | 0.99399649 | 0.996824347 | 0.583399835 |
| Dnajb14 | High | Q149L6 | DnaJ homolog subfamily B member 14 OS=Mus musculus OX=10090 GN=Dnajb14 PE=2 SV=1 | 0.001 | 5.969 | 4 | 1 | 1 | 1 | 379 | 42.3 | 8.59 | 1.036 | 1.322 | 0.784 | 0.05 | -0.05 | 0.4 | 0.13333333 | -0.225 | -0.35 | 0.930535463 | 0.42364385 | 0.197947642 | 0.999652205 | 0.99399649 | 0.996824347 | 0.78697088 |
| Sec22b | High | O08547 | Vesicle-trafficking protein SEC22b OS=Mus musculus OX=10090 GN=Sec22b PE=1 SV=3 | 0 | 58.934 | 47 | 9 | 16 | 9 | 215 | 24.7 | 8.51 | 1.036 | 1.332 | 0.778 | 0.05 | -0.05 | 0.41 | 0.13666667 | -0.23 | -0.36 | 0.93216326 | 0.427193432 | 0.138559764 | 0.999652205 | 0.99399649 | 0.996824347 | 0.701851981 |
| C2cd2l | High | Q80X80 | Phospholipid transfer protein C2CD2L OS=Mus musculus OX=10090 GN=C2cd2l PE=1 SV=3 | 0 | 50.602 | 25 | 12 | 14 | 12 | 706 | 76.3 | 7.21 | 1.11 | 1.337 | 0.83 | 0.15 | -0.15 | 0.42 | 0.14 | -0.285 | -0.27 | 0.762398741 | 0.420611269 | 0.253565657 | 0.999652205 | 0.99399649 | 0.996824347 | 0.852875009 |
| Tnr | High | Q8BYI9 | Tenascin-R OS=Mus musculus OX=10090 GN=Tnr PE=1 SV=2 | 0 | 20.328 | 6 | 6 | 6 | 6 | 1358 | 149.5 | 4.94 | 1.117 | 1.335 | 0.837 | 0.16 | -0.16 | 0.42 | 0.14 | -0.29 | -0.26 | 0.697481806 | 0.422931415 | 0.307442593 | 0.999652205 | 0.99399649 | 0.996824347 | 0.904295409 |
| Serbp1 | High | Q9CY58 | Plasminogen activator inhibitor 1 RNA-binding protein OS=Mus musculus OX=10090 GN=Serbp1 F | 0 | 88.108 | 39 | 14 | 23 | 14 | 407 | 44.7 | 8.54 | 1.135 | 1.333 | 0.852 | 0.18 | -0.18 | 0.41 | 0.13666667 | -0.295 | -0.23 | 0.667345086 | 0.426435447 | 0.350244806 | 0.999652205 | 0.99399649 | 0.996824347 | 0.909466648 |
| | High | F6VBV0 | Regulating synaptic membrane exocytosis protein 1 OS=Mus musculus OX=10090 GN=Rims1 PE= | 0.005 | 3.585 | 1 | 2 | 2 | 1 | 1666 | 185.7 | 9.63 | 1.141 | 1.34 | 0.852 | 0.19 | -0.19 | 0.42 | 0.14 | -0.305 | -0.23 | 0.685838097 | 0.416783956 | 0.320274544 | 0.999652205 | 0.99399649 | 0.996824347 | 0.905392802 |
| D10Jhu81e | High | Q9D172 | Glutamine amidotransferase-like class 1 domain-containing protein 3A, mitochondrial OS=Mus m | 0 | 37.757 | 47 | 8 | 10 | 8 | 266 | 28.1 | 8.78 | 1.217 | 1.333 | 0.913 | 0.28 | -0.28 | 0.41 | 0.13666667 | -0.345 | -0.13 | 0.478351783 | 0.426592993 | 0.628740383 | 0.999652205 | 0.99399649 | 0.996824347 | 0.981444134 |
| Rac1 | High | Q3TLP8 | Ras-related C3 botulinum toxin substrate 1 OS=Mus musculus OX=10090 GN=Rac1 PE=1 SV=1 | 0 | 77.215 | 38 | 10 | 42 | 9 | 211 | 23.4 | 8.69 | 0.795 | 0.801 | 0.992 | -0.33 | 0.33 | -0.32 | -0.1066667 | 0.325 | -0.01 | 0.442919992 | 0.444333368 | 0.959080118 | 0.999652205 | 0.994630833 | 0.997141519 | 0.992902921 |
| | High | A0A1W2P | Myosin light polypeptide 6 OS=Mus musculus OX=10090 GN=Myl6 PE=1 SV=1 | 0 | 37.287 | 51 | 7 | 12 | 7 | 152 | 17 | 4.5 | 0.805 | 0.799 | 1.007 | -0.31 | 0.31 | -0.32 | -0.1066667 | 0.315 | 0.01 | 0.493435877 | 0.439342811 | 0.883146322 | 0.999652205 | 0.994630833 | 0.997141519 | 0.992535481 |
| Hprt | High | P00493 | Hypoxanthine-guanine phosphoribosyltransferase OS=Mus musculus OX=10090 GN=Hprt1 PE=1 S | 0 | 97.741 | 73 | 12 | 29 | 12 | 218 | 24.6 | 6.68 | 0.847 | 0.8 | 1.059 | -0.24 | 0.24 | -0.32 | -0.1066667 | 0.28 | 0.08 | 0.579380683 | 0.441027665 | 0.637687454 | 0.999652205 | 0.994630833 | 0.997141519 | 0.983597794 |
| Bpnt1 | High | Q9Z0S1 | 3'(2'),5'-bisphosphate nucleotidase 1 OS=Mus musculus OX=10090 GN=Bpnt1 PE=1 SV=2 | 0 | 61.129 | 44 | 10 | 20 | 10 | 308 | 33.2 | 5.82 | 0.866 | 0.799 | 1.083 | -0.21 | 0.21 | -0.32 | -0.1066667 | 0.265 | 0.12 | 0.630996114 | 0.439972953 | 0.537668218 | 0.999652205 | 0.994630833 | 0.997141519 | 0.96694628 |
| | High | E9QAI5 | Aspartate carbamoyltransferase OS=Mus musculus OX=10090 GN=Cad PE=1 SV=1 | 0 | 13.604 | 2 | 5 | 5 | 5 | 2162 | 236.1 | 6.49 | 0.862 | 0.8 | 1.077 | -0.21 | 0.21 | -0.32 | -0.1066667 | 0.265 | 0.11 | 0.623081034 | 0.440889513 | 0.658903928 | 0.999652205 | 0.994630833 | 0.997141519 | 0.985010103 |
| Ap3s2 | Low | Q8BSZ2 | AP-3 complex subunit sigma-2 OS=Mus musculus OX=10090 GN=Ap3s2 PE=1 SV=1 | 0.075 | 1.247 | 4 | 1 | 1 | 1 | 193 | 22 | 5.22 | 0.851 | 0.812 | 1.048 | -0.23 | 0.23 | -0.3 | -0.1 | 0.265 | 0.07 | 0.447091848 | 0.443951676 | 0.713258119 | 0.999652205 | 0.994630833 | 0.997141519 | 0.992535481 |
| Cct2 | High | P80314 | T-complex protein 1 subunit beta OS=Mus musculus OX=10090 GN=Cct2 PE=1 SV=4 | 0 | 130.991 | 48 | 24 | 44 | 24 | 535 | 57.4 | 6.4 | 0.893 | 0.8 | 1.117 | -0.16 | 0.16 | -0.32 | -0.1066667 | 0.24 | 0.16 | 0.706964668 | 0.440302038 | 0.415003955 | 0.999652205 | 0.994630833 | 0.997141519 | 0.936397673 |
| | High | A0A494B9 | E3 ubiquitin-protein ligase OS=Mus musculus OX=10090 GN=Nedd4l PE=1 SV=1 | 0 | 23.96 | 9 | 8 | 8 | 6 | 912 | 105 | 5.58 | 0.897 | 0.803 | 1.117 | -0.16 | 0.16 | -0.32 | -0.1066667 | 0.24 | 0.16 | 0.632483204 | 0.448786274 | 0.543733794 | 0.999652205 | 0.994630833 | 0.997141519 | 0.96694628 |
| Ppcdc | High | A0A1L1ST | Phosphopantothenoylcysteine decarboxylase OS=Mus musculus OX=10090 GN=Ppcdc PE=1 SV=1 | 0.005 | 3.785 | 15 | 2 | 2 | 2 | 146 | 15.8 | 5.27 | 0.892 | 0.813 | 1.098 | -0.16 | 0.16 | -0.3 | -0.1 | 0.23 | 0.13 | 0.605076734 | 0.444678311 | 0.523807599 | 0.999652205 | 0.994630833 | 0.997141519 | 0.964378027 |
| Psmd2 | High | Q8VDM4 | 26S proteasome non-ATPase regulatory subunit 2 OS=Mus musculus OX=10090 GN=Psmd2 PE=1 | 0 | 52.054 | 16 | 11 | 14 | 11 | 908 | 100.1 | 5.17 | 0.955 | 0.798 | 1.197 | -0.07 | 0.07 | -0.33 | -0.11 | 0.2 | 0.26 | 0.838652498 | 0.436622443 | 0.262576965 | 0.999652205 | 0.994630833 | 0.997141519 | 0.859470827 |
| Pkm | High | P52480 | Pyruvate kinase PKM OS=Mus musculus OX=10090 GN=Pkm PE=1 SV=4 | 0 | 866.279 | 81 | 52 | 826 | 51 | 531 | 57.8 | 7.47 | 0.973 | 0.799 | 1.217 | -0.04 | 0.04 | -0.32 | -0.1066667 | 0.18 | 0.28 | 0.927788806 | 0.439549319 | 0.172485747 | 0.999652205 | 0.994630833 | 0.997141519 | 0.738844686 |
| Aarsd1 | High | Q3THG9 | Alanyl-tRNA editing protein Aarsd1 OS=Mus musculus OX=10090 GN=Aarsd1 PE=1 SV=2 | 0 | 18.016 | 14 | 4 | 5 | 4 | 412 | 44.9 | 6.42 | 0.972 | 0.801 | 1.213 | -0.04 | 0.04 | -0.32 | -0.1066667 | 0.18 | 0.28 | 0.904155261 | 0.444042896 | 0.227660879 | 0.999652205 | 0.994630833 | 0.997141519 | 0.816808422 |
| Rps27l | High | Q6ZWY3 | 40S ribosomal protein S27-like OS=Mus musculus OX=10090 GN=Rps27l PE=1 SV=3 | 0.001 | 4.305 | 23 | 2 | 4 | 1 | 84 | 9.5 | 9.45 | 1.069 | 0.812 | 1.316 | 0.1 | -0.1 | -0.3 | -0.1 | 0.1 | 0.4 | 0.868523021 | 0.445469469 | 0.180727642 | 0.999652205 | 0.994630833 | 0.997141519 | 0.759056098 |
| Lrsam1 | Low | Q80ZI6 | E3 ubiquitin-protein ligase LRSAM1 OS=Mus musculus OX=10090 GN=Lrsam1 PE=1 SV=1 | 0.054 | 1.416 | 1 | 1 | 1 | 1 | 727 | 83.9 | 6.14 | 1.082 | 0.814 | 1.329 | 0.11 | -0.11 | -0.3 | -0.1 | 0.095 | 0.41 | 0.808185967 | 0.449137848 | 0.081551792 | 0.999652205 | 0.994630833 | 0.997141519 | 0.550253785 |
| Pdap1 | High | Q3UHX2 | 28 kDa heat- and acid-stable phosphoprotein OS=Mus musculus OX=10090 GN=Pdap1 PE=1 SV=1 | 0 | 23.721 | 35 | 5 | 9 | 5 | 181 | 20.6 | 7.39 | 1.121 | 0.798 | 1.404 | 0.16 | -0.16 | -0.33 | -0.11 | 0.085 | 0.49 | 0.725448493 | 0.436963765 | 0.044256133 | 0.999652205 | 0.994630833 | 0.997141519 | 0.405968973 |
| | Low | F6YTS6 | Epoxide hydrolase 1 (Fragment) OS=Mus musculus OX=10090 GN=Ephx1 PE=1 SV=2 | 0.078 | 1.223 | 4 | 1 | 1 | 1 | 247 | 28 | 8.48 | 0.913 | 1.313 | 0.695 | -0.13 | 0.13 | 0.39 | 0.13 | -0.13 | -0.52 | 0.661926206 | 0.439563191 | 0.040277029 | 0.999652205 | 0.994630833 | 0.997141519 | 0.387364155 |
| Mxra7 | High | Q9CZH7 | Matrix-remodeling-associated protein 7 OS=Mus musculus OX=10090 GN=Mxra7 PE=1 SV=2 | 0 | 28.615 | 44 | 4 | 7 | 2 | 178 | 19.4 | 4.26 | 0.915 | 1.322 | 0.692 | -0.13 | 0.13 | 0.4 | 0.13333333 | -0.135 | -0.53 | 0.682438256 | 0.439040554 | 0.023849587 | 0.999652205 | 0.994630833 | 0.997141519 | 0.283916426 |
| Rgs7bp | Medium | Q8BQP9 | Regulator of G-protein signaling 7-binding protein OS=Mus musculus OX=10090 GN=Rgs7bp PE=1 | 0.032 | 1.848 | 3 | 1 | 1 | 1 | 257 | 29 | 8.46 | 0.927 | 1.323 | 0.701 | -0.11 | 0.11 | 0.4 | 0.13333333 | -0.145 | -0.51 | 0.698429566 | 0.438154433 | 0.074994603 | 0.999652205 | 0.994630833 | 0.997141519 | 0.520184986 |
| Actr3b | High | Q641P0 | Actin-related protein 3B OS=Mus musculus OX=10090 GN=Actr3b PE=1 SV=1 | 0 | 17.179 | 8 | 3 | 4 | 1 | 418 | 47.5 | 6.02 | 0.967 | 1.306 | 0.741 | -0.05 | 0.05 | 0.38 | 0.12666667 | -0.165 | -0.43 | 0.867053274 | 0.449153784 | 0.090311832 | 0.999652205 | 0.994630833 | 0.997141519 | 0.575430045 |
| Ppib | High | P24369 | Peptidyl-prolyl cis-trans isomerase B OS=Mus musculus OX=10090 GN=Ppib PE=1 SV=2 | 0 | 24.432 | 31 | 7 | 8 | 7 | 216 | 23.7 | 9.55 | 0.994 | 1.317 | 0.755 | -0.01 | 0.01 | 0.4 | 0.13333333 | -0.195 | -0.41 | 0.92685619 | 0.448591324 | 0.088930046 | 0.999652205 | 0.994630833 | 0.997141519 | 0.5746627 |
| Tmem106 | Medium | Q80X71 | Transmembrane protein 106B OS=Mus musculus OX=10090 GN=Tmem106b PE=1 SV=1 | 0.028 | 1.929 | 3 | 1 | 1 | 1 | 275 | 31.2 | 6.68 | 1.048 | 1.308 | 0.801 | 0.07 | -0.07 | 0.39 | 0.13 | -0.23 | -0.32 | 0.913273349 | 0.445541842 | 0.224671684 | 0.999652205 | 0.994630833 | 0.997141519 | 0.812731698 |
| | High | H3BJZ9 | 3-ketoacyl-CoA thiolase A, peroxisomal OS=Mus musculus OX=10090 GN=Acaa1a PE=1 SV=1 | 0 | 29.062 | 24 | 5 | 5 | 5 | 331 | 34.5 | 8.54 | 1.055 | 1.325 | 0.796 | 0.08 | -0.08 | 0.41 | 0.13666667 | -0.245 | -0.33 | 0.863156763 | 0.436527757 | 0.200483194 | 0.999652205 | 0.994630833 | 0.997141519 | 0.78697088 |
| Polr2b | Medium | Q8CFI7 | DNA-directed RNA polymerase II subunit RPB2 OS=Mus musculus OX=10090 GN=Polr2b PE=1 SV= | 0.017 | 2.134 | 1 | 1 | 1 | 1 | 1174 | 133.8 | 6.87 | 1.085 | 1.312 | 0.827 | 0.12 | -0.12 | 0.39 | 0.13 | -0.255 | -0.27 | 0.792575068 | 0.439572096 | 0.316630454 | 0.999652205 | 0.994630833 | 0.997141519 | 0.904943958 |
| Slc30a9 | High | F8WHL1 | Solute carrier family 30 member 9 OS=Mus musculus OX=10090 GN=Slc30a9 PE=1 SV=1 | 0.001 | 7.694 | 4 | 2 | 2 | 2 | 547 | 60.6 | 8.7 | 1.1 | 1.321 | 0.833 | 0.14 | -0.14 | 0.4 | 0.13333333 | -0.27 | -0.26 | 0.757236419 | 0.4403937 | 0.32830295 | 0.999652205 | 0.994630833 | 0.997141519 | 0.905686588 |
| | Medium | A0A0U1R | Hormone-sensitive lipase (Fragment) OS=Mus musculus OX=10090 GN=Lipe PE=1 SV=1 | 0.037 | 1.734 | 4 | 1 | 1 | 1 | 180 | 19.7 | 6.52 | 1.11 | 1.31 | 0.848 | 0.15 | -0.15 | 0.39 | 0.13 | -0.27 | -0.24 | 0.749136098 | 0.444002521 | 0.351005141 | 0.999652205 | 0.994630833 | 0.997141519 | 0.909466648 |
| Kdelr3 | Medium | Q8R1L4 | ER lumen protein-retaining receptor 3 OS=Mus musculus OX=10090 GN=Kdelr3 PE=1 SV=1 | 0.01 | 2.733 | 4 | 1 | 1 | 1 | 214 | 25.1 | 9.06 | 1.115 | 1.318 | 0.846 | 0.16 | -0.16 | 0.4 | 0.13333333 | -0.28 | -0.24 | 0.697627538 | 0.445334986 | 0.341727044 | 0.999652205 | 0.994630833 | 0.997141519 | 0.907770836 |
| Fxr2 | High | Q6P5B5 | Fragile X mental retardation syndrome-related protein 2 OS=Mus musculus OX=10090 GN=Fxr2 P | 0 | 23.593 | 10 | 5 | 5 | 4 | 674 | 74.2 | 6.23 | 1.135 | 1.317 | 0.862 | 0.18 | -0.18 | 0.4 | 0.13333333 | -0.29 | -0.21 | 0.608546884 | 0.447966608 | 0.409400595 | 0.999652205 | 0.994630833 | 0.997141519 | 0.936397673 |
| Cngb1 | High | E1AZ71 | Cyclic nucleotide-gated channel beta 1 OS=Mus musculus OX=10090 GN=Cngb1 PE=1 SV=1 | 0 | 743.501 | 52 | 62 | 459 | 57 | 1325 | 148.6 | 4.53 | 1.138 | 1.323 | 0.861 | 0.19 | -0.19 | 0.4 | 0.13333333 | -0.295 | -0.22 | 0.660902817 | 0.440390662 | 0.385848451 | 0.999652205 | 0.994630833 | 0.997141519 | 0.929303068 |
| Rom1 | High | P32958 | Rod outer segment membrane protein 1 OS=Mus musculus OX=10090 GN=Rom1 PE=1 SV=1 | 0 | 185.876 | 38 | 12 | 229 | 12 | 351 | 37.2 | 5.55 | 1.187 | 1.323 | 0.897 | 0.25 | -0.25 | 0.4 | 0.13333333 | -0.325 | -0.16 | 0.56214377 | 0.440186446 | 0.549945465 | 0.999652205 | 0.994630833 | 0.997141519 | 0.96694628 |
| Bcam | High | Q9R069 | Basal cell adhesion molecule OS=Mus musculus OX=10090 GN=Bcam PE=1 SV=1 | 0 | 11.851 | 6 | 3 | 3 | 3 | 622 | 67.6 | 6.25 | 1.201 | 1.321 | 0.909 | 0.26 | -0.26 | 0.4 | 0.13333333 | -0.33 | -0.14 | 0.528155715 | 0.443935868 | 0.621115047 | 0.999652205 | 0.994630833 | 0.997141519 | 0.97826833 |
| B3galt6 | High | Q91Z92 | Beta-1,3-galactosyltransferase 6 OS=Mus musculus OX=10090 GN=B3galt6 PE=2 SV=1 | 0.001 | 5.867 | 3 | 1 | 2 | 1 | 325 | 37 | 8.82 | 1.212 | 1.318 | 0.919 | 0.28 | -0.28 | 0.4 | 0.13333333 | -0.34 | -0.12 | 0.502443123 | 0.445946857 | 0.650418221 | 0.999652205 | 0.994630833 | 0.997141519 | 0.985010103 |
| Pcyt1b | High | Q811Q9 | Choline-phosphate cytidylyltransferase B OS=Mus musculus OX=10090 GN=Pcyt1b PE=1 SV=2 | 0 | 8.868 | 7 | 3 | 3 | 2 | 369 | 41.9 | 6.4 | 1.241 | 1.324 | 0.937 | 0.31 | -0.31 | 0.41 | 0.13666667 | -0.36 | -0.09 | 0.450594392 | 0.437881149 | 0.740433539 | 0.999652205 | 0.994630833 | 0.997141519 | 0.992535481 |
| Spna2; Spt | High | A3KGU9 | Spectrin alpha chain, non-erythrocytic 1 OS=Mus musculus OX=10090 GN=Sptan1 PE=1 SV=2 | 0 | 749.459 | 59 | 129 | 210 | 129 | 2498 | 287.4 | 5.35 | 0.793 | 0.812 | 0.977 | -0.33 | 0.33 | -0.3 | -0.1 | 0.315 | -0.03 | 0.437346971 | 0.468098258 | 0.960655653 | 0.999652205 | 0.995559182 | 0.997605693 | 0.99314725 |
| Ddah1 | High | Q9CWS0 | N(G),N(G)-dimethylarginine dimethylaminohydrolase 1 OS=Mus musculus OX=10090 GN=Ddah1 | 0 | 37.32 | 36 | 11 | 14 | 10 | 285 | 31.4 | 5.97 | 0.801 | 0.818 | 0.98 | -0.32 | 0.32 | -0.29 | -0.0966667 | 0.305 | -0.03 | 0.481167974 | 0.482068955 | 0.973180932 | 0.999652205 | 0.995559182 | 0.997605693 | 0.996848549 |
| | High | A0A5F8M | Septin-4 OS=Mus musculus OX=10090 GN=Septin4 PE=1 SV=1 | 0.001 | 7.164 | 3 | 3 | 4 | 2 | 971 | 110 | 8.35 | 0.83 | 0.808 | 1.028 | -0.27 | 0.27 | -0.31 | -0.1033333 | 0.29 | 0.04 | 0.516957563 | 0.458539907 | 0.869996967 | 0.999652205 | 0.995559182 | 0.997605693 | 0.992535481 |
| Add3 | High | Q9QYB5 | Gamma-adducin OS=Mus musculus OX=10090 GN=Add3 PE=1 SV=2 | 0.001 | 5.118 | 3 | 3 | 3 | 3 | 706 | 78.7 | 5.95 | 0.82 | 0.821 | 1 | -0.29 | 0.29 | -0.29 | -0.0966667 | 0.29 | 0 | 0.473323034 | 0.488690833 | 0.95275343 | 0.999652205 | 0.995559182 | 0.997605693 | 0.992535481 |
| Spnb2; Spt | High | Q62261 | Spectrin beta chain, non-erythrocytic 1 OS=Mus musculus OX=10090 GN=Sptbn1 PE=1 SV=2 | 0 | 390.137 | 41 | 86 | 127 | 76 | 2363 | 274.1 | 5.58 | 0.816 | 0.824 | 0.99 | -0.29 | 0.29 | -0.28 | -0.0933333 | 0.285 | -0.01 | 0.496337344 | 0.497327012 | 0.970604325 | 0.999652205 | 0.995559182 | 0.997605693 | 0.995824053 |
| Wdr1 | High | O88342 | WD repeat-containing protein 1 OS=Mus musculus OX=10090 GN=Wdr1 PE=1 SV=3 | 0 | 100.236 | 33 | 15 | 33 | 15 | 606 | 66.4 | 6.6 | 0.843 | 0.805 | 1.048 | -0.25 | 0.25 | -0.31 | -0.1033333 | 0.28 | 0.07 | 0.567382933 | 0.451524637 | 0.6889801 | 0.999652205 | 0.995559182 | 0.997605693 | 0.991285312 |
| S100a13 | High | P97352 | Protein S100-A13 OS=Mus musculus OX=10090 GN=S100a13 PE=1 SV=1 | 0.001 | 6.327 | 19 | 2 | 2 | 2 | 98 | 11.2 | 6.13 | 0.808 | 0.851 | 0.949 | -0.31 | 0.31 | -0.23 | -0.0766667 | 0.27 | -0.07 | 0.440585301 | 0.560670213 | 0.767689452 | 0.999652205 | 0.995559182 | 0.997605693 | 0.992535481 |
| Tsn | High | Q62348 | Translin OS=Mus musculus OX=10090 GN=Tsn PE=1 SV=1 | 0 | 29.809 | 31 | 8 | 11 | 8 | 228 | 26.2 | 6.44 | 0.854 | 0.808 | 1.057 | -0.23 | 0.23 | -0.31 | -0.1033333 | 0.27 | 0.08 | 0.517843365 | 0.459371195 | 0.772124234 | 0.999652205 | 0.995559182 | 0.997605693 | 0.992535481 |
| Prdx1 | High | P35700 | Peroxiredoxin-1 OS=Mus musculus OX=10090 GN=Prdx1 PE=1 SV=1 | 0 | 101.033 | 72 | 15 | 40 | 15 | 199 | 22.2 | 8.12 | 0.824 | 0.836 | 0.985 | -0.28 | 0.28 | -0.26 | -0.0866667 | 0.27 | -0.02 | 0.517420618 | 0.526222693 | 0.995567333 | 0.999652205 | 0.995559182 | 0.997605693 | 0.999917596 |
| Ranbp1 | High | P34022 | Ran-specific GTPase-activating protein OS=Mus musculus OX=10090 GN=Ranbp1 PE=1 SV=2 | 0 | 8.725 | 15 | 2 | 3 | 2 | 203 | 23.6 | 5.22 | 0.856 | 0.804 | 1.065 | -0.22 | 0.22 | -0.31 | -0.1033333 | 0.265 | 0.09 | 0.587947958 | 0.450388839 | 0.66225656 | 0.999652205 | 0.995559182 | 0.997605693 | 0.986260965 |
| Aldh1l1 | High | Q8R0Y6 | Cytosolic 10-formyltetrahydrofolate dehydrogenase OS=Mus musculus OX=10090 GN=Aldh1l1 PE= | 0 | 16.6 | 8 | 7 | 7 | 7 | 902 | 98.6 | 5.91 | 0.859 | 0.809 | 1.061 | -0.22 | 0.22 | -0.31 | -0.1033333 | 0.265 | 0.09 | 0.526928619 | 0.461956018 | 0.744144631 | 0.999652205 | 0.995559182 | 0.997605693 | 0.992535481 |
| Cops5 | Low | O35864 | COP9 signalosome complex subunit 5 OS=Mus musculus OX=10090 GN=Cops5 PE=1 SV=3 | 0.076 | 1.238 | 2 | 1 | 1 | 1 | 334 | 37.5 | 6.54 | 0.823 | 0.844 | 0.975 | -0.28 | 0.28 | -0.25 | -0.0833333 | 0.265 | -0.04 | 0.459296506 | 0.539497145 | 0.922952976 | 0.999652205 | 0.995559182 | 0.997605693 | 0.992535481 |
| Twf1 | High | Q91YR1 | Twinfilin-1 OS=Mus musculus OX=10090 GN=Twf1 PE=1 SV=2 | 0.006 | 3.462 | 4 | 2 | 2 | 2 | 350 | 40.1 | 6.67 | 0.828 | 0.834 | 0.993 | -0.27 | 0.27 | -0.26 | -0.0866667 | 0.265 | -0.01 | 0.540316953 | 0.521154088 | 0.995068476 | 0.999652205 | 0.995559182 | 0.997605693 | 0.999917596 |
| Pgm2 | High | Q9D0F9 | Phosphoglucomutase-1 OS=Mus musculus OX=10090 GN=Pgm1 PE=1 SV=4 | 0 | 73.886 | 40 | 18 | 22 | 18 | 562 | 61.4 | 6.57 | 0.835 | 0.838 | 0.996 | -0.26 | 0.26 | -0.26 | -0.0866667 | 0.26 | -0.01 | 0.545441594 | 0.52931595 | 0.938273232 | 0.999652205 | 0.995559182 | 0.997605693 | 0.992535481 |
| Map1lc3a | High | Q91VR7 | Microtubule-associated proteins 1A/1B light chain 3A OS=Mus musculus OX=10090 GN=Map1lc3 | 0.001 | 7.004 | 17 | 2 | 3 | 2 | 121 | 14.3 | 8.68 | 0.836 | 0.834 | 1.003 | -0.26 | 0.26 | -0.26 | -0.0866667 | 0.26 | 0 | 0.451527272 | 0.514887066 | 0.962316386 | 0.999652205 | 0.995559182 | 0.997605693 | 0.993960013 |
| Gnaz | High | O70443 | Guanine nucleotide-binding protein G(z) subunit alpha OS=Mus musculus OX=10090 GN=Gnaz PE | 0 | 110.649 | 52 | 17 | 44 | 15 | 355 | 40.8 | 7.61 | 0.827 | 0.84 | 0.984 | -0.27 | 0.27 | -0.25 | -0.0833333 | 0.26 | -0.02 | 0.52389557 | 0.534154355 | 0.999381054 | 0.999652205 | 0.995559182 | 0.997605693 | 0.999917596 |
| Car2 | High | P00920 | Carbonic anhydrase 2 OS=Mus musculus OX=10090 GN=Ca2 PE=1 SV=4 | 0 | 77.39 | 59 | 10 | 21 | 10 | 260 | 29 | 7.01 | 0.852 | 0.825 | 1.033 | -0.23 | 0.23 | -0.28 | -0.0933333 | 0.255 | 0.05 | 0.592715312 | 0.498224271 | 0.75481897 | 0.999652205 | 0.995559182 | 0.997605693 | 0.992535481 |
| Dync1li1 | High | Q8R1Q8 | Cytoplasmic dynein 1 light intermediate chain 1 OS=Mus musculus OX=10090 GN=Dync1li1 PE=1 | 0 | 102.631 | 44 | 18 | 32 | 18 | 523 | 56.6 | 6.42 | 0.838 | 0.839 | 0.999 | -0.26 | 0.26 | -0.25 | -0.0833333 | 0.255 | 0 | 0.55348212 | 0.531904761 | 0.926514952 | 0.999652205 | 0.995559182 | 0.997605693 | 0.992535481 |
| Sh3glb1 | High | Q9JK48 | Endophilin-B1 OS=Mus musculus OX=10090 GN=Sh3glb1 PE=1 SV=1 | 0 | 24.443 | 23 | 8 | 11 | 6 | 365 | 40.8 | 6.04 | 0.835 | 0.841 | 0.993 | -0.26 | 0.26 | -0.25 | -0.0833333 | 0.255 | -0.01 | 0.493565448 | 0.538329652 | 0.949929732 | 0.999652205 | 0.995559182 | 0.997605693 | 0.992535481 |
| Dlg1 | High | H7BWY4 | Disks large homolog 1 OS=Mus musculus OX=10090 GN=Dlg1 PE=1 SV=1 | 0 | 32.149 | 12 | 10 | 11 | 7 | 872 | 96.8 | 5.9 | 0.838 | 0.84 | 0.998 | -0.26 | 0.26 | -0.25 | -0.0833333 | 0.255 | 0 | 0.511085844 | 0.534461915 | 0.990035001 | 0.999652205 | 0.995559182 | 0.997605693 | 0.999783808 |
| Ptp4a2 | High | O70274 | Protein tyrosine phosphatase type IVA 2 OS=Mus musculus OX=10090 GN=Ptp4a2 PE=1 SV=1 | 0 | 26.497 | 26 | 5 | 11 | 2 | 167 | 19.1 | 8.37 | 0.835 | 0.84 | 0.994 | -0.26 | 0.26 | -0.25 | -0.0833333 | 0.255 | -0.01 | 0.471634647 | 0.530294242 | 0.997914794 | 0.999652205 | 0.995559182 | 0.997605693 | 0.999917596 |
| Ppia | High | P17742 | Peptidyl-prolyl cis-trans isomerase A OS=Mus musculus OX=10090 GN=Ppia PE=1 SV=2 | 0 | 142.001 | 64 | 12 | 65 | 12 | 164 | 18 | 7.9 | 0.857 | 0.823 | 1.04 | -0.22 | 0.22 | -0.28 | -0.0933333 | 0.25 | 0.06 | 0.60470414 | 0.495133236 | 0.722138859 | 0.999652205 | 0.995559182 | 0.997605693 | 0.992535481 |
| Cast | High | Q8CE04 | Calpain inhibitor OS=Mus musculus OX=10090 GN=Cast PE=1 SV=1 | 0.001 | 7.086 | 5 | 2 | 2 | 2 | 704 | 76.4 | 5.2 | 0.817 | 0.864 | 0.946 | -0.29 | 0.29 | -0.21 | -0.07 | 0.25 | -0.08 | 0.437966823 | 0.592783232 | 0.764035017 | 0.999652205 | 0.995559182 | 0.997605693 | 0.992535481 |
| Eci2 | Medium | Q9WUR2 | Enoyl-CoA delta isomerase 2 OS=Mus musculus OX=10090 GN=Eci2 PE=1 SV=2 | 0.048 | 1.525 | 2 | 1 | 1 | 1 | 391 | 43.2 | 8.92 | 0.853 | 0.829 | 1.029 | -0.23 | 0.23 | -0.27 | -0.09 | 0.25 | 0.04 | 0.490575926 | 0.482170999 | 0.806008347 | 0.999652205 | 0.995559182 | 0.997605693 | 0.992535481 |
| | High | F6UFG6 | Acidic leucine-rich nuclear phosphoprotein 32 family member A (Fragment) OS=Mus musculus O | 0 | 11.336 | 26 | 4 | 4 | 4 | 138 | 15.4 | 5.36 | 0.852 | 0.828 | 1.028 | -0.23 | 0.23 | -0.27 | -0.09 | 0.25 | 0.04 | 0.493684674 | 0.506697246 | 0.917889491 | 0.999652205 | 0.995559182 | 0.997605693 | 0.992535481 |
| Ddb1 | High | Q3U1J4 | DNA damage-binding protein 1 OS=Mus musculus OX=10090 GN=Ddb1 PE=1 SV=2 | 0 | 55.975 | 16 | 18 | 21 | 18 | 1140 | 126.8 | 5.26 | 0.881 | 0.808 | 1.09 | -0.18 | 0.18 | -0.31 | -0.1033333 | 0.245 | 0.12 | 0.715839653 | 0.459817406 | 0.512699511 | 0.999652205 | 0.995559182 | 0.997605693 | 0.964378027 |
| Klc4 | High | Q9DBS5 | Kinesin light chain 4 OS=Mus musculus OX=10090 GN=Klc4 PE=1 SV=1 | 0 | 14.653 | 10 | 5 | 5 | 3 | 619 | 68.6 | 6.09 | 0.879 | 0.81 | 1.086 | -0.19 | 0.19 | -0.3 | -0.1 | 0.245 | 0.12 | 0.648229802 | 0.464195772 | 0.579665347 | 0.999652205 | 0.995559182 | 0.997605693 | 0.972910028 |
| Sept2 | High | P42208 | Septin-2 OS=Mus musculus OX=10090 GN=Septin2 PE=1 SV=2 | 0 | 35.291 | 34 | 8 | 9 | 8 | 361 | 41.5 | 6.55 | 0.881 | 0.805 | 1.094 | -0.18 | 0.18 | -0.31 | -0.1033333 | 0.245 | 0.13 | 0.584704932 | 0.451732845 | 0.630739434 | 0.999652205 | 0.995559182 | 0.997605693 | 0.981916182 |
| Arpc1b | High | Q9WV32 | Actin-related protein 2/3 complex subunit 1B OS=Mus musculus OX=10090 GN=Arpc1b PE=1 SV= | 0 | 12.79 | 5 | 1 | 1 | 1 | 372 | 41 | 8.35 | 0.876 | 0.814 | 1.077 | -0.19 | 0.19 | -0.3 | -0.1 | 0.245 | 0.11 | 0.60025229 | 0.466169009 | 0.648242698 | 0.999652205 | 0.995559182 | 0.997605693 | 0.985010103 |
| | High | A2ATS6 | Reticulon-4 receptor-like 2 (Fragment) OS=Mus musculus OX=10090 GN=Rtn4rl2 PE=1 SV=1 | 0 | 29.728 | 22 | 8 | 10 | 8 | 426 | 46.8 | 7.81 | 0.85 | 0.837 | 1.017 | -0.23 | 0.23 | -0.26 | -0.0866667 | 0.245 | 0.02 | 0.497848505 | 0.526560058 | 0.92472084 | 0.999652205 | 0.995559182 | 0.997605693 | 0.992535481 |
| Phactr4 | High | Q501J7 | Phosphatase and actin regulator 4 OS=Mus musculus OX=10090 GN=Phactr4 PE=1 SV=2 | 0 | 4.726 | 3 | 2 | 2 | 2 | 694 | 76.6 | 6.48 | 0.834 | 0.852 | 0.979 | -0.26 | 0.26 | -0.23 | -0.0766667 | 0.245 | -0.03 | 0.450301122 | 0.557576529 | 0.924863153 | 0.999652205 | 0.995559182 | 0.997605693 | 0.992535481 |
| Phyhipl | High | F7D3N3 | Phytanoyl-CoA hydroxylase-interacting protein-like OS=Mus musculus OX=10090 GN=Phyhipl PE=1 | 0 | 24.819 | 22 | 7 | 7 | 7 | 330 | 37.5 | 6.93 | 0.845 | 0.842 | 1.003 | -0.24 | 0.24 | -0.25 | -0.0833333 | 0.245 | 0 | 0.496709113 | 0.540219443 | 0.983529875 | 0.999652205 | 0.995559182 | 0.997605693 | 0.997765878 |
| Mpp2 | High | Q9WV34 | MAGUK p55 subfamily member 2 OS=Mus musculus OX=10090 GN=Mpp2 PE=1 SV=1 | 0 | 76.287 | 36 | 18 | 25 | 16 | 552 | 61.5 | 6.44 | 0.786 | 0.916 | 0.859 | -0.35 | 0.35 | -0.13 | -0.0433333 | 0.24 | -0.22 | 0.451758533 | 0.721403984 | 0.379021564 | 0.999652205 | 0.995559182 | 0.997605693 | 0.922487352 |
| Sept8 | High | B7ZC46 | Septin OS=Mus musculus OX=10090 GN=Septin8 PE=1 SV=1 | 0 | 34.392 | 22 | 8 | 13 | 5 | 440 | 50.9 | 6.09 | 0.851 | 0.841 | 1.011 | -0.23 | 0.23 | -0.25 | -0.0833333 | 0.24 | 0.02 | 0.531774099 | 0.5374905 | 0.957431978 | 0.999652205 | 0.995559182 | 0.997605693 | 0.992902921 |
| Irgm1 | Low | Q60766 | Immunity-related GTPase family M protein 1 OS=Mus musculus OX=10090 GN=Irgm1 PE=1 SV=1 | 0.06 | 1.321 | 2 | 1 | 1 | 1 | 409 | 46.5 | 8.28 | 0.841 | 0.854 | 0.985 | -0.25 | 0.25 | -0.23 | -0.0766667 | 0.24 | -0.02 | 0.491230141 | 0.547161163 | 0.974380076 | 0.999652205 | 0.995559182 | 0.997605693 | 0.997221123 |
| Abca8b | High | Q8K440 | ABC-type organic anion transporter ABCA8B OS=Mus musculus OX=10090 GN=Abca8b PE=1 SV=2 | 0 | 23.805 | 4 | 6 | 6 | 4 | 1620 | 182.9 | 7.18 | 0.846 | 0.844 | 1.002 | -0.24 | 0.24 | -0.24 | -0.08 | 0.24 | 0 | 0.544393292 | 0.544589321 | 0.986025997 | 0.999652205 | 0.995559182 | 0.997605693 | 0.998671923 |
| Wdr47 | High | Q8CGF6 | WD repeat-containing protein 47 OS=Mus musculus OX=10090 GN=Wdr47 PE=1 SV=2 | 0 | 9.089 | 4 | 4 | 4 | 4 | 920 | 102.2 | 5.97 | 0.879 | 0.821 | 1.071 | -0.19 | 0.19 | -0.28 | -0.0933333 | 0.235 | 0.1 | 0.619123813 | 0.490819385 | 0.644139025 | 0.999652205 | 0.995559182 | 0.997605693 | 0.985010103 |
| Gng2 | High | P63213 | Guanine nucleotide-binding protein G(I)/G(S)/G(O) subunit gamma-2 OS=Mus musculus OX=100 | 0 | 23.176 | 62 | 4 | 10 | 4 | 71 | 7.8 | 7.99 | 0.831 | 0.869 | 0.957 | -0.27 | 0.27 | -0.2 | -0.0666667 | 0.235 | -0.06 | 0.490650636 | 0.604531387 | 0.760710337 | 0.999652205 | 0.995559182 | 0.997605693 | 0.992535481 |
| Tppp | High | Q7TQD2 | Tubulin polymerization-promoting protein OS=Mus musculus OX=10090 GN=Tppp PE=1 SV=1 | 0 | 53.2 | 37 | 9 | 17 | 9 | 218 | 23.6 | 9.42 | 0.826 | 0.878 | 0.942 | -0.28 | 0.28 | -0.19 | -0.0633333 | 0.235 | -0.09 | 0.52332804 | 0.627201456 | 0.773179144 | 0.999652205 | 0.995559182 | 0.997605693 | 0.992535481 |
| Ahnak | High | E9Q616 | AHNAK nucleoprotein (desmoyokin) OS=Mus musculus OX=10090 GN=Ahnak PE=1 SV=1 | 0.001 | 7.833 | 4 | 3 | 3 | 3 | 5656 | 603.9 | 6.3 | 0.882 | 0.819 | 1.077 | -0.18 | 0.18 | -0.29 | -0.0966667 | 0.235 | 0.11 | 0.540281114 | 0.474648331 | 0.602264705 | 0.999652205 | 0.995559182 | 0.997605693 | 0.97826833 |
| Fam129a | Medium | Q3UW53 | Protein Niban 1 OS=Mus musculus OX=10090 GN=Niban1 PE=1 SV=2 | 0.01 | 2.729 | 1 | 1 | 1 | 1 | 926 | 102.6 | 4.78 | 0.886 | 0.817 | 1.084 | -0.18 | 0.18 | -0.29 | -0.0966667 | 0.235 | 0.12 | 0.586823673 | 0.476061577 | 0.611773358 | 0.999652205 | 0.995559182 | 0.997605693 | 0.97826833 |
| Pafah1b1 | High | P63005 | Platelet-activating factor acetylhydrolase IB subunit beta OS=Mus musculus OX=10090 GN=Pafah | 0 | 27.312 | 21 | 8 | 10 | 8 | 410 | 46.6 | 7.37 | 0.884 | 0.817 | 1.082 | -0.18 | 0.18 | -0.29 | -0.0966667 | 0.235 | 0.11 | 0.603250078 | 0.479641925 | 0.6633528 | 0.999652205 | 0.995559182 | 0.997605693 | 0.986647506 |
| | High | E9Q4G8 | Activated leukocyte cell adhesion molecule OS=Mus musculus OX=10090 GN=Alcam PE=1 SV=1 | 0 | 13.772 | 8 | 5 | 5 | 5 | 555 | 61.9 | 6.06 | 0.864 | 0.833 | 1.037 | -0.21 | 0.21 | -0.26 | -0.0866667 | 0.235 | 0.05 | 0.555272058 | 0.517421627 | 0.852868041 | 0.999652205 | 0.995559182 | 0.997605693 | 0.992535481 |
| Tnpo2 | High | E9PV58 | Transportin-2 OS=Mus musculus OX=10090 GN=Tnpo2 PE=1 SV=1 | 0.001 | 5.551 | 3 | 3 | 5 | 1 | 897 | 101.4 | 5.01 | 0.863 | 0.834 | 1.035 | -0.21 | 0.21 | -0.26 | -0.0866667 | 0.235 | 0.05 | 0.528355978 | 0.52070151 | 0.886119269 | 0.999652205 | 0.995559182 | 0.997605693 | 0.992535481 |
| Sept11 | High | Q8C1B7 | Septin-11 OS=Mus musculus OX=10090 GN=Septin11 PE=1 SV=4 | 0 | 71.348 | 32 | 11 | 17 | 5 | 431 | 49.7 | 6.68 | 0.854 | 0.849 | 1.005 | -0.23 | 0.23 | -0.24 | -0.08 | 0.235 | 0.01 | 0.646296673 | 0.556850164 | 0.888014817 | 0.999652205 | 0.995559182 | 0.997605693 | 0.992535481 |
| Rab35 | High | Q6PHN9 | Ras-related protein Rab-35 OS=Mus musculus OX=10090 GN=Rab35 PE=1 SV=1 | 0 | 66.13 | 52 | 9 | 27 | 7 | 201 | 23 | 8.29 | 0.837 | 0.864 | 0.969 | -0.26 | 0.26 | -0.21 | -0.07 | 0.235 | -0.04 | 0.597366824 | 0.592356864 | 0.922712832 | 0.999652205 | 0.995559182 | 0.997605693 | 0.992535481 |
| | High | E9QMY1 | OX-2 membrane glycoprotein OS=Mus musculus OX=10090 GN=Cd200 PE=1 SV=1 | 0 | 14.468 | 13 | 3 | 4 | 3 | 269 | 30.3 | 8.51 | 0.827 | 0.879 | 0.942 | -0.27 | 0.27 | -0.19 | -0.0633333 | 0.23 | -0.09 | 0.541753495 | 0.629321194 | 0.759864532 | 0.999652205 | 0.995559182 | 0.997605693 | 0.992535481 |
| Ank2 | High | S4R2R5 | Ankyrin-2 OS=Mus musculus OX=10090 GN=Ank2 PE=1 SV=3 | 0 | 99.229 | 10 | 29 | 34 | 27 | 3900 | 426.2 | 5.12 | 0.833 | 0.868 | 0.959 | -0.26 | 0.26 | -0.2 | -0.0666667 | 0.23 | -0.06 | 0.5402783 | 0.603681264 | 0.866213537 | 0.999652205 | 0.995559182 | 0.997605693 | 0.992535481 |
| Syt9 | Medium | Q9R0N9 | Synaptotagmin-9 OS=Mus musculus OX=10090 GN=Syt9 PE=1 SV=2 | 0.01 | 2.59 | 2 | 1 | 1 | 1 | 491 | 56.2 | 7.24 | 0.887 | 0.82 | 1.082 | -0.17 | 0.17 | -0.29 | -0.0966667 | 0.23 | 0.11 | 0.622490722 | 0.481629058 | 0.650446926 | 0.999652205 | 0.995559182 | 0.997605693 | 0.985010103 |
| Dctn4 | High | Q8CBY8 | Dynactin subunit 4 OS=Mus musculus OX=10090 GN=Dctn4 PE=1 SV=1 | 0.001 | 4.499 | 4 | 1 | 1 | 1 | 467 | 53 | 7.72 | 0.82 | 0.89 | 0.922 | -0.29 | 0.29 | -0.17 | -0.0566667 | 0.23 | -0.12 | 0.447719591 | 0.638486336 | 0.73325943 | 0.999652205 | 0.995559182 | 0.997605693 | 0.992535481 |
| Pak7 | High | Q8C015 | Serine/threonine-protein kinase PAK 5 OS=Mus musculus OX=10090 GN=Pak5 PE=1 SV=1 | 0.005 | 3.55 | 2 | 1 | 1 | 1 | 719 | 80.9 | 8.21 | 0.856 | 0.847 | 1.012 | -0.22 | 0.22 | -0.24 | -0.08 | 0.23 | 0.02 | 0.531764343 | 0.529299537 | 0.929830014 | 0.999652205 | 0.995559182 | 0.997605693 | 0.992535481 |
| | High | E9PYD9 | Folate receptor alpha (Fragment) OS=Mus musculus OX=10090 GN=Folr1 PE=1 SV=1 | 0 | 25.742 | 42 | 5 | 8 | 5 | 137 | 16 | 6.83 | 0.886 | 0.821 | 1.08 | -0.17 | 0.17 | -0.28 | -0.0933333 | 0.225 | 0.11 | 0.597820841 | 0.48948575 | 0.684827315 | 0.999652205 | 0.995559182 | 0.997605693 | 0.991285312 |
| Stip1 | High | Q60864 | Stress-induced-phosphoprotein 1 OS=Mus musculus OX=10090 GN=Stip1 PE=1 SV=1 | 0 | 90.969 | 41 | 25 | 31 | 25 | 543 | 62.5 | 6.8 | 0.805 | 0.905 | 0.89 | -0.31 | 0.31 | -0.14 | -0.0466667 | 0.225 | -0.17 | 0.468136574 | 0.695221841 | 0.513954563 | 0.999652205 | 0.995559182 | 0.997605693 | 0.964378027 |
| Atp11a | High | P98197 | Phospholipid-transporting ATPase IH OS=Mus musculus OX=10090 GN=Atp11a PE=1 SV=1 | 0 | 20.067 | 6 | 6 | 7 | 6 | 1187 | 135.4 | 6.48 | 0.827 | 0.883 | 0.936 | -0.27 | 0.27 | -0.18 | -0.06 | 0.225 | -0.09 | 0.504585128 | 0.64027971 | 0.734484202 | 0.999652205 | 0.995559182 | 0.997605693 | 0.992535481 |
| | Medium | F7CFT1 | AN1-type zinc finger protein 1 (Fragment) OS=Mus musculus OX=10090 GN=Zfand1 PE=1 SV=1 | 0.045 | 1.585 | 3 | 1 | 1 | 1 | 247 | 27.6 | 8.75 | 0.878 | 0.837 | 1.049 | -0.19 | 0.19 | -0.26 | -0.0866667 | 0.225 | 0.07 | 0.54676519 | 0.522265987 | 0.764412822 | 0.999652205 | 0.995559182 | 0.997605693 | 0.992535481 |
| Npepps | High | Q11011 | Puromycin-sensitive aminopeptidase OS=Mus musculus OX=10090 GN=Npepps PE=1 SV=2 | 0 | 86.554 | 28 | 22 | 29 | 22 | 920 | 103.3 | 5.9 | 0.859 | 0.851 | 1.008 | -0.22 | 0.22 | -0.23 | -0.0766667 | 0.225 | 0.01 | 0.609959265 | 0.562557945 | 0.878142949 | 0.999652205 | 0.995559182 | 0.997605693 | 0.992535481 |
| Myh14 | High | Q6URW6 | Myosin-14 OS=Mus musculus OX=10090 GN=Myh14 PE=1 SV=1 | 0 | 41.511 | 6 | 12 | 13 | 9 | 2000 | 228.4 | 5.55 | 0.861 | 0.851 | 1.012 | -0.22 | 0.22 | -0.23 | -0.0766667 | 0.225 | 0.02 | 0.582650926 | 0.560829121 | 0.922932455 | 0.999652205 | 0.995559182 | 0.997605693 | 0.992535481 |
| Ahsa1 | High | Q8BK64 | Activator of 90 kDa heat shock protein ATPase homolog 1 OS=Mus musculus OX=10090 GN=Ahsa | 0 | 20.272 | 15 | 4 | 7 | 4 | 338 | 38.1 | 5.53 | 0.862 | 0.849 | 1.015 | -0.21 | 0.21 | -0.24 | -0.08 | 0.225 | 0.02 | 0.55182791 | 0.557155372 | 0.918269074 | 0.999652205 | 0.995559182 | 0.997605693 | 0.992535481 |
| Adprh | High | P54923 | [Protein ADP-ribosylarginine] hydrolase OS=Mus musculus OX=10090 GN=Adprh PE=1 SV=1 | 0 | 18.237 | 19 | 6 | 6 | 6 | 362 | 40 | 5.76 | 0.84 | 0.874 | 0.961 | -0.25 | 0.25 | -0.19 | -0.0633333 | 0.22 | -0.06 | 0.491768257 | 0.617819188 | 0.829670206 | 0.999652205 | 0.995559182 | 0.997605693 | 0.992535481 |
| Crtac1 | High | Q8R555 | Cartilage acidic protein 1 OS=Mus musculus OX=10090 GN=Crtac1 PE=2 SV=1 | 0.001 | 4.622 | 3 | 2 | 2 | 2 | 646 | 70.3 | 5.14 | 0.867 | 0.854 | 1.015 | -0.21 | 0.21 | -0.23 | -0.0766667 | 0.22 | 0.02 | 0.516687025 | 0.553634652 | 0.868848757 | 0.999652205 | 0.995559182 | 0.997605693 | 0.992535481 |
| Ndrg2 | High | Q9QYG0 | Protein NDRG2 OS=Mus musculus OX=10090 GN=Ndrg2 PE=1 SV=1 | 0 | 76.97 | 56 | 13 | 21 | 13 | 371 | 40.8 | 5.4 | 0.82 | 0.903 | 0.909 | -0.29 | 0.29 | -0.15 | -0.05 | 0.22 | -0.14 | 0.507229767 | 0.68943839 | 0.605305743 | 0.999652205 | 0.995559182 | 0.997605693 | 0.97826833 |
| Usp14 | High | E9PYI8 | Ubiquitin carboxyl-terminal hydrolase OS=Mus musculus OX=10090 GN=Usp14 PE=1 SV=1 | 0 | 14.762 | 6 | 2 | 3 | 2 | 458 | 52.3 | 5.66 | 0.82 | 0.9 | 0.912 | -0.29 | 0.29 | -0.15 | -0.05 | 0.22 | -0.13 | 0.471147335 | 0.681332495 | 0.620338266 | 0.999652205 | 0.995559182 | 0.997605693 | 0.97826833 |
| Cct3 | High | P80318 | T-complex protein 1 subunit gamma OS=Mus musculus OX=10090 GN=Cct3 PE=1 SV=1 | 0 | 122.508 | 55 | 24 | 46 | 24 | 545 | 60.6 | 6.7 | 0.916 | 0.81 | 1.13 | -0.13 | 0.13 | -0.3 | -0.1 | 0.215 | 0.18 | 0.768986976 | 0.464554486 | 0.374583209 | 0.999652205 | 0.995559182 | 0.997605693 | 0.920164724 |
| Cct5 | High | P80316 | T-complex protein 1 subunit epsilon OS=Mus musculus OX=10090 GN=Cct5 PE=1 SV=1 | 0 | 125.051 | 44 | 22 | 38 | 22 | 541 | 59.6 | 6.02 | 0.912 | 0.812 | 1.123 | -0.13 | 0.13 | -0.3 | -0.1 | 0.215 | 0.17 | 0.75876149 | 0.469609027 | 0.398165995 | 0.999652205 | 0.995559182 | 0.997605693 | 0.93262326 |
| Dlgap4 | High | B1AZP2 | Disks large-associated protein 4 OS=Mus musculus OX=10090 GN=Dlgap4 PE=1 SV=1 | 0 | 43.3 | 14 | 12 | 16 | 12 | 992 | 108 | 7.08 | 0.886 | 0.843 | 1.051 | -0.18 | 0.18 | -0.25 | -0.0833333 | 0.215 | 0.07 | 0.729725834 | 0.542277718 | 0.675893058 | 0.999652205 | 0.995559182 | 0.997605693 | 0.990250803 |

| | | | | | | | | | | | | | | | | | | | | | | | | | | | | |
|---|---|---|---|---|---|---|---|---|---|---|---|---|---|---|---|---|---|---|---|---|---|---|---|---|---|---|---|---|
| Psmd5 | High | Q8BJY1 | 26S proteasome non-ATPase regulatory subunit 5 OS=Mus musculus OX=10090 GN=Psmd5 PE=1 | 0 | 14.782 | 11 | 4 | 4 | 4 | 504 | 55.9 | 5.21 | 0.883 | 0.841 | 1.05 | -0.18 | 0.18 | -0.25 | -0.0833333 | 0.215 | 0.07 | 0.553174807 | 0.537788607 | 0.706209415 | 0.999652205 | 0.995559182 | 0.997605693 | 0.992485323 |
| Plcb4 | High | Q91UZ1 | 1-phosphatidylinositol 4,5-bisphosphate phosphodiesterase OS=Mus musculus OX=10090 GN=Plcl | 0 | 33.739 | 11 | 10 | 11 | 10 | 1175 | 134.4 | 6.9 | 0.846 | 0.877 | 0.964 | -0.24 | 0.24 | -0.19 | -0.0633333 | 0.215 | -0.05 | 0.486426807 | 0.624684707 | 0.803428065 | 0.999652205 | 0.995559182 | 0.997605693 | 0.992535481 |
| Ppp1ca | High | P62137 | Serine/threonine-protein phosphatase PP1-alpha catalytic subunit OS=Mus musculus OX=10090 G | 0 | 37.577 | 27 | 6 | 9 | 1 | 330 | 37.5 | 6.33 | 0.864 | 0.861 | 1.004 | -0.21 | 0.21 | -0.22 | -0.0733333 | 0.215 | 0.01 | 0.519990517 | 0.583760288 | 0.98763497 | 0.999652205 | 0.995559182 | 0.997605693 | 0.998907549 |
| Kif5b | High | Q61768 | Kinesin-1 heavy chain OS=Mus musculus OX=10090 GN=Kif5b PE=1 SV=3 | 0 | 61.334 | 20 | 16 | 20 | 14 | 963 | 109.5 | 6.44 | 0.894 | 0.835 | 1.07 | -0.16 | 0.16 | -0.26 | -0.0866667 | 0.21 | 0.1 | 0.726517409 | 0.523746402 | 0.610149627 | 0.999652205 | 0.995559182 | 0.997605693 | 0.97826833 |
| Gabrg2 | High | Q8C7N0 | GABA(A) receptor subunit gamma-2 OS=Mus musculus OX=10090 GN=Gabrg2 PE=1 SV=1 | 0 | 9.362 | 10 | 3 | 4 | 3 | 384 | 44.5 | 8.82 | 0.824 | 0.907 | 0.908 | -0.28 | 0.28 | -0.14 | -0.0466667 | 0.21 | -0.14 | 0.473501909 | 0.701009649 | 0.612783049 | 0.999652205 | 0.995559182 | 0.997605693 | 0.97826833 |
| Ccdc92 | High | Q8VDN4 | Coiled-coil domain-containing protein 92 OS=Mus musculus OX=10090 GN=Ccdc92 PE=1 SV=1 | 0.003 | 3.952 | 5 | 2 | 2 | 2 | 314 | 35.2 | 9.38 | 0.832 | 0.903 | 0.922 | -0.27 | 0.27 | -0.15 | -0.05 | 0.21 | -0.12 | 0.463386227 | 0.687673566 | 0.653484994 | 0.999652205 | 0.995559182 | 0.997605693 | 0.985010103 |
| Wars | High | P32921 | Tryptophan--tRNA ligase, cytoplasmic OS=Mus musculus OX=10090 GN=Wars1 PE=1 SV=2 | 0 | 44.748 | 26 | 11 | 12 | 11 | 481 | 54.3 | 6.89 | 0.887 | 0.843 | 1.052 | -0.17 | 0.17 | -0.25 | -0.0833333 | 0.21 | 0.07 | 0.650477927 | 0.542206861 | 0.731751947 | 0.999652205 | 0.995559182 | 0.997605693 | 0.992535481 |
| Igf1r | High | E9QNX9 | Tyrosine-protein kinase receptor OS=Mus musculus OX=10090 GN=Igf1r PE=1 SV=1 | 0 | 35.067 | 10 | 13 | 14 | 11 | 1369 | 155.2 | 5.83 | 0.835 | 0.893 | 0.935 | -0.26 | 0.26 | -0.16 | -0.0533333 | 0.21 | -0.1 | 0.518793426 | 0.664067288 | 0.753219898 | 0.999652205 | 0.995559182 | 0.997605693 | 0.992535481 |
| Acy1 | Medium | A0A0R4J0 | N-acyl-L-amino-acid amidohydrolase OS=Mus musculus OX=10090 GN=Acy1 PE=1 SV=1 | 0.045 | 1.576 | 2 | 1 | 1 | 1 | 408 | 45.8 | 6.21 | 0.878 | 0.855 | 1.027 | -0.19 | 0.19 | -0.23 | -0.0766667 | 0.21 | 0.04 | 0.547011827 | 0.566314669 | 0.882735983 | 0.999652205 | 0.995559182 | 0.997605693 | 0.992535481 |
| Pmm2 | High | Q9Z2M7 | Phosphomannomutase 2 OS=Mus musculus OX=10090 GN=Pmm2 PE=1 SV=1 | 0.006 | 3.328 | 7 | 2 | 2 | 1 | 242 | 27.6 | 6.42 | 0.871 | 0.86 | 1.014 | -0.2 | 0.2 | -0.22 | -0.0733333 | 0.21 | 0.02 | 0.532191702 | 0.579099309 | 0.944069687 | 0.999652205 | 0.995559182 | 0.997605693 | 0.992535481 |
| Bsn | High | O88737 | Protein bassoon OS=Mus musculus OX=10090 GN=Bsn PE=1 SV=4 | 0 | 79.382 | 8 | 20 | 24 | 19 | 3942 | 418.6 | 7.71 | 0.857 | 0.873 | 0.982 | -0.22 | 0.22 | -0.2 | -0.0666667 | 0.21 | -0.03 | 0.651717498 | 0.615149541 | 0.997270212 | 0.999652205 | 0.995559182 | 0.997605693 | 0.999917596 |
| Trappc6b | High | Q9D289 | Trafficking protein particle complex subunit 6B OS=Mus musculus OX=10090 GN=Trappc6b PE=1 | 0.001 | 7.279 | 15 | 3 | 3 | 3 | 158 | 17.9 | 8.68 | 0.914 | 0.817 | 1.119 | -0.13 | 0.13 | -0.29 | -0.0966667 | 0.21 | 0.16 | 0.79943528 | 0.481149151 | 0.468663685 | 0.999652205 | 0.995559182 | 0.997605693 | 0.961044326 |
| Cyfip1 | High | A0A0R4J1 | Cytoplasmic FMR1-interacting protein OS=Mus musculus OX=10090 GN=Cyfip1 PE=1 SV=1 | 0 | 26.378 | 7 | 9 | 11 | 3 | 1251 | 144.8 | 6.95 | 0.916 | 0.819 | 1.118 | -0.13 | 0.13 | -0.29 | -0.0966667 | 0.21 | 0.16 | 0.708082676 | 0.464108361 | 0.537756075 | 0.999652205 | 0.995559182 | 0.997605693 | 0.96694628 |
| Ctnna1 | High | P26231 | Catenin alpha-1 OS=Mus musculus OX=10090 GN=Ctnna1 PE=1 SV=1 | 0 | 68.452 | 23 | 15 | 22 | 11 | 906 | 100 | 6.23 | 0.806 | 0.934 | 0.863 | -0.31 | 0.31 | -0.1 | -0.0333333 | 0.205 | -0.21 | 0.478139532 | 0.766840659 | 0.3970488 | 0.999652205 | 0.995559182 | 0.997605693 | 0.93262326 |
| Ppp1r16a | High | Q923M0 | Protein phosphatase 1 regulatory subunit 16A OS=Mus musculus OX=10090 GN=Ppp1r16a PE=1 S | 0.001 | 5.607 | 2 | 1 | 2 | 1 | 524 | 57.5 | 5.81 | 0.821 | 0.913 | 0.9 | -0.28 | 0.28 | -0.13 | -0.0433333 | 0.205 | -0.15 | 0.458184661 | 0.711265324 | 0.577581536 | 0.999652205 | 0.995559182 | 0.997605693 | 0.972910028 |
| Ptp4a1; Gi | High | Q63739 | Protein tyrosine phosphatase type IVA 1 OS=Mus musculus OX=10090 GN=Ptp4a1 PE=1 SV=1 | 0 | 26.209 | 23 | 5 | 11 | 2 | 173 | 19.8 | 8.97 | 0.909 | 0.832 | 1.093 | -0.14 | 0.14 | -0.27 | -0.09 | 0.205 | 0.13 | 0.703556138 | 0.515619641 | 0.611947139 | 0.999652205 | 0.995559182 | 0.997605693 | 0.97826833 |
| Il17a | Medium | Q62386 | Interleukin-17A OS=Mus musculus OX=10090 GN=Il17a PE=1 SV=1 | 0.042 | 1.629 | 5 | 1 | 2 | 1 | 158 | 17.5 | 8.92 | 0.884 | 0.851 | 1.039 | -0.18 | 0.18 | -0.23 | -0.0766667 | 0.205 | 0.06 | 0.665681396 | 0.561506974 | 0.805333624 | 0.999652205 | 0.995559182 | 0.997605693 | 0.992535481 |
| Hip1r | High | Q9JKY5 | Huntingtin-interacting protein 1-related protein OS=Mus musculus OX=10090 GN=Hip1r PE=1 SV= | 0 | 9.953 | 3 | 3 | 3 | 2 | 1068 | 119.4 | 6.52 | 0.919 | 0.816 | 1.126 | -0.12 | 0.12 | -0.29 | -0.0966667 | 0.205 | 0.17 | 0.650175928 | 0.460142577 | 0.424039337 | 0.999652205 | 0.995559182 | 0.997605693 | 0.940873985 |
| Gdi2 | High | Q61598 | Rab GDP dissociation inhibitor beta OS=Mus musculus OX=10090 GN=Gdi2 PE=1 SV=1 | 0 | 97.818 | 51 | 20 | 30 | 15 | 445 | 50.5 | 6.25 | 0.887 | 0.855 | 1.038 | -0.17 | 0.17 | -0.23 | -0.0766667 | 0.2 | 0.05 | 0.689207658 | 0.571080461 | 0.73539707 | 0.999652205 | 0.995559182 | 0.997605693 | 0.992535481 |
| Hras1; Hra | High | Q61411 | GTPase HRas OS=Mus musculus OX=10090 GN=Hras PE=1 SV=2 | 0 | 189.504 | 79 | 14 | 50 | 5 | 189 | 21.3 | 5.31 | 0.866 | 0.879 | 0.985 | -0.21 | 0.21 | -0.19 | -0.0633333 | 0.2 | -0.02 | 0.629411315 | 0.630298306 | 0.999381054 | 0.999652205 | 0.995559182 | 0.997605693 | 0.999917596 |
| Ak1 | High | Q9R0Y5 | Adenylate kinase isoenzyme 1 OS=Mus musculus OX=10090 GN=Ak1 PE=1 SV=1 | 0 | 28.655 | 47 | 8 | 10 | 8 | 194 | 21.5 | 5.81 | 0.853 | 0.893 | 0.955 | -0.23 | 0.23 | -0.16 | -0.0533333 | 0.195 | -0.07 | 0.541704002 | 0.664221958 | 0.757238962 | 0.999652205 | 0.995559182 | 0.997605693 | 0.992535481 |
| Pebp1 | High | P70296 | Phosphatidylethanolamine-binding protein 1 OS=Mus musculus OX=10090 GN=Pebp1 PE=1 SV=3 | 0 | 92.276 | 73 | 10 | 31 | 10 | 187 | 20.8 | 5.4 | 0.882 | 0.866 | 1.018 | -0.18 | 0.18 | -0.21 | -0.07 | 0.195 | 0.03 | 0.675505875 | 0.599369834 | 0.829207487 | 0.999652205 | 0.995559182 | 0.997605693 | 0.992535481 |
| Marcks | High | P26645 | Myristoylated alanine-rich C-kinase substrate OS=Mus musculus OX=10090 GN=Marcks PE=1 SV= | 0 | 50.881 | 30 | 6 | 14 | 6 | 309 | 29.6 | 4.34 | 0.854 | 0.893 | 0.957 | -0.23 | 0.23 | -0.16 | -0.0533333 | 0.195 | -0.06 | 0.587909151 | 0.665343589 | 0.858443251 | 0.999652205 | 0.995559182 | 0.997605693 | 0.992535481 |
| Hspa8 | High | P63017 | Heat shock cognate 71 kDa protein OS=Mus musculus OX=10090 GN=Hspa8 PE=1 SV=1 | 0 | 556.08 | 75 | 49 | 282 | 32 | 646 | 70.8 | 5.52 | 0.875 | 0.871 | 1.005 | -0.19 | 0.19 | -0.2 | -0.0666667 | 0.195 | 0.01 | 0.656511721 | 0.609944221 | 0.893539072 | 0.999652205 | 0.995559182 | 0.997605693 | 0.992535481 |
| Vps35 | High | Q9EQH3 | Vacuolar protein sorting-associated protein 35 OS=Mus musculus OX=10090 GN=Vps35 PE=1 SV= | 0 | 41.622 | 15 | 10 | 11 | 10 | 796 | 91.7 | 5.44 | 0.866 | 0.881 | 0.982 | -0.21 | 0.21 | -0.18 | -0.06 | 0.195 | -0.03 | 0.536958612 | 0.636107946 | 0.914590864 | 0.999652205 | 0.995559182 | 0.997605693 | 0.992535481 |
| Gnl1 | High | P36916 | Guanine nucleotide-binding protein-like 1 OS=Mus musculus OX=10090 GN=Gnl1 PE=1 SV=4 | 0 | 20.668 | 10 | 6 | 6 | 6 | 607 | 68.7 | 5.68 | 0.872 | 0.879 | 0.992 | -0.2 | 0.2 | -0.19 | -0.0633333 | 0.195 | -0.01 | 0.585737765 | 0.630464163 | 0.990522885 | 0.999652205 | 0.995559182 | 0.997605693 | 0.999783808 |
| Aldoa | High | A6ZI44 | Fructose-bisphosphate aldolase OS=Mus musculus OX=10090 GN=Aldoa PE=1 SV=1 | 0 | 722.822 | 86 | 40 | 696 | 33 | 418 | 45.1 | 7.91 | 0.922 | 0.837 | 1.101 | -0.12 | 0.12 | -0.26 | -0.0866667 | 0.19 | 0.14 | 0.786038527 | 0.527145672 | 0.469981014 | 0.999652205 | 0.995559182 | 0.997605693 | 0.961044326 |
| | High | E0CX61 | Small glutamine-rich tetratricopeptide repeat-containing protein beta (Fragment) OS=Mus muscu | 0 | 11.773 | 18 | 4 | 4 | 4 | 244 | 26.8 | 5.44 | 0.924 | 0.832 | 1.111 | -0.11 | 0.11 | -0.27 | -0.09 | 0.19 | 0.15 | 0.720714211 | 0.514678459 | 0.543233012 | 0.999652205 | 0.995559182 | 0.997605693 | 0.96694628 |
| AW54987 | High | Q8BR90 | UPF0600 protein C5orf51 homolog OS=Mus musculus OX=10090 PE=1 SV=1 | 0 | 12.922 | 18 | 3 | 3 | 3 | 294 | 33.5 | 5.19 | 0.907 | 0.846 | 1.072 | -0.14 | 0.14 | -0.24 | -0.08 | 0.19 | 0.1 | 0.598521457 | 0.52495631 | 0.6094793 | 0.999652205 | 0.995559182 | 0.997605693 | 0.97826833 |
| Adssl1 | High | P28650 | Adenylosuccinate synthetase isozyme 1 OS=Mus musculus OX=10090 GN=Adss1 PE=1 SV=2 | 0 | 9.909 | 5 | 2 | 2 | 2 | 457 | 50.2 | 8.43 | 0.91 | 0.847 | 1.074 | -0.14 | 0.14 | -0.24 | -0.08 | 0.19 | 0.1 | 0.680709836 | 0.528351151 | 0.698536181 | 0.999652205 | 0.995559182 | 0.997605693 | 0.991285312 |
| Ppp2r4; Pt | High | P58389 | Serine/threonine-protein phosphatase 2A activator OS=Mus musculus OX=10090 GN=Ptpa PE=1 S | 0 | 10.961 | 10 | 4 | 4 | 4 | 323 | 36.7 | 6.39 | 0.881 | 0.873 | 1.009 | -0.18 | 0.18 | -0.2 | -0.0666667 | 0.19 | 0.01 | 0.58260726 | 0.615287263 | 0.985601918 | 0.999652205 | 0.995559182 | 0.997605693 | 0.998604519 |
| Kpna2 | High | P52293 | Importin subunit alpha-1 OS=Mus musculus OX=10090 GN=Kpna2 PE=1 SV=2 | 0 | 33.491 | 14 | 5 | 7 | 5 | 529 | 57.9 | 5.68 | 0.956 | 0.809 | 1.182 | -0.06 | 0.06 | -0.31 | -0.1033333 | 0.185 | 0.24 | 0.812197029 | 0.46096919 | 0.344530474 | 0.999652205 | 0.995559182 | 0.997605693 | 0.907770836 |
| | High | A0A571BE | Tropomyosin alpha-4 chain OS=Mus musculus OX=10090 GN=Tpm4 PE=1 SV=1 | 0 | 8.189 | 16 | 3 | 3 | 3 | 158 | 18.4 | 4.73 | 0.831 | 0.935 | 0.889 | -0.27 | 0.27 | -0.1 | -0.0333333 | 0.185 | -0.17 | 0.550576212 | 0.7696743 | 0.521695664 | 0.999652205 | 0.995559182 | 0.997605693 | 0.964378027 |
| Dbn1 | High | Q9QXS6 | Drebrin OS=Mus musculus OX=10090 GN=Dbn1 PE=1 SV=4 | 0 | 35.357 | 23 | 10 | 13 | 10 | 706 | 77.2 | 4.49 | 0.842 | 0.92 | 0.915 | -0.25 | 0.25 | -0.12 | -0.04 | 0.185 | -0.13 | 0.476046474 | 0.733305809 | 0.572412685 | 0.999652205 | 0.995559182 | 0.997605693 | 0.972910028 |
| Ctnnb1 | High | Q02248 | Catenin beta-1 OS=Mus musculus OX=10090 GN=Ctnnb1 PE=1 SV=1 | 0 | 45.469 | 17 | 12 | 18 | 1 | 781 | 85.4 | 5.86 | 0.834 | 0.925 | 0.902 | -0.26 | 0.26 | -0.11 | -0.0366667 | 0.185 | -0.15 | 0.572900648 | 0.744352665 | 0.572782421 | 0.999652205 | 0.995559182 | 0.997605693 | 0.972910028 |
| Rbbp7 | Medium | Q60973 | Histone-binding protein RBBP7 OS=Mus musculus OX=10090 GN=Rbbp7 PE=1 SV=1 | 0.016 | 2.145 | 3 | 1 | 1 | 1 | 425 | 47.8 | 5.05 | 0.91 | 0.853 | 1.066 | -0.14 | 0.14 | -0.23 | -0.0766667 | 0.185 | 0.09 | 0.666032715 | 0.533498308 | 0.659132853 | 0.999652205 | 0.995559182 | 0.997605693 | 0.985010103 |
| Llgl1 | Medium | Q80Y17 | Lethal(2) giant larvae protein homolog 1 OS=Mus musculus OX=10090 GN=Llgl1 PE=1 SV=1 | 0.011 | 2.5 | 1 | 1 | 1 | 1 | 1036 | 112.5 | 6.48 | 0.907 | 0.85 | 1.068 | -0.14 | 0.14 | -0.23 | -0.0766667 | 0.185 | 0.09 | 0.696896524 | 0.553444932 | 0.695587537 | 0.999652205 | 0.995559182 | 0.997605693 | 0.991285312 |
| | High | D6RFU9 | Synaptophysin-like protein OS=Mus musculus OX=10090 GN=Sypl PE=1 SV=1 | 0 | 9.235 | 14 | 2 | 3 | 2 | 79 | 8.9 | 9.29 | 0.864 | 0.896 | 0.964 | -0.21 | 0.21 | -0.16 | -0.0533333 | 0.185 | -0.05 | 0.518978631 | 0.673177784 | 0.777536646 | 0.999652205 | 0.995559182 | 0.997605693 | 0.992535481 |
| Rap2a | High | Q80ZJ1 | Ras-related protein Rap-2a OS=Mus musculus OX=10090 GN=Rap2a PE=1 SV=2 | 0 | 48.345 | 56 | 8 | 16 | 4 | 183 | 20.6 | 4.82 | 0.887 | 0.869 | 1.021 | -0.17 | 0.17 | -0.2 | -0.0666667 | 0.185 | 0.03 | 0.706965841 | 0.604759175 | 0.812510915 | 0.999652205 | 0.995559182 | 0.997605693 | 0.992535481 |
| Stk11ip | Medium | Q3TAA7 | Serine/threonine-protein kinase 11-interacting protein OS=Mus musculus OX=10090 GN=Stk11ip | 0.02 | 2.028 | 2 | 1 | 1 | 1 | 1072 | 117.9 | 4.96 | 0.86 | 0.899 | 0.957 | -0.22 | 0.22 | -0.15 | -0.05 | 0.185 | -0.06 | 0.554410446 | 0.654246173 | 0.868412622 | 0.999652205 | 0.995559182 | 0.997605693 | 0.992535481 |
| Sec14l1 | Medium | A8Y5H7 | SEC14-like protein 1 OS=Mus musculus OX=10090 GN=Sec14l1 PE=1 SV=1 | 0.028 | 1.936 | 1 | 1 | 1 | 1 | 715 | 81.2 | 6.34 | 0.875 | 0.881 | 0.993 | -0.19 | 0.19 | -0.18 | -0.06 | 0.185 | -0.01 | 0.524554468 | 0.610386219 | 0.991581393 | 0.999652205 | 0.995559182 | 0.997605693 | 0.999783808 |
| Gart | High | Q64737 | Trifunctional purine biosynthetic protein adenosine-3 OS=Mus musculus OX=10090 GN=Gart PE=1 | 0 | 62.697 | 16 | 14 | 18 | 14 | 1010 | 107.4 | 6.68 | 0.943 | 0.825 | 1.144 | -0.08 | 0.08 | -0.28 | -0.0933333 | 0.18 | 0.19 | 0.846751272 | 0.498393931 | 0.333996789 | 0.999652205 | 0.995559182 | 0.997605693 | 0.906329766 |
| Gipc1 | High | Q9Z0G0 | PDZ domain-containing protein GIPC1 OS=Mus musculus OX=10090 GN=Gipc1 PE=1 SV=1 | 0 | 11.992 | 12 | 3 | 3 | 3 | 333 | 36.1 | 5.91 | 0.943 | 0.823 | 1.146 | -0.08 | 0.08 | -0.28 | -0.0933333 | 0.18 | 0.2 | 0.844618643 | 0.493461818 | 0.417785546 | 0.999652205 | 0.995559182 | 0.997605693 | 0.936397673 |
| Tcp1 | High | P11983 | T-complex protein 1 subunit alpha OS=Mus musculus OX=10090 GN=Tcp1 PE=1 SV=3 | 0 | 90.481 | 40 | 18 | 28 | 18 | 556 | 60.4 | 6.16 | 0.951 | 0.817 | 1.163 | -0.07 | 0.07 | -0.29 | -0.0966667 | 0.18 | 0.22 | 0.866915711 | 0.481349971 | 0.283600086 | 0.999652205 | 0.995559182 | 0.997605693 | 0.886074068 |
| Arf5 | High | P84084 | ADP-ribosylation factor 5 OS=Mus musculus OX=10090 GN=Arf5 PE=1 SV=2 | 0 | 26.775 | 33 | 5 | 8 | 2 | 180 | 20.5 | 6.79 | 0.959 | 0.81 | 1.184 | -0.06 | 0.06 | -0.3 | -0.1 | 0.18 | 0.24 | 0.833469731 | 0.457980103 | 0.312680775 | 0.999652205 | 0.995559182 | 0.997605693 | 0.904943958 |
| Abca8a | High | F8WGE3 | ABC-type organic anion transporter ABCA8A OS=Mus musculus OX=10090 GN=Abca8a PE=1 SV=1 | 0 | 15.713 | 3 | 5 | 5 | 3 | 1619 | 183.7 | 7.36 | 0.942 | 0.828 | 1.137 | -0.09 | 0.09 | -0.27 | -0.09 | 0.18 | 0.19 | 0.730458086 | 0.502615164 | 0.383601484 | 0.999652205 | 0.995559182 | 0.997605693 | 0.926692597 |
| Abi2 | High | P62484 | Abl interactor 2 OS=Mus musculus OX=10090 GN=Abi2 PE=1 SV=1 | 0.001 | 5.514 | 4 | 2 | 4 | 1 | 446 | 49.4 | 6.01 | 0.934 | 0.837 | 1.116 | -0.1 | 0.1 | -0.26 | -0.0866667 | 0.18 | 0.16 | 0.720588792 | 0.521591085 | 0.539850888 | 0.999652205 | 0.995559182 | 0.997605693 | 0.96694628 |
| Agfg1 | High | A0A087W | Arf-GAP domain and FG repeat-containing protein 1 OS=Mus musculus OX=10090 GN=Agfg1 PE= | 0.001 | 7.426 | 7 | 3 | 4 | 2 | 521 | 53.9 | 8.92 | 0.848 | 0.917 | 0.925 | -0.24 | 0.24 | -0.12 | -0.04 | 0.18 | -0.11 | 0.483795801 | 0.714460633 | 0.678621129 | 0.999652205 | 0.995559182 | 0.997605693 | 0.990892882 |
| Cndp2 | High | Q9D1A2 | Cytosolic non-specific dipeptidase OS=Mus musculus OX=10090 GN=Cndp2 PE=1 SV=1 | 0 | 70.253 | 43 | 13 | 20 | 13 | 475 | 52.7 | 5.66 | 0.85 | 0.921 | 0.922 | -0.24 | 0.24 | -0.12 | -0.04 | 0.18 | -0.12 | 0.631040811 | 0.735357964 | 0.699980073 | 0.999652205 | 0.995559182 | 0.997605693 | 0.991285312 |
| Aldh2 | High | P47738 | Aldehyde dehydrogenase, mitochondrial OS=Mus musculus OX=10090 GN=Aldh2 PE=1 SV=1 | 0 | 18.99 | 11 | 6 | 6 | 5 | 519 | 56.5 | 7.62 | 0.908 | 0.857 | 1.058 | -0.14 | 0.14 | -0.22 | -0.0733333 | 0.18 | 0.08 | 0.746111007 | 0.577238579 | 0.742009795 | 0.999652205 | 0.995559182 | 0.997605693 | 0.992535481 |
| Wipf3 | High | P0C7L0 | WAS/WASL-interacting protein family member 3 OS=Mus musculus OX=10090 GN=Wipf3 PE=1 S | 0 | 11.842 | 8 | 3 | 3 | 3 | 485 | 49.4 | 10.18 | 0.898 | 0.863 | 1.041 | -0.15 | 0.15 | -0.21 | -0.07 | 0.18 | 0.06 | 0.607971644 | 0.566663507 | 0.806579466 | 0.999652205 | 0.995559182 | 0.997605693 | 0.992535481 |
| Gulo | High | P58710 | L-gulonolactone oxidase OS=Mus musculus OX=10090 GN=Gulo PE=1 SV=3 | 0.006 | 3.241 | 2 | 1 | 1 | 1 | 440 | 50.4 | 8.24 | 0.865 | 0.899 | 0.962 | -0.21 | 0.21 | -0.15 | -0.05 | 0.18 | -0.06 | 0.509219936 | 0.679859957 | 0.860354796 | 0.999652205 | 0.995559182 | 0.997605693 | 0.992535481 |
| Kctd8 | High | Q50H33 | BTB/POZ domain-containing protein KCTD8 OS=Mus musculus OX=10090 GN=Kctd8 PE=1 SV=1 | 0 | 12.639 | 9 | 2 | 2 | 2 | 476 | 52.7 | 8.38 | 0.866 | 0.899 | 0.963 | -0.21 | 0.21 | -0.15 | -0.05 | 0.18 | -0.05 | 0.471934617 | 0.657233098 | 0.871742311 | 0.999652205 | 0.995559182 | 0.997605693 | 0.992535481 |
| Nt5c2 | High | E9Q9M1 | Cytosolic purine 5'-nucleotidase OS=Mus musculus OX=10090 GN=Nt5c2 PE=1 SV=1 | 0 | 11.361 | 7 | 4 | 4 | 4 | 586 | 67.7 | 6.74 | 0.893 | 0.874 | 1.023 | -0.16 | 0.16 | -0.2 | -0.0666667 | 0.18 | 0.03 | 0.654101061 | 0.616662171 | 0.881311909 | 0.999652205 | 0.995559182 | 0.997605693 | 0.992535481 |
| Iglon5 | High | Q8HW98 | IgLON family member 5 OS=Mus musculus OX=10090 GN=Iglon5 PE=2 SV=2 | 0 | 9.215 | 11 | 3 | 3 | 3 | 336 | 36.7 | 7.69 | 0.864 | 0.899 | 0.961 | -0.21 | 0.21 | -0.15 | -0.05 | 0.18 | -0.06 | 0.53140603 | 0.657974199 | 0.895708996 | 0.999652205 | 0.995559182 | 0.997605693 | 0.992535481 |
| Nans | High | Q99J77 | Sialic acid synthase OS=Mus musculus OX=10090 GN=Nans PE=1 SV=1 | 0.001 | 5.142 | 7 | 2 | 2 | 2 | 359 | 40 | 7.06 | 0.874 | 0.895 | 0.976 | -0.2 | 0.2 | -0.16 | -0.0533333 | 0.18 | -0.03 | 0.541023415 | 0.669572903 | 0.927678117 | 0.999652205 | 0.995559182 | 0.997605693 | 0.992535481 |
| Prkar2b | High | P31324 | cAMP-dependent protein kinase type II-beta regulatory subunit OS=Mus musculus OX=10090 GN= | 0 | 15.845 | 10 | 3 | 4 | 2 | 416 | 46.1 | 4.98 | 0.964 | 0.814 | 1.184 | -0.05 | 0.05 | -0.3 | -0.1 | 0.175 | 0.24 | 0.838951917 | 0.456792802 | 0.407962763 | 0.999652205 | 0.995559182 | 0.997605693 | 0.936397673 |
| | Low | A0A3B2W | CD2-associated protein OS=Mus musculus OX=10090 GN=Cd2ap PE=1 SV=1 | 0.073 | 1.263 | 1 | 1 | 1 | 1 | 532 | 58.4 | 6.14 | 0.935 | 0.84 | 1.114 | -0.1 | 0.1 | -0.25 | -0.0833333 | 0.175 | 0.16 | 0.771934753 | 0.533907381 | 0.471715165 | 0.999652205 | 0.995559182 | 0.997605693 | 0.961211268 |
| Itgb1 | High | P09055 | Integrin beta-1 OS=Mus musculus OX=10090 GN=Itgb1 PE=1 SV=1 | 0 | 26.723 | 10 | 7 | 10 | 7 | 798 | 88.2 | 5.94 | 0.939 | 0.834 | 1.125 | -0.09 | 0.09 | -0.26 | -0.0866667 | 0.175 | 0.17 | 0.76337792 | 0.52146704 | 0.513134437 | 0.999652205 | 0.995559182 | 0.997605693 | 0.964378027 |
| Adal | High | Q80SY6 | Adenosine deaminase-like protein OS=Mus musculus OX=10090 GN=Adal PE=2 SV=1 | 0.001 | 6.799 | 7 | 2 | 2 | 2 | 360 | 41 | 6.35 | 0.906 | 0.863 | 1.05 | -0.14 | 0.14 | -0.21 | -0.07 | 0.175 | 0.07 | 0.642212243 | 0.585231944 | 0.76144937 | 0.999652205 | 0.995559182 | 0.997605693 | 0.992535481 |
| Myo6 | High | E9Q175 | Unconventional myosin-6 OS=Mus musculus OX=10090 GN=Myo6 PE=1 SV=1 | 0 | 119.442 | 32 | 34 | 46 | 34 | 1253 | 144.7 | 8.51 | 0.896 | 0.876 | 1.022 | -0.16 | 0.16 | -0.19 | -0.0633333 | 0.175 | 0.03 | 0.714370341 | 0.623837269 | 0.807864537 | 0.999652205 | 0.995559182 | 0.997605693 | 0.992535481 |
| | High | A0A494B9 | Matrin-3 (Fragment) OS=Mus musculus OX=10090 GN=Matr3 PE=1 SV=1 | 0 | 14.026 | 31 | 3 | 3 | 3 | 155 | 16.4 | 9.52 | 0.898 | 0.873 | 1.029 | -0.15 | 0.15 | -0.2 | -0.0666667 | 0.175 | 0.04 | 0.750219564 | 0.615732186 | 0.832659432 | 0.999652205 | 0.995559182 | 0.997605693 | 0.992535481 |
| | High | E9QLL2 | Dynamin GTPase OS=Mus musculus OX=10090 GN=Dnm3 PE=1 SV=1 | 0 | 133.706 | 33 | 28 | 46 | 14 | 863 | 97.2 | 8.56 | 0.867 | 0.907 | 0.956 | -0.21 | 0.21 | -0.14 | -0.0466667 | 0.175 | -0.06 | 0.634639095 | 0.700938653 | 0.849704809 | 0.999652205 | 0.995559182 | 0.997605693 | 0.992535481 |
| | High | Z4YKC4 | Eukaryotic translation initiation factor 4 gamma 3 OS=Mus musculus OX=10090 GN=Eif4g3 PE=1 | 0.001 | 7.245 | 2 | 4 | 4 | 3 | 1567 | 173.4 | 5.53 | 0.891 | 0.883 | 1.009 | -0.17 | 0.17 | -0.18 | -0.06 | 0.175 | 0.01 | 0.62764619 | 0.639777049 | 0.91429916 | 0.999652205 | 0.995559182 | 0.997605693 | 0.992535481 |
| | High | A0A0A0M | Phosphoinositide phospholipase C OS=Mus musculus OX=10090 GN=Plch1 PE=1 SV=1 | 0.001 | 6.789 | 2 | 2 | 2 | 1 | 1073 | 122.5 | 7.72 | 0.885 | 0.888 | 0.997 | -0.18 | 0.18 | -0.17 | -0.0566667 | 0.175 | 0 | 0.565625543 | 0.633700782 | 0.947387934 | 0.999652205 | 0.995559182 | 0.997605693 | 0.992535481 |
| Nqo2 | High | A0A1Y7VL | Ribosyldihydronicotinamide dehydrogenase [quinone] OS=Mus musculus OX=10090 GN=Nqo2 PE= | 0.005 | 3.589 | 6 | 1 | 1 | 1 | 187 | 21.5 | 6.2 | 0.955 | 0.827 | 1.154 | -0.07 | 0.07 | -0.27 | -0.09 | 0.17 | 0.21 | 0.817080494 | 0.484143902 | 0.417741509 | 0.999652205 | 0.995559182 | 0.997605693 | 0.936397673 |
| Slc6a11 | High | P31650 | Sodium- and chloride-dependent GABA transporter 3 OS=Mus musculus OX=10090 GN=Slc6a11 P | 0 | 62.842 | 12 | 9 | 30 | 7 | 627 | 69.9 | 6.98 | 0.908 | 0.87 | 1.044 | -0.14 | 0.14 | -0.2 | -0.0666667 | 0.17 | 0.06 | 0.747075721 | 0.607445112 | 0.707319997 | 0.999652205 | 0.995559182 | 0.997605693 | 0.992485323 |
| Lrrc4b | High | P0C192 | Leucine-rich repeat-containing protein 4B OS=Mus musculus OX=10090 GN=Lrrc4b PE=1 SV=1 | 0.003 | 3.896 | 3 | 2 | 2 | 2 | 709 | 76.1 | 7.24 | 0.907 | 0.869 | 1.044 | -0.14 | 0.14 | -0.2 | -0.0666667 | 0.17 | 0.06 | 0.590891831 | 0.581142193 | 0.739561804 | 0.999652205 | 0.995559182 | 0.997605693 | 0.992535481 |
| Phlpp2 | High | J3QM82 | PH domain leucine-rich repeat-containing protein phosphatase 2 OS=Mus musculus OX=10090 GN | 0 | 12.548 | 3 | 4 | 4 | 4 | 1355 | 149.5 | 5.95 | 0.89 | 0.886 | 1.004 | -0.17 | 0.17 | -0.17 | -0.0566667 | 0.17 | 0.01 | 0.688359916 | 0.648020567 | 0.930053133 | 0.999652205 | 0.995559182 | 0.997605693 | 0.992535481 |
| Atp6v1g1 | High | Q9CR51 | V-type proton ATPase subunit G 1 OS=Mus musculus OX=10090 GN=Atp6v1g1 PE=1 SV=3 | 0 | 16.397 | 29 | 3 | 6 | 3 | 118 | 13.7 | 7.97 | 0.969 | 0.819 | 1.183 | -0.05 | 0.05 | -0.29 | -0.0966667 | 0.17 | 0.24 | 0.939014098 | 0.485126316 | 0.322349835 | 0.999652205 | 0.995559182 | 0.997605693 | 0.905686588 |
| Ahcyl1 | High | Q80SW1 | S-adenosylhomocysteine hydrolase-like protein 1 OS=Mus musculus OX=10090 GN=Ahcyl1 PE=1 S | 0 | 78.195 | 33 | 19 | 29 | 9 | 530 | 58.9 | 6.89 | 0.938 | 0.843 | 1.113 | -0.09 | 0.09 | -0.25 | -0.0833333 | 0.17 | 0.15 | 0.831673234 | 0.542145069 | 0.431030439 | 0.999652205 | 0.995559182 | 0.997605693 | 0.945905156 |
| Stk25 | High | Q9Z2W1 | Serine/threonine-protein kinase 25 OS=Mus musculus OX=10090 GN=Stk25 PE=1 SV=2 | 0 | 17.675 | 13 | 5 | 5 | 3 | 426 | 48.1 | 6.81 | 0.932 | 0.846 | 1.102 | -0.1 | 0.1 | -0.24 | -0.08 | 0.17 | 0.14 | 0.671939697 | 0.524841937 | 0.490034267 | 0.999652205 | 0.995559182 | 0.997605693 | 0.964378027 |
| Abca13 | Medium | Q5SSE9 | ATP-binding cassette sub-family A member 13 OS=Mus musculus OX=10090 GN=Abca13 PE=1 SV | 0.029 | 1.914 | 0 | 1 | 2 | 1 | 5034 | 568.5 | 6.52 | 0.921 | 0.859 | 1.072 | -0.12 | 0.12 | -0.22 | -0.0733333 | 0.17 | 0.1 | 0.760504943 | 0.58190389 | 0.637626664 | 0.999652205 | 0.995559182 | 0.997605693 | 0.983597794 |
| Synpr | High | Q8BGN8 | Synaptoporin OS=Mus musculus OX=10090 GN=Synpr PE=1 SV=1 | 0 | 42.082 | 32 | 6 | 17 | 6 | 265 | 29.2 | 6.3 | 0.859 | 0.922 | 0.931 | -0.22 | 0.22 | -0.12 | -0.04 | 0.17 | -0.1 | 0.610608249 | 0.73874106 | 0.718255711 | 0.999652205 | 0.995559182 | 0.997605693 | 0.992535481 |
| | Low | D3YWK1 | WD repeat domain phosphoinositide-interacting protein 2 OS=Mus musculus OX=10090 GN=Wipi | 0.082 | 1.191 | 2 | 1 | 1 | 1 | 425 | 46.4 | 5.85 | 0.864 | 0.917 | 0.943 | -0.21 | 0.21 | -0.13 | -0.0433333 | 0.17 | -0.08 | 0.531001357 | 0.701259883 | 0.796973158 | 0.999652205 | 0.995559182 | 0.997605693 | 0.992535481 |
| Psmd11 | High | Q8BG32 | 26S proteasome non-ATPase regulatory subunit 11 OS=Mus musculus OX=10090 GN=Psmd11 PE= | 0 | 29.497 | 23 | 10 | 12 | 10 | 422 | 47.4 | 6.48 | 0.987 | 0.807 | 1.223 | -0.02 | 0.02 | -0.31 | -0.1033333 | 0.165 | 0.29 | 0.907012184 | 0.45701791 | 0.235187178 | 0.999652205 | 0.995559182 | 0.997605693 | 0.826767714 |
| Commd6 | High | Q3V4B5 | COMM domain-containing protein 6 OS=Mus musculus OX=10090 GN=Commd6 PE=1 SV=1 | 0.009 | 2.801 | 13 | 1 | 1 | 1 | 87 | 9.8 | 5.38 | 0.967 | 0.823 | 1.175 | -0.05 | 0.05 | -0.28 | -0.0933333 | 0.165 | 0.23 | 0.908590584 | 0.494164979 | 0.344709847 | 0.999652205 | 0.995559182 | 0.997605693 | 0.907770836 |
| Gfra2 | High | O08842 | GDNF family receptor alpha-2 OS=Mus musculus OX=10090 GN=Gfra2 PE=1 SV=2 | 0.005 | 3.807 | 2 | 1 | 1 | 1 | 464 | 51.7 | 7.75 | 0.961 | 0.828 | 1.16 | -0.06 | 0.06 | -0.27 | -0.09 | 0.165 | 0.21 | 0.829694889 | 0.483934956 | 0.394477921 | 0.999652205 | 0.995559182 | 0.997605693 | 0.93262326 |
| Arpc5 | High | Q9CPW4 | Actin-related protein 2/3 complex subunit 5 OS=Mus musculus OX=10090 GN=Arpc5 PE=1 SV=3 | 0 | 27.648 | 48 | 7 | 10 | 7 | 151 | 16.3 | 5.67 | 0.944 | 0.841 | 1.123 | -0.08 | 0.08 | -0.25 | -0.0833333 | 0.165 | 0.17 | 0.807665895 | 0.53711165 | 0.459696874 | 0.999652205 | 0.995559182 | 0.997605693 | 0.959744396 |
| Vat1 | High | Q62465 | Synaptic vesicle membrane protein VAT-1 homolog OS=Mus musculus OX=10090 GN=Vat1 PE=1 | 0 | 40.491 | 23 | 7 | 8 | 7 | 406 | 43.1 | 6.37 | 0.856 | 0.927 | 0.924 | -0.22 | 0.22 | -0.11 | -0.0366667 | 0.165 | -0.11 | 0.522311197 | 0.750951153 | 0.613185562 | 0.999652205 | 0.995559182 | 0.997605693 | 0.97826833 |
| Ppp2r5d | High | Q91V89 | Serine/threonine-protein phosphatase 2A 56 kDa regulatory subunit OS=Mus musculus OX=10090 | 0 | 16.32 | 11 | 6 | 6 | 4 | 594 | 69 | 7.96 | 0.926 | 0.856 | 1.081 | -0.11 | 0.11 | -0.22 | -0.0733333 | 0.165 | 0.11 | 0.716319795 | 0.574676314 | 0.653700481 | 0.999652205 | 0.995559182 | 0.997605693 | 0.985010103 |
| Maged1 | High | Q9QYH6 | Melanoma-associated antigen D1 OS=Mus musculus OX=10090 GN=Maged1 PE=1 SV=1 | 0.001 | 7.263 | 4 | 3 | 3 | 3 | 775 | 85.6 | 7.5 | 0.855 | 0.935 | 0.915 | -0.23 | 0.23 | -0.1 | -0.0333333 | 0.165 | -0.13 | 0.519987773 | 0.767887613 | 0.653874659 | 0.999652205 | 0.995559182 | 0.997605693 | 0.985010103 |
| Palm | High | Q9Z0P4 | Paralemmin-1 OS=Mus musculus OX=10090 GN=Palm PE=1 SV=1 | 0 | 76.606 | 43 | 12 | 20 | 12 | 383 | 41.6 | 4.84 | 0.861 | 0.927 | 0.93 | -0.22 | 0.22 | -0.11 | -0.0366667 | 0.165 | -0.11 | 0.577296533 | 0.749485736 | 0.698171718 | 0.999652205 | 0.995559182 | 0.997605693 | 0.991285312 |
| Spnb1; Spt | High | Q3UGX2 | Spectrin beta chain OS=Mus musculus OX=10090 GN=Sptb PE=1 SV=1 | 0 | 51.91 | 7 | 15 | 18 | 10 | 2329 | 267.9 | 5.49 | 0.911 | 0.871 | 1.046 | -0.13 | 0.13 | -0.2 | -0.0666667 | 0.165 | 0.07 | 0.716285219 | 0.609646487 | 0.779214935 | 0.999652205 | 0.995559182 | 0.997605693 | 0.992535481 |
| Hbxip; Lam | High | Q9D1L9 | Ragulator complex protein LAMTOR5 OS=Mus musculus OX=10090 GN=Lamtor5 PE=1 SV=1 | 0 | 26.135 | 81 | 4 | 8 | 4 | 91 | 9.6 | 4.87 | 0.87 | 0.916 | 0.95 | -0.2 | 0.2 | -0.13 | -0.0433333 | 0.165 | -0.07 | 0.618916584 | 0.723023558 | 0.800926355 | 0.999652205 | 0.995559182 | 0.997605693 | 0.992535481 |
| Fmnl2 | Medium | F8VPR2 | Formin-like protein 2 OS=Mus musculus OX=10090 GN=Fmnl2 PE=1 SV=1 | 0.035 | 1.77 | 1 | 1 | 1 | 1 | 1083 | 122.8 | 7.4 | 0.898 | 0.887 | 1.012 | -0.16 | 0.16 | -0.17 | -0.0566667 | 0.165 | 0.02 | 0.643497536 | 0.644787078 | 0.951868812 | 0.999652205 | 0.995559182 | 0.997605693 | 0.992535481 |
| | High | A0A2I3BR | Glia maturation factor beta (Fragment) OS=Mus musculus OX=10090 GN=Gmfb PE=1 SV=1 | 0.001 | 7.869 | 20 | 3 | 4 | 3 | 137 | 15.9 | 5.44 | 0.895 | 0.888 | 1.008 | -0.16 | 0.16 | -0.17 | -0.0566667 | 0.165 | 0.01 | 0.63104298 | 0.653827864 | 0.955107927 | 0.999652205 | 0.995559182 | 0.997605693 | 0.992884189 |
| Syn1 | High | O88935 | Synapsin-1 OS=Mus musculus OX=10090 GN=Syn1 PE=1 SV=2 | 0 | 147.01 | 45 | 25 | 61 | 23 | 706 | 74.1 | 9.8 | 0.863 | 0.922 | 0.936 | -0.21 | 0.21 | -0.12 | -0.04 | 0.165 | -0.1 | 0.622045757 | 0.737152114 | 0.745002038 | 0.999652205 | 0.995559182 | 0.997605693 | 0.992535481 |
| Tln2 | High | A0A1L1SQ | Talin-2 OS=Mus musculus OX=10090 GN=Tln2 PE=1 SV=1 | 0 | 87.211 | 11 | 25 | 28 | 18 | 2544 | 271.8 | 5.57 | 0.899 | 0.884 | 1.017 | -0.15 | 0.15 | -0.18 | -0.06 | 0.165 | 0.02 | 0.722582005 | 0.643175805 | 0.835886141 | 0.999652205 | 0.995559182 | 0.997605693 | 0.992535481 |
| Plekhf1 | Medium | Q3TB82 | Pleckstrin homology domain-containing family F member 1 OS=Mus musculus OX=10090 GN=Ple | 0.035 | 1.783 | 4 | 1 | 1 | 1 | 279 | 31.1 | 8.1 | 0.904 | 0.885 | 1.022 | -0.15 | 0.15 | -0.18 | -0.06 | 0.165 | 0.03 | 0.564986257 | 0.622424254 | 0.843652438 | 0.999652205 | 0.995559182 | 0.997605693 | 0.992535481 |
| Cct4 | High | P80315 | T-complex protein 1 subunit delta OS=Mus musculus OX=10090 GN=Cct4 PE=1 SV=3 | 0 | 119.74 | 48 | 22 | 39 | 22 | 539 | 58 | 8.02 | 0.957 | 0.837 | 1.143 | -0.06 | 0.06 | -0.26 | -0.0866667 | 0.16 | 0.19 | 0.885489244 | 0.528249609 | 0.335515846 | 0.999652205 | 0.995559182 | 0.997605693 | 0.906329766 |
| Psmc5 | High | P62196 | 26S proteasome regulatory subunit 8 OS=Mus musculus OX=10090 GN=Psmc5 PE=1 SV=1 | 0 | 21.014 | 16 | 5 | 5 | 4 | 406 | 45.6 | 7.55 | 0.943 | 0.845 | 1.115 | -0.08 | 0.08 | -0.24 | -0.08 | 0.16 | 0.16 | 0.659750614 | 0.523108077 | 0.454733438 | 0.999652205 | 0.995559182 | 0.997605693 | 0.959744396 |
| | High | A0A286YD | Myoferlin OS=Mus musculus OX=10090 GN=Myof PE=1 SV=1 | 0 | 25.583 | 6 | 9 | 9 | 9 | 2061 | 234.4 | 6.14 | 0.945 | 0.848 | 1.114 | -0.08 | 0.08 | -0.24 | -0.08 | 0.16 | 0.16 | 0.826398553 | 0.554005528 | 0.487968204 | 0.999652205 | 0.995559182 | 0.997605693 | 0.964378027 |
| Nbea | High | Q9EPN1 | Neurobeachin OS=Mus musculus OX=10090 GN=Nbea PE=1 SV=1 | 0 | 28.32 | 3 | 7 | 7 | 5 | 2936 | 326.5 | 6.2 | 0.943 | 0.846 | 1.114 | -0.08 | 0.08 | -0.24 | -0.08 | 0.16 | 0.16 | 0.765519093 | 0.550104203 | 0.558339407 | 0.999652205 | 0.995559182 | 0.997605693 | 0.971936993 |
| Ppp2r1a | High | Q76MZ3 | Serine/threonine-protein phosphatase 2A 65 kDa regulatory subunit A alpha isoform OS=Mus mu | 0 | 151.511 | 52 | 26 | 45 | 26 | 589 | 65.3 | 5.11 | 0.923 | 0.872 | 1.058 | -0.12 | 0.12 | -0.2 | -0.0666667 | 0.16 | 0.08 | 0.790361754 | 0.61401389 | 0.64244363 | 0.999652205 | 0.995559182 | 0.997605693 | 0.985010103 |
| Arfgef2 | High | A2A5R2 | Brefeldin A-inhibited guanine nucleotide-exchange protein 2 OS=Mus musculus OX=10090 GN=Ar | 0 | 16.364 | 5 | 6 | 8 | 6 | 1792 | 202.1 | 6.55 | 0.866 | 0.926 | 0.935 | -0.21 | 0.21 | -0.11 | -0.0366667 | 0.16 | -0.1 | 0.536918811 | 0.748146536 | 0.657544684 | 0.999652205 | 0.995559182 | 0.997605693 | 0.985010103 |
| Dpp9 | High | Q8BVG4 | Dipeptidyl peptidase 9 OS=Mus musculus OX=10090 GN=Dpp9 PE=1 SV=2 | 0 | 12.944 | 5 | 4 | 4 | 4 | 862 | 97.9 | 6.65 | 0.926 | 0.867 | 1.069 | -0.11 | 0.11 | -0.21 | -0.07 | 0.16 | 0.1 | 0.81342246 | 0.600214774 | 0.674587253 | 0.999652205 | 0.995559182 | 0.997605693 | 0.990250803 |
| Gnai2 | High | P08752 | Guanine nucleotide-binding protein G(i) subunit alpha-2 OS=Mus musculus OX=10090 GN=Gnai2 | 0 | 146.421 | 58 | 19 | 108 | 8 | 355 | 40.5 | 5.45 | 0.905 | 0.88 | 1.028 | -0.14 | 0.14 | -0.18 | -0.06 | 0.16 | 0.04 | 0.738324211 | 0.632024627 | 0.779422417 | 0.999652205 | 0.995559182 | 0.997605693 | 0.992535481 |
| Efhd2 | High | Q9D8Y0 | EF-hand domain-containing protein D2 OS=Mus musculus OX=10090 GN=Efhd2 PE=1 SV=1 | 0 | 14.457 | 18 | 4 | 5 | 4 | 240 | 26.8 | 5.06 | 0.874 | 0.916 | 0.954 | -0.19 | 0.19 | -0.13 | -0.0433333 | 0.16 | -0.07 | 0.656630439 | 0.722243778 | 0.807268849 | 0.999652205 | 0.995559182 | 0.997605693 | 0.992535481 |
| Lsm5; LOC | Low | P62322 | U6 snRNA-associated Sm-like protein LSm5 OS=Mus musculus OX=10090 GN=Lsm5 PE=3 SV=2 | 0.054 | 1.402 | 7 | 1 | 1 | 1 | 91 | 9.9 | 4.54 | 0.875 | 0.915 | 0.956 | -0.19 | 0.19 | -0.13 | -0.0433333 | 0.16 | -0.06 | 0.535201651 | 0.719633453 | 0.831552191 | 0.999652205 | 0.995559182 | 0.997605693 | 0.992535481 |
| Pgp | High | Q8CHP8 | Glycerol-3-phosphate phosphatase OS=Mus musculus OX=10090 GN=Pgp PE=1 SV=1 | 0.001 | 7.487 | 6 | 2 | 2 | 2 | 321 | 34.5 | 5.35 | 0.875 | 0.913 | 0.958 | -0.19 | 0.19 | -0.13 | -0.0433333 | 0.16 | -0.06 | 0.558208922 | 0.693318186 | 0.870057237 | 0.999652205 | 0.995559182 | 0.997605693 | 0.992535481 |
| Capza1 | High | P47753 | F-actin-capping protein subunit alpha-1 OS=Mus musculus OX=10090 GN=Capza1 PE=1 SV=4 | 0 | 17.788 | 20 | 4 | 6 | 3 | 286 | 32.9 | 5.55 | 0.907 | 0.885 | 1.024 | -0.14 | 0.14 | -0.18 | -0.06 | 0.16 | 0.03 | 0.669956283 | 0.645806144 | 0.885046864 | 0.999652205 | 0.995559182 | 0.997605693 | 0.992535481 |
| Actr1b | High | Q8R5C5 | Beta-centractin OS=Mus musculus OX=10090 GN=Actr1b PE=1 SV=1 | 0 | 42.295 | 24 | 7 | 8 | 2 | 376 | 42.3 | 6.4 | 0.894 | 0.898 | 0.996 | -0.16 | 0.16 | -0.16 | -0.0533333 | 0.16 | -0.01 | 0.614769854 | 0.647438829 | 0.95413708 | 0.999652205 | 0.995559182 | 0.997605693 | 0.992884189 |
| Rnf214 | Low | Q8BFU3 | RING finger protein 214 OS=Mus musculus OX=10090 GN=Rnf214 PE=1 SV=1 | 0.08 | 1.207 | 1 | 1 | 1 | 1 | 668 | 73.6 | 6.23 | 0.983 | 0.823 | 1.194 | -0.03 | 0.03 | -0.28 | -0.0933333 | 0.155 | 0.26 | 0.891904918 | 0.463945525 | 0.259969421 | 0.999652205 | 0.995559182 | 0.997605693 | 0.85616924 |
| Ppp3cc | High | P48455 | Serine/threonine-protein phosphatase 2B catalytic subunit gamma isoform OS=Mus musculus OX | 0 | 75.508 | 35 | 14 | 28 | 9 | 513 | 58.7 | 7.25 | 0.981 | 0.824 | 1.19 | -0.03 | 0.03 | -0.28 | -0.0933333 | 0.155 | 0.25 | 0.91546493 | 0.496785091 | 0.266304869 | 0.999652205 | 0.995559182 | 0.997605693 | 0.867114634 |
| Psmd7 | High | P26516 | 26S proteasome non-ATPase regulatory subunit 7 OS=Mus musculus OX=10090 GN=Psmd7 PE=1 | 0.001 | 5.97 | 8 | 2 | 2 | 2 | 321 | 36.5 | 6.77 | 0.909 | 0.889 | 1.022 | -0.14 | 0.14 | -0.17 | -0.0566667 | 0.155 | 0.03 | 0.606688442 | 0.637591865 | 0.859495064 | 0.999652205 | 0.995559182 | 0.997605693 | 0.992535481 |
| Copg; Cop | High | Q9QZE5 | Coatomer subunit gamma-1 OS=Mus musculus OX=10090 GN=Copg1 PE=1 SV=1 | 0 | 10.255 | 4 | 3 | 3 | 2 | 874 | 97.5 | 5.35 | 0.886 | 0.91 | 0.974 | -0.17 | 0.17 | -0.14 | -0.0466667 | 0.155 | -0.04 | 0.615801537 | 0.707284599 | 0.922168955 | 0.999652205 | 0.995559182 | 0.997605693 | 0.992535481 |
| Ywhaq; LC | High | P68254 | 14-3-3 protein theta OS=Mus musculus OX=10090 GN=Ywhaq PE=1 SV=1 | 0 | 147.143 | 63 | 20 | 63 | 14 | 245 | 27.8 | 4.78 | 0.887 | 0.91 | 0.975 | -0.17 | 0.17 | -0.14 | -0.0466667 | 0.155 | -0.04 | 0.688620797 | 0.707587975 | 0.947650735 | 0.999652205 | 0.995559182 | 0.997605693 | 0.992535481 |
| Sec13 | High | Q9D1M0 | Protein SEC13 homolog OS=Mus musculus OX=10090 GN=Sec13 PE=1 SV=3 | 0.001 | 7.284 | 6 | 2 | 2 | 2 | 322 | 35.5 | 5.38 | 0.965 | 0.838 | 1.152 | -0.05 | 0.05 | -0.26 | -0.0866667 | 0.155 | 0.2 | 0.813796294 | 0.503523099 | 0.395001945 | 0.999652205 | 0.995559182 | 0.997605693 | 0.93262326 |
| Smc1a | High | Q9CU62 | Structural maintenance of chromosomes protein 1A OS=Mus musculus OX=10090 GN=Smc1a PE= | 0 | 14.567 | 4 | 5 | 5 | 5 | 1233 | 143.1 | 7.64 | 0.919 | 0.879 | 1.045 | -0.12 | 0.12 | -0.19 | -0.0633333 | 0.155 | 0.06 | 0.624169423 | 0.609389938 | 0.718035572 | 0.999652205 | 0.995559182 | 0.997605693 | 0.992535481 |
| Pfkm | High | P47857 | ATP-dependent 6-phosphofructokinase, muscle type OS=Mus musculus OX=10090 GN=Pfkm PE=1 | 0 | 106.382 | 28 | 21 | 42 | 16 | 780 | 85.2 | 8 | 0.87 | 0.93 | 0.936 | -0.2 | 0.2 | -0.11 | -0.0366667 | 0.155 | -0.1 | 0.642816956 | 0.757245826 | 0.745130143 | 0.999652205 | 0.995559182 | 0.997605693 | 0.992535481 |
| Rhob | High | P62746 | Rho-related GTP-binding protein RhoB OS=Mus musculus OX=10090 GN=Rhob PE=1 SV=1 | 0 | 52.599 | 37 | 8 | 17 | 6 | 196 | 22.1 | 5.24 | 0.872 | 0.93 | 0.938 | -0.2 | 0.2 | -0.11 | -0.0366667 | 0.155 | -0.09 | 0.648209738 | 0.75711757 | 0.756315688 | 0.999652205 | 0.995559182 | 0.997605693 | 0.992535481 |
| Prpsap2 | High | Q8R574 | Phosphoribosyl pyrophosphate synthase-associated protein 2 OS=Mus musculus OX=10090 GN=Pr | 0 | 40.389 | 35 | 10 | 13 | 8 | 369 | 40.9 | 7.17 | 0.904 | 0.894 | 1.012 | -0.15 | 0.15 | -0.16 | -0.0533333 | 0.155 | 0.02 | 0.770750611 | 0.666592783 | 0.86106955 | 0.999652205 | 0.995559182 | 0.997605693 | 0.992535481 |
| Snx27 | High | Q3UHD6 | Sorting nexin-27 OS=Mus musculus OX=10090 GN=Snx27 PE=1 SV=2 | 0 | 26.619 | 16 | 7 | 10 | 7 | 539 | 61 | 6.4 | 0.892 | 0.902 | 0.99 | -0.16 | 0.16 | -0.15 | -0.05 | 0.155 | -0.02 | 0.613813285 | 0.686675191 | 0.922959859 | 0.999652205 | 0.995559182 | 0.997605693 | 0.992535481 |
| Pgm3 | High | Q9CYR6 | Phosphoacetylglucosamine mutase OS=Mus musculus OX=10090 GN=Pgm3 PE=1 SV=1 | 0.005 | 3.541 | 3 | 1 | 2 | 1 | 542 | 59.4 | 6.2 | 0.868 | 0.93 | 0.932 | -0.2 | 0.2 | -0.1 | -0.0333333 | 0.15 | -0.1 | 0.617069969 | 0.758867103 | 0.70682045 | 0.999652205 | 0.995559182 | 0.997605693 | 0.992485323 |

| | | | | | | | | | | | | | | | | | | | | | | | | | | | | |
|---|---|---|---|---|---|---|---|---|---|---|---|---|---|---|---|---|---|---|---|---|---|---|---|---|---|---|---|---|
| Lims1 | Medium | Q9D7B2 | LIM and senescent cell antigen-like-containing domain protein OS=Mus musculus OX=10090 GN= | 0.029 | 1.942 | 2 | 1 | 1 | 1 | 396 | 45.6 | 7.88 | 0.915 | 0.891 | 1.026 | -0.13 | 0.13 | -0.17 | -0.0566667 | 0.15 | 0.04 | 0.668830515 | 0.635196008 | 0.810065653 | 0.999652205 | 0.995559182 | 0.997605693 | 0.992535481 |
| Tssc1; Eipr | Medium | Q8K0G5 | EARP and GARP complex-interacting protein 1 OS=Mus musculus OX=10090 GN=Eipr1 PE=1 SV=2 | 0.036 | 1.761 | 2 | 1 | 1 | 1 | 386 | 43.1 | 5.14 | 0.915 | 0.891 | 1.027 | -0.13 | 0.13 | -0.17 | -0.0566667 | 0.15 | 0.04 | 0.718123085 | 0.658187221 | 0.849066752 | 0.999652205 | 0.995559182 | 0.997605693 | 0.992535481 |
| Gfpt1 | High | P47856 | Glutamine--fructose-6-phosphate aminotransferase [isomerizing] 1 OS=Mus musculus OX=10090 | 0 | 13.26 | 9 | 6 | 6 | 6 | 697 | 78.5 | 6.84 | 0.889 | 0.916 | 0.971 | -0.17 | 0.17 | -0.13 | -0.0433333 | 0.15 | -0.04 | 0.617366032 | 0.722925062 | 0.875537613 | 0.999652205 | 0.995559182 | 0.997605693 | 0.992535481 |
| Dab2ip | Medium | A2AUX5 | Disabled homolog 2-interacting protein OS=Mus musculus OX=10090 GN=Dab2ip PE=1 SV=1 | 0.03 | 1.9 | 1 | 1 | 1 | 1 | 1124 | 123.9 | 7.28 | 0.895 | 0.909 | 0.985 | -0.16 | 0.16 | -0.14 | -0.0466667 | 0.15 | -0.02 | 0.594409293 | 0.688910516 | 0.999504948 | 0.999652205 | 0.995559182 | 0.997605693 | 0.999917596 |
| Trim2 | High | Q9ESN6 | Tripartite motif-containing protein 2 OS=Mus musculus OX=10090 GN=Trim2 PE=1 SV=1 | 0 | 29.813 | 11 | 6 | 6 | 5 | 744 | 81.4 | 6.96 | 0.966 | 0.844 | 1.145 | -0.05 | 0.05 | -0.25 | -0.0833333 | 0.15 | 0.2 | 0.873301185 | 0.54377348 | 0.371098995 | 0.999652205 | 0.995559182 | 0.997605693 | 0.920164724 |
| | High | H3BJY2 | Arf-GAP with SH3 domain, ANK repeat and PH domain-containing protein 1 OS=Mus musculus OX | 0 | 12.17 | 2 | 2 | 3 | 2 | 1144 | 126.9 | 7.43 | 0.904 | 0.904 | 1 | -0.15 | 0.15 | -0.15 | -0.05 | 0.15 | 0 | 0.532628498 | 0.664740499 | 0.951143226 | 0.999652205 | 0.995559182 | 0.997605693 | 0.992535481 |
| Cct8 | High | P42932 | T-complex protein 1 subunit theta OS=Mus musculus OX=10090 GN=Cct8 PE=1 SV=3 | 0 | 144.059 | 53 | 28 | 52 | 28 | 548 | 59.5 | 5.62 | 0.939 | 0.873 | 1.076 | -0.09 | 0.09 | -0.2 | -0.0666667 | 0.145 | 0.11 | 0.834974889 | 0.614692914 | 0.567038977 | 0.999652205 | 0.995559182 | 0.997605693 | 0.972910028 |
| Usp4 | High | P35123 | Ubiquitin carboxyl-terminal hydrolase 4 OS=Mus musculus OX=10090 GN=Usp4 PE=1 SV=3 | 0 | 13.688 | 6 | 5 | 5 | 5 | 962 | 108.3 | 5.64 | 0.889 | 0.918 | 0.968 | -0.17 | 0.17 | -0.12 | -0.04 | 0.145 | -0.05 | 0.707220622 | 0.728453681 | 0.86228737 | 0.999652205 | 0.995559182 | 0.997605693 | 0.992535481 |
| Grlf1; Arhg | High | Q91YM2 | Rho GTPase-activating protein 35 OS=Mus musculus OX=10090 GN=Arhgap35 PE=1 SV=3 | 0 | 9.349 | 2 | 3 | 4 | 3 | 1499 | 170.3 | 6.61 | 0.91 | 0.903 | 1.008 | -0.14 | 0.14 | -0.15 | -0.05 | 0.145 | 0.01 | 0.768411275 | 0.689157388 | 0.924968989 | 0.999652205 | 0.995559182 | 0.997605693 | 0.992535481 |
| Bcas3 | High | Q8CCN5 | BCAS3 microtubule associated cell migration factor OS=Mus musculus OX=10090 GN=Bcas3 PE=1 | 0 | 11.423 | 5 | 4 | 4 | 4 | 928 | 101 | 6.7 | 0.908 | 0.9 | 1.008 | -0.14 | 0.14 | -0.15 | -0.05 | 0.145 | 0.01 | 0.691753034 | 0.683707308 | 0.925161245 | 0.999652205 | 0.995559182 | 0.997605693 | 0.992535481 |
| Gng7 | High | Q61016 | Guanine nucleotide-binding protein G(I)/G(S)/G(O) subunit gamma-7 OS=Mus musculus OX=100 | 0 | 23.832 | 57 | 3 | 7 | 3 | 68 | 7.5 | 8.51 | 0.897 | 0.912 | 0.984 | -0.16 | 0.16 | -0.13 | -0.0433333 | 0.145 | -0.02 | 0.662693393 | 0.71164435 | 0.927201855 | 0.999652205 | 0.995559182 | 0.997605693 | 0.992535481 |
| | High | G3XA25 | Acetyl-CoA acetyltransferase, cytosolic OS=Mus musculus OX=10090 GN=Acat2 PE=1 SV=1 | 0.001 | 5.846 | 6 | 3 | 4 | 3 | 365 | 38.1 | 7.71 | 0.909 | 0.903 | 1.007 | -0.14 | 0.14 | -0.15 | -0.05 | 0.145 | 0.01 | 0.738891348 | 0.690114956 | 0.955166277 | 0.999652205 | 0.995559182 | 0.997605693 | 0.992884189 |
| Sept7 | High | E9Q1G8 | Septin OS=Mus musculus OX=10090 GN=Septin7 PE=1 SV=2 | 0 | 103.983 | 52 | 22 | 36 | 20 | 437 | 50.6 | 8.57 | 0.895 | 0.914 | 0.98 | -0.16 | 0.16 | -0.13 | -0.0433333 | 0.145 | -0.03 | 0.712608778 | 0.716870627 | 0.976212366 | 0.999652205 | 0.995559182 | 0.997605693 | 0.997221123 |
| Gphn | High | Q8BUV3 | Gephyrin OS=Mus musculus OX=10090 GN=Gphn PE=1 SV=2 | 0 | 33.937 | 15 | 9 | 12 | 9 | 769 | 83.2 | 5.6 | 0.903 | 0.905 | 0.997 | -0.15 | 0.15 | -0.14 | -0.0466667 | 0.145 | 0 | 0.693867799 | 0.695780006 | 0.99265141 | 0.999652205 | 0.995559182 | 0.997605693 | 0.999783808 |
| Eif1 | High | P48024 | Eukaryotic translation initiation factor 1 OS=Mus musculus OX=10090 GN=Eif1 PE=1 SV=2 | 0 | 55.605 | 63 | 8 | 13 | 8 | 113 | 12.7 | 7.44 | 1.015 | 0.807 | 1.258 | 0.02 | -0.02 | -0.31 | -0.1033333 | 0.145 | 0.33 | 0.984967792 | 0.457842377 | 0.149669874 | 0.999652205 | 0.995559182 | 0.997605693 | 0.71722748 |
| Pdpk1 | High | Q9Z2A0 | 3-phosphoinositide-dependent protein kinase 1 OS=Mus musculus OX=10090 GN=Pdpk1 PE=1 SV= | 0 | 18.434 | 13 | 6 | 7 | 6 | 559 | 63.7 | 7.37 | 1.016 | 0.805 | 1.262 | 0.02 | -0.02 | -0.31 | -0.1033333 | 0.145 | 0.34 | 0.992233214 | 0.453611531 | 0.174685875 | 0.999652205 | 0.995559182 | 0.997605693 | 0.744450291 |
| Psmd8 | High | Q9CX56 | 26S proteasome non-ATPase regulatory subunit 8 OS=Mus musculus OX=10090 GN=Psmd8 PE=1 | 0.001 | 5.692 | 5 | 2 | 3 | 2 | 353 | 39.9 | 9.58 | 1 | 0.82 | 1.22 | 0 | 0 | -0.29 | -0.0966667 | 0.145 | 0.29 | 0.980758393 | 0.487146136 | 0.259930341 | 0.999652205 | 0.995559182 | 0.997605693 | 0.85616924 |
| Cd151 | High | O35566 | CD151 antigen OS=Mus musculus OX=10090 GN=Cd151 PE=1 SV=2 | 0.001 | 4.594 | 6 | 2 | 3 | 2 | 253 | 28.2 | 7.47 | 0.948 | 0.863 | 1.099 | -0.08 | 0.08 | -0.21 | -0.07 | 0.145 | 0.14 | 0.792224659 | 0.589954178 | 0.59902329 | 0.999652205 | 0.995559182 | 0.997605693 | 0.977451587 |
| Ate1 | High | J3QNU1 | Arginyl-tRNA--protein transferase 1 OS=Mus musculus OX=10090 GN=Ate1 PE=1 SV=1 | 0.001 | 6.857 | 8 | 3 | 3 | 3 | 509 | 58.2 | 7.25 | 0.924 | 0.885 | 1.044 | -0.11 | 0.11 | -0.18 | -0.06 | 0.145 | 0.06 | 0.718839763 | 0.645067528 | 0.797120219 | 0.999652205 | 0.995559182 | 0.997605693 | 0.992535481 |
| | High | P35235 | Tyrosine-protein phosphatase non-receptor type 11 OS=Mus musculus OX=10090 GN=Ptpn11 PE= | 0 | 52.554 | 28 | 15 | 17 | 15 | 593 | 68 | 7.3 | 0.992 | 0.829 | 1.197 | -0.01 | 0.01 | -0.27 | -0.09 | 0.14 | 0.26 | 0.997281384 | 0.50877959 | 0.208381977 | 0.999652205 | 0.995559182 | 0.997605693 | 0.801370196 |
| Echdc1 | High | Q9D9V3 | Ethylmalonyl-CoA decarboxylase OS=Mus musculus OX=10090 GN=Echdc1 PE=1 SV=2 | 0 | 13.362 | 18 | 5 | 5 | 5 | 322 | 35.4 | 7.01 | 0.985 | 0.837 | 1.177 | -0.02 | 0.02 | -0.26 | -0.0866667 | 0.14 | 0.24 | 0.855412383 | 0.500949184 | 0.287812829 | 0.999652205 | 0.995559182 | 0.997605693 | 0.887894721 |
| Nhlrc2 | High | Q8BZW8 | NHL repeat-containing protein 2 OS=Mus musculus OX=10090 GN=Nhlrc2 PE=1 SV=1 | 0 | 9.554 | 3 | 2 | 2 | 2 | 725 | 78.4 | 5.54 | 0.977 | 0.843 | 1.159 | -0.03 | 0.03 | -0.25 | -0.0833333 | 0.14 | 0.21 | 0.875452075 | 0.512737784 | 0.34300931 | 0.999652205 | 0.995559182 | 0.997605693 | 0.907770836 |
| Camk1d | High | Q8BW96 | Calcium/calmodulin-dependent protein kinase type 1D OS=Mus musculus OX=10090 GN=Camk1d | 0 | 33.31 | 21 | 7 | 7 | 7 | 385 | 42.9 | 7.17 | 0.981 | 0.842 | 1.166 | -0.03 | 0.03 | -0.25 | -0.0833333 | 0.14 | 0.22 | 0.901572179 | 0.538695872 | 0.373431539 | 0.999652205 | 0.995559182 | 0.997605693 | 0.920164724 |
| Cul4a | High | Q3TCH7 | Cullin-4A OS=Mus musculus OX=10090 GN=Cul4a PE=1 SV=1 | 0.001 | 7.874 | 3 | 2 | 3 | 2 | 759 | 87.7 | 8.35 | 0.941 | 0.879 | 1.071 | -0.09 | 0.09 | -0.19 | -0.0633333 | 0.14 | 0.1 | 0.738796743 | 0.606418989 | 0.617794503 | 0.999652205 | 0.995559182 | 0.997605693 | 0.97826833 |
| Tceb1; Elo | High | P83940 | Elongin-C OS=Mus musculus OX=10090 GN=Eloc PE=1 SV=1 | 0 | 54.176 | 58 | 5 | 12 | 5 | 112 | 12.5 | 4.78 | 0.933 | 0.885 | 1.054 | -0.1 | 0.1 | -0.18 | -0.06 | 0.14 | 0.08 | 0.876375601 | 0.646391411 | 0.660800071 | 0.999652205 | 0.995559182 | 0.997605693 | 0.98592496 |
| Dpysl5 | High | Q9EQF6 | Dihydropyrimidinase-related protein 5 OS=Mus musculus OX=10090 GN=Dpysl5 PE=1 SV=1 | 0 | 17.77 | 9 | 5 | 6 | 5 | 564 | 61.5 | 7.09 | 0.886 | 0.929 | 0.954 | -0.17 | 0.17 | -0.11 | -0.0366667 | 0.14 | -0.07 | 0.640359595 | 0.754982592 | 0.761705937 | 0.999652205 | 0.995559182 | 0.997605693 | 0.992535481 |
| Rab34 | High | B1AQD4 | Ras-related protein Rab-34 (Fragment) OS=Mus musculus OX=10090 GN=Rab34 PE=1 SV=1 | 0.005 | 3.49 | 5 | 1 | 1 | 1 | 265 | 30 | 8.69 | 0.92 | 0.896 | 1.027 | -0.12 | 0.12 | -0.16 | -0.0533333 | 0.14 | 0.04 | 0.673811327 | 0.672258525 | 0.826817861 | 0.999652205 | 0.995559182 | 0.997605693 | 0.992535481 |
| Oplah | High | Q8K010 | 5-oxoprolinase OS=Mus musculus OX=10090 GN=Oplah PE=1 SV=1 | 0.001 | 5.992 | 1 | 2 | 2 | 2 | 1288 | 137.5 | 6.28 | 0.911 | 0.899 | 1.013 | -0.13 | 0.13 | -0.15 | -0.05 | 0.14 | 0.02 | 0.613713989 | 0.666891261 | 0.896777523 | 0.999652205 | 0.995559182 | 0.997605693 | 0.992535481 |
| Arhgap17 | High | E9QAJ9 | Rho GTPase-activating protein 17 OS=Mus musculus OX=10090 GN=Arhgap17 PE=1 SV=1 | 0 | 10.513 | 5 | 3 | 3 | 2 | 818 | 88.9 | 8.12 | 0.91 | 0.91 | 1 | -0.14 | 0.14 | -0.14 | -0.0466667 | 0.14 | 0 | 0.720614991 | 0.707834592 | 0.952156688 | 0.999652205 | 0.995559182 | 0.997605693 | 0.992535481 |
| Gng3 | High | P63216 | Guanine nucleotide-binding protein G(I)/G(S)/G(O) subunit gamma-3 OS=Mus musculus OX=100 | 0 | 40.539 | 72 | 4 | 10 | 4 | 75 | 8.3 | 7.78 | 0.907 | 0.906 | 1.001 | -0.14 | 0.14 | -0.14 | -0.0466667 | 0.14 | 0 | 0.6590515 | 0.698805273 | 0.976851903 | 0.999652205 | 0.995559182 | 0.997605693 | 0.997221123 |
| | High | A0A3B2W | Serine/threonine-protein kinase 38 (Fragment) OS=Mus musculus OX=10090 GN=Stk38 PE=1 SV= | 0.001 | 4.249 | 5 | 1 | 1 | 1 | 261 | 30.1 | 5.82 | 0.965 | 0.86 | 1.122 | -0.05 | 0.05 | -0.22 | -0.0733333 | 0.135 | 0.17 | 0.82296045 | 0.564637479 | 0.495492179 | 0.999652205 | 0.995559182 | 0.997605693 | 0.964378027 |
| Acyp2 | High | P56375 | Acylphosphatase-2 OS=Mus musculus OX=10090 GN=Acyp2 PE=1 SV=2 | 0 | 18.09 | 42 | 5 | 6 | 5 | 106 | 11.9 | 9.26 | 0.963 | 0.862 | 1.117 | -0.06 | 0.06 | -0.21 | -0.07 | 0.135 | 0.16 | 0.870782484 | 0.588525439 | 0.519365944 | 0.999652205 | 0.995559182 | 0.997605693 | 0.964378027 |
| Xpo1 | High | Q6P5F9 | Exportin-1 OS=Mus musculus OX=10090 GN=Xpo1 PE=1 SV=1 | 0 | 10.263 | 4 | 3 | 3 | 3 | 1071 | 123 | 6.07 | 0.955 | 0.868 | 1.099 | -0.07 | 0.07 | -0.2 | -0.0666667 | 0.135 | 0.14 | 0.854522582 | 0.601301169 | 0.577184548 | 0.999652205 | 0.995559182 | 0.997605693 | 0.972910028 |
| Macf1 | High | B1ARU4 | Microtubule-actin cross-linking factor 1 OS=Mus musculus OX=10090 GN=Macf1 PE=1 SV=1 | 0 | 39.608 | 2 | 14 | 14 | 14 | 7353 | 831.1 | 5.41 | 0.932 | 0.89 | 1.047 | -0.1 | 0.1 | -0.17 | -0.0566667 | 0.135 | 0.07 | 0.750564319 | 0.658142038 | 0.808872326 | 0.999652205 | 0.995559182 | 0.997605693 | 0.992535481 |
| | High | E9PX30 | Mitogen-activated protein kinase kinase kinase kinase 5 OS=Mus musculus OX=10090 GN=Map4k | 0.001 | 4.44 | 2 | 1 | 1 | 1 | 847 | 94.9 | 7.91 | 0.894 | 0.926 | 0.965 | -0.16 | 0.16 | -0.11 | -0.0366667 | 0.135 | -0.05 | 0.664622468 | 0.742691851 | 0.865835718 | 0.999652205 | 0.995559182 | 0.997605693 | 0.992535481 |
| Ppp2cb | High | P62715 | Serine/threonine-protein phosphatase 2A catalytic subunit beta isoform OS=Mus musculus OX=1 | 0 | 54.494 | 42 | 11 | 16 | 1 | 309 | 35.6 | 5.43 | 0.914 | 0.91 | 1.004 | -0.13 | 0.13 | -0.14 | -0.0466667 | 0.135 | 0.01 | 0.753953754 | 0.708834108 | 0.909172291 | 0.999652205 | 0.995559182 | 0.997605693 | 0.992535481 |
| Pygb | High | Q8CI94 | Glycogen phosphorylase, brain form OS=Mus musculus OX=10090 GN=Pygb PE=1 SV=3 | 0 | 114.642 | 40 | 28 | 37 | 20 | 843 | 96.7 | 6.73 | 0.905 | 0.914 | 0.99 | -0.14 | 0.14 | -0.13 | -0.0433333 | 0.135 | -0.01 | 0.738255878 | 0.717332265 | 0.972580654 | 0.999652205 | 0.995559182 | 0.997605693 | 0.996575243 |
| Add2 | High | Q9QYB8 | Beta-adducin OS=Mus musculus OX=10090 GN=Add2 PE=1 SV=4 | 0 | 8.653 | 6 | 4 | 4 | 3 | 725 | 80.6 | 6.21 | 0.902 | 0.921 | 0.979 | -0.15 | 0.15 | -0.12 | -0.04 | 0.135 | -0.03 | 0.632393571 | 0.711738867 | 0.974639052 | 0.999652205 | 0.995559182 | 0.997605693 | 0.997221123 |
| Slc13a5 | Medium | Q67BT3 | Solute carrier family 13 member 5 OS=Mus musculus OX=10090 GN=Slc13a5 PE=2 SV=1 | 0.01 | 2.615 | 1 | 1 | 1 | 1 | 572 | 63.8 | 7.64 | 1.005 | 0.83 | 1.211 | 0.01 | -0.01 | -0.27 | -0.09 | 0.13 | 0.28 | 0.947413422 | 0.502997631 | 0.286109493 | 0.999652205 | 0.995559182 | 0.997605693 | 0.886074068 |
| | Low | E9QLS6 | Phosphatidylinositol 3-kinase catalytic subunit type 3 OS=Mus musculus OX=10090 GN=Pik3c3 PE | 0.074 | 1.259 | 1 | 1 | 1 | 1 | 848 | 96.8 | 7.2 | 0.976 | 0.85 | 1.149 | -0.03 | 0.03 | -0.23 | -0.0766667 | 0.13 | 0.2 | 0.809585187 | 0.539353293 | 0.343240252 | 0.999652205 | 0.995559182 | 0.997605693 | 0.907770836 |
| Cnp | High | P16330 | 2',3'-cyclic-nucleotide 3'-phosphodiesterase OS=Mus musculus OX=10090 GN=Cnp PE=1 SV=3 | 0 | 15.486 | 15 | 6 | 6 | 6 | 420 | 47.1 | 8.97 | 0.989 | 0.845 | 1.17 | -0.02 | 0.02 | -0.24 | -0.08 | 0.13 | 0.23 | 0.994401284 | 0.54810475 | 0.363700322 | 0.999652205 | 0.995559182 | 0.997605693 | 0.91781256 |
| Dpysl3 | High | E9PWE8 | Dihydropyrimidinase-related protein 3 OS=Mus musculus OX=10090 GN=Dpysl3 PE=1 SV=1 | 0 | 321.894 | 64 | 35 | 121 | 30 | 683 | 73.8 | 6.46 | 0.954 | 0.877 | 1.087 | -0.07 | 0.07 | -0.19 | -0.0633333 | 0.13 | 0.12 | 0.875451255 | 0.625902345 | 0.523569902 | 0.999652205 | 0.995559182 | 0.997605693 | 0.964378027 |
| Ppap2b; Pl | Medium | Q99JY8 | Phospholipid phosphatase 3 OS=Mus musculus OX=10090 GN=Plpp3 PE=1 SV=1 | 0.011 | 2.405 | 4 | 1 | 2 | 1 | 312 | 35.2 | 9.07 | 0.956 | 0.874 | 1.094 | -0.07 | 0.07 | -0.19 | -0.0633333 | 0.13 | 0.13 | 0.779778607 | 0.591412368 | 0.578408273 | 0.999652205 | 0.995559182 | 0.997605693 | 0.972910028 |
| Gga1 | High | Q8R0H9 | ADP-ribosylation factor-binding protein GGA1 OS=Mus musculus OX=10090 GN=Gga1 PE=1 SV=1 | 0 | 8.598 | 5 | 4 | 5 | 3 | 635 | 69.9 | 5.27 | 0.943 | 0.881 | 1.07 | -0.08 | 0.08 | -0.18 | -0.06 | 0.13 | 0.1 | 0.862870254 | 0.636386345 | 0.697352349 | 0.999652205 | 0.995559182 | 0.997605693 | 0.991285312 |
| Abi1 | High | Q8CBW3 | Abl interactor 1 OS=Mus musculus OX=10090 GN=Abi1 PE=1 SV=3 | 0 | 12.56 | 10 | 4 | 5 | 3 | 481 | 52.3 | 7.64 | 0.894 | 0.933 | 0.959 | -0.16 | 0.16 | -0.1 | -0.0333333 | 0.13 | -0.06 | 0.695456744 | 0.764679915 | 0.843475177 | 0.999652205 | 0.995559182 | 0.997605693 | 0.992535481 |
| BC023829; | Medium | Q3TYW3 | Transmembrane protein 185A OS=Mus musculus OX=10090 GN=Tmem185a PE=2 SV=1 | 0.011 | 2.468 | 3 | 1 | 1 | 1 | 322 | 37.2 | 7.2 | 0.921 | 0.906 | 1.017 | -0.12 | 0.12 | -0.14 | -0.0466667 | 0.13 | 0.02 | 0.664034705 | 0.69633702 | 0.861612579 | 0.999652205 | 0.995559182 | 0.997605693 | 0.992535481 |
| Grb2 | High | Q60631 | Growth factor receptor-bound protein 2 OS=Mus musculus OX=10090 GN=Grb2 PE=1 SV=1 | 0.001 | 7.355 | 11 | 2 | 2 | 2 | 217 | 25.2 | 6.32 | 0.903 | 0.93 | 0.972 | -0.15 | 0.15 | -0.11 | -0.0366667 | 0.13 | -0.04 | 0.62958545 | 0.755053498 | 0.904420999 | 0.999652205 | 0.995559182 | 0.997605693 | 0.992535481 |
| As3mt | High | Q91WU5 | Arsenite methyltransferase OS=Mus musculus OX=10090 GN=As3mt PE=1 SV=2 | 0 | 27.55 | 22 | 6 | 9 | 6 | 376 | 41.8 | 5.86 | 0.971 | 0.867 | 1.12 | -0.04 | 0.04 | -0.21 | -0.07 | 0.125 | 0.16 | 0.862811882 | 0.599841353 | 0.518098385 | 0.999652205 | 0.995559182 | 0.997605693 | 0.964378027 |
| Tceb2; Elo | High | P62869 | Elongin-B OS=Mus musculus OX=10090 GN=Elob PE=1 SV=1 | 0 | 51.438 | 77 | 8 | 13 | 8 | 118 | 13.2 | 5.01 | 0.945 | 0.891 | 1.061 | -0.08 | 0.08 | -0.17 | -0.0566667 | 0.125 | 0.09 | 0.821630153 | 0.659129916 | 0.680520596 | 0.999652205 | 0.995559182 | 0.997605693 | 0.991285312 |
| Aak1 | High | Q3UHJ0 | AP2-associated protein kinase 1 OS=Mus musculus OX=10090 GN=Aak1 PE=1 SV=2 | 0 | 151.914 | 41 | 28 | 38 | 22 | 959 | 103.3 | 6.7 | 0.937 | 0.897 | 1.044 | -0.09 | 0.09 | -0.16 | -0.0533333 | 0.125 | 0.06 | 0.828177669 | 0.676146811 | 0.707107504 | 0.999652205 | 0.995559182 | 0.997605693 | 0.992485323 |
| Spag9 | High | Q58A65 | C-Jun-amino-terminal kinase-interacting protein 4 OS=Mus musculus OX=10090 GN=Spag9 PE=1 | 0.001 | 4.373 | 1 | 2 | 2 | 2 | 1321 | 146.1 | 5.15 | 0.894 | 0.936 | 0.954 | -0.16 | 0.16 | -0.09 | -0.03 | 0.125 | -0.07 | 0.563086432 | 0.770619388 | 0.803139496 | 0.999652205 | 0.995559182 | 0.997605693 | 0.992535481 |
| Capza2 | High | P47754 | F-actin-capping protein subunit alpha-2 OS=Mus musculus OX=10090 GN=Capza2 PE=1 SV=3 | 0 | 37.517 | 30 | 6 | 10 | 5 | 286 | 32.9 | 5.85 | 0.937 | 0.894 | 1.047 | -0.09 | 0.09 | -0.16 | -0.0533333 | 0.125 | 0.07 | 0.751398703 | 0.668450038 | 0.810813965 | 0.999652205 | 0.995559182 | 0.997605693 | 0.992535481 |
| Ly6h | High | Q8K356 | Lymphocyte antigen 6 complex, locus H OS=Mus musculus OX=10090 GN=Ly6h PE=1 SV=1 | 0 | 8.918 | 19 | 3 | 3 | 3 | 160 | 17 | 8.25 | 0.92 | 0.911 | 1.01 | -0.12 | 0.12 | -0.13 | -0.0433333 | 0.125 | 0.01 | 0.692893661 | 0.686210668 | 0.885100313 | 0.999652205 | 0.995559182 | 0.997605693 | 0.992535481 |
| Osbpl9 | Low | Q5FWX7 | Oxysterol-binding protein OS=Mus musculus OX=10090 GN=Osbpl9 PE=1 SV=1 | 0.074 | 1.251 | 1 | 1 | 1 | 1 | 626 | 70.1 | 6.05 | 0.936 | 0.906 | 1.033 | -0.1 | 0.1 | -0.14 | -0.0466667 | 0.12 | 0.05 | 0.785490464 | 0.696262941 | 0.813777475 | 0.999652205 | 0.995559182 | 0.997605693 | 0.992535481 |
| Ppp2ca | High | P63330 | Serine/threonine-protein phosphatase 2A catalytic subunit alpha isoform OS=Mus musculus OX=1 | 0 | 64.811 | 43 | 11 | 18 | 1 | 309 | 35.6 | 5.54 | 0.906 | 0.934 | 0.971 | -0.14 | 0.14 | -0.1 | -0.0333333 | 0.12 | -0.04 | 0.656145232 | 0.749940028 | 0.880448711 | 0.999652205 | 0.995559182 | 0.997605693 | 0.992535481 |
| Psmd13 | High | Q9WVJ2 | 26S proteasome non-ATPase regulatory subunit 13 OS=Mus musculus OX=10090 GN=Psmd13 PE= | 0.001 | 7.869 | 6 | 3 | 4 | 3 | 376 | 42.8 | 5.71 | 0.977 | 0.862 | 1.133 | -0.03 | 0.03 | -0.21 | -0.07 | 0.12 | 0.18 | 0.951757937 | 0.58924996 | 0.456817391 | 0.999652205 | 0.995559182 | 0.997605693 | 0.959744396 |
| Cycs; Gm1 | High | P62897 | Cytochrome c, somatic OS=Mus musculus OX=10090 GN=Cycs PE=1 SV=2 | 0 | 52.641 | 64 | 9 | 16 | 9 | 105 | 11.6 | 9.58 | 0.963 | 0.876 | 1.1 | -0.05 | 0.05 | -0.19 | -0.0633333 | 0.12 | 0.14 | 0.901584581 | 0.622188151 | 0.476317195 | 0.999652205 | 0.995559182 | 0.997605693 | 0.962418203 |
| Plscr3 | High | Q9JIZ9 | Phospholipid scramblase 3 OS=Mus musculus OX=10090 GN=Plscr3 PE=1 SV=1 | 0.001 | 5.866 | 11 | 2 | 2 | 2 | 296 | 31.8 | 6.38 | 0.903 | 0.941 | 0.96 | -0.15 | 0.15 | -0.09 | -0.03 | 0.12 | -0.06 | 0.671595244 | 0.759665726 | 0.872093023 | 0.999652205 | 0.995559182 | 0.997605693 | 0.992535481 |
| | Medium | S4R2D5 | Ubiquitin-like modifier-activating enzyme ATG7 OS=Mus musculus OX=10090 GN=Atg7 PE=1 SV= | 0.029 | 1.921 | 1 | 1 | 1 | 1 | 618 | 68.2 | 6.54 | 0.928 | 0.914 | 1.015 | -0.11 | 0.11 | -0.13 | -0.0433333 | 0.12 | 0.02 | 0.728485432 | 0.714807108 | 0.938194539 | 0.999652205 | 0.995559182 | 0.997605693 | 0.992535481 |
| Fam105b; | Medium | Q3UCV8 | Ubiquitin thioesterase otulin OS=Mus musculus OX=10090 GN=Otulin PE=1 SV=1 | 0.04 | 1.688 | 2 | 1 | 1 | 1 | 352 | 40.3 | 5.68 | 1.022 | 0.834 | 1.225 | 0.03 | -0.03 | -0.26 | -0.0866667 | 0.115 | 0.29 | 0.99130195 | 0.499291094 | 0.252786394 | 0.999652205 | 0.995559182 | 0.997605693 | 0.851275262 |
| Tpi1 | High | P17751 | Triosephosphate isomerase OS=Mus musculus OX=10090 GN=Tpi1 PE=1 SV=5 | 0 | 252.649 | 84 | 17 | 119 | 17 | 249 | 26.7 | 7.3 | 0.951 | 0.897 | 1.06 | -0.07 | 0.07 | -0.16 | -0.0533333 | 0.115 | 0.08 | 0.869380359 | 0.675441297 | 0.632289049 | 0.999652205 | 0.995559182 | 0.997605693 | 0.982523274 |
| | High | F7BQW7 | Phosphoinositide 5-phosphatase (Fragment) OS=Mus musculus OX=10090 GN=Synj1 PE=1 SV=1 | 0 | 216.986 | 38 | 44 | 73 | 44 | 1324 | 146.4 | 7.9 | 0.937 | 0.905 | 1.036 | -0.09 | 0.09 | -0.14 | -0.0466667 | 0.115 | 0.05 | 0.829300964 | 0.694266699 | 0.743895617 | 0.999652205 | 0.995559182 | 0.997605693 | 0.992535481 |
| Trappc3 | High | O55013 | Trafficking protein particle complex subunit 3 OS=Mus musculus OX=10090 GN=Trappc3 PE=1 SV | 0.005 | 3.561 | 8 | 2 | 3 | 2 | 180 | 20.3 | 4.96 | 0.934 | 0.914 | 1.022 | -0.1 | 0.1 | -0.13 | -0.0433333 | 0.115 | 0.03 | 0.778376247 | 0.717783815 | 0.861107229 | 0.999652205 | 0.995559182 | 0.997605693 | 0.992535481 |
| Anxa4 | High | P97429 | Annexin A4 OS=Mus musculus OX=10090 GN=Anxa4 PE=1 SV=4 | 0 | 33.75 | 39 | 12 | 12 | 12 | 319 | 35.9 | 5.57 | 0.916 | 0.932 | 0.983 | -0.13 | 0.13 | -0.1 | -0.0333333 | 0.115 | -0.02 | 0.707289534 | 0.762307208 | 0.891371314 | 0.999652205 | 0.995559182 | 0.997605693 | 0.992535481 |
| Ppp2r5b | High | Q6PD28 | Serine/threonine-protein phosphatase 2A 56 kDa regulatory subunit beta isoform OS=Mus muscu | 0 | 17.472 | 13 | 6 | 6 | 5 | 497 | 57.3 | 6.84 | 0.928 | 0.918 | 1.011 | -0.11 | 0.11 | -0.12 | -0.04 | 0.115 | 0.02 | 0.765548117 | 0.727596872 | 0.940214412 | 0.999652205 | 0.995559182 | 0.997605693 | 0.992535481 |
| Aldoc | High | P05063 | Fructose-bisphosphate aldolase C OS=Mus musculus OX=10090 GN=Aldoc PE=1 SV=4 | 0 | 338.374 | 71 | 29 | 187 | 22 | 363 | 39.4 | 7.12 | 1.009 | 0.852 | 1.185 | 0.01 | -0.01 | -0.23 | -0.0766667 | 0.11 | 0.24 | 0.973417864 | 0.563043812 | 0.232978184 | 0.999652205 | 0.995559182 | 0.997605693 | 0.821061393 |
| 2610301B | High | B1AV75 | RIKEN cDNA 2610301B20 gene OS=Mus musculus OX=10090 GN=2610301B20Rik PE=1 SV=1 | 0 | 13.968 | 24 | 5 | 5 | 5 | 210 | 23.9 | 6.86 | 0.999 | 0.858 | 1.165 | 0 | 0 | -0.22 | -0.0733333 | 0.11 | 0.22 | 0.912441276 | 0.558678133 | 0.302529953 | 0.999652205 | 0.995559182 | 0.997605693 | 0.900241969 |
| Nrbp1 | High | D3YUV1 | Nuclear receptor-binding protein OS=Mus musculus OX=10090 GN=Nrbp1 PE=1 SV=1 | 0.001 | 4.467 | 3 | 1 | 1 | 1 | 543 | 60.8 | 5.16 | 1.007 | 0.855 | 1.177 | 0.01 | -0.01 | -0.23 | -0.0766667 | 0.11 | 0.24 | 0.94313139 | 0.550743619 | 0.355026893 | 0.999652205 | 0.995559182 | 0.997605693 | 0.913554746 |
| Cask | High | O70589 | Peripheral plasma membrane protein CASK OS=Mus musculus OX=10090 GN=Cask PE=1 SV=2 | 0 | 18.664 | 7 | 6 | 7 | 6 | 926 | 105 | 6.43 | 0.957 | 0.897 | 1.067 | -0.06 | 0.06 | -0.16 | -0.0533333 | 0.11 | 0.09 | 0.817686549 | 0.675192699 | 0.727648894 | 0.999652205 | 0.995559182 | 0.997605693 | 0.992535481 |
| Abca4 | High | O35600 | Retinal-specific phospholipid-transporting ATPase ABCA4 OS=Mus musculus OX=10090 GN=Abca | 0 | 1723.77 | 60 | 125 | 1001 | 124 | 2310 | 260 | 6.2 | 0.938 | 0.915 | 1.026 | -0.09 | 0.09 | -0.13 | -0.0433333 | 0.11 | 0.04 | 0.833292824 | 0.718945141 | 0.789671179 | 0.999652205 | 0.995559182 | 0.997605693 | 0.992535481 |
| Frmd5 | High | A2AR56 | FERM domain-containing protein 5 OS=Mus musculus OX=10090 GN=Frmd5 PE=1 SV=1 | 0 | 10.318 | 5 | 3 | 3 | 3 | 570 | 64.9 | 8.44 | 0.923 | 0.935 | 0.987 | -0.12 | 0.12 | -0.1 | -0.0333333 | 0.11 | -0.02 | 0.804674798 | 0.770043808 | 0.963882756 | 0.999652205 | 0.995559182 | 0.997605693 | 0.994203571 |
| Exoc7 | High | O35250 | Exocyst complex component 7 OS=Mus musculus OX=10090 GN=Exoc7 PE=1 SV=2 | 0.009 | 2.842 | 2 | 1 | 1 | 1 | 697 | 79.9 | 6.98 | 0.976 | 0.889 | 1.097 | -0.04 | 0.04 | -0.17 | -0.0566667 | 0.105 | 0.13 | 0.86924444 | 0.638032219 | 0.59148882 | 0.999652205 | 0.995559182 | 0.997605693 | 0.975147842 |
| Iqsec3 | High | Q3TES0 | IQ motif and SEC7 domain-containing protein 3 OS=Mus musculus OX=10090 GN=Iqsec3 PE=1 SV | 0 | 37.016 | 11 | 10 | 12 | 7 | 1195 | 129 | 6.19 | 0.927 | 0.93 | 0.997 | -0.11 | 0.11 | -0.1 | -0.0333333 | 0.105 | 0 | 0.736018363 | 0.758859659 | 0.94944042 | 0.999652205 | 0.995559182 | 0.997605693 | 0.992535481 |
| Ap2s1 | High | P62743 | AP-2 complex subunit sigma OS=Mus musculus OX=10090 GN=Ap2s1 PE=1 SV=1 | 0 | 20.58 | 27 | 5 | 8 | 5 | 142 | 17 | 6.18 | 0.929 | 0.931 | 0.998 | -0.11 | 0.11 | -0.1 | -0.0333333 | 0.105 | 0 | 0.725476337 | 0.759819112 | 0.957331484 | 0.999652205 | 0.995559182 | 0.997605693 | 0.992902921 |
| Ppp1r9b | High | Q6R891 | Neurabin-2 OS=Mus musculus OX=10090 GN=Ppp1r9b PE=1 SV=1 | 0 | 25.583 | 9 | 6 | 7 | 6 | 817 | 89.5 | 4.92 | 0.93 | 0.93 | 1 | -0.1 | 0.1 | -0.11 | -0.0366667 | 0.105 | 0 | 0.727081871 | 0.756668139 | 0.972063592 | 0.999652205 | 0.995559182 | 0.997605693 | 0.996575243 |
| Slc8a3 | Medium | E9PX50 | Sodium/calcium exchanger 3 OS=Mus musculus OX=10090 GN=Slc8a3 PE=1 SV=2 | 0.014 | 2.235 | 2 | 1 | 1 | 1 | 927 | 102.9 | 5.12 | 1.022 | 0.844 | 1.21 | 0.03 | -0.03 | -0.24 | -0.08 | 0.105 | 0.28 | 0.987293823 | 0.520592013 | 0.243860813 | 0.999652205 | 0.995559182 | 0.997605693 | 0.837310213 |
| | High | Q3TLI0 | Trafficking protein particle complex subunit 10 OS=Mus musculus OX=10090 GN=Trappc10 PE=1 | 0.001 | 7.081 | 1 | 1 | 2 | 1 | 1258 | 141.4 | 5.92 | 1.004 | 0.861 | 1.167 | 0.01 | -0.01 | -0.22 | -0.0733333 | 0.105 | 0.22 | 0.943256816 | 0.552431565 | 0.317781835 | 0.999652205 | 0.995559182 | 0.997605693 | 0.904943958 |
| Dhrs7b | Medium | Q99J47 | Dehydrogenase/reductase SDR family member 7B OS=Mus musculus OX=10090 GN=Dhrs7b PE=1 | 0.015 | 2.185 | 2 | 1 | 1 | 1 | 323 | 35 | 9.63 | 0.989 | 0.874 | 1.132 | -0.02 | 0.02 | -0.19 | -0.0633333 | 0.105 | 0.18 | 0.916256838 | 0.606083042 | 0.494402907 | 0.999652205 | 0.995559182 | 0.997605693 | 0.964378027 |
| Fasn | High | P19096 | Fatty acid synthase OS=Mus musculus OX=10090 GN=Fasn PE=1 SV=2 | 0 | 105.354 | 13 | 27 | 32 | 27 | 2504 | 272.3 | 6.58 | 0.971 | 0.894 | 1.087 | -0.04 | 0.04 | -0.16 | -0.0533333 | 0.1 | 0.12 | 0.924712397 | 0.667050775 | 0.524347417 | 0.999652205 | 0.995559182 | 0.997605693 | 0.964378027 |
| Ass1 | High | P16460 | Argininosuccinate synthase OS=Mus musculus OX=10090 GN=Ass1 PE=1 SV=1 | 0 | 18.641 | 11 | 5 | 8 | 5 | 412 | 46.6 | 8.22 | 0.949 | 0.913 | 1.04 | -0.07 | 0.07 | -0.13 | -0.0433333 | 0.1 | 0.06 | 0.843717674 | 0.714332949 | 0.737445547 | 0.999652205 | 0.995559182 | 0.997605693 | 0.992535481 |
| Adh5 | High | P28474 | Alcohol dehydrogenase class-3 OS=Mus musculus OX=10090 GN=Adh5 PE=1 SV=3 | 0 | 54.388 | 33 | 11 | 20 | 11 | 374 | 39.5 | 7.25 | 0.939 | 0.925 | 1.016 | -0.09 | 0.09 | -0.11 | -0.0366667 | 0.1 | 0.02 | 0.835451668 | 0.744911423 | 0.841857647 | 0.999652205 | 0.995559182 | 0.997605693 | 0.992535481 |
| F11r | High | O88792 | Junctional adhesion molecule A OS=Mus musculus OX=10090 GN=F11r PE=1 SV=2 | 0.005 | 3.728 | 5 | 2 | 2 | 2 | 300 | 32.4 | 6.77 | 0.944 | 0.92 | 1.025 | -0.08 | 0.08 | -0.12 | -0.04 | 0.1 | 0.04 | 0.751335422 | 0.717275485 | 0.867601549 | 0.999652205 | 0.995559182 | 0.997605693 | 0.992535481 |
| Cp | High | G3X8Q5 | Ceruloplasmin OS=Mus musculus OX=10090 GN=Cp PE=1 SV=1 | 0 | 161.552 | 34 | 32 | 60 | 1 | 1085 | 124.1 | 5.88 | 0.934 | 0.931 | 1.003 | -0.1 | 0.1 | -0.1 | -0.0333333 | 0.1 | 0 | 0.821147094 | 0.760487343 | 0.903489032 | 0.999652205 | 0.995559182 | 0.997605693 | 0.992535481 |
| Klc1 | High | Q5UE59 | Kinesin light chain OS=Mus musculus OX=10090 GN=Klc1 PE=1 SV=1 | 0 | 30.516 | 19 | 9 | 9 | 5 | 542 | 61.6 | 5.76 | 0.936 | 0.936 | 1 | -0.1 | 0.1 | -0.1 | -0.0333333 | 0.1 | 0 | 0.786705369 | 0.77241241 | 0.990928774 | 0.999652205 | 0.995559182 | 0.997605693 | 0.999783808 |
| Ehbp1 | High | Q69ZW3 | EH domain-binding protein 1 OS=Mus musculus OX=10090 GN=Ehbp1 PE=1 SV=3 | 0.006 | 3.248 | 1 | 1 | 1 | 1 | 1231 | 139 | 5.38 | 0.926 | 0.94 | 0.985 | -0.11 | 0.11 | -0.09 | -0.03 | 0.1 | -0.02 | 0.714797652 | 0.763071738 | 0.993327656 | 0.999652205 | 0.995559182 | 0.997605693 | 0.999917596 |
| Nme3 | Medium | Q9WV85 | Nucleoside diphosphate kinase 3 OS=Mus musculus OX=10090 GN=Nme3 PE=1 SV=3 | 0.011 | 2.487 | 6 | 1 | 1 | 1 | 169 | 19.1 | 6.68 | 0.99 | 0.88 | 1.125 | -0.02 | 0.02 | -0.18 | -0.06 | 0.1 | 0.17 | 0.924232647 | 0.60757568 | 0.501884763 | 0.999652205 | 0.995559182 | 0.997605693 | 0.964378027 |
| Wdr45b; V | Medium | Q9CR39 | WD repeat domain phosphoinositide-interacting protein 3 OS=Mus musculus OX=10090 GN=Wdr | 0.019 | 2.049 | 3 | 1 | 1 | 1 | 344 | 38 | 7.56 | 1.033 | 0.844 | 1.224 | 0.05 | -0.05 | -0.24 | -0.08 | 0.095 | 0.29 | 0.979042037 | 0.528449912 | 0.185523713 | 0.999652205 | 0.995559182 | 0.997605693 | 0.771113016 |
| Taok3 | High | Q8BYC6 | Serine/threonine-protein kinase TAO3 OS=Mus musculus OX=10090 GN=Taok3 PE=1 SV=2 | 0 | 26.083 | 9 | 7 | 8 | 6 | 898 | 105.3 | 7.36 | 1.045 | 0.842 | 1.242 | 0.06 | -0.06 | -0.25 | -0.0833333 | 0.095 | 0.31 | 0.933484954 | 0.538561515 | 0.19904004 | 0.999652205 | 0.995559182 | 0.997605693 | 0.78697088 |
| Maged2 | Low | Q9ER67 | MAGE family member D2 OS=Mus musculus OX=10090 GN=Maged2 PE=1 SV=1 | 0.053 | 1.422 | 1 | 1 | 1 | 1 | 616 | 65.4 | 9.14 | 0.973 | 0.901 | 1.08 | -0.04 | 0.04 | -0.15 | -0.05 | 0.095 | 0.11 | 0.885044474 | 0.683704363 | 0.635656664 | 0.999652205 | 0.995559182 | 0.997605693 | 0.983562471 |
| Ogdhl | High | B2RXT3 | Oxoglutarate dehydrogenase (succinyl-transferring) OS=Mus musculus OX=10090 GN=Ogdhl PE=1 | 0 | 34.261 | 10 | 10 | 12 | 4 | 1010 | 114.5 | 6.84 | 0.961 | 0.915 | 1.05 | -0.06 | 0.06 | -0.13 | -0.0433333 | 0.095 | 0.07 | 0.922238294 | 0.721025759 | 0.751633878 | 0.999652205 | 0.995559182 | 0.997605693 | 0.992535481 |
| Cpox | High | P36552 | Oxygen-dependent coproporphyrinogen-III oxidase, mitochondrial OS=Mus musculus OX=10090 G | 0.006 | 3.391 | 3 | 1 | 1 | 1 | 443 | 49.7 | 8.53 | 1.014 | 0.868 | 1.169 | 0.02 | -0.02 | -0.2 | -0.0666667 | 0.09 | 0.22 | 0.972181089 | 0.583071356 | 0.393432149 | 0.999652205 | 0.995559182 | 0.997605693 | 0.931576883 |
| Ahcy; Gm4 | High | P50247 | Adenosylhomocysteinase OS=Mus musculus OX=10090 GN=Ahcy PE=1 SV=3 | 0 | 42.614 | 22 | 10 | 17 | 10 | 432 | 47.7 | 6.54 | 0.994 | 0.89 | 1.117 | -0.01 | 0.01 | -0.17 | -0.0566667 | 0.09 | 0.16 | 0.996406846 | 0.657896255 | 0.41568987 | 0.999652205 | 0.995559182 | 0.997605693 | 0.936397673 |
| Ppp2r5e | High | Q61151 | Serine/threonine-protein phosphatase 2A 56 kDa regulatory subunit epsilon isoform OS=Mus mus | 0 | 16.281 | 11 | 5 | 5 | 3 | 467 | 54.7 | 6.95 | 0.996 | 0.886 | 1.124 | -0.01 | 0.01 | -0.17 | -0.0566667 | 0.09 | 0.17 | 0.912130703 | 0.637875803 | 0.434022314 | 0.999652205 | 0.995559182 | 0.997605693 | 0.948052758 |
| Pdcd10 | High | Q8VE70 | Programmed cell death protein 10 OS=Mus musculus OX=10090 GN=Pdcd10 PE=1 SV=1 | 0 | 12.124 | 15 | 3 | 3 | 3 | 212 | 24.7 | 8.19 | 0.971 | 0.905 | 1.074 | -0.04 | 0.04 | -0.14 | -0.0466667 | 0.09 | 0.1 | 0.81714732 | 0.676327046 | 0.612974474 | 0.999652205 | 0.995559182 | 0.997605693 | 0.97826833 |
| Acly | High | Q3V117 | ATP-citrate synthase OS=Mus musculus OX=10090 GN=Acly PE=1 SV=1 | 0 | 237.12 | 46 | 43 | 73 | 2 | 1101 | 120.7 | 7.44 | 1.085 | 0.813 | 1.334 | 0.12 | -0.12 | -0.3 | -0.1 | 0.09 | 0.42 | 0.781184894 | 0.471223326 | 0.050036907 | 0.999652205 | 0.995559182 | 0.997605693 | 0.427566646 |
| Ccs | High | Q9WU84 | Copper chaperone for superoxide dismutase OS=Mus musculus OX=10090 GN=Ccs PE=1 SV=1 | 0 | 4.75 | 6 | 2 | 2 | 2 | 274 | 28.9 | 6.1 | 1.002 | 0.881 | 1.137 | 0 | 0 | -0.18 | -0.06 | 0.09 | 0.19 | 0.98322364 | 0.635522316 | 0.46394774 | 0.999652205 | 0.995559182 | 0.997605693 | 0.959744396 |
| Rab8b | High | P61028 | Ras-related protein Rab-8B OS=Mus musculus OX=10090 GN=Rab8b PE=1 SV=1 | 0 | 37.765 | 43 | 9 | 19 | 3 | 207 | 23.6 | 9.07 | 0.98 | 0.903 | 1.085 | -0.03 | 0.03 | -0.15 | -0.05 | 0.09 | 0.12 | 0.964638507 | 0.690453518 | 0.624438668 | 0.999652205 | 0.995559182 | 0.997605693 | 0.978918782 |
| Capn2 | High | O08529 | Calpain-2 catalytic subunit OS=Mus musculus OX=10090 GN=Capn2 PE=1 SV=4 | 0.001 | 6.922 | 3 | 2 | 2 | 2 | 700 | 79.8 | 4.96 | 0.945 | 0.933 | 1.012 | -0.08 | 0.08 | -0.1 | -0.0333333 | 0.09 | 0.02 | 0.817409949 | 0.765250368 | 0.920635657 | 0.999652205 | 0.995559182 | 0.997605693 | 0.992535481 |
| Map2k4 | High | B1ATL6 | Dual-specificity mitogen-activated protein kinase kinase 4 (Fragment) OS=Mus musculus OX=100 | 0 | 19.406 | 23 | 4 | 5 | 4 | 206 | 22.5 | 8.65 | 1.042 | 0.853 | 1.222 | 0.06 | -0.06 | -0.23 | -0.0766667 | 0.085 | 0.29 | 0.87025623 | 0.565114237 | 0.244919779 | 0.999652205 | 0.995559182 | 0.997605693 | 0.838891386 |
| Smyd5 | High | Q3TYX3 | SET and MYND domain-containing protein 5 OS=Mus musculus OX=10090 GN=Smyd5 PE=1 SV=2 | 0 | 4.726 | 3 | 1 | 1 | 1 | 416 | 47.1 | 5.21 | 1.02 | 0.868 | 1.176 | 0.03 | -0.03 | -0.2 | -0.0666667 | 0.085 | 0.23 | 0.992657857 | 0.599070064 | 0.336018233 | 0.999652205 | 0.995559182 | 0.997605693 | 0.906329766 |
| | High | A0A6I8MX | L-lactate dehydrogenase OS=Mus musculus OX=10090 GN=Ldhb PE=1 SV=1 | 0 | 178.161 | 60 | 18 | 74 | 17 | 341 | 37.3 | 6.25 | 0.98 | 0.907 | 1.081 | -0.03 | 0.03 | -0.14 | -0.0466667 | 0.085 | 0.11 | 0.949510828 | 0.700316738 | 0.547719744 | 0.999652205 | 0.995559182 | 0.997605693 | 0.96694628 |
| Exog | High | Q8C163 | Nuclease EXOG, mitochondrial OS=Mus musculus OX=10090 GN=Exog PE=1 SV=1 | 0.001 | 6.64 | 6 | 2 | 2 | 2 | 368 | 41.4 | 8.12 | 0.99 | 0.897 | 1.104 | -0.01 | 0.01 | -0.16 | -0.0533333 | 0.085 | 0.14 | 0.908376261 | 0.662361673 | 0.586720206 | 0.999652205 | 0.995559182 | 0.997605693 | 0.973095093 |
| Pde4b | High | B1AWC8 | Phosphodiesterase OS=Mus musculus OX=10090 GN=Pde4b PE=1 SV=1 | 0.001 | 6.575 | 4 | 2 | 2 | 2 | 659 | 75 | 5.22 | 0.972 | 0.913 | 1.065 | -0.04 | 0.04 | -0.13 | -0.0433333 | 0.085 | 0.09 | 0.772567638 | 0.688116692 | 0.628581803 | 0.999652205 | 0.995559182 | 0.997605693 | 0.981444134 |
| Ugp2 | High | Q91ZJ5 | UTP--glucose-1-phosphate uridylyltransferase OS=Mus musculus OX=10090 GN=Ugp2 PE=1 SV=3 | 0 | 10.33 | 6 | 3 | 3 | 3 | 508 | 56.9 | 7.61 | 0.953 | 0.934 | 1.021 | -0.07 | 0.07 | -0.1 | -0.0333333 | 0.085 | 0.03 | 0.894876498 | 0.767472878 | 0.884956852 | 0.999652205 | 0.995559182 | 0.997605693 | 0.992535481 |
| Eif3e | High | P60229 | Eukaryotic translation initiation factor 3 subunit E OS=Mus musculus OX=10090 GN=Eif3e PE=1 S | 0 | 14.718 | 11 | 5 | 5 | 5 | 445 | 52.2 | 6.04 | 1.033 | 0.86 | 1.202 | 0.05 | -0.05 | -0.22 | -0.0733333 | 0.085 | 0.26 | 0.978069128 | 0.552836046 | 0.229510506 | 0.999652205 | 0.995559182 | 0.997605693 | 0.819186996 |
| | High | A0A0U1RF | FXYD domain-containing ion transport regulator OS=Mus musculus OX=10090 GN=Fxyd7 PE=1 SV | 0.001 | 6.886 | 29 | 2 | 2 | 2 | 77 | 8.1 | 7.84 | 1.004 | 0.885 | 1.135 | 0.01 | -0.01 | -0.18 | -0.06 | 0.085 | 0.18 | 0.989243248 | 0.639494658 | 0.457588122 | 0.999652205 | 0.995559182 | 0.997605693 | 0.959744396 |
| Pclo | High | Q9QYX7 | Protein piccolo OS=Mus musculus OX=10090 GN=Pclo PE=1 SV=4 | 0 | 60.133 | 4 | 17 | 22 | 17 | 5068 | 550.5 | 6.51 | 0.987 | 0.903 | 1.093 | -0.02 | 0.02 | -0.15 | -0.05 | 0.085 | 0.13 | 0.998331865 | 0.689071344 | 0.499870749 | 0.999652205 | 0.995559182 | 0.997605693 | 0.964378027 |
| Tpbg | Medium | Q9Z0L0 | Trophoblast glycoprotein OS=Mus musculus OX=10090 GN=Tpbg PE=1 SV=3 | 0.011 | 2.426 | 2 | 1 | 1 | 1 | 426 | 46.4 | 6.83 | 0.989 | 0.904 | 1.094 | -0.02 | 0.02 | -0.15 | -0.05 | 0.085 | 0.13 | 0.801225191 | 0.670221782 | 0.520652207 | 0.999652205 | 0.995559182 | 0.997605693 | 0.964378027 |
| Asap2 | High | Q7SIG6 | Arf-GAP with SH3 domain, ANK repeat and PH domain-containing protein 2 OS=Mus musculus OX | 0.001 | 7.85 | 3 | 3 | 4 | 3 | 958 | 106.7 | 6.65 | 1.026 | 0.872 | 1.176 | 0.04 | -0.04 | -0.2 | -0.0666667 | 0.08 | 0.23 | 0.878924181 | 0.613496157 | 0.309389635 | 0.999652205 | 0.995559182 | 0.997605693 | 0.904943958 |
| Arpc3 | High | Q9JM76 | Actin-related protein 2/3 complex subunit 3 OS=Mus musculus OX=10090 GN=Arpc3 PE=1 SV=3 | 0 | 24.238 | 25 | 5 | 10 | 5 | 178 | 20.5 | 8.59 | 1.012 | 0.883 | 1.146 | 0.02 | -0.02 | -0.18 | -0.06 | 0.08 | 0.2 | 0.906477384 | 0.639415494 | 0.329618491 | 0.999652205 | 0.995559182 | 0.997605693 | 0.906329766 |
| Dnm1l | High | E9PUD2 | Dynamin-1-like protein OS=Mus musculus OX=10090 GN=Dnm1l PE=1 SV=1 | 0 | 67.245 | 28 | 14 | 18 | 14 | 712 | 79.5 | 7.08 | 0.997 | 0.896 | 1.113 | 0 | 0 | -0.16 | -0.0533333 | 0.08 | 0.15 | 0.944269304 | 0.672423929 | 0.417643491 | 0.999652205 | 0.995559182 | 0.997605693 | 0.936397673 |
| Guca1a | High | P43081 | Guanylyl cyclase-activating protein 1 OS=Mus musculus OX=10090 GN=Guca1a PE=1 SV=3 | 0 | 54.5 | 56 | 8 | 19 | 8 | 202 | 23 | 4.55 | 1.002 | 0.895 | 1.119 | 0 | 0 | -0.16 | -0.0533333 | 0.08 | 0.16 | 0.991681161 | 0.670892054 | 0.428884517 | 0.999652205 | 0.995559182 | 0.997605693 | 0.943781588 |
| Trap1 | High | Q9CQN1 | Heat shock protein 75 kDa, mitochondrial OS=Mus musculus OX=10090 GN=Trap1 PE=1 SV=1 | 0 | 9.603 | 4 | 3 | 9 | 2 | 706 | 80.2 | 6.68 | 0.984 | 0.906 | 1.086 | -0.02 | 0.02 | -0.14 | -0.0466667 | 0.08 | 0.12 | 0.895811083 | 0.674439739 | 0.572565376 | 0.999652205 | 0.995559182 | 0.997605693 | 0.972910028 |
| Eml3 | High | Q8VC03 | Echinoderm microtubule-associated protein-like 3 OS=Mus musculus OX=10090 GN=Eml3 PE=1 S | 0.001 | 6.529 | 2 | 2 | 3 | 2 | 897 | 95.6 | 7.05 | 1.032 | 0.87 | 1.186 | 0.05 | -0.05 | -0.2 | -0.0666667 | 0.075 | 0.25 | 0.956823275 | 0.586541146 | 0.27927727 | 0.999652205 | 0.995559182 | 0.997605693 | 0.876067403 |
| | High | F6Q8N5 | UPF3 regulator of nonsense transcripts homolog B (yeast) (Fragment) OS=Mus musculus OX=100 | 0.006 | 3.282 | 3 | 1 | 1 | 1 | 260 | 32.4 | 9.74 | 1.066 | 0.848 | 1.258 | 0.09 | -0.09 | -0.24 | -0.08 | 0.075 | 0.33 | 0.886734941 | 0.528704384 | 0.142726734 | 0.999652205 | 0.995559182 | 0.997605693 | 0.703085292 |

| | | | | | | | | | | | | | | | | | | | | | | | | | | | | |
|---|---|---|---|---|---|---|---|---|---|---|---|---|---|---|---|---|---|---|---|---|---|---|---|---|---|---|---|---|
| Arpc4 | High | P59999 | Actin-related protein 2/3 complex subunit 4 OS=Mus musculus OX=10090 GN=Arpc4 PE=1 SV=3 | 0 | 23.537 | 39 | 6 | 7 | 6 | 168 | 19.7 | 8.43 | 1.018 | 0.885 | 1.15 | 0.03 | -0.03 | -0.18 | -0.06 | 0.075 | 0.2 | 0.984637032 | 0.645309897 | 0.411809525 | 0.999652205 | 0.995559182 | 0.997605693 | 0.936397673 |
| | High | E9PWR4 | Disco-interacting protein 2 homolog C OS=Mus musculus OX=10090 GN=Dip2c PE=1 SV=1 | 0 | 4.754 | 1 | 2 | 2 | 2 | 1557 | 171 | 7.62 | 0.998 | 0.901 | 1.108 | 0 | 0 | -0.15 | -0.05 | 0.075 | 0.15 | 0.925580093 | 0.662285345 | 0.495704303 | 0.999652205 | 0.995559182 | 0.997605693 | 0.964378027 |
| Fam126b | High | Q8C729 | Protein FAM126B OS=Mus musculus OX=10090 GN=Fam126b PE=1 SV=1 | 0 | 4.834 | 4 | 2 | 2 | 2 | 530 | 58.6 | 7.72 | 0.974 | 0.927 | 1.05 | -0.04 | 0.04 | -0.11 | -0.0366667 | 0.075 | 0.07 | 0.957630474 | 0.751282167 | 0.774192209 | 0.999652205 | 0.995559182 | 0.997605693 | 0.992535481 |
| Cul2 | High | Q9D4H8 | Cullin-2 OS=Mus musculus OX=10090 GN=Cul2 PE=1 SV=2 | 0 | 18.846 | 11 | 8 | 9 | 8 | 745 | 86.8 | 7.01 | 0.972 | 0.928 | 1.047 | -0.04 | 0.04 | -0.11 | -0.0366667 | 0.075 | 0.07 | 0.870160488 | 0.75216032 | 0.800397535 | 0.999652205 | 0.995559182 | 0.997605693 | 0.992535481 |
| Ldha | High | A0A1B0GS | L-lactate dehydrogenase OS=Mus musculus OX=10090 GN=Ldha PE=1 SV=1 | 0 | 382.515 | 68 | 29 | 234 | 28 | 361 | 39.7 | 8.35 | 1.101 | 0.826 | 1.332 | 0.14 | -0.14 | -0.28 | -0.0933333 | 0.07 | 0.41 | 0.744789961 | 0.502045218 | 0.051452976 | 0.999652205 | 0.995559182 | 0.997605693 | 0.429310767 |
| | High | A0A3Q4EC | COMM domain-containing protein 10 OS=Mus musculus OX=10090 GN=Commd10 PE=1 SV=1 | 0.006 | 3.341 | 5 | 1 | 2 | 1 | 165 | 18.6 | 9.5 | 1.102 | 0.826 | 1.334 | 0.14 | -0.14 | -0.28 | -0.0933333 | 0.07 | 0.42 | 0.808599824 | 0.476340071 | 0.067891404 | 0.999652205 | 0.995559182 | 0.997605693 | 0.499896415 |
| Grcc10 | High | O35127 | Protein C10 OS=Mus musculus OX=10090 GN=Grcc10 PE=1 SV=1 | 0 | 13.788 | 40 | 3 | 3 | 3 | 126 | 13.2 | 5.14 | 1.083 | 0.84 | 1.289 | 0.11 | -0.11 | -0.25 | -0.0833333 | 0.07 | 0.37 | 0.789230078 | 0.534076614 | 0.110220096 | 0.999652205 | 0.995559182 | 0.997605693 | 0.632640059 |
| Otud7b | High | B2RUR8 | OTU domain-containing protein 7B OS=Mus musculus OX=10090 GN=Otud7b PE=1 SV=1 | 0 | 35.386 | 11 | 7 | 9 | 7 | 840 | 91.9 | 6.86 | 0.998 | 0.909 | 1.097 | 0 | 0 | -0.14 | -0.0466667 | 0.07 | 0.13 | 0.952178412 | 0.706199209 | 0.592867921 | 0.999652205 | 0.995559182 | 0.997605693 | 0.976360704 |
| Pet112; Pe | High | Q99JT1 | Glutamyl-tRNA(Gln) amidotransferase subunit B, mitochondrial OS=Mus musculus OX=10090 GN | 0.006 | 3.454 | 1 | 1 | 1 | 1 | 557 | 62.1 | 8.54 | 0.982 | 0.925 | 1.061 | -0.03 | 0.03 | -0.11 | -0.0366667 | 0.07 | 0.09 | 0.874327913 | 0.734371399 | 0.745617633 | 0.999652205 | 0.995559182 | 0.997605693 | 0.992535481 |
| Arih2 | High | Q9Z1K6 | E3 ubiquitin-protein ligase ARIH2 OS=Mus musculus OX=10090 GN=Arih2 PE=1 SV=1 | 0.001 | 7.166 | 3 | 1 | 1 | 1 | 492 | 57.7 | 5.69 | 1.118 | 0.811 | 1.378 | 0.16 | -0.16 | -0.3 | -0.1 | 0.07 | 0.46 | 0.682350818 | 0.459977855 | 0.066749903 | 0.999652205 | 0.995559182 | 0.997605693 | 0.499896415 |
| Gmds | High | Q8K0C9 | GDP-mannose 4,6 dehydratase OS=Mus musculus OX=10090 GN=Gmds PE=1 SV=1 | 0.001 | 7.176 | 7 | 2 | 2 | 2 | 372 | 42 | 7.03 | 1.028 | 0.883 | 1.164 | 0.04 | -0.04 | -0.18 | -0.06 | 0.07 | 0.22 | 0.986320736 | 0.619737682 | 0.365237242 | 0.999652205 | 0.995559182 | 0.997605693 | 0.920124252 |
| | High | D3YU75 | Translationally-controlled tumor protein OS=Mus musculus OX=10090 GN=Tpt1 PE=1 SV=1 | 0 | 10.324 | 20 | 3 | 3 | 3 | 138 | 15.5 | 4.96 | 1.027 | 0.884 | 1.162 | 0.04 | -0.04 | -0.18 | -0.06 | 0.07 | 0.22 | 0.971244806 | 0.635947278 | 0.374663111 | 0.999652205 | 0.995559182 | 0.997605693 | 0.920164724 |
| Dstn | High | Q9R0P5 | Destrin OS=Mus musculus OX=10090 GN=Dstn PE=1 SV=3 | 0 | 73.667 | 69 | 15 | 29 | 14 | 165 | 18.5 | 7.97 | 1.055 | 0.866 | 1.218 | 0.08 | -0.08 | -0.21 | -0.07 | 0.065 | 0.28 | 0.855715981 | 0.597211917 | 0.169971344 | 0.999652205 | 0.995559182 | 0.997605693 | 0.738844686 |
| Arpc2 | High | Q9CVB6 | Actin-related protein 2/3 complex subunit 2 OS=Mus musculus OX=10090 GN=Arpc2 PE=1 SV=3 | 0 | 51.022 | 38 | 14 | 19 | 14 | 300 | 34.3 | 7.36 | 1.032 | 0.883 | 1.169 | 0.05 | -0.05 | -0.18 | -0.06 | 0.065 | 0.23 | 0.912179968 | 0.640216128 | 0.268267457 | 0.999652205 | 0.995559182 | 0.997605693 | 0.869723277 |
| Ddx6 | High | P54823 | Probable ATP-dependent RNA helicase DDX6 OS=Mus musculus OX=10090 GN=Ddx6 PE=1 SV=1 | 0 | 28.717 | 18 | 7 | 11 | 7 | 483 | 54.2 | 8.66 | 1.003 | 0.912 | 1.099 | 0 | 0 | -0.13 | -0.0433333 | 0.065 | 0.14 | 0.934931309 | 0.713628862 | 0.586057745 | 0.999652205 | 0.995559182 | 0.997605693 | 0.973095093 |
| Snx32 | Low | Q80ZJ7 | Sorting nexin-32 OS=Mus musculus OX=10090 GN=Snx32 PE=2 SV=1 | 0.087 | 1.173 | 1 | 1 | 1 | 1 | 404 | 46.6 | 6.96 | 0.973 | 0.94 | 1.035 | -0.04 | 0.04 | -0.09 | -0.03 | 0.065 | 0.05 | 0.840233025 | 0.770361469 | 0.807676414 | 0.999652205 | 0.995559182 | 0.997605693 | 0.992535481 |
| Mpp1 | High | P70290 | 55 kDa erythrocyte membrane protein OS=Mus musculus OX=10090 GN=Mpp1 PE=1 SV=1 | 0.006 | 3.338 | 3 | 1 | 1 | 1 | 466 | 52.2 | 7.2 | 1.06 | 0.873 | 1.213 | 0.08 | -0.08 | -0.2 | -0.0666667 | 0.06 | 0.28 | 0.892146222 | 0.593796472 | 0.277022006 | 0.999652205 | 0.995559182 | 0.997605693 | 0.876067403 |
| | Low | A0A1D5RL | Adenine phosphoribosyltransferase OS=Mus musculus OX=10090 GN=Aprt PE=1 SV=1 | 0.072 | 1.278 | 10 | 1 | 1 | 1 | 67 | 7.6 | 8.92 | 1.01 | 0.917 | 1.102 | 0.01 | -0.01 | -0.13 | -0.0433333 | 0.06 | 0.14 | 0.94760555 | 0.701432941 | 0.537632044 | 0.999652205 | 0.995559182 | 0.997605693 | 0.96694628 |
| Brk1 | Medium | Q91VR8 | Protein BRICK1 OS=Mus musculus OX=10090 GN=Brk1 PE=1 SV=1 | 0.012 | 2.365 | 11 | 1 | 1 | 1 | 75 | 8.8 | 5.45 | 0.989 | 0.935 | 1.057 | -0.02 | 0.02 | -0.1 | -0.0333333 | 0.06 | 0.08 | 0.965149765 | 0.770805444 | 0.72073422 | 0.999652205 | 0.995559182 | 0.997605693 | 0.992535481 |
| Psmd9 | High | Q9CR00 | 26S proteasome non-ATPase regulatory subunit 9 OS=Mus musculus OX=10090 GN=Psmd9 PE=1 | 0 | 37.581 | 32 | 6 | 8 | 6 | 222 | 24.7 | 6.43 | 1.084 | 0.849 | 1.276 | 0.12 | -0.12 | -0.24 | -0.08 | 0.06 | 0.35 | 0.820440773 | 0.556740757 | 0.141846916 | 0.999652205 | 0.995559182 | 0.997605693 | 0.701851981 |
| | High | A0A5K1VV | Protein unc-45 homolog A OS=Mus musculus OX=10090 GN=Unc45a PE=1 SV=1 | 0.007 | 3.058 | 1 | 1 | 1 | 1 | 1088 | 119.1 | 6.9 | 1.049 | 0.878 | 1.195 | 0.07 | -0.07 | -0.19 | -0.0633333 | 0.06 | 0.26 | 0.910846002 | 0.623991487 | 0.304576247 | 0.999652205 | 0.995559182 | 0.997605693 | 0.900241969 |
| Map2k1 | High | P31938 | Dual specificity mitogen-activated protein kinase kinase 1 OS=Mus musculus OX=10090 GN=Map | 0 | 37.324 | 27 | 7 | 13 | 6 | 393 | 43.4 | 6.7 | 1.03 | 0.894 | 1.152 | 0.04 | -0.04 | -0.16 | -0.0533333 | 0.06 | 0.2 | 0.863773069 | 0.666836071 | 0.305266078 | 0.999652205 | 0.995559182 | 0.997605693 | 0.900981762 |
| Pja1 | High | O55176 | E3 ubiquitin-protein ligase Praja-1 OS=Mus musculus OX=10090 GN=Pja1 PE=1 SV=3 | 0.001 | 4.587 | 3 | 2 | 2 | 2 | 578 | 63.9 | 4.97 | 1.044 | 0.882 | 1.185 | 0.06 | -0.06 | -0.18 | -0.06 | 0.06 | 0.24 | 0.913432224 | 0.63642271 | 0.322389851 | 0.999652205 | 0.995559182 | 0.997605693 | 0.905686588 |
| Actr2 | High | P61161 | Actin-related protein 2 OS=Mus musculus OX=10090 GN=Actr2 PE=1 SV=1 | 0 | 63.403 | 25 | 9 | 20 | 9 | 394 | 44.7 | 6.74 | 1.063 | 0.87 | 1.222 | 0.09 | -0.09 | -0.2 | -0.0666667 | 0.055 | 0.29 | 0.834191086 | 0.608917523 | 0.164626672 | 0.999652205 | 0.995559182 | 0.997605693 | 0.736602876 |
| Hk1 | High | G3UVV4 | Hexokinase OS=Mus musculus OX=10090 GN=Hk1 PE=1 SV=1 | 0 | 203.131 | 47 | 40 | 70 | 35 | 917 | 101.8 | 6.67 | 1.005 | 0.921 | 1.091 | 0.01 | -0.01 | -0.12 | -0.04 | 0.055 | 0.13 | 0.984424729 | 0.734827124 | 0.507445718 | 0.999652205 | 0.995559182 | 0.997605693 | 0.964378027 |
| Mff | Medium | Q6PCP5 | Mitochondrial fission factor OS=Mus musculus OX=10090 GN=Mff PE=1 SV=1 | 0.027 | 1.954 | 4 | 1 | 1 | 1 | 291 | 32.9 | 6.83 | 1.017 | 0.915 | 1.112 | 0.02 | -0.02 | -0.13 | -0.0433333 | 0.055 | 0.15 | 0.984598891 | 0.695301948 | 0.515286683 | 0.999652205 | 0.995559182 | 0.997605693 | 0.964378027 |
| Cacna2d2 | High | Q6PHS9 | Voltage-dependent calcium channel subunit alpha-2/delta-2 OS=Mus musculus OX=10090 GN=Ca | 0 | 57.899 | 14 | 14 | 18 | 13 | 1154 | 130.3 | 5.66 | 0.999 | 0.928 | 1.076 | 0 | 0 | -0.11 | -0.0366667 | 0.055 | 0.11 | 0.987774175 | 0.753475313 | 0.563715586 | 0.999652205 | 0.995559182 | 0.997605693 | 0.972910028 |
| | High | D3YYN8 | Non-specific serine/threonine protein kinase OS=Mus musculus OX=10090 GN=Cdc42bpa PE=1 SV | 0.001 | 6.729 | 1 | 2 | 2 | 2 | 1691 | 192.6 | 6.39 | 0.992 | 0.93 | 1.067 | -0.01 | 0.01 | -0.1 | -0.0333333 | 0.055 | 0.09 | 0.953191881 | 0.758018293 | 0.701146508 | 0.999652205 | 0.995559182 | 0.997605693 | 0.991285312 |
| Aipl1 | High | Q924K1 | Aryl-hydrocarbon-interacting protein-like 1 OS=Mus musculus OX=10090 GN=Aipl1 PE=2 SV=2 | 0 | 93.05 | 52 | 15 | 31 | 15 | 328 | 38.2 | 5.95 | 1.147 | 0.808 | 1.419 | 0.2 | -0.2 | -0.31 | -0.1033333 | 0.055 | 0.51 | 0.642018532 | 0.459919906 | 0.018429861 | 0.999652205 | 0.995559182 | 0.997605693 | 0.238069974 |
| Naa25 | High | Q8BWZ3 | N-alpha-acetyltransferase 25, NatB auxiliary subunit OS=Mus musculus OX=10090 GN=Naa25 PE | 0.001 | 6.181 | 3 | 3 | 3 | 3 | 972 | 111.6 | 6.52 | 1.155 | 0.805 | 1.434 | 0.21 | -0.21 | -0.31 | -0.1033333 | 0.05 | 0.52 | 0.60019981 | 0.453724195 | 0.035165902 | 0.999652205 | 0.995559182 | 0.997605693 | 0.367937483 |
| | High | E9PYX3 | Non-specific serine/threonine protein kinase OS=Mus musculus OX=10090 GN=Mark2 PE=1 SV=1 | 0 | 16.493 | 7 | 4 | 4 | 4 | 716 | 80.2 | 9.72 | 1.062 | 0.878 | 1.211 | 0.09 | -0.09 | -0.19 | -0.0633333 | 0.05 | 0.28 | 0.842127693 | 0.626682763 | 0.21424457 | 0.999652205 | 0.995559182 | 0.997605693 | 0.804011919 |
| Dennd6a; | High | Q8BH65 | Protein DENND6A OS=Mus musculus OX=10090 GN=Dennd6a PE=1 SV=1 | 0.001 | 6.021 | 5 | 2 | 2 | 2 | 605 | 68.9 | 7.93 | 1.046 | 0.887 | 1.18 | 0.07 | -0.07 | -0.17 | -0.0566667 | 0.05 | 0.24 | 0.914289293 | 0.628924787 | 0.286888366 | 0.999652205 | 0.995559182 | 0.997605693 | 0.886760237 |
| | High | A0A494BA | Cytosolic iron-sulfur assembly component 3 OS=Mus musculus OX=10090 GN=Ciao3 PE=1 SV=1 | 0.006 | 3.333 | 2 | 1 | 1 | 1 | 377 | 41.7 | 6.3 | 1.044 | 0.896 | 1.165 | 0.06 | -0.06 | -0.16 | -0.0533333 | 0.05 | 0.22 | 0.913644933 | 0.672254699 | 0.375326219 | 0.999652205 | 0.995559182 | 0.997605693 | 0.92018107 |
| | High | A0A3B2W | Tight junction protein ZO-2 OS=Mus musculus OX=10090 GN=Tjp2 PE=1 SV=1 | 0.005 | 3.527 | 1 | 2 | 2 | 2 | 1165 | 131 | 6.79 | 1.028 | 0.908 | 1.132 | 0.04 | -0.04 | -0.14 | -0.0466667 | 0.05 | 0.18 | 0.945191714 | 0.699919109 | 0.457234666 | 0.999652205 | 0.995559182 | 0.997605693 | 0.959744396 |
| Gcc2 | Medium | Q8CHG3 | GRIP and coiled-coil domain-containing protein 2 OS=Mus musculus OX=10090 GN=Gcc2 PE=1 SV | 0.013 | 2.266 | 0 | 1 | 1 | 1 | 1679 | 194.3 | 5.12 | 1.006 | 0.924 | 1.088 | 0.01 | -0.01 | -0.11 | -0.0366667 | 0.05 | 0.12 | 0.960655968 | 0.743364183 | 0.600033594 | 0.999652205 | 0.995559182 | 0.997605693 | 0.977451587 |
| Mapk3 | High | Q63844 | Mitogen-activated protein kinase 3 OS=Mus musculus OX=10090 GN=Mapk3 PE=1 SV=5 | 0 | 20.284 | 19 | 6 | 8 | 3 | 380 | 43 | 6.61 | 1.074 | 0.874 | 1.229 | 0.1 | -0.1 | -0.19 | -0.0633333 | 0.045 | 0.3 | 0.754286742 | 0.617764119 | 0.195744603 | 0.999652205 | 0.995559182 | 0.997605693 | 0.784517504 |
| Syn2 | High | Q64332 | Synapsin-2 OS=Mus musculus OX=10090 GN=Syn2 PE=1 SV=2 | 0 | 24.601 | 9 | 4 | 7 | 2 | 586 | 63.3 | 8.43 | 0.999 | 0.937 | 1.066 | 0 | 0 | -0.09 | -0.03 | 0.045 | 0.09 | 0.919907013 | 0.766814731 | 0.644697171 | 0.999652205 | 0.995559182 | 0.997605693 | 0.985010103 |
| Tiprl | High | Q8BH58 | TIP41-like protein OS=Mus musculus OX=10090 GN=Tiprl PE=1 SV=1 | 0.001 | 7.01 | 7 | 2 | 2 | 2 | 271 | 31.2 | 5.63 | 1.045 | 0.908 | 1.151 | 0.06 | -0.06 | -0.14 | -0.0466667 | 0.04 | 0.2 | 0.91409137 | 0.67962402 | 0.373570332 | 0.999652205 | 0.995559182 | 0.997605693 | 0.920164724 |
| | High | G3UZK1 | Guanine nucleotide-binding protein subunit gamma OS=Mus musculus OX=10090 GN=Gm20503 I | 0.008 | 2.96 | 12 | 1 | 1 | 1 | 150 | 16 | 8.6 | 1.02 | 0.927 | 1.1 | 0.03 | -0.03 | -0.11 | -0.0366667 | 0.04 | 0.14 | 0.988993313 | 0.739633433 | 0.598617648 | 0.999652205 | 0.995559182 | 0.997605693 | 0.977451587 |
| Ctdsp1 | Medium | P58466 | Carboxy-terminal domain RNA polymerase II polypeptide A small phosphatase 1 OS=Mus muscul | 0.017 | 2.127 | 3 | 1 | 1 | 1 | 261 | 29.2 | 6.13 | 1.027 | 0.919 | 1.117 | 0.04 | -0.04 | -0.12 | -0.04 | 0.04 | 0.16 | 0.983999199 | 0.705174225 | 0.446449729 | 0.999652205 | 0.995559182 | 0.997605693 | 0.958608248 |
| Opn1sw | High | P51491 | Short-wave-sensitive opsin 1 OS=Mus musculus OX=10090 GN=Opn1sw PE=1 SV=1 | 0 | 120.217 | 32 | 10 | 87 | 10 | 346 | 38.9 | 8.35 | 1.114 | 0.85 | 1.31 | 0.16 | -0.16 | -0.23 | -0.0766667 | 0.035 | 0.39 | 0.713541679 | 0.559950449 | 0.065522354 | 0.999652205 | 0.995559182 | 0.997605693 | 0.493132672 |
| Rcc2 | High | Q8BK67 | Protein RCC2 OS=Mus musculus OX=10090 GN=Rcc2 PE=1 SV=1 | 0.005 | 3.701 | 2 | 1 | 2 | 1 | 520 | 55.9 | 8.72 | 1.129 | 0.838 | 1.347 | 0.18 | -0.18 | -0.25 | -0.0833333 | 0.035 | 0.43 | 0.659684536 | 0.52796041 | 0.084749667 | 0.999652205 | 0.995559182 | 0.997605693 | 0.56102879 |
| | High | A0A286YD | Calcium/calmodulin-dependent protein kinase type II subunit gamma (Fragment) OS=Mus muscu | 0 | 47.154 | 46 | 9 | 18 | 1 | 241 | 26.1 | 8.73 | 1.018 | 0.933 | 1.091 | 0.03 | -0.03 | -0.1 | -0.0333333 | 0.035 | 0.13 | 0.919574523 | 0.74145371 | 0.5408168 | 0.999652205 | 0.995559182 | 0.997605693 | 0.96694628 |
| Tshr | High | P47750 | Thyrotropin receptor OS=Mus musculus OX=10090 GN=Tshr PE=2 SV=2 | 0 | 11.605 | 5 | 4 | 4 | 4 | 764 | 86.5 | 7.37 | 1.114 | 0.86 | 1.295 | 0.16 | -0.16 | -0.22 | -0.0733333 | 0.03 | 0.37 | 0.716370519 | 0.584279802 | 0.135872696 | 0.999652205 | 0.995559182 | 0.997605693 | 0.699265243 |
| Eif2b3 | High | B1AUN2 | Eukaryotic translation initiation factor 2B, subunit 3 OS=Mus musculus OX=10090 GN=Eif2b3 PE= | 0.001 | 5.343 | 3 | 1 | 1 | 1 | 452 | 50.5 | 6.74 | 1.101 | 0.874 | 1.26 | 0.14 | -0.14 | -0.2 | -0.0666667 | 0.03 | 0.33 | 0.740628767 | 0.616756031 | 0.150758794 | 0.999652205 | 0.995559182 | 0.997605693 | 0.719408135 |
| Ppp2r5a | High | Q6PD03 | Serine/threonine-protein phosphatase 2A 56 kDa regulatory subunit alpha isoform OS=Mus musc | 0 | 21.102 | 16 | 7 | 7 | 6 | 486 | 56.3 | 7.03 | 1.049 | 0.917 | 1.144 | 0.07 | -0.07 | -0.13 | -0.0433333 | 0.03 | 0.19 | 0.826117678 | 0.724716131 | 0.427715731 | 0.999652205 | 0.995559182 | 0.997605693 | 0.943389567 |
| Lsm14b | Medium | Q8BM41 | Protein LSM14 homolog B OS=Mus musculus OX=10090 GN=Lsm14b PE=1 SV=1 | 0.018 | 2.082 | 4 | 1 | 1 | 1 | 305 | 33.2 | 9.74 | 1.168 | 0.832 | 1.404 | 0.22 | -0.22 | -0.27 | -0.09 | 0.025 | 0.49 | 0.601336478 | 0.494724696 | 0.056059739 | 0.999652205 | 0.995559182 | 0.997605693 | 0.454949834 |
| Pacs1 | High | Q8K212 | Phosphofurin acidic cluster sorting protein 1 OS=Mus musculus OX=10090 GN=Pacs1 PE=1 SV=2 | 0 | 24.971 | 7 | 6 | 8 | 6 | 961 | 104.8 | 7.74 | 1.036 | 0.933 | 1.11 | 0.05 | -0.05 | -0.1 | -0.0333333 | 0.025 | 0.15 | 0.935989844 | 0.766425467 | 0.552358859 | 0.999652205 | 0.995559182 | 0.997605693 | 0.967231769 |
| 1700080E | High | E9Q9G1 | Nudix (nucleoside diphosphate-linked moiety X)-type motif 16-like 2 OS=Mus musculus OX=1009 | 0 | 4.944 | 5 | 1 | 2 | 1 | 206 | 22.5 | 6.21 | 1.072 | 0.901 | 1.189 | 0.1 | -0.1 | -0.15 | -0.05 | 0.025 | 0.25 | 0.857330204 | 0.664478789 | 0.296223059 | 0.999652205 | 0.995559182 | 0.997605693 | 0.894411787 |
| Hist4h4; H | High | P62806 | Histone H4 OS=Mus musculus OX=10090 GN=H4c1 PE=1 SV=2 | 0 | 10.052 | 29 | 3 | 4 | 3 | 103 | 11.4 | 11.36 | 0.825 | 1.182 | 0.698 | -0.28 | 0.28 | 0.24 | 0.08 | 0.02 | -0.52 | 0.426316109 | 0.677202849 | 0.043784883 | 0.999652205 | 0.995559182 | 0.997605693 | 0.404911892 |
| Ehd1 | High | Q9WVK4 | EH domain-containing protein 1 OS=Mus musculus OX=10090 GN=Ehd1 PE=1 SV=1 | 0 | 95.026 | 41 | 19 | 29 | 11 | 534 | 60.6 | 6.83 | 1.081 | 0.899 | 1.202 | 0.11 | -0.11 | -0.15 | -0.05 | 0.02 | 0.27 | 0.792001882 | 0.680154849 | 0.198773683 | 0.999652205 | 0.995559182 | 0.997605693 | 0.78697088 |
| Prkcq | High | Q02111 | Protein kinase C theta type OS=Mus musculus OX=10090 GN=Prkcq PE=1 SV=1 | 0 | 12.475 | 5 | 4 | 6 | 3 | 707 | 81.5 | 7.71 | 1.079 | 0.909 | 1.187 | 0.11 | -0.11 | -0.14 | -0.0466667 | 0.015 | 0.25 | 0.731853772 | 0.704303261 | 0.330405574 | 0.999652205 | 0.995559182 | 0.997605693 | 0.906329766 |
| Hars | High | Q61035 | Histidine--tRNA ligase, cytoplasmic OS=Mus musculus OX=10090 GN=Hars1 PE=1 SV=2 | 0 | 35.016 | 20 | 10 | 11 | 10 | 509 | 57.4 | 6 | 1.159 | 0.849 | 1.365 | 0.21 | -0.21 | -0.24 | -0.08 | 0.015 | 0.45 | 0.659531231 | 0.557223455 | 0.073186274 | 0.999652205 | 0.995559182 | 0.997605693 | 0.516302163 |
| Cryba1 | Medium | Q9QXC6 | Beta-A3/A1 crystallin protein OS=Mus musculus OX=10090 GN=Cryba1 PE=1 SV=1 | 0.013 | 2.261 | 4 | 1 | 1 | 1 | 215 | 25.2 | 6.42 | 0.847 | 1.156 | 0.733 | -0.24 | 0.24 | 0.21 | 0.07 | 0.015 | -0.45 | 0.481968695 | 0.70800042 | 0.081392671 | 0.999652205 | 0.995559182 | 0.997605693 | 0.549841817 |
| | High | E9Q4Z6 | Double zinc ribbon and ankyrin repeat-containing protein 1 OS=Mus musculus OX=10090 GN=Dza | 0 | 8.249 | 3 | 2 | 2 | 2 | 779 | 85.1 | 8.18 | 1.083 | 0.901 | 1.203 | 0.12 | -0.12 | -0.15 | -0.05 | 0.015 | 0.27 | 0.839626533 | 0.659818358 | 0.226015477 | 0.999652205 | 0.995559182 | 0.997605693 | 0.813917005 |
| | Medium | Z4YJJ5 | Deubiquitinating enzyme CYLD OS=Mus musculus OX=10090 GN=Cyld PE=1 SV=1 | 0.029 | 1.908 | 1 | 1 | 1 | 1 | 955 | 106.8 | 5.63 | 1.076 | 0.913 | 1.179 | 0.11 | -0.11 | -0.13 | -0.0433333 | 0.01 | 0.24 | 0.860066777 | 0.690303788 | 0.270074118 | 0.999652205 | 0.995559182 | 0.997605693 | 0.871793654 |
| Ehd3 | High | Q9QXY6 | EH domain-containing protein 3 OS=Mus musculus OX=10090 GN=Ehd3 PE=1 SV=2 | 0 | 70.971 | 40 | 18 | 25 | 10 | 535 | 60.8 | 6.46 | 1.058 | 0.933 | 1.135 | 0.08 | -0.08 | -0.1 | -0.0333333 | 0.01 | 0.18 | 0.914223599 | 0.764136147 | 0.457696255 | 0.999652205 | 0.995559182 | 0.997605693 | 0.959744396 |
| Zmat2 | Medium | Q9CPW7 | Zinc finger matrin-type protein 2 OS=Mus musculus OX=10090 GN=Zmat2 PE=2 SV=1 | 0.034 | 1.816 | 4 | 1 | 2 | 1 | 199 | 23.6 | 9.01 | 1.1 | 0.895 | 1.229 | 0.14 | -0.14 | -0.16 | -0.0533333 | 0.01 | 0.3 | 0.740296994 | 0.666853251 | 0.24355325 | 0.999652205 | 0.995559182 | 0.997605693 | 0.837310213 |
| | High | D3YVU0 | Ubiquitin carboxyl-terminal hydrolase 46 OS=Mus musculus OX=10090 GN=Usp46 PE=1 SV=1 | 0 | 4.791 | 5 | 1 | 1 | 1 | 339 | 39.5 | 7.36 | 1.052 | 0.942 | 1.117 | 0.07 | -0.07 | -0.09 | -0.03 | 0.01 | 0.16 | 0.916189434 | 0.763247186 | 0.569779854 | 0.999652205 | 0.995559182 | 0.997605693 | 0.972910028 |
| Herc3 | High | A6H6S0 | Hect domain and RLD 3 OS=Mus musculus OX=10090 GN=Herc3 PE=1 SV=1 | 0 | 15.003 | 6 | 6 | 6 | 6 | 1050 | 117.3 | 6.48 | 1.204 | 0.822 | 1.464 | 0.27 | -0.27 | -0.28 | -0.0933333 | 0.005 | 0.55 | 0.544972999 | 0.493201939 | 0.026299592 | 0.999652205 | 0.995559182 | 0.997605693 | 0.306573414 |
| Hspa1b | High | P17879 | Heat shock 70 kDa protein 1B OS=Mus musculus OX=10090 GN=Hspa1b PE=1 SV=3 | 0 | 310.882 | 58 | 38 | 139 | 1 | 642 | 70.1 | 5.72 | 1.18 | 0.839 | 1.406 | 0.24 | -0.24 | -0.25 | -0.0833333 | 0.005 | 0.49 | 0.555389246 | 0.507393747 | 0.027442038 | 0.999652205 | 0.995559182 | 0.997605693 | 0.315745518 |
| Slc1a2 | High | P43006 | Excitatory amino acid transporter 2 OS=Mus musculus OX=10090 GN=Slc1a2 PE=1 SV=1 | 0 | 28.846 | 14 | 7 | 9 | 7 | 572 | 62 | 6.68 | 0.84 | 1.184 | 0.709 | -0.25 | 0.25 | 0.24 | 0.08 | 0.005 | -0.5 | 0.459470309 | 0.665619123 | 0.035316809 | 0.999652205 | 0.995559182 | 0.997605693 | 0.367937483 |
| | High | V9GWX3 | RNA polymerase II-associated protein 1 OS=Mus musculus OX=10090 GN=Rpap1 PE=1 SV=1 | 0.001 | 6.127 | 2 | 2 | 2 | 2 | 869 | 96.3 | 6.37 | 0.828 | 1.199 | 0.69 | -0.27 | 0.27 | 0.26 | 0.08666667 | 0.005 | -0.53 | 0.444207635 | 0.638787774 | 0.062762438 | 0.999652205 | 0.995559182 | 0.997605693 | 0.48140765 |
| Canx | High | P35564 | Calnexin OS=Mus musculus OX=10090 GN=Canx PE=1 SV=1 | 0 | 106.683 | 42 | 24 | 46 | 24 | 591 | 67.2 | 4.64 | 0.834 | 1.201 | 0.694 | -0.26 | 0.26 | 0.26 | 0.08666667 | 0 | -0.53 | 0.542749083 | 0.633734573 | 0.024245104 | 0.999652205 | 0.995559182 | 0.997605693 | 0.285927032 |
| Dlg2 | High | D3YWU0 | Disks large homolog 2 OS=Mus musculus OX=10090 GN=Dlg2 PE=1 SV=1 | 0 | 11.875 | 9 | 4 | 4 | 1 | 481 | 54.7 | 6.71 | 0.864 | 1.153 | 0.749 | -0.21 | 0.21 | 0.21 | 0.07 | 0 | -0.42 | 0.527853087 | 0.735007311 | 0.108777133 | 0.999652205 | 0.995559182 | 0.997605693 | 0.631380318 |
| Prpf38b | High | Q80SY5 | Pre-mRNA-splicing factor 38B OS=Mus musculus OX=10090 GN=Prpf38b PE=1 SV=1 | 0.001 | 6.219 | 2 | 1 | 1 | 1 | 542 | 63.7 | 10.54 | 1.178 | 0.847 | 1.39 | 0.24 | -0.24 | -0.24 | -0.08 | 0 | 0.48 | 0.609175809 | 0.529208195 | 0.110271403 | 0.999652205 | 0.995559182 | 0.997605693 | 0.632640059 |
| Sec61a1 | Low | P61620 | Protein transport protein Sec61 subunit alpha isoform 1 OS=Mus musculus OX=10090 GN=Sec61a | 0.051 | 1.486 | 1 | 1 | 1 | 1 | 476 | 52.2 | 8.06 | 1.098 | 0.917 | 1.198 | 0.13 | -0.13 | -0.13 | -0.0433333 | 0 | 0.26 | 0.773161943 | 0.721580261 | 0.290183677 | 0.999652205 | 0.995559182 | 0.997605693 | 0.889103759 |
| Gab2 | Medium | Q3ZB59 | GRB2-associated-binding protein 2 OS=Mus musculus OX=10090 GN=Gab2 PE=1 SV=1 | 0.029 | 1.912 | 2 | 1 | 1 | 1 | 665 | 73.2 | 8.31 | 1.08 | 0.926 | 1.166 | 0.11 | -0.11 | -0.11 | -0.0366667 | 0 | 0.22 | 0.836959848 | 0.735084694 | 0.379379496 | 0.999652205 | 0.995559182 | 0.997605693 | 0.922487352 |
| Gng13 | High | Q9JMF3 | Guanine nucleotide-binding protein G(I)/G(S)/G(O) subunit gamma-13 OS=Mus musculus OX=10 | 0 | 10.436 | 58 | 4 | 6 | 4 | 67 | 8 | 5.45 | 0.854 | 1.185 | 0.721 | -0.23 | 0.23 | 0.24 | 0.08 | -0.005 | -0.47 | 0.510265583 | 0.663949746 | 0.042541818 | 0.999652205 | 0.995559182 | 0.997605693 | 0.401420039 |
| Fn3krp | High | Q8K274 | Ketosamine-3-kinase OS=Mus musculus OX=10090 GN=Fn3krp PE=1 SV=2 | 0 | 44.88 | 30 | 9 | 15 | 9 | 309 | 34.4 | 7.87 | 1.135 | 0.892 | 1.273 | 0.18 | -0.18 | -0.17 | -0.0566667 | -0.005 | 0.35 | 0.609948062 | 0.661607924 | 0.090457769 | 0.999652205 | 0.995559182 | 0.997605693 | 0.575705685 |
| Rapgef4 | High | Q9EQZ6 | Rap guanine nucleotide exchange factor 4 OS=Mus musculus OX=10090 GN=Rapgef4 PE=1 SV=1 | 0.005 | 3.743 | 3 | 2 | 2 | 2 | 1011 | 115.4 | 6.92 | 1.086 | 0.926 | 1.173 | 0.12 | -0.12 | -0.11 | -0.0366667 | -0.005 | 0.23 | 0.853112725 | 0.731612756 | 0.286219212 | 0.999652205 | 0.995559182 | 0.997605693 | 0.886074068 |
| Fn3k | High | Q9ER35 | Fructosamine-3-kinase OS=Mus musculus OX=10090 GN=Fn3k PE=1 SV=1 | 0 | 25.14 | 21 | 5 | 6 | 5 | 309 | 35 | 8.4 | 1.242 | 0.81 | 1.533 | 0.31 | -0.31 | -0.3 | -0.1 | -0.005 | 0.62 | 0.466570099 | 0.464891513 | 0.011352447 | 0.999652205 | 0.995559182 | 0.997605693 | 0.173441884 |
| Herc4 | High | Q6PAV2 | Probable E3 ubiquitin-protein ligase HERC4 OS=Mus musculus OX=10090 GN=Herc4 PE=1 SV=2 | 0.002 | 4.173 | 2 | 2 | 2 | 2 | 1057 | 118.3 | 6.29 | 1.217 | 0.828 | 1.47 | 0.28 | -0.28 | -0.27 | -0.09 | -0.005 | 0.56 | 0.495275833 | 0.47553262 | 0.013191853 | 0.999652205 | 0.995559182 | 0.997605693 | 0.190145812 |
| Lrrc57 | High | Q9D1G5 | Leucine-rich repeat-containing protein 57 OS=Mus musculus OX=10090 GN=Lrrc57 PE=1 SV=1 | 0 | 97.792 | 66 | 16 | 41 | 16 | 239 | 26.7 | 8.35 | 1.197 | 0.842 | 1.421 | 0.26 | -0.26 | -0.25 | -0.0833333 | -0.005 | 0.51 | 0.542791505 | 0.54030379 | 0.017977541 | 0.999652205 | 0.995559182 | 0.997605693 | 0.237176647 |
| Ptk7 | High | Q8BKG3 | Inactive tyrosine-protein kinase 7 OS=Mus musculus OX=10090 GN=Ptk7 PE=1 SV=1 | 0.001 | 5.384 | 2 | 1 | 1 | 1 | 1062 | 117.5 | 6.84 | 1.207 | 0.832 | 1.45 | 0.27 | -0.27 | -0.26 | -0.0866667 | -0.005 | 0.54 | 0.54768898 | 0.493070299 | 0.064147431 | 0.999652205 | 0.995559182 | 0.997605693 | 0.486704528 |
| | High | G3UWV4 | Serine/threonine-protein kinase BRSK2 OS=Mus musculus OX=10090 GN=Brsk2 PE=1 SV=1 | 0 | 34.832 | 15 | 10 | 14 | 7 | 697 | 77.7 | 8.82 | 1.092 | 0.923 | 1.183 | 0.13 | -0.13 | -0.12 | -0.04 | -0.005 | 0.24 | 0.706978698 | 0.739789945 | 0.236315765 | 0.999652205 | 0.995559182 | 0.997605693 | 0.828110564 |
| Abhd12 | High | Q99LR1 | Lysophosphatidylserine lipase ABHD12 OS=Mus musculus OX=10090 GN=Abhd12 PE=1 SV=2 | 0 | 15.142 | 19 | 6 | 7 | 6 | 398 | 45.2 | 8.72 | 0.861 | 1.177 | 0.731 | -0.22 | 0.22 | 0.24 | 0.08 | -0.01 | -0.45 | 0.552447366 | 0.678298732 | 0.069765074 | 0.999652205 | 0.995559182 | 0.997605693 | 0.50473906 |
| Uso1 | High | Q9Z1Z0 | General vesicular transport factor p115 OS=Mus musculus OX=10090 GN=Uso1 PE=1 SV=2 | 0 | 17.224 | 5 | 4 | 4 | 4 | 959 | 106.9 | 4.93 | 0.872 | 1.166 | 0.748 | -0.2 | 0.2 | 0.22 | 0.07333333 | -0.01 | -0.42 | 0.603787182 | 0.699522037 | 0.085248202 | 0.999652205 | 0.995559182 | 0.997605693 | 0.56316433 |
| D16Ertd47 | High | Q9D7G4 | Protein EURL homolog OS=Mus musculus OX=10090 GN=Eurl PE=1 SV=1 | 0 | 8.982 | 11 | 3 | 3 | 3 | 290 | 32.9 | 6.9 | 1.158 | 0.875 | 1.322 | 0.21 | -0.21 | -0.19 | -0.0633333 | -0.01 | 0.4 | 0.59432004 | 0.621564434 | 0.102828289 | 0.999652205 | 0.995559182 | 0.997605693 | 0.610114517 |
| 1700047I1 | High | Q8BR63 | Protein FAM177A1 OS=Mus musculus OX=10090 GN=Fam177a1 PE=1 SV=1 | 0.001 | 5.266 | 10 | 2 | 2 | 2 | 207 | 23.6 | 4.59 | 0.848 | 1.196 | 0.709 | -0.24 | 0.24 | 0.26 | 0.08666667 | -0.01 | -0.5 | 0.51867876 | 0.665638162 | 0.111332604 | 0.999652205 | 0.995559182 | 0.997605693 | 0.632640059 |
| Tardbp | High | Q8R0B4 | TAR DNA-binding protein 43 OS=Mus musculus OX=10090 GN=Tardbp PE=1 SV=1 | 0.001 | 7.592 | 8 | 2 | 2 | 2 | 295 | 33.3 | 7.06 | 1.146 | 0.89 | 1.287 | 0.2 | -0.2 | -0.17 | -0.0566667 | -0.015 | 0.36 | 0.647014502 | 0.654493015 | 0.14219794 | 0.999652205 | 0.995559182 | 0.997605693 | 0.701851981 |
| Mecp2 | High | Q9Z2D6 | Methyl-CpG-binding protein 2 OS=Mus musculus OX=10090 GN=Mecp2 PE=1 SV=1 | 0.003 | 3.858 | 4 | 2 | 2 | 2 | 484 | 52.3 | 9.96 | 1.148 | 0.891 | 1.288 | 0.2 | -0.2 | -0.17 | -0.0566667 | -0.015 | 0.36 | 0.618976903 | 0.661356964 | 0.154248373 | 0.999652205 | 0.995559182 | 0.997605693 | 0.728999569 |
| Eif5a | High | A0A0A0M | Eukaryotic translation initiation factor 5A (Fragment) OS=Mus musculus OX=10090 GN=Eif5a PE= | 0 | 73.596 | 58 | 12 | 28 | 12 | 149 | 16.3 | 5 | 1.217 | 0.841 | 1.447 | 0.28 | -0.28 | -0.25 | -0.0833333 | -0.015 | 0.53 | 0.506238333 | 0.53828062 | 0.013138883 | 0.999652205 | 0.995559182 | 0.997605693 | 0.189870411 |
| Prrc1 | Medium | Q3UPH1 | Protein PRRC1 OS=Mus musculus OX=10090 GN=Prrc1 PE=1 SV=1 | 0.01 | 2.585 | 2 | 1 | 1 | 1 | 443 | 46.3 | 5.95 | 0.837 | 1.228 | 0.682 | -0.26 | 0.26 | 0.3 | 0.1 | -0.02 | -0.55 | 0.42914582 | 0.577587339 | 0.035347537 | 0.999652205 | 0.995559182 | 0.997605693 | 0.367937483 |
| Nos1 | High | F8WGF2 | Constitutive NOS OS=Mus musculus OX=10090 GN=Nos1 PE=1 SV=1 | 0.001 | 4.269 | 1 | 1 | 1 | 1 | 1463 | 164.2 | 7.24 | 1.18 | 0.871 | 1.355 | 0.24 | -0.24 | -0.2 | -0.0666667 | -0.02 | 0.44 | 0.567995464 | 0.604039632 | 0.087976842 | 0.999652205 | 0.995559182 | 0.997605693 | 0.573625007 |
| Sec14l2 | High | Q99J08 | SEC14-like protein 2 OS=Mus musculus OX=10090 GN=Sec14l2 PE=1 SV=1 | 0 | 58.465 | 37 | 11 | 15 | 11 | 403 | 46.3 | 7.11 | 1.125 | 0.916 | 1.228 | 0.17 | -0.17 | -0.13 | -0.0433333 | -0.02 | 0.3 | 0.661634761 | 0.723583418 | 0.160468913 | 0.999652205 | 0.995559182 | 0.997605693 | 0.733684585 |
| Mxra7 | High | Q9D1T2 | Matrix-remodeling-associated protein 7 OS=Mus musculus OX=10090 GN=Mxra7 PE=1 SV=1 | 0 | 19.004 | 46 | 3 | 5 | 1 | 151 | 16.2 | 4.13 | 0.84 | 1.229 | 0.683 | -0.25 | 0.25 | 0.3 | 0.1 | -0.025 | -0.55 | 0.44546681 | 0.583420407 | 0.056418038 | 0.999652205 | 0.995559182 | 0.997605693 | 0.455815471 |
| Bzw1 | High | A0A087W | Basic leucine zipper and W2 domain-containing protein 1 OS=Mus musculus OX=10090 GN=Bzw1 | 0 | 24.825 | 20 | 8 | 9 | 8 | 451 | 51.2 | 6.95 | 1.14 | 0.906 | 1.259 | 0.19 | -0.19 | -0.14 | -0.0466667 | -0.025 | 0.33 | 0.680969436 | 0.696993083 | 0.171604044 | 0.999652205 | 0.995559182 | 0.997605693 | 0.738844686 |
| Dync1li2 | High | Q6PDL0 | Cytoplasmic dynein 1 light intermediate chain 2 OS=Mus musculus OX=10090 GN=Dync1li2 PE=1 | 0 | 32.466 | 11 | 5 | 8 | 5 | 492 | 54.2 | 6.28 | 1.113 | 0.933 | 1.193 | 0.15 | -0.15 | -0.1 | -0.0333333 | -0.025 | 0.25 | 0.642703 | 0.765875536 | 0.30777273 | 0.999652205 | 0.995559182 | 0.997605693 | 0.904445334 |
| Slc6a17 | High | A0A0R4J0 | Transporter OS=Mus musculus OX=10090 GN=Slc6a17 PE=1 SV=1 | 0.005 | 3.728 | 2 | 1 | 1 | 1 | 724 | 80.7 | 6.23 | 0.904 | 1.156 | 0.782 | -0.15 | 0.15 | 0.21 | 0.07 | -0.03 | -0.35 | 0.632558764 | 0.711006591 | 0.179997525 | 0.999652205 | 0.995559182 | 0.997605693 | 0.756556316 |
| Ruvbl1 | High | P60122 | RuvB-like 1 OS=Mus musculus OX=10090 GN=Ruvbl1 PE=1 SV=1 | 0 | 28.017 | 20 | 7 | 9 | 7 | 456 | 50.2 | 6.42 | 1.122 | 0.933 | 1.202 | 0.17 | -0.17 | -0.1 | -0.0333333 | -0.035 | 0.27 | 0.723114108 | 0.766282798 | 0.243776287 | 0.999652205 | 0.995559182 | 0.997605693 | 0.837310213 |
| Rspry1 | High | Q8BVR6 | RING finger and SPRY domain-containing protein 1 OS=Mus musculus OX=10090 GN=Rspry1 PE= | 0 | 11.448 | 4 | 3 | 3 | 3 | 576 | 64.3 | 5.5 | 0.917 | 1.155 | 0.793 | -0.13 | 0.13 | 0.21 | 0.07 | -0.04 | -0.33 | 0.706911559 | 0.712274341 | 0.219425696 | 0.999652205 | 0.995559182 | 0.997605693 | 0.805859554 |
| Hspa1l | High | P16627 | Heat shock 70 kDa protein 1-like OS=Mus musculus OX=10090 GN=Hspa1l PE=1 SV=4 | 0 | 147.779 | 29 | 21 | 90 | 1 | 641 | 70.6 | 6.24 | 1.141 | 0.93 | 1.227 | 0.19 | -0.19 | -0.11 | -0.0366667 | -0.04 | 0.3 | 0.695900015 | 0.735143779 | 0.175431261 | 0.999652205 | 0.995559182 | 0.997605693 | 0.74518415 |
| Glt28d2 | High | Q8BML3 | N-acetylglucosaminyldiphosphodolichol N-acetylglucosaminyltransferase OS=Mus musculus OX=1 | 0.001 | 4.605 | 11 | 2 | 2 | 2 | 165 | 18.4 | 6.67 | 1.201 | 0.882 | 1.361 | 0.26 | -0.26 | -0.18 | -0.06 | -0.04 | 0.44 | 0.528937852 | 0.620427565 | 0.086159021 | 0.999652205 | 0.995559182 | 0.997605693 | 0.567677591 |
| Pank4 | High | Q80YV4 | 4'-phosphopantetheine phosphatase OS=Mus musculus OX=10090 GN=Pank4 PE=1 SV=2 | 0 | 61.518 | 17 | 12 | 18 | 12 | 820 | 91.5 | 6.71 | 1.145 | 0.923 | 1.24 | 0.2 | -0.2 | -0.12 | -0.04 | -0.04 | 0.31 | 0.645778619 | 0.741042871 | 0.136811518 | 0.999652205 | 0.995559182 | 0.997605693 | 0.700189449 |
| Eif4g2 | High | Q62448 | Eukaryotic translation initiation factor 4 gamma 2 OS=Mus musculus OX=10090 GN=Eif4g2 PE=1 | 0 | 15.478 | 6 | 5 | 6 | 5 | 906 | 102 | 7.14 | 1.156 | 0.919 | 1.258 | 0.21 | -0.21 | -0.12 | -0.04 | -0.045 | 0.33 | 0.66342848 | 0.729753319 | 0.15933505 | 0.999652205 | 0.995559182 | 0.997605693 | 0.733684585 |
| Cpe | High | Q00493 | Carboxypeptidase E OS=Mus musculus OX=10090 GN=Cpe PE=1 SV=2 | 0 | 19.425 | 10 | 4 | 5 | 4 | 476 | 53.2 | 5.19 | 0.866 | 1.24 | 0.699 | -0.21 | 0.21 | 0.31 | 0.10333333 | -0.05 | -0.52 | 0.546086499 | 0.565431033 | 0.031546379 | 0.999652205 | 0.995559182 | 0.997605693 | 0.345469816 |
| Gstt2 | High | Q61133 | Glutathione S-transferase theta-2 OS=Mus musculus OX=10090 GN=Gstt2 PE=1 SV=4 | 0 | 12.125 | 15 | 3 | 3 | 2 | 244 | 27.6 | 7.55 | 1.141 | 0.937 | 1.218 | 0.19 | -0.19 | -0.09 | -0.03 | -0.05 | 0.28 | 0.688649255 | 0.749298168 | 0.213027393 | 0.999652205 | 0.995559182 | 0.997605693 | 0.804011919 |
| Sort1 | High | Q6PHU5 | Sortilin OS=Mus musculus OX=10090 GN=Sort1 PE=1 SV=1 | 0.001 | 5.341 | 1 | 1 | 1 | 1 | 825 | 91.1 | 5.88 | 0.906 | 1.192 | 0.76 | -0.14 | 0.14 | 0.25 | 0.08333333 | -0.055 | -0.4 | 0.653613314 | 0.654806699 | 0.12394691 | 0.999652205 | 0.995559182 | 0.997605693 | 0.677192798 |
| Larp1 | High | Z4YJT3 | La-related protein 1 OS=Mus musculus OX=10090 GN=Larp1 PE=1 SV=1 | 0 | 15.93 | 4 | 2 | 3 | 2 | 1072 | 121.1 | 8.79 | 1.245 | 0.873 | 1.427 | 0.32 | -0.32 | -0.2 | -0.0666667 | -0.06 | 0.51 | 0.451486444 | 0.599637738 | 0.029085531 | 0.999652205 | 0.995559182 | 0.997605693 | 0.324925684 |
| Ube2f | High | Q9CY34 | NEDD8-conjugating enzyme UBE2F OS=Mus musculus OX=10090 GN=Ube2f PE=1 SV=1 | 0 | 13.48 | 22 | 3 | 3 | 3 | 185 | 21.1 | 6.54 | 1.235 | 0.88 | 1.404 | 0.3 | -0.3 | -0.18 | -0.06 | -0.06 | 0.49 | 0.446996367 | 0.631859892 | 0.055291249 | 0.999652205 | 0.995559182 | 0.997605693 | 0.450649022 |
| 9630033F | High | Q8BZA9 | Fructose-2,6-bisphosphatase TIGAR OS=Mus musculus OX=10090 GN=Tigar PE=1 SV=1 | 0 | 10.596 | 15 | 4 | 4 | 4 | 269 | 29.2 | 8.18 | 1.18 | 0.921 | 1.282 | 0.24 | -0.24 | -0.12 | -0.04 | -0.06 | 0.36 | 0.488175565 | 0.734450711 | 0.157369707 | 0.999652205 | 0.995559182 | 0.997605693 | 0.733365826 |
| Sh3glb2 | High | A2AWI7 | Endophilin-B2 OS=Mus musculus OX=10090 GN=Sh3glb2 PE=1 SV=1 | 0 | 29.161 | 28 | 10 | 12 | 9 | 404 | 44.9 | 5.58 | 0.937 | 1.16 | 0.808 | -0.09 | 0.09 | 0.21 | 0.07 | -0.06 | -0.31 | 0.788443246 | 0.71159526 | 0.201115243 | 0.999652205 | 0.995559182 | 0.997605693 | 0.78697088 |
| 6430548M | High | Q3TA40 | RIKEN cDNA 6430548M08 gene OS=Mus musculus OX=10090 GN=6430548M08Rik PE=1 SV=1 | 0 | 73.027 | 37 | 14 | 20 | 14 | 397 | 45.2 | 5.07 | 0.959 | 1.135 | 0.845 | -0.06 | 0.06 | 0.18 | 0.06 | -0.06 | -0.24 | 0.90568256 | 0.761832046 | 0.344374542 | 0.999652205 | 0.995559182 | 0.997605693 | 0.907770836 |
| Ssr4 | High | Q9D8L3 | Signal sequence receptor subunit delta OS=Mus musculus OX=10090 GN=Ssr4 PE=1 SV=1 | 0.005 | 3.701 | 6 | 1 | 1 | 1 | 173 | 19 | 5.78 | 0.86 | 1.278 | 0.673 | -0.22 | 0.22 | 0.35 | 0.11666667 | -0.065 | -0.57 | 0.493138712 | 0.489856603 | 0.027576175 | 0.999652205 | 0.995559182 | 0.997605693 | 0.315745518 |
| Epb4.1; Ep | High | A2A842 | Band 4.1 OS=Mus musculus OX=10090 GN=Epb41 PE=1 SV=1 | 0 | 46.563 | 17 | 11 | 16 | 10 | 769 | 85.6 | 5.39 | 0.892 | 1.23 | 0.725 | -0.17 | 0.17 | 0.3 | 0.1 | -0.065 | -0.46 | 0.716833053 | 0.582233592 | 0.054867746 | 0.999652205 | 0.995559182 | 0.997605693 | 0.449114524 |
| Scrn1 | High | Q9CZC8 | Secernin-1 OS=Mus musculus OX=10090 GN=Scrn1 PE=1 SV=1 | 0 | 23.842 | 16 | 5 | 7 | 5 | 414 | 46.3 | 4.79 | 0.934 | 1.17 | 0.798 | -0.1 | 0.1 | 0.23 | 0.07666667 | -0.065 | -0.32 | 0.757825108 | 0.691600573 | 0.179646809 | 0.999652205 | 0.995559182 | 0.997605693 | 0.756556316 |
| Otud6b | Medium | Q8K2H2 | Deubiquitinase OTUD6B OS=Mus musculus OX=10090 GN=Otud6b PE=1 SV=1 | 0.01 | 2.542 | 3 | 1 | 1 | 1 | 294 | 33.7 | 5.53 | 1.167 | 0.94 | 1.241 | 0.22 | -0.22 | -0.09 | -0.03 | -0.065 | 0.31 | 0.621805041 | 0.760087087 | 0.230109217 | 0.999652205 | 0.995559182 | 0.997605693 | 0.819186996 |
| Pde8b | High | E9Q4S1 | High affinity cAMP-specific and IBMX-insensitive 3',5'-cyclic phosphodiesterase 8B OS=Mus musc | 0 | 51.444 | 13 | 12 | 19 | 10 | 865 | 96.7 | 6.86 | 0.959 | 1.141 | 0.841 | -0.06 | 0.06 | 0.19 | 0.06333333 | -0.065 | -0.25 | 0.903981369 | 0.750804748 | 0.331917016 | 0.999652205 | 0.995559182 | 0.997605693 | 0.906329766 |
| Msn | High | P26041 | Moesin OS=Mus musculus OX=10090 GN=Msn PE=1 SV=3 | 0 | 249.459 | 59 | 45 | 117 | 32 | 577 | 67.7 | 6.6 | 0.968 | 1.135 | 0.853 | -0.05 | 0.05 | 0.18 | 0.06 | -0.065 | -0.23 | 0.916517809 | 0.762562128 | 0.356617755 | 0.999652205 | 0.995559182 | 0.997605693 | 0.914600864 |
| Sorl1 | High | O88307 | Sortilin-related receptor OS=Mus musculus OX=10090 GN=Sorl1 PE=1 SV=3 | 0.001 | 4.292 | 0 | 1 | 1 | 1 | 2215 | 246.9 | 5.54 | 0.839 | 1.313 | 0.638 | -0.25 | 0.25 | 0.39 | 0.13 | -0.07 | -0.65 | 0.442210147 | 0.452622473 | 0.010641396 | 0.999652205 | 0.995559182 | 0.997605693 | 0.165739737 |
| Cpne1 | High | Q8C166 | Copine-1 OS=Mus musculus OX=10090 GN=Cpne1 PE=1 SV=1 | 0 | 36.025 | 18 | 11 | 15 | 10 | 536 | 58.8 | 5.66 | 0.917 | 1.205 | 0.76 | -0.13 | 0.13 | 0.27 | 0.09 | -0.07 | -0.4 | 0.801744648 | 0.626102206 | 0.095930776 | 0.999652205 | 0.995559182 | 0.997605693 | 0.590616329 |
| Arhgap26 | High | E9QAQ3 | Rho GTPase-activating protein 26 OS=Mus musculus OX=10090 GN=Arhgap26 PE=1 SV=1 | 0.001 | 7.827 | 3 | 3 | 3 | 3 | 759 | 85.9 | 6.74 | 1.187 | 0.927 | 1.281 | 0.25 | -0.25 | -0.11 | -0.0366667 | -0.07 | 0.36 | 0.545802265 | 0.72519872 | 0.105557618 | 0.999652205 | 0.995559182 | 0.997605693 | 0.619102056 |
| Sh3gl1 | High | Q62419 | Endophilin-A2 OS=Mus musculus OX=10090 GN=Sh3gl1 PE=1 SV=1 | 0 | 18.072 | 17 | 7 | 10 | 2 | 368 | 41.5 | 5.72 | 0.894 | 1.237 | 0.723 | -0.16 | 0.16 | 0.31 | 0.10333333 | -0.075 | -0.47 | 0.560688566 | 0.564418646 | 0.035521764 | 0.999652205 | 0.995559182 | 0.997605693 | 0.368152557 |
| Reep5 | High | G3X8R0 | Receptor expression-enhancing protein OS=Mus musculus OX=10090 GN=Reep5 PE=1 SV=1 | 0 | 12.225 | 21 | 5 | 7 | 5 | 189 | 21.4 | 8.12 | 0.932 | 1.19 | 0.784 | -0.1 | 0.1 | 0.25 | 0.08333333 | -0.075 | -0.35 | 0.752019075 | 0.65499049 | 0.134736658 | 0.999652205 | 0.995559182 | 0.997605693 | 0.697570122 |

| | | | | | | | | | | | | | | | | | | | | | | | | | | | |
|---|---|---|---|---|---|---|---|---|---|---|---|---|---|---|---|---|---|---|---|---|---|---|---|---|---|---|---|
| Epb4.1l2; I | High | Q80UE5 | Band 4.1-like protein 2 OS=Mus musculus OX=10090 GN=Epb41l2 PE=1 SV=1 | 0 | 225.991 | 41 | 37 | 78 | 1 | 918 | 101.8 | 5.33 | 0.932 | 1.19 | 0.784 | -0.1 | 0.1 | 0.25 | 0.08333333 | -0.075 | -0.35 | 0.712181437 | 0.654395406 | 0.144434872 | 0.999652205 | 0.995559182 | 0.997605693 | 0.706643254 |
| Preb | High | D3Z3S1 | Prolactin regulatory element-binding protein OS=Mus musculus OX=10090 GN=Preb PE=1 SV=1 | 0.001 | 6.541 | 6 | 2 | 2 | 2 | 350 | 37.9 | 8.56 | 0.981 | 1.133 | 0.866 | -0.03 | 0.03 | 0.18 | 0.06 | -0.075 | -0.21 | 0.865296695 | 0.766262044 | 0.40285599 | 0.999652205 | 0.995559182 | 0.997605693 | 0.93262326 |
| Isyna1 | High | Q9JHU9 | Inositol-3-phosphate synthase 1 OS=Mus musculus OX=10090 GN=Isyna1 PE=1 SV=1 | 0.001 | 5.097 | 4 | 1 | 3 | 1 | 557 | 60.9 | 6.42 | 0.97 | 1.151 | 0.843 | -0.04 | 0.04 | 0.2 | 0.06666667 | -0.08 | -0.25 | 0.808985687 | 0.729189125 | 0.314905009 | 0.999652205 | 0.995559182 | 0.997605693 | 0.904943958 |
| | High | A0A0U1RF | Non-specific serine/threonine protein kinase OS=Mus musculus OX=10090 GN=Csnk1g1 PE=1 SV= | 0 | 35.138 | 28 | 12 | 18 | 6 | 424 | 48.9 | 8.7 | 0.948 | 1.189 | 0.797 | -0.08 | 0.08 | 0.25 | 0.08333333 | -0.085 | -0.33 | 0.804783273 | 0.656113055 | 0.165486259 | 0.999652205 | 0.995559182 | 0.997605693 | 0.736602876 |
| Erp29 | High | P57759 | Endoplasmic reticulum resident protein 29 OS=Mus musculus OX=10090 GN=Erp29 PE=1 SV=2 | 0.001 | 4.551 | 12 | 3 | 3 | 3 | 262 | 28.8 | 6.15 | 0.994 | 1.133 | 0.877 | -0.01 | 0.01 | 0.18 | 0.06 | -0.085 | -0.19 | 0.905888129 | 0.765469722 | 0.450110704 | 0.999652205 | 0.995559182 | 0.997605693 | 0.959449555 |
| Calb2 | High | Q08331 | Calretinin OS=Mus musculus OX=10090 GN=Calb2 PE=1 SV=3 | 0 | 56.392 | 41 | 12 | 19 | 11 | 271 | 31.4 | 5.02 | 0.897 | 1.254 | 0.715 | -0.16 | 0.16 | 0.33 | 0.11 | -0.085 | -0.48 | 0.732046427 | 0.542902689 | 0.037615505 | 0.999652205 | 0.995559182 | 0.997605693 | 0.373579725 |
| Arl6ip1 | Medium | Q9JKW0 | ADP-ribosylation factor-like protein 6-interacting protein 1 OS=Mus musculus OX=10090 GN=Arl6 | 0.029 | 1.911 | 4 | 1 | 1 | 1 | 203 | 23.4 | 9.32 | 0.986 | 1.142 | 0.864 | -0.02 | 0.02 | 0.19 | 0.06333333 | -0.085 | -0.21 | 0.881299974 | 0.752511292 | 0.498193726 | 0.999652205 | 0.995559182 | 0.997605693 | 0.964378027 |
| Slc32a1 | High | O35633 | Vesicular inhibitory amino acid transporter OS=Mus musculus OX=10090 GN=Slc32a1 PE=1 SV=3 | 0 | 59.973 | 24 | 10 | 18 | 10 | 525 | 57.3 | 6.64 | 0.958 | 1.18 | 0.812 | -0.06 | 0.06 | 0.24 | 0.08 | -0.09 | -0.3 | 0.888812753 | 0.673006789 | 0.214685783 | 0.999652205 | 0.995559182 | 0.997605693 | 0.804011919 |
| Myo1d | High | Q5SYD0 | Unconventional myosin-Id OS=Mus musculus OX=10090 GN=Myo1d PE=1 SV=1 | 0 | 17.224 | 8 | 7 | 7 | 7 | 1006 | 116 | 9.41 | 0.975 | 1.168 | 0.835 | -0.04 | 0.04 | 0.22 | 0.07333333 | -0.09 | -0.26 | 0.948796661 | 0.696605029 | 0.311526941 | 0.999652205 | 0.995559182 | 0.997605693 | 0.904943958 |
| Cltc | High | Q68FD5 | Clathrin heavy chain 1 OS=Mus musculus OX=10090 GN=Cltc PE=1 SV=3 | 0 | 503.327 | 48 | 66 | 160 | 66 | 1675 | 191.4 | 5.69 | 0.99 | 1.145 | 0.865 | -0.01 | 0.01 | 0.19 | 0.06333333 | -0.09 | -0.21 | 0.974669242 | 0.74263758 | 0.402580925 | 0.999652205 | 0.995559182 | 0.997605693 | 0.93262326 |
| | High | F6T4M4 | Serine/arginine repetitive matrix protein 1 (Fragment) OS=Mus musculus OX=10090 GN=Srrm1 P | 0.001 | 6.513 | 6 | 1 | 12 | 1 | 140 | 16.2 | 5.38 | 0.94 | 1.206 | 0.78 | -0.09 | 0.09 | 0.27 | 0.09 | -0.09 | -0.36 | 0.785658643 | 0.625561785 | 0.128082523 | 0.999652205 | 0.995559182 | 0.997605693 | 0.680719152 |
| Snx1 | High | Q6NZD2 | Sorting nexin-1 OS=Mus musculus OX=10090 GN=Snx1 PE=1 SV=1 | 0 | 18.279 | 9 | 5 | 8 | 4 | 521 | 58.8 | 5.26 | 0.933 | 1.214 | 0.769 | -0.1 | 0.1 | 0.28 | 0.09333333 | -0.09 | -0.38 | 0.832687839 | 0.610244164 | 0.144776513 | 0.999652205 | 0.995559182 | 0.997605693 | 0.706643254 |
| 2810008M | Low | Q3UTD9 | Small integral membrane protein 15 OS=Mus musculus OX=10090 GN=Smim15 PE=1 SV=1 | 0.087 | 1.173 | 9 | 1 | 1 | 1 | 74 | 8.6 | 9.82 | 0.989 | 1.15 | 0.86 | -0.02 | 0.02 | 0.2 | 0.06666667 | -0.09 | -0.22 | 0.905231797 | 0.739374982 | 0.413696484 | 0.999652205 | 0.995559182 | 0.997605693 | 0.936397673 |
| Chid1 | Medium | Q922Q9 | Chitinase domain-containing protein 1 OS=Mus musculus OX=10090 GN=Chid1 PE=1 SV=1 | 0.036 | 1.762 | 2 | 1 | 1 | 1 | 393 | 44.9 | 7.97 | 0.911 | 1.248 | 0.73 | -0.13 | 0.13 | 0.32 | 0.10666667 | -0.095 | -0.45 | 0.661836215 | 0.546708459 | 0.050794913 | 0.999652205 | 0.995559182 | 0.997605693 | 0.427566646 |
| 2610034M | High | Q8BGR1 | PDZ and pleckstrin homology domains 1 OS=Mus musculus OX=10090 GN=Pdzph1 PE=2 SV=1 | 0 | 8.387 | 3 | 3 | 3 | 3 | 1238 | 138.5 | 6.76 | 0.988 | 1.158 | 0.853 | -0.02 | 0.02 | 0.21 | 0.07 | -0.095 | -0.23 | 0.88451671 | 0.715734944 | 0.35075502 | 0.999652205 | 0.995559182 | 0.997605693 | 0.909466648 |
| Prdx6 | High | Q6GT24 | Peroxiredoxin-6 OS=Mus musculus OX=10090 GN=Prdx6 PE=1 SV=1 | 0 | 100.615 | 77 | 17 | 32 | 17 | 224 | 24.8 | 6.37 | 0.992 | 1.149 | 0.863 | -0.01 | 0.01 | 0.2 | 0.06666667 | -0.095 | -0.21 | 0.981076023 | 0.733259114 | 0.396574835 | 0.999652205 | 0.995559182 | 0.997605693 | 0.93262326 |
| Ddah2 | High | Q99LD8 | N(G),N(G)-dimethylarginine dimethylaminohydrolase 2 OS=Mus musculus OX=10090 GN=Ddah2 | 0 | 73.354 | 72 | 14 | 22 | 13 | 285 | 29.6 | 6.01 | 1.004 | 1.132 | 0.887 | 0.01 | -0.01 | 0.18 | 0.06 | -0.095 | -0.17 | 0.987760863 | 0.768878355 | 0.500127986 | 0.999652205 | 0.995559182 | 0.997605693 | 0.964378027 |
| Ppm1l | Medium | Q8BHN0 | Protein phosphatase 1L OS=Mus musculus OX=10090 GN=Ppm1l PE=1 SV=1 | 0.01 | 2.721 | 3 | 1 | 1 | 1 | 360 | 41 | 5.99 | 1.01 | 1.136 | 0.889 | 0.01 | -0.01 | 0.18 | 0.06 | -0.095 | -0.17 | 0.962342659 | 0.768865855 | 0.531696026 | 0.999652205 | 0.995559182 | 0.997605693 | 0.96694628 |
| B3galtl; B3 | Medium | Q8BHT6 | Beta-1,3-glucosyltransferase OS=Mus musculus OX=10090 GN=B3glct PE=1 SV=3 | 0.042 | 1.628 | 2 | 1 | 1 | 1 | 489 | 55.3 | 6.92 | 1.011 | 1.136 | 0.89 | 0.02 | -0.02 | 0.18 | 0.06 | -0.1 | -0.17 | 0.968654006 | 0.762509969 | 0.511344491 | 0.999652205 | 0.995559182 | 0.997605693 | 0.964378027 |
| Ncald | High | Q91X97 | Neurocalcin-delta OS=Mus musculus OX=10090 GN=Ncald PE=1 SV=4 | 0 | 14.079 | 20 | 5 | 9 | 2 | 193 | 22.2 | 5.35 | 0.943 | 1.211 | 0.779 | -0.08 | 0.08 | 0.28 | 0.09333333 | -0.1 | -0.36 | 0.856967961 | 0.616745439 | 0.165014954 | 0.999652205 | 0.995559182 | 0.997605693 | 0.736602876 |
| Fermt2 | High | Q8CIB5 | Fermitin family homolog 2 OS=Mus musculus OX=10090 GN=Fermt2 PE=1 SV=1 | 0 | 14.926 | 9 | 6 | 6 | 6 | 680 | 77.8 | 6.7 | 1.006 | 1.144 | 0.879 | 0.01 | -0.01 | 0.19 | 0.06333333 | -0.1 | -0.19 | 0.998891218 | 0.744410714 | 0.479025071 | 0.999652205 | 0.995559182 | 0.997605693 | 0.964378027 |
| L3hypdh | High | Q9CXA2 | Trans-L-3-hydroxyproline dehydratase OS=Mus musculus OX=10090 GN=L3hypdh PE=1 SV=1 | 0.008 | 3.022 | 5 | 1 | 1 | 1 | 354 | 37.8 | 6.7 | 1.233 | 0.942 | 1.309 | 0.3 | -0.3 | -0.09 | -0.03 | -0.105 | 0.39 | 0.456493137 | 0.772121217 | 0.129216311 | 0.999652205 | 0.995559182 | 0.997605693 | 0.683250784 |
| Rps26; Rp: | High | P62855 | 40S ribosomal protein S26 OS=Mus musculus OX=10090 GN=Rps26 PE=1 SV=3 | 0.003 | 3.952 | 19 | 2 | 3 | 2 | 115 | 13 | 11 | 0.998 | 1.156 | 0.863 | 0 | 0 | 0.21 | 0.07 | -0.105 | -0.21 | 0.919223871 | 0.719448407 | 0.390796378 | 0.999652205 | 0.995559182 | 0.997605693 | 0.93064581 |
| | Low | A0A087W( | Regulator of G-protein-signaling 6 OS=Mus musculus OX=10090 GN=Rgs6 PE=1 SV=2 | 0.072 | 1.27 | 1 | 1 | 1 | 1 | 465 | 53.8 | 7.14 | 0.989 | 1.176 | 0.841 | -0.02 | 0.02 | 0.23 | 0.07666667 | -0.105 | -0.25 | 0.87798327 | 0.680943518 | 0.328715286 | 0.999652205 | 0.995559182 | 0.997605693 | 0.905686588 |
| Gad1 | High | P48318 | Glutamate decarboxylase 1 OS=Mus musculus OX=10090 GN=Gad1 PE=1 SV=2 | 0 | 37.038 | 14 | 8 | 10 | 7 | 593 | 66.6 | 7.17 | 0.946 | 1.229 | 0.769 | -0.08 | 0.08 | 0.3 | 0.1 | -0.11 | -0.38 | 0.81467365 | 0.584094902 | 0.124077055 | 0.999652205 | 0.995559182 | 0.997605693 | 0.677192798 |
| Txndc12 | High | Q9CQU0 | Thioredoxin domain-containing protein 12 OS=Mus musculus OX=10090 GN=Txndc12 PE=1 SV=1 | 0 | 12.667 | 24 | 3 | 3 | 3 | 170 | 19 | 5.26 | 0.945 | 1.229 | 0.769 | -0.08 | 0.08 | 0.3 | 0.1 | -0.11 | -0.38 | 0.891965962 | 0.584987931 | 0.144335916 | 0.999652205 | 0.995559182 | 0.997605693 | 0.706643254 |
| Asna1 | High | O54984 | ATPase GET3 OS=Mus musculus OX=10090 GN=Get3 PE=1 SV=2 | 0 | 12.221 | 16 | 4 | 4 | 4 | 348 | 38.8 | 4.91 | 0.975 | 1.198 | 0.814 | -0.04 | 0.04 | 0.26 | 0.08666667 | -0.11 | -0.3 | 0.846153828 | 0.640310939 | 0.24329481 | 0.999652205 | 0.995559182 | 0.997605693 | 0.837310213 |
| Slc4a10 | High | Q5DTL9 | Sodium-driven chloride bicarbonate exchanger OS=Mus musculus OX=10090 GN=Slc4a10 PE=1 SV | 0 | 28.44 | 8 | 7 | 9 | 5 | 1118 | 125.7 | 6.51 | 0.984 | 1.178 | 0.835 | -0.02 | 0.02 | 0.24 | 0.08 | -0.11 | -0.26 | 0.867904466 | 0.677013612 | 0.252444531 | 0.999652205 | 0.995559182 | 0.997605693 | 0.851275262 |
| Gyk; Gk | High | Q64516 | Glycerol kinase OS=Mus musculus OX=10090 GN=Gk PE=1 SV=2 | 0 | 13.701 | 8 | 4 | 5 | 4 | 559 | 61.2 | 5.87 | 0.997 | 1.162 | 0.859 | 0 | 0 | 0.22 | 0.07333333 | -0.11 | -0.22 | 0.91521816 | 0.70880026 | 0.397839274 | 0.999652205 | 0.995559182 | 0.997605693 | 0.93262326 |
| Acot7 | High | E9PYH2 | Cytosolic acyl coenzyme A thioester hydrolase OS=Mus musculus OX=10090 GN=Acot7 PE=1 SV=1 | 0 | 10.863 | 10 | 4 | 4 | 4 | 384 | 42.8 | 8.68 | 1.018 | 1.141 | 0.892 | 0.03 | -0.03 | 0.19 | 0.06333333 | -0.11 | -0.17 | 0.973261046 | 0.75609927 | 0.507023452 | 0.999652205 | 0.995559182 | 0.997605693 | 0.964378027 |
| Aldh3a2 | High | B1AV77 | Aldehyde dehydrogenase OS=Mus musculus OX=10090 GN=Aldh3a2 PE=1 SV=1 | 0 | 16.993 | 10 | 4 | 6 | 4 | 487 | 54.3 | 8.48 | 1.027 | 1.133 | 0.907 | 0.04 | -0.04 | 0.18 | 0.06 | -0.11 | -0.14 | 0.911965 | 0.767239985 | 0.588610145 | 0.999652205 | 0.995559182 | 0.997605693 | 0.973095093 |
| Clint1 | High | Q5SUH7 | Clathrin interactor 1 OS=Mus musculus OX=10090 GN=Clint1 PE=1 SV=1 | 0.001 | 7.629 | 4 | 2 | 2 | 2 | 623 | 67.7 | 6.42 | 0.951 | 1.228 | 0.775 | -0.07 | 0.07 | 0.3 | 0.1 | -0.115 | -0.37 | 0.795622825 | 0.590881957 | 0.15813078 | 0.999652205 | 0.995559182 | 0.997605693 | 0.733365826 |
| Pds5b | High | F8WHU5 | Sister chromatid cohesion protein PDS5 homolog B OS=Mus musculus OX=10090 GN=Pds5b PE=1 | 0.001 | 4.638 | 1 | 1 | 1 | 1 | 1449 | 164.7 | 8.54 | 0.991 | 1.181 | 0.839 | -0.01 | 0.01 | 0.24 | 0.08 | -0.115 | -0.25 | 0.898634394 | 0.658918071 | 0.326357562 | 0.999652205 | 0.995559182 | 0.997605693 | 0.905686588 |
| Ppm1h | High | Q3UYC0 | Protein phosphatase 1H OS=Mus musculus OX=10090 GN=Ppm1h PE=1 SV=1 | 0 | 13.526 | 8 | 3 | 3 | 3 | 513 | 56.3 | 6.73 | 1.032 | 1.137 | 0.908 | 0.05 | -0.05 | 0.18 | 0.06 | -0.115 | -0.14 | 0.955909172 | 0.772548243 | 0.629353051 | 0.999652205 | 0.995559182 | 0.997605693 | 0.981916182 |
| Ndufaf2 | High | Q59J78 | NADH dehydrogenase [ubiquinone] 1 alpha subcomplex assembly factor 2 OS=Mus musculus OX= | 0 | 15.348 | 33 | 5 | 5 | 5 | 168 | 19.6 | 8.25 | 0.939 | 1.248 | 0.753 | -0.09 | 0.09 | 0.32 | 0.10666667 | -0.115 | -0.41 | 0.775905218 | 0.552700024 | 0.088592717 | 0.999652205 | 0.995559182 | 0.997605693 | 0.57426516 |
| Atp1b1 | High | P14094 | Sodium/potassium-transporting ATPase subunit beta-1 OS=Mus musculus OX=10090 GN=Atp1b1 | 0 | 168.578 | 50 | 21 | 78 | 21 | 304 | 35.2 | 8.65 | 1.004 | 1.164 | 0.862 | 0.01 | -0.01 | 0.22 | 0.07333333 | -0.115 | -0.21 | 0.988023234 | 0.703282573 | 0.390988507 | 0.999652205 | 0.995559182 | 0.997605693 | 0.93064581 |
| Flad1 | High | Q8R123 | FAD synthase OS=Mus musculus OX=10090 GN=Flad1 PE=1 SV=1 | 0.001 | 4.363 | 2 | 1 | 1 | 1 | 492 | 54.7 | 6.64 | 1.005 | 1.162 | 0.865 | 0.01 | -0.01 | 0.22 | 0.07333333 | -0.115 | -0.21 | 0.98220929 | 0.696946329 | 0.415846646 | 0.999652205 | 0.995559182 | 0.997605693 | 0.936397673 |
| Rrh | High | Q80XL3 | Rrh protein OS=Mus musculus OX=10090 GN=Rrh PE=2 SV=1 | 0 | 14.537 | 10 | 3 | 3 | 3 | 379 | 41.8 | 8.65 | 1.264 | 0.924 | 1.368 | 0.34 | -0.34 | -0.11 | -0.0366667 | -0.115 | 0.45 | 0.459475851 | 0.742322084 | 0.072422134 | 0.999652205 | 0.995559182 | 0.997605693 | 0.512069238 |
| Clta | High | O08585 | Clathrin light chain A OS=Mus musculus OX=10090 GN=Clta PE=1 SV=2 | 0 | 20.394 | 21 | 8 | 15 | 1 | 235 | 25.6 | 4.58 | 0.968 | 1.22 | 0.794 | -0.05 | 0.05 | 0.29 | 0.09666667 | -0.12 | -0.33 | 0.870210644 | 0.600300629 | 0.142162083 | 0.999652205 | 0.995559182 | 0.997605693 | 0.701851981 |
| Kdsr | High | Q6GV12 | 3-ketodihydrosphingosine reductase OS=Mus musculus OX=10090 GN=Kdsr PE=1 SV=1 | 0.001 | 5.39 | 5 | 1 | 1 | 1 | 332 | 35.9 | 7.9 | 0.998 | 1.182 | 0.844 | 0 | 0 | 0.24 | 0.08 | -0.12 | -0.24 | 0.882823121 | 0.673024961 | 0.308215694 | 0.999652205 | 0.995559182 | 0.997605693 | 0.904943958 |
| Itgav | High | P43406 | Integrin alpha-V OS=Mus musculus OX=10090 GN=Itgav PE=1 SV=2 | 0 | 265.735 | 46 | 41 | 78 | 41 | 1044 | 115.3 | 5.63 | 1.014 | 1.164 | 0.871 | 0.02 | -0.02 | 0.22 | 0.07333333 | -0.12 | -0.2 | 0.96016263 | 0.703527267 | 0.429402956 | 0.999652205 | 0.995559182 | 0.997605693 | 0.944356224 |
| Uap1l1 | High | Q3TW96 | UDP-N-acetylhexosamine pyrophosphorylase-like protein 1 OS=Mus musculus OX=10090 GN=Uap | 0.001 | 5.035 | 5 | 3 | 3 | 3 | 507 | 56.6 | 5.43 | 1.014 | 1.161 | 0.873 | 0.02 | -0.02 | 0.22 | 0.07333333 | -0.12 | -0.2 | 0.961918647 | 0.709400769 | 0.431026934 | 0.999652205 | 0.995559182 | 0.997605693 | 0.945905156 |
| Atox1 | High | O08997 | Copper transport protein ATOX1 OS=Mus musculus OX=10090 GN=Atox1 PE=1 SV=1 | 0 | 24.221 | 59 | 4 | 7 | 4 | 68 | 7.3 | 6.51 | 1.037 | 1.137 | 0.912 | 0.05 | -0.05 | 0.19 | 0.06333333 | -0.12 | -0.13 | 0.943004361 | 0.757979441 | 0.557954495 | 0.999652205 | 0.995559182 | 0.997605693 | 0.971568588 |
| Opcml | High | Q6DFY2 | Opioid-binding protein/cell adhesion molecule-like OS=Mus musculus OX=10090 GN=Opcml PE=1 | 0 | 17.519 | 10 | 4 | 11 | 1 | 337 | 37.1 | 6.68 | 1.035 | 1.139 | 0.909 | 0.05 | -0.05 | 0.19 | 0.06333333 | -0.12 | -0.14 | 0.931779012 | 0.753050206 | 0.616422845 | 0.999652205 | 0.995559182 | 0.997605693 | 0.97826833 |
| Dhx29 | Medium | Q6PGC1 | ATP-dependent RNA helicase DHX29 OS=Mus musculus OX=10090 GN=Dhx29 PE=1 SV=1 | 0.018 | 2.103 | 1 | 1 | 1 | 1 | 1365 | 153.9 | 7.94 | 1.046 | 1.136 | 0.921 | 0.06 | -0.06 | 0.18 | 0.06 | -0.12 | -0.12 | 0.910067641 | 0.749886834 | 0.639991393 | 0.999652205 | 0.995559182 | 0.997605693 | 0.983766261 |
| Eef1a2 | High | P62631 | Elongation factor 1-alpha 2 OS=Mus musculus OX=10090 GN=Eef1a2 PE=1 SV=1 | 0 | 119.528 | 37 | 20 | 78 | 6 | 463 | 50.4 | 9.03 | 0.943 | 1.256 | 0.751 | -0.09 | 0.09 | 0.33 | 0.11 | -0.12 | -0.41 | 0.86523069 | 0.54041813 | 0.080556766 | 0.999652205 | 0.995559182 | 0.997605693 | 0.546169029 |
| Ankrd12 | Low | G5E893 | Ankyrin repeat domain 12 OS=Mus musculus OX=10090 GN=Ankrd12 PE=1 SV=1 | 0.054 | 1.396 | 1 | 1 | 1 | 1 | 2041 | 232.8 | 7.02 | 1.025 | 1.148 | 0.893 | 0.04 | -0.04 | 0.2 | 0.06666667 | -0.12 | -0.16 | 0.962229691 | 0.727826616 | 0.544195356 | 0.999652205 | 0.995559182 | 0.997605693 | 0.96694628 |
| Epb4.1l2; I | High | O70318 | Band 4.1-like protein 2 OS=Mus musculus OX=10090 GN=Epb41l2 PE=1 SV=2 | 0 | 226.512 | 39 | 38 | 77 | 2 | 988 | 109.9 | 5.43 | 0.963 | 1.234 | 0.78 | -0.05 | 0.05 | 0.3 | 0.1 | -0.125 | -0.36 | 0.90128538 | 0.576340237 | 0.13417547 | 0.999652205 | 0.995559182 | 0.997605693 | 0.697570122 |
| Hsd17b11 | Medium | Q9EQ06 | Estradiol 17-beta-dehydrogenase 11 OS=Mus musculus OX=10090 GN=Hsd17b11 PE=1 SV=1 | 0.01 | 2.536 | 4 | 1 | 1 | 1 | 298 | 32.9 | 8.66 | 1.011 | 1.169 | 0.865 | 0.02 | -0.02 | 0.23 | 0.07666667 | -0.125 | -0.21 | 0.99922417 | 0.682536263 | 0.455759961 | 0.999652205 | 0.995559182 | 0.997605693 | 0.959744396 |
| | Medium | A0A0N4SV | [F-actin]-monooxygenase MICAL3 (Fragment) OS=Mus musculus OX=10090 GN=Mical3 PE=1 SV= | 0.011 | 2.413 | 1 | 1 | 1 | 1 | 695 | 78.9 | 7.72 | 1.029 | 1.16 | 0.888 | 0.04 | -0.04 | 0.21 | 0.07 | -0.125 | -0.17 | 0.960968097 | 0.703358757 | 0.524822102 | 0.999652205 | 0.995559182 | 0.997605693 | 0.964378027 |
| Sord | High | Q64442 | Sorbitol dehydrogenase OS=Mus musculus OX=10090 GN=Sord PE=1 SV=3 | 0 | 17.972 | 14 | 5 | 7 | 5 | 357 | 38.2 | 7.02 | 1.038 | 1.147 | 0.905 | 0.05 | -0.05 | 0.2 | 0.06666667 | -0.125 | -0.14 | 0.935683948 | 0.737654139 | 0.52485339 | 0.999652205 | 0.995559182 | 0.997605693 | 0.964378027 |
| Fam54b; M | High | Q9CWE0 | Mitochondrial fission regulator 1-like OS=Mus musculus OX=10090 GN=Mtfr1l PE=1 SV=1 | 0.003 | 4.068 | 7 | 2 | 2 | 2 | 289 | 31.7 | 6.1 | 1.027 | 1.158 | 0.887 | 0.04 | -0.04 | 0.21 | 0.07 | -0.125 | -0.17 | 0.997229358 | 0.726252707 | 0.537089734 | 0.999652205 | 0.995559182 | 0.997605693 | 0.96694628 |
| Dclk1 | High | Q9JLM8 | Serine/threonine-protein kinase DCLK1 OS=Mus musculus OX=10090 GN=Dclk1 PE=1 SV=1 | 0 | 26.164 | 10 | 5 | 6 | 5 | 756 | 84.1 | 8.87 | 1.038 | 1.149 | 0.903 | 0.05 | -0.05 | 0.2 | 0.06666667 | -0.125 | -0.15 | 0.891776808 | 0.732931684 | 0.548673386 | 0.999652205 | 0.995559182 | 0.997605693 | 0.96694628 |
| Iqgap2 | High | Q3UQ44 | Ras GTPase-activating-like protein IQGAP2 OS=Mus musculus OX=10090 GN=Iqgap2 PE=1 SV=2 | 0 | 99.102 | 17 | 23 | 31 | 20 | 1575 | 180.4 | 5.64 | 1.046 | 1.136 | 0.921 | 0.07 | -0.07 | 0.18 | 0.06 | -0.125 | -0.12 | 0.876855481 | 0.760295188 | 0.666823498 | 0.999652205 | 0.995559182 | 0.997605693 | 0.987489386 |
| Comt | High | O88587 | Catechol O-methyltransferase OS=Mus musculus OX=10090 GN=Comt PE=1 SV=2 | 0 | 9.26 | 11 | 2 | 2 | 2 | 265 | 29.5 | 5.83 | 0.977 | 1.227 | 0.796 | -0.03 | 0.03 | 0.29 | 0.09666667 | -0.13 | -0.33 | 0.873053765 | 0.591071697 | 0.172634582 | 0.999652205 | 0.995559182 | 0.997605693 | 0.738844686 |
| Arfip2 | High | Q8K221 | Arfaptin-2 OS=Mus musculus OX=10090 GN=Arfip2 PE=1 SV=2 | 0 | 15.082 | 13 | 4 | 4 | 4 | 341 | 37.7 | 5.87 | 0.999 | 1.202 | 0.832 | 0 | 0 | 0.26 | 0.08666667 | -0.13 | -0.27 | 0.958180454 | 0.633031803 | 0.275308734 | 0.999652205 | 0.995559182 | 0.997605693 | 0.876067403 |
| Adam9 | High | A0A140LH | Disintegrin and metalloproteinase domain-containing protein 9 OS=Mus musculus OX=10090 GN= | 0 | 80.23 | 21 | 15 | 25 | 15 | 863 | 93.6 | 7.62 | 1.017 | 1.185 | 0.858 | 0.02 | -0.02 | 0.24 | 0.08 | -0.13 | -0.22 | 0.913568652 | 0.66442206 | 0.376617438 | 0.999652205 | 0.995559182 | 0.997605693 | 0.920529196 |
| Clic1 | High | Q9Z1Q5 | Chloride intracellular channel protein 1 OS=Mus musculus OX=10090 GN=Clic1 PE=1 SV=3 | 0 | 54.161 | 41 | 8 | 13 | 8 | 241 | 27 | 5.17 | 1.026 | 1.167 | 0.88 | 0.04 | -0.04 | 0.22 | 0.07333333 | -0.13 | -0.18 | 0.868682584 | 0.698959604 | 0.467923555 | 0.999652205 | 0.995559182 | 0.997605693 | 0.961042994 |
| Rtn1 | High | Q8K0T0 | Reticulon-1 OS=Mus musculus OX=10090 GN=Rtn1 PE=1 SV=1 | 0 | 63.115 | 17 | 11 | 21 | 7 | 780 | 83.5 | 4.58 | 1.029 | 1.164 | 0.884 | 0.04 | -0.04 | 0.22 | 0.07333333 | -0.13 | -0.18 | 0.920111391 | 0.703891207 | 0.488462337 | 0.999652205 | 0.995559182 | 0.997605693 | 0.964378027 |
| Exoc1 | High | Q6P1Y9 | Exocyst complex component 1 OS=Mus musculus OX=10090 GN=Exoc1 PE=1 SV=1 | 0.001 | 5.865 | 2 | 2 | 3 | 2 | 901 | 102.6 | 6.47 | 1.025 | 1.162 | 0.882 | 0.04 | -0.04 | 0.22 | 0.07333333 | -0.13 | -0.18 | 0.97964667 | 0.716801531 | 0.517379671 | 0.999652205 | 0.995559182 | 0.997605693 | 0.964378027 |
| Tspan13 | High | Q9D8C2 | Tetraspanin-13 OS=Mus musculus OX=10090 GN=Tspan13 PE=1 SV=1 | 0.006 | 3.078 | 11 | 2 | 2 | 2 | 204 | 22.2 | 8.37 | 1.046 | 1.142 | 0.916 | 0.07 | -0.07 | 0.19 | 0.06333333 | -0.13 | -0.13 | 0.916383156 | 0.761340777 | 0.639976814 | 0.999652205 | 0.995559182 | 0.997605693 | 0.983766261 |
| Naca | High | P70670 | Nascent polypeptide-associated complex subunit alpha, muscle-specific form OS=Mus musculus ( | 0 | 16.667 | 2 | 3 | 5 | 3 | 2187 | 220.4 | 9.35 | 1.058 | 1.135 | 0.932 | 0.08 | -0.08 | 0.18 | 0.06 | -0.13 | -0.1 | 0.826368671 | 0.76304177 | 0.711905361 | 0.999652205 | 0.995559182 | 0.997605693 | 0.992535481 |
| Shank2 | High | Q80Z38 | SH3 and multiple ankyrin repeat domains protein 2 OS=Mus musculus OX=10090 GN=Shank2 PE= | 0 | 18.537 | 4 | 4 | 4 | 4 | 1476 | 158.9 | 7.18 | 1.058 | 1.132 | 0.935 | 0.08 | -0.08 | 0.18 | 0.06 | -0.13 | -0.1 | 0.850498245 | 0.769233023 | 0.726593087 | 0.999652205 | 0.995559182 | 0.997605693 | 0.992535481 |
| Cbr4 | Medium | Q91VT4 | 3-oxoacyl-[acyl-carrier-protein] reductase OS=Mus musculus OX=10090 GN=Cbr4 PE=1 SV=2 | 0.014 | 2.238 | 4 | 1 | 1 | 1 | 236 | 25.4 | 9.69 | 0.991 | 1.212 | 0.817 | -0.01 | 0.01 | 0.28 | 0.09333333 | -0.135 | -0.29 | 0.959248098 | 0.60356207 | 0.258148496 | 0.999652205 | 0.995559182 | 0.997605693 | 0.85616924 |
| Pon2 | High | Q62086 | Serum paraoxonase/arylesterase 2 OS=Mus musculus OX=10090 GN=Pon2 PE=1 SV=2 | 0 | 8.262 | 6 | 2 | 3 | 2 | 354 | 39.6 | 5.83 | 1.031 | 1.17 | 0.881 | 0.04 | -0.04 | 0.23 | 0.07666667 | -0.135 | -0.18 | 0.896012245 | 0.691952726 | 0.464930164 | 0.999652205 | 0.995559182 | 0.997605693 | 0.960359242 |
| Atp6v0c; A | High | P63082 | V-type proton ATPase 16 kDa proteolipid subunit OS=Mus musculus OX=10090 GN=Atp6v0c PE=1 | 0 | 12.611 | 16 | 2 | 6 | 2 | 155 | 15.8 | 9.13 | 1.036 | 1.164 | 0.89 | 0.05 | -0.05 | 0.22 | 0.07333333 | -0.135 | -0.17 | 0.923443709 | 0.703758077 | 0.518623838 | 0.999652205 | 0.995559182 | 0.997605693 | 0.964378027 |
| Ogfrl1 | High | Q8VE52 | Opioid growth factor receptor-like protein 1 OS=Mus musculus OX=10090 GN=Ogfrl1 PE=1 SV=2 | 0.001 | 5.817 | 4 | 2 | 2 | 2 | 464 | 52.2 | 6.57 | 1.043 | 1.16 | 0.899 | 0.06 | -0.06 | 0.21 | 0.07 | -0.135 | -0.15 | 0.922991484 | 0.700345 | 0.544708212 | 0.999652205 | 0.995559182 | 0.997605693 | 0.96694628 |
| Fxyd6 | High | Q9D164 | FXYD domain-containing ion transport regulator 6 OS=Mus musculus OX=10090 GN=Fxyd6 PE=1 S | 0 | 11.441 | 35 | 3 | 3 | 3 | 94 | 10.4 | 5.11 | 1.056 | 1.137 | 0.928 | 0.08 | -0.08 | 0.19 | 0.06333333 | -0.135 | -0.11 | 0.906601424 | 0.757123801 | 0.679001534 | 0.999652205 | 0.995559182 | 0.997605693 | 0.99119021 |
| Atp8a2 | High | P98200 | Phospholipid-transporting ATPase IB OS=Mus musculus OX=10090 GN=Atp8a2 PE=1 SV=1 | 0 | 338.134 | 42 | 46 | 166 | 38 | 1148 | 129.3 | 7.33 | 1.066 | 1.134 | 0.94 | 0.09 | -0.09 | 0.18 | 0.06 | -0.135 | -0.09 | 0.828175403 | 0.765025248 | 0.765277509 | 0.999652205 | 0.995559182 | 0.997605693 | 0.992535481 |
| Ncln | Medium | Q8VCM8 | Nicalin OS=Mus musculus OX=10090 GN=Ncln PE=1 SV=2 | 0.042 | 1.644 | 2 | 1 | 1 | 1 | 563 | 62.9 | 6.49 | 0.944 | 1.28 | 0.737 | -0.08 | 0.08 | 0.36 | 0.12 | -0.14 | -0.44 | 0.749554893 | 0.499382575 | 0.135937064 | 0.999652205 | 0.995559182 | 0.997605693 | 0.699265243 |
| | High | H3BJ02 | Syntaxin 5A OS=Mus musculus OX=10090 GN=Stx5a PE=1 SV=1 | 0.001 | 6.382 | 7 | 1 | 1 | 1 | 259 | 29.4 | 7.56 | 1.027 | 1.182 | 0.868 | 0.04 | -0.04 | 0.24 | 0.08 | -0.14 | -0.2 | 0.971993321 | 0.672463056 | 0.389725779 | 0.999652205 | 0.995559182 | 0.997605693 | 0.93064581 |
| Arcn1 | High | Q5XJY5 | Coatomer subunit delta OS=Mus musculus OX=10090 GN=Arcn1 PE=1 SV=2 | 0 | 11.348 | 10 | 4 | 5 | 4 | 511 | 57.2 | 6.21 | 1.027 | 1.183 | 0.868 | 0.04 | -0.04 | 0.24 | 0.08 | -0.14 | -0.2 | 0.88540136 | 0.667050251 | 0.447561789 | 0.999652205 | 0.995559182 | 0.997605693 | 0.958608248 |
| Maoa | High | Q64133 | Amine oxidase [flavin-containing] A OS=Mus musculus OX=10090 GN=Maoa PE=1 SV=3 | 0.001 | 5.583 | 4 | 2 | 3 | 2 | 526 | 59.6 | 7.81 | 0.976 | 1.237 | 0.789 | -0.03 | 0.03 | 0.31 | 0.10333333 | -0.14 | -0.34 | 0.983453811 | 0.571485893 | 0.174404984 | 0.999652205 | 0.995559182 | 0.997605693 | 0.744450291 |
| Srpk2 | High | A0A0R4J1 | SRSF protein kinase 2 OS=Mus musculus OX=10090 GN=Srpk2 PE=1 SV=1 | 0.008 | 2.978 | 1 | 1 | 1 | 1 | 682 | 76.8 | 4.91 | 1.022 | 1.186 | 0.862 | 0.03 | -0.03 | 0.25 | 0.08333333 | -0.14 | -0.21 | 0.987152052 | 0.667525567 | 0.380348841 | 0.999652205 | 0.995559182 | 0.997605693 | 0.923104982 |
| Timm50 | High | Q9D880 | Mitochondrial import inner membrane translocase subunit TIM50 OS=Mus musculus OX=10090 G | 0.001 | 5.874 | 8 | 2 | 2 | 2 | 353 | 39.8 | 8.13 | 1.022 | 1.186 | 0.861 | 0.03 | -0.03 | 0.25 | 0.08333333 | -0.14 | -0.22 | 0.98918931 | 0.667505099 | 0.392124367 | 0.999652205 | 0.995559182 | 0.997605693 | 0.93064581 |
| Gjd2 | Medium | O54851 | Gap junction delta-2 protein OS=Mus musculus OX=10090 GN=Gjd2 PE=1 SV=2 | 0.047 | 1.544 | 2 | 1 | 1 | 1 | 321 | 36.1 | 8.68 | 1.03 | 1.19 | 0.865 | 0.04 | -0.04 | 0.25 | 0.08333333 | -0.145 | -0.21 | 0.940867106 | 0.645652495 | 0.418706326 | 0.999652205 | 0.995559182 | 0.997605693 | 0.936826165 |
| Cdv3 | High | Q4VAA2 | Protein CDV3 OS=Mus musculus OX=10090 GN=Cdv3 PE=1 SV=2 | 0 | 16.72 | 20 | 4 | 5 | 4 | 281 | 29.7 | 6.1 | 1.033 | 1.184 | 0.873 | 0.05 | -0.05 | 0.24 | 0.08 | -0.145 | -0.2 | 0.883667365 | 0.66629534 | 0.462220385 | 0.999652205 | 0.995559182 | 0.997605693 | 0.959744396 |
| Gdap1 | High | O88741 | Ganglioside-induced differentiation-associated protein 1 OS=Mus musculus OX=10090 GN=Gdap1 | 0 | 15.59 | 15 | 5 | 6 | 5 | 358 | 41.3 | 8.37 | 0.978 | 1.248 | 0.784 | -0.03 | 0.03 | 0.32 | 0.10666667 | -0.145 | -0.35 | 0.890407195 | 0.552826737 | 0.124158052 | 0.999652205 | 0.995559182 | 0.997605693 | 0.677192798 |
| Acsl4 | High | Q9QUJ7 | Long-chain-fatty-acid--CoA ligase 4 OS=Mus musculus OX=10090 GN=Acsl4 PE=1 SV=2 | 0.006 | 3.28 | 1 | 1 | 1 | 1 | 711 | 79 | 8.28 | 1.004 | 1.212 | 0.828 | 0.01 | -0.01 | 0.28 | 0.09333333 | -0.145 | -0.27 | 0.956177468 | 0.616753066 | 0.260194825 | 0.999652205 | 0.995559182 | 0.997605693 | 0.85616924 |
| Cadps | High | Q80TJ1 | Calcium-dependent secretion activator 1 OS=Mus musculus OX=10090 GN=Cadps PE=1 SV=3 | 0 | 104.193 | 23 | 22 | 29 | 22 | 1355 | 153 | 5.74 | 1.01 | 1.218 | 0.829 | 0.01 | -0.01 | 0.28 | 0.09333333 | -0.145 | -0.27 | 0.970013108 | 0.603152341 | 0.268469213 | 0.999652205 | 0.995559182 | 0.997605693 | 0.869723277 |
| Gapdhs | High | Q64467 | Glyceraldehyde-3-phosphate dehydrogenase, testis-specific OS=Mus musculus OX=10090 GN=Ga | 0 | 33.85 | 10 | 4 | 32 | 2 | 440 | 47.6 | 7.88 | 1.067 | 1.153 | 0.925 | 0.09 | -0.09 | 0.2 | 0.06666667 | -0.145 | -0.11 | 0.85592244 | 0.736358729 | 0.679978475 | 0.999652205 | 0.995559182 | 0.997605693 | 0.991285312 |
| Gng4 | High | P50153 | Guanine nucleotide-binding protein G(I)/G(S)/G(O) subunit gamma-4 OS=Mus musculus OX=100( | 0 | 10.527 | 32 | 2 | 4 | 2 | 75 | 8.4 | 7.08 | 1.031 | 1.2 | 0.859 | 0.04 | -0.04 | 0.26 | 0.08666667 | -0.15 | -0.22 | 0.988251654 | 0.63512162 | 0.335265023 | 0.999652205 | 0.995559182 | 0.997605693 | 0.906329766 |
| Itgb8 | High | Q0VBD0 | Integrin beta-8 OS=Mus musculus OX=10090 GN=Itgb8 PE=1 SV=1 | 0 | 157.247 | 36 | 27 | 62 | 27 | 767 | 84.5 | 7.09 | 1.03 | 1.197 | 0.86 | 0.04 | -0.04 | 0.26 | 0.08666667 | -0.15 | -0.22 | 0.918226677 | 0.640709344 | 0.384168449 | 0.999652205 | 0.995559182 | 0.997605693 | 0.927662573 |
| Manf | High | Q80ZP8 | Armet protein OS=Mus musculus OX=10090 GN=Manf PE=1 SV=1 | 0.001 | 7.552 | 15 | 2 | 2 | 2 | 165 | 19 | 8.13 | 0.974 | 1.266 | 0.769 | -0.04 | 0.04 | 0.34 | 0.11333333 | -0.15 | -0.38 | 0.852303228 | 0.519266411 | 0.16750917 | 0.999652205 | 0.995559182 | 0.997605693 | 0.738806217 |
| Ankrd33b | High | Q3U0L2 | Ankyrin repeat domain-containing protein 33B OS=Mus musculus OX=10090 GN=Ankrd33b PE=1 S | 0 | 272.355 | 65 | 34 | 155 | 34 | 486 | 53.2 | 8.68 | 1.024 | 1.209 | 0.847 | 0.03 | -0.03 | 0.27 | 0.09 | -0.15 | -0.24 | 0.933408256 | 0.619019047 | 0.331547715 | 0.999652205 | 0.995559182 | 0.997605693 | 0.906329766 |
| Cox17 | High | P56394 | Cytochrome c oxidase copper chaperone OS=Mus musculus OX=10090 GN=Cox17 PE=1 SV=2 | 0.001 | 5.509 | 21 | 1 | 1 | 1 | 63 | 6.8 | 7.69 | 1.022 | 1.205 | 0.848 | 0.03 | -0.03 | 0.27 | 0.09 | -0.15 | -0.24 | 0.984232881 | 0.634486523 | 0.371770898 | 0.999652205 | 0.995559182 | 0.997605693 | 0.920164724 |
| Lasp1 | High | Q61792 | LIM and SH3 domain protein 1 OS=Mus musculus OX=10090 GN=Lasp1 PE=1 SV=1 | 0 | 66.854 | 37 | 12 | 23 | 11 | 263 | 30 | 7.05 | 1.074 | 1.149 | 0.935 | 0.1 | -0.1 | 0.2 | 0.06666667 | -0.15 | -0.1 | 0.808078664 | 0.733818636 | 0.737241575 | 0.999652205 | 0.995559182 | 0.997605693 | 0.992535481 |
| Slc27a4 | Low | Q91VE0 | Long-chain fatty acid transport protein 4 OS=Mus musculus OX=10090 GN=Slc27a4 PE=1 SV=1 | 0.087 | 1.171 | 1 | 1 | 1 | 1 | 643 | 72.3 | 8.59 | 1.003 | 1.242 | 0.808 | 0 | 0 | 0.31 | 0.10333333 | -0.155 | -0.31 | 0.94346104 | 0.559028696 | 0.237279661 | 0.999652205 | 0.995559182 | 0.997605693 | 0.828110564 |
| Dnm1 | High | A0A0J9YU | Dynamin GTPase OS=Mus musculus OX=10090 GN=Dnm1 PE=1 SV=1 | 0 | 209.191 | 49 | 42 | 76 | 2 | 864 | 97.2 | 6.87 | 1.028 | 1.206 | 0.853 | 0.04 | -0.04 | 0.27 | 0.09 | -0.155 | -0.23 | 0.989561 | 0.62590716 | 0.350936929 | 0.999652205 | 0.995559182 | 0.997605693 | 0.909466648 |
| Plxna1 | High | P70206 | Plexin-A1 OS=Mus musculus OX=10090 GN=Plxna1 PE=1 SV=1 | 0 | 14.875 | 2 | 4 | 4 | 4 | 1894 | 211 | 6.9 | 1.051 | 1.185 | 0.887 | 0.07 | -0.07 | 0.24 | 0.08 | -0.155 | -0.17 | 0.942717103 | 0.661020317 | 0.487388431 | 0.999652205 | 0.995559182 | 0.997605693 | 0.964378027 |
| Dnajb11 | High | Q99KV1 | DnaJ homolog subfamily B member 11 OS=Mus musculus OX=10090 GN=Dnajb11 PE=1 SV=1 | 0.001 | 7.643 | 12 | 2 | 2 | 2 | 358 | 40.5 | 6.32 | 1.053 | 1.179 | 0.893 | 0.07 | -0.07 | 0.24 | 0.08 | -0.155 | -0.16 | 0.8930753 | 0.680692644 | 0.561654277 | 0.999652205 | 0.995559182 | 0.997605693 | 0.972910028 |
| Bcap29 | High | Q61334 | B-cell receptor-associated protein 29 OS=Mus musculus OX=10090 GN=Bcap29 PE=1 SV=1 | 0.009 | 2.775 | 3 | 1 | 1 | 1 | 240 | 27.9 | 9.72 | 1.052 | 1.178 | 0.893 | 0.07 | -0.07 | 0.24 | 0.08 | -0.155 | -0.16 | 0.876743838 | 0.663860849 | 0.567155146 | 0.999652205 | 0.995559182 | 0.997605693 | 0.972910028 |
| Ank1 | High | G5E8J2 | Ankyrin-1 OS=Mus musculus OX=10090 GN=Ank1 PE=1 SV=1 | 0.001 | 7.194 | 1 | 2 | 4 | 1 | 1848 | 202.4 | 6.65 | 1.082 | 1.151 | 0.94 | 0.11 | -0.11 | 0.2 | 0.06666667 | -0.155 | -0.09 | 0.778953738 | 0.72942838 | 0.736871527 | 0.999652205 | 0.995559182 | 0.997605693 | 0.992535481 |
| Ndufa12 | High | Q7TMF3 | NADH dehydrogenase [ubiquinone] 1 alpha subcomplex subunit 12 OS=Mus musculus OX=10090 ( | 0 | 17.849 | 39 | 5 | 5 | 5 | 145 | 17.1 | 9.36 | 1.096 | 1.135 | 0.966 | 0.13 | -0.13 | 0.18 | 0.06 | -0.155 | -0.05 | 0.782177041 | 0.761394905 | 0.817879006 | 0.999652205 | 0.995559182 | 0.997605693 | 0.992535481 |
| Dcakd | Medium | Q8BHC4 | Dephospho-CoA kinase domain-containing protein OS=Mus musculus OX=10090 GN=Dcakd PE=1 S | 0.033 | 1.84 | 6 | 1 | 1 | 1 | 231 | 26.5 | 9.58 | 1.023 | 1.211 | 0.845 | 0.03 | -0.03 | 0.28 | 0.09333333 | -0.155 | -0.24 | 0.995472861 | 0.622139685 | 0.391530251 | 0.999652205 | 0.995559182 | 0.997605693 | 0.93064581 |
| Sgpp1 | High | Q9JI99 | Sphingosine-1-phosphate phosphatase 1 OS=Mus musculus OX=10090 GN=Sgpp1 PE=1 SV=1 | 0.008 | 2.888 | 4 | 2 | 2 | 2 | 430 | 47.7 | 8 | 1.098 | 1.128 | 0.974 | 0.14 | -0.14 | 0.17 | 0.05666667 | -0.155 | -0.04 | 0.744880858 | 0.767879999 | 0.922200242 | 0.999652205 | 0.995559182 | 0.997605693 | 0.992535481 |
| Ubxn4 | High | A0A0R4J0 | UBX domain-containing protein 4 OS=Mus musculus OX=10090 GN=Ubxn4 PE=1 SV=1 | 0 | 23.076 | 16 | 5 | 7 | 5 | 506 | 56.4 | 6.61 | 1.014 | 1.23 | 0.825 | 0.02 | -0.02 | 0.3 | 0.1 | -0.16 | -0.28 | 0.995791297 | 0.582787103 | 0.226008658 | 0.999652205 | 0.995559182 | 0.997605693 | 0.813917005 |
| | High | E9Q517 | Sodium- and chloride-dependent glycine transporter 1 OS=Mus musculus OX=10090 GN=Slc6a9 P | 0 | 43.035 | 14 | 7 | 17 | 7 | 521 | 58.2 | 7.14 | 1.032 | 1.21 | 0.853 | 0.05 | -0.05 | 0.27 | 0.09 | -0.16 | -0.23 | 0.950187489 | 0.618174813 | 0.333602158 | 0.999652205 | 0.995559182 | 0.997605693 | 0.906329766 |
| Cspg4 | High | Q8VHY0 | Chondroitin sulfate proteoglycan 4 OS=Mus musculus OX=10090 GN=Cspg4 PE=1 SV=3 | 0 | 214.362 | 24 | 37 | 61 | 37 | 2327 | 252.2 | 5.44 | 1.093 | 1.14 | 0.959 | 0.13 | -0.13 | 0.19 | 0.06333333 | -0.16 | -0.06 | 0.762727684 | 0.751750892 | 0.86290786 | 0.999652205 | 0.995559182 | 0.997605693 | 0.992535481 |
| Stk24 | High | Q99KH8 | Serine/threonine-protein kinase 24 OS=Mus musculus OX=10090 GN=Stk24 PE=1 SV=1 | 0 | 14.002 | 10 | 3 | 3 | 1 | 431 | 47.9 | 5.43 | 1.1 | 1.134 | 0.97 | 0.14 | -0.14 | 0.18 | 0.06 | -0.16 | -0.04 | 0.79151812 | 0.771607314 | 0.900592108 | 0.999652205 | 0.995559182 | 0.997605693 | 0.992535481 |
| Asrgl1 | High | Q8C0M9 | Isoaspartyl peptidase/L-asparaginase OS=Mus musculus OX=10090 GN=Asrgl1 PE=1 SV=1 | 0 | 135.333 | 60 | 18 | 47 | 18 | 326 | 33.9 | 7.65 | 1.101 | 1.131 | 0.974 | 0.14 | -0.14 | 0.18 | 0.06 | -0.16 | -0.04 | 0.742670892 | 0.771457115 | 0.945997421 | 0.999652205 | 0.995559182 | 0.997605693 | 0.992535481 |
| Rab13 | High | Q9DD03 | Ras-related protein Rab-13 OS=Mus musculus OX=10090 GN=Rab13 PE=1 SV=1 | 0 | 10.786 | 19 | 4 | 8 | 1 | 202 | 22.8 | 9.5 | 1.02 | 1.235 | 0.826 | 0.03 | -0.03 | 0.3 | 0.1 | -0.165 | -0.28 | 0.988233727 | 0.582811769 | 0.280209791 | 0.999652205 | 0.995559182 | 0.997605693 | 0.8782204 |
| Golgb1 | High | E9QAH1 | Golgi autoantigen, golgin subfamily b, macrogolgin 1 OS=Mus musculus OX=10090 GN=Golgb1 PI | 0 | 16.46 | 1 | 4 | 5 | 4 | 3197 | 365.1 | 5.06 | 0.962 | 1.311 | 0.734 | -0.06 | 0.06 | 0.39 | 0.13 | -0.165 | -0.45 | 0.789509646 | 0.456959375 | 0.068920734 | 0.999652205 | 0.995559182 | 0.997605693 | 0.499920511 |
| Tm9sf2 | High | P58021 | Transmembrane 9 superfamily member 2 OS=Mus musculus OX=10090 GN=Tm9sf2 PE=1 SV=1 | 0 | 48.308 | 17 | 8 | 13 | 8 | 662 | 75.3 | 7.43 | 0.969 | 1.302 | 0.745 | -0.05 | 0.05 | 0.38 | 0.12666667 | -0.165 | -0.43 | 0.878871234 | 0.469776204 | 0.075132373 | 0.999652205 | 0.995559182 | 0.997605693 | 0.520184986 |
| Ric3 | High | D3YTZ9 | Protein RIC-3 OS=Mus musculus OX=10090 GN=Ric3 PE=1 SV=1 | 0.002 | 4.153 | 2 | 1 | 1 | 1 | 366 | 40.2 | 5.05 | 0.996 | 1.268 | 0.786 | -0.01 | 0.01 | 0.34 | 0.11333333 | -0.165 | -0.35 | 0.919608524 | 0.521342222 | 0.149918068 | 0.999652205 | 0.995559182 | 0.997605693 | 0.71722748 |
| Pdia3 | High | P27773 | Protein disulfide-isomerase A3 OS=Mus musculus OX=10090 GN=Pdia3 PE=1 SV=2 | 0 | 122.365 | 47 | 26 | 46 | 26 | 505 | 56.6 | 6.21 | 1.002 | 1.258 | 0.797 | 0 | 0 | 0.33 | 0.11 | -0.165 | -0.33 | 0.99220179 | 0.536725442 | 0.172070962 | 0.999652205 | 0.995559182 | 0.997605693 | 0.738844686 |
| Hnrnpl | High | Q8R081 | Heterogeneous nuclear ribonucleoprotein L OS=Mus musculus OX=10090 GN=Hnrnpl PE=1 SV=2 | 0 | 9.337 | 6 | 3 | 3 | 3 | 586 | 63.9 | 8.1 | 1.074 | 1.174 | 0.915 | 0.1 | -0.1 | 0.23 | 0.07666667 | -0.165 | -0.13 | 0.867952425 | 0.6827172 | 0.618417281 | 0.999652205 | 0.995559182 | 0.997605693 | 0.97826833 |
| Met | High | P16056 | Hepatocyte growth factor receptor OS=Mus musculus OX=10090 GN=Met PE=1 SV=1 | 0 | 26.206 | 7 | 10 | 11 | 9 | 1379 | 153.5 | 7.05 | 1.076 | 1.168 | 0.921 | 0.11 | -0.11 | 0.22 | 0.07333333 | -0.165 | -0.12 | 0.841904026 | 0.696799988 | 0.634901987 | 0.999652205 | 0.995559182 | 0.997605693 | 0.983562471 |
| AI597468; | High | Q9DAM7 | Transmembrane protein 263 OS=Mus musculus OX=10090 GN=Tmem263 PE=1 SV=1 | 0.005 | 3.772 | 9 | 1 | 1 | 1 | 115 | 11.5 | 9.32 | 1.077 | 1.166 | 0.923 | 0.11 | -0.11 | 0.22 | 0.07333333 | -0.165 | -0.12 | 0.81380363 | 0.691158717 | 0.677139726 | 0.999652205 | 0.995559182 | 0.997605693 | 0.990892882 |
| Csad | High | Q9DBE0 | Cysteine sulfinic acid decarboxylase OS=Mus musculus OX=10090 GN=Csad PE=1 SV=1 | 0 | 37.931 | 27 | 12 | 15 | 12 | 493 | 55.1 | 6.61 | 1.103 | 1.14 | 0.968 | 0.14 | -0.14 | 0.19 | 0.06333333 | -0.165 | -0.05 | 0.774671326 | 0.752506045 | 0.852247189 | 0.999652205 | 0.995559182 | 0.997605693 | 0.992535481 |
| Pdia6 | High | Q3TML0 | Protein disulfide-isomerase A6 OS=Mus musculus OX=10090 GN=Pdia6 PE=1 SV=1 | 0 | 32.938 | 28 | 8 | 12 | 8 | 445 | 48.7 | 5.19 | 0.996 | 1.273 | 0.783 | -0.01 | 0.01 | 0.35 | 0.11666667 | -0.17 | -0.35 | 0.944229188 | 0.51283564 | 0.141352334 | 0.999652205 | 0.995559182 | 0.997605693 | 0.701851981 |
| Opn1mw | High | O35599 | Medium-wave-sensitive opsin 1 OS=Mus musculus OX=10090 GN=Opn1mw PE=1 SV=1 | 0 | 60.37 | 20 | 10 | 48 | 9 | 359 | 40.2 | 8.37 | 1.034 | 1.225 | 0.844 | 0.05 | -0.05 | 0.29 | 0.09666667 | -0.17 | -0.24 | 0.908133381 | 0.59210896 | 0.321682883 | 0.999652205 | 0.995559182 | 0.997605693 | 0.905686588 |
| Cyb5r3 | High | Q9DCN2 | NADH-cytochrome b5 reductase 3 OS=Mus musculus OX=10090 GN=Cyb5r3 PE=1 SV=3 | 0 | 10.432 | 11 | 3 | 3 | 3 | 301 | 34.1 | 8.38 | 1.041 | 1.212 | 0.859 | 0.06 | -0.06 | 0.28 | 0.09333333 | -0.17 | -0.22 | 0.925115066 | 0.614862658 | 0.338644701 | 0.999652205 | 0.995559182 | 0.997605693 | 0.907205371 |
| Hs1bp3 | High | Q3TC93 | HCLS1-binding protein 3 OS=Mus musculus OX=10090 GN=Hs1bp3 PE=1 SV=2 | 0.001 | 6.609 | 5 | 1 | 1 | 1 | 395 | 43.7 | 4.98 | 1.06 | 1.199 | 0.884 | 0.08 | -0.08 | 0.26 | 0.08666667 | -0.17 | -0.18 | 0.878614817 | 0.654056724 | 0.576422348 | 0.999652205 | 0.995559182 | 0.997605693 | 0.972910028 |
| Csda; Ybx3 | High | Q9JKB3 | Y-box-binding protein 3 OS=Mus musculus OX=10090 GN=Ybx3 PE=1 SV=2 | 0.001 | 6.585 | 17 | 2 | 2 | 1 | 361 | 38.8 | 9.69 | 1.042 | 1.223 | 0.852 | 0.06 | -0.06 | 0.29 | 0.09666667 | -0.175 | -0.23 | 0.936742343 | 0.580686459 | 0.363477625 | 0.999652205 | 0.995559182 | 0.997605693 | 0.91781256 |

| | | | | | | | | | | | | | | | | | | | | | | | | | | | | |
|---|---|---|---|---|---|---|---|---|---|---|---|---|---|---|---|---|---|---|---|---|---|---|---|---|---|---|---|---|
| Araf | High | P04627 | Serine/threonine-protein kinase A-Raf OS=Mus musculus OX=10090 GN=Araf PE=1 SV=2 | 0 | 13.76 | 7 | 5 | 5 | 2 | 604 | 67.5 | 9.11 | 1.062 | 1.201 | 0.884 | 0.09 | -0.09 | 0.26 | 0.08666667 | -0.175 | -0.18 | 0.817658303 | 0.633945197 | 0.488402049 | 0.999652205 | 0.995559182 | 0.997605693 | 0.964378027 |
| Rdx | High | P26043 | Radixin OS=Mus musculus OX=10090 GN=Rdx PE=1 SV=3 | 0 | 235.672 | 50 | 35 | 101 | 22 | 583 | 68.5 | 6.2 | 1.069 | 1.193 | 0.896 | 0.1 | -0.1 | 0.25 | 0.08333333 | -0.175 | -0.16 | 0.82101159 | 0.648935797 | 0.542222485 | 0.999652205 | 0.995559182 | 0.997605693 | 0.96694628 |
| Sccpdh | High | Q8R127 | Saccharopine dehydrogenase-like oxidoreductase OS=Mus musculus OX=10090 GN=Sccpdh PE=1 S | 0 | 18.925 | 16 | 6 | 6 | 6 | 429 | 47.1 | 8.6 | 1.078 | 1.18 | 0.914 | 0.11 | -0.11 | 0.24 | 0.08 | -0.175 | -0.13 | 0.854591199 | 0.673080109 | 0.576212914 | 0.999652205 | 0.995559182 | 0.997605693 | 0.972910028 |
| | High | E9PV44 | ATP synthase F1 subunit epsilon OS=Mus musculus OX=10090 GN=Atpif1 PE=1 SV=1 | 0 | 9.509 | 35 | 4 | 6 | 4 | 74 | 8.8 | 9.31 | 1.09 | 1.175 | 0.927 | 0.12 | -0.12 | 0.23 | 0.07666667 | -0.175 | -0.11 | 0.786059504 | 0.682126184 | 0.610782733 | 0.999652205 | 0.995559182 | 0.997605693 | 0.97826833 |
| Rcn2 | High | Q8BP92 | Reticulocalbin-2 OS=Mus musculus OX=10090 GN=Rcn2 PE=1 SV=1 | 0 | 48.726 | 33 | 7 | 11 | 7 | 320 | 37.2 | 4.42 | 1.077 | 1.184 | 0.909 | 0.11 | -0.11 | 0.24 | 0.08 | -0.175 | -0.14 | 0.830859964 | 0.664796427 | 0.616443129 | 0.999652205 | 0.995559182 | 0.997605693 | 0.97826833 |
| Plxnb2 | High | B2RXS4 | Plexin-B2 OS=Mus musculus OX=10090 GN=Plxnb2 PE=1 SV=1 | 0 | 166.801 | 26 | 35 | 47 | 35 | 1842 | 206.1 | 5.87 | 1.095 | 1.161 | 0.943 | 0.13 | -0.13 | 0.22 | 0.07333333 | -0.175 | -0.09 | 0.758684166 | 0.709501198 | 0.779032353 | 0.999652205 | 0.995559182 | 0.997605693 | 0.992535481 |
| Lap3 | High | Q9CPY7 | Cytosol aminopeptidase OS=Mus musculus OX=10090 GN=Lap3 PE=1 SV=3 | 0 | 10.232 | 8 | 4 | 4 | 4 | 519 | 56.1 | 7.72 | 1.104 | 1.159 | 0.952 | 0.14 | -0.14 | 0.21 | 0.07 | -0.175 | -0.07 | 0.768250104 | 0.713711052 | 0.803847843 | 0.999652205 | 0.995559182 | 0.997605693 | 0.992535481 |
| | High | F7CZG3 | E3 ubiquitin-protein ligase MYCBP2 (Fragment) OS=Mus musculus OX=10090 GN=Mycbp2 PE=1 S | 0.001 | 4.544 | 2 | 1 | 1 | 1 | 871 | 95.1 | 9 | 1.014 | 1.258 | 0.806 | 0.02 | -0.02 | 0.33 | 0.11 | -0.175 | -0.31 | 0.993898514 | 0.552289446 | 0.303306895 | 0.999652205 | 0.995559182 | 0.997605693 | 0.900241969 |
| Plxnb1 | High | Q8CJH3 | Plexin-B1 OS=Mus musculus OX=10090 GN=Plxnb1 PE=1 SV=2 | 0 | 26.641 | 4 | 8 | 11 | 8 | 2119 | 231.2 | 5.55 | 1.075 | 1.196 | 0.899 | 0.1 | -0.1 | 0.26 | 0.08666667 | -0.18 | -0.15 | 0.837406238 | 0.644011577 | 0.539762913 | 0.999652205 | 0.995559182 | 0.997605693 | 0.96694628 |
| Hnrnpab | High | Q99020 | Heterogeneous nuclear ribonucleoprotein A/B OS=Mus musculus OX=10090 GN=Hnrnpab PE=1 SV | 0 | 7.949 | 9 | 2 | 2 | 2 | 285 | 30.8 | 7.91 | 1.069 | 1.199 | 0.892 | 0.1 | -0.1 | 0.26 | 0.08666667 | -0.18 | -0.17 | 0.863541864 | 0.642321845 | 0.552227133 | 0.999652205 | 0.995559182 | 0.997605693 | 0.967231769 |
| Tusc2 | High | Q9WVF8 | Tumor suppressor candidate 2 OS=Mus musculus OX=10090 GN=Tusc2 PE=1 SV=3 | 0.006 | 3.164 | 8 | 1 | 1 | 1 | 110 | 12.1 | 9.69 | 1.079 | 1.192 | 0.905 | 0.11 | -0.11 | 0.25 | 0.08333333 | -0.18 | -0.14 | 0.851603931 | 0.656955721 | 0.588486107 | 0.999652205 | 0.995559182 | 0.997605693 | 0.973095093 |
| Mertk | High | Q60805 | Tyrosine-protein kinase Mer OS=Mus musculus OX=10090 GN=Mertk PE=1 SV=1 | 0.001 | 6.621 | 2 | 3 | 4 | 3 | 994 | 110.1 | 5.5 | 1.096 | 1.175 | 0.933 | 0.13 | -0.13 | 0.23 | 0.07666667 | -0.18 | -0.1 | 0.802578617 | 0.683501757 | 0.647082822 | 0.999652205 | 0.995559182 | 0.997605693 | 0.985010103 |
| Mrpl9 | Medium | Q99N94 | 39S ribosomal protein L9, mitochondrial OS=Mus musculus OX=10090 GN=Mrpl9 PE=1 SV=2 | 0.013 | 2.279 | 3 | 1 | 1 | 1 | 265 | 30.2 | 10.08 | 1.095 | 1.174 | 0.933 | 0.13 | -0.13 | 0.23 | 0.07666667 | -0.18 | -0.1 | 0.781751052 | 0.693012219 | 0.722540938 | 0.999652205 | 0.995559182 | 0.997605693 | 0.992535481 |
| Psmg1 | High | Q9JK23 | Proteasome assembly chaperone 1 OS=Mus musculus OX=10090 GN=Psmg1 PE=1 SV=1 | 0.01 | 2.745 | 4 | 1 | 1 | 1 | 289 | 33.1 | 6.37 | 1.094 | 1.169 | 0.936 | 0.13 | -0.13 | 0.23 | 0.07666667 | -0.18 | -0.1 | 0.812955464 | 0.69011549 | 0.726623181 | 0.999652205 | 0.995559182 | 0.997605693 | 0.992535481 |
| Sacm1l | High | Q9EP69 | Phosphatidylinositol-3-phosphatase SAC1 OS=Mus musculus OX=10090 GN=Sacm1l PE=1 SV=1 | 0 | 19.289 | 10 | 6 | 6 | 6 | 587 | 66.9 | 7.3 | 1.027 | 1.253 | 0.82 | 0.04 | -0.04 | 0.33 | 0.11 | -0.185 | -0.29 | 0.873979099 | 0.54491176 | 0.255609261 | 0.999652205 | 0.995559182 | 0.997605693 | 0.855700667 |
| H2afx | High | P27661 | Histone H2AX OS=Mus musculus OX=10090 GN=H2ax PE=1 SV=2 | 0 | 10.208 | 13 | 3 | 3 | 2 | 143 | 15.1 | 10.74 | 1.038 | 1.251 | 0.83 | 0.05 | -0.05 | 0.32 | 0.10666667 | -0.185 | -0.27 | 0.954213355 | 0.531903773 | 0.295453949 | 0.999652205 | 0.995559182 | 0.997605693 | 0.894411787 |
| F3 | High | P20352 | Tissue factor OS=Mus musculus OX=10090 GN=F3 PE=1 SV=2 | 0 | 43.686 | 37 | 10 | 15 | 10 | 294 | 32.9 | 9.32 | 1.12 | 1.159 | 0.966 | 0.16 | -0.16 | 0.21 | 0.07 | -0.185 | -0.05 | 0.760503826 | 0.71394636 | 0.81561345 | 0.999652205 | 0.995559182 | 0.997605693 | 0.992535481 |
| Osbpl1a | High | Q3V156 | Oxysterol-binding protein OS=Mus musculus OX=10090 GN=Osbpl1a PE=1 SV=1 | 0.001 | 4.239 | 3 | 2 | 2 | 2 | 558 | 63.4 | 5.68 | 1.118 | 1.157 | 0.966 | 0.16 | -0.16 | 0.21 | 0.07 | -0.185 | -0.05 | 0.732094201 | 0.717815274 | 0.879142278 | 0.999652205 | 0.995559182 | 0.997605693 | 0.992535481 |
| Cnga1 | High | P29974 | cGMP-gated cation channel alpha-1 OS=Mus musculus OX=10090 GN=Cnga1 PE=2 SV=2 | 0 | 433.04 | 50 | 36 | 398 | 30 | 684 | 79.4 | 7.58 | 1.137 | 1.134 | 1.003 | 0.19 | -0.19 | 0.18 | 0.06 | -0.185 | 0 | 0.662408889 | 0.764842376 | 0.90333308 | 0.999652205 | 0.995559182 | 0.997605693 | 0.992535481 |
| Pcdh9 | Medium | A0A0A6YV | Protocadherin 9 OS=Mus musculus OX=10090 GN=Pcdh9 PE=1 SV=1 | 0.01 | 2.684 | 2 | 1 | 1 | 1 | 1203 | 132.2 | 5.47 | 1.123 | 1.148 | 0.978 | 0.17 | -0.17 | 0.2 | 0.06666667 | -0.185 | -0.03 | 0.681771728 | 0.723962563 | 0.925763638 | 0.999652205 | 0.995559182 | 0.997605693 | 0.992535481 |
| Tmx3 | High | Q8BXZ1 | Protein disulfide-isomerase TMX3 OS=Mus musculus OX=10090 GN=Tmx3 PE=1 SV=2 | 0 | 10.535 | 10 | 4 | 4 | 4 | 456 | 51.8 | 5.16 | 0.997 | 1.303 | 0.765 | 0 | 0 | 0.38 | 0.12666667 | -0.19 | -0.39 | 0.995486502 | 0.467563793 | 0.144117443 | 0.999652205 | 0.995559182 | 0.997605693 | 0.706351836 |
| Pdia4 | High | A0A0R4J0 | Protein disulfide-isomerase A4 OS=Mus musculus OX=10090 GN=Pdia4 PE=1 SV=1 | 0 | 38.137 | 21 | 13 | 16 | 13 | 641 | 72.3 | 5.22 | 1.052 | 1.238 | 0.849 | 0.07 | -0.07 | 0.31 | 0.10333333 | -0.19 | -0.24 | 0.862736385 | 0.569053792 | 0.341296503 | 0.999652205 | 0.995559182 | 0.997605693 | 0.907770836 |
| Rab9 | High | Q9R0M6 | Ras-related protein Rab-9A OS=Mus musculus OX=10090 GN=Rab9a PE=1 SV=1 | 0 | 24.614 | 13 | 2 | 5 | 2 | 201 | 22.9 | 5.66 | 1.09 | 1.202 | 0.907 | 0.12 | -0.12 | 0.26 | 0.08666667 | -0.19 | -0.14 | 0.707109099 | 0.633003952 | 0.598581411 | 0.999652205 | 0.995559182 | 0.997605693 | 0.977451587 |
| Thsd7a | High | E9QNR5 | Thrombospondin type-1 domain-containing protein 7A OS=Mus musculus OX=10090 GN=Thsd7a F | 0.001 | 5.175 | 1 | 2 | 3 | 2 | 1646 | 183.5 | 7.2 | 1.097 | 1.189 | 0.923 | 0.13 | -0.13 | 0.25 | 0.08333333 | -0.19 | -0.12 | 0.729292055 | 0.648016401 | 0.661968626 | 0.999652205 | 0.995559182 | 0.997605693 | 0.986260965 |
| Rho | High | P15409 | Rhodopsin OS=Mus musculus OX=10090 GN=Rho PE=1 SV=2 | 0 | 390.65 | 33 | 15 | 1101 | 14 | 348 | 39 | 6.65 | 1.104 | 1.179 | 0.936 | 0.14 | -0.14 | 0.24 | 0.08 | -0.19 | -0.09 | 0.737921673 | 0.675974155 | 0.746260449 | 0.999652205 | 0.995559182 | 0.997605693 | 0.992535481 |
| Dock9 | High | E9QMR2 | Dedicator of cytokinesis protein 9 OS=Mus musculus OX=10090 GN=Dock9 PE=1 SV=1 | 0 | 9.36 | 1 | 3 | 3 | 3 | 2058 | 234.8 | 7.56 | 1.11 | 1.176 | 0.944 | 0.15 | -0.15 | 0.23 | 0.07666667 | -0.19 | -0.08 | 0.735287424 | 0.684195365 | 0.812473541 | 0.999652205 | 0.995559182 | 0.997605693 | 0.992535481 |
| Wars2 | Medium | Q8BZQ9 | Tryptophanyl-tRNA synthetase OS=Mus musculus OX=10090 GN=Wars2 PE=1 SV=1 | 0.011 | 2.52 | 4 | 1 | 1 | 1 | 188 | 21.1 | 8.92 | 1.142 | 1.143 | 1 | 0.19 | -0.19 | 0.19 | 0.06333333 | -0.19 | 0 | 0.687526666 | 0.755341478 | 0.953414678 | 0.999652205 | 0.995559182 | 0.997605693 | 0.992535481 |
| | Low | A0A338P6 | Coiled-coil domain-containing protein 14 OS=Mus musculus OX=10090 GN=Ccdc14 PE=1 SV=1 | 0.056 | 1.365 | 3 | 1 | 1 | 1 | 886 | 97.1 | 6.65 | 1.132 | 1.149 | 0.985 | 0.18 | -0.18 | 0.2 | 0.06666667 | -0.19 | -0.02 | 0.688590596 | 0.736472462 | 0.959013792 | 0.999652205 | 0.995559182 | 0.997605693 | 0.992902921 |
| Ppif | High | Q99KR7 | Peptidyl-prolyl cis-trans isomerase F, mitochondrial OS=Mus musculus OX=10090 GN=Ppif PE=1 S | 0 | 8.889 | 11 | 3 | 4 | 2 | 206 | 21.7 | 9.16 | 1.141 | 1.149 | 0.994 | 0.19 | -0.19 | 0.2 | 0.06666667 | -0.195 | -0.01 | 0.65508362 | 0.733543446 | 0.981347957 | 0.999652205 | 0.995559182 | 0.997605693 | 0.997765878 |
| Zwint | High | Q9CQU5 | ZW10 interactor OS=Mus musculus OX=10090 GN=Zwint PE=1 SV=1 | 0.001 | 6.227 | 9 | 3 | 3 | 3 | 252 | 28.7 | 8.43 | 1.137 | 1.149 | 0.99 | 0.19 | -0.19 | 0.2 | 0.06666667 | -0.195 | -0.01 | 0.661753643 | 0.733934967 | 0.982371572 | 0.999652205 | 0.995559182 | 0.997605693 | 0.997765878 |
| | High | A0A2I3BR | Triple functional domain protein OS=Mus musculus OX=10090 GN=Trio PE=1 SV=1 | 0.001 | 5.061 | 1 | 2 | 3 | 2 | 1849 | 211.5 | 6.19 | 1.079 | 1.224 | 0.882 | 0.11 | -0.11 | 0.29 | 0.09666667 | -0.2 | -0.18 | 0.815414115 | 0.596554861 | 0.519512876 | 0.999652205 | 0.995559182 | 0.997605693 | 0.964378027 |
| Tmem65 | High | Q4VAE3 | Transmembrane protein 65 OS=Mus musculus OX=10090 GN=Tmem65 PE=1 SV=1 | 0 | 12.114 | 13 | 3 | 4 | 3 | 234 | 24.9 | 7.78 | 1.011 | 1.305 | 0.774 | 0.02 | -0.02 | 0.38 | 0.12666667 | -0.2 | -0.37 | 0.945126142 | 0.464537972 | 0.157808692 | 0.999652205 | 0.995559182 | 0.997605693 | 0.733365826 |
| Aldh1a1 | High | P24549 | Retinal dehydrogenase 1 OS=Mus musculus OX=10090 GN=Aldh1a1 PE=1 SV=5 | 0 | 42.676 | 29 | 12 | 15 | 11 | 501 | 54.4 | 7.8 | 1.044 | 1.268 | 0.824 | 0.06 | -0.06 | 0.34 | 0.11333333 | -0.2 | -0.28 | 0.932184612 | 0.52017316 | 0.223164859 | 0.999652205 | 0.995559182 | 0.997605693 | 0.80884639 |
| Cend1 | High | Q9JKC6 | Cell cycle exit and neuronal differentiation protein 1 OS=Mus musculus OX=10090 GN=Cend1 PE= | 0 | 47.089 | 58 | 9 | 19 | 9 | 149 | 15 | 8.97 | 1.039 | 1.265 | 0.821 | 0.06 | -0.06 | 0.34 | 0.11333333 | -0.2 | -0.28 | 0.894614597 | 0.525240782 | 0.2430572 | 0.999652205 | 0.995559182 | 0.997605693 | 0.837310213 |
| | High | A0A087W | Protein ABHD14B (Fragment) OS=Mus musculus OX=10090 GN=Abhd14b PE=1 SV=1 | 0 | 28.483 | 55 | 8 | 10 | 8 | 174 | 18.4 | 6.79 | 1.09 | 1.214 | 0.898 | 0.12 | -0.12 | 0.28 | 0.09333333 | -0.2 | -0.16 | 0.827721634 | 0.611377033 | 0.495649493 | 0.999652205 | 0.995559182 | 0.997605693 | 0.964378027 |
| Serpinb6a | High | F8WIV2 | Serine (or cysteine) peptidase inhibitor, clade B, member 6a OS=Mus musculus OX=10090 GN=Se | 0 | 17.523 | 12 | 4 | 5 | 4 | 399 | 44.7 | 6.38 | 1.112 | 1.187 | 0.937 | 0.15 | -0.15 | 0.25 | 0.08333333 | -0.2 | -0.09 | 0.718622038 | 0.659411326 | 0.66351695 | 0.999652205 | 0.995559182 | 0.997605693 | 0.986647506 |
| Ddx3x | High | Q62167 | ATP-dependent RNA helicase DDX3X OS=Mus musculus OX=10090 GN=Ddx3x PE=1 SV=3 | 0 | 101.194 | 36 | 22 | 33 | 22 | 662 | 73.1 | 7.18 | 1.109 | 1.189 | 0.933 | 0.15 | -0.15 | 0.25 | 0.08333333 | -0.2 | -0.1 | 0.725712656 | 0.656681727 | 0.727652833 | 0.999652205 | 0.995559182 | 0.997605693 | 0.992535481 |
| Mtch2 | High | Q791V5 | Mitochondrial carrier homolog 2 OS=Mus musculus OX=10090 GN=Mtch2 PE=1 SV=1 | 0 | 11.066 | 19 | 4 | 4 | 4 | 303 | 33.5 | 8.25 | 1.114 | 1.181 | 0.943 | 0.16 | -0.16 | 0.24 | 0.08 | -0.2 | -0.08 | 0.728469787 | 0.677178964 | 0.800228296 | 0.999652205 | 0.995559182 | 0.997605693 | 0.992535481 |
| Rab1; Rab | High | P62821 | Ras-related protein Rab-1A OS=Mus musculus OX=10090 GN=Rab1A PE=1 SV=3 | 0 | 166.989 | 79 | 17 | 96 | 10 | 205 | 22.7 | 6.21 | 1.045 | 1.278 | 0.818 | 0.06 | -0.06 | 0.35 | 0.11666667 | -0.205 | -0.29 | 0.8794327 | 0.505145047 | 0.231708112 | 0.999652205 | 0.995559182 | 0.997605693 | 0.819676582 |
| Tbc1d2 | High | B1AVH7 | TBC1 domain family member 2A OS=Mus musculus OX=10090 GN=Tbc1d2 PE=2 SV=1 | 0 | 15.417 | 7 | 6 | 6 | 6 | 922 | 104.2 | 6.43 | 1.09 | 1.222 | 0.892 | 0.12 | -0.12 | 0.29 | 0.09666667 | -0.205 | -0.17 | 0.714165725 | 0.59614184 | 0.521356663 | 0.999652205 | 0.995559182 | 0.997605693 | 0.964378027 |
| Tubb4a | High | Q9D6F9 | Tubulin beta-4A chain OS=Mus musculus OX=10090 GN=Tubb4a PE=1 SV=3 | 0 | 417.925 | 74 | 24 | 190 | 5 | 444 | 49.6 | 4.88 | 1.012 | 1.306 | 0.775 | 0.02 | -0.02 | 0.39 | 0.13 | -0.205 | -0.37 | 0.981055821 | 0.46320607 | 0.106914595 | 0.999652205 | 0.995559182 | 0.997605693 | 0.626405572 |
| Rab14 | High | Q91V41 | Ras-related protein Rab-14 OS=Mus musculus OX=10090 GN=Rab14 PE=1 SV=3 | 0 | 161.944 | 83 | 18 | 53 | 16 | 215 | 23.9 | 6.21 | 1.064 | 1.248 | 0.852 | 0.09 | -0.09 | 0.32 | 0.10666667 | -0.205 | -0.23 | 0.833335839 | 0.55287315 | 0.352280156 | 0.999652205 | 0.995559182 | 0.997605693 | 0.910898173 |
| Pla2g16 | Medium | Q8R3U1 | Phospholipase A and acyltransferase 3 OS=Mus musculus OX=10090 GN=Plaat3 PE=1 SV=2 | 0.047 | 1.559 | 6 | 1 | 1 | 1 | 162 | 17.9 | 8.72 | 1.107 | 1.196 | 0.926 | 0.15 | -0.15 | 0.26 | 0.08666667 | -0.205 | -0.11 | 0.776347006 | 0.646625171 | 0.673152054 | 0.999652205 | 0.995559182 | 0.997605693 | 0.990250803 |
| Rab39b | High | Q8BHC1 | Ras-related protein Rab-39B OS=Mus musculus OX=10090 GN=Rab39b PE=1 SV=1 | 0 | 49.914 | 59 | 13 | 21 | 11 | 213 | 24.6 | 7.83 | 1.16 | 1.152 | 1.006 | 0.21 | -0.21 | 0.2 | 0.06666667 | -0.205 | 0.01 | 0.615935602 | 0.727105205 | 0.887663178 | 0.999652205 | 0.995559182 | 0.997605693 | 0.992535481 |
| Faf2 | High | Q3TDN2 | FAS-associated factor 2 OS=Mus musculus OX=10090 GN=Faf2 PE=1 SV=2 | 0 | 20.209 | 15 | 4 | 4 | 4 | 445 | 52.4 | 5.47 | 1.139 | 1.163 | 0.979 | 0.19 | -0.19 | 0.22 | 0.07333333 | -0.205 | -0.03 | 0.653611023 | 0.705915513 | 0.924516708 | 0.999652205 | 0.995559182 | 0.997605693 | 0.992535481 |
| Plin3 | High | Q9DBG5 | Perilipin-3 OS=Mus musculus OX=10090 GN=Plin3 PE=1 SV=1 | 0 | 30.617 | 23 | 7 | 7 | 7 | 437 | 47.2 | 5.62 | 1.14 | 1.168 | 0.976 | 0.19 | -0.19 | 0.22 | 0.07333333 | -0.205 | -0.04 | 0.657266547 | 0.695421301 | 0.934863702 | 0.999652205 | 0.995559182 | 0.997605693 | 0.992535481 |
| Tnrc6b | High | Q8BKI2 | Trinucleotide repeat-containing gene 6B protein OS=Mus musculus OX=10090 GN=Tnrc6b PE=1 S | 0.001 | 4.307 | 1 | 1 | 1 | 1 | 1810 | 191.8 | 6.3 | 1.028 | 1.305 | 0.788 | 0.04 | -0.04 | 0.38 | 0.12666667 | -0.21 | -0.34 | 0.984286123 | 0.479652097 | 0.259410489 | 0.999652205 | 0.995559182 | 0.997605693 | 0.85616924 |
| Dmd | High | P11531 | Dystrophin OS=Mus musculus OX=10090 GN=Dmd PE=1 SV=3 | 0.001 | 6.424 | 1 | 2 | 2 | 1 | 3678 | 425.6 | 5.94 | 1.079 | 1.243 | 0.868 | 0.11 | -0.11 | 0.31 | 0.10333333 | -0.21 | -0.2 | 0.798905945 | 0.544353405 | 0.464043743 | 0.999652205 | 0.995559182 | 0.997605693 | 0.959744396 |
| Tmem33 | High | A0A0R4J1 | Transmembrane protein 33 OS=Mus musculus OX=10090 GN=Tmem33 PE=1 SV=1 | 0.003 | 3.978 | 4 | 1 | 1 | 1 | 246 | 27.9 | 9.66 | 1.059 | 1.266 | 0.836 | 0.08 | -0.08 | 0.34 | 0.11333333 | -0.21 | -0.26 | 0.878188356 | 0.518465621 | 0.319220965 | 0.999652205 | 0.995559182 | 0.997605693 | 0.904943958 |
| Pak3 | High | Q61036 | Serine/threonine-protein kinase PAK 3 OS=Mus musculus OX=10090 GN=Pak3 PE=1 SV=2 | 0 | 25.094 | 13 | 7 | 10 | 2 | 559 | 62.4 | 5.4 | 1.055 | 1.269 | 0.832 | 0.08 | -0.08 | 0.34 | 0.11333333 | -0.21 | -0.27 | 0.874888662 | 0.502946269 | 0.333269024 | 0.999652205 | 0.995559182 | 0.997605693 | 0.906329766 |
| Ift122 | High | Q6NWV3 | Intraflagellar transport protein 122 homolog OS=Mus musculus OX=10090 GN=Ift122 PE=1 SV=1 | 0 | 14.075 | 2 | 2 | 3 | 2 | 1182 | 134.7 | 6.98 | 1.141 | 1.174 | 0.971 | 0.19 | -0.19 | 0.23 | 0.07666667 | -0.21 | -0.04 | 0.634477367 | 0.673265419 | 0.892931794 | 0.999652205 | 0.995559182 | 0.997605693 | 0.992535481 |
| | High | F6VVY4 | Tricarboxylate transport protein, mitochondrial (Fragment) OS=Mus musculus OX=10090 GN=Slc2 | 0 | 8.61 | 10 | 2 | 2 | 2 | 197 | 21.5 | 9.8 | 1.048 | 1.281 | 0.818 | 0.07 | -0.07 | 0.36 | 0.12 | -0.215 | -0.29 | 0.932845266 | 0.500744276 | 0.236104756 | 0.999652205 | 0.995559182 | 0.997605693 | 0.827917053 |
| Dnajb4 | High | Q9D832 | DnaJ homolog subfamily B member 4 OS=Mus musculus OX=10090 GN=Dnajb4 PE=1 SV=1 | 0 | 24.808 | 17 | 5 | 6 | 4 | 337 | 37.8 | 8.59 | 1.098 | 1.227 | 0.895 | 0.13 | -0.13 | 0.3 | 0.1 | -0.215 | -0.16 | 0.812367282 | 0.587808721 | 0.500645908 | 0.999652205 | 0.995559182 | 0.997605693 | 0.964378027 |
| Gulp1 | High | Q8K2A1 | PTB domain-containing engulfment adapter protein 1 OS=Mus musculus OX=10090 GN=Gulp1 PE | 0 | 19.979 | 18 | 7 | 8 | 7 | 304 | 34.4 | 7.9 | 1.104 | 1.226 | 0.901 | 0.14 | -0.14 | 0.29 | 0.09666667 | -0.215 | -0.15 | 0.76513084 | 0.589256753 | 0.543497151 | 0.999652205 | 0.995559182 | 0.997605693 | 0.96694628 |
| Pip4k2c | High | Q91XU3 | Phosphatidylinositol 5-phosphate 4-kinase type-2 gamma OS=Mus musculus OX=10090 GN=Pip4k | 0 | 37.867 | 24 | 12 | 17 | 11 | 421 | 47.3 | 6.89 | 1.136 | 1.192 | 0.952 | 0.18 | -0.18 | 0.25 | 0.08333333 | -0.215 | -0.07 | 0.666230152 | 0.649964352 | 0.830825746 | 0.999652205 | 0.995559182 | 0.997605693 | 0.992535481 |
| Adrbk1; Gr | High | Q99MK8 | Beta-adrenergic receptor kinase 1 OS=Mus musculus OX=10090 GN=Grk2 PE=1 SV=2 | 0.008 | 3.01 | 4 | 2 | 2 | 2 | 689 | 79.6 | 7.28 | 1.142 | 1.181 | 0.966 | 0.19 | -0.19 | 0.24 | 0.08 | -0.215 | -0.05 | 0.668176932 | 0.666312518 | 0.878832228 | 0.999652205 | 0.995559182 | 0.997605693 | 0.992535481 |
| Eif2b1 | High | Q99LC8 | Translation initiation factor eIF-2B subunit alpha OS=Mus musculus OX=10090 GN=Eif2b1 PE=1 S | 0 | 13.157 | 8 | 2 | 2 | 2 | 305 | 33.8 | 8.32 | 1.162 | 1.153 | 1.008 | 0.22 | -0.22 | 0.21 | 0.07 | -0.215 | 0.01 | 0.612462236 | 0.725703998 | 0.941391327 | 0.999652205 | 0.995559182 | 0.997605693 | 0.992535481 |
| Tor1aip1 | High | Q921T2 | Torsin-1A-interacting protein 1 OS=Mus musculus OX=10090 GN=Tor1aip1 PE=1 SV=3 | 0.001 | 6.33 | 4 | 2 | 3 | 2 | 595 | 66.7 | 7.05 | 1.153 | 1.163 | 0.992 | 0.21 | -0.21 | 0.22 | 0.07333333 | -0.215 | -0.01 | 0.694935627 | 0.712981201 | 0.99930189 | 0.999652205 | 0.995559182 | 0.997605693 | 0.999917596 |
| Atxn2 | High | O70305 | Ataxin-2 OS=Mus musculus OX=10090 GN=Atxn2 PE=1 SV=1 | 0 | 22.231 | 4 | 6 | 6 | 4 | 1285 | 136.4 | 9.55 | 1.109 | 1.218 | 0.91 | 0.15 | -0.15 | 0.28 | 0.09333333 | -0.215 | -0.14 | 0.746064174 | 0.604460273 | 0.621857481 | 0.999652205 | 0.995559182 | 0.997605693 | 0.97826833 |
| Egln1 | High | Q91YE3 | Egl nine homolog 1 OS=Mus musculus OX=10090 GN=Egln1 PE=1 SV=2 | 0 | 9.466 | 14 | 3 | 3 | 3 | 400 | 43.1 | 8.29 | 1.119 | 1.204 | 0.929 | 0.16 | -0.16 | 0.27 | 0.09 | -0.215 | -0.11 | 0.695884234 | 0.617921402 | 0.684569746 | 0.999652205 | 0.995559182 | 0.997605693 | 0.991285312 |
| Pard6b | High | Q9JK83 | Partitioning defective 6 homolog beta OS=Mus musculus OX=10090 GN=Pard6b PE=1 SV=2 | 0 | 8.153 | 6 | 2 | 2 | 2 | 371 | 41 | 5.59 | 1.177 | 1.148 | 1.025 | 0.23 | -0.23 | 0.2 | 0.06666667 | -0.215 | 0.03 | 0.572433222 | 0.734848339 | 0.857711231 | 0.999652205 | 0.995559182 | 0.997605693 | 0.992535481 |
| Dld | High | O08749 | Dihydrolipoyl dehydrogenase, mitochondrial OS=Mus musculus OX=10090 GN=Dld PE=1 SV=2 | 0 | 16.078 | 9 | 4 | 4 | 4 | 509 | 54.2 | 7.9 | 1.068 | 1.275 | 0.837 | 0.09 | -0.09 | 0.35 | 0.11666667 | -0.22 | -0.26 | 0.823498863 | 0.509885545 | 0.295647504 | 0.999652205 | 0.995559182 | 0.997605693 | 0.894411787 |
| Llgl2 | High | Q3TJ91 | LLGL scribble cell polarity complex component 2 OS=Mus musculus OX=10090 GN=Llgl2 PE=1 SV= | 0.001 | 6.79 | 2 | 2 | 2 | 2 | 1027 | 114.3 | 7.46 | 1.09 | 1.245 | 0.875 | 0.12 | -0.12 | 0.32 | 0.10666667 | -0.22 | -0.19 | 0.812589595 | 0.572358701 | 0.556156982 | 0.999652205 | 0.995559182 | 0.997605693 | 0.970246484 |
| Slc25a46 | Low | Q9CQS4 | Solute carrier family 25 member 46 OS=Mus musculus OX=10090 GN=Slc25a46 PE=1 SV=1 | 0.056 | 1.373 | 2 | 1 | 1 | 1 | 418 | 46.2 | 7.64 | 1.102 | 1.229 | 0.897 | 0.14 | -0.14 | 0.3 | 0.1 | -0.22 | -0.16 | 0.770886245 | 0.588823329 | 0.557764564 | 0.999652205 | 0.995559182 | 0.997605693 | 0.971568588 |
| | High | E9Q740 | Signal recognition particle subunit SRP72 OS=Mus musculus OX=10090 GN=Srp72 PE=1 SV=1 | 0.006 | 3.262 | 3 | 1 | 1 | 1 | 610 | 67.9 | 9.38 | 1.201 | 1.137 | 1.057 | 0.26 | -0.26 | 0.18 | 0.06 | -0.22 | 0.08 | 0.51780853 | 0.770341581 | 0.673763784 | 0.999652205 | 0.995559182 | 0.997605693 | 0.990250803 |
| Auh | High | A0A0R4J0 | Methylglutaconyl-CoA hydratase, mitochondrial OS=Mus musculus OX=10090 GN=Auh PE=1 SV=1 | 0 | 22.868 | 21 | 7 | 10 | 7 | 314 | 33.3 | 9.42 | 1.131 | 1.2 | 0.942 | 0.18 | -0.18 | 0.26 | 0.08666667 | -0.22 | -0.09 | 0.69366864 | 0.63595916 | 0.788249924 | 0.999652205 | 0.995559182 | 0.997605693 | 0.992535481 |
| Arl3 | High | Q9WUL7 | ADP-ribosylation factor-like protein 3 OS=Mus musculus OX=10090 GN=Arl3 PE=1 SV=1 | 0 | 161.08 | 82 | 14 | 63 | 14 | 182 | 20.5 | 7.24 | 1.142 | 1.191 | 0.959 | 0.19 | -0.19 | 0.25 | 0.08333333 | -0.22 | -0.06 | 0.652143506 | 0.652483883 | 0.865778347 | 0.999652205 | 0.995559182 | 0.997605693 | 0.992535481 |
| Abat | High | P61922 | 4-aminobutyrate aminotransferase, mitochondrial OS=Mus musculus OX=10090 GN=Abat PE=1 S | 0 | 108.071 | 44 | 16 | 28 | 16 | 500 | 56.4 | 8.09 | 1.143 | 1.19 | 0.961 | 0.19 | -0.19 | 0.25 | 0.08333333 | -0.22 | -0.06 | 0.649984422 | 0.654355376 | 0.874590027 | 0.999652205 | 0.995559182 | 0.997605693 | 0.992535481 |
| | High | A0A3B2W | Spastin OS=Mus musculus OX=10090 GN=Spast PE=1 SV=1 | 0.002 | 4.162 | 4 | 2 | 3 | 2 | 581 | 62.7 | 9.76 | 1.169 | 1.154 | 1.013 | 0.23 | -0.23 | 0.21 | 0.07 | -0.22 | 0.02 | 0.602902083 | 0.724100369 | 0.889666389 | 0.999652205 | 0.995559182 | 0.997605693 | 0.992535481 |
| Camk2d | High | A0A0G2JG | Calcium/calmodulin-dependent protein kinase OS=Mus musculus OX=10090 GN=Camk2d PE=1 SV | 0 | 126.295 | 48 | 20 | 60 | 2 | 533 | 60 | 7.27 | 1.166 | 1.168 | 0.998 | 0.22 | -0.22 | 0.22 | 0.07333333 | -0.22 | 0 | 0.604094064 | 0.696886278 | 0.92952785 | 0.999652205 | 0.995559182 | 0.997605693 | 0.992535481 |
| Ndufb3 | High | Q9CQZ6 | NADH dehydrogenase [ubiquinone] 1 beta subcomplex subunit 3 OS=Mus musculus OX=10090 GN | 0.005 | 3.663 | 11 | 1 | 1 | 1 | 104 | 11.7 | 9.04 | 1.164 | 1.162 | 1.001 | 0.22 | -0.22 | 0.22 | 0.07333333 | -0.22 | 0 | 0.6365609 | 0.714675022 | 0.94435898 | 0.999652205 | 0.995559182 | 0.997605693 | 0.992535481 |
| Car12 | High | Q8K2J1 | Carbonic anhydrase OS=Mus musculus OX=10090 GN=Car12 PE=1 SV=1 | 0.008 | 2.973 | 3 | 1 | 1 | 1 | 344 | 38.7 | 8.31 | 1.157 | 1.175 | 0.985 | 0.21 | -0.21 | 0.23 | 0.07666667 | -0.22 | -0.02 | 0.600369262 | 0.675165157 | 0.945637957 | 0.999652205 | 0.995559182 | 0.997605693 | 0.992535481 |
| Ybx1 | High | P62960 | Y-box-binding protein 1 OS=Mus musculus OX=10090 GN=Ybx1 PE=1 SV=3 | 0 | 21.992 | 13 | 3 | 4 | 2 | 322 | 35.7 | 9.88 | 1.117 | 1.213 | 0.921 | 0.16 | -0.16 | 0.28 | 0.09333333 | -0.22 | -0.12 | 0.689556178 | 0.613220037 | 0.662253765 | 0.999652205 | 0.995559182 | 0.997605693 | 0.986260965 |
| Psd | Medium | Q5DTT2 | PH and SEC7 domain-containing protein 1 OS=Mus musculus OX=10090 GN=Psd PE=1 SV=2 | 0.048 | 1.517 | 1 | 1 | 1 | 1 | 1024 | 109.6 | 7.05 | 1.109 | 1.234 | 0.899 | 0.15 | -0.15 | 0.3 | 0.1 | -0.225 | -0.15 | 0.743327816 | 0.563481725 | 0.563477268 | 0.999652205 | 0.995559182 | 0.997605693 | 0.972910028 |
| Stra6 | High | A0A0R4J1 | Receptor for retinol uptake STRA6 OS=Mus musculus OX=10090 GN=Stra6 PE=1 SV=1 | 0 | 24.63 | 9 | 6 | 11 | 6 | 670 | 73.7 | 8.53 | 1.182 | 1.155 | 1.023 | 0.24 | -0.24 | 0.21 | 0.07 | -0.225 | 0.03 | 0.582157223 | 0.72109612 | 0.81716829 | 0.999652205 | 0.995559182 | 0.997605693 | 0.992535481 |
| Timm10 | High | P62073 | Mitochondrial import inner membrane translocase subunit Tim10 OS=Mus musculus OX=10090 G | 0 | 9.86 | 26 | 2 | 4 | 2 | 90 | 10.3 | 6.29 | 1.154 | 1.178 | 0.98 | 0.21 | -0.21 | 0.24 | 0.08 | -0.225 | -0.03 | 0.624920806 | 0.677045313 | 0.931057023 | 0.999652205 | 0.995559182 | 0.997605693 | 0.992535481 |
| Ndufs5 | High | B1ARW4 | Complex I-15 kDa (Fragment) OS=Mus musculus OX=10090 GN=Ndufs5 PE=1 SV=1 | 0 | 13.671 | 35 | 3 | 5 | 3 | 91 | 10.9 | 8.94 | 1.05 | 1.3 | 0.807 | 0.07 | -0.07 | 0.38 | 0.12666667 | -0.225 | -0.31 | 0.913879941 | 0.471435693 | 0.194421367 | 0.999652205 | 0.995559182 | 0.997605693 | 0.784517504 |
| D17Wsu1( | High | Q9CPT4 | Myeloid-derived growth factor OS=Mus musculus OX=10090 GN=Mydgf PE=1 SV=1 | 0 | 8.368 | 16 | 2 | 3 | 2 | 166 | 18 | 6.79 | 1.057 | 1.293 | 0.817 | 0.08 | -0.08 | 0.37 | 0.12333333 | -0.225 | -0.29 | 0.902941894 | 0.481816051 | 0.252950169 | 0.999652205 | 0.995559182 | 0.997605693 | 0.851315484 |
| | High | A0A571BE | InaD-like protein OS=Mus musculus OX=10090 GN=Patj PE=1 SV=1 | 0 | 8.315 | 2 | 3 | 3 | 2 | 1858 | 200.8 | 4.98 | 1.076 | 1.262 | 0.852 | 0.11 | -0.11 | 0.34 | 0.11333333 | -0.225 | -0.23 | 0.84102208 | 0.513558144 | 0.379721921 | 0.999652205 | 0.995559182 | 0.997605693 | 0.922487352 |
| Emc8 | High | O70378 | ER membrane protein complex subunit 8 OS=Mus musculus OX=10090 GN=Emc8 PE=1 SV=1 | 0 | 11.971 | 23 | 4 | 4 | 3 | 207 | 23.3 | 6.15 | 1.105 | 1.241 | 0.89 | 0.14 | -0.14 | 0.31 | 0.10333333 | -0.225 | -0.17 | 0.661947032 | 0.564162844 | 0.513239437 | 0.999652205 | 0.995559182 | 0.997605693 | 0.964378027 |
| Aldh5a1 | High | Q8BWF0 | Succinate-semialdehyde dehydrogenase, mitochondrial OS=Mus musculus OX=10090 GN=Aldh5a | 0 | 51.05 | 27 | 11 | 15 | 11 | 523 | 55.9 | 8.25 | 1.135 | 1.209 | 0.939 | 0.18 | -0.18 | 0.27 | 0.09 | -0.225 | -0.09 | 0.693216794 | 0.619708079 | 0.723582181 | 0.999652205 | 0.995559182 | 0.997605693 | 0.992535481 |
| Padi2 | High | Q08642 | Protein-arginine deiminase type-2 OS=Mus musculus OX=10090 GN=Padi2 PE=1 SV=2 | 0 | 44.1 | 15 | 8 | 11 | 8 | 673 | 76.2 | 5.53 | 1.191 | 1.149 | 1.036 | 0.25 | -0.25 | 0.2 | 0.06666667 | -0.225 | 0.05 | 0.580313698 | 0.732747025 | 0.799642225 | 0.999652205 | 0.995559182 | 0.997605693 | 0.992535481 |
| | Medium | A0A494B9 | Cytochrome c oxidase subunit 8A, mitochondrial OS=Mus musculus OX=10090 GN=Cox8a PE=3 SV | 0.011 | 2.513 | 21 | 1 | 1 | 1 | 43 | 4.7 | 12.18 | 1.148 | 1.188 | 0.966 | 0.2 | -0.2 | 0.25 | 0.08333333 | -0.225 | -0.05 | 0.621464817 | 0.648600323 | 0.869743107 | 0.999652205 | 0.995559182 | 0.997605693 | 0.992535481 |
| Slc17a7 | High | Q3TXX4 | Vesicular glutamate transporter 1 OS=Mus musculus OX=10090 GN=Slc17a7 PE=1 SV=2 | 0 | 17.067 | 6 | 5 | 5 | 5 | 560 | 61.6 | 7.34 | 1.125 | 1.222 | 0.921 | 0.17 | -0.17 | 0.29 | 0.09666667 | -0.23 | -0.12 | 0.743971045 | 0.596656495 | 0.585010357 | 0.999652205 | 0.995559182 | 0.997605693 | 0.973095093 |
| Abcb6 | Low | Q9DC29 | ATP-binding cassette sub-family B member 6 OS=Mus musculus OX=10090 GN=Abcb6 PE=1 SV=1 | 0.052 | 1.472 | 1 | 1 | 1 | 1 | 842 | 93.7 | 8.05 | 1.112 | 1.238 | 0.898 | 0.15 | -0.15 | 0.31 | 0.10333333 | -0.23 | -0.15 | 0.743297683 | 0.569416083 | 0.585705147 | 0.999652205 | 0.995559182 | 0.997605693 | 0.973095093 |
| 2810405K( | High | Q9DB60 | Prostamide/prostaglandin F synthase OS=Mus musculus OX=10090 GN=Prxl2b PE=1 SV=1 | 0 | 10.582 | 13 | 3 | 3 | 3 | 201 | 21.7 | 6.74 | 1.185 | 1.161 | 1.02 | 0.24 | -0.24 | 0.22 | 0.07333333 | -0.23 | 0.03 | 0.543390171 | 0.70898978 | 0.939064496 | 0.999652205 | 0.995559182 | 0.997605693 | 0.992535481 |
| Sec62 | High | Q8BU14 | Translocation protein SEC62 OS=Mus musculus OX=10090 GN=Sec62 PE=1 SV=1 | 0 | 21.654 | 17 | 7 | 8 | 7 | 398 | 45.6 | 7.31 | 1.168 | 1.179 | 0.99 | 0.22 | -0.22 | 0.24 | 0.08 | -0.23 | -0.01 | 0.639379745 | 0.674281502 | 0.939373969 | 0.999652205 | 0.995559182 | 0.997605693 | 0.992535481 |
| Lars | High | Q8BMJ2 | Leucine--tRNA ligase, cytoplasmic OS=Mus musculus OX=10090 GN=Lars1 PE=1 SV=2 | 0.001 | 6.997 | 2 | 3 | 3 | 3 | 1178 | 134.1 | 7.05 | 1.203 | 1.14 | 1.055 | 0.27 | -0.27 | 0.19 | 0.06333333 | -0.23 | 0.08 | 0.5075882 | 0.765586644 | 0.698432948 | 0.999652205 | 0.995559182 | 0.997605693 | 0.991285312 |
| Gucy2e | High | P52785 | Retinal guanylyl cyclase 1 OS=Mus musculus OX=10090 GN=Gucy2e PE=1 SV=2 | 0 | 924.734 | 64 | 64 | 664 | 56 | 1108 | 120.6 | 7.77 | 1.142 | 1.208 | 0.945 | 0.19 | -0.19 | 0.27 | 0.09 | -0.23 | -0.08 | 0.653478044 | 0.622210984 | 0.793577515 | 0.999652205 | 0.995559182 | 0.997605693 | 0.992535481 |
| Myo1b | High | P46735 | Unconventional myosin-Ib OS=Mus musculus OX=10090 GN=Myo1b PE=1 SV=3 | 0.001 | 6.955 | 2 | 3 | 3 | 3 | 1107 | 128.5 | 9.26 | 1.15 | 1.199 | 0.959 | 0.2 | -0.2 | 0.26 | 0.08666667 | -0.23 | -0.06 | 0.64534393 | 0.634330496 | 0.840497486 | 0.999652205 | 0.995559182 | 0.997605693 | 0.992535481 |
| Map1b; M | High | P14873 | Microtubule-associated protein 1B OS=Mus musculus OX=10090 GN=Map1b PE=1 SV=2 | 0 | 277.361 | 34 | 59 | 95 | 57 | 2464 | 270.1 | 4.83 | 1.175 | 1.173 | 1.002 | 0.23 | -0.23 | 0.23 | 0.07666667 | -0.23 | 0 | 0.584767793 | 0.687426508 | 0.908479153 | 0.999652205 | 0.995559182 | 0.997605693 | 0.992535481 |
| Rdh14 | High | Q9ERI6 | Retinol dehydrogenase 14 OS=Mus musculus OX=10090 GN=Rdh14 PE=1 SV=1 | 0 | 50.198 | 38 | 10 | 16 | 10 | 334 | 36.3 | 8.18 | 1.075 | 1.288 | 0.835 | 0.1 | -0.1 | 0.37 | 0.12333333 | -0.235 | -0.26 | 0.824203756 | 0.489842189 | 0.309990903 | 0.999652205 | 0.995559182 | 0.997605693 | 0.904943958 |
| | High | D3Z2Z6 | Phospholipid-transporting ATPase IB (Fragment) OS=Mus musculus OX=10090 GN=Atp8a2 PE=1 S | 0 | 37.026 | 64 | 5 | 18 | 3 | 73 | 8 | 9.72 | 1.088 | 1.276 | 0.853 | 0.12 | -0.12 | 0.35 | 0.11666667 | -0.235 | -0.23 | 0.819701887 | 0.508348154 | 0.310073789 | 0.999652205 | 0.995559182 | 0.997605693 | 0.904943958 |
| Gucy2g | Low | Q6TL19 | Guanylate cyclase 2G OS=Mus musculus OX=10090 GN=Gucy2g PE=1 SV=1 | 0.055 | 1.386 | 2 | 1 | 1 | 1 | 1100 | 122.5 | 6.39 | 1.108 | 1.252 | 0.885 | 0.15 | -0.15 | 0.32 | 0.10666667 | -0.235 | -0.18 | 0.742411651 | 0.533852048 | 0.492103176 | 0.999652205 | 0.995559182 | 0.997605693 | 0.964378027 |
| Coasy | High | Q9DBL7 | Bifunctional coenzyme A synthase OS=Mus musculus OX=10090 GN=Coasy PE=1 SV=2 | 0.001 | 5.737 | 4 | 2 | 2 | 2 | 563 | 62 | 7.11 | 1.151 | 1.207 | 0.953 | 0.2 | -0.2 | 0.27 | 0.09 | -0.235 | -0.07 | 0.691193112 | 0.620371836 | 0.803268946 | 0.999652205 | 0.995559182 | 0.997605693 | 0.992535481 |
| Rdh8 | High | D3Z6W3 | Retinol dehydrogenase OS=Mus musculus OX=10090 GN=Rdh8 PE=3 SV=1 | 0 | 114.404 | 59 | 18 | 65 | 18 | 317 | 34.7 | 8.65 | 1.074 | 1.301 | 0.825 | 0.1 | -0.1 | 0.38 | 0.12666667 | -0.24 | -0.28 | 0.807377934 | 0.470384392 | 0.255953592 | 0.999652205 | 0.995559182 | 0.997605693 | 0.855700667 |
| Arl6ip5 | Medium | Q8R5J9 | PRA1 family protein 3 OS=Mus musculus OX=10090 GN=Arl6ip5 PE=1 SV=2 | 0.035 | 1.779 | 4 | 1 | 1 | 1 | 188 | 21.5 | 9.61 | 1.079 | 1.294 | 0.834 | 0.11 | -0.11 | 0.37 | 0.12333333 | -0.24 | -0.26 | 0.807231799 | 0.469932226 | 0.32574793 | 0.999652205 | 0.995559182 | 0.997605693 | 0.905686588 |
| | High | A0A0U1RI | Membrane frizzled-related protein OS=Mus musculus OX=10090 GN=Mfrp PE=4 SV=1 | 0 | 45.992 | 19 | 8 | 12 | 8 | 584 | 63.6 | 5.29 | 1.122 | 1.238 | 0.906 | 0.17 | -0.17 | 0.31 | 0.10333333 | -0.24 | -0.14 | 0.707140825 | 0.569366563 | 0.524739881 | 0.999652205 | 0.995559182 | 0.997605693 | 0.964378027 |
| Prosc; Plpt | High | Q9Z2Y8 | Pyridoxal phosphate homeostasis protein OS=Mus musculus OX=10090 GN=Plpbp PE=1 SV=1 | 0 | 9.749 | 14 | 4 | 4 | 4 | 274 | 30 | 8.27 | 1.188 | 1.172 | 1.014 | 0.25 | -0.25 | 0.23 | 0.07666667 | -0.24 | 0.02 | 0.539958632 | 0.689131523 | 0.870834533 | 0.999652205 | 0.995559182 | 0.997605693 | 0.992535481 |
| Pcbp2 | High | Q61990 | Poly(rC)-binding protein 2 OS=Mus musculus OX=10090 GN=Pcbp2 PE=1 SV=1 | 0 | 83.889 | 41 | 11 | 28 | 5 | 362 | 38.2 | 6.79 | 1.183 | 1.181 | 1.002 | 0.24 | -0.24 | 0.24 | 0.08 | -0.24 | 0 | 0.563353936 | 0.670722967 | 0.911374403 | 0.999652205 | 0.995559182 | 0.997605693 | 0.992535481 |
| Gstt1 | High | Q64471 | Glutathione S-transferase theta-1 OS=Mus musculus OX=10090 GN=Gstt1 PE=1 SV=4 | 0 | 10.483 | 20 | 5 | 5 | 5 | 240 | 27.4 | 7.27 | 1.1 | 1.263 | 0.871 | 0.14 | -0.14 | 0.34 | 0.11333333 | -0.24 | -0.2 | 0.789073389 | 0.52860639 | 0.386007998 | 0.999652205 | 0.995559182 | 0.997605693 | 0.929303068 |
| Ipmk | Low | D3YWA2 | Kinase OS=Mus musculus OX=10090 GN=Ipmk PE=1 SV=1 | 0.056 | 1.378 | 2 | 1 | 1 | 1 | 397 | 44.5 | 8.69 | 1.215 | 1.146 | 1.061 | 0.28 | -0.28 | 0.2 | 0.06666667 | -0.24 | 0.08 | 0.506532084 | 0.750586883 | 0.659013434 | 0.999652205 | 0.995559182 | 0.997605693 | 0.985010103 |
| Timm9 | High | Q9WV98 | Mitochondrial import inner membrane translocase subunit Tim9 OS=Mus musculus OX=10090 GN | 0.005 | 3.757 | 18 | 2 | 3 | 2 | 89 | 10.3 | 7.21 | 1.151 | 1.215 | 0.947 | 0.2 | -0.2 | 0.28 | 0.09333333 | -0.24 | -0.08 | 0.604440357 | 0.608370911 | 0.787550321 | 0.999652205 | 0.995559182 | 0.997605693 | 0.992535481 |
| | Medium | D3Z0Q9 | Solute carrier family 9 (sodium/hydrogen exchanger), member 6 (Fragment) OS=Mus musculus C | 0.035 | 1.783 | 2 | 1 | 1 | 1 | 313 | 34.6 | 5.57 | 1.121 | 1.253 | 0.894 | 0.16 | -0.16 | 0.33 | 0.11 | -0.245 | -0.16 | 0.743821428 | 0.547180473 | 0.533910655 | 0.999652205 | 0.995559182 | 0.997605693 | 0.96694628 |
| Ndufb9 | High | Q9CQJ8 | NADH dehydrogenase [ubiquinone] 1 beta subcomplex subunit 9 OS=Mus musculus OX=10090 GN | 0 | 14.013 | 24 | 3 | 4 | 3 | 179 | 22 | 7.8 | 1.13 | 1.238 | 0.913 | 0.18 | -0.18 | 0.31 | 0.10333333 | -0.245 | -0.13 | 0.684631808 | 0.569410827 | 0.617362737 | 0.999652205 | 0.995559182 | 0.997605693 | 0.97826833 |
| Nhp2l1; Sr | High | Q9D0T1 | NHP2-like protein 1 OS=Mus musculus OX=10090 GN=Snu13 PE=1 SV=4 | 0 | 10.419 | 22 | 2 | 2 | 2 | 128 | 14.2 | 8.46 | 1.198 | 1.173 | 1.021 | 0.26 | -0.26 | 0.23 | 0.07666667 | -0.245 | 0.03 | 0.493840909 | 0.67520822 | 0.881134437 | 0.999652205 | 0.995559182 | 0.997605693 | 0.992535481 |
| Lrrc47 | High | Q505F5 | Leucine-rich repeat-containing protein 47 OS=Mus musculus OX=10090 GN=Lrrc47 PE=1 SV=1 | 0 | 51.77 | 26 | 12 | 15 | 12 | 581 | 63.6 | 8.1 | 1.184 | 1.188 | 0.996 | 0.24 | -0.24 | 0.25 | 0.08333333 | -0.245 | -0.01 | 0.579837915 | 0.657206209 | 0.946934597 | 0.999652205 | 0.995559182 | 0.997605693 | 0.992535481 |
| 4930506M | High | Q8K2Q9 | Shootin-1 OS=Mus musculus OX=10090 GN=Shtn1 PE=1 SV=1 | 0.001 | 5.409 | 3 | 2 | 2 | 2 | 631 | 71.3 | 5.44 | 1.077 | 1.314 | 0.819 | 0.11 | -0.11 | 0.39 | 0.13 | -0.25 | -0.29 | 0.845386344 | 0.450896171 | 0.288010359 | 0.999652205 | 0.995559182 | 0.997605693 | 0.887894721 |
| Rtn4 | High | Q99P72 | Reticulon-4 OS=Mus musculus OX=10090 GN=Rtn4 PE=1 SV=2 | 0 | 91.724 | 18 | 14 | 26 | 14 | 1162 | 126.5 | 4.54 | 1.134 | 1.246 | 0.91 | 0.18 | -0.18 | 0.32 | 0.10666667 | -0.25 | -0.14 | 0.670592226 | 0.555689203 | 0.610253442 | 0.999652205 | 0.995559182 | 0.997605693 | 0.97826833 |
| Guca1b | High | Q8VBV8 | Guanylyl cyclase-activating protein 2 OS=Mus musculus OX=10090 GN=Guca1b PE=1 SV=3 | 0 | 128.177 | 62 | 12 | 45 | 12 | 201 | 23.5 | 5.01 | 1.138 | 1.238 | 0.919 | 0.19 | -0.19 | 0.31 | 0.10333333 | -0.25 | -0.12 | 0.661261447 | 0.56855095 | 0.656279267 | 0.999652205 | 0.995559182 | 0.997605693 | 0.985010103 |
| Ptrh2 | High | Q8R2Y8 | Peptidyl-tRNA hydrolase 2, mitochondrial OS=Mus musculus OX=10090 GN=Ptrh2 PE=1 SV=1 | 0 | 8.563 | 19 | 2 | 2 | 2 | 181 | 19.5 | 7.42 | 1.226 | 1.16 | 1.057 | 0.29 | -0.29 | 0.21 | 0.07 | -0.25 | 0.08 | 0.452394651 | 0.709643448 | 0.681914228 | 0.999652205 | 0.995559182 | 0.997605693 | 0.991285312 |
| Rps10 | High | P63325 | 40S ribosomal protein S10 OS=Mus musculus OX=10090 GN=Rps10 PE=1 SV=1 | 0 | 36.603 | 51 | 10 | 18 | 2 | 165 | 18.9 | 10.15 | 1.203 | 1.172 | 1.027 | 0.27 | -0.27 | 0.23 | 0.07666667 | -0.25 | 0.04 | 0.531133209 | 0.68778994 | 0.78798841 | 0.999652205 | 0.995559182 | 0.997605693 | 0.992535481 |
| Ankfn1 | Medium | Q8CDJ6 | Ankyrin-repeat and fibronectin type III domain-containing 1 OS=Mus musculus OX=10090 GN=Ank | 0.033 | 1.837 | 3 | 1 | 1 | 1 | 403 | 44 | 6.11 | 1.157 | 1.218 | 0.95 | 0.21 | -0.21 | 0.29 | 0.09666667 | -0.25 | -0.07 | 0.617037335 | 0.602993878 | 0.792595394 | 0.999652205 | 0.995559182 | 0.997605693 | 0.992535481 |
| Mme | High | Q61391 | Neprilysin OS=Mus musculus OX=10090 GN=Mme PE=1 SV=3 | 0.001 | 5.689 | 3 | 3 | 3 | 3 | 750 | 85.6 | 5.81 | 1.206 | 1.174 | 1.027 | 0.27 | -0.27 | 0.23 | 0.07666667 | -0.25 | 0.04 | 0.49849601 | 0.681691897 | 0.815812436 | 0.999652205 | 0.995559182 | 0.997605693 | 0.992535481 |
| Tmem63b | High | Q3TWI9 | CSC1-like protein 2 OS=Mus musculus OX=10090 GN=Tmem63b PE=1 SV=1 | 0 | 97.157 | 19 | 18 | 47 | 4 | 832 | 94.7 | 7.49 | 1.179 | 1.196 | 0.986 | 0.24 | -0.24 | 0.26 | 0.08666667 | -0.25 | -0.02 | 0.576891541 | 0.642864263 | 0.993556034 | 0.999652205 | 0.995559182 | 0.997605693 | 0.999917596 |

| | | | | | | | | | | | | | | | | | | | | | | | | | | | | |
|---|---|---|---|---|---|---|---|---|---|---|---|---|---|---|---|---|---|---|---|---|---|---|---|---|---|---|---|---|
| Tubb4b | High | P68372 | Tubulin beta-4B chain OS=Mus musculus OX=10090 GN=Tubb4b PE=1 SV=1 | 0 | 421.098 | 73 | 25 | 216 | 1 | 445 | 49.8 | 4.89 | 1.115 | 1.273 | 0.876 | 0.16 | -0.16 | 0.35 | 0.11666667 | -0.255 | -0.19 | 0.712897671 | 0.512949533 | 0.449554223 | 0.999652205 | 0.995559182 | 0.997605693 | 0.959151647 |
| Dynll1 | High | P63168 | Dynein light chain 1, cytoplasmic OS=Mus musculus OX=10090 GN=Dynll1 PE=1 SV=1 | 0 | 41.208 | 57 | 6 | 18 | 3 | 89 | 10.4 | 7.4 | 1.224 | 1.162 | 1.053 | 0.29 | -0.29 | 0.22 | 0.07333333 | -0.255 | 0.07 | 0.494876999 | 0.708001074 | 0.664185939 | 0.999652205 | 0.995559182 | 0.997605693 | 0.986647506 |
| Usp7 | High | E9PXY8 | Ubiquitin carboxyl-terminal hydrolase 7 OS=Mus musculus OX=10090 GN=Usp7 PE=1 SV=1 | 0 | 50.685 | 15 | 15 | 17 | 15 | 1143 | 132.8 | 5.66 | 1.16 | 1.23 | 0.943 | 0.21 | -0.21 | 0.3 | 0.1 | -0.255 | -0.08 | 0.648240802 | 0.582988564 | 0.715732108 | 0.999652205 | 0.995559182 | 0.997605693 | 0.992535481 |
| Eif3b | High | Q8JZQ9 | Eukaryotic translation initiation factor 3 subunit B OS=Mus musculus OX=10090 GN=Eif3b PE=1 S | 0 | 9.162 | 6 | 3 | 3 | 3 | 803 | 91.3 | 5.02 | 1.219 | 1.168 | 1.044 | 0.29 | -0.29 | 0.22 | 0.07333333 | -0.255 | 0.06 | 0.487626831 | 0.696457197 | 0.797215402 | 0.999652205 | 0.995559182 | 0.997605693 | 0.992535481 |
| | Medium | D3Z670 | Complex I-B9 OS=Mus musculus OX=10090 GN=Ndufa3 PE=1 SV=1 | 0.035 | 1.789 | 7 | 1 | 2 | 1 | 100 | 10.9 | 8.76 | 1.163 | 1.226 | 0.949 | 0.22 | -0.22 | 0.29 | 0.09666667 | -0.255 | -0.08 | 0.598552135 | 0.590201799 | 0.811565713 | 0.999652205 | 0.995559182 | 0.997605693 | 0.992535481 |
| Hint2 | High | Q9D0S9 | Histidine triad nucleotide-binding protein 2, mitochondrial OS=Mus musculus OX=10090 GN=Hint | 0 | 13.016 | 27 | 3 | 4 | 3 | 163 | 17.3 | 9.82 | 1.17 | 1.216 | 0.962 | 0.23 | -0.23 | 0.28 | 0.09333333 | -0.255 | -0.06 | 0.589497109 | 0.60658158 | 0.84840075 | 0.999652205 | 0.995559182 | 0.997605693 | 0.992535481 |
| Mtx1 | High | P47802 | Metaxin-1 OS=Mus musculus OX=10090 GN=Mtx1 PE=1 SV=1 | 0 | 14.095 | 11 | 3 | 3 | 3 | 317 | 35.6 | 6.18 | 1.206 | 1.182 | 1.021 | 0.27 | -0.27 | 0.24 | 0.08 | -0.255 | 0.03 | 0.543281859 | 0.677788581 | 0.84840527 | 0.999652205 | 0.995559182 | 0.997605693 | 0.992535481 |
| Pcnp | High | Q3TM89 | PEST proteolytic signal-containing nuclear protein OS=Mus musculus OX=10090 GN=Pcnp PE=1 S | 0 | 9.292 | 17 | 2 | 2 | 2 | 147 | 15.6 | 6.81 | 1.119 | 1.285 | 0.87 | 0.16 | -0.16 | 0.36 | 0.12 | -0.26 | -0.2 | 0.713634606 | 0.494382785 | 0.453948927 | 0.999652205 | 0.995559182 | 0.997605693 | 0.959744396 |
| G3bp2 | High | P97379 | Ras GTPase-activating protein-binding protein 2 OS=Mus musculus OX=10090 GN=G3bp2 PE=1 S | 0 | 24.726 | 17 | 7 | 9 | 7 | 482 | 54.1 | 5.62 | 1.137 | 1.261 | 0.902 | 0.19 | -0.19 | 0.33 | 0.11 | -0.26 | -0.15 | 0.689386518 | 0.531791824 | 0.522043467 | 0.999652205 | 0.995559182 | 0.997605693 | 0.964378027 |
| | High | A0A0G2JD | ATP-binding cassette sub-family D member 3 OS=Mus musculus OX=10090 GN=Abcd3 PE=1 SV=1 | 0 | 14.202 | 9 | 4 | 4 | 4 | 549 | 62.7 | 9.13 | 1.139 | 1.256 | 0.907 | 0.19 | -0.19 | 0.33 | 0.11 | -0.26 | -0.14 | 0.599526276 | 0.539955144 | 0.586867033 | 0.999652205 | 0.995559182 | 0.997605693 | 0.973095093 |
| Araf | High | Q8CAD1 | Ras-binding protein DA-Raf OS=Mus musculus OX=10090 GN=Araf PE=1 SV=1 | 0 | 10.544 | 25 | 4 | 4 | 1 | 186 | 20.9 | 8.47 | 1.151 | 1.252 | 0.919 | 0.2 | -0.2 | 0.32 | 0.10666667 | -0.26 | -0.12 | 0.641695185 | 0.532613416 | 0.639845124 | 0.999652205 | 0.995559182 | 0.997605693 | 0.983766261 |
| | High | E9Q223 | Hemoglobin, beta adult s chain (Fragment) OS=Mus musculus OX=10090 GN=Hbb-bs PE=1 SV=1 | 0 | 14.072 | 37 | 4 | 5 | 4 | 103 | 11.1 | 6.37 | 1.157 | 1.242 | 0.932 | 0.21 | -0.21 | 0.31 | 0.10333333 | -0.26 | -0.1 | 0.645083868 | 0.563087915 | 0.696879599 | 0.999652205 | 0.995559182 | 0.997605693 | 0.991285312 |
| 0610010F | High | Q68FF0 | Uncharacterized protein KIAA1841 OS=Mus musculus OX=10090 GN=Kiaa1841 PE=1 SV=2 | 0.001 | 4.451 | 1 | 1 | 1 | 1 | 718 | 81.9 | 6.62 | 1.229 | 1.169 | 1.052 | 0.3 | -0.3 | 0.22 | 0.07333333 | -0.26 | 0.07 | 0.493594444 | 0.715969451 | 0.730409056 | 0.999652205 | 0.995559182 | 0.997605693 | 0.992535481 |
| Dhrs1 | High | Q99L04 | Dehydrogenase/reductase SDR family member 1 OS=Mus musculus OX=10090 GN=Dhrs1 PE=1 S | 0 | 13.058 | 11 | 2 | 2 | 2 | 313 | 34 | 8.35 | 1.178 | 1.214 | 0.971 | 0.24 | -0.24 | 0.28 | 0.09333333 | -0.26 | -0.04 | 0.536802249 | 0.597597972 | 0.893293675 | 0.999652205 | 0.995559182 | 0.997605693 | 0.992535481 |
| 6330407J2 | High | Q6NZL0 | Protein SOGA3 OS=Mus musculus OX=10090 GN=Soga3 PE=1 SV=2 | 0.006 | 3.375 | 1 | 1 | 2 | 1 | 945 | 103.4 | 6.14 | 1.117 | 1.291 | 0.865 | 0.16 | -0.16 | 0.37 | 0.12333333 | -0.265 | -0.21 | 0.761619565 | 0.482780693 | 0.376187464 | 0.999652205 | 0.995559182 | 0.997605693 | 0.92034959 |
| Pgam5 | High | Q8BX10 | Serine/threonine-protein phosphatase PGAM5, mitochondrial OS=Mus musculus OX=10090 GN=P | 0.01 | 2.751 | 3 | 1 | 1 | 1 | 288 | 32 | 9.04 | 1.139 | 1.265 | 0.901 | 0.19 | -0.19 | 0.34 | 0.11333333 | -0.265 | -0.15 | 0.674772739 | 0.519335494 | 0.59942997 | 0.999652205 | 0.995559182 | 0.997605693 | 0.977451587 |
| Gldc | Medium | Q91W43 | Glycine dehydrogenase (decarboxylating), mitochondrial OS=Mus musculus OX=10090 GN=Gldc P | 0.041 | 1.684 | 1 | 1 | 1 | 1 | 1025 | 113.2 | 7.56 | 1.261 | 1.15 | 1.097 | 0.33 | -0.33 | 0.2 | 0.06666667 | -0.265 | 0.13 | 0.448838336 | 0.750885866 | 0.600969504 | 0.999652205 | 0.995559182 | 0.997605693 | 0.978211931 |
| | High | F6RQ26 | Neurotrimin (Fragment) OS=Mus musculus OX=10090 GN=Ntm PE=1 SV=1 | 0 | 30.474 | 26 | 4 | 10 | 1 | 191 | 21 | 6.77 | 1.219 | 1.18 | 1.033 | 0.29 | -0.29 | 0.24 | 0.08 | -0.265 | 0.05 | 0.518433726 | 0.674324833 | 0.802907479 | 0.999652205 | 0.995559182 | 0.997605693 | 0.992535481 |
| Prkd3 | High | Q8K1Y2 | Serine/threonine-protein kinase D3 OS=Mus musculus OX=10090 GN=Prkd3 PE=1 SV=1 | 0.001 | 5.249 | 2 | 3 | 3 | 1 | 889 | 100 | 6.81 | 1.218 | 1.191 | 1.022 | 0.28 | -0.28 | 0.25 | 0.08333333 | -0.265 | 0.03 | 0.517607182 | 0.673209244 | 0.845134411 | 0.999652205 | 0.995559182 | 0.997605693 | 0.992535481 |
| Sash1 | Low | P59808 | SAM and SH3 domain-containing protein 1 OS=Mus musculus OX=10090 GN=Sash1 PE=1 SV=1 | 0.051 | 1.477 | 1 | 1 | 1 | 1 | 1230 | 135.5 | 6.2 | 1.213 | 1.189 | 1.02 | 0.28 | -0.28 | 0.25 | 0.08333333 | -0.265 | 0.03 | 0.504610095 | 0.650479076 | 0.914657068 | 0.999652205 | 0.995559182 | 0.997605693 | 0.992535481 |
| Pak1 | High | G5E884 | Non-specific serine/threonine protein kinase OS=Mus musculus OX=10090 GN=Pak1 PE=1 SV=1 | 0 | 13.769 | 7 | 4 | 7 | 1 | 544 | 60.6 | 5.86 | 1.194 | 1.209 | 0.987 | 0.26 | -0.26 | 0.27 | 0.09 | -0.265 | -0.02 | 0.531092186 | 0.624942958 | 0.991613563 | 0.999652205 | 0.995559182 | 0.997605693 | 0.999783808 |
| Slc9a3r1 | High | P70441 | Na(+)/H(+) exchange regulatory cofactor NHE-RF1 OS=Mus musculus OX=10090 GN=Slc9a3r1 PE= | 0 | 249.622 | 72 | 28 | 133 | 28 | 355 | 38.6 | 5.9 | 1.119 | 1.304 | 0.858 | 0.16 | -0.16 | 0.38 | 0.12666667 | -0.27 | -0.22 | 0.702573663 | 0.466315522 | 0.375859741 | 0.999652205 | 0.995559182 | 0.997605693 | 0.92034959 |
| Ktn1 | High | A0A087W | Kinectin OS=Mus musculus OX=10090 GN=Ktn1 PE=1 SV=1 | 0 | 23.439 | 6 | 7 | 8 | 7 | 1252 | 143.7 | 6.13 | 1.123 | 1.293 | 0.869 | 0.17 | -0.17 | 0.37 | 0.12333333 | -0.27 | -0.2 | 0.682533415 | 0.483182014 | 0.400754018 | 0.999652205 | 0.995559182 | 0.997605693 | 0.93262326 |
| Letm2 | Medium | Q7TNU7 | LETM1 domain-containing protein LETM2, mitochondrial OS=Mus musculus OX=10090 GN=Letm2 | 0.01 | 2.662 | 2 | 1 | 1 | 1 | 480 | 54.4 | 9.63 | 1.133 | 1.283 | 0.883 | 0.18 | -0.18 | 0.36 | 0.12 | -0.27 | -0.18 | 0.670297221 | 0.480893876 | 0.489541998 | 0.999652205 | 0.995559182 | 0.997605693 | 0.964378027 |
| C3 | High | P01027 | Complement C3 OS=Mus musculus OX=10090 GN=C3 PE=1 SV=3 | 0 | 7.91 | 1 | 2 | 3 | 2 | 1663 | 186.4 | 6.73 | 1.157 | 1.254 | 0.922 | 0.21 | -0.21 | 0.33 | 0.11 | -0.27 | -0.12 | 0.628604233 | 0.535862443 | 0.655811184 | 0.999652205 | 0.995559182 | 0.997605693 | 0.985010103 |
| Mgat2 | Low | Q921V5 | Alpha-1,6-mannosyl-glycoprotein 2-beta-N-acetylglucosaminyltransferase OS=Mus musculus OX= | 0.052 | 1.453 | 2 | 1 | 1 | 1 | 442 | 51 | 8.59 | 1.236 | 1.17 | 1.056 | 0.31 | -0.31 | 0.23 | 0.07666667 | -0.27 | 0.08 | 0.498365446 | 0.697513323 | 0.675229683 | 0.999652205 | 0.995559182 | 0.997605693 | 0.990250803 |
| B630019K | Low | Q7TNS5 | RIKEN cDNA B630019K06 gene OS=Mus musculus OX=10090 GN=B630019K06Rik PE=1 SV=1 | 0.051 | 1.495 | 6 | 1 | 1 | 1 | 333 | 34.8 | 6.65 | 1.25 | 1.164 | 1.074 | 0.32 | -0.32 | 0.22 | 0.07333333 | -0.27 | 0.1 | 0.436495437 | 0.702121156 | 0.690606077 | 0.999652205 | 0.995559182 | 0.997605693 | 0.991285312 |
| Cars | High | Q9ER72 | Cysteine--tRNA ligase, cytoplasmic OS=Mus musculus OX=10090 GN=Cars1 PE=1 SV=2 | 0 | 12.404 | 5 | 3 | 3 | 3 | 831 | 94.8 | 6.76 | 1.22 | 1.189 | 1.026 | 0.29 | -0.29 | 0.25 | 0.08333333 | -0.27 | 0.04 | 0.497023099 | 0.659626181 | 0.803739887 | 0.999652205 | 0.995559182 | 0.997605693 | 0.992535481 |
| Slc24a1 | High | Q91WD8 | Solute carrier family 24 (Sodium/potassium/calcium exchanger), member 1 OS=Mus musculus O | 0 | 391.639 | 32 | 35 | 309 | 34 | 1130 | 124.6 | 4.65 | 1.214 | 1.201 | 1.011 | 0.28 | -0.28 | 0.26 | 0.08666667 | -0.27 | 0.02 | 0.511639801 | 0.634878863 | 0.862662275 | 0.999652205 | 0.995559182 | 0.997605693 | 0.992535481 |
| Ddx1 | High | Q91VR5 | ATP-dependent RNA helicase DDX1 OS=Mus musculus OX=10090 GN=Ddx1 PE=1 SV=1 | 0 | 58.118 | 21 | 13 | 22 | 13 | 740 | 82.4 | 7.21 | 1.21 | 1.198 | 1.01 | 0.28 | -0.28 | 0.26 | 0.08666667 | -0.27 | 0.01 | 0.519181193 | 0.640106612 | 0.868156068 | 0.999652205 | 0.995559182 | 0.997605693 | 0.992535481 |
| Gpd2 | High | Q64521 | Glycerol-3-phosphate dehydrogenase, mitochondrial OS=Mus musculus OX=10090 GN=Gpd2 PE= | 0.005 | 3.71 | 2 | 2 | 2 | 2 | 727 | 80.9 | 6.61 | 1.189 | 1.227 | 0.969 | 0.25 | -0.25 | 0.29 | 0.09666667 | -0.27 | -0.05 | 0.59588336 | 0.585608843 | 0.883990838 | 0.999652205 | 0.995559182 | 0.997605693 | 0.992535481 |
| Zfp37 | Low | Q6P3Z4 | Zinc finger protein 37 OS=Mus musculus OX=10090 GN=Zfp37 PE=1 SV=1 | 0.051 | 1.476 | 1 | 1 | 1 | 1 | 590 | 66.7 | 8.75 | 1.228 | 1.188 | 1.034 | 0.3 | -0.3 | 0.25 | 0.08333333 | -0.275 | 0.05 | 0.476933217 | 0.660355786 | 0.827987761 | 0.999652205 | 0.995559182 | 0.997605693 | 0.992535481 |
| Tmem27 | High | Q9ESG4 | Collectrin OS=Mus musculus OX=10090 GN=Cltrn PE=1 SV=1 | 0 | 43.402 | 36 | 6 | 14 | 6 | 222 | 25.1 | 5.85 | 1.201 | 1.22 | 0.985 | 0.26 | -0.26 | 0.29 | 0.09666667 | -0.275 | -0.02 | 0.56984199 | 0.600725863 | 0.901235698 | 0.999652205 | 0.995559182 | 0.997605693 | 0.992535481 |
| Atad3a | High | Q925I1 | ATPase family AAA domain-containing protein 3 OS=Mus musculus OX=10090 GN=Atad3 PE=1 S | 0 | 12.451 | 6 | 4 | 4 | 4 | 591 | 66.7 | 9.29 | 1.196 | 1.224 | 0.977 | 0.26 | -0.26 | 0.29 | 0.09666667 | -0.275 | -0.03 | 0.542677142 | 0.593251607 | 0.921875769 | 0.999652205 | 0.995559182 | 0.997605693 | 0.992535481 |
| | High | E9Q3G5 | ADP-ribosylation factor interacting protein 1 OS=Mus musculus OX=10090 GN=Arfip1 PE=1 SV=1 | 0.001 | 6.064 | 5 | 2 | 2 | 2 | 366 | 40.8 | 6.92 | 1.166 | 1.264 | 0.922 | 0.22 | -0.22 | 0.34 | 0.11333333 | -0.28 | -0.12 | 0.636021411 | 0.52969534 | 0.668188939 | 0.999652205 | 0.995559182 | 0.997605693 | 0.987489386 |
| Casz1 | Low | B1AS46 | Zinc finger protein castor homolog 1 OS=Mus musculus OX=10090 GN=Casz1 PE=1 SV=1 | 0.055 | 1.383 | 1 | 1 | 1 | 1 | 1762 | 191.2 | 7.14 | 1.203 | 1.227 | 0.981 | 0.27 | -0.27 | 0.29 | 0.09666667 | -0.28 | -0.03 | 0.530947396 | 0.588305896 | 0.941052167 | 0.999652205 | 0.995559182 | 0.997605693 | 0.992535481 |
| Tomm70a | High | Q9CZW5 | Mitochondrial import receptor subunit TOM70 OS=Mus musculus OX=10090 GN=Tomm70 PE=1 S | 0 | 56.606 | 29 | 18 | 20 | 18 | 611 | 67.5 | 7.53 | 1.168 | 1.275 | 0.916 | 0.22 | -0.22 | 0.35 | 0.11666667 | -0.285 | -0.13 | 0.598706443 | 0.509350408 | 0.642125835 | 0.999652205 | 0.995559182 | 0.997605693 | 0.985010103 |
| Minos1 | Medium | Q7TNS2 | MICOS complex subunit Mic10 OS=Mus musculus OX=10090 GN=Micos10 PE=1 SV=1 | 0.011 | 2.445 | 11 | 1 | 1 | 1 | 76 | 8.6 | 8.94 | 1.14 | 1.3 | 0.877 | 0.19 | -0.19 | 0.38 | 0.12666667 | -0.285 | -0.19 | 0.633869066 | 0.472181896 | 0.44802334 | 0.999652205 | 0.995559182 | 0.997605693 | 0.958608248 |
| | High | D3Z396 | Neurotrimin OS=Mus musculus OX=10090 GN=Ntm PE=1 SV=1 | 0 | 66.459 | 30 | 8 | 26 | 2 | 316 | 34.9 | 6.96 | 1.145 | 1.291 | 0.886 | 0.2 | -0.2 | 0.37 | 0.12333333 | -0.285 | -0.17 | 0.646641544 | 0.484717558 | 0.498818089 | 0.999652205 | 0.995559182 | 0.997605693 | 0.964378027 |
| | High | E9PWC4 | Sodium- and chloride-dependent glycine transporter 1 OS=Mus musculus OX=10090 GN=Slc6a9 P | 0.001 | 6.943 | 10 | 1 | 3 | 1 | 126 | 13.9 | 9.72 | 1.176 | 1.262 | 0.932 | 0.23 | -0.23 | 0.34 | 0.11333333 | -0.285 | -0.1 | 0.574491111 | 0.530826732 | 0.638010567 | 0.999652205 | 0.995559182 | 0.997605693 | 0.983597794 |
| Pde8a | High | O88502 | High affinity cAMP-specific and IBMX-insensitive 3',5'-cyclic phosphodiesterase 8A OS=Mus musc | 0 | 198.355 | 44 | 28 | 59 | 26 | 823 | 93.1 | 5.85 | 1.252 | 1.187 | 1.054 | 0.32 | -0.32 | 0.25 | 0.08333333 | -0.285 | 0.08 | 0.448130005 | 0.659915893 | 0.658460266 | 0.999652205 | 0.995559182 | 0.997605693 | 0.985010103 |
| | High | J3QMG3 | Voltage-dependent anion-selective channel protein 3 OS=Mus musculus OX=10090 GN=Vdac3 PE= | 0 | 102.503 | 42 | 13 | 33 | 12 | 284 | 30.8 | 8.79 | 1.198 | 1.238 | 0.968 | 0.26 | -0.26 | 0.31 | 0.10333333 | -0.285 | -0.05 | 0.540980324 | 0.568776788 | 0.91027322 | 0.999652205 | 0.995559182 | 0.997605693 | 0.992535481 |
| Ebpl | High | Q9D0P0 | Emopamil-binding protein-like OS=Mus musculus OX=10090 GN=Ebpl PE=2 SV=1 | 0.005 | 3.558 | 4 | 1 | 2 | 1 | 206 | 23.3 | 6.89 | 1.184 | 1.263 | 0.937 | 0.24 | -0.24 | 0.34 | 0.11333333 | -0.29 | -0.09 | 0.565034953 | 0.527744461 | 0.720232068 | 0.999652205 | 0.995559182 | 0.997605693 | 0.992535481 |
| Grpel1 | High | Q99LP6 | GrpE protein homolog 1, mitochondrial OS=Mus musculus OX=10090 GN=Grpel1 PE=1 SV=1 | 0 | 18.771 | 23 | 4 | 5 | 4 | 217 | 24.3 | 8.38 | 1.246 | 1.198 | 1.039 | 0.32 | -0.32 | 0.26 | 0.08666667 | -0.29 | 0.06 | 0.453082243 | 0.638822433 | 0.783214089 | 0.999652205 | 0.995559182 | 0.997605693 | 0.992535481 |
| Aldh6a1 | High | Q9EQ20 | Methylmalonate-semialdehyde dehydrogenase [acylating], mitochondrial OS=Mus musculus OX= | 0 | 16.153 | 13 | 5 | 6 | 5 | 535 | 57.9 | 8.07 | 1.214 | 1.235 | 0.983 | 0.28 | -0.28 | 0.3 | 0.1 | -0.29 | -0.02 | 0.495248499 | 0.574290626 | 0.953810958 | 0.999652205 | 0.995559182 | 0.997605693 | 0.992763698 |
| Dopey1 | Low | Q8BL99 | Protein dopey-1 OS=Mus musculus OX=10090 GN=Dop1a PE=1 SV=2 | 0.072 | 1.285 | 0 | 1 | 2 | 1 | 2399 | 269.1 | 6.24 | 1.163 | 1.293 | 0.9 | 0.22 | -0.22 | 0.37 | 0.12333333 | -0.295 | -0.15 | 0.615499718 | 0.483125238 | 0.549744714 | 0.999652205 | 0.995559182 | 0.997605693 | 0.96694628 |
| | High | A0A087W | Fas apoptotic inhibitory molecule 1 (Fragment) OS=Mus musculus OX=10090 GN=Faim PE=1 SV= | 0 | 101.919 | 59 | 15 | 49 | 15 | 174 | 19.6 | 5.87 | 1.227 | 1.224 | 1.003 | 0.3 | -0.3 | 0.29 | 0.09666667 | -0.295 | 0 | 0.488528219 | 0.59287713 | 0.906869206 | 0.999652205 | 0.995559182 | 0.997605693 | 0.992535481 |
| Ptn | High | Q3UZI5 | Pleiotrophin OS=Mus musculus OX=10090 GN=Ptn PE=1 SV=1 | 0.002 | 4.165 | 16 | 3 | 3 | 3 | 156 | 17.4 | 9.57 | 1.259 | 1.196 | 1.052 | 0.33 | -0.33 | 0.26 | 0.08666667 | -0.295 | 0.07 | 0.441929765 | 0.649571628 | 0.696308386 | 0.999652205 | 0.995559182 | 0.997605693 | 0.991285312 |
| Mt2 | High | P02798 | Metallothionein-2 OS=Mus musculus OX=10090 GN=Mt2 PE=1 SV=2 | 0 | 19.774 | 34 | 3 | 9 | 3 | 61 | 6.1 | 7.83 | 1.238 | 1.212 | 1.021 | 0.31 | -0.31 | 0.28 | 0.09333333 | -0.295 | 0.03 | 0.492335205 | 0.615150239 | 0.792246572 | 0.999652205 | 0.995559182 | 0.997605693 | 0.992535481 |
| Shmt2 | High | Q9CZN7 | Serine hydroxymethyltransferase, mitochondrial OS=Mus musculus OX=10090 GN=Shmt2 PE=1 S | 0 | 12.228 | 6 | 3 | 4 | 3 | 504 | 55.7 | 8.47 | 1.197 | 1.259 | 0.951 | 0.26 | -0.26 | 0.33 | 0.11 | -0.295 | -0.07 | 0.534588186 | 0.534834855 | 0.795295968 | 0.999652205 | 0.995559182 | 0.997605693 | 0.992535481 |
| | High | E9Q6A9 | Catenin beta-1 OS=Mus musculus OX=10090 GN=Ctnnb1 PE=1 SV=1 | 0 | 39.93 | 20 | 12 | 20 | 1 | 607 | 66.5 | 7.59 | 1.197 | 1.257 | 0.953 | 0.26 | -0.26 | 0.33 | 0.11 | -0.295 | -0.07 | 0.527412004 | 0.540987687 | 0.852159023 | 0.999652205 | 0.995559182 | 0.997605693 | 0.992535481 |
| Gsta4 | High | P24472 | Glutathione S-transferase A4 OS=Mus musculus OX=10090 GN=Gsta4 PE=1 SV=3 | 0 | 30.591 | 36 | 8 | 14 | 8 | 222 | 25.5 | 7.39 | 1.212 | 1.238 | 0.979 | 0.28 | -0.28 | 0.31 | 0.10333333 | -0.295 | -0.03 | 0.462946699 | 0.569478667 | 0.972238747 | 0.999652205 | 0.995559182 | 0.997605693 | 0.996575243 |
| Vps18 | High | Q8R307 | Vacuolar protein sorting-associated protein 18 homolog OS=Mus musculus OX=10090 GN=Vps18 | 0.003 | 3.94 | 1 | 1 | 1 | 1 | 973 | 110.1 | 6.09 | 1.162 | 1.3 | 0.894 | 0.22 | -0.22 | 0.38 | 0.12666667 | -0.3 | -0.16 | 0.619842629 | 0.465645342 | 0.53395207 | 0.999652205 | 0.995559182 | 0.997605693 | 0.96694628 |
| Galm | High | Q8K157 | Galactose mutarotase OS=Mus musculus OX=10090 GN=Galm PE=1 SV=1 | 0 | 17.437 | 16 | 4 | 4 | 4 | 342 | 37.8 | 6.74 | 1.174 | 1.297 | 0.905 | 0.23 | -0.23 | 0.37 | 0.12333333 | -0.3 | -0.14 | 0.581085619 | 0.477030099 | 0.593122636 | 0.999652205 | 0.995559182 | 0.997605693 | 0.976405937 |
| Slc7a10 | High | P63115 | Asc-type amino acid transporter 1 OS=Mus musculus OX=10090 GN=Slc7a10 PE=1 SV=1 | 0 | 15.032 | 10 | 3 | 3 | 3 | 530 | 57.5 | 8.44 | 1.19 | 1.271 | 0.936 | 0.25 | -0.25 | 0.35 | 0.11666667 | -0.3 | -0.09 | 0.535342091 | 0.515790596 | 0.726096587 | 0.999652205 | 0.995559182 | 0.997605693 | 0.992535481 |
| Fbxl20 | High | Q9CZV8 | F-box/LRR-repeat protein 20 OS=Mus musculus OX=10090 GN=Fbxl20 PE=1 SV=3 | 0 | 76.295 | 34 | 14 | 27 | 14 | 436 | 48.4 | 7.49 | 1.234 | 1.232 | 1.001 | 0.3 | -0.3 | 0.3 | 0.1 | -0.3 | 0 | 0.477259664 | 0.578982123 | 0.913087478 | 0.999652205 | 0.995559182 | 0.997605693 | 0.992535481 |
| Rps11 | High | P62281 | 40S ribosomal protein S11 OS=Mus musculus OX=10090 GN=Rps11 PE=1 SV=3 | 0 | 18.8 | 34 | 7 | 10 | 7 | 158 | 18.4 | 10.3 | 1.295 | 1.177 | 1.1 | 0.37 | -0.37 | 0.24 | 0.08 | -0.305 | 0.14 | 0.440372835 | 0.678784103 | 0.567697805 | 0.999652205 | 0.995559182 | 0.997605693 | 0.972910028 |
| Prph2 | High | P15499 | Peripherin-2 OS=Mus musculus OX=10090 GN=Prph2 PE=1 SV=1 | 0 | 318.099 | 61 | 24 | 380 | 24 | 346 | 39.2 | 8.02 | 1.18 | 1.288 | 0.915 | 0.24 | -0.24 | 0.37 | 0.12333333 | -0.305 | -0.13 | 0.576256101 | 0.489187473 | 0.63906666 | 0.999652205 | 0.995559182 | 0.997605693 | 0.983597794 |
| Tab1 | Medium | Q8CF89 | TGF-beta-activated kinase 1 and MAP3K7-binding protein 1 OS=Mus musculus OX=10090 GN=Tab | 0.047 | 1.55 | 2 | 1 | 1 | 1 | 502 | 54.6 | 5.52 | 1.186 | 1.284 | 0.924 | 0.25 | -0.25 | 0.36 | 0.12 | -0.305 | -0.11 | 0.540381977 | 0.483668311 | 0.666196864 | 0.999652205 | 0.995559182 | 0.997605693 | 0.987489386 |
| | High | E9Q3M5 | Anion exchange protein OS=Mus musculus OX=10090 GN=Slc4a5 PE=1 SV=1 | 0 | 65.365 | 18 | 19 | 28 | 19 | 1116 | 124.1 | 8.18 | 1.188 | 1.29 | 0.921 | 0.25 | -0.25 | 0.37 | 0.12333333 | -0.31 | -0.12 | 0.560042845 | 0.486893448 | 0.666556315 | 0.999652205 | 0.995559182 | 0.997605693 | 0.987489386 |
| Fnbp1 | High | A2AQ43 | Formin-binding protein 1 OS=Mus musculus OX=10090 GN=Fnbp1 PE=1 SV=1 | 0.001 | 5.307 | 3 | 2 | 2 | 2 | 554 | 64.6 | 5.24 | 1.207 | 1.273 | 0.948 | 0.27 | -0.27 | 0.35 | 0.11666667 | -0.31 | -0.08 | 0.570160184 | 0.51220053 | 0.766169634 | 0.999652205 | 0.995559182 | 0.997605693 | 0.992535481 |
| Cntnap1 | High | O54991 | Contactin-associated protein 1 OS=Mus musculus OX=10090 GN=Cntnap1 PE=1 SV=2 | 0.001 | 4.581 | 2 | 2 | 2 | 2 | 1385 | 156.2 | 7.06 | 1.214 | 1.274 | 0.953 | 0.28 | -0.28 | 0.35 | 0.11666667 | -0.315 | -0.07 | 0.521286561 | 0.51264027 | 0.812466695 | 0.999652205 | 0.995559182 | 0.997605693 | 0.992535481 |
| Tpp2 | High | Q64514 | Tripeptidyl-peptidase 2 OS=Mus musculus OX=10090 GN=Tpp2 PE=1 SV=3 | 0 | 41.822 | 9 | 10 | 14 | 10 | 1262 | 139.8 | 6.58 | 1.212 | 1.276 | 0.95 | 0.28 | -0.28 | 0.35 | 0.11666667 | -0.315 | -0.07 | 0.484607526 | 0.508161917 | 0.817518222 | 0.999652205 | 0.995559182 | 0.997605693 | 0.992535481 |
| Smg7 | High | Q5RJH6 | Protein SMG7 OS=Mus musculus OX=10090 GN=Smg7 PE=2 SV=1 | 0.005 | 3.515 | 2 | 2 | 2 | 2 | 1138 | 126.8 | 7.99 | 1.215 | 1.272 | 0.955 | 0.28 | -0.28 | 0.35 | 0.11666667 | -0.315 | -0.07 | 0.482625753 | 0.513770799 | 0.819145569 | 0.999652205 | 0.995559182 | 0.997605693 | 0.992535481 |
| Bbs2 | High | Q9CWF6 | Bardet-Biedl syndrome 2 protein homolog OS=Mus musculus OX=10090 GN=Bbs2 PE=1 SV=1 | 0 | 399.217 | 50 | 35 | 216 | 35 | 721 | 79.9 | 5.86 | 1.223 | 1.276 | 0.959 | 0.29 | -0.29 | 0.35 | 0.11666667 | -0.32 | -0.06 | 0.496679048 | 0.508774635 | 0.862905052 | 0.999652205 | 0.995559182 | 0.997605693 | 0.992535481 |
| Rlbp1 | High | Q9Z275 | Retinaldehyde-binding protein 1 OS=Mus musculus OX=10090 GN=Rlbp1 PE=1 SV=3 | 0 | 223.749 | 68 | 18 | 114 | 18 | 317 | 36.4 | 5.06 | 1.191 | 1.315 | 0.906 | 0.25 | -0.25 | 0.39 | 0.13 | -0.32 | -0.14 | 0.553458839 | 0.451110073 | 0.592921027 | 0.999652205 | 0.995559182 | 0.997605693 | 0.976360704 |
| Fkbp2 | High | P45878 | Peptidyl-prolyl cis-trans isomerase FKBP2 OS=Mus musculus OX=10090 GN=Fkbp2 PE=1 SV=1 | 0.001 | 4.264 | 11 | 2 | 2 | 2 | 140 | 15.3 | 8.88 | 1.232 | 1.264 | 0.975 | 0.3 | -0.3 | 0.34 | 0.11333333 | -0.32 | -0.04 | 0.456131726 | 0.526222432 | 0.896103174 | 0.999652205 | 0.995559182 | 0.997605693 | 0.992535481 |
| Swsap1 | High | Q8VCI7 | ATPase SWSAP1 OS=Mus musculus OX=10090 GN=Swsap1 PE=2 SV=2 | 0.008 | 2.915 | 4 | 1 | 1 | 1 | 278 | 29.4 | 5.63 | 1.232 | 1.264 | 0.975 | 0.3 | -0.3 | 0.34 | 0.11333333 | -0.32 | -0.04 | 0.492831231 | 0.527529462 | 0.91840383 | 0.999652205 | 0.995559182 | 0.997605693 | 0.992535481 |
| Sema4a | High | Q62178 | Semaphorin-4A OS=Mus musculus OX=10090 GN=Sema4a PE=1 SV=2 | 0 | 37.586 | 16 | 10 | 12 | 10 | 760 | 83.4 | 7.72 | 1.245 | 1.248 | 0.998 | 0.32 | -0.32 | 0.32 | 0.10666667 | -0.32 | 0 | 0.484872399 | 0.552082034 | 0.958169433 | 0.999652205 | 0.995559182 | 0.997605693 | 0.992902921 |
| Opa1 | High | H7BX01 | Dynamin-like 120 kDa protein, form S1 OS=Mus musculus OX=10090 GN=Opa1 PE=1 SV=1 | 0 | 46.174 | 16 | 14 | 14 | 14 | 978 | 113.2 | 7.33 | 1.2 | 1.308 | 0.917 | 0.26 | -0.26 | 0.39 | 0.13 | -0.325 | -0.12 | 0.557297401 | 0.460405547 | 0.618758188 | 0.999652205 | 0.995559182 | 0.997605693 | 0.97826833 |
| Dynll2 | High | Q9D0M5 | Dynein light chain 2, cytoplasmic OS=Mus musculus OX=10090 GN=Dynll2 PE=1 SV=1 | 0 | 43.912 | 57 | 6 | 13 | 3 | 89 | 10.3 | 7.37 | 1.212 | 1.288 | 0.941 | 0.28 | -0.28 | 0.37 | 0.12333333 | -0.325 | -0.09 | 0.470164518 | 0.489458278 | 0.688299256 | 0.999652205 | 0.995559182 | 0.997605693 | 0.991285312 |
| Wdr41 | High | Q3UDP0 | WD repeat-containing protein 41 OS=Mus musculus OX=10090 GN=Wdr41 PE=1 SV=1 | 0 | 4.903 | 4 | 2 | 3 | 2 | 460 | 51.5 | 5.15 | 1.235 | 1.287 | 0.959 | 0.3 | -0.3 | 0.36 | 0.12 | -0.33 | -0.06 | 0.460810758 | 0.482136749 | 0.843017981 | 0.999652205 | 0.995559182 | 0.997605693 | 0.992535481 |
| Capns1 | Medium | O88456 | Calpain small subunit 1 OS=Mus musculus OX=10090 GN=Capns1 PE=1 SV=1 | 0.04 | 1.686 | 3 | 1 | 1 | 1 | 269 | 28.4 | 5.63 | 1.23 | 1.286 | 0.956 | 0.3 | -0.3 | 0.36 | 0.12 | -0.33 | -0.06 | 0.482561947 | 0.493417063 | 0.880873483 | 0.999652205 | 0.995559182 | 0.997605693 | 0.992535481 |
| Bbs4 | High | Q8C1Z7 | Bardet-Biedl syndrome 4 protein homolog OS=Mus musculus OX=10090 GN=Bbs4 PE=1 SV=1 | 0 | 423.245 | 70 | 39 | 131 | 39 | 520 | 58.2 | 8.1 | 1.265 | 1.247 | 1.014 | 0.34 | -0.34 | 0.32 | 0.10666667 | -0.33 | 0.02 | 0.42658929 | 0.553597117 | 0.847859454 | 0.999652205 | 0.995559182 | 0.997605693 | 0.992535481 |
| Ubxn6 | High | Q99PL6 | UBX domain-containing protein 6 OS=Mus musculus OX=10090 GN=Ubxn6 PE=1 SV=1 | 0 | 17.624 | 14 | 6 | 6 | 6 | 442 | 49.8 | 8.54 | 1.223 | 1.301 | 0.94 | 0.29 | -0.29 | 0.38 | 0.12666667 | -0.335 | -0.09 | 0.538118035 | 0.470706552 | 0.689984294 | 0.999652205 | 0.995559182 | 0.997605693 | 0.991285312 |
| Pitpna | High | P53810 | Phosphatidylinositol transfer protein alpha isoform OS=Mus musculus OX=10090 GN=Pitpna PE=1 | 0 | 88.196 | 66 | 19 | 32 | 17 | 271 | 31.9 | 6.37 | 1.257 | 1.262 | 0.996 | 0.33 | -0.33 | 0.34 | 0.11333333 | -0.335 | -0.01 | 0.439176473 | 0.530000093 | 0.939503784 | 0.999652205 | 0.995559182 | 0.997605693 | 0.992535481 |
| Ivd | High | Q9JHI5 | Isovaleryl-CoA dehydrogenase, mitochondrial OS=Mus musculus OX=10090 GN=Ivd PE=1 SV=1 | 0.001 | 5.836 | 4 | 2 | 2 | 2 | 424 | 46.3 | 8.29 | 1.231 | 1.304 | 0.944 | 0.3 | -0.3 | 0.38 | 0.12666667 | -0.34 | -0.08 | 0.449968824 | 0.465825681 | 0.803391798 | 0.999652205 | 0.995559182 | 0.997605693 | 0.992535481 |
| Rps15 | High | P62843 | 40S ribosomal protein S15 OS=Mus musculus OX=10090 GN=Rps15 PE=1 SV=2 | 0 | 24.141 | 27 | 4 | 10 | 4 | 145 | 17 | 10.39 | 1.292 | 1.262 | 1.024 | 0.37 | -0.37 | 0.34 | 0.11333333 | -0.355 | 0.03 | 0.446539758 | 0.529919191 | 0.898816401 | 0.999652205 | 0.995559182 | 0.997605693 | 0.992535481 |
| Ttyh1 | High | Q9D3A9 | Protein tweety homolog 1 OS=Mus musculus OX=10090 GN=Ttyh1 PE=1 SV=1 | 0.001 | 7.168 | 6 | 2 | 3 | 2 | 450 | 49 | 5.03 | 1.258 | 1.311 | 0.96 | 0.33 | -0.33 | 0.39 | 0.13 | -0.36 | -0.06 | 0.454189916 | 0.451485177 | 0.842506849 | 0.999652205 | 0.995559182 | 0.997605693 | 0.992535481 |
| Fig4 | Medium | Q91WF7 | Polyphosphoinositide phosphatase OS=Mus musculus OX=10090 GN=Fig4 PE=1 SV=1 | 0.011 | 2.423 | 2 | 1 | 1 | 1 | 907 | 103.4 | 6.98 | 0.95 | 0.941 | 1.01 | -0.07 | 0.07 | -0.09 | -0.03 | 0.08 | 0.01 | 0.769795127 | 0.772732829 | 0.949041275 | 0.999652205 | 0.995568238 | 0.997610221 | 0.992535481 |
| Casp6 | Medium | O08738 | Caspase-6 OS=Mus musculus OX=10090 GN=Casp6 PE=1 SV=1 | 0.028 | 1.929 | 3 | 1 | 1 | 1 | 276 | 31.6 | 6.89 | 0.963 | 1.124 | 0.857 | -0.05 | 0.05 | 0.17 | 0.05666667 | -0.06 | -0.22 | 0.814690808 | 0.774464717 | 0.39137044 | 0.999652205 | 0.997341219 | 0.998496712 | 0.93064581 |
| Psmd3 | High | P14685 | 26S proteasome non-ATPase regulatory subunit 3 OS=Mus musculus OX=10090 GN=Psmd3 PE=1 | 0 | 10.581 | 7 | 3 | 3 | 3 | 530 | 60.7 | 8.44 | 1.108 | 0.937 | 1.182 | 0.15 | -0.15 | -0.09 | -0.03 | -0.03 | 0.24 | 0.724090205 | 0.775007958 | 0.289238729 | 0.999652205 | 0.997756384 | 0.998704294 | 0.888636466 |
| Kif5c | High | P28738 | Kinesin heavy chain isoform 5C OS=Mus musculus OX=10090 GN=Kif5c PE=1 SV=3 | 0 | 27.865 | 11 | 9 | 10 | 7 | 956 | 109.2 | 6.19 | 1.06 | 1.129 | 0.939 | 0.08 | -0.08 | 0.17 | 0.05666667 | -0.125 | -0.09 | 0.86925241 | 0.775143001 | 0.708055519 | 0.999652205 | 0.997756384 | 0.998704294 | 0.992507514 |
| Epb4.1l3; I | High | D0VYV6 | Band 4.1-like protein 3 OS=Mus musculus OX=10090 GN=Epb41l3 PE=1 SV=1 | 0 | 144.113 | 35 | 29 | 44 | 5 | 911 | 101.4 | 5.33 | 0.818 | 0.949 | 0.861 | -0.29 | 0.29 | -0.08 | -0.0266667 | 0.185 | -0.22 | 0.500465732 | 0.806150152 | 0.389164691 | 0.999652205 | 0.998402594 | 0.9990274 | 0.93064581 |
| | High | D3Z2H9 | Tropomyosin 3, related sequence 7 OS=Mus musculus OX=10090 GN=Tpm3-rs7 PE=3 SV=1 | 0 | 28.673 | 37 | 10 | 15 | 8 | 248 | 29 | 4.81 | 0.819 | 0.962 | 0.851 | -0.29 | 0.29 | -0.06 | -0.02 | 0.175 | -0.23 | 0.478352174 | 0.837736029 | 0.370813163 | 0.999652205 | 0.998402594 | 0.9990274 | 0.920164724 |
| Vasn | High | Q9CZT5 | Vasorin OS=Mus musculus OX=10090 GN=Vasn PE=2 SV=2 | 0 | 42.523 | 14 | 9 | 13 | 9 | 673 | 72.2 | 7.08 | 0.834 | 0.943 | 0.884 | -0.26 | 0.26 | -0.08 | -0.0266667 | 0.17 | -0.18 | 0.500522512 | 0.791315974 | 0.469211069 | 0.999652205 | 0.998402594 | 0.9990274 | 0.961044326 |
| Cadm3 | High | Q99N28 | Cell adhesion molecule 3 OS=Mus musculus OX=10090 GN=Cadm3 PE=1 SV=1 | 0 | 51.081 | 34 | 9 | 12 | 9 | 396 | 42.9 | 5.8 | 0.808 | 0.979 | 0.826 | -0.31 | 0.31 | -0.03 | -0.01 | 0.17 | -0.28 | 0.433953882 | 0.880229984 | 0.231126073 | 0.999652205 | 0.998402594 | 0.9990274 | 0.819676582 |
| Strap | High | Q9Z1Z2 | Serine-threonine kinase receptor-associated protein OS=Mus musculus OX=10090 GN=Strap PE=1 | 0 | 8.127 | 7 | 2 | 2 | 2 | 350 | 38.4 | 5.12 | 0.843 | 0.938 | 0.898 | -0.25 | 0.25 | -0.09 | -0.03 | 0.17 | -0.15 | 0.439364293 | 0.77797146 | 0.543912554 | 0.999652205 | 0.998402594 | 0.9990274 | 0.96694628 |
| Lin7c | High | O88952 | Protein lin-7 homolog C OS=Mus musculus OX=10090 GN=Lin7c PE=1 SV=2 | 0 | 45.759 | 54 | 8 | 15 | 5 | 197 | 21.8 | 8.43 | 0.845 | 0.941 | 0.897 | -0.24 | 0.24 | -0.09 | -0.03 | 0.165 | -0.16 | 0.527161778 | 0.786543353 | 0.527494661 | 0.999652205 | 0.998402594 | 0.9990274 | 0.965609718 |
| Coro1b | High | Q9WUM3 | Coronin-1B OS=Mus musculus OX=10090 GN=Coro1b PE=1 SV=1 | 0.001 | 5.234 | 11 | 3 | 3 | 3 | 484 | 53.9 | 5.78 | 0.833 | 0.957 | 0.871 | -0.26 | 0.26 | -0.06 | -0.02 | 0.16 | -0.2 | 0.453896702 | 0.824279354 | 0.39287295 | 0.999652205 | 0.998402594 | 0.9990274 | 0.93064581 |
| Snap91 | High | Q61548 | Clathrin coat assembly protein AP180 OS=Mus musculus OX=10090 GN=Snap91 PE=1 SV=1 | 0 | 107.931 | 16 | 15 | 41 | 13 | 901 | 91.8 | 4.88 | 0.814 | 0.993 | 0.82 | -0.3 | 0.3 | -0.01 | -0.0033333 | 0.155 | -0.29 | 0.491616813 | 0.914597891 | 0.238716812 | 0.999652205 | 0.998402594 | 0.9990274 | 0.830051837 |
| Itgb5 | High | G5E8F8 | Integrin beta OS=Mus musculus OX=10090 GN=Itgb5 PE=1 SV=1 | 0.009 | 2.858 | 2 | 2 | 2 | 2 | 816 | 90 | 6.25 | 0.846 | 0.952 | 0.888 | -0.24 | 0.24 | -0.07 | -0.0233333 | 0.155 | -0.17 | 0.47392788 | 0.793460216 | 0.511207181 | 0.999652205 | 0.998402594 | 0.9990274 | 0.964378027 |
| Bsg | High | P18572 | Basigin OS=Mus musculus OX=10090 GN=Bsg PE=1 SV=2 | 0 | 205.321 | 43 | 18 | 127 | 18 | 389 | 42.4 | 5.85 | 0.863 | 0.943 | 0.915 | -0.21 | 0.21 | -0.09 | -0.03 | 0.15 | -0.13 | 0.621560072 | 0.789338975 | 0.638073049 | 0.999652205 | 0.998402594 | 0.9990274 | 0.983597794 |
| Itpa | High | Q9D892 | Inosine triphosphate pyrophosphatase OS=Mus musculus OX=10090 GN=Itpa PE=1 SV=2 | 0.006 | 3.341 | 6 | 1 | 1 | 1 | 198 | 21.9 | 5.87 | 0.861 | 0.951 | 0.906 | -0.22 | 0.22 | -0.07 | -0.0233333 | 0.145 | -0.14 | 0.512586469 | 0.789091466 | 0.608480135 | 0.999652205 | 0.998402594 | 0.9990274 | 0.97826833 |
| Serpinh1 | High | P19324 | Serpin H1 OS=Mus musculus OX=10090 GN=Serpinh1 PE=1 SV=3 | 0 | 12.202 | 7 | 4 | 5 | 4 | 417 | 46.5 | 8.82 | 0.842 | 0.975 | 0.864 | -0.25 | 0.25 | -0.04 | -0.0133333 | 0.145 | -0.21 | 0.475909864 | 0.869448178 | 0.359546992 | 0.999652205 | 0.998402594 | 0.9990274 | 0.915938202 |
| | High | A0A2R8VH | 3-mercaptopyruvate sulfurtransferase (Fragment) OS=Mus musculus OX=10090 GN=Mpst PE=1 S | 0 | 8.743 | 13 | 2 | 3 | 2 | 169 | 19 | 6.52 | 0.856 | 0.957 | 0.894 | -0.22 | 0.22 | -0.06 | -0.02 | 0.14 | -0.16 | 0.441508587 | 0.803043863 | 0.519971328 | 0.999652205 | 0.998402594 | 0.9990274 | 0.964378027 |
| Impg1 | High | Q8R1W8 | Interphotoreceptor matrix proteoglycan 1 OS=Mus musculus OX=10090 GN=Impg1 PE=1 SV=1 | 0 | 81.574 | 27 | 18 | 26 | 18 | 798 | 89.4 | 5.24 | 0.858 | 0.957 | 0.897 | -0.22 | 0.22 | -0.06 | -0.02 | 0.14 | -0.16 | 0.60927321 | 0.824144396 | 0.549634778 | 0.999652205 | 0.998402594 | 0.9990274 | 0.96694628 |
| Ywhae | High | P62259 | 14-3-3 protein epsilon OS=Mus musculus OX=10090 GN=Ywhae PE=1 SV=1 | 0 | 187.874 | 74 | 24 | 78 | 21 | 255 | 29.2 | 4.74 | 0.868 | 0.945 | 0.918 | -0.2 | 0.2 | -0.08 | -0.0266667 | 0.14 | -0.12 | 0.636450198 | 0.796048723 | 0.653781685 | 0.999652205 | 0.998402594 | 0.9990274 | 0.985010103 |
| | High | D3YWF6 | Ubiquitinyl hydrolase 1 OS=Mus musculus OX=10090 GN=Otub1 PE=1 SV=1 | 0 | 33.189 | 28 | 5 | 8 | 5 | 241 | 28 | 5.29 | 0.879 | 0.94 | 0.935 | -0.19 | 0.19 | -0.09 | -0.03 | 0.14 | -0.1 | 0.567250474 | 0.782628113 | 0.657226204 | 0.999652205 | 0.998402594 | 0.9990274 | 0.985010103 |
| Kpnb1 | High | P70168 | Importin subunit beta-1 OS=Mus musculus OX=10090 GN=Kpnb1 PE=1 SV=2 | 0 | 62.716 | 21 | 13 | 19 | 13 | 876 | 97.1 | 4.78 | 0.847 | 0.971 | 0.873 | -0.24 | 0.24 | -0.04 | -0.0133333 | 0.14 | -0.2 | 0.588312171 | 0.859041077 | 0.457625423 | 0.999652205 | 0.998402594 | 0.9990274 | 0.959744396 |
| Gprin3 | High | Q8BWS5 | G protein-regulated inducer of neurite outgrowth 3 OS=Mus musculus OX=10090 GN=Gprin3 PE= | 0 | 8.787 | 3 | 2 | 2 | 2 | 763 | 80.4 | 7.02 | 0.84 | 0.987 | 0.851 | -0.25 | 0.25 | -0.02 | -0.0066667 | 0.135 | -0.23 | 0.464887176 | 0.900093182 | 0.325347597 | 0.999652205 | 0.998402594 | 0.9990274 | 0.905686588 |
| Nmt2 | Medium | O70311 | Glycylpeptide N-tetradecanoyltransferase 2 OS=Mus musculus OX=10090 GN=Nmt2 PE=1 SV=1 | 0.042 | 1.626 | 2 | 1 | 1 | 1 | 529 | 60.4 | 7.8 | 0.825 | 1.008 | 0.819 | -0.28 | 0.28 | 0.01 | 0.00333333 | 0.135 | -0.29 | 0.449448929 | 0.938394204 | 0.337994568 | 0.999652205 | 0.998402594 | 0.9990274 | 0.906329766 |
| Flnb | High | Q80X90 | Filamin-B OS=Mus musculus OX=10090 GN=Flnb PE=1 SV=3 | 0.005 | 3.806 | 1 | 2 | 2 | 1 | 2602 | 277.7 | 5.71 | 0.838 | 0.991 | 0.846 | -0.26 | 0.26 | -0.01 | -0.0033333 | 0.135 | -0.24 | 0.481197921 | 0.885859622 | 0.447260376 | 0.999652205 | 0.998402594 | 0.9990274 | 0.958608248 |
| Gnao1 | High | P18872 | Guanine nucleotide-binding protein G(o) subunit alpha OS=Mus musculus OX=10090 GN=Gnao1 F | 0 | 207.06 | 60 | 21 | 121 | 16 | 354 | 40.1 | 5.53 | 0.86 | 0.968 | 0.888 | -0.22 | 0.22 | -0.05 | -0.0166667 | 0.135 | -0.17 | 0.613055834 | 0.852777469 | 0.506072829 | 0.999652205 | 0.998402594 | 0.9990274 | 0.964378027 |
| Vps29 | High | Q9QZ88 | Vacuolar protein sorting-associated protein 29 OS=Mus musculus OX=10090 GN=Vps29 PE=1 SV= | 0 | 9.79 | 18 | 4 | 4 | 4 | 182 | 20.5 | 6.79 | 0.885 | 0.939 | 0.942 | -0.18 | 0.18 | -0.09 | -0.03 | 0.135 | -0.09 | 0.634414029 | 0.781523759 | 0.690284111 | 0.999652205 | 0.998402594 | 0.9990274 | 0.991285312 |
| Gpr158 | High | Q8C419 | Probable G-protein coupled receptor 158 OS=Mus musculus OX=10090 GN=Gpr158 PE=1 SV=2 | 0 | 14.895 | 2 | 2 | 3 | 2 | 1200 | 134.3 | 8.09 | 0.883 | 0.938 | 0.942 | -0.18 | 0.18 | -0.09 | -0.03 | 0.135 | -0.09 | 0.681968009 | 0.777718375 | 0.754966839 | 0.999652205 | 0.998402594 | 0.9990274 | 0.992535481 |
| Anxa1 | High | P10107 | Annexin A1 OS=Mus musculus OX=10090 GN=Anxa1 PE=1 SV=2 | 0 | 97.705 | 59 | 18 | 35 | 18 | 346 | 38.7 | 7.37 | 0.819 | 1.011 | 0.81 | -0.29 | 0.29 | 0.02 | 0.00666667 | 0.135 | -0.3 | 0.502993566 | 0.957416672 | 0.208770953 | 0.999652205 | 0.998402594 | 0.9990274 | 0.801552719 |
| Anxa6 | High | P14824 | Annexin A6 OS=Mus musculus OX=10090 GN=Anxa6 PE=1 SV=3 | 0 | 184.142 | 63 | 38 | 60 | 37 | 673 | 75.8 | 5.5 | 0.827 | 1.01 | 0.819 | -0.27 | 0.27 | 0.01 | 0.00333333 | 0.13 | -0.29 | 0.524895235 | 0.955154991 | 0.235652432 | 0.999652205 | 0.998402594 | 0.9990274 | 0.82759705 |
| Wfs1 | High | Q3UN10 | Wolframin OS=Mus musculus OX=10090 GN=Wfs1 PE=1 SV=1 | 0 | 8.51 | 2 | 2 | 2 | 2 | 814 | 91.8 | 7.99 | 0.836 | 0.998 | 0.838 | -0.26 | 0.26 | 0 | 0 | 0.13 | -0.26 | 0.433894327 | 0.908880811 | 0.302206801 | 0.999652205 | 0.998402594 | 0.9990274 | 0.900241969 |
| | High | A0A1W2P | Neuronal cell adhesion molecule OS=Mus musculus OX=10090 GN=Nrcam PE=1 SV=1 | 0 | 31.743 | 8 | 7 | 7 | 7 | 1134 | 125.4 | 6.38 | 0.853 | 0.976 | 0.874 | -0.23 | 0.23 | -0.03 | -0.01 | 0.13 | -0.19 | 0.482997959 | 0.873286976 | 0.434038065 | 0.999652205 | 0.998402594 | 0.9990274 | 0.948052758 |
| | High | D3YYS6 | Monoglyceride lipase OS=Mus musculus OX=10090 GN=Mgll PE=1 SV=1 | 0 | 17.506 | 14 | 4 | 4 | 4 | 331 | 36.6 | 7.21 | 0.86 | 0.971 | 0.886 | -0.22 | 0.22 | -0.04 | -0.0133333 | 0.13 | -0.17 | 0.526207014 | 0.860203336 | 0.476051441 | 0.999652205 | 0.998402594 | 0.9990274 | 0.962227984 |
| Dnm1 | High | P39053 | Dynamin-1 OS=Mus musculus OX=10090 GN=Dnm1 PE=1 SV=2 | 0 | 212.603 | 50 | 43 | 75 | 3 | 867 | 97.7 | 7.74 | 0.876 | 0.952 | 0.921 | -0.19 | 0.19 | -0.07 | -0.0233333 | 0.13 | -0.12 | 0.658706467 | 0.811692944 | 0.665967774 | 0.999652205 | 0.998402594 | 0.9990274 | 0.987489386 |
| Lsamp | High | Q3TYE5 | Limbic system-associated membrane protein OS=Mus musculus OX=10090 GN=Lsamp PE=1 SV=1 | 0 | 82.809 | 44 | 13 | 33 | 13 | 338 | 37.3 | 6.68 | 0.881 | 0.947 | 0.93 | -0.18 | 0.18 | -0.08 | -0.0266667 | 0.13 | -0.11 | 0.671045622 | 0.801188183 | 0.710900612 | 0.999652205 | 0.998402594 | 0.9990274 | 0.992535481 |
| Gabrb2 | High | P63137 | Gamma-aminobutyric acid receptor subunit beta-2 OS=Mus musculus OX=10090 GN=Gabrb2 PE= | 0 | 5.002 | 3 | 2 | 2 | 2 | 512 | 59.2 | 9.31 | 0.884 | 0.945 | 0.936 | -0.18 | 0.18 | -0.08 | -0.0266667 | 0.13 | -0.1 | 0.55832056 | 0.794313409 | 0.726485537 | 0.999652205 | 0.998402594 | 0.9990274 | 0.992535481 |
| Anxa3 | High | O35639 | Annexin A3 OS=Mus musculus OX=10090 GN=Anxa3 PE=1 SV=4 | 0 | 8.768 | 11 | 3 | 3 | 3 | 323 | 36.4 | 5.76 | 0.842 | 0.998 | 0.843 | -0.25 | 0.25 | 0 | 0 | 0.125 | -0.25 | 0.4763422 | 0.931864574 | 0.345387735 | 0.999652205 | 0.998402594 | 0.9990274 | 0.907770836 |

| | | | | | | | | | | | | | | | | | | | | | | | | | | | | |
|---|---|---|---|---|---|---|---|---|---|---|---|---|---|---|---|---|---|---|---|---|---|---|---|---|---|---|---|---|
| Slc24a2 | High | Q14BI1 | Slc24a2 protein OS=Mus musculus OX=10090 GN=Slc24a2 PE=1 SV=1 | 0 | 15.342 | 10 | 4 | 5 | 3 | 711 | 78.9 | 6.2 | 0.853 | 0.989 | 0.862 | -0.23 | 0.23 | -0.02 | -0.0066667 | 0.125 | -0.21 | 0.452619459 | 0.905025896 | 0.384789628 | 0.999652205 | 0.998402594 | 0.9990274 | 0.927963632 |
| | High | A0A087W | Neurofascin OS=Mus musculus OX=10090 GN=Nfasc PE=1 SV=1 | 0 | 84.859 | 19 | 18 | 26 | 18 | 1347 | 150.3 | 6.68 | 0.855 | 0.987 | 0.867 | -0.23 | 0.23 | -0.02 | -0.0066667 | 0.125 | -0.21 | 0.609263914 | 0.899844322 | 0.411022385 | 0.999652205 | 0.998402594 | 0.9990274 | 0.936397673 |
| Slc6a1 | High | P31648 | Sodium- and chloride-dependent GABA transporter 1 OS=Mus musculus OX=10090 GN=Slc6a1 PE | 0 | 46.038 | 13 | 5 | 15 | 5 | 599 | 67 | 7.96 | 0.859 | 0.982 | 0.875 | -0.22 | 0.22 | -0.03 | -0.01 | 0.125 | -0.19 | 0.64132843 | 0.888226187 | 0.444871365 | 0.999652205 | 0.998402594 | 0.9990274 | 0.958608248 |
| Pltp | High | A2A5K4 | Phospholipid transfer protein (Fragment) OS=Mus musculus OX=10090 GN=Pltp PE=1 SV=1 | 0.003 | 3.959 | 7 | 1 | 1 | 1 | 165 | 18.8 | 7.5 | 0.871 | 0.968 | 0.9 | -0.2 | 0.2 | -0.05 | -0.0166667 | 0.125 | -0.15 | 0.546042145 | 0.832732395 | 0.574803159 | 0.999652205 | 0.998402594 | 0.9990274 | 0.972910028 |
| Rabl3 | High | Q9D4V7 | Rab-like protein 3 OS=Mus musculus OX=10090 GN=Rabl3 PE=1 SV=1 | 0.009 | 2.79 | 5 | 1 | 2 | 1 | 236 | 26.3 | 7.15 | 0.847 | 0.999 | 0.848 | -0.24 | 0.24 | 0 | 0 | 0.12 | -0.24 | 0.533113692 | 0.929712191 | 0.315382341 | 0.999652205 | 0.998402594 | 0.9990274 | 0.904943958 |
| Ywhab | High | Q9CQV8 | 14-3-3 protein beta/alpha OS=Mus musculus OX=10090 GN=Ywhab PE=1 SV=3 | 0 | 113.657 | 63 | 14 | 54 | 7 | 246 | 28.1 | 4.83 | 0.844 | 0.997 | 0.847 | -0.24 | 0.24 | 0 | 0 | 0.12 | -0.24 | 0.615241841 | 0.924320625 | 0.331206937 | 0.999652205 | 0.998402594 | 0.9990274 | 0.906329766 |
| Ush1g | Medium | Q80T11 | Usher syndrome type-1G protein homolog OS=Mus musculus OX=10090 GN=Ush1g PE=1 SV=1 | 0.01 | 2.733 | 2 | 1 | 2 | 1 | 461 | 51.5 | 6.99 | 0.854 | 1.001 | 0.853 | -0.23 | 0.23 | 0 | 0 | 0.115 | -0.23 | 0.508861146 | 0.934572762 | 0.371444266 | 0.999652205 | 0.998402594 | 0.9990274 | 0.920164724 |
| Vsnl1 | High | P62761 | Visinin-like protein 1 OS=Mus musculus OX=10090 GN=Vsnl1 PE=1 SV=2 | 0 | 28.102 | 41 | 8 | 10 | 5 | 191 | 22.1 | 5.15 | 0.859 | 0.995 | 0.863 | -0.22 | 0.22 | -0.01 | -0.0033333 | 0.115 | -0.21 | 0.604120177 | 0.91974983 | 0.409858207 | 0.999652205 | 0.998402594 | 0.9990274 | 0.936397673 |
| Syt1 | High | P46096 | Synaptotagmin-1 OS=Mus musculus OX=10090 GN=Syt1 PE=1 SV=1 | 0 | 154.145 | 47 | 23 | 64 | 23 | 421 | 47.4 | 8.53 | 0.893 | 0.954 | 0.936 | -0.16 | 0.16 | -0.07 | -0.0233333 | 0.115 | -0.1 | 0.706039163 | 0.818410443 | 0.744203318 | 0.999652205 | 0.998402594 | 0.9990274 | 0.992535481 |
| Cntn1 | High | P12960 | Contactin-1 OS=Mus musculus OX=10090 GN=Cntn1 PE=1 SV=1 | 0 | 88.06 | 24 | 20 | 28 | 20 | 1020 | 113.3 | 6.16 | 0.894 | 0.954 | 0.936 | -0.16 | 0.16 | -0.07 | -0.0233333 | 0.115 | -0.09 | 0.707893166 | 0.819101945 | 0.746403604 | 0.999652205 | 0.998402594 | 0.9990274 | 0.992535481 |
| Ppp2r5c | High | Q60996 | Serine/threonine-protein phosphatase 2A 56 kDa regulatory subunit gamma isoform OS=Mus mu | 0 | 4.799 | 6 | 3 | 3 | 1 | 524 | 60.8 | 6.64 | 0.901 | 0.947 | 0.951 | -0.15 | 0.15 | -0.08 | -0.0266667 | 0.115 | -0.07 | 0.66470493 | 0.799798033 | 0.82743102 | 0.999652205 | 0.998402594 | 0.9990274 | 0.992535481 |
| Fam40b; S | Medium | Q8C9H6 | Striatin-interacting proteins 2 OS=Mus musculus OX=10090 GN=Strip2 PE=1 SV=1 | 0.014 | 2.323 | 1 | 1 | 1 | 1 | 844 | 96.2 | 5.77 | 0.889 | 0.964 | 0.922 | -0.17 | 0.17 | -0.05 | -0.0166667 | 0.11 | -0.12 | 0.580525003 | 0.842493225 | 0.643676998 | 0.999652205 | 0.998402594 | 0.9990274 | 0.985010103 |
| Rab2a | High | P53994 | Ras-related protein Rab-2A OS=Mus musculus OX=10090 GN=Rab2a PE=1 SV=1 | 0 | 147.095 | 66 | 12 | 58 | 6 | 212 | 23.5 | 6.54 | 0.907 | 0.944 | 0.961 | -0.14 | 0.14 | -0.08 | -0.0266667 | 0.11 | -0.06 | 0.74363929 | 0.791684037 | 0.87470327 | 0.999652205 | 0.998402594 | 0.9990274 | 0.992535481 |
| Sms; Gm1 | High | P97355 | Spermine synthase OS=Mus musculus OX=10090 GN=Sms PE=1 SV=1 | 0.006 | 3.162 | 5 | 2 | 2 | 2 | 366 | 41.3 | 5.06 | 0.894 | 0.96 | 0.932 | -0.16 | 0.16 | -0.06 | -0.02 | 0.11 | -0.1 | 0.730324728 | 0.83231855 | 0.691490017 | 0.999652205 | 0.998402594 | 0.9990274 | 0.991285312 |
| | High | A0A1D5RL | Protein EFR3 homolog A OS=Mus musculus OX=10090 GN=Efr3a PE=1 SV=1 | 0 | 15.488 | 4 | 3 | 5 | 3 | 846 | 95.6 | 6.7 | 0.893 | 0.969 | 0.922 | -0.16 | 0.16 | -0.05 | -0.0166667 | 0.105 | -0.12 | 0.595234125 | 0.854202437 | 0.652403965 | 0.999652205 | 0.998402594 | 0.9990274 | 0.985010103 |
| Hcfc1 | Medium | Q61191 | Host cell factor 1 OS=Mus musculus OX=10090 GN=Hcfc1 PE=1 SV=2 | 0.015 | 2.181 | 0 | 1 | 1 | 1 | 2045 | 210.3 | 7.18 | 0.908 | 0.955 | 0.951 | -0.14 | 0.14 | -0.07 | -0.0233333 | 0.105 | -0.07 | 0.603936452 | 0.811232411 | 0.783486863 | 0.999652205 | 0.998402594 | 0.9990274 | 0.992535481 |
| Nub1 | High | P54729 | NEDD8 ultimate buster 1 OS=Mus musculus OX=10090 GN=Nub1 PE=1 SV=2 | 0.009 | 2.761 | 2 | 1 | 1 | 1 | 614 | 70.3 | 5.88 | 0.91 | 0.956 | 0.952 | -0.14 | 0.14 | -0.07 | -0.0233333 | 0.105 | -0.07 | 0.599773029 | 0.803357173 | 0.80823979 | 0.999652205 | 0.998402594 | 0.9990274 | 0.992535481 |
| Cdipt | High | A0A0U1RF | CDP-diacylglycerol--inositol 3-phosphatidyltransferase OS=Mus musculus OX=10090 GN=Cdipt PE | 0.008 | 2.933 | 6 | 1 | 1 | 1 | 170 | 18.9 | 7.03 | 0.857 | 1.005 | 0.853 | -0.22 | 0.22 | 0.01 | 0.00333333 | 0.105 | -0.23 | 0.536895373 | 0.920274022 | 0.469980584 | 0.999652205 | 0.998402594 | 0.9990274 | 0.961044326 |
| Slc2a1 | High | P17809 | Solute carrier family 2, facilitated glucose transporter member 1 OS=Mus musculus OX=10090 GN | 0 | 40.745 | 11 | 8 | 47 | 8 | 492 | 53.9 | 8.87 | 0.876 | 0.986 | 0.889 | -0.19 | 0.19 | -0.02 | -0.0066667 | 0.105 | -0.17 | 0.659465318 | 0.898109886 | 0.50847469 | 0.999652205 | 0.998402594 | 0.9990274 | 0.964378027 |
| | High | F6R6F1 | RalBP1-associated Eps domain-containing protein 1 (Fragment) OS=Mus musculus OX=10090 GN | 0.001 | 5.057 | 1 | 1 | 1 | 1 | 755 | 81.9 | 5.91 | 0.836 | 1.042 | 0.802 | -0.26 | 0.26 | 0.06 | 0.02 | 0.1 | -0.32 | 0.450858264 | 0.981048047 | 0.218734586 | 0.999652205 | 0.998402594 | 0.9990274 | 0.805811317 |
| Hpcal4 | High | Q8BGZ1 | Hippocalcin-like protein 4 OS=Mus musculus OX=10090 GN=Hpcal4 PE=1 SV=3 | 0 | 15.053 | 35 | 6 | 6 | 3 | 191 | 22.2 | 4.89 | 0.897 | 0.976 | 0.919 | -0.16 | 0.16 | -0.04 | -0.0133333 | 0.1 | -0.12 | 0.633163742 | 0.875321841 | 0.658848713 | 0.999652205 | 0.998402594 | 0.9990274 | 0.985010103 |
| Rab11a | High | P62492 | Ras-related protein Rab-11A OS=Mus musculus OX=10090 GN=Rab11a PE=1 SV=3 | 0 | 60.308 | 58 | 11 | 32 | 1 | 216 | 24.4 | 6.57 | 0.925 | 0.942 | 0.982 | -0.11 | 0.11 | -0.09 | -0.03 | 0.1 | -0.03 | 0.734576786 | 0.788642511 | 0.946116635 | 0.999652205 | 0.998402594 | 0.9990274 | 0.992535481 |
| Pi4k2a | High | Q2TBE6 | Phosphatidylinositol 4-kinase type 2-alpha OS=Mus musculus OX=10090 GN=Pi4k2a PE=1 SV=1 | 0 | 21.565 | 13 | 7 | 7 | 7 | 479 | 54.2 | 8.05 | 0.864 | 1.015 | 0.851 | -0.21 | 0.21 | 0.02 | 0.00666667 | 0.095 | -0.23 | 0.59370653 | 0.968211471 | 0.381727197 | 0.999652205 | 0.998402594 | 0.9990274 | 0.923955536 |
| Oxsr1 | High | Q6P9R2 | Serine/threonine-protein kinase OSR1 OS=Mus musculus OX=10090 GN=Oxsr1 PE=1 SV=1 | 0 | 10.354 | 7 | 3 | 3 | 3 | 527 | 58.2 | 6.43 | 0.872 | 1.009 | 0.865 | -0.2 | 0.2 | 0.01 | 0.00333333 | 0.095 | -0.21 | 0.671170584 | 0.953096125 | 0.415196454 | 0.999652205 | 0.998402594 | 0.9990274 | 0.936397673 |
| Sntb2 | High | Q61235 | Beta-2-syntrophin OS=Mus musculus OX=10090 GN=Sntb2 PE=1 SV=2 | 0 | 8.632 | 7 | 4 | 5 | 2 | 520 | 56.3 | 8.69 | 0.879 | 1.002 | 0.877 | -0.19 | 0.19 | 0 | 0 | 0.095 | -0.19 | 0.572725091 | 0.915765229 | 0.464201995 | 0.999652205 | 0.998402594 | 0.9990274 | 0.959744396 |
| | High | E9Q3I4 | Intersectin-1 OS=Mus musculus OX=10090 GN=Itsn1 PE=1 SV=1 | 0 | 13.722 | 5 | 6 | 6 | 6 | 1176 | 133.1 | 7.88 | 0.902 | 0.976 | 0.924 | -0.15 | 0.15 | -0.04 | -0.0133333 | 0.095 | -0.11 | 0.747610942 | 0.872257742 | 0.668581222 | 0.999652205 | 0.998402594 | 0.9990274 | 0.987489386 |
| Ttc7b | High | A0A1Y7VL | Tetratricopeptide repeat protein 7B OS=Mus musculus OX=10090 GN=Ttc7b PE=1 SV=1 | 0 | 10.399 | 5 | 4 | 4 | 4 | 860 | 96 | 6.87 | 0.9 | 0.972 | 0.926 | -0.15 | 0.15 | -0.04 | -0.0133333 | 0.095 | -0.11 | 0.73087915 | 0.862253657 | 0.688366217 | 0.999652205 | 0.998402594 | 0.9990274 | 0.991285312 |
| Rab33b | High | O35963 | Ras-related protein Rab-33B OS=Mus musculus OX=10090 GN=Rab33b PE=1 SV=1 | 0 | 25.084 | 20 | 4 | 8 | 2 | 229 | 25.8 | 7.69 | 0.925 | 0.946 | 0.977 | -0.11 | 0.11 | -0.08 | -0.0266667 | 0.095 | -0.03 | 0.62125319 | 0.776299404 | 0.929791087 | 0.999652205 | 0.998402594 | 0.9990274 | 0.992535481 |
| Cnn3 | High | Q9DAW9 | Calponin-3 OS=Mus musculus OX=10090 GN=Cnn3 PE=1 SV=1 | 0 | 45.772 | 33 | 8 | 14 | 6 | 330 | 36.4 | 5.72 | 0.87 | 1.012 | 0.86 | -0.2 | 0.2 | 0.02 | 0.00666667 | 0.09 | -0.22 | 0.596230755 | 0.960355565 | 0.358858909 | 0.999652205 | 0.998402594 | 0.9990274 | 0.914600864 |
| Sept9 | High | Q80UG5 | Septin-9 OS=Mus musculus OX=10090 GN=Septin9 PE=1 SV=1 | 0 | 14.994 | 8 | 5 | 6 | 5 | 583 | 65.5 | 8.9 | 0.85 | 1.044 | 0.814 | -0.24 | 0.24 | 0.06 | 0.02 | 0.09 | -0.3 | 0.512462908 | 0.963358345 | 0.246027255 | 0.999652205 | 0.998402594 | 0.9990274 | 0.840125423 |
| Tbc1d9b | High | Q5SVR0 | TBC1 domain family member 9B OS=Mus musculus OX=10090 GN=Tbc1d9b PE=1 SV=1 | 0.001 | 4.536 | 1 | 1 | 1 | 1 | 1263 | 141.7 | 5.29 | 0.837 | 1.058 | 0.791 | -0.26 | 0.26 | 0.08 | 0.02666667 | 0.09 | -0.34 | 0.492262173 | 0.951667543 | 0.28749984 | 0.999652205 | 0.998402594 | 0.9990274 | 0.887671587 |
| Ywhag | High | P61982 | 14-3-3 protein gamma OS=Mus musculus OX=10090 GN=Ywhag PE=1 SV=2 | 0 | 99.489 | 71 | 16 | 47 | 10 | 247 | 28.3 | 4.89 | 0.866 | 1.02 | 0.849 | -0.21 | 0.21 | 0.03 | 0.01 | 0.09 | -0.24 | 0.631810547 | 0.980388705 | 0.340887289 | 0.999652205 | 0.998402594 | 0.9990274 | 0.907770836 |
| Oxr1 | High | Q4KMM3 | Oxidation resistance protein 1 OS=Mus musculus OX=10090 GN=Oxr1 PE=1 SV=3 | 0 | 31.903 | 12 | 8 | 11 | 8 | 866 | 95.9 | 5.33 | 0.895 | 0.983 | 0.91 | -0.16 | 0.16 | -0.02 | -0.0066667 | 0.09 | -0.14 | 0.702898781 | 0.890628721 | 0.597712562 | 0.999652205 | 0.998402594 | 0.9990274 | 0.977451587 |
| Rtn1 | High | A3QM89 | Reticulon (Fragment) OS=Mus musculus OX=10090 GN=Rtn1 PE=1 SV=1 | 0 | 28.017 | 23 | 6 | 11 | 2 | 253 | 28.3 | 9.92 | 0.899 | 0.98 | 0.918 | -0.15 | 0.15 | -0.03 | -0.01 | 0.09 | -0.12 | 0.707862305 | 0.882453369 | 0.634509551 | 0.999652205 | 0.998402594 | 0.9990274 | 0.983562471 |
| Stx1b | High | P61264 | Syntaxin-1B OS=Mus musculus OX=10090 GN=Stx1b PE=1 SV=1 | 0 | 148.065 | 57 | 22 | 55 | 18 | 288 | 33.2 | 5.38 | 0.9 | 0.977 | 0.921 | -0.15 | 0.15 | -0.03 | -0.01 | 0.09 | -0.12 | 0.725096936 | 0.875082029 | 0.667566586 | 0.999652205 | 0.998402594 | 0.9990274 | 0.987489386 |
| Ywhah | High | P68510 | 14-3-3 protein eta OS=Mus musculus OX=10090 GN=Ywhah PE=1 SV=2 | 0 | 72.221 | 55 | 15 | 34 | 10 | 246 | 28.2 | 4.89 | 0.913 | 0.964 | 0.947 | -0.13 | 0.13 | -0.05 | -0.0166667 | 0.09 | -0.08 | 0.802068257 | 0.842702814 | 0.798737413 | 0.999652205 | 0.998402594 | 0.9990274 | 0.992535481 |
| Cdkn1b | High | P46414 | Cyclin-dependent kinase inhibitor 1B OS=Mus musculus OX=10090 GN=Cdkn1b PE=1 SV=2 | 0 | 33.411 | 49 | 7 | 9 | 7 | 197 | 22.2 | 7.02 | 0.917 | 0.956 | 0.959 | -0.12 | 0.12 | -0.06 | -0.02 | 0.09 | -0.06 | 0.744871737 | 0.823572303 | 0.830094078 | 0.999652205 | 0.998402594 | 0.9990274 | 0.992535481 |
| | High | E9PUL6 | Sodium/potassium/calcium exchanger 4 OS=Mus musculus OX=10090 GN=Slc24a4 PE=1 SV=1 | 0 | 17.836 | 10 | 4 | 11 | 4 | 564 | 62.2 | 7.9 | 0.924 | 0.952 | 0.971 | -0.11 | 0.11 | -0.07 | -0.0233333 | 0.09 | -0.04 | 0.725650299 | 0.813147184 | 0.839066756 | 0.999652205 | 0.998402594 | 0.9990274 | 0.992535481 |
| Dpp6 | High | Q80VM5 | Dipeptidyl aminopeptidase-like protein 6 OS=Mus musculus OX=10090 GN=Dpp6 PE=1 SV=1 | 0 | 14.775 | 9 | 7 | 7 | 6 | 803 | 91.2 | 6.48 | 0.871 | 1.021 | 0.853 | -0.2 | 0.2 | 0.03 | 0.01 | 0.085 | -0.23 | 0.559511427 | 0.982690134 | 0.318088802 | 0.999652205 | 0.998402594 | 0.9990274 | 0.904943958 |
| Ywhaz | High | P63101 | 14-3-3 protein zeta/delta OS=Mus musculus OX=10090 GN=Ywhaz PE=1 SV=1 | 0 | 156.368 | 71 | 19 | 77 | 13 | 245 | 27.8 | 4.79 | 0.869 | 1.022 | 0.85 | -0.2 | 0.2 | 0.03 | 0.01 | 0.085 | -0.23 | 0.638282049 | 0.984745574 | 0.343892473 | 0.999652205 | 0.998402594 | 0.9990274 | 0.907770836 |
| Prkca | High | Q4VA93 | Protein kinase C OS=Mus musculus OX=10090 GN=Prkca PE=1 SV=1 | 0 | 147.229 | 40 | 25 | 54 | 16 | 672 | 76.8 | 7.05 | 0.872 | 1.021 | 0.854 | -0.2 | 0.2 | 0.03 | 0.01 | 0.085 | -0.23 | 0.645910865 | 0.982313685 | 0.357732432 | 0.999652205 | 0.998402594 | 0.9990274 | 0.914600864 |
| Cct6b | High | Q61390 | T-complex protein 1 subunit zeta-2 OS=Mus musculus OX=10090 GN=Cct6b PE=1 SV=4 | 0 | 19.175 | 11 | 5 | 6 | 1 | 531 | 58.1 | 7.36 | 0.909 | 0.98 | 0.928 | -0.14 | 0.14 | -0.03 | -0.01 | 0.085 | -0.11 | 0.699884 | 0.882187802 | 0.705751791 | 0.999652205 | 0.998402594 | 0.9990274 | 0.992485323 |
| Anxa5 | High | P48036 | Annexin A5 OS=Mus musculus OX=10090 GN=Anxa5 PE=1 SV=1 | 0 | 96.12 | 70 | 23 | 50 | 22 | 319 | 35.7 | 4.96 | 0.804 | 1.102 | 0.73 | -0.31 | 0.31 | 0.14 | 0.04666667 | 0.085 | -0.45 | 0.466031365 | 0.8318464 | 0.053778662 | 0.999652205 | 0.998402594 | 0.9990274 | 0.444089776 |
| | High | A0A1L1SU | Thy-1 antigen (Fragment) OS=Mus musculus OX=10090 GN=Thy1 PE=1 SV=1 | 0 | 34.59 | 41 | 4 | 10 | 4 | 125 | 14 | 9.22 | 0.884 | 1.01 | 0.876 | -0.18 | 0.18 | 0.01 | 0.00333333 | 0.085 | -0.19 | 0.60731298 | 0.955217989 | 0.395006612 | 0.999652205 | 0.998402594 | 0.9990274 | 0.93262326 |
| Spr | High | Q91XH5 | Sepiapterin reductase OS=Mus musculus OX=10090 GN=Spr PE=1 SV=1 | 0 | 33.23 | 36 | 7 | 7 | 7 | 262 | 27.9 | 6.21 | 0.921 | 0.968 | 0.952 | -0.12 | 0.12 | -0.05 | -0.0166667 | 0.085 | -0.07 | 0.705343821 | 0.852465541 | 0.741007034 | 0.999652205 | 0.998402594 | 0.9990274 | 0.992535481 |
| Uchl1 | High | Q9R0P9 | Ubiquitin carboxyl-terminal hydrolase isozyme L1 OS=Mus musculus OX=10090 GN=Uchl1 PE=1 S | 0 | 32.736 | 34 | 6 | 9 | 6 | 223 | 24.8 | 5.24 | 0.921 | 0.966 | 0.954 | -0.12 | 0.12 | -0.05 | -0.0166667 | 0.085 | -0.07 | 0.711417757 | 0.847528135 | 0.752812803 | 0.999652205 | 0.998402594 | 0.9990274 | 0.992535481 |
| Scn2b | Medium | Q56A07 | Sodium channel subunit beta-2 OS=Mus musculus OX=10090 GN=Scn2b PE=1 SV=1 | 0.038 | 1.721 | 4 | 1 | 1 | 1 | 215 | 24.2 | 6.54 | 0.921 | 0.968 | 0.952 | -0.12 | 0.12 | -0.05 | -0.0166667 | 0.085 | -0.07 | 0.684953875 | 0.852622291 | 0.81126443 | 0.999652205 | 0.998402594 | 0.9990274 | 0.992535481 |
| 1110057K( | High | Q8BVA5 | Lipid droplet-associated hydrolase OS=Mus musculus OX=10090 GN=Ldah PE=1 SV=1 | 0 | 11.183 | 11 | 3 | 4 | 3 | 326 | 37.3 | 8.28 | 0.946 | 0.94 | 1.006 | -0.08 | 0.08 | -0.09 | -0.03 | 0.085 | 0.01 | 0.848191798 | 0.782447788 | 0.936883723 | 0.999652205 | 0.998402594 | 0.9990274 | 0.992535481 |
| | High | A0A668KN | Amphiphysin OS=Mus musculus OX=10090 GN=Amph PE=1 SV=1 | 0 | 90.379 | 24 | 17 | 31 | 15 | 820 | 89.8 | 4.59 | 0.859 | 1.043 | 0.824 | -0.22 | 0.22 | 0.06 | 0.02 | 0.08 | -0.28 | 0.639228707 | 0.966447652 | 0.252360545 | 0.999652205 | 0.998402594 | 0.9990274 | 0.851275262 |
| Abhd16a | High | Q9Z1Q2 | Phosphatidylserine lipase ABHD16A OS=Mus musculus OX=10090 GN=Abhd16a PE=1 SV=3 | 0 | 13.876 | 9 | 5 | 6 | 5 | 558 | 63 | 8.25 | 0.871 | 1.026 | 0.848 | -0.2 | 0.2 | 0.04 | 0.01333333 | 0.08 | -0.24 | 0.65729951 | 0.995375513 | 0.361782607 | 0.999652205 | 0.998402594 | 0.9990274 | 0.917494008 |
| | High | E9PV14 | Band 4.1-like protein 1 OS=Mus musculus OX=10090 GN=Epb41l1 PE=1 SV=2 | 0 | 33.414 | 6 | 8 | 13 | 7 | 1556 | 169.8 | 5.39 | 0.898 | 1.001 | 0.897 | -0.16 | 0.16 | 0 | 0 | 0.08 | -0.16 | 0.649335461 | 0.933142783 | 0.496286305 | 0.999652205 | 0.998402594 | 0.9990274 | 0.964378027 |
| Btf3l4 | Medium | Q9CQH7 | Transcription factor BTF3 homolog 4 OS=Mus musculus OX=10090 GN=Btf3l4 PE=1 SV=1 | 0.015 | 2.203 | 4 | 1 | 1 | 1 | 158 | 17.3 | 6.35 | 0.888 | 1.009 | 0.88 | -0.17 | 0.17 | 0.01 | 0.00333333 | 0.08 | -0.18 | 0.602913342 | 0.960449135 | 0.507895327 | 0.999652205 | 0.998402594 | 0.9990274 | 0.964378027 |
| Pex19 | High | Q8VCI5 | Peroxisomal biogenesis factor 19 OS=Mus musculus OX=10090 GN=Pex19 PE=1 SV=1 | 0 | 10.514 | 18 | 5 | 5 | 5 | 299 | 32.7 | 4.34 | 0.898 | 0.999 | 0.899 | -0.16 | 0.16 | 0 | 0 | 0.08 | -0.15 | 0.711611946 | 0.928885309 | 0.54411434 | 0.999652205 | 0.998402594 | 0.9990274 | 0.96694628 |
| Spnb3; Spt | High | Q68FG2 | Spectrin beta chain OS=Mus musculus OX=10090 GN=Sptbn2 PE=1 SV=1 | 0 | 54.981 | 6 | 15 | 20 | 6 | 2388 | 270.8 | 5.86 | 0.906 | 0.988 | 0.917 | -0.14 | 0.14 | -0.02 | -0.0066667 | 0.08 | -0.12 | 0.699180919 | 0.902233689 | 0.632410647 | 0.999652205 | 0.998402594 | 0.9990274 | 0.982523274 |
| Ap1b1 | High | Q5SVG5 | AP complex subunit beta OS=Mus musculus OX=10090 GN=Ap1b1 PE=1 SV=1 | 0 | 147.34 | 32 | 28 | 53 | 14 | 916 | 101.1 | 5.22 | 0.907 | 0.987 | 0.918 | -0.14 | 0.14 | -0.02 | -0.0066667 | 0.08 | -0.12 | 0.762669274 | 0.900645954 | 0.653916072 | 0.999652205 | 0.998402594 | 0.9990274 | 0.985010103 |
| Stxbp6 | High | Q3TYA4 | Syntaxin-binding protein 6 OS=Mus musculus OX=10090 GN=Stxbp6 PE=1 SV=1 | 0.001 | 6.687 | 8 | 2 | 2 | 2 | 210 | 23.7 | 9.19 | 0.911 | 0.98 | 0.93 | -0.13 | 0.13 | -0.03 | -0.01 | 0.08 | -0.11 | 0.787521735 | 0.881548404 | 0.698314304 | 0.999652205 | 0.998402594 | 0.9990274 | 0.991285312 |
| Map7d1; M | High | A2AJI0 | MAP7 domain-containing protein 1 OS=Mus musculus OX=10090 GN=Map7d1 PE=1 SV=1 | 0 | 8.277 | 3 | 2 | 2 | 2 | 846 | 93.2 | 10.15 | 0.903 | 0.991 | 0.911 | -0.15 | 0.15 | -0.01 | -0.0033333 | 0.08 | -0.13 | 0.671536015 | 0.885655088 | 0.703902609 | 0.999652205 | 0.998402594 | 0.9990274 | 0.992485323 |
| Map7d2; M | High | A2AG51 | MAP7 domain-containing protein 2 (Fragment) OS=Mus musculus OX=10090 GN=Map7d2 PE=1 S | 0.006 | 3.404 | 5 | 1 | 1 | 1 | 402 | 45.4 | 9.44 | 0.918 | 0.975 | 0.941 | -0.12 | 0.12 | -0.04 | -0.0133333 | 0.08 | -0.09 | 0.601946558 | 0.85756633 | 0.746929573 | 0.999652205 | 0.998402594 | 0.9990274 | 0.992535481 |
| Impa1 | High | Q924B0 | Inositol-1-monophosphatase OS=Mus musculus OX=10090 GN=Impa1 PE=1 SV=1 | 0 | 119.659 | 64 | 18 | 58 | 18 | 277 | 30.4 | 5.19 | 0.937 | 0.952 | 0.984 | -0.09 | 0.09 | -0.07 | -0.0233333 | 0.08 | -0.02 | 0.827983307 | 0.813602423 | 0.994389657 | 0.999652205 | 0.998402594 | 0.9990274 | 0.999917596 |
| Fam63a; M | High | Q76LS9 | Ubiquitin carboxyl-terminal hydrolase MINDY-1 OS=Mus musculus OX=10090 GN=Mindy1 PE=1 S\ | 0 | 35.248 | 21 | 8 | 11 | 7 | 468 | 51.2 | 4.73 | 0.867 | 1.035 | 0.838 | -0.21 | 0.21 | 0.05 | 0.01666667 | 0.08 | -0.25 | 0.544735656 | 0.98469115 | 0.289163498 | 0.999652205 | 0.998402594 | 0.9990274 | 0.888636466 |
| Copg2 | High | Q9QXK3 | Coatomer subunit gamma-2 OS=Mus musculus OX=10090 GN=Copg2 PE=1 SV=1 | 0.001 | 6.143 | 2 | 2 | 2 | 1 | 871 | 97.6 | 5.8 | 0.892 | 1.016 | 0.877 | -0.17 | 0.17 | 0.02 | 0.00666667 | 0.075 | -0.19 | 0.632062122 | 0.971154386 | 0.466164263 | 0.999652205 | 0.998402594 | 0.9990274 | 0.960359242 |
| Snx2 | High | Q9CWK8 | Sorting nexin-2 OS=Mus musculus OX=10090 GN=Snx2 PE=1 SV=2 | 0 | 24.373 | 12 | 6 | 10 | 5 | 519 | 58.4 | 5.12 | 0.908 | 0.99 | 0.917 | -0.14 | 0.14 | -0.01 | -0.0033333 | 0.075 | -0.13 | 0.669472337 | 0.907515136 | 0.57839311 | 0.999652205 | 0.998402594 | 0.9990274 | 0.972910028 |
| Lamtor2 | High | Q9JHS3 | Ragulator complex protein LAMTOR2 OS=Mus musculus OX=10090 GN=Lamtor2 PE=1 SV=1 | 0 | 42.176 | 52 | 5 | 13 | 5 | 125 | 13.5 | 5.4 | 0.907 | 0.995 | 0.912 | -0.14 | 0.14 | -0.01 | -0.0033333 | 0.075 | -0.13 | 0.701035164 | 0.918999174 | 0.588855297 | 0.999652205 | 0.998402594 | 0.9990274 | 0.973095093 |
| Zfp185 | High | G5E8W8 | Zinc finger protein 185 OS=Mus musculus OX=10090 GN=Zfp185 PE=1 SV=1 | 0 | 10.482 | 4 | 1 | 1 | 1 | 486 | 52.8 | 6.34 | 0.861 | 1.052 | 0.818 | -0.22 | 0.22 | 0.07 | 0.02333333 | 0.075 | -0.29 | 0.49732428 | 0.949923969 | 0.293623553 | 0.999652205 | 0.998402594 | 0.9990274 | 0.894267931 |
| | High | E9Q741 | Chloride transport protein 6 OS=Mus musculus OX=10090 GN=Clcn6 PE=1 SV=1 | 0.003 | 3.929 | 1 | 1 | 1 | 1 | 731 | 80.7 | 7.68 | 0.88 | 1.018 | 0.865 | -0.18 | 0.18 | 0.03 | 0.01 | 0.075 | -0.21 | 0.54845228 | 0.968095974 | 0.417268105 | 0.999652205 | 0.998402594 | 0.9990274 | 0.936397673 |
| Scamp4 | High | Q9JKV5 | Secretory carrier-associated membrane protein 4 OS=Mus musculus OX=10090 GN=Scamp4 PE=1 | 0.001 | 7.459 | 8 | 2 | 3 | 2 | 230 | 25.3 | 8.66 | 0.911 | 0.987 | 0.923 | -0.13 | 0.13 | -0.02 | -0.0066667 | 0.075 | -0.12 | 0.671612856 | 0.882443444 | 0.656339254 | 0.999652205 | 0.998402594 | 0.9990274 | 0.985010103 |
| Ppp1cb | High | P62141 | Serine/threonine-protein phosphatase PP1-beta catalytic subunit OS=Mus musculus OX=10090 GN | 0 | 36.738 | 29 | 7 | 10 | 2 | 327 | 37.2 | 6.19 | 0.928 | 0.971 | 0.955 | -0.11 | 0.11 | -0.04 | -0.0133333 | 0.075 | -0.07 | 0.762035664 | 0.86097651 | 0.801928085 | 0.999652205 | 0.998402594 | 0.9990274 | 0.992535481 |
| Adipor1 | High | Q91VH1 | Adiponectin receptor protein 1 OS=Mus musculus OX=10090 GN=Adipor1 PE=1 SV=1 | 0 | 28.39 | 14 | 4 | 7 | 4 | 375 | 42.3 | 7.02 | 0.957 | 0.942 | 1.016 | -0.06 | 0.06 | -0.09 | -0.03 | 0.075 | 0.02 | 0.818159702 | 0.788628011 | 0.956936122 | 0.999652205 | 0.998402594 | 0.9990274 | 0.992902921 |
| Tspan14 | Low | Q8QZY6 | Tetraspanin-14 OS=Mus musculus OX=10090 GN=Tspan14 PE=1 SV=1 | 0.078 | 1.22 | 4 | 1 | 1 | 1 | 270 | 30.7 | 6.51 | 0.817 | 1.105 | 0.739 | -0.29 | 0.29 | 0.14 | 0.04666667 | 0.075 | -0.44 | 0.436911732 | 0.845490952 | 0.162779316 | 0.999652205 | 0.998402594 | 0.9990274 | 0.733684585 |
| Hsp90ab1 | High | P11499 | Heat shock protein HSP 90-beta OS=Mus musculus OX=10090 GN=Hsp90ab1 PE=1 SV=3 | 0 | 235.105 | 48 | 34 | 91 | 16 | 724 | 83.2 | 5.03 | 0.857 | 1.057 | 0.811 | -0.22 | 0.22 | 0.08 | 0.02666667 | 0.07 | -0.3 | 0.605813968 | 0.933360383 | 0.211156223 | 0.999652205 | 0.998402594 | 0.9990274 | 0.804011919 |
| Rpe65 | High | Q91ZQ5 | Retinoid isomerohydrolase OS=Mus musculus OX=10090 GN=Rpe65 PE=1 SV=4 | 0.001 | 4.56 | 2 | 1 | 1 | 1 | 533 | 61 | 6.58 | 0.876 | 1.036 | 0.845 | -0.19 | 0.19 | 0.05 | 0.01666667 | 0.07 | -0.24 | 0.542784174 | 0.975095544 | 0.360171504 | 0.999652205 | 0.998402594 | 0.9990274 | 0.916290479 |
| | High | F2Z488 | Glutamate receptor OS=Mus musculus OX=10090 GN=Gria3 PE=1 SV=1 | 0.001 | 7.123 | 3 | 3 | 4 | 1 | 818 | 92.7 | 7.64 | 0.91 | 1.001 | 0.909 | -0.14 | 0.14 | 0 | 0 | 0.07 | -0.14 | 0.677765511 | 0.934932479 | 0.586377951 | 0.999652205 | 0.998402594 | 0.9990274 | 0.973095093 |
| Actc1 | High | P68033 | Actin, alpha cardiac muscle 1 OS=Mus musculus OX=10090 GN=Actc1 PE=1 SV=1 | 0 | 168.287 | 46 | 16 | 192 | 4 | 377 | 42 | 5.39 | 0.906 | 0.998 | 0.908 | -0.14 | 0.14 | 0 | 0 | 0.07 | -0.14 | 0.756706382 | 0.92794981 | 0.613417302 | 0.999652205 | 0.998402594 | 0.9990274 | 0.97826833 |
| | High | A0A338P6 | Protein HEG homolog 1 OS=Mus musculus OX=10090 GN=Heg1 PE=1 SV=1 | 0 | 17.558 | 3 | 4 | 7 | 4 | 1313 | 138.8 | 6.67 | 0.926 | 0.976 | 0.949 | -0.11 | 0.11 | -0.03 | -0.01 | 0.07 | -0.08 | 0.787516905 | 0.873791206 | 0.785215929 | 0.999652205 | 0.998402594 | 0.9990274 | 0.992535481 |
| F8a | High | Q00558 | 40-kDa huntingtin-associated protein OS=Mus musculus OX=10090 GN=F8a1 PE=1 SV=1 | 0 | 16.388 | 7 | 2 | 3 | 2 | 380 | 40.5 | 6.89 | 0.934 | 0.972 | 0.961 | -0.1 | 0.1 | -0.04 | -0.0133333 | 0.07 | -0.06 | 0.804031324 | 0.861707013 | 0.843692146 | 0.999652205 | 0.998402594 | 0.9990274 | 0.992535481 |
| Dnm2 | High | Q3TCR7 | Dynamin GTPase OS=Mus musculus OX=10090 GN=Dnm2 PE=1 SV=1 | 0 | 57.335 | 19 | 18 | 30 | 2 | 869 | 97.9 | 7.44 | 0.967 | 0.943 | 1.026 | -0.05 | 0.05 | -0.09 | -0.03 | 0.07 | 0.04 | 0.884708655 | 0.788982865 | 0.857252118 | 0.999652205 | 0.998402594 | 0.9990274 | 0.992535481 |
| Sgip1 | High | Q8VD37 | SH3-containing GRB2-like protein 3-interacting protein 1 OS=Mus musculus OX=10090 GN=Sgip1 | 0 | 60.426 | 23 | 14 | 15 | 14 | 806 | 86 | 7.87 | 0.956 | 0.945 | 1.011 | -0.06 | 0.06 | -0.08 | -0.0266667 | 0.07 | 0.02 | 0.853281718 | 0.796547657 | 0.9070694 | 0.999652205 | 0.998402594 | 0.9990274 | 0.992535481 |
| Srgap2 | High | Q91Z67 | SLIT-ROBO Rho GTPase-activating protein 2 OS=Mus musculus OX=10090 GN=Srgap2 PE=1 SV=2 | 0.005 | 3.711 | 1 | 2 | 2 | 2 | 1071 | 120.7 | 6.64 | 0.956 | 0.947 | 1.01 | -0.06 | 0.06 | -0.08 | -0.0266667 | 0.07 | 0.01 | 0.850010022 | 0.800937808 | 0.913040268 | 0.999652205 | 0.998402594 | 0.9990274 | 0.992535481 |
| Rras | High | P10833 | Ras-related protein R-Ras OS=Mus musculus OX=10090 GN=Rras PE=1 SV=1 | 0 | 33.401 | 32 | 7 | 11 | 5 | 218 | 23.7 | 6.79 | 0.95 | 0.949 | 1 | -0.07 | 0.07 | -0.07 | -0.0233333 | 0.07 | 0 | 0.841557572 | 0.806557743 | 0.928534992 | 0.999652205 | 0.998402594 | 0.9990274 | 0.992535481 |
| Pip4k2a | High | O70172 | Phosphatidylinositol 5-phosphate 4-kinase type-2 alpha OS=Mus musculus OX=10090 GN=Pip4k2a | 0 | 18.337 | 15 | 6 | 8 | 3 | 405 | 46.1 | 6.99 | 0.919 | 0.987 | 0.93 | -0.12 | 0.12 | -0.02 | -0.0066667 | 0.07 | -0.1 | 0.791309045 | 0.900447897 | 0.711950903 | 0.999652205 | 0.998402594 | 0.9990274 | 0.992535481 |
| Rab4a | High | P56371 | Ras-related protein Rab-4A OS=Mus musculus OX=10090 GN=Rab4a PE=1 SV=2 | 0 | 11.743 | 17 | 3 | 5 | 1 | 218 | 24.4 | 6.07 | 0.854 | 1.073 | 0.796 | -0.23 | 0.23 | 0.1 | 0.03333333 | 0.065 | -0.33 | 0.430290974 | 0.895955545 | 0.166238857 | 0.999652205 | 0.998402594 | 0.9990274 | 0.737935797 |
| Fam131b | High | Q3TY60 | Protein FAM131B OS=Mus musculus OX=10090 GN=Fam131b PE=1 SV=1 | 0.005 | 3.513 | 4 | 1 | 1 | 1 | 332 | 35.9 | 4.45 | 0.916 | 1.001 | 0.915 | -0.13 | 0.13 | 0 | 0 | 0.065 | -0.13 | 0.664229173 | 0.917293376 | 0.633871231 | 0.999652205 | 0.998402594 | 0.9990274 | 0.983562471 |
| Slc16a8 | Medium | O35308 | Monocarboxylate transporter 3 OS=Mus musculus OX=10090 GN=Slc16a8 PE=2 SV=2 | 0.032 | 1.845 | 1 | 1 | 1 | 1 | 492 | 51.5 | 6.68 | 0.96 | 0.95 | 1.011 | -0.06 | 0.06 | -0.07 | -0.0233333 | 0.065 | 0.02 | 0.797265828 | 0.786372966 | 0.876722744 | 0.999652205 | 0.998402594 | 0.9990274 | 0.992535481 |
| Rab21 | High | P35282 | Ras-related protein Rab-21 OS=Mus musculus OX=10090 GN=Rab21 PE=1 SV=4 | 0 | 67.29 | 55 | 11 | 23 | 11 | 222 | 24.1 | 7.94 | 0.938 | 0.97 | 0.967 | -0.09 | 0.09 | -0.04 | -0.0133333 | 0.065 | -0.05 | 0.885688669 | 0.85885298 | 0.929483118 | 0.999652205 | 0.998402594 | 0.9990274 | 0.992535481 |
| Cngb3 | High | Q9JJZ9 | Cyclic nucleotide-gated cation channel beta-3 OS=Mus musculus OX=10090 GN=Cngb3 PE=1 SV=1 | 0 | 9.893 | 6 | 4 | 7 | 3 | 694 | 79.7 | 9.11 | 0.959 | 0.955 | 1.004 | -0.06 | 0.06 | -0.07 | -0.0233333 | 0.065 | 0.01 | 0.860280404 | 0.820969503 | 0.975308186 | 0.999652205 | 0.998402594 | 0.9990274 | 0.997221123 |
| Ncam2 | High | O35136 | Neural cell adhesion molecule 2 OS=Mus musculus OX=10090 GN=Ncam2 PE=1 SV=1 | 0 | 50.585 | 18 | 15 | 22 | 15 | 837 | 93.1 | 6.07 | 0.889 | 1.034 | 0.86 | -0.17 | 0.17 | 0.05 | 0.01666667 | 0.06 | -0.22 | 0.726116942 | 0.985567767 | 0.382656087 | 0.999652205 | 0.998402594 | 0.9990274 | 0.925205986 |
| Iqgap1 | High | Q9JKF1 | Ras GTPase-activating-like protein IQGAP1 OS=Mus musculus OX=10090 GN=Iqgap1 PE=1 SV=2 | 0 | 42.178 | 9 | 13 | 15 | 10 | 1657 | 188.6 | 6.48 | 0.905 | 1.013 | 0.894 | -0.14 | 0.14 | 0.02 | 0.00666667 | 0.06 | -0.16 | 0.654855475 | 0.963170799 | 0.483537311 | 0.999652205 | 0.998402594 | 0.9990274 | 0.964378027 |
| Atp6v1h | High | Q8BVE3 | V-type proton ATPase subunit H OS=Mus musculus OX=10090 GN=Atp6v1h PE=1 SV=1 | 0 | 111.927 | 41 | 17 | 31 | 17 | 483 | 55.8 | 6.61 | 0.964 | 0.951 | 1.014 | -0.05 | 0.05 | -0.07 | -0.0233333 | 0.06 | 0.02 | 0.904563154 | 0.809334002 | 0.848470623 | 0.999652205 | 0.998402594 | 0.9990274 | 0.992535481 |
| Rab11b | High | P46638 | Ras-related protein Rab-11B OS=Mus musculus OX=10090 GN=Rab11b PE=1 SV=3 | 0 | 73.729 | 60 | 12 | 36 | 2 | 218 | 24.5 | 5.94 | 0.84 | 1.093 | 0.769 | -0.25 | 0.25 | 0.13 | 0.04333333 | 0.06 | -0.38 | 0.5607954 | 0.851595212 | 0.11106071 | 0.999652205 | 0.998402594 | 0.9990274 | 0.632640059 |
| Plekhf2 | High | Q91WB4 | Pleckstrin homology domain-containing family F member 2 OS=Mus musculus OX=10090 GN=Ple | 0.001 | 5.847 | 7 | 2 | 2 | 2 | 249 | 27.7 | 8.21 | 0.893 | 1.027 | 0.87 | -0.16 | 0.16 | 0.04 | 0.01333333 | 0.06 | -0.2 | 0.638954686 | 0.974363577 | 0.51626583 | 0.999652205 | 0.998402594 | 0.9990274 | 0.964378027 |
| Skp1a | High | Q9WTX5 | S-phase kinase-associated protein 1 OS=Mus musculus OX=10090 GN=Skp1 PE=1 SV=3 | 0 | 43.861 | 67 | 8 | 17 | 8 | 163 | 18.7 | 4.54 | 0.925 | 0.996 | 0.929 | -0.11 | 0.11 | -0.01 | -0.0033333 | 0.06 | -0.11 | 0.837693579 | 0.921268043 | 0.708913041 | 0.999652205 | 0.998402594 | 0.9990274 | 0.992507514 |
| | High | A0A0G2JE | RIKEN cDNA 5730480H06 gene OS=Mus musculus OX=10090 GN=5730480H06Rik PE=1 SV=1 | 0.008 | 2.921 | 9 | 1 | 1 | 1 | 93 | 10.1 | 11.55 | 0.926 | 0.993 | 0.932 | -0.11 | 0.11 | -0.01 | -0.0033333 | 0.06 | -0.1 | 0.724517132 | 0.916964459 | 0.728204713 | 0.999652205 | 0.998402594 | 0.9990274 | 0.992535481 |
| L1cam | High | A2AFG7 | Neural cell adhesion molecule L1 OS=Mus musculus OX=10090 GN=L1cam PE=1 SV=1 | 0 | 25.883 | 9 | 8 | 8 | 8 | 1250 | 139.7 | 5.85 | 0.947 | 0.972 | 0.975 | -0.08 | 0.08 | -0.04 | -0.0133333 | 0.06 | -0.04 | 0.797657149 | 0.862497902 | 0.864953133 | 0.999652205 | 0.998402594 | 0.9990274 | 0.992535481 |
| Tnfrsf21 | Low | Q9EPU5 | Tumor necrosis factor receptor superfamily member 21 OS=Mus musculus OX=10090 GN=Tnfrsf2 | 0.057 | 1.351 | 5 | 1 | 1 | 1 | 655 | 71.9 | 7.46 | 0.941 | 0.978 | 0.962 | -0.09 | 0.09 | -0.03 | -0.01 | 0.06 | -0.06 | 0.677159628 | 0.856305304 | 0.866307752 | 0.999652205 | 0.998402594 | 0.9990274 | 0.992535481 |
| Lgals1 | High | P16045 | Galectin-1 OS=Mus musculus OX=10090 GN=Lgals1 PE=1 SV=3 | 0.001 | 6.656 | 21 | 3 | 3 | 3 | 135 | 14.9 | 5.49 | 0.879 | 1.057 | 0.831 | -0.19 | 0.19 | 0.08 | 0.02666667 | 0.055 | -0.27 | 0.585824034 | 0.931953553 | 0.244414171 | 0.999652205 | 0.998402594 | 0.9990274 | 0.838696607 |
| Eif4g1 | High | E9Q9E1 | Eukaryotic translation initiation factor 4 gamma 1 OS=Mus musculus OX=10090 GN=Eif4g1 PE=1 | 0 | 8.684 | 2 | 3 | 3 | 2 | 1593 | 175.2 | 5.38 | 0.922 | 1.004 | 0.918 | -0.12 | 0.12 | 0.01 | 0.00333333 | 0.055 | -0.12 | 0.620222777 | 0.932022722 | 0.619776139 | 0.999652205 | 0.998402594 | 0.9990274 | 0.97826833 |
| Emd | High | O08579 | Emerin OS=Mus musculus OX=10090 GN=Emd PE=1 SV=1 | 0.002 | 4.124 | 4 | 1 | 2 | 1 | 259 | 29.4 | 5.01 | 0.928 | 0.997 | 0.931 | -0.11 | 0.11 | 0 | 0 | 0.055 | -0.1 | 0.733776871 | 0.923275572 | 0.620637585 | 0.999652205 | 0.998402594 | 0.9990274 | 0.97826833 |
| Pgm2l1 | High | Q8CAA7 | Glucose 1,6-bisphosphate synthase OS=Mus musculus OX=10090 GN=Pgm2l1 PE=1 SV=1 | 0 | 17.489 | 8 | 5 | 6 | 5 | 621 | 70.2 | 6.49 | 0.982 | 0.949 | 1.035 | -0.03 | 0.03 | -0.08 | -0.0266667 | 0.055 | 0.05 | 0.934202353 | 0.805136131 | 0.837706615 | 0.999652205 | 0.998402594 | 0.9990274 | 0.992535481 |
| Tsnax | High | Q9QZE7 | Translin-associated protein X OS=Mus musculus OX=10090 GN=Tsnax PE=1 SV=1 | 0.001 | 6.257 | 9 | 3 | 3 | 3 | 290 | 32.9 | 6.55 | 0.944 | 0.978 | 0.966 | -0.08 | 0.08 | -0.03 | -0.01 | 0.055 | -0.05 | 0.795207621 | 0.876592359 | 0.874337642 | 0.999652205 | 0.998402594 | 0.9990274 | 0.992535481 |
| Fam114a2 | High | Q8VE88 | Protein FAM114A2 OS=Mus musculus OX=10090 GN=Fam114a2 PE=1 SV=2 | 0.001 | 7.725 | 7 | 3 | 3 | 3 | 497 | 54 | 4.93 | 0.947 | 0.979 | 0.968 | -0.08 | 0.08 | -0.03 | -0.01 | 0.055 | -0.05 | 0.846349727 | 0.87979872 | 0.875630802 | 0.999652205 | 0.998402594 | 0.9990274 | 0.992535481 |
| Itpk1 | High | Q8BYN3 | Inositol-tetrakisphosphate 1-kinase OS=Mus musculus OX=10090 GN=Itpk1 PE=2 SV=1 | 0.001 | 4.537 | 5 | 2 | 2 | 2 | 419 | 46.1 | 6.29 | 0.959 | 0.968 | 0.991 | -0.06 | 0.06 | -0.05 | -0.0166667 | 0.055 | -0.01 | 0.807250531 | 0.832545423 | 0.96881148 | 0.999652205 | 0.998402594 | 0.9990274 | 0.99525943 |
| Igsf8 | High | G3UYZ1 | Immunoglobulin superfamily member 8 OS=Mus musculus OX=10090 GN=Igsf8 PE=1 SV=1 | 0 | 27.02 | 20 | 7 | 8 | 7 | 548 | 58.1 | 7.91 | 0.964 | 0.956 | 1.008 | -0.05 | 0.05 | -0.06 | -0.02 | 0.055 | 0.01 | 0.846453738 | 0.823703072 | 0.986646034 | 0.999652205 | 0.998402594 | 0.9990274 | 0.998758677 |
| Rab3c | High | P62823 | Ras-related protein Rab-3C OS=Mus musculus OX=10090 GN=Rab3c PE=1 SV=1 | 0 | 52.926 | 52 | 10 | 18 | 6 | 227 | 25.9 | 5.24 | 0.863 | 1.073 | 0.804 | -0.21 | 0.21 | 0.1 | 0.03333333 | 0.055 | -0.32 | 0.63162171 | 0.895555473 | 0.191201668 | 0.999652205 | 0.998402594 | 0.9990274 | 0.783675259 |
| | High | K3W4T3 | V-type proton ATPase subunit a OS=Mus musculus OX=10090 GN=Atp6v0a1 PE=1 SV=1 | 0 | 129.611 | 26 | 21 | 41 | 21 | 838 | 96.4 | 6.84 | 0.904 | 1.031 | 0.877 | -0.15 | 0.15 | 0.04 | 0.01333333 | 0.055 | -0.19 | 0.736982383 | 0.99377873 | 0.455714684 | 0.999652205 | 0.998402594 | 0.9990274 | 0.959744396 |
| Add1 | High | F8WGR0 | Alpha-adducin OS=Mus musculus OX=10090 GN=Add1 PE=1 SV=1 | 0 | 45.858 | 20 | 11 | 15 | 10 | 663 | 73.2 | 6.58 | 0.927 | 1.007 | 0.921 | -0.11 | 0.11 | 0.01 | 0.00333333 | 0.05 | -0.12 | 0.713446844 | 0.949297562 | 0.584868996 | 0.999652205 | 0.998402594 | 0.9990274 | 0.973095093 |
| Gpr37 | High | Q9QY42 | Prosaposin receptor GPR37 OS=Mus musculus OX=10090 GN=Gpr37 PE=2 SV=1 | 0 | 9.455 | 3 | 2 | 2 | 2 | 600 | 66.7 | 8.12 | 0.933 | 1.001 | 0.932 | -0.1 | 0.1 | 0 | 0 | 0.05 | -0.1 | 0.747271603 | 0.92622905 | 0.697123028 | 0.999652205 | 0.998402594 | 0.9990274 | 0.991285312 |
| Taok1 | High | Q5F2E8 | Serine/threonine-protein kinase TAO1 OS=Mus musculus OX=10090 GN=Taok1 PE=1 SV=1 | 0 | 8.039 | 3 | 3 | 3 | 2 | 1001 | 116 | 7.55 | 0.949 | 0.979 | 0.969 | -0.07 | 0.07 | -0.03 | -0.01 | 0.05 | -0.04 | 0.853009927 | 0.880905873 | 0.887665285 | 0.999652205 | 0.998402594 | 0.9990274 | 0.992535481 |
| Crygs | High | O35486 | Gamma-crystallin S OS=Mus musculus OX=10090 GN=Crygs PE=1 SV=3 | 0 | 12.486 | 19 | 3 | 5 | 3 | 178 | 20.8 | 7.28 | 0.884 | 1.056 | 0.837 | -0.18 | 0.18 | 0.08 | 0.02666667 | 0.05 | -0.26 | 0.629255475 | 0.934451138 | 0.254708017 | 0.999652205 | 0.998402594 | 0.9990274 | 0.855690743 |
| Cttn | High | Q60598 | Src substrate cortactin OS=Mus musculus OX=10090 GN=Cttn PE=1 SV=2 | 0 | 50.338 | 24 | 12 | 16 | 12 | 546 | 61.2 | 5.4 | 0.901 | 1.037 | 0.869 | -0.15 | 0.15 | 0.05 | 0.01666667 | 0.05 | -0.2 | 0.779470868 | 0.979823243 | 0.420988341 | 0.999652205 | 0.998402594 | 0.9990274 | 0.938934618 |
| Epn2 | High | J3QNT7 | Epsin-2 OS=Mus musculus OX=10090 GN=Epn2 PE=1 SV=1 | 0.001 | 5.648 | 2 | 1 | 1 | 1 | 640 | 68.3 | 7.46 | 0.903 | 1.032 | 0.875 | -0.15 | 0.15 | 0.05 | 0.01666667 | 0.05 | -0.19 | 0.623090492 | 0.986526256 | 0.472613016 | 0.999652205 | 0.998402594 | 0.9990274 | 0.961211268 |
| Slirp | High | Q9D8T7 | SRA stem-loop-interacting RNA-binding protein, mitochondrial OS=Mus musculus OX=10090 GN= | 0.003 | 3.886 | 11 | 1 | 2 | 1 | 112 | 12.6 | 9.82 | 0.903 | 1.034 | 0.874 | -0.15 | 0.15 | 0.05 | 0.01666667 | 0.05 | -0.19 | 0.672790247 | 0.989634201 | 0.567302281 | 0.999652205 | 0.998402594 | 0.9990274 | 0.972910028 |
| | High | A0A2R8W | Neuronal-specific septin-3 OS=Mus musculus OX=10090 GN=Septin3 PE=1 SV=1 | 0 | 27.626 | 17 | 6 | 10 | 6 | 341 | 39.1 | 8.43 | 0.941 | 0.992 | 0.949 | -0.09 | 0.09 | -0.01 | -0.0033333 | 0.05 | -0.07 | 0.798040797 | 0.910965201 | 0.763607382 | 0.999652205 | 0.998402594 | 0.9990274 | 0.992535481 |
| | High | A0A494BB | Synaptotagmin-7 OS=Mus musculus OX=10090 GN=Syt7 PE=1 SV=1 | 0 | 8.155 | 3 | 2 | 2 | 2 | 643 | 70.5 | 9.55 | 0.871 | 1.082 | 0.805 | -0.2 | 0.2 | 0.11 | 0.03666667 | 0.045 | -0.31 | 0.551093833 | 0.86575709 | 0.222206934 | 0.999652205 | 0.998402594 | 0.9990274 | 0.807447486 |
| Hsdl2 | High | Q2TPA8 | Hydroxysteroid dehydrogenase-like protein 2 OS=Mus musculus OX=10090 GN=Hsdl2 PE=1 SV=1 | 0.006 | 3.391 | 2 | 1 | 1 | 1 | 490 | 54.2 | 6.74 | 0.892 | 1.058 | 0.843 | -0.17 | 0.17 | 0.08 | 0.02666667 | 0.045 | -0.25 | 0.645282215 | 0.935976582 | 0.352694834 | 0.999652205 | 0.998402594 | 0.9990274 | 0.910898173 |
| Tpm1 | High | E9Q455 | Tropomyosin alpha-1 chain OS=Mus musculus OX=10090 GN=Tpm1 PE=1 SV=1 | 0 | 9.348 | 14 | 4 | 6 | 2 | 251 | 28.7 | 4.79 | 0.848 | 1.108 | 0.765 | -0.24 | 0.24 | 0.15 | 0.05 | 0.045 | -0.39 | 0.455767409 | 0.818244277 | 0.116386639 | 0.999652205 | 0.998402594 | 0.9990274 | 0.651277329 |

| | | | | | | | | | | | | | | | | | | | | | | | | | | | | |
|---|---|---|---|---|---|---|---|---|---|---|---|---|---|---|---|---|---|---|---|---|---|---|---|---|---|---|---|---|
| Rph3a | High | P47708 | Rabphilin-3A OS=Mus musculus OX=10090 GN=Rph3a PE=1 SV=2 | 0 | 81.482 | 26 | 13 | 21 | 12 | 681 | 75.4 | 8.27 | 0.892 | 1.047 | 0.852 | -0.16 | 0.16 | 0.07 | 0.02333333 | 0.045 | -0.23 | 0.746049142 | 0.956551408 | 0.35028356 | 0.999652205 | 0.998402594 | 0.9990274 | 0.909466648 |
| Dnajc11 | Medium | Q5U458 | DnaJ homolog subfamily C member 11 OS=Mus musculus OX=10090 GN=Dnajc11 PE=1 SV=2 | 0.033 | 1.84 | 2 | 1 | 1 | 1 | 559 | 63.2 | 8.32 | 0.911 | 1.029 | 0.886 | -0.13 | 0.13 | 0.04 | 0.01333333 | 0.045 | -0.18 | 0.671053384 | 0.981696197 | 0.497340324 | 0.999652205 | 0.998402594 | 0.9990274 | 0.964378027 |
| Mtor | High | Q9JLN9 | Serine/threonine-protein kinase mTOR OS=Mus musculus OX=10090 GN=Mtor PE=1 SV=2 | 0 | 8.125 | 1 | 2 | 2 | 2 | 2549 | 288.6 | 7.17 | 0.92 | 1.018 | 0.904 | -0.12 | 0.12 | 0.03 | 0.01 | 0.045 | -0.15 | 0.700281861 | 0.954643214 | 0.578718783 | 0.999652205 | 0.998402594 | 0.9990274 | 0.972910028 |
| Ahcyl2 | High | Q68FL4 | Putative adenosylhomocysteinase 3 OS=Mus musculus OX=10090 GN=Ahcyl2 PE=1 SV=1 | 0 | 33.46 | 15 | 11 | 12 | 1 | 613 | 66.9 | 7.36 | 0.918 | 1.02 | 0.901 | -0.12 | 0.12 | 0.03 | 0.01 | 0.045 | -0.15 | 0.710916369 | 0.986575982 | 0.598075957 | 0.999652205 | 0.998402594 | 0.9990274 | 0.977451587 |
| Stx1a | High | O35526 | Syntaxin-1A OS=Mus musculus OX=10090 GN=Stx1a PE=1 SV=3 | 0 | 18.62 | 17 | 6 | 10 | 2 | 288 | 33 | 5.24 | 0.938 | 0.999 | 0.939 | -0.09 | 0.09 | 0 | 0 | 0.045 | -0.09 | 0.843729772 | 0.929444193 | 0.738310035 | 0.999652205 | 0.998402594 | 0.9990274 | 0.992535481 |
| Hpcal1 | High | P62748 | Hippocalcin-like protein 1 OS=Mus musculus OX=10090 GN=Hpcal1 PE=1 SV=2 | 0 | 114.766 | 73 | 15 | 71 | 12 | 193 | 22.3 | 5.5 | 0.939 | 1.001 | 0.938 | -0.09 | 0.09 | 0 | 0 | 0.045 | -0.09 | 0.835408204 | 0.933829759 | 0.756742474 | 0.999652205 | 0.998402594 | 0.9990274 | 0.992535481 |
| Capzb | High | A2AMW0 | F-actin-capping protein subunit beta OS=Mus musculus OX=10090 GN=Capzb PE=1 SV=1 | 0 | 41.802 | 44 | 12 | 14 | 12 | 260 | 29.3 | 6.92 | 0.981 | 0.957 | 1.026 | -0.03 | 0.03 | -0.06 | -0.02 | 0.045 | 0.04 | 0.990753835 | 0.824711497 | 0.793705492 | 0.999652205 | 0.998402594 | 0.9990274 | 0.992535481 |
| D3Bwg05 | High | Q7TME0 | 2-lysophosphatidate phosphatase PLPPR4 OS=Mus musculus OX=10090 GN=Plppr4 PE=1 SV=2 | 0.001 | 4.368 | 1 | 1 | 1 | 1 | 766 | 83.2 | 8.84 | 0.978 | 0.959 | 1.02 | -0.03 | 0.03 | -0.06 | -0.02 | 0.045 | 0.03 | 0.884665027 | 0.807546894 | 0.863363349 | 0.999652205 | 0.998402594 | 0.9990274 | 0.992535481 |
| Map6; Mt | High | Q7TSJ2 | Microtubule-associated protein 6 OS=Mus musculus OX=10090 GN=Map6 PE=1 SV=2 | 0 | 192.339 | 48 | 33 | 49 | 33 | 906 | 96.4 | 9.5 | 0.956 | 0.982 | 0.973 | -0.06 | 0.06 | -0.03 | -0.01 | 0.045 | -0.04 | 0.882347167 | 0.888542762 | 0.940017409 | 0.999652205 | 0.998402594 | 0.9990274 | 0.992535481 |
| Ppp3cb | High | E0CZ78 | Serine/threonine-protein phosphatase OS=Mus musculus OX=10090 GN=Ppp3cb PE=1 SV=1 | 0 | 212.806 | 65 | 24 | 113 | 14 | 524 | 59 | 5.91 | 0.913 | 1.038 | 0.88 | -0.13 | 0.13 | 0.05 | 0.01666667 | 0.04 | -0.18 | 0.762978099 | 0.976333068 | 0.467707478 | 0.999652205 | 0.998402594 | 0.9990274 | 0.960951201 |
| Bmp5 | Low | P49003 | Bone morphogenetic protein 5 OS=Mus musculus OX=10090 GN=Bmp5 PE=2 SV=1 | 0.078 | 1.222 | 1 | 1 | 1 | 1 | 452 | 51.5 | 8.95 | 0.995 | 0.953 | 1.045 | -0.01 | 0.01 | -0.07 | -0.0233333 | 0.04 | 0.06 | 0.940576937 | 0.79301461 | 0.775197649 | 0.999652205 | 0.998402594 | 0.9990274 | 0.992535481 |
| Bin1 | High | O08539 | Myc box-dependent-interacting protein 1 OS=Mus musculus OX=10090 GN=Bin1 PE=1 SV=1 | 0 | 96.764 | 41 | 20 | 30 | 18 | 588 | 64.4 | 5.03 | 0.949 | 0.999 | 0.949 | -0.08 | 0.08 | 0 | 0 | 0.04 | -0.08 | 0.861911086 | 0.930394073 | 0.813520411 | 0.999652205 | 0.998402594 | 0.9990274 | 0.992535481 |
| Atp6v1d | High | P57746 | V-type proton ATPase subunit D OS=Mus musculus OX=10090 GN=Atp6v1d PE=1 SV=1 | 0 | 32.178 | 38 | 8 | 12 | 8 | 247 | 28.4 | 9.45 | 0.987 | 0.959 | 1.03 | -0.02 | 0.02 | -0.06 | -0.02 | 0.04 | 0.04 | 0.950768142 | 0.829797328 | 0.814535727 | 0.999652205 | 0.998402594 | 0.9990274 | 0.992535481 |
| Lhfpl2 | High | Q8BGA2 | LHFPL tetraspan subfamily member 2 protein OS=Mus musculus OX=10090 GN=Lhfpl2 PE=1 SV=1 | 0.001 | 5.244 | 8 | 2 | 2 | 2 | 222 | 23.9 | 6.48 | 0.983 | 0.96 | 1.024 | -0.02 | 0.02 | -0.06 | -0.02 | 0.04 | 0.03 | 0.918319131 | 0.834519683 | 0.86106362 | 0.999652205 | 0.998402594 | 0.9990274 | 0.992535481 |
| Elmo2 | Medium | Q8BHL5 | Engulfment and cell motility protein 2 OS=Mus musculus OX=10090 GN=Elmo2 PE=1 SV=1 | 0.041 | 1.679 | 2 | 1 | 1 | 1 | 732 | 83.8 | 5.95 | 0.956 | 0.995 | 0.961 | -0.07 | 0.07 | -0.01 | -0.0033333 | 0.04 | -0.06 | 0.783166249 | 0.918517576 | 0.861900438 | 0.999652205 | 0.998402594 | 0.9990274 | 0.992535481 |
| H2-Ke6 | Medium | P50171 | (3R)-3-hydroxyacyl-CoA dehydrogenase OS=Mus musculus OX=10090 GN=Hsd17b8 PE=1 SV=2 | 0.033 | 1.839 | 5 | 1 | 1 | 1 | 259 | 26.6 | 6.54 | 0.986 | 0.959 | 1.028 | -0.02 | 0.02 | -0.06 | -0.02 | 0.04 | 0.04 | 0.899135814 | 0.833055951 | 0.86371496 | 0.999652205 | 0.998402594 | 0.9990274 | 0.992535481 |
| | High | S4R1W1 | Glyceraldehyde-3-phosphate dehydrogenase OS=Mus musculus OX=10090 GN=Gm3839 PE=1 SV= | 0 | 753.011 | 91 | 32 | 874 | 1 | 333 | 35.8 | 8.03 | 0.972 | 0.971 | 1.001 | -0.04 | 0.04 | -0.04 | -0.0133333 | 0.04 | 0 | 0.926843792 | 0.860429332 | 0.914534247 | 0.999652205 | 0.998402594 | 0.9990274 | 0.992535481 |
| Cp | High | E9PZD8 | Ceruloplasmin OS=Mus musculus OX=10090 GN=Cp PE=1 SV=1 | 0 | 164.554 | 34 | 32 | 57 | 1 | 1086 | 124.1 | 5.88 | 0.971 | 0.973 | 0.998 | -0.04 | 0.04 | -0.04 | -0.0133333 | 0.04 | 0 | 0.888156823 | 0.865240714 | 0.960668024 | 0.999652205 | 0.998402594 | 0.9990274 | 0.99314725 |
| Wasl | High | Q91YD9 | Neural Wiskott-Aldrich syndrome protein OS=Mus musculus OX=10090 GN=Wasl PE=1 SV=1 | 0 | 13.763 | 7 | 3 | 3 | 3 | 501 | 54.2 | 7.93 | 0.969 | 0.981 | 0.988 | -0.05 | 0.05 | -0.03 | -0.01 | 0.04 | -0.02 | 0.814212684 | 0.876318122 | 0.986501861 | 0.999652205 | 0.998402594 | 0.9990274 | 0.998758677 |
| | Medium | E9Q7X7 | Neurexin-2 OS=Mus musculus OX=10090 GN=Nrxn2 PE=1 SV=1 | 0.01 | 2.658 | 0 | 1 | 1 | 1 | 1710 | 184.8 | 5.9 | 0.858 | 1.105 | 0.776 | -0.22 | 0.22 | 0.14 | 0.04666667 | 0.04 | -0.37 | 0.481952849 | 0.825687657 | 0.14158066 | 0.999652205 | 0.998402594 | 0.9990274 | 0.701851981 |
| Mfge8 | High | P21956 | Lactadherin OS=Mus musculus OX=10090 GN=Mfge8 PE=1 SV=3 | 0 | 128.974 | 37 | 17 | 45 | 17 | 463 | 51.2 | 6.52 | 0.864 | 1.096 | 0.788 | -0.21 | 0.21 | 0.13 | 0.04333333 | 0.04 | -0.34 | 0.625555998 | 0.844939383 | 0.152176881 | 0.999652205 | 0.998402594 | 0.9990274 | 0.724325782 |
| Apmap | Medium | Q9D7N9 | Adipocyte plasma membrane-associated protein OS=Mus musculus OX=10090 GN=Apmap PE=1 S | 0.01 | 2.55 | 3 | 1 | 1 | 1 | 415 | 46.4 | 6.32 | 0.858 | 1.1 | 0.781 | -0.22 | 0.22 | 0.14 | 0.04666667 | 0.04 | -0.36 | 0.484458148 | 0.830276303 | 0.170057739 | 0.999652205 | 0.998402594 | 0.9990274 | 0.738844686 |
| | Medium | F6TCV0 | Serine protease HTRA2, mitochondrial (Fragment) OS=Mus musculus OX=10090 GN=Htra2 PE=1 S | 0.016 | 2.136 | 3 | 1 | 1 | 1 | 294 | 31.8 | 8.46 | 0.854 | 1.116 | 0.766 | -0.23 | 0.23 | 0.16 | 0.05333333 | 0.035 | -0.38 | 0.533588598 | 0.823425182 | 0.202586679 | 0.999652205 | 0.998402594 | 0.9990274 | 0.78697088 |
| Cpne3 | High | Q8BT60 | Copine-3 OS=Mus musculus OX=10090 GN=Cpne3 PE=1 SV=2 | 0 | 67.243 | 32 | 15 | 23 | 13 | 533 | 59.5 | 5.78 | 0.915 | 1.045 | 0.876 | -0.13 | 0.13 | 0.06 | 0.02 | 0.035 | -0.19 | 0.767540578 | 0.961779899 | 0.451379975 | 0.999652205 | 0.998402594 | 0.9990274 | 0.959744396 |
| 1810020D | High | Q8R0P4 | Mth938 domain-containing protein OS=Mus musculus OX=10090 GN=Aamdc PE=1 SV=1 | 0 | 11.721 | 29 | 4 | 5 | 4 | 122 | 13.2 | 7.99 | 0.948 | 1.004 | 0.944 | -0.08 | 0.08 | 0.01 | 0.00333333 | 0.035 | -0.08 | 0.726578916 | 0.923828593 | 0.765503855 | 0.999652205 | 0.998402594 | 0.9990274 | 0.992535481 |
| Stk39 | High | Q9Z1W9 | STE20/SPS1-related proline-alanine-rich protein kinase OS=Mus musculus OX=10090 GN=Stk39 P | 0 | 34.12 | 15 | 8 | 9 | 8 | 556 | 60.3 | 6.29 | 1.003 | 0.953 | 1.053 | 0 | 0 | -0.07 | -0.0233333 | 0.035 | 0.07 | 0.96442811 | 0.815335188 | 0.779510646 | 0.999652205 | 0.998402594 | 0.9990274 | 0.992535481 |
| Wdr7 | High | Q920I9 | WD repeat-containing protein 7 OS=Mus musculus OX=10090 GN=Wdr7 PE=1 SV=3 | 0 | 25.749 | 5 | 6 | 8 | 6 | 1489 | 163.3 | 6.9 | 0.989 | 0.969 | 1.021 | -0.02 | 0.02 | -0.05 | -0.0166667 | 0.035 | 0.03 | 0.944102221 | 0.85451331 | 0.823152168 | 0.999652205 | 0.998402594 | 0.9990274 | 0.992535481 |
| Atp6v1f | High | Q9D1K2 | V-type proton ATPase subunit F OS=Mus musculus OX=10090 GN=Atp6v1f PE=1 SV=2 | 0 | 30.145 | 48 | 5 | 7 | 5 | 119 | 13.4 | 5.82 | 0.963 | 0.989 | 0.974 | -0.05 | 0.05 | -0.02 | -0.0066667 | 0.035 | -0.04 | 0.833579819 | 0.903548801 | 0.846265784 | 0.999652205 | 0.998402594 | 0.9990274 | 0.992535481 |
| Hip1 | High | A0A0J9YU | Huntingtin-interacting protein 1 (Fragment) OS=Mus musculus OX=10090 GN=Hip1 PE=1 SV=1 | 0.001 | 5.424 | 4 | 2 | 2 | 1 | 618 | 68.1 | 5 | 0.987 | 0.968 | 1.02 | -0.02 | 0.02 | -0.05 | -0.0166667 | 0.035 | 0.03 | 0.937433887 | 0.857694055 | 0.875155535 | 0.999652205 | 0.998402594 | 0.9990274 | 0.992535481 |
| Limk2 | Medium | Q5NC03 | LIM domain kinase 2 OS=Mus musculus OX=10090 GN=Limk2 PE=1 SV=1 | 0.01 | 2.713 | 1 | 1 | 1 | 1 | 623 | 70.5 | 7.28 | 0.963 | 0.989 | 0.974 | -0.05 | 0.05 | -0.02 | -0.0066667 | 0.035 | -0.04 | 0.830295608 | 0.907828054 | 0.899798599 | 0.999652205 | 0.998402594 | 0.9990274 | 0.992535481 |
| | High | E9Q0J5 | Kinesin-like protein KIF21A OS=Mus musculus OX=10090 GN=Kif21a PE=1 SV=2 | 0 | 15.514 | 3 | 6 | 6 | 6 | 1672 | 186.5 | 6.47 | 0.986 | 0.963 | 1.024 | -0.02 | 0.02 | -0.05 | -0.0166667 | 0.035 | 0.03 | 0.905576871 | 0.841570546 | 0.922105412 | 0.999652205 | 0.998402594 | 0.9990274 | 0.992535481 |
| Rasa3 | Low | Q60790 | Ras GTPase-activating protein 3 OS=Mus musculus OX=10090 GN=Rasa3 PE=1 SV=2 | 0.053 | 1.419 | 1 | 1 | 1 | 1 | 834 | 95.9 | 7.34 | 0.97 | 0.98 | 0.989 | -0.04 | 0.04 | -0.03 | -0.01 | 0.035 | -0.02 | 0.864986089 | 0.878372147 | 0.968454856 | 0.999652205 | 0.998402594 | 0.9990274 | 0.995075385 |
| Cr1l | High | A0A0A6YY | Complement component receptor 1-like protein OS=Mus musculus OX=10090 GN=Cr1l PE=1 SV=1 | 0 | 12.189 | 7 | 3 | 7 | 3 | 429 | 48 | 6.06 | 0.941 | 1.014 | 0.928 | -0.09 | 0.09 | 0.02 | 0.00666667 | 0.035 | -0.11 | 0.756895715 | 0.9648162 | 0.6227562 | 0.999652205 | 0.998402594 | 0.9990274 | 0.97826833 |
| Lgals3 | High | P16110 | Galectin-3 OS=Mus musculus OX=10090 GN=Lgals3 PE=1 SV=3 | 0.001 | 6.045 | 7 | 2 | 2 | 2 | 264 | 27.5 | 8.38 | 0.887 | 1.077 | 0.823 | -0.17 | 0.17 | 0.11 | 0.03666667 | 0.03 | -0.28 | 0.489994048 | 0.890963111 | 0.237248691 | 0.999652205 | 0.998402594 | 0.9990274 | 0.828110564 |
| Ppt1 | High | O88531 | Palmitoyl-protein thioesterase 1 OS=Mus musculus OX=10090 GN=Ppt1 PE=1 SV=2 | 0 | 22.661 | 19 | 6 | 7 | 6 | 306 | 34.5 | 8 | 0.905 | 1.056 | 0.857 | -0.14 | 0.14 | 0.08 | 0.02666667 | 0.03 | -0.22 | 0.664483173 | 0.935515244 | 0.323401374 | 0.999652205 | 0.998402594 | 0.9990274 | 0.905686588 |
| Rab1b | High | Q9D1G1 | Ras-related protein Rab-1B OS=Mus musculus OX=10090 GN=Rab1b PE=1 SV=1 | 0 | 119.285 | 78 | 15 | 52 | 8 | 201 | 22.2 | 5.73 | 0.91 | 1.056 | 0.862 | -0.14 | 0.14 | 0.08 | 0.02666667 | 0.03 | -0.21 | 0.75392559 | 0.935688837 | 0.39188416 | 0.999652205 | 0.998402594 | 0.9990274 | 0.93064581 |
| Nt5c3 | High | Q9D020 | Cytosolic 5'-nucleotidase 3A OS=Mus musculus OX=10090 GN=Nt5c3a PE=1 SV=4 | 0.009 | 2.815 | 3 | 1 | 1 | 1 | 331 | 37.2 | 6.65 | 0.955 | 1.008 | 0.947 | -0.07 | 0.07 | 0.01 | 0.00333333 | 0.03 | -0.08 | 0.781257916 | 0.951677802 | 0.777040186 | 0.999652205 | 0.998402594 | 0.9990274 | 0.992535481 |
| Msra | High | Q9D6Y7 | Mitochondrial peptide methionine sulfoxide reductase OS=Mus musculus OX=10090 GN=Msra PE | 0 | 14.747 | 18 | 4 | 6 | 4 | 233 | 26 | 8.41 | 0.863 | 1.107 | 0.78 | -0.21 | 0.21 | 0.15 | 0.05 | 0.03 | -0.36 | 0.518293656 | 0.821314595 | 0.132319317 | 0.999652205 | 0.998402594 | 0.9990274 | 0.695121227 |
| | High | D3Z3F8 | Spastic paraplegia 20, spartin (Troyer syndrome) homolog (human) OS=Mus musculus OX=10090 | 0 | 42.777 | 22 | 11 | 15 | 11 | 582 | 63 | 5.58 | 0.915 | 1.049 | 0.872 | -0.13 | 0.13 | 0.07 | 0.02333333 | 0.03 | -0.2 | 0.823798586 | 0.951428433 | 0.435635474 | 0.999652205 | 0.998402594 | 0.9990274 | 0.949084077 |
| Ap2a2 | High | P17427 | AP-2 complex subunit alpha-2 OS=Mus musculus OX=10090 GN=Ap2a2 PE=1 SV=2 | 0 | 175.365 | 47 | 38 | 62 | 23 | 938 | 104 | 6.93 | 0.958 | 0.997 | 0.961 | -0.06 | 0.06 | 0 | 0 | 0.03 | -0.06 | 0.887486777 | 0.924599828 | 0.874143073 | 0.999652205 | 0.998402594 | 0.9990274 | 0.992535481 |
| Rab27a | High | Q9ERI2 | Ras-related protein Rab-27A OS=Mus musculus OX=10090 GN=Rab27a PE=1 SV=1 | 0.001 | 4.469 | 5 | 1 | 1 | 1 | 221 | 25 | 5.36 | 0.959 | 0.999 | 0.96 | -0.06 | 0.06 | 0 | 0 | 0.03 | -0.06 | 0.818079235 | 0.911700746 | 0.878075753 | 0.999652205 | 0.998402594 | 0.9990274 | 0.992535481 |
| Brsk1 | High | Q5RJI5 | Serine/threonine-protein kinase BRSK1 OS=Mus musculus OX=10090 GN=Brsk1 PE=1 SV=1 | 0 | 9.325 | 5 | 4 | 4 | 1 | 778 | 85.1 | 9.32 | 0.959 | 1.001 | 0.958 | -0.06 | 0.06 | 0 | 0 | 0.03 | -0.06 | 0.831314943 | 0.909744695 | 0.881052091 | 0.999652205 | 0.998402594 | 0.9990274 | 0.992535481 |
| Cyb5b | High | Q9CQX2 | Cytochrome b5 type B OS=Mus musculus OX=10090 GN=Cyb5b PE=1 SV=1 | 0 | 4.694 | 8 | 1 | 3 | 1 | 146 | 16.3 | 4.89 | 0.979 | 0.981 | 0.998 | -0.03 | 0.03 | -0.03 | -0.01 | 0.03 | 0 | 0.864793037 | 0.872227845 | 0.944241079 | 0.999652205 | 0.998402594 | 0.9990274 | 0.992535481 |
| Utrn | High | E9Q6R7 | Utrophin OS=Mus musculus OX=10090 GN=Utrn PE=1 SV=1 | 0 | 12.421 | 1 | 6 | 6 | 5 | 3430 | 392.5 | 5.33 | 0.972 | 0.986 | 0.986 | -0.04 | 0.04 | -0.02 | -0.0066667 | 0.03 | -0.02 | 0.876915769 | 0.896516991 | 0.991429641 | 0.999652205 | 0.998402594 | 0.9990274 | 0.999783808 |
| Epm2aip1 | High | Q8VEH5 | EPM2A-interacting protein 1 OS=Mus musculus OX=10090 GN=Epm2aip1 PE=1 SV=1 | 0.003 | 3.85 | 2 | 1 | 1 | 1 | 606 | 70.1 | 5.87 | 1.024 | 0.949 | 1.079 | 0.03 | -0.03 | -0.08 | -0.0266667 | 0.025 | 0.11 | 0.962847235 | 0.80661591 | 0.654124857 | 0.999652205 | 0.998402594 | 0.9990274 | 0.985010103 |
| Ddx19a | High | Q61655 | ATP-dependent RNA helicase DDX19A OS=Mus musculus OX=10090 GN=Ddx19a PE=1 SV=2 | 0 | 42.46 | 27 | 12 | 12 | 12 | 478 | 53.9 | 6.67 | 0.998 | 0.966 | 1.034 | 0 | 0 | -0.05 | -0.0166667 | 0.025 | 0.05 | 0.991751276 | 0.846837535 | 0.746163873 | 0.999652205 | 0.998402594 | 0.9990274 | 0.992535481 |
| Tln1 | High | P26039 | Talin-1 OS=Mus musculus OX=10090 GN=Tln1 PE=1 SV=2 | 0 | 51.918 | 6 | 14 | 16 | 8 | 2541 | 269.7 | 6.18 | 1.011 | 0.955 | 1.059 | 0.02 | -0.02 | -0.07 | -0.0233333 | 0.025 | 0.08 | 0.973608134 | 0.819367496 | 0.76518964 | 0.999652205 | 0.998402594 | 0.9990274 | 0.992535481 |
| Atp6v1a | High | P50516 | V-type proton ATPase catalytic subunit A OS=Mus musculus OX=10090 GN=Atp6v1a PE=1 SV=2 | 0 | 246.576 | 68 | 35 | 84 | 35 | 617 | 68.3 | 5.58 | 0.996 | 0.972 | 1.025 | -0.01 | 0.01 | -0.04 | -0.0133333 | 0.025 | 0.04 | 0.99268373 | 0.862129174 | 0.79403499 | 0.999652205 | 0.998402594 | 0.9990274 | 0.992535481 |
| Nif3l1 | High | Q9EQ80 | NIF3-like protein 1 OS=Mus musculus OX=10090 GN=Nif3l1 PE=1 SV=4 | 0 | 10.547 | 12 | 3 | 3 | 3 | 376 | 41.7 | 6.76 | 0.965 | 1 | 0.965 | -0.05 | 0.05 | 0 | 0 | 0.025 | -0.05 | 0.776283259 | 0.921787162 | 0.852806389 | 0.999652205 | 0.998402594 | 0.9990274 | 0.992535481 |
| Snrpa1 | High | P57784 | U2 small nuclear ribonucleoprotein A' OS=Mus musculus OX=10090 GN=Snrpa1 PE=1 SV=2 | 0.006 | 3.125 | 5 | 1 | 1 | 1 | 255 | 28.3 | 8.62 | 0.986 | 0.982 | 1.004 | -0.02 | 0.02 | -0.03 | -0.01 | 0.025 | 0.01 | 0.927004393 | 0.89268863 | 0.948300026 | 0.999652205 | 0.998402594 | 0.9990274 | 0.992535481 |
| Atp1a1 | High | Q8VDN2 | Sodium/potassium-transporting ATPase subunit alpha-1 OS=Mus musculus OX=10090 GN=Atp1a1 | 0 | 775.555 | 53 | 61 | 386 | 37 | 1023 | 112.9 | 5.45 | 0.978 | 0.987 | 0.991 | -0.03 | 0.03 | -0.02 | -0.0066667 | 0.025 | -0.01 | 0.943156889 | 0.899268003 | 0.965583472 | 0.999652205 | 0.998402594 | 0.9990274 | 0.994494219 |
| Dmxl2 | High | Q8BPN8 | DmX-like protein 2 OS=Mus musculus OX=10090 GN=Dmxl2 PE=1 SV=3 | 0 | 36.708 | 4 | 12 | 13 | 12 | 3032 | 338 | 6.42 | 1.01 | 0.962 | 1.049 | 0.01 | -0.01 | -0.06 | -0.02 | 0.025 | 0.07 | 0.97094033 | 0.838494316 | 0.699154956 | 0.999652205 | 0.998402594 | 0.9990274 | 0.991285312 |
| Pitpnm3 | Low | Q3UHE1 | Membrane-associated phosphatidylinositol transfer protein 3 OS=Mus musculus OX=10090 GN=P | 0.051 | 1.478 | 1 | 1 | 1 | 1 | 974 | 106.4 | 7.12 | 0.914 | 1.061 | 0.861 | -0.13 | 0.13 | 0.09 | 0.03 | 0.02 | -0.22 | 0.651686946 | 0.932003142 | 0.481610915 | 0.999652205 | 0.998402594 | 0.9990274 | 0.964378027 |
| Cnga3 | High | A0A0A6YX | Cyclic nucleotide-gated cation channel alpha-3 OS=Mus musculus OX=10090 GN=Cnga3 PE=4 SV= | 0 | 148.372 | 23 | 15 | 137 | 9 | 669 | 76.5 | 7.02 | 1.021 | 0.952 | 1.072 | 0.03 | -0.03 | -0.07 | -0.0233333 | 0.02 | 0.1 | 0.882111909 | 0.813954813 | 0.583955577 | 0.999652205 | 0.998402594 | 0.9990274 | 0.973095093 |
| Pfn2 | High | Q9JJV2 | Profilin-2 OS=Mus musculus OX=10090 GN=Pfn2 PE=1 SV=3 | 0 | 34.548 | 43 | 5 | 8 | 5 | 140 | 15 | 6.99 | 1.02 | 0.955 | 1.068 | 0.03 | -0.03 | -0.07 | -0.0233333 | 0.02 | 0.09 | 0.997622795 | 0.820229949 | 0.729758076 | 0.999652205 | 0.998402594 | 0.9990274 | 0.992535481 |
| Rps17 | High | P63276 | 40S ribosomal protein S17 OS=Mus musculus OX=10090 GN=Rps17 PE=1 SV=2 | 0 | 55.631 | 56 | 8 | 16 | 8 | 135 | 15.5 | 9.85 | 1.03 | 0.946 | 1.089 | 0.04 | -0.04 | -0.08 | -0.0266667 | 0.02 | 0.12 | 0.917738973 | 0.797605933 | 0.515433901 | 0.999652205 | 0.998402594 | 0.9990274 | 0.964378027 |
| Ceacam20 | Low | Q9D2Z1 | Carcinoembryonic antigen-related cell adhesion molecule 20 OS=Mus musculus OX=10090 GN=Ce | 0.054 | 1.391 | 3 | 1 | 2 | 1 | 577 | 63.4 | 7.15 | 0.945 | 1.026 | 0.921 | -0.08 | 0.08 | 0.04 | 0.01333333 | 0.02 | -0.12 | 0.787451566 | 0.980852106 | 0.647072331 | 0.999652205 | 0.998402594 | 0.9990274 | 0.985010103 |
| Slc3a2 | High | P10852 | 4F2 cell-surface antigen heavy chain OS=Mus musculus OX=10090 GN=Slc3a2 PE=1 SV=1 | 0 | 108.889 | 43 | 20 | 43 | 20 | 526 | 58.3 | 5.91 | 0.964 | 1.009 | 0.956 | -0.05 | 0.05 | 0.01 | 0.00333333 | 0.02 | -0.06 | 0.905673503 | 0.952792576 | 0.850221589 | 0.999652205 | 0.998402594 | 0.9990274 | 0.992535481 |
| Rnf181 | Medium | Q9CY62 | E3 ubiquitin-protein ligase RNF181 OS=Mus musculus OX=10090 GN=Rnf181 PE=1 SV=1 | 0.034 | 1.822 | 4 | 1 | 1 | 1 | 165 | 19.1 | 6.04 | 0.994 | 0.978 | 1.017 | -0.01 | 0.01 | -0.03 | -0.01 | 0.02 | 0.02 | 0.856644512 | 0.860195454 | 0.86826068 | 0.999652205 | 0.998402594 | 0.9990274 | 0.992535481 |
| Atp8a1 | High | P70704 | Phospholipid-transporting ATPase IA OS=Mus musculus OX=10090 GN=Atp8a1 PE=1 SV=2 | 0 | 40.718 | 4 | 8 | 25 | 2 | 1164 | 131.3 | 6.84 | 0.975 | 0.998 | 0.977 | -0.04 | 0.04 | 0 | 0 | 0.02 | -0.03 | 0.861712771 | 0.927296145 | 0.924550711 | 0.999652205 | 0.998402594 | 0.9990274 | 0.992535481 |
| Rhog | High | P84096 | Rho-related GTP-binding protein RhoG OS=Mus musculus OX=10090 GN=Rhog PE=1 SV=1 | 0 | 30.832 | 31 | 6 | 8 | 5 | 191 | 21.3 | 8.12 | 0.92 | 1.057 | 0.87 | -0.12 | 0.12 | 0.08 | 0.02666667 | 0.02 | -0.2 | 0.696500391 | 0.933545071 | 0.38535813 | 0.999652205 | 0.998402594 | 0.9990274 | 0.928935096 |
| Sult1c1 | Medium | Q80VR3 | Sulfotransferase 1C1 OS=Mus musculus OX=10090 GN=Sult1c1 PE=1 SV=1 | 0.011 | 2.496 | 5 | 1 | 1 | 1 | 304 | 35.8 | 6.96 | 0.929 | 1.052 | 0.883 | -0.11 | 0.11 | 0.07 | 0.02333333 | 0.02 | -0.18 | 0.69951054 | 0.948142621 | 0.499045135 | 0.999652205 | 0.998402594 | 0.9990274 | 0.964378027 |
| Caskin1 | High | Q6P9K8 | Caskin-1 OS=Mus musculus OX=10090 GN=Caskin1 PE=1 SV=2 | 0 | 43.313 | 8 | 9 | 9 | 8 | 1431 | 150.4 | 9.17 | 0.961 | 1.016 | 0.946 | -0.06 | 0.06 | 0.02 | 0.00666667 | 0.02 | -0.08 | 0.835442203 | 0.970169149 | 0.724981718 | 0.999652205 | 0.998402594 | 0.9990274 | 0.992535481 |
| Fhl1 | High | A2AEX6 | Four and a half LIM domains protein 1 OS=Mus musculus OX=10090 GN=Fhl1 PE=1 SV=1 | 0 | 15.709 | 17 | 5 | 6 | 5 | 309 | 35.1 | 8.44 | 1.015 | 0.959 | 1.058 | 0.02 | -0.02 | -0.06 | -0.02 | 0.02 | 0.08 | 0.996676074 | 0.830698053 | 0.758066743 | 0.999652205 | 0.998402594 | 0.9990274 | 0.992535481 |
| Ppp3ca | High | P63328 | Serine/threonine-protein phosphatase 2B catalytic subunit alpha isoform OS=Mus musculus OX=1 | 0 | 212.178 | 61 | 24 | 107 | 14 | 521 | 58.6 | 5.86 | 0.957 | 1.014 | 0.943 | -0.06 | 0.06 | 0.02 | 0.00666667 | 0.02 | -0.08 | 0.884145369 | 0.966126548 | 0.782448718 | 0.999652205 | 0.998402594 | 0.9990274 | 0.992535481 |
| Sema7a | High | Q9QUR8 | Semaphorin-7A OS=Mus musculus OX=10090 GN=Sema7a PE=1 SV=1 | 0 | 10.774 | 6 | 3 | 3 | 3 | 664 | 74.9 | 7.74 | 0.905 | 1.079 | 0.839 | -0.14 | 0.14 | 0.11 | 0.03666667 | 0.015 | -0.25 | 0.663898672 | 0.897912194 | 0.317955099 | 0.999652205 | 0.998402594 | 0.9990274 | 0.904943958 |
| Rpap3 | High | Q9D706 | RNA polymerase II-associated protein 3 OS=Mus musculus OX=10090 GN=Rpap3 PE=1 SV=1 | 0 | 9.289 | 4 | 3 | 3 | 3 | 660 | 74.1 | 7.99 | 1.017 | 0.967 | 1.052 | 0.02 | -0.02 | -0.05 | -0.0166667 | 0.015 | 0.07 | 0.920103157 | 0.85043298 | 0.761147687 | 0.999652205 | 0.998402594 | 0.9990274 | 0.992535481 |
| Cnnm3 | High | Q32NY4 | Metal transporter CNNM3 OS=Mus musculus OX=10090 GN=Cnnm3 PE=1 SV=2 | 0.001 | 6.498 | 4 | 3 | 3 | 2 | 713 | 76.2 | 5.58 | 0.932 | 1.053 | 0.885 | -0.1 | 0.1 | 0.07 | 0.02333333 | 0.015 | -0.18 | 0.73201498 | 0.958303283 | 0.498736894 | 0.999652205 | 0.998402594 | 0.9990274 | 0.964378027 |
| Snx5 | High | Q9D8U8 | Sorting nexin-5 OS=Mus musculus OX=10090 GN=Snx5 PE=1 SV=1 | 0 | 8.593 | 8 | 3 | 4 | 3 | 404 | 46.8 | 6.62 | 0.939 | 1.043 | 0.9 | -0.09 | 0.09 | 0.06 | 0.02 | 0.015 | -0.15 | 0.768590954 | 0.964393813 | 0.532682378 | 0.999652205 | 0.998402594 | 0.9990274 | 0.96694628 |
| Atp2b2 | High | F8WHB1 | Calcium-transporting ATPase OS=Mus musculus OX=10090 GN=Atp2b2 PE=1 SV=1 | 0 | 260.666 | 40 | 43 | 74 | 28 | 1243 | 136.7 | 5.97 | 0.948 | 1.036 | 0.915 | -0.08 | 0.08 | 0.05 | 0.01666667 | 0.015 | -0.13 | 0.859148167 | 0.98194627 | 0.635710789 | 0.999652205 | 0.998402594 | 0.9990274 | 0.983562471 |
| Sppl2a | High | Q9JJF9 | Signal peptide peptidase-like 2A OS=Mus musculus OX=10090 GN=Sppl2a PE=1 SV=2 | 0 | 23.443 | 8 | 3 | 7 | 3 | 523 | 58.1 | 7.06 | 1.01 | 0.975 | 1.035 | 0.01 | -0.01 | -0.04 | -0.0133333 | 0.015 | 0.05 | 0.92313279 | 0.871091985 | 0.814995697 | 0.999652205 | 0.998402594 | 0.9990274 | 0.992535481 |
| Cyfip2 | High | Q5SQX6 | Cytoplasmic FMR1-interacting protein 2 OS=Mus musculus OX=10090 GN=Cyfip2 PE=1 SV=2 | 0 | 22.954 | 6 | 9 | 11 | 3 | 1253 | 145.6 | 7.05 | 1.003 | 0.981 | 1.022 | 0 | 0 | -0.03 | -0.01 | 0.015 | 0.03 | 0.981056705 | 0.886011285 | 0.817652872 | 0.999652205 | 0.998402594 | 0.9990274 | 0.992535481 |
| Sept6 | High | Q9R1T4 | Septin-6 OS=Mus musculus OX=10090 GN=Septin6 PE=1 SV=4 | 0 | 48.334 | 31 | 8 | 12 | 4 | 434 | 49.6 | 6.43 | 0.97 | 1.004 | 0.966 | -0.04 | 0.04 | 0.01 | 0.00333333 | 0.015 | -0.05 | 0.906886322 | 0.940945916 | 0.875600392 | 0.999652205 | 0.998402594 | 0.9990274 | 0.992535481 |
| Gnai1 | High | B2RSH2 | Guanine nucleotide-binding protein G(i) subunit alpha-1 OS=Mus musculus OX=10090 GN=Gnai1 | 0 | 84.797 | 38 | 15 | 93 | 4 | 354 | 40.3 | 5.97 | 1.002 | 0.98 | 1.021 | 0 | 0 | -0.03 | -0.01 | 0.015 | 0.03 | 0.972847392 | 0.883704954 | 0.877605495 | 0.999652205 | 0.998402594 | 0.9990274 | 0.992535481 |
| Cd81 | High | P35762 | CD81 antigen OS=Mus musculus OX=10090 GN=Cd81 PE=1 SV=2 | 0 | 72.389 | 18 | 2 | 16 | 2 | 236 | 25.8 | 5.83 | 0.999 | 0.979 | 1.021 | 0 | 0 | -0.03 | -0.01 | 0.015 | 0.03 | 0.960647621 | 0.879375515 | 0.890658589 | 0.999652205 | 0.998402594 | 0.9990274 | 0.992535481 |
| Pi4ka | High | E9Q3L2 | Phosphatidylinositol 4-kinase alpha OS=Mus musculus OX=10090 GN=Pi4ka PE=1 SV=2 | 0 | 43.109 | 6 | 10 | 11 | 10 | 2105 | 236.9 | 7.06 | 0.981 | 1 | 0.982 | -0.03 | 0.03 | 0 | 0 | 0.015 | -0.03 | 0.893742109 | 0.930659308 | 0.912979368 | 0.999652205 | 0.998402594 | 0.9990274 | 0.992535481 |
| Mtap | High | Q9CQ65 | S-methyl-5'-thioadenosine phosphorylase OS=Mus musculus OX=10090 GN=Mtap PE=1 SV=1 | 0.005 | 3.648 | 4 | 1 | 1 | 1 | 283 | 31 | 7.14 | 0.997 | 0.981 | 1.016 | 0 | 0 | -0.03 | -0.01 | 0.015 | 0.02 | 0.958569495 | 0.885963577 | 0.937471711 | 0.999652205 | 0.998402594 | 0.9990274 | 0.992535481 |
| Gkap1 | High | Q9JMB0 | G kinase-anchoring protein 1 OS=Mus musculus OX=10090 GN=Gkap1 PE=1 SV=1 | 0.001 | 4.616 | 4 | 2 | 2 | 2 | 366 | 41.7 | 8.82 | 0.944 | 1.041 | 0.906 | -0.08 | 0.08 | 0.06 | 0.02 | 0.01 | -0.14 | 0.812857283 | 0.967901402 | 0.588383389 | 0.999652205 | 0.998402594 | 0.9990274 | 0.973095093 |
| Calu | High | O35887 | Calumenin OS=Mus musculus OX=10090 GN=Calu PE=1 SV=1 | 0 | 12.383 | 16 | 3 | 3 | 3 | 315 | 37 | 4.67 | 0.947 | 1.039 | 0.911 | -0.08 | 0.08 | 0.06 | 0.02 | 0.01 | -0.13 | 0.776913121 | 0.967636573 | 0.59560612 | 0.999652205 | 0.998402594 | 0.9990274 | 0.977451587 |
| Dpysl2 | High | O08553 | Dihydropyrimidinase-related protein 2 OS=Mus musculus OX=10090 GN=Dpysl2 PE=1 SV=2 | 0 | 326.61 | 74 | 32 | 119 | 27 | 572 | 62.2 | 6.38 | 0.954 | 1.032 | 0.925 | -0.07 | 0.07 | 0.05 | 0.01666667 | 0.01 | -0.11 | 0.876182666 | 0.992069543 | 0.68505336 | 0.999652205 | 0.998402594 | 0.9990274 | 0.991285312 |
| | High | A0A1L1SV | Choline transporter-like protein 2 OS=Mus musculus OX=10090 GN=Slc44a2 PE=1 SV=1 | 0 | 20.787 | 12 | 6 | 8 | 6 | 554 | 62.6 | 8.21 | 0.966 | 1.018 | 0.949 | -0.05 | 0.05 | 0.03 | 0.01 | 0.01 | -0.08 | 0.840924182 | 0.975077036 | 0.718273455 | 0.999652205 | 0.998402594 | 0.9990274 | 0.992535481 |
| Exoc8 | High | Q6PGF7 | Exocyst complex component 8 OS=Mus musculus OX=10090 GN=Exoc8 PE=1 SV=1 | 0.009 | 2.78 | 2 | 2 | 2 | 2 | 716 | 81 | 5.4 | 0.974 | 1.013 | 0.961 | -0.04 | 0.04 | 0.02 | 0.00666667 | 0.01 | -0.06 | 0.816730325 | 0.954753762 | 0.83530482 | 0.999652205 | 0.998402594 | 0.9990274 | 0.992535481 |
| Bola2 | High | Q8BGS2 | BolA-like protein 2 OS=Mus musculus OX=10090 GN=Bola2 PE=1 SV=1 | 0.006 | 3.36 | 19 | 2 | 2 | 2 | 86 | 10.2 | 6.16 | 0.999 | 0.986 | 1.013 | 0 | 0 | -0.02 | -0.0066667 | 0.01 | 0.02 | 0.920037959 | 0.897417669 | 0.890984737 | 0.999652205 | 0.998402594 | 0.9990274 | 0.992535481 |
| Gnpda1 | High | O88958 | Glucosamine-6-phosphate isomerase 1 OS=Mus musculus OX=10090 GN=Gnpda1 PE=1 SV=3 | 0 | 21.451 | 27 | 7 | 8 | 5 | 289 | 32.5 | 6.6 | 0.991 | 0.993 | 0.998 | -0.01 | 0.01 | -0.01 | -0.0033333 | 0.01 | 0 | 0.915230917 | 0.914251133 | 0.95633115 | 0.999652205 | 0.998402594 | 0.9990274 | 0.992902921 |
| Eepd1 | High | Q3TGW2 | Endonuclease/exonuclease/phosphatase family domain-containing protein 1 OS=Mus musculus C | 0.001 | 5.649 | 2 | 1 | 1 | 1 | 569 | 62.9 | 8.35 | 0.986 | 1 | 0.986 | -0.02 | 0.02 | 0 | 0 | 0.01 | -0.02 | 0.893511985 | 0.911563076 | 0.99022943 | 0.999652205 | 0.998402594 | 0.9990274 | 0.999783808 |
| Top1 | Medium | Q04750 | DNA topoisomerase 1 OS=Mus musculus OX=10090 GN=Top1 PE=1 SV=2 | 0.02 | 2.03 | 1 | 1 | 1 | 1 | 767 | 90.8 | 9.33 | 1.029 | 0.961 | 1.071 | 0.04 | -0.04 | -0.06 | -0.02 | 0.01 | 0.1 | 0.94402736 | 0.837217052 | 0.683142237 | 0.999652205 | 0.998402594 | 0.9990274 | 0.991285312 |
| 2310035C | High | E9QM90 | RAB11-binding protein RELCH OS=Mus musculus OX=10090 GN=Relch PE=1 SV=2 | 0.001 | 5.591 | 1 | 1 | 2 | 1 | 1216 | 134.5 | 5.34 | 1.01 | 0.98 | 1.03 | 0.01 | -0.01 | -0.03 | -0.01 | 0.01 | 0.04 | 0.975676633 | 0.877420558 | 0.85673639 | 0.999652205 | 0.998402594 | 0.9990274 | 0.992535481 |
| Rp2h; Rp2 | High | Q9EPK2 | Protein XRP2 OS=Mus musculus OX=10090 GN=Rp2 PE=1 SV=3 | 0 | 16.178 | 14 | 5 | 6 | 5 | 347 | 39.4 | 5.24 | 1.008 | 0.982 | 1.026 | 0.01 | -0.01 | -0.03 | -0.01 | 0.01 | 0.04 | 0.961250736 | 0.887657174 | 0.904572526 | 0.999652205 | 0.998402594 | 0.9990274 | 0.992535481 |
| | High | E9QKR0 | Guanine nucleotide-binding protein G(I)/G(S)/G(T) subunit beta-2 OS=Mus musculus OX=10090 G | 0 | 170.907 | 37 | 14 | 539 | 4 | 382 | 41.4 | 6.6 | 0.977 | 1.009 | 0.968 | -0.03 | 0.03 | 0.01 | 0.00333333 | 0.01 | -0.05 | 0.956953139 | 0.953881723 | 0.915406659 | 0.999652205 | 0.998402594 | 0.9990274 | 0.992535481 |
| Gps1 | High | Q99LD4 | COP9 signalosome complex subunit 1 OS=Mus musculus OX=10090 GN=Gps1 PE=1 SV=1 | 0.001 | 7.709 | 4 | 2 | 3 | 2 | 471 | 53.4 | 6.84 | 0.877 | 1.125 | 0.78 | -0.19 | 0.19 | 0.17 | 0.05666667 | 0.01 | -0.36 | 0.538377898 | 0.783847129 | 0.137615262 | 0.999652205 | 0.998402594 | 0.9990274 | 0.700189449 |
| Hspa12a | High | Q8K0U4 | Heat shock 70 kDa protein 12A OS=Mus musculus OX=10090 GN=Hspa12a PE=1 SV=1 | 0 | 45.602 | 29 | 15 | 17 | 15 | 675 | 74.8 | 6.77 | 0.917 | 1.075 | 0.853 | -0.12 | 0.12 | 0.1 | 0.03333333 | 0.01 | -0.23 | 0.777048747 | 0.891290925 | 0.36678791 | 0.999652205 | 0.998402594 | 0.9990274 | 0.920164724 |
| Lamtor1 | High | Q9CQ22 | Ragulator complex protein LAMTOR1 OS=Mus musculus OX=10090 GN=Lamtor1 PE=1 SV=1 | 0 | 76.382 | 56 | 7 | 23 | 7 | 161 | 17.7 | 5.15 | 0.923 | 1.071 | 0.862 | -0.12 | 0.12 | 0.1 | 0.03333333 | 0.01 | -0.21 | 0.805142357 | 0.901854633 | 0.391360922 | 0.999652205 | 0.998402594 | 0.9990274 | 0.93064581 |
| Vamp5 | High | Q9Z2P8 | Vesicle-associated membrane protein 5 OS=Mus musculus OX=10090 GN=Vamp5 PE=1 SV=1 | 0.006 | 3.386 | 13 | 1 | 1 | 1 | 102 | 11.4 | 7.97 | 1.051 | 0.941 | 1.117 | 0.07 | -0.07 | -0.09 | -0.03 | 0.01 | 0.16 | 0.89939564 | 0.787044413 | 0.52128522 | 0.999652205 | 0.998402594 | 0.9990274 | 0.964378027 |
| Pip5k1b | Medium | P70181 | Phosphatidylinositol 4-phosphate 5-kinase type-1 beta OS=Mus musculus OX=10090 GN=Pip5k1b | 0.025 | 1.98 | 1 | 1 | 1 | 1 | 539 | 60.8 | 6.86 | 0.911 | 1.087 | 0.838 | -0.13 | 0.13 | 0.12 | 0.04 | 0.005 | -0.25 | 0.670873477 | 0.854508093 | 0.347581535 | 0.999652205 | 0.998402594 | 0.9990274 | 0.909421216 |
| Ccny | High | Q8BGU5 | Cyclin-Y OS=Mus musculus OX=10090 GN=Ccny PE=1 SV=1 | 0.001 | 5.284 | 7 | 2 | 2 | 1 | 341 | 39.4 | 7.2 | 0.933 | 1.062 | 0.879 | -0.1 | 0.1 | 0.09 | 0.03 | 0.005 | -0.19 | 0.748218479 | 0.94232677 | 0.569212179 | 0.999652205 | 0.998402594 | 0.9990274 | 0.972910028 |
| Mapt | High | A0A0A0M | Microtubule-associated protein OS=Mus musculus OX=10090 GN=Mapt PE=1 SV=1 | 0 | 61.592 | 27 | 18 | 31 | 18 | 733 | 76.2 | 6.79 | 0.95 | 1.045 | 0.909 | -0.07 | 0.07 | 0.06 | 0.02 | 0.005 | -0.14 | 0.866466224 | 0.960718553 | 0.608782728 | 0.999652205 | 0.998402594 | 0.9990274 | 0.97826833 |
| Kif1a | High | Q6TA13 | Kinesin-like protein KIF1A OS=Mus musculus OX=10090 GN=Kif1a PE=1 SV=1 | 0 | 21.018 | 4 | 7 | 8 | 2 | 1698 | 191.7 | 6.23 | 1.006 | 0.987 | 1.02 | 0.01 | -0.01 | -0.02 | -0.0066667 | 0.005 | 0.03 | 0.996866378 | 0.900868824 | 0.892124913 | 0.999652205 | 0.998402594 | 0.9990274 | 0.992535481 |
| Gstm1 | High | P10649 | Glutathione S-transferase Mu 1 OS=Mus musculus OX=10090 GN=Gstm1 PE=1 SV=2 | 0 | 24.548 | 33 | 7 | 11 | 4 | 218 | 26 | 7.94 | 0.992 | 1.003 | 0.989 | -0.01 | 0.01 | 0 | 0 | 0.005 | -0.02 | 0.933108522 | 0.938885042 | 0.913666694 | 0.999652205 | 0.998402594 | 0.9990274 | 0.992535481 |
| | High | A0A0G2JD | Phosphatidylcholine translocator ABCB4 (Fragment) OS=Mus musculus OX=10090 GN=Abcb4 PE= | 0.005 | 3.667 | 1 | 1 | 1 | 1 | 882 | 96.8 | 8.82 | 1.014 | 0.982 | 1.032 | 0.02 | -0.02 | -0.03 | -0.01 | 0.005 | 0.05 | 0.98547083 | 0.892971829 | 0.847979751 | 0.999652205 | 0.998402594 | 0.9990274 | 0.992535481 |
| Yes1 | High | Q04736 | Tyrosine-protein kinase Yes OS=Mus musculus OX=10090 GN=Yes1 PE=1 SV=3 | 0 | 39.239 | 27 | 11 | 13 | 8 | 541 | 60.6 | 6.64 | 0.926 | 1.07 | 0.865 | -0.11 | 0.11 | 0.1 | 0.03333333 | 0.005 | -0.21 | 0.750479247 | 0.902277026 | 0.353727495 | 0.999652205 | 0.998402594 | 0.9990274 | 0.911642008 |
| Ttc5 | Medium | Q99LG4 | Tetratricopeptide repeat protein 5 OS=Mus musculus OX=10090 GN=Ttc5 PE=1 SV=2 | 0.01 | 2.646 | 2 | 1 | 1 | 1 | 440 | 48.8 | 6.29 | 0.92 | 1.076 | 0.855 | -0.12 | 0.12 | 0.11 | 0.03666667 | 0.005 | -0.23 | 0.695384283 | 0.880362219 | 0.412200748 | 0.999652205 | 0.998402594 | 0.9990274 | 0.936397673 |
| Cltb | High | Q6IRU5 | Clathrin light chain B OS=Mus musculus OX=10090 GN=Cltb PE=1 SV=1 | 0 | 19.145 | 30 | 6 | 7 | 6 | 229 | 25.2 | 4.63 | 0.9 | 1.099 | 0.819 | -0.15 | 0.15 | 0.14 | 0.04666667 | 0.005 | -0.29 | 0.657306153 | 0.837529882 | 0.203168704 | 0.999652205 | 0.998402594 | 0.9990274 | 0.788350811 |
| Slc38a1 | Medium | Q8K2P7 | Sodium-coupled neutral amino acid transporter 1 OS=Mus musculus OX=10090 GN=Slc38a1 PE=1 | 0.037 | 1.737 | 1 | 1 | 1 | 1 | 485 | 53.8 | 7.08 | 0.905 | 1.1 | 0.823 | -0.14 | 0.14 | 0.14 | 0.04666667 | 0 | -0.28 | 0.652479466 | 0.846478487 | 0.291882143 | 0.999652205 | 0.998402594 | 0.9990274 | 0.892844069 |
| | High | A0A2I3BQ | Calcium/calmodulin-dependent protein kinase OS=Mus musculus OX=10090 GN=Camk2g PE=1 SV | 0 | 80.126 | 29 | 15 | 31 | 4 | 565 | 63.2 | 7.84 | 1.051 | 0.955 | 1.1 | 0.07 | -0.07 | -0.07 | -0.0233333 | 0 | 0.14 | 0.865236493 | 0.820890583 | 0.474915732 | 0.999652205 | 0.998402594 | 0.9990274 | 0.962227984 |
| Enoph1 | Medium | Q8BGB7 | Enolase-phosphatase E1 OS=Mus musculus OX=10090 GN=Enoph1 PE=1 SV=1 | 0.01 | 2.742 | 3 | 1 | 1 | 1 | 257 | 28.6 | 4.92 | 1.048 | 0.95 | 1.103 | 0.07 | -0.07 | -0.07 | -0.0233333 | 0 | 0.14 | 0.906852214 | 0.808207482 | 0.503652349 | 0.999652205 | 0.998402594 | 0.9990274 | 0.964378027 |
| Atp6v1c1 | High | Q9Z1G3 | V-type proton ATPase subunit C 1 OS=Mus musculus OX=10090 GN=Atp6v1c1 PE=1 SV=4 | 0 | 64.689 | 45 | 19 | 32 | 19 | 382 | 43.9 | 7.46 | 1.038 | 0.968 | 1.072 | 0.05 | -0.05 | -0.05 | -0.0166667 | 0 | 0.1 | 0.896711864 | 0.853411512 | 0.581899603 | 0.999652205 | 0.998402594 | 0.9990274 | 0.972910028 |
| Dynlrb1 | High | A2AVR9 | Dynein light chain roadblock OS=Mus musculus OX=10090 GN=Dynlrb1 PE=1 SV=1 | 0 | 18.757 | 47 | 3 | 5 | 3 | 104 | 11.9 | 6.06 | 1.045 | 0.958 | 1.091 | 0.06 | -0.06 | -0.06 | -0.02 | 0 | 0.13 | 0.854088637 | 0.827470462 | 0.604077004 | 0.999652205 | 0.998402594 | 0.9990274 | 0.97826833 |
| | High | E9PZ92 | Exocyst complex component 5 OS=Mus musculus OX=10090 GN=Exoc5 PE=1 SV=1 | 0.001 | 6.257 | 6 | 3 | 3 | 3 | 643 | 74.2 | 6.99 | 1.035 | 0.967 | 1.07 | 0.05 | -0.05 | -0.05 | -0.0166667 | 0 | 0.1 | 0.964792645 | 0.832183137 | 0.62227611 | 0.999652205 | 0.998402594 | 0.9990274 | 0.97826833 |
| Mfap3l | High | Q9D3X9 | Microfibrillar-associated protein 3-like OS=Mus musculus OX=10090 GN=Mfap3l PE=1 SV=1 | 0 | 22.575 | 18 | 7 | 11 | 7 | 409 | 45.3 | 5.08 | 1.025 | 0.97 | 1.057 | 0.04 | -0.04 | -0.04 | -0.0133333 | 0 | 0.08 | 0.919579349 | 0.857622154 | 0.64748606 | 0.999652205 | 0.998402594 | 0.9990274 | 0.985010103 |
| Coro2b | High | Q8BH44 | Coronin-2B OS=Mus musculus OX=10090 GN=Coro2b PE=1 SV=2 | 0 | 11.834 | 10 | 4 | 4 | 4 | 480 | 54.9 | 8.27 | 0.964 | 1.034 | 0.932 | -0.05 | 0.05 | 0.05 | 0.01666667 | 0 | -0.1 | 0.864163461 | 0.978004411 | 0.733784704 | 0.999652205 | 0.998402594 | 0.9990274 | 0.992535481 |
| Znrf2 | High | Q71FD5 | E3 ubiquitin-protein ligase ZNRF2 OS=Mus musculus OX=10090 GN=Znrf2 PE=1 SV=1 | 0 | 9.686 | 10 | 2 | 2 | 2 | 238 | 23.7 | 6.76 | 1.025 | 0.979 | 1.047 | 0.03 | -0.03 | -0.03 | -0.01 | 0 | 0.07 | 0.994137309 | 0.859840346 | 0.742437536 | 0.999652205 | 0.998402594 | 0.9990274 | 0.992535481 |
| | High | A0A0N4SV | Guanine nucleotide-binding protein subunit gamma OS=Mus musculus OX=10090 GN=Gng12 PE= | 0 | 22.011 | 58 | 4 | 10 | 4 | 83 | 9.2 | 9.25 | 0.974 | 1.03 | 0.945 | -0.04 | 0.04 | 0.04 | 0.01333333 | 0 | -0.08 | 0.96143561 | 0.995223679 | 0.773319361 | 0.999652205 | 0.998402594 | 0.9990274 | 0.992535481 |
| Cyp2d22 | High | Q9JKY7 | Cytochrome P450 CYP2D22 OS=Mus musculus OX=10090 GN=Cyp2d22 PE=1 SV=1 | 0 | 13.893 | 8 | 3 | 3 | 3 | 500 | 56.4 | 6.98 | 1.013 | 0.985 | 1.029 | 0.02 | -0.02 | -0.02 | -0.0066667 | 0 | 0.04 | 0.970867047 | 0.874476481 | 0.800015852 | 0.999652205 | 0.998402594 | 0.9990274 | 0.992535481 |
| Smap1 | High | Q91VZ6 | Stromal membrane-associated protein 1 OS=Mus musculus OX=10090 GN=Smap1 PE=1 SV=1 | 0 | 10.352 | 9 | 4 | 5 | 4 | 440 | 47.6 | 8.51 | 1.022 | 0.976 | 1.047 | 0.03 | -0.03 | -0.03 | -0.01 | 0 | 0.07 | 0.983332771 | 0.87324191 | 0.815043955 | 0.999652205 | 0.998402594 | 0.9990274 | 0.992535481 |
| Farp1 | High | F8VPU2 | FERM, ARHGEF and pleckstrin domain-containing protein 1 OS=Mus musculus OX=10090 GN=Far | 0.001 | 4.652 | 1 | 1 | 1 | 1 | 1048 | 118.8 | 7.88 | 1.017 | 0.984 | 1.033 | 0.02 | -0.02 | -0.02 | -0.0066667 | 0 | 0.05 | 0.978787872 | 0.896789936 | 0.816329291 | 0.999652205 | 0.998402594 | 0.9990274 | 0.992535481 |

| | | | | | | | | | | | | | | | | | | | | | | | | | | | | |
|---|---|---|---|---|---|---|---|---|---|---|---|---|---|---|---|---|---|---|---|---|---|---|---|---|---|---|---|---|
| Apoa1bp; | High | Q8K4Z3 | NAD(P)H-hydrate epimerase OS=Mus musculus OX=10090 GN=Naxe PE=1 SV=1 | 0 | 17.165 | 22 | 5 | 7 | 5 | 282 | 31 | 7.69 | 0.991 | 1.004 | 0.987 | -0.01 | 0.01 | 0.01 | 0.00333333 | 0 | -0.02 | 0.920297667 | 0.941625783 | 0.936902931 | 0.999652205 | 0.998402594 | 0.9990274 | 0.992535481 |
| Itpr1 | Low | P11881 | Inositol 1,4,5-trisphosphate receptor type 1 OS=Mus musculus OX=10090 GN=Itpr1 PE=1 SV=2 | 0.056 | 1.377 | 0 | 1 | 1 | 1 | 2749 | 313 | 6.04 | 1.003 | 1 | 1.003 | 0 | 0 | 0 | 0 | 0 | 0 | 0.942314184 | 0.918802331 | 0.941010446 | 0.999652205 | 0.998402594 | 0.9990274 | 0.992535481 |
| Gmpr | High | Q9DCZ1 | GMP reductase 1 OS=Mus musculus OX=10090 GN=Gmpr PE=1 SV=1 | 0 | 8.187 | 8 | 2 | 2 | 2 | 345 | 37.5 | 7.09 | 1.002 | 1.003 | 0.998 | 0 | 0 | 0 | 0 | 0 | 0 | 0.969823079 | 0.939798115 | 0.982123366 | 0.999652205 | 0.998402594 | 0.9990274 | 0.997765878 |
| Dock7 | High | A2A9M4 | Dedicator of cytokinesis protein 7 OS=Mus musculus OX=10090 GN=Dock7 PE=1 SV=2 | 0.001 | 6.475 | 1 | 3 | 3 | 3 | 2130 | 241.3 | 6.76 | 0.933 | 1.078 | 0.866 | -0.1 | 0.1 | 0.11 | 0.03666667 | -0.005 | -0.21 | 0.7176052 | 0.884371702 | 0.414583472 | 0.999652205 | 0.998402594 | 0.9990274 | 0.936397673 |
| | High | A0A2R8VK | Methyltransferase-like 7A1 OS=Mus musculus OX=10090 GN=Mettl7a1 PE=1 SV=1 | 0.005 | 3.74 | 5 | 1 | 2 | 1 | 193 | 22.5 | 8 | 0.943 | 1.063 | 0.887 | -0.08 | 0.08 | 0.09 | 0.03 | -0.005 | -0.17 | 0.711734281 | 0.929464705 | 0.486441223 | 0.999652205 | 0.998402594 | 0.9990274 | 0.964378027 |
| | Low | A0A3B2W | Mitogen-activated protein kinase OS=Mus musculus OX=10090 GN=Mapk14 PE=1 SV=1 | 0.056 | 1.367 | 8 | 1 | 1 | 1 | 307 | 35.3 | 8.97 | 0.949 | 1.065 | 0.891 | -0.08 | 0.08 | 0.09 | 0.03 | -0.005 | -0.17 | 0.711849441 | 0.927169909 | 0.491697467 | 0.999652205 | 0.998402594 | 0.9990274 | 0.964378027 |
| Crb2 | High | Q80YA8 | Protein crumbs homolog 2 OS=Mus musculus OX=10090 GN=Crb2 PE=1 SV=3 | 0 | 13.181 | 3 | 3 | 4 | 3 | 1282 | 134.7 | 5.22 | 0.948 | 1.064 | 0.89 | -0.08 | 0.08 | 0.09 | 0.03 | -0.005 | -0.17 | 0.753012248 | 0.91577417 | 0.508841204 | 0.999652205 | 0.998402594 | 0.9990274 | 0.964378027 |
| Yars | High | Q91WQ3 | Tyrosine--tRNA ligase, cytoplasmic OS=Mus musculus OX=10090 GN=Yars1 PE=1 SV=3 | 0 | 86.43 | 40 | 25 | 35 | 25 | 528 | 59.1 | 7.01 | 1.039 | 0.963 | 1.079 | 0.06 | -0.06 | -0.05 | -0.0166667 | -0.005 | 0.11 | 0.89503139 | 0.83950744 | 0.553857577 | 0.999652205 | 0.998402594 | 0.9990274 | 0.96775057 |
| Ano6 | High | Q6P9J9 | Anoctamin-6 OS=Mus musculus OX=10090 GN=Ano6 PE=1 SV=1 | 0.001 | 6.774 | 3 | 2 | 2 | 2 | 911 | 106.2 | 6.76 | 0.964 | 1.04 | 0.926 | -0.05 | 0.05 | 0.06 | 0.02 | -0.005 | -0.11 | 0.813354282 | 0.985930903 | 0.716910141 | 0.999652205 | 0.998402594 | 0.9990274 | 0.992535481 |
| Abr | High | Q5SSL4 | Active breakpoint cluster region-related protein OS=Mus musculus OX=10090 GN=Abr PE=1 SV=1 | 0.001 | 6.644 | 2 | 2 | 2 | 2 | 859 | 97.6 | 6.58 | 1.022 | 0.983 | 1.04 | 0.03 | -0.03 | -0.02 | -0.0066667 | -0.005 | 0.06 | 0.950164019 | 0.890635098 | 0.803766833 | 0.999652205 | 0.998402594 | 0.9990274 | 0.992535481 |
| Epb4.1l5; I | High | Q8BGS1 | Band 4.1-like protein 5 OS=Mus musculus OX=10090 GN=Epb41l5 PE=1 SV=1 | 0.006 | 3.211 | 1 | 1 | 1 | 1 | 731 | 81.6 | 6.39 | 1.004 | 1.003 | 1.001 | 0.01 | -0.01 | 0 | 0 | -0.005 | 0 | 0.93572084 | 0.919812208 | 0.923071475 | 0.999652205 | 0.998402594 | 0.9990274 | 0.992535481 |
| Inpp4b | High | Q3URI3 | Phosphatidylinositol-3,4-bisphosphate 4-phosphatase (Fragment) OS=Mus musculus OX=10090 G | 0.001 | 6.684 | 3 | 3 | 3 | 3 | 818 | 92.3 | 5.58 | 0.996 | 1.016 | 0.98 | -0.01 | 0.01 | 0.02 | 0.00666667 | -0.005 | -0.03 | 0.888401695 | 0.960907634 | 0.934471822 | 0.999652205 | 0.998402594 | 0.9990274 | 0.992535481 |
| Atp6v1b2 | High | P62814 | V-type proton ATPase subunit B, brain isoform OS=Mus musculus OX=10090 GN=Atp6v1b2 PE=1 S | 0 | 184.88 | 74 | 25 | 58 | 25 | 511 | 56.5 | 5.81 | 1.035 | 0.975 | 1.061 | 0.05 | -0.05 | -0.04 | -0.0133333 | -0.005 | 0.09 | 0.906550263 | 0.870163667 | 0.629789412 | 0.999652205 | 0.998402594 | 0.9990274 | 0.981916182 |
| Arhgef26 | High | D3YYY8 | Rho guanine nucleotide exchange factor (GEF) 26 OS=Mus musculus OX=10090 GN=Arhgef26 PE= | 0.001 | 7.402 | 2 | 2 | 2 | 2 | 869 | 97.3 | 8.72 | 1.038 | 0.971 | 1.069 | 0.05 | -0.05 | -0.04 | -0.0133333 | -0.005 | 0.1 | 0.939810434 | 0.833641021 | 0.636696617 | 0.999652205 | 0.998402594 | 0.9990274 | 0.983597794 |
| Lrrc73 | High | B2RWC4 | Gm88 protein OS=Mus musculus OX=10090 GN=Lrrc73 PE=1 SV=1 | 0 | 4.759 | 11 | 2 | 2 | 2 | 316 | 33.4 | 4.84 | 0.896 | 1.121 | 0.799 | -0.16 | 0.16 | 0.17 | 0.05666667 | -0.005 | -0.32 | 0.602169728 | 0.783608858 | 0.216714872 | 0.999652205 | 0.998402594 | 0.9990274 | 0.804947026 |
| Ndufs3 | High | Q9DCT2 | NADH dehydrogenase [ubiquinone] iron-sulfur protein 3, mitochondrial OS=Mus musculus OX=10( | 0 | 20.32 | 22 | 5 | 7 | 5 | 263 | 30.1 | 7.17 | 0.961 | 1.053 | 0.913 | -0.06 | 0.06 | 0.07 | 0.02333333 | -0.005 | -0.13 | 0.830981574 | 0.941768993 | 0.553938586 | 0.999652205 | 0.998402594 | 0.9990274 | 0.96775057 |
| | High | D3YYN7 | Sodium/potassium-transporting ATPase subunit alpha OS=Mus musculus OX=10090 GN=Atp1a2 F | 0 | 447.527 | 40 | 38 | 223 | 14 | 947 | 103.5 | 5.44 | 0.959 | 1.047 | 0.915 | -0.06 | 0.06 | 0.07 | 0.02333333 | -0.005 | -0.13 | 0.937351591 | 0.955585424 | 0.634884158 | 0.999652205 | 0.998402594 | 0.9990274 | 0.983562471 |
| | High | A0A2R8VH | GTPase RhebL1 (Fragment) OS=Mus musculus OX=10090 GN=Rhebl1 PE=4 SV=1 | 0.009 | 2.841 | 6 | 1 | 1 | 1 | 151 | 16.9 | 7.66 | 0.91 | 1.117 | 0.815 | -0.14 | 0.14 | 0.16 | 0.05333333 | -0.01 | -0.3 | 0.679545777 | 0.798963246 | 0.201053354 | 0.999652205 | 0.998402594 | 0.9990274 | 0.78697088 |
| Usp9x | High | Q4FE56 | Ubiquitinyl hydrolase 1 OS=Mus musculus OX=10090 GN=Usp9x PE=1 SV=1 | 0 | 20.677 | 3 | 6 | 6 | 6 | 2554 | 290 | 5.83 | 0.935 | 1.083 | 0.863 | -0.1 | 0.1 | 0.12 | 0.04 | -0.01 | -0.21 | 0.831163556 | 0.873398579 | 0.425687987 | 0.999652205 | 0.998402594 | 0.9990274 | 0.941180024 |
| | High | E9PZF5 | Acidic leucine-rich nuclear phosphoprotein 32 family member E (Fragment) OS=Mus musculus OX | 0 | 10.886 | 19 | 3 | 3 | 3 | 171 | 19.2 | 3.8 | 1.064 | 0.954 | 1.115 | 0.09 | -0.09 | -0.07 | -0.0233333 | -0.01 | 0.16 | 0.848065105 | 0.819099801 | 0.514039945 | 0.999652205 | 0.998402594 | 0.9990274 | 0.964378027 |
| Gsk3b | High | E9QAQ5 | Glycogen synthase kinase-3 beta OS=Mus musculus OX=10090 GN=Gsk3b PE=1 SV=1 | 0 | 43.784 | 37 | 10 | 12 | 8 | 433 | 48 | 8.78 | 1.062 | 0.954 | 1.113 | 0.09 | -0.09 | -0.07 | -0.0233333 | -0.01 | 0.15 | 0.878050138 | 0.818699571 | 0.519453978 | 0.999652205 | 0.998402594 | 0.9990274 | 0.964378027 |
| Actr3 | High | Q99JY9 | Actin-related protein 3 OS=Mus musculus OX=10090 GN=Actr3 PE=1 SV=3 | 0 | 104.25 | 55 | 19 | 40 | 17 | 418 | 47.3 | 5.88 | 1.044 | 0.974 | 1.072 | 0.06 | -0.06 | -0.04 | -0.0133333 | -0.01 | 0.1 | 0.882196971 | 0.86782625 | 0.584210902 | 0.999652205 | 0.998402594 | 0.9990274 | 0.973095093 |
| Ap1g1 | High | Q8CBB7 | AP-1 complex subunit gamma OS=Mus musculus OX=10090 GN=Ap1g1 PE=1 SV=1 | 0 | 13.034 | 5 | 4 | 4 | 4 | 825 | 91.7 | 6.8 | 0.987 | 1.025 | 0.963 | -0.02 | 0.02 | 0.04 | 0.01333333 | -0.01 | -0.05 | 0.800561397 | 0.976398379 | 0.854410316 | 0.999652205 | 0.998402594 | 0.9990274 | 0.992535481 |
| Nckap1 | High | P28660 | Nck-associated protein 1 OS=Mus musculus OX=10090 GN=Nckap1 PE=1 SV=2 | 0 | 27.882 | 9 | 11 | 14 | 11 | 1128 | 128.7 | 6.62 | 0.984 | 1.026 | 0.959 | -0.02 | 0.02 | 0.04 | 0.01333333 | -0.01 | -0.06 | 0.966709257 | 0.993493688 | 0.865582801 | 0.999652205 | 0.998402594 | 0.9990274 | 0.992535481 |
| Dcaf7 | High | P61963 | DDB1- and CUL4-associated factor 7 OS=Mus musculus OX=10090 GN=Dcaf7 PE=1 SV=1 | 0.001 | 6.088 | 5 | 2 | 2 | 2 | 342 | 38.9 | 5.52 | 1.013 | 1 | 1.013 | 0.02 | -0.02 | 0 | 0 | -0.01 | 0.02 | 0.994192467 | 0.934947566 | 0.908497933 | 0.999652205 | 0.998402594 | 0.9990274 | 0.992535481 |
| Crmp1 | High | P97427 | Dihydropyrimidinase-related protein 1 OS=Mus musculus OX=10090 GN=Crmp1 PE=1 SV=1 | 0 | 107.914 | 35 | 18 | 39 | 12 | 572 | 62.1 | 7.12 | 0.926 | 1.097 | 0.844 | -0.11 | 0.11 | 0.13 | 0.04333333 | -0.01 | -0.24 | 0.828126227 | 0.842511681 | 0.31889966 | 0.999652205 | 0.998402594 | 0.9990274 | 0.904943958 |
| 0610009B | Medium | Q8R3W2 | RIKEN cDNA 0610009B22 gene OS=Mus musculus OX=10090 GN=0610009B22Rik PE=2 SV=1 | 0.014 | 2.254 | 5 | 1 | 1 | 1 | 140 | 16.4 | 6.61 | 1.07 | 0.947 | 1.13 | 0.1 | -0.1 | -0.08 | -0.0266667 | -0.01 | 0.18 | 0.866739377 | 0.799925229 | 0.421761589 | 0.999652205 | 0.998402594 | 0.9990274 | 0.939192266 |
| Pip4k2b | High | Q80XI4 | Phosphatidylinositol 5-phosphate 4-kinase type-2 beta OS=Mus musculus OX=10090 GN=Pip4k2b | 0 | 20.132 | 16 | 6 | 9 | 3 | 416 | 47.3 | 7.33 | 0.938 | 1.078 | 0.871 | -0.09 | 0.09 | 0.11 | 0.03666667 | -0.01 | -0.2 | 0.725386975 | 0.88560842 | 0.433617659 | 0.999652205 | 0.998402594 | 0.9990274 | 0.948052758 |
| Actb | High | P60710 | Actin, cytoplasmic 1 OS=Mus musculus OX=10090 GN=Actb PE=1 SV=1 | 0 | 346.015 | 69 | 22 | 272 | 10 | 375 | 41.7 | 5.48 | 0.945 | 1.075 | 0.879 | -0.08 | 0.08 | 0.1 | 0.03333333 | -0.01 | -0.19 | 0.851476641 | 0.891335347 | 0.464210237 | 0.999652205 | 0.998402594 | 0.9990274 | 0.959744396 |
| Rufy3 | High | Q9D394 | Protein RUFY3 OS=Mus musculus OX=10090 GN=Rufy3 PE=1 SV=1 | 0 | 102.37 | 42 | 17 | 34 | 2 | 469 | 53 | 5.49 | 1.055 | 0.958 | 1.101 | 0.08 | -0.08 | -0.06 | -0.02 | -0.01 | 0.14 | 0.847376788 | 0.827698514 | 0.572772262 | 0.999652205 | 0.998402594 | 0.9990274 | 0.972910028 |
| Arhgdia | High | Q99PT1 | Rho GDP-dissociation inhibitor 1 OS=Mus musculus OX=10090 GN=Arhgdia PE=1 SV=3 | 0 | 23.244 | 20 | 4 | 7 | 4 | 204 | 23.4 | 5.2 | 0.966 | 1.049 | 0.921 | -0.05 | 0.05 | 0.07 | 0.02333333 | -0.01 | -0.12 | 0.854789474 | 0.951619959 | 0.628226303 | 0.999652205 | 0.998402594 | 0.9990274 | 0.981444134 |
| Ap2m1 | High | P84091 | AP-2 complex subunit mu OS=Mus musculus OX=10090 GN=Ap2m1 PE=1 SV=1 | 0 | 105.097 | 47 | 20 | 35 | 20 | 435 | 49.6 | 9.54 | 0.964 | 1.048 | 0.919 | -0.05 | 0.05 | 0.07 | 0.02333333 | -0.01 | -0.12 | 0.903679547 | 0.953004402 | 0.658578284 | 0.999652205 | 0.998402594 | 0.9990274 | 0.985010103 |
| Ap2a1 | High | P17426 | AP-2 complex subunit alpha-1 OS=Mus musculus OX=10090 GN=Ap2a1 PE=1 SV=1 | 0 | 128.677 | 40 | 36 | 51 | 21 | 977 | 107.6 | 7.03 | 0.967 | 1.046 | 0.924 | -0.05 | 0.05 | 0.07 | 0.02333333 | -0.01 | -0.11 | 0.913085016 | 0.957352764 | 0.683509154 | 0.999652205 | 0.998402594 | 0.9990274 | 0.991285312 |
| Sv2a | High | Q9JIS5 | Synaptic vesicle glycoprotein 2A OS=Mus musculus OX=10090 GN=Sv2a PE=1 SV=1 | 0 | 78.275 | 22 | 16 | 36 | 15 | 742 | 82.6 | 5.57 | 0.938 | 1.088 | 0.862 | -0.09 | 0.09 | 0.12 | 0.04 | -0.015 | -0.21 | 0.832506724 | 0.862678757 | 0.392682096 | 0.999652205 | 0.998402594 | 0.9990274 | 0.93064581 |
| Icmt | High | Q9EQK7 | Protein-S-isoprenylcysteine O-methyltransferase OS=Mus musculus OX=10090 GN=Icmt PE=2 SV= | 0.001 | 4.651 | 4 | 2 | 2 | 2 | 283 | 31.8 | 8.18 | 0.942 | 1.087 | 0.867 | -0.09 | 0.09 | 0.12 | 0.04 | -0.015 | -0.21 | 0.719532851 | 0.86605365 | 0.402203647 | 0.999652205 | 0.998402594 | 0.9990274 | 0.93262326 |
| | Medium | F6ZLN8 | Vacuolar protein sorting-associated protein 8 homolog (Fragment) OS=Mus musculus OX=10090 ( | 0.029 | 1.937 | 1 | 1 | 1 | 1 | 964 | 110.1 | 5.72 | 1.08 | 0.945 | 1.143 | 0.11 | -0.11 | -0.08 | -0.0266667 | -0.015 | 0.19 | 0.818863174 | 0.791990799 | 0.438982648 | 0.999652205 | 0.998402594 | 0.9990274 | 0.952819047 |
| | High | H3BJU7 | Guanine nucleotide exchange factor H1 OS=Mus musculus OX=10090 GN=Arhgef2 PE=1 SV=1 | 0.001 | 5.139 | 2 | 2 | 2 | 2 | 956 | 108.5 | 6.87 | 0.931 | 1.092 | 0.853 | -0.1 | 0.1 | 0.13 | 0.04333333 | -0.015 | -0.23 | 0.742271116 | 0.87584623 | 0.454851517 | 0.999652205 | 0.998402594 | 0.9990274 | 0.959744396 |
| Glra2 | Medium | Q7TNC8 | Glycine receptor subunit alpha-2 OS=Mus musculus OX=10090 GN=Glra2 PE=1 SV=1 | 0.041 | 1.675 | 3 | 1 | 1 | 1 | 452 | 52 | 8.9 | 1.071 | 0.955 | 1.121 | 0.1 | -0.1 | -0.07 | -0.0233333 | -0.015 | 0.16 | 0.848528524 | 0.810343734 | 0.506638117 | 0.999652205 | 0.998402594 | 0.9990274 | 0.964378027 |
| Sfxn1 | High | Q99JR1 | Sideroflexin-1 OS=Mus musculus OX=10090 GN=Sfxn1 PE=1 SV=3 | 0 | 23.743 | 28 | 5 | 8 | 5 | 322 | 35.6 | 9.23 | 0.955 | 1.069 | 0.894 | -0.07 | 0.07 | 0.1 | 0.03333333 | -0.015 | -0.16 | 0.847374302 | 0.905500274 | 0.527688382 | 0.999652205 | 0.998402594 | 0.9990274 | 0.965649072 |
| | High | A0A494BB | Eukaryotic translation initiation factor 4C OS=Mus musculus OX=10090 GN=Eif1a PE=1 SV=1 | 0.001 | 6.523 | 10 | 2 | 2 | 2 | 254 | 28.6 | 7.44 | 1.057 | 0.965 | 1.095 | 0.08 | -0.08 | -0.05 | -0.0166667 | -0.015 | 0.13 | 0.907768832 | 0.846298182 | 0.531991906 | 0.999652205 | 0.998402594 | 0.9990274 | 0.96694628 |
| | High | F6U6X3 | Dual-specificity kinase OS=Mus musculus OX=10090 GN=Dyrk1a PE=1 SV=1 | 0.001 | 6.273 | 2 | 1 | 1 | 1 | 725 | 81.5 | 8.79 | 1.06 | 0.968 | 1.095 | 0.08 | -0.08 | -0.05 | -0.0166667 | -0.015 | 0.13 | 0.895970544 | 0.841297933 | 0.577164505 | 0.999652205 | 0.998402594 | 0.9990274 | 0.972910028 |
| Rab3gap2 | High | A0A0A6YV | Rab3 GTPase-activating protein non-catalytic subunit OS=Mus musculus OX=10090 GN=Rab3gap2 | 0.001 | 6.231 | 1 | 2 | 2 | 2 | 1367 | 152.4 | 6.18 | 1.014 | 1.009 | 1.005 | 0.02 | -0.02 | 0.01 | 0.00333333 | -0.015 | 0.01 | 0.990674016 | 0.9352286 | 0.908881511 | 0.999652205 | 0.998402594 | 0.9990274 | 0.992535481 |
| | High | D6RGR9 | Double-stranded RNA-binding protein Staufen homolog 2 OS=Mus musculus OX=10090 GN=Stau2 | 0.001 | 5.016 | 8 | 2 | 2 | 2 | 112 | 12.5 | 9.8 | 1.015 | 1.005 | 1.01 | 0.02 | -0.02 | 0.01 | 0.00333333 | -0.015 | 0.01 | 0.983607693 | 0.948200854 | 0.918325944 | 0.999652205 | 0.998402594 | 0.9990274 | 0.992535481 |
| Slc5a5 | High | G3X8P5 | Sodium/iodide cotransporter OS=Mus musculus OX=10090 GN=Slc5a5 PE=3 SV=1 | 0.003 | 3.949 | 3 | 2 | 2 | 2 | 618 | 65.5 | 7.49 | 1.036 | 0.984 | 1.053 | 0.05 | -0.05 | -0.02 | -0.0066667 | -0.015 | 0.07 | 0.969287052 | 0.881990557 | 0.70851938 | 0.999652205 | 0.998402594 | 0.9990274 | 0.992507514 |
| Ap1s2 | Medium | Q3TIV9 | AP complex subunit sigma OS=Mus musculus OX=10090 GN=Ap1s2 PE=1 SV=1 | 0.01 | 2.621 | 6 | 1 | 1 | 1 | 157 | 18.6 | 5.47 | 1.038 | 0.989 | 1.05 | 0.05 | -0.05 | -0.02 | -0.0066667 | -0.015 | 0.07 | 0.924691887 | 0.907053677 | 0.762935709 | 0.999652205 | 0.998402594 | 0.9990274 | 0.992535481 |
| Psme1 | High | P97371 | Proteasome activator complex subunit 1 OS=Mus musculus OX=10090 GN=Psme1 PE=1 SV=2 | 0 | 10.986 | 20 | 4 | 4 | 4 | 249 | 28.7 | 5.97 | 1.048 | 0.975 | 1.075 | 0.07 | -0.07 | -0.04 | -0.0133333 | -0.015 | 0.1 | 0.846935353 | 0.87005919 | 0.678597212 | 0.999652205 | 0.998402594 | 0.9990274 | 0.990892882 |
| Slc1a1 | High | P51906 | Excitatory amino acid transporter 3 OS=Mus musculus OX=10090 GN=Slc1a1 PE=1 SV=2 | 0 | 48.479 | 20 | 7 | 13 | 7 | 523 | 56.7 | 6.11 | 0.949 | 1.088 | 0.873 | -0.08 | 0.08 | 0.12 | 0.04 | -0.02 | -0.2 | 0.826335267 | 0.863070827 | 0.461370278 | 0.999652205 | 0.998402594 | 0.9990274 | 0.959744396 |
| Insr | High | P15208 | Insulin receptor OS=Mus musculus OX=10090 GN=Insr PE=1 SV=2 | 0 | 39.51 | 9 | 13 | 14 | 11 | 1372 | 155.5 | 5.95 | 0.953 | 1.077 | 0.885 | -0.07 | 0.07 | 0.11 | 0.03666667 | -0.02 | -0.18 | 0.931770557 | 0.888141918 | 0.492027384 | 0.999652205 | 0.998402594 | 0.9990274 | 0.964378027 |
| | Medium | F6SH71 | BTB/POZ domain-containing protein KCTD2 (Fragment) OS=Mus musculus OX=10090 GN=Kctd2 F | 0.033 | 1.838 | 3 | 1 | 1 | 1 | 278 | 30.1 | 5.29 | 0.945 | 1.086 | 0.87 | -0.08 | 0.08 | 0.12 | 0.04 | -0.02 | -0.2 | 0.793118088 | 0.884753098 | 0.521896827 | 0.999652205 | 0.998402594 | 0.9990274 | 0.964378027 |
| Rab18 | High | P35293 | Ras-related protein Rab-18 OS=Mus musculus OX=10090 GN=Rab18 PE=1 SV=2 | 0 | 128.173 | 74 | 15 | 43 | 15 | 206 | 23 | 5.36 | 0.966 | 1.064 | 0.908 | -0.05 | 0.05 | 0.09 | 0.03 | -0.02 | -0.14 | 0.910065607 | 0.917175429 | 0.602286155 | 0.999652205 | 0.998402594 | 0.9990274 | 0.97826833 |
| Impact | High | O55091 | Protein IMPACT OS=Mus musculus OX=10090 GN=Impact PE=1 SV=2 | 0.001 | 5.391 | 3 | 1 | 1 | 1 | 318 | 36.3 | 5.05 | 1.044 | 0.985 | 1.061 | 0.06 | -0.06 | -0.02 | -0.0066667 | -0.02 | 0.08 | 0.931686979 | 0.876258323 | 0.667989474 | 0.999652205 | 0.998402594 | 0.9990274 | 0.987489386 |
| Lrp1 | High | A0A0R4J0 | Prolow-density lipoprotein receptor-related protein 1 OS=Mus musculus OX=10090 GN=Lrp1 PE=1 | 0 | 184.962 | 13 | 51 | 66 | 51 | 4545 | 504.4 | 5.36 | 0.986 | 1.042 | 0.946 | -0.02 | 0.02 | 0.06 | 0.02 | -0.02 | -0.08 | 0.963520948 | 0.968078868 | 0.796450162 | 0.999652205 | 0.998402594 | 0.9990274 | 0.992535481 |
| Cpne6 | High | Q9Z140 | Copine-6 OS=Mus musculus OX=10090 GN=Cpne6 PE=1 SV=1 | 0 | 18.181 | 9 | 5 | 7 | 4 | 557 | 61.7 | 5.59 | 0.971 | 1.06 | 0.916 | -0.04 | 0.04 | 0.08 | 0.02666667 | -0.02 | -0.13 | 0.893360043 | 0.924856992 | 0.639270607 | 0.999652205 | 0.998402594 | 0.9990274 | 0.983641683 |
| | High | F6RJV6 | LanC-like protein 2 (Fragment) OS=Mus musculus OX=10090 GN=Lancl2 PE=1 SV=1 | 0 | 77.441 | 44 | 14 | 21 | 14 | 441 | 49.7 | 7.14 | 1.033 | 0.996 | 1.037 | 0.05 | -0.05 | -0.01 | -0.0033333 | -0.02 | 0.05 | 0.947684338 | 0.920747735 | 0.802692934 | 0.999652205 | 0.998402594 | 0.9990274 | 0.992535481 |
| Dpysl4 | High | O35098 | Dihydropyrimidinase-related protein 4 OS=Mus musculus OX=10090 GN=Dpysl4 PE=1 SV=1 | 0 | 104.576 | 49 | 21 | 33 | 19 | 572 | 61.9 | 6.98 | 0.995 | 1.037 | 0.96 | -0.01 | 0.01 | 0.05 | 0.01666667 | -0.02 | -0.06 | 0.990329235 | 0.979736399 | 0.870641697 | 0.999652205 | 0.998402594 | 0.9990274 | 0.992535481 |
| Cdk5 | High | P49615 | Cyclin-dependent-like kinase 5 OS=Mus musculus OX=10090 GN=Cdk5 PE=1 SV=1 | 0 | 10.475 | 9 | 3 | 6 | 2 | 292 | 33.3 | 7.66 | 1.024 | 1.009 | 1.015 | 0.03 | -0.03 | 0.01 | 0.00333333 | -0.02 | 0.02 | 0.947976141 | 0.953571953 | 0.880232543 | 0.999652205 | 0.998402594 | 0.9990274 | 0.992535481 |
| Dtna | High | A0A1Y7VL | Dystrobrevin OS=Mus musculus OX=10090 GN=Dtna PE=1 SV=1 | 0 | 17.06 | 7 | 6 | 6 | 6 | 746 | 84 | 6.83 | 1 | 1.028 | 0.973 | 0 | 0 | 0.04 | 0.01333333 | -0.02 | -0.04 | 0.951895556 | 0.994105791 | 0.892967687 | 0.999652205 | 0.998402594 | 0.9990274 | 0.992535481 |
| Vps53 | High | Q8CCB4 | Vacuolar protein sorting-associated protein 53 homolog OS=Mus musculus OX=10090 GN=Vps53 | 0 | 16.547 | 7 | 5 | 6 | 5 | 832 | 94.4 | 6.61 | 0.913 | 1.121 | 0.815 | -0.13 | 0.13 | 0.17 | 0.05666667 | -0.02 | -0.3 | 0.667121207 | 0.800027753 | 0.257595715 | 0.999652205 | 0.998402594 | 0.9990274 | 0.85616924 |
| | High | A0A1D5RL | Glyceraldehyde-3-phosphate dehydrogenase OS=Mus musculus OX=10090 GN=Gm10358 PE=1 SV | 0 | 794.733 | 92 | 33 | 904 | 1 | 333 | 35.8 | 8.25 | 0.933 | 1.102 | 0.847 | -0.1 | 0.1 | 0.14 | 0.04666667 | -0.02 | -0.24 | 0.735152875 | 0.821857794 | 0.347479069 | 0.999652205 | 0.998402594 | 0.9990274 | 0.909421216 |
| | High | A0A1D5RN | Arf-GAP with GTPase, ANK repeat and PH domain-containing protein 3 OS=Mus musculus OX=10( | 0.001 | 6.774 | 5 | 2 | 2 | 1 | 396 | 42.4 | 8.06 | 1.069 | 0.962 | 1.112 | 0.1 | -0.1 | -0.06 | -0.02 | -0.02 | 0.15 | 0.793457222 | 0.836622683 | 0.506531073 | 0.999652205 | 0.998402594 | 0.9990274 | 0.964378027 |
| Pea15a | High | Q62048 | Astrocytic phosphoprotein PEA-15 OS=Mus musculus OX=10090 GN=Pea15 PE=1 SV=1 | 0 | 17.146 | 35 | 3 | 4 | 3 | 130 | 15 | 5.02 | 0.935 | 1.11 | 0.843 | -0.1 | 0.1 | 0.15 | 0.05 | -0.025 | -0.25 | 0.755470692 | 0.815487784 | 0.314187296 | 0.999652205 | 0.998402594 | 0.9990274 | 0.904943958 |
| Galnt7 | High | Q80VA0 | N-acetylgalactosaminyltransferase 7 OS=Mus musculus OX=10090 GN=Galnt7 PE=1 SV=2 | 0.006 | 3.12 | 2 | 1 | 1 | 1 | 657 | 75.4 | 7.93 | 1.067 | 0.97 | 1.099 | 0.09 | -0.09 | -0.04 | -0.0133333 | -0.025 | 0.14 | 0.839376126 | 0.858958493 | 0.563964607 | 0.999652205 | 0.998402594 | 0.9990274 | 0.972910028 |
| Rap2c | High | Q8BU31 | Ras-related protein Rap-2c OS=Mus musculus OX=10090 GN=Rap2c PE=1 SV=1 | 0 | 33.429 | 45 | 6 | 12 | 2 | 183 | 20.7 | 4.94 | 0.992 | 1.045 | 0.949 | -0.01 | 0.01 | 0.06 | 0.02 | -0.025 | -0.07 | 0.913763801 | 0.980441327 | 0.800137002 | 0.999652205 | 0.998402594 | 0.9990274 | 0.992535481 |
| Rab5b | High | P61021 | Ras-related protein Rab-5B OS=Mus musculus OX=10090 GN=Rab5b PE=1 SV=1 | 0 | 74.953 | 54 | 9 | 37 | 6 | 215 | 23.7 | 8.13 | 0.995 | 1.041 | 0.956 | -0.01 | 0.01 | 0.06 | 0.02 | -0.025 | -0.07 | 0.989064946 | 0.969704953 | 0.847252893 | 0.999652205 | 0.998402594 | 0.9990274 | 0.992535481 |
| Stxbp5 | High | D3Z079 | Syntaxin-binding protein 5 OS=Mus musculus OX=10090 GN=Stxbp5 PE=1 SV=1 | 0 | 38.172 | 11 | 9 | 10 | 9 | 1116 | 123.5 | 7.11 | 0.961 | 1.076 | 0.894 | -0.06 | 0.06 | 0.11 | 0.03666667 | -0.025 | -0.16 | 0.826346542 | 0.890097881 | 0.494946584 | 0.999652205 | 0.998402594 | 0.9990274 | 0.964378027 |
| Rab22a | High | A2ARZ7 | Ras-related protein Rab-22A OS=Mus musculus OX=10090 GN=Rab22a PE=1 SV=1 | 0 | 22.532 | 33 | 5 | 10 | 5 | 187 | 20.9 | 7.74 | 0.968 | 1.073 | 0.902 | -0.05 | 0.05 | 0.1 | 0.03333333 | -0.025 | -0.15 | 0.957923898 | 0.896859683 | 0.572391983 | 0.999652205 | 0.998402594 | 0.9990274 | 0.972910028 |
| Emc7 | Medium | Q9EP72 | ER membrane protein complex subunit 7 OS=Mus musculus OX=10090 GN=Emc7 PE=1 SV=1 | 0.01 | 2.581 | 4 | 1 | 1 | 1 | 241 | 26.3 | 9.23 | 1.056 | 0.981 | 1.077 | 0.08 | -0.08 | -0.03 | -0.01 | -0.025 | 0.11 | 0.896720521 | 0.862057094 | 0.665410758 | 0.999652205 | 0.998402594 | 0.9990274 | 0.987366731 |
| 4632415K | High | Q8K0P3 | MTOR-associated protein MEAK7 OS=Mus musculus OX=10090 GN=Meak7 PE=2 SV=1 | 0 | 15.855 | 7 | 2 | 3 | 2 | 455 | 50.8 | 6.52 | 1.033 | 0.999 | 1.033 | 0.05 | -0.05 | 0 | 0 | -0.025 | 0.05 | 0.937266774 | 0.929634937 | 0.801728553 | 0.999652205 | 0.998402594 | 0.9990274 | 0.992535481 |
| Atp6v1e1 | High | P50518 | V-type proton ATPase subunit E 1 OS=Mus musculus OX=10090 GN=Atp6v1e1 PE=1 SV=2 | 0 | 99.828 | 58 | 17 | 39 | 17 | 226 | 26.1 | 8.43 | 1.027 | 1.007 | 1.02 | 0.04 | -0.04 | 0.01 | 0.00333333 | -0.025 | 0.03 | 0.927154852 | 0.947548282 | 0.82036177 | 0.999652205 | 0.998402594 | 0.9990274 | 0.992535481 |
| | High | B2FDH0 | Ubiquitin-conjugating enzyme E2 E3 (Fragment) OS=Mus musculus OX=10090 GN=Ube2e3 PE=1 S | 0.005 | 3.682 | 8 | 1 | 1 | 1 | 103 | 11.1 | 5.27 | 1.008 | 1.032 | 0.977 | 0.01 | -0.01 | 0.04 | 0.01333333 | -0.025 | -0.03 | 0.968840776 | 0.986967523 | 0.893473616 | 0.999652205 | 0.998402594 | 0.9990274 | 0.992535481 |
| Tomm34 | High | Q9CYG7 | Mitochondrial import receptor subunit TOM34 OS=Mus musculus OX=10090 GN=Tomm34 PE=1 S | 0.005 | 3.57 | 3 | 1 | 1 | 1 | 309 | 34.3 | 9.14 | 1.009 | 1.031 | 0.978 | 0.01 | -0.01 | 0.04 | 0.01333333 | -0.025 | -0.03 | 0.972561998 | 0.982540933 | 0.983230453 | 0.999652205 | 0.998402594 | 0.9990274 | 0.997765878 |
| | High | D3Z5N2 | Stathmin OS=Mus musculus OX=10090 GN=Stmn1 PE=1 SV=1 | 0 | 14.364 | 28 | 4 | 5 | 4 | 147 | 17 | 9.09 | 0.932 | 1.121 | 0.832 | -0.1 | 0.1 | 0.16 | 0.05333333 | -0.03 | -0.27 | 0.737471573 | 0.792352846 | 0.27458552 | 0.999652205 | 0.998402594 | 0.9990274 | 0.876067403 |
| Atp6v0d1 | High | P51863 | V-type proton ATPase subunit d 1 OS=Mus musculus OX=10090 GN=Atp6v0d1 PE=1 SV=2 | 0 | 52.714 | 36 | 12 | 23 | 12 | 351 | 40.3 | 5 | 0.952 | 1.097 | 0.868 | -0.07 | 0.07 | 0.13 | 0.04333333 | -0.03 | -0.2 | 0.871532504 | 0.842924184 | 0.416589655 | 0.999652205 | 0.998402594 | 0.9990274 | 0.936397673 |
| Ubr4 | High | A2AN08 | E3 ubiquitin-protein ligase UBR4 OS=Mus musculus OX=10090 GN=Ubr4 PE=1 SV=1 | 0 | 8.57 | 1 | 3 | 3 | 3 | 5180 | 571.9 | 6.06 | 1.097 | 0.956 | 1.147 | 0.13 | -0.13 | -0.07 | -0.0233333 | -0.03 | 0.2 | 0.749948609 | 0.822766263 | 0.429589461 | 0.999652205 | 0.998402594 | 0.9990274 | 0.944356224 |
| Abcc1 | High | O35379 | Multidrug resistance-associated protein 1 OS=Mus musculus OX=10090 GN=Abcc1 PE=1 SV=1 | 0 | 33.053 | 5 | 6 | 9 | 6 | 1528 | 171.1 | 7.36 | 0.983 | 1.054 | 0.933 | -0.02 | 0.02 | 0.08 | 0.02666667 | -0.03 | -0.1 | 0.989845997 | 0.940351193 | 0.720632285 | 0.999652205 | 0.998402594 | 0.9990274 | 0.992535481 |
| Carkd; Nax | High | Q9CZ42 | ATP-dependent (S)-NAD(P)H-hydrate dehydratase OS=Mus musculus OX=10090 GN=Naxd PE=1 S | 0 | 14.312 | 16 | 4 | 5 | 4 | 343 | 36.7 | 7.77 | 1.043 | 1.001 | 1.041 | 0.06 | -0.06 | 0 | 0 | -0.03 | 0.06 | 0.83272739 | 0.934779256 | 0.79174933 | 0.999652205 | 0.998402594 | 0.9990274 | 0.992535481 |
| | High | H3BIW6 | Phosphoinositide phospholipase C OS=Mus musculus OX=10090 GN=Plch2 PE=1 SV=1 | 0 | 25.6 | 7 | 8 | 9 | 7 | 1137 | 125.7 | 6.79 | 1.075 | 0.973 | 1.105 | 0.1 | -0.1 | -0.04 | -0.0133333 | -0.03 | 0.14 | 0.834370597 | 0.864633644 | 0.565119144 | 0.999652205 | 0.998402594 | 0.9990274 | 0.972910028 |
| Arpc1a | High | Q9R0Q6 | Actin-related protein 2/3 complex subunit 1A OS=Mus musculus OX=10090 GN=Arpc1a PE=1 SV= | 0 | 32.96 | 28 | 10 | 17 | 10 | 370 | 41.6 | 8.18 | 1.061 | 0.986 | 1.076 | 0.09 | -0.09 | -0.02 | -0.0066667 | -0.035 | 0.11 | 0.834029289 | 0.898032164 | 0.569154182 | 0.999652205 | 0.998402594 | 0.9990274 | 0.972910028 |
| Erc2 | High | Q3UHT7 | ERC protein 2 OS=Mus musculus OX=10090 GN=Erc2 PE=1 SV=1 | 0.001 | 5.311 | 1 | 1 | 1 | 1 | 977 | 113 | 7.09 | 1.062 | 0.988 | 1.075 | 0.09 | -0.09 | -0.02 | -0.0066667 | -0.035 | 0.1 | 0.888451152 | 0.879996024 | 0.702919645 | 0.999652205 | 0.998402594 | 0.9990274 | 0.992013705 |
| Ppfia2 | High | Q8BSS9 | Liprin-alpha-2 OS=Mus musculus OX=10090 GN=Ppfia2 PE=1 SV=2 | 0 | 7.97 | 3 | 4 | 4 | 2 | 1257 | 143.1 | 6.11 | 1.043 | 1.007 | 1.036 | 0.06 | -0.06 | 0.01 | 0.00333333 | -0.035 | 0.05 | 0.894901119 | 0.948459298 | 0.78402477 | 0.999652205 | 0.998402594 | 0.9990274 | 0.992535481 |
| Mpp5 | High | Q9JLB2 | Protein PALS1 OS=Mus musculus OX=10090 GN=Pals1 PE=1 SV=1 | 0 | 28.986 | 14 | 7 | 8 | 7 | 675 | 77.2 | 6.09 | 1.008 | 1.041 | 0.968 | 0.01 | -0.01 | 0.06 | 0.02 | -0.035 | -0.05 | 0.977591001 | 0.970144407 | 0.814055792 | 0.999652205 | 0.998402594 | 0.9990274 | 0.992535481 |
| Agap1 | High | Q8BXK8 | Arf-GAP with GTPase, ANK repeat and PH domain-containing protein 1 OS=Mus musculus OX=10( | 0 | 8.291 | 4 | 3 | 3 | 2 | 857 | 94.4 | 7.94 | 1.006 | 1.045 | 0.963 | 0.01 | -0.01 | 0.06 | 0.02 | -0.035 | -0.05 | 0.95956268 | 0.960167065 | 0.855598329 | 0.999652205 | 0.998402594 | 0.9990274 | 0.992535481 |
| Snca | High | O55042 | Alpha-synuclein OS=Mus musculus OX=10090 GN=Snca PE=1 SV=2 | 0 | 44.101 | 60 | 8 | 19 | 5 | 140 | 14.5 | 4.77 | 1.005 | 1.044 | 0.963 | 0.01 | -0.01 | 0.06 | 0.02 | -0.035 | -0.05 | 0.979627811 | 0.964207435 | 0.886264299 | 0.999652205 | 0.998402594 | 0.9990274 | 0.992535481 |
| Gad2 | High | P48320 | Glutamate decarboxylase 2 OS=Mus musculus OX=10090 GN=Gad2 PE=1 SV=1 | 0 | 31.614 | 19 | 7 | 8 | 6 | 585 | 65.2 | 6.9 | 0.957 | 1.095 | 0.874 | -0.06 | 0.06 | 0.13 | 0.04333333 | -0.035 | -0.19 | 0.81484804 | 0.846418171 | 0.41153074 | 0.999652205 | 0.998402594 | 0.9990274 | 0.936397673 |
| Acot10 | High | Q32MW3 | Acyl-coenzyme A thioesterase 10, mitochondrial OS=Mus musculus OX=10090 GN=Acot10 PE=2 S | 0.006 | 3.432 | 4 | 2 | 2 | 2 | 439 | 50.5 | 7.88 | 0.958 | 1.092 | 0.877 | -0.06 | 0.06 | 0.13 | 0.04333333 | -0.035 | -0.19 | 0.803022367 | 0.864089582 | 0.499233332 | 0.999652205 | 0.998402594 | 0.9990274 | 0.964378027 |
| | High | E9Q1X8 | Voltage-dependent calcium channel subunit alpha-2/delta-1 OS=Mus musculus OX=10090 GN=Ca | 0 | 38.119 | 11 | 11 | 12 | 10 | 1091 | 123.1 | 5.17 | 0.973 | 1.076 | 0.905 | -0.04 | 0.04 | 0.11 | 0.03666667 | -0.035 | -0.14 | 0.868745094 | 0.889605381 | 0.520263676 | 0.999652205 | 0.998402594 | 0.9990274 | 0.964378027 |
| Cald1 | High | Q8VCQ8 | Caldesmon 1 OS=Mus musculus OX=10090 GN=Cald1 PE=1 SV=1 | 0 | 63.164 | 27 | 13 | 19 | 13 | 530 | 60.4 | 7.37 | 0.973 | 1.079 | 0.902 | -0.04 | 0.04 | 0.11 | 0.03666667 | -0.035 | -0.15 | 0.973142897 | 0.883092997 | 0.572281805 | 0.999652205 | 0.998402594 | 0.9990274 | 0.972910028 |
| Lsm14a | Medium | Q8K2F8 | Protein LSM14 homolog A OS=Mus musculus OX=10090 GN=Lsm14a PE=1 SV=1 | 0.014 | 2.187 | 2 | 1 | 1 | 1 | 462 | 50.5 | 9.52 | 1.071 | 0.978 | 1.095 | 0.1 | -0.1 | -0.03 | -0.01 | -0.035 | 0.13 | 0.830777014 | 0.879300102 | 0.587235657 | 0.999652205 | 0.998402594 | 0.9990274 | 0.973095093 |
| | Medium | D3Z600 | PTS1-BP OS=Mus musculus OX=10090 GN=Pex5 PE=1 SV=1 | 0.012 | 2.368 | 3 | 1 | 1 | 1 | 632 | 69.8 | 4.51 | 1.055 | 0.991 | 1.064 | 0.08 | -0.08 | -0.01 | -0.0033333 | -0.035 | 0.09 | 0.900275375 | 0.897216502 | 0.706391034 | 0.999652205 | 0.998402594 | 0.9990274 | 0.992485323 |
| | High | D3Z2D8 | CSC1-like protein 2 (Fragment) OS=Mus musculus OX=10090 GN=Tmem63b PE=1 SV=8 | 0 | 84.534 | 47 | 15 | 39 | 1 | 309 | 34.8 | 8.43 | 1.059 | 0.99 | 1.069 | 0.08 | -0.08 | -0.01 | -0.0033333 | -0.035 | 0.1 | 0.877792772 | 0.908481688 | 0.717421026 | 0.999652205 | 0.998402594 | 0.9990274 | 0.992535481 |
| Hnmt | High | Q91VF2 | Histamine N-methyltransferase OS=Mus musculus OX=10090 GN=Hnmt PE=1 SV=1 | 0.001 | 4.429 | 5 | 2 | 3 | 2 | 295 | 33.6 | 5.06 | 0.994 | 1.055 | 0.942 | -0.01 | 0.01 | 0.08 | 0.02666667 | -0.035 | -0.09 | 0.892453266 | 0.942204945 | 0.749323587 | 0.999652205 | 0.998402594 | 0.9990274 | 0.992535481 |
| Atp2b4 | High | Q6Q477 | Plasma membrane calcium-transporting ATPase 4 OS=Mus musculus OX=10090 GN=Atp2b4 PE=1 | 0 | 139.361 | 22 | 22 | 39 | 9 | 1205 | 133 | 6.13 | 0.999 | 1.05 | 0.951 | 0 | 0 | 0.07 | 0.02333333 | -0.035 | -0.07 | 0.929742685 | 0.948092297 | 0.75190813 | 0.999652205 | 0.998402594 | 0.9990274 | 0.992535481 |
| Dbnl | High | Q62418 | Drebrin-like protein OS=Mus musculus OX=10090 GN=Dbnl PE=1 SV=2 | 0 | 28.922 | 20 | 6 | 6 | 6 | 436 | 48.7 | 4.92 | 0.997 | 1.047 | 0.952 | 0 | 0 | 0.07 | 0.02333333 | -0.035 | -0.07 | 0.956141638 | 0.955827483 | 0.77511111 | 0.999652205 | 0.998402594 | 0.9990274 | 0.992535481 |
| Pgd | High | Q9DCD0 | 6-phosphogluconate dehydrogenase, decarboxylating OS=Mus musculus OX=10090 GN=Pgd PE=1 | 0 | 180.676 | 47 | 23 | 61 | 23 | 483 | 53.2 | 7.23 | 1.033 | 1.013 | 1.02 | 0.05 | -0.05 | 0.02 | 0.00666667 | -0.035 | 0.03 | 0.909925871 | 0.96370349 | 0.82116532 | 0.999652205 | 0.998402594 | 0.9990274 | 0.992535481 |
| Endod1 | High | Q8C522 | Endonuclease domain-containing 1 protein OS=Mus musculus OX=10090 GN=Endod1 PE=1 SV=2 | 0 | 14.846 | 13 | 4 | 4 | 4 | 501 | 55.2 | 6.16 | 1.034 | 1.011 | 1.022 | 0.05 | -0.05 | 0.02 | 0.00666667 | -0.035 | 0.03 | 0.978136707 | 0.94016026 | 0.823943461 | 0.999652205 | 0.998402594 | 0.9990274 | 0.992535481 |
| Negr1 | High | A0A4W9 | Neuronal growth regulator 1 OS=Mus musculus OX=10090 GN=Negr1 PE=1 SV=1 | 0 | 91.349 | 40 | 12 | 29 | 12 | 329 | 36.1 | 6.86 | 1.028 | 1.021 | 1.007 | 0.04 | -0.04 | 0.03 | 0.01 | -0.035 | 0.01 | 0.923490187 | 0.98119143 | 0.882955099 | 0.999652205 | 0.998402594 | 0.9990274 | 0.992535481 |
| Ipo7 | High | Q9EPL8 | Importin-7 OS=Mus musculus OX=10090 GN=Ipo7 PE=1 SV=2 | 0 | 33.083 | 9 | 8 | 10 | 8 | 1038 | 119.4 | 4.82 | 1.026 | 1.019 | 1.007 | 0.04 | -0.04 | 0.03 | 0.01 | -0.035 | 0.01 | 0.919867841 | 0.977798231 | 0.891242818 | 0.999652205 | 0.998402594 | 0.9990274 | 0.992535481 |
| Acsl3 | High | Q9CZW4 | Long-chain-fatty-acid--CoA ligase 3 OS=Mus musculus OX=10090 GN=Acsl3 PE=1 SV=2 | 0 | 8.133 | 4 | 3 | 4 | 3 | 720 | 80.4 | 8.54 | 1.032 | 1.014 | 1.018 | 0.05 | -0.05 | 0.02 | 0.00666667 | -0.035 | 0.03 | 0.965796386 | 0.965503996 | 0.952710272 | 0.999652205 | 0.998402594 | 0.9990274 | 0.992535481 |
| Cryz | High | P47199 | Quinone oxidoreductase OS=Mus musculus OX=10090 GN=Cryz PE=1 SV=1 | 0 | 78.834 | 58 | 16 | 23 | 16 | 331 | 35.2 | 8.07 | 1.112 | 0.95 | 1.171 | 0.15 | -0.15 | -0.07 | -0.0233333 | -0.04 | 0.23 | 0.717720611 | 0.807328781 | 0.263321227 | 0.999652205 | 0.998402594 | 0.9990274 | 0.859470827 |
| | Medium | A0A286YE | Ubiquitin-conjugating enzyme E2 E1 (Fragment) OS=Mus musculus OX=10090 GN=Ube2e1 PE=1 S | 0.047 | 1.536 | 5 | 1 | 1 | 1 | 148 | 16.4 | 8.07 | 0.97 | 1.089 | 0.891 | -0.04 | 0.04 | 0.12 | 0.04 | -0.04 | -0.17 | 0.948030508 | 0.860968556 | 0.535867586 | 0.999652205 | 0.998402594 | 0.9990274 | 0.96694628 |
| Wdr37 | High | Q8CBE3 | WD repeat-containing protein 37 OS=Mus musculus OX=10090 GN=Wdr37 PE=1 SV=1 | 0 | 10.884 | 7 | 4 | 4 | 4 | 496 | 55 | 7.23 | 1.081 | 0.981 | 1.101 | 0.11 | -0.11 | -0.03 | -0.01 | -0.04 | 0.14 | 0.731496597 | 0.885308542 | 0.551615564 | 0.999652205 | 0.998402594 | 0.9990274 | 0.96694628 |
| Coro1c | High | Q9WUM4 | Coronin-1C OS=Mus musculus OX=10090 GN=Coro1c PE=1 SV=2 | 0 | 58.19 | 29 | 15 | 22 | 14 | 474 | 53.1 | 7.08 | 0.977 | 1.082 | 0.903 | -0.03 | 0.03 | 0.11 | 0.03666667 | -0.04 | -0.15 | 0.940366947 | 0.87664196 | 0.578413511 | 0.999652205 | 0.998402594 | 0.9990274 | 0.972910028 |
| | High | A0A0G2JG | Eukaryotic translation initiation factor 4E (Fragment) OS=Mus musculus OX=10090 GN=Eif4e PE= | 0.008 | 3.008 | 8 | 2 | 2 | 2 | 183 | 21.4 | 5.2 | 1.064 | 0.996 | 1.068 | 0.09 | -0.09 | -0.01 | -0.0033333 | -0.04 | 0.09 | 0.859038407 | 0.922408061 | 0.656462955 | 0.999652205 | 0.998402594 | 0.9990274 | 0.985010103 |
| Rab5a | High | Q9CQD1 | Ras-related protein Rab-5A OS=Mus musculus OX=10090 GN=Rab5a PE=1 SV=1 | 0 | 88.594 | 68 | 11 | 35 | 7 | 215 | 23.6 | 8.15 | 0.993 | 1.065 | 0.933 | -0.01 | 0.01 | 0.09 | 0.03 | -0.04 | -0.1 | 0.984777049 | 0.913876609 | 0.726304017 | 0.999652205 | 0.998402594 | 0.9990274 | 0.992535481 |
| Cdc42ep4 | High | Q9JM96 | Cdc42 effector protein 4 OS=Mus musculus OX=10090 GN=Cdc42ep4 PE=1 SV=1 | 0 | 8.207 | 5 | 1 | 1 | 1 | 349 | 37.8 | 5.36 | 1.061 | 0.996 | 1.065 | 0.09 | -0.09 | -0.01 | -0.0033333 | -0.04 | 0.09 | 0.887280884 | 0.90781768 | 0.734902177 | 0.999652205 | 0.998402594 | 0.9990274 | 0.992535481 |
| | High | A0A571BE | Disks large homolog 2 OS=Mus musculus OX=10090 GN=Dlg2 PE=1 SV=1 | 0 | 20.334 | 7 | 5 | 5 | 2 | 901 | 99.3 | 6.6 | 0.993 | 1.062 | 0.936 | -0.01 | 0.01 | 0.09 | 0.03 | -0.04 | -0.1 | 0.908453489 | 0.941102336 | 0.759852138 | 0.999652205 | 0.998402594 | 0.9990274 | 0.992535481 |
| Eif2s3x | High | Q9Z0N1 | Eukaryotic translation initiation factor 2 subunit 3, X-linked OS=Mus musculus OX=10090 GN=Eif2 | 0 | 20.653 | 11 | 5 | 6 | 5 | 472 | 51 | 8.4 | 1.055 | 0.999 | 1.056 | 0.08 | -0.08 | 0 | 0 | -0.04 | 0.08 | 0.903875866 | 0.929469978 | 0.780000903 | 0.999652205 | 0.998402594 | 0.9990274 | 0.992535481 |
| Ago2; Eif2 | High | Q8CJG0 | Protein argonaute-2 OS=Mus musculus OX=10090 GN=Ago2 PE=1 SV=3 | 0 | 35.18 | 12 | 8 | 8 | 8 | 860 | 97.2 | 9.19 | 1.01 | 1.051 | 0.961 | 0.01 | -0.01 | 0.07 | 0.02333333 | -0.04 | -0.06 | 0.949596375 | 0.947181703 | 0.793159319 | 0.999652205 | 0.998402594 | 0.9990274 | 0.992535481 |
| Pde12 | High | Q3TIU4 | 2',5'-phosphodiesterase 12 OS=Mus musculus OX=10090 GN=Pde12 PE=1 SV=2 | 0.003 | 3.885 | 3 | 1 | 1 | 1 | 608 | 67.5 | 6.92 | 1.035 | 1.02 | 1.014 | 0.05 | -0.05 | 0.03 | 0.01 | -0.04 | 0.02 | 0.958900089 | 0.959805179 | 0.920994779 | 0.999652205 | 0.998402594 | 0.9990274 | 0.992535481 |
| Ndufb10 | High | Q9DCS9 | NADH dehydrogenase [ubiquinone] 1 beta subcomplex subunit 10 OS=Mus musculus OX=10090 G | 0 | 25.314 | 41 | 6 | 8 | 6 | 176 | 21 | 8.03 | 1.024 | 1.034 | 0.991 | 0.03 | -0.03 | 0.05 | 0.01666667 | -0.04 | -0.01 | 0.937634848 | 0.987659991 | 0.981863762 | 0.999652205 | 0.998402594 | 0.9990274 | 0.997765878 |
| Mrpl12 | High | Q9DB15 | 39S ribosomal protein L12, mitochondrial OS=Mus musculus OX=10090 GN=Mrpl12 PE=1 SV=2 | 0.001 | 7.072 | 13 | 2 | 2 | 2 | 201 | 21.7 | 9.29 | 1.107 | 0.959 | 1.155 | 0.15 | -0.15 | -0.06 | -0.02 | -0.045 | 0.21 | 0.746963145 | 0.83058709 | 0.357921922 | 0.999652205 | 0.998402594 | 0.9990274 | 0.914600864 |
| Eif3a | High | P23116 | Eukaryotic translation initiation factor 3 subunit A OS=Mus musculus OX=10090 GN=Eif3a PE=1 S | 0 | 32.38 | 8 | 9 | 11 | 9 | 1344 | 161.8 | 6.77 | 1.11 | 0.96 | 1.156 | 0.15 | -0.15 | -0.06 | -0.02 | -0.045 | 0.21 | 0.773354891 | 0.833940114 | 0.414372128 | 0.999652205 | 0.998402594 | 0.9990274 | 0.936397673 |
| | High | A0A087W | E3 ubiquitin-protein ligase TRIP12 OS=Mus musculus OX=10090 GN=Trip12 PE=1 SV=1 | 0 | 14.265 | 2 | 3 | 3 | 3 | 1992 | 220.5 | 8.44 | 1.085 | 0.976 | 1.111 | 0.12 | -0.12 | -0.03 | -0.01 | -0.045 | 0.15 | 0.811092268 | 0.850766983 | 0.485228175 | 0.999652205 | 0.998402594 | 0.9990274 | 0.964378027 |
| Pam16 | High | Q9CQV1 | Mitochondrial import inner membrane translocase subunit TIM16 OS=Mus musculus OX=10090 G | 0.001 | 5.084 | 9 | 1 | 1 | 1 | 125 | 13.8 | 9.64 | 1.09 | 0.98 | 1.113 | 0.12 | -0.12 | -0.03 | -0.01 | -0.045 | 0.15 | 0.785701357 | 0.885259679 | 0.522135343 | 0.999652205 | 0.998402594 | 0.9990274 | 0.964378027 |
| Map4; Mta | High | E9PZ43 | Microtubule-associated protein OS=Mus musculus OX=10090 GN=Map4 PE=1 SV=3 | 0 | 210.676 | 36 | 35 | 90 | 8 | 933 | 97.7 | 9.09 | 0.979 | 1.084 | 0.904 | -0.03 | 0.03 | 0.12 | 0.04 | -0.045 | -0.15 | 0.999232464 | 0.872415889 | 0.58006793 | 0.999652205 | 0.998402594 | 0.9990274 | 0.972910028 |

| | | | | | | | | | | | | | | | | | | | | | | | | | | | | |
|---|---|---|---|---|---|---|---|---|---|---|---|---|---|---|---|---|---|---|---|---|---|---|---|---|---|---|---|---|
| Slc5a6 | High | Q5U4D8 | Sodium-dependent multivitamin transporter OS=Mus musculus OX=10090 GN=Slc5a6 PE=1 SV=2 | 0 | 17.757 | 6 | 3 | 5 | 3 | 634 | 68.5 | 7.85 | 1.079 | 0.986 | 1.094 | 0.11 | -0.11 | -0.02 | -0.0066667 | -0.045 | 0.13 | 0.797264163 | 0.898108818 | 0.623320534 | 0.999652205 | 0.998402594 | 0.9990274 | 0.97826833 |
| Pik3r1 | High | P26450 | Phosphatidylinositol 3-kinase regulatory subunit alpha OS=Mus musculus OX=10090 GN=Pik3r1 PE | 0 | 12.879 | 6 | 4 | 4 | 4 | 724 | 83.5 | 6.28 | 1.057 | 1.008 | 1.048 | 0.08 | -0.08 | 0.01 | 0.00333333 | -0.045 | 0.07 | 0.929891479 | 0.932322724 | 0.734289442 | 0.999652205 | 0.998402594 | 0.9990274 | 0.992535481 |
| Pcnxl3; Pc | High | Q8VI59 | Pecanex-like protein 3 OS=Mus musculus OX=10090 GN=Pcnx3 PE=1 SV=2 | 0.006 | 3.203 | 1 | 2 | 2 | 2 | 2028 | 221.4 | 6.62 | 1.006 | 1.056 | 0.953 | 0.01 | -0.01 | 0.08 | 0.02666667 | -0.045 | -0.07 | 0.966664405 | 0.929488933 | 0.781731552 | 0.999652205 | 0.998402594 | 0.9990274 | 0.992535481 |
| Pdxk | High | Q8K183 | Pyridoxal kinase OS=Mus musculus OX=10090 GN=Pdxk PE=1 SV=1 | 0 | 23.967 | 29 | 7 | 10 | 7 | 312 | 35 | 6.29 | 1.02 | 1.041 | 0.98 | 0.03 | -0.03 | 0.06 | 0.02 | -0.045 | -0.03 | 0.985462252 | 0.970323176 | 0.878252112 | 0.999652205 | 0.998402594 | 0.9990274 | 0.992535481 |
| Bcat1 | High | Q8CBC8 | Branched-chain-amino-acid aminotransferase OS=Mus musculus OX=10090 GN=Bcat1 PE=1 SV=1 | 0 | 9.529 | 7 | 4 | 6 | 4 | 453 | 49.9 | 6.77 | 1.026 | 1.032 | 0.994 | 0.04 | -0.04 | 0.05 | 0.01666667 | -0.045 | -0.01 | 0.953769856 | 0.99108336 | 0.969520288 | 0.999652205 | 0.998402594 | 0.9990274 | 0.995598794 |
| Sag | High | P20443 | S-arrestin OS=Mus musculus OX=10090 GN=Sag PE=1 SV=1 | 0 | 1493.881 | 95 | 55 | 2415 | 52 | 403 | 44.9 | 5.8 | 1.105 | 0.965 | 1.146 | 0.14 | -0.14 | -0.05 | -0.0166667 | -0.045 | 0.2 | 0.733592965 | 0.844902077 | 0.328728114 | 0.999652205 | 0.998402594 | 0.9990274 | 0.905686588 |
| Sv2b | High | Q8BG39 | Synaptic vesicle glycoprotein 2B OS=Mus musculus OX=10090 GN=Sv2b PE=1 SV=1 | 0 | 32.786 | 8 | 7 | 12 | 6 | 683 | 77.4 | 5.57 | 0.963 | 1.098 | 0.877 | -0.05 | 0.05 | 0.14 | 0.04666667 | -0.045 | -0.19 | 0.852689145 | 0.840253876 | 0.424584584 | 0.999652205 | 0.998402594 | 0.9990274 | 0.941180024 |
| Iqsec1 | High | A0A1D5RN | IQ motif and SEC7 domain-containing protein 1 OS=Mus musculus OX=10090 GN=Iqsec1 PE=1 SV | 0 | 22.492 | 7 | 7 | 7 | 4 | 1051 | 117.9 | 7.68 | 1.103 | 0.966 | 1.142 | 0.14 | -0.14 | -0.05 | -0.0166667 | -0.045 | 0.19 | 0.709740369 | 0.848237151 | 0.428538118 | 0.999652205 | 0.998402594 | 0.9990274 | 0.943389567 |
| Ctnna2 | High | E0CXB9 | Alpha N-catenin OS=Mus musculus OX=10090 GN=Ctnna2 PE=1 SV=1 | 0 | 21.136 | 8 | 8 | 9 | 4 | 966 | 106.7 | 5.69 | 0.966 | 1.099 | 0.879 | -0.05 | 0.05 | 0.14 | 0.04666667 | -0.045 | -0.19 | 0.880253974 | 0.838109373 | 0.445809681 | 0.999652205 | 0.998402594 | 0.9990274 | 0.958608248 |
| Ap2b1 | High | Q9DBG3 | AP-2 complex subunit beta OS=Mus musculus OX=10090 GN=Ap2b1 PE=1 SV=1 | 0 | 127.719 | 33 | 29 | 47 | 15 | 937 | 104.5 | 5.38 | 0.963 | 1.1 | 0.876 | -0.05 | 0.05 | 0.14 | 0.04666667 | -0.045 | -0.19 | 0.901604091 | 0.837351738 | 0.450610841 | 0.999652205 | 0.998402594 | 0.9990274 | 0.959744396 |
| | Medium | A0A2I3BR | Acyl-CoA-binding domain-containing protein 7 OS=Mus musculus OX=10090 GN=Acbd7 PE=1 SV= | 0.01 | 2.654 | 9 | 1 | 1 | 1 | 89 | 10.1 | 5.55 | 1.087 | 0.986 | 1.102 | 0.12 | -0.12 | -0.02 | -0.0066667 | -0.05 | 0.14 | 0.775990819 | 0.899252681 | 0.553423597 | 0.999652205 | 0.998402594 | 0.9990274 | 0.96775057 |
| Mob4 | High | Q6PEB6 | MOB-like protein phocein OS=Mus musculus OX=10090 GN=Mob4 PE=1 SV=1 | 0.001 | 5.379 | 13 | 2 | 2 | 2 | 225 | 26 | 5.78 | 0.987 | 1.084 | 0.911 | -0.02 | 0.02 | 0.12 | 0.04 | -0.05 | -0.13 | 0.917224005 | 0.865471077 | 0.6159696 | 0.999652205 | 0.998402594 | 0.9990274 | 0.97826833 |
| Gtf3c5 | Medium | Q8R2T8 | General transcription factor 3C polypeptide 5 OS=Mus musculus OX=10090 GN=Gtf3c5 PE=2 SV=2 | 0.03 | 1.885 | 2 | 1 | 1 | 1 | 520 | 60.5 | 6.73 | 1.066 | 1.009 | 1.056 | 0.09 | -0.09 | 0.01 | 0.00333333 | -0.05 | 0.08 | 0.824295127 | 0.958926092 | 0.736725529 | 0.999652205 | 0.998402594 | 0.9990274 | 0.992535481 |
| Jam3 | High | Q9D8B7 | Junctional adhesion molecule C OS=Mus musculus OX=10090 GN=Jam3 PE=1 SV=2 | 0 | 20.796 | 17 | 6 | 7 | 6 | 310 | 34.8 | 7.03 | 1.068 | 1.01 | 1.058 | 0.09 | -0.09 | 0.01 | 0.00333333 | -0.05 | 0.08 | 0.863571642 | 0.955279717 | 0.756596759 | 0.999652205 | 0.998402594 | 0.9990274 | 0.992535481 |
| Prkar1a | High | Q9DBC7 | cAMP-dependent protein kinase type I-alpha regulatory subunit OS=Mus musculus OX=10090 GN= | 0 | 30.889 | 22 | 7 | 9 | 7 | 381 | 43.2 | 5.35 | 1.061 | 1.008 | 1.052 | 0.09 | -0.09 | 0.01 | 0.00333333 | -0.05 | 0.07 | 0.850323209 | 0.952243549 | 0.797131799 | 0.999652205 | 0.998402594 | 0.9990274 | 0.992535481 |
| Galk1 | High | Q9R0N0 | Galactokinase OS=Mus musculus OX=10090 GN=Galk1 PE=1 SV=2 | 0 | 12.382 | 11 | 4 | 4 | 4 | 392 | 42.3 | 5.26 | 0.956 | 1.128 | 0.848 | -0.07 | 0.07 | 0.17 | 0.05666667 | -0.05 | -0.24 | 0.845799823 | 0.777364814 | 0.353470136 | 0.999652205 | 0.998402594 | 0.9990274 | 0.911640779 |
| Stx6 | High | Q9JKK1 | Syntaxin-6 OS=Mus musculus OX=10090 GN=Stx6 PE=1 SV=1 | 0 | 10.519 | 13 | 2 | 2 | 2 | 255 | 29 | 4.92 | 0.954 | 1.123 | 0.849 | -0.07 | 0.07 | 0.17 | 0.05666667 | -0.05 | -0.24 | 0.778287629 | 0.791451215 | 0.44314778 | 0.999652205 | 0.998402594 | 0.9990274 | 0.958608248 |
| Gucy1a3 | High | Q9ERL9 | Guanylate cyclase soluble subunit alpha-1 OS=Mus musculus OX=10090 GN=Gucy1a1 PE=1 SV=2 | 0.001 | 6.801 | 5 | 2 | 2 | 2 | 691 | 77.5 | 7.17 | 0.983 | 1.098 | 0.895 | -0.03 | 0.03 | 0.13 | 0.04333333 | -0.05 | -0.16 | 0.891161434 | 0.851021248 | 0.571288022 | 0.999652205 | 0.998402594 | 0.9990274 | 0.972910028 |
| Lsm12 | High | Q9D0R8 | Protein LSM12 homolog OS=Mus musculus OX=10090 GN=Lsm12 PE=1 SV=1 | 0.001 | 4.525 | 11 | 2 | 2 | 2 | 195 | 21.7 | 7.74 | 1.078 | 0.995 | 1.084 | 0.11 | -0.11 | -0.01 | -0.0033333 | -0.05 | 0.12 | 0.819083484 | 0.901920553 | 0.577225517 | 0.999652205 | 0.998402594 | 0.9990274 | 0.972910028 |
| Surf6 | High | P70279 | Surfeit locus protein 6 OS=Mus musculus OX=10090 GN=Surf6 PE=1 SV=1 | 0.005 | 3.59 | 3 | 1 | 1 | 1 | 355 | 41.2 | 10.52 | 1.017 | 1.059 | 0.961 | 0.02 | -0.02 | 0.08 | 0.02666667 | -0.05 | -0.06 | 0.996107331 | 0.92356008 | 0.847611722 | 0.999652205 | 0.998402594 | 0.9990274 | 0.992535481 |
| Sparcl1 | High | P70663 | SPARC-like protein 1 OS=Mus musculus OX=10090 GN=Sparcl1 PE=1 SV=3 | 0.009 | 2.798 | 2 | 1 | 1 | 1 | 650 | 72.2 | 4.6 | 1.029 | 1.04 | 0.989 | 0.04 | -0.04 | 0.06 | 0.02 | -0.05 | -0.02 | 0.971460647 | 0.979529135 | 0.979674335 | 0.999652205 | 0.998402594 | 0.9990274 | 0.997765878 |
| Stxbp2 | High | Q64324 | Syntaxin-binding protein 2 OS=Mus musculus OX=10090 GN=Stxbp2 PE=1 SV=1 | 0 | 108.438 | 39 | 19 | 34 | 18 | 593 | 66.3 | 6.74 | 1.031 | 1.043 | 0.988 | 0.04 | -0.04 | 0.06 | 0.02 | -0.05 | -0.02 | 0.916858647 | 0.966341025 | 0.980035941 | 0.999652205 | 0.998402594 | 0.9990274 | 0.997765878 |
| Snx9 | Low | Q91VH2 | Sorting nexin-9 OS=Mus musculus OX=10090 GN=Snx9 PE=1 SV=1 | 0.075 | 1.238 | 1 | 1 | 1 | 1 | 595 | 66.5 | 5.52 | 0.954 | 1.129 | 0.845 | -0.07 | 0.07 | 0.18 | 0.06 | -0.055 | -0.24 | 0.781295709 | 0.782972464 | 0.320367277 | 0.999652205 | 0.998402594 | 0.9990274 | 0.905392802 |
| Emb | High | P21995 | Embigin OS=Mus musculus OX=10090 GN=Emb PE=1 SV=2 | 0 | 10.311 | 10 | 4 | 5 | 4 | 330 | 37 | 6.02 | 0.975 | 1.109 | 0.879 | -0.04 | 0.04 | 0.15 | 0.05 | -0.055 | -0.19 | 0.860220215 | 0.815955151 | 0.406983561 | 0.999652205 | 0.998402594 | 0.9990274 | 0.936397673 |
| Csrp1 | High | P97315 | Cysteine and glycine-rich protein 1 OS=Mus musculus OX=10090 GN=Csrp1 PE=1 SV=3 | 0 | 39.185 | 28 | 4 | 8 | 4 | 193 | 20.6 | 8.57 | 0.999 | 1.08 | 0.925 | 0 | 0 | 0.11 | 0.03666667 | -0.055 | -0.11 | 0.948457878 | 0.880904315 | 0.613027226 | 0.999652205 | 0.998402594 | 0.9990274 | 0.97826833 |
| Crkl | High | P47941 | Crk-like protein OS=Mus musculus OX=10090 GN=Crkl PE=1 SV=2 | 0 | 11.376 | 17 | 6 | 6 | 6 | 303 | 33.8 | 6.74 | 0.994 | 1.086 | 0.915 | -0.01 | 0.01 | 0.12 | 0.04 | -0.055 | -0.13 | 0.994469978 | 0.866432956 | 0.628025927 | 0.999652205 | 0.998402594 | 0.9990274 | 0.981444134 |
| Sh3pxd2b | High | A2AAY5 | SH3 and PX domain-containing protein 2B OS=Mus musculus OX=10090 GN=Sh3pxd2b PE=1 SV=1 | 0 | 4.871 | 2 | 2 | 2 | 2 | 908 | 101.5 | 8.66 | 1.003 | 1.076 | 0.932 | 0 | 0 | 0.11 | 0.03666667 | -0.055 | -0.1 | 0.87000428 | 0.897195309 | 0.682447216 | 0.999652205 | 0.998402594 | 0.9990274 | 0.991285312 |
| Arf4 | High | P61750 | ADP-ribosylation factor 4 OS=Mus musculus OX=10090 GN=Arf4 PE=1 SV=2 | 0 | 16.439 | 23 | 3 | 6 | 1 | 180 | 20.4 | 7.14 | 1.075 | 1.007 | 1.068 | 0.1 | -0.1 | 0.01 | 0.00333333 | -0.055 | 0.09 | 0.861529993 | 0.948214039 | 0.710266471 | 0.999652205 | 0.998402594 | 0.9990274 | 0.992535481 |
| Ap3d1 | High | O54774 | AP-3 complex subunit delta-1 OS=Mus musculus OX=10090 GN=Ap3d1 PE=1 SV=1 | 0 | 32.09 | 8 | 8 | 9 | 8 | 1199 | 135 | 7.37 | 1.063 | 1.017 | 1.046 | 0.09 | -0.09 | 0.02 | 0.00666667 | -0.055 | 0.06 | 0.834161317 | 0.972228848 | 0.722841909 | 0.999652205 | 0.998402594 | 0.9990274 | 0.992535481 |
| Psmf1 | High | Q8BHL8 | Proteasome inhibitor PI31 subunit OS=Mus musculus OX=10090 GN=Psmf1 PE=1 SV=1 | 0.001 | 4.492 | 5 | 1 | 1 | 1 | 271 | 29.6 | 5.25 | 1.007 | 1.071 | 0.94 | 0.01 | -0.01 | 0.1 | 0.03333333 | -0.055 | -0.09 | 0.966159925 | 0.89577839 | 0.751396557 | 0.999652205 | 0.998402594 | 0.9990274 | 0.992535481 |
| Gng5; Gm | High | Q80SZ7 | Guanine nucleotide-binding protein G(I)/G(S)/G(O) subunit gamma-5 OS=Mus musculus OX=100 | 0 | 31.561 | 76 | 5 | 17 | 5 | 68 | 7.3 | 9.85 | 1.013 | 1.067 | 0.95 | 0.02 | -0.02 | 0.09 | 0.03 | -0.055 | -0.07 | 0.962291272 | 0.909878391 | 0.815343146 | 0.999652205 | 0.998402594 | 0.9990274 | 0.992535481 |
| Tmem30a | High | Q8VEK0 | Cell cycle control protein 50A OS=Mus musculus OX=10090 GN=Tmem30a PE=1 SV=1 | 0 | 125.853 | 41 | 18 | 60 | 18 | 364 | 41 | 8.37 | 1.015 | 1.066 | 0.952 | 0.02 | -0.02 | 0.09 | 0.03 | -0.055 | -0.07 | 0.958226685 | 0.913170137 | 0.829730629 | 0.999652205 | 0.998402594 | 0.9990274 | 0.992535481 |
| Dgkz | Medium | A2AHJ7 | Diacylglycerol kinase OS=Mus musculus OX=10090 GN=Dgkz PE=1 SV=1 | 0.011 | 2.435 | 1 | 1 | 1 | 1 | 946 | 105.8 | 8.07 | 1.012 | 1.068 | 0.948 | 0.02 | -0.02 | 0.09 | 0.03 | -0.055 | -0.08 | 0.988365989 | 0.929862675 | 0.845274022 | 0.999652205 | 0.998402594 | 0.9990274 | 0.992535481 |
| Pik3r4 | High | Q8VD65 | Phosphoinositide 3-kinase regulatory subunit 4 OS=Mus musculus OX=10090 GN=Pik3r4 PE=1 SV= | 0.001 | 6.866 | 1 | 2 | 2 | 2 | 1358 | 152.5 | 7.12 | 0.962 | 1.121 | 0.858 | -0.06 | 0.06 | 0.17 | 0.05666667 | -0.055 | -0.22 | 0.80527548 | 0.790356717 | 0.383574232 | 0.999652205 | 0.998402594 | 0.9990274 | 0.926692597 |
| Mpi | Medium | Q924M7 | Mannose-6-phosphate isomerase OS=Mus musculus OX=10090 GN=Mpi PE=1 SV=1 | 0.026 | 1.954 | 3 | 1 | 1 | 1 | 423 | 46.5 | 5.95 | 1.12 | 0.972 | 1.152 | 0.16 | -0.16 | -0.04 | -0.0133333 | -0.06 | 0.2 | 0.72663511 | 0.863998731 | 0.429819348 | 0.999652205 | 0.998402594 | 0.9990274 | 0.944356224 |
| Ctu1 | High | Q99J10 | Cytoplasmic tRNA 2-thiolation protein 1 OS=Mus musculus OX=10090 GN=Ctu1 PE=2 SV=1 | 0.008 | 2.914 | 2 | 1 | 1 | 1 | 420 | 43.8 | 8.1 | 1.119 | 0.973 | 1.15 | 0.16 | -0.16 | -0.04 | -0.0133333 | -0.06 | 0.2 | 0.740523125 | 0.843253688 | 0.463444255 | 0.999652205 | 0.998402594 | 0.9990274 | 0.959744396 |
| Arpc5l | High | Q9D898 | Actin-related protein 2/3 complex subunit 5-like protein OS=Mus musculus OX=10090 GN=Arpc5l | 0 | 27.49 | 33 | 4 | 7 | 4 | 153 | 17 | 6.8 | 1.108 | 0.982 | 1.128 | 0.15 | -0.15 | -0.03 | -0.01 | -0.06 | 0.17 | 0.73118532 | 0.887678116 | 0.490405885 | 0.999652205 | 0.998402594 | 0.9990274 | 0.964378027 |
| Gpd1l | High | Q3ULJ0 | Glycerol-3-phosphate dehydrogenase 1-like protein OS=Mus musculus OX=10090 GN=Gpd1l PE=1 | 0 | 8.505 | 9 | 3 | 3 | 3 | 351 | 38.2 | 6.77 | 1.086 | 1.003 | 1.083 | 0.12 | -0.12 | 0 | 0 | -0.06 | 0.11 | 0.872293597 | 0.921112434 | 0.574200427 | 0.999652205 | 0.998402594 | 0.9990274 | 0.972910028 |
| Atpaf1 | High | H3BLL2 | ATP synthase mitochondrial F1 complex assembly factor 1 OS=Mus musculus OX=10090 GN=Atpa | 0 | 4.903 | 2 | 1 | 2 | 1 | 348 | 38.8 | 5.86 | 0.998 | 1.089 | 0.916 | 0 | 0 | 0.12 | 0.04 | -0.06 | -0.13 | 0.96265617 | 0.853697083 | 0.630452588 | 0.999652205 | 0.998402594 | 0.9990274 | 0.981916182 |
| Uhrf2 | Medium | Q7TMI3 | E3 ubiquitin-protein ligase UHRF2 OS=Mus musculus OX=10090 GN=Uhrf2 PE=1 SV=1 | 0.01 | 2.616 | 2 | 1 | 1 | 1 | 803 | 90 | 8.03 | 1.121 | 0.969 | 1.157 | 0.17 | -0.17 | -0.05 | -0.0166667 | -0.06 | 0.21 | 0.681997024 | 0.858043413 | 0.384580577 | 0.999652205 | 0.998402594 | 0.9990274 | 0.92785856 |
| Gm4907 | High | B1AXB9 | Testin OS=Mus musculus OX=10090 GN=Tesl1 PE=1 SV=1 | 0.003 | 3.867 | 2 | 1 | 1 | 1 | 411 | 46.8 | 8.62 | 1.123 | 0.963 | 1.166 | 0.17 | -0.17 | -0.05 | -0.0166667 | -0.06 | 0.22 | 0.681641772 | 0.839907836 | 0.39212335 | 0.999652205 | 0.998402594 | 0.9990274 | 0.93064581 |
| Snta1 | High | A2AKD7 | Alpha-1-syntrophin OS=Mus musculus OX=10090 GN=Snta1 PE=1 SV=1 | 0.006 | 3.47 | 3 | 2 | 2 | 1 | 499 | 53.2 | 6.8 | 1.128 | 0.97 | 1.162 | 0.17 | -0.17 | -0.04 | -0.0133333 | -0.065 | 0.22 | 0.760003587 | 0.839293492 | 0.311082648 | 0.999652205 | 0.998402594 | 0.9990274 | 0.904943958 |
| Arhgef7 | High | Q9ES28 | Rho guanine nucleotide exchange factor 7 OS=Mus musculus OX=10090 GN=Arhgef7 PE=1 SV=2 | 0 | 9.829 | 5 | 4 | 4 | 4 | 862 | 97 | 6.8 | 1.128 | 0.971 | 1.161 | 0.17 | -0.17 | -0.04 | -0.0133333 | -0.065 | 0.22 | 0.645590664 | 0.860615484 | 0.390330597 | 0.999652205 | 0.998402594 | 0.9990274 | 0.93064581 |
| Clasp2 | High | E9Q8N5 | CLIP-associating protein 2 OS=Mus musculus OX=10090 GN=Clasp2 PE=1 SV=1 | 0 | 74.702 | 19 | 19 | 20 | 13 | 1286 | 140.6 | 8.63 | 1.105 | 0.991 | 1.115 | 0.14 | -0.14 | -0.01 | -0.0033333 | -0.065 | 0.16 | 0.677586106 | 0.91057994 | 0.42296011 | 0.999652205 | 0.998402594 | 0.9990274 | 0.939860852 |
| Wdr11 | Medium | G5E8J3 | WD repeat-containing protein 11 OS=Mus musculus OX=10090 GN=Wdr11 PE=1 SV=1 | 0.01 | 2.681 | 1 | 1 | 1 | 1 | 1223 | 135.9 | 7.11 | 0.99 | 1.105 | 0.896 | -0.01 | 0.01 | 0.14 | 0.04666667 | -0.065 | -0.16 | 0.862951005 | 0.830296755 | 0.526270028 | 0.999652205 | 0.998402594 | 0.9990274 | 0.964378027 |
| Eif5b | High | Q05D44 | Eukaryotic translation initiation factor 5B OS=Mus musculus OX=10090 GN=Eif5b PE=1 SV=2 | 0 | 8.956 | 3 | 3 | 3 | 3 | 1216 | 137.5 | 5.59 | 1.092 | 1.001 | 1.091 | 0.13 | -0.13 | 0 | 0 | -0.065 | 0.13 | 0.770375421 | 0.935092971 | 0.564762371 | 0.999652205 | 0.998402594 | 0.9990274 | 0.972910028 |
| | High | E9Q0A3 | Rho guanine nucleotide exchange factor (GEF) 11 (Fragment) OS=Mus musculus OX=10090 GN=A | 0.001 | 5.251 | 1 | 2 | 2 | 1 | 1475 | 163.4 | 5.78 | 1.001 | 1.095 | 0.914 | 0 | 0 | 0.13 | 0.04333333 | -0.065 | -0.13 | 0.948478774 | 0.863609495 | 0.621025818 | 0.999652205 | 0.998402594 | 0.9990274 | 0.97826833 |
| Sipa1l3 | High | G3X9J0 | Signal-induced proliferation-associated 1-like protein 3 OS=Mus musculus OX=10090 GN=Sipa1l3 | 0 | 8.658 | 1 | 2 | 2 | 2 | 1776 | 194.9 | 8.32 | 1.084 | 1.008 | 1.075 | 0.12 | -0.12 | 0.01 | 0.00333333 | -0.065 | 0.1 | 0.784840142 | 0.955210685 | 0.657317904 | 0.999652205 | 0.998402594 | 0.9990274 | 0.985010103 |
| Camk2b | High | P28652 | Calcium/calmodulin-dependent protein kinase type II subunit beta OS=Mus musculus OX=10090 G | 0 | 67.818 | 21 | 10 | 29 | 4 | 542 | 60.4 | 7.28 | 1.083 | 1.006 | 1.076 | 0.12 | -0.12 | 0.01 | 0.00333333 | -0.065 | 0.11 | 0.851669814 | 0.946786063 | 0.675743941 | 0.999652205 | 0.998402594 | 0.9990274 | 0.990250803 |
| Puf60 | High | Q3UEB3 | Poly(U)-binding-splicing factor PUF60 OS=Mus musculus OX=10090 GN=Puf60 PE=1 SV=2 | 0.001 | 7.257 | 4 | 2 | 2 | 2 | 564 | 60.2 | 5.29 | 1.079 | 1.013 | 1.065 | 0.11 | -0.11 | 0.02 | 0.00666667 | -0.065 | 0.09 | 0.811147696 | 0.966328413 | 0.705688103 | 0.999652205 | 0.998402594 | 0.9990274 | 0.992485323 |
| Epha4 | High | Q03137 | Ephrin type-A receptor 4 OS=Mus musculus OX=10090 GN=Epha4 PE=1 SV=2 | 0.001 | 4.295 | 1 | 1 | 1 | 1 | 986 | 109.7 | 6.51 | 1.062 | 1.031 | 1.03 | 0.09 | -0.09 | 0.04 | 0.01333333 | -0.065 | 0.04 | 0.872164035 | 0.989133166 | 0.875393819 | 0.999652205 | 0.998402594 | 0.9990274 | 0.992535481 |
| Mpdz | High | Q8VBX6 | Multiple PDZ domain protein OS=Mus musculus OX=10090 GN=Mpdz PE=1 SV=2 | 0.001 | 7.516 | 2 | 3 | 3 | 2 | 2055 | 218.6 | 5.02 | 1.039 | 1.046 | 0.993 | 0.06 | -0.06 | 0.07 | 0.02333333 | -0.065 | -0.01 | 0.938877355 | 0.957502266 | 0.982878108 | 0.999652205 | 0.998402594 | 0.9990274 | 0.997765878 |
| Entpd2 | High | O55026 | Ectonucleoside triphosphate diphosphohydrolase 2 OS=Mus musculus OX=10090 GN=Entpd2 PE=1 | 0 | 9.46 | 5 | 2 | 2 | 2 | 495 | 54.3 | 8.37 | 1.04 | 1.051 | 0.989 | 0.06 | -0.06 | 0.07 | 0.02333333 | -0.065 | -0.02 | 0.909387786 | 0.936838652 | 0.98899984 | 0.999652205 | 0.998402594 | 0.9990274 | 0.999783808 |
| Prkd1 | High | Q62101 | Serine/threonine-protein kinase D1 OS=Mus musculus OX=10090 GN=Prkd1 PE=1 SV=2 | 0 | 12.852 | 6 | 6 | 6 | 3 | 918 | 102 | 6.58 | 1.09 | 1.017 | 1.072 | 0.12 | -0.12 | 0.02 | 0.00666667 | -0.07 | 0.1 | 0.744547259 | 0.972141301 | 0.682433879 | 0.999652205 | 0.998402594 | 0.9990274 | 0.991285312 |
| Hspa14 | Medium | Q99M31 | Heat shock 70 kDa protein 14 OS=Mus musculus OX=10090 GN=Hspa14 PE=1 SV=2 | 0.035 | 1.775 | 1 | 1 | 1 | 1 | 509 | 54.6 | 5.92 | 1.129 | 0.98 | 1.152 | 0.17 | -0.17 | -0.03 | -0.01 | -0.07 | 0.2 | 0.683874329 | 0.86359227 | 0.352321256 | 0.999652205 | 0.998402594 | 0.9990274 | 0.910898173 |
| Nptn | High | P97300 | Neuroplastin OS=Mus musculus OX=10090 GN=Nptn PE=1 SV=3 | 0 | 98.584 | 36 | 16 | 40 | 16 | 397 | 44.3 | 7.74 | 0.979 | 1.125 | 0.87 | -0.03 | 0.03 | 0.17 | 0.05666667 | -0.07 | -0.2 | 0.944553408 | 0.783538963 | 0.425636916 | 0.999652205 | 0.998402594 | 0.9990274 | 0.941180024 |
| Slk | High | O54988 | STE20-like serine/threonine-protein kinase OS=Mus musculus OX=10090 GN=Slk PE=1 SV=2 | 0 | 4.757 | 1 | 2 | 2 | 2 | 1233 | 141.4 | 5.14 | 0.989 | 1.115 | 0.887 | -0.02 | 0.02 | 0.16 | 0.05333333 | -0.07 | -0.17 | 0.884141456 | 0.805071982 | 0.515541246 | 0.999652205 | 0.998402594 | 0.9990274 | 0.964378027 |
| Sec31a | High | Q3UPL0 | Protein transport protein Sec31A OS=Mus musculus OX=10090 GN=Sec31a PE=1 SV=2 | 0 | 23.367 | 6 | 6 | 6 | 6 | 1230 | 133.5 | 6.76 | 1.009 | 1.098 | 0.919 | 0.01 | -0.01 | 0.13 | 0.04333333 | -0.07 | -0.12 | 0.986646474 | 0.841557156 | 0.634527923 | 0.999652205 | 0.998402594 | 0.9990274 | 0.983562471 |
| Prkar2a | High | Q8K1M3 | Protein kinase, cAMP dependent regulatory, type II alpha OS=Mus musculus OX=10090 GN=Prkar2 | 0 | 56.374 | 35 | 13 | 19 | 12 | 402 | 45.6 | 4.93 | 1.068 | 1.032 | 1.034 | 0.09 | -0.09 | 0.05 | 0.01666667 | -0.07 | 0.05 | 0.816397918 | 0.991055893 | 0.750106788 | 0.999652205 | 0.998402594 | 0.9990274 | 0.992535481 |
| Fktn | Low | A0A1Y7VN | Ribitol-5-phosphate transferase OS=Mus musculus OX=10090 GN=Fktn PE=1 SV=1 | 0.052 | 1.45 | 2 | 1 | 1 | 1 | 500 | 58.2 | 8.41 | 1.075 | 1.027 | 1.047 | 0.1 | -0.1 | 0.04 | 0.01333333 | -0.07 | 0.07 | 0.835560576 | 0.991265378 | 0.782414484 | 0.999652205 | 0.998402594 | 0.9990274 | 0.992535481 |
| | High | P05480 | Neuronal proto-oncogene tyrosine-protein kinase Src OS=Mus musculus OX=10090 GN=Src PE=1 S | 0 | 26.821 | 13 | 6 | 11 | 3 | 535 | 59.9 | 7.42 | 1.07 | 1.031 | 1.038 | 0.1 | -0.1 | 0.04 | 0.01333333 | -0.07 | 0.05 | 0.852625693 | 0.994165055 | 0.805388075 | 0.999652205 | 0.998402594 | 0.9990274 | 0.992535481 |
| Adsl | High | P54822 | Adenylosuccinate lyase OS=Mus musculus OX=10090 GN=Adsl PE=1 SV=2 | 0 | 34.598 | 23 | 7 | 7 | 7 | 484 | 54.8 | 7.27 | 1.026 | 1.073 | 0.957 | 0.04 | -0.04 | 0.1 | 0.03333333 | -0.07 | -0.06 | 0.882630063 | 0.897000589 | 0.821891497 | 0.999652205 | 0.998402594 | 0.9990274 | 0.992535481 |
| Cstb | High | Q62426 | Cystatin-B OS=Mus musculus OX=10090 GN=Cstb PE=1 SV=1 | 0 | 32.834 | 49 | 4 | 12 | 4 | 98 | 11 | 7.39 | 1.054 | 1.042 | 1.011 | 0.08 | -0.08 | 0.06 | 0.02 | -0.07 | 0.02 | 0.857952635 | 0.967106471 | 0.865083063 | 0.999652205 | 0.998402594 | 0.9990274 | 0.992535481 |
| 0610011F( | High | Q9DCS2 | Methyltransferase-like 26 OS=Mus musculus OX=10090 GN=Mettl26 PE=1 SV=1 | 0 | 8.112 | 9 | 1 | 1 | 1 | 204 | 22.7 | 6.52 | 1.035 | 1.065 | 0.971 | 0.05 | -0.05 | 0.09 | 0.03 | -0.07 | -0.04 | 0.965045309 | 0.920053587 | 0.901180699 | 0.999652205 | 0.998402594 | 0.9990274 | 0.992535481 |
| Itfg3; Fam | High | Q8C0Z1 | Protein FAM234A OS=Mus musculus OX=10090 GN=Fam234a PE=1 SV=1 | 0 | 13.453 | 9 | 5 | 6 | 5 | 555 | 60.5 | 5.68 | 1.062 | 1.038 | 1.023 | 0.09 | -0.09 | 0.05 | 0.01666667 | -0.07 | 0.03 | 0.868369193 | 0.977600052 | 0.924641187 | 0.999652205 | 0.998402594 | 0.9990274 | 0.992535481 |
| | High | E9Q1S3 | Protein transport protein SEC23 OS=Mus musculus OX=10090 GN=Sec23a PE=1 SV=1 | 0 | 20.846 | 11 | 6 | 6 | 5 | 736 | 82.9 | 7.46 | 1.046 | 1.06 | 0.986 | 0.06 | -0.06 | 0.08 | 0.02666667 | -0.07 | -0.02 | 0.836536874 | 0.926098684 | 0.958784667 | 0.999652205 | 0.998402594 | 0.9990274 | 0.992902921 |
| Tmem2 | High | Q5FWI3 | Cell surface hyaluronidase OS=Mus musculus OX=10090 GN=Cemip2 PE=1 SV=1 | 0 | 46.724 | 10 | 13 | 14 | 13 | 1383 | 153.7 | 7.78 | 1.042 | 1.059 | 0.983 | 0.06 | -0.06 | 0.08 | 0.02666667 | -0.07 | -0.02 | 0.888761558 | 0.927529535 | 0.989313414 | 0.999652205 | 0.998402594 | 0.9990274 | 0.999783808 |
| Fam57b | Medium | Q7TNV1 | Ceramide synthase OS=Mus musculus OX=10090 GN=Tlcd3b PE=1 SV=2 | 0.035 | 1.799 | 4 | 1 | 1 | 1 | 275 | 30.8 | 8.84 | 1.134 | 0.982 | 1.156 | 0.18 | -0.18 | -0.03 | -0.01 | -0.075 | 0.21 | 0.677609639 | 0.889563347 | 0.410411095 | 0.999652205 | 0.998402594 | 0.9990274 | 0.936397673 |
| Bdh2 | High | Q8JZV9 | 3-hydroxybutyrate dehydrogenase type 2 OS=Mus musculus OX=10090 GN=Bdh2 PE=1 SV=1 | 0 | 33.004 | 34 | 8 | 10 | 8 | 245 | 26.7 | 7.99 | 1.117 | 0.994 | 1.124 | 0.16 | -0.16 | -0.01 | -0.0033333 | -0.075 | 0.17 | 0.749677619 | 0.917583462 | 0.49874342 | 0.999652205 | 0.998402594 | 0.9990274 | 0.964378027 |
| Rab10 | High | P61027 | Ras-related protein Rab-10 OS=Mus musculus OX=10090 GN=Rab10 PE=1 SV=1 | 0 | 109.465 | 65 | 17 | 41 | 13 | 200 | 22.5 | 8.38 | 0.994 | 1.115 | 0.892 | -0.01 | 0.01 | 0.16 | 0.05333333 | -0.075 | -0.17 | 0.986194182 | 0.804516692 | 0.522528233 | 0.999652205 | 0.998402594 | 0.9990274 | 0.964378027 |
| Gtpbp1 | High | O08582 | GTP-binding protein 1 OS=Mus musculus OX=10090 GN=Gtpbp1 PE=1 SV=2 | 0 | 74.082 | 29 | 10 | 16 | 10 | 668 | 72.3 | 8.29 | 1.098 | 1.015 | 1.082 | 0.13 | -0.13 | 0.02 | 0.00666667 | -0.075 | 0.11 | 0.813364363 | 0.967462109 | 0.643181086 | 0.999652205 | 0.998402594 | 0.9990274 | 0.985010103 |
| Mras | High | O08989 | Ras-related protein M-Ras OS=Mus musculus OX=10090 GN=Mras PE=1 SV=1 | 0 | 25.61 | 30 | 6 | 9 | 6 | 208 | 23.9 | 8.78 | 1.02 | 1.089 | 0.937 | 0.03 | -0.03 | 0.12 | 0.04 | -0.075 | -0.09 | 0.997024083 | 0.860197073 | 0.658032627 | 0.999652205 | 0.998402594 | 0.9990274 | 0.985010103 |
| Anxa7 | High | Q07076 | Annexin A7 OS=Mus musculus OX=10090 GN=Anxa7 PE=1 SV=2 | 0 | 81.261 | 35 | 17 | 28 | 17 | 463 | 49.9 | 6.18 | 1.013 | 1.098 | 0.923 | 0.02 | -0.02 | 0.13 | 0.04333333 | -0.075 | -0.12 | 0.923266054 | 0.840819951 | 0.676238688 | 0.999652205 | 0.998402594 | 0.9990274 | 0.990250803 |
| Slmap | High | H7BX64 | Sarcolemmal membrane-associated protein OS=Mus musculus OX=10090 GN=Slmap PE=1 SV=1 | 0 | 60.535 | 17 | 15 | 24 | 15 | 828 | 95 | 5.35 | 1.076 | 1.031 | 1.044 | 0.11 | -0.11 | 0.04 | 0.01333333 | -0.075 | 0.06 | 0.801894659 | 0.993453671 | 0.706130603 | 0.999652205 | 0.998402594 | 0.9990274 | 0.992485323 |
| Tmem237 | High | A0A087W | Transmembrane protein 237 (Fragment) OS=Mus musculus OX=10090 GN=Tmem237 PE=1 SV=1 | 0 | 69.361 | 58 | 10 | 26 | 10 | 230 | 25.5 | 9.14 | 1.028 | 1.077 | 0.954 | 0.04 | -0.04 | 0.11 | 0.03666667 | -0.075 | -0.07 | 0.923980809 | 0.886643597 | 0.839118977 | 0.999652205 | 0.998402594 | 0.9990274 | 0.992535481 |
| Cotl1 | High | Q9CQI6 | Coactosin-like protein OS=Mus musculus OX=10090 GN=Cotl1 PE=1 SV=3 | 0 | 27.986 | 61 | 7 | 11 | 7 | 142 | 15.9 | 5.4 | 1.044 | 1.062 | 0.983 | 0.06 | -0.06 | 0.09 | 0.03 | -0.075 | -0.03 | 0.933991905 | 0.920778979 | 0.911320106 | 0.999652205 | 0.998402594 | 0.9990274 | 0.992535481 |
| Pdlim5 | High | Q8CI51 | PDZ and LIM domain protein 5 OS=Mus musculus OX=10090 GN=Pdlim5 PE=1 SV=4 | 0 | 47.638 | 24 | 10 | 11 | 10 | 591 | 63.3 | 8.25 | 1.065 | 1.044 | 1.02 | 0.09 | -0.09 | 0.06 | 0.02 | -0.075 | 0.03 | 0.870426228 | 0.962250204 | 0.942033101 | 0.999652205 | 0.998402594 | 0.9990274 | 0.992535481 |
| 2010012O | High | Q9CRC6 | BLOC-1-related complex subunit 7 OS=Mus musculus OX=10090 GN=Borcs7 PE=1 SV=1 | 0.001 | 6.415 | 16 | 1 | 1 | 1 | 105 | 11.5 | 6.79 | 0.989 | 1.128 | 0.877 | -0.02 | 0.02 | 0.17 | 0.05666667 | -0.075 | -0.19 | 0.894890052 | 0.777982796 | 0.496992169 | 0.999652205 | 0.998402594 | 0.9990274 | 0.964378027 |
| Fam49b | High | Q921M7 | CYFIP-related Rac1 interactor B OS=Mus musculus OX=10090 GN=Cyrib PE=1 SV=1 | 0.001 | 5.983 | 9 | 2 | 2 | 2 | 324 | 36.8 | 6.06 | 1.05 | 1.056 | 0.994 | 0.07 | -0.07 | 0.08 | 0.02666667 | -0.075 | -0.01 | 0.901535111 | 0.930567796 | 0.994660251 | 0.999652205 | 0.998402594 | 0.9990274 | 0.999917596 |
| Arfgap2 | High | Q99K28 | ADP-ribosylation factor GTPase-activating protein 2 OS=Mus musculus OX=10090 GN=Arfgap2 PE | 0 | 41.087 | 18 | 11 | 15 | 10 | 520 | 56.6 | 8.18 | 1.136 | 0.985 | 1.154 | 0.18 | -0.18 | -0.02 | -0.0066667 | -0.08 | 0.21 | 0.68124945 | 0.893771504 | 0.330585409 | 0.999652205 | 0.998402594 | 0.9990274 | 0.906329766 |
| | Low | A0A2R8W | Ran GTPase-activating protein 1 (Fragment) OS=Mus musculus OX=10090 GN=Rangap1 PE=1 SV= | 0.052 | 1.452 | 5 | 1 | 1 | 1 | 144 | 15.4 | 5.07 | 1.118 | 0.997 | 1.121 | 0.16 | -0.16 | 0 | 0 | -0.08 | 0.17 | 0.671335391 | 0.929570128 | 0.486682516 | 0.999652205 | 0.998402594 | 0.9990274 | 0.964378027 |
| Gmps | High | Q3THK7 | GMP synthase [glutamine-hydrolyzing] OS=Mus musculus OX=10090 GN=Gmps PE=1 SV=2 | 0.005 | 3.483 | 2 | 2 | 2 | 2 | 693 | 76.7 | 6.73 | 1.097 | 1.022 | 1.073 | 0.13 | -0.13 | 0.03 | 0.01 | -0.08 | 0.1 | 0.863155349 | 0.968432525 | 0.606465919 | 0.999652205 | 0.998402594 | 0.9990274 | 0.97826833 |
| Calm3; Ca | High | P0DP27 | Calmodulin-2 OS=Mus musculus OX=10090 GN=Calm2 PE=1 SV=1 | 0 | 141.517 | 77 | 11 | 145 | 11 | 149 | 16.8 | 4.22 | 1.018 | 1.096 | 0.929 | 0.03 | -0.03 | 0.13 | 0.04333333 | -0.08 | -0.11 | 0.949587559 | 0.845183131 | 0.707325014 | 0.999652205 | 0.998402594 | 0.9990274 | 0.992485323 |
| Arr3 | High | Q9EQP6 | Arrestin-C OS=Mus musculus OX=10090 GN=Arr3 PE=1 SV=1 | 0 | 27.559 | 20 | 5 | 7 | 5 | 381 | 41.9 | 6.44 | 1.022 | 1.094 | 0.934 | 0.03 | -0.03 | 0.13 | 0.04333333 | -0.08 | -0.1 | 0.885968691 | 0.84933506 | 0.723666949 | 0.999652205 | 0.998402594 | 0.9990274 | 0.992535481 |
| Git1 | High | Q68FF6 | ARF GTPase-activating protein GIT1 OS=Mus musculus OX=10090 GN=Git1 PE=1 SV=1 | 0 | 9.659 | 3 | 4 | 4 | 3 | 770 | 85.2 | 6.93 | 1.023 | 1.091 | 0.938 | 0.03 | -0.03 | 0.13 | 0.04333333 | -0.08 | -0.09 | 0.90759814 | 0.85693156 | 0.742418815 | 0.999652205 | 0.998402594 | 0.9990274 | 0.992535481 |
| Islr | Low | Q6GU68 | Immunoglobulin superfamily containing leucine-rich repeat protein OS=Mus musculus OX=10090 | 0.069 | 1.301 | 2 | 1 | 1 | 1 | 428 | 45.6 | 5.38 | 1.075 | 1.043 | 1.031 | 0.1 | -0.1 | 0.06 | 0.02 | -0.08 | 0.04 | 0.810257238 | 0.961288981 | 0.838340081 | 0.999652205 | 0.998402594 | 0.9990274 | 0.992535481 |
| Fam195b; | High | Q3UGS4 | Mapk-regulated corepressor-interacting protein 1 OS=Mus musculus OX=10090 GN=Mcrip1 PE=1 | 0.01 | 2.75 | 11 | 1 | 1 | 1 | 97 | 11.1 | 9.14 | 1.048 | 1.065 | 0.984 | 0.07 | -0.07 | 0.09 | 0.03 | -0.08 | -0.02 | 0.911426144 | 0.927859664 | 0.950803266 | 0.999652205 | 0.998402594 | 0.9990274 | 0.992535481 |
| Anxa11 | High | P97384 | Annexin A11 OS=Mus musculus OX=10090 GN=Anxa11 PE=1 SV=2 | 0 | 10.249 | 7 | 4 | 4 | 4 | 503 | 54 | 7.66 | 1.048 | 1.063 | 0.986 | 0.07 | -0.07 | 0.09 | 0.03 | -0.08 | -0.02 | 0.82495902 | 0.919757766 | 0.962848954 | 0.999652205 | 0.998402594 | 0.9990274 | 0.993960013 |
| Mpc2 | Low | Q9D023 | Mitochondrial pyruvate carrier 2 OS=Mus musculus OX=10090 GN=Mpc2 PE=1 SV=1 | 0.078 | 1.228 | 9 | 1 | 1 | 1 | 127 | 14.3 | 10.61 | 1.109 | 1.016 | 1.092 | 0.15 | -0.15 | 0.02 | 0.00666667 | -0.085 | 0.13 | 0.763250305 | 0.953922059 | 0.533975871 | 0.999652205 | 0.998402594 | 0.9990274 | 0.96694628 |
| Trpv4 | High | Q9EPK8 | Transient receptor potential cation channel subfamily V member 4 OS=Mus musculus OX=10090 ( | 0.008 | 2.906 | 1 | 1 | 1 | 1 | 871 | 98 | 7.33 | 1.039 | 1.081 | 0.962 | 0.06 | -0.06 | 0.11 | 0.03666667 | -0.085 | -0.06 | 0.921670669 | 0.87392147 | 0.854511066 | 0.999652205 | 0.998402594 | 0.9990274 | 0.992535481 |
| Arl15 | Medium | Q8BGR6 | ADP-ribosylation factor-like protein 15 OS=Mus musculus OX=10090 GN=Arl15 PE=1 SV=1 | 0.029 | 1.908 | 6 | 1 | 1 | 1 | 204 | 22.9 | 5.29 | 1.062 | 1.059 | 1.003 | 0.09 | -0.09 | 0.08 | 0.02666667 | -0.085 | 0 | 0.857849168 | 0.923455894 | 0.959435926 | 0.999652205 | 0.998402594 | 0.9990274 | 0.992962986 |
| Etf1 | High | Q8BWY3 | Eukaryotic peptide chain release factor subunit 1 OS=Mus musculus OX=10090 GN=Etf1 PE=1 SV= | 0 | 16.615 | 10 | 4 | 5 | 4 | 437 | 49 | 5.71 | 0.998 | 1.122 | 0.889 | 0 | 0 | 0.17 | 0.05666667 | -0.085 | -0.17 | 0.857515351 | 0.789110717 | 0.476950336 | 0.999652205 | 0.998402594 | 0.9990274 | 0.963350336 |
| Plekho2 | High | Q8K124 | Pleckstrin homology domain-containing family O member 2 OS=Mus musculus OX=10090 GN=Ple | 0 | 26.529 | 20 | 6 | 6 | 6 | 495 | 53.8 | 5.5 | 1.023 | 1.1 | 0.93 | 0.03 | -0.03 | 0.14 | 0.04666667 | -0.085 | -0.1 | 0.977534223 | 0.836944221 | 0.642304773 | 0.999652205 | 0.998402594 | 0.9990274 | 0.985010103 |
| Sntb1 | High | Q99L88 | Beta-1-syntrophin OS=Mus musculus OX=10090 GN=Sntb1 PE=1 SV=4 | 0 | 31.089 | 24 | 11 | 15 | 9 | 537 | 58 | 8.31 | 1.031 | 1.092 | 0.944 | 0.04 | -0.04 | 0.13 | 0.04333333 | -0.085 | -0.08 | 0.863814032 | 0.853980406 | 0.781936601 | 0.999652205 | 0.998402594 | 0.9990274 | 0.992535481 |
| | High | A0A7N9VF | AHNAK nucleoprotein 2 OS=Mus musculus OX=10090 GN=Ahnak2 PE=4 SV=1 | 0 | 36.792 | 37 | 8 | 11 | 8 | 3501 | 368.2 | 5.88 | 1.046 | 1.075 | 0.974 | 0.07 | -0.07 | 0.1 | 0.03333333 | -0.085 | -0.04 | 0.85619495 | 0.892648945 | 0.964454959 | 0.999652205 | 0.998402594 | 0.9990274 | 0.994302941 |
| Gid8 | Low | Q9D7M1 | Glucose-induced degradation protein 8 homolog OS=Mus musculus OX=10090 GN=Gid8 PE=1 SV= | 0.052 | 1.464 | 3 | 1 | 1 | 1 | 228 | 26.8 | 4.97 | 1.112 | 1.021 | 1.088 | 0.15 | -0.15 | 0.03 | 0.01 | -0.09 | 0.12 | 0.73980021 | 0.974992919 | 0.645142285 | 0.999652205 | 0.998402594 | 0.9990274 | 0.985010103 |
| 2210404O | High | Q0VG18 | Small integral membrane protein 24 OS=Mus musculus OX=10090 GN=Smim24 PE=1 SV=1 | 0.001 | 6.412 | 11 | 1 | 1 | 1 | 120 | 13.5 | 8.12 | 1.097 | 1.038 | 1.057 | 0.13 | -0.13 | 0.05 | 0.01666667 | -0.09 | 0.08 | 0.818727096 | 0.990880176 | 0.678575122 | 0.999652205 | 0.998402594 | 0.9990274 | 0.990892882 |
| Dpp10 | High | E9QN98 | Inactive dipeptidyl peptidase 10 OS=Mus musculus OX=10090 GN=Dpp10 PE=1 SV=1 | 0.001 | 6.115 | 2 | 2 | 2 | 1 | 800 | 91.1 | 6.48 | 1.033 | 1.098 | 0.941 | 0.05 | -0.05 | 0.13 | 0.04333333 | -0.09 | -0.09 | 0.916822453 | 0.834607438 | 0.738213659 | 0.999652205 | 0.998402594 | 0.9990274 | 0.992535481 |
| Fahd1 | High | Q8R0F8 | Acylpyruvase FAHD1, mitochondrial OS=Mus musculus OX=10090 GN=Fahd1 PE=1 SV=2 | 0.001 | 6.988 | 14 | 2 | 2 | 2 | 227 | 25.2 | 7.69 | 1.04 | 1.085 | 0.958 | 0.06 | -0.06 | 0.12 | 0.04 | -0.09 | -0.06 | 0.946257623 | 0.860713227 | 0.804897171 | 0.999652205 | 0.998402594 | 0.9990274 | 0.992535481 |
| Hmgcl | High | P38060 | Hydroxymethylglutaryl-CoA lyase, mitochondrial OS=Mus musculus OX=10090 GN=Hmgcl PE=1 SV | 0.001 | 7.083 | 6 | 2 | 2 | 2 | 325 | 34.2 | 8.41 | 1.074 | 1.059 | 1.014 | 0.1 | -0.1 | 0.08 | 0.02666667 | -0.09 | 0.02 | 0.897230996 | 0.939360188 | 0.896404667 | 0.999652205 | 0.998402594 | 0.9990274 | 0.992535481 |
| Snap23 | High | O09044 | Synaptosomal-associated protein 23 OS=Mus musculus OX=10090 GN=Snap23 PE=1 SV=1 | 0 | 51.406 | 48 | 7 | 18 | 6 | 210 | 23.2 | 4.98 | 1.125 | 1.007 | 1.118 | 0.17 | -0.17 | 0.01 | 0.00333333 | -0.09 | 0.16 | 0.738857861 | 0.947570778 | 0.536304478 | 0.999652205 | 0.998402594 | 0.9990274 | 0.96694628 |
| Gstm5 | High | P48774 | Glutathione S-transferase Mu 5 OS=Mus musculus OX=10090 GN=Gstm5 PE=1 SV=1 | 0 | 11.878 | 26 | 6 | 6 | 5 | 224 | 26.6 | 7.21 | 1.032 | 1.1 | 0.938 | 0.04 | -0.04 | 0.14 | 0.04666667 | -0.09 | -0.09 | 0.929986287 | 0.836988361 | 0.738763647 | 0.999652205 | 0.998402594 | 0.9990274 | 0.992535481 |
| Kit | High | P05532 | Mast/stem cell growth factor receptor Kit OS=Mus musculus OX=10090 GN=Kit PE=1 SV=3 | 0.001 | 5.684 | 2 | 2 | 2 | 1 | 979 | 109.3 | 7.21 | 1.112 | 1.025 | 1.085 | 0.15 | -0.15 | 0.04 | 0.01333333 | -0.095 | 0.12 | 0.746826177 | 0.985611254 | 0.562176871 | 0.999652205 | 0.998402594 | 0.9990274 | 0.972910028 |
| Golga7 | High | Q91W53 | Golgin subfamily A member 7 OS=Mus musculus OX=10090 GN=Golga7 PE=1 SV=1 | 0 | 41.698 | 50 | 7 | 21 | 7 | 137 | 15.8 | 7.05 | 1.016 | 1.125 | 0.903 | 0.02 | -0.02 | 0.17 | 0.05666667 | -0.095 | -0.15 | 0.955865576 | 0.783516238 | 0.577164486 | 0.999652205 | 0.998402594 | 0.9990274 | 0.972910028 |
| Adam10 | High | O35598 | Disintegrin and metalloproteinase domain-containing protein 10 OS=Mus musculus OX=10090 GN | 0 | 58.802 | 20 | 12 | 15 | 12 | 749 | 83.9 | 7.94 | 1.103 | 1.034 | 1.066 | 0.14 | -0.14 | 0.05 | 0.01666667 | -0.095 | 0.09 | 0.713517638 | 0.985496023 | 0.588185447 | 0.999652205 | 0.998402594 | 0.9990274 | 0.973095093 |
| Sarm1 | High | Q6PDS3 | NAD(+) hydrolase SARM1 OS=Mus musculus OX=10090 GN=Sarm1 PE=1 SV=1 | 0 | 95.065 | 32 | 20 | 28 | 20 | 724 | 79.6 | 6.27 | 1.102 | 1.032 | 1.068 | 0.14 | -0.14 | 0.05 | 0.01666667 | -0.095 | 0.1 | 0.741126466 | 0.991693064 | 0.599841923 | 0.999652205 | 0.998402594 | 0.9990274 | 0.977451587 |
| Mboat7 | High | Q8CHK3 | Lysophospholipid acyltransferase 7 OS=Mus musculus OX=10090 GN=Mboat7 PE=1 SV=1 | 0 | 12.223 | 3 | 2 | 5 | 2 | 473 | 53.4 | 8.69 | 1.018 | 1.118 | 0.911 | 0.03 | -0.03 | 0.16 | 0.05333333 | -0.095 | -0.13 | 0.992113734 | 0.798520121 | 0.636980397 | 0.999652205 | 0.998402594 | 0.9990274 | 0.983597794 |
| Dars | High | Q922B2 | Aspartate--tRNA ligase, cytoplasmic OS=Mus musculus OX=10090 GN=Dars1 PE=1 SV=2 | 0 | 33.04 | 20 | 9 | 10 | 9 | 501 | 57.1 | 6.49 | 1.045 | 1.098 | 0.952 | 0.06 | -0.06 | 0.13 | 0.04333333 | -0.095 | -0.07 | 0.945817008 | 0.840714536 | 0.728953245 | 0.999652205 | 0.998402594 | 0.9990274 | 0.992535481 |
| Mapre1 | High | Q61166 | Microtubule-associated protein RP/EB family member 1 OS=Mus musculus OX=10090 GN=Mapre | 0 | 45.813 | 51 | 9 | 10 | 8 | 268 | 30 | 5.22 | 1.1 | 1.035 | 1.062 | 0.14 | -0.14 | 0.05 | 0.01666667 | -0.095 | 0.09 | 0.800951535 | 0.983163912 | 0.74792346 | 0.999652205 | 0.998402594 | 0.9990274 | 0.992535481 |
| | High | G3UYV7 | 40S ribosomal protein S28 (Fragment) OS=Mus musculus OX=10090 GN=Rps28 PE=1 SV=1 | 0 | 15.219 | 57 | 3 | 10 | 3 | 56 | 6.3 | 9.96 | 1.041 | 1.097 | 0.949 | 0.06 | -0.06 | 0.13 | 0.04333333 | -0.095 | -0.08 | 0.906269266 | 0.842075925 | 0.786800101 | 0.999652205 | 0.998402594 | 0.9990274 | 0.992535481 |
| | High | G3UY75 | Protein-tyrosine-phosphatase OS=Mus musculus OX=10090 GN=Ptprd PE=1 SV=1 | 0 | 11.137 | 2 | 4 | 4 | 4 | 1890 | 211.7 | 6.73 | 1.084 | 1.051 | 1.032 | 0.12 | -0.12 | 0.07 | 0.02333333 | -0.095 | 0.05 | 0.823268787 | 0.947552725 | 0.80739167 | 0.999652205 | 0.998402594 | 0.9990274 | 0.992535481 |
| Rtn3 | High | Q9ES97 | Reticulon-3 OS=Mus musculus OX=10090 GN=Rtn3 PE=1 SV=2 | 0 | 15.917 | 3 | 3 | 5 | 2 | 964 | 103.8 | 4.92 | 1.05 | 1.086 | 0.967 | 0.07 | -0.07 | 0.12 | 0.04 | -0.095 | -0.05 | 0.813765736 | 0.866902318 | 0.87044581 | 0.999652205 | 0.998402594 | 0.9990274 | 0.992535481 |
| | High | G3UZU3 | Metabotropic glutamate receptor 7 OS=Mus musculus OX=10090 GN=Grm7 PE=1 SV=1 | 0.002 | 4.216 | 10 | 1 | 1 | 1 | 188 | 20.3 | 7.28 | 1.071 | 1.064 | 1.007 | 0.1 | -0.1 | 0.09 | 0.03 | -0.095 | 0.01 | 0.84539256 | 0.922063066 | 0.976237352 | 0.999652205 | 0.998402594 | 0.9990274 | 0.997221123 |
| Acot1 | High | O55137 | Acyl-coenzyme A thioesterase 1 OS=Mus musculus OX=10090 GN=Acot1 PE=1 SV=1 | 0 | 13.979 | 12 | 5 | 5 | 5 | 419 | 46.1 | 6.58 | 1.072 | 1.068 | 1.003 | 0.1 | -0.1 | 0.09 | 0.03 | -0.095 | 0 | 0.797051887 | 0.907720057 | 0.980945535 | 0.999652205 | 0.998402594 | 0.9990274 | 0.997765878 |
| | High | A0A0J9YV | Paxillin OS=Mus musculus OX=10090 GN=Pxn PE=1 SV=1 | 0 | 20.889 | 10 | 4 | 6 | 4 | 424 | 46.5 | 6.95 | 1.124 | 1.02 | 1.102 | 0.17 | -0.17 | 0.03 | 0.01 | -0.1 | 0.14 | 0.688041589 | 0.980034262 | 0.529726998 | 0.999652205 | 0.998402594 | 0.9990274 | 0.966226179 |

| | | | | | | | | | | | | | | | | | | | | | | | | | | | | |
|---|---|---|---|---|---|---|---|---|---|---|---|---|---|---|---|---|---|---|---|---|---|---|---|---|---|---|---|---|
| Aldh9a1 | High | Q3U367 | 4-trimethylaminobutyraldehyde dehydrogenase OS=Mus musculus OX=10090 GN=Aldh9a1 PE=1 S | 0 | 48.343 | 25 | 11 | 14 | 11 | 518 | 55.9 | 7.46 | 1.111 | 1.035 | 1.073 | 0.15 | -0.15 | 0.05 | 0.01666667 | -0.1 | 0.1 | 0.752084296 | 0.983483317 | 0.597362509 | 0.999652205 | 0.998402594 | 0.9990274 | 0.977451587 |
| Usp10 | High | P52479 | Ubiquitin carboxyl-terminal hydrolase 10 OS=Mus musculus OX=10090 GN=Usp10 PE=1 SV=3 | 0 | 4.694 | 2 | 2 | 2 | 2 | 792 | 87 | 5.17 | 1.099 | 1.04 | 1.057 | 0.14 | -0.14 | 0.06 | 0.02 | -0.1 | 0.08 | 0.777147092 | 0.96748957 | 0.759917945 | 0.999652205 | 0.998402594 | 0.9990274 | 0.992535481 |
| Commd7 | Medium | Q8BG94 | COMM domain-containing protein 7 OS=Mus musculus OX=10090 GN=Commd7 PE=1 SV=1 | 0.047 | 1.571 | 4 | 1 | 1 | 1 | 200 | 22.6 | 5.94 | 1.098 | 1.05 | 1.046 | 0.13 | -0.13 | 0.07 | 0.02333333 | -0.1 | 0.06 | 0.785270609 | 0.941816768 | 0.806637431 | 0.999652205 | 0.998402594 | 0.9990274 | 0.992535481 |
| Uqcr10 | High | Q8R1I1 | Cytochrome b-c1 complex subunit 9 OS=Mus musculus OX=10090 GN=Uqcr10 PE=1 SV=1 | 0 | 15.431 | 27 | 1 | 3 | 1 | 64 | 7.4 | 9.19 | 1.049 | 1.094 | 0.959 | 0.07 | -0.07 | 0.13 | 0.04333333 | -0.1 | -0.06 | 0.872620398 | 0.849590143 | 0.828532765 | 0.999652205 | 0.998402594 | 0.9990274 | 0.992535481 |
| Dnajc6 | High | Q80TZ3 | Putative tyrosine-protein phosphatase auxilin OS=Mus musculus OX=10090 GN=Dnajc6 PE=1 SV=2 | 0 | 47.788 | 16 | 12 | 15 | 12 | 938 | 102.2 | 7.23 | 1.053 | 1.091 | 0.965 | 0.07 | -0.07 | 0.13 | 0.04333333 | -0.1 | -0.05 | 0.801515466 | 0.85702212 | 0.898017925 | 0.999652205 | 0.998402594 | 0.9990274 | 0.992535481 |
| Celf4 | High | Q7TSY6 | CUGBP Elav-like family member 4 OS=Mus musculus OX=10090 GN=Celf4 PE=1 SV=2 | 0.005 | 3.646 | 3 | 2 | 2 | 2 | 486 | 51.9 | 8 | 1.064 | 1.079 | 0.986 | 0.09 | -0.09 | 0.11 | 0.03666667 | -0.1 | -0.02 | 0.871829069 | 0.88181049 | 0.981466887 | 0.999652205 | 0.998402594 | 0.9990274 | 0.997765878 |
| Amer2; Fa | High | Q8CCJ4 | APC membrane recruitment protein 2 OS=Mus musculus OX=10090 GN=Amer2 PE=1 SV=2 | 0 | 28.517 | 13 | 8 | 9 | 8 | 672 | 69.9 | 6.64 | 1.13 | 1.019 | 1.109 | 0.18 | -0.18 | 0.03 | 0.01 | -0.105 | 0.15 | 0.680235321 | 0.978399889 | 0.48547032 | 0.999652205 | 0.998402594 | 0.9990274 | 0.964378027 |
| Slc4a2 | Low | A0A0R4J1 | Anion exchange protein OS=Mus musculus OX=10090 GN=Slc4a2 PE=1 SV=1 | 0.054 | 1.39 | 1 | 1 | 1 | 1 | 1237 | 136.7 | 6.14 | 1.04 | 1.107 | 0.939 | 0.06 | -0.06 | 0.15 | 0.05 | -0.105 | -0.09 | 0.911935234 | 0.814670159 | 0.73848413 | 0.999652205 | 0.998402594 | 0.9990274 | 0.992535481 |
| | High | F7AMS7 | Misshapen-like kinase 1 (Fragment) OS=Mus musculus OX=10090 GN=Mink1 PE=1 SV=1 | 0 | 11.437 | 3 | 4 | 4 | 4 | 1198 | 135 | 8.48 | 1.09 | 1.065 | 1.023 | 0.12 | -0.12 | 0.09 | 0.03 | -0.105 | 0.03 | 0.805646509 | 0.914220534 | 0.836528933 | 0.999652205 | 0.998402594 | 0.9990274 | 0.992535481 |
| Cntfr | High | O88507 | Ciliary neurotrophic factor receptor subunit alpha OS=Mus musculus OX=10090 GN=Cntfr PE=1 SV | 0.001 | 6.679 | 11 | 3 | 3 | 3 | 372 | 40.8 | 6.83 | 1.084 | 1.067 | 1.017 | 0.12 | -0.12 | 0.09 | 0.03 | -0.105 | 0.02 | 0.816165864 | 0.925516998 | 0.850882499 | 0.999652205 | 0.998402594 | 0.9990274 | 0.992535481 |
| Rps6kb1 | High | Q8C3J7 | Ribosomal protein S6 kinase beta-1 OS=Mus musculus OX=10090 GN=Rps6kb1 PE=1 SV=1 | 0.005 | 3.599 | 3 | 1 | 1 | 1 | 316 | 35.8 | 5.59 | 1.063 | 1.083 | 0.981 | 0.09 | -0.09 | 0.12 | 0.04 | -0.105 | -0.03 | 0.882804806 | 0.887582793 | 0.985781347 | 0.999652205 | 0.998402594 | 0.9990274 | 0.998604519 |
| Idh3g | High | P70404 | Isocitrate dehydrogenase [NAD] subunit gamma 1, mitochondrial OS=Mus musculus OX=10090 G | 0 | 27.602 | 29 | 8 | 11 | 7 | 393 | 42.8 | 9.01 | 1.114 | 1.032 | 1.08 | 0.16 | -0.16 | 0.05 | 0.01666667 | -0.105 | 0.11 | 0.716265536 | 0.992009101 | 0.652036628 | 0.999652205 | 0.998402594 | 0.9990274 | 0.985010103 |
| Edc4 | High | A0A0R4J1 | Enhancer of mRNA-decapping protein 4 OS=Mus musculus OX=10090 GN=Edc4 PE=1 SV=1 | 0.001 | 5.435 | 2 | 2 | 2 | 2 | 1390 | 150.6 | 5.68 | 1.092 | 1.06 | 1.031 | 0.13 | -0.13 | 0.08 | 0.02666667 | -0.105 | 0.04 | 0.76260489 | 0.917571982 | 0.838963596 | 0.999652205 | 0.998402594 | 0.9990274 | 0.992535481 |
| Slc7a5 | High | Q9Z127 | Large neutral amino acids transporter small subunit 1 OS=Mus musculus OX=10090 GN=Slc7a5 Pl | 0.001 | 6.726 | 4 | 2 | 2 | 2 | 512 | 55.8 | 7.9 | 1.2 | 0.974 | 1.233 | 0.26 | -0.26 | -0.04 | -0.0133333 | -0.11 | 0.3 | 0.516219794 | 0.844492713 | 0.181071075 | 0.999652205 | 0.998402594 | 0.9990274 | 0.759531947 |
| Pabpc4 | High | Q91YZ8 | Polyadenylate-binding protein OS=Mus musculus OX=10090 GN=Pabpc4 PE=1 SV=1 | 0 | 40.758 | 10 | 5 | 11 | 1 | 615 | 67.8 | 9.5 | 1.167 | 1.002 | 1.165 | 0.22 | -0.22 | 0 | 0 | -0.11 | 0.22 | 0.596670918 | 0.925504157 | 0.31794977 | 0.999652205 | 0.998402594 | 0.9990274 | 0.904943958 |
| Pygl | High | Q9ET01 | Glycogen phosphorylase, liver form OS=Mus musculus OX=10090 GN=Pygl PE=1 SV=4 | 0 | 37.54 | 12 | 10 | 14 | 5 | 850 | 97.4 | 7.09 | 1.111 | 1.053 | 1.055 | 0.15 | -0.15 | 0.07 | 0.02333333 | -0.11 | 0.08 | 0.721151381 | 0.941954359 | 0.724858107 | 0.999652205 | 0.998402594 | 0.9990274 | 0.992535481 |
| | High | A0A571BE | AP-3 complex subunit beta OS=Mus musculus OX=10090 GN=Ap3b2 PE=1 SV=1 | 0 | 45.042 | 14 | 14 | 17 | 14 | 1050 | 115.4 | 5.54 | 1.088 | 1.07 | 1.017 | 0.12 | -0.12 | 0.1 | 0.03333333 | -0.11 | 0.02 | 0.756079463 | 0.902397454 | 0.814925667 | 0.999652205 | 0.998402594 | 0.9990274 | 0.992535481 |
| Tes | High | Q3ULB1 | Testin OS=Mus musculus OX=10090 GN=Tes PE=1 SV=1 | 0 | 55.167 | 36 | 13 | 16 | 13 | 410 | 46.6 | 7.87 | 1.083 | 1.076 | 1.006 | 0.11 | -0.11 | 0.11 | 0.03666667 | -0.11 | 0.01 | 0.812282439 | 0.888616524 | 0.896756997 | 0.999652205 | 0.998402594 | 0.9990274 | 0.992535481 |
| Wdr48 | High | Q8BH57 | WD repeat-containing protein 48 OS=Mus musculus OX=10090 GN=Wdr48 PE=1 SV=1 | 0 | 26.416 | 13 | 8 | 8 | 8 | 676 | 76 | 7.17 | 1.092 | 1.064 | 1.026 | 0.13 | -0.13 | 0.09 | 0.03 | -0.11 | 0.04 | 0.763817896 | 0.915723487 | 0.932496858 | 0.999652205 | 0.998402594 | 0.9990274 | 0.992535481 |
| | High | A0A1C7CY | Plastin-3 (Fragment) OS=Mus musculus OX=10090 GN=Pls3 PE=1 SV=1 | 0 | 17.934 | 12 | 7 | 7 | 6 | 629 | 70.6 | 5.62 | 1.076 | 1.077 | 0.999 | 0.11 | -0.11 | 0.11 | 0.03666667 | -0.11 | 0 | 0.80802582 | 0.888044784 | 0.947135608 | 0.999652205 | 0.998402594 | 0.9990274 | 0.992535481 |
| Lima1 | High | Q9ERG0 | LIM domain and actin-binding protein 1 OS=Mus musculus OX=10090 GN=Lima1 PE=1 SV=3 | 0 | 21.695 | 9 | 8 | 9 | 8 | 753 | 84 | 6.6 | 1.125 | 1.038 | 1.083 | 0.17 | -0.17 | 0.05 | 0.01666667 | -0.11 | 0.12 | 0.725616904 | 0.976473489 | 0.637003996 | 0.999652205 | 0.998402594 | 0.9990274 | 0.983597794 |
| Kars | High | Q8R2P8 | Lysine--tRNA ligase OS=Mus musculus OX=10090 GN=Kars PE=1 SV=1 | 0 | 28.032 | 13 | 8 | 10 | 8 | 624 | 71.2 | 6.43 | 1.06 | 1.104 | 0.96 | 0.08 | -0.08 | 0.14 | 0.04666667 | -0.11 | -0.06 | 0.877156989 | 0.827296474 | 0.778964684 | 0.999652205 | 0.998402594 | 0.9990274 | 0.992535481 |
| Lamtor3; ( | High | O88653 | Ragulator complex protein LAMTOR3 OS=Mus musculus OX=10090 GN=Lamtor3 PE=1 SV=1 | 0 | 28.004 | 50 | 3 | 8 | 3 | 124 | 13.5 | 7.34 | 1.054 | 1.112 | 0.948 | 0.08 | -0.08 | 0.15 | 0.05 | -0.115 | -0.08 | 0.899428065 | 0.811551937 | 0.744141076 | 0.999652205 | 0.998402594 | 0.9990274 | 0.992535481 |
| | Low | A0A0J9YU | FRY-like transcription coactivator (Fragment) OS=Mus musculus OX=10090 GN=Fryl PE=1 SV=3 | 0.075 | 1.244 | 0 | 1 | 1 | 1 | 1464 | 163.1 | 5.49 | 1.195 | 0.977 | 1.224 | 0.26 | -0.26 | -0.03 | -0.01 | -0.115 | 0.29 | 0.540935294 | 0.861508137 | 0.18811779 | 0.999652205 | 0.998402594 | 0.9990274 | 0.775001064 |
| | High | A0A338P7 | Dynein light chain Tctex-type 1A OS=Mus musculus OX=10090 GN=Dynlt1a PE=1 SV=1 | 0 | 12.496 | 30 | 2 | 2 | 2 | 112 | 12.2 | 5.12 | 1.176 | 1 | 1.177 | 0.23 | -0.23 | 0 | 0 | -0.115 | 0.23 | 0.651726815 | 0.913461792 | 0.252538343 | 0.999652205 | 0.998402594 | 0.9990274 | 0.851275262 |
| Abce1 | High | P61222 | ATP-binding cassette sub-family E member 1 OS=Mus musculus OX=10090 GN=Abce1 PE=1 SV=1 | 0 | 35.833 | 16 | 9 | 13 | 9 | 599 | 67.3 | 8.34 | 1.168 | 1.007 | 1.159 | 0.22 | -0.22 | 0.01 | 0.00333333 | -0.115 | 0.21 | 0.544451775 | 0.94959278 | 0.29278199 | 0.999652205 | 0.998402594 | 0.9990274 | 0.893646499 |
| Dnajc8 | High | Q6NZB0 | DnaJ homolog subfamily C member 8 OS=Mus musculus OX=10090 GN=Dnajc8 PE=1 SV=2 | 0 | 8.37 | 13 | 3 | 3 | 3 | 253 | 29.8 | 9.06 | 1.149 | 1.019 | 1.128 | 0.2 | -0.2 | 0.03 | 0.01 | -0.115 | 0.17 | 0.630709867 | 0.976834687 | 0.471767648 | 0.999652205 | 0.998402594 | 0.9990274 | 0.961211268 |
| Fam120b | Low | Q6RI63 | Constitutive coactivator of peroxisome proliferator-activated receptor gamma OS=Mus musculus | 0.054 | 1.417 | 1 | 1 | 1 | 1 | 786 | 89.3 | 6.52 | 1.119 | 1.049 | 1.067 | 0.16 | -0.16 | 0.07 | 0.02333333 | -0.115 | 0.09 | 0.72278463 | 0.954935224 | 0.727046524 | 0.999652205 | 0.998402594 | 0.9990274 | 0.992535481 |
| Tex2 | High | B1ATR2 | Testis-expressed protein 2 OS=Mus musculus OX=10090 GN=Tex2 PE=1 SV=1 | 0.005 | 3.738 | 1 | 1 | 1 | 1 | 844 | 92.7 | 6.2 | 1.052 | 1.118 | 0.941 | 0.07 | -0.07 | 0.16 | 0.05333333 | -0.115 | -0.09 | 0.916848591 | 0.786406827 | 0.744227508 | 0.999652205 | 0.998402594 | 0.9990274 | 0.992535481 |
| Dlg3 | High | A2BEE9 | Disks large homolog 3 OS=Mus musculus OX=10090 GN=Dlg3 PE=1 SV=1 | 0.001 | 7.162 | 3 | 2 | 2 | 1 | 835 | 92.1 | 6.86 | 1.1 | 1.064 | 1.033 | 0.14 | -0.14 | 0.09 | 0.03 | -0.115 | 0.05 | 0.769646047 | 0.92928979 | 0.781139108 | 0.999652205 | 0.998402594 | 0.9990274 | 0.992535481 |
| Cfl2 | High | P45591 | Cofilin-2 OS=Mus musculus OX=10090 GN=Cfl2 PE=1 SV=1 | 0 | 26.592 | 37 | 6 | 16 | 3 | 166 | 18.7 | 7.88 | 1.243 | 0.954 | 1.303 | 0.31 | -0.31 | -0.07 | -0.0233333 | -0.12 | 0.38 | 0.451575405 | 0.817010653 | 0.09816495 | 0.999652205 | 0.998402594 | 0.9990274 | 0.598923694 |
| | High | A0A3B2W | DNA-directed RNA polymerase II subunit GRINL1A OS=Mus musculus OX=10090 GN=Polr2m PE=4 | 0 | 14.039 | 57 | 2 | 2 | 2 | 84 | 8.6 | 10.48 | 1.114 | 1.055 | 1.056 | 0.16 | -0.16 | 0.08 | 0.02666667 | -0.12 | 0.08 | 0.696461714 | 0.931933567 | 0.731125878 | 0.999652205 | 0.998402594 | 0.9990274 | 0.992535481 |
| Mtpn | High | P62774 | Myotrophin OS=Mus musculus OX=10090 GN=Mtpn PE=1 SV=2 | 0 | 20.141 | 28 | 3 | 6 | 3 | 118 | 12.9 | 5.52 | 1.117 | 1.056 | 1.058 | 0.16 | -0.16 | 0.08 | 0.02666667 | -0.12 | 0.08 | 0.705002334 | 0.935310502 | 0.764630866 | 0.999652205 | 0.998402594 | 0.9990274 | 0.992535481 |
| Gapdh | High | A0A0A0M | Glyceraldehyde-3-phosphate dehydrogenase OS=Mus musculus OX=10090 GN=Gapdh PE=1 SV=1 | 0 | 820.529 | 87 | 35 | 943 | 1 | 359 | 38.6 | 8.97 | 1.055 | 1.118 | 0.943 | 0.08 | -0.08 | 0.16 | 0.05333333 | -0.12 | -0.08 | 0.854575976 | 0.797341839 | 0.783499878 | 0.999652205 | 0.998402594 | 0.9990274 | 0.992535481 |
| Fscn1 | High | Q61553 | Fascin OS=Mus musculus OX=10090 GN=Fscn1 PE=1 SV=4 | 0 | 53.531 | 35 | 14 | 19 | 14 | 493 | 54.5 | 6.89 | 1.186 | 0.999 | 1.187 | 0.25 | -0.25 | 0 | 0 | -0.125 | 0.25 | 0.52760479 | 0.928721622 | 0.22757013 | 0.999652205 | 0.998402594 | 0.9990274 | 0.816808422 |
| Gdpd5 | High | Q640M6 | Glycerophosphodiester phosphodiesterase domain-containing protein 5 OS=Mus musculus OX=10 | 0.001 | 7.125 | 2 | 1 | 1 | 1 | 607 | 68.8 | 8 | 1.19 | 1.002 | 1.187 | 0.25 | -0.25 | 0 | 0 | -0.125 | 0.25 | 0.535251717 | 0.92671136 | 0.336386686 | 0.999652205 | 0.998402594 | 0.9990274 | 0.906329766 |
| Sap18; Gm | Medium | E9Q317 | 18 kDa Sin3-associated polypeptide OS=Mus musculus OX=10090 GN=Sap18 PE=1 SV=1 | 0.019 | 2.053 | 4 | 1 | 1 | 1 | 172 | 19.6 | 9.8 | 1.137 | 1.04 | 1.093 | 0.19 | -0.19 | 0.06 | 0.02 | -0.125 | 0.13 | 0.697100377 | 0.995091965 | 0.623390894 | 0.999652205 | 0.998402594 | 0.9990274 | 0.97826833 |
| Pum2 | Medium | Q3TQ29 | Pumilio homolog 2 OS=Mus musculus OX=10090 GN=Pum2 PE=1 SV=1 | 0.01 | 2.591 | 1 | 1 | 1 | 1 | 980 | 105.6 | 6.86 | 1.068 | 1.111 | 0.962 | 0.1 | -0.1 | 0.15 | 0.05 | -0.125 | -0.06 | 0.863265789 | 0.801854993 | 0.843663643 | 0.999652205 | 0.998402594 | 0.9990274 | 0.992535481 |
| Map1a; M | High | A2ARP8 | Microtubule-associated protein 1A OS=Mus musculus OX=10090 GN=Map1a PE=1 SV=1 | 0 | 58.646 | 6 | 15 | 15 | 13 | 3014 | 325.7 | 5.01 | 1.106 | 1.081 | 1.023 | 0.14 | -0.14 | 0.11 | 0.03666667 | -0.125 | 0.03 | 0.761879737 | 0.878806113 | 0.869862669 | 0.999652205 | 0.998402594 | 0.9990274 | 0.992535481 |
| Hdac11 | High | Q91WA3 | Histone deacetylase 11 OS=Mus musculus OX=10090 GN=Hdac11 PE=1 SV=1 | 0 | 22.06 | 20 | 6 | 7 | 6 | 347 | 39.1 | 7.14 | 1.083 | 1.094 | 0.99 | 0.12 | -0.12 | 0.13 | 0.04333333 | -0.125 | -0.01 | 0.732301643 | 0.85031459 | 0.979999004 | 0.999652205 | 0.998402594 | 0.9990274 | 0.997765878 |
| Mthfd1l | High | Q3V3R1 | Monofunctional C1-tetrahydrofolate synthase, mitochondrial OS=Mus musculus OX=10090 GN=M | 0 | 8.734 | 2 | 2 | 2 | 2 | 977 | 105.7 | 7.02 | 1.083 | 1.099 | 0.985 | 0.11 | -0.11 | 0.14 | 0.04666667 | -0.125 | -0.02 | 0.824195451 | 0.838119242 | 0.984736953 | 0.999652205 | 0.998402594 | 0.9990274 | 0.998604519 |
| BC027072 | High | Q6PAC4 | Photoreceptor cilium actin regulator OS=Mus musculus OX=10090 GN=Pcare PE=2 SV=1 | 0 | 46.898 | 11 | 14 | 15 | 14 | 1279 | 139.2 | 8.06 | 1.243 | 0.966 | 1.286 | 0.31 | -0.31 | -0.05 | -0.0166667 | -0.13 | 0.36 | 0.467345492 | 0.848869397 | 0.113757572 | 0.999652205 | 0.998402594 | 0.9990274 | 0.643956085 |
| Hspa1a; H | High | Q61696 | Heat shock 70 kDa protein 1A OS=Mus musculus OX=10090 GN=Hspa1a PE=1 SV=2 | 0 | 326.878 | 58 | 38 | 140 | 1 | 641 | 70 | 5.72 | 1.175 | 1.018 | 1.154 | 0.23 | -0.23 | 0.03 | 0.01 | -0.13 | 0.21 | 0.58534298 | 0.976134036 | 0.306962847 | 0.999652205 | 0.998402594 | 0.9990274 | 0.903485924 |
| Stard7 | High | Q8R1R3 | StAR-related lipid transfer protein 7, mitochondrial OS=Mus musculus OX=10090 GN=Stard7 PE= | 0 | 10.709 | 10 | 3 | 4 | 3 | 373 | 43.1 | 8.95 | 1.167 | 1.025 | 1.139 | 0.22 | -0.22 | 0.04 | 0.01333333 | -0.13 | 0.19 | 0.535625701 | 0.992989132 | 0.436668188 | 0.999652205 | 0.998402594 | 0.9990274 | 0.949727902 |
| Eif3c | High | Q8R1B4 | Eukaryotic translation initiation factor 3 subunit C OS=Mus musculus OX=10090 GN=Eif3c PE=1 S | 0 | 19.893 | 7 | 7 | 8 | 7 | 911 | 105.5 | 5.78 | 1.086 | 1.103 | 0.985 | 0.12 | -0.12 | 0.14 | 0.04666667 | -0.13 | -0.02 | 0.829846501 | 0.830665709 | 0.892276081 | 0.999652205 | 0.998402594 | 0.9990274 | 0.992535481 |
| Gcdh | Medium | A0A0A0M | Glutaryl-CoA dehydrogenase, mitochondrial OS=Mus musculus OX=10090 GN=Gcdh PE=1 SV=1 | 0.015 | 2.212 | 2 | 1 | 1 | 1 | 447 | 49.5 | 8.72 | 1.179 | 1.02 | 1.156 | 0.24 | -0.24 | 0.03 | 0.01 | -0.135 | 0.21 | 0.599117178 | 0.964847554 | 0.313426982 | 0.999652205 | 0.998402594 | 0.9990274 | 0.904943958 |
| Atxn2l | High | Q7TQH0 | Ataxin-2-like protein OS=Mus musculus OX=10090 GN=Atxn2l PE=1 SV=1 | 0 | 19.728 | 6 | 7 | 8 | 5 | 1049 | 110.6 | 8.85 | 1.132 | 1.063 | 1.065 | 0.18 | -0.18 | 0.09 | 0.03 | -0.135 | 0.09 | 0.61336505 | 0.919532795 | 0.693603505 | 0.999652205 | 0.998402594 | 0.9990274 | 0.991285312 |
| Ufc1 | Low | Q9CR09 | Ubiquitin-fold modifier-conjugating enzyme 1 OS=Mus musculus OX=10090 GN=Ufc1 PE=1 SV=1 | 0.085 | 1.18 | 4 | 1 | 1 | 1 | 167 | 19.5 | 7.4 | 1.106 | 1.087 | 1.017 | 0.15 | -0.15 | 0.12 | 0.04 | -0.135 | 0.02 | 0.783746996 | 0.885253976 | 0.907601095 | 0.999652205 | 0.998402594 | 0.9990274 | 0.992535481 |
| Suclg1 | High | Q9WUM5 | Succinate--CoA ligase [ADP/GDP-forming] subunit alpha, mitochondrial OS=Mus musculus OX=10 | 0 | 45.169 | 29 | 9 | 14 | 9 | 346 | 36.1 | 9.39 | 1.09 | 1.112 | 0.981 | 0.12 | -0.12 | 0.15 | 0.05 | -0.135 | -0.03 | 0.768566366 | 0.811009934 | 0.979975512 | 0.999652205 | 0.998402594 | 0.9990274 | 0.997765878 |
| Upf1 | High | Q9EPU0 | Regulator of nonsense transcripts 1 OS=Mus musculus OX=10090 GN=Upf1 PE=1 SV=2 | 0 | 59.706 | 18 | 14 | 17 | 14 | 1124 | 123.9 | 6.61 | 1.167 | 1.04 | 1.122 | 0.22 | -0.22 | 0.06 | 0.02 | -0.14 | 0.17 | 0.622234571 | 0.972411576 | 0.451461719 | 0.999652205 | 0.998402594 | 0.9990274 | 0.959744396 |
| AW55491 | High | Q6NZK5 | Protein hinderin OS=Mus musculus OX=10090 GN=Kiaa1328 PE=1 SV=2 | 0.001 | 6.836 | 6 | 3 | 3 | 3 | 612 | 69.7 | 8.57 | 1.151 | 1.058 | 1.088 | 0.2 | -0.2 | 0.08 | 0.02666667 | -0.14 | 0.12 | 0.633348538 | 0.943938867 | 0.647947268 | 0.999652205 | 0.998402594 | 0.9990274 | 0.985010103 |
| Sar1b | High | Q9CQC9 | GTP-binding protein SAR1b OS=Mus musculus OX=10090 GN=Sar1b PE=1 SV=1 | 0 | 18.579 | 17 | 3 | 6 | 2 | 198 | 22.4 | 6.11 | 1.128 | 1.076 | 1.048 | 0.17 | -0.17 | 0.11 | 0.03666667 | -0.14 | 0.07 | 0.64956138 | 0.888690885 | 0.769869935 | 0.999652205 | 0.998402594 | 0.9990274 | 0.992535481 |
| | High | E9Q0H0 | Transporter OS=Mus musculus OX=10090 GN=Slc6a20a PE=3 SV=1 | 0 | 32.902 | 11 | 7 | 18 | 7 | 555 | 62 | 6.09 | 1.108 | 1.096 | 1.011 | 0.15 | -0.15 | 0.13 | 0.04333333 | -0.14 | 0.02 | 0.728050494 | 0.845028697 | 0.865760267 | 0.999652205 | 0.998402594 | 0.9990274 | 0.992535481 |
| Bloc1s6 | Medium | Q9R0C0 | Biogenesis of lysosome-related organelles complex 1 subunit 6 OS=Mus musculus OX=10090 GN= | 0.034 | 1.827 | 5 | 1 | 1 | 1 | 172 | 19.7 | 6.3 | 1.178 | 1.038 | 1.135 | 0.24 | -0.24 | 0.05 | 0.01666667 | -0.145 | 0.18 | 0.573343094 | 0.97440201 | 0.445836867 | 0.999652205 | 0.998402594 | 0.9990274 | 0.958608248 |
| | High | D3Z0Q3 | DCN1-like protein (Fragment) OS=Mus musculus OX=10090 GN=Dcun1d3 PE=1 SV=1 | 0.008 | 2.952 | 17 | 1 | 1 | 1 | 63 | 6.6 | 9.39 | 1.193 | 1.026 | 1.163 | 0.25 | -0.25 | 0.04 | 0.01333333 | -0.145 | 0.22 | 0.577398355 | 0.976198812 | 0.453402907 | 0.999652205 | 0.998402594 | 0.9990274 | 0.959744396 |
| Hdgf | High | P51859 | Hepatoma-derived growth factor OS=Mus musculus OX=10090 GN=Hdgf PE=1 SV=2 | 0 | 8.184 | 11 | 3 | 3 | 2 | 237 | 26.3 | 4.83 | 1.215 | 1.006 | 1.207 | 0.28 | -0.28 | 0.01 | 0.00333333 | -0.145 | 0.27 | 0.492894891 | 0.942320623 | 0.26850205 | 0.999652205 | 0.998402594 | 0.9990274 | 0.869723277 |
| Vamp8 | High | A0A0R4J0 | Vesicle-associated membrane protein 8 OS=Mus musculus OX=10090 GN=Vamp8 PE=1 SV=1 | 0 | 18.397 | 31 | 3 | 6 | 3 | 101 | 11.4 | 8.19 | 1.206 | 1.017 | 1.186 | 0.27 | -0.27 | 0.02 | 0.00666667 | -0.145 | 0.25 | 0.475616965 | 0.973272464 | 0.333520785 | 0.999652205 | 0.998402594 | 0.9990274 | 0.906329766 |
| Pabpc1 | High | P29341 | Polyadenylate-binding protein 1 OS=Mus musculus OX=10090 GN=Pabpc1 PE=1 SV=2 | 0 | 87.907 | 28 | 16 | 27 | 12 | 636 | 70.6 | 9.5 | 1.173 | 1.045 | 1.122 | 0.23 | -0.23 | 0.06 | 0.02 | -0.145 | 0.17 | 0.592929446 | 0.95981558 | 0.399441008 | 0.999652205 | 0.998402594 | 0.9990274 | 0.93262326 |
| Timm8b | High | P62077 | Mitochondrial import inner membrane translocase subunit Tim8 B OS=Mus musculus OX=10090 ( | 0.002 | 4.221 | 18 | 2 | 2 | 2 | 83 | 9.3 | 5.12 | 1.164 | 1.051 | 1.107 | 0.22 | -0.22 | 0.07 | 0.02333333 | -0.145 | 0.15 | 0.655902456 | 0.959257975 | 0.497295009 | 0.999652205 | 0.998402594 | 0.9990274 | 0.964378027 |
| Afg3l2 | High | Q8JZQ2 | AFG3-like protein 2 OS=Mus musculus OX=10090 GN=Afg3l2 PE=1 SV=1 | 0 | 32.903 | 11 | 8 | 10 | 8 | 802 | 89.5 | 8.6 | 1.144 | 1.072 | 1.067 | 0.19 | -0.19 | 0.1 | 0.03333333 | -0.145 | 0.09 | 0.663104408 | 0.897538257 | 0.638901055 | 0.999652205 | 0.998402594 | 0.9990274 | 0.983597794 |
| Trappc9 | High | Q3U0M1 | Trafficking protein particle complex subunit 9 OS=Mus musculus OX=10090 GN=Trappc9 PE=1 SV | 0.001 | 7.381 | 3 | 3 | 3 | 3 | 1148 | 128.2 | 6.47 | 1.137 | 1.075 | 1.058 | 0.19 | -0.19 | 0.1 | 0.03333333 | -0.145 | 0.08 | 0.666639773 | 0.891104188 | 0.686943758 | 0.999652205 | 0.998402594 | 0.9990274 | 0.991285312 |
| Wdr60 | Medium | Q8C761 | Cytoplasmic dynein 2 intermediate chain 1 OS=Mus musculus OX=10090 GN=Dync2i1 PE=1 SV=1 | 0.03 | 1.902 | 1 | 1 | 1 | 1 | 999 | 115.3 | 7.24 | 1.126 | 1.087 | 1.036 | 0.17 | -0.17 | 0.12 | 0.04 | -0.145 | 0.05 | 0.688130734 | 0.86520913 | 0.813177237 | 0.999652205 | 0.998402594 | 0.9990274 | 0.992535481 |
| Arl6 | High | O88848 | ADP-ribosylation factor-like protein 6 OS=Mus musculus OX=10090 GN=Arl6 PE=1 SV=1 | 0 | 201.975 | 88 | 19 | 71 | 19 | 186 | 20.9 | 8.25 | 1.137 | 1.078 | 1.055 | 0.19 | -0.19 | 0.11 | 0.03666667 | -0.15 | 0.08 | 0.663150809 | 0.88472305 | 0.657495401 | 0.999652205 | 0.998402594 | 0.9990274 | 0.985010103 |
| Fech | High | P22315 | Ferrochelatase, mitochondrial OS=Mus musculus OX=10090 GN=Fech PE=1 SV=2 | 0.001 | 5.328 | 6 | 2 | 2 | 2 | 420 | 47.1 | 8.91 | 1.177 | 1.054 | 1.116 | 0.23 | -0.23 | 0.08 | 0.02666667 | -0.155 | 0.16 | 0.564885497 | 0.951912693 | 0.445798619 | 0.999652205 | 0.998402594 | 0.9990274 | 0.958608248 |
| Dip2b | High | Q3UH60 | Disco-interacting protein 2 homolog B OS=Mus musculus OX=10090 GN=Dip2b PE=1 SV=1 | 0 | 85.788 | 16 | 19 | 26 | 5 | 1574 | 171 | 8.09 | 1.156 | 1.074 | 1.076 | 0.21 | -0.21 | 0.1 | 0.03333333 | -0.155 | 0.11 | 0.623733487 | 0.893804506 | 0.566736246 | 0.999652205 | 0.998402594 | 0.9990274 | 0.972910028 |
| 6530401N | High | Q8BHA3 | D-aminoacyl-tRNA deacylase 2 OS=Mus musculus OX=10090 GN=Dtd2 PE=1 SV=1 | 0 | 23.962 | 45 | 5 | 9 | 5 | 168 | 18.2 | 7.91 | 1.109 | 1.118 | 0.992 | 0.15 | -0.15 | 0.16 | 0.05333333 | -0.155 | -0.01 | 0.729063876 | 0.797859815 | 0.908784268 | 0.999652205 | 0.998402594 | 0.9990274 | 0.992535481 |
| Gpm6b | High | P35803 | Neuronal membrane glycoprotein M6-b OS=Mus musculus OX=10090 GN=Gpm6b PE=1 SV=2 | 0 | 25.984 | 15 | 6 | 12 | 6 | 328 | 36.2 | 6.14 | 1.236 | 1.008 | 1.227 | 0.31 | -0.31 | 0.01 | 0.00333333 | -0.16 | 0.29 | 0.473628705 | 0.950117783 | 0.156274736 | 0.999652205 | 0.998402594 | 0.9990274 | 0.731412725 |
| Rabl2 | High | E9Q9D5 | Rab-like protein 2A OS=Mus musculus OX=10090 GN=Rabl2 PE=1 SV=1 | 0.001 | 6.879 | 8 | 2 | 2 | 2 | 223 | 25.6 | 5.66 | 1.235 | 1.012 | 1.22 | 0.3 | -0.3 | 0.02 | 0.00666667 | -0.16 | 0.29 | 0.445561385 | 0.942933911 | 0.190940009 | 0.999652205 | 0.998402594 | 0.9990274 | 0.783175296 |
| Sec24b | Low | Q80ZX0 | Sec24-related gene family, member B (S. cerevisiae) OS=Mus musculus OX=10090 GN=Sec24b PE | 0.074 | 1.252 | 1 | 1 | 1 | 1 | 1251 | 135.5 | 6.96 | 1.164 | 1.075 | 1.083 | 0.22 | -0.22 | 0.1 | 0.03333333 | -0.16 | 0.11 | 0.578440259 | 0.885964511 | 0.624503742 | 0.999652205 | 0.998402594 | 0.9990274 | 0.978918782 |
| | High | K3W4S6 | Glycogenin OS=Mus musculus OX=10090 GN=Gyg PE=1 SV=1 | 0 | 14.461 | 6 | 2 | 3 | 2 | 377 | 41.9 | 6.02 | 1.157 | 1.08 | 1.071 | 0.21 | -0.21 | 0.11 | 0.03666667 | -0.16 | 0.1 | 0.614049261 | 0.880485075 | 0.699931973 | 0.999652205 | 0.998402594 | 0.9990274 | 0.991285312 |
| Dhx30 | High | A0A0G2JG | RNA helicase OS=Mus musculus OX=10090 GN=Dhx30 PE=1 SV=1 | 0 | 12.606 | 4 | 4 | 4 | 4 | 1186 | 132.9 | 8.59 | 1.13 | 1.101 | 1.026 | 0.18 | -0.18 | 0.14 | 0.04666667 | -0.16 | 0.04 | 0.658088098 | 0.833233099 | 0.873253001 | 0.999652205 | 0.998402594 | 0.9990274 | 0.992535481 |
| Tbc1d10a | High | Q5SPX8 | TBC1 domain family member 10A OS=Mus musculus OX=10090 GN=Tbc1d10a PE=1 SV=1 | 0 | 18.13 | 14 | 6 | 6 | 6 | 534 | 59.2 | 8.02 | 1.11 | 1.122 | 0.99 | 0.15 | -0.15 | 0.17 | 0.05666667 | -0.16 | -0.01 | 0.687086427 | 0.79012916 | 0.982889688 | 0.999652205 | 0.998402594 | 0.9990274 | 0.997765878 |
| Tppp3 | High | Q9CRB6 | Tubulin polymerization-promoting protein family member 3 OS=Mus musculus OX=10090 GN=Tp | 0.001 | 7.719 | 13 | 2 | 2 | 2 | 176 | 19 | 9.11 | 1.184 | 1.068 | 1.108 | 0.24 | -0.24 | 0.09 | 0.03 | -0.165 | 0.15 | 0.587334736 | 0.929983723 | 0.551850918 | 0.999652205 | 0.998402594 | 0.9990274 | 0.96694628 |
| | High | F7BJK1 | Protocadherin 1 (Fragment) OS=Mus musculus OX=10090 GN=Pcdh1 PE=1 SV=1 | 0.001 | 7.137 | 2 | 3 | 3 | 3 | 1077 | 116.4 | 5.08 | 1.131 | 1.112 | 1.017 | 0.18 | -0.18 | 0.15 | 0.05 | -0.165 | 0.02 | 0.68280128 | 0.820691912 | 0.849731602 | 0.999652205 | 0.998402594 | 0.9990274 | 0.992535481 |
| Rars | High | Q9D0I9 | Arginine--tRNA ligase, cytoplasmic OS=Mus musculus OX=10090 GN=Rars1 PE=1 SV=2 | 0 | 25.66 | 14 | 8 | 9 | 8 | 660 | 75.6 | 7.55 | 1.185 | 1.054 | 1.125 | 0.25 | -0.25 | 0.08 | 0.02666667 | -0.165 | 0.17 | 0.541320737 | 0.94082776 | 0.504111342 | 0.999652205 | 0.998402594 | 0.9990274 | 0.964378027 |
| Trim3 | High | Q3TDT0 | Tripartite motif-containing protein 3 OS=Mus musculus OX=10090 GN=Trim3 PE=1 SV=1 | 0 | 13.805 | 6 | 4 | 4 | 3 | 717 | 78.2 | 8.16 | 1.172 | 1.071 | 1.095 | 0.23 | -0.23 | 0.1 | 0.03333333 | -0.165 | 0.13 | 0.616691375 | 0.922005121 | 0.618425473 | 0.999652205 | 0.998402594 | 0.9990274 | 0.97826833 |
| Wasf3 | High | Q8VHI6 | Wiskott-Aldrich syndrome protein family member 3 OS=Mus musculus OX=10090 GN=Wasf3 PE= | 0 | 23.12 | 11 | 4 | 5 | 4 | 501 | 55.2 | 6.37 | 1.167 | 1.077 | 1.084 | 0.22 | -0.22 | 0.11 | 0.03666667 | -0.165 | 0.12 | 0.521323696 | 0.887753214 | 0.631600232 | 0.999652205 | 0.998402594 | 0.9990274 | 0.982425436 |
| Tbcc | High | Q8VCN9 | Tubulin-specific chaperone C OS=Mus musculus OX=10090 GN=Tbcc PE=1 SV=1 | 0.001 | 5.085 | 5 | 2 | 2 | 2 | 341 | 38.1 | 5.3 | 1.163 | 1.088 | 1.069 | 0.22 | -0.22 | 0.12 | 0.04 | -0.17 | 0.1 | 0.585523642 | 0.853769261 | 0.684541394 | 0.999652205 | 0.998402594 | 0.9990274 | 0.991285312 |
| Crip2 | High | Q9DCT8 | Cysteine-rich protein 2 OS=Mus musculus OX=10090 GN=Crip2 PE=1 SV=1 | 0.001 | 6.533 | 15 | 2 | 2 | 2 | 208 | 22.7 | 8.63 | 1.141 | 1.112 | 1.026 | 0.19 | -0.19 | 0.15 | 0.05 | -0.17 | 0.04 | 0.659949493 | 0.821953851 | 0.843400451 | 0.999652205 | 0.998402594 | 0.9990274 | 0.992535481 |
| Rpl37a | High | P61514 | 60S ribosomal protein L37a OS=Mus musculus OX=10090 GN=Rpl37a PE=1 SV=2 | 0 | 21.758 | 50 | 5 | 8 | 5 | 92 | 10.3 | 10.43 | 1.213 | 1.043 | 1.163 | 0.28 | -0.28 | 0.06 | 0.02 | -0.17 | 0.22 | 0.549948476 | 0.964766305 | 0.364068219 | 0.999652205 | 0.998402594 | 0.9990274 | 0.917864435 |
| | High | A0A1Y7VI | Cyclin-dependent kinase 20 OS=Mus musculus OX=10090 GN=Cdk20 PE=1 SV=1 | 0 | 8.416 | 13 | 2 | 2 | 1 | 188 | 21 | 8.75 | 1.201 | 1.06 | 1.133 | 0.26 | -0.26 | 0.08 | 0.02666667 | -0.17 | 0.18 | 0.500454792 | 0.925312844 | 0.410760502 | 0.999652205 | 0.998402594 | 0.9990274 | 0.936397673 |
| Lancl1 | High | O89112 | Glutathione S-transferase LANCL1 OS=Mus musculus OX=10090 GN=Lancl1 PE=1 SV=1 | 0 | 39.628 | 19 | 6 | 11 | 6 | 399 | 45.3 | 7.77 | 1.146 | 1.103 | 1.039 | 0.2 | -0.2 | 0.14 | 0.04666667 | -0.17 | 0.05 | 0.660647388 | 0.828857056 | 0.741024943 | 0.999652205 | 0.998402594 | 0.9990274 | 0.992535481 |
| Pcbp1 | High | P60335 | Poly(rC)-binding protein 1 OS=Mus musculus OX=10090 GN=Pcbp1 PE=1 SV=1 | 0 | 97.413 | 47 | 12 | 32 | 8 | 356 | 37.5 | 7.09 | 1.129 | 1.122 | 1.006 | 0.17 | -0.17 | 0.17 | 0.05666667 | -0.17 | 0.01 | 0.681283968 | 0.789466011 | 0.889147048 | 0.999652205 | 0.998402594 | 0.9990274 | 0.992535481 |
| Gstt3 | High | Q99L20 | Glutathione S-transferase theta-3 OS=Mus musculus OX=10090 GN=Gstt3 PE=1 SV=1 | 0 | 14.545 | 17 | 4 | 4 | 3 | 241 | 27.4 | 7.74 | 1.19 | 1.073 | 1.109 | 0.25 | -0.25 | 0.1 | 0.03333333 | -0.175 | 0.15 | 0.563250943 | 0.896666549 | 0.500903851 | 0.999652205 | 0.998402594 | 0.9990274 | 0.964378027 |
| Sbds | High | P70122 | Ribosome maturation protein SBDS OS=Mus musculus OX=10090 GN=Sbds PE=1 SV=4 | 0 | 10.875 | 15 | 4 | 6 | 4 | 250 | 28.8 | 8.76 | 1.268 | 1.009 | 1.257 | 0.34 | -0.34 | 0.01 | 0.00333333 | -0.175 | 0.33 | 0.449852931 | 0.954095355 | 0.174618663 | 0.999652205 | 0.998402594 | 0.9990274 | 0.744450291 |
| Eif3i | High | Q9QZD9 | Eukaryotic translation initiation factor 3 subunit I OS=Mus musculus OX=10090 GN=Eif3i PE=1 SV | 0 | 22.257 | 17 | 5 | 5 | 5 | 325 | 36.4 | 5.64 | 1.206 | 1.06 | 1.138 | 0.27 | -0.27 | 0.08 | 0.02666667 | -0.175 | 0.19 | 0.5190406 | 0.92697519 | 0.384394957 | 0.999652205 | 0.998402594 | 0.9990274 | 0.927809953 |
| Nudt21 | High | Q9CQF3 | Cleavage and polyadenylation specificity factor subunit 5 OS=Mus musculus OX=10090 GN=Nudt2 | 0 | 10.654 | 20 | 4 | 5 | 4 | 227 | 26.2 | 8.82 | 1.236 | 1.037 | 1.192 | 0.31 | -0.31 | 0.05 | 0.01666667 | -0.18 | 0.25 | 0.457871066 | 0.979111802 | 0.265399407 | 0.999652205 | 0.998402594 | 0.9990274 | 0.865171207 |
| Rfc4 | Low | Q3UI84 | Replication factor C subunit 4 OS=Mus musculus OX=10090 GN=Rfc4 PE=1 SV=1 | 0.055 | 1.382 | 3 | 1 | 1 | 1 | 370 | 40.9 | 8.53 | 1.182 | 1.088 | 1.086 | 0.24 | -0.24 | 0.12 | 0.04 | -0.18 | 0.12 | 0.601382207 | 0.874153874 | 0.559654464 | 0.999652205 | 0.998402594 | 0.9990274 | 0.972910028 |
| Ptprk | High | P35822 | Receptor-type tyrosine-protein phosphatase kappa OS=Mus musculus OX=10090 GN=Ptprk PE=1 S | 0.001 | 4.658 | 1 | 1 | 1 | 1 | 1457 | 164.1 | 5.95 | 1.169 | 1.098 | 1.064 | 0.22 | -0.22 | 0.14 | 0.04666667 | -0.18 | 0.09 | 0.604913926 | 0.861869553 | 0.686360998 | 0.999652205 | 0.998402594 | 0.9990274 | 0.991285312 |
| Npr1 | High | P18293 | Atrial natriuretic peptide receptor 1 OS=Mus musculus OX=10090 GN=Npr1 PE=1 SV=2 | 0 | 34.966 | 6 | 5 | 18 | 4 | 1057 | 119 | 6.89 | 1.159 | 1.106 | 1.047 | 0.21 | -0.21 | 0.15 | 0.05 | -0.18 | 0.07 | 0.630532671 | 0.826761941 | 0.733403121 | 0.999652205 | 0.998402594 | 0.9990274 | 0.992535481 |
| Gabbr1 | High | Q9WV18 | Gamma-aminobutyric acid type B receptor subunit 1 OS=Mus musculus OX=10090 GN=Gabbr1 PE | 0 | 8.936 | 3 | 2 | 3 | 2 | 960 | 108.1 | 8.21 | 1.151 | 1.115 | 1.032 | 0.2 | -0.2 | 0.16 | 0.05333333 | -0.18 | 0.05 | 0.645689341 | 0.82396461 | 0.800826724 | 0.999652205 | 0.998402594 | 0.9990274 | 0.992535481 |
| Fahd2a | High | A0A0R4J0 | Fumarylacetoacetate hydrolase domain-containing 2A OS=Mus musculus OX=10090 GN=Fahd2a I | 0 | 9.705 | 12 | 3 | 4 | 3 | 313 | 34.7 | 8.16 | 1.138 | 1.124 | 1.012 | 0.19 | -0.19 | 0.17 | 0.05666667 | -0.18 | 0.02 | 0.678660467 | 0.786135294 | 0.898360968 | 0.999652205 | 0.998402594 | 0.9990274 | 0.992535481 |
| Kif2a | High | P28740 | Kinesin-like protein KIF2A OS=Mus musculus OX=10090 GN=Kif2a PE=1 SV=2 | 0.001 | 5.212 | 4 | 3 | 3 | 3 | 705 | 79.7 | 6.73 | 1.137 | 1.128 | 1.008 | 0.19 | -0.19 | 0.17 | 0.05666667 | -0.18 | 0.01 | 0.675572578 | 0.781145557 | 0.908919258 | 0.999652205 | 0.998402594 | 0.9990274 | 0.992535481 |
| Tbc1d24 | High | Q3UUG6 | TBC1 domain family member 24 OS=Mus musculus OX=10090 GN=Tbc1d24 PE=1 SV=2 | 0 | 34.117 | 17 | 9 | 10 | 9 | 561 | 63.2 | 7.24 | 1.252 | 1.036 | 1.209 | 0.32 | -0.32 | 0.05 | 0.01666667 | -0.185 | 0.27 | 0.460756061 | 0.982582643 | 0.228030263 | 0.999652205 | 0.998402594 | 0.9990274 | 0.816974879 |
| Adam22 | Medium | Q9R1V6 | Disintegrin and metalloproteinase domain-containing protein 22 OS=Mus musculus OX=10090 GN | 0.03 | 1.903 | 1 | 1 | 1 | 1 | 904 | 99.7 | 7.83 | 1.211 | 1.065 | 1.136 | 0.28 | -0.28 | 0.09 | 0.03 | -0.185 | 0.18 | 0.493028266 | 0.930221527 | 0.403849562 | 0.999652205 | 0.998402594 | 0.9990274 | 0.934095017 |
| Aifm2 | High | Q8BUE4 | Ferroptosis suppressor protein 1 OS=Mus musculus OX=10090 GN=Aifm2 PE=1 SV=1 | 0 | 8.773 | 9 | 3 | 3 | 3 | 373 | 40.6 | 8.98 | 1.17 | 1.099 | 1.065 | 0.23 | -0.23 | 0.14 | 0.04666667 | -0.185 | 0.09 | 0.590085741 | 0.853317981 | 0.650199553 | 0.999652205 | 0.998402594 | 0.9990274 | 0.985010103 |
| Nim1; Nin | High | Q8BHI9 | Serine/threonine-protein kinase NIM1 OS=Mus musculus OX=10090 GN=Nim1k PE=2 SV=1 | 0 | 10.272 | 8 | 3 | 3 | 3 | 436 | 49.7 | 8.68 | 1.159 | 1.121 | 1.034 | 0.21 | -0.21 | 0.16 | 0.05333333 | -0.185 | 0.05 | 0.616952993 | 0.792855744 | 0.780184448 | 0.999652205 | 0.998402594 | 0.9990274 | 0.992535481 |
| Taldo1 | High | Q93092 | Transaldolase OS=Mus musculus OX=10090 GN=Taldo1 PE=1 SV=2 | 0 | 18.303 | 20 | 7 | 7 | 7 | 337 | 37.4 | 7.03 | 1.157 | 1.12 | 1.033 | 0.21 | -0.21 | 0.16 | 0.05333333 | -0.185 | 0.05 | 0.619366125 | 0.792954284 | 0.85067523 | 0.999652205 | 0.998402594 | 0.9990274 | 0.992535481 |
| | High | E9PY39 | Predicted gene 20431 OS=Mus musculus OX=10090 GN=Gm20431 PE=3 SV=1 | 0 | 16.059 | 18 | 5 | 5 | 5 | 371 | 42.1 | 6.76 | 1.14 | 1.132 | 1.007 | 0.19 | -0.19 | 0.18 | 0.06 | -0.185 | 0.01 | 0.658826984 | 0.779141934 | 0.919718961 | 0.999652205 | 0.998402594 | 0.9990274 | 0.992535481 |
| Vac14 | Medium | Q80WQ2 | Protein VAC14 homolog OS=Mus musculus OX=10090 GN=Vac14 PE=1 SV=1 | 0.034 | 1.819 | 1 | 1 | 2 | 1 | 782 | 88 | 6.13 | 1.175 | 1.112 | 1.056 | 0.23 | -0.23 | 0.15 | 0.05 | -0.19 | 0.08 | 0.580039689 | 0.814358936 | 0.68706102 | 0.999652205 | 0.998402594 | 0.9990274 | 0.991285312 |
| Dhx36 | High | Q8VHK9 | ATP-dependent DNA/RNA helicase DHX36 OS=Mus musculus OX=10090 GN=Dhx36 PE=1 SV=2 | 0.001 | 6.064 | 2 | 2 | 2 | 2 | 1001 | 113.8 | 8.29 | 1.171 | 1.113 | 1.052 | 0.23 | -0.23 | 0.15 | 0.05 | -0.19 | 0.07 | 0.590246673 | 0.819487919 | 0.700124839 | 0.999652205 | 0.998402594 | 0.9990274 | 0.991285312 |
| Gtf3c3 | High | Q3TMP1 | General transcription factor IIIC, polypeptide 3 OS=Mus musculus OX=10090 GN=Gtf3c3 PE=1 SV= | 0 | 11.851 | 3 | 3 | 3 | 3 | 882 | 100.6 | 5.15 | 1.254 | 1.042 | 1.204 | 0.33 | -0.33 | 0.06 | 0.02 | -0.195 | 0.27 | 0.435200443 | 0.968620318 | 0.238481009 | 0.999652205 | 0.998402594 | 0.9990274 | 0.830020494 |
| Rras2 | High | P62071 | Ras-related protein R-Ras2 OS=Mus musculus OX=10090 GN=Rras2 PE=1 SV=1 | 0 | 31.477 | 25 | 5 | 9 | 3 | 204 | 23.4 | 6.01 | 1.239 | 1.054 | 1.175 | 0.31 | -0.31 | 0.08 | 0.02666667 | -0.195 | 0.23 | 0.462181859 | 0.939102518 | 0.289907644 | 0.999652205 | 0.998402594 | 0.9990274 | 0.889103759 |
| Myo5a | High | D3YZ62 | Unconventional myosin-Va OS=Mus musculus OX=10090 GN=Myo5a PE=1 SV=1 | 0 | 183.559 | 26 | 48 | 66 | 41 | 1828 | 212.2 | 8.62 | 1.229 | 1.064 | 1.155 | 0.3 | -0.3 | 0.09 | 0.03 | -0.195 | 0.21 | 0.486306931 | 0.917096298 | 0.303718406 | 0.999652205 | 0.998402594 | 0.9990274 | 0.900241969 |
| | High | G3UWR2 | GRB10-interacting GYF protein 2 OS=Mus musculus OX=10090 GN=Gigyf2 PE=1 SV=1 | 0 | 8.12 | 1 | 1 | 1 | 1 | 1122 | 129.6 | 5.43 | 1.199 | 1.097 | 1.093 | 0.26 | -0.26 | 0.13 | 0.04333333 | -0.195 | 0.13 | 0.554262763 | 0.866464023 | 0.585995685 | 0.999652205 | 0.998402594 | 0.9990274 | 0.973095093 |
| Phgdh | High | Q61753 | D-3-phosphoglycerate dehydrogenase OS=Mus musculus OX=10090 GN=Phgdh PE=1 SV=3 | 0 | 113.521 | 37 | 18 | 31 | 18 | 533 | 56.5 | 6.54 | 1.164 | 1.128 | 1.033 | 0.22 | -0.22 | 0.17 | 0.05666667 | -0.195 | 0.05 | 0.606376341 | 0.777623377 | 0.758666936 | 0.999652205 | 0.998402594 | 0.9990274 | 0.992535481 |
| Pcnxl4; Pc | High | Q3UVY5 | Pecanex-like protein 4 OS=Mus musculus OX=10090 GN=Pcnx4 PE=2 SV=2 | 0 | 9.385 | 4 | 3 | 3 | 3 | 1174 | 131.2 | 6.54 | 1.225 | 1.082 | 1.132 | 0.29 | -0.29 | 0.11 | 0.03666667 | -0.2 | 0.18 | 0.460451812 | 0.886805619 | 0.423079863 | 0.999652205 | 0.998402594 | 0.9990274 | 0.939860852 |
| Sar1a | High | Q99JZ4 | GTP-binding protein SAR1a OS=Mus musculus OX=10090 GN=Sar1a PE=1 SV=1 | 0 | 54.861 | 39 | 6 | 12 | 5 | 198 | 22.4 | 6.68 | 1.211 | 1.087 | 1.114 | 0.28 | -0.28 | 0.12 | 0.04 | -0.2 | 0.16 | 0.542060054 | 0.864542208 | 0.451558909 | 0.999652205 | 0.998402594 | 0.9990274 | 0.959744396 |
| Camk2a | High | P11798 | Calcium/calmodulin-dependent protein kinase type II subunit alpha OS=Mus musculus OX=10090 | 0 | 29.703 | 13 | 5 | 12 | 2 | 478 | 54.1 | 7.08 | 1.216 | 1.087 | 1.119 | 0.28 | -0.28 | 0.12 | 0.04 | -0.2 | 0.16 | 0.472894449 | 0.856675551 | 0.493434905 | 0.999652205 | 0.998402594 | 0.9990274 | 0.964378027 |
| Aldh4a1 | Medium | Q8CHT0 | Delta-1-pyrroline-5-carboxylate dehydrogenase, mitochondrial OS=Mus musculus OX=10090 GN=/ | 0.026 | 1.968 | 1 | 1 | 1 | 1 | 562 | 61.8 | 8.24 | 1.186 | 1.109 | 1.069 | 0.25 | -0.25 | 0.15 | 0.05 | -0.2 | 0.1 | 0.588001828 | 0.825925093 | 0.627010531 | 0.999652205 | 0.998402594 | 0.9990274 | 0.980381496 |
| Lmln | High | Q8BMN4 | Leishmanolysin-like peptidase OS=Mus musculus OX=10090 GN=Lmln PE=2 SV=1 | 0 | 10.58 | 6 | 4 | 4 | 4 | 681 | 76.5 | 7.28 | 1.182 | 1.115 | 1.06 | 0.24 | -0.24 | 0.16 | 0.05333333 | -0.2 | 0.08 | 0.567973792 | 0.803336701 | 0.672763893 | 0.999652205 | 0.998402594 | 0.9990274 | 0.990250803 |
| Mid1 | Medium | O70583 | E3 ubiquitin-protein ligase Midline-1 OS=Mus musculus OX=10090 GN=Mid1 PE=1 SV=2 | 0.01 | 2.721 | 2 | 1 | 1 | 1 | 680 | 76.1 | 6.93 | 1.169 | 1.122 | 1.042 | 0.23 | -0.23 | 0.17 | 0.05666667 | -0.2 | 0.06 | 0.566691729 | 0.783365106 | 0.773666896 | 0.999652205 | 0.998402594 | 0.9990274 | 0.992535481 |
| Dnal4; Dna | High | Q8CDT8 | Dynein light chain OS=Mus musculus OX=10090 GN=Dnal4 PE=1 SV=1 | 0.002 | 4.19 | 7 | 1 | 1 | 1 | 139 | 15.9 | 6.54 | 1.232 | 1.078 | 1.143 | 0.3 | -0.3 | 0.11 | 0.03666667 | -0.205 | 0.19 | 0.447941672 | 0.89130579 | 0.462043725 | 0.999652205 | 0.998402594 | 0.9990274 | 0.959744396 |
| Ptplad1; H | High | Q8K2C9 | Very-long-chain (3R)-3-hydroxyacyl-CoA dehydratase 3 OS=Mus musculus OX=10090 GN=Hacd3 P | 0 | 11.542 | 14 | 5 | 5 | 5 | 362 | 43.1 | 9.13 | 1.236 | 1.071 | 1.154 | 0.31 | -0.31 | 0.1 | 0.03333333 | -0.205 | 0.21 | 0.446376559 | 0.900724525 | 0.416285386 | 0.999652205 | 0.998402594 | 0.9990274 | 0.936397673 |
| Ccz1 | Medium | Q8C1Y8 | Vacuolar fusion protein CCZ1 homolog OS=Mus musculus OX=10090 GN=Ccz1 PE=1 SV=1 | 0.013 | 2.259 | 3 | 1 | 1 | 1 | 480 | 55.5 | 6.02 | 1.229 | 1.089 | 1.129 | 0.3 | -0.3 | 0.12 | 0.04 | -0.21 | 0.17 | 0.50047716 | 0.883512146 | 0.480773482 | 0.999652205 | 0.998402594 | 0.9990274 | 0.964378027 |

| | | | | | | | | | | | | | | | | | | | | | | | | | | | | |
|---|---|---|---|---|---|---|---|---|---|---|---|---|---|---|---|---|---|---|---|---|---|---|---|---|---|---|---|---|
| Pacsin3 | High | Q99JB8 | Protein kinase C and casein kinase II substrate protein 3 OS=Mus musculus OX=10090 GN=Pacsin3 | 0 | 8.626 | 5 | 2 | 2 | 2 | 424 | 48.6 | 6.1 | 1.211 | 1.1 | 1.1 | 0.28 | -0.28 | 0.14 | 0.04666667 | -0.21 | 0.14 | 0.486400205 | 0.843080084 | 0.507593448 | 0.999652205 | 0.998402594 | 0.9990274 | 0.964378027 |
| Abcf2 | High | Q99LE6 | ATP-binding cassette sub-family F member 2 OS=Mus musculus OX=10090 GN=Abcf2 PE=1 SV=1 | 0 | 8.937 | 6 | 4 | 4 | 4 | 628 | 71.7 | 7.05 | 1.247 | 1.082 | 1.152 | 0.32 | -0.32 | 0.11 | 0.03666667 | -0.215 | 0.2 | 0.430054675 | 0.875075761 | 0.413809836 | 0.999652205 | 0.998402594 | 0.9990274 | 0.936397673 |
| Apc | High | B2RUG9 | Adenomatosis polyposis coli OS=Mus musculus OX=10090 GN=Apc PE=1 SV=1 | 0.001 | 5.045 | 1 | 2 | 2 | 2 | 2842 | 310.7 | 7.58 | 1.196 | 1.126 | 1.062 | 0.26 | -0.26 | 0.17 | 0.05666667 | -0.215 | 0.09 | 0.551517237 | 0.787159278 | 0.670681829 | 0.999652205 | 0.998402594 | 0.9990274 | 0.989088116 |
| | High | A0A1B0G | 60S ribosomal protein L18 OS=Mus musculus OX=10090 GN=Rpl18 PE=1 SV=1 | 0 | 29.768 | 36 | 6 | 10 | 6 | 159 | 18.1 | 11.91 | 1.204 | 1.118 | 1.076 | 0.27 | -0.27 | 0.16 | 0.05333333 | -0.215 | 0.11 | 0.571591745 | 0.796882192 | 0.675594808 | 0.999652205 | 0.998402594 | 0.9990274 | 0.990250803 |
| Fh1 | High | P97807 | Fumarate hydratase, mitochondrial OS=Mus musculus OX=10090 GN=Fh PE=1 SV=3 | 0 | 68.338 | 33 | 14 | 20 | 14 | 507 | 54.3 | 9.04 | 1.231 | 1.105 | 1.114 | 0.3 | -0.3 | 0.14 | 0.04666667 | -0.22 | 0.16 | 0.481631291 | 0.824545083 | 0.426199546 | 0.999652205 | 0.998402594 | 0.9990274 | 0.941822646 |
| Rpl7a | High | P12970 | 60S ribosomal protein L7a OS=Mus musculus OX=10090 GN=Rpl7a PE=1 SV=2 | 0 | 96.419 | 50 | 16 | 33 | 16 | 266 | 30 | 10.56 | 1.252 | 1.094 | 1.145 | 0.32 | -0.32 | 0.13 | 0.04333333 | -0.225 | 0.2 | 0.447448833 | 0.850067463 | 0.331246228 | 0.999652205 | 0.998402594 | 0.9990274 | 0.906329766 |
| Mt1 | High | P02802 | Metallothionein-1 OS=Mus musculus OX=10090 GN=Mt1 PE=1 SV=1 | 0 | 21.709 | 34 | 3 | 11 | 3 | 61 | 6 | 7.96 | 1.241 | 1.126 | 1.102 | 0.31 | -0.31 | 0.17 | 0.05666667 | -0.24 | 0.14 | 0.472036623 | 0.780339825 | 0.459155424 | 0.999652205 | 0.998402594 | 0.9990274 | 0.959744396 |
| Trim25 | High | Q5SU71 | E3 ubiquitin/ISG15 ligase TRIM25 OS=Mus musculus OX=10090 GN=Trim25 PE=1 SV=1 | 0.001 | 4.244 | 2 | 1 | 1 | 1 | 626 | 70.7 | 8.12 | 1.248 | 1.127 | 1.106 | 0.32 | -0.32 | 0.17 | 0.05666667 | -0.245 | 0.15 | 0.459468397 | 0.786160486 | 0.474403001 | 0.999652205 | 0.998402594 | 0.9990274 | 0.962227984 |
| Gbf1 | High | Q6DFZ1 | Golgi-specific brefeldin A-resistance factor 1 OS=Mus musculus OX=10090 GN=Gbf1 PE=1 SV=1 | 0.005 | 3.783 | 1 | 2 | 2 | 2 | 1861 | 206.7 | 5.8 | 0.926 | 1.031 | 0.897 | -0.11 | 0.11 | 0.04 | 0.01333333 | 0.035 | -0.16 | 0.599684474 | 0.997931774 | 0.518997471 | 0.999652205 | 0.998490794 | 0.999071499 | 0.964378027 |
| Rab30 | High | Q923S9 | Ras-related protein Rab-30 OS=Mus musculus OX=10090 GN=Rab30 PE=1 SV=1 | 0 | 20.917 | 24 | 4 | 10 | 3 | 203 | 23 | 4.97 | 0.927 | 1.027 | 0.902 | -0.11 | 0.11 | 0.04 | 0.01333333 | 0.035 | -0.15 | 0.754142242 | 0.996852663 | 0.528656978 | 0.999652205 | 0.998490794 | 0.999071499 | 0.965674043 |
| Sod1 | High | P08228 | Superoxide dismutase [Cu-Zn] OS=Mus musculus OX=10090 GN=Sod1 PE=1 SV=2 | 0 | 66.142 | 64 | 10 | 32 | 10 | 154 | 15.9 | 6.51 | 0.944 | 1.027 | 0.919 | -0.08 | 0.08 | 0.04 | 0.01333333 | 0.02 | -0.12 | 0.848276992 | 0.995804511 | 0.658740213 | 0.999652205 | 0.998490794 | 0.999071499 | 0.985010103 |
| Tecpr1 | High | Q80VP0 | Tectonin beta-propeller repeat-containing protein 1 OS=Mus musculus OX=10090 GN=Tecpr1 PE= | 0 | 14.773 | 4 | 4 | 4 | 4 | 1166 | 130.2 | 6.48 | 0.968 | 1.028 | 0.942 | -0.05 | 0.05 | 0.04 | 0.01333333 | 0.005 | -0.09 | 0.934974931 | 0.998381476 | 0.73775827 | 0.999652205 | 0.998490794 | 0.999071499 | 0.992535481 |
| Rab7 | High | P51150 | Ras-related protein Rab-7a OS=Mus musculus OX=10090 GN=Rab7a PE=1 SV=2 | 0 | 159.706 | 77 | 16 | 67 | 16 | 207 | 23.5 | 6.7 | 0.975 | 1.028 | 0.949 | -0.04 | 0.04 | 0.04 | 0.01333333 | 0 | -0.08 | 0.934213873 | 0.998490794 | 0.810029892 | 0.999652205 | 0.998490794 | 0.999071499 | 0.992535481 |
| Rab5c | High | P35278 | Ras-related protein Rab-5C OS=Mus musculus OX=10090 GN=Rab5c PE=1 SV=2 | 0 | 119.88 | 68 | 11 | 49 | 7 | 216 | 23.4 | 8.41 | 0.985 | 1.029 | 0.957 | -0.02 | 0.02 | 0.04 | 0.01333333 | -0.01 | -0.06 | 0.962469745 | 0.998490794 | 0.856869637 | 0.999652205 | 0.998490794 | 0.999071499 | 0.992535481 |
| Mat2a | High | Q3THS6 | S-adenosylmethionine synthase isoform type-2 OS=Mus musculus OX=10090 GN=Mat2a PE=1 SV | 0.001 | 6.245 | 6 | 3 | 3 | 3 | 395 | 43.7 | 6.48 | 1.017 | 1.027 | 0.99 | 0.02 | -0.02 | 0.04 | 0.01333333 | -0.03 | -0.01 | 0.982110179 | 0.998061037 | 0.990467612 | 0.999652205 | 0.998490794 | 0.999071499 | 0.999783808 |
| Plcd1 | High | G5DDB7 | Phosphoinositide phospholipase C OS=Mus musculus OX=10090 GN=Plcd1 PE=1 SV=1 | 0 | 15.909 | 9 | 6 | 6 | 6 | 782 | 88.8 | 6.65 | 1.029 | 1.03 | 0.999 | 0.04 | -0.04 | 0.04 | 0.01333333 | -0.04 | 0 | 0.887007024 | 0.996589846 | 0.987683868 | 0.999652205 | 0.998490794 | 0.999071499 | 0.998907549 |
| Clasp1 | High | A0A087W | CLIP-associating protein 1 OS=Mus musculus OX=10090 GN=Clasp1 PE=1 SV=1 | 0 | 34.035 | 9 | 11 | 11 | 7 | 1552 | 170.9 | 9.03 | 1.139 | 1.027 | 1.109 | 0.19 | -0.19 | 0.04 | 0.01333333 | -0.115 | 0.15 | 0.65775837 | 0.997664054 | 0.537453094 | 0.999652205 | 0.998490794 | 0.999071499 | 0.96694628 |
| Acot8 | Medium | P58137 | Acyl-coenzyme A thioesterase 8 OS=Mus musculus OX=10090 GN=Acot8 PE=1 SV=1 | 0.01 | 2.568 | 3 | 1 | 1 | 1 | 320 | 35.8 | 7.64 | 1.017 | 1.202 | 0.847 | 0.02 | -0.02 | 0.27 | 0.09 | -0.145 | -0.24 | 0.999613241 | 0.639329583 | 0.374252905 | 0.999707778 | 0.995559182 | 0.99763348 | 0.920164724 |
| Tbc1d10b | High | Q8BHL3 | TBC1 domain family member 10B OS=Mus musculus OX=10090 GN=Tbc1d10b PE=1 SV=2 | 0.001 | 4.585 | 5 | 3 | 3 | 3 | 798 | 87.2 | 8.9 | 1.018 | 1.02 | 0.998 | 0.03 | -0.03 | 0.03 | 0.01 | -0.03 | 0 | 0.999707778 | 0.965173585 | 0.937813441 | 0.999707778 | 0.998402594 | 0.999055186 | 0.992535481 |
