## Supplemental Table 2 for "Tectonic Complex Impedes Diffusion through the Ciliary Transition Zone to Ensure Proper Sorting of Membrane Proteins"

**Supplementary Table 2. List of Primers and Their Uses**

| Primer Name | Sequence from 5'-3' | Use |
|---|---|---|
| mTctn1-Exon2 F | CTGCCAACAGTCCTTCCTTG | For genotyping Tctn1Flox |
| mTctn1-Exon2 R | CAGGTCGCAGACACACAGAG | For genotyping Tctn1Flox |
| iCre75 F | TCAGTGCCTGGAGTTGCGCTGTGG | For genotyping rod-specific cre (iCre75) |
| iCre75 R | CTTAAAGGCCAGGGCCTGCTTGGC | For genotyping rod-specific cre (iCre75) |
| mTctn1-qPCR-Ex2-F | CTGCTCAGTCCCAGTTGTCA | Used to verify excision of Exon2 of *Tctn1* in mouse cDNA |
| mTctn1-qPCR-Ex2-R | ATGGATGCAGAAGACGGAAG | Used to verify excision of Exon2 of *Tctn1* in mouse cDNA |
| mTCTN1 seqF1 | ACGTTGCTGCTCTGTGTGTC | Sequencing primers for mouse *Tctn1* cDNA |
| mTCTN1 seqF2 | TTTATTCAGCATGGCACCAA | Sequencing primers for mouse *Tctn1* cDNA |
| mTCTN1 seqR1 | ATGGATGCAGAAGACGGAAG | Sequencing primers for mouse *Tctn1* cDNA |
| mTCTN1 seqR2 | CTGACAGCCAGACTGCACAT | Sequencing primers for mouse *Tctn1* cDNA |
| Age-mTCTN1 F | GGATCCACCGGTATGGGGTCGCGGGGTCTCCCG | Clone mouse *Tctn1* into pRho plasmid |
| mTCTN1-myc-NotI R | CTCTTCTGAGATAAGCTTTTGTTCAGCCACAAACGGGAAGAAGAAG | Add C-terminal myc tag to mouse *Tctn1* for pRho plasmid |
| Age-mKate2-F | ggatccaccggtGCCGCCACCatggtgagcgagctgattaag | Add N-terminal mKate tag to mouse *Cetn2* for pRho plasmid |
| mKate-Centrin2-R | CCATAGATCTGAGTCCGGAtctgtgccccagtttgctagg | Add N-terminal mKate tag to mouse *Cetn2* |
| mKate-Centrin2-F | agcaaactggggcacagaTCCGGACTCAGATCTATGGC | Add N-terminal mKate tag to mouse *Cetn2* |
| NotI-Centrin-R | ggagtgcggccgcTTAATAGAGGCTGGTCTTTTTCATGATGCG | Add N-terminal mKate tag to mouse *Cetn2* for pRho plasmid |
| SalI-hRK-F | GTCGACGGGCCCCAGAAGCCTGGTG | Replacing bovine rhodopin promoter with human rhodopsin kinase promoter + SV40 SD/SA |
| hRK-Kpn1-R | GCCCGCGGTACCCGGCGGGTACAATTCCGCAGC | Replacing bovine rhodopin promoter with human rhodopsin kinase promoter + SV40 SD/SA |
