## Supplemental Table 3 for "Tectonic Complex Impedes Diffusion through the Ciliary Transition Zone to Ensure Proper Sorting of Membrane Proteins"

**Supplementary Table 3. List of Antibodies**

| Name | Source | Catalog # | Dilution for Hard-fixed Eyecup IF | Dilution for Isolated Outer Segment TZ IF | Dilution for Fresh Retina TZ IF | Dilution for WB |
|---|---|---|---|---|---|---|
| Abca4 (Gt) | Everest | EB08615 | 1:1000 | NA | NA | NA |
| Cngβ1 (Rb) | Steven Pitler Lab | NA | 1:1000 | NA | NA | NA |
| Multi Ubiquitin (M) | MBL International | D058-3 | 1:1000 | NA | NA | NA |
| Myc (Rb) | Cell Signaling | 2278S | 1:1000 | NA | 1:100 | NA |
| Na/K ATPase (M) | Santa Cruz | SC58628 | 1:1000 | NA | NA | 1:5000 |
| Peripherin (Sh) | Vadim Arshavsky Lab | NA | 1:1000 | NA | NA | NA |
| Phalloidin-iFluor 647 | Abcam | ab176759 | 1:1000 | NA | NA | NA |
| R9AP (Rb) | Vadim Arshavsky Lab | NA | 1:1000 | NA | NA | NA |
| Rhodopsin-1D4 (M) | Abcam | ab5417 | 1:1000 | NA | NA | NA |
| Snap25 [SMI81] (M) | BioLegend | 836304 | 1:1000 | NA | NA | 1:2000 |
| Syntaxin3 (Rb) | Proteintech | 15556-1-AP | 1:1000 | NA | NA | 1:2000 |
| WGA-Alexa 594 | Thermo Fisher | W11262 | 1:2000 | NA | NA | NA |
| B9d1 (Rb) | Jeremy Reiter Lab | NA | NA | 1:100 | NA | NA |
| Spata7 (Rb) | Rui Chen Lab | NA | NA | 1:100 | NA | NA |
| Alpha Tubulin (M) | Sigma-Aldrich | T9026 | NA | 1:1000 | NA | NA |
| Centrin1 (M) | EMD Millipore | 04-1624 | NA | 1:1000 | 1:100 | NA |
| Cep290 (Rb) | Bathyl laboratories | A301-659A | NA | 1:1000 | 1:50 | NA |
| Bbs5 (M) | Clay Smith Lab | NA | NA | 1:200 | NA | NA |
| Ahi (Rb) | Joseph Gleeson Lab | NA | NA | 1:250 | NA | NA |
| Rpgr (Rb) | Proteintech | 16891-1-AP | NA | 1:500 | NA | NA |
| Phosducin (Sh) | Vadim Arshavsky Lab | NA | NA | NA | NA | 1:5000 |
